# Shared and distinct immunometabolic states link to divergent pathology in cutaneous mycobacterial infections

**DOI:** 10.64898/2026.10.19.760140

**Authors:** Huangtao Guo, Rui Zeng, Qin Wei, Jingshu Xiong, Haiqin Jiang, Ying Shi, Wenyue Zhang, Chun Pan, Chaojiang Cheng, Hongsheng Wang, Yetao Wang

## Abstract

Cutaneous mycobacterial infections provide a common tissue setting in which related intracellular pathogens elicit distinct pathological responses, but the innate immune basis of this divergence remains incompletely understood. Here, we combined single-cell profiling from 33 individuals with histological and functional analyses, examining healthy skin and lesions from multibacillary leprosy (MB), paucibacillary leprosy (PB), cutaneous tuberculosis (CTB) and Mycobacterium marinum infection (Mari). All infections showed lower mononuclear phagocyte and higher natural killer (NK) cell proportions within the CD45⁺ compartment, alongside shared and infection-associated dendritic cell, macrophage and NK cell programs. CTB and Mari displayed broader macrophage antimicrobial signatures, whereas MB showed lipid-associated programs and attenuated interferon-associated effector expression across multiple macrophage subsets. Reduced expression of fructose-1,6-bisphosphatase 1 (FBP1) was associated with the absence of Langhans giant cells in MB, and pharmacological FBP1 inhibition nearly abolished fusion of macrophages. Compared with NK cells from Mari, MB NK cells showed reduced central carbon and oxidative metabolism signatures alongside increased lipid-associated programs. Functional experiments showed that inhibition of glucose utilization, disruption of mitochondrial function, or co-culture with foamy macrophages abundant in MB lesions reduced NK cell IFN-γ production. These findings identify shared and distinct innate immune programs and suggest that their interplay with local metabolic environments shapes divergent pathology across cutaneous mycobacterial infections.

**GRAPHICAL ABSTRACT:** 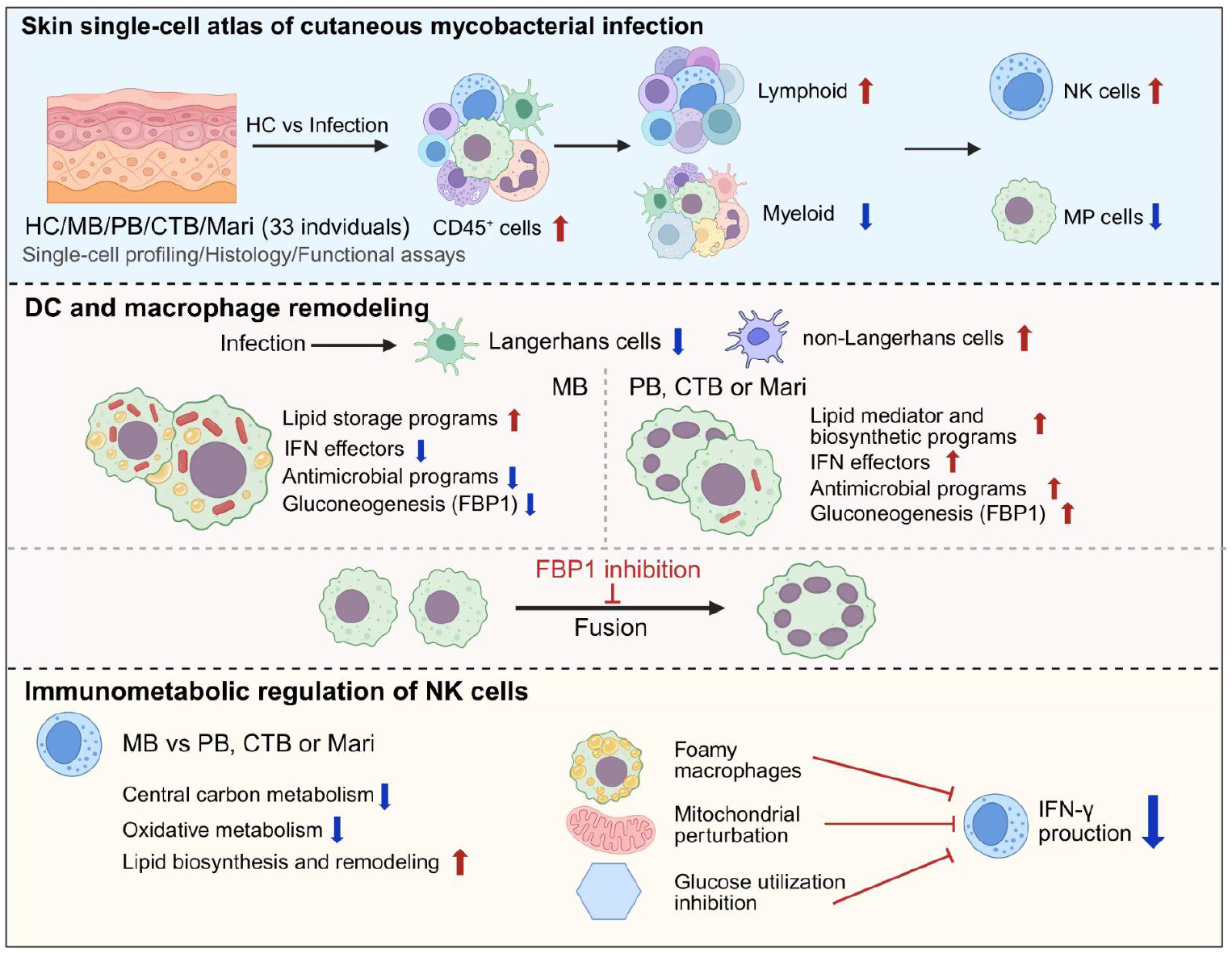

## INTRODUCTION

Mycobacterial infections cause life-threatening disease, chronic tissue injury, and permanent disability worldwide. For example, *Mycobacterium tuberculosis* infection causes tuberculosis (TB), with an estimated 10.7 million cases and 1.23 million deaths in 2024^1^, and can also involve the skin as cutaneous tuberculosis (CTB), leading to persistent lesions and scarring^2^. *Mycobacterium leprae* infection causes leprosy (Lep), with more than 170,000 new cases reported in 2024^3^; it is characterized by skin lesions and peripheral nerve damage that can lead to irreversible disability^4^. *Mycobacterium marinum* infection (Mari), a form of cutaneous nontuberculous mycobacterial disease, causes chronic skin lesions and can involve tendons, joints, and bone^5^. These infections therefore impose a substantial and lasting burden on human health.

Although these Mycobacterium species are all intracellular pathogens capable of causing cutaneous disease, they elicit distinct histopathological responses. CTB is characterized by epithelioid granulomatous inflammation, with the presence and extent of caseous necrosis varying among clinical forms^6^. *M. leprae* infection produces a broad pathological spectrum, from compact epithelioid granulomas at the tuberculoid pole to diffuse infiltrates of foamy macrophages at the lepromatous pole^7,8^. Mari exhibits suppurative and granulomatous inflammation^9,10^. However, the immunological basis of this pathological diversity remains incompletely understood. A deeper understanding may inform new approaches to clinical diagnosis and treatment.

Innate immunity plays a central role in host defense against mycobacteria by mediating early pathogen recognition and intracellular bacterial control, and by shaping adaptive immune responses that influence disease outcomes. Macrophages are major intracellular host cells for mycobacteria and exhibit distinct functional states across disease contexts^11–14^. For example, in leprosy, macrophages in tuberculoid lesions show stronger IFN-γ- and vitamin D-dependent antimicrobial activity, whereas lepromatous lesions are associated with lipid accumulation and the type I IFN–IL-10 axis that can restrain IFN-γ-mediated responses^15,16^. In Mari, macrophages can restrict early bacterial growth while also facilitating bacterial dissemination and granuloma formation and recruit permissive monocytes^17–19^. Dendritic cells (DCs) link pathogen sensing and antigen processing to adaptive immunity^20^, with stronger CD1-dependent antigen presentation in tuberculoid lesions, whereas LILRA2 activation in lepromatous disease impairs DC differentiation and *M. leprae*-specific antigen presentation^21,22^. NK cells contribute to host defense against mycobacterial infection through IFN-γ production and cytotoxicity, with stronger IL-12/IL-18-associated responses reported toward the tuberculoid or paucibacillary end of the leprosy spectrum^23,24^. Whether distinct innate immune cell states are associated with the divergent clinical manifestations of these infections warrants further investigation.

Cutaneous mycobacterial infections offer a shared tissue context for investigating how related intracellular pathogens elicit distinct pathological responses. Here, we compared macrophage, DC, and NK-cell states across leprosy, CTB, and Mari. Our analysis identified shared and infection-associated immune programs, linked fructose-1,6-bisphosphatase 1 (FBP1) expression to Langhans giant-cell formation, and showed that foamy macrophages suppressed NK-cell IFN-γ production. These findings provide insight into the cellular and metabolic basis of divergent tissue pathology and local immune responses across mycobacterial infections.

## RESULTS

### A single-cell atlas reveals shared immune remodeling across cutaneous mycobacterial infections

To define shared and disease-specific immune responses across cutaneous mycobacterial infections, we obtained skin specimens from 33 individuals, including normal skin from 6 healthy controls (HC) and lesional skin from 27 patients: 15 with *Mycobacterium leprae* infection, including 7 with multibacillary leprosy (MB) and 8 with paucibacillary leprosy (PB); 8 with *Mycobacterium tuberculosis* infection (CTB); and 4 with *Mycobacterium marinum* infection (Mari). Single-cell RNA sequencing (scRNA-seq) was performed using the 10x Genomics platform. A total of 266,203 cells passed quality control and were retained for downstream analyses. The study design is summarized in Figure 1A, and detailed participant and specimen characteristics are provided in Table S1.

**Figure 1.**
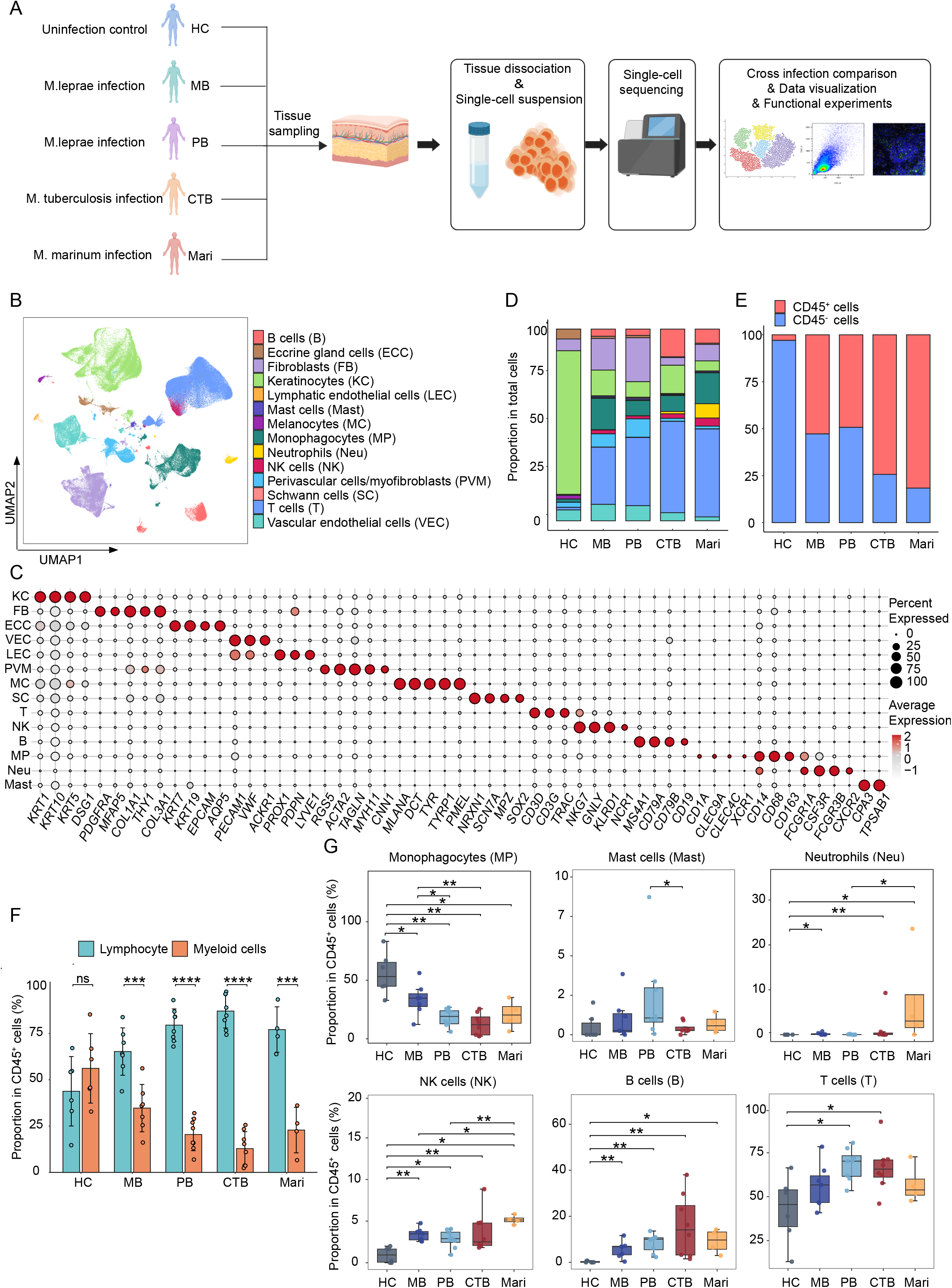
A single-cell atlas reveals shared immune remodeling across cutaneous mycobacterial infections. (A) Study design and single-cell RNA sequencing workflow for skin samples from healthy controls and patients with cutaneous mycobacterial infections. (B) UMAP of all skin cells, colored by the 14 annotated cell types. (C) Dotplot of representative cell-type markers. (D, E) Proportions of different cell types (D) and CD45⁺ versus CD45⁻ cells (E) in skin samples from HC and patients with MB, PB, CTB, or Mari. (F) Proportions of lymphoid (T, B, NK) and myeloid (MP, Mast, Neu) cells within CD45^+^ cells. (G) Proportions of indicated cell types within CD45^+^ compartment. For (F), unpaired t-test. For (G), Wilcoxon rank-sum test. Each dot represents one donor (HC, n = 6; MB, n = 7; PB, n = 8; CTB, n = 8; Mari, n = 4). ns, not significant, \**P* < 0.05, \*\**P* < 0.01, \*\*\**P* < 0.001, \*\*\*\**P* < 0.0001. Data are mean ± SD.

Following unsupervised graph-based clustering, 14 distinct clusters were resolved in uniform manifold approximation and projection (UMAP), representing keratinocytes (KC), fibroblasts (FB), eccrine gland cells (ECC), vascular endothelial cells (VEC), lymphatic endothelial cells (LEC), perivascular cells/myofibroblasts (PVM), melanocytes (MC), Schwann cells (SC), T cells (T), natural killer cells (NK), B cells (B), mononuclear phagocytes (MP), neutrophils (Neu) and mast cells (Mast) (Figures 1B and 1C).

Compared with HC skin, CD45⁺ hematopoietic cells represented a larger proportion of all cells in each infection group (Figures 1D and 1E). Within the CD45⁺ compartment, all infection groups showed a compositional shift from myeloid toward lymphoid cells (Figure 1F). MPs and NK cells showed the most consistent changes across infection groups, but in opposite directions: MPs, the dominant myeloid population, accounted for a lower proportion of CD45⁺ cells than in HC, whereas NK cells accounted for a higher proportion (Figure 1G).

### Shared and distinct dendritic cell programs across cutaneous mycobacterial infections

To compare MP composition and transcriptional states across cutaneous mycobacterial infections, we reclustered MP and identified 12 transcriptionally distinct subsets, comprising 7 macrophage and 5 dendritic cell (DC) populations. The DC compartment included cDC1s (*CLEC9A, XCR1*), cDC2s (*CD1C, CLEC10A*), LAMP3^+^ DCs (*LAMP3, FSCN1, CCR7*), pDCs (*LILRA4, GZMB*), and Langerhans cells (*CD207, CD1A, FCER1A*) (Figures 2A and 2B)^25,26^.

**Figure 2.**
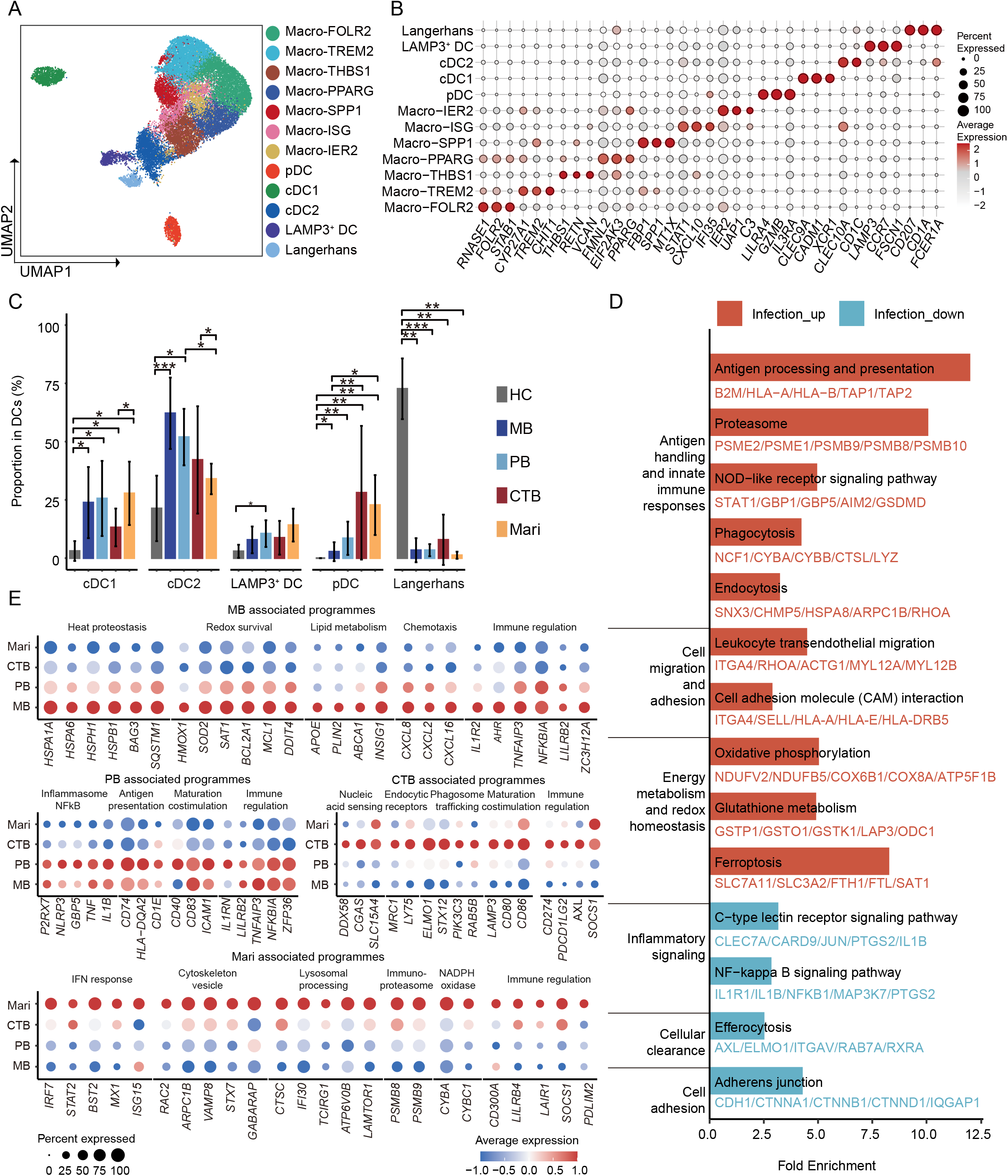
Shared and distinct dendritic cell programs across cutaneous mycobacterial infections. (A) UMAP of skin MP, colored by 7 macrophage and 5 DC subsets. (B) Dotplot of representative macrophage and DC subset markers. (C) Proportions of cDC1, cDC2, LAMP3^+^ DCs, pDCs, and Langerhans cells among total DCs in indicated clinical groups (HC, n = 6; MB, n = 7; PB, n = 8; CTB, n = 8; Mari, n = 4); Wilcoxon rank-sum test; \**P* < 0.05, \*\**P* < 0.01, \*\*\**P* < 0.001, \*\*\*\**P* < 0.0001; data are mean ± SD. (D) Genes upregulated or downregulated in non-Langerhans DCs from pooled infection groups relative to HC were enriched in the indicated KEGG pathways. (E) Dotplot of infection-associated marker genes of total DCs across MB, PB, CTB, and Mari.

Within the DC compartment, the proportions of pDCs, cDC1s, and cDC2s differed across infection groups (Figure 2C). Compared with HC skin, Langerhans cells were consistently less represented across the infection groups, whereas non-Langerhans DCs collectively showed the opposite trend (Figure 2C). Transcriptional changes were then assessed separately in Langerhans cells and non-Langerhans DCs by comparing pooled infection samples (MB, PB, CTB, and Mari) with HC.

Relative to HC, upregulated genes in Langerhans cells and non-Langerhans DCs from infected samples shared enrichment for antigen processing and presentation, phagosomal and endocytic processing, and cellular trafficking and adhesion (*PSME1, TAP1*, *and B2M*; *CYBB* and *SNX3; ITGA4*, respectively) (Figures 2D and S1A). In non-Langerhans DCs, upregulated genes were additionally enriched for innate immune signaling (*STAT1, GBP1, GBP5, AIM2,* and *GSDMD*), oxidative phosphorylation (*NDUFV2, COX6B1,* and *ATP5F1B*), and redox homeostasis (*GSTP1, GSTO1, SLC7A11, FTH1,* and *FTL*). Enrichment among downregulated genes was detected only in non-Langerhans DCs and involved C-type lectin receptor and NF-κB signaling (*CLEC7A* and *CARD9*; *IL1B*, *PTGS2*, and *NFKB1*, respectively), as well as cellular clearance and cell adhesion (*AXL, ELMO1,* and *RAB7A*; *CDH1, CTNNA1,* and *CTNNB1*, respectively) (Figures 2D and S1A), suggesting broader infection-associated transcriptional remodeling in non-Langerhans DCs.

Next, to define distinct infection-associated transcriptional features, we compared DC profiles across MB, PB, CTB, and Mari at both the overall and subset levels. Infection associated features were broadly shared across DC subsets in MB and Mari but were largely restricted to particular subsets in PB and CTB (Figures 2E and S1B-S1F). MB showed stress-response and cell-survival features across all DC subsets, with additional lipid-processing and chemotaxis in cDC2 (*HSPA1A*, *MCL1*; *APOE*, *CXCL8*, respectively) (Figures 2E and S1C). In PB, cDC2 and pDCs shared NF-κB- and MHC class II-related features (*NFKBIA*; *CD74* and *HLA-DQA1/2*, respectively), while cDC2 additionally showed inflammasome-associated and maturation features (*NLRP3* and *IL1B*; *CD83*, respectively) (Figures S1C and S1E). In CTB, cDC2 showed pathogen sensing, phagocytic processing, and cross-presentation, whereas LAMP3^+^ DCs shared sensing/processing features and additionally showed maturation and costimulation (*CGAS*, *ELMO1* and *WDFY4*; *LAMP3* and *CD80*, respectively) (Figures S1C and S1D). Mari showed broadly shared interferon and vesicular/lysosomal features across multiple DC subsets (*IRF7* and *ISG15*; *VAMP8,* respectively) (Figures 2E and S1B-S1F).

### Transcriptionally distinct macrophage states across cutaneous mycobacterial infections

Within the macrophage compartment, Macro-FOLR2 showed tissue-resident and homeostatic features (*CD163, MERTK, MRC1,* and *SELENOP*)^27^. Macro-TREM2 was characterized by lipid metabolism and lysosomal lipid-processing features (*APOE, APOC1, LIPA,* and *GPNMB*)^28^. Macro-THBS1 showed inflammatory-response features (*IL1B, PTGS2, NLRP3,* and *CCL20*), and Macro-PPARG showed features associated with metabolic regulation and stress responses (*PPARG, FOXO1, EIF2AK3,* and *OXSR1*). Macro-SPP1 was characterized by glycolysis-, metal homeostasis-, and redox-associated expression (*HK2, LDHA, SLC2A1, MT1G,* and *PRDX2*). Macro-ISG displayed a prominent interferon-response signature (*CXCL9, CXCL10, GBP1,* and *IFIT3*), and Macro-IER2 showed immediate-early and stress-response features (*IER2, JUN, DDIT4,* and *GADD45B*) (Figure 3A).

**Figure 3.**
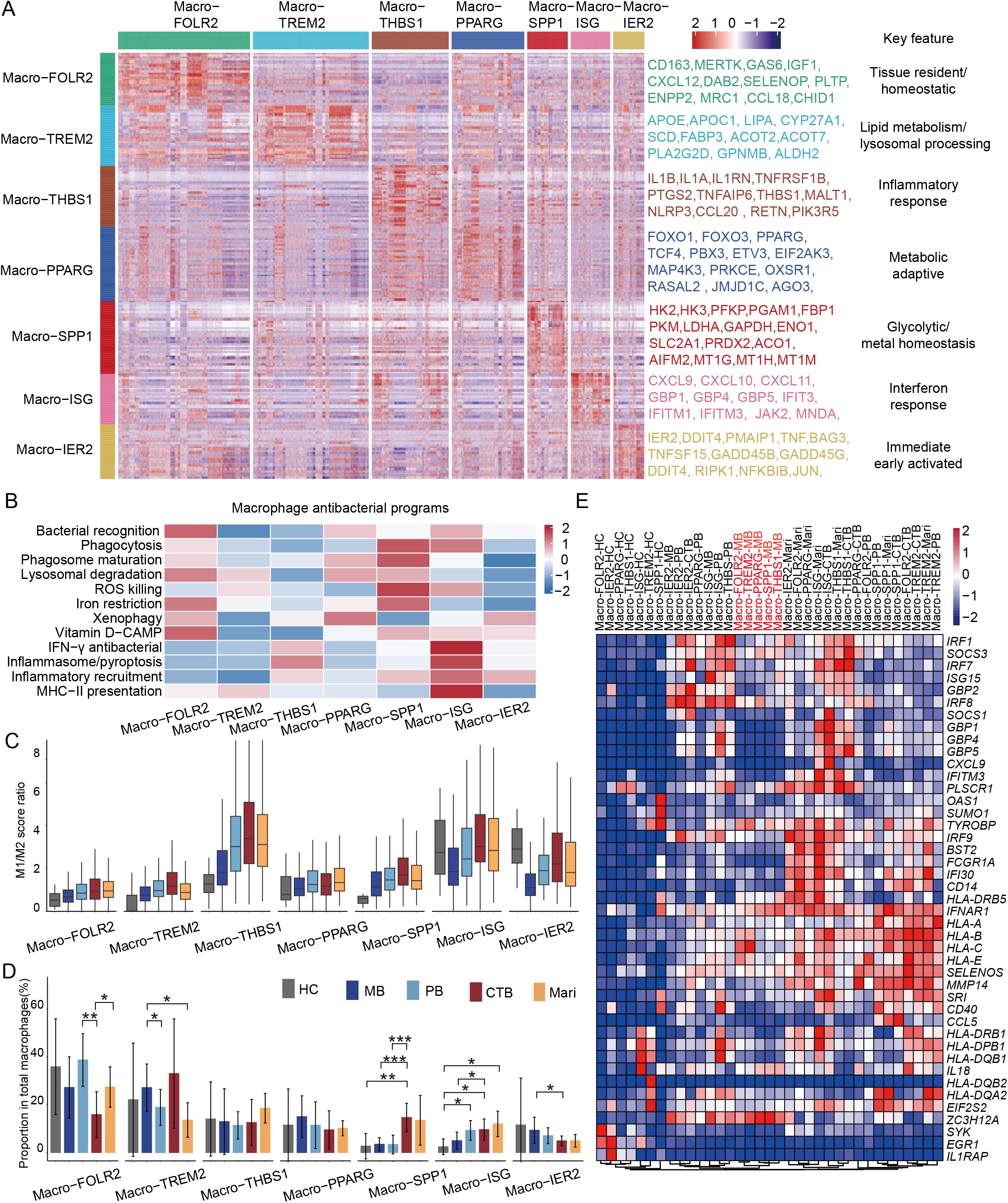
Transcriptionally distinct macrophage states across cutaneous mycobacterial infections. (A) Heatmap of signature gene expression across macrophage subsets. (B) Heatmap of antibacterial signature scores calculated using AUCell. (C) Single-cell M1/M2 AUCell score ratios across macrophage subsets and infection groups. (D) Proportions of macrophage subsets within total macrophages. (HC, n = 6; MB, n = 7; PB, n = 8; CTB, n = 8; Mari, n = 4). Wilcoxon rank-sum test; \**P* < 0.05, \*\**P* < 0.01, \*\*\**P* < 0.001; data are mean ± SD. (E) Heatmap of interferon- and inflammation-related gene expression across infection groups and macrophage subsets. Genes were selected based on differential expression between each infection group and HC.

As macrophages are major effector cells in mycobacterial control and their antimicrobial capacity may shape clinical manifestations^29,30^, we compared antimicrobial programs across macrophage subsets. Gene-set scoring revealed three major configurations: a sensing-and-restriction profile in Macro-FOLR2, involving bacterial recognition, lysosomal degradation, iron restriction, and vitamin D–CAMP; an uptake-and-killing profile in Macro-SPP1, coupling phagocytosis and phagosome maturation with ROS-mediated killing; and inflammatory profiles in Macro-THBS1 and Macro-ISG. Notably, Macro-ISG combined inflammasome/pyroptosis with IFN-γ responses, inflammatory recruitment, and MHC class II antigen presentation (Figure 3B). Analysis of M1 and M2 signatures showed an M1 bias in Macro-THBS1 and Macro-ISG (Figures S2A and S2B). M1-to-M2 score ratios were generally lower in HC and MB-derived macrophages than in those from PB, CTB, and Mari (Figures 3C and S2C).

The relative abundance of these macrophage states was compared across HC and the infection groups. CTB showed a lower proportion of Macro-FOLR2 than in PB and Mari and a higher proportion of Macro-SPP1 than in HC, MB, and PB. MB showed a higher proportion of Macro-TREM2 than in PB and Mari. The increase in Macro-ISG relative to HC observed in the other infections was not evident in MB (Figure 3D).

We next examined whether reduced M1 bias and the lack of increased Macro-ISG representation in MB were accompanied by lower IFN-associated gene expression across macrophage subsets. Hierarchical clustering revealed that five MB-derived subsets—Macro-FOLR2, Macro-TREM2, Macro-PPARG, Macro-SPP1, and Macro-THBS1—clustered together rather than with their corresponding subsets from other groups, suggesting that a shared MB-associated program partially overrode macrophage subset identity. A shared feature of these five MB-derived subsets was lower expression of *GBP1*, *GBP4*, *GBP5*, *CXCL9*, *ISG15*, and *FCGR1A*, indicating broad attenuation of IFN-associated effector expression (Figure 3E).

### Shared and distinct macrophage programs across cutaneous mycobacterial infections

To define infection-associated transcriptional features of macrophage subsets, we compared pathway activities across infection groups using gene set variation analysis (GSVA). Hierarchical clustering separated MB-derived Macro-TREM2 and Macro-SPP1 from their counterparts in PB, CTB, and Mari (Figure 4A). Gene-level analysis identified shared features that distinguished MB-derived Macro-TREM2 and Macro-SPP1 from their counterparts in other infections, including higher expression of TNF–NF-κB-associated genes and lower expression of TCA cycle, gluconeogenesis and cellular maintenance associated genes. Beyond these shared features, Macro-TREM2 in MB showed increased expression of hypoxia-, apoptosis-, and IL2–STAT5-associated genes, whereas Macro-SPP1 in MB showed increased expression of mTOR pathway-associated genes and predominantly reduced expression of redox-associated genes (Figures 4B and S3). Additionally, macrophages from MB and PB showed narrower antimicrobial gene-set score profiles, with higher scores largely confined to xenophagy, whereas those from CTB and Mari displayed broader profiles spanning pathogen recognition, intracellular bacterial processing, and effector killing (Figure 4C).

**Figure 4.**
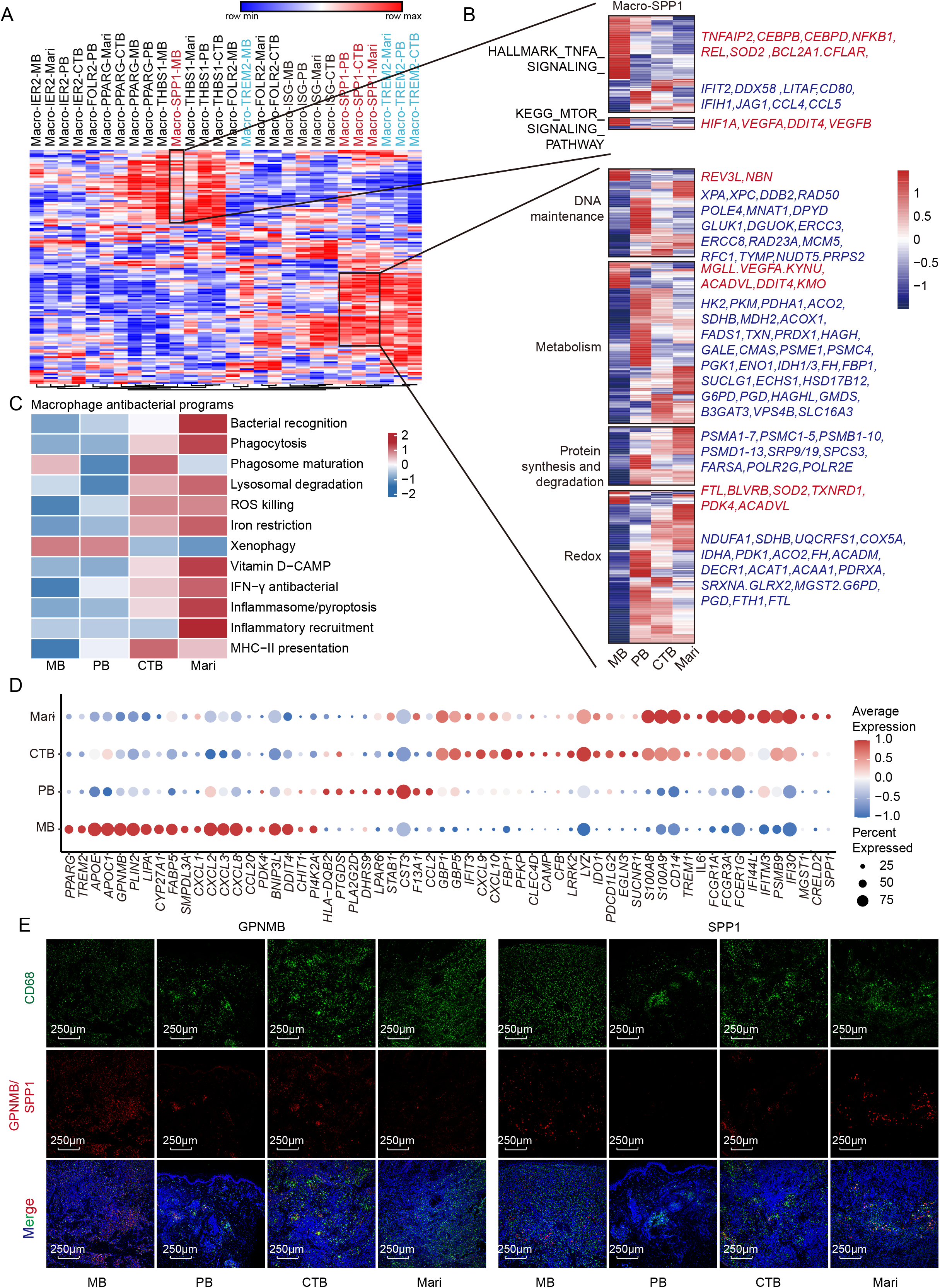
Shared and distinct macrophage programs across cutaneous mycobacterial infections. (A) Heatmap of GSVA scores for selected Hallmark and KEGG pathways across macrophage subsets and infection groups. (B) Heatmaps of selected pathway associated genes in Macro-SPP1 across infections, grouped by annotated function. (C) Heatmap of antibacterial AUCell scores across infections. (D) Dotplots of infection-associated marker genes across the total macrophages in MB, PB, CTB, and Mari. (E) Immunofluorescence of CD68 (green), GPNMB and SPP1 (red) in MB, PB, CTB and Mari skin biopsies. Images were acquired using 10x.

We further examined gene expression differences across infections at both the overall macrophage and subset levels. In MB, macrophage subsets shared features of lipid transport, storage, lysosomal adaptation (*APOE*; *PLIN2*; *GPNMB*, respectively), and chemotaxis (*CXCL2, CXCL3, CXCL8*), whereas several subsets showed lower expression of genes associated with Toll-like receptor signaling, IFN responses, and antigen processing and presentation (*TLR8*; *GBP1* and *CXCL10*; *CD74* and *IFI30*, respectively) than their counterparts from other infections. In PB, infection-associated features were more subset-specific, including lipid mediator metabolism in Macro-FOLR2 and Macro-TREM2 (*PTGDS, PLA2G2D*) and cholesterol biosynthesis and fatty acid remodeling in Macro-SPP1 (*SREBF2* and *SQLE*; *FADS1* and *ELOVL5*, respectively). CTB showed higher expression of fructose-1,6-bisphosphatase 1 (*FBP1*), a key enzyme in gluconeogenesis, across multiple macrophage subsets, alongside vitamin D-associated antimicrobial genes in Macro-TREM2 and Macro-SPP1 (*CAMP, CYP27B1, VDR*) and metal-homeostasis and stress-response genes in Macro-SPP1, Macro-PPARG, and Macro-ISG (*MT1E, MT1G, MT1X*). Mari showed tissue-remodeling gene expression in Macro-TREM2 and Macro-SPP1 (*SPP1, MMP12*) (Figures 4D and S4A-S4G).

Consistent with the scRNA-seq findings, immunofluorescence showed that GPNMB^+^ and SPP1^+^ cells were most abundant in MB and Mari, respectively, whereas FOLR2^+^ cells were more abundant in Mari than in PB and CTB (Figures 4E and S5).

### Reduced FBP1-associated metabolism is linked to impaired LGC formation in MB

Hematoxylin and eosin (H&E) staining revealed a distinct histological contrast between MB and the other infections: Langhans giant cells (LGCs) were present in PB, CTB, and Mari lesions but nearly absent from the examined MB sections, which predominantly contained mononuclear macrophages (Figures 5A-5D). We asked whether the absence of LGCs in MB could be explained by the macrophage differences identified above, particularly weaker Toll-like receptor (TLR) and IFN-associated features and altered expression of glucose metabolism-related genes in MB (Figures 4B, 4D, S4, and S6). We therefore tested innate immune stimulation and metabolic modulation during LGC induction.

**Figure 5.**
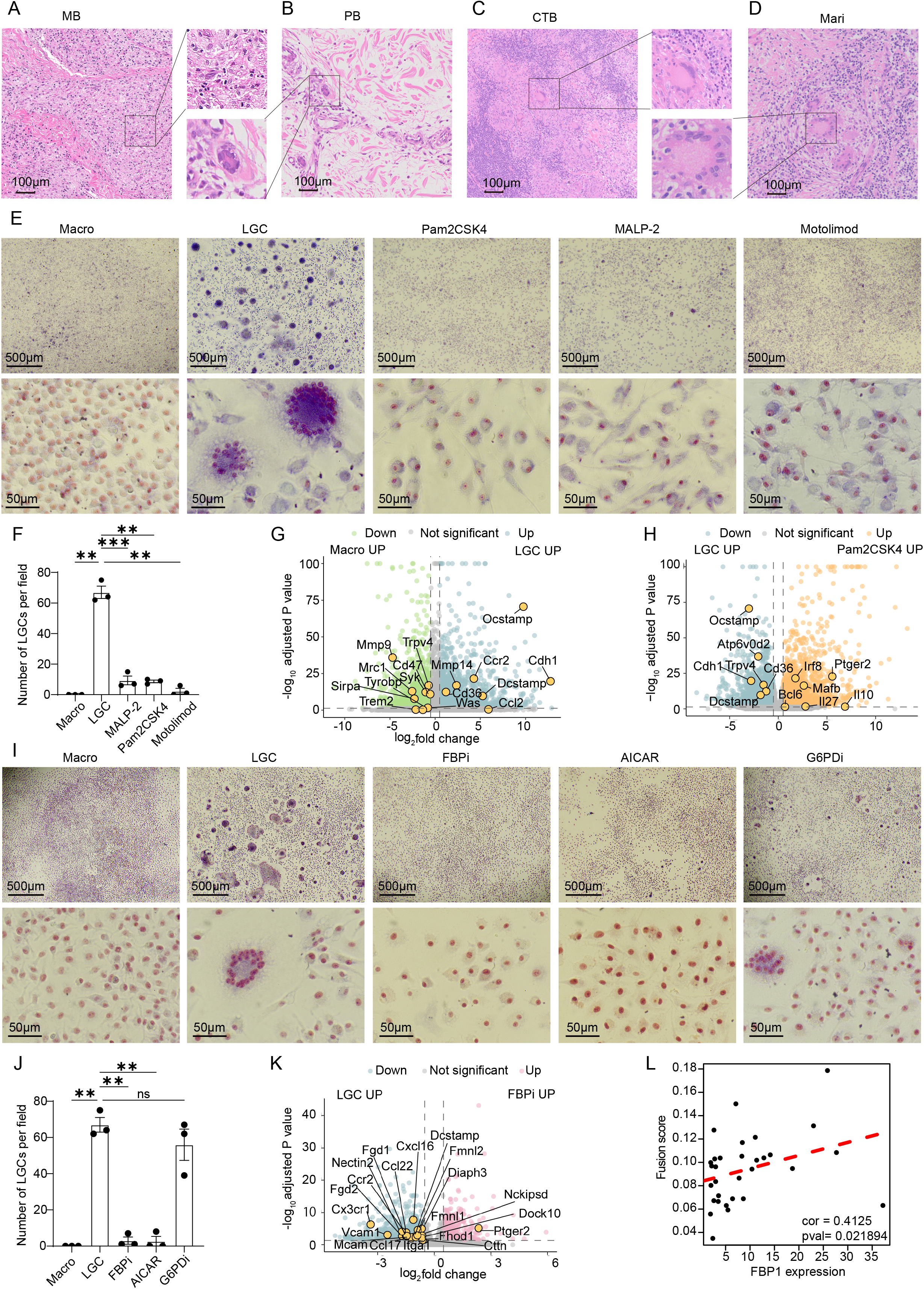
Reduced FBP1 associated metabolism is linked to impaired LGC formation in MB. (A–D) Histological images of lesional skin biopsies of MB (A), PB (B), CTB (C), and Mari (D). (E) Representative images of untreated macrophages (Macro), untreated Langhans giant cell (LGC), and LGC-inducing cultures treated with 60 ng/ml Pam2CSK4, 100 ng/ml MALP-2, or 800 nM motolimod. Images were acquired using 4x (upper panels) and 40x (lower panels) objectives. (F) Numbers of LGCs per field acquired using a 4x objective in (E) (n=3). (G, H) Volcano plots of differential gene expression between macrophages and LGCs (G) and between cells from untreated and Pam2CSK4-treated LGC-inducing cultures (H) (n=4, adjusted P < 0.05, |log₂FC| ≥ 0.5). (I) Representative images of untreated macrophage, untreated LGCs and LGC inducing cultures treated with a 100 μM FBP1 inhibitor (FBPi), 500 μM AICAR, or 50 μM G6PD inhibitor (G6PDi). Images were acquired using 4x (upper panels) and 40x (lower panels) objectives. (J) Numbers of LGCs per field acquired using a 4x objective in (I) (n=3). (K) Volcano plots of differential gene expression between cells from untreated and FBPi-treated LGC inducing cultures (n=4, adjusted *P* < 0.05, |log₂FC| ≥ 0.5). (L) Spearman correlation between donor-average FBP1 expression and fusion-associated AUCell scores in FBP1-expressing macrophages. Each dot represents one donor. For (F, J), paired t-test, ns, not significant, \**P* < 0.05, \*\**P* < 0.01, \*\*\**P* < 0.001.

Counterintuitively, treating mouse bone marrow-derived macrophages (BMDMs) with TLR2/6 agonists (Pam2CSK4 or MALP-2) or a TLR8 agonist (motolimod) at the onset of induction suppressed LGC formation (Figures 5E and 5F). RNA-seq analysis showed increased expression of fusion-, adhesion-, and chemotaxis-related genes during LGC induction (*Dcstamp*, *Ocstamp*; *Cdh1*; *Ccr2*, respectively), whereas Pam2CSK4 reduced fusion-associated transcripts (*Dcstamp, Ocstamp, Cdh1, Trpv4, Atp6v0d2*) (Figures 5G and 5H). These findings argue against reduced TLR signaling alone explaining the absence of LGCs in MB.

We next examined whether perturbing MB-associated metabolic features affected LGC formation. FBP1 inhibition, targeting a key gluconeogenic enzyme whose reduced expression distinguished MB from the other infections (Figures 4B and 4D), nearly abolished LGC formation. AICAR, an AMP mimetic and AMPK activator, produced a similar effect, whereas G6PD inhibition, despite its reduced expression in MB macrophages, did not significantly reduce cell fusion, although the resulting multinucleated cells showed altered morphology (Figures 5I and 5J). RNA-seq further showed that FBP1 inhibition reduced the expression of genes related to fusion, adhesion, chemotaxis, and cytoskeletal regulation, including *Dcstamp*, *Nectin2, Vcam1, Ccr2, Ccl22, Cxcl16, Fgd2,* and *Fmnl2* (Figure 5K), and this finding was further supported by a positive correlation between FBP1 expression and the macrophage fusion score in our scRNA-seq data (Figure 5L). Collectively, these results suggest that reduced FBP1 expression may constrain macrophage fusion and contribute to the absence of LGCs in MB.

### Distinct NK cell states across cutaneous mycobacterial infections

Within the NK cell compartment, TCF7 NK cells displayed transcriptional features of a precursor/memory-like state, with high expression of *TCF7, BACH2, SOX4*, and *IL7R*; GZMK NK cells displayed a transitional activation state, expressing *GZMK* and *EOMES* together with the immediate-early genes *EGR1, FOS,* and *JUNB*; FCGR3A NK cells displayed a mature cytotoxic state, characterized by high expression of *FCGR3A, FGFBP2, PRF1, GZMB,* and *NKG7*; TNFRSF9 NK cells displayed sustained activation accompanied by regulatory features, coexpressing activation-associated genes (*TNFRSF9, TRAF1,* and *NFKB1*) and immunoregulatory genes (*TGFBR2, SMAD3,* and *CBLB*) (Figures 6A-6C). Both MB and PB showed higher proportions of FCGR3A NK cells than CTB, and MB also showed a lower proportion of TCF7 NK cells than CTB and Mari (Figure 6D).

**Figure 6.**
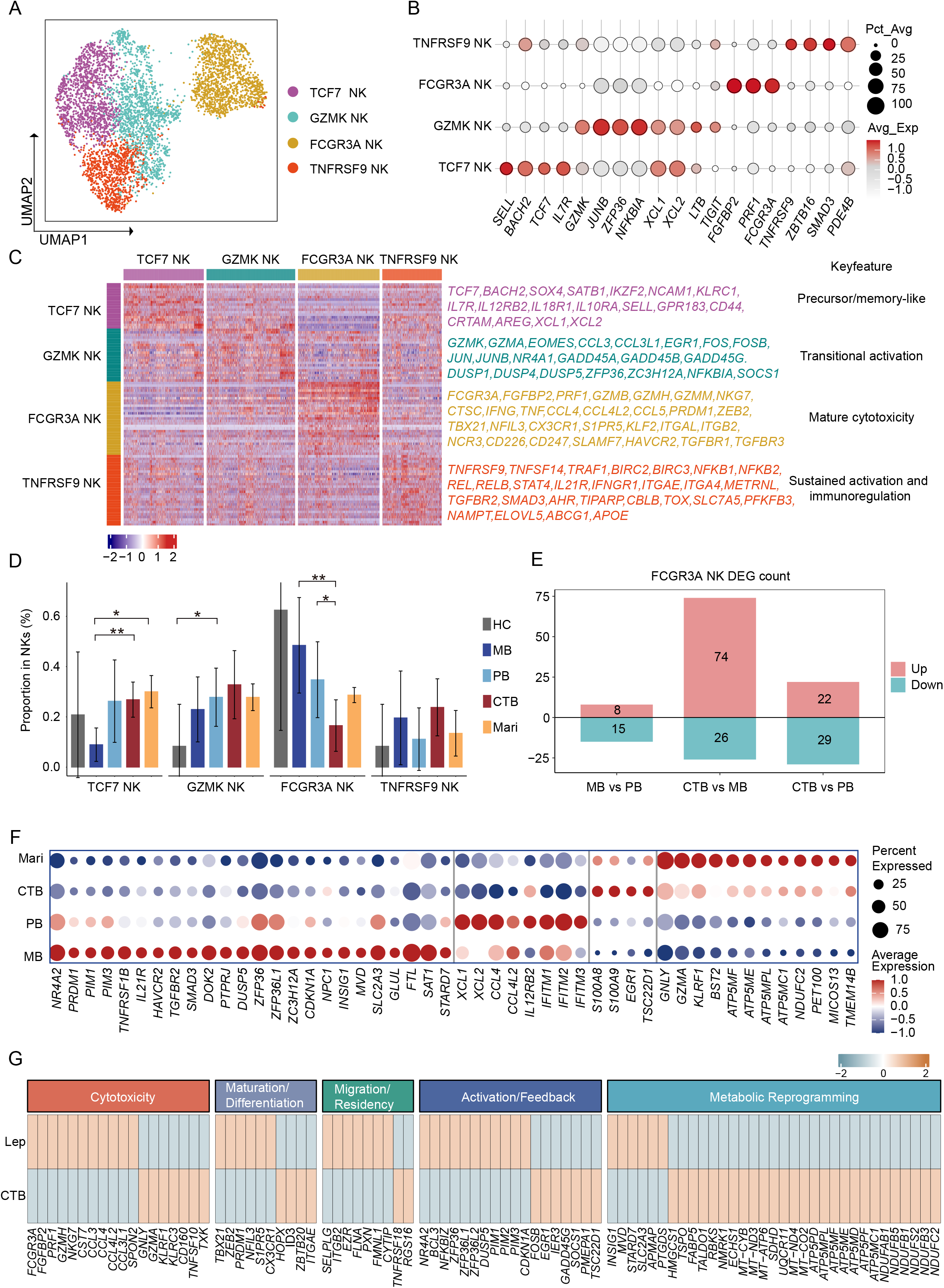
Distinct NK cell states across cutaneous mycobacterial infections. (A) UMAP of skin NK cells, colored by 4 annotated subsets. (B) Dotplot of representative NK cell subset markers. (C) Heatmap of signature genes across NK subsets. (D) Proportions of NK subsets within total NK cells. Wilcoxon rank-sum test; Data are mean ± SD. \**P* < 0.05, \*\**P* < 0.01; (E) Numbers of upregulated and downregulated genes in FCGR3A NK cells in indicated comparisons. (F) Dotplot of infection-associated marker genes. (G) Heatmap of featured DEGs between leprosy and CTB-derived FCGR3A NK cells.

In MB, NK cell subsets commonly showed immunoregulatory features (*RUNX3*, *HAVCR2*, *TGFBR2*, *PIM3,* and *FKBP5*). In PB, TCF7, GZMK, and TNFRSF9 NK cells shared cytokine- and interferon-responsive features (*IL12RB2*, *STAT4*, and *IL21R*; *IFITM1*, *IFITM2*, *IFITM3*, and *XAF1*, respectively). CTB and Mari showed similar cytotoxic-effector (*GNLY*, *GZMA* and *KLRF1*) and oxidative metabolism-associated features (*NDUFB2*, *NDUFS6*, and *ATP5PF*) in TCF7, GZMK, and FCGR3A NK cells, generally more pronounced in Mari. TNFRSF9 NK cells from both infections also shared activation- and cytotoxicity-associated features (*FCER1G*, *JUNB*, and *HOPX*; *CTSW* and *GNLY*, respectively) (Figures 6F and S7A-S7D).

The higher proportions of FCGR3A NK cells in both MB and PB than in CTB prompted a focused comparison of their transcriptional profiles. As only 23 genes were differentially expressed between MB and PB within this subset, cells from both groups were pooled to characterize leprosy-associated transcriptional features relative to CTB (Figure 6E). Compared with CTB, leprosy-derived FCGR3A NK cells expressed higher levels of maturation- and cytotoxicity-associated genes *(TBX21* and *PRDM1*; *FGFBP2, PRF1, and GZMH*, respectively), but lower levels of *GZMA* and the antibacterial effector *GNLY*^31^ (Figure 6G). These cells also showed higher expression of genes associated with sterol homeostasis (*INSIG1* and *MVD*), phospholipid transport (*STARD7*), and glucose uptake (*SLC2A3*). CTB-derived cells showed higher expression of *HMGCS1*, involved in mevalonate biosynthesis, *FABP5*, involved in fatty acid transport, and genes encoding respiratory chain and ATP synthase components (Figure 6G).

### Metabolic regulation of NK cell IFN-γ responses

Given the infection-associated differences in metabolism-related gene expression across NK-cell subsets, metabolic pathway scores in the overall NK-cell compartment were compared among infection groups. Mari showed higher scores for central carbon and oxidative energy metabolism (glycolysis/gluconeogenesis, pyruvate metabolism, the tricarboxylic acid cycle (TCA cycle), and oxidative phosphorylation), whereas MB showed higher scores for lipid biosynthesis and remodeling (fatty acid elongation, steroid biosynthesis, and ether lipid metabolism) (Figure 7A).

**Figure 7.**
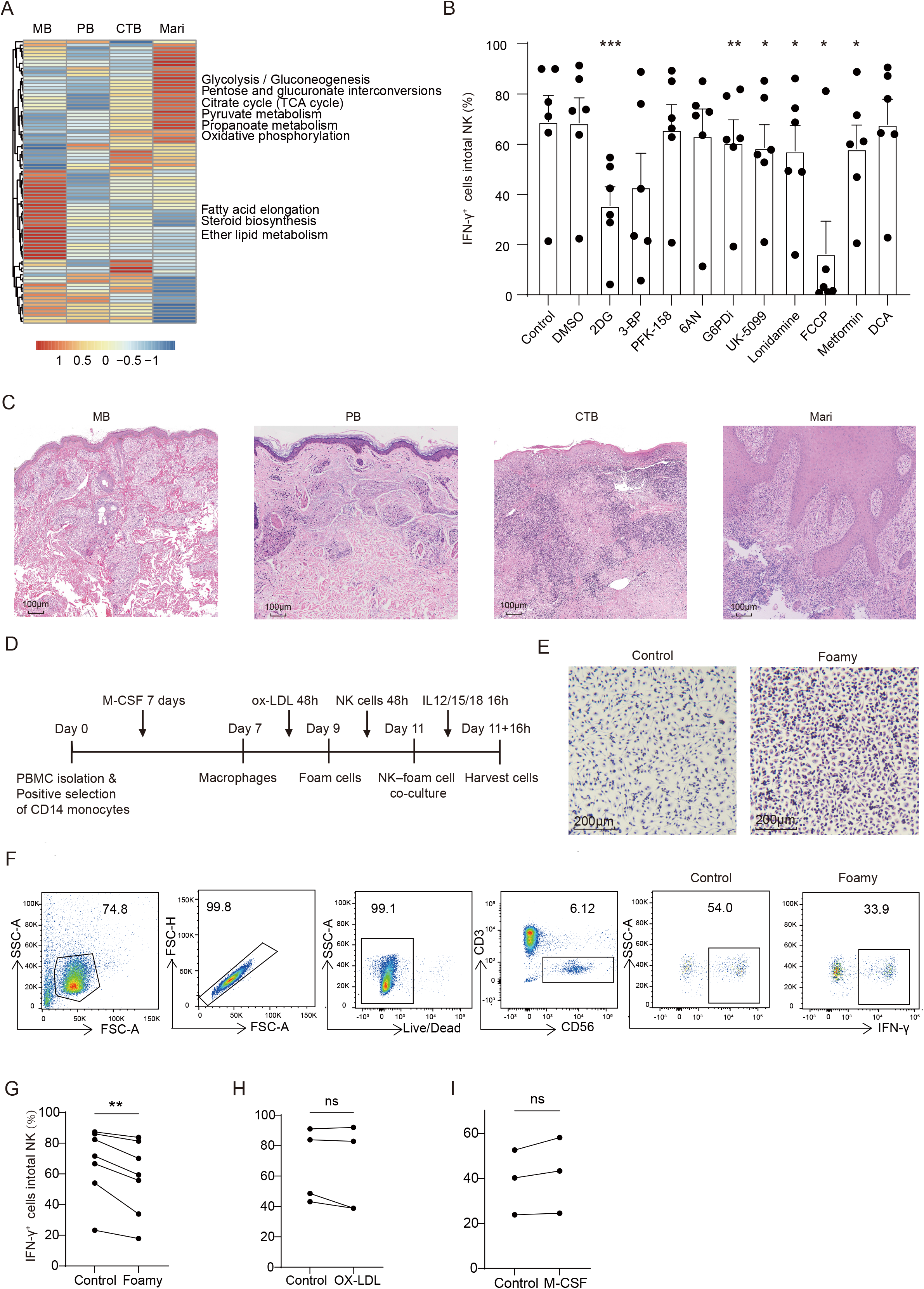
Metabolic regulation of NK cell IFN-γ responses. (A) Heatmap of KEGG metabolic signature scores in NK cells across infections. (B) Percentages of IFN-γ⁺ NK cells following treatment with the indicated metabolic modulators (n = 6). (C) Representative H&E images of lesional skin from MB, PB, CTB and Mari. (D) Experimental schematic of Macrophage-NK cell co-culture workflow. CD14^+^ monocytes were differentiated with 50 ng/ml M-CSF (7 days), treated with 50 μg/ml ox-LDL (48 hrs), and co-cultured with NK cells (48 hrs), followed by 10 ng/ml IL-12, 50 ng/ml IL-15, 50 ng/ml IL-18 stimulation (16 hrs). Culture and treatment conditions are described in Methods. (E) Representative images of control and foamy macrophages. (F) Flow cytometry of IFN-γ detection in NK cells co-cultured with normal or foamy macrophages. (G-I) Percentages of IFN-γ^+^ NK cells following co-culture with foamy macrophages (G, n = 6), or direct 50 μg/ml ox-LDL treatment (H, n = 4), or 50 ng/ml M-CSF treatment (I, n = 3). For (B, G-I), paired t-test; each dot represents one donor. ns, not significant, \**P* < 0.05, \*\**P* < 0.01, \*\*\**P* < 0.001; data are mean ± SEM.

We next tested how metabolic perturbations affect NK cell production of IFN-γ, a cytokine that promotes macrophage antimycobacterial activity^30^. Among compounds targeting glucose metabolism, 2-deoxy-D-glucose (2-DG) significantly reduced the proportion of IFN-γ⁺ NK cells, whereas the effects of 3-bromopyruvate (3-BP) and PFK-158 did not reach statistical significance. Targeting the pentose phosphate pathway with 6-aminonicotinamide (6-AN) and glucose-6-phosphate dehydrogenase inhibitor G6PDi showed no significant or only mild reduction of IFN-γ production by NK cells. These patterns raise the possibility that NK-cell IFN-γ production is differentially sensitive to perturbations at distinct steps of glucose metabolism. Compounds targeting mitochondrial metabolism also significantly reduced NK-cell IFN-γ production, including the pyruvate transport inhibitors UK-5099 and lonidamine, the uncoupler carbonyl cyanide *p*-trifluoromethoxyphenylhydrazone (FCCP), and the respiratory complex I inhibitor metformin. Dichloroacetate (DCA), which promotes pyruvate dehydrogenase activity through kinase inhibition, did not significantly alter NK-cell IFN-γ production (Figure 7B). The convergent effects of mitochondrial interventions support a role for mitochondrial pyruvate import and respiratory function in sustaining NK-cell IFN-γ production.

MB NK cells exhibited lipid-associated transcriptional features, and H&E staining revealed more extensive accumulation of foamy macrophages in MB lesions than in PB, CTB, and Mari lesions (Figure 7C). To examine whether macrophage lipid loading influenced NK cell IFN-γ production, we performed co-culture experiments. NK cells produced significantly less IFN-γ when co-cultured with oxidized low-density lipoprotein (ox-LDL)-induced foamy macrophages than with control macrophages (Figures 7D-7G), whereas direct exposure to ox-LDL or the macrophage differentiation factor M-CSF had no significant effect (Figures 7H and 7I). These findings suggest that foamy macrophage accumulation may contribute to local suppression of NK cell IFN-γ responses in MB lesions.

## DISCUSSION

Cutaneous mycobacterial infections provide a common tissue setting in which related intracellular pathogens elicit distinct pathological responses. By profiling more than 266,000 cells from 33 individuals spanning healthy controls and patients with MB, PB, CTB, and Mari within a unified analytical framework, we identified shared and distinct immune programs across these infections. Two main findings emerged: first, a shared relative contraction of myeloid cells and expansion of NK cells within the CD45⁺ compartment coexisted with distinct antimicrobial and metabolic programs: CTB and Mari showed broader macrophage antimicrobial signatures, whereas MB was characterized by lipid-associated macrophage states and attenuated interferon-associated effector expression; second, macrophage metabolic features were linked to tissue pathology and immune regulation. Reduced FBP1 expression was associated with the absence of Langhans giant cells in MB, and foamy macrophages suppressed NK-cell IFN-γ production in co-culture. Together, these findings point to metabolic reprogramming as a potential link between infection-associated immune states and divergent tissue pathology.

Across infections, the DC compartment showed lower Langerhans cells and higher non-Langerhans DC proportions. Compared with healthy skin, both populations showed enrichment of antigen-processing-related programs, indicating that lower Langerhans cells representation did not imply weaker antigen-processing-related transcription. Although MB and PB share the same causative pathogen, cDC2s from MB showed stress- and lipid-associated programs, whereas those from PB showed inflammatory, MHC class II, and maturation-related features, suggesting that pathogen identity alone does not explain DC heterogeneity. Evidence that oxidized phospholipids impair DC differentiation and antigen presentation^32^ raises the possibility that the local lipid environment helps shape these distinct DC states.

Distinct lipid metabolic programs may have different implications for host defense. MB combined broad lipid-storage programs with relatively weak IFN-associated effector expression, and previous studies showed that lipid-droplet accumulation supports *M. leprae* survival in Schwann cells^33,34^. By contrast, PB showed subset-restricted PTGDS expression, potentially relevant to host defense given that PTGDS-derived PGD₂ promotes bacterial clearance in other experimental settings^35,36^. Lipid-associated signaling may also help explain attenuated IFN responses in MB. For example, LXR-induced SMPDL3A restricts STING signaling by degrading 2′3′-cGAMP produced by cGAS^37^, which also promotes autophagic targeting of *M. tuberculosis*^38^. Whether these mechanisms contribute to persistent infection in MB warrants further investigation.

Our fusion experiments provide a more direct connection between macrophage metabolism and tissue organization. TLR stimulation at the onset of LGC induction suppressed fusion, suggesting that stronger innate stimulation does not necessarily favor giant-cell formation. Reduced FBP1 expression in MB and its positive association with the fusion score, together with the suppression of fusion by pharmacological FBP1 inhibition, support a potential contribution of FBP1-associated metabolism to the absence of LGCs in MB. The consequences may extend beyond multinucleation, given reported roles of LGCs in macrophage recruitment, antigen presentation and bacterial control^39–41^. Genetic perturbation and metabolic tracing in human macrophages will help clarify how FBP1 supports fusion and whether restoring FBP1 expression or activity can rescue impaired LGC formation.

Leprosy-derived FCGR3A NK cells combined stronger cytotoxic transcriptional signatures with lower GNLY expression. Given the role of granulysin in bacterial membrane disruption and granzyme delivery^31,42^, this pattern suggests that cytotoxic differentiation may coexist with limitations in direct antibacterial capacity. NK cell activity is also sensitive to metabolic regulation^43–45^. Our pharmacological experiments support roles for glucose utilization and mitochondrial function in sustaining NK cell IFN-γ production. Foamy macrophages also suppressed NK cell IFN-γ production, suggesting that macrophage lipid loading can influence neighboring immune cells. Murine evidence linking macrophage-derived lipid transfer to reduced mTORC1 activity and NK cell IFN-γ production^46^ raises the broader question of how lipid exchange between immune cells shapes local antimicrobial immunity.

### Limitations

This study has several limitations. The modest cohort size may limit the generalizability of some comparisons. Cross-sectional transcriptomic analyses reveal associations, while pathway signatures do not directly measure metabolic flux or antimicrobial activity. Our functional experiments support metabolic regulation of macrophage fusion and NK cell IFN-γ production, but do not fully reproduce the complexity of human lesions. The mechanisms connecting these experimental effects to disease-associated tissue states therefore remain an area for further investigation.

## RESOURCE AVAILABILITY

### Lead contact

Further information and requests for resources should be directed to the lead contact, Yetao Wang.

### Materials availability

This study did not generate new unique reagents.

### Data availability

The scRNA-seq data generated in this study, together with the bulk RNA-seq data presented in Figure 5, have been deposited in the Genome Sequence Archive for Humans (GSA-Human) and the Genome Sequence Archive (GSA) under accession numbers HRA021486, CRA051248 and CRA051249.

## Acknowledgments

This study was supported by National Natural Science Foundation of China (82471876), CAMS Innovation Fund for Medical Sciences (CIFMS) (2024-I2M-3-005) and Hospital for Skin Diseases, Institute of Dermatology, Chinese Academy of Medical Sciences and Peking Union Medical College grant (3301030103119) to Yetao Wang. The authors used BioRender and BioGDP^47^ to create graphical abstract and schematic illustrations.

## Author contributions

Yetao Wang: conceptualization; experiment design; data analysis; resources; supervision; validation; investigation; visualization; methodology; data curation; software; writing – original draft, review and editing; project administration; funding acquisition. Huangtao Guo: experiment design; performing experiments; data analysis; validation; visualization; methodology; data curation; writing – original draft, review and editing. Rui Zeng: performing experiments; data analysis; validation; visualization; methodology. Hongsheng Wang: resources; writing – review and editing. Qin Wei, Jingshu Xiong, Haiqin Jiang, Ying Shi, Wenyue Zhang, Chun Pan, Chaojiang Cheng: resource.

## FIGURE TITLES AND LEGENDS

**Figure S1.**
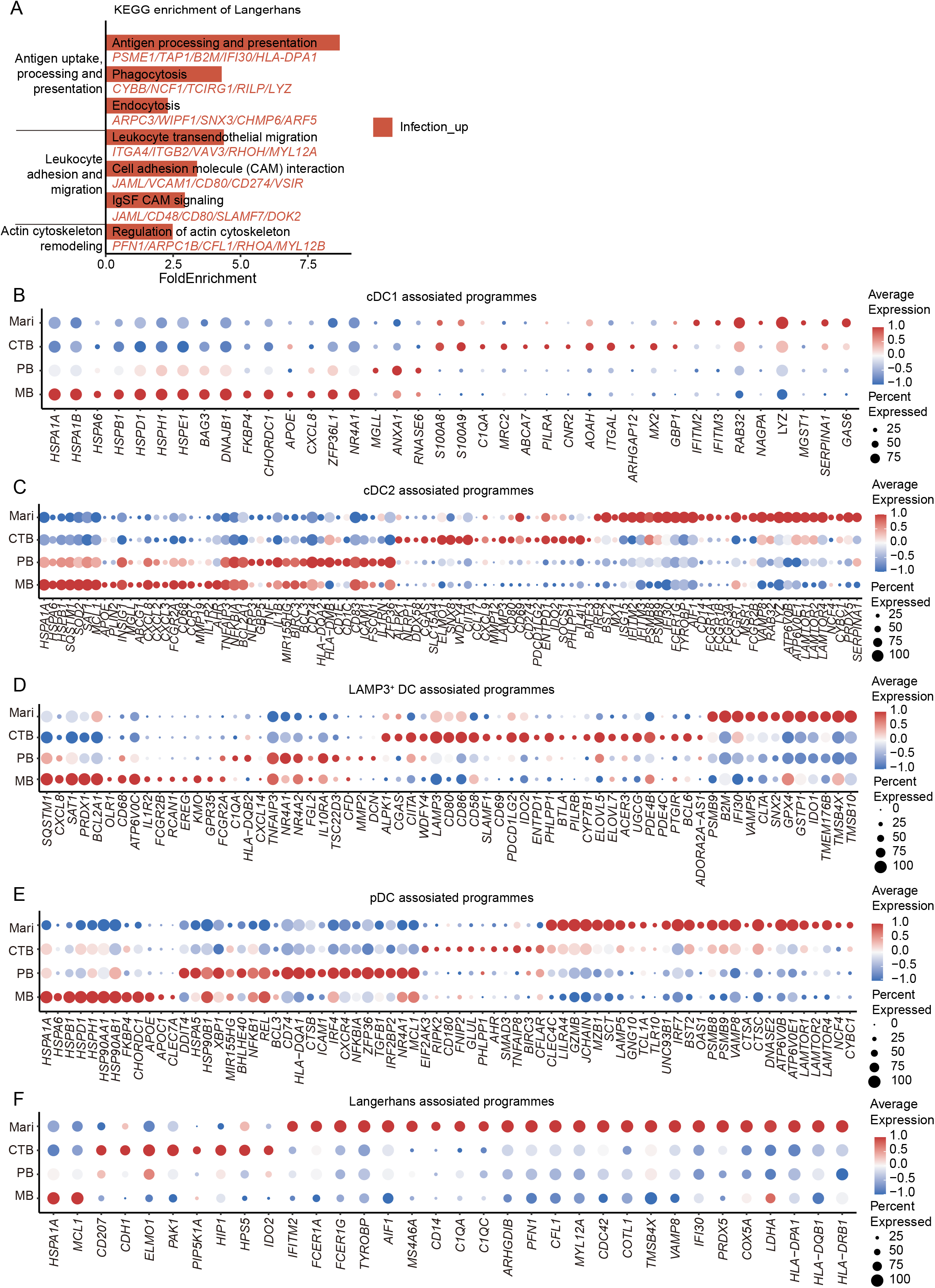
Infection-associated transcriptional programs in DCs. (A) Genes upregulated or downregulated in Langerhans cells from pooled infection groups relative to HC were enriched in the indicated KEGG pathways. (B-F) Dotplots of infection-associated marker genes in cDC1 (B), cDC2 (C), LAMP3^+^ DCs (D), pDCs (E), and Langerhans cells (F) across MB, PB, CTB, and Mari.

**Figure S2.**
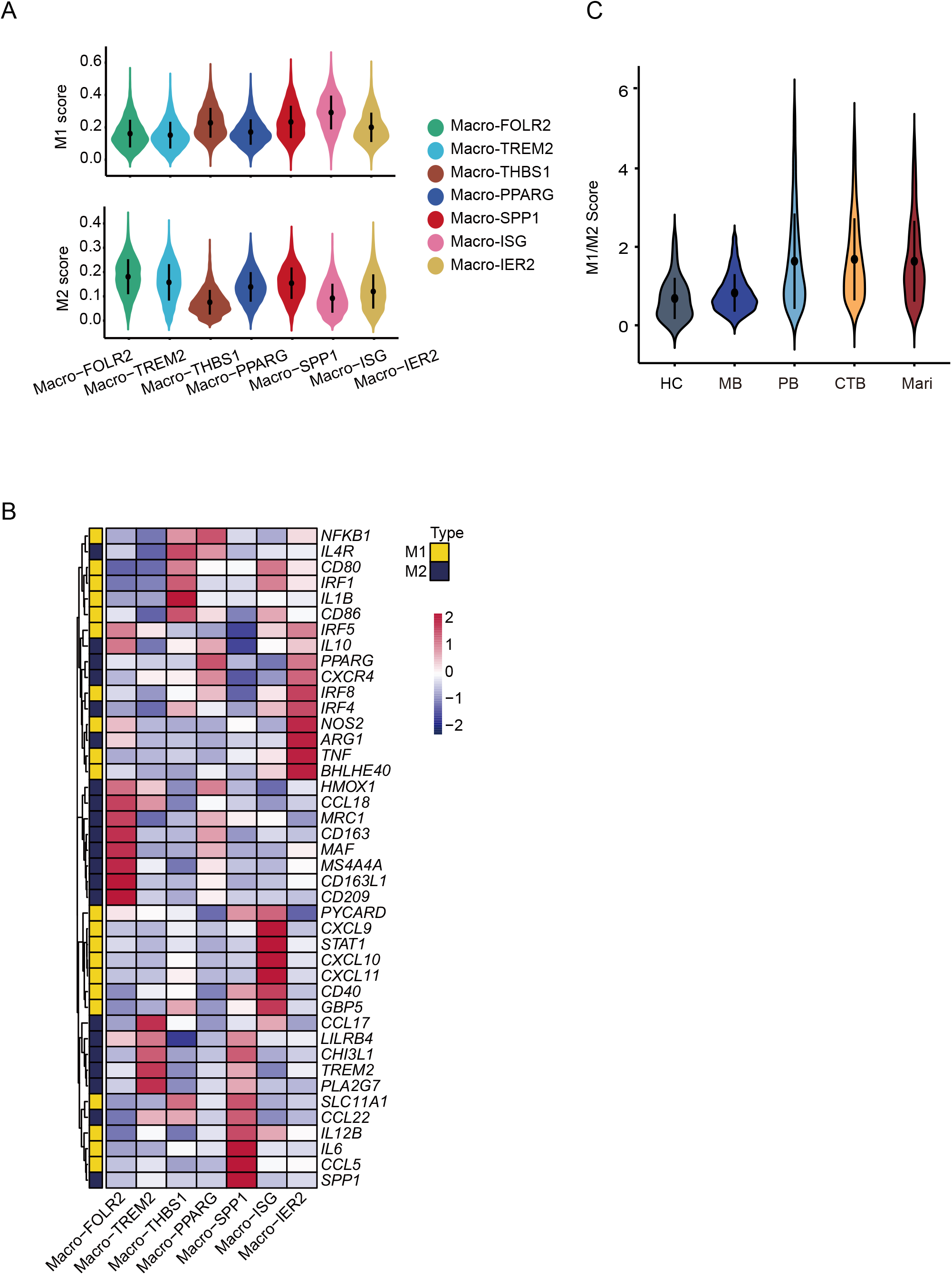
M1- and M2-associated transcriptional signatures in skin macrophages. (A) Violin plots of M1 and M2 AUCell scores across macrophage subsets. (B) Heatmap of M1- and M2-associated genes across macrophage subsets. (C) Violin plots of M1/M2 AUCell score ratios across HC and infection groups.

**Figure S3.**
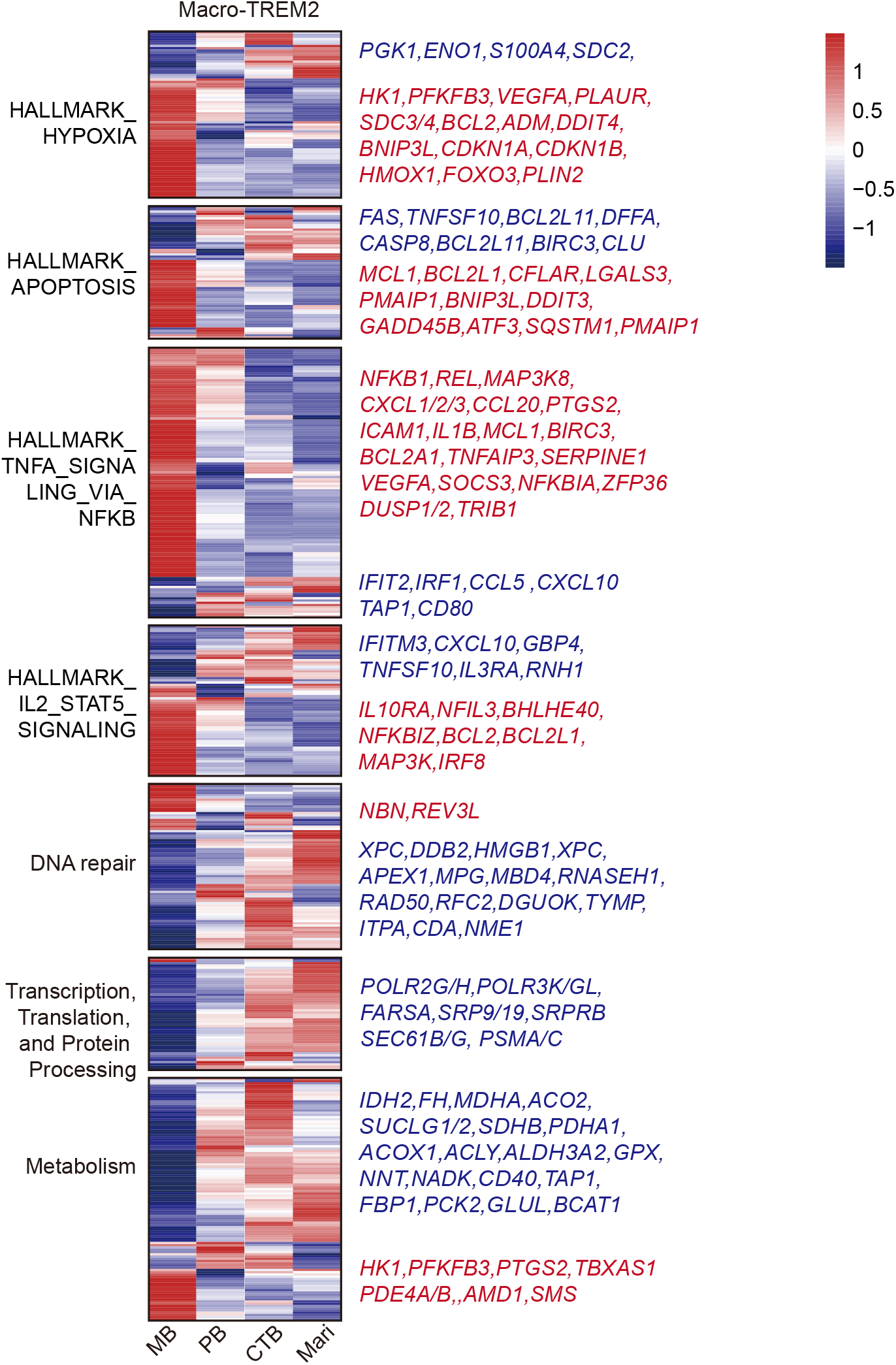
Infection-associated programs in Macro-TREM2. Heatmaps of selected pathway associated genes in Macro-TREM2 across infections, grouped by annotated function.

**Figure S4.**
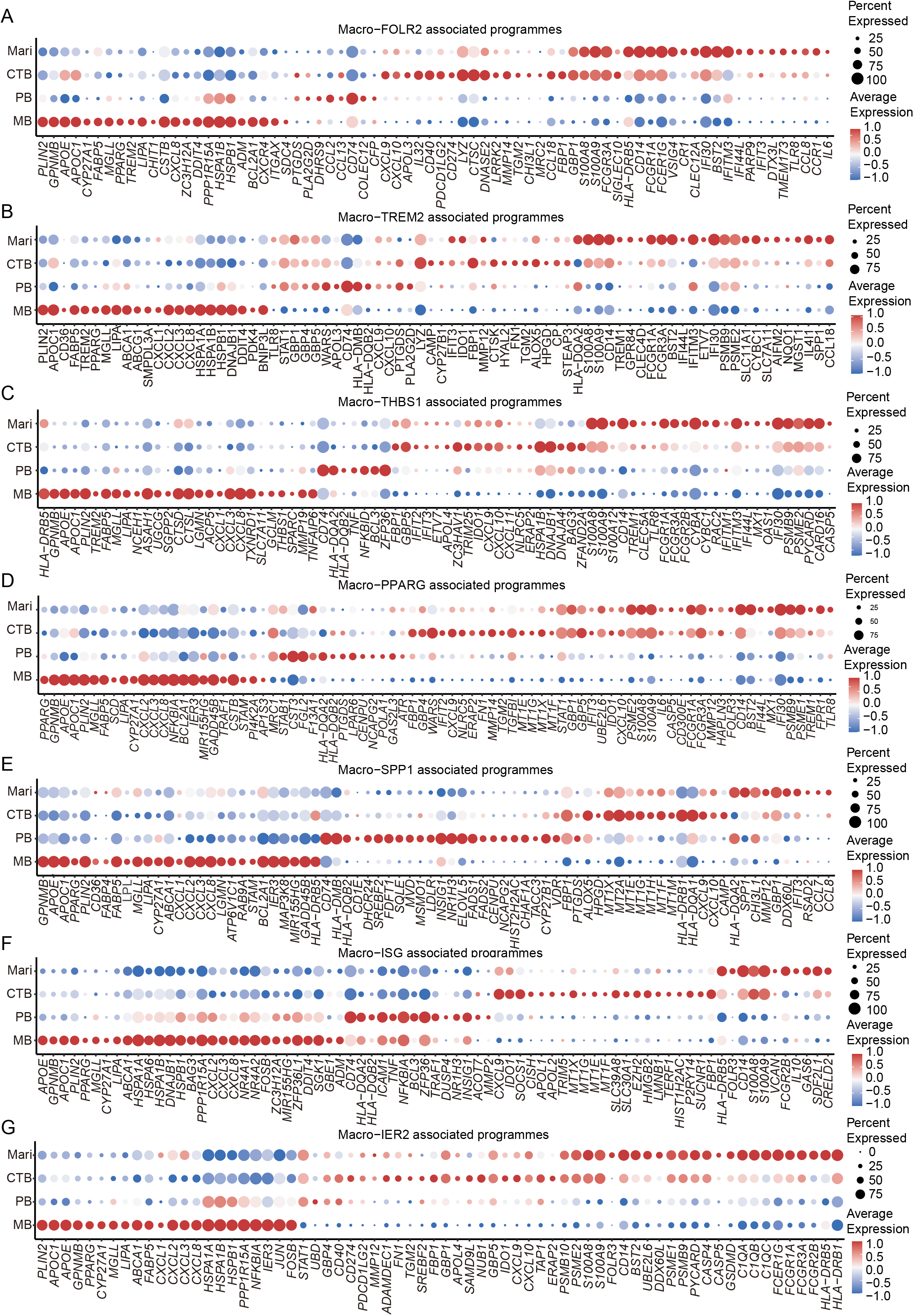
Infection-associated transcriptional programs within macrophage subsets. (A-G) Dotplots of infection-associated marker genes in Macro-FOLR2 (A), Macro-TREM2 (B), Macro-THBS1 (C), Macro-PPARG (D), Macro-SPP1 (E), Macro-ISG (F), and Macro-IER2 (G) across MB, PB, CTB, and Mari.

**Figure S5.**
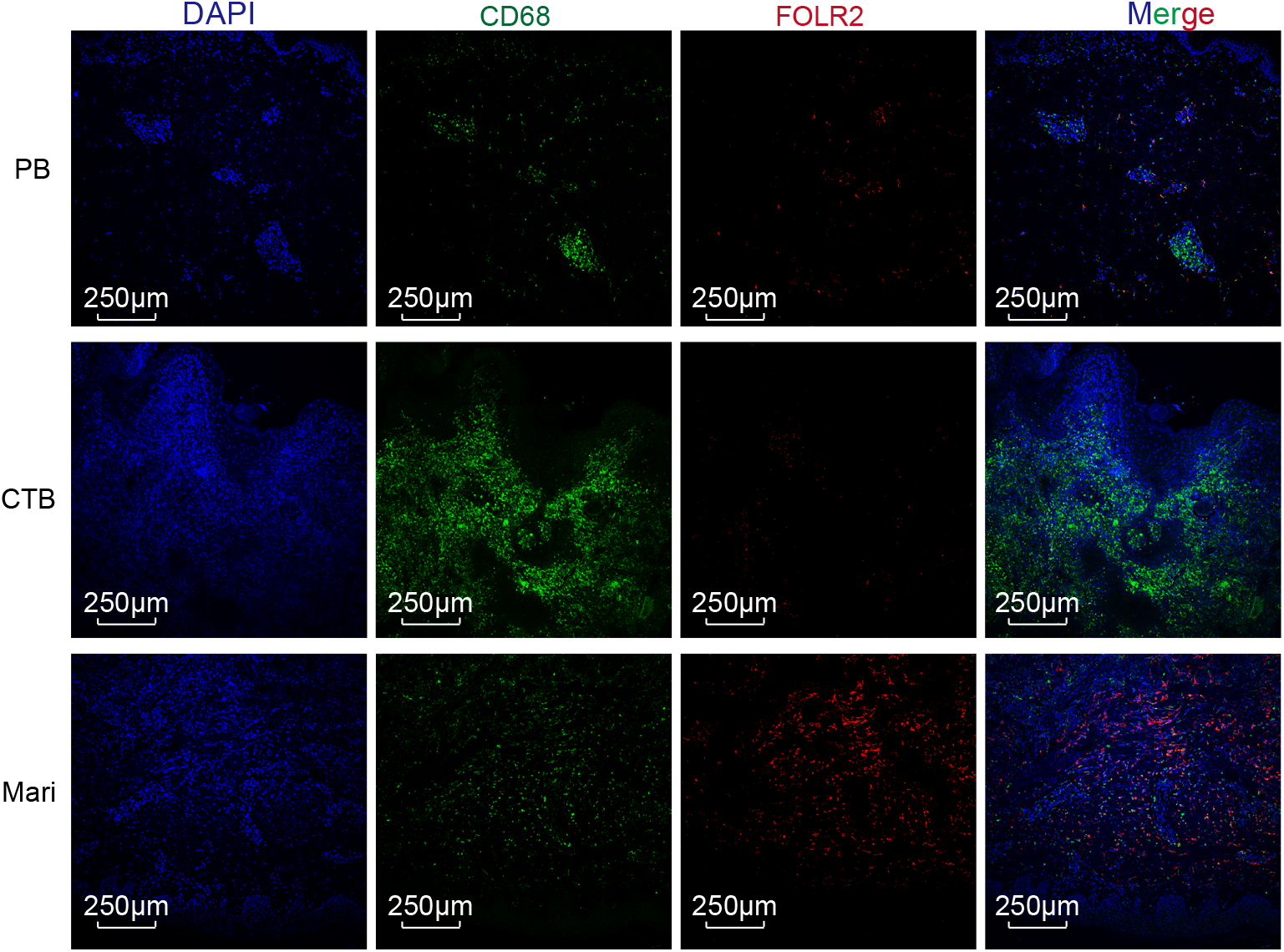
FOLR2 expression in infected skin. Immunofluorescence images of DAPI (blue), CD68 (green) and FOLR2 (red) in PB, CTB and Mari skin biopsies. Images were acquired using 10x.

**Figure S6.**
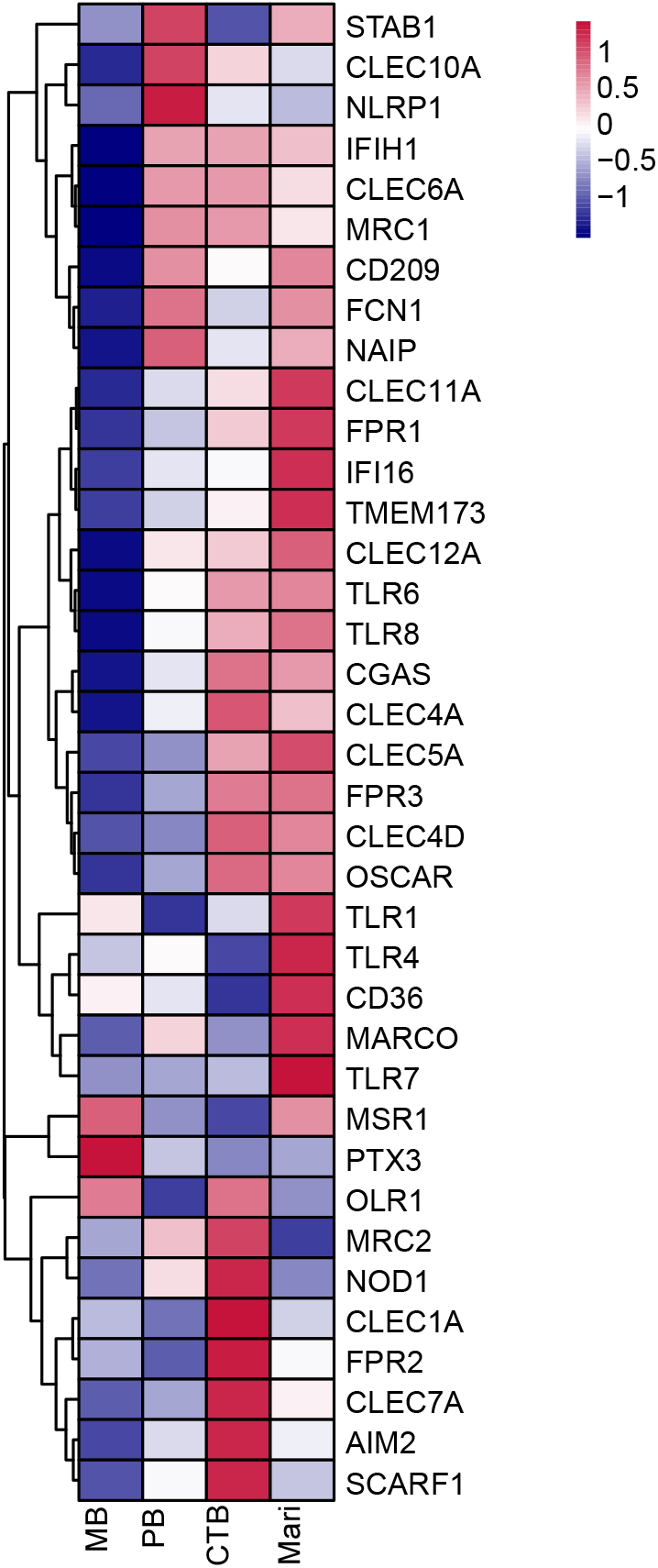
Pathogen-sensing and scavenger receptor gene expression in macrophages. Heatmap of pathogen-sensing and scavenger receptor genes across the total macrophages in MB, PB, CTB, and Mari.

**Figure S7.**
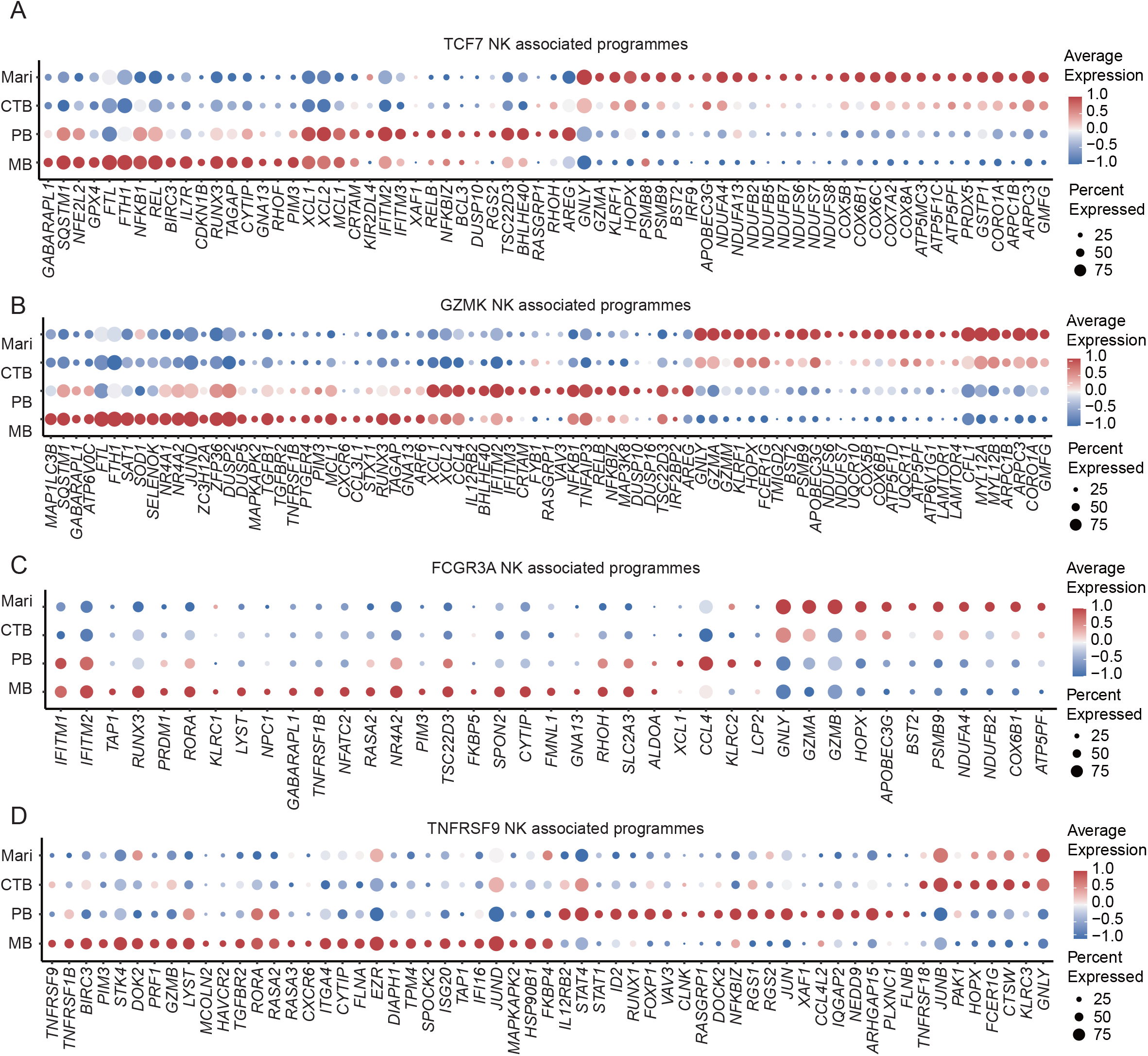
Infection-associated transcriptional programs within NK cell subsets. (A–D) Dotplots of infection-associated marker genes in TCF7 NK (A), GZMK NK (B), FCGR3A NK (C), and TNFRSF9 NK (D) across MB, PB, CTB, and Mari.

## METHODS

### Clinical samples

Skin biopsies from HC, MB, PB, CTB and Mari were obtained from the Biobank of the Institute of Dermatology, Chinese Academy of Medical Sciences, and the Jiangsu Biobank of Clinical Resources. The clinical characteristics were provided in Table S1.

Human peripheral blood mononuclear cells (PBMCs) from healthy donors were obtained from the Jiangsu Province Blood Center. All participants provided written informed consent. The study was approved by the Ethics Committee of the Institute of Dermatology, Chinese Academy of Medical Sciences and Peking Union Medical College (2022-KY-013).

### Reagents and software

Details of all reagents and software used in this study are provided in Table S2.

### Experimental animals

Female specific-pathogen-free (SPF) Institute of Cancer Research (ICR) mice aged 7 weeks were used in this study. Animal use complied with the guidelines of the Institutional Animal Care and Use Committee at the Hospital for Skin Diseases, Institute of Dermatology, Chinese Academy of Medical Sciences, and Peking Union Medical College (2023-DW-019).

### Preparation of skin cell suspensions

Following removal of subcutaneous fat, skin biopsies were minced into fragments (<4 mm²) and processed with the whole-skin dissociation kit (MACS, 130-101-540) according to the manufacturer’s protocol. Tissue digestion was performed at 37 °C for 3 h with shaking in a solution containing 435 μL buffer L, 12.5 μL enzyme P, 50 μL enzyme D, and 2.5 μL enzyme A. After digestion, samples were diluted with ice-cold DMEM supplemented with 2% BSA and passed through a 70 μm cell strainer. Cells were collected from the filtrate by centrifugation at 300 x g for 10 min at 4 °C, and the resulting pellets were resuspended in MACS buffer.

### Single cell RNA sequencing

#### Library preparation and sequencing

Libraries were prepared using Chromium Next GEM Single Cell 3′ Reagent Kits v3.1 (10x Genomics)^48^. Briefly, single-cell suspensions were assessed for cell number and viability before loading onto the Chromium system to generate gel bead-in-emulsion (GEM) droplets. Following reverse transcription, barcoded cDNA was purified and amplified by PCR, then used to construct 3′ gene expression libraries. Amplified cDNA and final libraries underwent quality assessment and quantification before sequencing. Libraries were prepared and were sequenced on Illumina NovaSeq 6000 platform in paired-end mode by Genergy and Genedenovo Biotechnology.

#### Data processing

Raw sequencing reads were processed using Cell Ranger (10x Genomics; version 5.0.0). Following alignment, subsequent analyses were performed using Seurat (version 5.0)^49^. Cells were retained according to the following criteria: (1) >200 detected genes; (2) ≤15% mitochondrial transcripts. Genes detected in fewer than 10 cells were excluded. Potential doublets and multiplets were identified and removed for each sample using Scrublet with an expected doublet rate of 0.05 and a doublet-score threshold of 0.20^50^. After quality control and doublet removal, 266,203 cells were retained for downstream analysis. Gene expression data were normalized using the LogNormalize method, and the top 2,000 highly variable genes were selected using the variance-stabilizing transformation method implemented in FindVariableFeatures for principal component analysis. Batch effects across samples were corrected using Harmony (version 1.2.3)^51^, with sample ID used as the batch variable.

#### Cell clustering and annotation

Following Harmony-based batch correction, a shared nearest-neighbor (SNN) graph was constructed using the FindNeighbors function in Seurat, and cell clustering was performed using the FindClusters function. UMAP was used to visualize the cell populations. For subclustering analyses, selected cell populations were extracted and subjected to a similar workflow, including normalization, highly variable gene selection, dimensionality reduction, Harmony-based batch correction, and clustering. Cluster marker genes were identified using the FindAllMarkers function in Seurat with the Wilcoxon rank-sum test. Genes detected in at least 25% of cells and with a log2 fold change (log_2_FC) ≥ 0.5 and an adjusted P value (p.adj) < 0.05 were considered candidate marker genes. NK cells, monocytes/macrophages and dendritic cells were identified based on the expression of their respective markers, with exclusion of markers associated with other cell types, as performed previously^52–54^. NK cell markers included NKG7, GNLY, GZMB, CST7, KLRD1, PRF1, and FCGR3A; monocyte/macrophage markers included CD14, CD68, CD163, FCGR1A, FCGR2A, FCGR3A, FCN1, S100A8, S100A9, LST1, IL1RN, ALOX5AP, STAB1, TREM2, APOE, C1QA, C1QC, and PLTP. DC subsets and their markers are as follows: cDC1 (CLEC9A, XCR1), cDC2 (CD1C, CLEC10A), LAMP3⁺ DCs (LAMP3, FSCN1, CCR7), plasmacytoid DCs (LILRA4, CLEC4C, GZMB), and Langerhans cells (CD207, CD1A, FCER1A). The exclusion markers for other cell types are as follows: T cells (CD3D, CD3G, TRAC), B cells (CD19, MS4A1, CD79A, CD79B), ILC3 (RORC, KIT, IL17A), ILC2 (GATA3, IL5, IL13, PTGDR2, HPGD, HPGDS), mast cells (CPA3, TPSAB1), neutrophils (CSF3R, FCGR3B, CXCR2), keratinocytes (KRT1, KRT5, KRT10, DSG1), fibroblasts (PDGFRA, MFAP5, COL1A1, COL3A1), endothelial cells (PECAM1, VWF, ACKR1), melanocytes (MLANA, DCT, TYR, TYRP1, PMEL), and Schwann cells (SOX10, S100B, MPZ, PLP1).

#### Cell-type composition analysis

Cell-type proportions were calculated separately for each sample, relative to the total cells in the corresponding parent population. Differences between disease groups were assessed using two-sided Wilcoxon rank-sum tests.

#### Differential expression and KEGG pathway analysis

Differential expression (DE) analysis was performed using the FindMarkers function in Seurat. Genes were considered differentially expressed at an |log_2_FC| ≥ 0.5, p.adj < 0.05, and pct.1 > 0.1 of cells in the target group.

DE genes detected in at least 20% of cells in either comparison group were used for KEGG pathway analysis using the enrichKEGG function in the clusterProfiler package, with Homo sapiens (hsa) specified as the reference organism^55^. Gene symbols were converted to Entrez Gene IDs using the org.Hs.eg.db annotation database. KEGG pathway enrichment analysis was performed separately for upregulated and downregulated genes. P values were adjusted using the Benjamini–Hochberg method, and pathways containing 10-500 genes with p.adj < 0.05 were considered significantly enriched.

#### Functional gene signature scoring

Functional gene signatures were scored in individual cells using the AUCell R package (version 1.28.0)^56^. Signature genes were selected from differentially expressed genes and are listed in Table S3.

#### Pathway analysis

Gene set variation analysis (GSVA; version 2.0.4) was performed on average log-normalized expression profiles grouped by cell type and disease group, using Hallmark gene sets and KEGG pathway gene sets from the Molecular Signatures Database (MSigDB)^57^. The gene sets included in the analysis are listed in Table S4. Metabolic pathway scores were calculated using scMetabolism (version 0.2.1) with the AUCell method and KEGG metabolic gene sets. Scores were compared across disease groups^58^.

#### Disease-associated features

Disease-associated genes were identified using the FindAllMarkers function in Seurat with the Wilcoxon rank-sum test by comparing each disease group with all other groups. Genes with log_2_FC ≥ 0.5 for DC and NK cells and > 1 for macrophages, a detection rate of at least 25% in either of the two comparison groups, and p.adj < 0.05 were retained. The resulting disease-associated gene lists are provided in Tables S5–S7.

### Bulk RNA sequencing

#### Library preparation and sequencing

Total RNA was extracted using TRIzol reagent (Invitrogen, 15596026). RNA concentration and purity were assessed using a NanoDrop One spectrophotometer (Thermo Fisher Scientific), and RNA integrity was evaluated by RNA-specific agarose gel electrophoresis or an Agilent Fragment Analyzer 5400 system. Polyadenylated mRNA was enriched using the NEBNext Poly(A) mRNA Magnetic Isolation Module (New England Biolabs) and fragmented using the NEBNext Ultra II RNA Library Prep Kit for Illumina. Double-stranded cDNA was synthesized using random oligonucleotide primers, followed by end repair, A-tailing, and adapter ligation. cDNA fragments of approximately 400–500 bp were size-selected using AMPure XP beads (Beckman Coulter), PCR-amplified, and purified. Library fragment sizes were assessed using an electrophoresis-based system, and library concentrations were determined using PicoGreen and quantitative PCR (qPCR). The libraries were pooled and sequenced on an Illumina NovaSeq X Plus platform using 150-bp paired-end reads.

#### Read processing and differential expression analysis

Raw sequencing reads were processed using fastp (v0.22.0) to remove 3′ adapter sequences and filter low-quality reads according to the specified quality criteria. Clean reads were aligned to the Mus musculus GRCm39 reference genome using HISAT2 (v2.1.0), with gene annotations from Ensembl release 109. Gene-level read counts were generated using HTSeq (v0.9.1). Differential expression analysis was performed using DESeq2^59^. Genes with |log2FC| ≥ 0.5 and an adjusted P value (p.adj) < 0.05 were considered differentially expressed.

#### Human PBMC isolation

Human peripheral blood was diluted with PBS, layered onto Lymphoprep (STEMCELL Technologies, 07851), and centrifuged at 500 x g for 30 min. PBMCs were collected from the interface and washed twice with PBS by centrifugation at 500 x g for 5 min per wash. The isolated PBMCs were cryopreserved in FBS containing 10% dimethyl sulfoxide (DMSO).

#### Flow cytometry

PBMCs were first stained with Fixable Viability Dye eFluor 780 (Invitrogen, 65-0865-18). For surface staining, cells were incubated in MACS buffer with a FITC-conjugated lineage antibody cocktail against CD11c, CD3, CD4, CD14, TCRαβ, TCRγδ, CD19, CD20, and CD22, together with BV650-conjugated anti-CD56 (BD Biosciences, 564057), at 4°C for 30 min in the dark. For intracellular staining, cells were fixed and permeabilized using the Foxp3 staining kit (eBioscience, 00-5523-00), followed by staining with PerCP-Cyanine5.5-conjugated anti-IFN-γ (eBioscience, 45-7319-42) in permeabilization buffer at 4°C for 30 min in the dark. After washing, cells were subjected to flow cytometric analysis, and data were analyzed using FlowJo (version 10.9.0). All antibodies were used at a 1:200 dilution.

#### Foamy macrophage differentiation and coculture with PBMCs

Monocytes were isolated from PBMCs using Starter CD14 Nanobeads (S0K0004-2mL). Cells were cultured in RPMI 1640 supplemented with 10% FBS and human M-CSF (50 ng/mL; PeproTech, 300-25-50UG) for 7 days to generate monocyte-derived macrophages. The medium was then replaced with fresh medium of the same composition supplemented with oxidized low-density lipoprotein (ox-LDL; 50 μg/mL; Yiyuan Biotechnologies, YB-002), and cells were cultured for another 2 days to generate foamy macrophages. PBMCs were then added to foamy macrophages at a 1:1 ratio and cocultured for 2 days. Human IL-12 (10 ng/mL; Novoprotein, CI58), IL-15 (50 ng/mL; Novoprotein, C016), and IL-18 (50 ng/mL; Novoprotein, CH29) were then added. After 16 hrs of stimulation, PBMCs were collected for flow cytometric analysis.

#### Metabolic perturbation of NK cell responses

PBMCs were pretreated for 2 days with 2-deoxy-D-glucose (2-DG, 5 mM), FCCP (5 μM), metformin (2 mM), 6-aminonicotinamide (6-AN, 10 μM), 3-bromopyruvate (3-BP, 50 μM), lonidamine (100 μM), PFK-158 (500 nM), UK-5099 (20 μM), sodium dichloroacetate (DCA, 5 mM), or G6PDi (50 μM). Cells were then stimulated with IL-12 (10 ng/mL), IL-15 (50 ng/mL), and IL-18 (50 ng/mL) for 16 hrs, and the IFN-γ produced by NK cells were detected by flow cytometry.

#### BMDM isolation, macrophage differentiation and LGC induction

Bone marrow cells were flushed from the femurs of ICR mice with PBS and passed through a 70 μm cell strainer. Cells were centrifuged at 300 x g for 10 min at 4°C, treated with red blood cell lysis buffer (Biolegend, 420301) for approximately 3 min at room temperature, and washed with PBS. Cells were seeded in six-well plates in DMEM supplemented with 10% FBS, 1% penicillin–streptomycin (PS), and 20 ng/mL M-CSF (Novoprotein, CB34) for macrophage differentiation. Fresh medium was added every 3 days. After 6-7 days of differentiation, cells were further cultured for 3 days in the presence of mouse GM-CSF (20 ng/mL; Novoprotein, CK15) and mouse IL-4 (10 ng/mL; Novoprotein, CK02), together with MALP-2 (100 ng/mL), Pam2CSK4 (60 ng/mL), motolimod (800 nM), AICAR (500 μM), G6PDi (50 μM), or FBPi (100 μM), as indicated. LGCs were defined as multinucleated macrophages containing three or more nuclei, and LGC formation was quantified as the number of LGCs per field under a 4x objective.

#### Histology and immunofluorescence

For hematoxylin and eosin (H&E) staining, paraffin sections were deparaffinized, rehydrated through graded ethanol, stained sequentially with hematoxylin and eosin, dehydrated, cleared, and mounted with neutral resin for bright-field microscopy.

For immunofluorescence staining, sections were baked at 60°C for 1 h, deparaffinized, rehydrated, and subjected to antigen retrieval in Tris–EDTA buffer (pH 9.0; Servicebio, G1203-250ML) at 98°C for 25 min. After cooling at room temperature for 30 min, sections were washed three times with PBS and permeabilized with 0.1% Triton X-100 for 10 min. Endogenous peroxidase activity was then quenched using an H₂O₂-containing blocking solution for 15 min, followed by three washes with PBS and blocking with protein-free rapid blocking buffer. Sections were incubated overnight at 4°C with mouse anti-CD68 (Abcam, 1:500) together with rabbit anti-FOLR2 (Abcam, 1:200), anti-GPNMB (Cell signalling technology, 1:400), or anti-SPP1 (HUABIO, 1:200), followed by Alexa Fluor Plus 488-conjugated goat anti-mouse IgG (Invitrogen, 1:500) and Alexa Fluor 555-conjugated goat anti-rabbit IgG (Invitrogen, 1:500) for 40 min in the dark. Nuclei were counterstained with DAPI for 5 min before imaging.

#### Giemsa staining

Cells were fixed with 4% paraformaldehyde for 30 min, washed with PBS, and stained with Giemsa working solution prepared by diluting the concentrated stain 1:9 with the supplied diluent (Solarbio, G1010) for 30 min. After washing, cells were examined by bright-field microscopy.

#### Oil Red O staining

After foamy-cell induction, cells were fixed with 4% paraformaldehyde for 30 min and stained with Oil Red O working solution prepared by mixing stock solution (OriCell, OILR-10001) with distilled water at a 3:2 volume ratio. After staining for 30 min, cells were washed, counterstained with Mayer’s hematoxylin (Solarbio, G1080) for 5 min, and examined by bright-field microscopy.

### QUANTIFICATION AND STATISTICAL ANALYSIS

Paired comparisons were performed using two-tailed paired t-tests or Wilcoxon matched-pairs signed-rank tests, whereas independent groups were compared using two-tailed unpaired t-tests or Wilcoxon rank-sum test, as appropriate. Cell-type proportions from single-cell analyses were compared between groups using two-sided Wilcoxon rank-sum tests in R. Correlations were assessed using Spearman’s rank correlation. Data are presented as mean ± SEM or mean ± SD, as specified in the corresponding figure legends.

## REFERENCES

1. World Health Organization. Global Tuberculosis Report 2025. https://www.who.int/publications/i/item/9789240116924 (2025).

2. Brito, A. C. de, Oliveira, C. M. M. de, Unger, D. A.-A. & Bittencourt, M. de J. S. Cutaneous tuberculosis: epidemiological, clinical, diagnostic and therapeutic update. An. Bras. Dermatol. 97, 129–144 (2022).

3. World Health Organization. Leprosy. https://www.who.int/news-room/fact-sheets/detail/leprosy (2026).

4. de Paula, H. L. et al. Risk Factors for Physical Disability in Patients With Leprosy. JAMA Dermatol. 155, 1120–1128 (2019).

5. Tsiolakkis, G. et al. Mycobacterium marinum: A Case-Based Narrative Review of Diagnosis and Management. Microorganisms 11, 1799 (2023).

6. dos Santos, J. B., et al. Cutaneous tuberculosis: diagnosis, histopathology and treatment - Part II. An. Bras. Dermatol. 89, 545–555 (2014).

7. Bhat, R. M. & Prakash, C. Leprosy: An Overview of Pathophysiology. Interdiscip. Perspect. Infect. Dis. 2012, 181089 (2012).

8. de Sousa, J. R., Sotto, M. N. & Simões Quaresma, J. A. Leprosy As a Complex Infection: Breakdown of the Th1 and Th2 Immune Paradigm in the Immunopathogenesis of the Disease. Front. Immunol. 8, 1635 (2017).

9. Wu, T.-S. et al. Fish Tank Granuloma Caused by Mycobacterium marinum. PLoS ONE 7, e41296 (2012).

10. Haberecht, H. B., Cull, D. B. & Wieland, C. N. Mycobacterium marinum. Mayo Clin. Proc. 99, 196–197 (2024).

11. Sturgill-Koszycki, S. et al. Lack of acidification in Mycobacterium phagosomes produced by exclusion of the vesicular proton-ATPase. Science 263, 678–681 (1994).

12. Chandra, P., Grigsby, S. J. & Philips, J. A. Immune evasion and provocation by Mycobacterium tuberculosis. Nat. Rev. Microbiol. 20, 750–766 (2022).

13. Huang, L., Nazarova, E. V., Tan, S., Liu, Y. & Russell, D. G. Growth of Mycobacterium tuberculosis in vivo segregates with host macrophage metabolism and ontogeny. J. Exp. Med. 215, 1135–1152 (2018).

14. Weiss, G. & Schaible, U. E. Macrophage defense mechanisms against intracellular bacteria. Immunol. Rev. 264, 182–203 (2015).

15. Teles, R. M. B. et al. Type I Interferon Suppresses Type II Interferon–Triggered Human Anti-Mycobacterial Responses. Science 339, 1448–1453 (2013).

16. Mattos, K. A. et al. Mycobacterium leprae intracellular survival relies on cholesterol accumulation in infected macrophages: a potential target for new drugs for leprosy treatment. Cell. Microbiol. 16, 797–815 (2014).

17. Clay, H. et al. Dichotomous role of the macrophage in early Mycobacterium marinum infection of the zebrafish. Cell Host Microbe 2, 29–39 (2007).

18. Cambier, C. J. et al. Mycobacteria manipulate macrophage recruitment through coordinated use of membrane lipids. Nature 505, 218–222 (2014).

19. Davis, J. M. et al. Real-Time Visualization of Mycobacterium-Macrophage Interactions Leading to Initiation of Granuloma Formation in Zebrafish Embryos. Immunity 17, 693–702 (2002).

20. Wolf, A. J. et al. Mycobacterium tuberculosis infects dendritic cells with high frequency and impairs their function in vivo. J. Immunol. Baltim. Md 1950 179, 2509–2519 (2007).

21. Lee, D. J. et al. LILRA2 Activation Inhibits Dendritic Cell Differentiation and Antigen Presentation to T Cells. J. Immunol. Baltim. Md 1950 179, 8128–8136 (2007).

22. Sieling, P. A. et al. CD1 Expression by Dendritic Cells in Human Leprosy Lesions: Correlation with Effective Host Immunity. J. Immunol. 162, 1851–1858 (1999).

23. Vankayalapati, R. et al. The NKp46 Receptor Contributes to NK Cell Lysis of Mononuclear Phagocytes Infected with an Intracellular Bacterium. J. Immunol. 168, 3451–3457 (2002).

24. García, V. E. et al. IL-18 Promotes Type 1 Cytokine Production from NK Cells and T Cells in Human Intracellular Infection. J. Immunol. 162, 6114–6121 (1999).

25. Maier, B. et al. A conserved dendritic-cell regulatory program limits antitumour immunity. Nature 580, 257–262 (2020).

26. Hunger, R. E. et al. Langerhans cells utilize CD1a and langerin to efficiently present nonpeptide antigens to T cells. J. Clin. Invest. 113, 701–708 (2004).

27. Dick, S. A., et al. Three tissue resident macrophage subsets coexist across organs with conserved origins and life cycles. Sci. Immunol. 7, eabf7777 (2022).

28. Jaitin, D. A. et al. Lipid-associated macrophages control metabolic homeostasis in a Trem2-dependent manner. Cell 178, 686–698.e14 (2019).

29. Montoya, D. et al. Divergence of macrophage phagocytic and antimicrobial programs in leprosy. Cell Host Microbe 6, 343–353 (2009).

30. Fabri, M. et al. Vitamin D Is Required for IFN-γ–Mediated Antimicrobial Activity of Human Macrophages. Sci. Transl. Med. 3, 104ra102 (2011).

31. Walch, M. et al. Cytotoxic Cells Kill Intracellular Bacteria through Granulysin-Mediated Delivery of Granzymes. Cell 157, 1309–1323 (2014).

32. Cruz, D. et al. Host-derived oxidized phospholipids and HDL regulate innate immunity in human leprosy. J. Clin. Invest. 118, 2917–2928 (2008).

33. Mattos, K. A. et al. Modulation of lipid droplets by Mycobacterium leprae in Schwann cells: a putative mechanism for host lipid acquisition and bacterial survival in phagosomes. Cell. Microbiol. 13, 259–273 (2011).

34. Jin, S.-H., An, S.-K. & Lee, S.-B. The formation of lipid droplets favors intracellular Mycobacterium leprae survival in SW-10, non-myelinating Schwann cells. PLoS Negl. Trop. Dis. 11, e0005687 (2017).

35. Joo, M. et al. Induction and Function of Lipocalin Prostaglandin D Synthase in Host Immunity. J. Immunol. 179, 2565–2575 (2007).

36. Bedi, B. et al. Enhanced Clearance of Pseudomonas aeruginosa by Peroxisome Proliferator-Activated Receptor Gamma. Infect. Immun. 84, 1975–1985 (2016).

37. Hou, Y. et al. SMPDL3A is a cGAMP-degrading enzyme induced by LXR-mediated lipid metabolism to restrict cGAS-STING DNA sensing. Immunity 56, 2492–2507.e10 (2023).

38. Watson, R. O. et al. The cytosolic sensor cGAS detects Mycobacterium tuberculosis DNA to induce type I interferons and activate autophagy. Cell Host Microbe 17, 811–819 (2015).

39. Jiang, Y. et al. Langhans giant cells regulate cutaneous immune responses during mycobacterial infection through CXCL1/CXCL2 secretion. Int. Immunopharmacol. 152, 114447 (2025).

40. Lay, G. et al. Langhans giant cells from M. tuberculosis-induced human granulomas cannot mediate mycobacterial uptake. J. Pathol. 211, 76–85 (2007).

41. Wang, Z. et al. Analysis of the efficacy of Langhans giant cells in controlling cutaneous Mycobacterium marinum infections. Int. Immunopharmacol. 148, 114003 (2025).

42. Stenger, S. et al. An antimicrobial activity of cytolytic T cells mediated by granulysin. Science 282, 121–125 (1998).

43. Keating, S. E. et al. Metabolic Reprogramming Supports IFN-γ Production by CD56bright NK Cells. J. Immunol. 196, 2552–2560 (2016).

44. Wang, Z. et al. Glycolysis and Oxidative Phosphorylation Play Critical Roles in Natural Killer Cell Receptor-Mediated Natural Killer Cell Functions. Front. Immunol. 11, 202 (2020).

45. Kern Coquillat, N., et al. Pivotal role of exogenous pyruvate in human natural killer cell metabolism. Nat. Metab. 7, 336–347 (2025).

46. Keane, C. et al. Macrophages tune NK cell IFNγ production through direct lipid transfer. Nat. Commun. 17, 9527 (2026).

47. Jiang, S. et al. Generic Diagramming Platform (GDP): a comprehensive database of high-quality biomedical graphics. Nucleic Acids Res. 53, D1670–D1676 (2025).

48. Zheng, G. X. Y. et al. Massively parallel digital transcriptional profiling of single cells. Nat. Commun. 8, 14049 (2017).

49. Hao, Y. et al. Dictionary learning for integrative, multimodal and scalable single-cell analysis. Nat. Biotechnol. 42, 293–304 (2024).

50. Wolock, S. L., Lopez, R. & Klein, A. M. Scrublet: Computational Identification of Cell Doublets in Single-Cell Transcriptomic Data. Cell Syst. 8, 281–291.e9 (2019).

51. Korsunsky, I. et al. Fast, sensitive and accurate integration of single-cell data with Harmony. Nat. Methods 16, 1289–1296 (2019).

52. Wei, Q. et al. Glucocorticoid Receptor Activation Reprograms NK Cells to Drive AREG-Mediated Immunosuppression: A Pan-Cancer Role for AREG. Adv. Sci. 13, e12620 (2026).

53. Liang, G., Zhao, C., Wei, Q., Feng, S. & Wang, Y. Single cell transcriptome profiling reveals pathogenesis of bullous pemphigoid. *Commun*. Biol. 8, 203 (2025).

54. Zhao, Y. et al. Natural killer cell dysfunction drives keloid pathogenesis. Cell Rep. 45, 117129 (2026).

55. Wu, T. et al. clusterProfiler 4.0: A universal enrichment tool for interpreting omics data. The Innovation 2, 100141 (2021).

56. Aibar, S. et al. SCENIC: single-cell regulatory network inference and clustering. Nat. Methods 14, 1083–1086 (2017).

57. Hänzelmann, S., Castelo, R. & Guinney, J. GSVA: gene set variation analysis for microarray and RNA-Seq data. BMC Bioinformatics 14, 7 (2013).

58. Wu, Y. et al. Spatiotemporal Immune Landscape of Colorectal Cancer Liver Metastasis at Single-Cell Level. Cancer Discov. 12, 134–153 (2022).

59. Love, M. I., Huber, W. & Anders, S. Moderated estimation of fold change and dispersion for RNA-seq data with DESeq2. Genome Biol. 15, 550 (2014).

