## Supplemental tables 1-7 for "Shared and distinct immunometabolic states link to divergent pathology in cutaneous mycobacterial infections"

| Table S1 \| Skin samples for scRNA-Seq | | | |
|---|---|---|---|
| Sample ID | Sex | Age | Primary site |
| HC 1 | Male | 48 | Upper limb |
| HC 2 | Female | 33 | Upper limb |
| HC 3 | Female | 33 | Back |
| HC 4 | Male | 70 | Upper limb |
| HC 5 | Male | 48 | Buttock |
| HC 6 | Female | 60 | Face |
| MB 1 | Male | 45 | Abdomen |
| MB 2 | Female | 22 | Upper limb |
| MB 3 | Male | 28 | Thigh |
| MB 4 | Male | 26 | Abdomen |
| MB 5 | Male | 31 | Upper limb |
| MB 6 | Female | 43 | Abdomen |
| MB 7 | Male | 40 | Thigh |
| PB 1 | Female | 80 | Thigh |
| PB 2 | Male | 58 | Thigh |
| PB 3 | Male | 33 | Upper limb |
| PB 4 | Male | 25 | Thigh |
| PB 5 | Female | 47 | Thigh |
| PB 6 | Female | 56 | Neck |
| PB 7 | Male | 59 | Face |
| PB 8 | Male | 67 | Upper limb |
| CTB 1 | Female | 51 | Lower back |
| CTB 2 | Female | 19 | Left ear |
| CTB 3 | Male | 69 | Buttock |
| CTB 4 | Female | 68 | Face |
| CTB 5 | Female | 30 | Upper limb |
| CTB 6 | Male | 63 | Upper limb |
| CTB 7 | Female | 66 | Neck |
| CTB 8 | Female | 47 | Face |
| Mari 1 | Female | 66 | Upper limb |
| Mari 2 | Female | 55 | Upper limb |
| Mari 3 | Male | 55 | Upper limb |
| Mari 4 | Female | 66 | Upper limb |

| Table S2 \| Reagent and resource | | |
|---|---|---|
| **Reagent or resource** | **Source** | **Identifier** |
| **Antibodies** | | |
| Anti-Human CD3 | BioLegend | Cat# 317306 |
| Anti-Human CD4 | BioLegend | Cat# 317408 |
| Anti-Human TCRα/β | BioLegend | Cat# 306706 |
| Anti-Human TCRγ/δ | BioLegend | Cat# 331208 |
| Anti-Human CD19 | BioLegend | Cat# 302206 |
| Anti-Human CD20 | BioLegend | Cat# 302304 |
| Anti-Human CD22 | BioLegend | Cat# 363508 |
| Anti-Human CD14 | BioLegend | Cat# 325604 |
| Anti-Human IFN-γ | eBioscience | Cat# 45-7319-42 |
| Anti-Human CD56 | BD | Cat# 564057 |
| GPNMB (E4D7P) Rabbit Monoclonal Antibody | Cell Signaling Technology | Cat# 38313 |
| Anti-CD68 antibody | Abcam | Cat# ab955 |
| Anti-FOLR2 antibody | Abcam | Cat# ab302532 |
| Osteopontin/SPP1 Recombinant Rabbit Monoclonal Antibody | HUABIO | Cat# HA723081 |
| Goat anti-Mouse IgG (H+L) Highly Cross-Adsorbed Secondary Antibody, Alexa Fluor™ Plus 488 | Invitrogen | Cat# A32723 |
| Goat anti-Rabbit IgG (H+L) Highly Cross-Adsorbed Secondary Antibody, Alexa Fluor™ 555 | Invitrogen | Cat# A21429 |
| Biological Samples | | |
| PBMCs | Jiangsu Province Blood Center | https://www.jsblood.com.cn/ |
| **Chemicals, Peptides, and Recombinant Proteins** | | |
| IL-12 (human) | Novoprotein | Cat# CI58 |
| IL-15 (human) | Novoprotein | Cat# C016 |
| IL-18 (human) | Novoprotein | Cat# CH29 |
| M-CSF (human) | Peprotech | Cat# 300-25-50UG |
| M-CSF (mouse) | Novoprotein | Cat# CB34 |
| IL-4 (mouse) | Novoprotein | Cat# CK02 |
| GM-CSF (mouse) | Novoprotein | Cat# CK15 |
| TRIzol reagent | Invitrogen | Cat# 15596026 |

| | | |
|---|---|---|
| Fixable Viability Dye efluor 780 | Invitrogen | Cat# 65-0865-18 |
| CD14 Nanobeads, human (RUO) | Starter | Cat# S0K0004-2mL |
| Brefeldin A | Beyotime | Cat# S1536-5mg |
| Monensin | Beyotime | Cat# S1753 |
| 20× Tris-EDTA Antigen Retrieval Solution (pH 9.0) | Servicebio | Cat# G1203-250ML |
| BioDewax and Clear Solution | Servicebio | Cat# G1128-1L |
| Tween-20 | Beyotime | Cat# ST825-100ml |
| 4% paraformaldehyde | Biosharp | Cat# BL539A |
| Triton X-100 | Servicebio | Cat# GC204003-100ml |
| DAPI Staining Solution | Beyotime | Cat# C1005 |
| Giemsa Stain Solution (Working Solution) | Solarbio | Cat# G1010 |
| Oil Red O Solution | OriCell | Cat# OILR-10001 |
| Mayer’s Hematoxylin Stain Solution | Solarbio | Cat# G1080 |
| ox-LDL | Yiyuanbiotech | Cat# YB-002 |
| Macrophage-activating lipopeptide 2 (MALP-2) | MedChemExpress | Cat# HY-P10586 |
| Pam2CSK4 | InvivoGen | Cat# tlrl-pm2s-1 |
| Motolimod | MedChemExpress | Cat# HY-13773 |
| FBPase-1 inhibitor-1 (FBPi) | MedChemExpress | Cat# HY-136717 |
| AICAR | MedChemExpress | Cat# HY-13417 |
| G6PDi-1 | MedChemExpress | Cat# HY-W107464 |
| 2-Deoxy-D-glucose (2-DG) | MedChemExpress | Cat# HY-13966 |
| Metformin | MedChemExpress | Cat# HY-B0627 |
| Sodium dichloroacetate (DCA) | MedChemExpress | Cat# HY-Y0445A |
| FCCP | MedChemExpress | Cat# HY-100410 |
| 6-Aminonicotinamide | MedChemExpress | Cat# HY-W010342 |
| Lonidamine | MedChemExpress | Cat# HY-B0486 |
| PFK-158 | MedChemExpress | Cat# HY-12203 |
| 3-Bromopyruvic acid | MedChemExpress | Cat# HY-19992 |

| | | |
|---|---|---|
| UK-5099 | MedChemExpress | Cat# HY-15475 |
| **Software and Algorithms** | | |
| FlowJo | FlowJo, LLC | https://www.flowjo.com/ |
| GraphPad Prism | GraphPad Software, Inc | https://www.graphpad.com/features |
| Cell Ranger | 10x Genomics | https://www.10xgenomics.com/support/cn/software/cell-ranger/latest/algorithms-overview/cr-gex-algorithm |
| R Statistical Computing Software | R Foundation for Statistical Computing | https://www.r-project.org/ |
| Seurat 5.1.0 | Satija Lab | https://satijalab.org/seurat/ |
| Harmony 1.2.3 | Harmony developers | https://github.com/immunogenomics/harmony |
| Scrublet | Scrublet developers | https://github.com/swolock/scrublet |
| AUCell 1.28.0 | Aerts Lab | https://bioconductor.org/packages/release/bioc/html/AUCell.html |
| GSVA 2.0.4 | Bioconductor | https://bioconductor.org/packages/release/bioc/html/GSVA.html |
| clusterProfiler 4.14.4 | Bioconductor | https://bioconductor.org/packages/clusterProfiler/ |
| scMetabolism 0.2.1 | Gao Lab | https://github.com/wu-yc/scMetabolism |
| DESeq2 1.46.0 | Bioconductor | https://bioconductor.org/packages/release/bioc/html/DESeq2.html |
| fastp | OpenGene | https://github.com/OpenGene/fastp |
| HISAT2 | Daehwan Kim Lab | https://daehwanki |

| | | |
|---|---|---|
| | | mlab.github.io/hisat2/ |
| HTSeq | HTSeq developers | https://htseq.readthedocs.io/en/latest/ |

| **Table S3 \| Functional genes used for AUCell scoring** | | | |
|---|---|---|---|
| **Bacterial recognition** | **Phagocytosis** | **Phagosome maturation** | **Lysosomal degradation** |
| TLR1 | FCGR1A | RAB5A | CTSB |
| TLR2 | FCGR2A | RAB7A | CTSD |
| TLR4 | FCGR3A | RILP | CTSL |
| TLR6 | ITGAM | VPS18 | CTSS |
| CD14 | ITGB2 | VPS33A | CTSZ |
| CLEC4D | CD14 | VPS41 | LGMN |
| CLEC4E | MSR1 | STX7 | LYZ |
| CLEC6A | MARCO | STX8 | ACP2 |
| CLEC7A | CLEC7A | VAMP7 | HEXA |
| MARCO | FCER1G | VAMP8 | HEXB |
| MSR1 | TYROBP | LAMP1 | GAA |
| MRC1 | SYK | CLCN7 | GBA |
| CD209 | RAC2 | MCOLN1 | PSAP |
| CD36 | | ATP6V1A | SCARB2 |
| NOD1 | | ATP6V1B2 | |
| NOD2 | | ATP6V1H | |
| RIPK2 | | | |
| CGAS | | | |
| TMEM173 | | | |
| IFI16 | | | |
| AIM2 | | | |
| **ROS killing** | **Iron restriction** | **Xenophagy** | **Vitamin D–CAMP** |
| CYBB | SLC11A1 | ATG3 | TLR1 |
| CYBA | SLC40A1 | ATG7 | TLR2 |
| NCF1 | CP | ATG10 | CD14 |
| NCF2 | FTH1 | ATG13 | CYP27B1 |
| NCF4 | FTL | MAP1LC3B | VDR |
| RAC2 | | WIPI1 | CAMP |
| | | WIPI2 | |
| | | SQSTM1 | |
| | | CALCOCO2 | |
| | | OPTN | |
| | | TAX1BP1 | |

| | | | |
|---|---|---|---|
| | | TBK1 | |
| | | UVRAG | |
| | | LRSAM1 | |
| | | SMURF1 | |
| **IFN-γ antibacterial response** | **Inflammasome/ pyroptosis** | **Inflammatory recruitment** | **MHC class II presentation** |
| STAT1 | AIM2 | TNF | CIITA |
| IRF1 | NLRP3 | IL1B | CD74 |
| GBP1 | PYCARD | IL6 | HLA-DMA |
| GBP2 | CASP1 | CXCL8 | HLA-DMB |
| GBP3 | CASP4 | CCL2 | HLA-DPA1 |
| GBP4 | CASP5 | CCL3 | HLA-DPB1 |
| GBP5 | GSDMD | CCL4 | HLA-DRA |
| FCGR1A | IL1B | CXCL9 | HLA-DRB1 |
| CXCL9 | IL18 | CXCL10 | |
| CXCL10 | | | |
| CXCL11 | | | |
| **Fusion** | | | |
| DCSTAMP | | | |
| OCSTAMP | | | |
| CD47 | | | |
| SIRPA | | | |
| KCNN4 | | | |
| MMP9 | | | |
| ATP6V0D2 | | | |
| ADAM9 | | | |
| ADAM12 | | | |
| ITGB1 | | | |

| Table S4\| Gene sets from MSigDB used for GSVA |
|---|
| **name** |
| HALLMARK_ANGIOGENESIS |
| HALLMARK_APOPTOSIS |
| HALLMARK_CHOLESTEROL_HOMEOSTASIS |
| HALLMARK_COAGULATION |
| HALLMARK_COMPLEMENT |
| HALLMARK_DNA_REPAIR |
| HALLMARK_E2F_TARGETS |
| HALLMARK_FATTY_ACID_METABOLISM |
| HALLMARK_G2M_CHECKPOINT |
| HALLMARK_GLYCOLYSIS |
| HALLMARK_HEME_METABOLISM |
| HALLMARK_HYPOXIA |
| HALLMARK_IL6_JAK_STAT3_SIGNALING |
| HALLMARK_INFLAMMATORY_RESPONSE |
| HALLMARK_INTERFERON_ALPHA_RESPONSE |
| HALLMARK_INTERFERON_GAMMA_RESPONSE |
| HALLMARK_KRAS_SIGNALING_DN |
| HALLMARK_KRAS_SIGNALING_UP |
| HALLMARK_MITOTIC_SPINDLE |
| HALLMARK_MTORC1_SIGNALING |
| HALLMARK_MYC_TARGETS_V1 |
| HALLMARK_MYC_TARGETS_V2 |
| HALLMARK_NOTCH_SIGNALING |
| HALLMARK_OXIDATIVE_PHOSPHORYLATION |
| HALLMARK_P53_PATHWAY |
| HALLMARK_PEROXISOME |
| HALLMARK_PI3K_AKT_MTOR_SIGNALING |
| HALLMARK_PROTEIN_SECRETION |
| HALLMARK_REACTIVE_OXYGEN_SPECIES_PATHWAY |
| HALLMARK_TGF_BETA_SIGNALING |
| HALLMARK_TNFA_SIGNALING_VIA_NFKB |
| HALLMARK_UNFOLDED_PROTEIN_RESPONSE |
| HALLMARK_WNT_BETA_CATENIN_SIGNALING |
| HALLMARK_XENOBIOTIC_METABOLISM |
| KEGG_ABC_TRANSPORTERS |
| KEGG_ADHERENS_JUNCTION |
| KEGG_ALANINE_ASPARTATE_AND_GLUTAMATE_METABOLISM |
| KEGG_ALPHA_LINOLENIC_ACID_METABOLISM |
| KEGG_AMINOACYL_TRNA_BIOSYNTHESIS |
| KEGG_AMINO_SUGAR_AND_NUCLEOTIDE_SUGAR_METABOLISM |
| KEGG_ANTIGEN_PROCESSING_AND_PRESENTATION |
| KEGG_ARACHIDONIC_ACID_METABOLISM |
| KEGG_ARGININE_AND_PROLINE_METABOLISM |
| KEGG_ASCORBATE_AND_ALDARATE_METABOLISM |
| KEGG_BASAL_TRANSCRIPTION_FACTORS |
| KEGG_BASE_EXCISION_REPAIR |
| KEGG_BETA_ALANINE_METABOLISM |
| KEGG_BIOSYNTHESIS_OF_UNSATURATED_FATTY_ACIDS |
| KEGG_BUTANOATE_METABOLISM |
| KEGG_CALCIUM_SIGNALING_PATHWAY |
| KEGG_CELL_ADHESION_MOLECULES_CAMS |
| KEGG_CELL_CYCLE |
| KEGG_CHEMOKINE_SIGNALING_PATHWAY |
| KEGG_COMPLEMENT_AND_COAGULATION_CASCADES |

| |
|---|
| KEGG_CYSTEINE_AND_METHIONINE_METABOLISM |
| KEGG_CYTOKINE_CYTOKINE_RECEPTOR_INTERACTION |
| KEGG_CYTOSOLIC_DNA_SENSING_PATHWAY |
| KEGG_DNA_REPLICATION |
| KEGG_ECM_RECEPTOR_INTERACTION |
| KEGG_ENDOCYTOSIS |
| KEGG_ERBB_SIGNALING_PATHWAY |
| KEGG_FC_EPSILON_RI_SIGNALING_PATHWAY |
| KEGG_FC_GAMMA_R_MEDIATED_PHAGOCYTOSIS |
| KEGG_FOCAL_ADHESION |
| KEGG_GLUTATHIONE_METABOLISM |
| KEGG_GLYCEROLIPID_METABOLISM |
| KEGG_GLYCEROPHOSPHOLIPID_METABOLISM |
| KEGG_GLYCINE_SERINE_AND_THREONINE_METABOLISM |
| KEGG_GLYCOLYSIS_GLUCONEOGENESIS |
| KEGG_GLYCOSAMINOGLYCAN_BIOSYNTHESIS_HEPARAN_SULFATE |
| KEGG_GLYCOSAMINOGLYCAN_DEGRADATION |
| KEGG_GLYCOSPHINGOLIPID_BIOSYNTHESIS_GLOBO_SERIES |
| KEGG_GLYCOSPHINGOLIPID_BIOSYNTHESIS_LACTO_AND_NEOLACTO_SERIES |
| KEGG_GLYCOSYLPHOSPHATIDYLINOSITOL_GPI_ANCHOR_BIOSYNTHESIS |
| KEGG_HEDGEHOG_SIGNALING_PATHWAY |
| KEGG_HEMATOPOIETIC_CELL_LINEAGE |
| KEGG_HISTIDINE_METABOLISM |
| KEGG_HOMOLOGOUS_RECOMBINATION |
| KEGG_INOSITOL_PHOSPHATE_METABOLISM |
| KEGG_JAK_STAT_SIGNALING_PATHWAY |
| KEGG_LEUKOCYTE_TRANSENDOTHELIAL_MIGRATION |
| KEGG_LINOLEIC_ACID_METABOLISM |
| KEGG_LYSINE_DEGRADATION |
| KEGG_LYSOSOME |
| KEGG_MAPK_SIGNALING_PATHWAY |
| KEGG_METABOLISM_OF_XENOBIOTICS_BY_CYTOCHROME_P450 |
| KEGG_MTOR_SIGNALING_PATHWAY |
| KEGG_NICOTINATE_AND_NICOTINAMIDE_METABOLISM |
| KEGG_NITROGEN_METABOLISM |
| KEGG_NOD_LIKE_RECEPTOR_SIGNALING_PATHWAY |
| KEGG_NON_HOMOLOGOUS_END_JOINING |
| KEGG_NUCLEOTIDE_EXCISION_REPAIR |
| KEGG_N_GLYCAN_BIOSYNTHESIS |
| KEGG_ONE_CARBON_POOL_BY_FOLATE |
| KEGG_OTHER_GLYCAN_DEGRADATION |
| KEGG_O_GLYCAN_BIOSYNTHESIS |
| KEGG_P53_SIGNALING_PATHWAY |
| KEGG_PANTOTHENATE_AND_COA_BIOSYNTHESIS |
| KEGG_PENTOSE_AND_GLUCURONATE_INTERCONVERSIONS |
| KEGG_PENTOSE_PHOSPHATE_PATHWAY |
| KEGG_PHENYLALANINE_METABOLISM |
| KEGG_PHOSPHATIDYLINOSITOL_SIGNALING_SYSTEM |
| KEGG_PORPHYRIN_AND_CHLOROPHYLL_METABOLISM |
| KEGG_PPAR_SIGNALING_PATHWAY |
| KEGG_PRIMARY_IMMUNODEFICIENCY |
| KEGG_PROPANOATE_METABOLISM |
| KEGG_PROTEASOME |
| KEGG_PROTEIN_EXPORT |
| KEGG_PURINE_METABOLISM |
| KEGG_PYRIMIDINE_METABOLISM |

| |
|---|
| KEGG_PYRUVATE_METABOLISM |
| KEGG_REGULATION_OF_ACTIN_CYTOSKELETON |
| KEGG_REGULATION_OF_AUTOPHAGY |
| KEGG_RETINOL_METABOLISM |
| KEGG_RIBOFLAVIN_METABOLISM |
| KEGG_RIBOSOME |
| KEGG_RIG_I_LIKE_RECEPTOR_SIGNALING_PATHWAY |
| KEGG_RNA_DEGRADATION |
| KEGG_RNA_POLYMERASE |
| KEGG_SELENOAMINO_ACID_METABOLISM |
| KEGG_SNARE_INTERACTIONS_IN_VESICULAR_TRANSPORT |
| KEGG_SPHINGOLIPID_METABOLISM |
| KEGG_SPLICEOSOME |
| KEGG_SULFUR_METABOLISM |
| KEGG_TAURINE_AND_HYPOTAURINE_METABOLISM |
| KEGG_TOLL_LIKE_RECEPTOR_SIGNALING_PATHWAY |
| KEGG_TRYPTOPHAN_METABOLISM |
| KEGG_TYROSINE_METABOLISM |
| KEGG_UBIQUITIN_MEDIATED_PROTEOLYSIS |
| KEGG_VALINE_LEUCINE_AND_ISOLEUCINE_DEGRADATION |
| KEGG_VEGF_SIGNALING_PATHWAY |
| KEGG_WNT_SIGNALING_PATHWAY |
| KEGG_CITRATE_CYCLE_TCA_CYCLE |
| HALLMARK_IL2_STAT5_SIGNALING |
| KEGG_FRUCTOSE_AND_MANNOSE_METABOLISM |
| KEGG_GALACTOSE_METABOLISM |

| Table S5 \| Disease-associated genes in DCs | | | | | | | |
|---|---|---|---|---|---|---|---|
| **Disease-associated genes in all DCs** | | | | | | | |
| **group** | **gene** | **p_val** | **avg_log2FC** | **pct.1** | **pct.2** | **p_val_adj** | **cluster** |
| allcell | APOE | 8.54E-30 | 2.52591241 | 0.436 | 0.273 | 1.68E-25 | Lep-L |
| allcell | APOC1 | 4.10E-10 | 2.213247562 | 0.331 | 0.247 | 8.06E-06 | Lep-L |
| allcell | PLIN2 | 1.67E-20 | 2.10511571 | 0.402 | 0.27 | 3.29E-16 | Lep-L |
| allcell | C5AR1 | 7.03E-19 | 2.081133459 | 0.285 | 0.16 | 1.38E-14 | Lep-L |
| allcell | IL1R2 | 2.22E-46 | 1.851453984 | 0.521 | 0.273 | 4.36E-42 | Lep-L |
| allcell | CXCL3 | 2.76E-23 | 1.848924132 | 0.365 | 0.202 | 5.44E-19 | Lep-L |
| allcell | HMOX1 | 1.79E-14 | 1.824431916 | 0.465 | 0.381 | 3.53E-10 | Lep-L |
| allcell | KCNN4 | 2.60E-23 | 1.791993319 | 0.297 | 0.151 | 5.12E-19 | Lep-L |
| allcell | PID1 | 1.79E-17 | 1.773613851 | 0.353 | 0.219 | 3.52E-13 | Lep-L |
| allcell | MID1IP1 | 8.28E-07 | 1.613503948 | 0.275 | 0.209 | 0.016306958 | Lep-L |
| allcell | CXCL8 | 2.16E-61 | 1.493964307 | 0.705 | 0.433 | 4.25E-57 | Lep-L |
| allcell | HSPA6 | 6.28E-37 | 1.492688372 | 0.626 | 0.417 | 1.24E-32 | Lep-L |
| allcell | BAG3 | 2.38E-40 | 1.434337747 | 0.725 | 0.547 | 4.68E-36 | Lep-L |
| allcell | PAF1 | 3.43E-22 | 1.429786908 | 0.447 | 0.304 | 6.76E-18 | Lep-L |
| allcell | ZC3H12A | 1.58E-29 | 1.422136812 | 0.608 | 0.472 | 3.11E-25 | Lep-L |
| allcell | ITGA5 | 2.07E-18 | 1.381866266 | 0.377 | 0.246 | 4.07E-14 | Lep-L |
| allcell | DDX3Y | 4.91E-34 | 1.355014907 | 0.494 | 0.269 | 9.66E-30 | Lep-L |
| allcell | BNIP3L | 6.97E-12 | 1.342181937 | 0.635 | 0.588 | 1.37E-07 | Lep-L |
| allcell | FKBP4 | 2.53E-47 | 1.320272538 | 0.65 | 0.42 | 4.99E-43 | Lep-L |
| allcell | ATP1A1 | 5.97E-40 | 1.299640851 | 0.747 | 0.641 | 1.17E-35 | Lep-L |
| allcell | TUBB2A | 3.86E-08 | 1.297918683 | 0.423 | 0.355 | 0.000760003 | Lep-L |
| allcell | CEBPB | 1.37E-14 | 1.297093835 | 0.538 | 0.458 | 2.70E-10 | Lep-L |
| allcell | SERPINH1 | 1.14E-33 | 1.293242419 | 0.453 | 0.251 | 2.24E-29 | Lep-L |
| allcell | HSPB1 | 3.15E-52 | 1.281881387 | 0.868 | 0.742 | 6.21E-48 | Lep-L |
| allcell | SLC25A37 | 2.29E-14 | 1.276321131 | 0.384 | 0.272 | 4.51E-10 | Lep-L |
| allcell | NEU1 | 5.21E-11 | 1.275759353 | 0.428 | 0.354 | 1.03E-06 | Lep-L |
| allcell | RILPL2 | 3.21E-33 | 1.275611187 | 0.694 | 0.62 | 6.32E-29 | Lep-L |
| allcell | PHLDA1 | 6.47E-13 | 1.263872452 | 0.294 | 0.182 | 1.27E-08 | Lep-L |
| allcell | HSPA1B | 2.61E-20 | 1.257636585 | 0.869 | 0.877 | 5.14E-16 | Lep-L |
| allcell | CCR7 | 6.70E-12 | 1.253885207 | 0.462 | 0.353 | 1.32E-07 | Lep-L |
| allcell | FCGR2A | 3.04E-19 | 1.252324918 | 0.518 | 0.403 | 5.99E-15 | Lep-L |
| allcell | PDE4DIP | 1.33E-27 | 1.246751497 | 0.553 | 0.403 | 2.61E-23 | Lep-L |
| allcell | MMP9 | 1.05E-08 | 1.236377641 | 0.358 | 0.273 | 0.000206867 | Lep-L |
| allcell | ZFP36L1 | 1.72E-24 | 1.231869548 | 0.783 | 0.742 | 3.38E-20 | Lep-L |
| allcell | AHR | 2.96E-27 | 1.224303633 | 0.701 | 0.63 | 5.84E-23 | Lep-L |
| allcell | SMAD3 | 3.57E-11 | 1.221667804 | 0.368 | 0.27 | 7.02E-07 | Lep-L |
| allcell | SOD2 | 6.41E-27 | 1.216993728 | 0.896 | 0.877 | 1.26E-22 | Lep-L |
| allcell | AMPD2 | 3.95E-08 | 1.215967317 | 0.278 | 0.205 | 0.000778137 | Lep-L |
| allcell | TIMP1 | 4.36E-28 | 1.209862433 | 0.754 | 0.649 | 8.59E-24 | Lep-L |
| allcell | CA2 | 7.42E-11 | 1.189300407 | 0.448 | 0.357 | 1.46E-06 | Lep-L |
| allcell | SAT1 | 1.91E-82 | 1.18769241 | 0.981 | 0.975 | 3.77E-78 | Lep-L |
| allcell | MFSD2A | 9.67E-08 | 1.180511122 | 0.29 | 0.217 | 0.001902586 | Lep-L |
| allcell | NRARP | 2.10E-15 | 1.159526881 | 0.593 | 0.519 | 4.14E-11 | Lep-L |
| allcell | CDC42EP3 | 1.32E-25 | 1.159514304 | 0.621 | 0.48 | 2.59E-21 | Lep-L |
| allcell | HLA-DRB5 | 9.51E-65 | 1.13852355 | 0.873 | 0.718 | 1.87E-60 | Lep-L |
| allcell | PTTG1 | 3.68E-16 | 1.128630618 | 0.513 | 0.414 | 7.24E-12 | Lep-L |
| allcell | MAFG | 4.30E-12 | 1.11800552 | 0.37 | 0.277 | 8.46E-08 | Lep-L |
| allcell | MAP2K3 | 1.15E-42 | 1.112687338 | 0.771 | 0.64 | 2.27E-38 | Lep-L |
| allcell | CAPN2 | 1.53E-19 | 1.102417224 | 0.542 | 0.431 | 3.00E-15 | Lep-L |
| allcell | B3GNT2 | 3.10E-07 | 1.08700414 | 0.323 | 0.258 | 0.006100577 | Lep-L |
| allcell | RBFOX2 | 5.17E-20 | 1.077433944 | 0.635 | 0.572 | 1.02E-15 | Lep-L |
| allcell | DOK2 | 2.73E-14 | 1.072430146 | 0.467 | 0.357 | 5.36E-10 | Lep-L |
| allcell | MAP7D1 | 2.68E-19 | 1.071378992 | 0.518 | 0.412 | 5.27E-15 | Lep-L |

| | | | | | | | |
|---|---|---|---|---|---|---|---|
| allcell | RIT1 | 8.68E-13 | 1.061718206 | 0.411 | 0.316 | 1.71E-08 | Lep-L |
| allcell | PLEKHA5 | 1.45E-09 | 1.058054976 | 0.438 | 0.359 | 2.86E-05 | Lep-L |
| allcell | ALDOA | 4.73E-17 | 1.051678128 | 0.433 | 0.304 | 9.30E-13 | Lep-L |
| allcell | RHOF | 4.53E-11 | 1.0507983 | 0.55 | 0.503 | 8.92E-07 | Lep-L |
| allcell | SLC5A3 | 1.31E-08 | 1.044361323 | 0.385 | 0.306 | 0.000257871 | Lep-L |
| allcell | NFKBIE | 2.42E-14 | 1.031677736 | 0.579 | 0.516 | 4.76E-10 | Lep-L |
| allcell | HSPD1 | 1.67E-72 | 1.028514187 | 0.961 | 0.907 | 3.29E-68 | Lep-L |
| allcell | STIP1 | 2.05E-25 | 1.010550936 | 0.63 | 0.52 | 4.04E-21 | Lep-L |
| allcell | ABCA1 | 7.90E-09 | 1.005437996 | 0.43 | 0.368 | 0.000155579 | Lep-L |
| allcell | CSGALNACT2 | 8.02E-15 | 1.002863493 | 0.591 | 0.55 | 1.58E-10 | Lep-L |
| allcell | ERGIC1 | 9.16E-08 | 1.002247281 | 0.414 | 0.367 | 0.001802582 | Lep-L |
| allcell | MRPS6 | 5.18E-18 | 0.996729319 | 0.628 | 0.56 | 1.02E-13 | Lep-L |
| allcell | INSIG1 | 2.51E-30 | 0.989538917 | 0.762 | 0.635 | 4.95E-26 | Lep-L |
| allcell | PRNP | 5.36E-10 | 0.987205167 | 0.48 | 0.413 | 1.05E-05 | Lep-L |
| allcell | DNAJB1 | 1.09E-32 | 0.980876598 | 0.917 | 0.873 | 2.14E-28 | Lep-L |
| allcell | ARHGAP21 | 5.10E-10 | 0.978549037 | 0.36 | 0.273 | 1.00E-05 | Lep-L |
| allcell | HSPA1A | 3.25E-53 | 0.97317945 | 0.986 | 0.94 | 6.40E-49 | Lep-L |
| allcell | BIN1 | 2.24E-10 | 0.966444084 | 0.382 | 0.3 | 4.42E-06 | Lep-L |
| allcell | RARA | 1.09E-12 | 0.957428306 | 0.48 | 0.389 | 2.14E-08 | Lep-L |
| allcell | PPP1R18 | 1.93E-16 | 0.956123781 | 0.635 | 0.591 | 3.79E-12 | Lep-L |
| allcell | TSPYL2 | 8.36E-14 | 0.955186343 | 0.555 | 0.448 | 1.65E-09 | Lep-L |
| allcell | SLC6A6 | 2.89E-08 | 0.947546388 | 0.387 | 0.318 | 0.000568246 | Lep-L |
| allcell | TRIP10 | 1.20E-07 | 0.934001746 | 0.343 | 0.268 | 0.00235653 | Lep-L |
| allcell | ETS2 | 1.55E-12 | 0.927058984 | 0.56 | 0.487 | 3.04E-08 | Lep-L |
| allcell | CXCL2 | 1.45E-12 | 0.925664232 | 0.435 | 0.311 | 2.85E-08 | Lep-L |
| allcell | CHORDC1 | 1.01E-24 | 0.920045994 | 0.715 | 0.628 | 1.98E-20 | Lep-L |
| allcell | CACYBP | 6.45E-35 | 0.910614684 | 0.784 | 0.709 | 1.27E-30 | Lep-L |
| allcell | CD44 | 8.37E-29 | 0.906735654 | 0.874 | 0.81 | 1.65E-24 | Lep-L |
| allcell | CD9 | 9.18E-07 | 0.902648865 | 0.338 | 0.271 | 0.018064765 | Lep-L |
| allcell | TXNRD1 | 7.95E-10 | 0.9021268 | 0.598 | 0.601 | 1.57E-05 | Lep-L |
| allcell | TNFAIP3 | 3.13E-19 | 0.897691684 | 0.805 | 0.767 | 6.17E-15 | Lep-L |
| allcell | METRNL | 1.71E-15 | 0.891417196 | 0.684 | 0.654 | 3.37E-11 | Lep-L |
| allcell | TCP1 | 4.66E-33 | 0.889908818 | 0.759 | 0.679 | 9.18E-29 | Lep-L |
| allcell | ANPEP | 7.98E-07 | 0.883956087 | 0.389 | 0.32 | 0.015705038 | Lep-L |
| allcell | HSPE1 | 7.25E-37 | 0.879810904 | 0.93 | 0.927 | 1.43E-32 | Lep-L |
| allcell | ITGAV | 1.72E-08 | 0.87976671 | 0.306 | 0.22 | 0.000338215 | Lep-L |
| allcell | SDCBP | 6.27E-28 | 0.879279406 | 0.796 | 0.764 | 1.23E-23 | Lep-L |
| allcell | GPAT3 | 1.20E-22 | 0.869877657 | 0.594 | 0.438 | 2.36E-18 | Lep-L |
| allcell | HSP90AB1 | 2.79E-63 | 0.864350649 | 0.983 | 0.973 | 5.49E-59 | Lep-L |
| allcell | SERPINB9 | 1.04E-24 | 0.855740575 | 0.846 | 0.809 | 2.04E-20 | Lep-L |
| allcell | IER3 | 1.09E-19 | 0.855477741 | 0.745 | 0.645 | 2.15E-15 | Lep-L |
| allcell | MTSS1 | 7.25E-10 | 0.84302639 | 0.331 | 0.234 | 1.43E-05 | Lep-L |
| allcell | CD55 | 2.71E-11 | 0.841202023 | 0.803 | 0.803 | 5.34E-07 | Lep-L |
| allcell | MGAT1 | 7.51E-16 | 0.837568663 | 0.684 | 0.662 | 1.48E-11 | Lep-L |
| allcell | PPIF | 2.38E-14 | 0.830654704 | 0.681 | 0.633 | 4.68E-10 | Lep-L |
| allcell | EMP3 | 9.64E-29 | 0.830045265 | 0.82 | 0.793 | 1.90E-24 | Lep-L |
| allcell | HSPH1 | 1.33E-52 | 0.828067846 | 0.922 | 0.862 | 2.62E-48 | Lep-L |
| allcell | CLEC2B | 3.86E-11 | 0.826539091 | 0.732 | 0.716 | 7.59E-07 | Lep-L |
| allcell | PNPLA8 | 6.21E-12 | 0.819575591 | 0.598 | 0.565 | 1.22E-07 | Lep-L |
| allcell | CTTNBP2NL | 1.31E-11 | 0.818499496 | 0.543 | 0.469 | 2.58E-07 | Lep-L |
| allcell | PPTC7 | 7.28E-09 | 0.81468912 | 0.448 | 0.38 | 0.000143266 | Lep-L |
| allcell | NFKBIA | 1.51E-28 | 0.809380137 | 0.93 | 0.903 | 2.97E-24 | Lep-L |
| allcell | AHNAK | 3.25E-16 | 0.80343016 | 0.754 | 0.689 | 6.39E-12 | Lep-L |
| allcell | SLC43A3 | 8.92E-07 | 0.803181333 | 0.338 | 0.271 | 0.017555663 | Lep-L |
| allcell | SQSTM1 | 1.01E-36 | 0.801464169 | 0.917 | 0.868 | 1.99E-32 | Lep-L |
| allcell | HBP1 | 9.72E-15 | 0.796347402 | 0.535 | 0.455 | 1.91E-10 | Lep-L |

| | | | | | | | |
|---|---|---|---|---|---|---|---|
| allcell | HSPA8 | 3.45E-24 | 0.795580457 | 0.952 | 0.955 | 6.79E-20 | Lep-L |
| allcell | HECA | 1.53E-10 | 0.790772348 | 0.633 | 0.572 | 3.01E-06 | Lep-L |
| allcell | PPP1R15A | 2.26E-10 | 0.786479581 | 0.935 | 0.925 | 4.45E-06 | Lep-L |
| allcell | DNAJA1 | 2.16E-27 | 0.783182812 | 0.935 | 0.949 | 4.24E-23 | Lep-L |
| allcell | NFKB2 | 6.90E-09 | 0.776515834 | 0.482 | 0.42 | 0.000135755 | Lep-L |
| allcell | MYO9B | 1.97E-13 | 0.769655863 | 0.683 | 0.642 | 3.87E-09 | Lep-L |
| allcell | UPP1 | 1.11E-11 | 0.768600084 | 0.567 | 0.499 | 2.19E-07 | Lep-L |
| allcell | UBE2R2 | 1.59E-19 | 0.766907068 | 0.664 | 0.593 | 3.13E-15 | Lep-L |
| allcell | AHSA1 | 9.89E-13 | 0.762191998 | 0.601 | 0.56 | 1.95E-08 | Lep-L |
| allcell | DEDD2 | 2.85E-12 | 0.74956275 | 0.516 | 0.436 | 5.62E-08 | Lep-L |
| allcell | FTL | 1.17E-30 | 0.746546369 | 0.988 | 0.99 | 2.31E-26 | Lep-L |
| allcell | MRPL18 | 6.30E-12 | 0.744305565 | 0.727 | 0.746 | 1.24E-07 | Lep-L |
| allcell | ATP13A3 | 4.29E-11 | 0.743416885 | 0.589 | 0.532 | 8.45E-07 | Lep-L |
| allcell | BCL2A1 | 3.94E-34 | 0.739645517 | 0.851 | 0.782 | 7.76E-30 | Lep-L |
| allcell | GLA | 1.41E-17 | 0.733447546 | 0.71 | 0.62 | 2.78E-13 | Lep-L |
| allcell | SLC38A2 | 6.91E-13 | 0.733377005 | 0.784 | 0.758 | 1.36E-08 | Lep-L |
| allcell | LGALS3 | 1.75E-10 | 0.731892866 | 0.713 | 0.67 | 3.44E-06 | Lep-L |
| allcell | BTG1 | 8.41E-21 | 0.721929803 | 0.89 | 0.896 | 1.65E-16 | Lep-L |
| allcell | SGK1 | 3.31E-09 | 0.713052633 | 0.754 | 0.712 | 6.52E-05 | Lep-L |
| allcell | ST13 | 6.44E-30 | 0.711745788 | 0.812 | 0.781 | 1.27E-25 | Lep-L |
| allcell | LMNA | 1.94E-24 | 0.709728604 | 0.837 | 0.689 | 3.82E-20 | Lep-L |
| allcell | MCL1 | 1.67E-32 | 0.707469232 | 0.907 | 0.887 | 3.29E-28 | Lep-L |
| allcell | PTGES3 | 1.21E-22 | 0.698235952 | 0.83 | 0.842 | 2.39E-18 | Lep-L |
| allcell | IER5 | 4.37E-13 | 0.697729527 | 0.827 | 0.782 | 8.59E-09 | Lep-L |
| allcell | NINJ1 | 1.79E-10 | 0.693226371 | 0.625 | 0.59 | 3.53E-06 | Lep-L |
| allcell | ANXA1 | 1.51E-16 | 0.690435026 | 0.772 | 0.664 | 2.97E-12 | Lep-L |
| allcell | CEBPD | 5.40E-11 | 0.681985243 | 0.703 | 0.622 | 1.06E-06 | Lep-L |
| allcell | ZFP36 | 7.16E-07 | 0.681218411 | 0.91 | 0.94 | 0.014093431 | Lep-L |
| allcell | CXCL16 | 4.34E-12 | 0.678111495 | 0.647 | 0.596 | 8.55E-08 | Lep-L |
| allcell | FTH1 | 3.25E-33 | 0.676790519 | 0.993 | 0.993 | 6.39E-29 | Lep-L |
| allcell | EIF4E | 5.91E-08 | 0.675501856 | 0.55 | 0.503 | 0.001164157 | Lep-L |
| allcell | MIR181A1HG | 3.43E-08 | 0.674375353 | 0.586 | 0.529 | 0.000674872 | Lep-L |
| allcell | CHD2 | 2.24E-18 | 0.667906604 | 0.815 | 0.812 | 4.41E-14 | Lep-L |
| allcell | IL10RA | 1.79E-09 | 0.667137521 | 0.649 | 0.635 | 3.52E-05 | Lep-L |
| allcell | SLC2A3 | 1.27E-16 | 0.666151401 | 0.713 | 0.598 | 2.49E-12 | Lep-L |
| allcell | SLC3A2 | 2.74E-09 | 0.660472858 | 0.706 | 0.717 | 5.38E-05 | Lep-L |
| allcell | DDAH2 | 1.42E-09 | 0.657803944 | 0.606 | 0.583 | 2.79E-05 | Lep-L |
| allcell | UBC | 9.93E-20 | 0.656678385 | 0.975 | 0.98 | 1.95E-15 | Lep-L |
| allcell | YY1 | 2.97E-10 | 0.653011772 | 0.645 | 0.64 | 5.84E-06 | Lep-L |
| allcell | ANXA5 | 3.59E-29 | 0.650665447 | 0.864 | 0.799 | 7.06E-25 | Lep-L |
| allcell | MTCH1 | 6.94E-12 | 0.644162668 | 0.662 | 0.645 | 1.37E-07 | Lep-L |
| allcell | G3BP1 | 2.11E-06 | 0.638287735 | 0.57 | 0.561 | 0.041456076 | Lep-L |
| allcell | DDIT4 | 6.45E-16 | 0.635954094 | 0.718 | 0.619 | 1.27E-11 | Lep-L |
| allcell | CDKN1A | 9.62E-07 | 0.632823356 | 0.839 | 0.829 | 0.018926484 | Lep-L |
| allcell | EEF2 | 8.73E-18 | 0.632527513 | 0.827 | 0.889 | 1.72E-13 | Lep-L |
| allcell | EZR | 1.73E-27 | 0.631940079 | 0.902 | 0.873 | 3.41E-23 | Lep-L |
| allcell | CTSZ | 1.50E-24 | 0.631558547 | 0.91 | 0.926 | 2.95E-20 | Lep-L |
| allcell | ACBD3 | 1.39E-07 | 0.624538262 | 0.457 | 0.408 | 0.002730205 | Lep-L |
| allcell | CKS2 | 4.25E-11 | 0.623012509 | 0.766 | 0.754 | 8.36E-07 | Lep-L |
| allcell | ARL8B | 3.29E-12 | 0.619443987 | 0.706 | 0.685 | 6.48E-08 | Lep-L |
| allcell | STX11 | 1.76E-11 | 0.616246464 | 0.708 | 0.643 | 3.46E-07 | Lep-L |
| allcell | IL18 | 9.38E-07 | 0.615126157 | 0.511 | 0.462 | 0.018460726 | Lep-L |
| allcell | HSP90AA1 | 2.20E-44 | 0.613811051 | 0.988 | 0.986 | 4.33E-40 | Lep-L |
| allcell | SNAP23 | 1.61E-09 | 0.604460257 | 0.679 | 0.642 | 3.18E-05 | Lep-L |
| allcell | MXD1 | 1.51E-07 | 0.585692586 | 0.674 | 0.648 | 0.002977872 | Lep-L |
| allcell | DNTTIP2 | 1.44E-09 | 0.581569544 | 0.598 | 0.537 | 2.83E-05 | Lep-L |

| | | | | | | | |
|---|---|---|---|---|---|---|---|
| allcell | FXYD5 | 2.61E-09 | 0.570013971 | 0.716 | 0.716 | 5.15E-05 | Lep-L |
| allcell | GNA13 | 1.50E-20 | 0.567046979 | 0.862 | 0.791 | 2.96E-16 | Lep-L |
| allcell | CYTIP | 9.93E-12 | 0.561265541 | 0.783 | 0.808 | 1.95E-07 | Lep-L |
| allcell | DYNC1H1 | 4.33E-08 | 0.555872727 | 0.696 | 0.689 | 0.000851862 | Lep-L |
| allcell | CHMP1B | 9.21E-07 | 0.551700695 | 0.694 | 0.693 | 0.018138499 | Lep-L |
| allcell | RUNX3 | 1.63E-09 | 0.548726107 | 0.666 | 0.604 | 3.21E-05 | Lep-L |
| allcell | IRAK2 | 3.87E-07 | 0.543978044 | 0.503 | 0.436 | 0.007621108 | Lep-L |
| allcell | ZFAND2A | 1.95E-26 | 0.541133572 | 0.723 | 0.592 | 3.84E-22 | Lep-L |
| allcell | HNRNPA0 | 4.81E-07 | 0.529584145 | 0.638 | 0.662 | 0.009460941 | Lep-L |
| allcell | BCAR3 | 2.18E-06 | 0.528593134 | 0.367 | 0.293 | 0.042956048 | Lep-L |
| allcell | CLK1 | 3.05E-10 | 0.521632469 | 0.762 | 0.757 | 6.01E-06 | Lep-L |
| allcell | G0S2 | 5.45E-09 | 0.516730517 | 0.642 | 0.569 | 0.000107343 | Lep-L |
| allcell | CD83 | 1.69E-21 | 0.510741223 | 0.971 | 0.964 | 3.33E-17 | Lep-L |
| allcell | TSPAN33 | 8.78E-09 | 0.508407783 | 0.647 | 0.597 | 0.000172777 | Lep-L |
| allcell | DNAJB6 | 7.06E-24 | 0.50371849 | 0.89 | 0.879 | 1.39E-19 | Lep-L |
| allcell | H2AFZ | 6.63E-15 | 0.502215738 | 0.89 | 0.907 | 1.30E-10 | Lep-L |
| allcell | PNRC1 | 1.32E-14 | 0.500933076 | 0.946 | 0.964 | 2.60E-10 | Lep-L |
| allcell | BEST1 | 1.01E-49 | 2.164420032 | 0.251 | 0.1 | 1.98E-45 | Lep-T |
| allcell | ALDOA1 | 3.22E-73 | 2.081525428 | 0.449 | 0.247 | 6.35E-69 | Lep-T |
| allcell | RBFOX21 | 2.82E-80 | 1.549907853 | 0.698 | 0.514 | 5.55E-76 | Lep-T |
| allcell | APOO1 | 1.66E-44 | 1.453828407 | 0.421 | 0.26 | 3.27E-40 | Lep-T |
| allcell | NLRP3 | 3.34E-25 | 1.383245177 | 0.365 | 0.252 | 6.57E-21 | Lep-T |
| allcell | NFKBIA1 | 3.39E-67 | 1.358352041 | 0.946 | 0.883 | 6.67E-63 | Lep-T |
| allcell | ZFP361 | 1.65E-40 | 1.338634752 | 0.954 | 0.926 | 3.24E-36 | Lep-T |
| allcell | HCAR3 | 1.26E-26 | 1.299584418 | 0.252 | 0.137 | 2.49E-22 | Lep-T |
| allcell | CD74 | 1.24E-131 | 1.258890798 | 0.997 | 0.998 | 2.44E-127 | Lep-T |
| allcell | IL1B | 2.87E-28 | 1.253313564 | 0.583 | 0.478 | 5.64E-24 | Lep-T |
| allcell | TNF | 2.36E-34 | 1.223416576 | 0.49 | 0.342 | 4.64E-30 | Lep-T |
| allcell | CXCL81 | 2.45E-34 | 1.190654749 | 0.567 | 0.409 | 4.83E-30 | Lep-T |
| allcell | ICAM1 | 1.30E-50 | 1.107614661 | 0.76 | 0.646 | 2.56E-46 | Lep-T |
| allcell | PNRC11 | 1.67E-140 | 1.106113716 | 0.98 | 0.953 | 3.29E-136 | Lep-T |
| allcell | BCL31 | 1.44E-46 | 1.087210703 | 0.617 | 0.466 | 2.84E-42 | Lep-T |
| allcell | CDKN1A1 | 4.63E-89 | 1.057951814 | 0.885 | 0.799 | 9.12E-85 | Lep-T |
| allcell | HLA-DQA2 | 2.29E-80 | 1.054026717 | 0.728 | 0.567 | 4.51E-76 | Lep-T |
| allcell | ITGB2-AS1 | 1.43E-18 | 1.050926911 | 0.261 | 0.166 | 2.81E-14 | Lep-T |
| allcell | EMD1 | 1.93E-34 | 1.041816465 | 0.445 | 0.297 | 3.79E-30 | Lep-T |
| allcell | LILRB21 | 1.85E-15 | 1.002964718 | 0.263 | 0.177 | 3.65E-11 | Lep-T |
| allcell | PLAUR | 2.25E-43 | 0.985188692 | 0.791 | 0.732 | 4.43E-39 | Lep-T |
| allcell | CSRNP11 | 2.73E-70 | 0.980218148 | 0.724 | 0.537 | 5.37E-66 | Lep-T |
| allcell | IL1RN | 6.93E-15 | 0.97838242 | 0.386 | 0.3 | 1.36E-10 | Lep-T |
| allcell | IER31 | 6.68E-12 | 0.977501768 | 0.682 | 0.644 | 1.31E-07 | Lep-T |
| allcell | G0S21 | 8.48E-28 | 0.964287501 | 0.656 | 0.535 | 1.67E-23 | Lep-T |
| allcell | CXCL21 | 5.49E-09 | 0.918037601 | 0.372 | 0.3 | 0.00010802 | Lep-T |
| allcell | CFP1 | 8.89E-24 | 0.898784585 | 0.38 | 0.256 | 1.75E-19 | Lep-T |
| allcell | HIST1H1E | 1.23E-12 | 0.897358368 | 0.315 | 0.233 | 2.43E-08 | Lep-T |
| allcell | CST3 | 3.84E-63 | 0.89411063 | 0.98 | 0.964 | 7.55E-59 | Lep-T |
| allcell | INSIG11 | 5.71E-32 | 0.892317554 | 0.712 | 0.616 | 1.12E-27 | Lep-T |
| allcell | ALAS1 | 1.11E-12 | 0.89035472 | 0.32 | 0.239 | 2.18E-08 | Lep-T |
| allcell | MB21D2 | 2.34E-18 | 0.889651538 | 0.384 | 0.283 | 4.61E-14 | Lep-T |
| allcell | NINJ11 | 6.72E-27 | 0.889645893 | 0.661 | 0.557 | 1.32E-22 | Lep-T |
| allcell | MMP91 | 2.49E-26 | 0.881482107 | 0.366 | 0.237 | 4.90E-22 | Lep-T |
| allcell | ATP5PB | 2.93E-26 | 0.863208749 | 0.561 | 0.46 | 5.76E-22 | Lep-T |
| allcell | LMNA1 | 4.74E-67 | 0.842295879 | 0.816 | 0.647 | 9.33E-63 | Lep-T |
| allcell | MT2A | 4.78E-14 | 0.839565925 | 0.625 | 0.559 | 9.41E-10 | Lep-T |
| allcell | LPAR6 | 3.91E-09 | 0.83922324 | 0.272 | 0.207 | 7.69E-05 | Lep-T |
| allcell | AVPI11 | 7.13E-14 | 0.826751332 | 0.378 | 0.296 | 1.40E-09 | Lep-T |

| | | | | | | | |
|---|---|---|---|---|---|---|---|
| allcell | SERPING1 | 1.56E-10 | 0.822610607 | 0.326 | 0.256 | 3.08E-06 | Lep-T |
| allcell | IFI27 | 1.79E-06 | 0.820809685 | 0.335 | 0.285 | 0.035185214 | Lep-T |
| allcell | DUSP1 | 2.41E-37 | 0.806113902 | 0.946 | 0.908 | 4.75E-33 | Lep-T |
| allcell | OTUB1 | 2.77E-12 | 0.79839109 | 0.4 | 0.326 | 5.46E-08 | Lep-T |
| allcell | CD1C | 3.28E-24 | 0.796702831 | 0.401 | 0.269 | 6.46E-20 | Lep-T |
| allcell | NUDT21 | 1.50E-11 | 0.786199788 | 0.274 | 0.199 | 2.96E-07 | Lep-T |
| allcell | TNFAIP31 | 2.56E-55 | 0.78003931 | 0.837 | 0.734 | 5.05E-51 | Lep-T |
| allcell | TIAM1 | 3.71E-08 | 0.773002244 | 0.363 | 0.301 | 0.000730958 | Lep-T |
| allcell | SLC2A31 | 9.22E-28 | 0.767494239 | 0.67 | 0.579 | 1.82E-23 | Lep-T |
| allcell | CD1E | 9.59E-26 | 0.759904517 | 0.362 | 0.226 | 1.89E-21 | Lep-T |
| allcell | SYNGR2 | 8.85E-34 | 0.753965513 | 0.783 | 0.715 | 1.74E-29 | Lep-T |
| allcell | ZFP36L11 | 1.62E-63 | 0.750487685 | 0.832 | 0.7 | 3.19E-59 | Lep-T |
| allcell | CDC42EP31 | 3.33E-41 | 0.741088205 | 0.604 | 0.437 | 6.55E-37 | Lep-T |
| allcell | PSAP | 9.04E-67 | 0.738397166 | 0.877 | 0.804 | 1.78E-62 | Lep-T |
| allcell | GBP5 | 5.75E-16 | 0.72854351 | 0.365 | 0.27 | 1.13E-11 | Lep-T |
| allcell | MAP2K31 | 1.74E-38 | 0.722301995 | 0.727 | 0.616 | 3.42E-34 | Lep-T |
| allcell | P2RX7 | 1.36E-11 | 0.717363469 | 0.355 | 0.27 | 2.67E-07 | Lep-T |
| allcell | TPRA11 | 1.81E-11 | 0.713996831 | 0.379 | 0.298 | 3.56E-07 | Lep-T |
| allcell | GRINA | 3.12E-15 | 0.707592643 | 0.511 | 0.427 | 6.14E-11 | Lep-T |
| allcell | IMPDH2 | 2.21E-16 | 0.703458808 | 0.357 | 0.26 | 4.36E-12 | Lep-T |
| allcell | GNB2 | 2.53E-30 | 0.699901782 | 0.714 | 0.636 | 4.99E-26 | Lep-T |
| allcell | GEM1 | 2.89E-09 | 0.699481052 | 0.43 | 0.367 | 5.69E-05 | Lep-T |
| allcell | TIMP11 | 2.67E-29 | 0.698801474 | 0.738 | 0.618 | 5.26E-25 | Lep-T |
| allcell | PLEKHA7 | 8.94E-13 | 0.697189374 | 0.302 | 0.219 | 1.76E-08 | Lep-T |
| allcell | TSPAN331 | 6.29E-26 | 0.697084258 | 0.67 | 0.565 | 1.24E-21 | Lep-T |
| allcell | ERGIC3 | 1.92E-15 | 0.688988072 | 0.443 | 0.353 | 3.78E-11 | Lep-T |
| allcell | SESTD11 | 5.62E-08 | 0.679544023 | 0.364 | 0.3 | 0.001106607 | Lep-T |
| allcell | SAT11 | 3.98E-39 | 0.678837369 | 0.981 | 0.973 | 7.84E-35 | Lep-T |
| allcell | CS | 1.68E-09 | 0.676248906 | 0.318 | 0.248 | 3.30E-05 | Lep-T |
| allcell | GYPC | 8.71E-15 | 0.664389699 | 0.474 | 0.377 | 1.71E-10 | Lep-T |
| allcell | GRASP | 2.23E-19 | 0.659716466 | 0.674 | 0.63 | 4.39E-15 | Lep-T |
| allcell | AHNAK1 | 2.23E-42 | 0.658853483 | 0.767 | 0.657 | 4.38E-38 | Lep-T |
| allcell | VEGFA1 | 1.60E-13 | 0.653599522 | 0.43 | 0.35 | 3.15E-09 | Lep-T |
| allcell | BAG31 | 2.01E-13 | 0.652414527 | 0.626 | 0.537 | 3.95E-09 | Lep-T |
| allcell | MGLL | 1.43E-20 | 0.651630444 | 0.388 | 0.263 | 2.82E-16 | Lep-T |
| allcell | DDX3Y1 | 1.59E-17 | 0.648795989 | 0.365 | 0.257 | 3.13E-13 | Lep-T |
| allcell | ZYX | 4.24E-22 | 0.648232289 | 0.666 | 0.579 | 8.35E-18 | Lep-T |
| allcell | CYP2S1 | 1.35E-15 | 0.647804683 | 0.387 | 0.29 | 2.66E-11 | Lep-T |
| allcell | EEF21 | 3.52E-74 | 0.645259758 | 0.906 | 0.868 | 6.93E-70 | Lep-T |
| allcell | UBALD21 | 3.16E-08 | 0.639961919 | 0.499 | 0.448 | 0.0006222 | Lep-T |
| allcell | RASSF4 | 2.95E-23 | 0.637107569 | 0.577 | 0.462 | 5.81E-19 | Lep-T |
| allcell | TRAF1 | 1.16E-29 | 0.634991145 | 0.713 | 0.594 | 2.28E-25 | Lep-T |
| allcell | SLAMF8 | 4.26E-25 | 0.634610361 | 0.561 | 0.434 | 8.38E-21 | Lep-T |
| allcell | SQSTM11 | 4.86E-63 | 0.631513521 | 0.917 | 0.849 | 9.57E-59 | Lep-T |
| allcell | BCL2A11 | 7.06E-30 | 0.628182317 | 0.842 | 0.762 | 1.39E-25 | Lep-T |
| allcell | PPP2R1A | 3.44E-10 | 0.625422588 | 0.354 | 0.286 | 6.78E-06 | Lep-T |
| allcell | HACD4 | 6.65E-08 | 0.620530927 | 0.33 | 0.27 | 0.001308162 | Lep-T |
| allcell | CRTAP1 | 6.85E-12 | 0.616333679 | 0.436 | 0.366 | 1.35E-07 | Lep-T |
| allcell | BIRC3 | 1.31E-31 | 0.613973978 | 0.765 | 0.638 | 2.58E-27 | Lep-T |
| allcell | TACSTD2 | 1.84E-07 | 0.605451875 | 0.256 | 0.196 | 0.003626784 | Lep-T |
| allcell | BHLHE40 | 2.27E-25 | 0.604591153 | 0.786 | 0.737 | 4.46E-21 | Lep-T |
| allcell | NFKBIE1 | 2.94E-15 | 0.599777365 | 0.572 | 0.497 | 5.80E-11 | Lep-T |
| allcell | PIM3 | 3.70E-17 | 0.597727871 | 0.746 | 0.723 | 7.29E-13 | Lep-T |
| allcell | FCGR2A1 | 1.98E-10 | 0.59771746 | 0.467 | 0.388 | 3.89E-06 | Lep-T |
| allcell | ZC3H12A1 | 9.19E-25 | 0.59390606 | 0.569 | 0.442 | 1.81E-20 | Lep-T |
| allcell | ACSL5 | 7.72E-11 | 0.59186145 | 0.369 | 0.292 | 1.52E-06 | Lep-T |

| | | | | | | | |
|---|---|---|---|---|---|---|---|
| allcell | SEC13 | 8.67E-15 | 0.584462292 | 0.434 | 0.342 | 1.71E-10 | Lep-T |
| allcell | CD831 | 2.77E-72 | 0.577414997 | 0.979 | 0.957 | 5.45E-68 | Lep-T |
| allcell | DAZAP2 | 2.34E-16 | 0.575794059 | 0.606 | 0.523 | 4.61E-12 | Lep-T |
| allcell | PLEK | 4.03E-36 | 0.572736628 | 0.909 | 0.881 | 7.93E-32 | Lep-T |
| allcell | NUDT3 | 5.92E-07 | 0.568961972 | 0.276 | 0.22 | 0.011645442 | Lep-T |
| allcell | CNBP | 1.34E-23 | 0.567627257 | 0.794 | 0.749 | 2.64E-19 | Lep-T |
| allcell | CAPN21 | 5.67E-18 | 0.566553492 | 0.509 | 0.408 | 1.12E-13 | Lep-T |
| allcell | RTN1 | 9.82E-17 | 0.566442436 | 0.422 | 0.313 | 1.93E-12 | Lep-T |
| allcell | DUSP2 | 4.71E-24 | 0.566310909 | 0.846 | 0.768 | 9.28E-20 | Lep-T |
| allcell | CD40 | 1.85E-22 | 0.566266959 | 0.663 | 0.568 | 3.64E-18 | Lep-T |
| allcell | CD52 | 3.59E-31 | 0.56526891 | 0.632 | 0.475 | 7.06E-27 | Lep-T |
| allcell | PPIF1 | 4.51E-23 | 0.56367698 | 0.701 | 0.604 | 8.88E-19 | Lep-T |
| allcell | AFDN | 1.72E-10 | 0.562247824 | 0.367 | 0.288 | 3.38E-06 | Lep-T |
| allcell | PPDPF | 2.01E-07 | 0.55982233 | 0.693 | 0.661 | 0.003957355 | Lep-T |
| allcell | CD441 | 3.73E-32 | 0.557873948 | 0.86 | 0.794 | 7.35E-28 | Lep-T |
| allcell | PPP1R15A1 | 1.32E-39 | 0.549867015 | 0.935 | 0.921 | 2.60E-35 | Lep-T |
| allcell | LGALS2 | 5.29E-45 | 0.548633953 | 0.826 | 0.67 | 1.04E-40 | Lep-T |
| allcell | PEA15 | 3.82E-20 | 0.548076407 | 0.663 | 0.578 | 7.52E-16 | Lep-T |
| allcell | MIR155HG | 1.30E-25 | 0.540117781 | 0.6 | 0.466 | 2.55E-21 | Lep-T |
| allcell | POLR2G | 8.43E-08 | 0.538452028 | 0.382 | 0.323 | 0.001658614 | Lep-T |
| allcell | UNC119 | 1.36E-07 | 0.530198939 | 0.514 | 0.472 | 0.00268569 | Lep-T |
| allcell | TRIM28 | 1.26E-06 | 0.528897129 | 0.437 | 0.388 | 0.024858581 | Lep-T |
| allcell | OXA1L | 2.40E-08 | 0.528375003 | 0.414 | 0.351 | 0.000472038 | Lep-T |
| allcell | TAGLN2 | 6.78E-31 | 0.52639414 | 0.913 | 0.89 | 1.34E-26 | Lep-T |
| allcell | HSPA61 | 6.97E-08 | 0.521975266 | 0.489 | 0.415 | 0.001372048 | Lep-T |
| allcell | RARA1 | 1.18E-10 | 0.519376289 | 0.448 | 0.373 | 2.33E-06 | Lep-T |
| allcell | ID2 | 2.76E-10 | 0.513315827 | 0.736 | 0.689 | 5.43E-06 | Lep-T |
| allcell | LY6E | 1.17E-07 | 0.513242123 | 0.524 | 0.472 | 0.00230632 | Lep-T |
| allcell | FTH11 | 2.55E-31 | 0.508200977 | 0.991 | 0.995 | 5.02E-27 | Lep-T |
| allcell | FMNL2 | 1.66E-12 | 0.507663051 | 0.64 | 0.554 | 3.27E-08 | Lep-T |
| allcell | RPN1 | 5.47E-07 | 0.504982477 | 0.415 | 0.363 | 0.010769802 | Lep-T |
| allcell | IRAK21 | 9.30E-13 | 0.502986601 | 0.501 | 0.412 | 1.83E-08 | Lep-T |
| allcell | GZMB | 2.58E-47 | 2.188393602 | 0.312 | 0.15 | 5.07E-43 | Mari |
| allcell | LRRC26 | 5.30E-54 | 2.143724057 | 0.26 | 0.093 | 1.04E-49 | Mari |
| allcell | PLAC8 | 6.28E-72 | 2.119300691 | 0.455 | 0.217 | 1.24E-67 | Mari |
| allcell | CLEC4C | 1.21E-56 | 2.033463385 | 0.298 | 0.115 | 2.39E-52 | Mari |
| allcell | SCT | 1.42E-46 | 1.998899251 | 0.28 | 0.117 | 2.80E-42 | Mari |
| allcell | JCHAIN | 1.37E-41 | 1.907962649 | 0.304 | 0.148 | 2.70E-37 | Mari |
| allcell | LAMP5 | 1.77E-46 | 1.885036237 | 0.25 | 0.094 | 3.48E-42 | Mari |
| allcell | CLIC3 | 1.67E-42 | 1.862493426 | 0.338 | 0.171 | 3.28E-38 | Mari |
| allcell | MZB1 | 3.99E-44 | 1.84444628 | 0.257 | 0.103 | 7.86E-40 | Mari |
| allcell | DERL3 | 3.95E-43 | 1.835817777 | 0.25 | 0.099 | 7.77E-39 | Mari |
| allcell | IRF7 | 5.92E-56 | 1.724083459 | 0.728 | 0.522 | 1.17E-51 | Mari |
| allcell | KRT161 | 4.03E-43 | 1.709669391 | 0.309 | 0.151 | 7.93E-39 | Mari |
| allcell | MS4A4A | 9.25E-40 | 1.67143548 | 0.294 | 0.14 | 1.82E-35 | Mari |
| allcell | CIB2 | 2.86E-36 | 1.662634115 | 0.281 | 0.135 | 5.63E-32 | Mari |
| allcell | HIGD1A | 2.70E-78 | 1.659402592 | 0.597 | 0.345 | 5.31E-74 | Mari |
| allcell | LILRA4 | 2.31E-40 | 1.640300834 | 0.303 | 0.145 | 4.55E-36 | Mari |
| allcell | SELL | 6.18E-70 | 1.640129888 | 0.462 | 0.222 | 1.22E-65 | Mari |
| allcell | MGAT4A | 2.93E-37 | 1.569051669 | 0.305 | 0.153 | 5.77E-33 | Mari |
| allcell | ANK3 | 8.46E-39 | 1.531952887 | 0.301 | 0.145 | 1.67E-34 | Mari |
| allcell | GAS6 | 7.47E-119 | 1.531311003 | 0.765 | 0.475 | 1.47E-114 | Mari |
| allcell | ITM2C | 1.45E-28 | 1.52714356 | 0.431 | 0.296 | 2.86E-24 | Mari |
| allcell | PPP1R14B | 1.95E-44 | 1.525681964 | 0.825 | 0.727 | 3.84E-40 | Mari |
| allcell | SNRPN | 6.40E-35 | 1.479553397 | 0.408 | 0.252 | 1.26E-30 | Mari |
| allcell | ST6GALNAC4 | 1.40E-31 | 1.477194763 | 0.299 | 0.16 | 2.76E-27 | Mari |

| | | | | | | | |
|---|---|---|---|---|---|---|---|
| allcell | PCED1B-AS1 | 8.03E-34 | 1.475955022 | 0.299 | 0.154 | 1.58E-29 | Mari |
| allcell | CD36 | 4.92E-12 | 1.472966077 | 0.259 | 0.178 | 9.69E-08 | Mari |
| allcell | SDHAF3 | 5.57E-45 | 1.470816536 | 0.358 | 0.18 | 1.10E-40 | Mari |
| allcell | PPP1R14A1 | 1.75E-33 | 1.433830335 | 0.308 | 0.158 | 3.45E-29 | Mari |
| allcell | CCDC50 | 3.99E-39 | 1.410790039 | 0.615 | 0.464 | 7.86E-35 | Mari |
| allcell | ISCA2 | 2.64E-39 | 1.398708754 | 0.339 | 0.182 | 5.19E-35 | Mari |
| allcell | HLA-DRB51 | 2.95E-37 | 1.397952438 | 0.761 | 0.729 | 5.81E-33 | Mari |
| allcell | LIME1 | 4.46E-32 | 1.397395668 | 0.256 | 0.121 | 8.78E-28 | Mari |
| allcell | MSRB2 | 4.04E-49 | 1.39063268 | 0.37 | 0.184 | 7.96E-45 | Mari |
| allcell | PLAAT3 | 3.77E-27 | 1.375370248 | 0.356 | 0.224 | 7.43E-23 | Mari |
| allcell | PUS10 | 5.27E-08 | 1.360985577 | 0.323 | 0.262 | 0.001037835 | Mari |
| allcell | S100A81 | 9.20E-44 | 1.355529685 | 0.497 | 0.312 | 1.81E-39 | Mari |
| allcell | NUDT22 | 3.12E-57 | 1.350529558 | 0.533 | 0.328 | 6.13E-53 | Mari |
| allcell | S100A91 | 1.19E-32 | 1.337608325 | 0.535 | 0.376 | 2.35E-28 | Mari |
| allcell | SUSD1 | 1.62E-23 | 1.330992761 | 0.37 | 0.247 | 3.18E-19 | Mari |
| allcell | COBLL1 | 6.58E-41 | 1.317248376 | 0.278 | 0.12 | 1.29E-36 | Mari |
| allcell | AOPEP1 | 3.04E-13 | 1.30911337 | 0.258 | 0.175 | 5.98E-09 | Mari |
| allcell | VPS25 | 2.95E-43 | 1.302787371 | 0.359 | 0.185 | 5.81E-39 | Mari |
| allcell | ODC1 | 2.95E-31 | 1.297082036 | 0.549 | 0.41 | 5.81E-27 | Mari |
| allcell | POLE4 | 2.39E-59 | 1.293301003 | 0.516 | 0.308 | 4.71E-55 | Mari |
| allcell | OCIAD2 | 2.10E-24 | 1.268361787 | 0.264 | 0.145 | 4.13E-20 | Mari |
| allcell | TSPAN131 | 2.31E-39 | 1.26320421 | 0.467 | 0.291 | 4.54E-35 | Mari |
| allcell | GNG10 | 2.90E-82 | 1.244900872 | 0.713 | 0.511 | 5.70E-78 | Mari |
| allcell | ALOX5AP | 9.71E-14 | 1.236519301 | 0.436 | 0.371 | 1.91E-09 | Mari |
| allcell | PTPMT1 | 4.21E-38 | 1.235565915 | 0.42 | 0.257 | 8.30E-34 | Mari |
| allcell | ZBTB18 | 7.27E-14 | 1.234933767 | 0.29 | 0.205 | 1.43E-09 | Mari |
| allcell | RWDD3 | 4.04E-24 | 1.226240258 | 0.259 | 0.144 | 7.96E-20 | Mari |
| allcell | CD69 | 1.40E-40 | 1.220850736 | 0.574 | 0.404 | 2.76E-36 | Mari |
| allcell | CCDC186 | 4.42E-26 | 1.219180964 | 0.426 | 0.292 | 8.71E-22 | Mari |
| allcell | ITGAE | 1.93E-34 | 1.218122572 | 0.412 | 0.254 | 3.81E-30 | Mari |
| allcell | HINT2 | 3.68E-41 | 1.211419097 | 0.462 | 0.292 | 7.25E-37 | Mari |
| allcell | CCL4L2 | 2.87E-07 | 1.196164417 | 0.511 | 0.468 | 0.005648909 | Mari |
| allcell | TATDN3 | 1.15E-28 | 1.19319453 | 0.322 | 0.188 | 2.27E-24 | Mari |
| allcell | VEGFB | 6.17E-41 | 1.174580503 | 0.651 | 0.481 | 1.21E-36 | Mari |
| allcell | RASGRP2 | 1.95E-26 | 1.167709075 | 0.321 | 0.19 | 3.84E-22 | Mari |
| allcell | DCPS | 2.29E-30 | 1.151308942 | 0.373 | 0.229 | 4.51E-26 | Mari |
| allcell | SERPINF1 | 9.57E-75 | 1.146067411 | 0.851 | 0.684 | 1.88E-70 | Mari |
| allcell | MRPL58 | 1.78E-30 | 1.145659553 | 0.312 | 0.171 | 3.50E-26 | Mari |
| allcell | MTLN | 9.34E-26 | 1.129288617 | 0.327 | 0.201 | 1.84E-21 | Mari |
| allcell | NAE1 | 1.89E-20 | 1.118190452 | 0.253 | 0.148 | 3.73E-16 | Mari |
| allcell | HIST1H1C | 2.30E-10 | 1.115959098 | 0.325 | 0.251 | 4.54E-06 | Mari |
| allcell | RNASEH2C | 2.01E-50 | 1.114591948 | 0.533 | 0.336 | 3.96E-46 | Mari |
| allcell | SPCS1 | 2.61E-68 | 1.111755035 | 0.861 | 0.721 | 5.14E-64 | Mari |
| allcell | GCA | 2.50E-74 | 1.111116012 | 0.697 | 0.484 | 4.92E-70 | Mari |
| allcell | SVIP | 2.15E-19 | 1.106114708 | 0.274 | 0.169 | 4.23E-15 | Mari |
| allcell | ACTR6 | 1.89E-21 | 1.104383712 | 0.298 | 0.185 | 3.72E-17 | Mari |
| allcell | PLD4 | 4.42E-27 | 1.102544659 | 0.374 | 0.232 | 8.69E-23 | Mari |
| allcell | NSMCE3 | 1.29E-33 | 1.101193479 | 0.437 | 0.275 | 2.54E-29 | Mari |
| allcell | IDH3A | 1.22E-33 | 1.100798856 | 0.483 | 0.325 | 2.41E-29 | Mari |
| allcell | SEC61G | 2.49E-111 | 1.096225256 | 0.899 | 0.731 | 4.90E-107 | Mari |
| allcell | MEF2C | 8.18E-77 | 1.092499251 | 0.776 | 0.556 | 1.61E-72 | Mari |
| allcell | TMEM107 | 3.99E-14 | 1.091540468 | 0.321 | 0.23 | 7.85E-10 | Mari |
| allcell | NDUFC2 | 1.39E-113 | 1.088566111 | 0.851 | 0.605 | 2.73E-109 | Mari |
| allcell | COA5 | 6.84E-17 | 1.082204272 | 0.285 | 0.19 | 1.35E-12 | Mari |
| allcell | HIST1H1D | 3.85E-14 | 1.081803037 | 0.283 | 0.194 | 7.58E-10 | Mari |
| allcell | COMMD8 | 2.52E-46 | 1.077122161 | 0.478 | 0.289 | 4.96E-42 | Mari |

| | | | | | | | |
|---|---|---|---|---|---|---|---|
| allcell | MS4A6A | 3.64E-40 | 1.067417165 | 0.65 | 0.512 | 7.17E-36 | Mari |
| allcell | HRH2 | 5.13E-15 | 1.067313246 | 0.269 | 0.178 | 1.01E-10 | Mari |
| allcell | NDUFAF8 | 5.29E-44 | 1.061014435 | 0.501 | 0.32 | 1.04E-39 | Mari |
| allcell | CHCHD1 | 2.80E-40 | 1.053986302 | 0.471 | 0.303 | 5.51E-36 | Mari |
| allcell | CYBC1 | 7.98E-47 | 1.052742923 | 0.538 | 0.359 | 1.57E-42 | Mari |
| allcell | MX1 | 7.57E-33 | 1.050643665 | 0.482 | 0.318 | 1.49E-28 | Mari |
| allcell | METTL23 | 5.16E-36 | 1.047816628 | 0.449 | 0.285 | 1.02E-31 | Mari |
| allcell | ZNF511 | 4.37E-30 | 1.047001042 | 0.387 | 0.243 | 8.60E-26 | Mari |
| allcell | CNPY3 | 5.70E-44 | 1.046510735 | 0.656 | 0.487 | 1.12E-39 | Mari |
| allcell | SNRNP25 | 8.41E-50 | 1.044708567 | 0.597 | 0.405 | 1.65E-45 | Mari |
| allcell | HIST1H2AC | 2.77E-11 | 1.04454039 | 0.343 | 0.264 | 5.46E-07 | Mari |
| allcell | JKAMP | 3.83E-28 | 1.040067774 | 0.339 | 0.202 | 7.54E-24 | Mari |
| allcell | SRP9 | 8.09E-86 | 1.037861945 | 0.755 | 0.534 | 1.59E-81 | Mari |
| allcell | TMCO1 | 6.51E-46 | 1.035630321 | 0.574 | 0.388 | 1.28E-41 | Mari |
| allcell | POLR2I | 1.49E-42 | 1.03422983 | 0.516 | 0.338 | 2.93E-38 | Mari |
| allcell | CTSC | 2.48E-39 | 1.034224813 | 0.766 | 0.624 | 4.89E-35 | Mari |
| allcell | PAXX | 6.33E-51 | 1.033898075 | 0.736 | 0.562 | 1.25E-46 | Mari |
| allcell | LAIR1 | 5.41E-19 | 1.031773092 | 0.286 | 0.177 | 1.06E-14 | Mari |
| allcell | MYL12A | 2.77E-152 | 1.024034942 | 0.959 | 0.903 | 5.45E-148 | Mari |
| allcell | MRPS18C | 1.53E-36 | 1.022596578 | 0.462 | 0.296 | 3.01E-32 | Mari |
| allcell | RAC2 | 9.35E-25 | 1.021901403 | 0.47 | 0.34 | 1.84E-20 | Mari |
| allcell | FCER1G | 1.81E-61 | 1.021772643 | 0.835 | 0.726 | 3.56E-57 | Mari |
| allcell | SELENOS | 1.48E-34 | 1.021513325 | 0.663 | 0.527 | 2.92E-30 | Mari |
| allcell | FAM207A | 1.98E-18 | 1.018872362 | 0.301 | 0.196 | 3.89E-14 | Mari |
| allcell | SPCS3 | 3.27E-80 | 1.015434615 | 0.795 | 0.578 | 6.44E-76 | Mari |
| allcell | CYB561D2 | 3.53E-23 | 1.014898552 | 0.309 | 0.189 | 6.95E-19 | Mari |
| allcell | AGPAT5 | 1.78E-13 | 1.014776842 | 0.264 | 0.177 | 3.50E-09 | Mari |
| allcell | USF2 | 5.62E-24 | 1.012739173 | 0.473 | 0.353 | 1.11E-19 | Mari |
| allcell | TMEM141 | 2.26E-34 | 1.00739516 | 0.478 | 0.321 | 4.45E-30 | Mari |
| allcell | RHEX | 3.88E-27 | 1.006886746 | 0.427 | 0.282 | 7.63E-23 | Mari |
| allcell | FAM89B | 1.07E-50 | 1.006452656 | 0.607 | 0.414 | 2.10E-46 | Mari |
| allcell | MRPS23 | 1.97E-33 | 1.003947149 | 0.418 | 0.26 | 3.88E-29 | Mari |
| allcell | LYSMD2 | 9.11E-42 | 1.002809445 | 0.589 | 0.418 | 1.79E-37 | Mari |
| allcell | ZDHHC171 | 1.45E-25 | 1.000335562 | 0.441 | 0.307 | 2.84E-21 | Mari |
| allcell | PARK7 | 9.04E-126 | 0.997465634 | 0.931 | 0.809 | 1.78E-121 | Mari |
| allcell | BORCS8 | 4.30E-18 | 0.996378507 | 0.273 | 0.173 | 8.47E-14 | Mari |
| allcell | PAIP1 | 5.62E-24 | 0.993241369 | 0.39 | 0.264 | 1.11E-19 | Mari |
| allcell | RDH14 | 4.16E-20 | 0.992429186 | 0.252 | 0.146 | 8.18E-16 | Mari |
| allcell | PPP2R5C | 7.44E-30 | 0.991610179 | 0.642 | 0.496 | 1.46E-25 | Mari |
| allcell | NUCB21 | 2.03E-15 | 0.987877154 | 0.317 | 0.219 | 4.00E-11 | Mari |
| allcell | SOCS11 | 2.88E-41 | 0.982880007 | 0.646 | 0.465 | 5.67E-37 | Mari |
| allcell | CYB561A3 | 1.21E-17 | 0.98275532 | 0.381 | 0.279 | 2.38E-13 | Mari |
| allcell | 2-Mar | 4.96E-34 | 0.982566481 | 0.488 | 0.334 | 9.77E-30 | Mari |
| allcell | ARRDC3 | 8.62E-20 | 0.982083168 | 0.47 | 0.362 | 1.70E-15 | Mari |
| allcell | NUBP2 | 7.48E-27 | 0.977863624 | 0.386 | 0.253 | 1.47E-22 | Mari |
| allcell | CCS | 6.99E-25 | 0.976547819 | 0.398 | 0.267 | 1.38E-20 | Mari |
| allcell | UCHL3 | 6.33E-49 | 0.97610765 | 0.56 | 0.354 | 1.25E-44 | Mari |
| allcell | ZBTB8OS | 2.20E-30 | 0.971967847 | 0.423 | 0.275 | 4.32E-26 | Mari |
| allcell | ALG5 | 1.56E-37 | 0.971808615 | 0.474 | 0.306 | 3.08E-33 | Mari |
| allcell | TMX1 | 1.37E-34 | 0.971109862 | 0.491 | 0.336 | 2.69E-30 | Mari |
| allcell | UBLCP1 | 1.30E-17 | 0.969435094 | 0.322 | 0.219 | 2.55E-13 | Mari |
| allcell | RASD1 | 1.75E-14 | 0.968164618 | 0.262 | 0.171 | 3.44E-10 | Mari |
| allcell | TRAPPC1 | 1.23E-85 | 0.967681204 | 0.784 | 0.552 | 2.43E-81 | Mari |
| allcell | POP4 | 2.99E-23 | 0.967254035 | 0.343 | 0.22 | 5.88E-19 | Mari |
| allcell | PHB | 1.77E-48 | 0.966903313 | 0.698 | 0.525 | 3.48E-44 | Mari |
| allcell | ARL2 | 6.12E-32 | 0.963884119 | 0.431 | 0.283 | 1.21E-27 | Mari |

| | | | | | | | |
|---|---|---|---|---|---|---|---|
| allcell | TIMM8B | 3.43E-59 | 0.962654455 | 0.651 | 0.444 | 6.76E-55 | Mari |
| allcell | THYN1 | 7.88E-33 | 0.959326589 | 0.418 | 0.262 | 1.55E-28 | Mari |
| allcell | DTWD1 | 1.17E-17 | 0.956044832 | 0.276 | 0.174 | 2.30E-13 | Mari |
| allcell | RBX1 | 8.23E-120 | 0.953685681 | 0.893 | 0.723 | 1.62E-115 | Mari |
| allcell | GPR65 | 1.27E-37 | 0.953085449 | 0.623 | 0.441 | 2.49E-33 | Mari |
| allcell | SNRNP27 | 1.64E-13 | 0.952808403 | 0.27 | 0.188 | 3.22E-09 | Mari |
| allcell | GALK1 | 2.18E-23 | 0.951587461 | 0.38 | 0.256 | 4.29E-19 | Mari |
| allcell | COPZ1 | 1.72E-46 | 0.95116401 | 0.561 | 0.378 | 3.38E-42 | Mari |
| allcell | CZIB | 4.99E-20 | 0.948191392 | 0.318 | 0.207 | 9.82E-16 | Mari |
| allcell | PLP2 | 2.15E-30 | 0.94804525 | 0.663 | 0.54 | 4.24E-26 | Mari |
| allcell | MRPL20 | 7.44E-65 | 0.946512316 | 0.738 | 0.535 | 1.47E-60 | Mari |
| allcell | MRPL19 | 2.08E-18 | 0.945853035 | 0.32 | 0.216 | 4.09E-14 | Mari |
| allcell | TOMM5 | 2.80E-59 | 0.945563121 | 0.676 | 0.467 | 5.50E-55 | Mari |
| allcell | UNC50 | 4.84E-22 | 0.943942372 | 0.33 | 0.211 | 9.52E-18 | Mari |
| allcell | GTF3C6 | 3.09E-52 | 0.943119413 | 0.646 | 0.453 | 6.08E-48 | Mari |
| allcell | NDUFAB1 | 1.44E-81 | 0.942194091 | 0.773 | 0.546 | 2.83E-77 | Mari |
| allcell | TMEM59 | 4.14E-70 | 0.940056414 | 0.801 | 0.62 | 8.14E-66 | Mari |
| allcell | ERP29 | 1.29E-43 | 0.939004185 | 0.823 | 0.71 | 2.54E-39 | Mari |
| allcell | CYBA | 1.37E-148 | 0.938086769 | 0.975 | 0.939 | 2.71E-144 | Mari |
| allcell | RGS18 | 1.18E-25 | 0.936962825 | 0.376 | 0.241 | 2.33E-21 | Mari |
| allcell | PYURF | 1.42E-43 | 0.93366405 | 0.548 | 0.371 | 2.80E-39 | Mari |
| allcell | TXNDC17 | 3.06E-66 | 0.933463845 | 0.689 | 0.473 | 6.02E-62 | Mari |
| allcell | BST2 | 1.37E-94 | 0.933167643 | 0.868 | 0.713 | 2.70E-90 | Mari |
| allcell | YIF1A | 5.08E-28 | 0.931182504 | 0.44 | 0.3 | 1.00E-23 | Mari |
| allcell | CDKN2D | 1.63E-34 | 0.928599425 | 0.509 | 0.34 | 3.21E-30 | Mari |
| allcell | MRPL42 | 1.26E-13 | 0.927811804 | 0.252 | 0.167 | 2.47E-09 | Mari |
| allcell | PRDX5 | 6.32E-80 | 0.925555343 | 0.817 | 0.636 | 1.24E-75 | Mari |
| allcell | TMEM35B | 1.46E-19 | 0.92450378 | 0.347 | 0.237 | 2.87E-15 | Mari |
| allcell | UQCR11 | 1.18E-138 | 0.924064706 | 0.921 | 0.775 | 2.32E-134 | Mari |
| allcell | SLC39A10 | 2.79E-33 | 0.923306722 | 0.455 | 0.294 | 5.49E-29 | Mari |
| allcell | CIAO2A | 1.71E-53 | 0.922155838 | 0.663 | 0.462 | 3.36E-49 | Mari |
| allcell | NTAN1 | 7.33E-18 | 0.92132765 | 0.323 | 0.218 | 1.44E-13 | Mari |
| allcell | LSM5 | 3.53E-65 | 0.917268132 | 0.721 | 0.519 | 6.94E-61 | Mari |
| allcell | MRPS10 | 4.01E-19 | 0.914875571 | 0.36 | 0.252 | 7.88E-15 | Mari |
| allcell | SEC11C | 2.60E-42 | 0.914744118 | 0.631 | 0.449 | 5.11E-38 | Mari |
| allcell | NDUFA9 | 5.78E-29 | 0.914086643 | 0.483 | 0.344 | 1.14E-24 | Mari |
| allcell | VAMP8 | 1.05E-108 | 0.912134339 | 0.924 | 0.842 | 2.06E-104 | Mari |
| allcell | ADI1 | 5.41E-27 | 0.909731505 | 0.486 | 0.348 | 1.06E-22 | Mari |
| allcell | MRPL13 | 3.85E-35 | 0.909691359 | 0.467 | 0.309 | 7.57E-31 | Mari |
| allcell | TRIAP1 | 7.38E-16 | 0.908513167 | 0.273 | 0.177 | 1.45E-11 | Mari |
| allcell | CRELD2 | 3.35E-31 | 0.907508687 | 0.44 | 0.281 | 6.60E-27 | Mari |
| allcell | PRELID1 | 1.19E-61 | 0.907477714 | 0.865 | 0.741 | 2.34E-57 | Mari |
| allcell | MED30 | 5.90E-26 | 0.906851533 | 0.43 | 0.289 | 1.16E-21 | Mari |
| allcell | GTF2H5 | 9.71E-35 | 0.906321152 | 0.499 | 0.339 | 1.91E-30 | Mari |
| allcell | GSDMD | 1.60E-27 | 0.905985879 | 0.463 | 0.327 | 3.16E-23 | Mari |
| allcell | RPP30 | 9.37E-16 | 0.905913525 | 0.318 | 0.22 | 1.84E-11 | Mari |
| allcell | NHP2 | 2.94E-43 | 0.905077903 | 0.618 | 0.445 | 5.79E-39 | Mari |
| allcell | SRI | 8.13E-59 | 0.905000068 | 0.725 | 0.54 | 1.60E-54 | Mari |
| allcell | MRPL57 | 1.76E-50 | 0.904618888 | 0.65 | 0.453 | 3.47E-46 | Mari |
| allcell | RBM7 | 1.23E-07 | 0.90094222 | 0.279 | 0.222 | 0.002429471 | Mari |
| allcell | BLOC1S2 | 4.23E-39 | 0.900766958 | 0.567 | 0.399 | 8.33E-35 | Mari |
| allcell | ANAPC16 | 5.99E-47 | 0.900280462 | 0.702 | 0.525 | 1.18E-42 | Mari |
| allcell | CCND3 | 3.76E-30 | 0.899318866 | 0.544 | 0.391 | 7.41E-26 | Mari |
| allcell | LSM10 | 3.57E-35 | 0.897767436 | 0.525 | 0.369 | 7.02E-31 | Mari |
| allcell | SIVA1 | 4.79E-53 | 0.897365702 | 0.722 | 0.539 | 9.42E-49 | Mari |
| allcell | PIN1 | 4.06E-35 | 0.895443875 | 0.544 | 0.394 | 7.99E-31 | Mari |

| | | | | | | | |
|---|---|---|---|---|---|---|---|
| allcell | TMEM109 | 4.00E-31 | 0.894828102 | 0.498 | 0.341 | 7.87E-27 | Mari |
| allcell | GTF3A | 3.43E-51 | 0.894815985 | 0.723 | 0.558 | 6.76E-47 | Mari |
| allcell | COA4 | 5.68E-31 | 0.89420189 | 0.488 | 0.34 | 1.12E-26 | Mari |
| allcell | NDUFB7 | 1.02E-66 | 0.893632439 | 0.744 | 0.546 | 2.02E-62 | Mari |
| allcell | ZNF428 | 6.20E-16 | 0.8933471 | 0.425 | 0.324 | 1.22E-11 | Mari |
| allcell | NCBP2AS2 | 2.93E-21 | 0.892868141 | 0.347 | 0.229 | 5.76E-17 | Mari |
| allcell | POLR3K | 4.11E-15 | 0.892836759 | 0.286 | 0.192 | 8.10E-11 | Mari |
| allcell | MAGOHB | 1.51E-13 | 0.891434598 | 0.298 | 0.21 | 2.96E-09 | Mari |
| allcell | LAMTOR1 | 3.19E-74 | 0.890294867 | 0.807 | 0.647 | 6.28E-70 | Mari |
| allcell | GTF2A2 | 7.07E-49 | 0.889911854 | 0.669 | 0.497 | 1.39E-44 | Mari |
| allcell | SMIM7 | 9.43E-34 | 0.889493717 | 0.531 | 0.375 | 1.86E-29 | Mari |
| allcell | CCDC167 | 3.21E-21 | 0.88853483 | 0.331 | 0.211 | 6.32E-17 | Mari |
| allcell | NME4 | 2.90E-30 | 0.888173586 | 0.492 | 0.342 | 5.72E-26 | Mari |
| allcell | NSMCE4A | 5.32E-25 | 0.887494537 | 0.386 | 0.253 | 1.05E-20 | Mari |
| allcell | PPA1 | 7.52E-27 | 0.887342534 | 0.795 | 0.747 | 1.48E-22 | Mari |
| allcell | TMEM230 | 1.20E-50 | 0.887277743 | 0.673 | 0.492 | 2.36E-46 | Mari |
| allcell | NCF4 | 5.13E-24 | 0.886799108 | 0.407 | 0.271 | 1.01E-19 | Mari |
| allcell | DAD1 | 1.68E-72 | 0.886595223 | 0.798 | 0.61 | 3.31E-68 | Mari |
| allcell | TM2D1 | 1.44E-18 | 0.886245276 | 0.345 | 0.235 | 2.84E-14 | Mari |
| allcell | TYMP1 | 7.58E-59 | 0.882775947 | 0.887 | 0.813 | 1.49E-54 | Mari |
| allcell | SNF8 | 8.96E-53 | 0.880866291 | 0.677 | 0.484 | 1.76E-48 | Mari |
| allcell | MRPL40 | 6.26E-29 | 0.880306514 | 0.452 | 0.306 | 1.23E-24 | Mari |
| allcell | CCNDBP1 | 9.07E-28 | 0.880106816 | 0.451 | 0.311 | 1.78E-23 | Mari |
| allcell | VAMP5 | 4.06E-28 | 0.877134137 | 0.639 | 0.516 | 7.99E-24 | Mari |
| allcell | SNHG9 | 8.41E-27 | 0.876912052 | 0.344 | 0.206 | 1.65E-22 | Mari |
| allcell | SELENOT | 8.77E-54 | 0.876126464 | 0.719 | 0.528 | 1.73E-49 | Mari |
| allcell | TSTA3 | 3.72E-20 | 0.875155165 | 0.352 | 0.234 | 7.32E-16 | Mari |
| allcell | IER3IP1 | 1.25E-32 | 0.874442988 | 0.475 | 0.315 | 2.46E-28 | Mari |
| allcell | UROS | 1.28E-18 | 0.873033538 | 0.382 | 0.274 | 2.53E-14 | Mari |
| allcell | MIF4GD | 5.37E-33 | 0.872743828 | 0.566 | 0.414 | 1.06E-28 | Mari |
| allcell | SELENOH | 5.92E-82 | 0.870028291 | 0.879 | 0.704 | 1.16E-77 | Mari |
| allcell | VBP1 | 5.24E-32 | 0.869949705 | 0.482 | 0.324 | 1.03E-27 | Mari |
| allcell | LSM6 | 2.62E-46 | 0.869559419 | 0.644 | 0.468 | 5.16E-42 | Mari |
| allcell | IER2 | 7.04E-57 | 0.867726801 | 0.904 | 0.817 | 1.39E-52 | Mari |
| allcell | BOLA2 | 5.81E-28 | 0.867443582 | 0.445 | 0.295 | 1.14E-23 | Mari |
| allcell | NDUFB2 | 1.60E-100 | 0.865989724 | 0.894 | 0.722 | 3.14E-96 | Mari |
| allcell | TXNDC9 | 1.51E-20 | 0.865890911 | 0.374 | 0.257 | 2.97E-16 | Mari |
| allcell | UQCC2 | 1.46E-28 | 0.865259228 | 0.428 | 0.279 | 2.87E-24 | Mari |
| allcell | MRPL28 | 1.16E-29 | 0.864790781 | 0.446 | 0.299 | 2.28E-25 | Mari |
| allcell | CUTC | 2.28E-18 | 0.862406293 | 0.291 | 0.184 | 4.48E-14 | Mari |
| allcell | EEF1E1 | 1.57E-23 | 0.860792668 | 0.376 | 0.246 | 3.08E-19 | Mari |
| allcell | MIEN1 | 9.81E-27 | 0.860491545 | 0.422 | 0.28 | 1.93E-22 | Mari |
| allcell | DGUOK | 2.97E-50 | 0.85980842 | 0.679 | 0.494 | 5.85E-46 | Mari |
| allcell | EIF4EBP1 | 3.53E-30 | 0.857406346 | 0.533 | 0.398 | 6.95E-26 | Mari |
| allcell | UGCG | 1.58E-22 | 0.855934316 | 0.687 | 0.594 | 3.11E-18 | Mari |
| allcell | PLGRKT | 2.27E-26 | 0.855926334 | 0.481 | 0.342 | 4.47E-22 | Mari |
| allcell | LLPH | 5.42E-21 | 0.85553046 | 0.4 | 0.283 | 1.07E-16 | Mari |
| allcell | POP5 | 5.81E-14 | 0.854053029 | 0.29 | 0.199 | 1.14E-09 | Mari |
| allcell | SSU72 | 8.13E-39 | 0.85359077 | 0.627 | 0.48 | 1.60E-34 | Mari |
| allcell | PDHB | 4.91E-19 | 0.851802382 | 0.352 | 0.239 | 9.67E-15 | Mari |
| allcell | NUDT16L1 | 6.95E-12 | 0.851217412 | 0.254 | 0.175 | 1.37E-07 | Mari |
| allcell | IAH1 | 1.15E-40 | 0.85090354 | 0.62 | 0.444 | 2.26E-36 | Mari |
| allcell | SFT2D1 | 7.03E-57 | 0.84857464 | 0.732 | 0.531 | 1.38E-52 | Mari |
| allcell | SRBD11 | 5.62E-12 | 0.847831275 | 0.26 | 0.178 | 1.11E-07 | Mari |
| allcell | TMEM167A | 7.88E-47 | 0.8471868 | 0.651 | 0.474 | 1.55E-42 | Mari |
| allcell | SPCS2 | 5.19E-67 | 0.844630858 | 0.817 | 0.634 | 1.02E-62 | Mari |

| | | | | | | | |
|---|---|---|---|---|---|---|---|
| allcell | NOL12 | 7.33E-11 | 0.843325275 | 0.255 | 0.181 | 1.44E-06 | Mari |
| allcell | NUDT1 | 9.98E-47 | 0.843121212 | 0.615 | 0.417 | 1.96E-42 | Mari |
| allcell | ANTKMT | 3.64E-25 | 0.842660997 | 0.447 | 0.314 | 7.17E-21 | Mari |
| allcell | MRPL36 | 1.62E-20 | 0.841378274 | 0.405 | 0.281 | 3.19E-16 | Mari |
| allcell | SNRNP40 | 1.19E-21 | 0.840972149 | 0.406 | 0.285 | 2.33E-17 | Mari |
| allcell | ISCU | 1.77E-33 | 0.840370622 | 0.669 | 0.527 | 3.49E-29 | Mari |
| allcell | POLR2K | 2.03E-37 | 0.839493577 | 0.569 | 0.402 | 4.00E-33 | Mari |
| allcell | RER1 | 2.89E-54 | 0.83644973 | 0.715 | 0.533 | 5.68E-50 | Mari |
| allcell | C1D | 6.54E-36 | 0.835511762 | 0.519 | 0.358 | 1.29E-31 | Mari |
| allcell | MCTS1 | 7.28E-25 | 0.834330097 | 0.457 | 0.325 | 1.43E-20 | Mari |
| allcell | PEX2 | 3.98E-17 | 0.833969022 | 0.293 | 0.192 | 7.84E-13 | Mari |
| allcell | ETFB | 5.27E-43 | 0.833756172 | 0.609 | 0.431 | 1.04E-38 | Mari |
| allcell | EBNA1BP2 | 2.73E-15 | 0.832684409 | 0.307 | 0.21 | 5.37E-11 | Mari |
| allcell | PEX16 | 5.89E-17 | 0.832616685 | 0.324 | 0.221 | 1.16E-12 | Mari |
| allcell | RBKS | 7.28E-34 | 0.832268247 | 0.496 | 0.333 | 1.43E-29 | Mari |
| allcell | KRT141 | 4.48E-07 | 0.830694607 | 0.367 | 0.34 | 0.008820823 | Mari |
| allcell | TMEM208 | 2.28E-39 | 0.828861957 | 0.589 | 0.412 | 4.49E-35 | Mari |
| allcell | REXO2 | 5.85E-15 | 0.828294467 | 0.287 | 0.194 | 1.15E-10 | Mari |
| allcell | ZDHHC12 | 2.47E-21 | 0.828215215 | 0.386 | 0.267 | 4.86E-17 | Mari |
| allcell | HSPB111 | 3.62E-40 | 0.827566196 | 0.617 | 0.454 | 7.13E-36 | Mari |
| allcell | MVB12A | 1.05E-13 | 0.827347294 | 0.275 | 0.185 | 2.08E-09 | Mari |
| allcell | ORMDL2 | 2.51E-15 | 0.82646427 | 0.354 | 0.261 | 4.94E-11 | Mari |
| allcell | SMIM19 | 1.02E-28 | 0.826376534 | 0.488 | 0.342 | 2.01E-24 | Mari |
| allcell | TMBIM4 | 4.35E-62 | 0.826258549 | 0.786 | 0.593 | 8.55E-58 | Mari |
| allcell | GPN3 | 9.78E-18 | 0.825945208 | 0.349 | 0.241 | 1.92E-13 | Mari |
| allcell | STOML2 | 4.13E-33 | 0.825700084 | 0.511 | 0.351 | 8.12E-29 | Mari |
| allcell | SS18L2 | 1.66E-30 | 0.824309628 | 0.46 | 0.305 | 3.27E-26 | Mari |
| allcell | DCXR | 1.90E-44 | 0.823750915 | 0.622 | 0.441 | 3.73E-40 | Mari |
| allcell | DERA | 4.71E-22 | 0.823722266 | 0.385 | 0.257 | 9.26E-18 | Mari |
| allcell | HMGN1 | 4.76E-93 | 0.823254428 | 0.916 | 0.783 | 9.37E-89 | Mari |
| allcell | STMN1 | 1.53E-10 | 0.822890407 | 0.445 | 0.381 | 3.02E-06 | Mari |
| allcell | JTB | 1.59E-42 | 0.821570728 | 0.68 | 0.525 | 3.13E-38 | Mari |
| allcell | TIMM13 | 8.28E-38 | 0.8214492 | 0.662 | 0.504 | 1.63E-33 | Mari |
| allcell | DCK | 5.58E-20 | 0.820188714 | 0.445 | 0.327 | 1.10E-15 | Mari |
| allcell | OSTC | 3.09E-66 | 0.818767183 | 0.838 | 0.689 | 6.09E-62 | Mari |
| allcell | SLIRP | 1.22E-44 | 0.817530972 | 0.616 | 0.432 | 2.40E-40 | Mari |
| allcell | CHMP2A | 2.37E-55 | 0.817458356 | 0.717 | 0.527 | 4.67E-51 | Mari |
| allcell | METTL5 | 1.86E-24 | 0.816490101 | 0.429 | 0.298 | 3.66E-20 | Mari |
| allcell | SEC61B | 4.30E-79 | 0.815435162 | 0.945 | 0.884 | 8.46E-75 | Mari |
| allcell | FOS | 6.77E-81 | 0.814921435 | 0.924 | 0.813 | 1.33E-76 | Mari |
| allcell | MRPS12 | 3.14E-28 | 0.814306612 | 0.49 | 0.345 | 6.18E-24 | Mari |
| allcell | BLOC1S1 | 1.05E-76 | 0.813811915 | 0.844 | 0.691 | 2.06E-72 | Mari |
| allcell | IFI30 | 1.55E-10 | 0.811607986 | 0.779 | 0.828 | 3.06E-06 | Mari |
| allcell | PSENEN | 6.63E-40 | 0.811363015 | 0.639 | 0.468 | 1.30E-35 | Mari |
| allcell | AP1S1 | 1.58E-10 | 0.810406688 | 0.267 | 0.196 | 3.11E-06 | Mari |
| allcell | RPA3 | 4.10E-29 | 0.810215484 | 0.5 | 0.344 | 8.06E-25 | Mari |
| allcell | H3F3B | 2.12E-94 | 0.809817499 | 0.992 | 0.981 | 4.17E-90 | Mari |
| allcell | COMMD3 | 4.55E-28 | 0.808731607 | 0.509 | 0.375 | 8.96E-24 | Mari |
| allcell | RABAC1 | 2.61E-55 | 0.808053802 | 0.751 | 0.562 | 5.14E-51 | Mari |
| allcell | CYBB | 5.54E-21 | 0.808015556 | 0.562 | 0.457 | 1.09E-16 | Mari |
| allcell | TMED9 | 2.29E-40 | 0.804145565 | 0.662 | 0.499 | 4.50E-36 | Mari |
| allcell | ACP1 | 1.45E-39 | 0.803433416 | 0.611 | 0.443 | 2.86E-35 | Mari |
| allcell | FDPS | 9.88E-27 | 0.803367689 | 0.494 | 0.353 | 1.94E-22 | Mari |
| allcell | SLC35C2 | 3.78E-14 | 0.802678278 | 0.325 | 0.234 | 7.45E-10 | Mari |
| allcell | ZSWIM7 | 1.71E-23 | 0.801370297 | 0.461 | 0.336 | 3.37E-19 | Mari |
| allcell | TRABD | 3.02E-33 | 0.80103466 | 0.613 | 0.457 | 5.94E-29 | Mari |

| | | | | | | | |
|---|---|---|---|---|---|---|---|
| allcell | ANAPC15 | 3.54E-21 | 0.800849774 | 0.389 | 0.263 | 6.98E-17 | Mari |
| allcell | SRSF9 | 3.10E-78 | 0.800278552 | 0.878 | 0.734 | 6.11E-74 | Mari |
| allcell | UBE2V2 | 4.76E-30 | 0.799497079 | 0.467 | 0.313 | 9.37E-26 | Mari |
| allcell | PALD11 | 2.25E-12 | 0.799483623 | 0.303 | 0.217 | 4.43E-08 | Mari |
| allcell | IMPA2 | 5.59E-13 | 0.798643049 | 0.267 | 0.182 | 1.10E-08 | Mari |
| allcell | NDUFA4 | 7.58E-113 | 0.797624392 | 0.932 | 0.843 | 1.49E-108 | Mari |
| allcell | RNASET2 | 3.26E-34 | 0.797607412 | 0.821 | 0.758 | 6.42E-30 | Mari |
| allcell | JOSD2 | 1.28E-23 | 0.797570107 | 0.438 | 0.308 | 2.52E-19 | Mari |
| allcell | CNPY2 | 4.66E-30 | 0.796044809 | 0.505 | 0.361 | 9.17E-26 | Mari |
| allcell | CHMP5 | 5.92E-33 | 0.795621156 | 0.621 | 0.484 | 1.16E-28 | Mari |
| allcell | SDF2L1 | 1.22E-25 | 0.794543408 | 0.56 | 0.432 | 2.41E-21 | Mari |
| allcell | SMCO4 | 1.61E-46 | 0.792795812 | 0.719 | 0.554 | 3.18E-42 | Mari |
| allcell | SMIM20 | 2.96E-17 | 0.792205395 | 0.436 | 0.327 | 5.83E-13 | Mari |
| allcell | PAFAH1B3 | 1.41E-14 | 0.791013629 | 0.331 | 0.234 | 2.78E-10 | Mari |
| allcell | NDUFS6 | 2.86E-70 | 0.790951629 | 0.826 | 0.638 | 5.63E-66 | Mari |
| allcell | TUSC2 | 4.37E-12 | 0.790624526 | 0.296 | 0.212 | 8.61E-08 | Mari |
| allcell | SCAMP2 | 1.54E-24 | 0.789778482 | 0.505 | 0.38 | 3.02E-20 | Mari |
| allcell | AIF1 | 6.40E-10 | 0.788870701 | 0.734 | 0.746 | 1.26E-05 | Mari |
| allcell | DHRS7 | 4.53E-39 | 0.788443754 | 0.657 | 0.486 | 8.92E-35 | Mari |
| allcell | THOC7 | 1.50E-39 | 0.788092976 | 0.604 | 0.441 | 2.95E-35 | Mari |
| allcell | ATP5MF | 4.48E-99 | 0.786903834 | 0.907 | 0.766 | 8.83E-95 | Mari |
| allcell | PET100 | 2.44E-56 | 0.786884013 | 0.746 | 0.568 | 4.80E-52 | Mari |
| allcell | DNAJC19 | 6.44E-24 | 0.786534531 | 0.437 | 0.303 | 1.27E-19 | Mari |
| allcell | PRKACB | 2.00E-12 | 0.785885334 | 0.317 | 0.229 | 3.94E-08 | Mari |
| allcell | LDHB | 8.00E-46 | 0.785791156 | 0.776 | 0.61 | 1.58E-41 | Mari |
| allcell | SNRPG | 2.13E-86 | 0.78455884 | 0.878 | 0.72 | 4.19E-82 | Mari |
| allcell | LSM4 | 1.13E-48 | 0.784438106 | 0.692 | 0.509 | 2.22E-44 | Mari |
| allcell | MRPL51 | 4.33E-53 | 0.783661753 | 0.727 | 0.552 | 8.53E-49 | Mari |
| allcell | PPM1K1 | 1.28E-17 | 0.783567687 | 0.445 | 0.337 | 2.51E-13 | Mari |
| allcell | HMGN3 | 2.48E-63 | 0.78355646 | 0.845 | 0.685 | 4.89E-59 | Mari |
| allcell | NAA20 | 1.48E-26 | 0.783241904 | 0.511 | 0.371 | 2.92E-22 | Mari |
| allcell | CHMP4A | 3.01E-43 | 0.780862825 | 0.664 | 0.492 | 5.93E-39 | Mari |
| allcell | LSM8 | 5.33E-58 | 0.780651507 | 0.753 | 0.555 | 1.05E-53 | Mari |
| allcell | SRM | 3.20E-26 | 0.78038955 | 0.478 | 0.338 | 6.30E-22 | Mari |
| allcell | TMEM126A | 1.55E-24 | 0.779449451 | 0.418 | 0.284 | 3.05E-20 | Mari |
| allcell | NIPSNAP3A | 3.51E-09 | 0.778744301 | 0.253 | 0.187 | 6.90E-05 | Mari |
| allcell | AKR7A2 | 2.04E-25 | 0.778502676 | 0.495 | 0.361 | 4.02E-21 | Mari |
| allcell | NDUFA3 | 4.06E-53 | 0.777890055 | 0.697 | 0.507 | 7.99E-49 | Mari |
| allcell | NDUFB3 | 2.09E-47 | 0.777809686 | 0.681 | 0.511 | 4.12E-43 | Mari |
| allcell | DBNL | 8.66E-45 | 0.777783776 | 0.711 | 0.536 | 1.70E-40 | Mari |
| allcell | TIMM17B | 1.31E-27 | 0.777225906 | 0.515 | 0.373 | 2.57E-23 | Mari |
| allcell | A1BG | 2.20E-16 | 0.777079616 | 0.456 | 0.357 | 4.32E-12 | Mari |
| allcell | NEDD8 | 3.06E-76 | 0.776930153 | 0.863 | 0.696 | 6.02E-72 | Mari |
| allcell | STMP1 | 1.80E-42 | 0.776420509 | 0.641 | 0.479 | 3.55E-38 | Mari |
| allcell | CIB1 | 3.69E-42 | 0.776306706 | 0.78 | 0.668 | 7.26E-38 | Mari |
| allcell | SRP14 | 1.18E-113 | 0.775862272 | 0.951 | 0.909 | 2.32E-109 | Mari |
| allcell | NADK | 8.16E-07 | 0.775488825 | 0.285 | 0.235 | 0.016056673 | Mari |
| allcell | LAMTOR5 | 1.62E-46 | 0.775401723 | 0.713 | 0.537 | 3.19E-42 | Mari |
| allcell | PPCS | 4.71E-27 | 0.774476512 | 0.492 | 0.352 | 9.28E-23 | Mari |
| allcell | PRDX2 | 1.11E-20 | 0.773857778 | 0.446 | 0.326 | 2.18E-16 | Mari |
| allcell | TRADD | 7.21E-21 | 0.773636835 | 0.483 | 0.369 | 1.42E-16 | Mari |
| allcell | TRAPPC2L | 2.69E-26 | 0.773628839 | 0.492 | 0.356 | 5.30E-22 | Mari |
| allcell | PSMA2 | 8.39E-75 | 0.773057755 | 0.867 | 0.737 | 1.65E-70 | Mari |
| allcell | LMAN2 | 2.50E-42 | 0.771524005 | 0.679 | 0.509 | 4.91E-38 | Mari |
| allcell | COPS9 | 2.19E-55 | 0.771122818 | 0.75 | 0.556 | 4.30E-51 | Mari |
| allcell | NUDCD2 | 3.47E-18 | 0.771075637 | 0.416 | 0.307 | 6.82E-14 | Mari |

| | | | | | | | |
|---|---|---|---|---|---|---|---|
| allcell | SCP2 | 3.23E-42 | 0.770743523 | 0.664 | 0.495 | 6.35E-38 | Mari |
| allcell | SNX17 | 2.05E-39 | 0.769457199 | 0.615 | 0.438 | 4.03E-35 | Mari |
| allcell | HES4 | 6.32E-16 | 0.769040485 | 0.345 | 0.237 | 1.24E-11 | Mari |
| allcell | MRPL3 | 6.35E-27 | 0.769021832 | 0.515 | 0.375 | 1.25E-22 | Mari |
| allcell | EBPL | 8.19E-14 | 0.768709156 | 0.339 | 0.248 | 1.61E-09 | Mari |
| allcell | NMI | 5.06E-26 | 0.767395727 | 0.51 | 0.384 | 9.96E-22 | Mari |
| allcell | EMC6 | 1.29E-22 | 0.767286735 | 0.44 | 0.318 | 2.55E-18 | Mari |
| allcell | ROGDI | 1.14E-10 | 0.766981542 | 0.284 | 0.209 | 2.24E-06 | Mari |
| allcell | GMFG | 5.88E-60 | 0.766070769 | 0.811 | 0.671 | 1.16E-55 | Mari |
| allcell | MRPL17 | 5.53E-10 | 0.766025194 | 0.256 | 0.185 | 1.09E-05 | Mari |
| allcell | ORMDL1 | 1.68E-29 | 0.765046816 | 0.596 | 0.455 | 3.31E-25 | Mari |
| allcell | TXNL4A | 8.67E-44 | 0.764597334 | 0.71 | 0.556 | 1.71E-39 | Mari |
| allcell | ERH | 1.93E-63 | 0.763666229 | 0.81 | 0.644 | 3.79E-59 | Mari |
| allcell | COMMD6 | 6.82E-66 | 0.763310985 | 0.92 | 0.844 | 1.34E-61 | Mari |
| allcell | ERI3 | 3.46E-14 | 0.762559431 | 0.292 | 0.203 | 6.80E-10 | Mari |
| allcell | MRPL55 | 1.94E-25 | 0.762203835 | 0.517 | 0.384 | 3.82E-21 | Mari |
| allcell | UQCR10 | 1.97E-78 | 0.760002527 | 0.871 | 0.701 | 3.87E-74 | Mari |
| allcell | PPIB | 5.55E-82 | 0.758216531 | 0.927 | 0.863 | 1.09E-77 | Mari |
| allcell | RBCK1 | 2.37E-26 | 0.758199448 | 0.553 | 0.421 | 4.67E-22 | Mari |
| allcell | ATRAID | 9.03E-36 | 0.758098305 | 0.573 | 0.42 | 1.78E-31 | Mari |
| allcell | TRMT112 | 3.79E-58 | 0.757072399 | 0.838 | 0.692 | 7.47E-54 | Mari |
| allcell | ATP6V0B | 1.72E-95 | 0.756609572 | 0.921 | 0.797 | 3.38E-91 | Mari |
| allcell | CDC123 | 6.85E-27 | 0.75611694 | 0.489 | 0.352 | 1.35E-22 | Mari |
| allcell | UCP2 | 3.33E-55 | 0.755695767 | 0.848 | 0.698 | 6.56E-51 | Mari |
| allcell | LY96 | 7.00E-34 | 0.755551862 | 0.627 | 0.486 | 1.38E-29 | Mari |
| allcell | MTIF3 | 3.57E-25 | 0.754328613 | 0.503 | 0.368 | 7.03E-21 | Mari |
| allcell | MRPL2 | 3.13E-11 | 0.75402003 | 0.254 | 0.177 | 6.16E-07 | Mari |
| allcell | SLC35B1 | 3.89E-14 | 0.753774806 | 0.335 | 0.243 | 7.66E-10 | Mari |
| allcell | COX7A2 | 4.08E-90 | 0.753733779 | 0.916 | 0.796 | 8.02E-86 | Mari |
| allcell | UBL5 | 7.32E-116 | 0.752069773 | 0.936 | 0.831 | 1.44E-111 | Mari |
| allcell | EMC7 | 1.02E-25 | 0.751546542 | 0.54 | 0.396 | 2.01E-21 | Mari |
| allcell | HPS3 | 3.00E-11 | 0.751474108 | 0.32 | 0.236 | 5.90E-07 | Mari |
| allcell | FAM53C | 1.24E-09 | 0.751424364 | 0.388 | 0.313 | 2.44E-05 | Mari |
| allcell | SMARCB1 | 1.12E-21 | 0.75030028 | 0.462 | 0.335 | 2.20E-17 | Mari |
| allcell | SMC61 | 5.75E-09 | 0.7501045 | 0.266 | 0.199 | 0.00011316 | Mari |
| allcell | POLR2C | 1.03E-14 | 0.749872562 | 0.306 | 0.211 | 2.03E-10 | Mari |
| allcell | COMMD7 | 1.97E-24 | 0.749143315 | 0.461 | 0.328 | 3.88E-20 | Mari |
| allcell | RCN2 | 8.54E-15 | 0.748195575 | 0.363 | 0.268 | 1.68E-10 | Mari |
| allcell | UBE2E3 | 1.64E-14 | 0.746528547 | 0.411 | 0.317 | 3.24E-10 | Mari |
| allcell | ARPC1B | 2.16E-96 | 0.746193154 | 0.948 | 0.86 | 4.25E-92 | Mari |
| allcell | PLRG1 | 2.49E-10 | 0.744612938 | 0.265 | 0.19 | 4.91E-06 | Mari |
| allcell | TBCB | 2.88E-48 | 0.744179348 | 0.722 | 0.551 | 5.66E-44 | Mari |
| allcell | TNFSF13 | 6.68E-11 | 0.743525936 | 0.335 | 0.257 | 1.31E-06 | Mari |
| allcell | MRPS11 | 4.55E-15 | 0.74332527 | 0.355 | 0.254 | 8.96E-11 | Mari |
| allcell | CHD9 | 3.81E-40 | 0.742575365 | 0.669 | 0.489 | 7.50E-36 | Mari |
| allcell | MYL12B | 2.69E-120 | 0.742409448 | 0.941 | 0.874 | 5.30E-116 | Mari |
| allcell | MDP1 | 6.16E-12 | 0.741853533 | 0.296 | 0.213 | 1.21E-07 | Mari |
| allcell | MGST2 | 8.80E-25 | 0.741499537 | 0.549 | 0.419 | 1.73E-20 | Mari |
| allcell | MALSU1 | 1.80E-16 | 0.741409713 | 0.307 | 0.205 | 3.55E-12 | Mari |
| allcell | SETBP11 | 2.67E-15 | 0.740739477 | 0.26 | 0.164 | 5.26E-11 | Mari |
| allcell | AURKAIP1 | 3.15E-62 | 0.740609997 | 0.808 | 0.642 | 6.20E-58 | Mari |
| allcell | NDUFC1 | 4.55E-37 | 0.740347676 | 0.586 | 0.416 | 8.95E-33 | Mari |
| allcell | U2AF1L4 | 9.99E-13 | 0.739625443 | 0.323 | 0.235 | 1.97E-08 | Mari |
| allcell | PLEKHJ1 | 6.61E-15 | 0.739244861 | 0.419 | 0.327 | 1.30E-10 | Mari |
| allcell | ZNHIT1 | 1.28E-43 | 0.738213246 | 0.697 | 0.547 | 2.51E-39 | Mari |
| allcell | APOBEC3G | 1.67E-12 | 0.737644237 | 0.317 | 0.232 | 3.29E-08 | Mari |

| | | | | | | | |
|---|---|---|---|---|---|---|---|
| allcell | IFNAR1 | 8.13E-18 | 0.737402638 | 0.489 | 0.382 | 1.60E-13 | Mari |
| allcell | CENPX | 2.19E-31 | 0.73739679 | 0.542 | 0.394 | 4.32E-27 | Mari |
| allcell | PSMB2 | 7.44E-47 | 0.737157395 | 0.72 | 0.549 | 1.46E-42 | Mari |
| allcell | ABRACL | 8.64E-38 | 0.736901512 | 0.674 | 0.519 | 1.70E-33 | Mari |
| allcell | MCM5 | 1.02E-16 | 0.73629577 | 0.403 | 0.298 | 2.01E-12 | Mari |
| allcell | NSMCE1 | 4.17E-16 | 0.735753946 | 0.424 | 0.321 | 8.21E-12 | Mari |
| allcell | EXOSC7 | 1.51E-12 | 0.73504282 | 0.289 | 0.204 | 2.98E-08 | Mari |
| allcell | TMEM160 | 2.22E-37 | 0.734159083 | 0.625 | 0.473 | 4.37E-33 | Mari |
| allcell | CIAO2B | 4.75E-52 | 0.733653397 | 0.784 | 0.623 | 9.35E-48 | Mari |
| allcell | IMP4 | 7.53E-18 | 0.733281986 | 0.41 | 0.3 | 1.48E-13 | Mari |
| allcell | COX14 | 1.12E-42 | 0.731585612 | 0.667 | 0.496 | 2.20E-38 | Mari |
| allcell | ZFAT1 | 1.42E-20 | 0.73114468 | 0.294 | 0.175 | 2.80E-16 | Mari |
| allcell | P2RY61 | 1.87E-27 | 0.73105362 | 0.428 | 0.27 | 3.69E-23 | Mari |
| allcell | BOLA3 | 3.42E-15 | 0.730775994 | 0.357 | 0.257 | 6.73E-11 | Mari |
| allcell | WDR45 | 6.05E-12 | 0.728976545 | 0.294 | 0.213 | 1.19E-07 | Mari |
| allcell | DYNLRB1 | 3.91E-47 | 0.728889487 | 0.707 | 0.53 | 7.69E-43 | Mari |
| allcell | RPA2 | 3.49E-15 | 0.727214924 | 0.392 | 0.291 | 6.86E-11 | Mari |
| allcell | FKBP3 | 6.86E-17 | 0.726661906 | 0.405 | 0.3 | 1.35E-12 | Mari |
| allcell | MIIP | 1.59E-11 | 0.724999196 | 0.331 | 0.251 | 3.14E-07 | Mari |
| allcell | IFNAR2 | 1.51E-26 | 0.724495614 | 0.598 | 0.46 | 2.97E-22 | Mari |
| allcell | PSMA3 | 1.71E-47 | 0.724428376 | 0.768 | 0.616 | 3.37E-43 | Mari |
| allcell | BTG2 | 2.81E-32 | 0.724401281 | 0.794 | 0.685 | 5.52E-28 | Mari |
| allcell | ACADM | 8.33E-13 | 0.724349115 | 0.347 | 0.259 | 1.64E-08 | Mari |
| allcell | ATP5PF | 1.77E-81 | 0.723626407 | 0.881 | 0.732 | 3.48E-77 | Mari |
| allcell | GMPR2 | 1.92E-09 | 0.722243839 | 0.278 | 0.208 | 3.77E-05 | Mari |
| allcell | PDCD6 | 1.41E-47 | 0.722160453 | 0.717 | 0.526 | 2.78E-43 | Mari |
| allcell | YPEL3 | 1.78E-10 | 0.722104856 | 0.3 | 0.225 | 3.50E-06 | Mari |
| allcell | CPQ | 1.95E-20 | 0.721733878 | 0.523 | 0.398 | 3.85E-16 | Mari |
| allcell | SCAND1 | 1.74E-47 | 0.721569044 | 0.739 | 0.564 | 3.42E-43 | Mari |
| allcell | TNFRSF21 | 3.01E-13 | 0.720706755 | 0.333 | 0.239 | 5.93E-09 | Mari |
| allcell | CDC40 | 4.65E-16 | 0.719495819 | 0.437 | 0.341 | 9.16E-12 | Mari |
| allcell | NDUFA13 | 7.34E-92 | 0.719193213 | 0.923 | 0.783 | 1.44E-87 | Mari |
| allcell | TSN | 2.57E-09 | 0.71780517 | 0.257 | 0.189 | 5.07E-05 | Mari |
| allcell | FEZ2 | 1.42E-17 | 0.716958981 | 0.427 | 0.319 | 2.79E-13 | Mari |
| allcell | COX7A2L | 1.23E-42 | 0.716837953 | 0.803 | 0.696 | 2.42E-38 | Mari |
| allcell | NDUFB1 | 9.76E-65 | 0.716362554 | 0.82 | 0.624 | 1.92E-60 | Mari |
| allcell | SPINT2 | 5.59E-51 | 0.716156151 | 0.803 | 0.615 | 1.10E-46 | Mari |
| allcell | DERL2 | 5.36E-16 | 0.716132776 | 0.371 | 0.268 | 1.05E-11 | Mari |
| allcell | PHPT1 | 5.66E-47 | 0.715722518 | 0.781 | 0.634 | 1.11E-42 | Mari |
| allcell | DUSP22 | 1.27E-14 | 0.714501428 | 0.405 | 0.305 | 2.50E-10 | Mari |
| allcell | BAD | 3.68E-12 | 0.713361982 | 0.334 | 0.249 | 7.23E-08 | Mari |
| allcell | PSMA4 | 2.19E-47 | 0.712997813 | 0.773 | 0.603 | 4.32E-43 | Mari |
| allcell | SUB1 | 8.91E-100 | 0.712727203 | 0.965 | 0.904 | 1.75E-95 | Mari |
| allcell | FAM174C | 2.97E-27 | 0.712073877 | 0.558 | 0.422 | 5.85E-23 | Mari |
| allcell | DDX18 | 4.07E-26 | 0.7114811 | 0.655 | 0.517 | 8.02E-22 | Mari |
| allcell | ECI2 | 4.32E-10 | 0.710975077 | 0.27 | 0.198 | 8.50E-06 | Mari |
| allcell | PDLIM2 | 1.78E-12 | 0.710865472 | 0.342 | 0.252 | 3.49E-08 | Mari |
| allcell | SURF1 | 2.46E-26 | 0.710673299 | 0.551 | 0.412 | 4.84E-22 | Mari |
| allcell | CHCHD5 | 5.16E-25 | 0.710617726 | 0.538 | 0.401 | 1.02E-20 | Mari |
| allcell | MCRS1 | 9.20E-09 | 0.710071869 | 0.254 | 0.188 | 0.000181186 | Mari |
| allcell | IFI27L2 | 1.01E-23 | 0.710051579 | 0.5 | 0.367 | 1.98E-19 | Mari |
| allcell | ANAPC13 | 3.07E-11 | 0.707401067 | 0.278 | 0.198 | 6.04E-07 | Mari |
| allcell | SSBP1 | 5.43E-47 | 0.707006427 | 0.762 | 0.604 | 1.07E-42 | Mari |
| allcell | ERGIC2 | 3.45E-14 | 0.706921168 | 0.395 | 0.298 | 6.80E-10 | Mari |
| allcell | B3GAT3 | 8.62E-13 | 0.706602959 | 0.291 | 0.203 | 1.70E-08 | Mari |
| allcell | GADD45GIP1 | 3.22E-35 | 0.706547585 | 0.644 | 0.496 | 6.34E-31 | Mari |

| | | | | | | | |
|---|---|---|---|---|---|---|---|
| allcell | RBIS | 3.27E-52 | 0.706462263 | 0.804 | 0.651 | 6.45E-48 | Mari |
| allcell | ARL16 | 2.96E-11 | 0.705443456 | 0.287 | 0.206 | 5.83E-07 | Mari |
| allcell | TAF9 | 1.67E-16 | 0.705115583 | 0.393 | 0.287 | 3.29E-12 | Mari |
| allcell | CKLF | 1.33E-25 | 0.704229653 | 0.699 | 0.598 | 2.62E-21 | Mari |
| allcell | MAD2L2 | 1.25E-16 | 0.703598118 | 0.393 | 0.283 | 2.45E-12 | Mari |
| allcell | SNAPIN | 6.10E-14 | 0.703057494 | 0.285 | 0.194 | 1.20E-09 | Mari |
| allcell | SNRPA1 | 7.01E-30 | 0.702772131 | 0.549 | 0.391 | 1.38E-25 | Mari |
| allcell | MSRB1 | 2.77E-12 | 0.702673914 | 0.337 | 0.254 | 5.46E-08 | Mari |
| allcell | NDUFS2 | 9.08E-24 | 0.702170872 | 0.546 | 0.421 | 1.79E-19 | Mari |
| allcell | DCTN3 | 2.64E-35 | 0.702114442 | 0.653 | 0.507 | 5.19E-31 | Mari |
| allcell | EDF1 | 1.86E-107 | 0.701697873 | 0.949 | 0.854 | 3.65E-103 | Mari |
| allcell | NUDT5 | 7.14E-17 | 0.70134786 | 0.404 | 0.298 | 1.41E-12 | Mari |
| allcell | YIPF5 | 7.04E-08 | 0.701006667 | 0.26 | 0.2 | 0.001384984 | Mari |
| allcell | ATP6V0E1 | 9.06E-85 | 0.700788987 | 0.921 | 0.82 | 1.78E-80 | Mari |
| allcell | UFD1 | 7.50E-20 | 0.700105517 | 0.517 | 0.399 | 1.48E-15 | Mari |
| allcell | NUTF2 | 1.99E-30 | 0.699859035 | 0.561 | 0.411 | 3.91E-26 | Mari |
| allcell | POLR3GL | 6.91E-12 | 0.699222907 | 0.309 | 0.227 | 1.36E-07 | Mari |
| allcell | HCCS | 5.80E-17 | 0.698294601 | 0.347 | 0.239 | 1.14E-12 | Mari |
| allcell | MRPL54 | 1.84E-30 | 0.698231516 | 0.606 | 0.462 | 3.62E-26 | Mari |
| allcell | MYDGF | 1.15E-45 | 0.697930483 | 0.774 | 0.611 | 2.27E-41 | Mari |
| allcell | SF3B6 | 3.89E-54 | 0.697824488 | 0.794 | 0.642 | 7.66E-50 | Mari |
| allcell | MPEG1 | 2.81E-16 | 0.697510965 | 0.486 | 0.372 | 5.54E-12 | Mari |
| allcell | PRKRA | 1.39E-12 | 0.697193751 | 0.375 | 0.289 | 2.74E-08 | Mari |
| allcell | MICOS10 | 4.08E-62 | 0.69694547 | 0.855 | 0.708 | 8.03E-58 | Mari |
| allcell | SSNA1 | 2.16E-29 | 0.696722423 | 0.622 | 0.496 | 4.25E-25 | Mari |
| allcell | NMT1 | 1.56E-15 | 0.696283086 | 0.451 | 0.356 | 3.06E-11 | Mari |
| allcell | TMEM147 | 1.46E-24 | 0.696098405 | 0.481 | 0.345 | 2.87E-20 | Mari |
| allcell | ARL1 | 6.84E-13 | 0.695777826 | 0.333 | 0.245 | 1.35E-08 | Mari |
| allcell | RNASEH2B | 2.74E-24 | 0.695318824 | 0.502 | 0.36 | 5.39E-20 | Mari |
| allcell | ESD | 6.63E-28 | 0.694871905 | 0.599 | 0.466 | 1.30E-23 | Mari |
| allcell | GET3 | 2.15E-20 | 0.694728257 | 0.496 | 0.383 | 4.23E-16 | Mari |
| allcell | CORO1A | 6.62E-55 | 0.694503486 | 0.869 | 0.725 | 1.30E-50 | Mari |
| allcell | MRPS16 | 1.30E-14 | 0.693449687 | 0.399 | 0.307 | 2.56E-10 | Mari |
| allcell | MTHFD2L | 1.19E-07 | 0.693306294 | 0.253 | 0.191 | 0.00234245 | Mari |
| allcell | DCTN6 | 2.63E-12 | 0.69152326 | 0.34 | 0.255 | 5.17E-08 | Mari |
| allcell | TSR3 | 1.70E-12 | 0.691399708 | 0.346 | 0.261 | 3.34E-08 | Mari |
| allcell | FBXO6 | 1.48E-19 | 0.689524698 | 0.461 | 0.341 | 2.90E-15 | Mari |
| allcell | SATB1-AS11 | 1.41E-07 | 0.688409712 | 0.254 | 0.19 | 0.002771125 | Mari |
| allcell | MTCH2 | 8.37E-22 | 0.687771054 | 0.48 | 0.356 | 1.65E-17 | Mari |
| allcell | CARHSP1 | 2.70E-13 | 0.687630769 | 0.325 | 0.234 | 5.31E-09 | Mari |
| allcell | CLECL1 | 3.61E-15 | 0.686800209 | 0.414 | 0.309 | 7.10E-11 | Mari |
| allcell | UNC93B1 | 1.11E-17 | 0.686758247 | 0.505 | 0.4 | 2.19E-13 | Mari |
| allcell | FIBP | 3.49E-27 | 0.686746988 | 0.582 | 0.455 | 6.87E-23 | Mari |
| allcell | YDJC | 4.72E-10 | 0.686470506 | 0.293 | 0.217 | 9.29E-06 | Mari |
| allcell | LAMTOR2 | 1.09E-35 | 0.686395332 | 0.679 | 0.553 | 2.14E-31 | Mari |
| allcell | SUMO1 | 1.41E-49 | 0.68637413 | 0.773 | 0.609 | 2.77E-45 | Mari |
| allcell | SUMO3 | 2.99E-34 | 0.684747712 | 0.668 | 0.513 | 5.88E-30 | Mari |
| allcell | OSTF1 | 3.08E-36 | 0.6834347 | 0.773 | 0.637 | 6.06E-32 | Mari |
| allcell | MRPL52 | 1.48E-35 | 0.68248758 | 0.672 | 0.535 | 2.91E-31 | Mari |
| allcell | TAF12 | 1.76E-12 | 0.682155018 | 0.343 | 0.255 | 3.45E-08 | Mari |
| allcell | FLOT1 | 5.08E-37 | 0.68194386 | 0.644 | 0.482 | 1.00E-32 | Mari |
| allcell | AIMP1 | 1.91E-26 | 0.681493557 | 0.54 | 0.403 | 3.77E-22 | Mari |
| allcell | BAX | 4.43E-41 | 0.681334998 | 0.729 | 0.566 | 8.72E-37 | Mari |
| allcell | MRPL27 | 9.29E-15 | 0.680908445 | 0.336 | 0.241 | 1.83E-10 | Mari |
| allcell | LEPROTL1 | 2.95E-27 | 0.680184844 | 0.557 | 0.411 | 5.80E-23 | Mari |
| allcell | RNASEH1 | 2.81E-08 | 0.679286424 | 0.269 | 0.206 | 0.000553795 | Mari |

| | | | | | | | |
|---|---|---|---|---|---|---|---|
| allcell | CCDC115 | 2.57E-15 | 0.679052049 | 0.365 | 0.265 | 5.06E-11 | Mari |
| allcell | CENPC | 1.47E-11 | 0.678582045 | 0.424 | 0.343 | 2.90E-07 | Mari |
| allcell | GLRX5 | 4.29E-30 | 0.678327493 | 0.549 | 0.396 | 8.44E-26 | Mari |
| allcell | FLYWCH2 | 6.93E-09 | 0.678107145 | 0.265 | 0.198 | 0.000136465 | Mari |
| allcell | DPY30 | 2.28E-16 | 0.678084747 | 0.354 | 0.25 | 4.50E-12 | Mari |
| allcell | VAMP2 | 1.16E-20 | 0.677838155 | 0.486 | 0.365 | 2.28E-16 | Mari |
| allcell | XPA | 8.32E-13 | 0.677168359 | 0.287 | 0.203 | 1.64E-08 | Mari |
| allcell | UQCRQ | 3.24E-52 | 0.676569996 | 0.79 | 0.635 | 6.37E-48 | Mari |
| allcell | MRPL22 | 8.24E-19 | 0.676111044 | 0.418 | 0.3 | 1.62E-14 | Mari |
| allcell | CALHM6 | 8.77E-18 | 0.675187927 | 0.742 | 0.67 | 1.73E-13 | Mari |
| allcell | PSMD9 | 3.67E-30 | 0.674764029 | 0.62 | 0.479 | 7.22E-26 | Mari |
| allcell | RGS19 | 2.42E-30 | 0.674742283 | 0.637 | 0.494 | 4.76E-26 | Mari |
| allcell | SDHAF2 | 2.21E-12 | 0.673526963 | 0.348 | 0.262 | 4.35E-08 | Mari |
| allcell | TXNDC15 | 2.05E-09 | 0.672447074 | 0.265 | 0.196 | 4.03E-05 | Mari |
| allcell | TIMM10 | 3.70E-24 | 0.672256543 | 0.559 | 0.429 | 7.28E-20 | Mari |
| allcell | LSM3 | 4.05E-32 | 0.671665555 | 0.642 | 0.489 | 7.97E-28 | Mari |
| allcell | MRPL12 | 3.84E-13 | 0.670650937 | 0.362 | 0.27 | 7.56E-09 | Mari |
| allcell | ATP5MC1 | 8.21E-43 | 0.669848223 | 0.737 | 0.578 | 1.62E-38 | Mari |
| allcell | CCDC69 | 8.81E-08 | 0.669396254 | 0.335 | 0.275 | 0.001734652 | Mari |
| allcell | MNDA | 2.06E-16 | 0.669211484 | 0.631 | 0.566 | 4.05E-12 | Mari |
| allcell | POLR2L | 4.58E-56 | 0.669159413 | 0.837 | 0.685 | 9.02E-52 | Mari |
| allcell | LRR1 | 1.79E-08 | 0.668707742 | 0.285 | 0.215 | 0.00035153 | Mari |
| allcell | BLNK | 2.26E-10 | 0.668594001 | 0.252 | 0.174 | 4.45E-06 | Mari |
| allcell | DYNLT1 | 1.06E-27 | 0.667943635 | 0.578 | 0.437 | 2.08E-23 | Mari |
| allcell | STUB1 | 1.19E-24 | 0.667459732 | 0.561 | 0.441 | 2.33E-20 | Mari |
| allcell | LILRB4 | 5.43E-18 | 0.667080217 | 0.46 | 0.334 | 1.07E-13 | Mari |
| allcell | COMMD4 | 2.87E-13 | 0.666924383 | 0.385 | 0.3 | 5.64E-09 | Mari |
| allcell | TMED3 | 3.88E-17 | 0.664156583 | 0.469 | 0.361 | 7.64E-13 | Mari |
| allcell | MRPL41 | 4.96E-41 | 0.663959298 | 0.676 | 0.532 | 9.76E-37 | Mari |
| allcell | COX16 | 5.28E-22 | 0.663805805 | 0.519 | 0.396 | 1.04E-17 | Mari |
| allcell | PDIA6 | 5.28E-37 | 0.662850377 | 0.728 | 0.577 | 1.04E-32 | Mari |
| allcell | TRMT1 | 3.37E-13 | 0.662128112 | 0.328 | 0.236 | 6.63E-09 | Mari |
| allcell | ARPC1A | 2.59E-12 | 0.660535366 | 0.344 | 0.259 | 5.10E-08 | Mari |
| allcell | AK6 | 2.23E-13 | 0.660089133 | 0.281 | 0.191 | 4.39E-09 | Mari |
| allcell | CAT | 6.69E-19 | 0.660028474 | 0.566 | 0.459 | 1.32E-14 | Mari |
| allcell | POLR2J | 3.38E-27 | 0.660023698 | 0.57 | 0.44 | 6.66E-23 | Mari |
| allcell | EMB | 8.22E-10 | 0.659579142 | 0.361 | 0.288 | 1.62E-05 | Mari |
| allcell | NDUFAF3 | 1.74E-24 | 0.658480928 | 0.555 | 0.433 | 3.42E-20 | Mari |
| allcell | ITGB1BP1 | 6.36E-17 | 0.658096338 | 0.43 | 0.323 | 1.25E-12 | Mari |
| allcell | PSMA5 | 1.76E-44 | 0.658065932 | 0.77 | 0.642 | 3.47E-40 | Mari |
| allcell | EXOSC5 | 2.92E-09 | 0.657818342 | 0.276 | 0.205 | 5.74E-05 | Mari |
| allcell | CLPP | 8.36E-14 | 0.657025375 | 0.378 | 0.285 | 1.65E-09 | Mari |
| allcell | SKA2 | 7.09E-09 | 0.656746329 | 0.254 | 0.189 | 0.000139522 | Mari |
| allcell | DHRS4L2 | 1.70E-08 | 0.656408983 | 0.294 | 0.23 | 0.000333777 | Mari |
| allcell | TXNDC12 | 6.38E-21 | 0.655847725 | 0.497 | 0.376 | 1.25E-16 | Mari |
| allcell | PAM16 | 1.32E-08 | 0.655808458 | 0.3 | 0.235 | 0.000259735 | Mari |
| allcell | UBE2L3 | 1.33E-45 | 0.655469856 | 0.773 | 0.637 | 2.62E-41 | Mari |
| allcell | NDUFB10 | 2.45E-49 | 0.654433901 | 0.801 | 0.641 | 4.82E-45 | Mari |
| allcell | PDCD10 | 6.69E-29 | 0.654379191 | 0.584 | 0.444 | 1.32E-24 | Mari |
| allcell | MRPL33 | 2.95E-31 | 0.654022257 | 0.62 | 0.461 | 5.82E-27 | Mari |
| allcell | HS3ST3B11 | 1.24E-08 | 0.653936789 | 0.281 | 0.209 | 0.000243303 | Mari |
| allcell | JAGN1 | 1.79E-11 | 0.653828819 | 0.338 | 0.258 | 3.53E-07 | Mari |
| allcell | TCF41 | 3.28E-12 | 0.653679364 | 0.585 | 0.514 | 6.45E-08 | Mari |
| allcell | MRPS28 | 4.68E-13 | 0.652492227 | 0.348 | 0.261 | 9.21E-09 | Mari |
| allcell | PIP4P1 | 1.02E-11 | 0.652392045 | 0.302 | 0.217 | 2.01E-07 | Mari |
| allcell | AP2S1 | 8.75E-64 | 0.652383654 | 0.881 | 0.785 | 1.72E-59 | Mari |

| | | | | | | | |
|---|---|---|---|---|---|---|---|
| allcell | SSR4 | 3.57E-56 | 0.652378509 | 0.914 | 0.845 | 7.02E-52 | Mari |
| allcell | LSM2 | 2.42E-25 | 0.65164873 | 0.551 | 0.413 | 4.77E-21 | Mari |
| allcell | CD2AP1 | 3.22E-14 | 0.651457659 | 0.44 | 0.343 | 6.34E-10 | Mari |
| allcell | HSH2D1 | 4.35E-09 | 0.651263837 | 0.263 | 0.191 | 8.57E-05 | Mari |
| allcell | CKAP2 | 2.06E-06 | 0.65084756 | 0.292 | 0.235 | 0.040451421 | Mari |
| allcell | VDAC3 | 3.23E-28 | 0.65070466 | 0.64 | 0.506 | 6.36E-24 | Mari |
| allcell | MRPS7 | 5.78E-18 | 0.64964398 | 0.459 | 0.352 | 1.14E-13 | Mari |
| allcell | NDUFS3 | 6.84E-20 | 0.649472939 | 0.528 | 0.418 | 1.35E-15 | Mari |
| allcell | HTATIP2 | 4.12E-08 | 0.648533127 | 0.295 | 0.228 | 0.000811346 | Mari |
| allcell | PPIE | 6.29E-09 | 0.648479314 | 0.262 | 0.195 | 0.000123716 | Mari |
| allcell | RNF126 | 1.46E-16 | 0.648282453 | 0.462 | 0.36 | 2.87E-12 | Mari |
| allcell | TP53I13 | 1.73E-13 | 0.64803042 | 0.38 | 0.286 | 3.40E-09 | Mari |
| allcell | GPAA1 | 2.86E-11 | 0.647855286 | 0.332 | 0.252 | 5.63E-07 | Mari |
| allcell | SLC50A1 | 1.03E-13 | 0.646892692 | 0.349 | 0.258 | 2.04E-09 | Mari |
| allcell | LARP7 | 3.45E-13 | 0.645508136 | 0.389 | 0.3 | 6.79E-09 | Mari |
| allcell | MED8 | 1.64E-13 | 0.64521822 | 0.358 | 0.263 | 3.22E-09 | Mari |
| allcell | ATP5ME | 1.43E-57 | 0.643636357 | 0.836 | 0.692 | 2.82E-53 | Mari |
| allcell | CFL1 | 1.85E-96 | 0.643248824 | 0.97 | 0.944 | 3.65E-92 | Mari |
| allcell | HVCN1 | 4.57E-11 | 0.643046912 | 0.391 | 0.305 | 8.99E-07 | Mari |
| allcell | PSMB9 | 2.27E-56 | 0.642759431 | 0.893 | 0.806 | 4.47E-52 | Mari |
| allcell | CD300A | 1.48E-16 | 0.642684961 | 0.394 | 0.284 | 2.92E-12 | Mari |
| allcell | TMEM179B | 1.63E-24 | 0.64257515 | 0.525 | 0.379 | 3.20E-20 | Mari |
| allcell | MRPL23 | 6.38E-20 | 0.642014335 | 0.462 | 0.345 | 1.26E-15 | Mari |
| allcell | BORCS7 | 5.05E-09 | 0.641865337 | 0.331 | 0.261 | 9.95E-05 | Mari |
| allcell | NDUFB5 | 3.12E-38 | 0.641330658 | 0.735 | 0.6 | 6.15E-34 | Mari |
| allcell | TOMM6 | 3.87E-51 | 0.641253337 | 0.837 | 0.71 | 7.61E-47 | Mari |
| allcell | IFI35 | 1.45E-10 | 0.640614479 | 0.357 | 0.281 | 2.86E-06 | Mari |
| allcell | SRP19 | 1.95E-36 | 0.640265562 | 0.714 | 0.553 | 3.84E-32 | Mari |
| allcell | CHURC1 | 1.97E-36 | 0.639605357 | 0.733 | 0.59 | 3.88E-32 | Mari |
| allcell | DUSP23 | 1.06E-11 | 0.639489477 | 0.39 | 0.312 | 2.08E-07 | Mari |
| allcell | NDUFA5 | 3.34E-16 | 0.638341001 | 0.461 | 0.362 | 6.58E-12 | Mari |
| allcell | TMA7 | 1.76E-89 | 0.637906861 | 0.939 | 0.866 | 3.47E-85 | Mari |
| allcell | RANBP1 | 7.31E-29 | 0.637795901 | 0.648 | 0.514 | 1.44E-24 | Mari |
| allcell | BANF1 | 3.20E-35 | 0.637123893 | 0.689 | 0.542 | 6.31E-31 | Mari |
| allcell | MRPS26 | 6.10E-09 | 0.636781105 | 0.283 | 0.215 | 0.000120136 | Mari |
| allcell | SERF2 | 8.97E-141 | 0.636554479 | 0.975 | 0.955 | 1.76E-136 | Mari |
| allcell | NT5DC1 | 7.16E-07 | 0.635917354 | 0.27 | 0.209 | 0.014100592 | Mari |
| allcell | SLC25A39 | 2.14E-15 | 0.635863395 | 0.438 | 0.341 | 4.21E-11 | Mari |
| allcell | MDH2 | 2.25E-42 | 0.634372944 | 0.752 | 0.598 | 4.44E-38 | Mari |
| allcell | TRAPPC6A | 8.60E-21 | 0.634208154 | 0.495 | 0.369 | 1.69E-16 | Mari |
| allcell | MPG | 1.56E-24 | 0.633816648 | 0.596 | 0.462 | 3.07E-20 | Mari |
| allcell | NDUFB8 | 4.34E-49 | 0.633744235 | 0.844 | 0.696 | 8.54E-45 | Mari |
| allcell | PHF11 | 1.27E-16 | 0.633725109 | 0.446 | 0.345 | 2.50E-12 | Mari |
| allcell | MDH1 | 1.16E-28 | 0.633356995 | 0.628 | 0.477 | 2.28E-24 | Mari |
| allcell | PPP1R35 | 6.23E-08 | 0.63269757 | 0.255 | 0.195 | 0.001226731 | Mari |
| allcell | TGFBR21 | 8.41E-21 | 0.632669634 | 0.537 | 0.403 | 1.66E-16 | Mari |
| allcell | EXOSC8 | 1.14E-16 | 0.632600723 | 0.4 | 0.291 | 2.23E-12 | Mari |
| allcell | MDFIC | 6.65E-15 | 0.632479359 | 0.511 | 0.407 | 1.31E-10 | Mari |
| allcell | POMP | 5.10E-65 | 0.631326416 | 0.907 | 0.826 | 1.00E-60 | Mari |
| allcell | ARHGDIB | 1.33E-32 | 0.631131241 | 0.824 | 0.775 | 2.62E-28 | Mari |
| allcell | DPP7 | 1.52E-31 | 0.630622799 | 0.71 | 0.583 | 2.99E-27 | Mari |
| allcell | PRMT91 | 4.54E-27 | 0.630542693 | 0.582 | 0.426 | 8.93E-23 | Mari |
| allcell | EMC2 | 5.93E-10 | 0.630146669 | 0.306 | 0.23 | 1.17E-05 | Mari |
| allcell | COPE | 3.42E-67 | 0.629134422 | 0.905 | 0.782 | 6.74E-63 | Mari |
| allcell | SNRPB2 | 7.50E-34 | 0.628664846 | 0.678 | 0.512 | 1.48E-29 | Mari |
| allcell | UCHL5 | 1.96E-10 | 0.628661574 | 0.338 | 0.264 | 3.85E-06 | Mari |

| | | | | | | | |
|---|---|---|---|---|---|---|---|
| allcell | VPS29 | 4.77E-55 | 0.628399036 | 0.839 | 0.709 | 9.38E-51 | Mari |
| allcell | GLOD4 | 3.91E-08 | 0.627989325 | 0.293 | 0.234 | 0.0007692 | Mari |
| allcell | OST4 | 3.05E-77 | 0.627669642 | 0.924 | 0.851 | 5.99E-73 | Mari |
| allcell | TMEM14B | 1.90E-43 | 0.627604604 | 0.774 | 0.592 | 3.75E-39 | Mari |
| allcell | MTX1 | 8.07E-08 | 0.626923691 | 0.252 | 0.191 | 0.001588545 | Mari |
| allcell | ECH1 | 1.14E-22 | 0.626713175 | 0.556 | 0.43 | 2.24E-18 | Mari |
| allcell | COA1 | 2.82E-14 | 0.626073176 | 0.436 | 0.342 | 5.55E-10 | Mari |
| allcell | NSL1 | 6.23E-11 | 0.623558636 | 0.309 | 0.228 | 1.23E-06 | Mari |
| allcell | S100A11 | 2.66E-33 | 0.623254701 | 0.94 | 0.901 | 5.23E-29 | Mari |
| allcell | FAM50A | 1.21E-16 | 0.623011356 | 0.489 | 0.39 | 2.39E-12 | Mari |
| allcell | ATP5F1E | 4.61E-161 | 0.622848725 | 0.977 | 0.967 | 9.08E-157 | Mari |
| allcell | SNHG8 | 7.68E-40 | 0.62282333 | 0.866 | 0.759 | 1.51E-35 | Mari |
| allcell | EIF2AK2 | 1.41E-08 | 0.622737255 | 0.308 | 0.238 | 0.00027735 | Mari |
| allcell | FBXL15 | 1.84E-07 | 0.621873918 | 0.281 | 0.219 | 0.003621582 | Mari |
| allcell | COPS6 | 2.42E-14 | 0.621730594 | 0.429 | 0.335 | 4.77E-10 | Mari |
| allcell | CMC1 | 1.10E-13 | 0.620902545 | 0.391 | 0.299 | 2.17E-09 | Mari |
| allcell | PDCD2 | 1.23E-17 | 0.620370067 | 0.467 | 0.359 | 2.43E-13 | Mari |
| allcell | PIGP | 2.10E-08 | 0.619985537 | 0.334 | 0.267 | 0.000412809 | Mari |
| allcell | AGPAT2 | 5.91E-14 | 0.619463466 | 0.424 | 0.324 | 1.16E-09 | Mari |
| allcell | DPM2 | 2.70E-13 | 0.619305301 | 0.298 | 0.205 | 5.32E-09 | Mari |
| allcell | PSMB1 | 2.04E-58 | 0.619186647 | 0.878 | 0.766 | 4.02E-54 | Mari |
| allcell | GOSR2 | 4.23E-09 | 0.619032163 | 0.26 | 0.192 | 8.33E-05 | Mari |
| allcell | ROMO1 | 4.55E-38 | 0.618676128 | 0.727 | 0.585 | 8.95E-34 | Mari |
| allcell | IFITM2 | 2.13E-17 | 0.617895474 | 0.624 | 0.536 | 4.18E-13 | Mari |
| allcell | JPT1 | 3.30E-33 | 0.617858952 | 0.734 | 0.597 | 6.50E-29 | Mari |
| allcell | EIF5A | 1.08E-38 | 0.617673271 | 0.811 | 0.715 | 2.13E-34 | Mari |
| allcell | FAM32A | 6.50E-17 | 0.61647573 | 0.462 | 0.355 | 1.28E-12 | Mari |
| allcell | VTI1B | 7.94E-12 | 0.61618746 | 0.419 | 0.346 | 1.56E-07 | Mari |
| allcell | MFNG | 3.61E-07 | 0.615706693 | 0.275 | 0.216 | 0.007101378 | Mari |
| allcell | IFI16 | 1.04E-34 | 0.615559507 | 0.762 | 0.615 | 2.04E-30 | Mari |
| allcell | CLTB | 1.28E-33 | 0.615102454 | 0.713 | 0.581 | 2.51E-29 | Mari |
| allcell | SEM1 | 6.02E-56 | 0.613491581 | 0.846 | 0.737 | 1.19E-51 | Mari |
| allcell | HSBP1 | 5.88E-30 | 0.613107327 | 0.68 | 0.561 | 1.16E-25 | Mari |
| allcell | BCAS2 | 4.92E-22 | 0.612957792 | 0.59 | 0.475 | 9.68E-18 | Mari |
| allcell | MRPL43 | 3.74E-24 | 0.612947655 | 0.517 | 0.383 | 7.36E-20 | Mari |
| allcell | UQCRFS1 | 5.56E-33 | 0.612110724 | 0.702 | 0.565 | 1.10E-28 | Mari |
| allcell | BEX4 | 7.28E-14 | 0.611792835 | 0.385 | 0.288 | 1.43E-09 | Mari |
| allcell | SUCLA2 | 4.46E-08 | 0.610455485 | 0.25 | 0.187 | 0.000877197 | Mari |
| allcell | HSD17B10 | 8.69E-19 | 0.610377823 | 0.482 | 0.367 | 1.71E-14 | Mari |
| allcell | CALCRL | 3.50E-20 | 0.610275173 | 0.446 | 0.311 | 6.89E-16 | Mari |
| allcell | TMEM134 | 4.57E-11 | 0.609044505 | 0.359 | 0.279 | 8.99E-07 | Mari |
| allcell | IL10RB | 1.20E-12 | 0.6089908 | 0.381 | 0.291 | 2.37E-08 | Mari |
| allcell | MIA2 | 2.53E-08 | 0.608664415 | 0.358 | 0.288 | 0.000498565 | Mari |
| allcell | COMMD2 | 5.46E-14 | 0.607007351 | 0.381 | 0.275 | 1.07E-09 | Mari |
| allcell | YPEL5 | 8.40E-32 | 0.606923207 | 0.827 | 0.702 | 1.65E-27 | Mari |
| allcell | DHRS4 | 1.91E-07 | 0.606649078 | 0.259 | 0.198 | 0.003751622 | Mari |
| allcell | PIN4 | 2.75E-12 | 0.60624948 | 0.35 | 0.26 | 5.41E-08 | Mari |
| allcell | SMIM15 | 7.09E-10 | 0.605743619 | 0.262 | 0.192 | 1.39E-05 | Mari |
| allcell | SNRPE | 7.52E-32 | 0.605563135 | 0.678 | 0.54 | 1.48E-27 | Mari |
| allcell | SPIB | 1.44E-21 | 0.605497243 | 0.49 | 0.352 | 2.84E-17 | Mari |
| allcell | TPRKB | 3.76E-22 | 0.605476516 | 0.523 | 0.4 | 7.40E-18 | Mari |
| allcell | PLSCR1 | 2.32E-25 | 0.603608968 | 0.749 | 0.629 | 4.57E-21 | Mari |
| allcell | PSMD6 | 1.17E-14 | 0.602045187 | 0.425 | 0.334 | 2.31E-10 | Mari |
| allcell | NDUFS8 | 1.50E-36 | 0.601900825 | 0.713 | 0.566 | 2.95E-32 | Mari |
| allcell | TSG101 | 6.45E-13 | 0.601581371 | 0.414 | 0.327 | 1.27E-08 | Mari |
| allcell | RAD51C | 3.87E-09 | 0.601313678 | 0.298 | 0.225 | 7.62E-05 | Mari |

| | | | | | | | |
|---|---|---|---|---|---|---|---|
| allcell | BCAP29 | 8.82E-10 | 0.6003253 | 0.316 | 0.242 | 1.74E-05 | Mari |
| allcell | TMEM126B | 1.27E-08 | 0.599924162 | 0.285 | 0.22 | 0.000250715 | Mari |
| allcell | BAG1 | 3.66E-32 | 0.599560276 | 0.739 | 0.599 | 7.20E-28 | Mari |
| allcell | CCT8 | 1.06E-38 | 0.599391734 | 0.768 | 0.646 | 2.09E-34 | Mari |
| allcell | S100A71 | 3.90E-09 | 0.598630957 | 0.254 | 0.186 | 7.69E-05 | Mari |
| allcell | PSMB6 | 1.11E-47 | 0.598244757 | 0.82 | 0.692 | 2.19E-43 | Mari |
| allcell | TFPT | 2.45E-11 | 0.59821584 | 0.345 | 0.262 | 4.82E-07 | Mari |
| allcell | PFDN1 | 2.36E-21 | 0.597723067 | 0.478 | 0.338 | 4.65E-17 | Mari |
| allcell | TXN2 | 1.15E-25 | 0.597123539 | 0.623 | 0.5 | 2.27E-21 | Mari |
| allcell | PSMC5 | 9.22E-27 | 0.596922882 | 0.629 | 0.499 | 1.81E-22 | Mari |
| allcell | DTD1 | 4.36E-11 | 0.59676573 | 0.341 | 0.254 | 8.59E-07 | Mari |
| allcell | STX7 | 8.70E-24 | 0.596209035 | 0.647 | 0.525 | 1.71E-19 | Mari |
| allcell | LYRM2 | 2.02E-09 | 0.596188034 | 0.331 | 0.259 | 3.98E-05 | Mari |
| allcell | ZNRF2 | 8.07E-21 | 0.595873128 | 0.54 | 0.406 | 1.59E-16 | Mari |
| allcell | MICOS13 | 2.33E-40 | 0.595247493 | 0.762 | 0.611 | 4.59E-36 | Mari |
| allcell | ZDHHC24 | 1.57E-08 | 0.594681282 | 0.341 | 0.274 | 0.000309694 | Mari |
| allcell | SREK1IP1 | 5.96E-12 | 0.594586207 | 0.33 | 0.243 | 1.17E-07 | Mari |
| allcell | SASH3 | 5.61E-07 | 0.593986564 | 0.271 | 0.214 | 0.011049494 | Mari |
| allcell | MZT1 | 2.46E-08 | 0.593187138 | 0.314 | 0.253 | 0.00048384 | Mari |
| allcell | NOP10 | 1.26E-34 | 0.592975101 | 0.736 | 0.597 | 2.48E-30 | Mari |
| allcell | MPC2 | 2.63E-24 | 0.592658068 | 0.612 | 0.484 | 5.18E-20 | Mari |
| allcell | MRPL9 | 6.34E-11 | 0.592302491 | 0.311 | 0.23 | 1.25E-06 | Mari |
| allcell | WASHC3 | 7.27E-22 | 0.591861366 | 0.48 | 0.349 | 1.43E-17 | Mari |
| allcell | HSD17B41 | 1.45E-06 | 0.591803848 | 0.297 | 0.241 | 0.028625666 | Mari |
| allcell | AHI11 | 1.48E-15 | 0.591518537 | 0.471 | 0.359 | 2.92E-11 | Mari |
| allcell | TNFSF9 | 8.07E-11 | 0.590954033 | 0.432 | 0.352 | 1.59E-06 | Mari |
| allcell | CD2BP2 | 3.24E-11 | 0.590930136 | 0.356 | 0.278 | 6.37E-07 | Mari |
| allcell | NIPSNAP2 | 7.52E-12 | 0.590671385 | 0.391 | 0.306 | 1.48E-07 | Mari |
| allcell | TRAM1 | 2.96E-26 | 0.589927775 | 0.745 | 0.637 | 5.83E-22 | Mari |
| allcell | SEPHS11 | 3.45E-09 | 0.589407776 | 0.291 | 0.222 | 6.78E-05 | Mari |
| allcell | NR3C1 | 5.72E-12 | 0.588689885 | 0.728 | 0.649 | 1.13E-07 | Mari |
| allcell | BRK1 | 1.54E-45 | 0.588602929 | 0.839 | 0.714 | 3.03E-41 | Mari |
| allcell | SDHD | 2.26E-24 | 0.588535803 | 0.603 | 0.479 | 4.44E-20 | Mari |
| allcell | TMED10 | 3.90E-21 | 0.588140783 | 0.688 | 0.576 | 7.68E-17 | Mari |
| allcell | KRTCAP2 | 3.72E-41 | 0.587745503 | 0.811 | 0.676 | 7.32E-37 | Mari |
| allcell | PWP1 | 2.28E-08 | 0.587299919 | 0.307 | 0.242 | 0.000449541 | Mari |
| allcell | NOL7 | 7.53E-31 | 0.587233056 | 0.701 | 0.561 | 1.48E-26 | Mari |
| allcell | SHARPIN | 4.66E-09 | 0.586186567 | 0.325 | 0.254 | 9.17E-05 | Mari |
| allcell | SP110 | 1.12E-21 | 0.585794805 | 0.587 | 0.451 | 2.20E-17 | Mari |
| allcell | APRT | 2.34E-63 | 0.585425742 | 0.905 | 0.803 | 4.61E-59 | Mari |
| allcell | MRPL11 | 1.13E-21 | 0.584662375 | 0.531 | 0.408 | 2.22E-17 | Mari |
| allcell | RRP36 | 8.77E-13 | 0.584621072 | 0.359 | 0.268 | 1.73E-08 | Mari |
| allcell | LSM1 | 3.32E-14 | 0.584065094 | 0.455 | 0.361 | 6.54E-10 | Mari |
| allcell | NUP42 | 1.31E-10 | 0.583773756 | 0.345 | 0.266 | 2.57E-06 | Mari |
| allcell | DIMT1 | 3.96E-08 | 0.583697785 | 0.306 | 0.241 | 0.000780168 | Mari |
| allcell | MED10 | 1.83E-21 | 0.583586715 | 0.544 | 0.418 | 3.60E-17 | Mari |
| allcell | CTSA | 1.13E-08 | 0.583501552 | 0.3 | 0.232 | 0.00022264 | Mari |
| allcell | PSMD8 | 4.32E-43 | 0.582933912 | 0.817 | 0.679 | 8.50E-39 | Mari |
| allcell | PPP1R7 | 4.29E-14 | 0.58278489 | 0.462 | 0.373 | 8.45E-10 | Mari |
| allcell | OTUD6B-AS1 | 2.31E-15 | 0.582540957 | 0.375 | 0.272 | 4.55E-11 | Mari |
| allcell | ATP5MD | 4.02E-48 | 0.580970766 | 0.866 | 0.77 | 7.91E-44 | Mari |
| allcell | MRPL21 | 4.43E-11 | 0.580736449 | 0.407 | 0.329 | 8.72E-07 | Mari |
| allcell | SVBP | 2.96E-13 | 0.580398881 | 0.343 | 0.249 | 5.83E-09 | Mari |
| allcell | CEBPZOS | 1.21E-09 | 0.579844631 | 0.298 | 0.223 | 2.39E-05 | Mari |
| allcell | IRF9 | 2.55E-21 | 0.579592519 | 0.548 | 0.423 | 5.02E-17 | Mari |
| allcell | ATG101 | 7.49E-14 | 0.579038666 | 0.525 | 0.426 | 1.48E-09 | Mari |

| | | | | | | | |
|---|---|---|---|---|---|---|---|
| allcell | SLC39A3 | 2.24E-12 | 0.578912021 | 0.313 | 0.224 | 4.40E-08 | Mari |
| allcell | RFXANK | 1.00E-06 | 0.578431415 | 0.292 | 0.24 | 0.019780313 | Mari |
| allcell | FIS1 | 7.28E-30 | 0.578397153 | 0.699 | 0.567 | 1.43E-25 | Mari |
| allcell | TSTD1 | 1.67E-15 | 0.577788876 | 0.442 | 0.337 | 3.30E-11 | Mari |
| allcell | PSMB8 | 2.11E-43 | 0.577485197 | 0.866 | 0.753 | 4.16E-39 | Mari |
| allcell | PSMG2 | 4.87E-34 | 0.57745272 | 0.711 | 0.577 | 9.59E-30 | Mari |
| allcell | MPV17 | 9.83E-16 | 0.576945253 | 0.41 | 0.304 | 1.94E-11 | Mari |
| allcell | GUK1 | 2.04E-51 | 0.57637478 | 0.895 | 0.794 | 4.02E-47 | Mari |
| allcell | 11-Mar | 9.01E-24 | 0.575857806 | 0.571 | 0.417 | 1.77E-19 | Mari |
| allcell | MRPS34 | 2.24E-24 | 0.575752827 | 0.563 | 0.425 | 4.41E-20 | Mari |
| allcell | RNF166 | 2.59E-10 | 0.574975295 | 0.39 | 0.31 | 5.10E-06 | Mari |
| allcell | TMEM256 | 5.06E-26 | 0.574859977 | 0.641 | 0.524 | 9.96E-22 | Mari |
| allcell | CLNS1A | 1.03E-23 | 0.574833995 | 0.653 | 0.538 | 2.04E-19 | Mari |
| allcell | PAIP2 | 8.20E-26 | 0.574812457 | 0.72 | 0.597 | 1.61E-21 | Mari |
| allcell | DGCR6L | 1.71E-09 | 0.574767442 | 0.325 | 0.249 | 3.37E-05 | Mari |
| allcell | RNASE6 | 1.56E-16 | 0.574517678 | 0.516 | 0.413 | 3.07E-12 | Mari |
| allcell | MRPL16 | 1.28E-08 | 0.574149698 | 0.319 | 0.249 | 0.00025157 | Mari |
| allcell | SRSF7 | 1.87E-38 | 0.573901854 | 0.929 | 0.821 | 3.68E-34 | Mari |
| allcell | MOB1A | 1.51E-45 | 0.573885245 | 0.869 | 0.759 | 2.98E-41 | Mari |
| allcell | NDUFA8 | 1.03E-11 | 0.573872591 | 0.363 | 0.28 | 2.03E-07 | Mari |
| allcell | EEPD11 | 6.86E-09 | 0.573825907 | 0.26 | 0.19 | 0.000135011 | Mari |
| allcell | PPIH | 9.45E-14 | 0.573171059 | 0.396 | 0.297 | 1.86E-09 | Mari |
| allcell | CD47 | 2.33E-30 | 0.572654675 | 0.773 | 0.637 | 4.58E-26 | Mari |
| allcell | TNFSF10 | 1.78E-08 | 0.57248985 | 0.451 | 0.384 | 0.000350384 | Mari |
| allcell | FUNDC2 | 2.30E-16 | 0.572035032 | 0.475 | 0.369 | 4.52E-12 | Mari |
| allcell | TRAPPC4 | 7.16E-09 | 0.571813461 | 0.308 | 0.236 | 0.000140949 | Mari |
| allcell | DDA1 | 4.28E-08 | 0.571709528 | 0.303 | 0.237 | 0.000842315 | Mari |
| allcell | NAXE | 1.35E-06 | 0.570487953 | 0.27 | 0.217 | 0.026488251 | Mari |
| allcell | UFC1 | 1.10E-33 | 0.570434932 | 0.783 | 0.649 | 2.16E-29 | Mari |
| allcell | FKBP1A | 2.99E-35 | 0.569741968 | 0.802 | 0.686 | 5.89E-31 | Mari |
| allcell | STXBP2 | 9.53E-11 | 0.56955723 | 0.436 | 0.354 | 1.88E-06 | Mari |
| allcell | MEAF6 | 5.63E-14 | 0.569394949 | 0.399 | 0.304 | 1.11E-09 | Mari |
| allcell | ADH5 | 5.35E-09 | 0.569299681 | 0.337 | 0.268 | 0.000105293 | Mari |
| allcell | SCO2 | 2.31E-19 | 0.567915302 | 0.587 | 0.461 | 4.55E-15 | Mari |
| allcell | EAPP | 2.54E-10 | 0.567738614 | 0.348 | 0.272 | 4.99E-06 | Mari |
| allcell | ARL4D | 1.29E-11 | 0.567494317 | 0.275 | 0.185 | 2.55E-07 | Mari |
| allcell | SLC25A11 | 2.71E-12 | 0.566858435 | 0.419 | 0.335 | 5.34E-08 | Mari |
| allcell | TYROBP | 3.87E-30 | 0.566505442 | 0.863 | 0.816 | 7.62E-26 | Mari |
| allcell | OCIAD1 | 5.97E-24 | 0.566490991 | 0.648 | 0.513 | 1.18E-19 | Mari |
| allcell | DCUN1D5 | 1.47E-10 | 0.566302912 | 0.331 | 0.253 | 2.89E-06 | Mari |
| allcell | APEX1 | 2.27E-19 | 0.56596649 | 0.689 | 0.595 | 4.46E-15 | Mari |
| allcell | NDUFA1 | 9.04E-56 | 0.565641247 | 0.872 | 0.748 | 1.78E-51 | Mari |
| allcell | PYCARD | 7.84E-34 | 0.565639594 | 0.816 | 0.737 | 1.54E-29 | Mari |
| allcell | FAM192A | 4.74E-11 | 0.5655272 | 0.402 | 0.322 | 9.34E-07 | Mari |
| allcell | SELENOF | 7.05E-21 | 0.565289564 | 0.662 | 0.557 | 1.39E-16 | Mari |
| allcell | SMIM14 | 5.33E-21 | 0.565176754 | 0.594 | 0.465 | 1.05E-16 | Mari |
| allcell | ARPC3 | 6.80E-88 | 0.56481002 | 0.952 | 0.921 | 1.34E-83 | Mari |
| allcell | ECHDC1 | 2.26E-19 | 0.563894038 | 0.565 | 0.443 | 4.46E-15 | Mari |
| allcell | GABARAP | 1.96E-83 | 0.563128606 | 0.952 | 0.918 | 3.86E-79 | Mari |
| allcell | PARL | 1.65E-13 | 0.56287629 | 0.449 | 0.356 | 3.24E-09 | Mari |
| allcell | TRIM22 | 5.25E-15 | 0.562119245 | 0.601 | 0.511 | 1.03E-10 | Mari |
| allcell | SUMO2 | 1.97E-81 | 0.562064775 | 0.936 | 0.884 | 3.88E-77 | Mari |
| allcell | ITGA41 | 1.11E-27 | 0.560507674 | 0.674 | 0.518 | 2.18E-23 | Mari |
| allcell | LAMTOR4 | 7.90E-45 | 0.55923185 | 0.84 | 0.727 | 1.55E-40 | Mari |
| allcell | TLE5 | 3.14E-25 | 0.558786047 | 0.784 | 0.663 | 6.18E-21 | Mari |
| allcell | GGA21 | 5.42E-10 | 0.558297624 | 0.372 | 0.295 | 1.07E-05 | Mari |

| | | | | | | | |
|---|---|---|---|---|---|---|---|
| allcell | TWF2 | 9.45E-29 | 0.558167837 | 0.693 | 0.569 | 1.86E-24 | Mari |
| allcell | PUF60 | 1.95E-15 | 0.55775084 | 0.466 | 0.366 | 3.84E-11 | Mari |
| allcell | PEBP1 | 1.04E-28 | 0.55740837 | 0.786 | 0.667 | 2.04E-24 | Mari |
| allcell | TMEM50B | 1.72E-14 | 0.557237504 | 0.397 | 0.292 | 3.38E-10 | Mari |
| allcell | ISOC2 | 2.97E-11 | 0.556712501 | 0.413 | 0.328 | 5.85E-07 | Mari |
| allcell | COX17 | 4.32E-33 | 0.556219468 | 0.748 | 0.598 | 8.51E-29 | Mari |
| allcell | LRPAP1 | 1.44E-18 | 0.555967392 | 0.506 | 0.387 | 2.83E-14 | Mari |
| allcell | PSMD13 | 2.39E-18 | 0.555788502 | 0.554 | 0.451 | 4.70E-14 | Mari |
| allcell | POLR2F | 2.91E-15 | 0.555674738 | 0.442 | 0.338 | 5.73E-11 | Mari |
| allcell | PDAP1 | 1.75E-17 | 0.555579115 | 0.51 | 0.4 | 3.44E-13 | Mari |
| allcell | ATOX1 | 8.26E-22 | 0.554750438 | 0.813 | 0.721 | 1.63E-17 | Mari |
| allcell | PARVG | 9.37E-21 | 0.554342205 | 0.567 | 0.446 | 1.85E-16 | Mari |
| allcell | BROX | 3.41E-08 | 0.554022081 | 0.315 | 0.252 | 0.00067176 | Mari |
| allcell | AP3S1 | 9.41E-11 | 0.553756281 | 0.478 | 0.397 | 1.85E-06 | Mari |
| allcell | HEBP2 | 5.57E-15 | 0.553220167 | 0.451 | 0.344 | 1.10E-10 | Mari |
| allcell | SMIM12 | 9.99E-10 | 0.552997813 | 0.372 | 0.3 | 1.97E-05 | Mari |
| allcell | CAPZA2 | 6.51E-28 | 0.552978179 | 0.69 | 0.57 | 1.28E-23 | Mari |
| allcell | DUT | 1.31E-13 | 0.552585612 | 0.505 | 0.419 | 2.58E-09 | Mari |
| allcell | MRPL15 | 1.23E-10 | 0.552561787 | 0.329 | 0.249 | 2.43E-06 | Mari |
| allcell | DCTPP1 | 4.46E-11 | 0.552333955 | 0.402 | 0.323 | 8.79E-07 | Mari |
| allcell | NAA38 | 8.75E-22 | 0.552186354 | 0.587 | 0.459 | 1.72E-17 | Mari |
| allcell | HMGN2 | 1.28E-37 | 0.55177543 | 0.858 | 0.748 | 2.53E-33 | Mari |
| allcell | APP | 2.26E-08 | 0.551116203 | 0.506 | 0.45 | 0.000444198 | Mari |
| allcell | COMMD9 | 1.30E-13 | 0.550910501 | 0.433 | 0.337 | 2.56E-09 | Mari |
| allcell | PSMB3 | 5.92E-45 | 0.55062983 | 0.857 | 0.739 | 1.17E-40 | Mari |
| allcell | VMA21 | 1.92E-17 | 0.550222435 | 0.53 | 0.419 | 3.77E-13 | Mari |
| allcell | SNHG12 | 1.03E-11 | 0.549949701 | 0.456 | 0.377 | 2.03E-07 | Mari |
| allcell | NPC2 | 5.01E-19 | 0.549533598 | 0.844 | 0.782 | 9.86E-15 | Mari |
| allcell | ABI3 | 4.27E-07 | 0.549411996 | 0.363 | 0.303 | 0.008413448 | Mari |
| allcell | NDUFV2 | 1.62E-59 | 0.548223242 | 0.901 | 0.805 | 3.20E-55 | Mari |
| allcell | CASP4 | 3.95E-11 | 0.547945596 | 0.52 | 0.454 | 7.78E-07 | Mari |
| allcell | TMEM50A | 4.15E-35 | 0.54682892 | 0.809 | 0.691 | 8.16E-31 | Mari |
| allcell | MRPS33 | 1.84E-07 | 0.54651991 | 0.366 | 0.309 | 0.003628008 | Mari |
| allcell | FUOM | 7.76E-15 | 0.545556758 | 0.541 | 0.445 | 1.53E-10 | Mari |
| allcell | ECHS1 | 5.29E-19 | 0.544747871 | 0.561 | 0.448 | 1.04E-14 | Mari |
| allcell | RPAIN | 5.07E-14 | 0.544325752 | 0.43 | 0.329 | 9.97E-10 | Mari |
| allcell | TCTN3 | 2.35E-07 | 0.544065453 | 0.316 | 0.252 | 0.004616766 | Mari |
| allcell | TCIRG1 | 2.75E-09 | 0.543825344 | 0.383 | 0.308 | 5.41E-05 | Mari |
| allcell | HSCB | 1.36E-06 | 0.54316069 | 0.259 | 0.2 | 0.026772377 | Mari |
| allcell | MYL6 | 6.54E-76 | 0.541587772 | 0.972 | 0.953 | 1.29E-71 | Mari |
| allcell | SRA1 | 2.73E-23 | 0.541484288 | 0.605 | 0.486 | 5.37E-19 | Mari |
| allcell | ZNF706 | 1.51E-30 | 0.5410916 | 0.79 | 0.671 | 2.96E-26 | Mari |
| allcell | MESD | 4.12E-14 | 0.540426051 | 0.494 | 0.399 | 8.10E-10 | Mari |
| allcell | VKORC1 | 4.11E-13 | 0.540039763 | 0.42 | 0.326 | 8.08E-09 | Mari |
| allcell | COMT | 2.46E-25 | 0.539767922 | 0.687 | 0.569 | 4.83E-21 | Mari |
| allcell | ZSCAN16-AS1 | 7.23E-07 | 0.539017933 | 0.271 | 0.211 | 0.014238326 | Mari |
| allcell | COX6C | 6.30E-66 | 0.538935272 | 0.921 | 0.835 | 1.24E-61 | Mari |
| allcell | ATP5F1C | 2.33E-42 | 0.538685781 | 0.855 | 0.742 | 4.58E-38 | Mari |
| allcell | PPP1R11 | 5.22E-07 | 0.538457266 | 0.287 | 0.229 | 0.010265874 | Mari |
| allcell | NME1 | 1.88E-16 | 0.538136658 | 0.514 | 0.412 | 3.70E-12 | Mari |
| allcell | UBE2J1 | 1.32E-12 | 0.53802249 | 0.623 | 0.547 | 2.60E-08 | Mari |
| allcell | DTNBP1 | 1.51E-11 | 0.537063511 | 0.531 | 0.441 | 2.98E-07 | Mari |
| allcell | BCL7C | 1.51E-10 | 0.536914138 | 0.432 | 0.356 | 2.96E-06 | Mari |
| allcell | SRSF5 | 2.18E-40 | 0.535377254 | 0.839 | 0.681 | 4.29E-36 | Mari |
| allcell | TALDO1 | 3.56E-36 | 0.535360444 | 0.827 | 0.721 | 7.02E-32 | Mari |
| allcell | ISG15 | 2.91E-21 | 0.534912797 | 0.66 | 0.547 | 5.73E-17 | Mari |

| | | | | | | | |
|---|---|---|---|---|---|---|---|
| allcell | UXT | 1.24E-40 | 0.534640591 | 0.857 | 0.755 | 2.43E-36 | Mari |
| allcell | LSM7 | 4.73E-36 | 0.534157252 | 0.814 | 0.711 | 9.32E-32 | Mari |
| allcell | CAPG | 1.43E-23 | 0.533826821 | 0.776 | 0.693 | 2.82E-19 | Mari |
| allcell | GLO1 | 1.09E-11 | 0.533334865 | 0.409 | 0.324 | 2.15E-07 | Mari |
| allcell | PSME2 | 6.12E-32 | 0.533120204 | 0.911 | 0.852 | 1.20E-27 | Mari |
| allcell | NDUFA7 | 1.55E-25 | 0.532113947 | 0.7 | 0.587 | 3.06E-21 | Mari |
| allcell | COPS5 | 7.03E-12 | 0.531965879 | 0.415 | 0.331 | 1.38E-07 | Mari |
| allcell | BUD31 | 5.32E-25 | 0.531689122 | 0.672 | 0.551 | 1.05E-20 | Mari |
| allcell | ATP5MPL | 3.72E-33 | 0.529554379 | 0.797 | 0.642 | 7.32E-29 | Mari |
| allcell | NXT1 | 2.40E-22 | 0.529176021 | 0.511 | 0.369 | 4.73E-18 | Mari |
| allcell | SLC25A26 | 1.50E-07 | 0.528691351 | 0.295 | 0.23 | 0.002951854 | Mari |
| allcell | COX8A | 1.33E-57 | 0.528446141 | 0.909 | 0.841 | 2.62E-53 | Mari |
| allcell | MITD11 | 5.91E-07 | 0.527598418 | 0.276 | 0.218 | 0.011634882 | Mari |
| allcell | ZNF593 | 3.66E-13 | 0.527224204 | 0.354 | 0.26 | 7.20E-09 | Mari |
| allcell | TSSC4 | 1.42E-07 | 0.526788651 | 0.273 | 0.211 | 0.002796063 | Mari |
| allcell | PHYKPL | 4.47E-09 | 0.526217804 | 0.389 | 0.323 | 8.79E-05 | Mari |
| allcell | SNRPD3 | 4.37E-22 | 0.526173467 | 0.644 | 0.529 | 8.60E-18 | Mari |
| allcell | SNRPD1 | 2.86E-26 | 0.524993129 | 0.702 | 0.57 | 5.64E-22 | Mari |
| allcell | LMAN1 | 4.62E-08 | 0.524633788 | 0.377 | 0.311 | 0.000909441 | Mari |
| allcell | MRPL34 | 9.61E-11 | 0.524524162 | 0.445 | 0.366 | 1.89E-06 | Mari |
| allcell | PRMT1 | 1.26E-19 | 0.524340279 | 0.629 | 0.513 | 2.48E-15 | Mari |
| allcell | ZNHIT3 | 5.59E-08 | 0.524331374 | 0.347 | 0.282 | 0.001099709 | Mari |
| allcell | MRPS21 | 2.92E-33 | 0.524214979 | 0.762 | 0.636 | 5.75E-29 | Mari |
| allcell | C1QBP | 3.00E-30 | 0.524207767 | 0.768 | 0.648 | 5.90E-26 | Mari |
| allcell | ACAA1 | 6.86E-23 | 0.523993728 | 0.657 | 0.545 | 1.35E-18 | Mari |
| allcell | TRAPPC5 | 1.87E-23 | 0.523808517 | 0.668 | 0.558 | 3.69E-19 | Mari |
| allcell | RUNX21 | 2.32E-19 | 0.523665662 | 0.399 | 0.269 | 4.57E-15 | Mari |
| allcell | ATP5MC3 | 6.70E-52 | 0.522936657 | 0.902 | 0.806 | 1.32E-47 | Mari |
| allcell | TMEM258 | 5.94E-45 | 0.522598866 | 0.865 | 0.755 | 1.17E-40 | Mari |
| allcell | RNF187 | 9.14E-07 | 0.522591004 | 0.309 | 0.25 | 0.017982561 | Mari |
| allcell | PPCDC1 | 7.17E-16 | 0.521506063 | 0.341 | 0.233 | 1.41E-11 | Mari |
| allcell | PFN1 | 4.62E-64 | 0.520915044 | 0.977 | 0.941 | 9.10E-60 | Mari |
| allcell | SLC15A41 | 9.86E-09 | 0.520879929 | 0.44 | 0.374 | 0.000194153 | Mari |
| allcell | BBIP1 | 1.59E-06 | 0.520277021 | 0.25 | 0.196 | 0.031259709 | Mari |
| allcell | NDUFA12 | 3.90E-30 | 0.519621858 | 0.773 | 0.64 | 7.68E-26 | Mari |
| allcell | RMDN1 | 4.49E-08 | 0.51889863 | 0.361 | 0.295 | 0.000884653 | Mari |
| allcell | ATP6V1G1 | 5.75E-39 | 0.518119821 | 0.866 | 0.759 | 1.13E-34 | Mari |
| allcell | PIK3CG | 2.25E-07 | 0.517172255 | 0.375 | 0.31 | 0.004424937 | Mari |
| allcell | ERCC1 | 6.01E-14 | 0.515721951 | 0.575 | 0.494 | 1.18E-09 | Mari |
| allcell | DHPS | 9.24E-11 | 0.515454909 | 0.396 | 0.313 | 1.82E-06 | Mari |
| allcell | SHKBP1 | 1.04E-10 | 0.515043183 | 0.392 | 0.311 | 2.05E-06 | Mari |
| allcell | PPP4C | 7.69E-23 | 0.514863989 | 0.648 | 0.532 | 1.51E-18 | Mari |
| allcell | ASPH | 8.81E-09 | 0.514374442 | 0.403 | 0.327 | 0.000173418 | Mari |
| allcell | SYPL1 | 1.17E-06 | 0.514250416 | 0.328 | 0.275 | 0.023048182 | Mari |
| allcell | NDUFS7 | 1.76E-30 | 0.514232811 | 0.73 | 0.586 | 3.47E-26 | Mari |
| allcell | HAGH | 4.14E-08 | 0.5120685 | 0.38 | 0.318 | 0.000814038 | Mari |
| allcell | CD99 | 3.75E-11 | 0.511860821 | 0.662 | 0.595 | 7.39E-07 | Mari |
| allcell | FHIT | 1.11E-08 | 0.511259269 | 0.301 | 0.224 | 0.000219081 | Mari |
| allcell | MPC1 | 4.26E-16 | 0.510719153 | 0.587 | 0.484 | 8.38E-12 | Mari |
| allcell | RRP7A | 1.32E-09 | 0.510706173 | 0.395 | 0.323 | 2.60E-05 | Mari |
| allcell | DDRGK1 | 2.90E-10 | 0.510670424 | 0.405 | 0.329 | 5.70E-06 | Mari |
| allcell | SERPINB1 | 4.27E-23 | 0.510647345 | 0.86 | 0.791 | 8.41E-19 | Mari |
| allcell | PRPF31 | 6.68E-08 | 0.510453308 | 0.333 | 0.269 | 0.001314709 | Mari |
| allcell | PIK3AP1 | 3.98E-14 | 0.509485254 | 0.445 | 0.34 | 7.83E-10 | Mari |
| allcell | SMDT1 | 2.74E-37 | 0.509113573 | 0.863 | 0.756 | 5.39E-33 | Mari |
| allcell | RAB34 | 5.70E-12 | 0.508804022 | 0.462 | 0.368 | 1.12E-07 | Mari |

| | | | | | | | |
|---|---|---|---|---|---|---|---|
| allcell | ACTR10 | 6.50E-11 | 0.508666052 | 0.437 | 0.357 | 1.28E-06 | Mari |
| allcell | RNF7 | 3.43E-34 | 0.50832063 | 0.78 | 0.647 | 6.74E-30 | Mari |
| allcell | GSTK1 | 9.89E-35 | 0.507968177 | 0.819 | 0.72 | 1.95E-30 | Mari |
| allcell | OSGEP | 7.84E-10 | 0.507837611 | 0.358 | 0.282 | 1.54E-05 | Mari |
| allcell | TRNAU1AP | 4.38E-08 | 0.506726263 | 0.299 | 0.234 | 0.000861249 | Mari |
| allcell | UBE2I | 1.19E-22 | 0.504877586 | 0.735 | 0.634 | 2.35E-18 | Mari |
| allcell | ZNRD2 | 7.28E-10 | 0.504589559 | 0.346 | 0.272 | 1.43E-05 | Mari |
| allcell | VPS28 | 1.16E-29 | 0.504373242 | 0.772 | 0.661 | 2.29E-25 | Mari |
| allcell | STAT2 | 3.81E-09 | 0.503842705 | 0.373 | 0.302 | 7.51E-05 | Mari |
| allcell | MRPS36 | 2.23E-21 | 0.503705393 | 0.569 | 0.436 | 4.39E-17 | Mari |
| allcell | ERVK3-1 | 1.45E-11 | 0.503307543 | 0.415 | 0.327 | 2.85E-07 | Mari |
| allcell | HINT1 | 3.22E-46 | 0.502704397 | 0.943 | 0.912 | 6.35E-42 | Mari |
| allcell | ATXN10 | 1.31E-11 | 0.502685573 | 0.434 | 0.352 | 2.59E-07 | Mari |
| allcell | DNAJC15 | 1.51E-31 | 0.502394276 | 0.777 | 0.648 | 2.97E-27 | Mari |
| allcell | DPM3 | 2.01E-08 | 0.502068379 | 0.369 | 0.298 | 0.000395336 | Mari |
| allcell | NQO2 | 4.89E-22 | 0.501976591 | 0.567 | 0.441 | 9.63E-18 | Mari |
| allcell | ELOB | 7.41E-47 | 0.501822163 | 0.889 | 0.803 | 1.46E-42 | Mari |
| allcell | WDR74 | 8.03E-07 | 0.501732693 | 0.347 | 0.287 | 0.015814352 | Mari |
| allcell | IRAK4 | 1.32E-07 | 0.501626003 | 0.338 | 0.275 | 0.002599235 | Mari |
| allcell | NDUFA2 | 4.20E-31 | 0.500311285 | 0.755 | 0.618 | 8.27E-27 | Mari |
| allcell | NPHS1 | 5.31E-59 | 2.81251785 | 0.3 | 0.124 | 1.04E-54 | Tb |
| allcell | GCNT2 | 6.80E-40 | 2.145110032 | 0.272 | 0.122 | 1.34E-35 | Tb |
| allcell | SLC35F1 | 3.42E-75 | 1.981334723 | 0.366 | 0.145 | 6.72E-71 | Tb |
| allcell | S100A7 | 2.17E-115 | 1.923481291 | 0.411 | 0.118 | 4.27E-111 | Tb |
| allcell | ARL17A | 4.66E-80 | 1.920700459 | 0.307 | 0.093 | 9.18E-76 | Tb |
| allcell | PSMD5 | 2.57E-38 | 1.808933765 | 0.3 | 0.144 | 5.06E-34 | Tb |
| allcell | KRTDAP | 2.89E-32 | 1.800319102 | 0.265 | 0.132 | 5.69E-28 | Tb |
| allcell | FBP1 | 1.88E-10 | 1.778695766 | 0.395 | 0.311 | 3.69E-06 | Tb |
| allcell | S100A2 | 1.64E-107 | 1.769736524 | 0.365 | 0.097 | 3.24E-103 | Tb |
| allcell | HOMER1 | 6.95E-90 | 1.732081034 | 0.575 | 0.294 | 1.37E-85 | Tb |
| allcell | AQP3 | 3.00E-65 | 1.651992077 | 0.281 | 0.092 | 5.91E-61 | Tb |
| allcell | KRT6A | 9.84E-107 | 1.639178607 | 0.372 | 0.1 | 1.94E-102 | Tb |
| allcell | CCR6 | 5.64E-44 | 1.594604193 | 0.272 | 0.114 | 1.11E-39 | Tb |
| allcell | TAF4B | 9.72E-57 | 1.507025072 | 0.442 | 0.221 | 1.91E-52 | Tb |
| allcell | RHOH | 3.77E-82 | 1.388056185 | 0.586 | 0.31 | 7.43E-78 | Tb |
| allcell | ZNF609 | 1.94E-66 | 1.378191472 | 0.455 | 0.217 | 3.81E-62 | Tb |
| allcell | KRT17 | 9.98E-70 | 1.375176199 | 0.28 | 0.083 | 1.96E-65 | Tb |
| allcell | PHTF1 | 7.57E-47 | 1.35826151 | 0.395 | 0.201 | 1.49E-42 | Tb |
| allcell | MCF2L2 | 2.76E-41 | 1.357118494 | 0.418 | 0.235 | 5.43E-37 | Tb |
| allcell | RGS13 | 4.25E-44 | 1.34677274 | 0.361 | 0.181 | 8.37E-40 | Tb |
| allcell | FAM222B | 1.89E-52 | 1.34279309 | 0.295 | 0.115 | 3.72E-48 | Tb |
| allcell | CLEC16A | 7.24E-64 | 1.341060811 | 0.528 | 0.279 | 1.42E-59 | Tb |
| allcell | TBC1D32 | 1.59E-43 | 1.340750861 | 0.275 | 0.113 | 3.13E-39 | Tb |
| allcell | FAM118A | 7.02E-54 | 1.332111269 | 0.431 | 0.217 | 1.38E-49 | Tb |
| allcell | SFN | 6.33E-49 | 1.298165922 | 0.302 | 0.123 | 1.25E-44 | Tb |
| allcell | L3MBTL4 | 2.84E-22 | 1.297324638 | 0.304 | 0.179 | 5.59E-18 | Tb |
| allcell | IGKC | 6.36E-15 | 1.293682541 | 0.393 | 0.282 | 1.25E-10 | Tb |
| allcell | GAB1 | 8.52E-31 | 1.274970594 | 0.355 | 0.201 | 1.68E-26 | Tb |
| allcell | NEK1 | 9.49E-26 | 1.259723419 | 0.336 | 0.195 | 1.87E-21 | Tb |
| allcell | SLC33A1 | 3.23E-38 | 1.244753179 | 0.281 | 0.125 | 6.35E-34 | Tb |
| allcell | ADAM19 | 4.44E-82 | 1.219355118 | 0.663 | 0.377 | 8.75E-78 | Tb |
| allcell | NCR3LG1 | 5.83E-33 | 1.217723538 | 0.253 | 0.116 | 1.15E-28 | Tb |
| allcell | S100A8 | 1.56E-141 | 1.201221103 | 0.638 | 0.243 | 3.07E-137 | Tb |
| allcell | XIST | 5.57E-101 | 1.186932054 | 0.619 | 0.284 | 1.10E-96 | Tb |
| allcell | NLK | 3.04E-49 | 1.177793198 | 0.321 | 0.136 | 5.99E-45 | Tb |
| allcell | GYS1 | 4.37E-44 | 1.177170067 | 0.303 | 0.131 | 8.60E-40 | Tb |

| | | | | | | | |
|---|---|---|---|---|---|---|---|
| allcell | NARS2 | 1.78E-28 | 1.17702859 | 0.327 | 0.18 | 3.50E-24 | Tb |
| allcell | CLIC4 | 8.90E-38 | 1.170233096 | 0.548 | 0.374 | 1.75E-33 | Tb |
| allcell | INHBA | 1.47E-09 | 1.169449916 | 0.285 | 0.212 | 2.89E-05 | Tb |
| allcell | KRT6B | 2.24E-61 | 1.16230466 | 0.261 | 0.078 | 4.41E-57 | Tb |
| allcell | GBF1 | 7.93E-69 | 1.160018125 | 0.499 | 0.241 | 1.56E-64 | Tb |
| allcell | BRCA2 | 3.29E-40 | 1.157821232 | 0.446 | 0.254 | 6.48E-36 | Tb |
| allcell | PHLPP1 | 2.38E-51 | 1.149962232 | 0.549 | 0.335 | 4.69E-47 | Tb |
| allcell | CYTH3 | 4.75E-29 | 1.146532829 | 0.264 | 0.129 | 9.36E-25 | Tb |
| allcell | KRT14 | 4.35E-121 | 1.145999854 | 0.61 | 0.241 | 8.56E-117 | Tb |
| allcell | RASAL2 | 4.80E-34 | 1.145954887 | 0.376 | 0.215 | 9.44E-30 | Tb |
| allcell | SUCNR1 | 4.39E-37 | 1.145887722 | 0.32 | 0.157 | 8.63E-33 | Tb |
| allcell | PRAG1 | 2.44E-36 | 1.145882197 | 0.259 | 0.113 | 4.79E-32 | Tb |
| allcell | STAT4 | 9.69E-47 | 1.143277509 | 0.506 | 0.3 | 1.91E-42 | Tb |
| allcell | PFKP | 1.21E-37 | 1.139913647 | 0.413 | 0.236 | 2.38E-33 | Tb |
| allcell | MOB3B | 1.59E-49 | 1.137632431 | 0.525 | 0.312 | 3.12E-45 | Tb |
| allcell | MT-ATP6 | 6.01E-159 | 1.136149733 | 0.991 | 0.897 | 1.18E-154 | Tb |
| allcell | PATL2 | 3.74E-44 | 1.130614217 | 0.286 | 0.118 | 7.36E-40 | Tb |
| allcell | CD109 | 1.25E-12 | 1.128001151 | 0.3 | 0.211 | 2.46E-08 | Tb |
| allcell | PDCD1LG2 | 6.42E-30 | 1.120880662 | 0.278 | 0.14 | 1.26E-25 | Tb |
| allcell | UBAP2 | 2.91E-54 | 1.119882991 | 0.442 | 0.217 | 5.73E-50 | Tb |
| allcell | DENND5B | 8.98E-32 | 1.119171569 | 0.299 | 0.152 | 1.77E-27 | Tb |
| allcell | PLEKHG2 | 8.79E-36 | 1.109121583 | 0.411 | 0.241 | 1.73E-31 | Tb |
| allcell | TFCP2 | 9.99E-52 | 1.103410859 | 0.41 | 0.201 | 1.97E-47 | Tb |
| allcell | RAP1GAP2 | 1.04E-27 | 1.099089722 | 0.26 | 0.13 | 2.04E-23 | Tb |
| allcell | RUNX2 | 1.96E-15 | 1.098945048 | 0.378 | 0.268 | 3.86E-11 | Tb |
| allcell | AZIN1-AS1 | 2.52E-29 | 1.09600538 | 0.297 | 0.155 | 4.95E-25 | Tb |
| allcell | LRCH3 | 1.12E-55 | 1.08522305 | 0.494 | 0.257 | 2.21E-51 | Tb |
| allcell | CREB3L2 | 5.24E-26 | 1.079980016 | 0.311 | 0.174 | 1.03E-21 | Tb |
| allcell | DUSP16 | 1.45E-40 | 1.078057317 | 0.359 | 0.178 | 2.85E-36 | Tb |
| allcell | DLEU2 | 2.38E-64 | 1.077658129 | 0.769 | 0.567 | 4.69E-60 | Tb |
| allcell | PPARD | 5.37E-40 | 1.072692816 | 0.474 | 0.279 | 1.06E-35 | Tb |
| allcell | EEF2K | 5.16E-42 | 1.072529702 | 0.365 | 0.18 | 1.02E-37 | Tb |
| allcell | KIF1B | 2.15E-53 | 1.072281106 | 0.542 | 0.318 | 4.23E-49 | Tb |
| allcell | MT-CO3 | 7.42E-117 | 1.069938541 | 0.978 | 0.911 | 1.46E-112 | Tb |
| allcell | LRRK1 | 2.51E-59 | 1.066001562 | 0.6 | 0.361 | 4.94E-55 | Tb |
| allcell | SPECC1L | 8.96E-45 | 1.065069884 | 0.387 | 0.192 | 1.76E-40 | Tb |
| allcell | PRKCA | 3.53E-51 | 1.065067153 | 0.492 | 0.272 | 6.95E-47 | Tb |
| allcell | CCDC138 | 1.05E-41 | 1.065028116 | 0.418 | 0.224 | 2.07E-37 | Tb |
| allcell | JMY | 2.92E-34 | 1.062585045 | 0.503 | 0.322 | 5.76E-30 | Tb |
| allcell | MCU | 2.07E-29 | 1.059693295 | 0.318 | 0.17 | 4.07E-25 | Tb |
| allcell | ZNF10 | 2.22E-49 | 1.058234886 | 0.446 | 0.229 | 4.37E-45 | Tb |
| allcell | TOGARAM1 | 4.21E-39 | 1.056850057 | 0.282 | 0.125 | 8.29E-35 | Tb |
| allcell | ENPP2 | 1.38E-39 | 1.056747898 | 0.407 | 0.219 | 2.72E-35 | Tb |
| allcell | TMEM184C | 3.36E-43 | 1.05484298 | 0.383 | 0.193 | 6.62E-39 | Tb |
| allcell | CARD11 | 1.42E-32 | 1.053448251 | 0.389 | 0.223 | 2.79E-28 | Tb |
| allcell | ANKRD28 | 1.93E-28 | 1.052900053 | 0.407 | 0.247 | 3.81E-24 | Tb |
| allcell | EBF4 | 4.00E-31 | 1.051829052 | 0.265 | 0.128 | 7.86E-27 | Tb |
| allcell | ARHGAP25 | 2.41E-40 | 1.049113217 | 0.516 | 0.32 | 4.75E-36 | Tb |
| allcell | CLEC2D | 2.83E-20 | 1.048716671 | 0.253 | 0.141 | 5.57E-16 | Tb |
| allcell | CASP10 | 1.20E-27 | 1.039011945 | 0.262 | 0.132 | 2.36E-23 | Tb |
| allcell | IL1RAP | 7.87E-32 | 1.036824868 | 0.293 | 0.146 | 1.55E-27 | Tb |
| allcell | PALD1 | 3.27E-49 | 1.034494545 | 0.383 | 0.179 | 6.43E-45 | Tb |
| allcell | SFMBT2 | 5.28E-59 | 1.029381231 | 0.635 | 0.384 | 1.04E-54 | Tb |
| allcell | KRT16 | 4.83E-67 | 1.028505044 | 0.351 | 0.123 | 9.51E-63 | Tb |
| allcell | DIPK1A | 1.41E-31 | 1.025002946 | 0.351 | 0.193 | 2.77E-27 | Tb |
| allcell | NAA25 | 1.37E-21 | 1.019515986 | 0.33 | 0.198 | 2.70E-17 | Tb |

| | | | | | | | |
|---|---|---|---|---|---|---|---|
| allcell | ZNF516 | 2.85E-46 | 1.019412106 | 0.519 | 0.306 | 5.60E-42 | Tb |
| allcell | MGA | 1.12E-32 | 1.018908472 | 0.318 | 0.163 | 2.21E-28 | Tb |
| allcell | RPRD2 | 1.41E-25 | 1.017316588 | 0.267 | 0.14 | 2.78E-21 | Tb |
| allcell | CEP192 | 1.11E-38 | 1.016303543 | 0.311 | 0.146 | 2.19E-34 | Tb |
| allcell | SMG6 | 3.61E-45 | 1.016106777 | 0.388 | 0.19 | 7.10E-41 | Tb |
| allcell | NSMCE2 | 9.07E-67 | 1.009968912 | 0.678 | 0.418 | 1.78E-62 | Tb |
| allcell | CLCN3 | 3.58E-25 | 1.005874127 | 0.355 | 0.208 | 7.04E-21 | Tb |
| allcell | CBLB | 6.13E-41 | 1.005300284 | 0.411 | 0.226 | 1.21E-36 | Tb |
| allcell | RASA1 | 1.45E-42 | 1.002037116 | 0.387 | 0.194 | 2.86E-38 | Tb |
| allcell | WDR37 | 1.86E-50 | 1.000243826 | 0.476 | 0.252 | 3.67E-46 | Tb |
| allcell | N4BP2 | 6.36E-45 | 1.000157789 | 0.507 | 0.291 | 1.25E-40 | Tb |
| allcell | MT-CYB | 2.77E-89 | 0.997374237 | 0.963 | 0.887 | 5.46E-85 | Tb |
| allcell | UST | 3.71E-44 | 0.992180694 | 0.615 | 0.419 | 7.31E-40 | Tb |
| allcell | ARID2 | 6.33E-50 | 0.99172258 | 0.612 | 0.39 | 1.25E-45 | Tb |
| allcell | PLPP1 | 3.39E-26 | 0.989673522 | 0.371 | 0.219 | 6.68E-22 | Tb |
| allcell | BMPR2 | 4.12E-43 | 0.987702276 | 0.43 | 0.232 | 8.11E-39 | Tb |
| allcell | SPATA13 | 4.56E-29 | 0.984520167 | 0.307 | 0.162 | 8.98E-25 | Tb |
| allcell | PHLPP2 | 6.33E-27 | 0.982012799 | 0.295 | 0.157 | 1.25E-22 | Tb |
| allcell | CRY1 | 1.59E-27 | 0.980246765 | 0.297 | 0.16 | 3.14E-23 | Tb |
| allcell | CTDP1 | 5.77E-45 | 0.979449284 | 0.376 | 0.183 | 1.14E-40 | Tb |
| allcell | LRRC23 | 9.42E-34 | 0.978653328 | 0.421 | 0.246 | 1.85E-29 | Tb |
| allcell | DENND4A | 2.30E-68 | 0.975154906 | 0.852 | 0.67 | 4.52E-64 | Tb |
| allcell | B3GNT5 | 3.17E-28 | 0.974605434 | 0.545 | 0.386 | 6.24E-24 | Tb |
| allcell | P2RY14 | 1.42E-44 | 0.973971583 | 0.614 | 0.4 | 2.79E-40 | Tb |
| allcell | SPG11 | 6.01E-46 | 0.972873629 | 0.498 | 0.287 | 1.18E-41 | Tb |
| allcell | FUT8 | 1.67E-23 | 0.971047155 | 0.323 | 0.19 | 3.29E-19 | Tb |
| allcell | IQCG | 1.68E-27 | 0.970324771 | 0.26 | 0.131 | 3.30E-23 | Tb |
| allcell | GALNT11 | 1.04E-29 | 0.970213652 | 0.341 | 0.184 | 2.05E-25 | Tb |
| allcell | SPRED1 | 1.45E-21 | 0.969420958 | 0.275 | 0.156 | 2.84E-17 | Tb |
| allcell | THADA | 1.11E-37 | 0.96921633 | 0.378 | 0.197 | 2.19E-33 | Tb |
| allcell | ZNF366 | 3.25E-30 | 0.965501315 | 0.346 | 0.191 | 6.39E-26 | Tb |
| allcell | STIM1 | 1.38E-40 | 0.963850072 | 0.387 | 0.2 | 2.71E-36 | Tb |
| allcell | KSR1 | 3.32E-28 | 0.961873069 | 0.282 | 0.147 | 6.54E-24 | Tb |
| allcell | PAG1 | 7.65E-33 | 0.959970596 | 0.491 | 0.309 | 1.51E-28 | Tb |
| allcell | PHF19 | 4.26E-14 | 0.959067262 | 0.408 | 0.305 | 8.38E-10 | Tb |
| allcell | WWOX | 1.50E-25 | 0.958171208 | 0.395 | 0.249 | 2.94E-21 | Tb |
| allcell | AXL | 8.32E-31 | 0.958112195 | 0.4 | 0.237 | 1.64E-26 | Tb |
| allcell | RALGAPA1 | 5.53E-42 | 0.957338076 | 0.656 | 0.449 | 1.09E-37 | Tb |
| allcell | CEP83 | 4.61E-39 | 0.95562435 | 0.469 | 0.272 | 9.07E-35 | Tb |
| allcell | KDM6A | 1.26E-66 | 0.955416955 | 0.662 | 0.406 | 2.48E-62 | Tb |
| allcell | ARIH1 | 7.39E-67 | 0.95467437 | 0.861 | 0.667 | 1.46E-62 | Tb |
| allcell | RASGRP3 | 3.23E-31 | 0.953046942 | 0.387 | 0.225 | 6.36E-27 | Tb |
| allcell | MED12L | 4.10E-28 | 0.950543421 | 0.285 | 0.146 | 8.07E-24 | Tb |
| allcell | RAB30 | 1.06E-40 | 0.950219795 | 0.449 | 0.254 | 2.09E-36 | Tb |
| allcell | ABL1 | 1.07E-33 | 0.949901468 | 0.373 | 0.208 | 2.10E-29 | Tb |
| allcell | SENP2 | 4.98E-32 | 0.9490928 | 0.281 | 0.135 | 9.80E-28 | Tb |
| allcell | GALC | 2.17E-27 | 0.948747136 | 0.33 | 0.186 | 4.27E-23 | Tb |
| allcell | MX2 | 7.25E-20 | 0.948012483 | 0.457 | 0.325 | 1.43E-15 | Tb |
| allcell | HDAC4 | 3.14E-38 | 0.94532414 | 0.362 | 0.182 | 6.18E-34 | Tb |
| allcell | MSI2 | 1.07E-28 | 0.942545544 | 0.355 | 0.202 | 2.11E-24 | Tb |
| allcell | TBC1D17 | 9.44E-33 | 0.941805137 | 0.263 | 0.121 | 1.86E-28 | Tb |
| allcell | CAMK1D | 4.45E-53 | 0.940498118 | 0.732 | 0.526 | 8.76E-49 | Tb |
| allcell | CD80 | 4.73E-23 | 0.940416432 | 0.434 | 0.294 | 9.31E-19 | Tb |
| allcell | MBTD1 | 3.88E-32 | 0.940109901 | 0.318 | 0.164 | 7.63E-28 | Tb |
| allcell | TASP1 | 5.17E-25 | 0.937203566 | 0.29 | 0.157 | 1.02E-20 | Tb |
| allcell | SLC39A11 | 1.46E-43 | 0.936955031 | 0.406 | 0.208 | 2.88E-39 | Tb |

| | | | | | | | |
|---|---|---|---|---|---|---|---|
| allcell | TMTC2 | 3.25E-25 | 0.935142434 | 0.3 | 0.167 | 6.40E-21 | Tb |
| allcell | SCAF8 | 1.17E-48 | 0.934722982 | 0.426 | 0.209 | 2.30E-44 | Tb |
| allcell | UBXN2A | 1.59E-38 | 0.934013349 | 0.497 | 0.304 | 3.14E-34 | Tb |
| allcell | LRRC8D | 2.85E-41 | 0.932763784 | 0.419 | 0.228 | 5.60E-37 | Tb |
| allcell | ST6GAL1 | 2.55E-45 | 0.932072073 | 0.468 | 0.257 | 5.02E-41 | Tb |
| allcell | TOP3A | 1.58E-32 | 0.931743874 | 0.266 | 0.122 | 3.11E-28 | Tb |
| allcell | PPP1R16B | 4.53E-36 | 0.931123828 | 0.43 | 0.248 | 8.91E-32 | Tb |
| allcell | POLR2J3 | 1.21E-39 | 0.929966257 | 0.344 | 0.166 | 2.38E-35 | Tb |
| allcell | MRC1 | 6.13E-14 | 0.928602263 | 0.337 | 0.238 | 1.21E-09 | Tb |
| allcell | NT5C2 | 1.51E-49 | 0.926648566 | 0.54 | 0.303 | 2.98E-45 | Tb |
| allcell | ARAP2 | 3.95E-44 | 0.924594333 | 0.553 | 0.342 | 7.78E-40 | Tb |
| allcell | FAM13A | 2.08E-27 | 0.923854607 | 0.301 | 0.161 | 4.09E-23 | Tb |
| allcell | VAV1 | 6.78E-38 | 0.923464936 | 0.4 | 0.215 | 1.34E-33 | Tb |
| allcell | CERS5 | 3.35E-36 | 0.922682378 | 0.321 | 0.154 | 6.60E-32 | Tb |
| allcell | AKAP10 | 1.56E-30 | 0.920729357 | 0.27 | 0.13 | 3.08E-26 | Tb |
| allcell | SETBP1 | 5.89E-19 | 0.920291083 | 0.266 | 0.155 | 1.16E-14 | Tb |
| allcell | NFX1 | 1.81E-34 | 0.918284436 | 0.456 | 0.268 | 3.57E-30 | Tb |
| allcell | KLF7 | 9.09E-31 | 0.912605722 | 0.402 | 0.235 | 1.79E-26 | Tb |
| allcell | STEAP1B | 1.23E-18 | 0.910661623 | 0.394 | 0.256 | 2.42E-14 | Tb |
| allcell | SGPL1 | 1.38E-31 | 0.909999254 | 0.302 | 0.151 | 2.72E-27 | Tb |
| allcell | SYTL3 | 1.95E-26 | 0.908965959 | 0.374 | 0.227 | 3.83E-22 | Tb |
| allcell | TSPOAP1 | 1.08E-27 | 0.907179319 | 0.253 | 0.124 | 2.12E-23 | Tb |
| allcell | ZHX2 | 3.53E-44 | 0.906610915 | 0.603 | 0.381 | 6.94E-40 | Tb |
| allcell | CTNND1 | 3.41E-35 | 0.903611649 | 0.482 | 0.291 | 6.71E-31 | Tb |
| allcell | CMSS1 | 4.96E-12 | 0.901827615 | 0.42 | 0.307 | 9.77E-08 | Tb |
| allcell | GMDS-DT | 2.62E-28 | 0.901470541 | 0.383 | 0.232 | 5.16E-24 | Tb |
| allcell | USP25 | 7.96E-34 | 0.901102779 | 0.459 | 0.276 | 1.57E-29 | Tb |
| allcell | KRIT1 | 3.68E-41 | 0.898618399 | 0.362 | 0.175 | 7.25E-37 | Tb |
| allcell | SLC43A2 | 2.40E-36 | 0.897994747 | 0.547 | 0.355 | 4.72E-32 | Tb |
| allcell | ZNF33A | 2.34E-30 | 0.897001434 | 0.373 | 0.21 | 4.60E-26 | Tb |
| allcell | SESN1 | 1.37E-24 | 0.89688921 | 0.321 | 0.181 | 2.70E-20 | Tb |
| allcell | FAF2 | 6.64E-34 | 0.896828378 | 0.304 | 0.147 | 1.31E-29 | Tb |
| allcell | VPS13B | 2.65E-54 | 0.893049117 | 0.607 | 0.369 | 5.22E-50 | Tb |
| allcell | ADAP2 | 2.62E-27 | 0.892168872 | 0.361 | 0.208 | 5.16E-23 | Tb |
| allcell | BCL11A | 8.92E-51 | 0.892022324 | 0.659 | 0.431 | 1.76E-46 | Tb |
| allcell | STX17-AS1 | 2.30E-24 | 0.891381408 | 0.467 | 0.308 | 4.52E-20 | Tb |
| allcell | SMYD3 | 4.99E-23 | 0.89019075 | 0.341 | 0.208 | 9.82E-19 | Tb |
| allcell | FTX | 1.05E-60 | 0.889706637 | 0.596 | 0.335 | 2.07E-56 | Tb |
| allcell | SRGAP2 | 4.28E-59 | 0.886970288 | 0.621 | 0.373 | 8.42E-55 | Tb |
| allcell | CDK19 | 7.32E-33 | 0.885425628 | 0.373 | 0.203 | 1.44E-28 | Tb |
| allcell | WDPCP | 2.07E-28 | 0.884835107 | 0.282 | 0.142 | 4.07E-24 | Tb |
| allcell | TMCC1 | 7.87E-34 | 0.883778772 | 0.383 | 0.212 | 1.55E-29 | Tb |
| allcell | PRRG4 | 9.40E-28 | 0.882097825 | 0.307 | 0.165 | 1.85E-23 | Tb |
| allcell | SIK2 | 1.08E-28 | 0.881414835 | 0.446 | 0.278 | 2.13E-24 | Tb |
| allcell | RNF19A | 1.36E-40 | 0.88036768 | 0.549 | 0.343 | 2.67E-36 | Tb |
| allcell | TRAK1 | 1.09E-47 | 0.87993297 | 0.667 | 0.457 | 2.14E-43 | Tb |
| allcell | PITPNA | 7.82E-47 | 0.878182687 | 0.672 | 0.475 | 1.54E-42 | Tb |
| allcell | TAOK1 | 3.36E-45 | 0.877573741 | 0.538 | 0.309 | 6.62E-41 | Tb |
| allcell | FTO | 4.70E-32 | 0.874309142 | 0.297 | 0.145 | 9.24E-28 | Tb |
| allcell | ITCH | 4.21E-43 | 0.873904286 | 0.519 | 0.296 | 8.29E-39 | Tb |
| allcell | NBEAL1 | 5.71E-35 | 0.872741129 | 0.669 | 0.474 | 1.12E-30 | Tb |
| allcell | PRMT9 | 7.79E-44 | 0.87244899 | 0.618 | 0.401 | 1.53E-39 | Tb |
| allcell | MT-ND4 | 3.67E-132 | 0.871777147 | 0.989 | 0.933 | 7.23E-128 | Tb |
| allcell | DENND1B | 7.03E-57 | 0.869186896 | 0.776 | 0.584 | 1.38E-52 | Tb |
| allcell | BTBD7 | 2.46E-28 | 0.868572027 | 0.524 | 0.346 | 4.84E-24 | Tb |
| allcell | KANSL1 | 2.61E-50 | 0.866920159 | 0.533 | 0.299 | 5.14E-46 | Tb |

| | | | | | | | |
|---|---|---|---|---|---|---|---|
| allcell | MREG | 5.86E-26 | 0.866154102 | 0.381 | 0.23 | 1.15E-21 | Tb |
| allcell | BRAF | 5.78E-56 | 0.865232682 | 0.738 | 0.501 | 1.14E-51 | Tb |
| allcell | ELL2 | 6.30E-48 | 0.864367655 | 0.711 | 0.51 | 1.24E-43 | Tb |
| allcell | SPRED2 | 4.14E-29 | 0.863588082 | 0.544 | 0.381 | 8.15E-25 | Tb |
| allcell | CCDC97 | 5.68E-29 | 0.863358203 | 0.308 | 0.16 | 1.12E-24 | Tb |
| allcell | RAB30-DT | 2.75E-40 | 0.862446752 | 0.339 | 0.158 | 5.42E-36 | Tb |
| allcell | SEC23B | 1.99E-37 | 0.862283972 | 0.437 | 0.245 | 3.91E-33 | Tb |
| allcell | ZFAT | 2.78E-14 | 0.861994844 | 0.271 | 0.176 | 5.48E-10 | Tb |
| allcell | CSNK1G1 | 4.18E-33 | 0.861888437 | 0.338 | 0.177 | 8.23E-29 | Tb |
| allcell | IRS2 | 4.15E-21 | 0.861171548 | 0.273 | 0.155 | 8.16E-17 | Tb |
| allcell | VASH1 | 2.55E-40 | 0.861167825 | 0.433 | 0.235 | 5.03E-36 | Tb |
| allcell | FOXK2 | 1.08E-33 | 0.858707455 | 0.414 | 0.23 | 2.12E-29 | Tb |
| allcell | SCLT1 | 1.06E-49 | 0.858534299 | 0.711 | 0.494 | 2.08E-45 | Tb |
| allcell | STIM2 | 6.14E-39 | 0.855838077 | 0.5 | 0.303 | 1.21E-34 | Tb |
| allcell | NAF1 | 4.70E-17 | 0.854791933 | 0.459 | 0.331 | 9.25E-13 | Tb |
| allcell | FAF1 | 2.39E-47 | 0.854477289 | 0.596 | 0.374 | 4.71E-43 | Tb |
| allcell | GCNT1 | 6.84E-24 | 0.854095129 | 0.265 | 0.14 | 1.35E-19 | Tb |
| allcell | PAN3 | 4.96E-48 | 0.853028393 | 0.633 | 0.402 | 9.76E-44 | Tb |
| allcell | FBXL17 | 8.76E-41 | 0.850412536 | 0.502 | 0.292 | 1.73E-36 | Tb |
| allcell | WDR59 | 1.14E-30 | 0.849896388 | 0.336 | 0.177 | 2.25E-26 | Tb |
| allcell | GRAMD1B | 4.17E-27 | 0.849824504 | 0.372 | 0.222 | 8.21E-23 | Tb |
| allcell | TBC1D22A | 1.01E-54 | 0.849768521 | 0.704 | 0.489 | 1.99E-50 | Tb |
| allcell | ABTB2 | 1.17E-19 | 0.849573484 | 0.448 | 0.31 | 2.30E-15 | Tb |
| allcell | NRF1 | 5.43E-28 | 0.847879883 | 0.333 | 0.182 | 1.07E-23 | Tb |
| allcell | ILF3-DT | 2.26E-32 | 0.847409792 | 0.304 | 0.149 | 4.45E-28 | Tb |
| allcell | ABHD18 | 5.62E-20 | 0.847405371 | 0.25 | 0.137 | 1.11E-15 | Tb |
| allcell | SPDYA | 1.13E-19 | 0.84657305 | 0.283 | 0.169 | 2.22E-15 | Tb |
| allcell | SREBF2 | 3.91E-38 | 0.845193272 | 0.376 | 0.19 | 7.69E-34 | Tb |
| allcell | TNS3 | 2.00E-40 | 0.843596868 | 0.441 | 0.245 | 3.93E-36 | Tb |
| allcell | RAPGEF2 | 2.34E-35 | 0.843400361 | 0.693 | 0.517 | 4.61E-31 | Tb |
| allcell | LIMS1 | 1.46E-48 | 0.842945032 | 0.765 | 0.613 | 2.87E-44 | Tb |
| allcell | METTL16 | 8.76E-24 | 0.842200306 | 0.253 | 0.131 | 1.72E-19 | Tb |
| allcell | DAAM1 | 2.20E-21 | 0.841593536 | 0.348 | 0.213 | 4.32E-17 | Tb |
| allcell | ADNP2 | 7.95E-31 | 0.840837491 | 0.314 | 0.161 | 1.56E-26 | Tb |
| allcell | IPCEF1 | 1.24E-14 | 0.839760513 | 0.267 | 0.171 | 2.44E-10 | Tb |
| allcell | UBE4B | 1.08E-24 | 0.83873148 | 0.254 | 0.13 | 2.12E-20 | Tb |
| allcell | BABAM2 | 6.70E-38 | 0.838240217 | 0.569 | 0.375 | 1.32E-33 | Tb |
| allcell | MFSD14C | 5.01E-45 | 0.837067881 | 0.463 | 0.248 | 9.86E-41 | Tb |
| allcell | AHCYL2 | 1.60E-28 | 0.836206018 | 0.376 | 0.221 | 3.16E-24 | Tb |
| allcell | SKI | 1.34E-29 | 0.836038089 | 0.35 | 0.19 | 2.63E-25 | Tb |
| allcell | ITFG1 | 2.28E-40 | 0.83546699 | 0.523 | 0.316 | 4.49E-36 | Tb |
| allcell | RABGAP1L | 3.20E-41 | 0.833611047 | 0.768 | 0.57 | 6.30E-37 | Tb |
| allcell | CASS4 | 8.89E-27 | 0.833427189 | 0.37 | 0.217 | 1.75E-22 | Tb |
| allcell | ARHGAP10 | 5.59E-22 | 0.83271575 | 0.422 | 0.283 | 1.10E-17 | Tb |
| allcell | TNRC18 | 6.01E-33 | 0.832561057 | 0.417 | 0.237 | 1.18E-28 | Tb |
| allcell | LATS2 | 1.85E-20 | 0.831830251 | 0.341 | 0.21 | 3.64E-16 | Tb |
| allcell | RGS12 | 4.54E-31 | 0.831362434 | 0.324 | 0.17 | 8.94E-27 | Tb |
| allcell | S100PBP | 6.05E-24 | 0.830115537 | 0.253 | 0.131 | 1.19E-19 | Tb |
| allcell | RNF122 | 1.66E-29 | 0.829825482 | 0.269 | 0.132 | 3.27E-25 | Tb |
| allcell | ASB3 | 1.57E-24 | 0.829740328 | 0.379 | 0.226 | 3.08E-20 | Tb |
| allcell | LRBA | 1.94E-39 | 0.827591474 | 0.586 | 0.375 | 3.82E-35 | Tb |
| allcell | NFATC3 | 3.68E-25 | 0.827415996 | 0.314 | 0.172 | 7.25E-21 | Tb |
| allcell | ANAPC10 | 4.27E-32 | 0.826576779 | 0.404 | 0.225 | 8.40E-28 | Tb |
| allcell | ELMO1 | 4.31E-53 | 0.826571 | 0.816 | 0.668 | 8.49E-49 | Tb |
| allcell | STAU2 | 6.44E-32 | 0.82365668 | 0.336 | 0.175 | 1.27E-27 | Tb |
| allcell | PCYT1A | 1.40E-37 | 0.823638704 | 0.467 | 0.274 | 2.76E-33 | Tb |

| | | | | | | | |
|---|---|---|---|---|---|---|---|
| allcell | VPS54 | 2.16E-20 | 0.822674218 | 0.274 | 0.155 | 4.25E-16 | Tb |
| allcell | S100A9 | 6.61E-114 | 0.82226275 | 0.672 | 0.31 | 1.30E-109 | Tb |
| allcell | ALPK1 | 5.39E-36 | 0.821617554 | 0.453 | 0.268 | 1.06E-31 | Tb |
| allcell | ZBTB20 | 7.68E-46 | 0.821167935 | 0.619 | 0.406 | 1.51E-41 | Tb |
| allcell | GALNT7 | 2.31E-30 | 0.821104241 | 0.395 | 0.226 | 4.55E-26 | Tb |
| allcell | ATG7 | 5.63E-46 | 0.821055625 | 0.509 | 0.29 | 1.11E-41 | Tb |
| allcell | TSPAN14 | 2.46E-25 | 0.820224163 | 0.254 | 0.128 | 4.84E-21 | Tb |
| allcell | MFSD14B | 5.57E-33 | 0.82019518 | 0.457 | 0.272 | 1.10E-28 | Tb |
| allcell | MFN1 | 6.96E-32 | 0.819975226 | 0.363 | 0.195 | 1.37E-27 | Tb |
| allcell | PBRM1 | 4.64E-43 | 0.817571328 | 0.686 | 0.461 | 9.13E-39 | Tb |
| allcell | NOL10 | 4.21E-23 | 0.817499949 | 0.282 | 0.154 | 8.28E-19 | Tb |
| allcell | TET2 | 5.35E-30 | 0.817439218 | 0.825 | 0.72 | 1.05E-25 | Tb |
| allcell | CASP8 | 6.79E-40 | 0.815964018 | 0.547 | 0.331 | 1.34E-35 | Tb |
| allcell | ST3GAL2 | 1.83E-40 | 0.815803595 | 0.412 | 0.217 | 3.60E-36 | Tb |
| allcell | UBR1 | 2.04E-32 | 0.814736174 | 0.426 | 0.247 | 4.02E-28 | Tb |
| allcell | SLC12A2 | 1.44E-24 | 0.814181017 | 0.337 | 0.194 | 2.83E-20 | Tb |
| allcell | EVI5 | 2.84E-32 | 0.81398135 | 0.463 | 0.281 | 5.59E-28 | Tb |
| allcell | CDK6 | 1.84E-25 | 0.81330721 | 0.535 | 0.375 | 3.63E-21 | Tb |
| allcell | ESYT2 | 2.65E-45 | 0.813116873 | 0.781 | 0.628 | 5.21E-41 | Tb |
| allcell | RREB1 | 4.92E-37 | 0.810672666 | 0.561 | 0.355 | 9.68E-33 | Tb |
| allcell | SRGAP2C | 2.44E-36 | 0.810201283 | 0.352 | 0.181 | 4.80E-32 | Tb |
| allcell | ERCC6L2 | 8.25E-22 | 0.809057874 | 0.261 | 0.141 | 1.62E-17 | Tb |
| allcell | LPIN1 | 6.38E-20 | 0.808870573 | 0.392 | 0.252 | 1.26E-15 | Tb |
| allcell | SLC30A7 | 8.44E-27 | 0.80885964 | 0.332 | 0.182 | 1.66E-22 | Tb |
| allcell | RNF111 | 5.36E-33 | 0.808793807 | 0.43 | 0.25 | 1.05E-28 | Tb |
| allcell | FAM214A | 1.68E-27 | 0.80800806 | 0.33 | 0.182 | 3.30E-23 | Tb |
| allcell | CERS6 | 1.11E-44 | 0.807989452 | 0.735 | 0.521 | 2.18E-40 | Tb |
| allcell | ABR | 1.01E-34 | 0.807834708 | 0.659 | 0.479 | 2.00E-30 | Tb |
| allcell | POGZ | 1.24E-38 | 0.807641482 | 0.446 | 0.247 | 2.44E-34 | Tb |
| allcell | EPB41L2 | 1.11E-33 | 0.805894293 | 0.565 | 0.39 | 2.19E-29 | Tb |
| allcell | RFX7 | 1.27E-15 | 0.803487185 | 0.253 | 0.155 | 2.50E-11 | Tb |
| allcell | ARHGEF3 | 1.07E-30 | 0.802030053 | 0.353 | 0.192 | 2.11E-26 | Tb |
| allcell | CIITA | 3.17E-47 | 0.8010329 | 0.81 | 0.651 | 6.24E-43 | Tb |
| allcell | TPT1-AS1 | 1.62E-31 | 0.800832294 | 0.253 | 0.113 | 3.18E-27 | Tb |
| allcell | SAMD9L | 2.69E-47 | 0.799713197 | 0.528 | 0.298 | 5.30E-43 | Tb |
| allcell | CHML | 3.14E-21 | 0.799385033 | 0.357 | 0.218 | 6.17E-17 | Tb |
| allcell | MSL2 | 1.58E-33 | 0.798925874 | 0.368 | 0.194 | 3.11E-29 | Tb |
| allcell | ZCCHC7 | 1.70E-42 | 0.798100795 | 0.584 | 0.374 | 3.34E-38 | Tb |
| allcell | JAK3 | 1.68E-26 | 0.798011213 | 0.286 | 0.15 | 3.30E-22 | Tb |
| allcell | SLC20A2 | 1.46E-24 | 0.797380156 | 0.279 | 0.149 | 2.86E-20 | Tb |
| allcell | CCDC91 | 3.42E-45 | 0.797247156 | 0.639 | 0.408 | 6.72E-41 | Tb |
| allcell | RBM6 | 2.64E-43 | 0.796838897 | 0.614 | 0.393 | 5.20E-39 | Tb |
| allcell | EZH2 | 3.55E-49 | 0.796722304 | 0.663 | 0.438 | 6.98E-45 | Tb |
| allcell | TMEM39A | 9.28E-31 | 0.796434064 | 0.527 | 0.35 | 1.83E-26 | Tb |
| allcell | PPP3CC | 1.85E-31 | 0.795704979 | 0.402 | 0.228 | 3.64E-27 | Tb |
| allcell | UBL3 | 9.81E-36 | 0.794249093 | 0.671 | 0.48 | 1.93E-31 | Tb |
| allcell | ABCC1 | 1.13E-39 | 0.794092156 | 0.397 | 0.205 | 2.21E-35 | Tb |
| allcell | MYO9A | 1.43E-40 | 0.793836999 | 0.475 | 0.276 | 2.81E-36 | Tb |
| allcell | CLCN5 | 6.73E-29 | 0.793358394 | 0.412 | 0.25 | 1.33E-24 | Tb |
| allcell | DDX58 | 1.08E-20 | 0.792190795 | 0.251 | 0.137 | 2.12E-16 | Tb |
| allcell | CFLAR-AS1 | 2.82E-33 | 0.792047382 | 0.36 | 0.19 | 5.55E-29 | Tb |
| allcell | SLMAP | 2.31E-37 | 0.791945617 | 0.409 | 0.218 | 4.54E-33 | Tb |
| allcell | PPCDC | 1.08E-23 | 0.791938356 | 0.362 | 0.217 | 2.13E-19 | Tb |
| allcell | RERE | 1.16E-40 | 0.791212742 | 0.717 | 0.528 | 2.28E-36 | Tb |
| allcell | LRRC8C | 1.40E-22 | 0.791101193 | 0.385 | 0.244 | 2.76E-18 | Tb |
| allcell | GATAD2B | 3.18E-45 | 0.790571003 | 0.53 | 0.309 | 6.26E-41 | Tb |

| | | | | | | | |
|---|---|---|---|---|---|---|---|
| allcell | MAP3K14 | 6.47E-26 | 0.790331979 | 0.431 | 0.269 | 1.27E-21 | Tb |
| allcell | CHKA | 2.37E-25 | 0.789881679 | 0.421 | 0.265 | 4.66E-21 | Tb |
| allcell | TYW1B | 5.10E-30 | 0.787597786 | 0.283 | 0.141 | 1.00E-25 | Tb |
| allcell | GPHN | 8.18E-21 | 0.786207613 | 0.337 | 0.206 | 1.61E-16 | Tb |
| allcell | SLC7A6 | 4.33E-23 | 0.785949498 | 0.281 | 0.154 | 8.53E-19 | Tb |
| allcell | P2RY6 | 1.36E-18 | 0.785861267 | 0.4 | 0.27 | 2.68E-14 | Tb |
| allcell | SNRK | 1.69E-21 | 0.784081056 | 0.294 | 0.168 | 3.33E-17 | Tb |
| allcell | CD2AP | 1.15E-27 | 0.78349371 | 0.489 | 0.316 | 2.26E-23 | Tb |
| allcell | TCF4 | 2.60E-25 | 0.783448817 | 0.652 | 0.482 | 5.12E-21 | Tb |
| allcell | ITPR1 | 4.32E-46 | 0.782006793 | 0.57 | 0.349 | 8.50E-42 | Tb |
| allcell | SLC9A7 | 6.92E-30 | 0.781855973 | 0.564 | 0.389 | 1.36E-25 | Tb |
| allcell | UBE2O | 4.40E-25 | 0.781227076 | 0.295 | 0.158 | 8.65E-21 | Tb |
| allcell | BICRAL | 2.22E-33 | 0.781063899 | 0.29 | 0.137 | 4.37E-29 | Tb |
| allcell | DCAF10 | 2.15E-22 | 0.780706745 | 0.272 | 0.148 | 4.23E-18 | Tb |
| allcell | CDKAL1 | 7.27E-38 | 0.779889281 | 0.476 | 0.281 | 1.43E-33 | Tb |
| allcell | PHTF2 | 1.06E-32 | 0.779498651 | 0.462 | 0.279 | 2.08E-28 | Tb |
| allcell | AP1S3 | 4.43E-29 | 0.778430397 | 0.452 | 0.286 | 8.71E-25 | Tb |
| allcell | ARFGEF2 | 6.21E-38 | 0.77830805 | 0.432 | 0.237 | 1.22E-33 | Tb |
| allcell | WIPF2 | 3.91E-28 | 0.778083865 | 0.285 | 0.146 | 7.70E-24 | Tb |
| allcell | FAM168A | 5.27E-30 | 0.778010012 | 0.335 | 0.179 | 1.04E-25 | Tb |
| allcell | SOAT1 | 9.11E-26 | 0.777873784 | 0.393 | 0.237 | 1.79E-21 | Tb |
| allcell | LAMP3 | 1.91E-20 | 0.777552485 | 0.468 | 0.327 | 3.76E-16 | Tb |
| allcell | APOL2 | 4.82E-27 | 0.776954501 | 0.279 | 0.142 | 9.49E-23 | Tb |
| allcell | ERBIN | 8.77E-39 | 0.776667218 | 0.645 | 0.446 | 1.73E-34 | Tb |
| allcell | RNF38 | 1.03E-32 | 0.776633389 | 0.439 | 0.259 | 2.03E-28 | Tb |
| allcell | GRK3 | 3.26E-39 | 0.776486707 | 0.663 | 0.466 | 6.41E-35 | Tb |
| allcell | ABL2 | 2.93E-33 | 0.776272195 | 0.516 | 0.332 | 5.76E-29 | Tb |
| allcell | LYRM1 | 1.46E-24 | 0.775865638 | 0.268 | 0.138 | 2.88E-20 | Tb |
| allcell | CBFB | 2.67E-42 | 0.775561372 | 0.469 | 0.258 | 5.26E-38 | Tb |
| allcell | MALT1 | 5.67E-33 | 0.774803649 | 0.804 | 0.676 | 1.12E-28 | Tb |
| allcell | FBXL20 | 3.42E-22 | 0.773074941 | 0.301 | 0.173 | 6.74E-18 | Tb |
| allcell | ZCCHC2 | 3.02E-30 | 0.772839745 | 0.414 | 0.241 | 5.95E-26 | Tb |
| allcell | ALOX12-AS1 | 4.13E-26 | 0.77167239 | 0.255 | 0.127 | 8.14E-22 | Tb |
| allcell | KDM5B | 1.22E-25 | 0.771586074 | 0.344 | 0.197 | 2.40E-21 | Tb |
| allcell | QRICH1 | 7.33E-30 | 0.771318058 | 0.316 | 0.164 | 1.44E-25 | Tb |
| allcell | UBAC2 | 4.53E-41 | 0.770521335 | 0.719 | 0.528 | 8.91E-37 | Tb |
| allcell | GSK3B | 1.50E-40 | 0.77023073 | 0.595 | 0.392 | 2.96E-36 | Tb |
| allcell | ATR | 2.30E-32 | 0.769855797 | 0.402 | 0.226 | 4.52E-28 | Tb |
| allcell | HIP1 | 3.47E-30 | 0.76839631 | 0.584 | 0.42 | 6.83E-26 | Tb |
| allcell | P2RY8 | 8.94E-20 | 0.767217074 | 0.254 | 0.144 | 1.76E-15 | Tb |
| allcell | FCHSD2 | 1.24E-46 | 0.766914676 | 0.847 | 0.672 | 2.45E-42 | Tb |
| allcell | NPC1 | 6.69E-24 | 0.766122095 | 0.359 | 0.212 | 1.32E-19 | Tb |
| allcell | PDE7A | 7.55E-36 | 0.763724891 | 0.667 | 0.474 | 1.49E-31 | Tb |
| allcell | CD226 | 9.30E-22 | 0.763615088 | 0.353 | 0.221 | 1.83E-17 | Tb |
| allcell | MTHFR | 6.71E-29 | 0.762993236 | 0.406 | 0.235 | 1.32E-24 | Tb |
| allcell | HS3ST3B1 | 2.13E-18 | 0.762699694 | 0.31 | 0.193 | 4.19E-14 | Tb |
| allcell | TERF1 | 4.12E-19 | 0.762190464 | 0.355 | 0.22 | 8.11E-15 | Tb |
| allcell | OXNAD1 | 1.47E-28 | 0.761987809 | 0.28 | 0.14 | 2.89E-24 | Tb |
| allcell | DMXL1 | 1.42E-37 | 0.761299868 | 0.541 | 0.332 | 2.79E-33 | Tb |
| allcell | INPP5D | 1.23E-23 | 0.759977364 | 0.509 | 0.351 | 2.42E-19 | Tb |
| allcell | XPO1 | 3.87E-47 | 0.759507724 | 0.684 | 0.45 | 7.62E-43 | Tb |
| allcell | CDK17 | 5.73E-28 | 0.758998261 | 0.445 | 0.277 | 1.13E-23 | Tb |
| allcell | UTRN | 3.68E-36 | 0.75890789 | 0.663 | 0.462 | 7.24E-32 | Tb |
| allcell | TXLNA | 1.99E-28 | 0.7575859 | 0.33 | 0.176 | 3.91E-24 | Tb |
| allcell | ABHD3 | 2.26E-30 | 0.757151393 | 0.56 | 0.383 | 4.45E-26 | Tb |
| allcell | EYA3 | 2.49E-31 | 0.756839574 | 0.383 | 0.213 | 4.90E-27 | Tb |

| | | | | | | | |
|---|---|---|---|---|---|---|---|
| allcell | ZMYM2 | 4.44E-29 | 0.756544084 | 0.663 | 0.487 | 8.74E-25 | Tb |
| allcell | FOXP1 | 3.48E-47 | 0.756052324 | 0.832 | 0.673 | 6.84E-43 | Tb |
| allcell | JPX | 4.07E-38 | 0.755535225 | 0.5 | 0.29 | 8.00E-34 | Tb |
| allcell | STXBP5 | 1.22E-19 | 0.75536554 | 0.255 | 0.144 | 2.40E-15 | Tb |
| allcell | SKIL | 4.83E-53 | 0.754852417 | 0.789 | 0.569 | 9.51E-49 | Tb |
| allcell | PLEKHM3 | 3.37E-18 | 0.754504251 | 0.359 | 0.236 | 6.63E-14 | Tb |
| allcell | MLLT10 | 6.46E-34 | 0.754123507 | 0.437 | 0.253 | 1.27E-29 | Tb |
| allcell | FRS2 | 1.28E-30 | 0.753734174 | 0.335 | 0.177 | 2.51E-26 | Tb |
| allcell | VPS8 | 4.67E-20 | 0.753712606 | 0.339 | 0.21 | 9.19E-16 | Tb |
| allcell | LHFPL2 | 6.96E-16 | 0.753162567 | 0.262 | 0.16 | 1.37E-11 | Tb |
| allcell | ITPR2 | 3.43E-47 | 0.752392665 | 0.657 | 0.418 | 6.76E-43 | Tb |
| allcell | ARSB | 4.27E-27 | 0.751803415 | 0.417 | 0.251 | 8.41E-23 | Tb |
| allcell | CLASP1 | 2.55E-35 | 0.751787627 | 0.417 | 0.23 | 5.02E-31 | Tb |
| allcell | ZMYND8 | 9.93E-34 | 0.751250801 | 0.382 | 0.206 | 1.95E-29 | Tb |
| allcell | TOPORS | 5.06E-32 | 0.750919394 | 0.397 | 0.222 | 9.96E-28 | Tb |
| allcell | TCOF1 | 8.40E-21 | 0.750511655 | 0.318 | 0.187 | 1.65E-16 | Tb |
| allcell | ARMH3 | 8.93E-43 | 0.749805647 | 0.467 | 0.252 | 1.76E-38 | Tb |
| allcell | TRPM2 | 6.40E-22 | 0.749529934 | 0.259 | 0.143 | 1.26E-17 | Tb |
| allcell | PSEN1 | 1.10E-37 | 0.748758087 | 0.541 | 0.338 | 2.16E-33 | Tb |
| allcell | MOSMO | 1.29E-26 | 0.748524776 | 0.29 | 0.151 | 2.53E-22 | Tb |
| allcell | NR2C2 | 7.69E-25 | 0.747180374 | 0.286 | 0.153 | 1.51E-20 | Tb |
| allcell | THOC1 | 3.55E-20 | 0.746599136 | 0.287 | 0.167 | 6.99E-16 | Tb |
| allcell | TWISTNB | 1.45E-18 | 0.742903305 | 0.616 | 0.455 | 2.86E-14 | Tb |
| allcell | CSF2RB | 4.26E-42 | 0.742759437 | 0.518 | 0.296 | 8.38E-38 | Tb |
| allcell | TYW1 | 2.61E-30 | 0.742609838 | 0.351 | 0.189 | 5.14E-26 | Tb |
| allcell | BACH2 | 1.84E-19 | 0.741801181 | 0.297 | 0.179 | 3.62E-15 | Tb |
| allcell | ADD3 | 3.82E-19 | 0.741377955 | 0.346 | 0.221 | 7.52E-15 | Tb |
| allcell | COP1 | 1.79E-50 | 0.740724194 | 0.68 | 0.441 | 3.53E-46 | Tb |
| allcell | CBFA2T2 | 3.23E-30 | 0.740718144 | 0.32 | 0.168 | 6.35E-26 | Tb |
| allcell | MAN1A1 | 1.33E-50 | 0.740555302 | 0.79 | 0.569 | 2.61E-46 | Tb |
| allcell | MTMR9 | 7.90E-20 | 0.740531268 | 0.291 | 0.168 | 1.55E-15 | Tb |
| allcell | SRGAP2B | 6.31E-33 | 0.74051055 | 0.411 | 0.231 | 1.24E-28 | Tb |
| allcell | MBD5 | 1.87E-31 | 0.739906029 | 0.436 | 0.261 | 3.68E-27 | Tb |
| allcell | RIPK1 | 5.72E-37 | 0.739804193 | 0.538 | 0.332 | 1.13E-32 | Tb |
| allcell | TUT4 | 5.02E-19 | 0.739649551 | 0.278 | 0.163 | 9.89E-15 | Tb |
| allcell | CPED1 | 1.51E-14 | 0.738968712 | 0.275 | 0.175 | 2.97E-10 | Tb |
| allcell | NRP1 | 4.97E-19 | 0.738510349 | 0.302 | 0.185 | 9.77E-15 | Tb |
| allcell | PMM2 | 1.91E-25 | 0.738203739 | 0.342 | 0.193 | 3.76E-21 | Tb |
| allcell | TRIM38 | 2.93E-42 | 0.737568352 | 0.622 | 0.408 | 5.76E-38 | Tb |
| allcell | R3HDM2 | 6.71E-41 | 0.737518797 | 0.601 | 0.397 | 1.32E-36 | Tb |
| allcell | RBMS1 | 7.98E-30 | 0.737250111 | 0.499 | 0.316 | 1.57E-25 | Tb |
| allcell | NCOA1 | 3.61E-40 | 0.737131018 | 0.632 | 0.411 | 7.10E-36 | Tb |
| allcell | COMMD10 | 4.39E-29 | 0.7367169 | 0.607 | 0.414 | 8.64E-25 | Tb |
| allcell | ATP2C1 | 2.67E-31 | 0.735969048 | 0.675 | 0.484 | 5.26E-27 | Tb |
| allcell | TRAPPC9 | 3.54E-46 | 0.734037487 | 0.539 | 0.311 | 6.98E-42 | Tb |
| allcell | YAF2 | 3.27E-27 | 0.733641596 | 0.445 | 0.272 | 6.44E-23 | Tb |
| allcell | PPP1R14A | 2.32E-19 | 0.733098199 | 0.275 | 0.161 | 4.56E-15 | Tb |
| allcell | ANKRD44 | 1.26E-50 | 0.731994985 | 0.752 | 0.547 | 2.48E-46 | Tb |
| allcell | PDS5B | 3.47E-39 | 0.731646295 | 0.474 | 0.268 | 6.83E-35 | Tb |
| allcell | TENT2 | 3.39E-50 | 0.731552486 | 0.631 | 0.381 | 6.67E-46 | Tb |
| allcell | PRKCB | 7.61E-33 | 0.731028878 | 0.701 | 0.517 | 1.50E-28 | Tb |
| allcell | KDM7A-DT | 6.57E-22 | 0.729886834 | 0.25 | 0.13 | 1.29E-17 | Tb |
| allcell | ADAMTSL4-AS | 2.64E-36 | 0.729862764 | 0.49 | 0.29 | 5.21E-32 | Tb |
| allcell | CDC42SE2 | 2.89E-58 | 0.729850815 | 0.871 | 0.68 | 5.68E-54 | Tb |
| allcell | SP140 | 2.26E-17 | 0.729666002 | 0.353 | 0.232 | 4.45E-13 | Tb |
| allcell | SLC45A4 | 2.86E-24 | 0.729363837 | 0.318 | 0.18 | 5.63E-20 | Tb |

| | | | | | | | |
|---|---|---|---|---|---|---|---|
| allcell | CDR2 | 1.25E-30 | 0.728634252 | 0.502 | 0.32 | 2.46E-26 | Tb |
| allcell | SHLD2 | 5.49E-20 | 0.72835001 | 0.353 | 0.217 | 1.08E-15 | Tb |
| allcell | DOCK8 | 1.60E-39 | 0.726353722 | 0.751 | 0.58 | 3.15E-35 | Tb |
| allcell | PRORP | 1.78E-25 | 0.726304573 | 0.314 | 0.171 | 3.50E-21 | Tb |
| allcell | PTPN9 | 1.19E-19 | 0.726128223 | 0.269 | 0.152 | 2.34E-15 | Tb |
| allcell | ZZEF1 | 1.07E-30 | 0.726117683 | 0.403 | 0.227 | 2.10E-26 | Tb |
| allcell | RANBP2 | 2.09E-28 | 0.726116352 | 0.782 | 0.611 | 4.12E-24 | Tb |
| allcell | SH3KBP1 | 1.14E-26 | 0.724944628 | 0.66 | 0.502 | 2.24E-22 | Tb |
| allcell | CAMKMT | 4.98E-15 | 0.724890169 | 0.264 | 0.164 | 9.80E-11 | Tb |
| allcell | IBTK | 2.14E-29 | 0.723597223 | 0.393 | 0.223 | 4.22E-25 | Tb |
| allcell | PDSS2 | 4.66E-26 | 0.723214862 | 0.284 | 0.148 | 9.18E-22 | Tb |
| allcell | PICALM | 2.99E-52 | 0.721966071 | 0.852 | 0.719 | 5.88E-48 | Tb |
| allcell | DOCK10 | 1.12E-36 | 0.721637679 | 0.791 | 0.648 | 2.20E-32 | Tb |
| allcell | ADIPOR2 | 2.07E-32 | 0.72127276 | 0.379 | 0.207 | 4.07E-28 | Tb |
| allcell | DOP1A | 3.18E-19 | 0.721255365 | 0.316 | 0.195 | 6.26E-15 | Tb |
| allcell | RUBCN | 1.31E-24 | 0.720710484 | 0.484 | 0.323 | 2.58E-20 | Tb |
| allcell | SNX14 | 5.80E-27 | 0.720679383 | 0.378 | 0.219 | 1.14E-22 | Tb |
| allcell | ATXN7 | 3.51E-29 | 0.71977714 | 0.441 | 0.269 | 6.91E-25 | Tb |
| allcell | SLC44A1 | 5.19E-17 | 0.719684445 | 0.346 | 0.222 | 1.02E-12 | Tb |
| allcell | ULK4 | 1.33E-26 | 0.718211948 | 0.313 | 0.17 | 2.62E-22 | Tb |
| allcell | KIAA2026 | 2.83E-33 | 0.718099244 | 0.418 | 0.238 | 5.57E-29 | Tb |
| allcell | CLASP2 | 2.70E-42 | 0.717555732 | 0.644 | 0.427 | 5.32E-38 | Tb |
| allcell | SUPT3H | 1.55E-19 | 0.717494714 | 0.279 | 0.161 | 3.05E-15 | Tb |
| allcell | MICAL2 | 9.65E-21 | 0.717361969 | 0.323 | 0.195 | 1.90E-16 | Tb |
| allcell | ALOX5 | 6.66E-22 | 0.71689432 | 0.439 | 0.296 | 1.31E-17 | Tb |
| allcell | FAM172A | 1.21E-34 | 0.716152062 | 0.508 | 0.315 | 2.39E-30 | Tb |
| allcell | SMURF1 | 4.21E-35 | 0.715808344 | 0.421 | 0.238 | 8.28E-31 | Tb |
| allcell | SLC38A1 | 1.52E-33 | 0.714129648 | 0.626 | 0.44 | 2.99E-29 | Tb |
| allcell | ATP1B3 | 8.56E-29 | 0.713912273 | 0.9 | 0.837 | 1.68E-24 | Tb |
| allcell | MCCC1 | 2.77E-25 | 0.713628881 | 0.276 | 0.143 | 5.45E-21 | Tb |
| allcell | INTS6L | 5.02E-28 | 0.713183096 | 0.302 | 0.158 | 9.88E-24 | Tb |
| allcell | ENTPD1 | 9.81E-22 | 0.712945827 | 0.522 | 0.375 | 1.93E-17 | Tb |
| allcell | SNX8 | 5.61E-35 | 0.711913891 | 0.679 | 0.528 | 1.11E-30 | Tb |
| allcell | MT-CO1 | 2.46E-80 | 0.711510088 | 0.997 | 0.985 | 4.84E-76 | Tb |
| allcell | PHF21A | 1.41E-23 | 0.711248974 | 0.399 | 0.247 | 2.78E-19 | Tb |
| allcell | MICU1 | 2.91E-25 | 0.711245956 | 0.368 | 0.219 | 5.73E-21 | Tb |
| allcell | DIP2B | 6.52E-34 | 0.711008895 | 0.439 | 0.251 | 1.28E-29 | Tb |
| allcell | GLG1 | 4.52E-29 | 0.710305669 | 0.522 | 0.338 | 8.89E-25 | Tb |
| allcell | MITD1 | 5.09E-22 | 0.709076769 | 0.331 | 0.192 | 1.00E-17 | Tb |
| allcell | AKT3 | 2.45E-21 | 0.708932791 | 0.356 | 0.218 | 4.83E-17 | Tb |
| allcell | ZXDC | 7.43E-23 | 0.708215667 | 0.293 | 0.162 | 1.46E-18 | Tb |
| allcell | ANKIB1 | 1.54E-29 | 0.707634515 | 0.411 | 0.243 | 3.03E-25 | Tb |
| allcell | HMG20A | 2.80E-19 | 0.707621876 | 0.255 | 0.141 | 5.51E-15 | Tb |
| allcell | AP3S2 | 2.91E-23 | 0.706973053 | 0.316 | 0.177 | 5.72E-19 | Tb |
| allcell | ITPRIPL2 | 9.22E-23 | 0.706517979 | 0.281 | 0.152 | 1.82E-18 | Tb |
| allcell | DLEU1 | 1.26E-42 | 0.705327662 | 0.495 | 0.283 | 2.49E-38 | Tb |
| allcell | SEMA3C | 8.86E-17 | 0.705203864 | 0.259 | 0.155 | 1.74E-12 | Tb |
| allcell | RHBDD1 | 3.41E-20 | 0.705017954 | 0.298 | 0.176 | 6.71E-16 | Tb |
| allcell | SIPA1L1 | 4.29E-42 | 0.70492995 | 0.884 | 0.758 | 8.44E-38 | Tb |
| allcell | DIS3L2 | 4.39E-21 | 0.703970752 | 0.279 | 0.157 | 8.65E-17 | Tb |
| allcell | TLK1 | 1.61E-29 | 0.703618577 | 0.481 | 0.294 | 3.16E-25 | Tb |
| allcell | RLF | 1.51E-27 | 0.703581398 | 0.652 | 0.48 | 2.98E-23 | Tb |
| allcell | CCDC18 | 4.68E-29 | 0.703398136 | 0.353 | 0.195 | 9.21E-25 | Tb |
| allcell | ST3GAL1 | 2.17E-29 | 0.703045307 | 0.458 | 0.283 | 4.27E-25 | Tb |
| allcell | LARP1B | 1.87E-20 | 0.702857235 | 0.312 | 0.183 | 3.67E-16 | Tb |
| allcell | TMEM131 | 1.72E-37 | 0.702708446 | 0.717 | 0.528 | 3.39E-33 | Tb |

| | | | | | | | |
|---|---|---|---|---|---|---|---|
| allcell | UBP1 | 3.52E-25 | 0.702226878 | 0.317 | 0.175 | 6.93E-21 | Tb |
| allcell | GAB2 | 3.06E-33 | 0.702145534 | 0.658 | 0.47 | 6.03E-29 | Tb |
| allcell | PIP5K1A | 1.47E-32 | 0.701695351 | 0.427 | 0.247 | 2.90E-28 | Tb |
| allcell | EXT1 | 3.01E-30 | 0.701215589 | 0.479 | 0.303 | 5.93E-26 | Tb |
| allcell | SSH1 | 6.90E-20 | 0.70106283 | 0.3 | 0.175 | 1.36E-15 | Tb |
| allcell | MT-ND3 | 6.65E-97 | 0.700564326 | 0.983 | 0.899 | 1.31E-92 | Tb |
| allcell | USP32 | 1.77E-42 | 0.699917194 | 0.638 | 0.421 | 3.49E-38 | Tb |
| allcell | ZNF410 | 4.23E-26 | 0.699909659 | 0.381 | 0.223 | 8.33E-22 | Tb |
| allcell | RICTOR | 1.23E-27 | 0.699882077 | 0.432 | 0.262 | 2.42E-23 | Tb |
| allcell | SUSD6 | 7.14E-36 | 0.69855056 | 0.586 | 0.39 | 1.41E-31 | Tb |
| allcell | MED13L | 8.96E-60 | 0.698436392 | 0.88 | 0.7 | 1.76E-55 | Tb |
| allcell | MYO1C | 5.32E-28 | 0.697586422 | 0.302 | 0.158 | 1.05E-23 | Tb |
| allcell | INTS6 | 2.13E-39 | 0.697555882 | 0.731 | 0.545 | 4.20E-35 | Tb |
| allcell | FOXO1 | 4.52E-17 | 0.697337039 | 0.4 | 0.277 | 8.90E-13 | Tb |
| allcell | SLC16A3 | 2.83E-27 | 0.696944026 | 0.425 | 0.259 | 5.57E-23 | Tb |
| allcell | SEMA4D | 1.96E-19 | 0.696348657 | 0.266 | 0.151 | 3.85E-15 | Tb |
| allcell | SYNJ1 | 1.15E-24 | 0.694981799 | 0.307 | 0.17 | 2.27E-20 | Tb |
| allcell | NUBPL | 1.80E-22 | 0.69479366 | 0.428 | 0.282 | 3.55E-18 | Tb |
| allcell | IKZF1 | 2.11E-39 | 0.694735683 | 0.728 | 0.556 | 4.16E-35 | Tb |
| allcell | PRKCE | 2.26E-20 | 0.693536807 | 0.43 | 0.295 | 4.46E-16 | Tb |
| allcell | TBL1XR1 | 1.99E-36 | 0.693478969 | 0.667 | 0.457 | 3.92E-32 | Tb |
| allcell | RBBP8 | 5.46E-09 | 0.6924252 | 0.342 | 0.248 | 0.000107425 | Tb |
| allcell | FKBP5 | 1.04E-48 | 0.692082444 | 0.707 | 0.487 | 2.04E-44 | Tb |
| allcell | ATP6V0A1 | 8.57E-27 | 0.691812593 | 0.358 | 0.203 | 1.69E-22 | Tb |
| allcell | FNDC3B | 4.28E-36 | 0.691656543 | 0.669 | 0.479 | 8.42E-32 | Tb |
| allcell | RIPOR2 | 2.45E-22 | 0.691414635 | 0.532 | 0.367 | 4.82E-18 | Tb |
| allcell | STAT6 | 6.16E-41 | 0.691167909 | 0.527 | 0.314 | 1.21E-36 | Tb |
| allcell | MDN1 | 4.90E-21 | 0.690580283 | 0.282 | 0.158 | 9.64E-17 | Tb |
| allcell | CHST11 | 1.59E-40 | 0.690514941 | 0.741 | 0.557 | 3.13E-36 | Tb |
| allcell | DENND4B | 6.14E-23 | 0.690411696 | 0.284 | 0.154 | 1.21E-18 | Tb |
| allcell | STRADA | 2.40E-25 | 0.690086531 | 0.27 | 0.139 | 4.73E-21 | Tb |
| allcell | MCM9 | 2.82E-19 | 0.68947687 | 0.255 | 0.144 | 5.55E-15 | Tb |
| allcell | ARHGAP18 | 2.68E-26 | 0.688798243 | 0.574 | 0.412 | 5.28E-22 | Tb |
| allcell | ANKRD13C | 1.66E-19 | 0.688786406 | 0.252 | 0.14 | 3.28E-15 | Tb |
| allcell | ANKRD17 | 1.14E-38 | 0.688009315 | 0.637 | 0.425 | 2.24E-34 | Tb |
| allcell | CREB5 | 2.00E-12 | 0.686402193 | 0.447 | 0.342 | 3.93E-08 | Tb |
| allcell | ELMSAN1 | 3.27E-38 | 0.685953821 | 0.515 | 0.306 | 6.43E-34 | Tb |
| allcell | NUCB2 | 3.16E-15 | 0.685693913 | 0.318 | 0.212 | 6.21E-11 | Tb |
| allcell | NMRK1 | 9.73E-16 | 0.684601228 | 0.287 | 0.179 | 1.92E-11 | Tb |
| allcell | PPA2 | 8.77E-32 | 0.68448263 | 0.516 | 0.325 | 1.73E-27 | Tb |
| allcell | TGIF2 | 4.20E-25 | 0.684341498 | 0.419 | 0.259 | 8.27E-21 | Tb |
| allcell | B3GALNT2 | 1.23E-20 | 0.684321345 | 0.263 | 0.146 | 2.42E-16 | Tb |
| allcell | LRMP | 4.24E-15 | 0.683266667 | 0.374 | 0.257 | 8.34E-11 | Tb |
| allcell | PPP2R5A | 2.14E-30 | 0.682382702 | 0.329 | 0.17 | 4.21E-26 | Tb |
| allcell | RAB5B | 8.59E-20 | 0.682013399 | 0.329 | 0.198 | 1.69E-15 | Tb |
| allcell | KDM4C | 8.83E-47 | 0.681958588 | 0.67 | 0.44 | 1.74E-42 | Tb |
| allcell | CDK13 | 5.19E-30 | 0.68181147 | 0.551 | 0.366 | 1.02E-25 | Tb |
| allcell | DCLRE1C | 2.85E-20 | 0.68164573 | 0.251 | 0.135 | 5.61E-16 | Tb |
| allcell | CYP20A1 | 1.25E-17 | 0.681435573 | 0.394 | 0.261 | 2.47E-13 | Tb |
| allcell | FGD2 | 2.03E-34 | 0.681279488 | 0.512 | 0.313 | 3.99E-30 | Tb |
| allcell | SOS2 | 8.13E-24 | 0.681092422 | 0.379 | 0.23 | 1.60E-19 | Tb |
| allcell | SH2B3 | 3.53E-42 | 0.680724546 | 0.712 | 0.53 | 6.95E-38 | Tb |
| allcell | ACOX1 | 1.70E-16 | 0.68061862 | 0.251 | 0.146 | 3.36E-12 | Tb |
| allcell | SCFD2 | 6.11E-21 | 0.680580816 | 0.33 | 0.199 | 1.20E-16 | Tb |
| allcell | CDYL | 1.90E-35 | 0.680444612 | 0.714 | 0.521 | 3.75E-31 | Tb |
| allcell | NLRC5 | 5.93E-34 | 0.680307547 | 0.47 | 0.28 | 1.17E-29 | Tb |

| | | | | | | | |
|---|---|---|---|---|---|---|---|
| allcell | VPS53 | 8.84E-35 | 0.680225963 | 0.436 | 0.246 | 1.74E-30 | Tb |
| allcell | NFAT5 | 9.59E-33 | 0.679901958 | 0.825 | 0.712 | 1.89E-28 | Tb |
| allcell | SPATS2 | 3.83E-33 | 0.679761079 | 0.542 | 0.342 | 7.53E-29 | Tb |
| allcell | TBL1X | 1.40E-17 | 0.679359825 | 0.382 | 0.255 | 2.76E-13 | Tb |
| allcell | CRYBG1 | 8.20E-19 | 0.67894459 | 0.614 | 0.477 | 1.61E-14 | Tb |
| allcell | ELF2 | 3.67E-33 | 0.678649613 | 0.677 | 0.489 | 7.22E-29 | Tb |
| allcell | GABPB1-IT1 | 2.78E-21 | 0.678114211 | 0.284 | 0.157 | 5.47E-17 | Tb |
| allcell | 8-Mar | 9.10E-27 | 0.678036652 | 0.323 | 0.175 | 1.79E-22 | Tb |
| allcell | ENTPD1-AS1 | 1.04E-36 | 0.677539024 | 0.459 | 0.262 | 2.04E-32 | Tb |
| allcell | SELENOI | 2.53E-20 | 0.677380015 | 0.293 | 0.168 | 4.99E-16 | Tb |
| allcell | MT-ND1 | 1.07E-87 | 0.677273708 | 0.97 | 0.898 | 2.10E-83 | Tb |
| allcell | AHI1 | 4.28E-22 | 0.676277751 | 0.5 | 0.339 | 8.42E-18 | Tb |
| allcell | ZBTB44 | 5.33E-21 | 0.675323866 | 0.269 | 0.148 | 1.05E-16 | Tb |
| allcell | NUP107 | 3.38E-21 | 0.674891159 | 0.282 | 0.158 | 6.65E-17 | Tb |
| allcell | PHC3 | 1.62E-21 | 0.674741264 | 0.376 | 0.234 | 3.18E-17 | Tb |
| allcell | YARS | 3.78E-24 | 0.67461575 | 0.35 | 0.201 | 7.44E-20 | Tb |
| allcell | MED27 | 6.30E-23 | 0.673916665 | 0.3 | 0.167 | 1.24E-18 | Tb |
| allcell | ZFX | 4.34E-24 | 0.673796313 | 0.425 | 0.269 | 8.54E-20 | Tb |
| allcell | GPBP1L1 | 3.40E-37 | 0.673479585 | 0.533 | 0.316 | 6.69E-33 | Tb |
| allcell | KCNK6 | 8.14E-24 | 0.673376467 | 0.367 | 0.224 | 1.60E-19 | Tb |
| allcell | OSBPL3 | 1.17E-18 | 0.673238644 | 0.473 | 0.336 | 2.31E-14 | Tb |
| allcell | IP6K1 | 1.95E-18 | 0.673160999 | 0.27 | 0.155 | 3.83E-14 | Tb |
| allcell | FBXO34 | 2.54E-48 | 0.672551256 | 0.683 | 0.458 | 5.00E-44 | Tb |
| allcell | RCOR1 | 3.13E-28 | 0.671984133 | 0.731 | 0.582 | 6.17E-24 | Tb |
| allcell | EXOC6 | 9.71E-28 | 0.671557368 | 0.382 | 0.223 | 1.91E-23 | Tb |
| allcell | NLRP1 | 1.30E-14 | 0.671209881 | 0.316 | 0.205 | 2.55E-10 | Tb |
| allcell | EML4 | 2.33E-40 | 0.671132299 | 0.717 | 0.523 | 4.59E-36 | Tb |
| allcell | AGO3 | 1.14E-32 | 0.671120234 | 0.53 | 0.334 | 2.25E-28 | Tb |
| allcell | CEP128 | 2.49E-23 | 0.670803254 | 0.363 | 0.22 | 4.91E-19 | Tb |
| allcell | CCDC18-AS1 | 7.00E-24 | 0.67069831 | 0.328 | 0.186 | 1.38E-19 | Tb |
| allcell | MT-CO2 | 7.65E-96 | 0.670482558 | 0.989 | 0.982 | 1.51E-91 | Tb |
| allcell | STK3 | 1.98E-28 | 0.670262964 | 0.528 | 0.353 | 3.89E-24 | Tb |
| allcell | 3-Mar | 7.95E-12 | 0.670064766 | 0.313 | 0.215 | 1.57E-07 | Tb |
| allcell | MR1 | 6.18E-20 | 0.670062062 | 0.263 | 0.146 | 1.22E-15 | Tb |
| allcell | RAB3GAP1 | 8.40E-33 | 0.669569416 | 0.471 | 0.283 | 1.65E-28 | Tb |
| allcell | HAVCR2 | 5.49E-13 | 0.66928621 | 0.528 | 0.405 | 1.08E-08 | Tb |
| allcell | SEL1L3 | 1.35E-23 | 0.668076089 | 0.444 | 0.288 | 2.66E-19 | Tb |
| allcell | HOMER2 | 3.48E-28 | 0.668070909 | 0.286 | 0.143 | 6.86E-24 | Tb |
| allcell | EXOC2 | 3.00E-22 | 0.667938797 | 0.281 | 0.154 | 5.90E-18 | Tb |
| allcell | ZBTB46 | 8.00E-31 | 0.667374029 | 0.488 | 0.308 | 1.58E-26 | Tb |
| allcell | ZNF124 | 4.86E-34 | 0.666903107 | 0.395 | 0.215 | 9.56E-30 | Tb |
| allcell | PHACTR2 | 1.97E-21 | 0.666710663 | 0.44 | 0.292 | 3.88E-17 | Tb |
| allcell | FLT1 | 2.39E-10 | 0.666270552 | 0.554 | 0.471 | 4.70E-06 | Tb |
| allcell | TRIM13 | 2.81E-29 | 0.666167982 | 0.393 | 0.222 | 5.53E-25 | Tb |
| allcell | ACSL1 | 9.73E-12 | 0.66597617 | 0.586 | 0.461 | 1.92E-07 | Tb |
| allcell | PPP1R21 | 9.48E-20 | 0.665932594 | 0.276 | 0.159 | 1.87E-15 | Tb |
| allcell | YME1L1 | 5.82E-34 | 0.665802343 | 0.801 | 0.627 | 1.15E-29 | Tb |
| allcell | TEX14 | 2.06E-20 | 0.665539499 | 0.562 | 0.415 | 4.05E-16 | Tb |
| allcell | MRTFA | 2.12E-30 | 0.665371127 | 0.674 | 0.491 | 4.17E-26 | Tb |
| allcell | FARP2 | 4.58E-26 | 0.664867342 | 0.552 | 0.385 | 9.02E-22 | Tb |
| allcell | GFPT1 | 4.57E-29 | 0.663961183 | 0.416 | 0.244 | 9.00E-25 | Tb |
| allcell | FPR3 | 1.14E-10 | 0.663960603 | 0.416 | 0.323 | 2.25E-06 | Tb |
| allcell | SC5D | 3.04E-16 | 0.663140794 | 0.251 | 0.148 | 5.99E-12 | Tb |
| allcell | PDE4A | 1.84E-18 | 0.663101733 | 0.663 | 0.539 | 3.62E-14 | Tb |
| allcell | TRIM24 | 9.80E-18 | 0.662872887 | 0.321 | 0.2 | 1.93E-13 | Tb |
| allcell | ZC3H4 | 5.25E-28 | 0.662550774 | 0.303 | 0.155 | 1.03E-23 | Tb |

| | | | | | | | |
|---|---|---|---|---|---|---|---|
| allcell | MLLT6 | 1.46E-18 | 0.6624337 | 0.297 | 0.174 | 2.88E-14 | Tb |
| allcell | PCBD2 | 2.95E-21 | 0.661381091 | 0.257 | 0.138 | 5.81E-17 | Tb |
| allcell | BCL2L14 | 2.06E-12 | 0.660786162 | 0.279 | 0.188 | 4.06E-08 | Tb |
| allcell | APOL6 | 1.35E-16 | 0.660559531 | 0.281 | 0.17 | 2.66E-12 | Tb |
| allcell | TRAFD1 | 5.29E-24 | 0.660186388 | 0.334 | 0.193 | 1.04E-19 | Tb |
| allcell | KLF12 | 1.71E-13 | 0.660096779 | 0.25 | 0.158 | 3.37E-09 | Tb |
| allcell | ZNF407 | 6.93E-32 | 0.660095274 | 0.458 | 0.271 | 1.36E-27 | Tb |
| allcell | ST3GAL5 | 4.84E-40 | 0.659991348 | 0.544 | 0.321 | 9.52E-36 | Tb |
| allcell | FOXN3 | 3.87E-31 | 0.658933434 | 0.777 | 0.644 | 7.63E-27 | Tb |
| allcell | CNOT10 | 3.02E-25 | 0.658601862 | 0.418 | 0.261 | 5.95E-21 | Tb |
| allcell | GTF2H1 | 7.92E-23 | 0.657995315 | 0.455 | 0.298 | 1.56E-18 | Tb |
| allcell | ZEB1 | 3.22E-27 | 0.657985247 | 0.585 | 0.429 | 6.34E-23 | Tb |
| allcell | IDO2 | 7.45E-24 | 0.657532318 | 0.492 | 0.33 | 1.47E-19 | Tb |
| allcell | MTIF2 | 4.41E-26 | 0.657348091 | 0.286 | 0.148 | 8.69E-22 | Tb |
| allcell | MAP3K5 | 8.10E-26 | 0.657316495 | 0.468 | 0.301 | 1.59E-21 | Tb |
| allcell | SLC7A1 | 4.99E-29 | 0.656264814 | 0.468 | 0.287 | 9.83E-25 | Tb |
| allcell | STK38L | 1.51E-28 | 0.65593523 | 0.596 | 0.42 | 2.98E-24 | Tb |
| allcell | LTB | 6.49E-28 | 0.655669065 | 0.572 | 0.397 | 1.28E-23 | Tb |
| allcell | FAM133B | 4.11E-41 | 0.65503473 | 0.582 | 0.356 | 8.08E-37 | Tb |
| allcell | YEATS2 | 6.51E-24 | 0.654972831 | 0.316 | 0.179 | 1.28E-19 | Tb |
| allcell | SOCS1 | 2.95E-30 | 0.654863477 | 0.645 | 0.453 | 5.80E-26 | Tb |
| allcell | RAPH1 | 2.15E-11 | 0.654629107 | 0.297 | 0.209 | 4.23E-07 | Tb |
| allcell | OGA | 1.02E-31 | 0.654543383 | 0.486 | 0.298 | 2.01E-27 | Tb |
| allcell | ABI2 | 3.50E-21 | 0.654534851 | 0.314 | 0.183 | 6.88E-17 | Tb |
| allcell | ZNF791 | 8.07E-23 | 0.654343429 | 0.322 | 0.185 | 1.59E-18 | Tb |
| allcell | CNOT6L | 2.82E-28 | 0.653967127 | 0.619 | 0.43 | 5.54E-24 | Tb |
| allcell | HERC4 | 7.72E-26 | 0.653853432 | 0.407 | 0.247 | 1.52E-21 | Tb |
| allcell | STAT3 | 1.41E-46 | 0.653582294 | 0.798 | 0.607 | 2.77E-42 | Tb |
| allcell | 1-Mar | 1.34E-23 | 0.653548509 | 0.566 | 0.408 | 2.64E-19 | Tb |
| allcell | POU2F1 | 3.02E-24 | 0.653543787 | 0.346 | 0.201 | 5.95E-20 | Tb |
| allcell | MBNL1 | 1.40E-41 | 0.653495851 | 0.896 | 0.79 | 2.76E-37 | Tb |
| allcell | AP5Z1 | 5.24E-33 | 0.653494878 | 0.355 | 0.183 | 1.03E-28 | Tb |
| allcell | SETDB2 | 2.41E-22 | 0.653200354 | 0.442 | 0.285 | 4.74E-18 | Tb |
| allcell | TCF20 | 4.34E-22 | 0.652660152 | 0.283 | 0.156 | 8.54E-18 | Tb |
| allcell | GNPTAB | 1.65E-28 | 0.652605275 | 0.453 | 0.275 | 3.24E-24 | Tb |
| allcell | GABPB1 | 1.87E-21 | 0.652160282 | 0.322 | 0.19 | 3.68E-17 | Tb |
| allcell | PSD3 | 4.85E-09 | 0.651450288 | 0.257 | 0.178 | 9.55E-05 | Tb |
| allcell | IQCB1 | 3.07E-25 | 0.650536575 | 0.384 | 0.229 | 6.04E-21 | Tb |
| allcell | UBE2E1 | 8.31E-28 | 0.649676614 | 0.656 | 0.5 | 1.63E-23 | Tb |
| allcell | ADPGK | 9.13E-36 | 0.649311036 | 0.505 | 0.295 | 1.80E-31 | Tb |
| allcell | ZKSCAN1 | 9.10E-25 | 0.649015123 | 0.34 | 0.191 | 1.79E-20 | Tb |
| allcell | RABGAP1 | 3.56E-28 | 0.64847823 | 0.433 | 0.26 | 7.01E-24 | Tb |
| allcell | NIN | 3.02E-22 | 0.648302142 | 0.546 | 0.388 | 5.94E-18 | Tb |
| allcell | SLA | 4.56E-19 | 0.647976335 | 0.276 | 0.16 | 8.98E-15 | Tb |
| allcell | ZNF490 | 1.65E-29 | 0.647443343 | 0.273 | 0.133 | 3.25E-25 | Tb |
| allcell | KPNA6 | 3.51E-20 | 0.647426964 | 0.294 | 0.168 | 6.91E-16 | Tb |
| allcell | PCSK7 | 5.47E-31 | 0.64740579 | 0.311 | 0.157 | 1.08E-26 | Tb |
| allcell | KLHL15 | 5.71E-16 | 0.64658069 | 0.287 | 0.174 | 1.12E-11 | Tb |
| allcell | EIF4G3 | 1.35E-22 | 0.646541091 | 0.517 | 0.367 | 2.67E-18 | Tb |
| allcell | TFEC | 7.36E-08 | 0.646372802 | 0.476 | 0.382 | 0.001449657 | Tb |
| allcell | EPC2 | 1.86E-19 | 0.646299743 | 0.334 | 0.203 | 3.67E-15 | Tb |
| allcell | SNX29 | 1.21E-34 | 0.646004887 | 0.619 | 0.435 | 2.38E-30 | Tb |
| allcell | CGAS | 1.35E-28 | 0.645662367 | 0.469 | 0.29 | 2.66E-24 | Tb |
| allcell | PCNX4 | 1.47E-19 | 0.645449617 | 0.346 | 0.217 | 2.90E-15 | Tb |
| allcell | STRN3 | 3.45E-30 | 0.645424811 | 0.404 | 0.23 | 6.79E-26 | Tb |
| allcell | PRIM2 | 2.59E-16 | 0.645141392 | 0.259 | 0.154 | 5.10E-12 | Tb |

| | | | | | | | |
|---|---|---|---|---|---|---|---|
| allcell | PARP9 | 1.67E-24 | 0.645062448 | 0.383 | 0.23 | 3.28E-20 | Tb |
| allcell | MICU2 | 1.10E-21 | 0.644953666 | 0.403 | 0.251 | 2.16E-17 | Tb |
| allcell | SINHCAF | 6.94E-22 | 0.644710198 | 0.607 | 0.449 | 1.37E-17 | Tb |
| allcell | RBM23 | 8.50E-29 | 0.644580278 | 0.425 | 0.249 | 1.67E-24 | Tb |
| allcell | RFTN1 | 8.68E-31 | 0.644016549 | 0.777 | 0.658 | 1.71E-26 | Tb |
| allcell | MGAT5 | 4.66E-34 | 0.643880032 | 0.506 | 0.307 | 9.16E-30 | Tb |
| allcell | NEMP1 | 3.05E-20 | 0.642191683 | 0.25 | 0.134 | 6.00E-16 | Tb |
| allcell | BAZ2B | 2.21E-25 | 0.642076746 | 0.549 | 0.382 | 4.36E-21 | Tb |
| allcell | KRT10 | 2.92E-14 | 0.641434195 | 0.686 | 0.543 | 5.75E-10 | Tb |
| allcell | RAPGEF1 | 1.74E-48 | 0.641334699 | 0.89 | 0.768 | 3.43E-44 | Tb |
| allcell | USP36 | 5.16E-35 | 0.640054671 | 0.626 | 0.407 | 1.02E-30 | Tb |
| allcell | TCF12 | 9.03E-43 | 0.639900792 | 0.66 | 0.44 | 1.78E-38 | Tb |
| allcell | TPD52 | 8.41E-16 | 0.639370464 | 0.313 | 0.198 | 1.66E-11 | Tb |
| allcell | CMTM7 | 3.94E-24 | 0.638962904 | 0.575 | 0.397 | 7.76E-20 | Tb |
| allcell | WDR70 | 7.23E-26 | 0.638687577 | 0.459 | 0.289 | 1.42E-21 | Tb |
| allcell | DDX10 | 7.81E-25 | 0.638146474 | 0.346 | 0.197 | 1.54E-20 | Tb |
| allcell | AKAP8 | 3.29E-21 | 0.637855647 | 0.325 | 0.187 | 6.49E-17 | Tb |
| allcell | NSD2 | 2.41E-23 | 0.637647427 | 0.285 | 0.154 | 4.75E-19 | Tb |
| allcell | SMURF2 | 3.10E-34 | 0.637288674 | 0.502 | 0.303 | 6.09E-30 | Tb |
| allcell | PPP2CB | 2.18E-26 | 0.63688949 | 0.387 | 0.226 | 4.29E-22 | Tb |
| allcell | HIVEP2 | 5.20E-28 | 0.636860542 | 0.726 | 0.563 | 1.02E-23 | Tb |
| allcell | NEK9 | 5.80E-20 | 0.636654026 | 0.258 | 0.141 | 1.14E-15 | Tb |
| allcell | RYBP | 2.02E-38 | 0.635986629 | 0.69 | 0.481 | 3.99E-34 | Tb |
| allcell | ARID1A | 1.34E-34 | 0.635576742 | 0.494 | 0.294 | 2.65E-30 | Tb |
| allcell | KDM4B | 2.89E-37 | 0.634994285 | 0.503 | 0.291 | 5.70E-33 | Tb |
| allcell | EPB41 | 1.98E-19 | 0.634938125 | 0.438 | 0.294 | 3.91E-15 | Tb |
| allcell | ATP2B4 | 1.44E-18 | 0.634745533 | 0.329 | 0.204 | 2.83E-14 | Tb |
| allcell | PACS1 | 3.25E-22 | 0.634591832 | 0.538 | 0.385 | 6.39E-18 | Tb |
| allcell | CPEB2 | 2.23E-14 | 0.634160596 | 0.372 | 0.253 | 4.39E-10 | Tb |
| allcell | AQR | 3.51E-21 | 0.6337895 | 0.321 | 0.188 | 6.92E-17 | Tb |
| allcell | SRBD1 | 1.39E-14 | 0.633644426 | 0.27 | 0.168 | 2.75E-10 | Tb |
| allcell | XPO6 | 2.65E-26 | 0.632806164 | 0.362 | 0.205 | 5.22E-22 | Tb |
| allcell | SEPHS1 | 1.88E-20 | 0.632667502 | 0.334 | 0.199 | 3.70E-16 | Tb |
| allcell | TXNDC5 | 3.57E-13 | 0.632587696 | 0.259 | 0.165 | 7.02E-09 | Tb |
| allcell | WDR44 | 9.01E-20 | 0.632541154 | 0.334 | 0.203 | 1.77E-15 | Tb |
| allcell | FGD4 | 2.94E-19 | 0.632244429 | 0.453 | 0.316 | 5.79E-15 | Tb |
| allcell | CRLF3 | 4.12E-21 | 0.632032794 | 0.361 | 0.22 | 8.11E-17 | Tb |
| allcell | ZNF518A | 2.54E-21 | 0.631719515 | 0.327 | 0.193 | 5.00E-17 | Tb |
| allcell | BCL2L13 | 2.27E-21 | 0.631384457 | 0.452 | 0.3 | 4.47E-17 | Tb |
| allcell | NCOA3 | 1.14E-29 | 0.631334581 | 0.671 | 0.498 | 2.23E-25 | Tb |
| allcell | ANKRD33B | 6.76E-08 | 0.631168991 | 0.349 | 0.277 | 0.00133062 | Tb |
| allcell | PHLDB3 | 4.60E-22 | 0.630732726 | 0.344 | 0.206 | 9.06E-18 | Tb |
| allcell | PKNOX1 | 1.11E-20 | 0.630623793 | 0.295 | 0.168 | 2.18E-16 | Tb |
| allcell | PTPN22 | 6.28E-12 | 0.630367893 | 0.418 | 0.31 | 1.24E-07 | Tb |
| allcell | RSRC1 | 3.70E-33 | 0.63027996 | 0.53 | 0.327 | 7.27E-29 | Tb |
| allcell | DIAPH2 | 1.12E-25 | 0.630245813 | 0.5 | 0.326 | 2.21E-21 | Tb |
| allcell | AFF1 | 4.49E-34 | 0.630089053 | 0.742 | 0.549 | 8.84E-30 | Tb |
| allcell | VEZT | 6.73E-27 | 0.629907549 | 0.429 | 0.259 | 1.32E-22 | Tb |
| allcell | PLXNC1 | 1.83E-24 | 0.629459757 | 0.535 | 0.366 | 3.59E-20 | Tb |
| allcell | ZSWIM6 | 5.03E-43 | 0.62930254 | 0.838 | 0.674 | 9.90E-39 | Tb |
| allcell | ZNF710 | 4.21E-27 | 0.629133635 | 0.575 | 0.401 | 8.29E-23 | Tb |
| allcell | UBE2E2 | 1.83E-39 | 0.628813224 | 0.827 | 0.694 | 3.61E-35 | Tb |
| allcell | TET3 | 6.16E-29 | 0.628076379 | 0.412 | 0.24 | 1.21E-24 | Tb |
| allcell | HERC3 | 2.14E-19 | 0.62752948 | 0.269 | 0.154 | 4.22E-15 | Tb |
| allcell | PCNX1 | 6.29E-26 | 0.627264561 | 0.567 | 0.391 | 1.24E-21 | Tb |
| allcell | TPK1 | 1.39E-20 | 0.626647631 | 0.403 | 0.256 | 2.74E-16 | Tb |

| | | | | | | | |
|---|---|---|---|---|---|---|---|
| allcell | KRT5 | 1.64E-45 | 0.625609678 | 0.32 | 0.138 | 3.23E-41 | Tb |
| allcell | HEATR5B | 2.11E-19 | 0.625590894 | 0.331 | 0.2 | 4.16E-15 | Tb |
| allcell | TMEM131L | 2.59E-23 | 0.625566387 | 0.458 | 0.299 | 5.09E-19 | Tb |
| allcell | GNB4 | 2.25E-26 | 0.625355642 | 0.534 | 0.359 | 4.43E-22 | Tb |
| allcell | RIOK1 | 2.49E-22 | 0.625170454 | 0.369 | 0.22 | 4.91E-18 | Tb |
| allcell | SIPA1L3 | 1.13E-39 | 0.624901327 | 0.569 | 0.365 | 2.22E-35 | Tb |
| allcell | RBM47 | 1.15E-31 | 0.624516435 | 0.738 | 0.578 | 2.26E-27 | Tb |
| allcell | APPBP2 | 1.55E-28 | 0.624189189 | 0.335 | 0.177 | 3.06E-24 | Tb |
| allcell | MGAT2 | 5.87E-23 | 0.624157649 | 0.282 | 0.15 | 1.15E-18 | Tb |
| allcell | R3HCC1L | 2.30E-23 | 0.624118499 | 0.363 | 0.214 | 4.52E-19 | Tb |
| allcell | SF3B3 | 8.05E-19 | 0.624094297 | 0.27 | 0.153 | 1.58E-14 | Tb |
| allcell | NSD3 | 1.90E-22 | 0.622353448 | 0.656 | 0.483 | 3.75E-18 | Tb |
| allcell | TRPM7 | 2.21E-29 | 0.621566069 | 0.481 | 0.297 | 4.35E-25 | Tb |
| allcell | MAPKAP1 | 1.55E-28 | 0.621561234 | 0.561 | 0.369 | 3.05E-24 | Tb |
| allcell | GOLPH3 | 3.61E-29 | 0.621535627 | 0.442 | 0.261 | 7.11E-25 | Tb |
| allcell | COX19 | 5.51E-20 | 0.621171582 | 0.351 | 0.213 | 1.09E-15 | Tb |
| allcell | PLCG2 | 1.85E-24 | 0.620898444 | 0.475 | 0.306 | 3.65E-20 | Tb |
| allcell | SMC6 | 4.81E-16 | 0.620454317 | 0.293 | 0.184 | 9.48E-12 | Tb |
| allcell | NF1 | 4.45E-26 | 0.620342013 | 0.544 | 0.373 | 8.77E-22 | Tb |
| allcell | B4GALT4 | 7.75E-19 | 0.620278777 | 0.337 | 0.209 | 1.53E-14 | Tb |
| allcell | ASH1L | 3.59E-40 | 0.620107877 | 0.756 | 0.556 | 7.06E-36 | Tb |
| allcell | USP37 | 6.10E-20 | 0.620064911 | 0.422 | 0.279 | 1.20E-15 | Tb |
| allcell | STRBP | 9.90E-19 | 0.619846289 | 0.441 | 0.303 | 1.95E-14 | Tb |
| allcell | MFSD4B | 1.62E-22 | 0.619581782 | 0.404 | 0.249 | 3.18E-18 | Tb |
| allcell | GGA2 | 2.58E-25 | 0.618948782 | 0.428 | 0.267 | 5.07E-21 | Tb |
| allcell | CPB2-AS1 | 2.70E-26 | 0.618627977 | 0.393 | 0.229 | 5.31E-22 | Tb |
| allcell | RBM26 | 5.75E-25 | 0.618538781 | 0.421 | 0.257 | 1.13E-20 | Tb |
| allcell | TRMO | 3.64E-18 | 0.618480965 | 0.266 | 0.152 | 7.17E-14 | Tb |
| allcell | EPG5 | 1.14E-21 | 0.618088613 | 0.267 | 0.144 | 2.25E-17 | Tb |
| allcell | CLOCK | 3.29E-12 | 0.61796071 | 0.255 | 0.164 | 6.48E-08 | Tb |
| allcell | YTHDC2 | 1.27E-23 | 0.617304164 | 0.513 | 0.338 | 2.51E-19 | Tb |
| allcell | ASAP1 | 2.13E-21 | 0.616861296 | 0.782 | 0.713 | 4.19E-17 | Tb |
| allcell | BCAS3 | 1.11E-26 | 0.616861208 | 0.399 | 0.241 | 2.19E-22 | Tb |
| allcell | CYREN | 1.52E-19 | 0.61666429 | 0.301 | 0.175 | 3.00E-15 | Tb |
| allcell | KAT6A | 3.64E-22 | 0.616310438 | 0.48 | 0.322 | 7.17E-18 | Tb |
| allcell | GALM | 4.67E-11 | 0.614968751 | 0.372 | 0.273 | 9.19E-07 | Tb |
| allcell | SLC35E1 | 5.85E-24 | 0.614837962 | 0.337 | 0.19 | 1.15E-19 | Tb |
| allcell | FBXO28 | 2.14E-20 | 0.614524444 | 0.269 | 0.149 | 4.20E-16 | Tb |
| allcell | ZC3H8 | 5.09E-20 | 0.614132225 | 0.26 | 0.144 | 1.00E-15 | Tb |
| allcell | PAK1 | 2.58E-27 | 0.613470408 | 0.848 | 0.729 | 5.08E-23 | Tb |
| allcell | LPIN2 | 1.95E-28 | 0.612969001 | 0.467 | 0.285 | 3.83E-24 | Tb |
| allcell | RALGAPB | 6.64E-15 | 0.612916942 | 0.253 | 0.151 | 1.31E-10 | Tb |
| allcell | LIMD1 | 1.14E-22 | 0.612684458 | 0.323 | 0.187 | 2.24E-18 | Tb |
| allcell | ABCC4 | 3.36E-19 | 0.612637561 | 0.28 | 0.162 | 6.62E-15 | Tb |
| allcell | TENT4A | 1.47E-28 | 0.612306296 | 0.471 | 0.287 | 2.90E-24 | Tb |
| allcell | TRAF3 | 2.78E-44 | 0.611966445 | 0.727 | 0.511 | 5.47E-40 | Tb |
| allcell | TYMP | 2.92E-47 | 0.611431545 | 0.913 | 0.797 | 5.75E-43 | Tb |
| allcell | PPP6R2 | 4.40E-33 | 0.6114 | 0.51 | 0.31 | 8.66E-29 | Tb |
| allcell | EXOC4 | 7.44E-39 | 0.611251699 | 0.663 | 0.445 | 1.47E-34 | Tb |
| allcell | MT-ND2 | 3.13E-73 | 0.610338315 | 0.973 | 0.882 | 6.17E-69 | Tb |
| allcell | CPEB4 | 7.27E-34 | 0.610324414 | 0.589 | 0.389 | 1.43E-29 | Tb |
| allcell | UBE2W | 1.36E-27 | 0.610286701 | 0.65 | 0.487 | 2.68E-23 | Tb |
| allcell | SGMS2 | 2.08E-14 | 0.610084489 | 0.274 | 0.172 | 4.09E-10 | Tb |
| allcell | HECTD4 | 8.83E-16 | 0.609977784 | 0.306 | 0.195 | 1.74E-11 | Tb |
| allcell | CHD8 | 4.82E-23 | 0.609918413 | 0.332 | 0.192 | 9.48E-19 | Tb |
| allcell | THUMPD3-AS1 | 1.84E-32 | 0.609905617 | 0.542 | 0.341 | 3.63E-28 | Tb |

| | | | | | | | |
|---|---|---|---|---|---|---|---|
| allcell | P2RY13 | 7.04E-21 | 0.609769782 | 0.328 | 0.195 | 1.39E-16 | Tb |
| allcell | WASHC5 | 5.78E-13 | 0.609546118 | 0.277 | 0.179 | 1.14E-08 | Tb |
| allcell | TTC9C | 2.72E-28 | 0.609392464 | 0.297 | 0.149 | 5.35E-24 | Tb |
| allcell | STAG1 | 5.16E-38 | 0.608852543 | 0.647 | 0.43 | 1.02E-33 | Tb |
| allcell | IDI1 | 1.88E-23 | 0.608661359 | 0.67 | 0.501 | 3.71E-19 | Tb |
| allcell | DAPP1 | 7.31E-26 | 0.608550206 | 0.766 | 0.64 | 1.44E-21 | Tb |
| allcell | MON2 | 5.69E-24 | 0.608343426 | 0.442 | 0.283 | 1.12E-19 | Tb |
| allcell | SPOP | 1.55E-27 | 0.608246375 | 0.498 | 0.316 | 3.05E-23 | Tb |
| allcell | TP53 | 9.12E-21 | 0.608075612 | 0.282 | 0.157 | 1.80E-16 | Tb |
| allcell | KIAA0232 | 1.55E-21 | 0.607829754 | 0.367 | 0.222 | 3.05E-17 | Tb |
| allcell | KDM2B | 4.83E-11 | 0.607818191 | 0.493 | 0.379 | 9.50E-07 | Tb |
| allcell | ZNF146 | 1.42E-23 | 0.607362305 | 0.281 | 0.147 | 2.80E-19 | Tb |
| allcell | AEBP2 | 2.41E-18 | 0.607233594 | 0.386 | 0.252 | 4.74E-14 | Tb |
| allcell | KDM3A | 1.41E-22 | 0.607137914 | 0.305 | 0.171 | 2.77E-18 | Tb |
| allcell | STAM2 | 1.14E-18 | 0.606944604 | 0.289 | 0.17 | 2.24E-14 | Tb |
| allcell | IGF2BP2 | 1.44E-22 | 0.606931035 | 0.358 | 0.219 | 2.84E-18 | Tb |
| allcell | IL21R | 2.91E-16 | 0.606523096 | 0.303 | 0.192 | 5.73E-12 | Tb |
| allcell | GALNT1 | 1.50E-30 | 0.606088746 | 0.485 | 0.295 | 2.94E-26 | Tb |
| allcell | MNAT1 | 3.29E-20 | 0.606014628 | 0.342 | 0.208 | 6.47E-16 | Tb |
| allcell | SKAP2 | 4.93E-30 | 0.605911564 | 0.633 | 0.449 | 9.71E-26 | Tb |
| allcell | CEP135 | 1.42E-19 | 0.605074069 | 0.306 | 0.182 | 2.79E-15 | Tb |
| allcell | FBXO11 | 1.08E-37 | 0.604939301 | 0.782 | 0.6 | 2.13E-33 | Tb |
| allcell | USP6NL | 1.83E-18 | 0.604634329 | 0.316 | 0.193 | 3.60E-14 | Tb |
| allcell | HEATR5A | 3.17E-20 | 0.604382655 | 0.293 | 0.169 | 6.23E-16 | Tb |
| allcell | SATB1-AS1 | 1.45E-17 | 0.603749526 | 0.286 | 0.172 | 2.86E-13 | Tb |
| allcell | LCOR | 2.25E-29 | 0.603100288 | 0.588 | 0.402 | 4.42E-25 | Tb |
| allcell | RIC1 | 4.86E-25 | 0.60259104 | 0.589 | 0.419 | 9.56E-21 | Tb |
| allcell | PLXDC2 | 1.10E-17 | 0.601945307 | 0.556 | 0.432 | 2.17E-13 | Tb |
| allcell | EHMT1 | 8.70E-40 | 0.601027583 | 0.542 | 0.322 | 1.71E-35 | Tb |
| allcell | PPP2R5E | 3.17E-26 | 0.600969085 | 0.462 | 0.292 | 6.24E-22 | Tb |
| allcell | INO80 | 9.65E-24 | 0.600805291 | 0.406 | 0.248 | 1.90E-19 | Tb |
| allcell | TNRC6B | 1.20E-27 | 0.600447711 | 0.664 | 0.483 | 2.36E-23 | Tb |
| allcell | CCNT1 | 1.71E-22 | 0.600259854 | 0.386 | 0.238 | 3.37E-18 | Tb |
| allcell | PRPSAP1 | 3.39E-18 | 0.600205592 | 0.286 | 0.166 | 6.68E-14 | Tb |
| allcell | ATP11B | 1.46E-29 | 0.599422929 | 0.562 | 0.37 | 2.88E-25 | Tb |
| allcell | MIB1 | 3.01E-25 | 0.599347473 | 0.293 | 0.156 | 5.92E-21 | Tb |
| allcell | HMBOX1 | 1.02E-24 | 0.599329741 | 0.417 | 0.255 | 2.01E-20 | Tb |
| allcell | NSRP1 | 1.08E-26 | 0.599186592 | 0.459 | 0.279 | 2.12E-22 | Tb |
| allcell | CSTF3 | 4.47E-22 | 0.599186522 | 0.28 | 0.151 | 8.80E-18 | Tb |
| allcell | ERI1 | 4.12E-19 | 0.598272034 | 0.316 | 0.19 | 8.10E-15 | Tb |
| allcell | MEGF9 | 6.12E-10 | 0.598034234 | 0.262 | 0.18 | 1.21E-05 | Tb |
| allcell | SLC27A4 | 3.59E-28 | 0.597951257 | 0.437 | 0.259 | 7.06E-24 | Tb |
| allcell | PTPN7 | 2.80E-15 | 0.597713893 | 0.264 | 0.16 | 5.52E-11 | Tb |
| allcell | TGFBR2 | 5.29E-14 | 0.597353999 | 0.526 | 0.398 | 1.04E-09 | Tb |
| allcell | PHF14 | 5.05E-26 | 0.596713596 | 0.451 | 0.279 | 9.95E-22 | Tb |
| allcell | CARS2 | 3.67E-23 | 0.596639034 | 0.328 | 0.184 | 7.22E-19 | Tb |
| allcell | LY75 | 2.08E-18 | 0.595959796 | 0.491 | 0.341 | 4.09E-14 | Tb |
| allcell | CEBPA | 3.29E-15 | 0.595191343 | 0.272 | 0.169 | 6.48E-11 | Tb |
| allcell | SRSF4 | 8.91E-35 | 0.594614646 | 0.668 | 0.448 | 1.75E-30 | Tb |
| allcell | MAP3K13 | 1.38E-23 | 0.594129596 | 0.643 | 0.485 | 2.72E-19 | Tb |
| allcell | LNPEP | 8.37E-24 | 0.593751898 | 0.469 | 0.297 | 1.65E-19 | Tb |
| allcell | EED | 1.96E-22 | 0.593608509 | 0.421 | 0.261 | 3.86E-18 | Tb |
| allcell | FOXK1 | 1.27E-17 | 0.59353238 | 0.303 | 0.184 | 2.51E-13 | Tb |
| allcell | HSH2D | 1.25E-20 | 0.593164548 | 0.299 | 0.171 | 2.47E-16 | Tb |
| allcell | PLEKHA5 | 3.73E-07 | 0.593013898 | 0.43 | 0.343 | 0.007334339 | Tb |
| allcell | RALGAPA2 | 1.15E-23 | 0.592805053 | 0.3 | 0.166 | 2.27E-19 | Tb |

| | | | | | | | |
|---|---|---|---|---|---|---|---|
| allcell | N4BP1 | 5.33E-29 | 0.592743131 | 0.607 | 0.425 | 1.05E-24 | Tb |
| allcell | SRPK2 | 2.12E-22 | 0.592637269 | 0.539 | 0.377 | 4.17E-18 | Tb |
| allcell | SGMS1 | 1.84E-21 | 0.592628024 | 0.345 | 0.208 | 3.62E-17 | Tb |
| allcell | PMEPA1 | 9.35E-26 | 0.592498567 | 0.375 | 0.219 | 1.84E-21 | Tb |
| allcell | RUNX1 | 1.42E-33 | 0.592310192 | 0.805 | 0.651 | 2.79E-29 | Tb |
| allcell | RNF19B | 2.07E-17 | 0.592225997 | 0.442 | 0.307 | 4.07E-13 | Tb |
| allcell | CDK8 | 3.43E-16 | 0.591938145 | 0.267 | 0.16 | 6.76E-12 | Tb |
| allcell | FAM107B | 1.01E-28 | 0.591738007 | 0.762 | 0.601 | 1.99E-24 | Tb |
| allcell | SHB | 1.19E-14 | 0.591670266 | 0.314 | 0.202 | 2.33E-10 | Tb |
| allcell | SCARF1 | 4.42E-22 | 0.591149045 | 0.361 | 0.217 | 8.70E-18 | Tb |
| allcell | PBX3 | 4.52E-28 | 0.591039518 | 0.603 | 0.424 | 8.89E-24 | Tb |
| allcell | RFX3 | 5.49E-19 | 0.590650812 | 0.34 | 0.21 | 1.08E-14 | Tb |
| allcell | ENTPD7 | 1.35E-19 | 0.590569753 | 0.306 | 0.183 | 2.65E-15 | Tb |
| allcell | SLC15A4 | 1.47E-26 | 0.590246107 | 0.517 | 0.338 | 2.89E-22 | Tb |
| allcell | NFATC1 | 6.23E-21 | 0.590219397 | 0.434 | 0.286 | 1.23E-16 | Tb |
| allcell | KHDC4 | 4.52E-17 | 0.589747213 | 0.29 | 0.175 | 8.89E-13 | Tb |
| allcell | UIMC1 | 1.42E-24 | 0.589602508 | 0.353 | 0.2 | 2.80E-20 | Tb |
| allcell | RNF34 | 2.69E-28 | 0.589464557 | 0.369 | 0.204 | 5.30E-24 | Tb |
| allcell | AOPEP | 6.88E-23 | 0.589443477 | 0.286 | 0.158 | 1.35E-18 | Tb |
| allcell | BPTF | 1.30E-35 | 0.589357178 | 0.675 | 0.467 | 2.55E-31 | Tb |
| allcell | CELF2 | 8.54E-37 | 0.58898184 | 0.843 | 0.673 | 1.68E-32 | Tb |
| allcell | ADNP | 3.85E-27 | 0.588843289 | 0.485 | 0.308 | 7.58E-23 | Tb |
| allcell | SLC31A1 | 4.14E-16 | 0.588761138 | 0.307 | 0.195 | 8.14E-12 | Tb |
| allcell | KMT5B | 5.75E-21 | 0.587948306 | 0.334 | 0.196 | 1.13E-16 | Tb |
| allcell | NR4A3 | 8.28E-22 | 0.587431645 | 0.847 | 0.784 | 1.63E-17 | Tb |
| allcell | PIBF1 | 4.29E-18 | 0.587399337 | 0.31 | 0.191 | 8.45E-14 | Tb |
| allcell | TRERF1 | 8.28E-11 | 0.586967108 | 0.378 | 0.277 | 1.63E-06 | Tb |
| allcell | ADK | 2.74E-29 | 0.586741756 | 0.617 | 0.441 | 5.40E-25 | Tb |
| allcell | DNAJC5 | 2.99E-26 | 0.586358598 | 0.443 | 0.268 | 5.89E-22 | Tb |
| allcell | NAA15 | 4.72E-21 | 0.586154346 | 0.35 | 0.21 | 9.28E-17 | Tb |
| allcell | ANKRD11 | 2.25E-52 | 0.585755667 | 0.87 | 0.693 | 4.43E-48 | Tb |
| allcell | PI4KA | 3.64E-18 | 0.585718891 | 0.389 | 0.249 | 7.16E-14 | Tb |
| allcell | FOXJ3 | 5.27E-23 | 0.585642502 | 0.401 | 0.247 | 1.04E-18 | Tb |
| allcell | INSR | 3.59E-25 | 0.585354369 | 0.283 | 0.153 | 7.06E-21 | Tb |
| allcell | TGFBR1 | 4.90E-30 | 0.585286522 | 0.582 | 0.39 | 9.64E-26 | Tb |
| allcell | TCAIM | 2.12E-16 | 0.584692593 | 0.264 | 0.154 | 4.17E-12 | Tb |
| allcell | SCYL2 | 4.30E-28 | 0.5844111 | 0.447 | 0.267 | 8.47E-24 | Tb |
| allcell | ATF7IP | 7.13E-28 | 0.583990723 | 0.612 | 0.435 | 1.40E-23 | Tb |
| allcell | IL15 | 6.13E-17 | 0.583975788 | 0.508 | 0.379 | 1.21E-12 | Tb |
| allcell | TSPAN13 | 1.15E-15 | 0.583692386 | 0.417 | 0.299 | 2.26E-11 | Tb |
| allcell | ARID1B | 4.59E-48 | 0.583506469 | 0.792 | 0.569 | 9.03E-44 | Tb |
| allcell | FRYL | 1.05E-32 | 0.582773595 | 0.63 | 0.423 | 2.07E-28 | Tb |
| allcell | PARN | 1.42E-18 | 0.582398943 | 0.304 | 0.181 | 2.79E-14 | Tb |
| allcell | GNAQ | 1.05E-46 | 0.582298698 | 0.813 | 0.629 | 2.06E-42 | Tb |
| allcell | TTC14 | 1.42E-20 | 0.582236246 | 0.268 | 0.146 | 2.79E-16 | Tb |
| allcell | UHRF2 | 4.56E-18 | 0.580885088 | 0.271 | 0.157 | 8.97E-14 | Tb |
| allcell | SETD5 | 1.65E-34 | 0.580493596 | 0.57 | 0.361 | 3.24E-30 | Tb |
| allcell | FBXW7 | 2.84E-33 | 0.579786591 | 0.718 | 0.547 | 5.59E-29 | Tb |
| allcell | AKT1S1 | 1.38E-17 | 0.577906372 | 0.255 | 0.144 | 2.73E-13 | Tb |
| allcell | GPATCH11 | 8.11E-14 | 0.577526379 | 0.281 | 0.177 | 1.60E-09 | Tb |
| allcell | FOXO3 | 5.23E-19 | 0.576634208 | 0.49 | 0.348 | 1.03E-14 | Tb |
| allcell | AHCYL1 | 2.51E-21 | 0.576571153 | 0.46 | 0.302 | 4.94E-17 | Tb |
| allcell | MAN1A2 | 1.52E-23 | 0.576423201 | 0.448 | 0.285 | 2.98E-19 | Tb |
| allcell | FLT3 | 4.88E-08 | 0.575911926 | 0.5 | 0.42 | 0.000960262 | Tb |
| allcell | ZBTB38 | 2.96E-08 | 0.575680814 | 0.31 | 0.227 | 0.000583135 | Tb |
| allcell | CSF2RA | 3.02E-24 | 0.575469531 | 0.78 | 0.664 | 5.95E-20 | Tb |

| | | | | | | | |
|---|---|---|---|---|---|---|---|
| allcell | STX5 | 7.77E-28 | 0.575442633 | 0.507 | 0.321 | 1.53E-23 | Tb |
| allcell | ACOT13 | 6.55E-27 | 0.575354875 | 0.328 | 0.175 | 1.29E-22 | Tb |
| allcell | STX8 | 2.00E-26 | 0.575255239 | 0.562 | 0.376 | 3.94E-22 | Tb |
| allcell | RELL1 | 6.18E-24 | 0.575176567 | 0.391 | 0.24 | 1.22E-19 | Tb |
| allcell | RNF213 | 1.30E-35 | 0.574959258 | 0.702 | 0.482 | 2.56E-31 | Tb |
| allcell | PDCD4 | 2.95E-19 | 0.574923755 | 0.46 | 0.31 | 5.81E-15 | Tb |
| allcell | MED14 | 3.01E-28 | 0.574871308 | 0.331 | 0.176 | 5.92E-24 | Tb |
| allcell | NUP88 | 2.52E-19 | 0.574463125 | 0.251 | 0.138 | 4.96E-15 | Tb |
| allcell | PITPNB | 1.81E-28 | 0.574398051 | 0.55 | 0.358 | 3.57E-24 | Tb |
| allcell | GAS2L3 | 8.26E-09 | 0.573632768 | 0.268 | 0.19 | 0.000162647 | Tb |
| allcell | EGR2 | 2.02E-25 | 0.573602248 | 0.277 | 0.143 | 3.98E-21 | Tb |
| allcell | UBE3C | 6.24E-22 | 0.573595992 | 0.397 | 0.245 | 1.23E-17 | Tb |
| allcell | STX12 | 6.27E-22 | 0.573539015 | 0.628 | 0.474 | 1.23E-17 | Tb |
| allcell | OPA3 | 9.88E-23 | 0.573278774 | 0.363 | 0.214 | 1.94E-18 | Tb |
| allcell | HPS5 | 8.24E-36 | 0.572676503 | 0.671 | 0.462 | 1.62E-31 | Tb |
| allcell | TMX2 | 8.00E-25 | 0.572050543 | 0.299 | 0.157 | 1.57E-20 | Tb |
| allcell | PTPRA | 2.01E-19 | 0.571381034 | 0.44 | 0.292 | 3.96E-15 | Tb |
| allcell | CYFIP2 | 4.42E-14 | 0.570655197 | 0.352 | 0.24 | 8.70E-10 | Tb |
| allcell | MYOF | 1.27E-24 | 0.570480267 | 0.528 | 0.362 | 2.49E-20 | Tb |
| allcell | PRKAG1 | 4.80E-23 | 0.570128633 | 0.361 | 0.211 | 9.45E-19 | Tb |
| allcell | DNAJC13 | 4.14E-18 | 0.570018465 | 0.318 | 0.195 | 8.14E-14 | Tb |
| allcell | CEP85L | 6.44E-14 | 0.569538656 | 0.321 | 0.211 | 1.27E-09 | Tb |
| allcell | PRKAR2A | 1.72E-20 | 0.569223145 | 0.432 | 0.281 | 3.38E-16 | Tb |
| allcell | LRIF1 | 7.59E-37 | 0.56910037 | 0.696 | 0.479 | 1.49E-32 | Tb |
| allcell | RNF103 | 6.15E-19 | 0.569049335 | 0.329 | 0.2 | 1.21E-14 | Tb |
| allcell | CUX1 | 4.53E-33 | 0.568612561 | 0.75 | 0.573 | 8.92E-29 | Tb |
| allcell | CLSTN1 | 1.58E-21 | 0.568353882 | 0.276 | 0.152 | 3.10E-17 | Tb |
| allcell | EEPD1 | 1.77E-10 | 0.568267973 | 0.267 | 0.182 | 3.49E-06 | Tb |
| allcell | IL1R1 | 5.25E-09 | 0.568197817 | 0.348 | 0.262 | 0.000103304 | Tb |
| allcell | TLE4 | 1.15E-13 | 0.567755343 | 0.47 | 0.349 | 2.26E-09 | Tb |
| allcell | TAF1 | 1.37E-18 | 0.567314407 | 0.274 | 0.157 | 2.69E-14 | Tb |
| allcell | WDFY1 | 5.58E-18 | 0.567068211 | 0.279 | 0.164 | 1.10E-13 | Tb |
| allcell | SS18 | 5.27E-20 | 0.566742522 | 0.367 | 0.223 | 1.04E-15 | Tb |
| allcell | NAA35 | 2.68E-22 | 0.565986543 | 0.274 | 0.149 | 5.27E-18 | Tb |
| allcell | TTC3 | 6.67E-30 | 0.565932078 | 0.621 | 0.426 | 1.31E-25 | Tb |
| allcell | SEC31A | 1.09E-30 | 0.56572449 | 0.627 | 0.425 | 2.14E-26 | Tb |
| allcell | PPM1A | 2.94E-19 | 0.564946186 | 0.308 | 0.181 | 5.79E-15 | Tb |
| allcell | CHAF1A | 4.76E-22 | 0.564911419 | 0.479 | 0.321 | 9.37E-18 | Tb |
| allcell | CWC27 | 5.20E-14 | 0.564592124 | 0.255 | 0.157 | 1.02E-09 | Tb |
| allcell | ZNF292 | 1.00E-32 | 0.564127592 | 0.586 | 0.379 | 1.98E-28 | Tb |
| allcell | IKBKB | 2.91E-28 | 0.564091448 | 0.344 | 0.185 | 5.73E-24 | Tb |
| allcell | SLC41A2 | 6.00E-14 | 0.563670506 | 0.531 | 0.403 | 1.18E-09 | Tb |
| allcell | MSH3 | 2.13E-18 | 0.563609002 | 0.274 | 0.158 | 4.19E-14 | Tb |
| allcell | LPCAT3 | 2.71E-25 | 0.563272007 | 0.251 | 0.123 | 5.33E-21 | Tb |
| allcell | KRCC1 | 1.15E-23 | 0.563220997 | 0.292 | 0.158 | 2.27E-19 | Tb |
| allcell | PHKB | 3.05E-21 | 0.562994702 | 0.411 | 0.26 | 6.00E-17 | Tb |
| allcell | PARP11 | 5.93E-18 | 0.562416782 | 0.277 | 0.161 | 1.17E-13 | Tb |
| allcell | FAM193A | 9.02E-19 | 0.561585847 | 0.335 | 0.206 | 1.78E-14 | Tb |
| allcell | RNGTT | 4.66E-20 | 0.561472556 | 0.365 | 0.226 | 9.17E-16 | Tb |
| allcell | FAM241A | 3.23E-18 | 0.561396177 | 0.512 | 0.365 | 6.36E-14 | Tb |
| allcell | EAF2 | 2.09E-19 | 0.560759711 | 0.479 | 0.324 | 4.11E-15 | Tb |
| allcell | SLC1A4 | 3.85E-21 | 0.560547853 | 0.321 | 0.188 | 7.58E-17 | Tb |
| allcell | RAB11A | 2.20E-29 | 0.560107106 | 0.819 | 0.697 | 4.32E-25 | Tb |
| allcell | ATP11C | 3.17E-18 | 0.559735977 | 0.307 | 0.187 | 6.24E-14 | Tb |
| allcell | EXOSC6 | 1.75E-14 | 0.559112034 | 0.272 | 0.168 | 3.45E-10 | Tb |
| allcell | CD86 | 2.71E-09 | 0.558754359 | 0.726 | 0.675 | 5.34E-05 | Tb |

| | | | | | | | |
|---|---|---|---|---|---|---|---|
| allcell | MAP4 | 4.77E-28 | 0.558199874 | 0.632 | 0.446 | 9.39E-24 | Tb |
| allcell | RPF2 | 1.69E-16 | 0.557790967 | 0.295 | 0.178 | 3.33E-12 | Tb |
| allcell | PDE8A | 5.23E-17 | 0.557055412 | 0.339 | 0.219 | 1.03E-12 | Tb |
| allcell | PTPRJ | 4.00E-40 | 0.556788399 | 0.749 | 0.572 | 7.88E-36 | Tb |
| allcell | CCNYL1 | 7.98E-21 | 0.556436246 | 0.453 | 0.297 | 1.57E-16 | Tb |
| allcell | TNPO3 | 8.79E-23 | 0.556339758 | 0.38 | 0.23 | 1.73E-18 | Tb |
| allcell | ATF6 | 7.31E-21 | 0.556322106 | 0.636 | 0.473 | 1.44E-16 | Tb |
| allcell | TIAM11 | 5.35E-18 | 0.555094708 | 0.418 | 0.285 | 1.05E-13 | Tb |
| allcell | MAML3 | 1.71E-17 | 0.554707126 | 0.529 | 0.397 | 3.37E-13 | Tb |
| allcell | NR6A1 | 1.89E-15 | 0.554359054 | 0.262 | 0.158 | 3.73E-11 | Tb |
| allcell | PPM1K | 2.57E-26 | 0.554118671 | 0.491 | 0.311 | 5.06E-22 | Tb |
| allcell | SPECC1 | 2.91E-13 | 0.553830481 | 0.458 | 0.333 | 5.74E-09 | Tb |
| allcell | PIK3R5 | 1.81E-22 | 0.553651931 | 0.622 | 0.471 | 3.56E-18 | Tb |
| allcell | APTX | 4.16E-22 | 0.553640524 | 0.385 | 0.234 | 8.18E-18 | Tb |
| allcell | MCPH1 | 9.29E-14 | 0.553432964 | 0.28 | 0.18 | 1.83E-09 | Tb |
| allcell | SULF2 | 2.14E-28 | 0.552819065 | 0.735 | 0.564 | 4.20E-24 | Tb |
| allcell | USO1 | 7.34E-26 | 0.552174996 | 0.498 | 0.322 | 1.45E-21 | Tb |
| allcell | TSPOAP1-AS1 | 6.26E-21 | 0.552014278 | 0.345 | 0.206 | 1.23E-16 | Tb |
| allcell | HMGB2 | 2.24E-19 | 0.551415962 | 0.556 | 0.392 | 4.40E-15 | Tb |
| allcell | ESCO1 | 8.62E-21 | 0.551233052 | 0.386 | 0.238 | 1.70E-16 | Tb |
| allcell | SPEN | 8.16E-20 | 0.550907458 | 0.599 | 0.445 | 1.61E-15 | Tb |
| allcell | WNK1 | 3.57E-26 | 0.550777885 | 0.672 | 0.502 | 7.03E-22 | Tb |
| allcell | SLC4A7 | 7.75E-20 | 0.550292684 | 0.479 | 0.32 | 1.52E-15 | Tb |
| allcell | CTCF | 3.29E-21 | 0.54977157 | 0.36 | 0.215 | 6.47E-17 | Tb |
| allcell | FAM13B | 1.01E-22 | 0.549159628 | 0.495 | 0.332 | 1.99E-18 | Tb |
| allcell | AMBRA1 | 6.59E-26 | 0.549075249 | 0.526 | 0.343 | 1.30E-21 | Tb |
| allcell | MIR4435-2HG | 7.24E-29 | 0.549036012 | 0.658 | 0.473 | 1.43E-24 | Tb |
| allcell | EFL1 | 2.10E-15 | 0.548758439 | 0.254 | 0.15 | 4.13E-11 | Tb |
| allcell | KANSL1L | 1.20E-16 | 0.548422469 | 0.355 | 0.227 | 2.36E-12 | Tb |
| allcell | DMTF1 | 6.19E-20 | 0.548371888 | 0.283 | 0.16 | 1.22E-15 | Tb |
| allcell | MTF2 | 1.03E-24 | 0.548329738 | 0.383 | 0.225 | 2.03E-20 | Tb |
| allcell | PPP6R3 | 8.99E-29 | 0.548241367 | 0.633 | 0.439 | 1.77E-24 | Tb |
| allcell | SPPL3 | 8.93E-25 | 0.548091987 | 0.6 | 0.431 | 1.76E-20 | Tb |
| allcell | CASC4 | 7.08E-11 | 0.547893931 | 0.328 | 0.233 | 1.39E-06 | Tb |
| allcell | MAPK8 | 6.81E-28 | 0.547079865 | 0.569 | 0.391 | 1.34E-23 | Tb |
| allcell | CAAP1 | 2.17E-17 | 0.546252621 | 0.324 | 0.201 | 4.27E-13 | Tb |
| allcell | GPATCH8 | 4.17E-20 | 0.54595072 | 0.465 | 0.317 | 8.21E-16 | Tb |
| allcell | TVP23C | 3.68E-18 | 0.545443931 | 0.25 | 0.142 | 7.24E-14 | Tb |
| allcell | PWWP2A | 1.38E-16 | 0.545125518 | 0.269 | 0.159 | 2.71E-12 | Tb |
| allcell | COG5 | 5.71E-25 | 0.545056959 | 0.5 | 0.323 | 1.12E-20 | Tb |
| allcell | EIPR1 | 1.39E-18 | 0.545010996 | 0.262 | 0.148 | 2.74E-14 | Tb |
| allcell | ZNF644 | 2.18E-28 | 0.544886939 | 0.635 | 0.447 | 4.29E-24 | Tb |
| allcell | TAOK3 | 2.59E-35 | 0.543905327 | 0.849 | 0.721 | 5.10E-31 | Tb |
| allcell | CCNG2 | 3.21E-20 | 0.54385253 | 0.463 | 0.312 | 6.32E-16 | Tb |
| allcell | DNAJC1 | 6.17E-19 | 0.543527507 | 0.616 | 0.47 | 1.21E-14 | Tb |
| allcell | PRRC2B | 4.29E-21 | 0.543462062 | 0.437 | 0.283 | 8.44E-17 | Tb |
| allcell | GLB1 | 1.54E-21 | 0.542819081 | 0.361 | 0.214 | 3.02E-17 | Tb |
| allcell | RAB6A | 4.07E-18 | 0.542541862 | 0.436 | 0.29 | 8.02E-14 | Tb |
| allcell | ASMTL | 6.30E-22 | 0.542482692 | 0.461 | 0.306 | 1.24E-17 | Tb |
| allcell | PIP4K2A | 6.24E-20 | 0.541681517 | 0.579 | 0.417 | 1.23E-15 | Tb |
| allcell | ZDHHC17 | 1.63E-17 | 0.541584957 | 0.438 | 0.299 | 3.20E-13 | Tb |
| allcell | SLC8A1 | 2.19E-15 | 0.541291994 | 0.691 | 0.622 | 4.31E-11 | Tb |
| allcell | APBB1IP | 6.37E-16 | 0.541235837 | 0.652 | 0.517 | 1.25E-11 | Tb |
| allcell | WSB2 | 7.49E-17 | 0.541211327 | 0.311 | 0.19 | 1.47E-12 | Tb |
| allcell | NETO2 | 1.81E-07 | 0.540834661 | 0.262 | 0.192 | 0.003557885 | Tb |
| allcell | PPM1B | 3.42E-19 | 0.540797806 | 0.446 | 0.304 | 6.74E-15 | Tb |

| | | | | | | | |
|---|---|---|---|---|---|---|---|
| allcell | RABGEF1 | 1.19E-24 | 0.54054796 | 0.658 | 0.497 | 2.34E-20 | Tb |
| allcell | LYST | 2.54E-15 | 0.540247327 | 0.548 | 0.415 | 5.00E-11 | Tb |
| allcell | GALNT2 | 1.17E-15 | 0.539977833 | 0.403 | 0.278 | 2.30E-11 | Tb |
| allcell | UBR3 | 2.10E-22 | 0.539096654 | 0.348 | 0.205 | 4.14E-18 | Tb |
| allcell | DCTN4 | 7.54E-21 | 0.538835626 | 0.465 | 0.308 | 1.48E-16 | Tb |
| allcell | CTDSPL2 | 1.46E-18 | 0.538715444 | 0.354 | 0.221 | 2.87E-14 | Tb |
| allcell | PTEN | 2.72E-28 | 0.538705946 | 0.661 | 0.473 | 5.36E-24 | Tb |
| allcell | PIK3C3 | 2.88E-18 | 0.538275063 | 0.297 | 0.175 | 5.66E-14 | Tb |
| allcell | ATP6V1A | 4.74E-18 | 0.537920765 | 0.576 | 0.439 | 9.34E-14 | Tb |
| allcell | GGA3 | 3.35E-21 | 0.53786444 | 0.25 | 0.131 | 6.59E-17 | Tb |
| allcell | SCAPER | 3.28E-18 | 0.537674849 | 0.362 | 0.23 | 6.45E-14 | Tb |
| allcell | ZNF131 | 3.90E-24 | 0.537586068 | 0.453 | 0.289 | 7.68E-20 | Tb |
| allcell | CUL1 | 7.77E-20 | 0.536991761 | 0.624 | 0.468 | 1.53E-15 | Tb |
| allcell | SUMF1 | 9.41E-28 | 0.536928033 | 0.533 | 0.348 | 1.85E-23 | Tb |
| allcell | ABHD17B | 6.37E-14 | 0.536270341 | 0.34 | 0.227 | 1.25E-09 | Tb |
| allcell | TM9SF4 | 6.81E-19 | 0.535864344 | 0.252 | 0.139 | 1.34E-14 | Tb |
| allcell | DCAF6 | 1.53E-20 | 0.535511571 | 0.389 | 0.246 | 3.02E-16 | Tb |
| allcell | BTAF1 | 6.69E-28 | 0.535248482 | 0.615 | 0.42 | 1.32E-23 | Tb |
| allcell | PTRH2 | 1.59E-17 | 0.535098687 | 0.308 | 0.185 | 3.12E-13 | Tb |
| allcell | LARP4B | 1.48E-35 | 0.534925635 | 0.635 | 0.415 | 2.91E-31 | Tb |
| allcell | RNF149 | 9.86E-16 | 0.534842976 | 0.777 | 0.671 | 1.94E-11 | Tb |
| allcell | NOCT | 1.10E-15 | 0.534682532 | 0.289 | 0.18 | 2.17E-11 | Tb |
| allcell | LPP | 3.20E-40 | 0.534667799 | 0.78 | 0.585 | 6.29E-36 | Tb |
| allcell | KIF2A | 4.18E-13 | 0.534002467 | 0.629 | 0.496 | 8.23E-09 | Tb |
| allcell | KLHL28 | 1.23E-20 | 0.533887815 | 0.302 | 0.173 | 2.43E-16 | Tb |
| allcell | RSPRY1 | 4.81E-18 | 0.533660006 | 0.292 | 0.172 | 9.48E-14 | Tb |
| allcell | XIAP | 3.76E-19 | 0.533471261 | 0.361 | 0.223 | 7.40E-15 | Tb |
| allcell | MTMR6 | 7.96E-21 | 0.533273346 | 0.407 | 0.251 | 1.57E-16 | Tb |
| allcell | HIVEP1 | 2.13E-23 | 0.533179019 | 0.752 | 0.635 | 4.19E-19 | Tb |
| allcell | HECTD1 | 4.84E-27 | 0.533051867 | 0.512 | 0.326 | 9.53E-23 | Tb |
| allcell | PDCD7 | 7.23E-17 | 0.532845595 | 0.321 | 0.199 | 1.42E-12 | Tb |
| allcell | TUG1 | 3.13E-17 | 0.532706855 | 0.253 | 0.146 | 6.17E-13 | Tb |
| allcell | GCC2 | 5.74E-26 | 0.53179126 | 0.586 | 0.402 | 1.13E-21 | Tb |
| allcell | FAM117B | 1.17E-07 | 0.53141782 | 0.279 | 0.207 | 0.002309066 | Tb |
| allcell | APC | 1.66E-17 | 0.531365456 | 0.348 | 0.22 | 3.26E-13 | Tb |
| allcell | ATF5 | 6.79E-13 | 0.531062185 | 0.659 | 0.539 | 1.34E-08 | Tb |
| allcell | ATP8A1 | 1.79E-12 | 0.531053199 | 0.273 | 0.179 | 3.52E-08 | Tb |
| allcell | SNAPC1 | 1.06E-07 | 0.530979133 | 0.257 | 0.183 | 0.002084509 | Tb |
| allcell | PDXDC1 | 6.98E-24 | 0.530899893 | 0.383 | 0.226 | 1.37E-19 | Tb |
| allcell | WDFY2 | 2.50E-30 | 0.530789462 | 0.601 | 0.414 | 4.92E-26 | Tb |
| allcell | MBTPS1 | 2.37E-10 | 0.530488413 | 0.253 | 0.167 | 4.67E-06 | Tb |
| allcell | OFD1 | 3.53E-28 | 0.530461783 | 0.572 | 0.393 | 6.95E-24 | Tb |
| allcell | ETNK1 | 2.51E-30 | 0.529897865 | 0.6 | 0.397 | 4.95E-26 | Tb |
| allcell | TBC1D5 | 8.25E-24 | 0.529815066 | 0.626 | 0.464 | 1.62E-19 | Tb |
| allcell | KLHL18 | 3.38E-14 | 0.529483262 | 0.281 | 0.175 | 6.65E-10 | Tb |
| allcell | SOS1 | 9.20E-26 | 0.529135304 | 0.542 | 0.365 | 1.81E-21 | Tb |
| allcell | RC3H2 | 4.59E-20 | 0.528887431 | 0.348 | 0.214 | 9.04E-16 | Tb |
| allcell | RAPGEF6 | 6.07E-22 | 0.528853996 | 0.57 | 0.402 | 1.20E-17 | Tb |
| allcell | NBAS | 2.04E-21 | 0.528553854 | 0.344 | 0.204 | 4.01E-17 | Tb |
| allcell | TRAPPC8 | 4.30E-20 | 0.528418819 | 0.338 | 0.204 | 8.46E-16 | Tb |
| allcell | CCAR1 | 1.06E-27 | 0.528034543 | 0.517 | 0.331 | 2.08E-23 | Tb |
| allcell | CD274 | 4.11E-15 | 0.527091474 | 0.346 | 0.234 | 8.09E-11 | Tb |
| allcell | IL4R | 3.92E-20 | 0.527026753 | 0.397 | 0.252 | 7.71E-16 | Tb |
| allcell | USP14 | 1.07E-18 | 0.526872295 | 0.486 | 0.332 | 2.10E-14 | Tb |
| allcell | ZNF91 | 7.75E-19 | 0.52671278 | 0.302 | 0.175 | 1.53E-14 | Tb |
| allcell | MTMR12 | 1.52E-19 | 0.526380274 | 0.311 | 0.182 | 2.98E-15 | Tb |

| | | | | | | | |
|---|---|---|---|---|---|---|---|
| allcell | TLK2 | 4.24E-23 | 0.526290563 | 0.425 | 0.262 | 8.34E-19 | Tb |
| allcell | FNIP1 | 5.39E-31 | 0.526167294 | 0.604 | 0.401 | 1.06E-26 | Tb |
| allcell | AFF4 | 5.08E-34 | 0.526001273 | 0.815 | 0.662 | 1.00E-29 | Tb |
| allcell | DGKH | 3.99E-17 | 0.525750716 | 0.279 | 0.167 | 7.85E-13 | Tb |
| allcell | UBASH3B | 1.55E-11 | 0.525706271 | 0.437 | 0.325 | 3.05E-07 | Tb |
| allcell | NFE2L3 | 1.63E-15 | 0.525460937 | 0.547 | 0.405 | 3.21E-11 | Tb |
| allcell | TSC22D2 | 4.75E-21 | 0.525334507 | 0.621 | 0.461 | 9.34E-17 | Tb |
| allcell | OTUD5 | 1.79E-18 | 0.52529715 | 0.36 | 0.227 | 3.52E-14 | Tb |
| allcell | ATP9B | 1.00E-19 | 0.525122348 | 0.559 | 0.404 | 1.97E-15 | Tb |
| allcell | MINK1 | 1.41E-22 | 0.524958986 | 0.299 | 0.166 | 2.78E-18 | Tb |
| allcell | TFDP2 | 2.75E-20 | 0.524958645 | 0.421 | 0.268 | 5.41E-16 | Tb |
| allcell | SLC30A5 | 5.43E-15 | 0.524208894 | 0.276 | 0.17 | 1.07E-10 | Tb |
| allcell | ARID3A | 5.89E-16 | 0.523920815 | 0.441 | 0.307 | 1.16E-11 | Tb |
| allcell | CASK | 1.05E-15 | 0.523778916 | 0.304 | 0.191 | 2.07E-11 | Tb |
| allcell | ACO2 | 4.56E-22 | 0.523750667 | 0.419 | 0.261 | 8.98E-18 | Tb |
| allcell | FH | 3.31E-14 | 0.523502504 | 0.353 | 0.232 | 6.52E-10 | Tb |
| allcell | NUP153 | 7.93E-23 | 0.52329109 | 0.411 | 0.253 | 1.56E-18 | Tb |
| allcell | SLC30A9 | 5.41E-22 | 0.523252899 | 0.355 | 0.21 | 1.07E-17 | Tb |
| allcell | VPS13D | 5.26E-20 | 0.522247033 | 0.451 | 0.307 | 1.04E-15 | Tb |
| allcell | GMDS | 1.95E-19 | 0.52198051 | 0.562 | 0.406 | 3.84E-15 | Tb |
| allcell | UBE2H | 3.28E-28 | 0.521687733 | 0.693 | 0.521 | 6.45E-24 | Tb |
| allcell | TBC1D14 | 8.13E-18 | 0.521300505 | 0.341 | 0.213 | 1.60E-13 | Tb |
| allcell | IGF2R | 1.60E-33 | 0.521242688 | 0.696 | 0.479 | 3.15E-29 | Tb |
| allcell | JAK2 | 2.04E-23 | 0.521074082 | 0.714 | 0.555 | 4.01E-19 | Tb |
| allcell | HERC1 | 1.51E-24 | 0.520671666 | 0.627 | 0.453 | 2.97E-20 | Tb |
| allcell | ZNF506 | 3.20E-16 | 0.52062814 | 0.285 | 0.171 | 6.29E-12 | Tb |
| allcell | FAM49B | 1.70E-37 | 0.520458956 | 0.916 | 0.783 | 3.35E-33 | Tb |
| allcell | ATM | 8.08E-18 | 0.520241954 | 0.357 | 0.23 | 1.59E-13 | Tb |
| allcell | ST3GAL4 | 7.81E-18 | 0.519700353 | 0.277 | 0.161 | 1.54E-13 | Tb |
| allcell | ACSL3 | 8.56E-16 | 0.519674704 | 0.64 | 0.491 | 1.69E-11 | Tb |
| allcell | CYTOR | 2.72E-16 | 0.519662396 | 0.691 | 0.584 | 5.36E-12 | Tb |
| allcell | MTR | 9.57E-20 | 0.519582108 | 0.316 | 0.188 | 1.88E-15 | Tb |
| allcell | MRPL1 | 2.50E-19 | 0.519360505 | 0.308 | 0.182 | 4.92E-15 | Tb |
| allcell | AP2B1 | 9.57E-23 | 0.519294254 | 0.5 | 0.326 | 1.88E-18 | Tb |
| allcell | KCTD5 | 6.37E-16 | 0.519273216 | 0.346 | 0.223 | 1.25E-11 | Tb |
| allcell | NUS1 | 1.23E-16 | 0.519169618 | 0.301 | 0.182 | 2.42E-12 | Tb |
| allcell | CELF1 | 1.63E-21 | 0.519133997 | 0.486 | 0.32 | 3.20E-17 | Tb |
| allcell | ZBTB10 | 1.57E-16 | 0.519062313 | 0.593 | 0.462 | 3.08E-12 | Tb |
| allcell | WDR7 | 2.30E-24 | 0.519048833 | 0.368 | 0.215 | 4.52E-20 | Tb |
| allcell | ATP11A | 5.05E-21 | 0.519034532 | 0.358 | 0.215 | 9.94E-17 | Tb |
| allcell | N4BP2L2 | 2.47E-34 | 0.517693321 | 0.74 | 0.529 | 4.85E-30 | Tb |
| allcell | SESN3 | 2.49E-10 | 0.517654682 | 0.397 | 0.3 | 4.91E-06 | Tb |
| allcell | TNKS | 7.45E-17 | 0.5173738 | 0.378 | 0.25 | 1.47E-12 | Tb |
| allcell | SWAP70 | 4.03E-20 | 0.516846516 | 0.485 | 0.338 | 7.93E-16 | Tb |
| allcell | TRIM69 | 5.65E-24 | 0.51620785 | 0.586 | 0.422 | 1.11E-19 | Tb |
| allcell | CEP68 | 1.64E-23 | 0.515618197 | 0.438 | 0.275 | 3.23E-19 | Tb |
| allcell | KAT2B | 7.20E-24 | 0.515220199 | 0.578 | 0.404 | 1.42E-19 | Tb |
| allcell | ITGA4 | 5.20E-17 | 0.515109135 | 0.651 | 0.517 | 1.02E-12 | Tb |
| allcell | OSGIN2 | 1.08E-17 | 0.514926587 | 0.261 | 0.149 | 2.12E-13 | Tb |
| allcell | TASOR | 1.43E-18 | 0.514878553 | 0.346 | 0.215 | 2.81E-14 | Tb |
| allcell | SNX13 | 8.88E-16 | 0.514479823 | 0.365 | 0.237 | 1.75E-11 | Tb |
| allcell | HSD17B4 | 5.49E-17 | 0.514119042 | 0.346 | 0.217 | 1.08E-12 | Tb |
| allcell | ASXL2 | 1.31E-19 | 0.514093159 | 0.369 | 0.232 | 2.57E-15 | Tb |
| allcell | CNTLN | 2.66E-15 | 0.513762865 | 0.356 | 0.237 | 5.24E-11 | Tb |
| allcell | MAPK14 | 7.93E-21 | 0.513748322 | 0.398 | 0.25 | 1.56E-16 | Tb |
| allcell | BANP | 5.26E-20 | 0.513461859 | 0.646 | 0.491 | 1.04E-15 | Tb |

| | | | | | | | |
|---|---|---|---|---|---|---|---|
| allcell | PHF20L1 | 1.32E-16 | 0.512629396 | 0.507 | 0.358 | 2.61E-12 | Tb |
| allcell | PARP8 | 5.20E-19 | 0.512141798 | 0.463 | 0.321 | 1.02E-14 | Tb |
| allcell | YLPM1 | 9.93E-16 | 0.511849042 | 0.291 | 0.178 | 1.95E-11 | Tb |
| allcell | CCDC66 | 5.68E-15 | 0.511690338 | 0.271 | 0.165 | 1.12E-10 | Tb |
| allcell | HNRNPUL1 | 6.58E-25 | 0.511673249 | 0.649 | 0.462 | 1.30E-20 | Tb |
| allcell | ZCCHC17 | 1.21E-19 | 0.511592418 | 0.359 | 0.218 | 2.38E-15 | Tb |
| allcell | TBC1D9 | 5.10E-16 | 0.511326277 | 0.487 | 0.352 | 1.00E-11 | Tb |
| allcell | KIF13B | 3.02E-11 | 0.510623743 | 0.391 | 0.283 | 5.95E-07 | Tb |
| allcell | AAK1 | 3.81E-17 | 0.51025441 | 0.376 | 0.245 | 7.50E-13 | Tb |
| allcell | CEP63 | 2.13E-14 | 0.510031435 | 0.325 | 0.21 | 4.19E-10 | Tb |
| allcell | SND1 | 2.67E-31 | 0.509692193 | 0.742 | 0.53 | 5.26E-27 | Tb |
| allcell | STAU1 | 6.08E-23 | 0.509661569 | 0.547 | 0.364 | 1.20E-18 | Tb |
| allcell | CALU | 7.28E-17 | 0.509500401 | 0.343 | 0.217 | 1.43E-12 | Tb |
| allcell | CSPP1 | 8.39E-13 | 0.509367187 | 0.26 | 0.164 | 1.65E-08 | Tb |
| allcell | QSOX1 | 2.37E-15 | 0.508328692 | 0.295 | 0.184 | 4.67E-11 | Tb |
| allcell | SSH2 | 2.20E-22 | 0.508119024 | 0.609 | 0.446 | 4.33E-18 | Tb |
| allcell | FNDC3A | 1.85E-25 | 0.507955964 | 0.665 | 0.491 | 3.64E-21 | Tb |
| allcell | BBX | 7.91E-26 | 0.507816608 | 0.617 | 0.437 | 1.56E-21 | Tb |
| allcell | IMMP2L | 6.93E-11 | 0.507662367 | 0.385 | 0.287 | 1.36E-06 | Tb |
| allcell | FRMD4B | 3.93E-14 | 0.507392155 | 0.476 | 0.353 | 7.74E-10 | Tb |
| allcell | SPN | 6.85E-23 | 0.507288563 | 0.393 | 0.242 | 1.35E-18 | Tb |
| allcell | PPM1D | 5.20E-18 | 0.50681776 | 0.39 | 0.252 | 1.02E-13 | Tb |
| allcell | FLNB | 2.37E-15 | 0.50681145 | 0.281 | 0.178 | 4.66E-11 | Tb |
| allcell | BCOR | 2.60E-19 | 0.506611641 | 0.276 | 0.155 | 5.12E-15 | Tb |
| allcell | ADGRE5 | 2.40E-20 | 0.506518465 | 0.601 | 0.442 | 4.73E-16 | Tb |
| allcell | SIL1 | 3.61E-19 | 0.506242148 | 0.462 | 0.315 | 7.11E-15 | Tb |
| allcell | AKAP13 | 4.86E-38 | 0.505723582 | 0.952 | 0.878 | 9.56E-34 | Tb |
| allcell | SLC7A5 | 1.80E-16 | 0.505536835 | 0.586 | 0.447 | 3.55E-12 | Tb |
| allcell | PXK | 2.92E-17 | 0.504897646 | 0.29 | 0.173 | 5.76E-13 | Tb |
| allcell | HMGCS1 | 1.58E-14 | 0.504773622 | 0.432 | 0.299 | 3.11E-10 | Tb |
| allcell | DHRSX | 8.65E-23 | 0.504689416 | 0.463 | 0.295 | 1.70E-18 | Tb |
| allcell | ATG10 | 9.77E-16 | 0.504548508 | 0.251 | 0.149 | 1.92E-11 | Tb |
| allcell | ZC3H13 | 3.65E-23 | 0.50438064 | 0.481 | 0.307 | 7.18E-19 | Tb |
| allcell | ST14 | 1.70E-13 | 0.504254592 | 0.252 | 0.159 | 3.35E-09 | Tb |
| allcell | MDM4 | 1.65E-20 | 0.50376335 | 0.495 | 0.331 | 3.24E-16 | Tb |
| allcell | ATAD2B | 1.64E-27 | 0.503216212 | 0.648 | 0.456 | 3.23E-23 | Tb |
| allcell | LRRC8B | 1.17E-20 | 0.503193931 | 0.319 | 0.185 | 2.31E-16 | Tb |
| allcell | DENND10 | 1.37E-14 | 0.503115094 | 0.255 | 0.151 | 2.69E-10 | Tb |
| allcell | SLC35F5 | 1.57E-15 | 0.503039884 | 0.281 | 0.17 | 3.08E-11 | Tb |
| allcell | AKNA | 9.57E-20 | 0.503032004 | 0.42 | 0.276 | 1.88E-15 | Tb |
| allcell | RAB4B | 1.14E-23 | 0.503014501 | 0.472 | 0.301 | 2.24E-19 | Tb |
| allcell | CHSY1 | 1.05E-12 | 0.502738927 | 0.254 | 0.161 | 2.06E-08 | Tb |
| allcell | MKNK1 | 8.51E-18 | 0.502620705 | 0.304 | 0.183 | 1.67E-13 | Tb |
| allcell | TAF2 | 3.37E-14 | 0.502156232 | 0.283 | 0.179 | 6.63E-10 | Tb |
| allcell | XPNPEP1 | 2.14E-19 | 0.501002316 | 0.317 | 0.186 | 4.21E-15 | Tb |
| allcell | POLG2 | 5.31E-10 | 0.500138467 | 0.304 | 0.213 | 1.05E-05 | Tb |
| group | | p_val | avg_log2FC | pct.1 | pct.2 | p_val_adj | cluster |
| allcell | APOE | 8.54E-30 | 2.52591241 | 0.436 | 0.273 | 1.68E-25 | Lep-L |
| allcell | APOC1 | 4.10E-10 | 2.213247562 | 0.331 | 0.247 | 8.06E-06 | Lep-L |
| allcell | PLIN2 | 1.67E-20 | 2.10511571 | 0.402 | 0.27 | 3.29E-16 | Lep-L |
| allcell | C5AR1 | 7.03E-19 | 2.081133459 | 0.285 | 0.16 | 1.38E-14 | Lep-L |
| allcell | IL1R2 | 2.22E-46 | 1.851453984 | 0.521 | 0.273 | 4.36E-42 | Lep-L |
| allcell | CXCL3 | 2.76E-23 | 1.848924132 | 0.365 | 0.202 | 5.44E-19 | Lep-L |
| allcell | HMOX1 | 1.79E-14 | 1.824431916 | 0.465 | 0.381 | 3.53E-10 | Lep-L |
| allcell | KCNN4 | 2.60E-23 | 1.791993319 | 0.297 | 0.151 | 5.12E-19 | Lep-L |
| allcell | PID1 | 1.79E-17 | 1.773613851 | 0.353 | 0.219 | 3.52E-13 | Lep-L |

| | | | | | | | |
|---|---|---|---|---|---|---|---|
| allcell | MID1IP1 | 8.28E-07 | 1.613503948 | 0.275 | 0.209 | 0.016306958 | Lep-L |
| allcell | CXCL8 | 2.16E-61 | 1.493964307 | 0.705 | 0.433 | 4.25E-57 | Lep-L |
| allcell | HSPA6 | 6.28E-37 | 1.492688372 | 0.626 | 0.417 | 1.24E-32 | Lep-L |
| allcell | BAG3 | 2.38E-40 | 1.434337747 | 0.725 | 0.547 | 4.68E-36 | Lep-L |
| allcell | PAF1 | 3.43E-22 | 1.429786908 | 0.447 | 0.304 | 6.76E-18 | Lep-L |
| allcell | ZC3H12A | 1.58E-29 | 1.422136812 | 0.608 | 0.472 | 3.11E-25 | Lep-L |
| allcell | ITGA5 | 2.07E-18 | 1.381866266 | 0.377 | 0.246 | 4.07E-14 | Lep-L |
| allcell | DDX3Y | 4.91E-34 | 1.355014907 | 0.494 | 0.269 | 9.66E-30 | Lep-L |
| allcell | BNIP3L | 6.97E-12 | 1.342181937 | 0.635 | 0.588 | 1.37E-07 | Lep-L |
| allcell | FKBP4 | 2.53E-47 | 1.320272538 | 0.65 | 0.42 | 4.99E-43 | Lep-L |
| allcell | ATP1A1 | 5.97E-40 | 1.299640851 | 0.747 | 0.641 | 1.17E-35 | Lep-L |
| allcell | TUBB2A | 3.86E-08 | 1.297918683 | 0.423 | 0.355 | 0.000760003 | Lep-L |
| allcell | CEBPB | 1.37E-14 | 1.297093835 | 0.538 | 0.458 | 2.70E-10 | Lep-L |
| allcell | SERPINH1 | 1.14E-33 | 1.293242419 | 0.453 | 0.251 | 2.24E-29 | Lep-L |
| allcell | HSPB1 | 3.15E-52 | 1.281881387 | 0.868 | 0.742 | 6.21E-48 | Lep-L |
| allcell | SLC25A37 | 2.29E-14 | 1.276321131 | 0.384 | 0.272 | 4.51E-10 | Lep-L |
| allcell | NEU1 | 5.21E-11 | 1.275759353 | 0.428 | 0.354 | 1.03E-06 | Lep-L |
| allcell | RILPL2 | 3.21E-33 | 1.275611187 | 0.694 | 0.62 | 6.32E-29 | Lep-L |
| allcell | PHLDA1 | 6.47E-13 | 1.263872452 | 0.294 | 0.182 | 1.27E-08 | Lep-L |
| allcell | HSPA1B | 2.61E-20 | 1.257636585 | 0.869 | 0.877 | 5.14E-16 | Lep-L |
| allcell | CCR7 | 6.70E-12 | 1.253885207 | 0.462 | 0.353 | 1.32E-07 | Lep-L |
| allcell | FCGR2A | 3.04E-19 | 1.252324918 | 0.518 | 0.403 | 5.99E-15 | Lep-L |
| allcell | PDE4DIP | 1.33E-27 | 1.246751497 | 0.553 | 0.403 | 2.61E-23 | Lep-L |
| allcell | MMP9 | 1.05E-08 | 1.236377641 | 0.358 | 0.273 | 0.000206867 | Lep-L |
| allcell | ZFP36L1 | 1.72E-24 | 1.231869548 | 0.783 | 0.742 | 3.38E-20 | Lep-L |
| allcell | AHR | 2.96E-27 | 1.224303633 | 0.701 | 0.63 | 5.84E-23 | Lep-L |
| allcell | SMAD3 | 3.57E-11 | 1.221667804 | 0.368 | 0.27 | 7.02E-07 | Lep-L |
| allcell | SOD2 | 6.41E-27 | 1.216993728 | 0.896 | 0.877 | 1.26E-22 | Lep-L |
| allcell | AMPD2 | 3.95E-08 | 1.215967317 | 0.278 | 0.205 | 0.000778137 | Lep-L |
| allcell | TIMP1 | 4.36E-28 | 1.209862433 | 0.754 | 0.649 | 8.59E-24 | Lep-L |
| allcell | CA2 | 7.42E-11 | 1.189300407 | 0.448 | 0.357 | 1.46E-06 | Lep-L |
| allcell | SAT1 | 1.91E-82 | 1.18769241 | 0.981 | 0.975 | 3.77E-78 | Lep-L |
| allcell | MFSD2A | 9.67E-08 | 1.180511122 | 0.29 | 0.217 | 0.001902586 | Lep-L |
| allcell | NRARP | 2.10E-15 | 1.159526881 | 0.593 | 0.519 | 4.14E-11 | Lep-L |
| allcell | CDC42EP3 | 1.32E-25 | 1.159514304 | 0.621 | 0.48 | 2.59E-21 | Lep-L |
| allcell | HLA-DRB5 | 9.51E-65 | 1.13852355 | 0.873 | 0.718 | 1.87E-60 | Lep-L |
| allcell | PTTG1 | 3.68E-16 | 1.128630618 | 0.513 | 0.414 | 7.24E-12 | Lep-L |
| allcell | MAFG | 4.30E-12 | 1.11800552 | 0.37 | 0.277 | 8.46E-08 | Lep-L |
| allcell | MAP2K3 | 1.15E-42 | 1.112687338 | 0.771 | 0.64 | 2.27E-38 | Lep-L |
| allcell | CAPN2 | 1.53E-19 | 1.102417224 | 0.542 | 0.431 | 3.00E-15 | Lep-L |
| allcell | B3GNT2 | 3.10E-07 | 1.08700414 | 0.323 | 0.258 | 0.006100577 | Lep-L |
| allcell | RBFOX2 | 5.17E-20 | 1.077433944 | 0.635 | 0.572 | 1.02E-15 | Lep-L |
| allcell | DOK2 | 2.73E-14 | 1.072430146 | 0.467 | 0.357 | 5.36E-10 | Lep-L |
| allcell | MAP7D1 | 2.68E-19 | 1.071378992 | 0.518 | 0.412 | 5.27E-15 | Lep-L |
| allcell | RIT1 | 8.68E-13 | 1.061718206 | 0.411 | 0.316 | 1.71E-08 | Lep-L |
| allcell | PLEKHA5 | 1.45E-09 | 1.058054976 | 0.438 | 0.359 | 2.86E-05 | Lep-L |
| allcell | ALDOA | 4.73E-17 | 1.051678128 | 0.433 | 0.304 | 9.30E-13 | Lep-L |
| allcell | RHOF | 4.53E-11 | 1.0507983 | 0.55 | 0.503 | 8.92E-07 | Lep-L |
| allcell | SLC5A3 | 1.31E-08 | 1.044361323 | 0.385 | 0.306 | 0.000257871 | Lep-L |
| allcell | NFKBIE | 2.42E-14 | 1.031677736 | 0.579 | 0.516 | 4.76E-10 | Lep-L |
| allcell | HSPD1 | 1.67E-72 | 1.028514187 | 0.961 | 0.907 | 3.29E-68 | Lep-L |
| allcell | STIP1 | 2.05E-25 | 1.010550936 | 0.63 | 0.52 | 4.04E-21 | Lep-L |
| allcell | ABCA1 | 7.90E-09 | 1.005437996 | 0.43 | 0.368 | 0.000155579 | Lep-L |
| allcell | CSGALNACT2 | 8.02E-15 | 1.002863493 | 0.591 | 0.55 | 1.58E-10 | Lep-L |
| allcell | ERGIC1 | 9.16E-08 | 1.002247281 | 0.414 | 0.367 | 0.001802582 | Lep-L |
| allcell | MRPS6 | 5.18E-18 | 0.996729319 | 0.628 | 0.56 | 1.02E-13 | Lep-L |

| | | | | | | | |
|---|---|---|---|---|---|---|---|
| allcell | INSIG1 | 2.51E-30 | 0.989538917 | 0.762 | 0.635 | 4.95E-26 | Lep-L |
| allcell | PRNP | 5.36E-10 | 0.987205167 | 0.48 | 0.413 | 1.05E-05 | Lep-L |
| allcell | DNAJB1 | 1.09E-32 | 0.980876598 | 0.917 | 0.873 | 2.14E-28 | Lep-L |
| allcell | ARHGAP21 | 5.10E-10 | 0.978549037 | 0.36 | 0.273 | 1.00E-05 | Lep-L |
| allcell | HSPA1A | 3.25E-53 | 0.97317945 | 0.986 | 0.94 | 6.40E-49 | Lep-L |
| allcell | BIN1 | 2.24E-10 | 0.966444084 | 0.382 | 0.3 | 4.42E-06 | Lep-L |
| allcell | RARA | 1.09E-12 | 0.957428306 | 0.48 | 0.389 | 2.14E-08 | Lep-L |
| allcell | PPP1R18 | 1.93E-16 | 0.956123781 | 0.635 | 0.591 | 3.79E-12 | Lep-L |
| allcell | TSPYL2 | 8.36E-14 | 0.955186343 | 0.555 | 0.448 | 1.65E-09 | Lep-L |
| allcell | SLC6A6 | 2.89E-08 | 0.947546388 | 0.387 | 0.318 | 0.000568246 | Lep-L |
| allcell | TRIP10 | 1.20E-07 | 0.934001746 | 0.343 | 0.268 | 0.00235653 | Lep-L |
| allcell | ETS2 | 1.55E-12 | 0.927058984 | 0.56 | 0.487 | 3.04E-08 | Lep-L |
| allcell | CXCL2 | 1.45E-12 | 0.925664232 | 0.435 | 0.311 | 2.85E-08 | Lep-L |
| allcell | CHORDC1 | 1.01E-24 | 0.920045994 | 0.715 | 0.628 | 1.98E-20 | Lep-L |
| allcell | CACYBP | 6.45E-35 | 0.910614684 | 0.784 | 0.709 | 1.27E-30 | Lep-L |
| allcell | CD44 | 8.37E-29 | 0.906735654 | 0.874 | 0.81 | 1.65E-24 | Lep-L |
| allcell | CD9 | 9.18E-07 | 0.902648865 | 0.338 | 0.271 | 0.018064765 | Lep-L |
| allcell | TXNRD1 | 7.95E-10 | 0.9021268 | 0.598 | 0.601 | 1.57E-05 | Lep-L |
| allcell | TNFAIP3 | 3.13E-19 | 0.897691684 | 0.805 | 0.767 | 6.17E-15 | Lep-L |
| allcell | METRNL | 1.71E-15 | 0.891417196 | 0.684 | 0.654 | 3.37E-11 | Lep-L |
| allcell | TCP1 | 4.66E-33 | 0.889908818 | 0.759 | 0.679 | 9.18E-29 | Lep-L |
| allcell | ANPEP | 7.98E-07 | 0.883956087 | 0.389 | 0.32 | 0.015705038 | Lep-L |
| allcell | HSPE1 | 7.25E-37 | 0.879810904 | 0.93 | 0.927 | 1.43E-32 | Lep-L |
| allcell | ITGAV | 1.72E-08 | 0.87976671 | 0.306 | 0.22 | 0.000338215 | Lep-L |
| allcell | SDCBP | 6.27E-28 | 0.879279406 | 0.796 | 0.764 | 1.23E-23 | Lep-L |
| allcell | GPAT3 | 1.20E-22 | 0.869877657 | 0.594 | 0.438 | 2.36E-18 | Lep-L |
| allcell | HSP90AB1 | 2.79E-63 | 0.864350649 | 0.983 | 0.973 | 5.49E-59 | Lep-L |
| allcell | SERPINB9 | 1.04E-24 | 0.855740575 | 0.846 | 0.809 | 2.04E-20 | Lep-L |
| allcell | IER3 | 1.09E-19 | 0.855477741 | 0.745 | 0.645 | 2.15E-15 | Lep-L |
| allcell | MTSS1 | 7.25E-10 | 0.84302639 | 0.331 | 0.234 | 1.43E-05 | Lep-L |
| allcell | CD55 | 2.71E-11 | 0.841202023 | 0.803 | 0.803 | 5.34E-07 | Lep-L |
| allcell | MGAT1 | 7.51E-16 | 0.837568663 | 0.684 | 0.662 | 1.48E-11 | Lep-L |
| allcell | PPIF | 2.38E-14 | 0.830654704 | 0.681 | 0.633 | 4.68E-10 | Lep-L |
| allcell | EMP3 | 9.64E-29 | 0.830045265 | 0.82 | 0.793 | 1.90E-24 | Lep-L |
| allcell | HSPH1 | 1.33E-52 | 0.828067846 | 0.922 | 0.862 | 2.62E-48 | Lep-L |
| allcell | CLEC2B | 3.86E-11 | 0.826539091 | 0.732 | 0.716 | 7.59E-07 | Lep-L |
| allcell | PNPLA8 | 6.21E-12 | 0.819575591 | 0.598 | 0.565 | 1.22E-07 | Lep-L |
| allcell | CTTNBP2NL | 1.31E-11 | 0.818499496 | 0.543 | 0.469 | 2.58E-07 | Lep-L |
| allcell | PPTC7 | 7.28E-09 | 0.81468912 | 0.448 | 0.38 | 0.000143266 | Lep-L |
| allcell | NFKBIA | 1.51E-28 | 0.809380137 | 0.93 | 0.903 | 2.97E-24 | Lep-L |
| allcell | AHNAK | 3.25E-16 | 0.80343016 | 0.754 | 0.689 | 6.39E-12 | Lep-L |
| allcell | SLC43A3 | 8.92E-07 | 0.803181333 | 0.338 | 0.271 | 0.017555663 | Lep-L |
| allcell | SQSTM1 | 1.01E-36 | 0.801464169 | 0.917 | 0.868 | 1.99E-32 | Lep-L |
| allcell | HBP1 | 9.72E-15 | 0.796347402 | 0.535 | 0.455 | 1.91E-10 | Lep-L |
| allcell | HSPA8 | 3.45E-24 | 0.795580457 | 0.952 | 0.955 | 6.79E-20 | Lep-L |
| allcell | HECA | 1.53E-10 | 0.790772348 | 0.633 | 0.572 | 3.01E-06 | Lep-L |
| allcell | PPP1R15A | 2.26E-10 | 0.786479581 | 0.935 | 0.925 | 4.45E-06 | Lep-L |
| allcell | DNAJA1 | 2.16E-27 | 0.783182812 | 0.935 | 0.949 | 4.24E-23 | Lep-L |
| allcell | NFKB2 | 6.90E-09 | 0.776515834 | 0.482 | 0.42 | 0.000135755 | Lep-L |
| allcell | MYO9B | 1.97E-13 | 0.769655863 | 0.683 | 0.642 | 3.87E-09 | Lep-L |
| allcell | UPP1 | 1.11E-11 | 0.768600084 | 0.567 | 0.499 | 2.19E-07 | Lep-L |
| allcell | UBE2R2 | 1.59E-19 | 0.766907068 | 0.664 | 0.593 | 3.13E-15 | Lep-L |
| allcell | AHSA1 | 9.89E-13 | 0.762191998 | 0.601 | 0.56 | 1.95E-08 | Lep-L |
| allcell | DEDD2 | 2.85E-12 | 0.74956275 | 0.516 | 0.436 | 5.62E-08 | Lep-L |
| allcell | FTL | 1.17E-30 | 0.746546369 | 0.988 | 0.99 | 2.31E-26 | Lep-L |
| allcell | MRPL18 | 6.30E-12 | 0.744305565 | 0.727 | 0.746 | 1.24E-07 | Lep-L |

| | | | | | | | |
|---|---|---|---|---|---|---|---|
| allcell | ATP13A3 | 4.29E-11 | 0.743416885 | 0.589 | 0.532 | 8.45E-07 | Lep-L |
| allcell | BCL2A1 | 3.94E-34 | 0.739645517 | 0.851 | 0.782 | 7.76E-30 | Lep-L |
| allcell | GLA | 1.41E-17 | 0.733447546 | 0.71 | 0.62 | 2.78E-13 | Lep-L |
| allcell | SLC38A2 | 6.91E-13 | 0.733377005 | 0.784 | 0.758 | 1.36E-08 | Lep-L |
| allcell | LGALS3 | 1.75E-10 | 0.731892866 | 0.713 | 0.67 | 3.44E-06 | Lep-L |
| allcell | BTG1 | 8.41E-21 | 0.721929803 | 0.89 | 0.896 | 1.65E-16 | Lep-L |
| allcell | SGK1 | 3.31E-09 | 0.713052633 | 0.754 | 0.712 | 6.52E-05 | Lep-L |
| allcell | ST13 | 6.44E-30 | 0.711745788 | 0.812 | 0.781 | 1.27E-25 | Lep-L |
| allcell | LMNA | 1.94E-24 | 0.709728604 | 0.837 | 0.689 | 3.82E-20 | Lep-L |
| allcell | MCL1 | 1.67E-32 | 0.707469232 | 0.907 | 0.887 | 3.29E-28 | Lep-L |
| allcell | PTGES3 | 1.21E-22 | 0.698235952 | 0.83 | 0.842 | 2.39E-18 | Lep-L |
| allcell | IER5 | 4.37E-13 | 0.697729527 | 0.827 | 0.782 | 8.59E-09 | Lep-L |
| allcell | NINJ1 | 1.79E-10 | 0.693226371 | 0.625 | 0.59 | 3.53E-06 | Lep-L |
| allcell | ANXA1 | 1.51E-16 | 0.690435026 | 0.772 | 0.664 | 2.97E-12 | Lep-L |
| allcell | CEBPD | 5.40E-11 | 0.681985243 | 0.703 | 0.622 | 1.06E-06 | Lep-L |
| allcell | ZFP36 | 7.16E-07 | 0.681218411 | 0.91 | 0.94 | 0.014093431 | Lep-L |
| allcell | CXCL16 | 4.34E-12 | 0.678111495 | 0.647 | 0.596 | 8.55E-08 | Lep-L |
| allcell | FTH1 | 3.25E-33 | 0.676790519 | 0.993 | 0.993 | 6.39E-29 | Lep-L |
| allcell | EIF4E | 5.91E-08 | 0.675501856 | 0.55 | 0.503 | 0.001164157 | Lep-L |
| allcell | MIR181A1HG | 3.43E-08 | 0.674375353 | 0.586 | 0.529 | 0.000674872 | Lep-L |
| allcell | CHD2 | 2.24E-18 | 0.667906604 | 0.815 | 0.812 | 4.41E-14 | Lep-L |
| allcell | IL10RA | 1.79E-09 | 0.667137521 | 0.649 | 0.635 | 3.52E-05 | Lep-L |
| allcell | SLC2A3 | 1.27E-16 | 0.666151401 | 0.713 | 0.598 | 2.49E-12 | Lep-L |
| allcell | SLC3A2 | 2.74E-09 | 0.660472858 | 0.706 | 0.717 | 5.38E-05 | Lep-L |
| allcell | DDAH2 | 1.42E-09 | 0.657803944 | 0.606 | 0.583 | 2.79E-05 | Lep-L |
| allcell | UBC | 9.93E-20 | 0.656678385 | 0.975 | 0.98 | 1.95E-15 | Lep-L |
| allcell | YY1 | 2.97E-10 | 0.653011772 | 0.645 | 0.64 | 5.84E-06 | Lep-L |
| allcell | ANXA5 | 3.59E-29 | 0.650665447 | 0.864 | 0.799 | 7.06E-25 | Lep-L |
| allcell | MTCH1 | 6.94E-12 | 0.644162668 | 0.662 | 0.645 | 1.37E-07 | Lep-L |
| allcell | G3BP1 | 2.11E-06 | 0.638287735 | 0.57 | 0.561 | 0.041456076 | Lep-L |
| allcell | DDIT4 | 6.45E-16 | 0.635954094 | 0.718 | 0.619 | 1.27E-11 | Lep-L |
| allcell | CDKN1A | 9.62E-07 | 0.632823356 | 0.839 | 0.829 | 0.018926484 | Lep-L |
| allcell | EEF2 | 8.73E-18 | 0.632527513 | 0.827 | 0.889 | 1.72E-13 | Lep-L |
| allcell | EZR | 1.73E-27 | 0.631940079 | 0.902 | 0.873 | 3.41E-23 | Lep-L |
| allcell | CTSZ | 1.50E-24 | 0.631558547 | 0.91 | 0.926 | 2.95E-20 | Lep-L |
| allcell | ACBD3 | 1.39E-07 | 0.624538262 | 0.457 | 0.408 | 0.002730205 | Lep-L |
| allcell | CKS2 | 4.25E-11 | 0.623012509 | 0.766 | 0.754 | 8.36E-07 | Lep-L |
| allcell | ARL8B | 3.29E-12 | 0.619443987 | 0.706 | 0.685 | 6.48E-08 | Lep-L |
| allcell | STX11 | 1.76E-11 | 0.616246464 | 0.708 | 0.643 | 3.46E-07 | Lep-L |
| allcell | IL18 | 9.38E-07 | 0.615126157 | 0.511 | 0.462 | 0.018460726 | Lep-L |
| allcell | HSP90AA1 | 2.20E-44 | 0.613811051 | 0.988 | 0.986 | 4.33E-40 | Lep-L |
| allcell | SNAP23 | 1.61E-09 | 0.604460257 | 0.679 | 0.642 | 3.18E-05 | Lep-L |
| allcell | MXD1 | 1.51E-07 | 0.585692586 | 0.674 | 0.648 | 0.002977872 | Lep-L |
| allcell | DNTTIP2 | 1.44E-09 | 0.581569544 | 0.598 | 0.537 | 2.83E-05 | Lep-L |
| allcell | FXYD5 | 2.61E-09 | 0.570013971 | 0.716 | 0.716 | 5.15E-05 | Lep-L |
| allcell | GNA13 | 1.50E-20 | 0.567046979 | 0.862 | 0.791 | 2.96E-16 | Lep-L |
| allcell | CYTIP | 9.93E-12 | 0.561265541 | 0.783 | 0.808 | 1.95E-07 | Lep-L |
| allcell | DYNC1H1 | 4.33E-08 | 0.555872727 | 0.696 | 0.689 | 0.000851862 | Lep-L |
| allcell | CHMP1B | 9.21E-07 | 0.551700695 | 0.694 | 0.693 | 0.018138499 | Lep-L |
| allcell | RUNX3 | 1.63E-09 | 0.548726107 | 0.666 | 0.604 | 3.21E-05 | Lep-L |
| allcell | IRAK2 | 3.87E-07 | 0.543978044 | 0.503 | 0.436 | 0.007621108 | Lep-L |
| allcell | ZFAND2A | 1.95E-26 | 0.541133572 | 0.723 | 0.592 | 3.84E-22 | Lep-L |
| allcell | HNRNPA0 | 4.81E-07 | 0.529584145 | 0.638 | 0.662 | 0.009460941 | Lep-L |
| allcell | BCAR3 | 2.18E-06 | 0.528593134 | 0.367 | 0.293 | 0.042956048 | Lep-L |
| allcell | CLK1 | 3.05E-10 | 0.521632469 | 0.762 | 0.757 | 6.01E-06 | Lep-L |
| allcell | G0S2 | 5.45E-09 | 0.516730517 | 0.642 | 0.569 | 0.000107343 | Lep-L |

| | | | | | | | |
|---|---|---|---|---|---|---|---|
| allcell | CD83 | 1.69E-21 | 0.510741223 | 0.971 | 0.964 | 3.33E-17 | Lep-L |
| allcell | TSPAN33 | 8.78E-09 | 0.508407783 | 0.647 | 0.597 | 0.000172777 | Lep-L |
| allcell | DNAJB6 | 7.06E-24 | 0.50371849 | 0.89 | 0.879 | 1.39E-19 | Lep-L |
| allcell | H2AFZ | 6.63E-15 | 0.502215738 | 0.89 | 0.907 | 1.30E-10 | Lep-L |
| allcell | PNRC1 | 1.32E-14 | 0.500933076 | 0.946 | 0.964 | 2.60E-10 | Lep-L |
| allcell | BEST1 | 1.01E-49 | 2.164420032 | 0.251 | 0.1 | 1.98E-45 | Lep-T |
| allcell | ALDOA1 | 3.22E-73 | 2.081525428 | 0.449 | 0.247 | 6.35E-69 | Lep-T |
| allcell | RBFOX21 | 2.82E-80 | 1.549907853 | 0.698 | 0.514 | 5.55E-76 | Lep-T |
| allcell | APOO1 | 1.66E-44 | 1.453828407 | 0.421 | 0.26 | 3.27E-40 | Lep-T |
| allcell | NLRP3 | 3.34E-25 | 1.383245177 | 0.365 | 0.252 | 6.57E-21 | Lep-T |
| allcell | NFKBIA1 | 3.39E-67 | 1.358352041 | 0.946 | 0.883 | 6.67E-63 | Lep-T |
| allcell | ZFP361 | 1.65E-40 | 1.338634752 | 0.954 | 0.926 | 3.24E-36 | Lep-T |
| allcell | HCAR3 | 1.26E-26 | 1.299584418 | 0.252 | 0.137 | 2.49E-22 | Lep-T |
| allcell | CD74 | 1.24E-131 | 1.258890798 | 0.997 | 0.998 | 2.44E-127 | Lep-T |
| allcell | IL1B | 2.87E-28 | 1.253313564 | 0.583 | 0.478 | 5.64E-24 | Lep-T |
| allcell | TNF | 2.36E-34 | 1.223416576 | 0.49 | 0.342 | 4.64E-30 | Lep-T |
| allcell | CXCL81 | 2.45E-34 | 1.190654749 | 0.567 | 0.409 | 4.83E-30 | Lep-T |
| allcell | ICAM1 | 1.30E-50 | 1.107614661 | 0.76 | 0.646 | 2.56E-46 | Lep-T |
| allcell | PNRC11 | 1.67E-140 | 1.106113716 | 0.98 | 0.953 | 3.29E-136 | Lep-T |
| allcell | BCL31 | 1.44E-46 | 1.087210703 | 0.617 | 0.466 | 2.84E-42 | Lep-T |
| allcell | CDKN1A1 | 4.63E-89 | 1.057951814 | 0.885 | 0.799 | 9.12E-85 | Lep-T |
| allcell | HLA-DQA2 | 2.29E-80 | 1.054026717 | 0.728 | 0.567 | 4.51E-76 | Lep-T |
| allcell | ITGB2-AS1 | 1.43E-18 | 1.050926911 | 0.261 | 0.166 | 2.81E-14 | Lep-T |
| allcell | EMD1 | 1.93E-34 | 1.041816465 | 0.445 | 0.297 | 3.79E-30 | Lep-T |
| allcell | LILRB21 | 1.85E-15 | 1.002964718 | 0.263 | 0.177 | 3.65E-11 | Lep-T |
| allcell | PLAUR | 2.25E-43 | 0.985188692 | 0.791 | 0.732 | 4.43E-39 | Lep-T |
| allcell | CSRNP11 | 2.73E-70 | 0.980218148 | 0.724 | 0.537 | 5.37E-66 | Lep-T |
| allcell | IL1RN | 6.93E-15 | 0.97838242 | 0.386 | 0.3 | 1.36E-10 | Lep-T |
| allcell | IER31 | 6.68E-12 | 0.977501768 | 0.682 | 0.644 | 1.31E-07 | Lep-T |
| allcell | G0S21 | 8.48E-28 | 0.964287501 | 0.656 | 0.535 | 1.67E-23 | Lep-T |
| allcell | CXCL21 | 5.49E-09 | 0.918037601 | 0.372 | 0.3 | 0.00010802 | Lep-T |
| allcell | CFP1 | 8.89E-24 | 0.898784585 | 0.38 | 0.256 | 1.75E-19 | Lep-T |
| allcell | HIST1H1E | 1.23E-12 | 0.897358368 | 0.315 | 0.233 | 2.43E-08 | Lep-T |
| allcell | CST3 | 3.84E-63 | 0.89411063 | 0.98 | 0.964 | 7.55E-59 | Lep-T |
| allcell | INSIG11 | 5.71E-32 | 0.892317554 | 0.712 | 0.616 | 1.12E-27 | Lep-T |
| allcell | ALAS1 | 1.11E-12 | 0.89035472 | 0.32 | 0.239 | 2.18E-08 | Lep-T |
| allcell | MB21D2 | 2.34E-18 | 0.889651538 | 0.384 | 0.283 | 4.61E-14 | Lep-T |
| allcell | NINJ11 | 6.72E-27 | 0.889645893 | 0.661 | 0.557 | 1.32E-22 | Lep-T |
| allcell | MMP91 | 2.49E-26 | 0.881482107 | 0.366 | 0.237 | 4.90E-22 | Lep-T |
| allcell | ATP5PB | 2.93E-26 | 0.863208749 | 0.561 | 0.46 | 5.76E-22 | Lep-T |
| allcell | LMNA1 | 4.74E-67 | 0.842295879 | 0.816 | 0.647 | 9.33E-63 | Lep-T |
| allcell | MT2A | 4.78E-14 | 0.839565925 | 0.625 | 0.559 | 9.41E-10 | Lep-T |
| allcell | LPAR6 | 3.91E-09 | 0.83922324 | 0.272 | 0.207 | 7.69E-05 | Lep-T |
| allcell | AVPI11 | 7.13E-14 | 0.826751332 | 0.378 | 0.296 | 1.40E-09 | Lep-T |
| allcell | SERPING1 | 1.56E-10 | 0.822610607 | 0.326 | 0.256 | 3.08E-06 | Lep-T |
| allcell | IFI27 | 1.79E-06 | 0.820809685 | 0.335 | 0.285 | 0.035185214 | Lep-T |
| allcell | DUSP1 | 2.41E-37 | 0.806113902 | 0.946 | 0.908 | 4.75E-33 | Lep-T |
| allcell | OTUB1 | 2.77E-12 | 0.79839109 | 0.4 | 0.326 | 5.46E-08 | Lep-T |
| allcell | CD1C | 3.28E-24 | 0.796702831 | 0.401 | 0.269 | 6.46E-20 | Lep-T |
| allcell | NUDT21 | 1.50E-11 | 0.786199788 | 0.274 | 0.199 | 2.96E-07 | Lep-T |
| allcell | TNFAIP31 | 2.56E-55 | 0.78003931 | 0.837 | 0.734 | 5.05E-51 | Lep-T |
| allcell | TIAM1 | 3.71E-08 | 0.773002244 | 0.363 | 0.301 | 0.000730958 | Lep-T |
| allcell | SLC2A31 | 9.22E-28 | 0.767494239 | 0.67 | 0.579 | 1.82E-23 | Lep-T |
| allcell | CD1E | 9.59E-26 | 0.759904517 | 0.362 | 0.226 | 1.89E-21 | Lep-T |
| allcell | SYNGR2 | 8.85E-34 | 0.753965513 | 0.783 | 0.715 | 1.74E-29 | Lep-T |
| allcell | ZFP36L11 | 1.62E-63 | 0.750487685 | 0.832 | 0.7 | 3.19E-59 | Lep-T |

| | | | | | | | |
|---|---|---|---|---|---|---|---|
| allcell | CDC42EP31 | 3.33E-41 | 0.741088205 | 0.604 | 0.437 | 6.55E-37 | Lep-T |
| allcell | PSAP | 9.04E-67 | 0.738397166 | 0.877 | 0.804 | 1.78E-62 | Lep-T |
| allcell | GBP5 | 5.75E-16 | 0.72854351 | 0.365 | 0.27 | 1.13E-11 | Lep-T |
| allcell | MAP2K31 | 1.74E-38 | 0.722301995 | 0.727 | 0.616 | 3.42E-34 | Lep-T |
| allcell | P2RX7 | 1.36E-11 | 0.717363469 | 0.355 | 0.27 | 2.67E-07 | Lep-T |
| allcell | TPRA11 | 1.81E-11 | 0.713996831 | 0.379 | 0.298 | 3.56E-07 | Lep-T |
| allcell | GRINA | 3.12E-15 | 0.707592643 | 0.511 | 0.427 | 6.14E-11 | Lep-T |
| allcell | IMPDH2 | 2.21E-16 | 0.703458808 | 0.357 | 0.26 | 4.36E-12 | Lep-T |
| allcell | GNB2 | 2.53E-30 | 0.699901782 | 0.714 | 0.636 | 4.99E-26 | Lep-T |
| allcell | GEM1 | 2.89E-09 | 0.699481052 | 0.43 | 0.367 | 5.69E-05 | Lep-T |
| allcell | TIMP11 | 2.67E-29 | 0.698801474 | 0.738 | 0.618 | 5.26E-25 | Lep-T |
| allcell | PLEKHA7 | 8.94E-13 | 0.697189374 | 0.302 | 0.219 | 1.76E-08 | Lep-T |
| allcell | TSPAN331 | 6.29E-26 | 0.697084258 | 0.67 | 0.565 | 1.24E-21 | Lep-T |
| allcell | ERGIC3 | 1.92E-15 | 0.688988072 | 0.443 | 0.353 | 3.78E-11 | Lep-T |
| allcell | SESTD11 | 5.62E-08 | 0.679544023 | 0.364 | 0.3 | 0.001106607 | Lep-T |
| allcell | SAT11 | 3.98E-39 | 0.678837369 | 0.981 | 0.973 | 7.84E-35 | Lep-T |
| allcell | CS | 1.68E-09 | 0.676248906 | 0.318 | 0.248 | 3.30E-05 | Lep-T |
| allcell | GYPC | 8.71E-15 | 0.664389699 | 0.474 | 0.377 | 1.71E-10 | Lep-T |
| allcell | GRASP | 2.23E-19 | 0.659716466 | 0.674 | 0.63 | 4.39E-15 | Lep-T |
| allcell | AHNAK1 | 2.23E-42 | 0.658853483 | 0.767 | 0.657 | 4.38E-38 | Lep-T |
| allcell | VEGFA1 | 1.60E-13 | 0.653599522 | 0.43 | 0.35 | 3.15E-09 | Lep-T |
| allcell | BAG31 | 2.01E-13 | 0.652414527 | 0.626 | 0.537 | 3.95E-09 | Lep-T |
| allcell | MGLL | 1.43E-20 | 0.651630444 | 0.388 | 0.263 | 2.82E-16 | Lep-T |
| allcell | DDX3Y1 | 1.59E-17 | 0.648795989 | 0.365 | 0.257 | 3.13E-13 | Lep-T |
| allcell | ZYX | 4.24E-22 | 0.648232289 | 0.666 | 0.579 | 8.35E-18 | Lep-T |
| allcell | CYP2S1 | 1.35E-15 | 0.647804683 | 0.387 | 0.29 | 2.66E-11 | Lep-T |
| allcell | EEF21 | 3.52E-74 | 0.645259758 | 0.906 | 0.868 | 6.93E-70 | Lep-T |
| allcell | UBALD21 | 3.16E-08 | 0.639961919 | 0.499 | 0.448 | 0.0006222 | Lep-T |
| allcell | RASSF4 | 2.95E-23 | 0.637107569 | 0.577 | 0.462 | 5.81E-19 | Lep-T |
| allcell | TRAF1 | 1.16E-29 | 0.634991145 | 0.713 | 0.594 | 2.28E-25 | Lep-T |
| allcell | SLAMF8 | 4.26E-25 | 0.634610361 | 0.561 | 0.434 | 8.38E-21 | Lep-T |
| allcell | SQSTM11 | 4.86E-63 | 0.631513521 | 0.917 | 0.849 | 9.57E-59 | Lep-T |
| allcell | BCL2A11 | 7.06E-30 | 0.628182317 | 0.842 | 0.762 | 1.39E-25 | Lep-T |
| allcell | PPP2R1A | 3.44E-10 | 0.625422588 | 0.354 | 0.286 | 6.78E-06 | Lep-T |
| allcell | HACD4 | 6.65E-08 | 0.620530927 | 0.33 | 0.27 | 0.001308162 | Lep-T |
| allcell | CRTAP1 | 6.85E-12 | 0.616333679 | 0.436 | 0.366 | 1.35E-07 | Lep-T |
| allcell | BIRC3 | 1.31E-31 | 0.613973978 | 0.765 | 0.638 | 2.58E-27 | Lep-T |
| allcell | TACSTD2 | 1.84E-07 | 0.605451875 | 0.256 | 0.196 | 0.003626784 | Lep-T |
| allcell | BHLHE40 | 2.27E-25 | 0.604591153 | 0.786 | 0.737 | 4.46E-21 | Lep-T |
| allcell | NFKBIE1 | 2.94E-15 | 0.599777365 | 0.572 | 0.497 | 5.80E-11 | Lep-T |
| allcell | PIM3 | 3.70E-17 | 0.597727871 | 0.746 | 0.723 | 7.29E-13 | Lep-T |
| allcell | FCGR2A1 | 1.98E-10 | 0.59771746 | 0.467 | 0.388 | 3.89E-06 | Lep-T |
| allcell | ZC3H12A1 | 9.19E-25 | 0.59390606 | 0.569 | 0.442 | 1.81E-20 | Lep-T |
| allcell | ACSL5 | 7.72E-11 | 0.59186145 | 0.369 | 0.292 | 1.52E-06 | Lep-T |
| allcell | SEC13 | 8.67E-15 | 0.584462292 | 0.434 | 0.342 | 1.71E-10 | Lep-T |
| allcell | CD831 | 2.77E-72 | 0.577414997 | 0.979 | 0.957 | 5.45E-68 | Lep-T |
| allcell | DAZAP2 | 2.34E-16 | 0.575794059 | 0.606 | 0.523 | 4.61E-12 | Lep-T |
| allcell | PLEK | 4.03E-36 | 0.572736628 | 0.909 | 0.881 | 7.93E-32 | Lep-T |
| allcell | NUDT3 | 5.92E-07 | 0.568961972 | 0.276 | 0.22 | 0.011645442 | Lep-T |
| allcell | CNBP | 1.34E-23 | 0.567627257 | 0.794 | 0.749 | 2.64E-19 | Lep-T |
| allcell | CAPN21 | 5.67E-18 | 0.566553492 | 0.509 | 0.408 | 1.12E-13 | Lep-T |
| allcell | RTN1 | 9.82E-17 | 0.566442436 | 0.422 | 0.313 | 1.93E-12 | Lep-T |
| allcell | DUSP2 | 4.71E-24 | 0.566310909 | 0.846 | 0.768 | 9.28E-20 | Lep-T |
| allcell | CD40 | 1.85E-22 | 0.566266959 | 0.663 | 0.568 | 3.64E-18 | Lep-T |
| allcell | CD52 | 3.59E-31 | 0.56526891 | 0.632 | 0.475 | 7.06E-27 | Lep-T |
| allcell | PPIF1 | 4.51E-23 | 0.56367698 | 0.701 | 0.604 | 8.88E-19 | Lep-T |

| | | | | | | | |
|---|---|---|---|---|---|---|---|
| allcell | AFDN | 1.72E-10 | 0.562247824 | 0.367 | 0.288 | 3.38E-06 | Lep-T |
| allcell | PPDPF | 2.01E-07 | 0.55982233 | 0.693 | 0.661 | 0.003957355 | Lep-T |
| allcell | CD441 | 3.73E-32 | 0.557873948 | 0.86 | 0.794 | 7.35E-28 | Lep-T |
| allcell | PPP1R15A1 | 1.32E-39 | 0.549867015 | 0.935 | 0.921 | 2.60E-35 | Lep-T |
| allcell | LGALS2 | 5.29E-45 | 0.548633953 | 0.826 | 0.67 | 1.04E-40 | Lep-T |
| allcell | PEA15 | 3.82E-20 | 0.548076407 | 0.663 | 0.578 | 7.52E-16 | Lep-T |
| allcell | MIR155HG | 1.30E-25 | 0.540117781 | 0.6 | 0.466 | 2.55E-21 | Lep-T |
| allcell | POLR2G | 8.43E-08 | 0.538452028 | 0.382 | 0.323 | 0.001658614 | Lep-T |
| allcell | UNC119 | 1.36E-07 | 0.530198939 | 0.514 | 0.472 | 0.00268569 | Lep-T |
| allcell | TRIM28 | 1.26E-06 | 0.528897129 | 0.437 | 0.388 | 0.024858581 | Lep-T |
| allcell | OXA1L | 2.40E-08 | 0.528375003 | 0.414 | 0.351 | 0.000472038 | Lep-T |
| allcell | TAGLN2 | 6.78E-31 | 0.52639414 | 0.913 | 0.89 | 1.34E-26 | Lep-T |
| allcell | HSPA61 | 6.97E-08 | 0.521975266 | 0.489 | 0.415 | 0.001372048 | Lep-T |
| allcell | RARA1 | 1.18E-10 | 0.519376289 | 0.448 | 0.373 | 2.33E-06 | Lep-T |
| allcell | ID2 | 2.76E-10 | 0.513315827 | 0.736 | 0.689 | 5.43E-06 | Lep-T |
| allcell | LY6E | 1.17E-07 | 0.513242123 | 0.524 | 0.472 | 0.00230632 | Lep-T |
| allcell | FTH11 | 2.55E-31 | 0.508200977 | 0.991 | 0.995 | 5.02E-27 | Lep-T |
| allcell | FMNL2 | 1.66E-12 | 0.507663051 | 0.64 | 0.554 | 3.27E-08 | Lep-T |
| allcell | RPN1 | 5.47E-07 | 0.504982477 | 0.415 | 0.363 | 0.010769802 | Lep-T |
| allcell | IRAK21 | 9.30E-13 | 0.502986601 | 0.501 | 0.412 | 1.83E-08 | Lep-T |
| allcell | GZMB | 2.58E-47 | 2.188393602 | 0.312 | 0.15 | 5.07E-43 | Mari |
| allcell | LRRC26 | 5.30E-54 | 2.143724057 | 0.26 | 0.093 | 1.04E-49 | Mari |
| allcell | PLAC8 | 6.28E-72 | 2.119300691 | 0.455 | 0.217 | 1.24E-67 | Mari |
| allcell | CLEC4C | 1.21E-56 | 2.033463385 | 0.298 | 0.115 | 2.39E-52 | Mari |
| allcell | SCT | 1.42E-46 | 1.998899251 | 0.28 | 0.117 | 2.80E-42 | Mari |
| allcell | JCHAIN | 1.37E-41 | 1.907962649 | 0.304 | 0.148 | 2.70E-37 | Mari |
| allcell | LAMP5 | 1.77E-46 | 1.885036237 | 0.25 | 0.094 | 3.48E-42 | Mari |
| allcell | CLIC3 | 1.67E-42 | 1.862493426 | 0.338 | 0.171 | 3.28E-38 | Mari |
| allcell | MZB1 | 3.99E-44 | 1.84444628 | 0.257 | 0.103 | 7.86E-40 | Mari |
| allcell | DERL3 | 3.95E-43 | 1.835817777 | 0.25 | 0.099 | 7.77E-39 | Mari |
| allcell | IRF7 | 5.92E-56 | 1.724083459 | 0.728 | 0.522 | 1.17E-51 | Mari |
| allcell | KRT161 | 4.03E-43 | 1.709669391 | 0.309 | 0.151 | 7.93E-39 | Mari |
| allcell | MS4A4A | 9.25E-40 | 1.67143548 | 0.294 | 0.14 | 1.82E-35 | Mari |
| allcell | CIB2 | 2.86E-36 | 1.662634115 | 0.281 | 0.135 | 5.63E-32 | Mari |
| allcell | HIGD1A | 2.70E-78 | 1.659402592 | 0.597 | 0.345 | 5.31E-74 | Mari |
| allcell | LILRA4 | 2.31E-40 | 1.640300834 | 0.303 | 0.145 | 4.55E-36 | Mari |
| allcell | SELL | 6.18E-70 | 1.640129888 | 0.462 | 0.222 | 1.22E-65 | Mari |
| allcell | MGAT4A | 2.93E-37 | 1.569051669 | 0.305 | 0.153 | 5.77E-33 | Mari |
| allcell | ANK3 | 8.46E-39 | 1.531952887 | 0.301 | 0.145 | 1.67E-34 | Mari |
| allcell | GAS6 | 7.47E-119 | 1.531311003 | 0.765 | 0.475 | 1.47E-114 | Mari |
| allcell | ITM2C | 1.45E-28 | 1.52714356 | 0.431 | 0.296 | 2.86E-24 | Mari |
| allcell | PPP1R14B | 1.95E-44 | 1.525681964 | 0.825 | 0.727 | 3.84E-40 | Mari |
| allcell | SNRPN | 6.40E-35 | 1.479553397 | 0.408 | 0.252 | 1.26E-30 | Mari |
| allcell | ST6GALNAC4 | 1.40E-31 | 1.477194763 | 0.299 | 0.16 | 2.76E-27 | Mari |
| allcell | PCED1B-AS1 | 8.03E-34 | 1.475955022 | 0.299 | 0.154 | 1.58E-29 | Mari |
| allcell | CD36 | 4.92E-12 | 1.472966077 | 0.259 | 0.178 | 9.69E-08 | Mari |
| allcell | SDHAF3 | 5.57E-45 | 1.470816536 | 0.358 | 0.18 | 1.10E-40 | Mari |
| allcell | PPP1R14A1 | 1.75E-33 | 1.433830335 | 0.308 | 0.158 | 3.45E-29 | Mari |
| allcell | CCDC50 | 3.99E-39 | 1.410790039 | 0.615 | 0.464 | 7.86E-35 | Mari |
| allcell | ISCA2 | 2.64E-39 | 1.398708754 | 0.339 | 0.182 | 5.19E-35 | Mari |
| allcell | HLA-DRB51 | 2.95E-37 | 1.397952438 | 0.761 | 0.729 | 5.81E-33 | Mari |
| allcell | LIME1 | 4.46E-32 | 1.397395668 | 0.256 | 0.121 | 8.78E-28 | Mari |
| allcell | MSRB2 | 4.04E-49 | 1.39063268 | 0.37 | 0.184 | 7.96E-45 | Mari |
| allcell | PLAAT3 | 3.77E-27 | 1.375370248 | 0.356 | 0.224 | 7.43E-23 | Mari |
| allcell | PUS10 | 5.27E-08 | 1.360985577 | 0.323 | 0.262 | 0.001037835 | Mari |
| allcell | S100A81 | 9.20E-44 | 1.355529685 | 0.497 | 0.312 | 1.81E-39 | Mari |

| | | | | | | | |
|---|---|---|---|---|---|---|---|
| allcell | NUDT22 | 3.12E-57 | 1.350529558 | 0.533 | 0.328 | 6.13E-53 | Mari |
| allcell | S100A91 | 1.19E-32 | 1.337608325 | 0.535 | 0.376 | 2.35E-28 | Mari |
| allcell | SUSD1 | 1.62E-23 | 1.330992761 | 0.37 | 0.247 | 3.18E-19 | Mari |
| allcell | COBLL1 | 6.58E-41 | 1.317248376 | 0.278 | 0.12 | 1.29E-36 | Mari |
| allcell | AOPEP1 | 3.04E-13 | 1.30911337 | 0.258 | 0.175 | 5.98E-09 | Mari |
| allcell | VPS25 | 2.95E-43 | 1.302787371 | 0.359 | 0.185 | 5.81E-39 | Mari |
| allcell | ODC1 | 2.95E-31 | 1.297082036 | 0.549 | 0.41 | 5.81E-27 | Mari |
| allcell | POLE4 | 2.39E-59 | 1.293301003 | 0.516 | 0.308 | 4.71E-55 | Mari |
| allcell | OCIAD2 | 2.10E-24 | 1.268361787 | 0.264 | 0.145 | 4.13E-20 | Mari |
| allcell | TSPAN131 | 2.31E-39 | 1.26320421 | 0.467 | 0.291 | 4.54E-35 | Mari |
| allcell | GNG10 | 2.90E-82 | 1.244900872 | 0.713 | 0.511 | 5.70E-78 | Mari |
| allcell | ALOX5AP | 9.71E-14 | 1.236519301 | 0.436 | 0.371 | 1.91E-09 | Mari |
| allcell | PTPMT1 | 4.21E-38 | 1.235565915 | 0.42 | 0.257 | 8.30E-34 | Mari |
| allcell | ZBTB18 | 7.27E-14 | 1.234933767 | 0.29 | 0.205 | 1.43E-09 | Mari |
| allcell | RWDD3 | 4.04E-24 | 1.226240258 | 0.259 | 0.144 | 7.96E-20 | Mari |
| allcell | CD69 | 1.40E-40 | 1.220850736 | 0.574 | 0.404 | 2.76E-36 | Mari |
| allcell | CCDC186 | 4.42E-26 | 1.219180964 | 0.426 | 0.292 | 8.71E-22 | Mari |
| allcell | ITGAE | 1.93E-34 | 1.218122572 | 0.412 | 0.254 | 3.81E-30 | Mari |
| allcell | HINT2 | 3.68E-41 | 1.211419097 | 0.462 | 0.292 | 7.25E-37 | Mari |
| allcell | CCL4L2 | 2.87E-07 | 1.196164417 | 0.511 | 0.468 | 0.005648909 | Mari |
| allcell | TATDN3 | 1.15E-28 | 1.19319453 | 0.322 | 0.188 | 2.27E-24 | Mari |
| allcell | VEGFB | 6.17E-41 | 1.174580503 | 0.651 | 0.481 | 1.21E-36 | Mari |
| allcell | RASGRP2 | 1.95E-26 | 1.167709075 | 0.321 | 0.19 | 3.84E-22 | Mari |
| allcell | DCPS | 2.29E-30 | 1.151308942 | 0.373 | 0.229 | 4.51E-26 | Mari |
| allcell | SERPINF1 | 9.57E-75 | 1.146067411 | 0.851 | 0.684 | 1.88E-70 | Mari |
| allcell | MRPL58 | 1.78E-30 | 1.145659553 | 0.312 | 0.171 | 3.50E-26 | Mari |
| allcell | MTLN | 9.34E-26 | 1.129288617 | 0.327 | 0.201 | 1.84E-21 | Mari |
| allcell | NAE1 | 1.89E-20 | 1.118190452 | 0.253 | 0.148 | 3.73E-16 | Mari |
| allcell | HIST1H1C | 2.30E-10 | 1.115959098 | 0.325 | 0.251 | 4.54E-06 | Mari |
| allcell | RNASEH2C | 2.01E-50 | 1.114591948 | 0.533 | 0.336 | 3.96E-46 | Mari |
| allcell | SPCS1 | 2.61E-68 | 1.111755035 | 0.861 | 0.721 | 5.14E-64 | Mari |
| allcell | GCA | 2.50E-74 | 1.111116012 | 0.697 | 0.484 | 4.92E-70 | Mari |
| allcell | SVIP | 2.15E-19 | 1.106114708 | 0.274 | 0.169 | 4.23E-15 | Mari |
| allcell | ACTR6 | 1.89E-21 | 1.104383712 | 0.298 | 0.185 | 3.72E-17 | Mari |
| allcell | PLD4 | 4.42E-27 | 1.102544659 | 0.374 | 0.232 | 8.69E-23 | Mari |
| allcell | NSMCE3 | 1.29E-33 | 1.101193479 | 0.437 | 0.275 | 2.54E-29 | Mari |
| allcell | IDH3A | 1.22E-33 | 1.100798856 | 0.483 | 0.325 | 2.41E-29 | Mari |
| allcell | SEC61G | 2.49E-111 | 1.096225256 | 0.899 | 0.731 | 4.90E-107 | Mari |
| allcell | MEF2C | 8.18E-77 | 1.092499251 | 0.776 | 0.556 | 1.61E-72 | Mari |
| allcell | TMEM107 | 3.99E-14 | 1.091540468 | 0.321 | 0.23 | 7.85E-10 | Mari |
| allcell | NDUFC2 | 1.39E-113 | 1.088566111 | 0.851 | 0.605 | 2.73E-109 | Mari |
| allcell | COA5 | 6.84E-17 | 1.082204272 | 0.285 | 0.19 | 1.35E-12 | Mari |
| allcell | HIST1H1D | 3.85E-14 | 1.081803037 | 0.283 | 0.194 | 7.58E-10 | Mari |
| allcell | COMMD8 | 2.52E-46 | 1.077122161 | 0.478 | 0.289 | 4.96E-42 | Mari |
| allcell | MS4A6A | 3.64E-40 | 1.067417165 | 0.65 | 0.512 | 7.17E-36 | Mari |
| allcell | HRH2 | 5.13E-15 | 1.067313246 | 0.269 | 0.178 | 1.01E-10 | Mari |
| allcell | NDUFAF8 | 5.29E-44 | 1.061014435 | 0.501 | 0.32 | 1.04E-39 | Mari |
| allcell | CHCHD1 | 2.80E-40 | 1.053986302 | 0.471 | 0.303 | 5.51E-36 | Mari |
| allcell | CYBC1 | 7.98E-47 | 1.052742923 | 0.538 | 0.359 | 1.57E-42 | Mari |
| allcell | MX1 | 7.57E-33 | 1.050643665 | 0.482 | 0.318 | 1.49E-28 | Mari |
| allcell | METTL23 | 5.16E-36 | 1.047816628 | 0.449 | 0.285 | 1.02E-31 | Mari |
| allcell | ZNF511 | 4.37E-30 | 1.047001042 | 0.387 | 0.243 | 8.60E-26 | Mari |
| allcell | CNPY3 | 5.70E-44 | 1.046510735 | 0.656 | 0.487 | 1.12E-39 | Mari |
| allcell | SNRNP25 | 8.41E-50 | 1.044708567 | 0.597 | 0.405 | 1.65E-45 | Mari |
| allcell | HIST1H2AC | 2.77E-11 | 1.04454039 | 0.343 | 0.264 | 5.46E-07 | Mari |
| allcell | JKAMP | 3.83E-28 | 1.040067774 | 0.339 | 0.202 | 7.54E-24 | Mari |

| | | | | | | | |
|---|---|---|---|---|---|---|---|
| allcell | SRP9 | 8.09E-86 | 1.037861945 | 0.755 | 0.534 | 1.59E-81 | Mari |
| allcell | TMCO1 | 6.51E-46 | 1.035630321 | 0.574 | 0.388 | 1.28E-41 | Mari |
| allcell | POLR2I | 1.49E-42 | 1.03422983 | 0.516 | 0.338 | 2.93E-38 | Mari |
| allcell | CTSC | 2.48E-39 | 1.034224813 | 0.766 | 0.624 | 4.89E-35 | Mari |
| allcell | PAXX | 6.33E-51 | 1.033898075 | 0.736 | 0.562 | 1.25E-46 | Mari |
| allcell | LAIR1 | 5.41E-19 | 1.031773092 | 0.286 | 0.177 | 1.06E-14 | Mari |
| allcell | MYL12A | 2.77E-152 | 1.024034942 | 0.959 | 0.903 | 5.45E-148 | Mari |
| allcell | MRPS18C | 1.53E-36 | 1.022596578 | 0.462 | 0.296 | 3.01E-32 | Mari |
| allcell | RAC2 | 9.35E-25 | 1.021901403 | 0.47 | 0.34 | 1.84E-20 | Mari |
| allcell | FCER1G | 1.81E-61 | 1.021772643 | 0.835 | 0.726 | 3.56E-57 | Mari |
| allcell | SELENOS | 1.48E-34 | 1.021513325 | 0.663 | 0.527 | 2.92E-30 | Mari |
| allcell | FAM207A | 1.98E-18 | 1.018872362 | 0.301 | 0.196 | 3.89E-14 | Mari |
| allcell | SPCS3 | 3.27E-80 | 1.015434615 | 0.795 | 0.578 | 6.44E-76 | Mari |
| allcell | CYB561D2 | 3.53E-23 | 1.014898552 | 0.309 | 0.189 | 6.95E-19 | Mari |
| allcell | AGPAT5 | 1.78E-13 | 1.014776842 | 0.264 | 0.177 | 3.50E-09 | Mari |
| allcell | USF2 | 5.62E-24 | 1.012739173 | 0.473 | 0.353 | 1.11E-19 | Mari |
| allcell | TMEM141 | 2.26E-34 | 1.00739516 | 0.478 | 0.321 | 4.45E-30 | Mari |
| allcell | RHEX | 3.88E-27 | 1.006886746 | 0.427 | 0.282 | 7.63E-23 | Mari |
| allcell | FAM89B | 1.07E-50 | 1.006452656 | 0.607 | 0.414 | 2.10E-46 | Mari |
| allcell | MRPS23 | 1.97E-33 | 1.003947149 | 0.418 | 0.26 | 3.88E-29 | Mari |
| allcell | LYSMD2 | 9.11E-42 | 1.002809445 | 0.589 | 0.418 | 1.79E-37 | Mari |
| allcell | ZDHHC171 | 1.45E-25 | 1.000335562 | 0.441 | 0.307 | 2.84E-21 | Mari |
| allcell | PARK7 | 9.04E-126 | 0.997465634 | 0.931 | 0.809 | 1.78E-121 | Mari |
| allcell | BORCS8 | 4.30E-18 | 0.996378507 | 0.273 | 0.173 | 8.47E-14 | Mari |
| allcell | PAIP1 | 5.62E-24 | 0.993241369 | 0.39 | 0.264 | 1.11E-19 | Mari |
| allcell | RDH14 | 4.16E-20 | 0.992429186 | 0.252 | 0.146 | 8.18E-16 | Mari |
| allcell | PPP2R5C | 7.44E-30 | 0.991610179 | 0.642 | 0.496 | 1.46E-25 | Mari |
| allcell | NUCB21 | 2.03E-15 | 0.987877154 | 0.317 | 0.219 | 4.00E-11 | Mari |
| allcell | SOCS11 | 2.88E-41 | 0.982880007 | 0.646 | 0.465 | 5.67E-37 | Mari |
| allcell | CYB561A3 | 1.21E-17 | 0.98275532 | 0.381 | 0.279 | 2.38E-13 | Mari |
| allcell | 2-Mar | 4.96E-34 | 0.982566481 | 0.488 | 0.334 | 9.77E-30 | Mari |
| allcell | ARRDC3 | 8.62E-20 | 0.982083168 | 0.47 | 0.362 | 1.70E-15 | Mari |
| allcell | NUBP2 | 7.48E-27 | 0.977863624 | 0.386 | 0.253 | 1.47E-22 | Mari |
| allcell | CCS | 6.99E-25 | 0.976547819 | 0.398 | 0.267 | 1.38E-20 | Mari |
| allcell | UCHL3 | 6.33E-49 | 0.97610765 | 0.56 | 0.354 | 1.25E-44 | Mari |
| allcell | ZBTB8OS | 2.20E-30 | 0.971967847 | 0.423 | 0.275 | 4.32E-26 | Mari |
| allcell | ALG5 | 1.56E-37 | 0.971808615 | 0.474 | 0.306 | 3.08E-33 | Mari |
| allcell | TMX1 | 1.37E-34 | 0.971109862 | 0.491 | 0.336 | 2.69E-30 | Mari |
| allcell | UBLCP1 | 1.30E-17 | 0.969435094 | 0.322 | 0.219 | 2.55E-13 | Mari |
| allcell | RASD1 | 1.75E-14 | 0.968164618 | 0.262 | 0.171 | 3.44E-10 | Mari |
| allcell | TRAPPC1 | 1.23E-85 | 0.967681204 | 0.784 | 0.552 | 2.43E-81 | Mari |
| allcell | POP4 | 2.99E-23 | 0.967254035 | 0.343 | 0.22 | 5.88E-19 | Mari |
| allcell | PHB | 1.77E-48 | 0.966903313 | 0.698 | 0.525 | 3.48E-44 | Mari |
| allcell | ARL2 | 6.12E-32 | 0.963884119 | 0.431 | 0.283 | 1.21E-27 | Mari |
| allcell | TIMM8B | 3.43E-59 | 0.962654455 | 0.651 | 0.444 | 6.76E-55 | Mari |
| allcell | THYN1 | 7.88E-33 | 0.959326589 | 0.418 | 0.262 | 1.55E-28 | Mari |
| allcell | DTWD1 | 1.17E-17 | 0.956044832 | 0.276 | 0.174 | 2.30E-13 | Mari |
| allcell | RBX1 | 8.23E-120 | 0.953685681 | 0.893 | 0.723 | 1.62E-115 | Mari |
| allcell | GPR65 | 1.27E-37 | 0.953085449 | 0.623 | 0.441 | 2.49E-33 | Mari |
| allcell | SNRNP27 | 1.64E-13 | 0.952808403 | 0.27 | 0.188 | 3.22E-09 | Mari |
| allcell | GALK1 | 2.18E-23 | 0.951587461 | 0.38 | 0.256 | 4.29E-19 | Mari |
| allcell | COPZ1 | 1.72E-46 | 0.95116401 | 0.561 | 0.378 | 3.38E-42 | Mari |
| allcell | CZIB | 4.99E-20 | 0.948191392 | 0.318 | 0.207 | 9.82E-16 | Mari |
| allcell | PLP2 | 2.15E-30 | 0.94804525 | 0.663 | 0.54 | 4.24E-26 | Mari |
| allcell | MRPL20 | 7.44E-65 | 0.946512316 | 0.738 | 0.535 | 1.47E-60 | Mari |
| allcell | MRPL19 | 2.08E-18 | 0.945853035 | 0.32 | 0.216 | 4.09E-14 | Mari |

| | | | | | | | |
|---|---|---|---|---|---|---|---|
| allcell | TOMM5 | 2.80E-59 | 0.945563121 | 0.676 | 0.467 | 5.50E-55 | Mari |
| allcell | UNC50 | 4.84E-22 | 0.943942372 | 0.33 | 0.211 | 9.52E-18 | Mari |
| allcell | GTF3C6 | 3.09E-52 | 0.943119413 | 0.646 | 0.453 | 6.08E-48 | Mari |
| allcell | NDUFAB1 | 1.44E-81 | 0.942194091 | 0.773 | 0.546 | 2.83E-77 | Mari |
| allcell | TMEM59 | 4.14E-70 | 0.940056414 | 0.801 | 0.62 | 8.14E-66 | Mari |
| allcell | ERP29 | 1.29E-43 | 0.939004185 | 0.823 | 0.71 | 2.54E-39 | Mari |
| allcell | CYBA | 1.37E-148 | 0.938086769 | 0.975 | 0.939 | 2.71E-144 | Mari |
| allcell | RGS18 | 1.18E-25 | 0.936962825 | 0.376 | 0.241 | 2.33E-21 | Mari |
| allcell | PYURF | 1.42E-43 | 0.93366405 | 0.548 | 0.371 | 2.80E-39 | Mari |
| allcell | TXNDC17 | 3.06E-66 | 0.933463845 | 0.689 | 0.473 | 6.02E-62 | Mari |
| allcell | BST2 | 1.37E-94 | 0.933167643 | 0.868 | 0.713 | 2.70E-90 | Mari |
| allcell | YIF1A | 5.08E-28 | 0.931182504 | 0.44 | 0.3 | 1.00E-23 | Mari |
| allcell | CDKN2D | 1.63E-34 | 0.928599425 | 0.509 | 0.34 | 3.21E-30 | Mari |
| allcell | MRPL42 | 1.26E-13 | 0.927811804 | 0.252 | 0.167 | 2.47E-09 | Mari |
| allcell | PRDX5 | 6.32E-80 | 0.925555343 | 0.817 | 0.636 | 1.24E-75 | Mari |
| allcell | TMEM35B | 1.46E-19 | 0.92450378 | 0.347 | 0.237 | 2.87E-15 | Mari |
| allcell | UQCR11 | 1.18E-138 | 0.924064706 | 0.921 | 0.775 | 2.32E-134 | Mari |
| allcell | SLC39A10 | 2.79E-33 | 0.923306722 | 0.455 | 0.294 | 5.49E-29 | Mari |
| allcell | CIAO2A | 1.71E-53 | 0.922155838 | 0.663 | 0.462 | 3.36E-49 | Mari |
| allcell | NTAN1 | 7.33E-18 | 0.92132765 | 0.323 | 0.218 | 1.44E-13 | Mari |
| allcell | LSM5 | 3.53E-65 | 0.917268132 | 0.721 | 0.519 | 6.94E-61 | Mari |
| allcell | MRPS10 | 4.01E-19 | 0.914875571 | 0.36 | 0.252 | 7.88E-15 | Mari |
| allcell | SEC11C | 2.60E-42 | 0.914744118 | 0.631 | 0.449 | 5.11E-38 | Mari |
| allcell | NDUFA9 | 5.78E-29 | 0.914086643 | 0.483 | 0.344 | 1.14E-24 | Mari |
| allcell | VAMP8 | 1.05E-108 | 0.912134339 | 0.924 | 0.842 | 2.06E-104 | Mari |
| allcell | ADI1 | 5.41E-27 | 0.909731505 | 0.486 | 0.348 | 1.06E-22 | Mari |
| allcell | MRPL13 | 3.85E-35 | 0.909691359 | 0.467 | 0.309 | 7.57E-31 | Mari |
| allcell | TRIAP1 | 7.38E-16 | 0.908513167 | 0.273 | 0.177 | 1.45E-11 | Mari |
| allcell | CRELD2 | 3.35E-31 | 0.907508687 | 0.44 | 0.281 | 6.60E-27 | Mari |
| allcell | PRELID1 | 1.19E-61 | 0.907477714 | 0.865 | 0.741 | 2.34E-57 | Mari |
| allcell | MED30 | 5.90E-26 | 0.906851533 | 0.43 | 0.289 | 1.16E-21 | Mari |
| allcell | GTF2H5 | 9.71E-35 | 0.906321152 | 0.499 | 0.339 | 1.91E-30 | Mari |
| allcell | GSDMD | 1.60E-27 | 0.905985879 | 0.463 | 0.327 | 3.16E-23 | Mari |
| allcell | RPP30 | 9.37E-16 | 0.905913525 | 0.318 | 0.22 | 1.84E-11 | Mari |
| allcell | NHP2 | 2.94E-43 | 0.905077903 | 0.618 | 0.445 | 5.79E-39 | Mari |
| allcell | SRI | 8.13E-59 | 0.905000068 | 0.725 | 0.54 | 1.60E-54 | Mari |
| allcell | MRPL57 | 1.76E-50 | 0.904618888 | 0.65 | 0.453 | 3.47E-46 | Mari |
| allcell | RBM7 | 1.23E-07 | 0.90094222 | 0.279 | 0.222 | 0.002429471 | Mari |
| allcell | BLOC1S2 | 4.23E-39 | 0.900766958 | 0.567 | 0.399 | 8.33E-35 | Mari |
| allcell | ANAPC16 | 5.99E-47 | 0.900280462 | 0.702 | 0.525 | 1.18E-42 | Mari |
| allcell | CCND3 | 3.76E-30 | 0.899318866 | 0.544 | 0.391 | 7.41E-26 | Mari |
| allcell | LSM10 | 3.57E-35 | 0.897767436 | 0.525 | 0.369 | 7.02E-31 | Mari |
| allcell | SIVA1 | 4.79E-53 | 0.897365702 | 0.722 | 0.539 | 9.42E-49 | Mari |
| allcell | PIN1 | 4.06E-35 | 0.895443875 | 0.544 | 0.394 | 7.99E-31 | Mari |
| allcell | TMEM109 | 4.00E-31 | 0.894828102 | 0.498 | 0.341 | 7.87E-27 | Mari |
| allcell | GTF3A | 3.43E-51 | 0.894815985 | 0.723 | 0.558 | 6.76E-47 | Mari |
| allcell | COA4 | 5.68E-31 | 0.89420189 | 0.488 | 0.34 | 1.12E-26 | Mari |
| allcell | NDUFB7 | 1.02E-66 | 0.893632439 | 0.744 | 0.546 | 2.02E-62 | Mari |
| allcell | ZNF428 | 6.20E-16 | 0.8933471 | 0.425 | 0.324 | 1.22E-11 | Mari |
| allcell | NCBP2AS2 | 2.93E-21 | 0.892868141 | 0.347 | 0.229 | 5.76E-17 | Mari |
| allcell | POLR3K | 4.11E-15 | 0.892836759 | 0.286 | 0.192 | 8.10E-11 | Mari |
| allcell | MAGOHB | 1.51E-13 | 0.891434598 | 0.298 | 0.21 | 2.96E-09 | Mari |
| allcell | LAMTOR1 | 3.19E-74 | 0.890294867 | 0.807 | 0.647 | 6.28E-70 | Mari |
| allcell | GTF2A2 | 7.07E-49 | 0.889911854 | 0.669 | 0.497 | 1.39E-44 | Mari |
| allcell | SMIM7 | 9.43E-34 | 0.889493717 | 0.531 | 0.375 | 1.86E-29 | Mari |
| allcell | CCDC167 | 3.21E-21 | 0.88853483 | 0.331 | 0.211 | 6.32E-17 | Mari |

| | | | | | | | |
|---|---|---|---|---|---|---|---|
| allcell | NME4 | 2.90E-30 | 0.888173586 | 0.492 | 0.342 | 5.72E-26 | Mari |
| allcell | NSMCE4A | 5.32E-25 | 0.887494537 | 0.386 | 0.253 | 1.05E-20 | Mari |
| allcell | PPA1 | 7.52E-27 | 0.887342534 | 0.795 | 0.747 | 1.48E-22 | Mari |
| allcell | TMEM230 | 1.20E-50 | 0.887277743 | 0.673 | 0.492 | 2.36E-46 | Mari |
| allcell | NCF4 | 5.13E-24 | 0.886799108 | 0.407 | 0.271 | 1.01E-19 | Mari |
| allcell | DAD1 | 1.68E-72 | 0.886595223 | 0.798 | 0.61 | 3.31E-68 | Mari |
| allcell | TM2D1 | 1.44E-18 | 0.886245276 | 0.345 | 0.235 | 2.84E-14 | Mari |
| allcell | TYMP1 | 7.58E-59 | 0.882775947 | 0.887 | 0.813 | 1.49E-54 | Mari |
| allcell | SNF8 | 8.96E-53 | 0.880866291 | 0.677 | 0.484 | 1.76E-48 | Mari |
| allcell | MRPL40 | 6.26E-29 | 0.880306514 | 0.452 | 0.306 | 1.23E-24 | Mari |
| allcell | CCNDBP1 | 9.07E-28 | 0.880106816 | 0.451 | 0.311 | 1.78E-23 | Mari |
| allcell | VAMP5 | 4.06E-28 | 0.877134137 | 0.639 | 0.516 | 7.99E-24 | Mari |
| allcell | SNHG9 | 8.41E-27 | 0.876912052 | 0.344 | 0.206 | 1.65E-22 | Mari |
| allcell | SELENOT | 8.77E-54 | 0.876126464 | 0.719 | 0.528 | 1.73E-49 | Mari |
| allcell | TSTA3 | 3.72E-20 | 0.875155165 | 0.352 | 0.234 | 7.32E-16 | Mari |
| allcell | IER3IP1 | 1.25E-32 | 0.874442988 | 0.475 | 0.315 | 2.46E-28 | Mari |
| allcell | UROS | 1.28E-18 | 0.873033538 | 0.382 | 0.274 | 2.53E-14 | Mari |
| allcell | MIF4GD | 5.37E-33 | 0.872743828 | 0.566 | 0.414 | 1.06E-28 | Mari |
| allcell | SELENOH | 5.92E-82 | 0.870028291 | 0.879 | 0.704 | 1.16E-77 | Mari |
| allcell | VBP1 | 5.24E-32 | 0.869949705 | 0.482 | 0.324 | 1.03E-27 | Mari |
| allcell | LSM6 | 2.62E-46 | 0.869559419 | 0.644 | 0.468 | 5.16E-42 | Mari |
| allcell | IER2 | 7.04E-57 | 0.867726801 | 0.904 | 0.817 | 1.39E-52 | Mari |
| allcell | BOLA2 | 5.81E-28 | 0.867443582 | 0.445 | 0.295 | 1.14E-23 | Mari |
| allcell | NDUFB2 | 1.60E-100 | 0.865989724 | 0.894 | 0.722 | 3.14E-96 | Mari |
| allcell | TXNDC9 | 1.51E-20 | 0.865890911 | 0.374 | 0.257 | 2.97E-16 | Mari |
| allcell | UQCC2 | 1.46E-28 | 0.865259228 | 0.428 | 0.279 | 2.87E-24 | Mari |
| allcell | MRPL28 | 1.16E-29 | 0.864790781 | 0.446 | 0.299 | 2.28E-25 | Mari |
| allcell | CUTC | 2.28E-18 | 0.862406293 | 0.291 | 0.184 | 4.48E-14 | Mari |
| allcell | EEF1E1 | 1.57E-23 | 0.860792668 | 0.376 | 0.246 | 3.08E-19 | Mari |
| allcell | MIEN1 | 9.81E-27 | 0.860491545 | 0.422 | 0.28 | 1.93E-22 | Mari |
| allcell | DGUOK | 2.97E-50 | 0.85980842 | 0.679 | 0.494 | 5.85E-46 | Mari |
| allcell | EIF4EBP1 | 3.53E-30 | 0.857406346 | 0.533 | 0.398 | 6.95E-26 | Mari |
| allcell | UGCG | 1.58E-22 | 0.855934316 | 0.687 | 0.594 | 3.11E-18 | Mari |
| allcell | PLGRKT | 2.27E-26 | 0.855926334 | 0.481 | 0.342 | 4.47E-22 | Mari |
| allcell | LLPH | 5.42E-21 | 0.85553046 | 0.4 | 0.283 | 1.07E-16 | Mari |
| allcell | POP5 | 5.81E-14 | 0.854053029 | 0.29 | 0.199 | 1.14E-09 | Mari |
| allcell | SSU72 | 8.13E-39 | 0.85359077 | 0.627 | 0.48 | 1.60E-34 | Mari |
| allcell | PDHB | 4.91E-19 | 0.851802382 | 0.352 | 0.239 | 9.67E-15 | Mari |
| allcell | NUDT16L1 | 6.95E-12 | 0.851217412 | 0.254 | 0.175 | 1.37E-07 | Mari |
| allcell | IAH1 | 1.15E-40 | 0.85090354 | 0.62 | 0.444 | 2.26E-36 | Mari |
| allcell | SFT2D1 | 7.03E-57 | 0.84857464 | 0.732 | 0.531 | 1.38E-52 | Mari |
| allcell | SRBD11 | 5.62E-12 | 0.847831275 | 0.26 | 0.178 | 1.11E-07 | Mari |
| allcell | TMEM167A | 7.88E-47 | 0.8471868 | 0.651 | 0.474 | 1.55E-42 | Mari |
| allcell | SPCS2 | 5.19E-67 | 0.844630858 | 0.817 | 0.634 | 1.02E-62 | Mari |
| allcell | NOL12 | 7.33E-11 | 0.843325275 | 0.255 | 0.181 | 1.44E-06 | Mari |
| allcell | NUDT1 | 9.98E-47 | 0.843121212 | 0.615 | 0.417 | 1.96E-42 | Mari |
| allcell | ANTKMT | 3.64E-25 | 0.842660997 | 0.447 | 0.314 | 7.17E-21 | Mari |
| allcell | MRPL36 | 1.62E-20 | 0.841378274 | 0.405 | 0.281 | 3.19E-16 | Mari |
| allcell | SNRNP40 | 1.19E-21 | 0.840972149 | 0.406 | 0.285 | 2.33E-17 | Mari |
| allcell | ISCU | 1.77E-33 | 0.840370622 | 0.669 | 0.527 | 3.49E-29 | Mari |
| allcell | POLR2K | 2.03E-37 | 0.839493577 | 0.569 | 0.402 | 4.00E-33 | Mari |
| allcell | RER1 | 2.89E-54 | 0.83644973 | 0.715 | 0.533 | 5.68E-50 | Mari |
| allcell | C1D | 6.54E-36 | 0.835511762 | 0.519 | 0.358 | 1.29E-31 | Mari |
| allcell | MCTS1 | 7.28E-25 | 0.834330097 | 0.457 | 0.325 | 1.43E-20 | Mari |
| allcell | PEX2 | 3.98E-17 | 0.833969022 | 0.293 | 0.192 | 7.84E-13 | Mari |
| allcell | ETFB | 5.27E-43 | 0.833756172 | 0.609 | 0.431 | 1.04E-38 | Mari |

| | | | | | | | |
|---|---|---|---|---|---|---|---|
| allcell | EBNA1BP2 | 2.73E-15 | 0.832684409 | 0.307 | 0.21 | 5.37E-11 | Mari |
| allcell | PEX16 | 5.89E-17 | 0.832616685 | 0.324 | 0.221 | 1.16E-12 | Mari |
| allcell | RBKS | 7.28E-34 | 0.832268247 | 0.496 | 0.333 | 1.43E-29 | Mari |
| allcell | KRT141 | 4.48E-07 | 0.830694607 | 0.367 | 0.34 | 0.008820823 | Mari |
| allcell | TMEM208 | 2.28E-39 | 0.828861957 | 0.589 | 0.412 | 4.49E-35 | Mari |
| allcell | REXO2 | 5.85E-15 | 0.828294467 | 0.287 | 0.194 | 1.15E-10 | Mari |
| allcell | ZDHHC12 | 2.47E-21 | 0.828215215 | 0.386 | 0.267 | 4.86E-17 | Mari |
| allcell | HSPB111 | 3.62E-40 | 0.827566196 | 0.617 | 0.454 | 7.13E-36 | Mari |
| allcell | MVB12A | 1.05E-13 | 0.827347294 | 0.275 | 0.185 | 2.08E-09 | Mari |
| allcell | ORMDL2 | 2.51E-15 | 0.82646427 | 0.354 | 0.261 | 4.94E-11 | Mari |
| allcell | SMIM19 | 1.02E-28 | 0.826376534 | 0.488 | 0.342 | 2.01E-24 | Mari |
| allcell | TMBIM4 | 4.35E-62 | 0.826258549 | 0.786 | 0.593 | 8.55E-58 | Mari |
| allcell | GPN3 | 9.78E-18 | 0.825945208 | 0.349 | 0.241 | 1.92E-13 | Mari |
| allcell | STOML2 | 4.13E-33 | 0.825700084 | 0.511 | 0.351 | 8.12E-29 | Mari |
| allcell | SS18L2 | 1.66E-30 | 0.824309628 | 0.46 | 0.305 | 3.27E-26 | Mari |
| allcell | DCXR | 1.90E-44 | 0.823750915 | 0.622 | 0.441 | 3.73E-40 | Mari |
| allcell | DERA | 4.71E-22 | 0.823722266 | 0.385 | 0.257 | 9.26E-18 | Mari |
| allcell | HMGN1 | 4.76E-93 | 0.823254428 | 0.916 | 0.783 | 9.37E-89 | Mari |
| allcell | STMN1 | 1.53E-10 | 0.822890407 | 0.445 | 0.381 | 3.02E-06 | Mari |
| allcell | JTB | 1.59E-42 | 0.821570728 | 0.68 | 0.525 | 3.13E-38 | Mari |
| allcell | TIMM13 | 8.28E-38 | 0.8214492 | 0.662 | 0.504 | 1.63E-33 | Mari |
| allcell | DCK | 5.58E-20 | 0.820188714 | 0.445 | 0.327 | 1.10E-15 | Mari |
| allcell | OSTC | 3.09E-66 | 0.818767183 | 0.838 | 0.689 | 6.09E-62 | Mari |
| allcell | SLIRP | 1.22E-44 | 0.817530972 | 0.616 | 0.432 | 2.40E-40 | Mari |
| allcell | CHMP2A | 2.37E-55 | 0.817458356 | 0.717 | 0.527 | 4.67E-51 | Mari |
| allcell | METTL5 | 1.86E-24 | 0.816490101 | 0.429 | 0.298 | 3.66E-20 | Mari |
| allcell | SEC61B | 4.30E-79 | 0.815435162 | 0.945 | 0.884 | 8.46E-75 | Mari |
| allcell | FOS | 6.77E-81 | 0.814921435 | 0.924 | 0.813 | 1.33E-76 | Mari |
| allcell | MRPS12 | 3.14E-28 | 0.814306612 | 0.49 | 0.345 | 6.18E-24 | Mari |
| allcell | BLOC1S1 | 1.05E-76 | 0.813811915 | 0.844 | 0.691 | 2.06E-72 | Mari |
| allcell | IFI30 | 1.55E-10 | 0.811607986 | 0.779 | 0.828 | 3.06E-06 | Mari |
| allcell | PSENEN | 6.63E-40 | 0.811363015 | 0.639 | 0.468 | 1.30E-35 | Mari |
| allcell | AP1S1 | 1.58E-10 | 0.810406688 | 0.267 | 0.196 | 3.11E-06 | Mari |
| allcell | RPA3 | 4.10E-29 | 0.810215484 | 0.5 | 0.344 | 8.06E-25 | Mari |
| allcell | H3F3B | 2.12E-94 | 0.809817499 | 0.992 | 0.981 | 4.17E-90 | Mari |
| allcell | COMMD3 | 4.55E-28 | 0.808731607 | 0.509 | 0.375 | 8.96E-24 | Mari |
| allcell | RABAC1 | 2.61E-55 | 0.808053802 | 0.751 | 0.562 | 5.14E-51 | Mari |
| allcell | CYBB | 5.54E-21 | 0.808015556 | 0.562 | 0.457 | 1.09E-16 | Mari |
| allcell | TMED9 | 2.29E-40 | 0.804145565 | 0.662 | 0.499 | 4.50E-36 | Mari |
| allcell | ACP1 | 1.45E-39 | 0.803433416 | 0.611 | 0.443 | 2.86E-35 | Mari |
| allcell | FDPS | 9.88E-27 | 0.803367689 | 0.494 | 0.353 | 1.94E-22 | Mari |
| allcell | SLC35C2 | 3.78E-14 | 0.802678278 | 0.325 | 0.234 | 7.45E-10 | Mari |
| allcell | ZSWIM7 | 1.71E-23 | 0.801370297 | 0.461 | 0.336 | 3.37E-19 | Mari |
| allcell | TRABD | 3.02E-33 | 0.80103466 | 0.613 | 0.457 | 5.94E-29 | Mari |
| allcell | ANAPC15 | 3.54E-21 | 0.800849774 | 0.389 | 0.263 | 6.98E-17 | Mari |
| allcell | SRSF9 | 3.10E-78 | 0.800278552 | 0.878 | 0.734 | 6.11E-74 | Mari |
| allcell | UBE2V2 | 4.76E-30 | 0.799497079 | 0.467 | 0.313 | 9.37E-26 | Mari |
| allcell | PALD11 | 2.25E-12 | 0.799483623 | 0.303 | 0.217 | 4.43E-08 | Mari |
| allcell | IMPA2 | 5.59E-13 | 0.798643049 | 0.267 | 0.182 | 1.10E-08 | Mari |
| allcell | NDUFA4 | 7.58E-113 | 0.797624392 | 0.932 | 0.843 | 1.49E-108 | Mari |
| allcell | RNASET2 | 3.26E-34 | 0.797607412 | 0.821 | 0.758 | 6.42E-30 | Mari |
| allcell | JOSD2 | 1.28E-23 | 0.797570107 | 0.438 | 0.308 | 2.52E-19 | Mari |
| allcell | CNPY2 | 4.66E-30 | 0.796044809 | 0.505 | 0.361 | 9.17E-26 | Mari |
| allcell | CHMP5 | 5.92E-33 | 0.795621156 | 0.621 | 0.484 | 1.16E-28 | Mari |
| allcell | SDF2L1 | 1.22E-25 | 0.794543408 | 0.56 | 0.432 | 2.41E-21 | Mari |
| allcell | SMCO4 | 1.61E-46 | 0.792795812 | 0.719 | 0.554 | 3.18E-42 | Mari |

| | | | | | | | |
|---|---|---|---|---|---|---|---|
| allcell | SMIM20 | 2.96E-17 | 0.792205395 | 0.436 | 0.327 | 5.83E-13 | Mari |
| allcell | PAFAH1B3 | 1.41E-14 | 0.791013629 | 0.331 | 0.234 | 2.78E-10 | Mari |
| allcell | NDUFS6 | 2.86E-70 | 0.790951629 | 0.826 | 0.638 | 5.63E-66 | Mari |
| allcell | TUSC2 | 4.37E-12 | 0.790624526 | 0.296 | 0.212 | 8.61E-08 | Mari |
| allcell | SCAMP2 | 1.54E-24 | 0.789778482 | 0.505 | 0.38 | 3.02E-20 | Mari |
| allcell | AIF1 | 6.40E-10 | 0.788870701 | 0.734 | 0.746 | 1.26E-05 | Mari |
| allcell | DHRS7 | 4.53E-39 | 0.788443754 | 0.657 | 0.486 | 8.92E-35 | Mari |
| allcell | THOC7 | 1.50E-39 | 0.788092976 | 0.604 | 0.441 | 2.95E-35 | Mari |
| allcell | ATP5MF | 4.48E-99 | 0.786903834 | 0.907 | 0.766 | 8.83E-95 | Mari |
| allcell | PET100 | 2.44E-56 | 0.786884013 | 0.746 | 0.568 | 4.80E-52 | Mari |
| allcell | DNAJC19 | 6.44E-24 | 0.786534531 | 0.437 | 0.303 | 1.27E-19 | Mari |
| allcell | PRKACB | 2.00E-12 | 0.785885334 | 0.317 | 0.229 | 3.94E-08 | Mari |
| allcell | LDHB | 8.00E-46 | 0.785791156 | 0.776 | 0.61 | 1.58E-41 | Mari |
| allcell | SNRPG | 2.13E-86 | 0.78455884 | 0.878 | 0.72 | 4.19E-82 | Mari |
| allcell | LSM4 | 1.13E-48 | 0.784438106 | 0.692 | 0.509 | 2.22E-44 | Mari |
| allcell | MRPL51 | 4.33E-53 | 0.783661753 | 0.727 | 0.552 | 8.53E-49 | Mari |
| allcell | PPM1K1 | 1.28E-17 | 0.783567687 | 0.445 | 0.337 | 2.51E-13 | Mari |
| allcell | HMGN3 | 2.48E-63 | 0.78355646 | 0.845 | 0.685 | 4.89E-59 | Mari |
| allcell | NAA20 | 1.48E-26 | 0.783241904 | 0.511 | 0.371 | 2.92E-22 | Mari |
| allcell | CHMP4A | 3.01E-43 | 0.780862825 | 0.664 | 0.492 | 5.93E-39 | Mari |
| allcell | LSM8 | 5.33E-58 | 0.780651507 | 0.753 | 0.555 | 1.05E-53 | Mari |
| allcell | SRM | 3.20E-26 | 0.78038955 | 0.478 | 0.338 | 6.30E-22 | Mari |
| allcell | TMEM126A | 1.55E-24 | 0.779449451 | 0.418 | 0.284 | 3.05E-20 | Mari |
| allcell | NIPSNAP3A | 3.51E-09 | 0.778744301 | 0.253 | 0.187 | 6.90E-05 | Mari |
| allcell | AKR7A2 | 2.04E-25 | 0.778502676 | 0.495 | 0.361 | 4.02E-21 | Mari |
| allcell | NDUFA3 | 4.06E-53 | 0.777890055 | 0.697 | 0.507 | 7.99E-49 | Mari |
| allcell | NDUFB3 | 2.09E-47 | 0.777809686 | 0.681 | 0.511 | 4.12E-43 | Mari |
| allcell | DBNL | 8.66E-45 | 0.777783776 | 0.711 | 0.536 | 1.70E-40 | Mari |
| allcell | TIMM17B | 1.31E-27 | 0.777225906 | 0.515 | 0.373 | 2.57E-23 | Mari |
| allcell | A1BG | 2.20E-16 | 0.777079616 | 0.456 | 0.357 | 4.32E-12 | Mari |
| allcell | NEDD8 | 3.06E-76 | 0.776930153 | 0.863 | 0.696 | 6.02E-72 | Mari |
| allcell | STMP1 | 1.80E-42 | 0.776420509 | 0.641 | 0.479 | 3.55E-38 | Mari |
| allcell | CIB1 | 3.69E-42 | 0.776306706 | 0.78 | 0.668 | 7.26E-38 | Mari |
| allcell | SRP14 | 1.18E-113 | 0.775862272 | 0.951 | 0.909 | 2.32E-109 | Mari |
| allcell | NADK | 8.16E-07 | 0.775488825 | 0.285 | 0.235 | 0.016056673 | Mari |
| allcell | LAMTOR5 | 1.62E-46 | 0.775401723 | 0.713 | 0.537 | 3.19E-42 | Mari |
| allcell | PPCS | 4.71E-27 | 0.774476512 | 0.492 | 0.352 | 9.28E-23 | Mari |
| allcell | PRDX2 | 1.11E-20 | 0.773857778 | 0.446 | 0.326 | 2.18E-16 | Mari |
| allcell | TRADD | 7.21E-21 | 0.773636835 | 0.483 | 0.369 | 1.42E-16 | Mari |
| allcell | TRAPPC2L | 2.69E-26 | 0.773628839 | 0.492 | 0.356 | 5.30E-22 | Mari |
| allcell | PSMA2 | 8.39E-75 | 0.773057755 | 0.867 | 0.737 | 1.65E-70 | Mari |
| allcell | LMAN2 | 2.50E-42 | 0.771524005 | 0.679 | 0.509 | 4.91E-38 | Mari |
| allcell | COPS9 | 2.19E-55 | 0.771122818 | 0.75 | 0.556 | 4.30E-51 | Mari |
| allcell | NUDCD2 | 3.47E-18 | 0.771075637 | 0.416 | 0.307 | 6.82E-14 | Mari |
| allcell | SCP2 | 3.23E-42 | 0.770743523 | 0.664 | 0.495 | 6.35E-38 | Mari |
| allcell | SNX17 | 2.05E-39 | 0.769457199 | 0.615 | 0.438 | 4.03E-35 | Mari |
| allcell | HES4 | 6.32E-16 | 0.769040485 | 0.345 | 0.237 | 1.24E-11 | Mari |
| allcell | MRPL3 | 6.35E-27 | 0.769021832 | 0.515 | 0.375 | 1.25E-22 | Mari |
| allcell | EBPL | 8.19E-14 | 0.768709156 | 0.339 | 0.248 | 1.61E-09 | Mari |
| allcell | NMI | 5.06E-26 | 0.767395727 | 0.51 | 0.384 | 9.96E-22 | Mari |
| allcell | EMC6 | 1.29E-22 | 0.767286735 | 0.44 | 0.318 | 2.55E-18 | Mari |
| allcell | ROGDI | 1.14E-10 | 0.766981542 | 0.284 | 0.209 | 2.24E-06 | Mari |
| allcell | GMFG | 5.88E-60 | 0.766070769 | 0.811 | 0.671 | 1.16E-55 | Mari |
| allcell | MRPL17 | 5.53E-10 | 0.766025194 | 0.256 | 0.185 | 1.09E-05 | Mari |
| allcell | ORMDL1 | 1.68E-29 | 0.765046816 | 0.596 | 0.455 | 3.31E-25 | Mari |
| allcell | TXNL4A | 8.67E-44 | 0.764597334 | 0.71 | 0.556 | 1.71E-39 | Mari |

| | | | | | | | |
|---|---|---|---|---|---|---|---|
| allcell | ERH | 1.93E-63 | 0.763666229 | 0.81 | 0.644 | 3.79E-59 | Mari |
| allcell | COMMD6 | 6.82E-66 | 0.763310985 | 0.92 | 0.844 | 1.34E-61 | Mari |
| allcell | ERI3 | 3.46E-14 | 0.762559431 | 0.292 | 0.203 | 6.80E-10 | Mari |
| allcell | MRPL55 | 1.94E-25 | 0.762203835 | 0.517 | 0.384 | 3.82E-21 | Mari |
| allcell | UQCR10 | 1.97E-78 | 0.760002527 | 0.871 | 0.701 | 3.87E-74 | Mari |
| allcell | PPIB | 5.55E-82 | 0.758216531 | 0.927 | 0.863 | 1.09E-77 | Mari |
| allcell | RBCK1 | 2.37E-26 | 0.758199448 | 0.553 | 0.421 | 4.67E-22 | Mari |
| allcell | ATRAID | 9.03E-36 | 0.758098305 | 0.573 | 0.42 | 1.78E-31 | Mari |
| allcell | TRMT112 | 3.79E-58 | 0.757072399 | 0.838 | 0.692 | 7.47E-54 | Mari |
| allcell | ATP6V0B | 1.72E-95 | 0.756609572 | 0.921 | 0.797 | 3.38E-91 | Mari |
| allcell | CDC123 | 6.85E-27 | 0.75611694 | 0.489 | 0.352 | 1.35E-22 | Mari |
| allcell | UCP2 | 3.33E-55 | 0.755695767 | 0.848 | 0.698 | 6.56E-51 | Mari |
| allcell | LY96 | 7.00E-34 | 0.755551862 | 0.627 | 0.486 | 1.38E-29 | Mari |
| allcell | MTIF3 | 3.57E-25 | 0.754328613 | 0.503 | 0.368 | 7.03E-21 | Mari |
| allcell | MRPL2 | 3.13E-11 | 0.75402003 | 0.254 | 0.177 | 6.16E-07 | Mari |
| allcell | SLC35B1 | 3.89E-14 | 0.753774806 | 0.335 | 0.243 | 7.66E-10 | Mari |
| allcell | COX7A2 | 4.08E-90 | 0.753733779 | 0.916 | 0.796 | 8.02E-86 | Mari |
| allcell | UBL5 | 7.32E-116 | 0.752069773 | 0.936 | 0.831 | 1.44E-111 | Mari |
| allcell | EMC7 | 1.02E-25 | 0.751546542 | 0.54 | 0.396 | 2.01E-21 | Mari |
| allcell | HPS3 | 3.00E-11 | 0.751474108 | 0.32 | 0.236 | 5.90E-07 | Mari |
| allcell | FAM53C | 1.24E-09 | 0.751424364 | 0.388 | 0.313 | 2.44E-05 | Mari |
| allcell | SMARCB1 | 1.12E-21 | 0.75030028 | 0.462 | 0.335 | 2.20E-17 | Mari |
| allcell | SMC61 | 5.75E-09 | 0.7501045 | 0.266 | 0.199 | 0.00011316 | Mari |
| allcell | POLR2C | 1.03E-14 | 0.749872562 | 0.306 | 0.211 | 2.03E-10 | Mari |
| allcell | COMMD7 | 1.97E-24 | 0.749143315 | 0.461 | 0.328 | 3.88E-20 | Mari |
| allcell | RCN2 | 8.54E-15 | 0.748195575 | 0.363 | 0.268 | 1.68E-10 | Mari |
| allcell | UBE2E3 | 1.64E-14 | 0.746528547 | 0.411 | 0.317 | 3.24E-10 | Mari |
| allcell | ARPC1B | 2.16E-96 | 0.746193154 | 0.948 | 0.86 | 4.25E-92 | Mari |
| allcell | PLRG1 | 2.49E-10 | 0.744612938 | 0.265 | 0.19 | 4.91E-06 | Mari |
| allcell | TBCB | 2.88E-48 | 0.744179348 | 0.722 | 0.551 | 5.66E-44 | Mari |
| allcell | TNFSF13 | 6.68E-11 | 0.743525936 | 0.335 | 0.257 | 1.31E-06 | Mari |
| allcell | MRPS11 | 4.55E-15 | 0.74332527 | 0.355 | 0.254 | 8.96E-11 | Mari |
| allcell | CHD9 | 3.81E-40 | 0.742575365 | 0.669 | 0.489 | 7.50E-36 | Mari |
| allcell | MYL12B | 2.69E-120 | 0.742409448 | 0.941 | 0.874 | 5.30E-116 | Mari |
| allcell | MDP1 | 6.16E-12 | 0.741853533 | 0.296 | 0.213 | 1.21E-07 | Mari |
| allcell | MGST2 | 8.80E-25 | 0.741499537 | 0.549 | 0.419 | 1.73E-20 | Mari |
| allcell | MALSU1 | 1.80E-16 | 0.741409713 | 0.307 | 0.205 | 3.55E-12 | Mari |
| allcell | SETBP11 | 2.67E-15 | 0.740739477 | 0.26 | 0.164 | 5.26E-11 | Mari |
| allcell | AURKAIP1 | 3.15E-62 | 0.740609997 | 0.808 | 0.642 | 6.20E-58 | Mari |
| allcell | NDUFC1 | 4.55E-37 | 0.740347676 | 0.586 | 0.416 | 8.95E-33 | Mari |
| allcell | U2AF1L4 | 9.99E-13 | 0.739625443 | 0.323 | 0.235 | 1.97E-08 | Mari |
| allcell | PLEKHJ1 | 6.61E-15 | 0.739244861 | 0.419 | 0.327 | 1.30E-10 | Mari |
| allcell | ZNHIT1 | 1.28E-43 | 0.738213246 | 0.697 | 0.547 | 2.51E-39 | Mari |
| allcell | APOBEC3G | 1.67E-12 | 0.737644237 | 0.317 | 0.232 | 3.29E-08 | Mari |
| allcell | IFNAR1 | 8.13E-18 | 0.737402638 | 0.489 | 0.382 | 1.60E-13 | Mari |
| allcell | CENPX | 2.19E-31 | 0.73739679 | 0.542 | 0.394 | 4.32E-27 | Mari |
| allcell | PSMB2 | 7.44E-47 | 0.737157395 | 0.72 | 0.549 | 1.46E-42 | Mari |
| allcell | ABRACL | 8.64E-38 | 0.736901512 | 0.674 | 0.519 | 1.70E-33 | Mari |
| allcell | MCM5 | 1.02E-16 | 0.73629577 | 0.403 | 0.298 | 2.01E-12 | Mari |
| allcell | NSMCE1 | 4.17E-16 | 0.735753946 | 0.424 | 0.321 | 8.21E-12 | Mari |
| allcell | EXOSC7 | 1.51E-12 | 0.73504282 | 0.289 | 0.204 | 2.98E-08 | Mari |
| allcell | TMEM160 | 2.22E-37 | 0.734159083 | 0.625 | 0.473 | 4.37E-33 | Mari |
| allcell | CIAO2B | 4.75E-52 | 0.733653397 | 0.784 | 0.623 | 9.35E-48 | Mari |
| allcell | IMP4 | 7.53E-18 | 0.733281986 | 0.41 | 0.3 | 1.48E-13 | Mari |
| allcell | COX14 | 1.12E-42 | 0.731585612 | 0.667 | 0.496 | 2.20E-38 | Mari |
| allcell | ZFAT1 | 1.42E-20 | 0.73114468 | 0.294 | 0.175 | 2.80E-16 | Mari |

| | | | | | | | |
|---|---|---|---|---|---|---|---|
| allcell | P2RY61 | 1.87E-27 | 0.73105362 | 0.428 | 0.27 | 3.69E-23 | Mari |
| allcell | BOLA3 | 3.42E-15 | 0.730775994 | 0.357 | 0.257 | 6.73E-11 | Mari |
| allcell | WDR45 | 6.05E-12 | 0.728976545 | 0.294 | 0.213 | 1.19E-07 | Mari |
| allcell | DYNLRB1 | 3.91E-47 | 0.728889487 | 0.707 | 0.53 | 7.69E-43 | Mari |
| allcell | RPA2 | 3.49E-15 | 0.727214924 | 0.392 | 0.291 | 6.86E-11 | Mari |
| allcell | FKBP3 | 6.86E-17 | 0.726661906 | 0.405 | 0.3 | 1.35E-12 | Mari |
| allcell | MIIP | 1.59E-11 | 0.724999196 | 0.331 | 0.251 | 3.14E-07 | Mari |
| allcell | IFNAR2 | 1.51E-26 | 0.724495614 | 0.598 | 0.46 | 2.97E-22 | Mari |
| allcell | PSMA3 | 1.71E-47 | 0.724428376 | 0.768 | 0.616 | 3.37E-43 | Mari |
| allcell | BTG2 | 2.81E-32 | 0.724401281 | 0.794 | 0.685 | 5.52E-28 | Mari |
| allcell | ACADM | 8.33E-13 | 0.724349115 | 0.347 | 0.259 | 1.64E-08 | Mari |
| allcell | ATP5PF | 1.77E-81 | 0.723626407 | 0.881 | 0.732 | 3.48E-77 | Mari |
| allcell | GMPR2 | 1.92E-09 | 0.722243839 | 0.278 | 0.208 | 3.77E-05 | Mari |
| allcell | PDCD6 | 1.41E-47 | 0.722160453 | 0.717 | 0.526 | 2.78E-43 | Mari |
| allcell | YPEL3 | 1.78E-10 | 0.722104856 | 0.3 | 0.225 | 3.50E-06 | Mari |
| allcell | CPQ | 1.95E-20 | 0.721733878 | 0.523 | 0.398 | 3.85E-16 | Mari |
| allcell | SCAND1 | 1.74E-47 | 0.721569044 | 0.739 | 0.564 | 3.42E-43 | Mari |
| allcell | TNFRSF21 | 3.01E-13 | 0.720706755 | 0.333 | 0.239 | 5.93E-09 | Mari |
| allcell | CDC40 | 4.65E-16 | 0.719495819 | 0.437 | 0.341 | 9.16E-12 | Mari |
| allcell | NDUFA13 | 7.34E-92 | 0.719193213 | 0.923 | 0.783 | 1.44E-87 | Mari |
| allcell | TSN | 2.57E-09 | 0.71780517 | 0.257 | 0.189 | 5.07E-05 | Mari |
| allcell | FEZ2 | 1.42E-17 | 0.716958981 | 0.427 | 0.319 | 2.79E-13 | Mari |
| allcell | COX7A2L | 1.23E-42 | 0.716837953 | 0.803 | 0.696 | 2.42E-38 | Mari |
| allcell | NDUFB1 | 9.76E-65 | 0.716362554 | 0.82 | 0.624 | 1.92E-60 | Mari |
| allcell | SPINT2 | 5.59E-51 | 0.716156151 | 0.803 | 0.615 | 1.10E-46 | Mari |
| allcell | DERL2 | 5.36E-16 | 0.716132776 | 0.371 | 0.268 | 1.05E-11 | Mari |
| allcell | PHPT1 | 5.66E-47 | 0.715722518 | 0.781 | 0.634 | 1.11E-42 | Mari |
| allcell | DUSP22 | 1.27E-14 | 0.714501428 | 0.405 | 0.305 | 2.50E-10 | Mari |
| allcell | BAD | 3.68E-12 | 0.713361982 | 0.334 | 0.249 | 7.23E-08 | Mari |
| allcell | PSMA4 | 2.19E-47 | 0.712997813 | 0.773 | 0.603 | 4.32E-43 | Mari |
| allcell | SUB1 | 8.91E-100 | 0.712727203 | 0.965 | 0.904 | 1.75E-95 | Mari |
| allcell | FAM174C | 2.97E-27 | 0.712073877 | 0.558 | 0.422 | 5.85E-23 | Mari |
| allcell | DDX18 | 4.07E-26 | 0.7114811 | 0.655 | 0.517 | 8.02E-22 | Mari |
| allcell | ECI2 | 4.32E-10 | 0.710975077 | 0.27 | 0.198 | 8.50E-06 | Mari |
| allcell | PDLIM2 | 1.78E-12 | 0.710865472 | 0.342 | 0.252 | 3.49E-08 | Mari |
| allcell | SURF1 | 2.46E-26 | 0.710673299 | 0.551 | 0.412 | 4.84E-22 | Mari |
| allcell | CHCHD5 | 5.16E-25 | 0.710617726 | 0.538 | 0.401 | 1.02E-20 | Mari |
| allcell | MCRS1 | 9.20E-09 | 0.710071869 | 0.254 | 0.188 | 0.000181186 | Mari |
| allcell | IFI27L2 | 1.01E-23 | 0.710051579 | 0.5 | 0.367 | 1.98E-19 | Mari |
| allcell | ANAPC13 | 3.07E-11 | 0.707401067 | 0.278 | 0.198 | 6.04E-07 | Mari |
| allcell | SSBP1 | 5.43E-47 | 0.707006427 | 0.762 | 0.604 | 1.07E-42 | Mari |
| allcell | ERGIC2 | 3.45E-14 | 0.706921168 | 0.395 | 0.298 | 6.80E-10 | Mari |
| allcell | B3GAT3 | 8.62E-13 | 0.706602959 | 0.291 | 0.203 | 1.70E-08 | Mari |
| allcell | GADD45GIP1 | 3.22E-35 | 0.706547585 | 0.644 | 0.496 | 6.34E-31 | Mari |
| allcell | RBIS | 3.27E-52 | 0.706462263 | 0.804 | 0.651 | 6.45E-48 | Mari |
| allcell | ARL16 | 2.96E-11 | 0.705443456 | 0.287 | 0.206 | 5.83E-07 | Mari |
| allcell | TAF9 | 1.67E-16 | 0.705115583 | 0.393 | 0.287 | 3.29E-12 | Mari |
| allcell | CKLF | 1.33E-25 | 0.704229653 | 0.699 | 0.598 | 2.62E-21 | Mari |
| allcell | MAD2L2 | 1.25E-16 | 0.703598118 | 0.393 | 0.283 | 2.45E-12 | Mari |
| allcell | SNAPIN | 6.10E-14 | 0.703057494 | 0.285 | 0.194 | 1.20E-09 | Mari |
| allcell | SNRPA1 | 7.01E-30 | 0.702772131 | 0.549 | 0.391 | 1.38E-25 | Mari |
| allcell | MSRB1 | 2.77E-12 | 0.702673914 | 0.337 | 0.254 | 5.46E-08 | Mari |
| allcell | NDUFS2 | 9.08E-24 | 0.702170872 | 0.546 | 0.421 | 1.79E-19 | Mari |
| allcell | DCTN3 | 2.64E-35 | 0.702114442 | 0.653 | 0.507 | 5.19E-31 | Mari |
| allcell | EDF1 | 1.86E-107 | 0.701697873 | 0.949 | 0.854 | 3.65E-103 | Mari |
| allcell | NUDT5 | 7.14E-17 | 0.70134786 | 0.404 | 0.298 | 1.41E-12 | Mari |

| | | | | | | | |
|---|---|---|---|---|---|---|---|
| allcell | YIPF5 | 7.04E-08 | 0.701006667 | 0.26 | 0.2 | 0.001384984 | Mari |
| allcell | ATP6V0E1 | 9.06E-85 | 0.700788987 | 0.921 | 0.82 | 1.78E-80 | Mari |
| allcell | UFD1 | 7.50E-20 | 0.700105517 | 0.517 | 0.399 | 1.48E-15 | Mari |
| allcell | NUTF2 | 1.99E-30 | 0.699859035 | 0.561 | 0.411 | 3.91E-26 | Mari |
| allcell | POLR3GL | 6.91E-12 | 0.699222907 | 0.309 | 0.227 | 1.36E-07 | Mari |
| allcell | HCCS | 5.80E-17 | 0.698294601 | 0.347 | 0.239 | 1.14E-12 | Mari |
| allcell | MRPL54 | 1.84E-30 | 0.698231516 | 0.606 | 0.462 | 3.62E-26 | Mari |
| allcell | MYDGF | 1.15E-45 | 0.697930483 | 0.774 | 0.611 | 2.27E-41 | Mari |
| allcell | SF3B6 | 3.89E-54 | 0.697824488 | 0.794 | 0.642 | 7.66E-50 | Mari |
| allcell | MPEG1 | 2.81E-16 | 0.697510965 | 0.486 | 0.372 | 5.54E-12 | Mari |
| allcell | PRKRA | 1.39E-12 | 0.697193751 | 0.375 | 0.289 | 2.74E-08 | Mari |
| allcell | MICOS10 | 4.08E-62 | 0.69694547 | 0.855 | 0.708 | 8.03E-58 | Mari |
| allcell | SSNA1 | 2.16E-29 | 0.696722423 | 0.622 | 0.496 | 4.25E-25 | Mari |
| allcell | NMT1 | 1.56E-15 | 0.696283086 | 0.451 | 0.356 | 3.06E-11 | Mari |
| allcell | TMEM147 | 1.46E-24 | 0.696098405 | 0.481 | 0.345 | 2.87E-20 | Mari |
| allcell | ARL1 | 6.84E-13 | 0.695777826 | 0.333 | 0.245 | 1.35E-08 | Mari |
| allcell | RNASEH2B | 2.74E-24 | 0.695318824 | 0.502 | 0.36 | 5.39E-20 | Mari |
| allcell | ESD | 6.63E-28 | 0.694871905 | 0.599 | 0.466 | 1.30E-23 | Mari |
| allcell | GET3 | 2.15E-20 | 0.694728257 | 0.496 | 0.383 | 4.23E-16 | Mari |
| allcell | CORO1A | 6.62E-55 | 0.694503486 | 0.869 | 0.725 | 1.30E-50 | Mari |
| allcell | MRPS16 | 1.30E-14 | 0.693449687 | 0.399 | 0.307 | 2.56E-10 | Mari |
| allcell | MTHFD2L | 1.19E-07 | 0.693306294 | 0.253 | 0.191 | 0.00234245 | Mari |
| allcell | DCTN6 | 2.63E-12 | 0.69152326 | 0.34 | 0.255 | 5.17E-08 | Mari |
| allcell | TSR3 | 1.70E-12 | 0.691399708 | 0.346 | 0.261 | 3.34E-08 | Mari |
| allcell | FBXO6 | 1.48E-19 | 0.689524698 | 0.461 | 0.341 | 2.90E-15 | Mari |
| allcell | SATB1-AS11 | 1.41E-07 | 0.688409712 | 0.254 | 0.19 | 0.002771125 | Mari |
| allcell | MTCH2 | 8.37E-22 | 0.687771054 | 0.48 | 0.356 | 1.65E-17 | Mari |
| allcell | CARHSP1 | 2.70E-13 | 0.687630769 | 0.325 | 0.234 | 5.31E-09 | Mari |
| allcell | CLECL1 | 3.61E-15 | 0.686800209 | 0.414 | 0.309 | 7.10E-11 | Mari |
| allcell | UNC93B1 | 1.11E-17 | 0.686758247 | 0.505 | 0.4 | 2.19E-13 | Mari |
| allcell | FIBP | 3.49E-27 | 0.686746988 | 0.582 | 0.455 | 6.87E-23 | Mari |
| allcell | YDJC | 4.72E-10 | 0.686470506 | 0.293 | 0.217 | 9.29E-06 | Mari |
| allcell | LAMTOR2 | 1.09E-35 | 0.686395332 | 0.679 | 0.553 | 2.14E-31 | Mari |
| allcell | SUMO1 | 1.41E-49 | 0.68637413 | 0.773 | 0.609 | 2.77E-45 | Mari |
| allcell | SUMO3 | 2.99E-34 | 0.684747712 | 0.668 | 0.513 | 5.88E-30 | Mari |
| allcell | OSTF1 | 3.08E-36 | 0.6834347 | 0.773 | 0.637 | 6.06E-32 | Mari |
| allcell | MRPL52 | 1.48E-35 | 0.68248758 | 0.672 | 0.535 | 2.91E-31 | Mari |
| allcell | TAF12 | 1.76E-12 | 0.682155018 | 0.343 | 0.255 | 3.45E-08 | Mari |
| allcell | FLOT1 | 5.08E-37 | 0.68194386 | 0.644 | 0.482 | 1.00E-32 | Mari |
| allcell | AIMP1 | 1.91E-26 | 0.681493557 | 0.54 | 0.403 | 3.77E-22 | Mari |
| allcell | BAX | 4.43E-41 | 0.681334998 | 0.729 | 0.566 | 8.72E-37 | Mari |
| allcell | MRPL27 | 9.29E-15 | 0.680908445 | 0.336 | 0.241 | 1.83E-10 | Mari |
| allcell | LEPROTL1 | 2.95E-27 | 0.680184844 | 0.557 | 0.411 | 5.80E-23 | Mari |
| allcell | RNASEH1 | 2.81E-08 | 0.679286424 | 0.269 | 0.206 | 0.000553795 | Mari |
| allcell | CCDC115 | 2.57E-15 | 0.679052049 | 0.365 | 0.265 | 5.06E-11 | Mari |
| allcell | CENPC | 1.47E-11 | 0.678582045 | 0.424 | 0.343 | 2.90E-07 | Mari |
| allcell | GLRX5 | 4.29E-30 | 0.678327493 | 0.549 | 0.396 | 8.44E-26 | Mari |
| allcell | FLYWCH2 | 6.93E-09 | 0.678107145 | 0.265 | 0.198 | 0.000136465 | Mari |
| allcell | DPY30 | 2.28E-16 | 0.678084747 | 0.354 | 0.25 | 4.50E-12 | Mari |
| allcell | VAMP2 | 1.16E-20 | 0.677838155 | 0.486 | 0.365 | 2.28E-16 | Mari |
| allcell | XPA | 8.32E-13 | 0.677168359 | 0.287 | 0.203 | 1.64E-08 | Mari |
| allcell | UQCRQ | 3.24E-52 | 0.676569996 | 0.79 | 0.635 | 6.37E-48 | Mari |
| allcell | MRPL22 | 8.24E-19 | 0.676111044 | 0.418 | 0.3 | 1.62E-14 | Mari |
| allcell | CALHM6 | 8.77E-18 | 0.675187927 | 0.742 | 0.67 | 1.73E-13 | Mari |
| allcell | PSMD9 | 3.67E-30 | 0.674764029 | 0.62 | 0.479 | 7.22E-26 | Mari |
| allcell | RGS19 | 2.42E-30 | 0.674742283 | 0.637 | 0.494 | 4.76E-26 | Mari |

| | | | | | | | |
|---|---|---|---|---|---|---|---|
| allcell | SDHAF2 | 2.21E-12 | 0.673526963 | 0.348 | 0.262 | 4.35E-08 | Mari |
| allcell | TXNDC15 | 2.05E-09 | 0.672447074 | 0.265 | 0.196 | 4.03E-05 | Mari |
| allcell | TIMM10 | 3.70E-24 | 0.672256543 | 0.559 | 0.429 | 7.28E-20 | Mari |
| allcell | LSM3 | 4.05E-32 | 0.671665555 | 0.642 | 0.489 | 7.97E-28 | Mari |
| allcell | MRPL12 | 3.84E-13 | 0.670650937 | 0.362 | 0.27 | 7.56E-09 | Mari |
| allcell | ATP5MC1 | 8.21E-43 | 0.669848223 | 0.737 | 0.578 | 1.62E-38 | Mari |
| allcell | CCDC69 | 8.81E-08 | 0.669396254 | 0.335 | 0.275 | 0.001734652 | Mari |
| allcell | MNDA | 2.06E-16 | 0.669211484 | 0.631 | 0.566 | 4.05E-12 | Mari |
| allcell | POLR2L | 4.58E-56 | 0.669159413 | 0.837 | 0.685 | 9.02E-52 | Mari |
| allcell | LRR1 | 1.79E-08 | 0.668707742 | 0.285 | 0.215 | 0.00035153 | Mari |
| allcell | BLNK | 2.26E-10 | 0.668594001 | 0.252 | 0.174 | 4.45E-06 | Mari |
| allcell | DYNLT1 | 1.06E-27 | 0.667943635 | 0.578 | 0.437 | 2.08E-23 | Mari |
| allcell | STUB1 | 1.19E-24 | 0.667459732 | 0.561 | 0.441 | 2.33E-20 | Mari |
| allcell | LILRB4 | 5.43E-18 | 0.667080217 | 0.46 | 0.334 | 1.07E-13 | Mari |
| allcell | COMMD4 | 2.87E-13 | 0.666924383 | 0.385 | 0.3 | 5.64E-09 | Mari |
| allcell | TMED3 | 3.88E-17 | 0.664156583 | 0.469 | 0.361 | 7.64E-13 | Mari |
| allcell | MRPL41 | 4.96E-41 | 0.663959298 | 0.676 | 0.532 | 9.76E-37 | Mari |
| allcell | COX16 | 5.28E-22 | 0.663805805 | 0.519 | 0.396 | 1.04E-17 | Mari |
| allcell | PDIA6 | 5.28E-37 | 0.662850377 | 0.728 | 0.577 | 1.04E-32 | Mari |
| allcell | TRMT1 | 3.37E-13 | 0.662128112 | 0.328 | 0.236 | 6.63E-09 | Mari |
| allcell | ARPC1A | 2.59E-12 | 0.660535366 | 0.344 | 0.259 | 5.10E-08 | Mari |
| allcell | AK6 | 2.23E-13 | 0.660089133 | 0.281 | 0.191 | 4.39E-09 | Mari |
| allcell | CAT | 6.69E-19 | 0.660028474 | 0.566 | 0.459 | 1.32E-14 | Mari |
| allcell | POLR2J | 3.38E-27 | 0.660023698 | 0.57 | 0.44 | 6.66E-23 | Mari |
| allcell | EMB | 8.22E-10 | 0.659579142 | 0.361 | 0.288 | 1.62E-05 | Mari |
| allcell | NDUFAF3 | 1.74E-24 | 0.658480928 | 0.555 | 0.433 | 3.42E-20 | Mari |
| allcell | ITGB1BP1 | 6.36E-17 | 0.658096338 | 0.43 | 0.323 | 1.25E-12 | Mari |
| allcell | PSMA5 | 1.76E-44 | 0.658065932 | 0.77 | 0.642 | 3.47E-40 | Mari |
| allcell | EXOSC5 | 2.92E-09 | 0.657818342 | 0.276 | 0.205 | 5.74E-05 | Mari |
| allcell | CLPP | 8.36E-14 | 0.657025375 | 0.378 | 0.285 | 1.65E-09 | Mari |
| allcell | SKA2 | 7.09E-09 | 0.656746329 | 0.254 | 0.189 | 0.000139522 | Mari |
| allcell | DHRS4L2 | 1.70E-08 | 0.656408983 | 0.294 | 0.23 | 0.000333777 | Mari |
| allcell | TXNDC12 | 6.38E-21 | 0.655847725 | 0.497 | 0.376 | 1.25E-16 | Mari |
| allcell | PAM16 | 1.32E-08 | 0.655808458 | 0.3 | 0.235 | 0.000259735 | Mari |
| allcell | UBE2L3 | 1.33E-45 | 0.655469856 | 0.773 | 0.637 | 2.62E-41 | Mari |
| allcell | NDUFB10 | 2.45E-49 | 0.654433901 | 0.801 | 0.641 | 4.82E-45 | Mari |
| allcell | PDCD10 | 6.69E-29 | 0.654379191 | 0.584 | 0.444 | 1.32E-24 | Mari |
| allcell | MRPL33 | 2.95E-31 | 0.654022257 | 0.62 | 0.461 | 5.82E-27 | Mari |
| allcell | HS3ST3B11 | 1.24E-08 | 0.653936789 | 0.281 | 0.209 | 0.000243303 | Mari |
| allcell | JAGN1 | 1.79E-11 | 0.653828819 | 0.338 | 0.258 | 3.53E-07 | Mari |
| allcell | TCF41 | 3.28E-12 | 0.653679364 | 0.585 | 0.514 | 6.45E-08 | Mari |
| allcell | MRPS28 | 4.68E-13 | 0.652492227 | 0.348 | 0.261 | 9.21E-09 | Mari |
| allcell | PIP4P1 | 1.02E-11 | 0.652392045 | 0.302 | 0.217 | 2.01E-07 | Mari |
| allcell | AP2S1 | 8.75E-64 | 0.652383654 | 0.881 | 0.785 | 1.72E-59 | Mari |
| allcell | SSR4 | 3.57E-56 | 0.652378509 | 0.914 | 0.845 | 7.02E-52 | Mari |
| allcell | LSM2 | 2.42E-25 | 0.65164873 | 0.551 | 0.413 | 4.77E-21 | Mari |
| allcell | CD2AP1 | 3.22E-14 | 0.651457659 | 0.44 | 0.343 | 6.34E-10 | Mari |
| allcell | HSH2D1 | 4.35E-09 | 0.651263837 | 0.263 | 0.191 | 8.57E-05 | Mari |
| allcell | CKAP2 | 2.06E-06 | 0.65084756 | 0.292 | 0.235 | 0.040451421 | Mari |
| allcell | VDAC3 | 3.23E-28 | 0.65070466 | 0.64 | 0.506 | 6.36E-24 | Mari |
| allcell | MRPS7 | 5.78E-18 | 0.64964398 | 0.459 | 0.352 | 1.14E-13 | Mari |
| allcell | NDUFS3 | 6.84E-20 | 0.649472939 | 0.528 | 0.418 | 1.35E-15 | Mari |
| allcell | HTATIP2 | 4.12E-08 | 0.648533127 | 0.295 | 0.228 | 0.000811346 | Mari |
| allcell | PPIE | 6.29E-09 | 0.648479314 | 0.262 | 0.195 | 0.000123716 | Mari |
| allcell | RNF126 | 1.46E-16 | 0.648282453 | 0.462 | 0.36 | 2.87E-12 | Mari |
| allcell | TP53I13 | 1.73E-13 | 0.64803042 | 0.38 | 0.286 | 3.40E-09 | Mari |

| | | | | | | | |
|---|---|---|---|---|---|---|---|
| allcell | GPAA1 | 2.86E-11 | 0.647855286 | 0.332 | 0.252 | 5.63E-07 | Mari |
| allcell | SLC50A1 | 1.03E-13 | 0.646892692 | 0.349 | 0.258 | 2.04E-09 | Mari |
| allcell | LARP7 | 3.45E-13 | 0.645508136 | 0.389 | 0.3 | 6.79E-09 | Mari |
| allcell | MED8 | 1.64E-13 | 0.64521822 | 0.358 | 0.263 | 3.22E-09 | Mari |
| allcell | ATP5ME | 1.43E-57 | 0.643636357 | 0.836 | 0.692 | 2.82E-53 | Mari |
| allcell | CFL1 | 1.85E-96 | 0.643248824 | 0.97 | 0.944 | 3.65E-92 | Mari |
| allcell | HVCN1 | 4.57E-11 | 0.643046912 | 0.391 | 0.305 | 8.99E-07 | Mari |
| allcell | PSMB9 | 2.27E-56 | 0.642759431 | 0.893 | 0.806 | 4.47E-52 | Mari |
| allcell | CD300A | 1.48E-16 | 0.642684961 | 0.394 | 0.284 | 2.92E-12 | Mari |
| allcell | TMEM179B | 1.63E-24 | 0.64257515 | 0.525 | 0.379 | 3.20E-20 | Mari |
| allcell | MRPL23 | 6.38E-20 | 0.642014335 | 0.462 | 0.345 | 1.26E-15 | Mari |
| allcell | BORCS7 | 5.05E-09 | 0.641865337 | 0.331 | 0.261 | 9.95E-05 | Mari |
| allcell | NDUFB5 | 3.12E-38 | 0.641330658 | 0.735 | 0.6 | 6.15E-34 | Mari |
| allcell | TOMM6 | 3.87E-51 | 0.641253337 | 0.837 | 0.71 | 7.61E-47 | Mari |
| allcell | IFI35 | 1.45E-10 | 0.640614479 | 0.357 | 0.281 | 2.86E-06 | Mari |
| allcell | SRP19 | 1.95E-36 | 0.640265562 | 0.714 | 0.553 | 3.84E-32 | Mari |
| allcell | CHURC1 | 1.97E-36 | 0.639605357 | 0.733 | 0.59 | 3.88E-32 | Mari |
| allcell | DUSP23 | 1.06E-11 | 0.639489477 | 0.39 | 0.312 | 2.08E-07 | Mari |
| allcell | NDUFA5 | 3.34E-16 | 0.638341001 | 0.461 | 0.362 | 6.58E-12 | Mari |
| allcell | TMA7 | 1.76E-89 | 0.637906861 | 0.939 | 0.866 | 3.47E-85 | Mari |
| allcell | RANBP1 | 7.31E-29 | 0.637795901 | 0.648 | 0.514 | 1.44E-24 | Mari |
| allcell | BANF1 | 3.20E-35 | 0.637123893 | 0.689 | 0.542 | 6.31E-31 | Mari |
| allcell | MRPS26 | 6.10E-09 | 0.636781105 | 0.283 | 0.215 | 0.000120136 | Mari |
| allcell | SERF2 | 8.97E-141 | 0.636554479 | 0.975 | 0.955 | 1.76E-136 | Mari |
| allcell | NT5DC1 | 7.16E-07 | 0.635917354 | 0.27 | 0.209 | 0.014100592 | Mari |
| allcell | SLC25A39 | 2.14E-15 | 0.635863395 | 0.438 | 0.341 | 4.21E-11 | Mari |
| allcell | MDH2 | 2.25E-42 | 0.634372944 | 0.752 | 0.598 | 4.44E-38 | Mari |
| allcell | TRAPPC6A | 8.60E-21 | 0.634208154 | 0.495 | 0.369 | 1.69E-16 | Mari |
| allcell | MPG | 1.56E-24 | 0.633816648 | 0.596 | 0.462 | 3.07E-20 | Mari |
| allcell | NDUFB8 | 4.34E-49 | 0.633744235 | 0.844 | 0.696 | 8.54E-45 | Mari |
| allcell | PHF11 | 1.27E-16 | 0.633725109 | 0.446 | 0.345 | 2.50E-12 | Mari |
| allcell | MDH1 | 1.16E-28 | 0.633356995 | 0.628 | 0.477 | 2.28E-24 | Mari |
| allcell | PPP1R35 | 6.23E-08 | 0.63269757 | 0.255 | 0.195 | 0.001226731 | Mari |
| allcell | TGFBR21 | 8.41E-21 | 0.632669634 | 0.537 | 0.403 | 1.66E-16 | Mari |
| allcell | EXOSC8 | 1.14E-16 | 0.632600723 | 0.4 | 0.291 | 2.23E-12 | Mari |
| allcell | MDFIC | 6.65E-15 | 0.632479359 | 0.511 | 0.407 | 1.31E-10 | Mari |
| allcell | POMP | 5.10E-65 | 0.631326416 | 0.907 | 0.826 | 1.00E-60 | Mari |
| allcell | ARHGDIB | 1.33E-32 | 0.631131241 | 0.824 | 0.775 | 2.62E-28 | Mari |
| allcell | DPP7 | 1.52E-31 | 0.630622799 | 0.71 | 0.583 | 2.99E-27 | Mari |
| allcell | PRMT91 | 4.54E-27 | 0.630542693 | 0.582 | 0.426 | 8.93E-23 | Mari |
| allcell | EMC2 | 5.93E-10 | 0.630146669 | 0.306 | 0.23 | 1.17E-05 | Mari |
| allcell | COPE | 3.42E-67 | 0.629134422 | 0.905 | 0.782 | 6.74E-63 | Mari |
| allcell | SNRPB2 | 7.50E-34 | 0.628664846 | 0.678 | 0.512 | 1.48E-29 | Mari |
| allcell | UCHL5 | 1.96E-10 | 0.628661574 | 0.338 | 0.264 | 3.85E-06 | Mari |
| allcell | VPS29 | 4.77E-55 | 0.628399036 | 0.839 | 0.709 | 9.38E-51 | Mari |
| allcell | GLOD4 | 3.91E-08 | 0.627989325 | 0.293 | 0.234 | 0.0007692 | Mari |
| allcell | OST4 | 3.05E-77 | 0.627669642 | 0.924 | 0.851 | 5.99E-73 | Mari |
| allcell | TMEM14B | 1.90E-43 | 0.627604604 | 0.774 | 0.592 | 3.75E-39 | Mari |
| allcell | MTX1 | 8.07E-08 | 0.626923691 | 0.252 | 0.191 | 0.001588545 | Mari |
| allcell | ECH1 | 1.14E-22 | 0.626713175 | 0.556 | 0.43 | 2.24E-18 | Mari |
| allcell | COA1 | 2.82E-14 | 0.626073176 | 0.436 | 0.342 | 5.55E-10 | Mari |
| allcell | NSL1 | 6.23E-11 | 0.623558636 | 0.309 | 0.228 | 1.23E-06 | Mari |
| allcell | S100A11 | 2.66E-33 | 0.623254701 | 0.94 | 0.901 | 5.23E-29 | Mari |
| allcell | FAM50A | 1.21E-16 | 0.623011356 | 0.489 | 0.39 | 2.39E-12 | Mari |
| allcell | ATP5F1E | 4.61E-161 | 0.622848725 | 0.977 | 0.967 | 9.08E-157 | Mari |
| allcell | SNHG8 | 7.68E-40 | 0.62282333 | 0.866 | 0.759 | 1.51E-35 | Mari |

| | | | | | | | |
|---|---|---|---|---|---|---|---|
| allcell | EIF2AK2 | 1.41E-08 | 0.622737255 | 0.308 | 0.238 | 0.00027735 | Mari |
| allcell | FBXL15 | 1.84E-07 | 0.621873918 | 0.281 | 0.219 | 0.003621582 | Mari |
| allcell | COPS6 | 2.42E-14 | 0.621730594 | 0.429 | 0.335 | 4.77E-10 | Mari |
| allcell | CMC1 | 1.10E-13 | 0.620902545 | 0.391 | 0.299 | 2.17E-09 | Mari |
| allcell | PDCD2 | 1.23E-17 | 0.620370067 | 0.467 | 0.359 | 2.43E-13 | Mari |
| allcell | PIGP | 2.10E-08 | 0.619985537 | 0.334 | 0.267 | 0.000412809 | Mari |
| allcell | AGPAT2 | 5.91E-14 | 0.619463466 | 0.424 | 0.324 | 1.16E-09 | Mari |
| allcell | DPM2 | 2.70E-13 | 0.619305301 | 0.298 | 0.205 | 5.32E-09 | Mari |
| allcell | PSMB1 | 2.04E-58 | 0.619186647 | 0.878 | 0.766 | 4.02E-54 | Mari |
| allcell | GOSR2 | 4.23E-09 | 0.619032163 | 0.26 | 0.192 | 8.33E-05 | Mari |
| allcell | ROMO1 | 4.55E-38 | 0.618676128 | 0.727 | 0.585 | 8.95E-34 | Mari |
| allcell | IFITM2 | 2.13E-17 | 0.617895474 | 0.624 | 0.536 | 4.18E-13 | Mari |
| allcell | JPT1 | 3.30E-33 | 0.617858952 | 0.734 | 0.597 | 6.50E-29 | Mari |
| allcell | EIF5A | 1.08E-38 | 0.617673271 | 0.811 | 0.715 | 2.13E-34 | Mari |
| allcell | FAM32A | 6.50E-17 | 0.61647573 | 0.462 | 0.355 | 1.28E-12 | Mari |
| allcell | VTI1B | 7.94E-12 | 0.61618746 | 0.419 | 0.346 | 1.56E-07 | Mari |
| allcell | MFNG | 3.61E-07 | 0.615706693 | 0.275 | 0.216 | 0.007101378 | Mari |
| allcell | IFI16 | 1.04E-34 | 0.615559507 | 0.762 | 0.615 | 2.04E-30 | Mari |
| allcell | CLTB | 1.28E-33 | 0.615102454 | 0.713 | 0.581 | 2.51E-29 | Mari |
| allcell | SEM1 | 6.02E-56 | 0.613491581 | 0.846 | 0.737 | 1.19E-51 | Mari |
| allcell | HSBP1 | 5.88E-30 | 0.613107327 | 0.68 | 0.561 | 1.16E-25 | Mari |
| allcell | BCAS2 | 4.92E-22 | 0.612957792 | 0.59 | 0.475 | 9.68E-18 | Mari |
| allcell | MRPL43 | 3.74E-24 | 0.612947655 | 0.517 | 0.383 | 7.36E-20 | Mari |
| allcell | UQCRFS1 | 5.56E-33 | 0.612110724 | 0.702 | 0.565 | 1.10E-28 | Mari |
| allcell | BEX4 | 7.28E-14 | 0.611792835 | 0.385 | 0.288 | 1.43E-09 | Mari |
| allcell | SUCLA2 | 4.46E-08 | 0.610455485 | 0.25 | 0.187 | 0.000877197 | Mari |
| allcell | HSD17B10 | 8.69E-19 | 0.610377823 | 0.482 | 0.367 | 1.71E-14 | Mari |
| allcell | CALCRL | 3.50E-20 | 0.610275173 | 0.446 | 0.311 | 6.89E-16 | Mari |
| allcell | TMEM134 | 4.57E-11 | 0.609044505 | 0.359 | 0.279 | 8.99E-07 | Mari |
| allcell | IL10RB | 1.20E-12 | 0.6089908 | 0.381 | 0.291 | 2.37E-08 | Mari |
| allcell | MIA2 | 2.53E-08 | 0.608664415 | 0.358 | 0.288 | 0.000498565 | Mari |
| allcell | COMMD2 | 5.46E-14 | 0.607007351 | 0.381 | 0.275 | 1.07E-09 | Mari |
| allcell | YPEL5 | 8.40E-32 | 0.606923207 | 0.827 | 0.702 | 1.65E-27 | Mari |
| allcell | DHRS4 | 1.91E-07 | 0.606649078 | 0.259 | 0.198 | 0.003751622 | Mari |
| allcell | PIN4 | 2.75E-12 | 0.60624948 | 0.35 | 0.26 | 5.41E-08 | Mari |
| allcell | SMIM15 | 7.09E-10 | 0.605743619 | 0.262 | 0.192 | 1.39E-05 | Mari |
| allcell | SNRPE | 7.52E-32 | 0.605563135 | 0.678 | 0.54 | 1.48E-27 | Mari |
| allcell | SPIB | 1.44E-21 | 0.605497243 | 0.49 | 0.352 | 2.84E-17 | Mari |
| allcell | TPRKB | 3.76E-22 | 0.605476516 | 0.523 | 0.4 | 7.40E-18 | Mari |
| allcell | PLSCR1 | 2.32E-25 | 0.603608968 | 0.749 | 0.629 | 4.57E-21 | Mari |
| allcell | PSMD6 | 1.17E-14 | 0.602045187 | 0.425 | 0.334 | 2.31E-10 | Mari |
| allcell | NDUFS8 | 1.50E-36 | 0.601900825 | 0.713 | 0.566 | 2.95E-32 | Mari |
| allcell | TSG101 | 6.45E-13 | 0.601581371 | 0.414 | 0.327 | 1.27E-08 | Mari |
| allcell | RAD51C | 3.87E-09 | 0.601313678 | 0.298 | 0.225 | 7.62E-05 | Mari |
| allcell | BCAP29 | 8.82E-10 | 0.6003253 | 0.316 | 0.242 | 1.74E-05 | Mari |
| allcell | TMEM126B | 1.27E-08 | 0.599924162 | 0.285 | 0.22 | 0.000250715 | Mari |
| allcell | BAG1 | 3.66E-32 | 0.599560276 | 0.739 | 0.599 | 7.20E-28 | Mari |
| allcell | CCT8 | 1.06E-38 | 0.599391734 | 0.768 | 0.646 | 2.09E-34 | Mari |
| allcell | S100A71 | 3.90E-09 | 0.598630957 | 0.254 | 0.186 | 7.69E-05 | Mari |
| allcell | PSMB6 | 1.11E-47 | 0.598244757 | 0.82 | 0.692 | 2.19E-43 | Mari |
| allcell | TFPT | 2.45E-11 | 0.59821584 | 0.345 | 0.262 | 4.82E-07 | Mari |
| allcell | PFDN1 | 2.36E-21 | 0.597723067 | 0.478 | 0.338 | 4.65E-17 | Mari |
| allcell | TXN2 | 1.15E-25 | 0.597123539 | 0.623 | 0.5 | 2.27E-21 | Mari |
| allcell | PSMC5 | 9.22E-27 | 0.596922882 | 0.629 | 0.499 | 1.81E-22 | Mari |
| allcell | DTD1 | 4.36E-11 | 0.59676573 | 0.341 | 0.254 | 8.59E-07 | Mari |
| allcell | STX7 | 8.70E-24 | 0.596209035 | 0.647 | 0.525 | 1.71E-19 | Mari |

| | | | | | | | |
|---|---|---|---|---|---|---|---|
| allcell | LYRM2 | 2.02E-09 | 0.596188034 | 0.331 | 0.259 | 3.98E-05 | Mari |
| allcell | ZNRF2 | 8.07E-21 | 0.595873128 | 0.54 | 0.406 | 1.59E-16 | Mari |
| allcell | MICOS13 | 2.33E-40 | 0.595247493 | 0.762 | 0.611 | 4.59E-36 | Mari |
| allcell | ZDHHC24 | 1.57E-08 | 0.594681282 | 0.341 | 0.274 | 0.000309694 | Mari |
| allcell | SREK1IP1 | 5.96E-12 | 0.594586207 | 0.33 | 0.243 | 1.17E-07 | Mari |
| allcell | SASH3 | 5.61E-07 | 0.593986564 | 0.271 | 0.214 | 0.011049494 | Mari |
| allcell | MZT1 | 2.46E-08 | 0.593187138 | 0.314 | 0.253 | 0.00048384 | Mari |
| allcell | NOP10 | 1.26E-34 | 0.592975101 | 0.736 | 0.597 | 2.48E-30 | Mari |
| allcell | MPC2 | 2.63E-24 | 0.592658068 | 0.612 | 0.484 | 5.18E-20 | Mari |
| allcell | MRPL9 | 6.34E-11 | 0.592302491 | 0.311 | 0.23 | 1.25E-06 | Mari |
| allcell | WASHC3 | 7.27E-22 | 0.591861366 | 0.48 | 0.349 | 1.43E-17 | Mari |
| allcell | HSD17B41 | 1.45E-06 | 0.591803848 | 0.297 | 0.241 | 0.028625666 | Mari |
| allcell | AHI11 | 1.48E-15 | 0.591518537 | 0.471 | 0.359 | 2.92E-11 | Mari |
| allcell | TNFSF9 | 8.07E-11 | 0.590954033 | 0.432 | 0.352 | 1.59E-06 | Mari |
| allcell | CD2BP2 | 3.24E-11 | 0.590930136 | 0.356 | 0.278 | 6.37E-07 | Mari |
| allcell | NIPSNAP2 | 7.52E-12 | 0.590671385 | 0.391 | 0.306 | 1.48E-07 | Mari |
| allcell | TRAM1 | 2.96E-26 | 0.589927775 | 0.745 | 0.637 | 5.83E-22 | Mari |
| allcell | SEPHS11 | 3.45E-09 | 0.589407776 | 0.291 | 0.222 | 6.78E-05 | Mari |
| allcell | NR3C1 | 5.72E-12 | 0.588689885 | 0.728 | 0.649 | 1.13E-07 | Mari |
| allcell | BRK1 | 1.54E-45 | 0.588602929 | 0.839 | 0.714 | 3.03E-41 | Mari |
| allcell | SDHD | 2.26E-24 | 0.588535803 | 0.603 | 0.479 | 4.44E-20 | Mari |
| allcell | TMED10 | 3.90E-21 | 0.588140783 | 0.688 | 0.576 | 7.68E-17 | Mari |
| allcell | KRTCAP2 | 3.72E-41 | 0.587745503 | 0.811 | 0.676 | 7.32E-37 | Mari |
| allcell | PWP1 | 2.28E-08 | 0.587299919 | 0.307 | 0.242 | 0.000449541 | Mari |
| allcell | NOL7 | 7.53E-31 | 0.587233056 | 0.701 | 0.561 | 1.48E-26 | Mari |
| allcell | SHARPIN | 4.66E-09 | 0.586186567 | 0.325 | 0.254 | 9.17E-05 | Mari |
| allcell | SP110 | 1.12E-21 | 0.585794805 | 0.587 | 0.451 | 2.20E-17 | Mari |
| allcell | APRT | 2.34E-63 | 0.585425742 | 0.905 | 0.803 | 4.61E-59 | Mari |
| allcell | MRPL11 | 1.13E-21 | 0.584662375 | 0.531 | 0.408 | 2.22E-17 | Mari |
| allcell | RRP36 | 8.77E-13 | 0.584621072 | 0.359 | 0.268 | 1.73E-08 | Mari |
| allcell | LSM1 | 3.32E-14 | 0.584065094 | 0.455 | 0.361 | 6.54E-10 | Mari |
| allcell | NUP42 | 1.31E-10 | 0.583773756 | 0.345 | 0.266 | 2.57E-06 | Mari |
| allcell | DIMT1 | 3.96E-08 | 0.583697785 | 0.306 | 0.241 | 0.000780168 | Mari |
| allcell | MED10 | 1.83E-21 | 0.583586715 | 0.544 | 0.418 | 3.60E-17 | Mari |
| allcell | CTSA | 1.13E-08 | 0.583501552 | 0.3 | 0.232 | 0.00022264 | Mari |
| allcell | PSMD8 | 4.32E-43 | 0.582933912 | 0.817 | 0.679 | 8.50E-39 | Mari |
| allcell | PPP1R7 | 4.29E-14 | 0.58278489 | 0.462 | 0.373 | 8.45E-10 | Mari |
| allcell | OTUD6B-AS1 | 2.31E-15 | 0.582540957 | 0.375 | 0.272 | 4.55E-11 | Mari |
| allcell | ATP5MD | 4.02E-48 | 0.580970766 | 0.866 | 0.77 | 7.91E-44 | Mari |
| allcell | MRPL21 | 4.43E-11 | 0.580736449 | 0.407 | 0.329 | 8.72E-07 | Mari |
| allcell | SVBP | 2.96E-13 | 0.580398881 | 0.343 | 0.249 | 5.83E-09 | Mari |
| allcell | CEBPZOS | 1.21E-09 | 0.579844631 | 0.298 | 0.223 | 2.39E-05 | Mari |
| allcell | IRF9 | 2.55E-21 | 0.579592519 | 0.548 | 0.423 | 5.02E-17 | Mari |
| allcell | ATG101 | 7.49E-14 | 0.579038666 | 0.525 | 0.426 | 1.48E-09 | Mari |
| allcell | SLC39A3 | 2.24E-12 | 0.578912021 | 0.313 | 0.224 | 4.40E-08 | Mari |
| allcell | RFXANK | 1.00E-06 | 0.578431415 | 0.292 | 0.24 | 0.019780313 | Mari |
| allcell | FIS1 | 7.28E-30 | 0.578397153 | 0.699 | 0.567 | 1.43E-25 | Mari |
| allcell | TSTD1 | 1.67E-15 | 0.577788876 | 0.442 | 0.337 | 3.30E-11 | Mari |
| allcell | PSMB8 | 2.11E-43 | 0.577485197 | 0.866 | 0.753 | 4.16E-39 | Mari |
| allcell | PSMG2 | 4.87E-34 | 0.57745272 | 0.711 | 0.577 | 9.59E-30 | Mari |
| allcell | MPV17 | 9.83E-16 | 0.576945253 | 0.41 | 0.304 | 1.94E-11 | Mari |
| allcell | GUK1 | 2.04E-51 | 0.57637478 | 0.895 | 0.794 | 4.02E-47 | Mari |
| allcell | 11-Mar | 9.01E-24 | 0.575857806 | 0.571 | 0.417 | 1.77E-19 | Mari |
| allcell | MRPS34 | 2.24E-24 | 0.575752827 | 0.563 | 0.425 | 4.41E-20 | Mari |
| allcell | RNF166 | 2.59E-10 | 0.574975295 | 0.39 | 0.31 | 5.10E-06 | Mari |
| allcell | TMEM256 | 5.06E-26 | 0.574859977 | 0.641 | 0.524 | 9.96E-22 | Mari |

| | | | | | | | |
|---|---|---|---|---|---|---|---|
| allcell | CLNS1A | 1.03E-23 | 0.574833995 | 0.653 | 0.538 | 2.04E-19 | Mari |
| allcell | PAIP2 | 8.20E-26 | 0.574812457 | 0.72 | 0.597 | 1.61E-21 | Mari |
| allcell | DGCR6L | 1.71E-09 | 0.574767442 | 0.325 | 0.249 | 3.37E-05 | Mari |
| allcell | RNASE6 | 1.56E-16 | 0.574517678 | 0.516 | 0.413 | 3.07E-12 | Mari |
| allcell | MRPL16 | 1.28E-08 | 0.574149698 | 0.319 | 0.249 | 0.00025157 | Mari |
| allcell | SRSF7 | 1.87E-38 | 0.573901854 | 0.929 | 0.821 | 3.68E-34 | Mari |
| allcell | MOB1A | 1.51E-45 | 0.573885245 | 0.869 | 0.759 | 2.98E-41 | Mari |
| allcell | NDUFA8 | 1.03E-11 | 0.573872591 | 0.363 | 0.28 | 2.03E-07 | Mari |
| allcell | EEPD11 | 6.86E-09 | 0.573825907 | 0.26 | 0.19 | 0.000135011 | Mari |
| allcell | PPIH | 9.45E-14 | 0.573171059 | 0.396 | 0.297 | 1.86E-09 | Mari |
| allcell | CD47 | 2.33E-30 | 0.572654675 | 0.773 | 0.637 | 4.58E-26 | Mari |
| allcell | TNFSF10 | 1.78E-08 | 0.57248985 | 0.451 | 0.384 | 0.000350384 | Mari |
| allcell | FUNDC2 | 2.30E-16 | 0.572035032 | 0.475 | 0.369 | 4.52E-12 | Mari |
| allcell | TRAPPC4 | 7.16E-09 | 0.571813461 | 0.308 | 0.236 | 0.000140949 | Mari |
| allcell | DDA1 | 4.28E-08 | 0.571709528 | 0.303 | 0.237 | 0.000842315 | Mari |
| allcell | NAXE | 1.35E-06 | 0.570487953 | 0.27 | 0.217 | 0.026488251 | Mari |
| allcell | UFC1 | 1.10E-33 | 0.570434932 | 0.783 | 0.649 | 2.16E-29 | Mari |
| allcell | FKBP1A | 2.99E-35 | 0.569741968 | 0.802 | 0.686 | 5.89E-31 | Mari |
| allcell | STXBP2 | 9.53E-11 | 0.56955723 | 0.436 | 0.354 | 1.88E-06 | Mari |
| allcell | MEAF6 | 5.63E-14 | 0.569394949 | 0.399 | 0.304 | 1.11E-09 | Mari |
| allcell | ADH5 | 5.35E-09 | 0.569299681 | 0.337 | 0.268 | 0.000105293 | Mari |
| allcell | SCO2 | 2.31E-19 | 0.567915302 | 0.587 | 0.461 | 4.55E-15 | Mari |
| allcell | EAPP | 2.54E-10 | 0.567738614 | 0.348 | 0.272 | 4.99E-06 | Mari |
| allcell | ARL4D | 1.29E-11 | 0.567494317 | 0.275 | 0.185 | 2.55E-07 | Mari |
| allcell | SLC25A11 | 2.71E-12 | 0.566858435 | 0.419 | 0.335 | 5.34E-08 | Mari |
| allcell | TYROBP | 3.87E-30 | 0.566505442 | 0.863 | 0.816 | 7.62E-26 | Mari |
| allcell | OCIAD1 | 5.97E-24 | 0.566490991 | 0.648 | 0.513 | 1.18E-19 | Mari |
| allcell | DCUN1D5 | 1.47E-10 | 0.566302912 | 0.331 | 0.253 | 2.89E-06 | Mari |
| allcell | APEX1 | 2.27E-19 | 0.56596649 | 0.689 | 0.595 | 4.46E-15 | Mari |
| allcell | NDUFA1 | 9.04E-56 | 0.565641247 | 0.872 | 0.748 | 1.78E-51 | Mari |
| allcell | PYCARD | 7.84E-34 | 0.565639594 | 0.816 | 0.737 | 1.54E-29 | Mari |
| allcell | FAM192A | 4.74E-11 | 0.5655272 | 0.402 | 0.322 | 9.34E-07 | Mari |
| allcell | SELENOF | 7.05E-21 | 0.565289564 | 0.662 | 0.557 | 1.39E-16 | Mari |
| allcell | SMIM14 | 5.33E-21 | 0.565176754 | 0.594 | 0.465 | 1.05E-16 | Mari |
| allcell | ARPC3 | 6.80E-88 | 0.56481002 | 0.952 | 0.921 | 1.34E-83 | Mari |
| allcell | ECHDC1 | 2.26E-19 | 0.563894038 | 0.565 | 0.443 | 4.46E-15 | Mari |
| allcell | GABARAP | 1.96E-83 | 0.563128606 | 0.952 | 0.918 | 3.86E-79 | Mari |
| allcell | PARL | 1.65E-13 | 0.56287629 | 0.449 | 0.356 | 3.24E-09 | Mari |
| allcell | TRIM22 | 5.25E-15 | 0.562119245 | 0.601 | 0.511 | 1.03E-10 | Mari |
| allcell | SUMO2 | 1.97E-81 | 0.562064775 | 0.936 | 0.884 | 3.88E-77 | Mari |
| allcell | ITGA41 | 1.11E-27 | 0.560507674 | 0.674 | 0.518 | 2.18E-23 | Mari |
| allcell | LAMTOR4 | 7.90E-45 | 0.55923185 | 0.84 | 0.727 | 1.55E-40 | Mari |
| allcell | TLE5 | 3.14E-25 | 0.558786047 | 0.784 | 0.663 | 6.18E-21 | Mari |
| allcell | GGA21 | 5.42E-10 | 0.558297624 | 0.372 | 0.295 | 1.07E-05 | Mari |
| allcell | TWF2 | 9.45E-29 | 0.558167837 | 0.693 | 0.569 | 1.86E-24 | Mari |
| allcell | PUF60 | 1.95E-15 | 0.55775084 | 0.466 | 0.366 | 3.84E-11 | Mari |
| allcell | PEBP1 | 1.04E-28 | 0.55740837 | 0.786 | 0.667 | 2.04E-24 | Mari |
| allcell | TMEM50B | 1.72E-14 | 0.557237504 | 0.397 | 0.292 | 3.38E-10 | Mari |
| allcell | ISOC2 | 2.97E-11 | 0.556712501 | 0.413 | 0.328 | 5.85E-07 | Mari |
| allcell | COX17 | 4.32E-33 | 0.556219468 | 0.748 | 0.598 | 8.51E-29 | Mari |
| allcell | LRPAP1 | 1.44E-18 | 0.555967392 | 0.506 | 0.387 | 2.83E-14 | Mari |
| allcell | PSMD13 | 2.39E-18 | 0.555788502 | 0.554 | 0.451 | 4.70E-14 | Mari |
| allcell | POLR2F | 2.91E-15 | 0.555674738 | 0.442 | 0.338 | 5.73E-11 | Mari |
| allcell | PDAP1 | 1.75E-17 | 0.555579115 | 0.51 | 0.4 | 3.44E-13 | Mari |
| allcell | ATOX1 | 8.26E-22 | 0.554750438 | 0.813 | 0.721 | 1.63E-17 | Mari |
| allcell | PARVG | 9.37E-21 | 0.554342205 | 0.567 | 0.446 | 1.85E-16 | Mari |

| | | | | | | | |
|---|---|---|---|---|---|---|---|
| allcell | BROX | 3.41E-08 | 0.554022081 | 0.315 | 0.252 | 0.00067176 | Mari |
| allcell | AP3S1 | 9.41E-11 | 0.553756281 | 0.478 | 0.397 | 1.85E-06 | Mari |
| allcell | HEBP2 | 5.57E-15 | 0.553220167 | 0.451 | 0.344 | 1.10E-10 | Mari |
| allcell | SMIM12 | 9.99E-10 | 0.552997813 | 0.372 | 0.3 | 1.97E-05 | Mari |
| allcell | CAPZA2 | 6.51E-28 | 0.552978179 | 0.69 | 0.57 | 1.28E-23 | Mari |
| allcell | DUT | 1.31E-13 | 0.552585612 | 0.505 | 0.419 | 2.58E-09 | Mari |
| allcell | MRPL15 | 1.23E-10 | 0.552561787 | 0.329 | 0.249 | 2.43E-06 | Mari |
| allcell | DCTPP1 | 4.46E-11 | 0.552333955 | 0.402 | 0.323 | 8.79E-07 | Mari |
| allcell | NAA38 | 8.75E-22 | 0.552186354 | 0.587 | 0.459 | 1.72E-17 | Mari |
| allcell | HMGN2 | 1.28E-37 | 0.55177543 | 0.858 | 0.748 | 2.53E-33 | Mari |
| allcell | APP | 2.26E-08 | 0.551116203 | 0.506 | 0.45 | 0.000444198 | Mari |
| allcell | COMMD9 | 1.30E-13 | 0.550910501 | 0.433 | 0.337 | 2.56E-09 | Mari |
| allcell | PSMB3 | 5.92E-45 | 0.55062983 | 0.857 | 0.739 | 1.17E-40 | Mari |
| allcell | VMA21 | 1.92E-17 | 0.550222435 | 0.53 | 0.419 | 3.77E-13 | Mari |
| allcell | SNHG12 | 1.03E-11 | 0.549949701 | 0.456 | 0.377 | 2.03E-07 | Mari |
| allcell | NPC2 | 5.01E-19 | 0.549533598 | 0.844 | 0.782 | 9.86E-15 | Mari |
| allcell | ABI3 | 4.27E-07 | 0.549411996 | 0.363 | 0.303 | 0.008413448 | Mari |
| allcell | NDUFV2 | 1.62E-59 | 0.548223242 | 0.901 | 0.805 | 3.20E-55 | Mari |
| allcell | CASP4 | 3.95E-11 | 0.547945596 | 0.52 | 0.454 | 7.78E-07 | Mari |
| allcell | TMEM50A | 4.15E-35 | 0.54682892 | 0.809 | 0.691 | 8.16E-31 | Mari |
| allcell | MRPS33 | 1.84E-07 | 0.54651991 | 0.366 | 0.309 | 0.003628008 | Mari |
| allcell | FUOM | 7.76E-15 | 0.545556758 | 0.541 | 0.445 | 1.53E-10 | Mari |
| allcell | ECHS1 | 5.29E-19 | 0.544747871 | 0.561 | 0.448 | 1.04E-14 | Mari |
| allcell | RPAIN | 5.07E-14 | 0.544325752 | 0.43 | 0.329 | 9.97E-10 | Mari |
| allcell | TCTN3 | 2.35E-07 | 0.544065453 | 0.316 | 0.252 | 0.004616766 | Mari |
| allcell | TCIRG1 | 2.75E-09 | 0.543825344 | 0.383 | 0.308 | 5.41E-05 | Mari |
| allcell | HSCB | 1.36E-06 | 0.54316069 | 0.259 | 0.2 | 0.026772377 | Mari |
| allcell | MYL6 | 6.54E-76 | 0.541587772 | 0.972 | 0.953 | 1.29E-71 | Mari |
| allcell | SRA1 | 2.73E-23 | 0.541484288 | 0.605 | 0.486 | 5.37E-19 | Mari |
| allcell | ZNF706 | 1.51E-30 | 0.5410916 | 0.79 | 0.671 | 2.96E-26 | Mari |
| allcell | MESD | 4.12E-14 | 0.540426051 | 0.494 | 0.399 | 8.10E-10 | Mari |
| allcell | VKORC1 | 4.11E-13 | 0.540039763 | 0.42 | 0.326 | 8.08E-09 | Mari |
| allcell | COMT | 2.46E-25 | 0.539767922 | 0.687 | 0.569 | 4.83E-21 | Mari |
| allcell | ZSCAN16-AS1 | 7.23E-07 | 0.539017933 | 0.271 | 0.211 | 0.014238326 | Mari |
| allcell | COX6C | 6.30E-66 | 0.538935272 | 0.921 | 0.835 | 1.24E-61 | Mari |
| allcell | ATP5F1C | 2.33E-42 | 0.538685781 | 0.855 | 0.742 | 4.58E-38 | Mari |
| allcell | PPP1R11 | 5.22E-07 | 0.538457266 | 0.287 | 0.229 | 0.010265874 | Mari |
| allcell | NME1 | 1.88E-16 | 0.538136658 | 0.514 | 0.412 | 3.70E-12 | Mari |
| allcell | UBE2J1 | 1.32E-12 | 0.53802249 | 0.623 | 0.547 | 2.60E-08 | Mari |
| allcell | DTNBP1 | 1.51E-11 | 0.537063511 | 0.531 | 0.441 | 2.98E-07 | Mari |
| allcell | BCL7C | 1.51E-10 | 0.536914138 | 0.432 | 0.356 | 2.96E-06 | Mari |
| allcell | SRSF5 | 2.18E-40 | 0.535377254 | 0.839 | 0.681 | 4.29E-36 | Mari |
| allcell | TALDO1 | 3.56E-36 | 0.535360444 | 0.827 | 0.721 | 7.02E-32 | Mari |
| allcell | ISG15 | 2.91E-21 | 0.534912797 | 0.66 | 0.547 | 5.73E-17 | Mari |
| allcell | UXT | 1.24E-40 | 0.534640591 | 0.857 | 0.755 | 2.43E-36 | Mari |
| allcell | LSM7 | 4.73E-36 | 0.534157252 | 0.814 | 0.711 | 9.32E-32 | Mari |
| allcell | CAPG | 1.43E-23 | 0.533826821 | 0.776 | 0.693 | 2.82E-19 | Mari |
| allcell | GLO1 | 1.09E-11 | 0.533334865 | 0.409 | 0.324 | 2.15E-07 | Mari |
| allcell | PSME2 | 6.12E-32 | 0.533120204 | 0.911 | 0.852 | 1.20E-27 | Mari |
| allcell | NDUFA7 | 1.55E-25 | 0.532113947 | 0.7 | 0.587 | 3.06E-21 | Mari |
| allcell | COPS5 | 7.03E-12 | 0.531965879 | 0.415 | 0.331 | 1.38E-07 | Mari |
| allcell | BUD31 | 5.32E-25 | 0.531689122 | 0.672 | 0.551 | 1.05E-20 | Mari |
| allcell | ATP5MPL | 3.72E-33 | 0.529554379 | 0.797 | 0.642 | 7.32E-29 | Mari |
| allcell | NXT1 | 2.40E-22 | 0.529176021 | 0.511 | 0.369 | 4.73E-18 | Mari |
| allcell | SLC25A26 | 1.50E-07 | 0.528691351 | 0.295 | 0.23 | 0.002951854 | Mari |
| allcell | COX8A | 1.33E-57 | 0.528446141 | 0.909 | 0.841 | 2.62E-53 | Mari |

| | | | | | | | |
|---|---|---|---|---|---|---|---|
| allcell | MITD11 | 5.91E-07 | 0.527598418 | 0.276 | 0.218 | 0.011634882 | Mari |
| allcell | ZNF593 | 3.66E-13 | 0.527224204 | 0.354 | 0.26 | 7.20E-09 | Mari |
| allcell | TSSC4 | 1.42E-07 | 0.526788651 | 0.273 | 0.211 | 0.002796063 | Mari |
| allcell | PHYKPL | 4.47E-09 | 0.526217804 | 0.389 | 0.323 | 8.79E-05 | Mari |
| allcell | SNRPD3 | 4.37E-22 | 0.526173467 | 0.644 | 0.529 | 8.60E-18 | Mari |
| allcell | SNRPD1 | 2.86E-26 | 0.524993129 | 0.702 | 0.57 | 5.64E-22 | Mari |
| allcell | LMAN1 | 4.62E-08 | 0.524633788 | 0.377 | 0.311 | 0.000909441 | Mari |
| allcell | MRPL34 | 9.61E-11 | 0.524524162 | 0.445 | 0.366 | 1.89E-06 | Mari |
| allcell | PRMT1 | 1.26E-19 | 0.524340279 | 0.629 | 0.513 | 2.48E-15 | Mari |
| allcell | ZNHIT3 | 5.59E-08 | 0.524331374 | 0.347 | 0.282 | 0.001099709 | Mari |
| allcell | MRPS21 | 2.92E-33 | 0.524214979 | 0.762 | 0.636 | 5.75E-29 | Mari |
| allcell | C1QBP | 3.00E-30 | 0.524207767 | 0.768 | 0.648 | 5.90E-26 | Mari |
| allcell | ACAA1 | 6.86E-23 | 0.523993728 | 0.657 | 0.545 | 1.35E-18 | Mari |
| allcell | TRAPPC5 | 1.87E-23 | 0.523808517 | 0.668 | 0.558 | 3.69E-19 | Mari |
| allcell | RUNX21 | 2.32E-19 | 0.523665662 | 0.399 | 0.269 | 4.57E-15 | Mari |
| allcell | ATP5MC3 | 6.70E-52 | 0.522936657 | 0.902 | 0.806 | 1.32E-47 | Mari |
| allcell | TMEM258 | 5.94E-45 | 0.522598866 | 0.865 | 0.755 | 1.17E-40 | Mari |
| allcell | RNF187 | 9.14E-07 | 0.522591004 | 0.309 | 0.25 | 0.017982561 | Mari |
| allcell | PPCDC1 | 7.17E-16 | 0.521506063 | 0.341 | 0.233 | 1.41E-11 | Mari |
| allcell | PFN1 | 4.62E-64 | 0.520915044 | 0.977 | 0.941 | 9.10E-60 | Mari |
| allcell | SLC15A41 | 9.86E-09 | 0.520879929 | 0.44 | 0.374 | 0.000194153 | Mari |
| allcell | BBIP1 | 1.59E-06 | 0.520277021 | 0.25 | 0.196 | 0.031259709 | Mari |
| allcell | NDUFA12 | 3.90E-30 | 0.519621858 | 0.773 | 0.64 | 7.68E-26 | Mari |
| allcell | RMDN1 | 4.49E-08 | 0.51889863 | 0.361 | 0.295 | 0.000884653 | Mari |
| allcell | ATP6V1G1 | 5.75E-39 | 0.518119821 | 0.866 | 0.759 | 1.13E-34 | Mari |
| allcell | PIK3CG | 2.25E-07 | 0.517172255 | 0.375 | 0.31 | 0.004424937 | Mari |
| allcell | ERCC1 | 6.01E-14 | 0.515721951 | 0.575 | 0.494 | 1.18E-09 | Mari |
| allcell | DHPS | 9.24E-11 | 0.515454909 | 0.396 | 0.313 | 1.82E-06 | Mari |
| allcell | SHKBP1 | 1.04E-10 | 0.515043183 | 0.392 | 0.311 | 2.05E-06 | Mari |
| allcell | PPP4C | 7.69E-23 | 0.514863989 | 0.648 | 0.532 | 1.51E-18 | Mari |
| allcell | ASPH | 8.81E-09 | 0.514374442 | 0.403 | 0.327 | 0.000173418 | Mari |
| allcell | SYPL1 | 1.17E-06 | 0.514250416 | 0.328 | 0.275 | 0.023048182 | Mari |
| allcell | NDUFS7 | 1.76E-30 | 0.514232811 | 0.73 | 0.586 | 3.47E-26 | Mari |
| allcell | HAGH | 4.14E-08 | 0.5120685 | 0.38 | 0.318 | 0.000814038 | Mari |
| allcell | CD99 | 3.75E-11 | 0.511860821 | 0.662 | 0.595 | 7.39E-07 | Mari |
| allcell | FHIT | 1.11E-08 | 0.511259269 | 0.301 | 0.224 | 0.000219081 | Mari |
| allcell | MPC1 | 4.26E-16 | 0.510719153 | 0.587 | 0.484 | 8.38E-12 | Mari |
| allcell | RRP7A | 1.32E-09 | 0.510706173 | 0.395 | 0.323 | 2.60E-05 | Mari |
| allcell | DDRGK1 | 2.90E-10 | 0.510670424 | 0.405 | 0.329 | 5.70E-06 | Mari |
| allcell | SERPINB1 | 4.27E-23 | 0.510647345 | 0.86 | 0.791 | 8.41E-19 | Mari |
| allcell | PRPF31 | 6.68E-08 | 0.510453308 | 0.333 | 0.269 | 0.001314709 | Mari |
| allcell | PIK3AP1 | 3.98E-14 | 0.509485254 | 0.445 | 0.34 | 7.83E-10 | Mari |
| allcell | SMDT1 | 2.74E-37 | 0.509113573 | 0.863 | 0.756 | 5.39E-33 | Mari |
| allcell | RAB34 | 5.70E-12 | 0.508804022 | 0.462 | 0.368 | 1.12E-07 | Mari |
| allcell | ACTR10 | 6.50E-11 | 0.508666052 | 0.437 | 0.357 | 1.28E-06 | Mari |
| allcell | RNF7 | 3.43E-34 | 0.50832063 | 0.78 | 0.647 | 6.74E-30 | Mari |
| allcell | GSTK1 | 9.89E-35 | 0.507968177 | 0.819 | 0.72 | 1.95E-30 | Mari |
| allcell | OSGEP | 7.84E-10 | 0.507837611 | 0.358 | 0.282 | 1.54E-05 | Mari |
| allcell | TRNAU1AP | 4.38E-08 | 0.506726263 | 0.299 | 0.234 | 0.000861249 | Mari |
| allcell | UBE2I | 1.19E-22 | 0.504877586 | 0.735 | 0.634 | 2.35E-18 | Mari |
| allcell | ZNRD2 | 7.28E-10 | 0.504589559 | 0.346 | 0.272 | 1.43E-05 | Mari |
| allcell | VPS28 | 1.16E-29 | 0.504373242 | 0.772 | 0.661 | 2.29E-25 | Mari |
| allcell | STAT2 | 3.81E-09 | 0.503842705 | 0.373 | 0.302 | 7.51E-05 | Mari |
| allcell | MRPS36 | 2.23E-21 | 0.503705393 | 0.569 | 0.436 | 4.39E-17 | Mari |
| allcell | ERVK3-1 | 1.45E-11 | 0.503307543 | 0.415 | 0.327 | 2.85E-07 | Mari |
| allcell | HINT1 | 3.22E-46 | 0.502704397 | 0.943 | 0.912 | 6.35E-42 | Mari |

| | | | | | | | |
|---|---|---|---|---|---|---|---|
| allcell | ATXN10 | 1.31E-11 | 0.502685573 | 0.434 | 0.352 | 2.59E-07 | Mari |
| allcell | DNAJC15 | 1.51E-31 | 0.502394276 | 0.777 | 0.648 | 2.97E-27 | Mari |
| allcell | DPM3 | 2.01E-08 | 0.502068379 | 0.369 | 0.298 | 0.000395336 | Mari |
| allcell | NQO2 | 4.89E-22 | 0.501976591 | 0.567 | 0.441 | 9.63E-18 | Mari |
| allcell | ELOB | 7.41E-47 | 0.501822163 | 0.889 | 0.803 | 1.46E-42 | Mari |
| allcell | WDR74 | 8.03E-07 | 0.501732693 | 0.347 | 0.287 | 0.015814352 | Mari |
| allcell | IRAK4 | 1.32E-07 | 0.501626003 | 0.338 | 0.275 | 0.002599235 | Mari |
| allcell | NDUFA2 | 4.20E-31 | 0.500311285 | 0.755 | 0.618 | 8.27E-27 | Mari |
| allcell | NPHS1 | 5.31E-59 | 2.81251785 | 0.3 | 0.124 | 1.04E-54 | Tb |
| allcell | GCNT2 | 6.80E-40 | 2.145110032 | 0.272 | 0.122 | 1.34E-35 | Tb |
| allcell | SLC35F1 | 3.42E-75 | 1.981334723 | 0.366 | 0.145 | 6.72E-71 | Tb |
| allcell | S100A7 | 2.17E-115 | 1.923481291 | 0.411 | 0.118 | 4.27E-111 | Tb |
| allcell | ARL17A | 4.66E-80 | 1.920700459 | 0.307 | 0.093 | 9.18E-76 | Tb |
| allcell | PSMD5 | 2.57E-38 | 1.808933765 | 0.3 | 0.144 | 5.06E-34 | Tb |
| allcell | KRTDAP | 2.89E-32 | 1.800319102 | 0.265 | 0.132 | 5.69E-28 | Tb |
| allcell | FBP1 | 1.88E-10 | 1.778695766 | 0.395 | 0.311 | 3.69E-06 | Tb |
| allcell | S100A2 | 1.64E-107 | 1.769736524 | 0.365 | 0.097 | 3.24E-103 | Tb |
| allcell | HOMER1 | 6.95E-90 | 1.732081034 | 0.575 | 0.294 | 1.37E-85 | Tb |
| allcell | AQP3 | 3.00E-65 | 1.651992077 | 0.281 | 0.092 | 5.91E-61 | Tb |
| allcell | KRT6A | 9.84E-107 | 1.639178607 | 0.372 | 0.1 | 1.94E-102 | Tb |
| allcell | CCR6 | 5.64E-44 | 1.594604193 | 0.272 | 0.114 | 1.11E-39 | Tb |
| allcell | TAF4B | 9.72E-57 | 1.507025072 | 0.442 | 0.221 | 1.91E-52 | Tb |
| allcell | RHOH | 3.77E-82 | 1.388056185 | 0.586 | 0.31 | 7.43E-78 | Tb |
| allcell | ZNF609 | 1.94E-66 | 1.378191472 | 0.455 | 0.217 | 3.81E-62 | Tb |
| allcell | KRT17 | 9.98E-70 | 1.375176199 | 0.28 | 0.083 | 1.96E-65 | Tb |
| allcell | PHTF1 | 7.57E-47 | 1.35826151 | 0.395 | 0.201 | 1.49E-42 | Tb |
| allcell | MCF2L2 | 2.76E-41 | 1.357118494 | 0.418 | 0.235 | 5.43E-37 | Tb |
| allcell | RGS13 | 4.25E-44 | 1.34677274 | 0.361 | 0.181 | 8.37E-40 | Tb |
| allcell | FAM222B | 1.89E-52 | 1.34279309 | 0.295 | 0.115 | 3.72E-48 | Tb |
| allcell | CLEC16A | 7.24E-64 | 1.341060811 | 0.528 | 0.279 | 1.42E-59 | Tb |
| allcell | TBC1D32 | 1.59E-43 | 1.340750861 | 0.275 | 0.113 | 3.13E-39 | Tb |
| allcell | FAM118A | 7.02E-54 | 1.332111269 | 0.431 | 0.217 | 1.38E-49 | Tb |
| allcell | SFN | 6.33E-49 | 1.298165922 | 0.302 | 0.123 | 1.25E-44 | Tb |
| allcell | L3MBTL4 | 2.84E-22 | 1.297324638 | 0.304 | 0.179 | 5.59E-18 | Tb |
| allcell | IGKC | 6.36E-15 | 1.293682541 | 0.393 | 0.282 | 1.25E-10 | Tb |
| allcell | GAB1 | 8.52E-31 | 1.274970594 | 0.355 | 0.201 | 1.68E-26 | Tb |
| allcell | NEK1 | 9.49E-26 | 1.259723419 | 0.336 | 0.195 | 1.87E-21 | Tb |
| allcell | SLC33A1 | 3.23E-38 | 1.244753179 | 0.281 | 0.125 | 6.35E-34 | Tb |
| allcell | ADAM19 | 4.44E-82 | 1.219355118 | 0.663 | 0.377 | 8.75E-78 | Tb |
| allcell | NCR3LG1 | 5.83E-33 | 1.217723538 | 0.253 | 0.116 | 1.15E-28 | Tb |
| allcell | S100A8 | 1.56E-141 | 1.201221103 | 0.638 | 0.243 | 3.07E-137 | Tb |
| allcell | XIST | 5.57E-101 | 1.186932054 | 0.619 | 0.284 | 1.10E-96 | Tb |
| allcell | NLK | 3.04E-49 | 1.177793198 | 0.321 | 0.136 | 5.99E-45 | Tb |
| allcell | GYS1 | 4.37E-44 | 1.177170067 | 0.303 | 0.131 | 8.60E-40 | Tb |
| allcell | NARS2 | 1.78E-28 | 1.17702859 | 0.327 | 0.18 | 3.50E-24 | Tb |
| allcell | CLIC4 | 8.90E-38 | 1.170233096 | 0.548 | 0.374 | 1.75E-33 | Tb |
| allcell | INHBA | 1.47E-09 | 1.169449916 | 0.285 | 0.212 | 2.89E-05 | Tb |
| allcell | KRT6B | 2.24E-61 | 1.16230466 | 0.261 | 0.078 | 4.41E-57 | Tb |
| allcell | GBF1 | 7.93E-69 | 1.160018125 | 0.499 | 0.241 | 1.56E-64 | Tb |
| allcell | BRCA2 | 3.29E-40 | 1.157821232 | 0.446 | 0.254 | 6.48E-36 | Tb |
| allcell | PHLPP1 | 2.38E-51 | 1.149962232 | 0.549 | 0.335 | 4.69E-47 | Tb |
| allcell | CYTH3 | 4.75E-29 | 1.146532829 | 0.264 | 0.129 | 9.36E-25 | Tb |
| allcell | KRT14 | 4.35E-121 | 1.145999854 | 0.61 | 0.241 | 8.56E-117 | Tb |
| allcell | RASAL2 | 4.80E-34 | 1.145954887 | 0.376 | 0.215 | 9.44E-30 | Tb |
| allcell | SUCNR1 | 4.39E-37 | 1.145887722 | 0.32 | 0.157 | 8.63E-33 | Tb |
| allcell | PRAG1 | 2.44E-36 | 1.145882197 | 0.259 | 0.113 | 4.79E-32 | Tb |

| | | | | | | | |
|---|---|---|---|---|---|---|---|
| allcell | STAT4 | 9.69E-47 | 1.143277509 | 0.506 | 0.3 | 1.91E-42 | Tb |
| allcell | PFKP | 1.21E-37 | 1.139913647 | 0.413 | 0.236 | 2.38E-33 | Tb |
| allcell | MOB3B | 1.59E-49 | 1.137632431 | 0.525 | 0.312 | 3.12E-45 | Tb |
| allcell | MT-ATP6 | 6.01E-159 | 1.136149733 | 0.991 | 0.897 | 1.18E-154 | Tb |
| allcell | PATL2 | 3.74E-44 | 1.130614217 | 0.286 | 0.118 | 7.36E-40 | Tb |
| allcell | CD109 | 1.25E-12 | 1.128001151 | 0.3 | 0.211 | 2.46E-08 | Tb |
| allcell | PDCD1LG2 | 6.42E-30 | 1.120880662 | 0.278 | 0.14 | 1.26E-25 | Tb |
| allcell | UBAP2 | 2.91E-54 | 1.119882991 | 0.442 | 0.217 | 5.73E-50 | Tb |
| allcell | DENND5B | 8.98E-32 | 1.119171569 | 0.299 | 0.152 | 1.77E-27 | Tb |
| allcell | PLEKHG2 | 8.79E-36 | 1.109121583 | 0.411 | 0.241 | 1.73E-31 | Tb |
| allcell | TFCP2 | 9.99E-52 | 1.103410859 | 0.41 | 0.201 | 1.97E-47 | Tb |
| allcell | RAP1GAP2 | 1.04E-27 | 1.099089722 | 0.26 | 0.13 | 2.04E-23 | Tb |
| allcell | RUNX2 | 1.96E-15 | 1.098945048 | 0.378 | 0.268 | 3.86E-11 | Tb |
| allcell | AZIN1-AS1 | 2.52E-29 | 1.09600538 | 0.297 | 0.155 | 4.95E-25 | Tb |
| allcell | LRCH3 | 1.12E-55 | 1.08522305 | 0.494 | 0.257 | 2.21E-51 | Tb |
| allcell | CREB3L2 | 5.24E-26 | 1.079980016 | 0.311 | 0.174 | 1.03E-21 | Tb |
| allcell | DUSP16 | 1.45E-40 | 1.078057317 | 0.359 | 0.178 | 2.85E-36 | Tb |
| allcell | DLEU2 | 2.38E-64 | 1.077658129 | 0.769 | 0.567 | 4.69E-60 | Tb |
| allcell | PPARD | 5.37E-40 | 1.072692816 | 0.474 | 0.279 | 1.06E-35 | Tb |
| allcell | EEF2K | 5.16E-42 | 1.072529702 | 0.365 | 0.18 | 1.02E-37 | Tb |
| allcell | KIF1B | 2.15E-53 | 1.072281106 | 0.542 | 0.318 | 4.23E-49 | Tb |
| allcell | MT-CO3 | 7.42E-117 | 1.069938541 | 0.978 | 0.911 | 1.46E-112 | Tb |
| allcell | LRRK1 | 2.51E-59 | 1.066001562 | 0.6 | 0.361 | 4.94E-55 | Tb |
| allcell | SPECC1L | 8.96E-45 | 1.065069884 | 0.387 | 0.192 | 1.76E-40 | Tb |
| allcell | PRKCA | 3.53E-51 | 1.065067153 | 0.492 | 0.272 | 6.95E-47 | Tb |
| allcell | CCDC138 | 1.05E-41 | 1.065028116 | 0.418 | 0.224 | 2.07E-37 | Tb |
| allcell | JMY | 2.92E-34 | 1.062585045 | 0.503 | 0.322 | 5.76E-30 | Tb |
| allcell | MCU | 2.07E-29 | 1.059693295 | 0.318 | 0.17 | 4.07E-25 | Tb |
| allcell | ZNF10 | 2.22E-49 | 1.058234886 | 0.446 | 0.229 | 4.37E-45 | Tb |
| allcell | TOGARAM1 | 4.21E-39 | 1.056850057 | 0.282 | 0.125 | 8.29E-35 | Tb |
| allcell | ENPP2 | 1.38E-39 | 1.056747898 | 0.407 | 0.219 | 2.72E-35 | Tb |
| allcell | TMEM184C | 3.36E-43 | 1.05484298 | 0.383 | 0.193 | 6.62E-39 | Tb |
| allcell | CARD11 | 1.42E-32 | 1.053448251 | 0.389 | 0.223 | 2.79E-28 | Tb |
| allcell | ANKRD28 | 1.93E-28 | 1.052900053 | 0.407 | 0.247 | 3.81E-24 | Tb |
| allcell | EBF4 | 4.00E-31 | 1.051829052 | 0.265 | 0.128 | 7.86E-27 | Tb |
| allcell | ARHGAP25 | 2.41E-40 | 1.049113217 | 0.516 | 0.32 | 4.75E-36 | Tb |
| allcell | CLEC2D | 2.83E-20 | 1.048716671 | 0.253 | 0.141 | 5.57E-16 | Tb |
| allcell | CASP10 | 1.20E-27 | 1.039011945 | 0.262 | 0.132 | 2.36E-23 | Tb |
| allcell | IL1RAP | 7.87E-32 | 1.036824868 | 0.293 | 0.146 | 1.55E-27 | Tb |
| allcell | PALD1 | 3.27E-49 | 1.034494545 | 0.383 | 0.179 | 6.43E-45 | Tb |
| allcell | SFMBT2 | 5.28E-59 | 1.029381231 | 0.635 | 0.384 | 1.04E-54 | Tb |
| allcell | KRT16 | 4.83E-67 | 1.028505044 | 0.351 | 0.123 | 9.51E-63 | Tb |
| allcell | DIPK1A | 1.41E-31 | 1.025002946 | 0.351 | 0.193 | 2.77E-27 | Tb |
| allcell | NAA25 | 1.37E-21 | 1.019515986 | 0.33 | 0.198 | 2.70E-17 | Tb |
| allcell | ZNF516 | 2.85E-46 | 1.019412106 | 0.519 | 0.306 | 5.60E-42 | Tb |
| allcell | MGA | 1.12E-32 | 1.018908472 | 0.318 | 0.163 | 2.21E-28 | Tb |
| allcell | RPRD2 | 1.41E-25 | 1.017316588 | 0.267 | 0.14 | 2.78E-21 | Tb |
| allcell | CEP192 | 1.11E-38 | 1.016303543 | 0.311 | 0.146 | 2.19E-34 | Tb |
| allcell | SMG6 | 3.61E-45 | 1.016106777 | 0.388 | 0.19 | 7.10E-41 | Tb |
| allcell | NSMCE2 | 9.07E-67 | 1.009968912 | 0.678 | 0.418 | 1.78E-62 | Tb |
| allcell | CLCN3 | 3.58E-25 | 1.005874127 | 0.355 | 0.208 | 7.04E-21 | Tb |
| allcell | CBLB | 6.13E-41 | 1.005300284 | 0.411 | 0.226 | 1.21E-36 | Tb |
| allcell | RASA1 | 1.45E-42 | 1.002037116 | 0.387 | 0.194 | 2.86E-38 | Tb |
| allcell | WDR37 | 1.86E-50 | 1.000243826 | 0.476 | 0.252 | 3.67E-46 | Tb |
| allcell | N4BP2 | 6.36E-45 | 1.000157789 | 0.507 | 0.291 | 1.25E-40 | Tb |
| allcell | MT-CYB | 2.77E-89 | 0.997374237 | 0.963 | 0.887 | 5.46E-85 | Tb |

| | | | | | | | |
|---|---|---|---|---|---|---|---|
| allcell | UST | 3.71E-44 | 0.992180694 | 0.615 | 0.419 | 7.31E-40 | Tb |
| allcell | ARID2 | 6.33E-50 | 0.99172258 | 0.612 | 0.39 | 1.25E-45 | Tb |
| allcell | PLPP1 | 3.39E-26 | 0.989673522 | 0.371 | 0.219 | 6.68E-22 | Tb |
| allcell | BMPR2 | 4.12E-43 | 0.987702276 | 0.43 | 0.232 | 8.11E-39 | Tb |
| allcell | SPATA13 | 4.56E-29 | 0.984520167 | 0.307 | 0.162 | 8.98E-25 | Tb |
| allcell | PHLPP2 | 6.33E-27 | 0.982012799 | 0.295 | 0.157 | 1.25E-22 | Tb |
| allcell | CRY1 | 1.59E-27 | 0.980246765 | 0.297 | 0.16 | 3.14E-23 | Tb |
| allcell | CTDP1 | 5.77E-45 | 0.979449284 | 0.376 | 0.183 | 1.14E-40 | Tb |
| allcell | LRRC23 | 9.42E-34 | 0.978653328 | 0.421 | 0.246 | 1.85E-29 | Tb |
| allcell | DENND4A | 2.30E-68 | 0.975154906 | 0.852 | 0.67 | 4.52E-64 | Tb |
| allcell | B3GNT5 | 3.17E-28 | 0.974605434 | 0.545 | 0.386 | 6.24E-24 | Tb |
| allcell | P2RY14 | 1.42E-44 | 0.973971583 | 0.614 | 0.4 | 2.79E-40 | Tb |
| allcell | SPG11 | 6.01E-46 | 0.972873629 | 0.498 | 0.287 | 1.18E-41 | Tb |
| allcell | FUT8 | 1.67E-23 | 0.971047155 | 0.323 | 0.19 | 3.29E-19 | Tb |
| allcell | IQCG | 1.68E-27 | 0.970324771 | 0.26 | 0.131 | 3.30E-23 | Tb |
| allcell | GALNT11 | 1.04E-29 | 0.970213652 | 0.341 | 0.184 | 2.05E-25 | Tb |
| allcell | SPRED1 | 1.45E-21 | 0.969420958 | 0.275 | 0.156 | 2.84E-17 | Tb |
| allcell | THADA | 1.11E-37 | 0.96921633 | 0.378 | 0.197 | 2.19E-33 | Tb |
| allcell | ZNF366 | 3.25E-30 | 0.965501315 | 0.346 | 0.191 | 6.39E-26 | Tb |
| allcell | STIM1 | 1.38E-40 | 0.963850072 | 0.387 | 0.2 | 2.71E-36 | Tb |
| allcell | KSR1 | 3.32E-28 | 0.961873069 | 0.282 | 0.147 | 6.54E-24 | Tb |
| allcell | PAG1 | 7.65E-33 | 0.959970596 | 0.491 | 0.309 | 1.51E-28 | Tb |
| allcell | PHF19 | 4.26E-14 | 0.959067262 | 0.408 | 0.305 | 8.38E-10 | Tb |
| allcell | WWOX | 1.50E-25 | 0.958171208 | 0.395 | 0.249 | 2.94E-21 | Tb |
| allcell | AXL | 8.32E-31 | 0.958112195 | 0.4 | 0.237 | 1.64E-26 | Tb |
| allcell | RALGAPA1 | 5.53E-42 | 0.957338076 | 0.656 | 0.449 | 1.09E-37 | Tb |
| allcell | CEP83 | 4.61E-39 | 0.95562435 | 0.469 | 0.272 | 9.07E-35 | Tb |
| allcell | KDM6A | 1.26E-66 | 0.955416955 | 0.662 | 0.406 | 2.48E-62 | Tb |
| allcell | ARIH1 | 7.39E-67 | 0.95467437 | 0.861 | 0.667 | 1.46E-62 | Tb |
| allcell | RASGRP3 | 3.23E-31 | 0.953046942 | 0.387 | 0.225 | 6.36E-27 | Tb |
| allcell | MED12L | 4.10E-28 | 0.950543421 | 0.285 | 0.146 | 8.07E-24 | Tb |
| allcell | RAB30 | 1.06E-40 | 0.950219795 | 0.449 | 0.254 | 2.09E-36 | Tb |
| allcell | ABL1 | 1.07E-33 | 0.949901468 | 0.373 | 0.208 | 2.10E-29 | Tb |
| allcell | SENP2 | 4.98E-32 | 0.9490928 | 0.281 | 0.135 | 9.80E-28 | Tb |
| allcell | GALC | 2.17E-27 | 0.948747136 | 0.33 | 0.186 | 4.27E-23 | Tb |
| allcell | MX2 | 7.25E-20 | 0.948012483 | 0.457 | 0.325 | 1.43E-15 | Tb |
| allcell | HDAC4 | 3.14E-38 | 0.94532414 | 0.362 | 0.182 | 6.18E-34 | Tb |
| allcell | MSI2 | 1.07E-28 | 0.942545544 | 0.355 | 0.202 | 2.11E-24 | Tb |
| allcell | TBC1D17 | 9.44E-33 | 0.941805137 | 0.263 | 0.121 | 1.86E-28 | Tb |
| allcell | CAMK1D | 4.45E-53 | 0.940498118 | 0.732 | 0.526 | 8.76E-49 | Tb |
| allcell | CD80 | 4.73E-23 | 0.940416432 | 0.434 | 0.294 | 9.31E-19 | Tb |
| allcell | MBTD1 | 3.88E-32 | 0.940109901 | 0.318 | 0.164 | 7.63E-28 | Tb |
| allcell | TASP1 | 5.17E-25 | 0.937203566 | 0.29 | 0.157 | 1.02E-20 | Tb |
| allcell | SLC39A11 | 1.46E-43 | 0.936955031 | 0.406 | 0.208 | 2.88E-39 | Tb |
| allcell | TMTC2 | 3.25E-25 | 0.935142434 | 0.3 | 0.167 | 6.40E-21 | Tb |
| allcell | SCAF8 | 1.17E-48 | 0.934722982 | 0.426 | 0.209 | 2.30E-44 | Tb |
| allcell | UBXN2A | 1.59E-38 | 0.934013349 | 0.497 | 0.304 | 3.14E-34 | Tb |
| allcell | LRRC8D | 2.85E-41 | 0.932763784 | 0.419 | 0.228 | 5.60E-37 | Tb |
| allcell | ST6GAL1 | 2.55E-45 | 0.932072073 | 0.468 | 0.257 | 5.02E-41 | Tb |
| allcell | TOP3A | 1.58E-32 | 0.931743874 | 0.266 | 0.122 | 3.11E-28 | Tb |
| allcell | PPP1R16B | 4.53E-36 | 0.931123828 | 0.43 | 0.248 | 8.91E-32 | Tb |
| allcell | POLR2J3 | 1.21E-39 | 0.929966257 | 0.344 | 0.166 | 2.38E-35 | Tb |
| allcell | MRC1 | 6.13E-14 | 0.928602263 | 0.337 | 0.238 | 1.21E-09 | Tb |
| allcell | NT5C2 | 1.51E-49 | 0.926648566 | 0.54 | 0.303 | 2.98E-45 | Tb |
| allcell | ARAP2 | 3.95E-44 | 0.924594333 | 0.553 | 0.342 | 7.78E-40 | Tb |
| allcell | FAM13A | 2.08E-27 | 0.923854607 | 0.301 | 0.161 | 4.09E-23 | Tb |

| | | | | | | | |
|---|---|---|---|---|---|---|---|
| allcell | VAV1 | 6.78E-38 | 0.923464936 | 0.4 | 0.215 | 1.34E-33 | Tb |
| allcell | CERS5 | 3.35E-36 | 0.922682378 | 0.321 | 0.154 | 6.60E-32 | Tb |
| allcell | AKAP10 | 1.56E-30 | 0.920729357 | 0.27 | 0.13 | 3.08E-26 | Tb |
| allcell | SETBP1 | 5.89E-19 | 0.920291083 | 0.266 | 0.155 | 1.16E-14 | Tb |
| allcell | NFX1 | 1.81E-34 | 0.918284436 | 0.456 | 0.268 | 3.57E-30 | Tb |
| allcell | KLF7 | 9.09E-31 | 0.912605722 | 0.402 | 0.235 | 1.79E-26 | Tb |
| allcell | STEAP1B | 1.23E-18 | 0.910661623 | 0.394 | 0.256 | 2.42E-14 | Tb |
| allcell | SGPL1 | 1.38E-31 | 0.909999254 | 0.302 | 0.151 | 2.72E-27 | Tb |
| allcell | SYTL3 | 1.95E-26 | 0.908965959 | 0.374 | 0.227 | 3.83E-22 | Tb |
| allcell | TSPOAP1 | 1.08E-27 | 0.907179319 | 0.253 | 0.124 | 2.12E-23 | Tb |
| allcell | ZHX2 | 3.53E-44 | 0.906610915 | 0.603 | 0.381 | 6.94E-40 | Tb |
| allcell | CTNND1 | 3.41E-35 | 0.903611649 | 0.482 | 0.291 | 6.71E-31 | Tb |
| allcell | CMSS1 | 4.96E-12 | 0.901827615 | 0.42 | 0.307 | 9.77E-08 | Tb |
| allcell | GMDS-DT | 2.62E-28 | 0.901470541 | 0.383 | 0.232 | 5.16E-24 | Tb |
| allcell | USP25 | 7.96E-34 | 0.901102779 | 0.459 | 0.276 | 1.57E-29 | Tb |
| allcell | KRIT1 | 3.68E-41 | 0.898618399 | 0.362 | 0.175 | 7.25E-37 | Tb |
| allcell | SLC43A2 | 2.40E-36 | 0.897994747 | 0.547 | 0.355 | 4.72E-32 | Tb |
| allcell | ZNF33A | 2.34E-30 | 0.897001434 | 0.373 | 0.21 | 4.60E-26 | Tb |
| allcell | SESN1 | 1.37E-24 | 0.89688921 | 0.321 | 0.181 | 2.70E-20 | Tb |
| allcell | FAF2 | 6.64E-34 | 0.896828378 | 0.304 | 0.147 | 1.31E-29 | Tb |
| allcell | VPS13B | 2.65E-54 | 0.893049117 | 0.607 | 0.369 | 5.22E-50 | Tb |
| allcell | ADAP2 | 2.62E-27 | 0.892168872 | 0.361 | 0.208 | 5.16E-23 | Tb |
| allcell | BCL11A | 8.92E-51 | 0.892022324 | 0.659 | 0.431 | 1.76E-46 | Tb |
| allcell | STX17-AS1 | 2.30E-24 | 0.891381408 | 0.467 | 0.308 | 4.52E-20 | Tb |
| allcell | SMYD3 | 4.99E-23 | 0.89019075 | 0.341 | 0.208 | 9.82E-19 | Tb |
| allcell | FTX | 1.05E-60 | 0.889706637 | 0.596 | 0.335 | 2.07E-56 | Tb |
| allcell | SRGAP2 | 4.28E-59 | 0.886970288 | 0.621 | 0.373 | 8.42E-55 | Tb |
| allcell | CDK19 | 7.32E-33 | 0.885425628 | 0.373 | 0.203 | 1.44E-28 | Tb |
| allcell | WDPCP | 2.07E-28 | 0.884835107 | 0.282 | 0.142 | 4.07E-24 | Tb |
| allcell | TMCC1 | 7.87E-34 | 0.883778772 | 0.383 | 0.212 | 1.55E-29 | Tb |
| allcell | PRRG4 | 9.40E-28 | 0.882097825 | 0.307 | 0.165 | 1.85E-23 | Tb |
| allcell | SIK2 | 1.08E-28 | 0.881414835 | 0.446 | 0.278 | 2.13E-24 | Tb |
| allcell | RNF19A | 1.36E-40 | 0.88036768 | 0.549 | 0.343 | 2.67E-36 | Tb |
| allcell | TRAK1 | 1.09E-47 | 0.87993297 | 0.667 | 0.457 | 2.14E-43 | Tb |
| allcell | PITPNA | 7.82E-47 | 0.878182687 | 0.672 | 0.475 | 1.54E-42 | Tb |
| allcell | TAOK1 | 3.36E-45 | 0.877573741 | 0.538 | 0.309 | 6.62E-41 | Tb |
| allcell | FTO | 4.70E-32 | 0.874309142 | 0.297 | 0.145 | 9.24E-28 | Tb |
| allcell | ITCH | 4.21E-43 | 0.873904286 | 0.519 | 0.296 | 8.29E-39 | Tb |
| allcell | NBEAL1 | 5.71E-35 | 0.872741129 | 0.669 | 0.474 | 1.12E-30 | Tb |
| allcell | PRMT9 | 7.79E-44 | 0.87244899 | 0.618 | 0.401 | 1.53E-39 | Tb |
| allcell | MT-ND4 | 3.67E-132 | 0.871777147 | 0.989 | 0.933 | 7.23E-128 | Tb |
| allcell | DENND1B | 7.03E-57 | 0.869186896 | 0.776 | 0.584 | 1.38E-52 | Tb |
| allcell | BTBD7 | 2.46E-28 | 0.868572027 | 0.524 | 0.346 | 4.84E-24 | Tb |
| allcell | KANSL1 | 2.61E-50 | 0.866920159 | 0.533 | 0.299 | 5.14E-46 | Tb |
| allcell | MREG | 5.86E-26 | 0.866154102 | 0.381 | 0.23 | 1.15E-21 | Tb |
| allcell | BRAF | 5.78E-56 | 0.865232682 | 0.738 | 0.501 | 1.14E-51 | Tb |
| allcell | ELL2 | 6.30E-48 | 0.864367655 | 0.711 | 0.51 | 1.24E-43 | Tb |
| allcell | SPRED2 | 4.14E-29 | 0.863588082 | 0.544 | 0.381 | 8.15E-25 | Tb |
| allcell | CCDC97 | 5.68E-29 | 0.863358203 | 0.308 | 0.16 | 1.12E-24 | Tb |
| allcell | RAB30-DT | 2.75E-40 | 0.862446752 | 0.339 | 0.158 | 5.42E-36 | Tb |
| allcell | SEC23B | 1.99E-37 | 0.862283972 | 0.437 | 0.245 | 3.91E-33 | Tb |
| allcell | ZFAT | 2.78E-14 | 0.861994844 | 0.271 | 0.176 | 5.48E-10 | Tb |
| allcell | CSNK1G1 | 4.18E-33 | 0.861888437 | 0.338 | 0.177 | 8.23E-29 | Tb |
| allcell | IRS2 | 4.15E-21 | 0.861171548 | 0.273 | 0.155 | 8.16E-17 | Tb |
| allcell | VASH1 | 2.55E-40 | 0.861167825 | 0.433 | 0.235 | 5.03E-36 | Tb |
| allcell | FOXK2 | 1.08E-33 | 0.858707455 | 0.414 | 0.23 | 2.12E-29 | Tb |

| | | | | | | | |
|---|---|---|---|---|---|---|---|
| allcell | SCLT1 | 1.06E-49 | 0.858534299 | 0.711 | 0.494 | 2.08E-45 | Tb |
| allcell | STIM2 | 6.14E-39 | 0.855838077 | 0.5 | 0.303 | 1.21E-34 | Tb |
| allcell | NAF1 | 4.70E-17 | 0.854791933 | 0.459 | 0.331 | 9.25E-13 | Tb |
| allcell | FAF1 | 2.39E-47 | 0.854477289 | 0.596 | 0.374 | 4.71E-43 | Tb |
| allcell | GCNT1 | 6.84E-24 | 0.854095129 | 0.265 | 0.14 | 1.35E-19 | Tb |
| allcell | PAN3 | 4.96E-48 | 0.853028393 | 0.633 | 0.402 | 9.76E-44 | Tb |
| allcell | FBXL17 | 8.76E-41 | 0.850412536 | 0.502 | 0.292 | 1.73E-36 | Tb |
| allcell | WDR59 | 1.14E-30 | 0.849896388 | 0.336 | 0.177 | 2.25E-26 | Tb |
| allcell | GRAMD1B | 4.17E-27 | 0.849824504 | 0.372 | 0.222 | 8.21E-23 | Tb |
| allcell | TBC1D22A | 1.01E-54 | 0.849768521 | 0.704 | 0.489 | 1.99E-50 | Tb |
| allcell | ABTB2 | 1.17E-19 | 0.849573484 | 0.448 | 0.31 | 2.30E-15 | Tb |
| allcell | NRF1 | 5.43E-28 | 0.847879883 | 0.333 | 0.182 | 1.07E-23 | Tb |
| allcell | ILF3-DT | 2.26E-32 | 0.847409792 | 0.304 | 0.149 | 4.45E-28 | Tb |
| allcell | ABHD18 | 5.62E-20 | 0.847405371 | 0.25 | 0.137 | 1.11E-15 | Tb |
| allcell | SPDYA | 1.13E-19 | 0.84657305 | 0.283 | 0.169 | 2.22E-15 | Tb |
| allcell | SREBF2 | 3.91E-38 | 0.845193272 | 0.376 | 0.19 | 7.69E-34 | Tb |
| allcell | TNS3 | 2.00E-40 | 0.843596868 | 0.441 | 0.245 | 3.93E-36 | Tb |
| allcell | RAPGEF2 | 2.34E-35 | 0.843400361 | 0.693 | 0.517 | 4.61E-31 | Tb |
| allcell | LIMS1 | 1.46E-48 | 0.842945032 | 0.765 | 0.613 | 2.87E-44 | Tb |
| allcell | METTL16 | 8.76E-24 | 0.842200306 | 0.253 | 0.131 | 1.72E-19 | Tb |
| allcell | DAAM1 | 2.20E-21 | 0.841593536 | 0.348 | 0.213 | 4.32E-17 | Tb |
| allcell | ADNP2 | 7.95E-31 | 0.840837491 | 0.314 | 0.161 | 1.56E-26 | Tb |
| allcell | IPCEF1 | 1.24E-14 | 0.839760513 | 0.267 | 0.171 | 2.44E-10 | Tb |
| allcell | UBE4B | 1.08E-24 | 0.83873148 | 0.254 | 0.13 | 2.12E-20 | Tb |
| allcell | BABAM2 | 6.70E-38 | 0.838240217 | 0.569 | 0.375 | 1.32E-33 | Tb |
| allcell | MFSD14C | 5.01E-45 | 0.837067881 | 0.463 | 0.248 | 9.86E-41 | Tb |
| allcell | AHCYL2 | 1.60E-28 | 0.836206018 | 0.376 | 0.221 | 3.16E-24 | Tb |
| allcell | SKI | 1.34E-29 | 0.836038089 | 0.35 | 0.19 | 2.63E-25 | Tb |
| allcell | ITFG1 | 2.28E-40 | 0.83546699 | 0.523 | 0.316 | 4.49E-36 | Tb |
| allcell | RABGAP1L | 3.20E-41 | 0.833611047 | 0.768 | 0.57 | 6.30E-37 | Tb |
| allcell | CASS4 | 8.89E-27 | 0.833427189 | 0.37 | 0.217 | 1.75E-22 | Tb |
| allcell | ARHGAP10 | 5.59E-22 | 0.83271575 | 0.422 | 0.283 | 1.10E-17 | Tb |
| allcell | TNRC18 | 6.01E-33 | 0.832561057 | 0.417 | 0.237 | 1.18E-28 | Tb |
| allcell | LATS2 | 1.85E-20 | 0.831830251 | 0.341 | 0.21 | 3.64E-16 | Tb |
| allcell | RGS12 | 4.54E-31 | 0.831362434 | 0.324 | 0.17 | 8.94E-27 | Tb |
| allcell | S100PBP | 6.05E-24 | 0.830115537 | 0.253 | 0.131 | 1.19E-19 | Tb |
| allcell | RNF122 | 1.66E-29 | 0.829825482 | 0.269 | 0.132 | 3.27E-25 | Tb |
| allcell | ASB3 | 1.57E-24 | 0.829740328 | 0.379 | 0.226 | 3.08E-20 | Tb |
| allcell | LRBA | 1.94E-39 | 0.827591474 | 0.586 | 0.375 | 3.82E-35 | Tb |
| allcell | NFATC3 | 3.68E-25 | 0.827415996 | 0.314 | 0.172 | 7.25E-21 | Tb |
| allcell | ANAPC10 | 4.27E-32 | 0.826576779 | 0.404 | 0.225 | 8.40E-28 | Tb |
| allcell | ELMO1 | 4.31E-53 | 0.826571 | 0.816 | 0.668 | 8.49E-49 | Tb |
| allcell | STAU2 | 6.44E-32 | 0.82365668 | 0.336 | 0.175 | 1.27E-27 | Tb |
| allcell | PCYT1A | 1.40E-37 | 0.823638704 | 0.467 | 0.274 | 2.76E-33 | Tb |
| allcell | VPS54 | 2.16E-20 | 0.822674218 | 0.274 | 0.155 | 4.25E-16 | Tb |
| allcell | S100A9 | 6.61E-114 | 0.82226275 | 0.672 | 0.31 | 1.30E-109 | Tb |
| allcell | ALPK1 | 5.39E-36 | 0.821617554 | 0.453 | 0.268 | 1.06E-31 | Tb |
| allcell | ZBTB20 | 7.68E-46 | 0.821167935 | 0.619 | 0.406 | 1.51E-41 | Tb |
| allcell | GALNT7 | 2.31E-30 | 0.821104241 | 0.395 | 0.226 | 4.55E-26 | Tb |
| allcell | ATG7 | 5.63E-46 | 0.821055625 | 0.509 | 0.29 | 1.11E-41 | Tb |
| allcell | TSPAN14 | 2.46E-25 | 0.820224163 | 0.254 | 0.128 | 4.84E-21 | Tb |
| allcell | MFSD14B | 5.57E-33 | 0.82019518 | 0.457 | 0.272 | 1.10E-28 | Tb |
| allcell | MFN1 | 6.96E-32 | 0.819975226 | 0.363 | 0.195 | 1.37E-27 | Tb |
| allcell | PBRM1 | 4.64E-43 | 0.817571328 | 0.686 | 0.461 | 9.13E-39 | Tb |
| allcell | NOL10 | 4.21E-23 | 0.817499949 | 0.282 | 0.154 | 8.28E-19 | Tb |
| allcell | TET2 | 5.35E-30 | 0.817439218 | 0.825 | 0.72 | 1.05E-25 | Tb |

| | | | | | | | |
|---|---|---|---|---|---|---|---|
| allcell | CASP8 | 6.79E-40 | 0.815964018 | 0.547 | 0.331 | 1.34E-35 | Tb |
| allcell | ST3GAL2 | 1.83E-40 | 0.815803595 | 0.412 | 0.217 | 3.60E-36 | Tb |
| allcell | UBR1 | 2.04E-32 | 0.814736174 | 0.426 | 0.247 | 4.02E-28 | Tb |
| allcell | SLC12A2 | 1.44E-24 | 0.814181017 | 0.337 | 0.194 | 2.83E-20 | Tb |
| allcell | EVI5 | 2.84E-32 | 0.81398135 | 0.463 | 0.281 | 5.59E-28 | Tb |
| allcell | CDK6 | 1.84E-25 | 0.81330721 | 0.535 | 0.375 | 3.63E-21 | Tb |
| allcell | ESYT2 | 2.65E-45 | 0.813116873 | 0.781 | 0.628 | 5.21E-41 | Tb |
| allcell | RREB1 | 4.92E-37 | 0.810672666 | 0.561 | 0.355 | 9.68E-33 | Tb |
| allcell | SRGAP2C | 2.44E-36 | 0.810201283 | 0.352 | 0.181 | 4.80E-32 | Tb |
| allcell | ERCC6L2 | 8.25E-22 | 0.809057874 | 0.261 | 0.141 | 1.62E-17 | Tb |
| allcell | LPIN1 | 6.38E-20 | 0.808870573 | 0.392 | 0.252 | 1.26E-15 | Tb |
| allcell | SLC30A7 | 8.44E-27 | 0.80885964 | 0.332 | 0.182 | 1.66E-22 | Tb |
| allcell | RNF111 | 5.36E-33 | 0.808793807 | 0.43 | 0.25 | 1.05E-28 | Tb |
| allcell | FAM214A | 1.68E-27 | 0.80800806 | 0.33 | 0.182 | 3.30E-23 | Tb |
| allcell | CERS6 | 1.11E-44 | 0.807989452 | 0.735 | 0.521 | 2.18E-40 | Tb |
| allcell | ABR | 1.01E-34 | 0.807834708 | 0.659 | 0.479 | 2.00E-30 | Tb |
| allcell | POGZ | 1.24E-38 | 0.807641482 | 0.446 | 0.247 | 2.44E-34 | Tb |
| allcell | EPB41L2 | 1.11E-33 | 0.805894293 | 0.565 | 0.39 | 2.19E-29 | Tb |
| allcell | RFX7 | 1.27E-15 | 0.803487185 | 0.253 | 0.155 | 2.50E-11 | Tb |
| allcell | ARHGEF3 | 1.07E-30 | 0.802030053 | 0.353 | 0.192 | 2.11E-26 | Tb |
| allcell | CIITA | 3.17E-47 | 0.8010329 | 0.81 | 0.651 | 6.24E-43 | Tb |
| allcell | TPT1-AS1 | 1.62E-31 | 0.800832294 | 0.253 | 0.113 | 3.18E-27 | Tb |
| allcell | SAMD9L | 2.69E-47 | 0.799713197 | 0.528 | 0.298 | 5.30E-43 | Tb |
| allcell | CHML | 3.14E-21 | 0.799385033 | 0.357 | 0.218 | 6.17E-17 | Tb |
| allcell | MSL2 | 1.58E-33 | 0.798925874 | 0.368 | 0.194 | 3.11E-29 | Tb |
| allcell | ZCCHC7 | 1.70E-42 | 0.798100795 | 0.584 | 0.374 | 3.34E-38 | Tb |
| allcell | JAK3 | 1.68E-26 | 0.798011213 | 0.286 | 0.15 | 3.30E-22 | Tb |
| allcell | SLC20A2 | 1.46E-24 | 0.797380156 | 0.279 | 0.149 | 2.86E-20 | Tb |
| allcell | CCDC91 | 3.42E-45 | 0.797247156 | 0.639 | 0.408 | 6.72E-41 | Tb |
| allcell | RBM6 | 2.64E-43 | 0.796838897 | 0.614 | 0.393 | 5.20E-39 | Tb |
| allcell | EZH2 | 3.55E-49 | 0.796722304 | 0.663 | 0.438 | 6.98E-45 | Tb |
| allcell | TMEM39A | 9.28E-31 | 0.796434064 | 0.527 | 0.35 | 1.83E-26 | Tb |
| allcell | PPP3CC | 1.85E-31 | 0.795704979 | 0.402 | 0.228 | 3.64E-27 | Tb |
| allcell | UBL3 | 9.81E-36 | 0.794249093 | 0.671 | 0.48 | 1.93E-31 | Tb |
| allcell | ABCC1 | 1.13E-39 | 0.794092156 | 0.397 | 0.205 | 2.21E-35 | Tb |
| allcell | MYO9A | 1.43E-40 | 0.793836999 | 0.475 | 0.276 | 2.81E-36 | Tb |
| allcell | CLCN5 | 6.73E-29 | 0.793358394 | 0.412 | 0.25 | 1.33E-24 | Tb |
| allcell | DDX58 | 1.08E-20 | 0.792190795 | 0.251 | 0.137 | 2.12E-16 | Tb |
| allcell | CFLAR-AS1 | 2.82E-33 | 0.792047382 | 0.36 | 0.19 | 5.55E-29 | Tb |
| allcell | SLMAP | 2.31E-37 | 0.791945617 | 0.409 | 0.218 | 4.54E-33 | Tb |
| allcell | PPCDC | 1.08E-23 | 0.791938356 | 0.362 | 0.217 | 2.13E-19 | Tb |
| allcell | RERE | 1.16E-40 | 0.791212742 | 0.717 | 0.528 | 2.28E-36 | Tb |
| allcell | LRRC8C | 1.40E-22 | 0.791101193 | 0.385 | 0.244 | 2.76E-18 | Tb |
| allcell | GATAD2B | 3.18E-45 | 0.790571003 | 0.53 | 0.309 | 6.26E-41 | Tb |
| allcell | MAP3K14 | 6.47E-26 | 0.790331979 | 0.431 | 0.269 | 1.27E-21 | Tb |
| allcell | CHKA | 2.37E-25 | 0.789881679 | 0.421 | 0.265 | 4.66E-21 | Tb |
| allcell | TYW1B | 5.10E-30 | 0.787597786 | 0.283 | 0.141 | 1.00E-25 | Tb |
| allcell | GPHN | 8.18E-21 | 0.786207613 | 0.337 | 0.206 | 1.61E-16 | Tb |
| allcell | SLC7A6 | 4.33E-23 | 0.785949498 | 0.281 | 0.154 | 8.53E-19 | Tb |
| allcell | P2RY6 | 1.36E-18 | 0.785861267 | 0.4 | 0.27 | 2.68E-14 | Tb |
| allcell | SNRK | 1.69E-21 | 0.784081056 | 0.294 | 0.168 | 3.33E-17 | Tb |
| allcell | CD2AP | 1.15E-27 | 0.78349371 | 0.489 | 0.316 | 2.26E-23 | Tb |
| allcell | TCF4 | 2.60E-25 | 0.783448817 | 0.652 | 0.482 | 5.12E-21 | Tb |
| allcell | ITPR1 | 4.32E-46 | 0.782006793 | 0.57 | 0.349 | 8.50E-42 | Tb |
| allcell | SLC9A7 | 6.92E-30 | 0.781855973 | 0.564 | 0.389 | 1.36E-25 | Tb |
| allcell | UBE2O | 4.40E-25 | 0.781227076 | 0.295 | 0.158 | 8.65E-21 | Tb |

| | | | | | | | |
|---|---|---|---|---|---|---|---|
| allcell | BICRAL | 2.22E-33 | 0.781063899 | 0.29 | 0.137 | 4.37E-29 | Tb |
| allcell | DCAF10 | 2.15E-22 | 0.780706745 | 0.272 | 0.148 | 4.23E-18 | Tb |
| allcell | CDKAL1 | 7.27E-38 | 0.779889281 | 0.476 | 0.281 | 1.43E-33 | Tb |
| allcell | PHTF2 | 1.06E-32 | 0.779498651 | 0.462 | 0.279 | 2.08E-28 | Tb |
| allcell | AP1S3 | 4.43E-29 | 0.778430397 | 0.452 | 0.286 | 8.71E-25 | Tb |
| allcell | ARFGEF2 | 6.21E-38 | 0.77830805 | 0.432 | 0.237 | 1.22E-33 | Tb |
| allcell | WIPF2 | 3.91E-28 | 0.778083865 | 0.285 | 0.146 | 7.70E-24 | Tb |
| allcell | FAM168A | 5.27E-30 | 0.778010012 | 0.335 | 0.179 | 1.04E-25 | Tb |
| allcell | SOAT1 | 9.11E-26 | 0.777873784 | 0.393 | 0.237 | 1.79E-21 | Tb |
| allcell | LAMP3 | 1.91E-20 | 0.777552485 | 0.468 | 0.327 | 3.76E-16 | Tb |
| allcell | APOL2 | 4.82E-27 | 0.776954501 | 0.279 | 0.142 | 9.49E-23 | Tb |
| allcell | ERBIN | 8.77E-39 | 0.776667218 | 0.645 | 0.446 | 1.73E-34 | Tb |
| allcell | RNF38 | 1.03E-32 | 0.776633389 | 0.439 | 0.259 | 2.03E-28 | Tb |
| allcell | GRK3 | 3.26E-39 | 0.776486707 | 0.663 | 0.466 | 6.41E-35 | Tb |
| allcell | ABL2 | 2.93E-33 | 0.776272195 | 0.516 | 0.332 | 5.76E-29 | Tb |
| allcell | LYRM1 | 1.46E-24 | 0.775865638 | 0.268 | 0.138 | 2.88E-20 | Tb |
| allcell | CBFB | 2.67E-42 | 0.775561372 | 0.469 | 0.258 | 5.26E-38 | Tb |
| allcell | MALT1 | 5.67E-33 | 0.774803649 | 0.804 | 0.676 | 1.12E-28 | Tb |
| allcell | FBXL20 | 3.42E-22 | 0.773074941 | 0.301 | 0.173 | 6.74E-18 | Tb |
| allcell | ZCCHC2 | 3.02E-30 | 0.772839745 | 0.414 | 0.241 | 5.95E-26 | Tb |
| allcell | ALOX12-AS1 | 4.13E-26 | 0.77167239 | 0.255 | 0.127 | 8.14E-22 | Tb |
| allcell | KDM5B | 1.22E-25 | 0.771586074 | 0.344 | 0.197 | 2.40E-21 | Tb |
| allcell | QRICH1 | 7.33E-30 | 0.771318058 | 0.316 | 0.164 | 1.44E-25 | Tb |
| allcell | UBAC2 | 4.53E-41 | 0.770521335 | 0.719 | 0.528 | 8.91E-37 | Tb |
| allcell | GSK3B | 1.50E-40 | 0.77023073 | 0.595 | 0.392 | 2.96E-36 | Tb |
| allcell | ATR | 2.30E-32 | 0.769855797 | 0.402 | 0.226 | 4.52E-28 | Tb |
| allcell | HIP1 | 3.47E-30 | 0.76839631 | 0.584 | 0.42 | 6.83E-26 | Tb |
| allcell | P2RY8 | 8.94E-20 | 0.767217074 | 0.254 | 0.144 | 1.76E-15 | Tb |
| allcell | FCHSD2 | 1.24E-46 | 0.766914676 | 0.847 | 0.672 | 2.45E-42 | Tb |
| allcell | NPC1 | 6.69E-24 | 0.766122095 | 0.359 | 0.212 | 1.32E-19 | Tb |
| allcell | PDE7A | 7.55E-36 | 0.763724891 | 0.667 | 0.474 | 1.49E-31 | Tb |
| allcell | CD226 | 9.30E-22 | 0.763615088 | 0.353 | 0.221 | 1.83E-17 | Tb |
| allcell | MTHFR | 6.71E-29 | 0.762993236 | 0.406 | 0.235 | 1.32E-24 | Tb |
| allcell | HS3ST3B1 | 2.13E-18 | 0.762699694 | 0.31 | 0.193 | 4.19E-14 | Tb |
| allcell | TERF1 | 4.12E-19 | 0.762190464 | 0.355 | 0.22 | 8.11E-15 | Tb |
| allcell | OXNAD1 | 1.47E-28 | 0.761987809 | 0.28 | 0.14 | 2.89E-24 | Tb |
| allcell | DMXL1 | 1.42E-37 | 0.761299868 | 0.541 | 0.332 | 2.79E-33 | Tb |
| allcell | INPP5D | 1.23E-23 | 0.759977364 | 0.509 | 0.351 | 2.42E-19 | Tb |
| allcell | XPO1 | 3.87E-47 | 0.759507724 | 0.684 | 0.45 | 7.62E-43 | Tb |
| allcell | CDK17 | 5.73E-28 | 0.758998261 | 0.445 | 0.277 | 1.13E-23 | Tb |
| allcell | UTRN | 3.68E-36 | 0.75890789 | 0.663 | 0.462 | 7.24E-32 | Tb |
| allcell | TXLNA | 1.99E-28 | 0.7575859 | 0.33 | 0.176 | 3.91E-24 | Tb |
| allcell | ABHD3 | 2.26E-30 | 0.757151393 | 0.56 | 0.383 | 4.45E-26 | Tb |
| allcell | EYA3 | 2.49E-31 | 0.756839574 | 0.383 | 0.213 | 4.90E-27 | Tb |
| allcell | ZMYM2 | 4.44E-29 | 0.756544084 | 0.663 | 0.487 | 8.74E-25 | Tb |
| allcell | FOXP1 | 3.48E-47 | 0.756052324 | 0.832 | 0.673 | 6.84E-43 | Tb |
| allcell | JPX | 4.07E-38 | 0.755535225 | 0.5 | 0.29 | 8.00E-34 | Tb |
| allcell | STXBP5 | 1.22E-19 | 0.75536554 | 0.255 | 0.144 | 2.40E-15 | Tb |
| allcell | SKIL | 4.83E-53 | 0.754852417 | 0.789 | 0.569 | 9.51E-49 | Tb |
| allcell | PLEKHM3 | 3.37E-18 | 0.754504251 | 0.359 | 0.236 | 6.63E-14 | Tb |
| allcell | MLLT10 | 6.46E-34 | 0.754123507 | 0.437 | 0.253 | 1.27E-29 | Tb |
| allcell | FRS2 | 1.28E-30 | 0.753734174 | 0.335 | 0.177 | 2.51E-26 | Tb |
| allcell | VPS8 | 4.67E-20 | 0.753712606 | 0.339 | 0.21 | 9.19E-16 | Tb |
| allcell | LHFPL2 | 6.96E-16 | 0.753162567 | 0.262 | 0.16 | 1.37E-11 | Tb |
| allcell | ITPR2 | 3.43E-47 | 0.752392665 | 0.657 | 0.418 | 6.76E-43 | Tb |
| allcell | ARSB | 4.27E-27 | 0.751803415 | 0.417 | 0.251 | 8.41E-23 | Tb |

| | | | | | | | |
|---|---|---|---|---|---|---|---|
| allcell | CLASP1 | 2.55E-35 | 0.751787627 | 0.417 | 0.23 | 5.02E-31 | Tb |
| allcell | ZMYND8 | 9.93E-34 | 0.751250801 | 0.382 | 0.206 | 1.95E-29 | Tb |
| allcell | TOPORS | 5.06E-32 | 0.750919394 | 0.397 | 0.222 | 9.96E-28 | Tb |
| allcell | TCOF1 | 8.40E-21 | 0.750511655 | 0.318 | 0.187 | 1.65E-16 | Tb |
| allcell | ARMH3 | 8.93E-43 | 0.749805647 | 0.467 | 0.252 | 1.76E-38 | Tb |
| allcell | TRPM2 | 6.40E-22 | 0.749529934 | 0.259 | 0.143 | 1.26E-17 | Tb |
| allcell | PSEN1 | 1.10E-37 | 0.748758087 | 0.541 | 0.338 | 2.16E-33 | Tb |
| allcell | MOSMO | 1.29E-26 | 0.748524776 | 0.29 | 0.151 | 2.53E-22 | Tb |
| allcell | NR2C2 | 7.69E-25 | 0.747180374 | 0.286 | 0.153 | 1.51E-20 | Tb |
| allcell | THOC1 | 3.55E-20 | 0.746599136 | 0.287 | 0.167 | 6.99E-16 | Tb |
| allcell | TWISTNB | 1.45E-18 | 0.742903305 | 0.616 | 0.455 | 2.86E-14 | Tb |
| allcell | CSF2RB | 4.26E-42 | 0.742759437 | 0.518 | 0.296 | 8.38E-38 | Tb |
| allcell | TYW1 | 2.61E-30 | 0.742609838 | 0.351 | 0.189 | 5.14E-26 | Tb |
| allcell | BACH2 | 1.84E-19 | 0.741801181 | 0.297 | 0.179 | 3.62E-15 | Tb |
| allcell | ADD3 | 3.82E-19 | 0.741377955 | 0.346 | 0.221 | 7.52E-15 | Tb |
| allcell | COP1 | 1.79E-50 | 0.740724194 | 0.68 | 0.441 | 3.53E-46 | Tb |
| allcell | CBFA2T2 | 3.23E-30 | 0.740718144 | 0.32 | 0.168 | 6.35E-26 | Tb |
| allcell | MAN1A1 | 1.33E-50 | 0.740555302 | 0.79 | 0.569 | 2.61E-46 | Tb |
| allcell | MTMR9 | 7.90E-20 | 0.740531268 | 0.291 | 0.168 | 1.55E-15 | Tb |
| allcell | SRGAP2B | 6.31E-33 | 0.74051055 | 0.411 | 0.231 | 1.24E-28 | Tb |
| allcell | MBD5 | 1.87E-31 | 0.739906029 | 0.436 | 0.261 | 3.68E-27 | Tb |
| allcell | RIPK1 | 5.72E-37 | 0.739804193 | 0.538 | 0.332 | 1.13E-32 | Tb |
| allcell | TUT4 | 5.02E-19 | 0.739649551 | 0.278 | 0.163 | 9.89E-15 | Tb |
| allcell | CPED1 | 1.51E-14 | 0.738968712 | 0.275 | 0.175 | 2.97E-10 | Tb |
| allcell | NRP1 | 4.97E-19 | 0.738510349 | 0.302 | 0.185 | 9.77E-15 | Tb |
| allcell | PMM2 | 1.91E-25 | 0.738203739 | 0.342 | 0.193 | 3.76E-21 | Tb |
| allcell | TRIM38 | 2.93E-42 | 0.737568352 | 0.622 | 0.408 | 5.76E-38 | Tb |
| allcell | R3HDM2 | 6.71E-41 | 0.737518797 | 0.601 | 0.397 | 1.32E-36 | Tb |
| allcell | RBMS1 | 7.98E-30 | 0.737250111 | 0.499 | 0.316 | 1.57E-25 | Tb |
| allcell | NCOA1 | 3.61E-40 | 0.737131018 | 0.632 | 0.411 | 7.10E-36 | Tb |
| allcell | COMMD10 | 4.39E-29 | 0.7367169 | 0.607 | 0.414 | 8.64E-25 | Tb |
| allcell | ATP2C1 | 2.67E-31 | 0.735969048 | 0.675 | 0.484 | 5.26E-27 | Tb |
| allcell | TRAPPC9 | 3.54E-46 | 0.734037487 | 0.539 | 0.311 | 6.98E-42 | Tb |
| allcell | YAF2 | 3.27E-27 | 0.733641596 | 0.445 | 0.272 | 6.44E-23 | Tb |
| allcell | PPP1R14A | 2.32E-19 | 0.733098199 | 0.275 | 0.161 | 4.56E-15 | Tb |
| allcell | ANKRD44 | 1.26E-50 | 0.731994985 | 0.752 | 0.547 | 2.48E-46 | Tb |
| allcell | PDS5B | 3.47E-39 | 0.731646295 | 0.474 | 0.268 | 6.83E-35 | Tb |
| allcell | TENT2 | 3.39E-50 | 0.731552486 | 0.631 | 0.381 | 6.67E-46 | Tb |
| allcell | PRKCB | 7.61E-33 | 0.731028878 | 0.701 | 0.517 | 1.50E-28 | Tb |
| allcell | KDM7A-DT | 6.57E-22 | 0.729886834 | 0.25 | 0.13 | 1.29E-17 | Tb |
| allcell | ADAMTSL4-AS | 2.64E-36 | 0.729862764 | 0.49 | 0.29 | 5.21E-32 | Tb |
| allcell | CDC42SE2 | 2.89E-58 | 0.729850815 | 0.871 | 0.68 | 5.68E-54 | Tb |
| allcell | SP140 | 2.26E-17 | 0.729666002 | 0.353 | 0.232 | 4.45E-13 | Tb |
| allcell | SLC45A4 | 2.86E-24 | 0.729363837 | 0.318 | 0.18 | 5.63E-20 | Tb |
| allcell | CDR2 | 1.25E-30 | 0.728634252 | 0.502 | 0.32 | 2.46E-26 | Tb |
| allcell | SHLD2 | 5.49E-20 | 0.72835001 | 0.353 | 0.217 | 1.08E-15 | Tb |
| allcell | DOCK8 | 1.60E-39 | 0.726353722 | 0.751 | 0.58 | 3.15E-35 | Tb |
| allcell | PRORP | 1.78E-25 | 0.726304573 | 0.314 | 0.171 | 3.50E-21 | Tb |
| allcell | PTPN9 | 1.19E-19 | 0.726128223 | 0.269 | 0.152 | 2.34E-15 | Tb |
| allcell | ZZEF1 | 1.07E-30 | 0.726117683 | 0.403 | 0.227 | 2.10E-26 | Tb |
| allcell | RANBP2 | 2.09E-28 | 0.726116352 | 0.782 | 0.611 | 4.12E-24 | Tb |
| allcell | SH3KBP1 | 1.14E-26 | 0.724944628 | 0.66 | 0.502 | 2.24E-22 | Tb |
| allcell | CAMKMT | 4.98E-15 | 0.724890169 | 0.264 | 0.164 | 9.80E-11 | Tb |
| allcell | IBTK | 2.14E-29 | 0.723597223 | 0.393 | 0.223 | 4.22E-25 | Tb |
| allcell | PDSS2 | 4.66E-26 | 0.723214862 | 0.284 | 0.148 | 9.18E-22 | Tb |
| allcell | PICALM | 2.99E-52 | 0.721966071 | 0.852 | 0.719 | 5.88E-48 | Tb |

| | | | | | | | |
|---|---|---|---|---|---|---|---|
| allcell | DOCK10 | 1.12E-36 | 0.721637679 | 0.791 | 0.648 | 2.20E-32 | Tb |
| allcell | ADIPOR2 | 2.07E-32 | 0.72127276 | 0.379 | 0.207 | 4.07E-28 | Tb |
| allcell | DOP1A | 3.18E-19 | 0.721255365 | 0.316 | 0.195 | 6.26E-15 | Tb |
| allcell | RUBCN | 1.31E-24 | 0.720710484 | 0.484 | 0.323 | 2.58E-20 | Tb |
| allcell | SNX14 | 5.80E-27 | 0.720679383 | 0.378 | 0.219 | 1.14E-22 | Tb |
| allcell | ATXN7 | 3.51E-29 | 0.71977714 | 0.441 | 0.269 | 6.91E-25 | Tb |
| allcell | SLC44A1 | 5.19E-17 | 0.719684445 | 0.346 | 0.222 | 1.02E-12 | Tb |
| allcell | ULK4 | 1.33E-26 | 0.718211948 | 0.313 | 0.17 | 2.62E-22 | Tb |
| allcell | KIAA2026 | 2.83E-33 | 0.718099244 | 0.418 | 0.238 | 5.57E-29 | Tb |
| allcell | CLASP2 | 2.70E-42 | 0.717555732 | 0.644 | 0.427 | 5.32E-38 | Tb |
| allcell | SUPT3H | 1.55E-19 | 0.717494714 | 0.279 | 0.161 | 3.05E-15 | Tb |
| allcell | MICAL2 | 9.65E-21 | 0.717361969 | 0.323 | 0.195 | 1.90E-16 | Tb |
| allcell | ALOX5 | 6.66E-22 | 0.71689432 | 0.439 | 0.296 | 1.31E-17 | Tb |
| allcell | FAM172A | 1.21E-34 | 0.716152062 | 0.508 | 0.315 | 2.39E-30 | Tb |
| allcell | SMURF1 | 4.21E-35 | 0.715808344 | 0.421 | 0.238 | 8.28E-31 | Tb |
| allcell | SLC38A1 | 1.52E-33 | 0.714129648 | 0.626 | 0.44 | 2.99E-29 | Tb |
| allcell | ATP1B3 | 8.56E-29 | 0.713912273 | 0.9 | 0.837 | 1.68E-24 | Tb |
| allcell | MCCC1 | 2.77E-25 | 0.713628881 | 0.276 | 0.143 | 5.45E-21 | Tb |
| allcell | INTS6L | 5.02E-28 | 0.713183096 | 0.302 | 0.158 | 9.88E-24 | Tb |
| allcell | ENTPD1 | 9.81E-22 | 0.712945827 | 0.522 | 0.375 | 1.93E-17 | Tb |
| allcell | SNX8 | 5.61E-35 | 0.711913891 | 0.679 | 0.528 | 1.11E-30 | Tb |
| allcell | MT-CO1 | 2.46E-80 | 0.711510088 | 0.997 | 0.985 | 4.84E-76 | Tb |
| allcell | PHF21A | 1.41E-23 | 0.711248974 | 0.399 | 0.247 | 2.78E-19 | Tb |
| allcell | MICU1 | 2.91E-25 | 0.711245956 | 0.368 | 0.219 | 5.73E-21 | Tb |
| allcell | DIP2B | 6.52E-34 | 0.711008895 | 0.439 | 0.251 | 1.28E-29 | Tb |
| allcell | GLG1 | 4.52E-29 | 0.710305669 | 0.522 | 0.338 | 8.89E-25 | Tb |
| allcell | MITD1 | 5.09E-22 | 0.709076769 | 0.331 | 0.192 | 1.00E-17 | Tb |
| allcell | AKT3 | 2.45E-21 | 0.708932791 | 0.356 | 0.218 | 4.83E-17 | Tb |
| allcell | ZXDC | 7.43E-23 | 0.708215667 | 0.293 | 0.162 | 1.46E-18 | Tb |
| allcell | ANKIB1 | 1.54E-29 | 0.707634515 | 0.411 | 0.243 | 3.03E-25 | Tb |
| allcell | HMG20A | 2.80E-19 | 0.707621876 | 0.255 | 0.141 | 5.51E-15 | Tb |
| allcell | AP3S2 | 2.91E-23 | 0.706973053 | 0.316 | 0.177 | 5.72E-19 | Tb |
| allcell | ITPRIPL2 | 9.22E-23 | 0.706517979 | 0.281 | 0.152 | 1.82E-18 | Tb |
| allcell | DLEU1 | 1.26E-42 | 0.705327662 | 0.495 | 0.283 | 2.49E-38 | Tb |
| allcell | SEMA3C | 8.86E-17 | 0.705203864 | 0.259 | 0.155 | 1.74E-12 | Tb |
| allcell | RHBDD1 | 3.41E-20 | 0.705017954 | 0.298 | 0.176 | 6.71E-16 | Tb |
| allcell | SIPA1L1 | 4.29E-42 | 0.70492995 | 0.884 | 0.758 | 8.44E-38 | Tb |
| allcell | DIS3L2 | 4.39E-21 | 0.703970752 | 0.279 | 0.157 | 8.65E-17 | Tb |
| allcell | TLK1 | 1.61E-29 | 0.703618577 | 0.481 | 0.294 | 3.16E-25 | Tb |
| allcell | RLF | 1.51E-27 | 0.703581398 | 0.652 | 0.48 | 2.98E-23 | Tb |
| allcell | CCDC18 | 4.68E-29 | 0.703398136 | 0.353 | 0.195 | 9.21E-25 | Tb |
| allcell | ST3GAL1 | 2.17E-29 | 0.703045307 | 0.458 | 0.283 | 4.27E-25 | Tb |
| allcell | LARP1B | 1.87E-20 | 0.702857235 | 0.312 | 0.183 | 3.67E-16 | Tb |
| allcell | TMEM131 | 1.72E-37 | 0.702708446 | 0.717 | 0.528 | 3.39E-33 | Tb |
| allcell | UBP1 | 3.52E-25 | 0.702226878 | 0.317 | 0.175 | 6.93E-21 | Tb |
| allcell | GAB2 | 3.06E-33 | 0.702145534 | 0.658 | 0.47 | 6.03E-29 | Tb |
| allcell | PIP5K1A | 1.47E-32 | 0.701695351 | 0.427 | 0.247 | 2.90E-28 | Tb |
| allcell | EXT1 | 3.01E-30 | 0.701215589 | 0.479 | 0.303 | 5.93E-26 | Tb |
| allcell | SSH1 | 6.90E-20 | 0.70106283 | 0.3 | 0.175 | 1.36E-15 | Tb |
| allcell | MT-ND3 | 6.65E-97 | 0.700564326 | 0.983 | 0.899 | 1.31E-92 | Tb |
| allcell | USP32 | 1.77E-42 | 0.699917194 | 0.638 | 0.421 | 3.49E-38 | Tb |
| allcell | ZNF410 | 4.23E-26 | 0.699909659 | 0.381 | 0.223 | 8.33E-22 | Tb |
| allcell | RICTOR | 1.23E-27 | 0.699882077 | 0.432 | 0.262 | 2.42E-23 | Tb |
| allcell | SUSD6 | 7.14E-36 | 0.69855056 | 0.586 | 0.39 | 1.41E-31 | Tb |
| allcell | MED13L | 8.96E-60 | 0.698436392 | 0.88 | 0.7 | 1.76E-55 | Tb |
| allcell | MYO1C | 5.32E-28 | 0.697586422 | 0.302 | 0.158 | 1.05E-23 | Tb |

| | | | | | | | |
|---|---|---|---|---|---|---|---|
| allcell | INTS6 | 2.13E-39 | 0.697555882 | 0.731 | 0.545 | 4.20E-35 | Tb |
| allcell | FOXO1 | 4.52E-17 | 0.697337039 | 0.4 | 0.277 | 8.90E-13 | Tb |
| allcell | SLC16A3 | 2.83E-27 | 0.696944026 | 0.425 | 0.259 | 5.57E-23 | Tb |
| allcell | SEMA4D | 1.96E-19 | 0.696348657 | 0.266 | 0.151 | 3.85E-15 | Tb |
| allcell | SYNJ1 | 1.15E-24 | 0.694981799 | 0.307 | 0.17 | 2.27E-20 | Tb |
| allcell | NUBPL | 1.80E-22 | 0.69479366 | 0.428 | 0.282 | 3.55E-18 | Tb |
| allcell | IKZF1 | 2.11E-39 | 0.694735683 | 0.728 | 0.556 | 4.16E-35 | Tb |
| allcell | PRKCE | 2.26E-20 | 0.693536807 | 0.43 | 0.295 | 4.46E-16 | Tb |
| allcell | TBL1XR1 | 1.99E-36 | 0.693478969 | 0.667 | 0.457 | 3.92E-32 | Tb |
| allcell | RBBP8 | 5.46E-09 | 0.6924252 | 0.342 | 0.248 | 0.000107425 | Tb |
| allcell | FKBP5 | 1.04E-48 | 0.692082444 | 0.707 | 0.487 | 2.04E-44 | Tb |
| allcell | ATP6V0A1 | 8.57E-27 | 0.691812593 | 0.358 | 0.203 | 1.69E-22 | Tb |
| allcell | FNDC3B | 4.28E-36 | 0.691656543 | 0.669 | 0.479 | 8.42E-32 | Tb |
| allcell | RIPOR2 | 2.45E-22 | 0.691414635 | 0.532 | 0.367 | 4.82E-18 | Tb |
| allcell | STAT6 | 6.16E-41 | 0.691167909 | 0.527 | 0.314 | 1.21E-36 | Tb |
| allcell | MDN1 | 4.90E-21 | 0.690580283 | 0.282 | 0.158 | 9.64E-17 | Tb |
| allcell | CHST11 | 1.59E-40 | 0.690514941 | 0.741 | 0.557 | 3.13E-36 | Tb |
| allcell | DENND4B | 6.14E-23 | 0.690411696 | 0.284 | 0.154 | 1.21E-18 | Tb |
| allcell | STRADA | 2.40E-25 | 0.690086531 | 0.27 | 0.139 | 4.73E-21 | Tb |
| allcell | MCM9 | 2.82E-19 | 0.68947687 | 0.255 | 0.144 | 5.55E-15 | Tb |
| allcell | ARHGAP18 | 2.68E-26 | 0.688798243 | 0.574 | 0.412 | 5.28E-22 | Tb |
| allcell | ANKRD13C | 1.66E-19 | 0.688786406 | 0.252 | 0.14 | 3.28E-15 | Tb |
| allcell | ANKRD17 | 1.14E-38 | 0.688009315 | 0.637 | 0.425 | 2.24E-34 | Tb |
| allcell | CREB5 | 2.00E-12 | 0.686402193 | 0.447 | 0.342 | 3.93E-08 | Tb |
| allcell | ELMSAN1 | 3.27E-38 | 0.685953821 | 0.515 | 0.306 | 6.43E-34 | Tb |
| allcell | NUCB2 | 3.16E-15 | 0.685693913 | 0.318 | 0.212 | 6.21E-11 | Tb |
| allcell | NMRK1 | 9.73E-16 | 0.684601228 | 0.287 | 0.179 | 1.92E-11 | Tb |
| allcell | PPA2 | 8.77E-32 | 0.68448263 | 0.516 | 0.325 | 1.73E-27 | Tb |
| allcell | TGIF2 | 4.20E-25 | 0.684341498 | 0.419 | 0.259 | 8.27E-21 | Tb |
| allcell | B3GALNT2 | 1.23E-20 | 0.684321345 | 0.263 | 0.146 | 2.42E-16 | Tb |
| allcell | LRMP | 4.24E-15 | 0.683266667 | 0.374 | 0.257 | 8.34E-11 | Tb |
| allcell | PPP2R5A | 2.14E-30 | 0.682382702 | 0.329 | 0.17 | 4.21E-26 | Tb |
| allcell | RAB5B | 8.59E-20 | 0.682013399 | 0.329 | 0.198 | 1.69E-15 | Tb |
| allcell | KDM4C | 8.83E-47 | 0.681958588 | 0.67 | 0.44 | 1.74E-42 | Tb |
| allcell | CDK13 | 5.19E-30 | 0.68181147 | 0.551 | 0.366 | 1.02E-25 | Tb |
| allcell | DCLRE1C | 2.85E-20 | 0.68164573 | 0.251 | 0.135 | 5.61E-16 | Tb |
| allcell | CYP20A1 | 1.25E-17 | 0.681435573 | 0.394 | 0.261 | 2.47E-13 | Tb |
| allcell | FGD2 | 2.03E-34 | 0.681279488 | 0.512 | 0.313 | 3.99E-30 | Tb |
| allcell | SOS2 | 8.13E-24 | 0.681092422 | 0.379 | 0.23 | 1.60E-19 | Tb |
| allcell | SH2B3 | 3.53E-42 | 0.680724546 | 0.712 | 0.53 | 6.95E-38 | Tb |
| allcell | ACOX1 | 1.70E-16 | 0.68061862 | 0.251 | 0.146 | 3.36E-12 | Tb |
| allcell | SCFD2 | 6.11E-21 | 0.680580816 | 0.33 | 0.199 | 1.20E-16 | Tb |
| allcell | CDYL | 1.90E-35 | 0.680444612 | 0.714 | 0.521 | 3.75E-31 | Tb |
| allcell | NLRC5 | 5.93E-34 | 0.680307547 | 0.47 | 0.28 | 1.17E-29 | Tb |
| allcell | VPS53 | 8.84E-35 | 0.680225963 | 0.436 | 0.246 | 1.74E-30 | Tb |
| allcell | NFAT5 | 9.59E-33 | 0.679901958 | 0.825 | 0.712 | 1.89E-28 | Tb |
| allcell | SPATS2 | 3.83E-33 | 0.679761079 | 0.542 | 0.342 | 7.53E-29 | Tb |
| allcell | TBL1X | 1.40E-17 | 0.679359825 | 0.382 | 0.255 | 2.76E-13 | Tb |
| allcell | CRYBG1 | 8.20E-19 | 0.67894459 | 0.614 | 0.477 | 1.61E-14 | Tb |
| allcell | ELF2 | 3.67E-33 | 0.678649613 | 0.677 | 0.489 | 7.22E-29 | Tb |
| allcell | GABPB1-IT1 | 2.78E-21 | 0.678114211 | 0.284 | 0.157 | 5.47E-17 | Tb |
| allcell | 8-Mar | 9.10E-27 | 0.678036652 | 0.323 | 0.175 | 1.79E-22 | Tb |
| allcell | ENTPD1-AS1 | 1.04E-36 | 0.677539024 | 0.459 | 0.262 | 2.04E-32 | Tb |
| allcell | SELENOI | 2.53E-20 | 0.677380015 | 0.293 | 0.168 | 4.99E-16 | Tb |
| allcell | MT-ND1 | 1.07E-87 | 0.677273708 | 0.97 | 0.898 | 2.10E-83 | Tb |
| allcell | AHI1 | 4.28E-22 | 0.676277751 | 0.5 | 0.339 | 8.42E-18 | Tb |

| | | | | | | | |
|---|---|---|---|---|---|---|---|
| allcell | ZBTB44 | 5.33E-21 | 0.675323866 | 0.269 | 0.148 | 1.05E-16 | Tb |
| allcell | NUP107 | 3.38E-21 | 0.674891159 | 0.282 | 0.158 | 6.65E-17 | Tb |
| allcell | PHC3 | 1.62E-21 | 0.674741264 | 0.376 | 0.234 | 3.18E-17 | Tb |
| allcell | YARS | 3.78E-24 | 0.67461575 | 0.35 | 0.201 | 7.44E-20 | Tb |
| allcell | MED27 | 6.30E-23 | 0.673916665 | 0.3 | 0.167 | 1.24E-18 | Tb |
| allcell | ZFX | 4.34E-24 | 0.673796313 | 0.425 | 0.269 | 8.54E-20 | Tb |
| allcell | GPBP1L1 | 3.40E-37 | 0.673479585 | 0.533 | 0.316 | 6.69E-33 | Tb |
| allcell | KCNK6 | 8.14E-24 | 0.673376467 | 0.367 | 0.224 | 1.60E-19 | Tb |
| allcell | OSBPL3 | 1.17E-18 | 0.673238644 | 0.473 | 0.336 | 2.31E-14 | Tb |
| allcell | IP6K1 | 1.95E-18 | 0.673160999 | 0.27 | 0.155 | 3.83E-14 | Tb |
| allcell | FBXO34 | 2.54E-48 | 0.672551256 | 0.683 | 0.458 | 5.00E-44 | Tb |
| allcell | RCOR1 | 3.13E-28 | 0.671984133 | 0.731 | 0.582 | 6.17E-24 | Tb |
| allcell | EXOC6 | 9.71E-28 | 0.671557368 | 0.382 | 0.223 | 1.91E-23 | Tb |
| allcell | NLRP1 | 1.30E-14 | 0.671209881 | 0.316 | 0.205 | 2.55E-10 | Tb |
| allcell | EML4 | 2.33E-40 | 0.671132299 | 0.717 | 0.523 | 4.59E-36 | Tb |
| allcell | AGO3 | 1.14E-32 | 0.671120234 | 0.53 | 0.334 | 2.25E-28 | Tb |
| allcell | CEP128 | 2.49E-23 | 0.670803254 | 0.363 | 0.22 | 4.91E-19 | Tb |
| allcell | CCDC18-AS1 | 7.00E-24 | 0.67069831 | 0.328 | 0.186 | 1.38E-19 | Tb |
| allcell | MT-CO2 | 7.65E-96 | 0.670482558 | 0.989 | 0.982 | 1.51E-91 | Tb |
| allcell | STK3 | 1.98E-28 | 0.670262964 | 0.528 | 0.353 | 3.89E-24 | Tb |
| allcell | 3-Mar | 7.95E-12 | 0.670064766 | 0.313 | 0.215 | 1.57E-07 | Tb |
| allcell | MR1 | 6.18E-20 | 0.670062062 | 0.263 | 0.146 | 1.22E-15 | Tb |
| allcell | RAB3GAP1 | 8.40E-33 | 0.669569416 | 0.471 | 0.283 | 1.65E-28 | Tb |
| allcell | HAVCR2 | 5.49E-13 | 0.66928621 | 0.528 | 0.405 | 1.08E-08 | Tb |
| allcell | SEL1L3 | 1.35E-23 | 0.668076089 | 0.444 | 0.288 | 2.66E-19 | Tb |
| allcell | HOMER2 | 3.48E-28 | 0.668070909 | 0.286 | 0.143 | 6.86E-24 | Tb |
| allcell | EXOC2 | 3.00E-22 | 0.667938797 | 0.281 | 0.154 | 5.90E-18 | Tb |
| allcell | ZBTB46 | 8.00E-31 | 0.667374029 | 0.488 | 0.308 | 1.58E-26 | Tb |
| allcell | ZNF124 | 4.86E-34 | 0.666903107 | 0.395 | 0.215 | 9.56E-30 | Tb |
| allcell | PHACTR2 | 1.97E-21 | 0.666710663 | 0.44 | 0.292 | 3.88E-17 | Tb |
| allcell | FLT1 | 2.39E-10 | 0.666270552 | 0.554 | 0.471 | 4.70E-06 | Tb |
| allcell | TRIM13 | 2.81E-29 | 0.666167982 | 0.393 | 0.222 | 5.53E-25 | Tb |
| allcell | ACSL1 | 9.73E-12 | 0.66597617 | 0.586 | 0.461 | 1.92E-07 | Tb |
| allcell | PPP1R21 | 9.48E-20 | 0.665932594 | 0.276 | 0.159 | 1.87E-15 | Tb |
| allcell | YME1L1 | 5.82E-34 | 0.665802343 | 0.801 | 0.627 | 1.15E-29 | Tb |
| allcell | TEX14 | 2.06E-20 | 0.665539499 | 0.562 | 0.415 | 4.05E-16 | Tb |
| allcell | MRTFA | 2.12E-30 | 0.665371127 | 0.674 | 0.491 | 4.17E-26 | Tb |
| allcell | FARP2 | 4.58E-26 | 0.664867342 | 0.552 | 0.385 | 9.02E-22 | Tb |
| allcell | GFPT1 | 4.57E-29 | 0.663961183 | 0.416 | 0.244 | 9.00E-25 | Tb |
| allcell | FPR3 | 1.14E-10 | 0.663960603 | 0.416 | 0.323 | 2.25E-06 | Tb |
| allcell | SC5D | 3.04E-16 | 0.663140794 | 0.251 | 0.148 | 5.99E-12 | Tb |
| allcell | PDE4A | 1.84E-18 | 0.663101733 | 0.663 | 0.539 | 3.62E-14 | Tb |
| allcell | TRIM24 | 9.80E-18 | 0.662872887 | 0.321 | 0.2 | 1.93E-13 | Tb |
| allcell | ZC3H4 | 5.25E-28 | 0.662550774 | 0.303 | 0.155 | 1.03E-23 | Tb |
| allcell | MLLT6 | 1.46E-18 | 0.6624337 | 0.297 | 0.174 | 2.88E-14 | Tb |
| allcell | PCBD2 | 2.95E-21 | 0.661381091 | 0.257 | 0.138 | 5.81E-17 | Tb |
| allcell | BCL2L14 | 2.06E-12 | 0.660786162 | 0.279 | 0.188 | 4.06E-08 | Tb |
| allcell | APOL6 | 1.35E-16 | 0.660559531 | 0.281 | 0.17 | 2.66E-12 | Tb |
| allcell | TRAFD1 | 5.29E-24 | 0.660186388 | 0.334 | 0.193 | 1.04E-19 | Tb |
| allcell | KLF12 | 1.71E-13 | 0.660096779 | 0.25 | 0.158 | 3.37E-09 | Tb |
| allcell | ZNF407 | 6.93E-32 | 0.660095274 | 0.458 | 0.271 | 1.36E-27 | Tb |
| allcell | ST3GAL5 | 4.84E-40 | 0.659991348 | 0.544 | 0.321 | 9.52E-36 | Tb |
| allcell | FOXN3 | 3.87E-31 | 0.658933434 | 0.777 | 0.644 | 7.63E-27 | Tb |
| allcell | CNOT10 | 3.02E-25 | 0.658601862 | 0.418 | 0.261 | 5.95E-21 | Tb |
| allcell | GTF2H1 | 7.92E-23 | 0.657995315 | 0.455 | 0.298 | 1.56E-18 | Tb |
| allcell | ZEB1 | 3.22E-27 | 0.657985247 | 0.585 | 0.429 | 6.34E-23 | Tb |

| | | | | | | | |
|---|---|---|---|---|---|---|---|
| allcell | IDO2 | 7.45E-24 | 0.657532318 | 0.492 | 0.33 | 1.47E-19 | Tb |
| allcell | MTIF2 | 4.41E-26 | 0.657348091 | 0.286 | 0.148 | 8.69E-22 | Tb |
| allcell | MAP3K5 | 8.10E-26 | 0.657316495 | 0.468 | 0.301 | 1.59E-21 | Tb |
| allcell | SLC7A1 | 4.99E-29 | 0.656264814 | 0.468 | 0.287 | 9.83E-25 | Tb |
| allcell | STK38L | 1.51E-28 | 0.65593523 | 0.596 | 0.42 | 2.98E-24 | Tb |
| allcell | LTB | 6.49E-28 | 0.655669065 | 0.572 | 0.397 | 1.28E-23 | Tb |
| allcell | FAM133B | 4.11E-41 | 0.65503473 | 0.582 | 0.356 | 8.08E-37 | Tb |
| allcell | YEATS2 | 6.51E-24 | 0.654972831 | 0.316 | 0.179 | 1.28E-19 | Tb |
| allcell | SOCS1 | 2.95E-30 | 0.654863477 | 0.645 | 0.453 | 5.80E-26 | Tb |
| allcell | RAPH1 | 2.15E-11 | 0.654629107 | 0.297 | 0.209 | 4.23E-07 | Tb |
| allcell | OGA | 1.02E-31 | 0.654543383 | 0.486 | 0.298 | 2.01E-27 | Tb |
| allcell | ABI2 | 3.50E-21 | 0.654534851 | 0.314 | 0.183 | 6.88E-17 | Tb |
| allcell | ZNF791 | 8.07E-23 | 0.654343429 | 0.322 | 0.185 | 1.59E-18 | Tb |
| allcell | CNOT6L | 2.82E-28 | 0.653967127 | 0.619 | 0.43 | 5.54E-24 | Tb |
| allcell | HERC4 | 7.72E-26 | 0.653853432 | 0.407 | 0.247 | 1.52E-21 | Tb |
| allcell | STAT3 | 1.41E-46 | 0.653582294 | 0.798 | 0.607 | 2.77E-42 | Tb |
| allcell | 1-Mar | 1.34E-23 | 0.653548509 | 0.566 | 0.408 | 2.64E-19 | Tb |
| allcell | POU2F1 | 3.02E-24 | 0.653543787 | 0.346 | 0.201 | 5.95E-20 | Tb |
| allcell | MBNL1 | 1.40E-41 | 0.653495851 | 0.896 | 0.79 | 2.76E-37 | Tb |
| allcell | AP5Z1 | 5.24E-33 | 0.653494878 | 0.355 | 0.183 | 1.03E-28 | Tb |
| allcell | SETDB2 | 2.41E-22 | 0.653200354 | 0.442 | 0.285 | 4.74E-18 | Tb |
| allcell | TCF20 | 4.34E-22 | 0.652660152 | 0.283 | 0.156 | 8.54E-18 | Tb |
| allcell | GNPTAB | 1.65E-28 | 0.652605275 | 0.453 | 0.275 | 3.24E-24 | Tb |
| allcell | GABPB1 | 1.87E-21 | 0.652160282 | 0.322 | 0.19 | 3.68E-17 | Tb |
| allcell | PSD3 | 4.85E-09 | 0.651450288 | 0.257 | 0.178 | 9.55E-05 | Tb |
| allcell | IQCB1 | 3.07E-25 | 0.650536575 | 0.384 | 0.229 | 6.04E-21 | Tb |
| allcell | UBE2E1 | 8.31E-28 | 0.649676614 | 0.656 | 0.5 | 1.63E-23 | Tb |
| allcell | ADPGK | 9.13E-36 | 0.649311036 | 0.505 | 0.295 | 1.80E-31 | Tb |
| allcell | ZKSCAN1 | 9.10E-25 | 0.649015123 | 0.34 | 0.191 | 1.79E-20 | Tb |
| allcell | RABGAP1 | 3.56E-28 | 0.64847823 | 0.433 | 0.26 | 7.01E-24 | Tb |
| allcell | NIN | 3.02E-22 | 0.648302142 | 0.546 | 0.388 | 5.94E-18 | Tb |
| allcell | SLA | 4.56E-19 | 0.647976335 | 0.276 | 0.16 | 8.98E-15 | Tb |
| allcell | ZNF490 | 1.65E-29 | 0.647443343 | 0.273 | 0.133 | 3.25E-25 | Tb |
| allcell | KPNA6 | 3.51E-20 | 0.647426964 | 0.294 | 0.168 | 6.91E-16 | Tb |
| allcell | PCSK7 | 5.47E-31 | 0.64740579 | 0.311 | 0.157 | 1.08E-26 | Tb |
| allcell | KLHL15 | 5.71E-16 | 0.64658069 | 0.287 | 0.174 | 1.12E-11 | Tb |
| allcell | EIF4G3 | 1.35E-22 | 0.646541091 | 0.517 | 0.367 | 2.67E-18 | Tb |
| allcell | TFEC | 7.36E-08 | 0.646372802 | 0.476 | 0.382 | 0.001449657 | Tb |
| allcell | EPC2 | 1.86E-19 | 0.646299743 | 0.334 | 0.203 | 3.67E-15 | Tb |
| allcell | SNX29 | 1.21E-34 | 0.646004887 | 0.619 | 0.435 | 2.38E-30 | Tb |
| allcell | CGAS | 1.35E-28 | 0.645662367 | 0.469 | 0.29 | 2.66E-24 | Tb |
| allcell | PCNX4 | 1.47E-19 | 0.645449617 | 0.346 | 0.217 | 2.90E-15 | Tb |
| allcell | STRN3 | 3.45E-30 | 0.645424811 | 0.404 | 0.23 | 6.79E-26 | Tb |
| allcell | PRIM2 | 2.59E-16 | 0.645141392 | 0.259 | 0.154 | 5.10E-12 | Tb |
| allcell | PARP9 | 1.67E-24 | 0.645062448 | 0.383 | 0.23 | 3.28E-20 | Tb |
| allcell | MICU2 | 1.10E-21 | 0.644953666 | 0.403 | 0.251 | 2.16E-17 | Tb |
| allcell | SINHCAF | 6.94E-22 | 0.644710198 | 0.607 | 0.449 | 1.37E-17 | Tb |
| allcell | RBM23 | 8.50E-29 | 0.644580278 | 0.425 | 0.249 | 1.67E-24 | Tb |
| allcell | RFTN1 | 8.68E-31 | 0.644016549 | 0.777 | 0.658 | 1.71E-26 | Tb |
| allcell | MGAT5 | 4.66E-34 | 0.643880032 | 0.506 | 0.307 | 9.16E-30 | Tb |
| allcell | NEMP1 | 3.05E-20 | 0.642191683 | 0.25 | 0.134 | 6.00E-16 | Tb |
| allcell | BAZ2B | 2.21E-25 | 0.642076746 | 0.549 | 0.382 | 4.36E-21 | Tb |
| allcell | KRT10 | 2.92E-14 | 0.641434195 | 0.686 | 0.543 | 5.75E-10 | Tb |
| allcell | RAPGEF1 | 1.74E-48 | 0.641334699 | 0.89 | 0.768 | 3.43E-44 | Tb |
| allcell | USP36 | 5.16E-35 | 0.640054671 | 0.626 | 0.407 | 1.02E-30 | Tb |
| allcell | TCF12 | 9.03E-43 | 0.639900792 | 0.66 | 0.44 | 1.78E-38 | Tb |

| | | | | | | | |
|---|---|---|---|---|---|---|---|
| allcell | TPD52 | 8.41E-16 | 0.639370464 | 0.313 | 0.198 | 1.66E-11 | Tb |
| allcell | CMTM7 | 3.94E-24 | 0.638962904 | 0.575 | 0.397 | 7.76E-20 | Tb |
| allcell | WDR70 | 7.23E-26 | 0.638687577 | 0.459 | 0.289 | 1.42E-21 | Tb |
| allcell | DDX10 | 7.81E-25 | 0.638146474 | 0.346 | 0.197 | 1.54E-20 | Tb |
| allcell | AKAP8 | 3.29E-21 | 0.637855647 | 0.325 | 0.187 | 6.49E-17 | Tb |
| allcell | NSD2 | 2.41E-23 | 0.637647427 | 0.285 | 0.154 | 4.75E-19 | Tb |
| allcell | SMURF2 | 3.10E-34 | 0.637288674 | 0.502 | 0.303 | 6.09E-30 | Tb |
| allcell | PPP2CB | 2.18E-26 | 0.63688949 | 0.387 | 0.226 | 4.29E-22 | Tb |
| allcell | HIVEP2 | 5.20E-28 | 0.636860542 | 0.726 | 0.563 | 1.02E-23 | Tb |
| allcell | NEK9 | 5.80E-20 | 0.636654026 | 0.258 | 0.141 | 1.14E-15 | Tb |
| allcell | RYBP | 2.02E-38 | 0.635986629 | 0.69 | 0.481 | 3.99E-34 | Tb |
| allcell | ARID1A | 1.34E-34 | 0.635576742 | 0.494 | 0.294 | 2.65E-30 | Tb |
| allcell | KDM4B | 2.89E-37 | 0.634994285 | 0.503 | 0.291 | 5.70E-33 | Tb |
| allcell | EPB41 | 1.98E-19 | 0.634938125 | 0.438 | 0.294 | 3.91E-15 | Tb |
| allcell | ATP2B4 | 1.44E-18 | 0.634745533 | 0.329 | 0.204 | 2.83E-14 | Tb |
| allcell | PACS1 | 3.25E-22 | 0.634591832 | 0.538 | 0.385 | 6.39E-18 | Tb |
| allcell | CPEB2 | 2.23E-14 | 0.634160596 | 0.372 | 0.253 | 4.39E-10 | Tb |
| allcell | AQR | 3.51E-21 | 0.6337895 | 0.321 | 0.188 | 6.92E-17 | Tb |
| allcell | SRBD1 | 1.39E-14 | 0.633644426 | 0.27 | 0.168 | 2.75E-10 | Tb |
| allcell | XPO6 | 2.65E-26 | 0.632806164 | 0.362 | 0.205 | 5.22E-22 | Tb |
| allcell | SEPHS1 | 1.88E-20 | 0.632667502 | 0.334 | 0.199 | 3.70E-16 | Tb |
| allcell | TXNDC5 | 3.57E-13 | 0.632587696 | 0.259 | 0.165 | 7.02E-09 | Tb |
| allcell | WDR44 | 9.01E-20 | 0.632541154 | 0.334 | 0.203 | 1.77E-15 | Tb |
| allcell | FGD4 | 2.94E-19 | 0.632244429 | 0.453 | 0.316 | 5.79E-15 | Tb |
| allcell | CRLF3 | 4.12E-21 | 0.632032794 | 0.361 | 0.22 | 8.11E-17 | Tb |
| allcell | ZNF518A | 2.54E-21 | 0.631719515 | 0.327 | 0.193 | 5.00E-17 | Tb |
| allcell | BCL2L13 | 2.27E-21 | 0.631384457 | 0.452 | 0.3 | 4.47E-17 | Tb |
| allcell | NCOA3 | 1.14E-29 | 0.631334581 | 0.671 | 0.498 | 2.23E-25 | Tb |
| allcell | ANKRD33B | 6.76E-08 | 0.631168991 | 0.349 | 0.277 | 0.00133062 | Tb |
| allcell | PHLDB3 | 4.60E-22 | 0.630732726 | 0.344 | 0.206 | 9.06E-18 | Tb |
| allcell | PKNOX1 | 1.11E-20 | 0.630623793 | 0.295 | 0.168 | 2.18E-16 | Tb |
| allcell | PTPN22 | 6.28E-12 | 0.630367893 | 0.418 | 0.31 | 1.24E-07 | Tb |
| allcell | RSRC1 | 3.70E-33 | 0.63027996 | 0.53 | 0.327 | 7.27E-29 | Tb |
| allcell | DIAPH2 | 1.12E-25 | 0.630245813 | 0.5 | 0.326 | 2.21E-21 | Tb |
| allcell | AFF1 | 4.49E-34 | 0.630089053 | 0.742 | 0.549 | 8.84E-30 | Tb |
| allcell | VEZT | 6.73E-27 | 0.629907549 | 0.429 | 0.259 | 1.32E-22 | Tb |
| allcell | PLXNC1 | 1.83E-24 | 0.629459757 | 0.535 | 0.366 | 3.59E-20 | Tb |
| allcell | ZSWIM6 | 5.03E-43 | 0.62930254 | 0.838 | 0.674 | 9.90E-39 | Tb |
| allcell | ZNF710 | 4.21E-27 | 0.629133635 | 0.575 | 0.401 | 8.29E-23 | Tb |
| allcell | UBE2E2 | 1.83E-39 | 0.628813224 | 0.827 | 0.694 | 3.61E-35 | Tb |
| allcell | TET3 | 6.16E-29 | 0.628076379 | 0.412 | 0.24 | 1.21E-24 | Tb |
| allcell | HERC3 | 2.14E-19 | 0.62752948 | 0.269 | 0.154 | 4.22E-15 | Tb |
| allcell | PCNX1 | 6.29E-26 | 0.627264561 | 0.567 | 0.391 | 1.24E-21 | Tb |
| allcell | TPK1 | 1.39E-20 | 0.626647631 | 0.403 | 0.256 | 2.74E-16 | Tb |
| allcell | KRT5 | 1.64E-45 | 0.625609678 | 0.32 | 0.138 | 3.23E-41 | Tb |
| allcell | HEATR5B | 2.11E-19 | 0.625590894 | 0.331 | 0.2 | 4.16E-15 | Tb |
| allcell | TMEM131L | 2.59E-23 | 0.625566387 | 0.458 | 0.299 | 5.09E-19 | Tb |
| allcell | GNB4 | 2.25E-26 | 0.625355642 | 0.534 | 0.359 | 4.43E-22 | Tb |
| allcell | RIOK1 | 2.49E-22 | 0.625170454 | 0.369 | 0.22 | 4.91E-18 | Tb |
| allcell | SIPA1L3 | 1.13E-39 | 0.624901327 | 0.569 | 0.365 | 2.22E-35 | Tb |
| allcell | RBM47 | 1.15E-31 | 0.624516435 | 0.738 | 0.578 | 2.26E-27 | Tb |
| allcell | APPBP2 | 1.55E-28 | 0.624189189 | 0.335 | 0.177 | 3.06E-24 | Tb |
| allcell | MGAT2 | 5.87E-23 | 0.624157649 | 0.282 | 0.15 | 1.15E-18 | Tb |
| allcell | R3HCC1L | 2.30E-23 | 0.624118499 | 0.363 | 0.214 | 4.52E-19 | Tb |
| allcell | SF3B3 | 8.05E-19 | 0.624094297 | 0.27 | 0.153 | 1.58E-14 | Tb |
| allcell | NSD3 | 1.90E-22 | 0.622353448 | 0.656 | 0.483 | 3.75E-18 | Tb |

| | | | | | | | |
|---|---|---|---|---|---|---|---|
| allcell | TRPM7 | 2.21E-29 | 0.621566069 | 0.481 | 0.297 | 4.35E-25 | Tb |
| allcell | MAPKAP1 | 1.55E-28 | 0.621561234 | 0.561 | 0.369 | 3.05E-24 | Tb |
| allcell | GOLPH3 | 3.61E-29 | 0.621535627 | 0.442 | 0.261 | 7.11E-25 | Tb |
| allcell | COX19 | 5.51E-20 | 0.621171582 | 0.351 | 0.213 | 1.09E-15 | Tb |
| allcell | PLCG2 | 1.85E-24 | 0.620898444 | 0.475 | 0.306 | 3.65E-20 | Tb |
| allcell | SMC6 | 4.81E-16 | 0.620454317 | 0.293 | 0.184 | 9.48E-12 | Tb |
| allcell | NF1 | 4.45E-26 | 0.620342013 | 0.544 | 0.373 | 8.77E-22 | Tb |
| allcell | B4GALT4 | 7.75E-19 | 0.620278777 | 0.337 | 0.209 | 1.53E-14 | Tb |
| allcell | ASH1L | 3.59E-40 | 0.620107877 | 0.756 | 0.556 | 7.06E-36 | Tb |
| allcell | USP37 | 6.10E-20 | 0.620064911 | 0.422 | 0.279 | 1.20E-15 | Tb |
| allcell | STRBP | 9.90E-19 | 0.619846289 | 0.441 | 0.303 | 1.95E-14 | Tb |
| allcell | MFSD4B | 1.62E-22 | 0.619581782 | 0.404 | 0.249 | 3.18E-18 | Tb |
| allcell | GGA2 | 2.58E-25 | 0.618948782 | 0.428 | 0.267 | 5.07E-21 | Tb |
| allcell | CPB2-AS1 | 2.70E-26 | 0.618627977 | 0.393 | 0.229 | 5.31E-22 | Tb |
| allcell | RBM26 | 5.75E-25 | 0.618538781 | 0.421 | 0.257 | 1.13E-20 | Tb |
| allcell | TRMO | 3.64E-18 | 0.618480965 | 0.266 | 0.152 | 7.17E-14 | Tb |
| allcell | EPG5 | 1.14E-21 | 0.618088613 | 0.267 | 0.144 | 2.25E-17 | Tb |
| allcell | CLOCK | 3.29E-12 | 0.61796071 | 0.255 | 0.164 | 6.48E-08 | Tb |
| allcell | YTHDC2 | 1.27E-23 | 0.617304164 | 0.513 | 0.338 | 2.51E-19 | Tb |
| allcell | ASAP1 | 2.13E-21 | 0.616861296 | 0.782 | 0.713 | 4.19E-17 | Tb |
| allcell | BCAS3 | 1.11E-26 | 0.616861208 | 0.399 | 0.241 | 2.19E-22 | Tb |
| allcell | CYREN | 1.52E-19 | 0.61666429 | 0.301 | 0.175 | 3.00E-15 | Tb |
| allcell | KAT6A | 3.64E-22 | 0.616310438 | 0.48 | 0.322 | 7.17E-18 | Tb |
| allcell | GALM | 4.67E-11 | 0.614968751 | 0.372 | 0.273 | 9.19E-07 | Tb |
| allcell | SLC35E1 | 5.85E-24 | 0.614837962 | 0.337 | 0.19 | 1.15E-19 | Tb |
| allcell | FBXO28 | 2.14E-20 | 0.614524444 | 0.269 | 0.149 | 4.20E-16 | Tb |
| allcell | ZC3H8 | 5.09E-20 | 0.614132225 | 0.26 | 0.144 | 1.00E-15 | Tb |
| allcell | PAK1 | 2.58E-27 | 0.613470408 | 0.848 | 0.729 | 5.08E-23 | Tb |
| allcell | LPIN2 | 1.95E-28 | 0.612969001 | 0.467 | 0.285 | 3.83E-24 | Tb |
| allcell | RALGAPB | 6.64E-15 | 0.612916942 | 0.253 | 0.151 | 1.31E-10 | Tb |
| allcell | LIMD1 | 1.14E-22 | 0.612684458 | 0.323 | 0.187 | 2.24E-18 | Tb |
| allcell | ABCC4 | 3.36E-19 | 0.612637561 | 0.28 | 0.162 | 6.62E-15 | Tb |
| allcell | TENT4A | 1.47E-28 | 0.612306296 | 0.471 | 0.287 | 2.90E-24 | Tb |
| allcell | TRAF3 | 2.78E-44 | 0.611966445 | 0.727 | 0.511 | 5.47E-40 | Tb |
| allcell | TYMP | 2.92E-47 | 0.611431545 | 0.913 | 0.797 | 5.75E-43 | Tb |
| allcell | PPP6R2 | 4.40E-33 | 0.6114 | 0.51 | 0.31 | 8.66E-29 | Tb |
| allcell | EXOC4 | 7.44E-39 | 0.611251699 | 0.663 | 0.445 | 1.47E-34 | Tb |
| allcell | MT-ND2 | 3.13E-73 | 0.610338315 | 0.973 | 0.882 | 6.17E-69 | Tb |
| allcell | CPEB4 | 7.27E-34 | 0.610324414 | 0.589 | 0.389 | 1.43E-29 | Tb |
| allcell | UBE2W | 1.36E-27 | 0.610286701 | 0.65 | 0.487 | 2.68E-23 | Tb |
| allcell | SGMS2 | 2.08E-14 | 0.610084489 | 0.274 | 0.172 | 4.09E-10 | Tb |
| allcell | HECTD4 | 8.83E-16 | 0.609977784 | 0.306 | 0.195 | 1.74E-11 | Tb |
| allcell | CHD8 | 4.82E-23 | 0.609918413 | 0.332 | 0.192 | 9.48E-19 | Tb |
| allcell | THUMPD3-AS1 | 1.84E-32 | 0.609905617 | 0.542 | 0.341 | 3.63E-28 | Tb |
| allcell | P2RY13 | 7.04E-21 | 0.609769782 | 0.328 | 0.195 | 1.39E-16 | Tb |
| allcell | WASHC5 | 5.78E-13 | 0.609546118 | 0.277 | 0.179 | 1.14E-08 | Tb |
| allcell | TTC9C | 2.72E-28 | 0.609392464 | 0.297 | 0.149 | 5.35E-24 | Tb |
| allcell | STAG1 | 5.16E-38 | 0.608852543 | 0.647 | 0.43 | 1.02E-33 | Tb |
| allcell | IDI1 | 1.88E-23 | 0.608661359 | 0.67 | 0.501 | 3.71E-19 | Tb |
| allcell | DAPP1 | 7.31E-26 | 0.608550206 | 0.766 | 0.64 | 1.44E-21 | Tb |
| allcell | MON2 | 5.69E-24 | 0.608343426 | 0.442 | 0.283 | 1.12E-19 | Tb |
| allcell | SPOP | 1.55E-27 | 0.608246375 | 0.498 | 0.316 | 3.05E-23 | Tb |
| allcell | TP53 | 9.12E-21 | 0.608075612 | 0.282 | 0.157 | 1.80E-16 | Tb |
| allcell | KIAA0232 | 1.55E-21 | 0.607829754 | 0.367 | 0.222 | 3.05E-17 | Tb |
| allcell | KDM2B | 4.83E-11 | 0.607818191 | 0.493 | 0.379 | 9.50E-07 | Tb |
| allcell | ZNF146 | 1.42E-23 | 0.607362305 | 0.281 | 0.147 | 2.80E-19 | Tb |

| | | | | | | | |
|---|---|---|---|---|---|---|---|
| allcell | AEBP2 | 2.41E-18 | 0.607233594 | 0.386 | 0.252 | 4.74E-14 | Tb |
| allcell | KDM3A | 1.41E-22 | 0.607137914 | 0.305 | 0.171 | 2.77E-18 | Tb |
| allcell | STAM2 | 1.14E-18 | 0.606944604 | 0.289 | 0.17 | 2.24E-14 | Tb |
| allcell | IGF2BP2 | 1.44E-22 | 0.606931035 | 0.358 | 0.219 | 2.84E-18 | Tb |
| allcell | IL21R | 2.91E-16 | 0.606523096 | 0.303 | 0.192 | 5.73E-12 | Tb |
| allcell | GALNT1 | 1.50E-30 | 0.606088746 | 0.485 | 0.295 | 2.94E-26 | Tb |
| allcell | MNAT1 | 3.29E-20 | 0.606014628 | 0.342 | 0.208 | 6.47E-16 | Tb |
| allcell | SKAP2 | 4.93E-30 | 0.605911564 | 0.633 | 0.449 | 9.71E-26 | Tb |
| allcell | CEP135 | 1.42E-19 | 0.605074069 | 0.306 | 0.182 | 2.79E-15 | Tb |
| allcell | FBXO11 | 1.08E-37 | 0.604939301 | 0.782 | 0.6 | 2.13E-33 | Tb |
| allcell | USP6NL | 1.83E-18 | 0.604634329 | 0.316 | 0.193 | 3.60E-14 | Tb |
| allcell | HEATR5A | 3.17E-20 | 0.604382655 | 0.293 | 0.169 | 6.23E-16 | Tb |
| allcell | SATB1-AS1 | 1.45E-17 | 0.603749526 | 0.286 | 0.172 | 2.86E-13 | Tb |
| allcell | LCOR | 2.25E-29 | 0.603100288 | 0.588 | 0.402 | 4.42E-25 | Tb |
| allcell | RIC1 | 4.86E-25 | 0.60259104 | 0.589 | 0.419 | 9.56E-21 | Tb |
| allcell | PLXDC2 | 1.10E-17 | 0.601945307 | 0.556 | 0.432 | 2.17E-13 | Tb |
| allcell | EHMT1 | 8.70E-40 | 0.601027583 | 0.542 | 0.322 | 1.71E-35 | Tb |
| allcell | PPP2R5E | 3.17E-26 | 0.600969085 | 0.462 | 0.292 | 6.24E-22 | Tb |
| allcell | INO80 | 9.65E-24 | 0.600805291 | 0.406 | 0.248 | 1.90E-19 | Tb |
| allcell | TNRC6B | 1.20E-27 | 0.600447711 | 0.664 | 0.483 | 2.36E-23 | Tb |
| allcell | CCNT1 | 1.71E-22 | 0.600259854 | 0.386 | 0.238 | 3.37E-18 | Tb |
| allcell | PRPSAP1 | 3.39E-18 | 0.600205592 | 0.286 | 0.166 | 6.68E-14 | Tb |
| allcell | ATP11B | 1.46E-29 | 0.599422929 | 0.562 | 0.37 | 2.88E-25 | Tb |
| allcell | MIB1 | 3.01E-25 | 0.599347473 | 0.293 | 0.156 | 5.92E-21 | Tb |
| allcell | HMBOX1 | 1.02E-24 | 0.599329741 | 0.417 | 0.255 | 2.01E-20 | Tb |
| allcell | NSRP1 | 1.08E-26 | 0.599186592 | 0.459 | 0.279 | 2.12E-22 | Tb |
| allcell | CSTF3 | 4.47E-22 | 0.599186522 | 0.28 | 0.151 | 8.80E-18 | Tb |
| allcell | ERI1 | 4.12E-19 | 0.598272034 | 0.316 | 0.19 | 8.10E-15 | Tb |
| allcell | MEGF9 | 6.12E-10 | 0.598034234 | 0.262 | 0.18 | 1.21E-05 | Tb |
| allcell | SLC27A4 | 3.59E-28 | 0.597951257 | 0.437 | 0.259 | 7.06E-24 | Tb |
| allcell | PTPN7 | 2.80E-15 | 0.597713893 | 0.264 | 0.16 | 5.52E-11 | Tb |
| allcell | TGFBR2 | 5.29E-14 | 0.597353999 | 0.526 | 0.398 | 1.04E-09 | Tb |
| allcell | PHF14 | 5.05E-26 | 0.596713596 | 0.451 | 0.279 | 9.95E-22 | Tb |
| allcell | CARS2 | 3.67E-23 | 0.596639034 | 0.328 | 0.184 | 7.22E-19 | Tb |
| allcell | LY75 | 2.08E-18 | 0.595959796 | 0.491 | 0.341 | 4.09E-14 | Tb |
| allcell | CEBPA | 3.29E-15 | 0.595191343 | 0.272 | 0.169 | 6.48E-11 | Tb |
| allcell | SRSF4 | 8.91E-35 | 0.594614646 | 0.668 | 0.448 | 1.75E-30 | Tb |
| allcell | MAP3K13 | 1.38E-23 | 0.594129596 | 0.643 | 0.485 | 2.72E-19 | Tb |
| allcell | LNPEP | 8.37E-24 | 0.593751898 | 0.469 | 0.297 | 1.65E-19 | Tb |
| allcell | EED | 1.96E-22 | 0.593608509 | 0.421 | 0.261 | 3.86E-18 | Tb |
| allcell | FOXK1 | 1.27E-17 | 0.59353238 | 0.303 | 0.184 | 2.51E-13 | Tb |
| allcell | HSH2D | 1.25E-20 | 0.593164548 | 0.299 | 0.171 | 2.47E-16 | Tb |
| allcell | PLEKHA51 | 3.73E-07 | 0.593013898 | 0.43 | 0.343 | 0.007334339 | Tb |
| allcell | RALGAPA2 | 1.15E-23 | 0.592805053 | 0.3 | 0.166 | 2.27E-19 | Tb |
| allcell | N4BP1 | 5.33E-29 | 0.592743131 | 0.607 | 0.425 | 1.05E-24 | Tb |
| allcell | SRPK2 | 2.12E-22 | 0.592637269 | 0.539 | 0.377 | 4.17E-18 | Tb |
| allcell | SGMS1 | 1.84E-21 | 0.592628024 | 0.345 | 0.208 | 3.62E-17 | Tb |
| allcell | PMEPA1 | 9.35E-26 | 0.592498567 | 0.375 | 0.219 | 1.84E-21 | Tb |
| allcell | RUNX1 | 1.42E-33 | 0.592310192 | 0.805 | 0.651 | 2.79E-29 | Tb |
| allcell | RNF19B | 2.07E-17 | 0.592225997 | 0.442 | 0.307 | 4.07E-13 | Tb |
| allcell | CDK8 | 3.43E-16 | 0.591938145 | 0.267 | 0.16 | 6.76E-12 | Tb |
| allcell | FAM107B | 1.01E-28 | 0.591738007 | 0.762 | 0.601 | 1.99E-24 | Tb |
| allcell | SHB | 1.19E-14 | 0.591670266 | 0.314 | 0.202 | 2.33E-10 | Tb |
| allcell | SCARF1 | 4.42E-22 | 0.591149045 | 0.361 | 0.217 | 8.70E-18 | Tb |
| allcell | PBX3 | 4.52E-28 | 0.591039518 | 0.603 | 0.424 | 8.89E-24 | Tb |
| allcell | RFX3 | 5.49E-19 | 0.590650812 | 0.34 | 0.21 | 1.08E-14 | Tb |

| | | | | | | | |
|---|---|---|---|---|---|---|---|
| allcell | ENTPD7 | 1.35E-19 | 0.590569753 | 0.306 | 0.183 | 2.65E-15 | Tb |
| allcell | SLC15A4 | 1.47E-26 | 0.590246107 | 0.517 | 0.338 | 2.89E-22 | Tb |
| allcell | NFATC1 | 6.23E-21 | 0.590219397 | 0.434 | 0.286 | 1.23E-16 | Tb |
| allcell | KHDC4 | 4.52E-17 | 0.589747213 | 0.29 | 0.175 | 8.89E-13 | Tb |
| allcell | UIMC1 | 1.42E-24 | 0.589602508 | 0.353 | 0.2 | 2.80E-20 | Tb |
| allcell | RNF34 | 2.69E-28 | 0.589464557 | 0.369 | 0.204 | 5.30E-24 | Tb |
| allcell | AOPEP | 6.88E-23 | 0.589443477 | 0.286 | 0.158 | 1.35E-18 | Tb |
| allcell | BPTF | 1.30E-35 | 0.589357178 | 0.675 | 0.467 | 2.55E-31 | Tb |
| allcell | CELF2 | 8.54E-37 | 0.58898184 | 0.843 | 0.673 | 1.68E-32 | Tb |
| allcell | ADNP | 3.85E-27 | 0.588843289 | 0.485 | 0.308 | 7.58E-23 | Tb |
| allcell | SLC31A1 | 4.14E-16 | 0.588761138 | 0.307 | 0.195 | 8.14E-12 | Tb |
| allcell | KMT5B | 5.75E-21 | 0.587948306 | 0.334 | 0.196 | 1.13E-16 | Tb |
| allcell | NR4A3 | 8.28E-22 | 0.587431645 | 0.847 | 0.784 | 1.63E-17 | Tb |
| allcell | PIBF1 | 4.29E-18 | 0.587399337 | 0.31 | 0.191 | 8.45E-14 | Tb |
| allcell | TRERF1 | 8.28E-11 | 0.586967108 | 0.378 | 0.277 | 1.63E-06 | Tb |
| allcell | ADK | 2.74E-29 | 0.586741756 | 0.617 | 0.441 | 5.40E-25 | Tb |
| allcell | DNAJC5 | 2.99E-26 | 0.586358598 | 0.443 | 0.268 | 5.89E-22 | Tb |
| allcell | NAA15 | 4.72E-21 | 0.586154346 | 0.35 | 0.21 | 9.28E-17 | Tb |
| allcell | ANKRD11 | 2.25E-52 | 0.585755667 | 0.87 | 0.693 | 4.43E-48 | Tb |
| allcell | PI4KA | 3.64E-18 | 0.585718891 | 0.389 | 0.249 | 7.16E-14 | Tb |
| allcell | FOXJ3 | 5.27E-23 | 0.585642502 | 0.401 | 0.247 | 1.04E-18 | Tb |
| allcell | INSR | 3.59E-25 | 0.585354369 | 0.283 | 0.153 | 7.06E-21 | Tb |
| allcell | TGFBR1 | 4.90E-30 | 0.585286522 | 0.582 | 0.39 | 9.64E-26 | Tb |
| allcell | TCAIM | 2.12E-16 | 0.584692593 | 0.264 | 0.154 | 4.17E-12 | Tb |
| allcell | SCYL2 | 4.30E-28 | 0.5844111 | 0.447 | 0.267 | 8.47E-24 | Tb |
| allcell | ATF7IP | 7.13E-28 | 0.583990723 | 0.612 | 0.435 | 1.40E-23 | Tb |
| allcell | IL15 | 6.13E-17 | 0.583975788 | 0.508 | 0.379 | 1.21E-12 | Tb |
| allcell | TSPAN13 | 1.15E-15 | 0.583692386 | 0.417 | 0.299 | 2.26E-11 | Tb |
| allcell | ARID1B | 4.59E-48 | 0.583506469 | 0.792 | 0.569 | 9.03E-44 | Tb |
| allcell | FRYL | 1.05E-32 | 0.582773595 | 0.63 | 0.423 | 2.07E-28 | Tb |
| allcell | PARN | 1.42E-18 | 0.582398943 | 0.304 | 0.181 | 2.79E-14 | Tb |
| allcell | GNAQ | 1.05E-46 | 0.582298698 | 0.813 | 0.629 | 2.06E-42 | Tb |
| allcell | TTC14 | 1.42E-20 | 0.582236246 | 0.268 | 0.146 | 2.79E-16 | Tb |
| allcell | UHRF2 | 4.56E-18 | 0.580885088 | 0.271 | 0.157 | 8.97E-14 | Tb |
| allcell | SETD5 | 1.65E-34 | 0.580493596 | 0.57 | 0.361 | 3.24E-30 | Tb |
| allcell | FBXW7 | 2.84E-33 | 0.579786591 | 0.718 | 0.547 | 5.59E-29 | Tb |
| allcell | AKT1S1 | 1.38E-17 | 0.577906372 | 0.255 | 0.144 | 2.73E-13 | Tb |
| allcell | GPATCH11 | 8.11E-14 | 0.577526379 | 0.281 | 0.177 | 1.60E-09 | Tb |
| allcell | FOXO3 | 5.23E-19 | 0.576634208 | 0.49 | 0.348 | 1.03E-14 | Tb |
| allcell | AHCYL1 | 2.51E-21 | 0.576571153 | 0.46 | 0.302 | 4.94E-17 | Tb |
| allcell | MAN1A2 | 1.52E-23 | 0.576423201 | 0.448 | 0.285 | 2.98E-19 | Tb |
| allcell | FLT3 | 4.88E-08 | 0.575911926 | 0.5 | 0.42 | 0.000960262 | Tb |
| allcell | ZBTB38 | 2.96E-08 | 0.575680814 | 0.31 | 0.227 | 0.000583135 | Tb |
| allcell | CSF2RA | 3.02E-24 | 0.575469531 | 0.78 | 0.664 | 5.95E-20 | Tb |
| allcell | STX5 | 7.77E-28 | 0.575442633 | 0.507 | 0.321 | 1.53E-23 | Tb |
| allcell | ACOT13 | 6.55E-27 | 0.575354875 | 0.328 | 0.175 | 1.29E-22 | Tb |
| allcell | STX8 | 2.00E-26 | 0.575255239 | 0.562 | 0.376 | 3.94E-22 | Tb |
| allcell | RELL1 | 6.18E-24 | 0.575176567 | 0.391 | 0.24 | 1.22E-19 | Tb |
| allcell | RNF213 | 1.30E-35 | 0.574959258 | 0.702 | 0.482 | 2.56E-31 | Tb |
| allcell | PDCD4 | 2.95E-19 | 0.574923755 | 0.46 | 0.31 | 5.81E-15 | Tb |
| allcell | MED14 | 3.01E-28 | 0.574871308 | 0.331 | 0.176 | 5.92E-24 | Tb |
| allcell | NUP88 | 2.52E-19 | 0.574463125 | 0.251 | 0.138 | 4.96E-15 | Tb |
| allcell | PITPNB | 1.81E-28 | 0.574398051 | 0.55 | 0.358 | 3.57E-24 | Tb |
| allcell | GAS2L3 | 8.26E-09 | 0.573632768 | 0.268 | 0.19 | 0.000162647 | Tb |
| allcell | EGR2 | 2.02E-25 | 0.573602248 | 0.277 | 0.143 | 3.98E-21 | Tb |
| allcell | UBE3C | 6.24E-22 | 0.573595992 | 0.397 | 0.245 | 1.23E-17 | Tb |

| | | | | | | | |
|---|---|---|---|---|---|---|---|
| allcell | STX12 | 6.27E-22 | 0.573539015 | 0.628 | 0.474 | 1.23E-17 | Tb |
| allcell | OPA3 | 9.88E-23 | 0.573278774 | 0.363 | 0.214 | 1.94E-18 | Tb |
| allcell | HPS5 | 8.24E-36 | 0.572676503 | 0.671 | 0.462 | 1.62E-31 | Tb |
| allcell | TMX2 | 8.00E-25 | 0.572050543 | 0.299 | 0.157 | 1.57E-20 | Tb |
| allcell | PTPRA | 2.01E-19 | 0.571381034 | 0.44 | 0.292 | 3.96E-15 | Tb |
| allcell | CYFIP2 | 4.42E-14 | 0.570655197 | 0.352 | 0.24 | 8.70E-10 | Tb |
| allcell | MYOF | 1.27E-24 | 0.570480267 | 0.528 | 0.362 | 2.49E-20 | Tb |
| allcell | PRKAG1 | 4.80E-23 | 0.570128633 | 0.361 | 0.211 | 9.45E-19 | Tb |
| allcell | DNAJC13 | 4.14E-18 | 0.570018465 | 0.318 | 0.195 | 8.14E-14 | Tb |
| allcell | CEP85L | 6.44E-14 | 0.569538656 | 0.321 | 0.211 | 1.27E-09 | Tb |
| allcell | PRKAR2A | 1.72E-20 | 0.569223145 | 0.432 | 0.281 | 3.38E-16 | Tb |
| allcell | LRIF1 | 7.59E-37 | 0.56910037 | 0.696 | 0.479 | 1.49E-32 | Tb |
| allcell | RNF103 | 6.15E-19 | 0.569049335 | 0.329 | 0.2 | 1.21E-14 | Tb |
| allcell | CUX1 | 4.53E-33 | 0.568612561 | 0.75 | 0.573 | 8.92E-29 | Tb |
| allcell | CLSTN1 | 1.58E-21 | 0.568353882 | 0.276 | 0.152 | 3.10E-17 | Tb |
| allcell | EEPD1 | 1.77E-10 | 0.568267973 | 0.267 | 0.182 | 3.49E-06 | Tb |
| allcell | IL1R1 | 5.25E-09 | 0.568197817 | 0.348 | 0.262 | 0.000103304 | Tb |
| allcell | TLE4 | 1.15E-13 | 0.567755343 | 0.47 | 0.349 | 2.26E-09 | Tb |
| allcell | TAF1 | 1.37E-18 | 0.567314407 | 0.274 | 0.157 | 2.69E-14 | Tb |
| allcell | WDFY1 | 5.58E-18 | 0.567068211 | 0.279 | 0.164 | 1.10E-13 | Tb |
| allcell | SS18 | 5.27E-20 | 0.566742522 | 0.367 | 0.223 | 1.04E-15 | Tb |
| allcell | NAA35 | 2.68E-22 | 0.565986543 | 0.274 | 0.149 | 5.27E-18 | Tb |
| allcell | TTC3 | 6.67E-30 | 0.565932078 | 0.621 | 0.426 | 1.31E-25 | Tb |
| allcell | SEC31A | 1.09E-30 | 0.56572449 | 0.627 | 0.425 | 2.14E-26 | Tb |
| allcell | PPM1A | 2.94E-19 | 0.564946186 | 0.308 | 0.181 | 5.79E-15 | Tb |
| allcell | CHAF1A | 4.76E-22 | 0.564911419 | 0.479 | 0.321 | 9.37E-18 | Tb |
| allcell | CWC27 | 5.20E-14 | 0.564592124 | 0.255 | 0.157 | 1.02E-09 | Tb |
| allcell | ZNF292 | 1.00E-32 | 0.564127592 | 0.586 | 0.379 | 1.98E-28 | Tb |
| allcell | IKBKB | 2.91E-28 | 0.564091448 | 0.344 | 0.185 | 5.73E-24 | Tb |
| allcell | SLC41A2 | 6.00E-14 | 0.563670506 | 0.531 | 0.403 | 1.18E-09 | Tb |
| allcell | MSH3 | 2.13E-18 | 0.563609002 | 0.274 | 0.158 | 4.19E-14 | Tb |
| allcell | LPCAT3 | 2.71E-25 | 0.563272007 | 0.251 | 0.123 | 5.33E-21 | Tb |
| allcell | KRCC1 | 1.15E-23 | 0.563220997 | 0.292 | 0.158 | 2.27E-19 | Tb |
| allcell | PHKB | 3.05E-21 | 0.562994702 | 0.411 | 0.26 | 6.00E-17 | Tb |
| allcell | PARP11 | 5.93E-18 | 0.562416782 | 0.277 | 0.161 | 1.17E-13 | Tb |
| allcell | FAM193A | 9.02E-19 | 0.561585847 | 0.335 | 0.206 | 1.78E-14 | Tb |
| allcell | RNGTT | 4.66E-20 | 0.561472556 | 0.365 | 0.226 | 9.17E-16 | Tb |
| allcell | FAM241A | 3.23E-18 | 0.561396177 | 0.512 | 0.365 | 6.36E-14 | Tb |
| allcell | EAF2 | 2.09E-19 | 0.560759711 | 0.479 | 0.324 | 4.11E-15 | Tb |
| allcell | SLC1A4 | 3.85E-21 | 0.560547853 | 0.321 | 0.188 | 7.58E-17 | Tb |
| allcell | RAB11A | 2.20E-29 | 0.560107106 | 0.819 | 0.697 | 4.32E-25 | Tb |
| allcell | ATP11C | 3.17E-18 | 0.559735977 | 0.307 | 0.187 | 6.24E-14 | Tb |
| allcell | EXOSC6 | 1.75E-14 | 0.559112034 | 0.272 | 0.168 | 3.45E-10 | Tb |
| allcell | CD86 | 2.71E-09 | 0.558754359 | 0.726 | 0.675 | 5.34E-05 | Tb |
| allcell | MAP4 | 4.77E-28 | 0.558199874 | 0.632 | 0.446 | 9.39E-24 | Tb |
| allcell | RPF2 | 1.69E-16 | 0.557790967 | 0.295 | 0.178 | 3.33E-12 | Tb |
| allcell | PDE8A | 5.23E-17 | 0.557055412 | 0.339 | 0.219 | 1.03E-12 | Tb |
| allcell | PTPRJ | 4.00E-40 | 0.556788399 | 0.749 | 0.572 | 7.88E-36 | Tb |
| allcell | CCNYL1 | 7.98E-21 | 0.556436246 | 0.453 | 0.297 | 1.57E-16 | Tb |
| allcell | TNPO3 | 8.79E-23 | 0.556339758 | 0.38 | 0.23 | 1.73E-18 | Tb |
| allcell | ATF6 | 7.31E-21 | 0.556322106 | 0.636 | 0.473 | 1.44E-16 | Tb |
| allcell | TIAM11 | 5.35E-18 | 0.555094708 | 0.418 | 0.285 | 1.05E-13 | Tb |
| allcell | MAML3 | 1.71E-17 | 0.554707126 | 0.529 | 0.397 | 3.37E-13 | Tb |
| allcell | NR6A1 | 1.89E-15 | 0.554359054 | 0.262 | 0.158 | 3.73E-11 | Tb |
| allcell | PPM1K | 2.57E-26 | 0.554118671 | 0.491 | 0.311 | 5.06E-22 | Tb |
| allcell | SPECC1 | 2.91E-13 | 0.553830481 | 0.458 | 0.333 | 5.74E-09 | Tb |

| | | | | | | | |
|---|---|---|---|---|---|---|---|
| allcell | PIK3R5 | 1.81E-22 | 0.553651931 | 0.622 | 0.471 | 3.56E-18 | Tb |
| allcell | APTX | 4.16E-22 | 0.553640524 | 0.385 | 0.234 | 8.18E-18 | Tb |
| allcell | MCPH1 | 9.29E-14 | 0.553432964 | 0.28 | 0.18 | 1.83E-09 | Tb |
| allcell | SULF2 | 2.14E-28 | 0.552819065 | 0.735 | 0.564 | 4.20E-24 | Tb |
| allcell | USO1 | 7.34E-26 | 0.552174996 | 0.498 | 0.322 | 1.45E-21 | Tb |
| allcell | TSPOAP1-AS1 | 6.26E-21 | 0.552014278 | 0.345 | 0.206 | 1.23E-16 | Tb |
| allcell | HMGB2 | 2.24E-19 | 0.551415962 | 0.556 | 0.392 | 4.40E-15 | Tb |
| allcell | ESCO1 | 8.62E-21 | 0.551233052 | 0.386 | 0.238 | 1.70E-16 | Tb |
| allcell | SPEN | 8.16E-20 | 0.550907458 | 0.599 | 0.445 | 1.61E-15 | Tb |
| allcell | WNK1 | 3.57E-26 | 0.550777885 | 0.672 | 0.502 | 7.03E-22 | Tb |
| allcell | SLC4A7 | 7.75E-20 | 0.550292684 | 0.479 | 0.32 | 1.52E-15 | Tb |
| allcell | CTCF | 3.29E-21 | 0.54977157 | 0.36 | 0.215 | 6.47E-17 | Tb |
| allcell | FAM13B | 1.01E-22 | 0.549159628 | 0.495 | 0.332 | 1.99E-18 | Tb |
| allcell | AMBRA1 | 6.59E-26 | 0.549075249 | 0.526 | 0.343 | 1.30E-21 | Tb |
| allcell | MIR4435-2HG | 7.24E-29 | 0.549036012 | 0.658 | 0.473 | 1.43E-24 | Tb |
| allcell | EFL1 | 2.10E-15 | 0.548758439 | 0.254 | 0.15 | 4.13E-11 | Tb |
| allcell | KANSL1L | 1.20E-16 | 0.548422469 | 0.355 | 0.227 | 2.36E-12 | Tb |
| allcell | DMTF1 | 6.19E-20 | 0.548371888 | 0.283 | 0.16 | 1.22E-15 | Tb |
| allcell | MTF2 | 1.03E-24 | 0.548329738 | 0.383 | 0.225 | 2.03E-20 | Tb |
| allcell | PPP6R3 | 8.99E-29 | 0.548241367 | 0.633 | 0.439 | 1.77E-24 | Tb |
| allcell | SPPL3 | 8.93E-25 | 0.548091987 | 0.6 | 0.431 | 1.76E-20 | Tb |
| allcell | CASC4 | 7.08E-11 | 0.547893931 | 0.328 | 0.233 | 1.39E-06 | Tb |
| allcell | MAPK8 | 6.81E-28 | 0.547079865 | 0.569 | 0.391 | 1.34E-23 | Tb |
| allcell | CAAP1 | 2.17E-17 | 0.546252621 | 0.324 | 0.201 | 4.27E-13 | Tb |
| allcell | GPATCH8 | 4.17E-20 | 0.54595072 | 0.465 | 0.317 | 8.21E-16 | Tb |
| allcell | TVP23C | 3.68E-18 | 0.545443931 | 0.25 | 0.142 | 7.24E-14 | Tb |
| allcell | PWWP2A | 1.38E-16 | 0.545125518 | 0.269 | 0.159 | 2.71E-12 | Tb |
| allcell | COG5 | 5.71E-25 | 0.545056959 | 0.5 | 0.323 | 1.12E-20 | Tb |
| allcell | EIPR1 | 1.39E-18 | 0.545010996 | 0.262 | 0.148 | 2.74E-14 | Tb |
| allcell | ZNF644 | 2.18E-28 | 0.544886939 | 0.635 | 0.447 | 4.29E-24 | Tb |
| allcell | TAOK3 | 2.59E-35 | 0.543905327 | 0.849 | 0.721 | 5.10E-31 | Tb |
| allcell | CCNG2 | 3.21E-20 | 0.54385253 | 0.463 | 0.312 | 6.32E-16 | Tb |
| allcell | DNAJC1 | 6.17E-19 | 0.543527507 | 0.616 | 0.47 | 1.21E-14 | Tb |
| allcell | PRRC2B | 4.29E-21 | 0.543462062 | 0.437 | 0.283 | 8.44E-17 | Tb |
| allcell | GLB1 | 1.54E-21 | 0.542819081 | 0.361 | 0.214 | 3.02E-17 | Tb |
| allcell | RAB6A | 4.07E-18 | 0.542541862 | 0.436 | 0.29 | 8.02E-14 | Tb |
| allcell | ASMTL | 6.30E-22 | 0.542482692 | 0.461 | 0.306 | 1.24E-17 | Tb |
| allcell | PIP4K2A | 6.24E-20 | 0.541681517 | 0.579 | 0.417 | 1.23E-15 | Tb |
| allcell | ZDHHC17 | 1.63E-17 | 0.541584957 | 0.438 | 0.299 | 3.20E-13 | Tb |
| allcell | SLC8A1 | 2.19E-15 | 0.541291994 | 0.691 | 0.622 | 4.31E-11 | Tb |
| allcell | APBB1IP | 6.37E-16 | 0.541235837 | 0.652 | 0.517 | 1.25E-11 | Tb |
| allcell | WSB2 | 7.49E-17 | 0.541211327 | 0.311 | 0.19 | 1.47E-12 | Tb |
| allcell | NETO2 | 1.81E-07 | 0.540834661 | 0.262 | 0.192 | 0.003557885 | Tb |
| allcell | PPM1B | 3.42E-19 | 0.540797806 | 0.446 | 0.304 | 6.74E-15 | Tb |
| allcell | RABGEF1 | 1.19E-24 | 0.54054796 | 0.658 | 0.497 | 2.34E-20 | Tb |
| allcell | LYST | 2.54E-15 | 0.540247327 | 0.548 | 0.415 | 5.00E-11 | Tb |
| allcell | GALNT2 | 1.17E-15 | 0.539977833 | 0.403 | 0.278 | 2.30E-11 | Tb |
| allcell | UBR3 | 2.10E-22 | 0.539096654 | 0.348 | 0.205 | 4.14E-18 | Tb |
| allcell | DCTN4 | 7.54E-21 | 0.538835626 | 0.465 | 0.308 | 1.48E-16 | Tb |
| allcell | CTDSPL2 | 1.46E-18 | 0.538715444 | 0.354 | 0.221 | 2.87E-14 | Tb |
| allcell | PTEN | 2.72E-28 | 0.538705946 | 0.661 | 0.473 | 5.36E-24 | Tb |
| allcell | PIK3C3 | 2.88E-18 | 0.538275063 | 0.297 | 0.175 | 5.66E-14 | Tb |
| allcell | ATP6V1A | 4.74E-18 | 0.537920765 | 0.576 | 0.439 | 9.34E-14 | Tb |
| allcell | GGA3 | 3.35E-21 | 0.53786444 | 0.25 | 0.131 | 6.59E-17 | Tb |
| allcell | SCAPER | 3.28E-18 | 0.537674849 | 0.362 | 0.23 | 6.45E-14 | Tb |
| allcell | ZNF131 | 3.90E-24 | 0.537586068 | 0.453 | 0.289 | 7.68E-20 | Tb |

| | | | | | | | |
|---|---|---|---|---|---|---|---|
| allcell | CUL1 | 7.77E-20 | 0.536991761 | 0.624 | 0.468 | 1.53E-15 | Tb |
| allcell | SUMF1 | 9.41E-28 | 0.536928033 | 0.533 | 0.348 | 1.85E-23 | Tb |
| allcell | ABHD17B | 6.37E-14 | 0.536270341 | 0.34 | 0.227 | 1.25E-09 | Tb |
| allcell | TM9SF4 | 6.81E-19 | 0.535864344 | 0.252 | 0.139 | 1.34E-14 | Tb |
| allcell | DCAF6 | 1.53E-20 | 0.535511571 | 0.389 | 0.246 | 3.02E-16 | Tb |
| allcell | BTAF1 | 6.69E-28 | 0.535248482 | 0.615 | 0.42 | 1.32E-23 | Tb |
| allcell | PTRH2 | 1.59E-17 | 0.535098687 | 0.308 | 0.185 | 3.12E-13 | Tb |
| allcell | LARP4B | 1.48E-35 | 0.534925635 | 0.635 | 0.415 | 2.91E-31 | Tb |
| allcell | RNF149 | 9.86E-16 | 0.534842976 | 0.777 | 0.671 | 1.94E-11 | Tb |
| allcell | NOCT | 1.10E-15 | 0.534682532 | 0.289 | 0.18 | 2.17E-11 | Tb |
| allcell | LPP | 3.20E-40 | 0.534667799 | 0.78 | 0.585 | 6.29E-36 | Tb |
| allcell | KIF2A | 4.18E-13 | 0.534002467 | 0.629 | 0.496 | 8.23E-09 | Tb |
| allcell | KLHL28 | 1.23E-20 | 0.533887815 | 0.302 | 0.173 | 2.43E-16 | Tb |
| allcell | RSPRY1 | 4.81E-18 | 0.533660006 | 0.292 | 0.172 | 9.48E-14 | Tb |
| allcell | XIAP | 3.76E-19 | 0.533471261 | 0.361 | 0.223 | 7.40E-15 | Tb |
| allcell | MTMR6 | 7.96E-21 | 0.533273346 | 0.407 | 0.251 | 1.57E-16 | Tb |
| allcell | HIVEP1 | 2.13E-23 | 0.533179019 | 0.752 | 0.635 | 4.19E-19 | Tb |
| allcell | HECTD1 | 4.84E-27 | 0.533051867 | 0.512 | 0.326 | 9.53E-23 | Tb |
| allcell | PDCD7 | 7.23E-17 | 0.532845595 | 0.321 | 0.199 | 1.42E-12 | Tb |
| allcell | TUG1 | 3.13E-17 | 0.532706855 | 0.253 | 0.146 | 6.17E-13 | Tb |
| allcell | GCC2 | 5.74E-26 | 0.53179126 | 0.586 | 0.402 | 1.13E-21 | Tb |
| allcell | FAM117B | 1.17E-07 | 0.53141782 | 0.279 | 0.207 | 0.002309066 | Tb |
| allcell | APC | 1.66E-17 | 0.531365456 | 0.348 | 0.22 | 3.26E-13 | Tb |
| allcell | ATF5 | 6.79E-13 | 0.531062185 | 0.659 | 0.539 | 1.34E-08 | Tb |
| allcell | ATP8A1 | 1.79E-12 | 0.531053199 | 0.273 | 0.179 | 3.52E-08 | Tb |
| allcell | SNAPC1 | 1.06E-07 | 0.530979133 | 0.257 | 0.183 | 0.002084509 | Tb |
| allcell | PDXDC1 | 6.98E-24 | 0.530899893 | 0.383 | 0.226 | 1.37E-19 | Tb |
| allcell | WDFY2 | 2.50E-30 | 0.530789462 | 0.601 | 0.414 | 4.92E-26 | Tb |
| allcell | MBTPS1 | 2.37E-10 | 0.530488413 | 0.253 | 0.167 | 4.67E-06 | Tb |
| allcell | OFD1 | 3.53E-28 | 0.530461783 | 0.572 | 0.393 | 6.95E-24 | Tb |
| allcell | ETNK1 | 2.51E-30 | 0.529897865 | 0.6 | 0.397 | 4.95E-26 | Tb |
| allcell | TBC1D5 | 8.25E-24 | 0.529815066 | 0.626 | 0.464 | 1.62E-19 | Tb |
| allcell | KLHL18 | 3.38E-14 | 0.529483262 | 0.281 | 0.175 | 6.65E-10 | Tb |
| allcell | SOS1 | 9.20E-26 | 0.529135304 | 0.542 | 0.365 | 1.81E-21 | Tb |
| allcell | RC3H2 | 4.59E-20 | 0.528887431 | 0.348 | 0.214 | 9.04E-16 | Tb |
| allcell | RAPGEF6 | 6.07E-22 | 0.528853996 | 0.57 | 0.402 | 1.20E-17 | Tb |
| allcell | NBAS | 2.04E-21 | 0.528553854 | 0.344 | 0.204 | 4.01E-17 | Tb |
| allcell | TRAPPC8 | 4.30E-20 | 0.528418819 | 0.338 | 0.204 | 8.46E-16 | Tb |
| allcell | CCAR1 | 1.06E-27 | 0.528034543 | 0.517 | 0.331 | 2.08E-23 | Tb |
| allcell | CD274 | 4.11E-15 | 0.527091474 | 0.346 | 0.234 | 8.09E-11 | Tb |
| allcell | IL4R | 3.92E-20 | 0.527026753 | 0.397 | 0.252 | 7.71E-16 | Tb |
| allcell | USP14 | 1.07E-18 | 0.526872295 | 0.486 | 0.332 | 2.10E-14 | Tb |
| allcell | ZNF91 | 7.75E-19 | 0.52671278 | 0.302 | 0.175 | 1.53E-14 | Tb |
| allcell | MTMR12 | 1.52E-19 | 0.526380274 | 0.311 | 0.182 | 2.98E-15 | Tb |
| allcell | TLK2 | 4.24E-23 | 0.526290563 | 0.425 | 0.262 | 8.34E-19 | Tb |
| allcell | FNIP1 | 5.39E-31 | 0.526167294 | 0.604 | 0.401 | 1.06E-26 | Tb |
| allcell | AFF4 | 5.08E-34 | 0.526001273 | 0.815 | 0.662 | 1.00E-29 | Tb |
| allcell | DGKH | 3.99E-17 | 0.525750716 | 0.279 | 0.167 | 7.85E-13 | Tb |
| allcell | UBASH3B | 1.55E-11 | 0.525706271 | 0.437 | 0.325 | 3.05E-07 | Tb |
| allcell | NFE2L3 | 1.63E-15 | 0.525460937 | 0.547 | 0.405 | 3.21E-11 | Tb |
| allcell | TSC22D2 | 4.75E-21 | 0.525334507 | 0.621 | 0.461 | 9.34E-17 | Tb |
| allcell | OTUD5 | 1.79E-18 | 0.52529715 | 0.36 | 0.227 | 3.52E-14 | Tb |
| allcell | ATP9B | 1.00E-19 | 0.525122348 | 0.559 | 0.404 | 1.97E-15 | Tb |
| allcell | MINK1 | 1.41E-22 | 0.524958986 | 0.299 | 0.166 | 2.78E-18 | Tb |
| allcell | TFDP2 | 2.75E-20 | 0.524958645 | 0.421 | 0.268 | 5.41E-16 | Tb |
| allcell | SLC30A5 | 5.43E-15 | 0.524208894 | 0.276 | 0.17 | 1.07E-10 | Tb |

| | | | | | | | |
|---|---|---|---|---|---|---|---|
| allcell | ARID3A | 5.89E-16 | 0.523920815 | 0.441 | 0.307 | 1.16E-11 | Tb |
| allcell | CASK | 1.05E-15 | 0.523778916 | 0.304 | 0.191 | 2.07E-11 | Tb |
| allcell | ACO2 | 4.56E-22 | 0.523750667 | 0.419 | 0.261 | 8.98E-18 | Tb |
| allcell | FH | 3.31E-14 | 0.523502504 | 0.353 | 0.232 | 6.52E-10 | Tb |
| allcell | NUP153 | 7.93E-23 | 0.52329109 | 0.411 | 0.253 | 1.56E-18 | Tb |
| allcell | SLC30A9 | 5.41E-22 | 0.523252899 | 0.355 | 0.21 | 1.07E-17 | Tb |
| allcell | VPS13D | 5.26E-20 | 0.522247033 | 0.451 | 0.307 | 1.04E-15 | Tb |
| allcell | GMDS | 1.95E-19 | 0.52198051 | 0.562 | 0.406 | 3.84E-15 | Tb |
| allcell | UBE2H | 3.28E-28 | 0.521687733 | 0.693 | 0.521 | 6.45E-24 | Tb |
| allcell | TBC1D14 | 8.13E-18 | 0.521300505 | 0.341 | 0.213 | 1.60E-13 | Tb |
| allcell | IGF2R | 1.60E-33 | 0.521242688 | 0.696 | 0.479 | 3.15E-29 | Tb |
| allcell | JAK2 | 2.04E-23 | 0.521074082 | 0.714 | 0.555 | 4.01E-19 | Tb |
| allcell | HERC1 | 1.51E-24 | 0.520671666 | 0.627 | 0.453 | 2.97E-20 | Tb |
| allcell | ZNF506 | 3.20E-16 | 0.52062814 | 0.285 | 0.171 | 6.29E-12 | Tb |
| allcell | FAM49B | 1.70E-37 | 0.520458956 | 0.916 | 0.783 | 3.35E-33 | Tb |
| allcell | ATM | 8.08E-18 | 0.520241954 | 0.357 | 0.23 | 1.59E-13 | Tb |
| allcell | ST3GAL4 | 7.81E-18 | 0.519700353 | 0.277 | 0.161 | 1.54E-13 | Tb |
| allcell | ACSL3 | 8.56E-16 | 0.519674704 | 0.64 | 0.491 | 1.69E-11 | Tb |
| allcell | CYTOR | 2.72E-16 | 0.519662396 | 0.691 | 0.584 | 5.36E-12 | Tb |
| allcell | MTR | 9.57E-20 | 0.519582108 | 0.316 | 0.188 | 1.88E-15 | Tb |
| allcell | MRPL1 | 2.50E-19 | 0.519360505 | 0.308 | 0.182 | 4.92E-15 | Tb |
| allcell | AP2B1 | 9.57E-23 | 0.519294254 | 0.5 | 0.326 | 1.88E-18 | Tb |
| allcell | KCTD5 | 6.37E-16 | 0.519273216 | 0.346 | 0.223 | 1.25E-11 | Tb |
| allcell | NUS1 | 1.23E-16 | 0.519169618 | 0.301 | 0.182 | 2.42E-12 | Tb |
| allcell | CELF1 | 1.63E-21 | 0.519133997 | 0.486 | 0.32 | 3.20E-17 | Tb |
| allcell | ZBTB10 | 1.57E-16 | 0.519062313 | 0.593 | 0.462 | 3.08E-12 | Tb |
| allcell | WDR7 | 2.30E-24 | 0.519048833 | 0.368 | 0.215 | 4.52E-20 | Tb |
| allcell | ATP11A | 5.05E-21 | 0.519034532 | 0.358 | 0.215 | 9.94E-17 | Tb |
| allcell | N4BP2L2 | 2.47E-34 | 0.517693321 | 0.74 | 0.529 | 4.85E-30 | Tb |
| allcell | SESN3 | 2.49E-10 | 0.517654682 | 0.397 | 0.3 | 4.91E-06 | Tb |
| allcell | TNKS | 7.45E-17 | 0.5173738 | 0.378 | 0.25 | 1.47E-12 | Tb |
| allcell | SWAP70 | 4.03E-20 | 0.516846516 | 0.485 | 0.338 | 7.93E-16 | Tb |
| allcell | TRIM69 | 5.65E-24 | 0.51620785 | 0.586 | 0.422 | 1.11E-19 | Tb |
| allcell | CEP68 | 1.64E-23 | 0.515618197 | 0.438 | 0.275 | 3.23E-19 | Tb |
| allcell | KAT2B | 7.20E-24 | 0.515220199 | 0.578 | 0.404 | 1.42E-19 | Tb |
| allcell | ITGA4 | 5.20E-17 | 0.515109135 | 0.651 | 0.517 | 1.02E-12 | Tb |
| allcell | OSGIN2 | 1.08E-17 | 0.514926587 | 0.261 | 0.149 | 2.12E-13 | Tb |
| allcell | TASOR | 1.43E-18 | 0.514878553 | 0.346 | 0.215 | 2.81E-14 | Tb |
| allcell | SNX13 | 8.88E-16 | 0.514479823 | 0.365 | 0.237 | 1.75E-11 | Tb |
| allcell | HSD17B4 | 5.49E-17 | 0.514119042 | 0.346 | 0.217 | 1.08E-12 | Tb |
| allcell | ASXL2 | 1.31E-19 | 0.514093159 | 0.369 | 0.232 | 2.57E-15 | Tb |
| allcell | CNTLN | 2.66E-15 | 0.513762865 | 0.356 | 0.237 | 5.24E-11 | Tb |
| allcell | MAPK14 | 7.93E-21 | 0.513748322 | 0.398 | 0.25 | 1.56E-16 | Tb |
| allcell | BANP | 5.26E-20 | 0.513461859 | 0.646 | 0.491 | 1.04E-15 | Tb |
| allcell | PHF20L1 | 1.32E-16 | 0.512629396 | 0.507 | 0.358 | 2.61E-12 | Tb |
| allcell | PARP8 | 5.20E-19 | 0.512141798 | 0.463 | 0.321 | 1.02E-14 | Tb |
| allcell | YLPM1 | 9.93E-16 | 0.511849042 | 0.291 | 0.178 | 1.95E-11 | Tb |
| allcell | CCDC66 | 5.68E-15 | 0.511690338 | 0.271 | 0.165 | 1.12E-10 | Tb |
| allcell | HNRNPUL1 | 6.58E-25 | 0.511673249 | 0.649 | 0.462 | 1.30E-20 | Tb |
| allcell | ZCCHC17 | 1.21E-19 | 0.511592418 | 0.359 | 0.218 | 2.38E-15 | Tb |
| allcell | TBC1D9 | 5.10E-16 | 0.511326277 | 0.487 | 0.352 | 1.00E-11 | Tb |
| allcell | KIF13B | 3.02E-11 | 0.510623743 | 0.391 | 0.283 | 5.95E-07 | Tb |
| allcell | AAK1 | 3.81E-17 | 0.51025441 | 0.376 | 0.245 | 7.50E-13 | Tb |
| allcell | CEP63 | 2.13E-14 | 0.510031435 | 0.325 | 0.21 | 4.19E-10 | Tb |
| allcell | SND1 | 2.67E-31 | 0.509692193 | 0.742 | 0.53 | 5.26E-27 | Tb |
| allcell | STAU1 | 6.08E-23 | 0.509661569 | 0.547 | 0.364 | 1.20E-18 | Tb |

| | | | | | | | |
|---|---|---|---|---|---|---|---|
| allcell | CALU | 7.28E-17 | 0.509500401 | 0.343 | 0.217 | 1.43E-12 | Tb |
| allcell | CSPP1 | 8.39E-13 | 0.509367187 | 0.26 | 0.164 | 1.65E-08 | Tb |
| allcell | QSOX1 | 2.37E-15 | 0.508328692 | 0.295 | 0.184 | 4.67E-11 | Tb |
| allcell | SSH2 | 2.20E-22 | 0.508119024 | 0.609 | 0.446 | 4.33E-18 | Tb |
| allcell | FNDC3A | 1.85E-25 | 0.507955964 | 0.665 | 0.491 | 3.64E-21 | Tb |
| allcell | BBX | 7.91E-26 | 0.507816608 | 0.617 | 0.437 | 1.56E-21 | Tb |
| allcell | IMMP2L | 6.93E-11 | 0.507662367 | 0.385 | 0.287 | 1.36E-06 | Tb |
| allcell | FRMD4B | 3.93E-14 | 0.507392155 | 0.476 | 0.353 | 7.74E-10 | Tb |
| allcell | SPN | 6.85E-23 | 0.507288563 | 0.393 | 0.242 | 1.35E-18 | Tb |
| allcell | PPM1D | 5.20E-18 | 0.50681776 | 0.39 | 0.252 | 1.02E-13 | Tb |
| allcell | FLNB | 2.37E-15 | 0.50681145 | 0.281 | 0.178 | 4.66E-11 | Tb |
| allcell | BCOR | 2.60E-19 | 0.506611641 | 0.276 | 0.155 | 5.12E-15 | Tb |
| allcell | ADGRE5 | 2.40E-20 | 0.506518465 | 0.601 | 0.442 | 4.73E-16 | Tb |
| allcell | SIL1 | 3.61E-19 | 0.506242148 | 0.462 | 0.315 | 7.11E-15 | Tb |
| allcell | AKAP13 | 4.86E-38 | 0.505723582 | 0.952 | 0.878 | 9.56E-34 | Tb |
| allcell | SLC7A5 | 1.80E-16 | 0.505536835 | 0.586 | 0.447 | 3.55E-12 | Tb |
| allcell | PXK | 2.92E-17 | 0.504897646 | 0.29 | 0.173 | 5.76E-13 | Tb |
| allcell | HMGCS1 | 1.58E-14 | 0.504773622 | 0.432 | 0.299 | 3.11E-10 | Tb |
| allcell | DHRSX | 8.65E-23 | 0.504689416 | 0.463 | 0.295 | 1.70E-18 | Tb |
| allcell | ATG10 | 9.77E-16 | 0.504548508 | 0.251 | 0.149 | 1.92E-11 | Tb |
| allcell | ZC3H13 | 3.65E-23 | 0.50438064 | 0.481 | 0.307 | 7.18E-19 | Tb |
| allcell | ST14 | 1.70E-13 | 0.504254592 | 0.252 | 0.159 | 3.35E-09 | Tb |
| allcell | MDM4 | 1.65E-20 | 0.50376335 | 0.495 | 0.331 | 3.24E-16 | Tb |
| allcell | ATAD2B | 1.64E-27 | 0.503216212 | 0.648 | 0.456 | 3.23E-23 | Tb |
| allcell | LRRC8B | 1.17E-20 | 0.503193931 | 0.319 | 0.185 | 2.31E-16 | Tb |
| allcell | DENND10 | 1.37E-14 | 0.503115094 | 0.255 | 0.151 | 2.69E-10 | Tb |
| allcell | SLC35F5 | 1.57E-15 | 0.503039884 | 0.281 | 0.17 | 3.08E-11 | Tb |
| allcell | AKNA | 9.57E-20 | 0.503032004 | 0.42 | 0.276 | 1.88E-15 | Tb |
| allcell | RAB4B | 1.14E-23 | 0.503014501 | 0.472 | 0.301 | 2.24E-19 | Tb |
| allcell | CHSY1 | 1.05E-12 | 0.502738927 | 0.254 | 0.161 | 2.06E-08 | Tb |
| allcell | MKNK1 | 8.51E-18 | 0.502620705 | 0.304 | 0.183 | 1.67E-13 | Tb |
| allcell | TAF2 | 3.37E-14 | 0.502156232 | 0.283 | 0.179 | 6.63E-10 | Tb |
| allcell | XPNPEP1 | 2.14E-19 | 0.501002316 | 0.317 | 0.186 | 4.21E-15 | Tb |
| allcell | POLG2 | 5.31E-10 | 0.500138467 | 0.304 | 0.213 | 1.05E-05 | Tb |
| **Disease-associated genes in cDC1** | | | | | | | |
| cDC1 | SERPINH1 | 1.48E-11 | 1.690831063 | 0.509 | 0.31 | 2.91E-07 | Lep-L |
| cDC1 | HSPA1B | 3.75E-14 | 1.62050307 | 0.947 | 0.918 | 7.38E-10 | Lep-L |
| cDC1 | BAG3 | 1.68E-16 | 1.541454211 | 0.825 | 0.711 | 3.32E-12 | Lep-L |
| cDC1 | DDX3Y | 2.78E-10 | 1.471482892 | 0.515 | 0.309 | 5.47E-06 | Lep-L |
| cDC1 | CXCL8 | 1.21E-09 | 1.439038556 | 0.561 | 0.37 | 2.39E-05 | Lep-L |
| cDC1 | TSPYL2 | 2.59E-09 | 1.429663545 | 0.602 | 0.461 | 5.10E-05 | Lep-L |
| cDC1 | NR4A1 | 9.10E-12 | 1.429229741 | 0.883 | 0.88 | 1.79E-07 | Lep-L |
| cDC1 | FKBP4 | 1.48E-12 | 1.421490354 | 0.591 | 0.383 | 2.91E-08 | Lep-L |
| cDC1 | MRPS6 | 2.59E-09 | 1.350028043 | 0.708 | 0.647 | 5.09E-05 | Lep-L |
| cDC1 | MOB4 | 1.41E-09 | 1.348897963 | 0.69 | 0.624 | 2.77E-05 | Lep-L |
| cDC1 | SLC5A3 | 7.97E-07 | 1.311038118 | 0.532 | 0.393 | 0.015679961 | Lep-L |
| cDC1 | SOX4 | 1.91E-08 | 1.302890053 | 0.667 | 0.522 | 0.000375591 | Lep-L |
| cDC1 | HSPA6 | 2.58E-09 | 1.276427828 | 0.526 | 0.328 | 5.07E-05 | Lep-L |
| cDC1 | NRARP | 4.32E-12 | 1.264630133 | 0.766 | 0.705 | 8.50E-08 | Lep-L |
| cDC1 | ZFP36L1 | 1.16E-09 | 1.231857051 | 0.772 | 0.806 | 2.29E-05 | Lep-L |
| cDC1 | THAP9-AS1 | 1.38E-07 | 1.218559513 | 0.538 | 0.407 | 0.002707187 | Lep-L |
| cDC1 | RILPL2 | 2.93E-09 | 1.213001006 | 0.702 | 0.664 | 5.77E-05 | Lep-L |
| cDC1 | HLA-DRB5 | 6.25E-16 | 1.204294838 | 0.825 | 0.824 | 1.23E-11 | Lep-L |
| cDC1 | HSPH1 | 1.17E-27 | 1.200520167 | 0.93 | 0.897 | 2.29E-23 | Lep-L |
| cDC1 | ATP1A1 | 3.63E-15 | 1.176424984 | 0.871 | 0.875 | 7.14E-11 | Lep-L |
| cDC1 | BRD2 | 2.23E-06 | 1.143927579 | 0.789 | 0.812 | 0.04397468 | Lep-L |

| | | | | | | | |
|---|---|---|---|---|---|---|---|
| cDC1 | CHORDC1 | 5.67E-14 | 1.141157663 | 0.754 | 0.648 | 1.12E-09 | Lep-L |
| cDC1 | HSPB1 | 2.59E-12 | 1.128395827 | 0.778 | 0.72 | 5.10E-08 | Lep-L |
| cDC1 | HSPD1 | 1.22E-23 | 1.127618233 | 0.918 | 0.924 | 2.39E-19 | Lep-L |
| cDC1 | CD55 | 3.81E-08 | 1.121813751 | 0.83 | 0.821 | 0.00075001 | Lep-L |
| cDC1 | APOE | 2.01E-09 | 1.118936972 | 0.427 | 0.265 | 3.96E-05 | Lep-L |
| cDC1 | DNAJB1 | 2.02E-22 | 1.090010728 | 0.953 | 0.917 | 3.98E-18 | Lep-L |
| cDC1 | DNAJA1 | 2.35E-23 | 1.087325156 | 0.936 | 0.943 | 4.63E-19 | Lep-L |
| cDC1 | DEDD2 | 4.18E-07 | 1.057981449 | 0.55 | 0.442 | 0.008237296 | Lep-L |
| cDC1 | IER5 | 2.88E-14 | 1.04462231 | 0.895 | 0.872 | 5.67E-10 | Lep-L |
| cDC1 | HSPA1A | 1.29E-20 | 1.041265833 | 0.982 | 0.971 | 2.54E-16 | Lep-L |
| cDC1 | HSPE1 | 5.66E-16 | 1.032044618 | 0.906 | 0.93 | 1.11E-11 | Lep-L |
| cDC1 | PTGES3 | 6.88E-08 | 1.01126449 | 0.76 | 0.812 | 0.001353677 | Lep-L |
| cDC1 | SLC38A2 | 7.02E-08 | 1.008194419 | 0.784 | 0.833 | 0.001381165 | Lep-L |
| cDC1 | EZR | 2.31E-17 | 1.000319116 | 0.877 | 0.885 | 4.55E-13 | Lep-L |
| cDC1 | HSP90AA1 | 4.94E-25 | 0.965557071 | 0.977 | 0.982 | 9.73E-21 | Lep-L |
| cDC1 | HSP90AB1 | 2.87E-23 | 0.919136515 | 0.959 | 0.973 | 5.64E-19 | Lep-L |
| cDC1 | CACYBP | 1.40E-13 | 0.915380986 | 0.819 | 0.819 | 2.76E-09 | Lep-L |
| cDC1 | MCL1 | 1.43E-20 | 0.903712102 | 0.901 | 0.932 | 2.81E-16 | Lep-L |
| cDC1 | UBC | 8.27E-19 | 0.903435482 | 0.971 | 0.983 | 1.63E-14 | Lep-L |
| cDC1 | PPP1R15A | 1.68E-06 | 0.894872831 | 0.918 | 0.939 | 0.033018955 | Lep-L |
| cDC1 | SAT1 | 2.24E-20 | 0.875967778 | 0.988 | 0.992 | 4.41E-16 | Lep-L |
| cDC1 | CHD2 | 4.15E-07 | 0.875631824 | 0.766 | 0.821 | 0.00817476 | Lep-L |
| cDC1 | GPBP1 | 4.64E-12 | 0.865725489 | 0.877 | 0.877 | 9.14E-08 | Lep-L |
| cDC1 | ZC3HAV1 | 5.08E-08 | 0.853930298 | 0.725 | 0.664 | 0.000999474 | Lep-L |
| cDC1 | CKS2 | 3.79E-07 | 0.824800628 | 0.819 | 0.847 | 0.007452253 | Lep-L |
| cDC1 | MAP2K3 | 7.78E-08 | 0.823847286 | 0.725 | 0.667 | 0.001531845 | Lep-L |
| cDC1 | BCL2A1 | 3.27E-08 | 0.807505323 | 0.836 | 0.859 | 0.00064377 | Lep-L |
| cDC1 | TCP1 | 4.39E-07 | 0.805912938 | 0.725 | 0.745 | 0.008632603 | Lep-L |
| cDC1 | ZFAND2A | 4.92E-10 | 0.803759567 | 0.749 | 0.626 | 9.69E-06 | Lep-L |
| cDC1 | MRPL18 | 1.17E-06 | 0.794735409 | 0.778 | 0.817 | 0.022947854 | Lep-L |
| cDC1 | JMJD1C | 5.62E-10 | 0.785821108 | 0.86 | 0.892 | 1.11E-05 | Lep-L |
| cDC1 | CLK1 | 3.45E-09 | 0.784184799 | 0.819 | 0.793 | 6.79E-05 | Lep-L |
| cDC1 | DNAJB6 | 2.78E-10 | 0.738023526 | 0.83 | 0.869 | 5.47E-06 | Lep-L |
| cDC1 | JUND | 9.64E-20 | 0.718762098 | 0.953 | 0.969 | 1.90E-15 | Lep-L |
| cDC1 | ST13 | 2.79E-07 | 0.714994029 | 0.772 | 0.825 | 0.005495222 | Lep-L |
| cDC1 | HSPA8 | 2.07E-08 | 0.707845263 | 0.953 | 0.961 | 0.000407873 | Lep-L |
| cDC1 | SQSTM1 | 8.62E-07 | 0.701478959 | 0.918 | 0.902 | 0.016963135 | Lep-L |
| cDC1 | DDIT4 | 2.48E-06 | 0.697737552 | 0.667 | 0.593 | 0.048764183 | Lep-L |
| cDC1 | CD83 | 2.12E-15 | 0.679887458 | 0.953 | 0.967 | 4.18E-11 | Lep-L |
| cDC1 | SNHG5 | 7.80E-08 | 0.661498487 | 0.795 | 0.761 | 0.001534951 | Lep-L |
| cDC1 | SFPQ | 1.84E-07 | 0.575057973 | 0.86 | 0.897 | 0.003619946 | Lep-L |
| cDC1 | MGLL | 6.98E-11 | 1.398983237 | 0.383 | 0.214 | 1.37E-06 | Lep-T |
| cDC1 | HIST2H2AC | 1.60E-08 | 1.274336432 | 0.306 | 0.178 | 0.000315508 | Lep-T |
| cDC1 | ANXA1 | 4.23E-20 | 1.032511415 | 0.761 | 0.574 | 8.33E-16 | Lep-T |
| cDC1 | RNASE6 | 1.18E-07 | 1.015474677 | 0.398 | 0.261 | 0.002330313 | Lep-T |
| cDC1 | TACSTD2 | 7.83E-18 | 0.968667005 | 0.707 | 0.479 | 1.54E-13 | Lep-T |
| cDC1 | CD74 | 3.52E-15 | 0.898273503 | 0.996 | 1 | 6.93E-11 | Lep-T |
| cDC1 | TGFBI | 9.45E-11 | 0.831903546 | 0.672 | 0.546 | 1.86E-06 | Lep-T |
| cDC1 | PAK3 | 9.05E-10 | 0.813572705 | 0.582 | 0.418 | 1.78E-05 | Lep-T |
| cDC1 | TCEA3 | 4.12E-07 | 0.784563563 | 0.435 | 0.301 | 0.008102553 | Lep-T |
| cDC1 | SNX9 | 1.27E-06 | 0.772501086 | 0.755 | 0.7 | 0.025076169 | Lep-T |
| cDC1 | PLAUR | 1.66E-08 | 0.766463941 | 0.742 | 0.699 | 0.000327254 | Lep-T |
| cDC1 | HLA-DQA2 | 8.03E-21 | 0.763408855 | 0.777 | 0.635 | 1.58E-16 | Lep-T |
| cDC1 | CDKN1A | 9.77E-12 | 0.755274988 | 0.786 | 0.666 | 1.92E-07 | Lep-T |
| cDC1 | BCL3 | 1.52E-08 | 0.709360942 | 0.595 | 0.464 | 0.000299721 | Lep-T |
| cDC1 | SYNGR2 | 4.39E-08 | 0.691232486 | 0.744 | 0.639 | 0.000863624 | Lep-T |

| | | | | | | | |
|---|---|---|---|---|---|---|---|
| cDC1 | DNASE1L3 | 1.13E-12 | 0.673050964 | 0.84 | 0.736 | 2.23E-08 | Lep-T |
| cDC1 | CD52 | 1.07E-09 | 0.671216251 | 0.687 | 0.555 | 2.12E-05 | Lep-T |
| cDC1 | CD40 | 2.66E-09 | 0.652215686 | 0.709 | 0.645 | 5.24E-05 | Lep-T |
| cDC1 | FBL | 7.98E-07 | 0.637275937 | 0.554 | 0.448 | 0.015704009 | Lep-T |
| cDC1 | PSAP | 1.30E-24 | 0.624776272 | 0.947 | 0.921 | 2.56E-20 | Lep-T |
| cDC1 | LMNA | 3.15E-18 | 0.623562313 | 0.934 | 0.907 | 6.20E-14 | Lep-T |
| cDC1 | PNRC1 | 5.58E-19 | 0.622868925 | 0.952 | 0.941 | 1.10E-14 | Lep-T |
| cDC1 | USP12 | 5.10E-09 | 0.60949833 | 0.766 | 0.681 | 0.000100412 | Lep-T |
| cDC1 | LGALS3 | 2.85E-09 | 0.600523561 | 0.779 | 0.708 | 5.61E-05 | Lep-T |
| cDC1 | ICAM1 | 4.24E-08 | 0.596876119 | 0.748 | 0.674 | 0.000834576 | Lep-T |
| cDC1 | GNB2 | 5.76E-08 | 0.593844141 | 0.779 | 0.721 | 0.001132947 | Lep-T |
| cDC1 | PLEK | 2.38E-13 | 0.577838508 | 0.939 | 0.948 | 4.69E-09 | Lep-T |
| cDC1 | MAP2K31 | 5.68E-08 | 0.571233108 | 0.713 | 0.65 | 0.001118071 | Lep-T |
| cDC1 | CST3 | 3.67E-13 | 0.561474368 | 0.991 | 0.99 | 7.22E-09 | Lep-T |
| cDC1 | CDC42EP31 | 4.04E-10 | 0.559374036 | 0.768 | 0.668 | 7.94E-06 | Lep-T |
| cDC1 | ATF4 | 6.17E-09 | 0.543343385 | 0.742 | 0.688 | 0.000121381 | Lep-T |
| cDC1 | EEF21 | 1.25E-16 | 0.534503869 | 0.886 | 0.849 | 2.46E-12 | Lep-T |
| cDC1 | DDX3Y1 | 1.42E-09 | 0.533594862 | 0.449 | 0.267 | 2.79E-05 | Lep-T |
| cDC1 | SQSTM11 | 8.79E-16 | 0.525302804 | 0.926 | 0.89 | 1.73E-11 | Lep-T |
| cDC1 | FSCN1 | 2.59E-10 | 0.519720608 | 0.722 | 0.543 | 5.10E-06 | Lep-T |
| cDC1 | ANXA5 | 1.47E-08 | 0.507975926 | 0.882 | 0.868 | 0.000290255 | Lep-T |
| cDC1 | KRT161 | 1.42E-10 | 1.970757302 | 0.254 | 0.107 | 2.80E-06 | Mari |
| cDC1 | GAS6 | 2.37E-36 | 1.793427046 | 0.683 | 0.299 | 4.66E-32 | Mari |
| cDC1 | IFITM3 | 4.87E-09 | 1.664673077 | 0.365 | 0.204 | 9.58E-05 | Mari |
| cDC1 | SELL1 | 3.08E-26 | 1.479762015 | 0.683 | 0.341 | 6.06E-22 | Mari |
| cDC1 | S100A81 | 1.60E-09 | 1.478946554 | 0.401 | 0.239 | 3.14E-05 | Mari |
| cDC1 | MGST1 | 1.01E-09 | 1.463646879 | 0.325 | 0.165 | 1.99E-05 | Mari |
| cDC1 | GCA | 2.15E-32 | 1.441269228 | 0.802 | 0.512 | 4.23E-28 | Mari |
| cDC1 | HLA-DRB51 | 7.78E-18 | 1.438706513 | 0.905 | 0.801 | 1.53E-13 | Mari |
| cDC1 | GNG10 | 1.49E-32 | 1.414342361 | 0.825 | 0.581 | 2.94E-28 | Mari |
| cDC1 | POLE4 | 2.70E-14 | 1.407085142 | 0.456 | 0.245 | 5.31E-10 | Mari |
| cDC1 | NAALADL1 | 8.41E-09 | 1.314506324 | 0.345 | 0.187 | 0.000165459 | Mari |
| cDC1 | MSRB2 | 2.61E-10 | 1.289395687 | 0.262 | 0.108 | 5.15E-06 | Mari |
| cDC1 | MTLN | 3.04E-14 | 1.269544677 | 0.405 | 0.188 | 5.98E-10 | Mari |
| cDC1 | MRPL58 | 1.32E-12 | 1.255516846 | 0.361 | 0.163 | 2.60E-08 | Mari |
| cDC1 | IFITM2 | 5.52E-11 | 1.250267488 | 0.452 | 0.254 | 1.09E-06 | Mari |
| cDC1 | NAGPA | 3.27E-07 | 1.230791463 | 0.254 | 0.127 | 0.006443238 | Mari |
| cDC1 | MRPS28 | 5.35E-11 | 1.225086754 | 0.381 | 0.199 | 1.05E-06 | Mari |
| cDC1 | CYB561D2 | 9.59E-08 | 1.213975317 | 0.262 | 0.127 | 0.001887154 | Mari |
| cDC1 | CD691 | 5.53E-09 | 1.198462799 | 0.452 | 0.278 | 0.000108838 | Mari |
| cDC1 | SERPINA1 | 8.58E-13 | 1.171235066 | 0.516 | 0.298 | 1.69E-08 | Mari |
| cDC1 | TATDN3 | 6.55E-08 | 1.154008921 | 0.274 | 0.135 | 0.001289502 | Mari |
| cDC1 | CUTC | 1.45E-10 | 1.146868401 | 0.357 | 0.174 | 2.85E-06 | Mari |
| cDC1 | TNFSF10 | 8.12E-11 | 1.14020501 | 0.476 | 0.284 | 1.60E-06 | Mari |
| cDC1 | MRPS18C | 2.20E-16 | 1.116239222 | 0.536 | 0.281 | 4.33E-12 | Mari |
| cDC1 | CEACAM3 | 1.12E-09 | 1.11354758 | 0.258 | 0.109 | 2.20E-05 | Mari |
| cDC1 | ISCA2 | 1.35E-09 | 1.106213986 | 0.365 | 0.192 | 2.66E-05 | Mari |
| cDC1 | RAB32 | 4.14E-27 | 1.098508731 | 0.893 | 0.751 | 8.16E-23 | Mari |
| cDC1 | HSCB | 1.04E-08 | 1.095834283 | 0.369 | 0.2 | 0.000205042 | Mari |
| cDC1 | METTL23 | 5.44E-10 | 1.089318654 | 0.437 | 0.249 | 1.07E-05 | Mari |
| cDC1 | ANKRD49 | 2.26E-07 | 1.079936436 | 0.25 | 0.121 | 0.004443661 | Mari |
| cDC1 | CIAO2A | 9.45E-16 | 1.07876887 | 0.659 | 0.418 | 1.86E-11 | Mari |
| cDC1 | RNASEH2C1 | 7.81E-12 | 1.078097054 | 0.575 | 0.363 | 1.54E-07 | Mari |
| cDC1 | DIMT11 | 1.00E-12 | 1.066287048 | 0.595 | 0.367 | 1.97E-08 | Mari |
| cDC1 | UNC50 | 1.42E-08 | 1.058650472 | 0.337 | 0.179 | 0.000279051 | Mari |
| cDC1 | PPP1R11 | 9.72E-10 | 1.056830929 | 0.405 | 0.222 | 1.91E-05 | Mari |

| | | | | | | | |
|---|---|---|---|---|---|---|---|
| cDC1 | JKAMP | 5.95E-10 | 1.055654373 | 0.405 | 0.225 | 1.17E-05 | Mari |
| cDC1 | HPRT1 | 1.77E-09 | 1.053461787 | 0.5 | 0.33 | 3.49E-05 | Mari |
| cDC1 | TLR101 | 1.18E-06 | 1.049361414 | 0.393 | 0.256 | 0.023273496 | Mari |
| cDC1 | COMMD8 | 1.43E-14 | 1.041184636 | 0.552 | 0.316 | 2.81E-10 | Mari |
| cDC1 | LYZ | 7.58E-30 | 1.034358845 | 0.996 | 0.934 | 1.49E-25 | Mari |
| cDC1 | EIF4EBP1 | 7.37E-19 | 1.02609023 | 0.742 | 0.509 | 1.45E-14 | Mari |
| cDC1 | RPA2 | 1.12E-08 | 1.02487158 | 0.44 | 0.261 | 0.00021984 | Mari |
| cDC1 | SRI | 1.06E-24 | 1.023705391 | 0.849 | 0.606 | 2.08E-20 | Mari |
| cDC1 | PTPMT1 | 1.46E-09 | 1.021048571 | 0.504 | 0.317 | 2.88E-05 | Mari |
| cDC1 | DBNDD2 | 7.49E-14 | 1.017811401 | 0.659 | 0.428 | 1.47E-09 | Mari |
| cDC1 | MRPL15 | 9.28E-12 | 1.009413325 | 0.448 | 0.235 | 1.83E-07 | Mari |
| cDC1 | ARL2 | 2.30E-12 | 1.000918991 | 0.5 | 0.295 | 4.52E-08 | Mari |
| cDC1 | ABT1 | 9.10E-07 | 1.000388952 | 0.25 | 0.125 | 0.017922169 | Mari |
| cDC1 | MS4A6A | 2.17E-07 | 0.996672649 | 0.409 | 0.253 | 0.004262623 | Mari |
| cDC1 | MRPL13 | 2.53E-12 | 0.992695401 | 0.532 | 0.324 | 4.97E-08 | Mari |
| cDC1 | SOCS1 | 1.56E-13 | 0.983011952 | 0.659 | 0.427 | 3.07E-09 | Mari |
| cDC1 | ORMDL2 | 1.76E-08 | 0.982754622 | 0.437 | 0.271 | 0.000345754 | Mari |
| cDC1 | VPS25 | 1.77E-10 | 0.980643941 | 0.369 | 0.183 | 3.49E-06 | Mari |
| cDC1 | TRIAP1 | 4.09E-07 | 0.979296829 | 0.325 | 0.183 | 0.008050629 | Mari |
| cDC1 | MRPS23 | 1.13E-11 | 0.978668587 | 0.5 | 0.29 | 2.22E-07 | Mari |
| cDC1 | FUCA1 | 2.82E-09 | 0.97643071 | 0.5 | 0.313 | 5.55E-05 | Mari |
| cDC1 | S100A91 | 3.90E-07 | 0.971619556 | 0.484 | 0.33 | 0.007675935 | Mari |
| cDC1 | SS18L2 | 2.14E-17 | 0.969960914 | 0.635 | 0.347 | 4.21E-13 | Mari |
| cDC1 | SERPINB1 | 9.96E-30 | 0.96717963 | 0.968 | 0.849 | 1.96E-25 | Mari |
| cDC1 | FCER1G | 1.61E-09 | 0.965951948 | 0.627 | 0.451 | 3.17E-05 | Mari |
| cDC1 | ROGDI | 2.03E-10 | 0.96452017 | 0.508 | 0.314 | 3.99E-06 | Mari |
| cDC1 | FAM89B | 1.88E-18 | 0.955846746 | 0.746 | 0.473 | 3.70E-14 | Mari |
| cDC1 | PLGRKT | 7.35E-10 | 0.954482912 | 0.401 | 0.221 | 1.45E-05 | Mari |
| cDC1 | NSMCE1 | 3.29E-10 | 0.953884687 | 0.591 | 0.396 | 6.48E-06 | Mari |
| cDC1 | CYBC1 | 1.10E-12 | 0.953310189 | 0.587 | 0.398 | 2.17E-08 | Mari |
| cDC1 | APOBEC3G | 4.12E-07 | 0.949028765 | 0.393 | 0.239 | 0.008117168 | Mari |
| cDC1 | CLPP | 2.20E-11 | 0.947617883 | 0.5 | 0.28 | 4.34E-07 | Mari |
| cDC1 | SRP9 | 1.65E-17 | 0.942956831 | 0.774 | 0.545 | 3.24E-13 | Mari |
| cDC1 | CKLF | 5.64E-25 | 0.942394769 | 0.913 | 0.754 | 1.11E-20 | Mari |
| cDC1 | F8A1 | 4.79E-07 | 0.939376574 | 0.294 | 0.159 | 0.00943073 | Mari |
| cDC1 | SNAPIN | 2.94E-08 | 0.939031349 | 0.329 | 0.179 | 0.000578604 | Mari |
| cDC1 | MRPL171 | 1.91E-07 | 0.936388815 | 0.385 | 0.229 | 0.003764898 | Mari |
| cDC1 | TXNDC17 | 3.05E-18 | 0.926107874 | 0.75 | 0.515 | 6.01E-14 | Mari |
| cDC1 | MDP1 | 2.50E-06 | 0.924013523 | 0.349 | 0.212 | 0.049114459 | Mari |
| cDC1 | PPCS | 3.78E-09 | 0.923671292 | 0.512 | 0.319 | 7.45E-05 | Mari |
| cDC1 | DPY30 | 2.20E-08 | 0.918372731 | 0.405 | 0.241 | 0.000432799 | Mari |
| cDC1 | TMEM167A | 2.94E-15 | 0.918307906 | 0.71 | 0.477 | 5.80E-11 | Mari |
| cDC1 | LAMTOR2 | 8.14E-18 | 0.915632939 | 0.774 | 0.568 | 1.60E-13 | Mari |
| cDC1 | YIF1A | 4.19E-09 | 0.913934695 | 0.484 | 0.312 | 8.25E-05 | Mari |
| cDC1 | ACTR61 | 3.81E-07 | 0.912628832 | 0.325 | 0.184 | 0.007501602 | Mari |
| cDC1 | SLIRP | 3.88E-18 | 0.912000423 | 0.722 | 0.436 | 7.63E-14 | Mari |
| cDC1 | NDUFC2 | 9.85E-26 | 0.90937714 | 0.893 | 0.739 | 1.94E-21 | Mari |
| cDC1 | TYMP1 | 1.69E-29 | 0.908429964 | 0.964 | 0.808 | 3.33E-25 | Mari |
| cDC1 | COA4 | 4.94E-11 | 0.906849703 | 0.544 | 0.344 | 9.72E-07 | Mari |
| cDC1 | MRPL57 | 6.47E-14 | 0.906354811 | 0.647 | 0.405 | 1.27E-09 | Mari |
| cDC1 | SERPINF11 | 1.42E-17 | 0.903828923 | 0.853 | 0.678 | 2.80E-13 | Mari |
| cDC1 | GCSAM | 3.02E-10 | 0.892484899 | 0.556 | 0.365 | 5.94E-06 | Mari |
| cDC1 | CCDC28A | 5.80E-07 | 0.890542579 | 0.353 | 0.204 | 0.011413695 | Mari |
| cDC1 | TMCO1 | 9.40E-11 | 0.884855485 | 0.56 | 0.353 | 1.85E-06 | Mari |
| cDC1 | MIF4GD | 6.17E-11 | 0.884304969 | 0.643 | 0.454 | 1.21E-06 | Mari |
| cDC1 | GALK1 | 1.14E-06 | 0.880885225 | 0.421 | 0.275 | 0.022347711 | Mari |

| | | | | | | | |
|---|---|---|---|---|---|---|---|
| cDC1 | EXOSC7 | 2.16E-10 | 0.87790068 | 0.401 | 0.209 | 4.26E-06 | Mari |
| cDC1 | LY96 | 9.33E-12 | 0.877697091 | 0.631 | 0.433 | 1.84E-07 | Mari |
| cDC1 | NDUFAB1 | 5.48E-20 | 0.875088971 | 0.845 | 0.61 | 1.08E-15 | Mari |
| cDC1 | DERA | 1.24E-07 | 0.874855655 | 0.341 | 0.188 | 0.002443835 | Mari |
| cDC1 | SMIM12 | 1.86E-06 | 0.873575228 | 0.444 | 0.305 | 0.036514885 | Mari |
| cDC1 | PYURF | 3.71E-12 | 0.871903315 | 0.587 | 0.372 | 7.30E-08 | Mari |
| cDC1 | HSBP1 | 1.59E-16 | 0.871805546 | 0.758 | 0.509 | 3.13E-12 | Mari |
| cDC1 | VBP1 | 8.91E-14 | 0.871283469 | 0.611 | 0.372 | 1.75E-09 | Mari |
| cDC1 | LLPH | 1.39E-06 | 0.86852219 | 0.413 | 0.268 | 0.027312725 | Mari |
| cDC1 | POLR2I | 1.21E-09 | 0.86556018 | 0.56 | 0.361 | 2.38E-05 | Mari |
| cDC1 | ACP1 | 8.97E-12 | 0.864428189 | 0.631 | 0.418 | 1.77E-07 | Mari |
| cDC1 | PEX16 | 2.03E-08 | 0.861972226 | 0.401 | 0.226 | 0.000400027 | Mari |
| cDC1 | DTWD1 | 3.62E-07 | 0.85844226 | 0.262 | 0.131 | 0.007124559 | Mari |
| cDC1 | PAFAH1B3 | 3.51E-08 | 0.858128677 | 0.512 | 0.33 | 0.00068993 | Mari |
| cDC1 | CLEC12A | 3.36E-07 | 0.857902162 | 0.365 | 0.217 | 0.006606162 | Mari |
| cDC1 | CHCHD1 | 6.21E-10 | 0.857002411 | 0.484 | 0.297 | 1.22E-05 | Mari |
| cDC1 | MED8 | 2.75E-08 | 0.855852085 | 0.448 | 0.276 | 0.000541515 | Mari |
| cDC1 | PRDX2 | 7.99E-13 | 0.855366056 | 0.714 | 0.523 | 1.57E-08 | Mari |
| cDC1 | SMCO4 | 7.57E-18 | 0.851522787 | 0.821 | 0.63 | 1.49E-13 | Mari |
| cDC1 | CALHM6 | 8.79E-12 | 0.851286512 | 0.833 | 0.636 | 1.73E-07 | Mari |
| cDC1 | UQCC2 | 2.09E-12 | 0.850291247 | 0.528 | 0.297 | 4.12E-08 | Mari |
| cDC1 | MALSU1 | 5.03E-10 | 0.848873087 | 0.381 | 0.196 | 9.89E-06 | Mari |
| cDC1 | LAMTOR1 | 2.38E-19 | 0.842901985 | 0.869 | 0.677 | 4.68E-15 | Mari |
| cDC1 | MRPL55 | 6.32E-10 | 0.842854817 | 0.563 | 0.366 | 1.24E-05 | Mari |
| cDC1 | PMM1 | 1.68E-07 | 0.842246343 | 0.409 | 0.249 | 0.003298213 | Mari |
| cDC1 | MSRB1 | 1.51E-08 | 0.839267108 | 0.56 | 0.398 | 0.00029659 | Mari |
| cDC1 | EBPL | 5.90E-10 | 0.836437717 | 0.583 | 0.404 | 1.16E-05 | Mari |
| cDC1 | RDH14 | 4.09E-07 | 0.834634197 | 0.258 | 0.125 | 0.008040937 | Mari |
| cDC1 | NDUFB2 | 1.03E-24 | 0.833265171 | 0.905 | 0.738 | 2.02E-20 | Mari |
| cDC1 | NCF4 | 4.27E-07 | 0.832769944 | 0.401 | 0.245 | 0.008414564 | Mari |
| cDC1 | NDUFA9 | 9.28E-10 | 0.832041867 | 0.571 | 0.391 | 1.83E-05 | Mari |
| cDC1 | POLR3K | 2.35E-08 | 0.831087178 | 0.385 | 0.215 | 0.000461941 | Mari |
| cDC1 | GLRX2 | 9.90E-08 | 0.830802479 | 0.353 | 0.198 | 0.001947837 | Mari |
| cDC1 | LSM6 | 1.71E-17 | 0.830279348 | 0.825 | 0.641 | 3.36E-13 | Mari |
| cDC1 | RPA3 | 2.59E-09 | 0.828113551 | 0.556 | 0.359 | 5.09E-05 | Mari |
| cDC1 | CIB1 | 9.68E-16 | 0.823823866 | 0.806 | 0.636 | 1.90E-11 | Mari |
| cDC1 | TMX1 | 7.18E-13 | 0.823489357 | 0.631 | 0.414 | 1.41E-08 | Mari |
| cDC1 | SNRNP40 | 1.30E-08 | 0.822976852 | 0.472 | 0.294 | 0.000255072 | Mari |
| cDC1 | P2RY61 | 1.65E-12 | 0.820432232 | 0.611 | 0.362 | 3.24E-08 | Mari |
| cDC1 | NUDT22 | 2.87E-11 | 0.817719509 | 0.536 | 0.314 | 5.65E-07 | Mari |
| cDC1 | HIGD1A | 2.94E-08 | 0.817575185 | 0.48 | 0.301 | 0.000577996 | Mari |
| cDC1 | ZNF593 | 5.86E-10 | 0.81689451 | 0.496 | 0.3 | 1.15E-05 | Mari |
| cDC1 | FUOM | 5.48E-11 | 0.81626046 | 0.615 | 0.395 | 1.08E-06 | Mari |
| cDC1 | NMI | 8.06E-12 | 0.815626918 | 0.647 | 0.44 | 1.59E-07 | Mari |
| cDC1 | MRPL20 | 3.42E-14 | 0.813996699 | 0.77 | 0.564 | 6.74E-10 | Mari |
| cDC1 | TOMM5 | 9.00E-13 | 0.81370913 | 0.698 | 0.494 | 1.77E-08 | Mari |
| cDC1 | LSM10 | 1.08E-10 | 0.811800193 | 0.579 | 0.379 | 2.13E-06 | Mari |
| cDC1 | TMBIM4 | 6.59E-17 | 0.808811997 | 0.758 | 0.522 | 1.30E-12 | Mari |
| cDC1 | SLC39A10 | 2.43E-07 | 0.801125182 | 0.405 | 0.25 | 0.004786062 | Mari |
| cDC1 | CPQ | 2.02E-10 | 0.795972754 | 0.655 | 0.437 | 3.98E-06 | Mari |
| cDC1 | FEZ2 | 1.75E-06 | 0.795151126 | 0.385 | 0.243 | 0.034429312 | Mari |
| cDC1 | SNRNP25 | 6.94E-11 | 0.794275389 | 0.687 | 0.495 | 1.37E-06 | Mari |
| cDC1 | MRPL40 | 4.32E-10 | 0.792715202 | 0.544 | 0.336 | 8.50E-06 | Mari |
| cDC1 | MRPL12 | 1.64E-06 | 0.792367268 | 0.472 | 0.32 | 0.032345285 | Mari |
| cDC1 | JOSD2 | 7.12E-09 | 0.79193929 | 0.46 | 0.274 | 0.000140244 | Mari |
| cDC1 | ABI31 | 2.58E-07 | 0.790760966 | 0.512 | 0.354 | 0.005070755 | Mari |

| | | | | | | | |
|---|---|---|---|---|---|---|---|
| cDC1 | PLSCR1 | 5.41E-11 | 0.790465483 | 0.714 | 0.489 | 1.06E-06 | Mari |
| cDC1 | PRPF31 | 9.92E-07 | 0.790128273 | 0.437 | 0.28 | 0.019517845 | Mari |
| cDC1 | BLOC1S1 | 3.04E-18 | 0.786778745 | 0.885 | 0.737 | 5.98E-14 | Mari |
| cDC1 | EXOSC9 | 3.62E-07 | 0.783598939 | 0.31 | 0.165 | 0.007133641 | Mari |
| cDC1 | MNDA | 5.58E-15 | 0.781627145 | 0.81 | 0.611 | 1.10E-10 | Mari |
| cDC1 | PPA1 | 3.31E-25 | 0.780078001 | 0.944 | 0.865 | 6.52E-21 | Mari |
| cDC1 | S100A11 | 4.41E-29 | 0.779705933 | 0.956 | 0.884 | 8.68E-25 | Mari |
| cDC1 | RAD51C | 2.73E-08 | 0.7796464 | 0.381 | 0.213 | 0.000536693 | Mari |
| cDC1 | POLD2 | 2.85E-07 | 0.77935313 | 0.373 | 0.214 | 0.005602899 | Mari |
| cDC1 | MYL12A | 2.49E-29 | 0.778835833 | 0.952 | 0.875 | 4.91E-25 | Mari |
| cDC1 | NSMCE4A | 7.47E-10 | 0.778511755 | 0.496 | 0.295 | 1.47E-05 | Mari |
| cDC1 | MPV17 | 1.81E-08 | 0.775601554 | 0.456 | 0.27 | 0.000357196 | Mari |
| cDC1 | TRMT1 | 6.83E-08 | 0.773598733 | 0.377 | 0.218 | 0.00134511 | Mari |
| cDC1 | MGST3 | 9.10E-14 | 0.772268829 | 0.79 | 0.577 | 1.79E-09 | Mari |
| cDC1 | BLOC1S2 | 3.75E-11 | 0.77179816 | 0.702 | 0.509 | 7.38E-07 | Mari |
| cDC1 | YDJC | 1.15E-07 | 0.77160891 | 0.44 | 0.273 | 0.002266662 | Mari |
| cDC1 | DUSP11 | 6.74E-07 | 0.771561546 | 0.294 | 0.156 | 0.01326766 | Mari |
| cDC1 | TMEM208 | 8.69E-12 | 0.771184229 | 0.679 | 0.451 | 1.71E-07 | Mari |
| cDC1 | FLOT1 | 4.52E-11 | 0.77076612 | 0.635 | 0.424 | 8.90E-07 | Mari |
| cDC1 | POP4 | 7.11E-07 | 0.770682091 | 0.365 | 0.216 | 0.014002023 | Mari |
| cDC1 | CNPY2 | 6.42E-10 | 0.767848111 | 0.56 | 0.378 | 1.26E-05 | Mari |
| cDC1 | CENPX | 5.20E-09 | 0.763684096 | 0.595 | 0.422 | 0.00010241 | Mari |
| cDC1 | TAF12 | 9.61E-09 | 0.76192433 | 0.464 | 0.279 | 0.000189082 | Mari |
| cDC1 | PSMA2 | 4.61E-18 | 0.760935441 | 0.885 | 0.737 | 9.08E-14 | Mari |
| cDC1 | FBXO6 | 4.73E-09 | 0.757533992 | 0.452 | 0.262 | 9.31E-05 | Mari |
| cDC1 | THYN1 | 4.88E-11 | 0.755604393 | 0.56 | 0.347 | 9.60E-07 | Mari |
| cDC1 | LSM8 | 8.11E-18 | 0.755432559 | 0.825 | 0.563 | 1.60E-13 | Mari |
| cDC1 | SRSF9 | 1.22E-18 | 0.754708582 | 0.917 | 0.757 | 2.40E-14 | Mari |
| cDC1 | SNRPG | 2.32E-18 | 0.753664809 | 0.877 | 0.722 | 4.57E-14 | Mari |
| cDC1 | GADD45GIP1 | 8.13E-11 | 0.751543835 | 0.687 | 0.481 | 1.60E-06 | Mari |
| cDC1 | CCNDBP1 | 2.80E-08 | 0.751390124 | 0.52 | 0.337 | 0.000550686 | Mari |
| cDC1 | TBCB | 1.07E-13 | 0.750150071 | 0.79 | 0.598 | 2.11E-09 | Mari |
| cDC1 | TSTD1 | 7.04E-10 | 0.749558282 | 0.548 | 0.344 | 1.39E-05 | Mari |
| cDC1 | GET3 | 3.23E-07 | 0.749555542 | 0.575 | 0.415 | 0.006357037 | Mari |
| cDC1 | BEX4 | 8.64E-07 | 0.74937525 | 0.341 | 0.196 | 0.017009238 | Mari |
| cDC1 | TMEM230 | 2.94E-09 | 0.74861813 | 0.69 | 0.53 | 5.79E-05 | Mari |
| cDC1 | TRAPPC1 | 1.09E-15 | 0.74834728 | 0.81 | 0.622 | 2.14E-11 | Mari |
| cDC1 | TRMT112 | 1.05E-22 | 0.746210605 | 0.901 | 0.821 | 2.07E-18 | Mari |
| cDC1 | SDHD | 2.33E-10 | 0.746001053 | 0.667 | 0.497 | 4.58E-06 | Mari |
| cDC1 | TSPAN13 | 2.17E-08 | 0.744330667 | 0.504 | 0.33 | 0.000428048 | Mari |
| cDC1 | GTF3C6 | 6.51E-10 | 0.742734499 | 0.698 | 0.501 | 1.28E-05 | Mari |
| cDC1 | LSM4 | 9.20E-13 | 0.742658916 | 0.75 | 0.555 | 1.81E-08 | Mari |
| cDC1 | PLAAT4 | 8.56E-08 | 0.742451318 | 0.476 | 0.299 | 0.001685345 | Mari |
| cDC1 | VAMP51 | 5.91E-11 | 0.742435003 | 0.671 | 0.474 | 1.16E-06 | Mari |
| cDC1 | NAA20 | 1.05E-09 | 0.74240651 | 0.571 | 0.37 | 2.07E-05 | Mari |
| cDC1 | HSPB11 | 1.45E-12 | 0.741748651 | 0.758 | 0.596 | 2.86E-08 | Mari |
| cDC1 | SPCS3 | 2.84E-15 | 0.741565995 | 0.837 | 0.63 | 5.59E-11 | Mari |
| cDC1 | ATRAID | 9.97E-10 | 0.740504663 | 0.579 | 0.397 | 1.96E-05 | Mari |
| cDC1 | BLOC1S4 | 1.22E-06 | 0.737228546 | 0.361 | 0.213 | 0.023959128 | Mari |
| cDC1 | COX14 | 4.09E-13 | 0.736179359 | 0.726 | 0.512 | 8.06E-09 | Mari |
| cDC1 | COPZ1 | 3.12E-10 | 0.734564593 | 0.619 | 0.408 | 6.15E-06 | Mari |
| cDC1 | LDHA | 6.00E-21 | 0.733750322 | 0.944 | 0.852 | 1.18E-16 | Mari |
| cDC1 | PSMB2 | 1.59E-14 | 0.733521412 | 0.79 | 0.578 | 3.12E-10 | Mari |
| cDC1 | UBE2V2 | 1.34E-11 | 0.732341619 | 0.643 | 0.404 | 2.65E-07 | Mari |
| cDC1 | TMEM59 | 1.62E-14 | 0.732016865 | 0.813 | 0.636 | 3.20E-10 | Mari |
| cDC1 | PSENEN | 3.09E-10 | 0.731129346 | 0.706 | 0.506 | 6.08E-06 | Mari |

| | | | | | | | |
|---|---|---|---|---|---|---|---|
| cDC1 | NUTF2 | 6.33E-09 | 0.727085512 | 0.639 | 0.457 | 0.000124624 | Mari |
| cDC1 | ANTKMT | 5.25E-09 | 0.726702081 | 0.524 | 0.332 | 0.000103256 | Mari |
| cDC1 | RANBP1 | 3.17E-10 | 0.725215473 | 0.738 | 0.565 | 6.24E-06 | Mari |
| cDC1 | U2AF1L4 | 7.85E-07 | 0.724779474 | 0.361 | 0.213 | 0.015449468 | Mari |
| cDC1 | BANF1 | 1.62E-15 | 0.724117986 | 0.794 | 0.534 | 3.18E-11 | Mari |
| cDC1 | ANAPC13 | 5.32E-08 | 0.724041424 | 0.361 | 0.203 | 0.001047142 | Mari |
| cDC1 | ECI2 | 4.43E-08 | 0.722379152 | 0.405 | 0.237 | 0.000872766 | Mari |
| cDC1 | NME1 | 4.76E-13 | 0.721989824 | 0.75 | 0.531 | 9.36E-09 | Mari |
| cDC1 | PRELID1 | 8.51E-15 | 0.721320789 | 0.885 | 0.719 | 1.67E-10 | Mari |
| cDC1 | SNX17 | 4.99E-11 | 0.720721494 | 0.683 | 0.477 | 9.81E-07 | Mari |
| cDC1 | GTF2A2 | 7.72E-10 | 0.717645796 | 0.687 | 0.529 | 1.52E-05 | Mari |
| cDC1 | SDF2L1 | 4.57E-07 | 0.71683612 | 0.655 | 0.511 | 0.008997422 | Mari |
| cDC1 | EIF5A | 1.20E-14 | 0.715741797 | 0.905 | 0.76 | 2.36E-10 | Mari |
| cDC1 | TIMM8B | 2.78E-11 | 0.713354778 | 0.73 | 0.551 | 5.47E-07 | Mari |
| cDC1 | SVBP | 1.55E-06 | 0.711419973 | 0.421 | 0.268 | 0.030512288 | Mari |
| cDC1 | UQCC3 | 7.22E-08 | 0.710675132 | 0.389 | 0.229 | 0.001421738 | Mari |
| cDC1 | NDUFC1 | 2.71E-12 | 0.708963797 | 0.702 | 0.481 | 5.33E-08 | Mari |
| cDC1 | CHURC1 | 8.37E-14 | 0.708700383 | 0.802 | 0.593 | 1.65E-09 | Mari |
| cDC1 | ATP6V0E1 | 7.63E-22 | 0.706543877 | 0.937 | 0.819 | 1.50E-17 | Mari |
| cDC1 | DBNL | 6.64E-09 | 0.706336188 | 0.73 | 0.575 | 0.000130689 | Mari |
| cDC1 | MCM5 | 3.69E-07 | 0.706041221 | 0.579 | 0.404 | 0.007254234 | Mari |
| cDC1 | DCAF13 | 1.61E-07 | 0.705805182 | 0.532 | 0.364 | 0.003173228 | Mari |
| cDC1 | DUT | 2.82E-07 | 0.705279362 | 0.603 | 0.444 | 0.005558781 | Mari |
| cDC1 | PSMG4 | 5.49E-07 | 0.703287876 | 0.373 | 0.22 | 0.01080773 | Mari |
| cDC1 | MRPS12 | 1.07E-06 | 0.702954142 | 0.536 | 0.372 | 0.021019128 | Mari |
| cDC1 | PIN1 | 5.86E-08 | 0.702352478 | 0.556 | 0.386 | 0.00115395 | Mari |
| cDC1 | ARPC1B | 5.00E-29 | 0.701737157 | 0.972 | 0.874 | 9.84E-25 | Mari |
| cDC1 | FKBP3 | 5.60E-07 | 0.701518843 | 0.464 | 0.313 | 0.01102107 | Mari |
| cDC1 | RPP30 | 1.45E-07 | 0.701227007 | 0.417 | 0.251 | 0.002851614 | Mari |
| cDC1 | SSU72 | 3.80E-09 | 0.700895517 | 0.663 | 0.497 | 7.47E-05 | Mari |
| cDC1 | CRELD2 | 1.90E-10 | 0.700883584 | 0.587 | 0.359 | 3.74E-06 | Mari |
| cDC1 | MDH1 | 4.34E-12 | 0.699801819 | 0.702 | 0.481 | 8.53E-08 | Mari |
| cDC1 | BATF3 | 4.95E-11 | 0.698866361 | 0.825 | 0.661 | 9.75E-07 | Mari |
| cDC1 | UQCR10 | 6.98E-19 | 0.697884732 | 0.889 | 0.728 | 1.37E-14 | Mari |
| cDC1 | BST2 | 1.03E-17 | 0.697409234 | 0.905 | 0.73 | 2.04E-13 | Mari |
| cDC1 | NDUFB5 | 2.02E-11 | 0.696730642 | 0.813 | 0.661 | 3.97E-07 | Mari |
| cDC1 | HMGN1 | 7.23E-18 | 0.693555151 | 0.94 | 0.829 | 1.42E-13 | Mari |
| cDC1 | APTX | 1.19E-06 | 0.693157938 | 0.425 | 0.258 | 0.023342305 | Mari |
| cDC1 | PSMA5 | 8.49E-15 | 0.691027591 | 0.817 | 0.648 | 1.67E-10 | Mari |
| cDC1 | GLRX5 | 1.96E-08 | 0.690780877 | 0.571 | 0.387 | 0.00038674 | Mari |
| cDC1 | RBCK1 | 4.82E-09 | 0.690268559 | 0.603 | 0.406 | 9.49E-05 | Mari |
| cDC1 | TIMM17B | 2.13E-09 | 0.690122102 | 0.667 | 0.49 | 4.19E-05 | Mari |
| cDC1 | ISOC2 | 4.84E-07 | 0.684549959 | 0.552 | 0.397 | 0.0095307 | Mari |
| cDC1 | SELENOS | 2.20E-06 | 0.683347988 | 0.607 | 0.458 | 0.043259461 | Mari |
| cDC1 | MRPL19 | 7.94E-07 | 0.683256058 | 0.393 | 0.246 | 0.015636614 | Mari |
| cDC1 | NDUFB3 | 1.72E-09 | 0.682793568 | 0.687 | 0.549 | 3.38E-05 | Mari |
| cDC1 | MDH2 | 5.85E-14 | 0.681772814 | 0.829 | 0.629 | 1.15E-09 | Mari |
| cDC1 | TPRKB | 1.06E-07 | 0.679683969 | 0.552 | 0.382 | 0.00209198 | Mari |
| cDC1 | CHMP4A | 5.42E-09 | 0.679325758 | 0.623 | 0.437 | 0.000106649 | Mari |
| cDC1 | ESD | 2.31E-09 | 0.67895204 | 0.631 | 0.445 | 4.56E-05 | Mari |
| cDC1 | DCTPP1 | 3.83E-09 | 0.677788187 | 0.647 | 0.478 | 7.54E-05 | Mari |
| cDC1 | IMP4 | 4.60E-09 | 0.672222029 | 0.5 | 0.311 | 9.06E-05 | Mari |
| cDC1 | SRP19 | 9.90E-16 | 0.671805898 | 0.861 | 0.696 | 1.95E-11 | Mari |
| cDC1 | FOS | 1.73E-13 | 0.670659049 | 0.937 | 0.863 | 3.41E-09 | Mari |
| cDC1 | GSTO1 | 4.21E-14 | 0.670257877 | 0.813 | 0.67 | 8.29E-10 | Mari |
| cDC1 | MRPS34 | 4.95E-09 | 0.670032221 | 0.655 | 0.472 | 9.74E-05 | Mari |

| | | | | | | | |
|---|---|---|---|---|---|---|---|
| cDC1 | ATP6V0B | 4.61E-20 | 0.6692442 | 0.917 | 0.775 | 9.07E-16 | Mari |
| cDC1 | AIMP1 | 8.74E-08 | 0.66870397 | 0.556 | 0.399 | 0.001719664 | Mari |
| cDC1 | CDC123 | 9.99E-07 | 0.668159304 | 0.488 | 0.339 | 0.019668846 | Mari |
| cDC1 | SNF8 | 3.50E-11 | 0.665736325 | 0.738 | 0.519 | 6.89E-07 | Mari |
| cDC1 | RBX1 | 2.97E-18 | 0.66511398 | 0.897 | 0.742 | 5.84E-14 | Mari |
| cDC1 | MICOS10 | 1.12E-15 | 0.663964032 | 0.893 | 0.749 | 2.20E-11 | Mari |
| cDC1 | CLTB | 1.16E-14 | 0.663610995 | 0.845 | 0.659 | 2.28E-10 | Mari |
| cDC1 | FKBP1A | 7.79E-14 | 0.663103915 | 0.889 | 0.695 | 1.53E-09 | Mari |
| cDC1 | SPINT2 | 7.51E-12 | 0.66240273 | 0.857 | 0.644 | 1.48E-07 | Mari |
| cDC1 | CFL1 | 4.29E-28 | 0.661042879 | 0.964 | 0.929 | 8.44E-24 | Mari |
| cDC1 | SELENOT | 1.18E-08 | 0.660855928 | 0.683 | 0.494 | 0.000233225 | Mari |
| cDC1 | NDUFA8 | 2.05E-07 | 0.660058578 | 0.468 | 0.295 | 0.004026032 | Mari |
| cDC1 | MIEN1 | 1.02E-06 | 0.659975754 | 0.409 | 0.25 | 0.02011841 | Mari |
| cDC1 | SEC61G | 1.43E-10 | 0.65945488 | 0.865 | 0.703 | 2.82E-06 | Mari |
| cDC1 | BTLA | 3.07E-08 | 0.659443665 | 0.516 | 0.342 | 0.000603509 | Mari |
| cDC1 | DPM2 | 3.68E-07 | 0.659187979 | 0.381 | 0.226 | 0.007242002 | Mari |
| cDC1 | GSTK1 | 1.72E-16 | 0.658912657 | 0.893 | 0.743 | 3.38E-12 | Mari |
| cDC1 | TMED9 | 1.00E-08 | 0.657880677 | 0.655 | 0.476 | 0.000197314 | Mari |
| cDC1 | TXNDC9 | 8.99E-07 | 0.657612254 | 0.468 | 0.308 | 0.017688846 | Mari |
| cDC1 | IGFBP7 | 3.16E-08 | 0.657176869 | 0.667 | 0.509 | 0.000621602 | Mari |
| cDC1 | FDPS | 2.47E-07 | 0.657099495 | 0.563 | 0.392 | 0.004866916 | Mari |
| cDC1 | HLA-DQB1 | 1.19E-27 | 0.656788868 | 0.996 | 0.974 | 2.34E-23 | Mari |
| cDC1 | COMMD3 | 1.02E-06 | 0.656657256 | 0.528 | 0.374 | 0.020134522 | Mari |
| cDC1 | TOMM6 | 2.31E-17 | 0.655170368 | 0.897 | 0.729 | 4.55E-13 | Mari |
| cDC1 | UQCR11 | 1.22E-18 | 0.655154998 | 0.913 | 0.777 | 2.40E-14 | Mari |
| cDC1 | SUMO1 | 7.61E-12 | 0.654148201 | 0.77 | 0.594 | 1.50E-07 | Mari |
| cDC1 | BIRC2 | 8.18E-08 | 0.650562009 | 0.575 | 0.372 | 0.001610969 | Mari |
| cDC1 | DGUOK | 3.24E-09 | 0.650290181 | 0.687 | 0.495 | 6.38E-05 | Mari |
| cDC1 | GABARAPL2 | 5.51E-15 | 0.649016332 | 0.889 | 0.737 | 1.08E-10 | Mari |
| cDC1 | SRM | 2.58E-09 | 0.647901395 | 0.635 | 0.428 | 5.07E-05 | Mari |
| cDC1 | STMP1 | 3.09E-14 | 0.645489054 | 0.774 | 0.554 | 6.09E-10 | Mari |
| cDC1 | UCHL3 | 1.29E-10 | 0.645077876 | 0.647 | 0.414 | 2.53E-06 | Mari |
| cDC1 | SNX3 | 1.06E-24 | 0.644481396 | 0.968 | 0.925 | 2.10E-20 | Mari |
| cDC1 | ECHS1 | 1.49E-07 | 0.644432528 | 0.647 | 0.484 | 0.002927935 | Mari |
| cDC1 | NEDD8 | 5.48E-16 | 0.643690192 | 0.893 | 0.719 | 1.08E-11 | Mari |
| cDC1 | NDUFB7 | 2.41E-10 | 0.6425822 | 0.762 | 0.563 | 4.75E-06 | Mari |
| cDC1 | ERH | 2.34E-13 | 0.642429031 | 0.821 | 0.681 | 4.60E-09 | Mari |
| cDC1 | ATP5MF | 1.30E-19 | 0.641894859 | 0.929 | 0.8 | 2.57E-15 | Mari |
| cDC1 | LSM2 | 5.61E-09 | 0.641836404 | 0.655 | 0.481 | 0.00011049 | Mari |
| cDC1 | IAH1 | 3.46E-08 | 0.641810553 | 0.667 | 0.474 | 0.000681131 | Mari |
| cDC1 | ADI1 | 5.84E-07 | 0.640093141 | 0.544 | 0.379 | 0.011504109 | Mari |
| cDC1 | MRPL52 | 9.57E-11 | 0.639471394 | 0.718 | 0.536 | 1.88E-06 | Mari |
| cDC1 | MYDGF | 1.53E-08 | 0.637410212 | 0.75 | 0.564 | 0.000301902 | Mari |
| cDC1 | PSMD7 | 9.67E-10 | 0.636529059 | 0.71 | 0.518 | 1.90E-05 | Mari |
| cDC1 | PFN1 | 5.67E-28 | 0.635570392 | 0.98 | 0.912 | 1.12E-23 | Mari |
| cDC1 | PSMB9 | 5.96E-18 | 0.634506764 | 0.929 | 0.852 | 1.17E-13 | Mari |
| cDC1 | STOML2 | 1.58E-06 | 0.634070564 | 0.536 | 0.371 | 0.031103355 | Mari |
| cDC1 | SF3B6 | 2.32E-12 | 0.633403749 | 0.833 | 0.676 | 4.58E-08 | Mari |
| cDC1 | PSMB6 | 2.20E-15 | 0.63314918 | 0.889 | 0.721 | 4.33E-11 | Mari |
| cDC1 | TRAPPC2L | 1.04E-08 | 0.632428648 | 0.591 | 0.386 | 0.000204192 | Mari |
| cDC1 | UBE2L3 | 1.88E-10 | 0.631412185 | 0.802 | 0.648 | 3.71E-06 | Mari |
| cDC1 | ZNHIT3 | 2.33E-06 | 0.630536628 | 0.417 | 0.265 | 0.045837662 | Mari |
| cDC1 | 11-Mar | 1.65E-09 | 0.629378555 | 0.75 | 0.531 | 3.26E-05 | Mari |
| cDC1 | ETFA | 3.43E-11 | 0.628465999 | 0.75 | 0.529 | 6.75E-07 | Mari |
| cDC1 | SERPINF2 | 1.30E-07 | 0.627659722 | 0.54 | 0.358 | 0.002553042 | Mari |
| cDC1 | PHPT1 | 6.21E-10 | 0.627448075 | 0.75 | 0.58 | 1.22E-05 | Mari |

| | | | | | | | |
|---|---|---|---|---|---|---|---|
| cDC1 | UBL5 | 1.12E-20 | 0.627140577 | 0.933 | 0.821 | 2.21E-16 | Mari |
| cDC1 | UBE2M | 2.13E-06 | 0.627054123 | 0.587 | 0.44 | 0.041936491 | Mari |
| cDC1 | C1D | 1.06E-10 | 0.626684989 | 0.571 | 0.359 | 2.09E-06 | Mari |
| cDC1 | POLD4 | 3.76E-11 | 0.625332368 | 0.794 | 0.621 | 7.40E-07 | Mari |
| cDC1 | POLR2K | 3.49E-09 | 0.624416424 | 0.611 | 0.436 | 6.88E-05 | Mari |
| cDC1 | SUCLG1 | 1.10E-08 | 0.623187313 | 0.651 | 0.479 | 0.000216024 | Mari |
| cDC1 | AP2S1 | 3.66E-19 | 0.623034019 | 0.921 | 0.816 | 7.20E-15 | Mari |
| cDC1 | TMEM256 | 4.01E-10 | 0.62201086 | 0.762 | 0.575 | 7.90E-06 | Mari |
| cDC1 | SELENOH | 3.05E-12 | 0.621373063 | 0.861 | 0.692 | 6.00E-08 | Mari |
| cDC1 | NDUFA13 | 1.00E-18 | 0.620320742 | 0.948 | 0.804 | 1.98E-14 | Mari |
| cDC1 | NCF1 | 6.45E-07 | 0.620271321 | 0.429 | 0.274 | 0.012705553 | Mari |
| cDC1 | ELP5 | 6.18E-07 | 0.618274131 | 0.464 | 0.291 | 0.012164947 | Mari |
| cDC1 | MRPL3 | 1.20E-08 | 0.61645891 | 0.659 | 0.486 | 0.000236788 | Mari |
| cDC1 | PDIA6 | 1.30E-10 | 0.615598348 | 0.837 | 0.67 | 2.55E-06 | Mari |
| cDC1 | PARK7 | 2.25E-18 | 0.615007478 | 0.901 | 0.791 | 4.43E-14 | Mari |
| cDC1 | OXLD1 | 1.52E-06 | 0.614906651 | 0.31 | 0.176 | 0.029845067 | Mari |
| cDC1 | MRPL41 | 1.53E-10 | 0.613949523 | 0.694 | 0.513 | 3.02E-06 | Mari |
| cDC1 | DAD1 | 1.65E-12 | 0.613601177 | 0.817 | 0.617 | 3.24E-08 | Mari |
| cDC1 | MTCH2 | 2.35E-07 | 0.612943548 | 0.556 | 0.385 | 0.004624534 | Mari |
| cDC1 | ZSWIM7 | 4.54E-07 | 0.612150959 | 0.488 | 0.32 | 0.008935413 | Mari |
| cDC1 | AURKAIP1 | 4.55E-14 | 0.609295831 | 0.845 | 0.662 | 8.96E-10 | Mari |
| cDC1 | NDUFV3 | 2.60E-07 | 0.609160007 | 0.512 | 0.341 | 0.005122947 | Mari |
| cDC1 | CDKN2D | 1.80E-06 | 0.608976284 | 0.619 | 0.468 | 0.035421878 | Mari |
| cDC1 | IFI30 | 3.68E-10 | 0.608907955 | 0.929 | 0.801 | 7.25E-06 | Mari |
| cDC1 | RAB34 | 4.27E-07 | 0.608693041 | 0.603 | 0.429 | 0.008406885 | Mari |
| cDC1 | SCP2 | 2.65E-08 | 0.608032466 | 0.675 | 0.49 | 0.000520928 | Mari |
| cDC1 | H3F3B | 9.57E-11 | 0.606681335 | 0.996 | 0.972 | 1.88E-06 | Mari |
| cDC1 | VPS29 | 7.97E-15 | 0.60539728 | 0.909 | 0.733 | 1.57E-10 | Mari |
| cDC1 | NQO2 | 3.42E-07 | 0.604557725 | 0.567 | 0.382 | 0.006725984 | Mari |
| cDC1 | PSME2 | 2.66E-17 | 0.602284047 | 0.96 | 0.854 | 5.23E-13 | Mari |
| cDC1 | SIVA1 | 2.27E-07 | 0.601886864 | 0.643 | 0.487 | 0.004473811 | Mari |
| cDC1 | MYL6 | 6.61E-26 | 0.60184141 | 0.972 | 0.934 | 1.30E-21 | Mari |
| cDC1 | MRPL54 | 1.74E-07 | 0.601592529 | 0.619 | 0.446 | 0.003432272 | Mari |
| cDC1 | ATP5PF | 7.67E-16 | 0.601313894 | 0.901 | 0.744 | 1.51E-11 | Mari |
| cDC1 | MRPL51 | 8.25E-09 | 0.599431476 | 0.718 | 0.592 | 0.000162338 | Mari |
| cDC1 | TMSB4X | 9.90E-24 | 0.599103144 | 1 | 0.984 | 1.95E-19 | Mari |
| cDC1 | MEF2C | 1.50E-08 | 0.598380401 | 0.714 | 0.536 | 0.000295903 | Mari |
| cDC1 | MRPL23 | 3.76E-07 | 0.597926894 | 0.575 | 0.391 | 0.007403595 | Mari |
| cDC1 | VAMP8 | 1.74E-16 | 0.596620671 | 0.905 | 0.826 | 3.43E-12 | Mari |
| cDC1 | UQCRFS1 | 2.15E-10 | 0.595396644 | 0.774 | 0.585 | 4.24E-06 | Mari |
| cDC1 | CUEDC2 | 1.41E-06 | 0.592675489 | 0.456 | 0.288 | 0.027789696 | Mari |
| cDC1 | MRPS24 | 1.22E-09 | 0.591381636 | 0.75 | 0.563 | 2.39E-05 | Mari |
| cDC1 | PDCD6 | 3.99E-09 | 0.589221109 | 0.75 | 0.563 | 7.85E-05 | Mari |
| cDC1 | TMEM179B | 3.39E-07 | 0.587878427 | 0.532 | 0.348 | 0.006679757 | Mari |
| cDC1 | TWF2 | 5.23E-11 | 0.587629701 | 0.821 | 0.677 | 1.03E-06 | Mari |
| cDC1 | MGST2 | 2.58E-09 | 0.586059778 | 0.655 | 0.466 | 5.07E-05 | Mari |
| cDC1 | PHB | 8.17E-08 | 0.585704256 | 0.687 | 0.522 | 0.001608747 | Mari |
| cDC1 | NDUFA4 | 5.06E-20 | 0.585517343 | 0.929 | 0.835 | 9.96E-16 | Mari |
| cDC1 | DCXR | 9.50E-11 | 0.585165273 | 0.798 | 0.606 | 1.87E-06 | Mari |
| cDC1 | CLNS1A | 2.63E-08 | 0.584543327 | 0.687 | 0.521 | 0.000517561 | Mari |
| cDC1 | FIBP | 3.80E-08 | 0.584374751 | 0.694 | 0.536 | 0.000747386 | Mari |
| cDC1 | ATP6V1G1 | 1.55E-12 | 0.583471224 | 0.849 | 0.685 | 3.05E-08 | Mari |
| cDC1 | NDUFS6 | 1.31E-13 | 0.582501849 | 0.889 | 0.694 | 2.57E-09 | Mari |
| cDC1 | HNRNPH2 | 2.29E-06 | 0.581365314 | 0.448 | 0.292 | 0.045151824 | Mari |
| cDC1 | COX7A2 | 9.80E-14 | 0.58056232 | 0.905 | 0.752 | 1.93E-09 | Mari |
| cDC1 | RNF7 | 7.75E-12 | 0.579857719 | 0.825 | 0.629 | 1.53E-07 | Mari |

| | | | | | | | |
|---|---|---|---|---|---|---|---|
| cDC1 | IDH3A | 3.45E-08 | 0.578655 | 0.615 | 0.415 | 0.000679998 | Mari |
| cDC1 | UQCRQ | 1.77E-10 | 0.5785629 | 0.81 | 0.681 | 3.48E-06 | Mari |
| cDC1 | CIAO2B | 1.64E-08 | 0.575242634 | 0.758 | 0.613 | 0.000323728 | Mari |
| cDC1 | TALDO1 | 2.94E-11 | 0.57380495 | 0.869 | 0.708 | 5.78E-07 | Mari |
| cDC1 | PSMA4 | 6.82E-08 | 0.572329596 | 0.786 | 0.638 | 0.001342512 | Mari |
| cDC1 | PSMB1 | 1.06E-13 | 0.571649707 | 0.889 | 0.784 | 2.08E-09 | Mari |
| cDC1 | CYBA | 1.08E-21 | 0.571480268 | 0.964 | 0.902 | 2.13E-17 | Mari |
| cDC1 | ANAPC15 | 3.28E-07 | 0.570137974 | 0.48 | 0.296 | 0.006452261 | Mari |
| cDC1 | LSM5 | 2.73E-09 | 0.567623864 | 0.714 | 0.572 | 5.37E-05 | Mari |
| cDC1 | JPT1 | 1.71E-13 | 0.567424123 | 0.905 | 0.783 | 3.37E-09 | Mari |
| cDC1 | BRK1 | 2.21E-12 | 0.566543187 | 0.865 | 0.722 | 4.35E-08 | Mari |
| cDC1 | SPCS2 | 1.30E-11 | 0.566162763 | 0.833 | 0.667 | 2.55E-07 | Mari |
| cDC1 | POMP | 7.04E-14 | 0.564970459 | 0.913 | 0.833 | 1.39E-09 | Mari |
| cDC1 | IER2 | 4.00E-08 | 0.564661049 | 0.905 | 0.836 | 0.000788287 | Mari |
| cDC1 | NDUFA3 | 5.42E-12 | 0.564592977 | 0.762 | 0.548 | 1.07E-07 | Mari |
| cDC1 | SNRPD3 | 7.44E-08 | 0.56414757 | 0.734 | 0.56 | 0.001464792 | Mari |
| cDC1 | MICOS13 | 2.04E-12 | 0.563839309 | 0.821 | 0.645 | 4.01E-08 | Mari |
| cDC1 | MAP3K13 | 5.96E-13 | 0.563221079 | 0.865 | 0.642 | 1.17E-08 | Mari |
| cDC1 | ZNRD1 | 1.45E-07 | 0.562295719 | 0.575 | 0.396 | 0.002847426 | Mari |
| cDC1 | PYCARD | 3.72E-12 | 0.562245427 | 0.893 | 0.77 | 7.32E-08 | Mari |
| cDC1 | GMFG | 2.22E-10 | 0.562167131 | 0.849 | 0.712 | 4.37E-06 | Mari |
| cDC1 | JTB | 4.51E-07 | 0.562029155 | 0.671 | 0.511 | 0.008875539 | Mari |
| cDC1 | COMMD2 | 6.49E-07 | 0.560771909 | 0.464 | 0.294 | 0.012777035 | Mari |
| cDC1 | NDUFA12 | 2.00E-09 | 0.559893892 | 0.762 | 0.57 | 3.94E-05 | Mari |
| cDC1 | UCP2 | 4.10E-09 | 0.55985935 | 0.937 | 0.835 | 8.07E-05 | Mari |
| cDC1 | SSBP1 | 9.88E-08 | 0.558751752 | 0.77 | 0.615 | 0.001945625 | Mari |
| cDC1 | BAX | 3.48E-09 | 0.558532789 | 0.77 | 0.581 | 6.86E-05 | Mari |
| cDC1 | CHMP2A | 2.70E-08 | 0.558459301 | 0.718 | 0.544 | 0.000531802 | Mari |
| cDC1 | MPC2 | 3.03E-08 | 0.554613158 | 0.734 | 0.563 | 0.000595967 | Mari |
| cDC1 | TMEM160 | 3.95E-08 | 0.554168636 | 0.639 | 0.465 | 0.000777763 | Mari |
| cDC1 | NDUFAF8 | 6.90E-07 | 0.553918548 | 0.544 | 0.381 | 0.013581842 | Mari |
| cDC1 | SP1101 | 6.82E-07 | 0.552902761 | 0.675 | 0.489 | 0.013417303 | Mari |
| cDC1 | EIF6 | 3.89E-12 | 0.552576743 | 0.865 | 0.713 | 7.66E-08 | Mari |
| cDC1 | PDCD10 | 2.61E-07 | 0.551250768 | 0.623 | 0.463 | 0.005133987 | Mari |
| cDC1 | SUB1 | 1.80E-20 | 0.549728213 | 0.968 | 0.919 | 3.55E-16 | Mari |
| cDC1 | DCTN3 | 1.64E-07 | 0.547567853 | 0.73 | 0.553 | 0.00323207 | Mari |
| cDC1 | SNRPD1 | 7.69E-09 | 0.547263936 | 0.794 | 0.634 | 0.00015144 | Mari |
| cDC1 | EFHD2 | 1.77E-13 | 0.546512524 | 0.94 | 0.835 | 3.49E-09 | Mari |
| cDC1 | MYL12B | 2.45E-17 | 0.544535057 | 0.913 | 0.841 | 4.82E-13 | Mari |
| cDC1 | LAMTOR5 | 1.25E-08 | 0.544178814 | 0.702 | 0.513 | 0.000245782 | Mari |
| cDC1 | HMGN3 | 1.19E-11 | 0.542115254 | 0.829 | 0.694 | 2.34E-07 | Mari |
| cDC1 | SUMO2 | 1.94E-17 | 0.541256934 | 0.921 | 0.848 | 3.83E-13 | Mari |
| cDC1 | DYNLRB1 | 3.91E-08 | 0.540829264 | 0.714 | 0.554 | 0.00076892 | Mari |
| cDC1 | HSD17B10 | 1.06E-06 | 0.540090722 | 0.583 | 0.42 | 0.020805395 | Mari |
| cDC1 | TPMT | 1.07E-08 | 0.539846353 | 0.774 | 0.617 | 0.00021035 | Mari |
| cDC1 | ARPC5L | 2.06E-07 | 0.538903474 | 0.79 | 0.614 | 0.004064211 | Mari |
| cDC1 | MRPS7 | 1.16E-07 | 0.535724371 | 0.544 | 0.364 | 0.002277257 | Mari |
| cDC1 | SCAND1 | 3.56E-10 | 0.533732506 | 0.754 | 0.577 | 7.01E-06 | Mari |
| cDC1 | COMMD7 | 2.06E-06 | 0.533723442 | 0.5 | 0.333 | 0.040541466 | Mari |
| cDC1 | TNFSF13B | 3.43E-07 | 0.533437417 | 0.631 | 0.477 | 0.006748327 | Mari |
| cDC1 | PSMD8 | 2.62E-11 | 0.532304801 | 0.869 | 0.694 | 5.15E-07 | Mari |
| cDC1 | NDUFB1 | 2.63E-11 | 0.532052684 | 0.865 | 0.659 | 5.19E-07 | Mari |
| cDC1 | GHITM | 1.77E-08 | 0.53190612 | 0.782 | 0.647 | 0.000348588 | Mari |
| cDC1 | GABARAP | 1.79E-18 | 0.531742955 | 0.956 | 0.883 | 3.53E-14 | Mari |
| cDC1 | LAP3 | 9.24E-12 | 0.530922739 | 0.952 | 0.826 | 1.82E-07 | Mari |
| cDC1 | SRP14 | 1.34E-22 | 0.529914501 | 0.96 | 0.887 | 2.63E-18 | Mari |

| | | | | | | | |
|---|---|---|---|---|---|---|---|
| cDC1 | MPG | 9.28E-08 | 0.527989813 | 0.738 | 0.588 | 0.00182708 | Mari |
| cDC1 | GSTP1 | 1.21E-16 | 0.525236839 | 0.976 | 0.902 | 2.38E-12 | Mari |
| cDC1 | RGS19 | 2.32E-06 | 0.525193822 | 0.687 | 0.547 | 0.045694735 | Mari |
| cDC1 | ST3GAL51 | 9.73E-07 | 0.523438279 | 0.619 | 0.437 | 0.019156885 | Mari |
| cDC1 | HLA-DMA | 5.42E-18 | 0.522850089 | 0.968 | 0.917 | 1.07E-13 | Mari |
| cDC1 | ATP5F1C | 1.16E-12 | 0.520976377 | 0.897 | 0.762 | 2.29E-08 | Mari |
| cDC1 | MCUR1 | 1.32E-06 | 0.519989361 | 0.627 | 0.456 | 0.026041678 | Mari |
| cDC1 | CLEC9A | 1.97E-11 | 0.519925561 | 0.94 | 0.843 | 3.88E-07 | Mari |
| cDC1 | RABAC1 | 1.13E-07 | 0.519689951 | 0.742 | 0.576 | 0.002224132 | Mari |
| cDC1 | PSMB3 | 2.88E-12 | 0.517288943 | 0.877 | 0.751 | 5.68E-08 | Mari |
| cDC1 | RAN | 6.95E-13 | 0.515417371 | 0.929 | 0.801 | 1.37E-08 | Mari |
| cDC1 | FGL2 | 6.85E-10 | 0.514386986 | 0.929 | 0.809 | 1.35E-05 | Mari |
| cDC1 | LMAN2 | 1.30E-08 | 0.513110567 | 0.71 | 0.526 | 0.000254944 | Mari |
| cDC1 | TUBA1B | 1.68E-08 | 0.510537395 | 0.96 | 0.886 | 0.000330302 | Mari |
| cDC1 | PSMA1 | 3.69E-09 | 0.507872085 | 0.845 | 0.687 | 7.26E-05 | Mari |
| cDC1 | ATP5MC1 | 7.38E-09 | 0.505853289 | 0.782 | 0.623 | 0.000145365 | Mari |
| cDC1 | MRPL43 | 9.59E-07 | 0.505756887 | 0.583 | 0.414 | 0.018883575 | Mari |
| cDC1 | TXNL4A | 1.59E-06 | 0.503778731 | 0.71 | 0.575 | 0.0312507 | Mari |
| cDC1 | CORO1A | 1.36E-12 | 0.500877489 | 0.952 | 0.84 | 2.67E-08 | Mari |
| cDC1 | S100A2 | 4.26E-37 | 2.222799149 | 0.398 | 0.074 | 8.38E-33 | Tb |
| cDC1 | NPHS1 | 8.56E-14 | 2.190675493 | 0.343 | 0.152 | 1.68E-09 | Tb |
| cDC1 | MRNIP | 1.71E-11 | 2.157250223 | 0.283 | 0.112 | 3.36E-07 | Tb |
| cDC1 | CDKL2 | 4.41E-13 | 2.155397349 | 0.251 | 0.083 | 8.69E-09 | Tb |
| cDC1 | S100A7 | 2.57E-27 | 2.105023122 | 0.386 | 0.093 | 5.06E-23 | Tb |
| cDC1 | GCNT2 | 5.71E-16 | 1.897664359 | 0.283 | 0.088 | 1.12E-11 | Tb |
| cDC1 | SLFN12L | 4.80E-14 | 1.837811236 | 0.299 | 0.105 | 9.45E-10 | Tb |
| cDC1 | CNR2 | 3.58E-18 | 1.725197684 | 0.287 | 0.078 | 7.05E-14 | Tb |
| cDC1 | S100A8 | 4.28E-45 | 1.700610595 | 0.641 | 0.17 | 8.42E-41 | Tb |
| cDC1 | RHOH | 8.70E-14 | 1.678044355 | 0.371 | 0.156 | 1.71E-09 | Tb |
| cDC1 | PILRA | 6.63E-19 | 1.670074413 | 0.311 | 0.091 | 1.31E-14 | Tb |
| cDC1 | KRT6A | 1.23E-30 | 1.669385534 | 0.382 | 0.082 | 2.41E-26 | Tb |
| cDC1 | KRT6B | 1.34E-22 | 1.592159614 | 0.275 | 0.055 | 2.64E-18 | Tb |
| cDC1 | DUSP16 | 2.41E-16 | 1.572613051 | 0.335 | 0.114 | 4.74E-12 | Tb |
| cDC1 | FUT10 | 3.46E-19 | 1.568799178 | 0.259 | 0.059 | 6.82E-15 | Tb |
| cDC1 | TPST1 | 9.33E-12 | 1.565280692 | 0.267 | 0.099 | 1.84E-07 | Tb |
| cDC1 | GRAMD1B | 6.88E-15 | 1.550592448 | 0.299 | 0.102 | 1.35E-10 | Tb |
| cDC1 | SFN | 1.90E-18 | 1.54085102 | 0.319 | 0.092 | 3.75E-14 | Tb |
| cDC1 | S100A9 | 1.79E-32 | 1.540737335 | 0.685 | 0.273 | 3.53E-28 | Tb |
| cDC1 | PSMD5 | 2.32E-09 | 1.519442984 | 0.279 | 0.12 | 4.56E-05 | Tb |
| cDC1 | ARHGAP25 | 9.04E-15 | 1.511099905 | 0.442 | 0.203 | 1.78E-10 | Tb |
| cDC1 | PLEKHG2 | 8.31E-19 | 1.491884717 | 0.53 | 0.243 | 1.64E-14 | Tb |
| cDC1 | INPP4B | 3.65E-08 | 1.486347847 | 0.382 | 0.209 | 0.000718682 | Tb |
| cDC1 | SLC22A23 | 1.69E-23 | 1.470493951 | 0.414 | 0.127 | 3.32E-19 | Tb |
| cDC1 | AXL | 8.93E-15 | 1.463483157 | 0.414 | 0.183 | 1.76E-10 | Tb |
| cDC1 | SLC35F1 | 4.54E-12 | 1.44674247 | 0.351 | 0.153 | 8.94E-08 | Tb |
| cDC1 | ROBO1 | 7.73E-12 | 1.418079709 | 0.418 | 0.208 | 1.52E-07 | Tb |
| cDC1 | ATP8B1 | 4.80E-10 | 1.397738172 | 0.251 | 0.098 | 9.44E-06 | Tb |
| cDC1 | PALD1 | 2.92E-17 | 1.395682735 | 0.323 | 0.101 | 5.75E-13 | Tb |
| cDC1 | XIST | 9.27E-29 | 1.39280979 | 0.649 | 0.253 | 1.82E-24 | Tb |
| cDC1 | CLIC4 | 2.32E-18 | 1.38642997 | 0.61 | 0.327 | 4.57E-14 | Tb |
| cDC1 | KRT17 | 1.94E-26 | 1.366169425 | 0.319 | 0.064 | 3.81E-22 | Tb |
| cDC1 | ENPP2 | 2.27E-09 | 1.358233355 | 0.414 | 0.223 | 4.47E-05 | Tb |
| cDC1 | SLC43A2 | 8.75E-23 | 1.318853858 | 0.625 | 0.3 | 1.72E-18 | Tb |
| cDC1 | ZNF44 | 1.31E-12 | 1.315666028 | 0.303 | 0.112 | 2.57E-08 | Tb |
| cDC1 | CCDC97 | 2.45E-09 | 1.313401537 | 0.327 | 0.153 | 4.81E-05 | Tb |
| cDC1 | KIF1B | 9.75E-14 | 1.300903621 | 0.486 | 0.245 | 1.92E-09 | Tb |

| | | | | | | | |
|---|---|---|---|---|---|---|---|
| cDC1 | PRKCB | 3.19E-27 | 1.300736002 | 0.729 | 0.364 | 6.28E-23 | Tb |
| cDC1 | FAM118A | 4.35E-18 | 1.289538987 | 0.49 | 0.222 | 8.56E-14 | Tb |
| cDC1 | KRT16 | 1.34E-16 | 1.286775403 | 0.307 | 0.092 | 2.65E-12 | Tb |
| cDC1 | PPP1R16B | 1.34E-13 | 1.283636809 | 0.315 | 0.114 | 2.63E-09 | Tb |
| cDC1 | EYA3 | 8.85E-12 | 1.267947107 | 0.378 | 0.17 | 1.74E-07 | Tb |
| cDC1 | SIMC1 | 4.63E-13 | 1.266698472 | 0.287 | 0.097 | 9.12E-09 | Tb |
| cDC1 | NYAP2 | 8.82E-17 | 1.265706712 | 0.622 | 0.328 | 1.74E-12 | Tb |
| cDC1 | C1QA | 8.04E-08 | 1.264750906 | 0.339 | 0.176 | 0.00158233 | Tb |
| cDC1 | PLPP1 | 5.75E-14 | 1.24127825 | 0.677 | 0.415 | 1.13E-09 | Tb |
| cDC1 | JMY | 2.48E-06 | 1.230338023 | 0.339 | 0.197 | 0.048907742 | Tb |
| cDC1 | AOAH | 3.07E-15 | 1.224600885 | 0.59 | 0.335 | 6.03E-11 | Tb |
| cDC1 | SENP2 | 8.77E-10 | 1.219976646 | 0.303 | 0.133 | 1.73E-05 | Tb |
| cDC1 | INPP5B | 4.76E-11 | 1.215957512 | 0.291 | 0.115 | 9.37E-07 | Tb |
| cDC1 | PRIM2 | 2.93E-14 | 1.207038016 | 0.382 | 0.152 | 5.77E-10 | Tb |
| cDC1 | ATXN7L1 | 4.10E-17 | 1.199506676 | 0.311 | 0.094 | 8.08E-13 | Tb |
| cDC1 | ARHGAP12 | 3.94E-10 | 1.190284302 | 0.275 | 0.11 | 7.75E-06 | Tb |
| cDC1 | PPARD | 5.75E-13 | 1.1888028 | 0.462 | 0.226 | 1.13E-08 | Tb |
| cDC1 | ZNF609 | 7.62E-16 | 1.186446361 | 0.438 | 0.185 | 1.50E-11 | Tb |
| cDC1 | HOMER1 | 6.43E-08 | 1.186148731 | 0.538 | 0.339 | 0.001265836 | Tb |
| cDC1 | FUT8 | 1.42E-16 | 1.184872863 | 0.677 | 0.392 | 2.80E-12 | Tb |
| cDC1 | MRC2 | 3.32E-12 | 1.1770463 | 0.454 | 0.224 | 6.53E-08 | Tb |
| cDC1 | DDI2 | 6.74E-09 | 1.176350522 | 0.335 | 0.161 | 0.000132707 | Tb |
| cDC1 | ERBIN | 4.58E-22 | 1.174998788 | 0.837 | 0.541 | 9.02E-18 | Tb |
| cDC1 | CDK19 | 6.13E-11 | 1.173252543 | 0.442 | 0.23 | 1.21E-06 | Tb |
| cDC1 | USF3 | 1.98E-06 | 1.172622763 | 0.283 | 0.144 | 0.038961412 | Tb |
| cDC1 | CLEC16A | 5.16E-14 | 1.170032907 | 0.474 | 0.227 | 1.01E-09 | Tb |
| cDC1 | PLEKHM3 | 2.68E-18 | 1.167712602 | 0.757 | 0.484 | 5.28E-14 | Tb |
| cDC1 | RFX7 | 1.92E-09 | 1.156832345 | 0.319 | 0.155 | 3.78E-05 | Tb |
| cDC1 | ANKDD1A | 5.27E-09 | 1.15539025 | 0.275 | 0.123 | 0.000103667 | Tb |
| cDC1 | BCL2L14 | 1.05E-14 | 1.150382994 | 0.466 | 0.223 | 2.07E-10 | Tb |
| cDC1 | KDM7A-DT | 3.99E-11 | 1.146190216 | 0.319 | 0.13 | 7.85E-07 | Tb |
| cDC1 | ADAP2 | 1.42E-13 | 1.145805158 | 0.522 | 0.278 | 2.79E-09 | Tb |
| cDC1 | SMYD3 | 1.32E-13 | 1.14330762 | 0.486 | 0.249 | 2.59E-09 | Tb |
| cDC1 | P3H2 | 3.98E-07 | 1.139995286 | 0.255 | 0.123 | 0.007839013 | Tb |
| cDC1 | ABCC1 | 2.05E-11 | 1.134692356 | 0.307 | 0.12 | 4.03E-07 | Tb |
| cDC1 | TMEM39A | 6.26E-11 | 1.132443084 | 0.542 | 0.324 | 1.23E-06 | Tb |
| cDC1 | EEF2K | 1.80E-08 | 1.128913492 | 0.267 | 0.116 | 0.000353801 | Tb |
| cDC1 | RPRD2 | 8.17E-09 | 1.119901998 | 0.303 | 0.143 | 0.000160743 | Tb |
| cDC1 | TENT2 | 6.06E-17 | 1.116688513 | 0.594 | 0.301 | 1.19E-12 | Tb |
| cDC1 | DLEU2 | 6.16E-20 | 1.110153582 | 0.853 | 0.614 | 1.21E-15 | Tb |
| cDC1 | ASCL2 | 1.21E-10 | 1.099373404 | 0.287 | 0.116 | 2.38E-06 | Tb |
| cDC1 | TOP3A | 1.55E-10 | 1.096731939 | 0.331 | 0.145 | 3.06E-06 | Tb |
| cDC1 | KRT14 | 1.85E-34 | 1.0960602 | 0.633 | 0.202 | 3.65E-30 | Tb |
| cDC1 | RNF213 | 3.65E-09 | 1.09404834 | 0.315 | 0.144 | 7.19E-05 | Tb |
| cDC1 | SLC10A1 | 1.05E-08 | 1.086732705 | 0.263 | 0.114 | 0.000206406 | Tb |
| cDC1 | GNAI1 | 9.81E-10 | 1.078283031 | 0.315 | 0.14 | 1.93E-05 | Tb |
| cDC1 | USP25 | 7.40E-14 | 1.069083347 | 0.558 | 0.291 | 1.46E-09 | Tb |
| cDC1 | LPAR6 | 7.08E-10 | 1.06793637 | 0.323 | 0.144 | 1.39E-05 | Tb |
| cDC1 | SUSD3 | 5.20E-18 | 1.067562142 | 0.59 | 0.288 | 1.02E-13 | Tb |
| cDC1 | TAF4B | 1.76E-17 | 1.05885856 | 0.526 | 0.247 | 3.45E-13 | Tb |
| cDC1 | LRRC23 | 1.09E-11 | 1.055263181 | 0.506 | 0.278 | 2.14E-07 | Tb |
| cDC1 | TMCC1 | 2.24E-10 | 1.054010701 | 0.394 | 0.195 | 4.41E-06 | Tb |
| cDC1 | BMPR2 | 5.83E-11 | 1.053000757 | 0.426 | 0.218 | 1.15E-06 | Tb |
| cDC1 | BET1 | 1.27E-16 | 1.052813756 | 0.39 | 0.14 | 2.50E-12 | Tb |
| cDC1 | FNDC3B | 1.06E-12 | 1.050188629 | 0.582 | 0.326 | 2.09E-08 | Tb |
| cDC1 | STK38L | 1.72E-14 | 1.045789572 | 0.598 | 0.322 | 3.39E-10 | Tb |

| | | | | | | | |
|---|---|---|---|---|---|---|---|
| cDC1 | PHTF1 | 2.12E-14 | 1.042670319 | 0.482 | 0.222 | 4.17E-10 | Tb |
| cDC1 | GALC | 2.11E-08 | 1.040968018 | 0.339 | 0.172 | 0.000415312 | Tb |
| cDC1 | FAM76A | 5.61E-09 | 1.039648172 | 0.259 | 0.109 | 0.000110377 | Tb |
| cDC1 | ADIPOR2 | 1.56E-13 | 1.037884701 | 0.382 | 0.159 | 3.07E-09 | Tb |
| cDC1 | FTX | 2.62E-14 | 1.037359142 | 0.486 | 0.226 | 5.15E-10 | Tb |
| cDC1 | SRGAP2 | 7.60E-18 | 1.035547553 | 0.657 | 0.382 | 1.50E-13 | Tb |
| cDC1 | TYW1 | 2.33E-11 | 1.034864288 | 0.418 | 0.202 | 4.58E-07 | Tb |
| cDC1 | ITGAL | 7.34E-16 | 1.032353008 | 0.61 | 0.336 | 1.45E-11 | Tb |
| cDC1 | ABCA7 | 3.34E-10 | 1.031550281 | 0.283 | 0.114 | 6.58E-06 | Tb |
| cDC1 | MT-ATP6 | 5.26E-26 | 1.03049054 | 1 | 0.91 | 1.04E-21 | Tb |
| cDC1 | KCNK13 | 3.48E-09 | 1.029302642 | 0.331 | 0.156 | 6.84E-05 | Tb |
| cDC1 | GBP1 | 1.02E-11 | 1.022915959 | 0.295 | 0.116 | 2.01E-07 | Tb |
| cDC1 | STIM2 | 9.52E-12 | 1.022473978 | 0.606 | 0.381 | 1.87E-07 | Tb |
| cDC1 | ZNF33A | 5.18E-08 | 1.022152345 | 0.359 | 0.182 | 0.001020095 | Tb |
| cDC1 | TYW1B | 6.87E-13 | 1.021787695 | 0.367 | 0.15 | 1.35E-08 | Tb |
| cDC1 | FAM222B | 1.06E-09 | 1.021123306 | 0.275 | 0.111 | 2.08E-05 | Tb |
| cDC1 | CTNND1 | 3.00E-07 | 1.019177856 | 0.271 | 0.13 | 0.005905843 | Tb |
| cDC1 | DLEU1 | 3.56E-19 | 1.018763066 | 0.594 | 0.283 | 7.00E-15 | Tb |
| cDC1 | TBC1D17 | 2.37E-07 | 1.014928158 | 0.263 | 0.122 | 0.004661828 | Tb |
| cDC1 | SNX14 | 8.48E-10 | 1.011801732 | 0.434 | 0.23 | 1.67E-05 | Tb |
| cDC1 | SIK2 | 9.35E-10 | 1.011655679 | 0.534 | 0.31 | 1.84E-05 | Tb |
| cDC1 | MFSD14B | 1.71E-14 | 1.010043226 | 0.622 | 0.339 | 3.36E-10 | Tb |
| cDC1 | MX2 | 5.23E-07 | 1.002040647 | 0.522 | 0.356 | 0.010299569 | Tb |
| cDC1 | MAP1S | 7.10E-07 | 1.000841767 | 0.275 | 0.132 | 0.013980005 | Tb |
| cDC1 | ENTPD7 | 3.76E-07 | 0.997995763 | 0.287 | 0.143 | 0.007402768 | Tb |
| cDC1 | RAP1GAP2 | 5.08E-15 | 0.997318164 | 0.331 | 0.119 | 1.00E-10 | Tb |
| cDC1 | LPIN2 | 6.62E-15 | 0.996967754 | 0.454 | 0.202 | 1.30E-10 | Tb |
| cDC1 | LTB | 5.60E-13 | 0.994281567 | 0.637 | 0.381 | 1.10E-08 | Tb |
| cDC1 | PARP9 | 1.28E-07 | 0.993451373 | 0.327 | 0.168 | 0.002517323 | Tb |
| cDC1 | ELF2 | 1.86E-15 | 0.992364985 | 0.697 | 0.411 | 3.66E-11 | Tb |
| cDC1 | ACOT7 | 1.89E-11 | 0.991189253 | 0.51 | 0.272 | 3.71E-07 | Tb |
| cDC1 | IFI27 | 2.72E-08 | 0.989625429 | 0.319 | 0.15 | 0.00053637 | Tb |
| cDC1 | TBC1D9 | 5.34E-12 | 0.988999196 | 0.598 | 0.351 | 1.05E-07 | Tb |
| cDC1 | CEP192 | 1.33E-11 | 0.988261702 | 0.335 | 0.136 | 2.61E-07 | Tb |
| cDC1 | PPP3CC | 9.50E-12 | 0.987559805 | 0.494 | 0.251 | 1.87E-07 | Tb |
| cDC1 | DOP1A | 5.80E-12 | 0.986012421 | 0.558 | 0.319 | 1.14E-07 | Tb |
| cDC1 | TLR10 | 4.79E-15 | 0.983372992 | 0.498 | 0.226 | 9.44E-11 | Tb |
| cDC1 | MAP3K21 | 2.35E-08 | 0.983237116 | 0.303 | 0.144 | 0.000463509 | Tb |
| cDC1 | TCOF1 | 2.05E-10 | 0.982519507 | 0.406 | 0.2 | 4.03E-06 | Tb |
| cDC1 | BCOR | 3.12E-12 | 0.98210062 | 0.406 | 0.18 | 6.14E-08 | Tb |
| cDC1 | MBTD1 | 1.84E-10 | 0.978562924 | 0.39 | 0.188 | 3.62E-06 | Tb |
| cDC1 | LRRC8D | 4.03E-08 | 0.977575954 | 0.303 | 0.145 | 0.000792298 | Tb |
| cDC1 | KDM7A | 2.72E-18 | 0.974311071 | 0.673 | 0.331 | 5.36E-14 | Tb |
| cDC1 | SEC23B | 5.48E-10 | 0.973138362 | 0.406 | 0.202 | 1.08E-05 | Tb |
| cDC1 | NFATC3 | 4.15E-11 | 0.97093486 | 0.406 | 0.189 | 8.17E-07 | Tb |
| cDC1 | BIN3 | 1.16E-07 | 0.968437197 | 0.267 | 0.122 | 0.002281867 | Tb |
| cDC1 | TBC1D22A | 1.67E-16 | 0.963152155 | 0.745 | 0.5 | 3.29E-12 | Tb |
| cDC1 | LRRC8C | 1.85E-08 | 0.961683865 | 0.375 | 0.203 | 0.000363819 | Tb |
| cDC1 | NOL10 | 2.76E-08 | 0.96094946 | 0.355 | 0.174 | 0.000543666 | Tb |
| cDC1 | SAMD9L | 5.55E-19 | 0.960747624 | 0.566 | 0.252 | 1.09E-14 | Tb |
| cDC1 | GNAO1 | 1.37E-17 | 0.960729075 | 0.602 | 0.299 | 2.70E-13 | Tb |
| cDC1 | STAT5A | 6.42E-13 | 0.959810403 | 0.442 | 0.2 | 1.26E-08 | Tb |
| cDC1 | TET3 | 1.17E-09 | 0.957656826 | 0.39 | 0.193 | 2.29E-05 | Tb |
| cDC1 | EBF4 | 1.46E-13 | 0.957193793 | 0.414 | 0.185 | 2.88E-09 | Tb |
| cDC1 | YPEL2 | 1.06E-08 | 0.956865149 | 0.426 | 0.227 | 0.0002086 | Tb |
| cDC1 | PSEN1 | 4.83E-13 | 0.955270464 | 0.598 | 0.34 | 9.50E-09 | Tb |

| | | | | | | | |
|---|---|---|---|---|---|---|---|
| cDC1 | SPDYA | 5.68E-07 | 0.951589971 | 0.339 | 0.188 | 0.01119014 | Tb |
| cDC1 | PHF21A | 1.19E-12 | 0.951048743 | 0.446 | 0.214 | 2.34E-08 | Tb |
| cDC1 | AKAP10 | 2.59E-07 | 0.951039623 | 0.299 | 0.151 | 0.00510382 | Tb |
| cDC1 | JPX | 2.25E-10 | 0.950718759 | 0.506 | 0.28 | 4.42E-06 | Tb |
| cDC1 | ADAM19 | 4.77E-21 | 0.949928978 | 0.781 | 0.415 | 9.39E-17 | Tb |
| cDC1 | KDM5B | 4.59E-11 | 0.947977833 | 0.375 | 0.169 | 9.03E-07 | Tb |
| cDC1 | UBE2W | 9.20E-14 | 0.947501788 | 0.673 | 0.405 | 1.81E-09 | Tb |
| cDC1 | ZNF700 | 2.04E-11 | 0.947189141 | 0.343 | 0.143 | 4.02E-07 | Tb |
| cDC1 | PPARGC1B | 5.18E-09 | 0.94527942 | 0.291 | 0.127 | 0.000101905 | Tb |
| cDC1 | MSL2 | 5.72E-08 | 0.944625046 | 0.287 | 0.136 | 0.001126621 | Tb |
| cDC1 | GNB4 | 6.29E-20 | 0.944352145 | 0.805 | 0.485 | 1.24E-15 | Tb |
| cDC1 | UBR1 | 8.59E-15 | 0.943739642 | 0.446 | 0.197 | 1.69E-10 | Tb |
| cDC1 | SP140 | 1.89E-13 | 0.942871536 | 0.602 | 0.345 | 3.72E-09 | Tb |
| cDC1 | CDKL5 | 7.62E-08 | 0.942677916 | 0.418 | 0.232 | 0.001499385 | Tb |
| cDC1 | ZNF532 | 3.62E-16 | 0.942014921 | 0.359 | 0.124 | 7.12E-12 | Tb |
| cDC1 | MPZL1 | 3.25E-10 | 0.939348185 | 0.398 | 0.192 | 6.39E-06 | Tb |
| cDC1 | UBAP2 | 3.55E-10 | 0.938135517 | 0.486 | 0.276 | 6.98E-06 | Tb |
| cDC1 | TRERF1 | 5.07E-13 | 0.936950181 | 0.809 | 0.594 | 9.97E-09 | Tb |
| cDC1 | SMG6 | 6.77E-10 | 0.934443189 | 0.323 | 0.142 | 1.33E-05 | Tb |
| cDC1 | GBF1 | 3.61E-13 | 0.934369895 | 0.526 | 0.261 | 7.10E-09 | Tb |
| cDC1 | ANKRD44 | 3.61E-18 | 0.93203908 | 0.829 | 0.539 | 7.11E-14 | Tb |
| cDC1 | FCF1 | 1.02E-08 | 0.931191738 | 0.259 | 0.11 | 0.000200614 | Tb |
| cDC1 | KIAA1958 | 9.85E-15 | 0.930918549 | 0.657 | 0.36 | 1.94E-10 | Tb |
| cDC1 | MT-CYB | 1.58E-16 | 0.930912823 | 0.992 | 0.928 | 3.10E-12 | Tb |
| cDC1 | YAF2 | 1.04E-08 | 0.930513431 | 0.57 | 0.338 | 0.000204904 | Tb |
| cDC1 | TNRC18 | 2.30E-16 | 0.930168785 | 0.526 | 0.234 | 4.53E-12 | Tb |
| cDC1 | CRLF3 | 3.23E-11 | 0.929303698 | 0.43 | 0.21 | 6.37E-07 | Tb |
| cDC1 | GGA2 | 1.03E-11 | 0.926946067 | 0.454 | 0.219 | 2.03E-07 | Tb |
| cDC1 | SLC8A1-AS1 | 1.45E-08 | 0.926771133 | 0.331 | 0.158 | 0.000284492 | Tb |
| cDC1 | BRF1 | 1.08E-07 | 0.92563049 | 0.315 | 0.15 | 0.002124765 | Tb |
| cDC1 | MYO1F | 1.54E-16 | 0.923908375 | 0.713 | 0.444 | 3.02E-12 | Tb |
| cDC1 | GNPTAB | 8.61E-12 | 0.922745949 | 0.486 | 0.245 | 1.70E-07 | Tb |
| cDC1 | SLC25A17 | 7.07E-19 | 0.919738849 | 0.554 | 0.24 | 1.39E-14 | Tb |
| cDC1 | ABL1 | 2.83E-15 | 0.914288116 | 0.633 | 0.351 | 5.56E-11 | Tb |
| cDC1 | ZNF366 | 4.14E-12 | 0.914191385 | 0.669 | 0.395 | 8.16E-08 | Tb |
| cDC1 | MED14 | 2.86E-10 | 0.913322058 | 0.367 | 0.168 | 5.63E-06 | Tb |
| cDC1 | KDM1B | 1.07E-12 | 0.912406281 | 0.602 | 0.326 | 2.11E-08 | Tb |
| cDC1 | ADAMTSL4-AS | 8.41E-12 | 0.910104571 | 0.558 | 0.316 | 1.66E-07 | Tb |
| cDC1 | N4BP2 | 3.28E-13 | 0.909681643 | 0.641 | 0.382 | 6.47E-09 | Tb |
| cDC1 | MAST2 | 1.13E-11 | 0.908601021 | 0.287 | 0.107 | 2.22E-07 | Tb |
| cDC1 | ZNF10 | 2.57E-11 | 0.908573909 | 0.382 | 0.181 | 5.07E-07 | Tb |
| cDC1 | SLC27A4 | 2.49E-17 | 0.908344065 | 0.697 | 0.397 | 4.91E-13 | Tb |
| cDC1 | PITPNA | 1.01E-17 | 0.906396714 | 0.813 | 0.569 | 1.98E-13 | Tb |
| cDC1 | CCDC138 | 1.52E-10 | 0.90217178 | 0.315 | 0.132 | 2.99E-06 | Tb |
| cDC1 | GPHN | 4.97E-09 | 0.896439799 | 0.402 | 0.211 | 9.78E-05 | Tb |
| cDC1 | MOB3B | 3.96E-08 | 0.895394916 | 0.311 | 0.15 | 0.000779196 | Tb |
| cDC1 | ITFG1 | 6.84E-14 | 0.893757714 | 0.554 | 0.282 | 1.35E-09 | Tb |
| cDC1 | FBXL20 | 2.85E-08 | 0.893032042 | 0.39 | 0.211 | 0.000561935 | Tb |
| cDC1 | MIR29B2CHG | 1.45E-13 | 0.891317453 | 0.319 | 0.112 | 2.85E-09 | Tb |
| cDC1 | ILF3-DT | 1.20E-13 | 0.891302894 | 0.323 | 0.117 | 2.35E-09 | Tb |
| cDC1 | PPP2R3C | 2.18E-12 | 0.889427916 | 0.418 | 0.18 | 4.30E-08 | Tb |
| cDC1 | RMI2 | 7.83E-07 | 0.887604845 | 0.275 | 0.134 | 0.015422061 | Tb |
| cDC1 | INPP5D | 5.75E-12 | 0.88525794 | 0.745 | 0.523 | 1.13E-07 | Tb |
| cDC1 | CSNK1G1 | 1.37E-11 | 0.885168386 | 0.382 | 0.173 | 2.69E-07 | Tb |
| cDC1 | ENPP1 | 2.34E-15 | 0.88439361 | 0.797 | 0.559 | 4.61E-11 | Tb |
| cDC1 | PPA2 | 7.85E-15 | 0.883651345 | 0.649 | 0.374 | 1.54E-10 | Tb |

| | | | | | | | |
|---|---|---|---|---|---|---|---|
| cDC1 | DENND5B | 6.33E-12 | 0.88212715 | 0.367 | 0.156 | 1.25E-07 | Tb |
| cDC1 | NECTIN2 | 1.95E-11 | 0.88197995 | 0.578 | 0.331 | 3.84E-07 | Tb |
| cDC1 | MT-CO3 | 7.31E-19 | 0.877844162 | 1 | 0.938 | 1.44E-14 | Tb |
| cDC1 | LATS2 | 3.89E-07 | 0.876732129 | 0.315 | 0.161 | 0.007659668 | Tb |
| cDC1 | HSDL2 | 2.27E-13 | 0.872919759 | 0.673 | 0.398 | 4.46E-09 | Tb |
| cDC1 | ELMO1 | 1.09E-20 | 0.87234105 | 0.896 | 0.683 | 2.14E-16 | Tb |
| cDC1 | VAMP5 | 4.45E-14 | 0.871195922 | 0.733 | 0.457 | 8.75E-10 | Tb |
| cDC1 | MON2 | 5.42E-08 | 0.869088787 | 0.51 | 0.312 | 0.001066884 | Tb |
| cDC1 | CLCN6 | 2.05E-06 | 0.866912513 | 0.255 | 0.124 | 0.040350732 | Tb |
| cDC1 | PLXNC1 | 2.31E-08 | 0.866783819 | 0.414 | 0.226 | 0.000455163 | Tb |
| cDC1 | DOCK11 | 6.01E-13 | 0.866444287 | 0.558 | 0.288 | 1.18E-08 | Tb |
| cDC1 | PAN3 | 5.33E-14 | 0.863168378 | 0.649 | 0.376 | 1.05E-09 | Tb |
| cDC1 | VPS13B | 4.37E-13 | 0.862277282 | 0.637 | 0.365 | 8.60E-09 | Tb |
| cDC1 | AHCYL2 | 1.77E-11 | 0.85950531 | 0.542 | 0.306 | 3.49E-07 | Tb |
| cDC1 | RAB11FIP3 | 1.13E-08 | 0.856885997 | 0.331 | 0.16 | 0.000222711 | Tb |
| cDC1 | LDLRAD3 | 2.07E-12 | 0.856491187 | 0.677 | 0.411 | 4.08E-08 | Tb |
| cDC1 | SLC39A11 | 2.73E-12 | 0.856055123 | 0.394 | 0.172 | 5.37E-08 | Tb |
| cDC1 | SRGAP2C | 1.83E-14 | 0.855866791 | 0.398 | 0.159 | 3.61E-10 | Tb |
| cDC1 | VAV1 | 2.77E-12 | 0.855662677 | 0.414 | 0.19 | 5.45E-08 | Tb |
| cDC1 | COP1 | 3.47E-15 | 0.855448765 | 0.721 | 0.426 | 6.83E-11 | Tb |
| cDC1 | NLK | 2.26E-10 | 0.855294724 | 0.303 | 0.122 | 4.45E-06 | Tb |
| cDC1 | CCDC91 | 2.17E-16 | 0.853818664 | 0.789 | 0.457 | 4.28E-12 | Tb |
| cDC1 | PARP14 | 5.52E-16 | 0.853729916 | 0.574 | 0.28 | 1.09E-11 | Tb |
| cDC1 | NCOA1 | 1.17E-14 | 0.852663982 | 0.725 | 0.44 | 2.30E-10 | Tb |
| cDC1 | ZFYVE1 | 6.36E-09 | 0.851210639 | 0.295 | 0.131 | 0.000125159 | Tb |
| cDC1 | ZBTB20 | 6.08E-14 | 0.850890527 | 0.618 | 0.353 | 1.20E-09 | Tb |
| cDC1 | FAF1 | 3.34E-15 | 0.850072866 | 0.701 | 0.425 | 6.58E-11 | Tb |
| cDC1 | PPM1H | 5.58E-11 | 0.849965306 | 0.482 | 0.261 | 1.10E-06 | Tb |
| cDC1 | RGS12 | 3.03E-14 | 0.849454565 | 0.546 | 0.272 | 5.96E-10 | Tb |
| cDC1 | TNS3 | 1.66E-07 | 0.849411855 | 0.267 | 0.122 | 0.003267693 | Tb |
| cDC1 | CD274 | 5.05E-07 | 0.849053737 | 0.259 | 0.122 | 0.009932097 | Tb |
| cDC1 | NT5DC3 | 3.63E-07 | 0.848521879 | 0.299 | 0.149 | 0.007149088 | Tb |
| cDC1 | RIPK2 | 6.76E-10 | 0.847589152 | 0.757 | 0.535 | 1.33E-05 | Tb |
| cDC1 | MIR924HG | 6.41E-13 | 0.847211773 | 0.558 | 0.301 | 1.26E-08 | Tb |
| cDC1 | EVI5 | 1.00E-11 | 0.846651734 | 0.39 | 0.174 | 1.98E-07 | Tb |
| cDC1 | ATP2B4 | 6.22E-07 | 0.843732955 | 0.45 | 0.265 | 0.012246806 | Tb |
| cDC1 | SELENOI | 2.40E-10 | 0.843541431 | 0.43 | 0.214 | 4.72E-06 | Tb |
| cDC1 | CHAF1A | 5.27E-07 | 0.842590059 | 0.442 | 0.262 | 0.01036577 | Tb |
| cDC1 | LRRK1 | 6.72E-13 | 0.842347418 | 0.594 | 0.348 | 1.32E-08 | Tb |
| cDC1 | BCL11A | 3.80E-11 | 0.839932914 | 0.558 | 0.328 | 7.49E-07 | Tb |
| cDC1 | 8-Mar | 1.98E-10 | 0.83797539 | 0.49 | 0.256 | 3.90E-06 | Tb |
| cDC1 | SLMAP | 2.86E-09 | 0.836838667 | 0.434 | 0.226 | 5.63E-05 | Tb |
| cDC1 | UBE2Q2 | 1.70E-08 | 0.836707684 | 0.446 | 0.25 | 0.000335244 | Tb |
| cDC1 | PHLPP1 | 4.70E-11 | 0.835125039 | 0.625 | 0.412 | 9.26E-07 | Tb |
| cDC1 | RSRC1 | 3.47E-13 | 0.83507366 | 0.542 | 0.274 | 6.84E-09 | Tb |
| cDC1 | TFCP2 | 2.94E-09 | 0.83496299 | 0.355 | 0.173 | 5.80E-05 | Tb |
| cDC1 | LIMD1 | 7.24E-07 | 0.834893397 | 0.343 | 0.189 | 0.014245952 | Tb |
| cDC1 | RCBTB2 | 9.12E-08 | 0.834757323 | 0.295 | 0.141 | 0.001794655 | Tb |
| cDC1 | FAM168A | 4.82E-10 | 0.834698934 | 0.311 | 0.132 | 9.49E-06 | Tb |
| cDC1 | MGME1 | 2.02E-07 | 0.83358345 | 0.359 | 0.188 | 0.003972756 | Tb |
| cDC1 | NUP88 | 3.49E-10 | 0.832742923 | 0.367 | 0.167 | 6.87E-06 | Tb |
| cDC1 | CDR2 | 8.01E-09 | 0.832528497 | 0.478 | 0.274 | 0.000157721 | Tb |
| cDC1 | LRBA | 8.30E-15 | 0.83102705 | 0.853 | 0.586 | 1.63E-10 | Tb |
| cDC1 | NFX1 | 2.80E-12 | 0.830279566 | 0.47 | 0.23 | 5.51E-08 | Tb |
| cDC1 | BCAT1 | 1.42E-14 | 0.829585972 | 0.689 | 0.419 | 2.79E-10 | Tb |
| cDC1 | TYMP | 1.08E-22 | 0.828071971 | 0.972 | 0.806 | 2.12E-18 | Tb |

| | | | | | | | |
|---|---|---|---|---|---|---|---|
| cDC1 | STX12 | 7.13E-17 | 0.827550684 | 0.876 | 0.641 | 1.40E-12 | Tb |
| cDC1 | MLLT10 | 1.67E-13 | 0.827318462 | 0.502 | 0.249 | 3.30E-09 | Tb |
| cDC1 | TESK2 | 9.95E-10 | 0.825808937 | 0.307 | 0.134 | 1.96E-05 | Tb |
| cDC1 | KLF7 | 7.85E-08 | 0.825343794 | 0.41 | 0.236 | 0.001544408 | Tb |
| cDC1 | APOL1 | 1.67E-09 | 0.824736812 | 0.335 | 0.153 | 3.28E-05 | Tb |
| cDC1 | PLD3 | 3.23E-12 | 0.824055134 | 0.49 | 0.241 | 6.35E-08 | Tb |
| cDC1 | CARS2 | 7.59E-13 | 0.821869912 | 0.478 | 0.231 | 1.49E-08 | Tb |
| cDC1 | MTF2 | 5.97E-10 | 0.821091191 | 0.422 | 0.208 | 1.18E-05 | Tb |
| cDC1 | ATG7 | 1.73E-12 | 0.820861559 | 0.442 | 0.21 | 3.41E-08 | Tb |
| cDC1 | MFSD14C | 2.71E-13 | 0.820820824 | 0.538 | 0.277 | 5.33E-09 | Tb |
| cDC1 | TMEM38B | 8.90E-12 | 0.820066172 | 0.622 | 0.366 | 1.75E-07 | Tb |
| cDC1 | STIM1 | 2.12E-09 | 0.818337328 | 0.41 | 0.217 | 4.18E-05 | Tb |
| cDC1 | SMC4 | 7.23E-07 | 0.817214815 | 0.47 | 0.278 | 0.014228948 | Tb |
| cDC1 | ARSB | 6.23E-09 | 0.814523712 | 0.494 | 0.285 | 0.000122614 | Tb |
| cDC1 | P2RY14 | 2.44E-14 | 0.814426679 | 0.801 | 0.527 | 4.80E-10 | Tb |
| cDC1 | DIPK1A | 8.19E-09 | 0.814033344 | 0.462 | 0.264 | 0.000161216 | Tb |
| cDC1 | SELL | 7.40E-11 | 0.813303929 | 0.59 | 0.368 | 1.46E-06 | Tb |
| cDC1 | PPP1R12B | 1.92E-08 | 0.812944894 | 0.303 | 0.141 | 0.00037735 | Tb |
| cDC1 | GALNT7 | 8.55E-15 | 0.812703931 | 0.578 | 0.299 | 1.68E-10 | Tb |
| cDC1 | NRF1 | 3.37E-08 | 0.812594657 | 0.319 | 0.153 | 0.000662756 | Tb |
| cDC1 | ULK4 | 1.04E-09 | 0.811415479 | 0.371 | 0.18 | 2.04E-05 | Tb |
| cDC1 | ADGRE5 | 6.94E-14 | 0.809848535 | 0.586 | 0.315 | 1.37E-09 | Tb |
| cDC1 | CD80 | 2.83E-08 | 0.809396701 | 0.458 | 0.261 | 0.000556925 | Tb |
| cDC1 | PIP5K1A | 6.60E-11 | 0.808724039 | 0.462 | 0.241 | 1.30E-06 | Tb |
| cDC1 | ZNF516 | 1.03E-16 | 0.808029908 | 0.801 | 0.518 | 2.02E-12 | Tb |
| cDC1 | TRAK1 | 1.48E-19 | 0.807888551 | 0.876 | 0.584 | 2.92E-15 | Tb |
| cDC1 | RNF19A | 5.69E-11 | 0.805470485 | 0.586 | 0.351 | 1.12E-06 | Tb |
| cDC1 | SLC11A2 | 7.89E-10 | 0.80492621 | 0.283 | 0.116 | 1.55E-05 | Tb |
| cDC1 | HERC4 | 2.18E-11 | 0.804104723 | 0.51 | 0.266 | 4.30E-07 | Tb |
| cDC1 | PHACTR2 | 1.39E-06 | 0.803629867 | 0.319 | 0.173 | 0.027331065 | Tb |
| cDC1 | ESYT2 | 7.53E-12 | 0.802625101 | 0.817 | 0.623 | 1.48E-07 | Tb |
| cDC1 | HIP1 | 1.12E-11 | 0.802408705 | 0.657 | 0.43 | 2.20E-07 | Tb |
| cDC1 | STAT3 | 2.44E-18 | 0.802050404 | 0.821 | 0.558 | 4.81E-14 | Tb |
| cDC1 | NUMB | 6.78E-13 | 0.801342477 | 0.689 | 0.412 | 1.33E-08 | Tb |
| cDC1 | ST3GAL1 | 5.14E-10 | 0.800264277 | 0.466 | 0.261 | 1.01E-05 | Tb |
| cDC1 | CSF2RB | 4.59E-10 | 0.799917037 | 0.406 | 0.195 | 9.03E-06 | Tb |
| cDC1 | PRPSAP1 | 1.27E-09 | 0.797565794 | 0.315 | 0.135 | 2.50E-05 | Tb |
| cDC1 | MBD5 | 8.00E-10 | 0.794733965 | 0.442 | 0.231 | 1.57E-05 | Tb |
| cDC1 | SPATS2 | 8.73E-11 | 0.794692197 | 0.697 | 0.469 | 1.72E-06 | Tb |
| cDC1 | LAMP3 | 4.39E-08 | 0.794216889 | 0.307 | 0.15 | 0.000863961 | Tb |
| cDC1 | ABI3 | 6.08E-15 | 0.792451176 | 0.622 | 0.323 | 1.20E-10 | Tb |
| cDC1 | IKZF1 | 1.73E-10 | 0.792267286 | 0.717 | 0.503 | 3.41E-06 | Tb |
| cDC1 | PTPN12 | 3.08E-11 | 0.791347103 | 0.625 | 0.37 | 6.06E-07 | Tb |
| cDC1 | MRPL42 | 1.55E-07 | 0.791051562 | 0.311 | 0.151 | 0.003055589 | Tb |
| cDC1 | SPECC1L | 3.88E-11 | 0.78923475 | 0.406 | 0.194 | 7.63E-07 | Tb |
| cDC1 | C1QB | 1.98E-11 | 0.788763846 | 0.47 | 0.247 | 3.90E-07 | Tb |
| cDC1 | CTDSP2 | 1.12E-08 | 0.788594964 | 0.367 | 0.18 | 0.000221256 | Tb |
| cDC1 | IDO2 | 4.55E-13 | 0.786935937 | 0.629 | 0.35 | 8.96E-09 | Tb |
| cDC1 | PIK3CG | 3.82E-10 | 0.78542426 | 0.554 | 0.312 | 7.51E-06 | Tb |
| cDC1 | ARHGEF7 | 8.33E-11 | 0.783545285 | 0.454 | 0.226 | 1.64E-06 | Tb |
| cDC1 | 1-Mar | 2.27E-14 | 0.783429872 | 0.801 | 0.517 | 4.47E-10 | Tb |
| cDC1 | AZIN1-AS1 | 5.76E-07 | 0.783199926 | 0.251 | 0.119 | 0.011343876 | Tb |
| cDC1 | ARRDC2 | 2.10E-10 | 0.780957532 | 0.307 | 0.126 | 4.13E-06 | Tb |
| cDC1 | GPR82 | 5.15E-09 | 0.78074028 | 0.255 | 0.103 | 0.000101428 | Tb |
| cDC1 | FAM49B | 3.52E-21 | 0.780445025 | 0.976 | 0.85 | 6.92E-17 | Tb |
| cDC1 | ATF7IP2 | 3.63E-10 | 0.779293149 | 0.542 | 0.318 | 7.14E-06 | Tb |

| | | | | | | | |
|---|---|---|---|---|---|---|---|
| cDC1 | POU2F1 | 2.17E-08 | 0.778359997 | 0.371 | 0.193 | 0.000426181 | Tb |
| cDC1 | MPHOSPH6 | 5.86E-08 | 0.776263161 | 0.279 | 0.127 | 0.00115426 | Tb |
| cDC1 | PAM | 1.86E-09 | 0.775752914 | 0.402 | 0.201 | 3.67E-05 | Tb |
| cDC1 | IGF2BP2 | 1.60E-06 | 0.775038981 | 0.307 | 0.166 | 0.031467551 | Tb |
| cDC1 | MGA | 6.13E-08 | 0.774442518 | 0.319 | 0.156 | 0.001206777 | Tb |
| cDC1 | FCHSD2 | 4.70E-17 | 0.774019813 | 0.888 | 0.65 | 9.25E-13 | Tb |
| cDC1 | RNGTT | 5.46E-08 | 0.772602936 | 0.422 | 0.235 | 0.001075176 | Tb |
| cDC1 | NUBPL | 1.09E-13 | 0.772001927 | 0.805 | 0.539 | 2.15E-09 | Tb |
| cDC1 | B4GALT4 | 4.11E-11 | 0.770955379 | 0.598 | 0.347 | 8.09E-07 | Tb |
| cDC1 | SUGP2 | 4.83E-07 | 0.770579749 | 0.378 | 0.205 | 0.009512289 | Tb |
| cDC1 | RASA1 | 7.04E-07 | 0.768362118 | 0.335 | 0.182 | 0.013859498 | Tb |
| cDC1 | TMX2 | 7.90E-10 | 0.768161411 | 0.339 | 0.148 | 1.56E-05 | Tb |
| cDC1 | KCNK6 | 3.88E-10 | 0.768127075 | 0.637 | 0.416 | 7.63E-06 | Tb |
| cDC1 | KDM6A | 1.15E-12 | 0.766302109 | 0.673 | 0.43 | 2.26E-08 | Tb |
| cDC1 | SQOR | 7.51E-08 | 0.7649388 | 0.39 | 0.206 | 0.001478225 | Tb |
| cDC1 | VRK2 | 6.36E-11 | 0.764837283 | 0.697 | 0.45 | 1.25E-06 | Tb |
| cDC1 | MTHFR | 4.40E-09 | 0.764720688 | 0.43 | 0.223 | 8.66E-05 | Tb |
| cDC1 | ANKIB1 | 4.83E-08 | 0.764683984 | 0.498 | 0.302 | 0.0009511 | Tb |
| cDC1 | CLASP1 | 3.92E-10 | 0.764617173 | 0.41 | 0.201 | 7.72E-06 | Tb |
| cDC1 | ATXN7 | 5.22E-08 | 0.764462607 | 0.406 | 0.223 | 0.001026969 | Tb |
| cDC1 | LYRM1 | 1.35E-06 | 0.760807292 | 0.307 | 0.157 | 0.026546793 | Tb |
| cDC1 | AK3 | 6.64E-08 | 0.760742451 | 0.339 | 0.17 | 0.00130685 | Tb |
| cDC1 | ST6GAL1 | 6.90E-11 | 0.759375429 | 0.526 | 0.299 | 1.36E-06 | Tb |
| cDC1 | TWSG1 | 4.21E-08 | 0.759230599 | 0.339 | 0.169 | 0.000828238 | Tb |
| cDC1 | KIAA0232 | 5.00E-11 | 0.757988407 | 0.486 | 0.253 | 9.84E-07 | Tb |
| cDC1 | EXOSC6 | 1.02E-06 | 0.757891428 | 0.351 | 0.189 | 0.020005714 | Tb |
| cDC1 | SLC20A2 | 2.81E-09 | 0.756279548 | 0.406 | 0.212 | 5.53E-05 | Tb |
| cDC1 | HDAC9 | 5.48E-21 | 0.755822495 | 0.992 | 0.948 | 1.08E-16 | Tb |
| cDC1 | FGD2 | 2.04E-12 | 0.755795033 | 0.653 | 0.398 | 4.02E-08 | Tb |
| cDC1 | USP20 | 1.14E-10 | 0.755203658 | 0.259 | 0.093 | 2.25E-06 | Tb |
| cDC1 | N4BP2L1 | 5.78E-08 | 0.753695595 | 0.502 | 0.289 | 0.001138667 | Tb |
| cDC1 | ATF5 | 3.01E-09 | 0.752503784 | 0.689 | 0.445 | 5.93E-05 | Tb |
| cDC1 | PCYT1A | 8.27E-10 | 0.75219048 | 0.506 | 0.291 | 1.63E-05 | Tb |
| cDC1 | CNOT10 | 8.02E-14 | 0.749428985 | 0.637 | 0.357 | 1.58E-09 | Tb |
| cDC1 | PRKX | 1.75E-09 | 0.74902453 | 0.502 | 0.288 | 3.44E-05 | Tb |
| cDC1 | RAB30 | 1.24E-11 | 0.748832954 | 0.637 | 0.389 | 2.44E-07 | Tb |
| cDC1 | EED | 1.16E-09 | 0.748700645 | 0.602 | 0.353 | 2.28E-05 | Tb |
| cDC1 | TET2 | 9.54E-09 | 0.748352436 | 0.924 | 0.806 | 0.000187721 | Tb |
| cDC1 | METTL16 | 1.28E-06 | 0.747971815 | 0.283 | 0.141 | 0.0252169 | Tb |
| cDC1 | MRTFA | 1.94E-14 | 0.747910513 | 0.773 | 0.5 | 3.81E-10 | Tb |
| cDC1 | NSMCE2 | 3.14E-12 | 0.747096087 | 0.701 | 0.44 | 6.18E-08 | Tb |
| cDC1 | IL15 | 4.61E-13 | 0.746021328 | 0.637 | 0.376 | 9.07E-09 | Tb |
| cDC1 | GLB1 | 1.13E-09 | 0.745867605 | 0.454 | 0.234 | 2.22E-05 | Tb |
| cDC1 | VCL | 1.56E-06 | 0.745857675 | 0.299 | 0.158 | 0.030671614 | Tb |
| cDC1 | ITGA4 | 5.00E-08 | 0.743696013 | 0.741 | 0.56 | 0.000985153 | Tb |
| cDC1 | ENTPD1-AS1 | 4.50E-09 | 0.743273797 | 0.458 | 0.249 | 8.87E-05 | Tb |
| cDC1 | STX5 | 1.22E-09 | 0.742076671 | 0.534 | 0.291 | 2.41E-05 | Tb |
| cDC1 | HMBOX1 | 4.79E-09 | 0.742065862 | 0.414 | 0.219 | 9.43E-05 | Tb |
| cDC1 | MYO9A | 3.35E-16 | 0.740354213 | 0.765 | 0.46 | 6.59E-12 | Tb |
| cDC1 | ABHD12 | 1.09E-08 | 0.736341602 | 0.351 | 0.169 | 0.000213763 | Tb |
| cDC1 | SEL1L3 | 2.05E-06 | 0.736177502 | 0.375 | 0.217 | 0.040335834 | Tb |
| cDC1 | CUX1 | 4.25E-15 | 0.735609711 | 0.853 | 0.651 | 8.36E-11 | Tb |
| cDC1 | GALNT11 | 3.13E-07 | 0.734844833 | 0.299 | 0.151 | 0.006158626 | Tb |
| cDC1 | CLNK | 1.04E-11 | 0.734697425 | 0.892 | 0.748 | 2.05E-07 | Tb |
| cDC1 | PDSS2 | 6.80E-09 | 0.734654995 | 0.343 | 0.167 | 0.00013386 | Tb |
| cDC1 | SOS1 | 7.11E-11 | 0.733952115 | 0.625 | 0.386 | 1.40E-06 | Tb |

| | | | | | | | |
|---|---|---|---|---|---|---|---|
| cDC1 | TEX2 | 3.66E-09 | 0.733765474 | 0.363 | 0.175 | 7.21E-05 | Tb |
| cDC1 | NLRP1 | 2.43E-06 | 0.733744151 | 0.371 | 0.218 | 0.047736077 | Tb |
| cDC1 | NCOA3 | 5.37E-11 | 0.733042325 | 0.705 | 0.473 | 1.06E-06 | Tb |
| cDC1 | TSC22D2 | 4.54E-07 | 0.731811067 | 0.653 | 0.452 | 0.008946181 | Tb |
| cDC1 | ARMH3 | 3.40E-12 | 0.730327669 | 0.49 | 0.244 | 6.68E-08 | Tb |
| cDC1 | ARHGAP27 | 2.39E-06 | 0.729828459 | 0.331 | 0.177 | 0.047074907 | Tb |
| cDC1 | ZCCHC2 | 9.16E-10 | 0.729644228 | 0.574 | 0.35 | 1.80E-05 | Tb |
| cDC1 | UST | 2.00E-10 | 0.727662445 | 0.594 | 0.388 | 3.93E-06 | Tb |
| cDC1 | MED27 | 7.58E-09 | 0.726848187 | 0.363 | 0.175 | 0.000149204 | Tb |
| cDC1 | CARD11 | 4.99E-07 | 0.72660922 | 0.474 | 0.29 | 0.009815681 | Tb |
| cDC1 | PPP2R5E | 1.41E-07 | 0.726370602 | 0.522 | 0.318 | 0.002776847 | Tb |
| cDC1 | TRIM13 | 3.66E-10 | 0.726080673 | 0.43 | 0.212 | 7.21E-06 | Tb |
| cDC1 | SPPL3 | 5.53E-16 | 0.725342744 | 0.841 | 0.576 | 1.09E-11 | Tb |
| cDC1 | KDM5C | 5.47E-07 | 0.724369099 | 0.279 | 0.135 | 0.010768893 | Tb |
| cDC1 | GTF2H1 | 1.18E-09 | 0.723508114 | 0.598 | 0.361 | 2.32E-05 | Tb |
| cDC1 | CIITA | 1.32E-16 | 0.723485961 | 0.861 | 0.6 | 2.61E-12 | Tb |
| cDC1 | RPP38 | 3.75E-07 | 0.723147504 | 0.255 | 0.115 | 0.007385932 | Tb |
| cDC1 | PPP6R2 | 5.44E-11 | 0.723145507 | 0.602 | 0.345 | 1.07E-06 | Tb |
| cDC1 | FTO | 3.57E-10 | 0.723098473 | 0.283 | 0.114 | 7.02E-06 | Tb |
| cDC1 | ABL2 | 4.12E-07 | 0.72301787 | 0.45 | 0.266 | 0.008112491 | Tb |
| cDC1 | CDH2 | 1.25E-07 | 0.722961299 | 0.331 | 0.17 | 0.002469099 | Tb |
| cDC1 | USP32 | 4.84E-16 | 0.722378834 | 0.737 | 0.47 | 9.52E-12 | Tb |
| cDC1 | KLF2 | 7.63E-10 | 0.721796417 | 0.287 | 0.118 | 1.50E-05 | Tb |
| cDC1 | PACS1 | 4.03E-13 | 0.721323757 | 0.757 | 0.51 | 7.93E-09 | Tb |
| cDC1 | YEATS2 | 5.66E-11 | 0.720634926 | 0.482 | 0.252 | 1.11E-06 | Tb |
| cDC1 | FAM133B | 7.55E-13 | 0.720415294 | 0.649 | 0.377 | 1.49E-08 | Tb |
| cDC1 | ARIH1 | 3.19E-15 | 0.720319976 | 0.884 | 0.673 | 6.28E-11 | Tb |
| cDC1 | EMC2 | 1.81E-07 | 0.719952796 | 0.382 | 0.201 | 0.003556442 | Tb |
| cDC1 | RIPK1 | 1.50E-14 | 0.718673546 | 0.661 | 0.362 | 2.96E-10 | Tb |
| cDC1 | PCNX4 | 1.13E-06 | 0.717917004 | 0.367 | 0.211 | 0.022307397 | Tb |
| cDC1 | ATP6V1A | 2.09E-07 | 0.716772526 | 0.633 | 0.475 | 0.004108718 | Tb |
| cDC1 | WIPF2 | 6.64E-09 | 0.716672938 | 0.311 | 0.148 | 0.000130651 | Tb |
| cDC1 | NBEAL1 | 5.60E-08 | 0.716567617 | 0.677 | 0.46 | 0.001102521 | Tb |
| cDC1 | TRAPPC9 | 5.72E-12 | 0.714739135 | 0.57 | 0.322 | 1.13E-07 | Tb |
| cDC1 | ATR | 1.43E-13 | 0.714465873 | 0.59 | 0.311 | 2.81E-09 | Tb |
| cDC1 | TIAM1 | 9.96E-07 | 0.714018878 | 0.327 | 0.17 | 0.01960033 | Tb |
| cDC1 | NAV1 | 2.89E-13 | 0.713803452 | 0.773 | 0.481 | 5.69E-09 | Tb |
| cDC1 | FBXL17 | 6.54E-08 | 0.712332408 | 0.506 | 0.306 | 0.001287501 | Tb |
| cDC1 | SERPINF1 | 5.83E-09 | 0.711888143 | 0.845 | 0.681 | 0.000114672 | Tb |
| cDC1 | SOGA1 | 3.10E-07 | 0.711405224 | 0.363 | 0.2 | 0.006107552 | Tb |
| cDC1 | KIAA2026 | 1.94E-10 | 0.710245604 | 0.49 | 0.256 | 3.82E-06 | Tb |
| cDC1 | FLNB | 3.43E-08 | 0.709385511 | 0.478 | 0.284 | 0.000675931 | Tb |
| cDC1 | ZSWIM6 | 2.76E-18 | 0.709207401 | 0.936 | 0.716 | 5.44E-14 | Tb |
| cDC1 | SRGAP2B | 3.67E-12 | 0.708753886 | 0.418 | 0.195 | 7.22E-08 | Tb |
| cDC1 | TPT1-AS1 | 1.28E-08 | 0.708442301 | 0.251 | 0.103 | 0.000252918 | Tb |
| cDC1 | TBC1D12 | 3.71E-07 | 0.707745392 | 0.39 | 0.219 | 0.007294129 | Tb |
| cDC1 | PTEN | 3.99E-11 | 0.707536003 | 0.729 | 0.491 | 7.86E-07 | Tb |
| cDC1 | THADA | 1.25E-07 | 0.706703494 | 0.315 | 0.153 | 0.002464059 | Tb |
| cDC1 | GSAP | 1.53E-10 | 0.706067468 | 0.502 | 0.272 | 3.02E-06 | Tb |
| cDC1 | CCPG1 | 3.90E-14 | 0.704579133 | 0.693 | 0.407 | 7.67E-10 | Tb |
| cDC1 | FARP2 | 4.01E-12 | 0.704285771 | 0.785 | 0.531 | 7.90E-08 | Tb |
| cDC1 | GSN | 3.52E-11 | 0.703575528 | 0.833 | 0.664 | 6.92E-07 | Tb |
| cDC1 | CD84 | 5.18E-09 | 0.703024145 | 0.51 | 0.285 | 0.00010199 | Tb |
| cDC1 | VEZT | 2.89E-09 | 0.702008012 | 0.514 | 0.282 | 5.68E-05 | Tb |
| cDC1 | DCAF6 | 2.56E-08 | 0.701695527 | 0.351 | 0.174 | 0.00050459 | Tb |
| cDC1 | SNX29 | 2.51E-15 | 0.701573591 | 0.769 | 0.505 | 4.95E-11 | Tb |

| | | | | | | | |
|---|---|---|---|---|---|---|---|
| cDC1 | HSH2D | 3.76E-11 | 0.700101059 | 0.554 | 0.293 | 7.41E-07 | Tb |
| cDC1 | TPK1 | 2.61E-10 | 0.69870378 | 0.53 | 0.295 | 5.13E-06 | Tb |
| cDC1 | FKBP5 | 1.77E-19 | 0.698335949 | 0.817 | 0.506 | 3.48E-15 | Tb |
| cDC1 | MTIF2 | 2.10E-08 | 0.696886282 | 0.315 | 0.149 | 0.00041315 | Tb |
| cDC1 | GPR132 | 9.80E-13 | 0.696112056 | 0.61 | 0.355 | 1.93E-08 | Tb |
| cDC1 | DHRSX | 4.48E-11 | 0.694462006 | 0.506 | 0.26 | 8.81E-07 | Tb |
| cDC1 | DLG1 | 1.50E-06 | 0.693283822 | 0.582 | 0.376 | 0.029460395 | Tb |
| cDC1 | EZH2 | 1.43E-15 | 0.692513402 | 0.789 | 0.536 | 2.82E-11 | Tb |
| cDC1 | SMURF2 | 6.75E-09 | 0.691365668 | 0.53 | 0.312 | 0.000132802 | Tb |
| cDC1 | KANSL1 | 6.78E-14 | 0.689595754 | 0.594 | 0.3 | 1.33E-09 | Tb |
| cDC1 | PDE7A | 7.32E-09 | 0.688271131 | 0.657 | 0.447 | 0.00014413 | Tb |
| cDC1 | FAM172A | 8.80E-10 | 0.687900403 | 0.518 | 0.301 | 1.73E-05 | Tb |
| cDC1 | SPPL2A | 1.16E-12 | 0.687086338 | 0.637 | 0.37 | 2.29E-08 | Tb |
| cDC1 | CFLAR-AS1 | 2.14E-09 | 0.687058618 | 0.367 | 0.176 | 4.20E-05 | Tb |
| cDC1 | ZDHHC4 | 1.13E-06 | 0.685908614 | 0.263 | 0.124 | 0.022211306 | Tb |
| cDC1 | CTDP1 | 1.85E-11 | 0.685032032 | 0.398 | 0.185 | 3.64E-07 | Tb |
| cDC1 | CD226 | 4.78E-11 | 0.684671756 | 0.757 | 0.494 | 9.40E-07 | Tb |
| cDC1 | CHSY1 | 6.55E-08 | 0.684650319 | 0.291 | 0.134 | 0.001289076 | Tb |
| cDC1 | KPNA6 | 1.85E-08 | 0.684275239 | 0.335 | 0.157 | 0.000364889 | Tb |
| cDC1 | TBC1D8 | 6.28E-10 | 0.684062344 | 0.546 | 0.324 | 1.24E-05 | Tb |
| cDC1 | DOCK8 | 9.14E-13 | 0.683748252 | 0.817 | 0.597 | 1.80E-08 | Tb |
| cDC1 | P2RY6 | 1.04E-11 | 0.683215838 | 0.622 | 0.359 | 2.04E-07 | Tb |
| cDC1 | RBMS1 | 7.35E-07 | 0.681923954 | 0.398 | 0.226 | 0.014470741 | Tb |
| cDC1 | DIS3L2 | 3.45E-07 | 0.681866153 | 0.283 | 0.139 | 0.006781884 | Tb |
| cDC1 | BTN3A1 | 2.25E-08 | 0.681728946 | 0.275 | 0.119 | 0.000442185 | Tb |
| cDC1 | MBNL1 | 5.88E-10 | 0.68036673 | 0.912 | 0.775 | 1.16E-05 | Tb |
| cDC1 | PTPRJ | 1.01E-17 | 0.680114277 | 0.896 | 0.613 | 1.99E-13 | Tb |
| cDC1 | MYO1C | 2.56E-08 | 0.679562115 | 0.371 | 0.198 | 0.000504854 | Tb |
| cDC1 | USP42 | 9.93E-07 | 0.678995411 | 0.307 | 0.157 | 0.019555926 | Tb |
| cDC1 | SCYL2 | 1.02E-07 | 0.678610329 | 0.458 | 0.269 | 0.002000907 | Tb |
| cDC1 | CDC42SE2 | 9.00E-12 | 0.678386694 | 0.876 | 0.673 | 1.77E-07 | Tb |
| cDC1 | GRK3 | 1.82E-09 | 0.67837261 | 0.677 | 0.449 | 3.59E-05 | Tb |
| cDC1 | GPR107 | 1.96E-06 | 0.678162712 | 0.39 | 0.22 | 0.038546184 | Tb |
| cDC1 | NUP42 | 3.54E-11 | 0.678086376 | 0.53 | 0.264 | 6.97E-07 | Tb |
| cDC1 | DENND4A | 7.59E-20 | 0.676588034 | 0.853 | 0.561 | 1.49E-15 | Tb |
| cDC1 | CUL2 | 1.54E-06 | 0.675757761 | 0.327 | 0.174 | 0.030295715 | Tb |
| cDC1 | GRAMD4 | 5.30E-07 | 0.675675167 | 0.386 | 0.21 | 0.010429396 | Tb |
| cDC1 | CCNG2 | 7.29E-10 | 0.674942328 | 0.633 | 0.386 | 1.44E-05 | Tb |
| cDC1 | CERT1 | 8.96E-08 | 0.674926714 | 0.625 | 0.441 | 0.001764589 | Tb |
| cDC1 | NFE2L3 | 3.13E-10 | 0.674032007 | 0.721 | 0.486 | 6.16E-06 | Tb |
| cDC1 | RICTOR | 4.63E-09 | 0.672931018 | 0.57 | 0.332 | 9.11E-05 | Tb |
| cDC1 | MAML2 | 1.02E-11 | 0.671541128 | 0.829 | 0.588 | 2.01E-07 | Tb |
| cDC1 | UBL3 | 5.73E-07 | 0.671166977 | 0.689 | 0.497 | 0.011287989 | Tb |
| cDC1 | XPO1 | 3.53E-10 | 0.670623348 | 0.733 | 0.48 | 6.95E-06 | Tb |
| cDC1 | PHLDB3 | 1.12E-08 | 0.670489591 | 0.339 | 0.165 | 0.000219842 | Tb |
| cDC1 | FAM135A | 1.05E-09 | 0.670122562 | 0.737 | 0.491 | 2.07E-05 | Tb |
| cDC1 | GCC2 | 1.04E-11 | 0.669168988 | 0.721 | 0.459 | 2.04E-07 | Tb |
| cDC1 | TSPOAP1 | 9.43E-07 | 0.668953528 | 0.363 | 0.207 | 0.018552567 | Tb |
| cDC1 | SPG11 | 9.46E-07 | 0.668321411 | 0.454 | 0.281 | 0.018620937 | Tb |
| cDC1 | ENPP4 | 2.28E-07 | 0.668151567 | 0.263 | 0.119 | 0.004485422 | Tb |
| cDC1 | MED23 | 2.21E-08 | 0.664570469 | 0.287 | 0.13 | 0.000435435 | Tb |
| cDC1 | HNRNPLL | 1.29E-07 | 0.663685493 | 0.681 | 0.458 | 0.002529417 | Tb |
| cDC1 | STAG1 | 1.96E-09 | 0.663314741 | 0.606 | 0.391 | 3.86E-05 | Tb |
| cDC1 | MFN1 | 2.34E-07 | 0.662889228 | 0.486 | 0.289 | 0.004612739 | Tb |
| cDC1 | PICALM | 3.20E-17 | 0.662137286 | 0.956 | 0.817 | 6.30E-13 | Tb |
| cDC1 | CAMK2D | 1.40E-11 | 0.661470402 | 0.952 | 0.864 | 2.76E-07 | Tb |

| | | | | | | | |
|---|---|---|---|---|---|---|---|
| cDC1 | HMGCL | 6.48E-07 | 0.661217338 | 0.259 | 0.119 | 0.012762095 | Tb |
| cDC1 | CST7 | 1.28E-09 | 0.660827988 | 0.717 | 0.488 | 2.51E-05 | Tb |
| cDC1 | ADAM17 | 1.87E-09 | 0.660668948 | 0.57 | 0.34 | 3.69E-05 | Tb |
| cDC1 | UTP3 | 5.25E-08 | 0.659845149 | 0.359 | 0.178 | 0.001033903 | Tb |
| cDC1 | MYOF | 3.50E-10 | 0.659183345 | 0.518 | 0.285 | 6.89E-06 | Tb |
| cDC1 | ACTR6 | 1.28E-07 | 0.659034793 | 0.351 | 0.177 | 0.002524603 | Tb |
| cDC1 | SNX27 | 3.27E-08 | 0.658936638 | 0.378 | 0.191 | 0.000643602 | Tb |
| cDC1 | ZDHHC17 | 1.77E-08 | 0.658598227 | 0.578 | 0.356 | 0.000349368 | Tb |
| cDC1 | TAB3 | 1.87E-08 | 0.657984514 | 0.291 | 0.128 | 0.000367143 | Tb |
| cDC1 | ZNF462 | 1.90E-07 | 0.656345749 | 0.394 | 0.218 | 0.003749761 | Tb |
| cDC1 | GPR89A | 3.05E-07 | 0.655950871 | 0.327 | 0.165 | 0.006010113 | Tb |
| cDC1 | SH3BP2 | 2.25E-11 | 0.655359736 | 0.386 | 0.168 | 4.43E-07 | Tb |
| cDC1 | TOR2A | 2.03E-06 | 0.655279255 | 0.315 | 0.162 | 0.039884331 | Tb |
| cDC1 | ARNTL | 1.94E-08 | 0.654817254 | 0.375 | 0.19 | 0.000382134 | Tb |
| cDC1 | DGKH | 3.92E-08 | 0.654166424 | 0.291 | 0.139 | 0.000772464 | Tb |
| cDC1 | TLK1 | 1.93E-10 | 0.654011242 | 0.534 | 0.29 | 3.79E-06 | Tb |
| cDC1 | LPCAT3 | 1.21E-12 | 0.653800043 | 0.359 | 0.143 | 2.39E-08 | Tb |
| cDC1 | DMXL1 | 1.20E-09 | 0.652930428 | 0.462 | 0.249 | 2.35E-05 | Tb |
| cDC1 | CCL3L1 | 7.47E-07 | 0.652471608 | 0.359 | 0.199 | 0.014709027 | Tb |
| cDC1 | TTC9C | 4.91E-10 | 0.652145558 | 0.359 | 0.16 | 9.67E-06 | Tb |
| cDC1 | RIC1 | 9.04E-12 | 0.652071935 | 0.649 | 0.384 | 1.78E-07 | Tb |
| cDC1 | SNX8 | 2.95E-20 | 0.652018246 | 0.884 | 0.615 | 5.80E-16 | Tb |
| cDC1 | ASAP2 | 1.42E-09 | 0.652000155 | 0.434 | 0.234 | 2.80E-05 | Tb |
| cDC1 | SNX20 | 9.60E-10 | 0.650287957 | 0.574 | 0.331 | 1.89E-05 | Tb |
| cDC1 | NCOA2 | 1.58E-10 | 0.650109571 | 0.582 | 0.336 | 3.11E-06 | Tb |
| cDC1 | NIPA2 | 6.56E-08 | 0.649014773 | 0.454 | 0.252 | 0.001291154 | Tb |
| cDC1 | SENP7 | 9.90E-09 | 0.648978212 | 0.283 | 0.12 | 0.000194783 | Tb |
| cDC1 | PNPLA6 | 1.94E-08 | 0.648282559 | 0.331 | 0.152 | 0.000381619 | Tb |
| cDC1 | RNF111 | 1.15E-06 | 0.647704363 | 0.422 | 0.258 | 0.022583634 | Tb |
| cDC1 | LIMS1 | 1.05E-09 | 0.647654018 | 0.793 | 0.577 | 2.07E-05 | Tb |
| cDC1 | TOX | 1.22E-08 | 0.646965433 | 0.773 | 0.565 | 0.000240112 | Tb |
| cDC1 | CEP83 | 2.04E-06 | 0.646902405 | 0.406 | 0.249 | 0.040235277 | Tb |
| cDC1 | RBM47 | 4.06E-08 | 0.646303255 | 0.661 | 0.481 | 0.000798944 | Tb |
| cDC1 | HIST1H2AC | 1.84E-10 | 0.645752317 | 0.434 | 0.217 | 3.63E-06 | Tb |
| cDC1 | ATF7IP | 1.14E-13 | 0.643882705 | 0.685 | 0.427 | 2.25E-09 | Tb |
| cDC1 | MLXIP | 2.49E-07 | 0.643789545 | 0.402 | 0.223 | 0.004901145 | Tb |
| cDC1 | PLCG2 | 4.80E-10 | 0.64373047 | 0.653 | 0.419 | 9.45E-06 | Tb |
| cDC1 | UBP1 | 5.21E-07 | 0.64350179 | 0.331 | 0.173 | 0.010259177 | Tb |
| cDC1 | VASH1 | 9.10E-14 | 0.643424985 | 0.602 | 0.314 | 1.79E-09 | Tb |
| cDC1 | SH2D3C | 4.04E-08 | 0.642839008 | 0.359 | 0.182 | 0.00079506 | Tb |
| cDC1 | POLR2J3 | 1.78E-08 | 0.642800234 | 0.291 | 0.13 | 0.000350073 | Tb |
| cDC1 | GATAD2B | 5.26E-08 | 0.642637131 | 0.51 | 0.312 | 0.001035938 | Tb |
| cDC1 | TMF1 | 2.01E-11 | 0.641422125 | 0.61 | 0.352 | 3.96E-07 | Tb |
| cDC1 | RNF38 | 1.97E-08 | 0.641083059 | 0.494 | 0.286 | 0.000387091 | Tb |
| cDC1 | PHLPP2 | 3.67E-08 | 0.640927851 | 0.347 | 0.174 | 0.000723202 | Tb |
| cDC1 | ST7 | 1.41E-07 | 0.640809781 | 0.39 | 0.208 | 0.002779224 | Tb |
| cDC1 | ARHGAP32 | 4.85E-08 | 0.64061866 | 0.339 | 0.168 | 0.0009542 | Tb |
| cDC1 | ANKRD13D | 6.52E-07 | 0.640281649 | 0.371 | 0.205 | 0.012838528 | Tb |
| cDC1 | STOM | 3.05E-12 | 0.639952048 | 0.685 | 0.42 | 6.01E-08 | Tb |
| cDC1 | AP3M2 | 2.10E-08 | 0.639570504 | 0.418 | 0.219 | 0.000414232 | Tb |
| cDC1 | ABHD2 | 2.03E-07 | 0.639412913 | 0.598 | 0.383 | 0.003993065 | Tb |
| cDC1 | DENND1B | 3.39E-16 | 0.637340101 | 0.952 | 0.798 | 6.68E-12 | Tb |
| cDC1 | COMMD10 | 1.46E-08 | 0.637155071 | 0.633 | 0.399 | 0.000287755 | Tb |
| cDC1 | ZEB1 | 8.75E-13 | 0.636939155 | 0.837 | 0.666 | 1.72E-08 | Tb |
| cDC1 | OSBPL3 | 3.82E-07 | 0.636740155 | 0.637 | 0.464 | 0.007511293 | Tb |
| cDC1 | PHF8 | 1.24E-06 | 0.63618571 | 0.307 | 0.161 | 0.024437806 | Tb |

| | | | | | | | |
|---|---|---|---|---|---|---|---|
| cDC1 | PRKRIP1 | 2.37E-08 | 0.636045441 | 0.474 | 0.266 | 0.000467136 | Tb |
| cDC1 | TTC14 | 3.55E-07 | 0.635875511 | 0.267 | 0.12 | 0.006989874 | Tb |
| cDC1 | APOL2 | 2.23E-11 | 0.635863683 | 0.43 | 0.2 | 4.40E-07 | Tb |
| cDC1 | RAB11FIP4 | 2.21E-06 | 0.635717554 | 0.386 | 0.222 | 0.043412753 | Tb |
| cDC1 | MT-ND4 | 3.47E-19 | 0.634434326 | 1 | 0.944 | 6.84E-15 | Tb |
| cDC1 | DOCK10 | 3.99E-11 | 0.633904498 | 0.896 | 0.699 | 7.85E-07 | Tb |
| cDC1 | NT5C2 | 9.33E-08 | 0.633064414 | 0.454 | 0.258 | 0.001837071 | Tb |
| cDC1 | SLC25A33 | 4.24E-07 | 0.633046284 | 0.777 | 0.619 | 0.008346147 | Tb |
| cDC1 | PRKAG1 | 3.63E-10 | 0.63289314 | 0.434 | 0.211 | 7.14E-06 | Tb |
| cDC1 | CLASP2 | 3.73E-08 | 0.632466226 | 0.693 | 0.476 | 0.000734581 | Tb |
| cDC1 | CASS4 | 3.03E-07 | 0.631553547 | 0.522 | 0.315 | 0.005956795 | Tb |
| cDC1 | CLEC1A | 3.27E-08 | 0.630966603 | 0.434 | 0.24 | 0.000644282 | Tb |
| cDC1 | PHTF2 | 6.38E-09 | 0.630707047 | 0.558 | 0.333 | 0.000125637 | Tb |
| cDC1 | ARID1A | 1.81E-08 | 0.629720206 | 0.578 | 0.358 | 0.00035545 | Tb |
| cDC1 | SEC24B | 1.93E-06 | 0.62944144 | 0.462 | 0.282 | 0.038074269 | Tb |
| cDC1 | PRKAR2A | 1.13E-08 | 0.62856654 | 0.502 | 0.286 | 0.000223374 | Tb |
| cDC1 | PIK3R5 | 1.24E-12 | 0.628298786 | 0.665 | 0.415 | 2.45E-08 | Tb |
| cDC1 | ZHX2 | 1.06E-08 | 0.62706122 | 0.49 | 0.267 | 0.000209191 | Tb |
| cDC1 | SKI | 4.60E-07 | 0.62689131 | 0.367 | 0.198 | 0.009045234 | Tb |
| cDC1 | CD2AP | 1.13E-09 | 0.626644079 | 0.49 | 0.26 | 2.23E-05 | Tb |
| cDC1 | ZCCHC17 | 2.85E-07 | 0.626561753 | 0.371 | 0.197 | 0.005618777 | Tb |
| cDC1 | ARHGAP15 | 1.00E-08 | 0.626196551 | 0.669 | 0.453 | 0.000196799 | Tb |
| cDC1 | ALDH2 | 8.03E-13 | 0.626158069 | 0.857 | 0.659 | 1.58E-08 | Tb |
| cDC1 | HERC3 | 3.21E-08 | 0.625694392 | 0.363 | 0.185 | 0.000630899 | Tb |
| cDC1 | CCSER2 | 3.86E-09 | 0.624196463 | 0.598 | 0.361 | 7.59E-05 | Tb |
| cDC1 | CADM1 | 3.86E-12 | 0.623732576 | 0.936 | 0.814 | 7.60E-08 | Tb |
| cDC1 | ELMSAN1 | 1.66E-10 | 0.622959273 | 0.645 | 0.399 | 3.27E-06 | Tb |
| cDC1 | UBE2O | 2.24E-06 | 0.62214373 | 0.251 | 0.117 | 0.044037361 | Tb |
| cDC1 | CCM2 | 9.73E-08 | 0.622135936 | 0.406 | 0.212 | 0.001915174 | Tb |
| cDC1 | ST3GAL5 | 3.73E-12 | 0.62200949 | 0.693 | 0.416 | 7.35E-08 | Tb |
| cDC1 | VAC14 | 2.37E-16 | 0.621541046 | 0.944 | 0.736 | 4.67E-12 | Tb |
| cDC1 | FOXP1 | 8.44E-12 | 0.619544204 | 0.873 | 0.647 | 1.66E-07 | Tb |
| cDC1 | GCLC | 4.22E-10 | 0.619139609 | 0.47 | 0.242 | 8.31E-06 | Tb |
| cDC1 | PPP2R5A | 1.67E-09 | 0.618647629 | 0.378 | 0.181 | 3.30E-05 | Tb |
| cDC1 | UBAC2 | 1.14E-06 | 0.618443507 | 0.518 | 0.339 | 0.022355832 | Tb |
| cDC1 | TMTC2 | 2.47E-08 | 0.616028697 | 0.351 | 0.182 | 0.000485562 | Tb |
| cDC1 | GLG1 | 1.13E-08 | 0.615600331 | 0.566 | 0.341 | 0.000223348 | Tb |
| cDC1 | SDHC | 6.46E-09 | 0.615433051 | 0.518 | 0.291 | 0.000127161 | Tb |
| cDC1 | RBM6 | 7.35E-08 | 0.614992901 | 0.586 | 0.382 | 0.001446349 | Tb |
| cDC1 | EXOC4 | 8.99E-11 | 0.614171248 | 0.653 | 0.405 | 1.77E-06 | Tb |
| cDC1 | TP53 | 2.08E-06 | 0.614113882 | 0.359 | 0.195 | 0.040858492 | Tb |
| cDC1 | AGO3 | 2.44E-09 | 0.613463685 | 0.586 | 0.353 | 4.81E-05 | Tb |
| cDC1 | PDLIM1 | 7.64E-07 | 0.612840782 | 0.386 | 0.216 | 0.015047732 | Tb |
| cDC1 | AP3B1 | 1.55E-14 | 0.612653562 | 0.729 | 0.427 | 3.05E-10 | Tb |
| cDC1 | MPHOSPH9 | 1.25E-07 | 0.612282135 | 0.315 | 0.152 | 0.002451789 | Tb |
| cDC1 | NPLOC4 | 5.92E-07 | 0.612010092 | 0.462 | 0.274 | 0.011643192 | Tb |
| cDC1 | OFD1 | 2.58E-13 | 0.610763586 | 0.693 | 0.411 | 5.07E-09 | Tb |
| cDC1 | SUSD6 | 3.43E-09 | 0.610643044 | 0.717 | 0.49 | 6.76E-05 | Tb |
| cDC1 | HMGB2 | 2.44E-07 | 0.609443372 | 0.578 | 0.369 | 0.004795057 | Tb |
| cDC1 | OXCT1 | 1.52E-06 | 0.609327822 | 0.343 | 0.188 | 0.029847326 | Tb |
| cDC1 | RB1 | 6.18E-11 | 0.608241924 | 0.701 | 0.465 | 1.22E-06 | Tb |
| cDC1 | PRKAR2B | 2.05E-09 | 0.607243589 | 0.805 | 0.566 | 4.03E-05 | Tb |
| cDC1 | ZCCHC7 | 2.41E-10 | 0.60696624 | 0.661 | 0.42 | 4.74E-06 | Tb |
| cDC1 | N4BP2L2 | 3.73E-09 | 0.60611207 | 0.57 | 0.334 | 7.34E-05 | Tb |
| cDC1 | POGZ | 1.09E-08 | 0.605654009 | 0.482 | 0.274 | 0.000215095 | Tb |
| cDC1 | DENND4B | 1.28E-08 | 0.605554863 | 0.382 | 0.189 | 0.000252473 | Tb |

| | | | | | | | |
|---|---|---|---|---|---|---|---|
| cDC1 | HS3ST1 | 1.60E-08 | 0.604671112 | 0.327 | 0.151 | 0.000315554 | Tb |
| cDC1 | ZUP1 | 1.62E-09 | 0.602378073 | 0.414 | 0.211 | 3.19E-05 | Tb |
| cDC1 | NEDD9 | 5.52E-08 | 0.60167214 | 0.689 | 0.477 | 0.001086191 | Tb |
| cDC1 | CNTLN | 3.42E-07 | 0.601039877 | 0.45 | 0.261 | 0.006731147 | Tb |
| cDC1 | EAPP | 1.54E-07 | 0.601013837 | 0.426 | 0.236 | 0.003030846 | Tb |
| cDC1 | PTPN22 | 1.93E-07 | 0.600991474 | 0.701 | 0.485 | 0.003795313 | Tb |
| cDC1 | SLCO5A1 | 1.89E-09 | 0.600241391 | 0.43 | 0.223 | 3.72E-05 | Tb |
| cDC1 | SCLT1 | 3.91E-11 | 0.599730632 | 0.753 | 0.501 | 7.69E-07 | Tb |
| cDC1 | BRCA2 | 1.09E-07 | 0.599616149 | 0.382 | 0.2 | 0.002148755 | Tb |
| cDC1 | EML4 | 5.94E-10 | 0.599482179 | 0.833 | 0.61 | 1.17E-05 | Tb |
| cDC1 | TIGAR | 2.02E-08 | 0.599298596 | 0.562 | 0.345 | 0.000396918 | Tb |
| cDC1 | UBE2E1 | 2.80E-07 | 0.599042379 | 0.637 | 0.434 | 0.005507613 | Tb |
| cDC1 | IRF2 | 7.54E-08 | 0.598099174 | 0.598 | 0.368 | 0.001484945 | Tb |
| cDC1 | HIVEP3 | 1.00E-10 | 0.597548034 | 0.542 | 0.303 | 1.97E-06 | Tb |
| cDC1 | ZNF407 | 8.98E-08 | 0.597258112 | 0.49 | 0.289 | 0.001766901 | Tb |
| cDC1 | FAM53C | 2.09E-08 | 0.596751678 | 0.498 | 0.294 | 0.000410467 | Tb |
| cDC1 | TLK2 | 1.11E-08 | 0.59557936 | 0.474 | 0.258 | 0.000219066 | Tb |
| cDC1 | RFC1 | 7.04E-09 | 0.594602477 | 0.486 | 0.27 | 0.00013848 | Tb |
| cDC1 | UIMC1 | 9.43E-10 | 0.593453702 | 0.41 | 0.202 | 1.86E-05 | Tb |
| cDC1 | THOC6 | 1.50E-07 | 0.592702248 | 0.394 | 0.207 | 0.002961666 | Tb |
| cDC1 | VMP1 | 1.02E-10 | 0.591909236 | 0.92 | 0.698 | 2.00E-06 | Tb |
| cDC1 | FUBP3 | 2.97E-07 | 0.591876372 | 0.275 | 0.128 | 0.005847106 | Tb |
| cDC1 | MT-CO2 | 6.65E-20 | 0.591212082 | 0.996 | 0.992 | 1.31E-15 | Tb |
| cDC1 | APOL3 | 1.41E-09 | 0.591184725 | 0.562 | 0.324 | 2.78E-05 | Tb |
| cDC1 | PBX3 | 3.77E-07 | 0.59006949 | 0.57 | 0.372 | 0.00742816 | Tb |
| cDC1 | FAM120B | 2.02E-08 | 0.588895823 | 0.279 | 0.122 | 0.00039737 | Tb |
| cDC1 | BBX | 2.49E-09 | 0.587646014 | 0.673 | 0.434 | 4.90E-05 | Tb |
| cDC1 | TVP23C | 2.39E-09 | 0.587632714 | 0.406 | 0.205 | 4.71E-05 | Tb |
| cDC1 | SCOC | 5.81E-07 | 0.587350529 | 0.359 | 0.183 | 0.011433138 | Tb |
| cDC1 | SNX5 | 1.13E-08 | 0.58714685 | 0.57 | 0.342 | 0.000221742 | Tb |
| cDC1 | JAK2 | 8.25E-12 | 0.586628983 | 0.853 | 0.591 | 1.62E-07 | Tb |
| cDC1 | MAP4 | 5.15E-12 | 0.585429115 | 0.789 | 0.532 | 1.01E-07 | Tb |
| cDC1 | HAPLN3 | 8.76E-07 | 0.585359105 | 0.43 | 0.245 | 0.017242223 | Tb |
| cDC1 | ATXN1 | 4.53E-09 | 0.585339607 | 0.785 | 0.568 | 8.91E-05 | Tb |
| cDC1 | IQCB1 | 1.65E-07 | 0.585291301 | 0.39 | 0.211 | 0.003240242 | Tb |
| cDC1 | MBOAT7 | 1.24E-08 | 0.58433632 | 0.375 | 0.183 | 0.000244812 | Tb |
| cDC1 | CASK | 1.60E-07 | 0.584021102 | 0.506 | 0.3 | 0.003154372 | Tb |
| cDC1 | PDXDC1 | 1.91E-08 | 0.584008154 | 0.414 | 0.216 | 0.00037557 | Tb |
| cDC1 | IL17RA | 2.34E-07 | 0.583426609 | 0.315 | 0.159 | 0.004599407 | Tb |
| cDC1 | CAMK1D | 2.79E-11 | 0.582535917 | 0.825 | 0.559 | 5.50E-07 | Tb |
| cDC1 | GSK3B | 1.08E-07 | 0.582374795 | 0.594 | 0.397 | 0.002135035 | Tb |
| cDC1 | SMURF1 | 7.83E-10 | 0.581423387 | 0.422 | 0.217 | 1.54E-05 | Tb |
| cDC1 | SLC2A13 | 7.96E-07 | 0.581295201 | 0.311 | 0.162 | 0.015666857 | Tb |
| cDC1 | PHKB | 1.37E-07 | 0.580801591 | 0.434 | 0.24 | 0.002693213 | Tb |
| cDC1 | SEMA4A | 9.08E-08 | 0.580451459 | 0.637 | 0.433 | 0.001787204 | Tb |
| cDC1 | CD4 | 1.19E-08 | 0.58021961 | 0.586 | 0.358 | 0.000234254 | Tb |
| cDC1 | MAGEF1 | 2.93E-09 | 0.580183266 | 0.514 | 0.29 | 5.77E-05 | Tb |
| cDC1 | GFPT1 | 6.32E-08 | 0.579330134 | 0.482 | 0.277 | 0.001243714 | Tb |
| cDC1 | CDKAL1 | 2.27E-10 | 0.578342853 | 0.462 | 0.24 | 4.47E-06 | Tb |
| cDC1 | RRP1B | 1.87E-07 | 0.578333663 | 0.331 | 0.164 | 0.003685776 | Tb |
| cDC1 | QRICH1 | 3.20E-08 | 0.576476124 | 0.355 | 0.173 | 0.000629067 | Tb |
| cDC1 | RUNX1 | 2.77E-10 | 0.57433751 | 0.916 | 0.773 | 5.45E-06 | Tb |
| cDC1 | KDM2A | 2.52E-09 | 0.573880359 | 0.841 | 0.666 | 4.96E-05 | Tb |
| cDC1 | STRN3 | 6.72E-08 | 0.573853305 | 0.446 | 0.245 | 0.001323412 | Tb |
| cDC1 | TXNDC11 | 6.79E-11 | 0.573670439 | 0.518 | 0.294 | 1.34E-06 | Tb |
| cDC1 | VPS8 | 1.43E-06 | 0.572994349 | 0.323 | 0.172 | 0.028239746 | Tb |

| | | | | | | | |
|---|---|---|---|---|---|---|---|
| cDC1 | BRAF | 2.40E-09 | 0.572928403 | 0.713 | 0.48 | 4.72E-05 | Tb |
| cDC1 | CHD8 | 3.73E-07 | 0.572891588 | 0.39 | 0.212 | 0.007332608 | Tb |
| cDC1 | TRIO | 5.41E-13 | 0.572240245 | 0.845 | 0.591 | 1.07E-08 | Tb |
| cDC1 | SLCO3A1 | 5.86E-14 | 0.570819763 | 0.884 | 0.65 | 1.15E-09 | Tb |
| cDC1 | NFATC2 | 6.63E-14 | 0.570728599 | 0.781 | 0.517 | 1.30E-09 | Tb |
| cDC1 | LEPROTL1 | 4.54E-09 | 0.568962186 | 0.629 | 0.386 | 8.93E-05 | Tb |
| cDC1 | MAPKAP1 | 2.09E-07 | 0.567078821 | 0.59 | 0.367 | 0.004123423 | Tb |
| cDC1 | ARFGEF2 | 5.44E-08 | 0.566380934 | 0.446 | 0.24 | 0.001071011 | Tb |
| cDC1 | ATXN10 | 8.77E-09 | 0.565136979 | 0.594 | 0.352 | 0.000172583 | Tb |
| cDC1 | LMBR1 | 2.20E-07 | 0.564865368 | 0.426 | 0.241 | 0.004327051 | Tb |
| cDC1 | STK3 | 1.11E-07 | 0.564785044 | 0.526 | 0.331 | 0.002190028 | Tb |
| cDC1 | AUTS2 | 3.26E-10 | 0.564250362 | 0.928 | 0.768 | 6.42E-06 | Tb |
| cDC1 | WDR37 | 5.17E-07 | 0.564082343 | 0.41 | 0.231 | 0.010184713 | Tb |
| cDC1 | TMEM131L | 2.17E-10 | 0.563590348 | 0.454 | 0.225 | 4.27E-06 | Tb |
| cDC1 | ACSS2 | 6.19E-11 | 0.5627508 | 0.59 | 0.327 | 1.22E-06 | Tb |
| cDC1 | ST3GAL2 | 5.48E-07 | 0.562468581 | 0.434 | 0.245 | 0.010778939 | Tb |
| cDC1 | RAD9A | 6.06E-07 | 0.562352071 | 0.295 | 0.147 | 0.011924349 | Tb |
| cDC1 | NAGK | 8.33E-07 | 0.562091327 | 0.709 | 0.523 | 0.016402286 | Tb |
| cDC1 | DIMT1 | 1.19E-08 | 0.5618019 | 0.582 | 0.372 | 0.00023384 | Tb |
| cDC1 | HPS5 | 2.57E-13 | 0.561579518 | 0.896 | 0.619 | 5.06E-09 | Tb |
| cDC1 | SLC31A2 | 2.56E-07 | 0.561570386 | 0.414 | 0.23 | 0.005029571 | Tb |
| cDC1 | STK38 | 1.76E-08 | 0.561404199 | 0.371 | 0.182 | 0.000346121 | Tb |
| cDC1 | DSE | 5.44E-10 | 0.561247172 | 0.928 | 0.767 | 1.07E-05 | Tb |
| cDC1 | CLSTN1 | 4.58E-08 | 0.559216088 | 0.371 | 0.192 | 0.000901094 | Tb |
| cDC1 | TMEM184B | 2.12E-10 | 0.558970221 | 0.251 | 0.09 | 4.17E-06 | Tb |
| cDC1 | ARL6IP5 | 1.09E-09 | 0.55890079 | 0.88 | 0.706 | 2.15E-05 | Tb |
| cDC1 | DYNC1LI2 | 1.98E-09 | 0.557748896 | 0.57 | 0.318 | 3.89E-05 | Tb |
| cDC1 | ELL | 1.67E-07 | 0.55586746 | 0.359 | 0.19 | 0.003287309 | Tb |
| cDC1 | SP110 | 1.81E-11 | 0.555033001 | 0.741 | 0.47 | 3.56E-07 | Tb |
| cDC1 | TM9SF4 | 7.79E-10 | 0.554922019 | 0.355 | 0.16 | 1.53E-05 | Tb |
| cDC1 | TAF1 | 1.48E-06 | 0.55439844 | 0.327 | 0.17 | 0.029222828 | Tb |
| cDC1 | HMG20A | 1.33E-06 | 0.553832495 | 0.251 | 0.117 | 0.026137101 | Tb |
| cDC1 | RIOK1 | 5.62E-07 | 0.553698931 | 0.398 | 0.214 | 0.011054947 | Tb |
| cDC1 | TUBB6 | 1.63E-08 | 0.553473989 | 0.677 | 0.45 | 0.00032118 | Tb |
| cDC1 | MT-ND3 | 7.00E-12 | 0.55309559 | 0.992 | 0.907 | 1.38E-07 | Tb |
| cDC1 | CEP63 | 1.99E-07 | 0.552910742 | 0.406 | 0.218 | 0.00392311 | Tb |
| cDC1 | PRRG4 | 1.32E-07 | 0.552861076 | 0.454 | 0.256 | 0.002599102 | Tb |
| cDC1 | VPS53 | 1.76E-06 | 0.552843751 | 0.43 | 0.255 | 0.034660097 | Tb |
| cDC1 | KRIT1 | 1.19E-07 | 0.552658757 | 0.323 | 0.16 | 0.002342988 | Tb |
| cDC1 | SKIL | 1.03E-11 | 0.551259371 | 0.761 | 0.492 | 2.04E-07 | Tb |
| cDC1 | ZMYM2 | 7.16E-09 | 0.55061345 | 0.681 | 0.472 | 0.000140974 | Tb |
| cDC1 | ZKSCAN1 | 9.15E-08 | 0.549873659 | 0.386 | 0.201 | 0.001802069 | Tb |
| cDC1 | INIP | 1.90E-07 | 0.549822228 | 0.466 | 0.261 | 0.003730539 | Tb |
| cDC1 | LARP4B | 3.29E-10 | 0.549821573 | 0.749 | 0.461 | 6.48E-06 | Tb |
| cDC1 | LNPEP | 2.25E-08 | 0.548790027 | 0.55 | 0.323 | 0.000443236 | Tb |
| cDC1 | EHMT1 | 7.56E-12 | 0.54811574 | 0.614 | 0.339 | 1.49E-07 | Tb |
| cDC1 | TTC3 | 6.97E-07 | 0.547916766 | 0.657 | 0.448 | 0.013713032 | Tb |
| cDC1 | NCK1 | 2.29E-06 | 0.547814384 | 0.398 | 0.225 | 0.045055787 | Tb |
| cDC1 | AP1S1 | 3.74E-07 | 0.547191581 | 0.355 | 0.184 | 0.00736427 | Tb |
| cDC1 | MT-CO1 | 9.45E-12 | 0.546841946 | 1 | 0.989 | 1.86E-07 | Tb |
| cDC1 | AP5Z1 | 2.73E-10 | 0.545116652 | 0.39 | 0.176 | 5.37E-06 | Tb |
| cDC1 | IPMK | 4.41E-07 | 0.544164941 | 0.649 | 0.438 | 0.008690008 | Tb |
| cDC1 | FAF2 | 5.47E-08 | 0.543636526 | 0.367 | 0.184 | 0.001077163 | Tb |
| cDC1 | IFNAR2 | 5.26E-08 | 0.543276838 | 0.61 | 0.39 | 0.001036126 | Tb |
| cDC1 | ABHD31 | 2.32E-06 | 0.542365395 | 0.653 | 0.49 | 0.04558039 | Tb |
| cDC1 | RABGAP1L | 3.00E-09 | 0.5420814 | 0.837 | 0.681 | 5.91E-05 | Tb |

| | | | | | | | |
|---|---|---|---|---|---|---|---|
| cDC1 | AFF4 | 9.88E-09 | 0.541583408 | 0.865 | 0.672 | 0.000194416 | Tb |
| cDC1 | CELF1 | 2.38E-06 | 0.541372213 | 0.534 | 0.342 | 0.046825984 | Tb |
| cDC1 | CHD7 | 4.20E-10 | 0.541011984 | 0.49 | 0.258 | 8.26E-06 | Tb |
| cDC1 | STAT6 | 2.10E-08 | 0.540275652 | 0.562 | 0.361 | 0.00041284 | Tb |
| cDC1 | LY75 | 1.19E-06 | 0.540092824 | 0.649 | 0.452 | 0.023503219 | Tb |
| cDC1 | VAV3 | 5.88E-10 | 0.539900481 | 0.857 | 0.668 | 1.16E-05 | Tb |
| cDC1 | IBTK | 1.90E-07 | 0.539591667 | 0.454 | 0.255 | 0.003735037 | Tb |
| cDC1 | UVRAG | 1.20E-12 | 0.539583868 | 0.912 | 0.705 | 2.35E-08 | Tb |
| cDC1 | ZZEF1 | 8.99E-11 | 0.538919834 | 0.466 | 0.228 | 1.77E-06 | Tb |
| cDC1 | MT-ND1 | 1.95E-15 | 0.538814398 | 0.992 | 0.92 | 3.84E-11 | Tb |
| cDC1 | POM121 | 4.91E-08 | 0.5387566 | 0.534 | 0.325 | 0.000965716 | Tb |
| cDC1 | MAP3K8 | 2.31E-14 | 0.538670126 | 0.888 | 0.726 | 4.54E-10 | Tb |
| cDC1 | RABGEF1 | 3.21E-07 | 0.538652764 | 0.821 | 0.685 | 0.006317819 | Tb |
| cDC1 | COA1 | 1.88E-06 | 0.538278558 | 0.514 | 0.307 | 0.037065107 | Tb |
| cDC1 | NSD1 | 4.69E-07 | 0.538232897 | 0.578 | 0.368 | 0.009238291 | Tb |
| cDC1 | RAB5B | 7.67E-08 | 0.538201916 | 0.323 | 0.158 | 0.001509896 | Tb |
| cDC1 | EIF4ENIF1 | 5.20E-09 | 0.537882842 | 0.251 | 0.098 | 0.000102401 | Tb |
| cDC1 | SMAD2 | 6.18E-09 | 0.537310023 | 0.458 | 0.243 | 0.000121588 | Tb |
| cDC1 | PHC2 | 6.58E-08 | 0.536368553 | 0.578 | 0.358 | 0.001295797 | Tb |
| cDC1 | MBD4 | 8.48E-08 | 0.535987987 | 0.566 | 0.339 | 0.001668951 | Tb |
| cDC1 | VTI1A | 5.47E-10 | 0.535260289 | 0.669 | 0.422 | 1.08E-05 | Tb |
| cDC1 | CD38 | 1.51E-08 | 0.535098212 | 0.534 | 0.322 | 0.000297695 | Tb |
| cDC1 | GRAMD1A | 1.26E-09 | 0.535016587 | 0.486 | 0.26 | 2.48E-05 | Tb |
| cDC1 | GALNT1 | 1.78E-08 | 0.534895918 | 0.462 | 0.255 | 0.000350315 | Tb |
| cDC1 | TAF15 | 7.04E-10 | 0.53462468 | 0.689 | 0.45 | 1.39E-05 | Tb |
| cDC1 | SNX30 | 2.56E-07 | 0.534195276 | 0.514 | 0.308 | 0.005039488 | Tb |
| cDC1 | CHMP3 | 8.52E-09 | 0.533485066 | 0.502 | 0.278 | 0.000167788 | Tb |
| cDC1 | IL3RA | 2.39E-09 | 0.533370906 | 0.287 | 0.122 | 4.70E-05 | Tb |
| cDC1 | PPP6R3 | 3.12E-07 | 0.533120115 | 0.649 | 0.436 | 0.006134515 | Tb |
| cDC1 | GOLIM4 | 6.53E-07 | 0.532427558 | 0.562 | 0.358 | 0.012851263 | Tb |
| cDC1 | CREBRF | 2.45E-07 | 0.532222927 | 0.53 | 0.312 | 0.004823686 | Tb |
| cDC1 | ACLY | 8.64E-08 | 0.531158462 | 0.375 | 0.193 | 0.001700038 | Tb |
| cDC1 | CAP1 | 7.90E-13 | 0.530585622 | 0.92 | 0.777 | 1.55E-08 | Tb |
| cDC1 | RIPOR2 | 1.19E-09 | 0.529963407 | 0.705 | 0.434 | 2.35E-05 | Tb |
| cDC1 | SREBF2 | 1.09E-08 | 0.528338746 | 0.39 | 0.193 | 0.000213651 | Tb |
| cDC1 | EXOC6 | 1.89E-08 | 0.52792141 | 0.454 | 0.25 | 0.000371225 | Tb |
| cDC1 | GPCPD1 | 8.12E-11 | 0.527604052 | 0.777 | 0.544 | 1.60E-06 | Tb |
| cDC1 | CLN3 | 1.84E-08 | 0.526885423 | 0.49 | 0.283 | 0.000362465 | Tb |
| cDC1 | ACSL4 | 4.97E-07 | 0.526728451 | 0.625 | 0.412 | 0.009778568 | Tb |
| cDC1 | EIPR1 | 9.05E-08 | 0.526618883 | 0.378 | 0.199 | 0.00178159 | Tb |
| cDC1 | WWP2 | 1.86E-06 | 0.526199645 | 0.51 | 0.328 | 0.036645665 | Tb |
| cDC1 | IL10RB | 4.21E-07 | 0.525775462 | 0.375 | 0.207 | 0.008278579 | Tb |
| cDC1 | CARMIL1 | 1.73E-07 | 0.525194243 | 0.518 | 0.318 | 0.003403831 | Tb |
| cDC1 | SLC9A7 | 8.04E-10 | 0.524871081 | 0.721 | 0.498 | 1.58E-05 | Tb |
| cDC1 | SRSF4 | 5.50E-07 | 0.523767767 | 0.673 | 0.468 | 0.010830583 | Tb |
| cDC1 | USP9X | 5.76E-07 | 0.522392662 | 0.681 | 0.46 | 0.011340687 | Tb |
| cDC1 | OGA | 2.44E-10 | 0.520449818 | 0.474 | 0.25 | 4.81E-06 | Tb |
| cDC1 | RFTN1 | 6.74E-11 | 0.520380499 | 0.761 | 0.568 | 1.33E-06 | Tb |
| cDC1 | ZXDC | 9.22E-07 | 0.519723134 | 0.363 | 0.197 | 0.018144167 | Tb |
| cDC1 | NEMF | 2.08E-07 | 0.519490191 | 0.582 | 0.36 | 0.004098865 | Tb |
| cDC1 | RELB | 1.72E-07 | 0.518678293 | 0.705 | 0.514 | 0.003389431 | Tb |
| cDC1 | ZNF641 | 1.59E-07 | 0.518050017 | 0.299 | 0.138 | 0.003124958 | Tb |
| cDC1 | ILKAP | 5.65E-07 | 0.517640819 | 0.438 | 0.243 | 0.011125288 | Tb |
| cDC1 | ORAI2 | 3.02E-12 | 0.516898062 | 0.438 | 0.201 | 5.94E-08 | Tb |
| cDC1 | EIF4G3 | 4.77E-08 | 0.516762498 | 0.59 | 0.393 | 0.000938905 | Tb |
| cDC1 | CD69 | 1.84E-10 | 0.516266345 | 0.506 | 0.262 | 3.63E-06 | Tb |

| | | | | | | | |
|---|---|---|---|---|---|---|---|
| cDC1 | SS18 | 2.52E-06 | 0.515607112 | 0.394 | 0.225 | 0.049622763 | Tb |
| cDC1 | CEP68 | 5.23E-12 | 0.515564466 | 0.614 | 0.335 | 1.03E-07 | Tb |
| cDC1 | IL2RG | 7.21E-08 | 0.515130523 | 0.498 | 0.281 | 0.001418773 | Tb |
| cDC1 | GLIPR2 | 3.53E-08 | 0.514935567 | 0.789 | 0.569 | 0.000694277 | Tb |
| cDC1 | HEATR5B | 3.97E-08 | 0.514882904 | 0.375 | 0.197 | 0.000781576 | Tb |
| cDC1 | TCF12 | 2.65E-09 | 0.514301376 | 0.697 | 0.465 | 5.22E-05 | Tb |
| cDC1 | ADD3 | 7.53E-07 | 0.513998396 | 0.486 | 0.299 | 0.01481799 | Tb |
| cDC1 | ASAP1 | 1.33E-13 | 0.512896588 | 0.98 | 0.887 | 2.61E-09 | Tb |
| cDC1 | KCMF1 | 4.54E-07 | 0.51208205 | 0.705 | 0.493 | 0.008944906 | Tb |
| cDC1 | SLC16A3 | 5.83E-09 | 0.511978616 | 0.398 | 0.197 | 0.00011472 | Tb |
| cDC1 | TMC6 | 5.37E-07 | 0.511143235 | 0.363 | 0.19 | 0.010565088 | Tb |
| cDC1 | INPP5F | 9.40E-09 | 0.510894697 | 0.809 | 0.603 | 0.000184994 | Tb |
| cDC1 | SCARF1 | 1.68E-09 | 0.510778405 | 0.637 | 0.376 | 3.30E-05 | Tb |
| cDC1 | NAAA | 2.07E-12 | 0.510354019 | 0.956 | 0.806 | 4.07E-08 | Tb |
| cDC1 | HDAC1 | 3.60E-08 | 0.509514781 | 0.494 | 0.267 | 0.000708414 | Tb |
| cDC1 | TBL1XR1 | 2.41E-07 | 0.509397995 | 0.721 | 0.49 | 0.004741887 | Tb |
| cDC1 | ITPR2 | 3.08E-08 | 0.508608539 | 0.375 | 0.191 | 0.000606978 | Tb |
| cDC1 | ETNK1 | 1.98E-07 | 0.508194162 | 0.542 | 0.333 | 0.003897974 | Tb |
| cDC1 | MTMR4 | 6.58E-07 | 0.508156492 | 0.303 | 0.151 | 0.012956531 | Tb |
| cDC1 | RUFY3 | 3.69E-09 | 0.507504793 | 0.717 | 0.512 | 7.26E-05 | Tb |
| cDC1 | UBR2 | 1.73E-09 | 0.506823476 | 0.637 | 0.378 | 3.41E-05 | Tb |
| cDC1 | MRPL17 | 3.35E-07 | 0.506140549 | 0.402 | 0.224 | 0.006587801 | Tb |
| cDC1 | EPB41L2 | 2.22E-07 | 0.505077132 | 0.614 | 0.44 | 0.004378833 | Tb |
| cDC1 | R3HDM2 | 2.33E-07 | 0.504756603 | 0.59 | 0.408 | 0.004594328 | Tb |
| cDC1 | PRR14L | 3.89E-09 | 0.503577877 | 0.534 | 0.314 | 7.65E-05 | Tb |
| cDC1 | ATXN2 | 5.87E-07 | 0.502667838 | 0.47 | 0.284 | 0.011554005 | Tb |
| cDC1 | PCSK7 | 3.44E-07 | 0.502524519 | 0.371 | 0.198 | 0.006768498 | Tb |
| cDC1 | RNASEH2C | 7.48E-07 | 0.501863344 | 0.562 | 0.367 | 0.014720921 | Tb |
| cDC1 | LRCH3 | 7.25E-08 | 0.500893522 | 0.371 | 0.192 | 0.001426944 | Tb |
| **Disease-associated genes in cDC2** | | | | | | | |
| cDC2 | APOE | 5.31E-14 | 2.604117383 | 0.41 | 0.27 | 1.04E-09 | Lep-L |
| cDC2 | MMP19 | 1.37E-10 | 2.29764461 | 0.301 | 0.172 | 2.69E-06 | Lep-L |
| cDC2 | NBPF19 | 6.60E-14 | 1.95076022 | 0.295 | 0.15 | 1.30E-09 | Lep-L |
| cDC2 | PLIN2 | 3.30E-13 | 1.90623971 | 0.497 | 0.373 | 6.50E-09 | Lep-L |
| cDC2 | CCR7 | 1.24E-22 | 1.903688913 | 0.59 | 0.377 | 2.44E-18 | Lep-L |
| cDC2 | CHST7 | 8.09E-23 | 1.831932217 | 0.295 | 0.105 | 1.59E-18 | Lep-L |
| cDC2 | CXCL3 | 4.76E-14 | 1.774513442 | 0.476 | 0.308 | 9.37E-10 | Lep-L |
| cDC2 | MTSS1 | 8.44E-22 | 1.767033932 | 0.476 | 0.269 | 1.66E-17 | Lep-L |
| cDC2 | C5AR1 | 7.86E-12 | 1.72453908 | 0.41 | 0.269 | 1.55E-07 | Lep-L |
| cDC2 | ADA | 4.41E-18 | 1.591349859 | 0.437 | 0.249 | 8.68E-14 | Lep-L |
| cDC2 | IL1R2 | 3.79E-25 | 1.580580154 | 0.66 | 0.434 | 7.46E-21 | Lep-L |
| cDC2 | KCNN4 | 5.18E-16 | 1.491170046 | 0.485 | 0.314 | 1.02E-11 | Lep-L |
| cDC2 | SDS | 7.05E-07 | 1.475373524 | 0.34 | 0.241 | 0.01387023 | Lep-L |
| cDC2 | FKBP4 | 9.94E-30 | 1.455609269 | 0.681 | 0.451 | 1.96E-25 | Lep-L |
| cDC2 | HSPA6 | 8.43E-17 | 1.437213523 | 0.702 | 0.569 | 1.66E-12 | Lep-L |
| cDC2 | MAFG | 1.28E-16 | 1.406521956 | 0.476 | 0.312 | 2.52E-12 | Lep-L |
| cDC2 | VSIG4 | 2.38E-11 | 1.371746419 | 0.283 | 0.145 | 4.69E-07 | Lep-L |
| cDC2 | PIM2 | 9.74E-08 | 1.37070503 | 0.262 | 0.158 | 0.001916631 | Lep-L |
| cDC2 | PID1 | 2.10E-09 | 1.360693393 | 0.581 | 0.485 | 4.14E-05 | Lep-L |
| cDC2 | MGLL | 1.92E-13 | 1.351512456 | 0.401 | 0.236 | 3.78E-09 | Lep-L |
| cDC2 | PAF1 | 1.92E-13 | 1.346179839 | 0.47 | 0.326 | 3.79E-09 | Lep-L |
| cDC2 | SLCO4A1 | 4.14E-09 | 1.339602773 | 0.268 | 0.151 | 8.14E-05 | Lep-L |
| cDC2 | TRAF4 | 2.04E-09 | 1.303079583 | 0.301 | 0.182 | 4.01E-05 | Lep-L |
| cDC2 | AHR | 4.20E-24 | 1.299404325 | 0.795 | 0.733 | 8.28E-20 | Lep-L |
| cDC2 | SERPINH1 | 3.03E-18 | 1.290058185 | 0.452 | 0.252 | 5.96E-14 | Lep-L |
| cDC2 | ZC3H12A | 2.01E-20 | 1.288575221 | 0.711 | 0.566 | 3.95E-16 | Lep-L |

| | | | | | | | |
|---|---|---|---|---|---|---|---|
| cDC2 | B3GNT2 | 3.71E-07 | 1.281212362 | 0.358 | 0.263 | 0.00731002 | Lep-L |
| cDC2 | ANKLE2 | 1.26E-11 | 1.271865353 | 0.476 | 0.37 | 2.48E-07 | Lep-L |
| cDC2 | TRIP10 | 7.43E-11 | 1.253549141 | 0.383 | 0.245 | 1.46E-06 | Lep-L |
| cDC2 | HSPB1 | 3.17E-29 | 1.232228159 | 0.913 | 0.807 | 6.23E-25 | Lep-L |
| cDC2 | MFSD2A | 1.05E-07 | 1.184461341 | 0.343 | 0.238 | 0.002060609 | Lep-L |
| cDC2 | SLC6A6 | 4.66E-09 | 1.176248403 | 0.431 | 0.322 | 9.17E-05 | Lep-L |
| cDC2 | RHOF | 6.04E-11 | 1.158349255 | 0.587 | 0.493 | 1.19E-06 | Lep-L |
| cDC2 | CXCL8 | 1.32E-34 | 1.154748133 | 0.834 | 0.64 | 2.60E-30 | Lep-L |
| cDC2 | ERGIC1 | 2.10E-07 | 1.145961304 | 0.455 | 0.385 | 0.004135848 | Lep-L |
| cDC2 | SLC25A37 | 1.08E-08 | 1.129271677 | 0.47 | 0.37 | 0.000213124 | Lep-L |
| cDC2 | RILPL2 | 7.67E-22 | 1.124737792 | 0.747 | 0.696 | 1.51E-17 | Lep-L |
| cDC2 | PDE4DIP | 2.16E-15 | 1.107841544 | 0.587 | 0.436 | 4.24E-11 | Lep-L |
| cDC2 | BTG1 | 3.31E-28 | 1.107326929 | 0.94 | 0.908 | 6.52E-24 | Lep-L |
| cDC2 | MAP7D1 | 4.85E-14 | 1.104553283 | 0.59 | 0.494 | 9.55E-10 | Lep-L |
| cDC2 | ZFP36L1 | 6.91E-11 | 1.096229408 | 0.825 | 0.826 | 1.36E-06 | Lep-L |
| cDC2 | DDX3Y | 1.16E-12 | 1.090145371 | 0.503 | 0.321 | 2.28E-08 | Lep-L |
| cDC2 | ATP1A1 | 5.42E-21 | 1.088573832 | 0.741 | 0.653 | 1.07E-16 | Lep-L |
| cDC2 | SOD2 | 1.86E-18 | 1.078722508 | 0.958 | 0.956 | 3.66E-14 | Lep-L |
| cDC2 | ABCG1 | 7.62E-08 | 1.076024991 | 0.419 | 0.309 | 0.001500497 | Lep-L |
| cDC2 | SERPINB9 | 4.10E-29 | 1.068645095 | 0.895 | 0.855 | 8.06E-25 | Lep-L |
| cDC2 | ITGA5 | 1.01E-09 | 1.066679718 | 0.476 | 0.367 | 1.99E-05 | Lep-L |
| cDC2 | MAP2K3 | 3.29E-24 | 1.052770269 | 0.825 | 0.743 | 6.47E-20 | Lep-L |
| cDC2 | CSGALNACT2 | 5.22E-11 | 1.043767701 | 0.651 | 0.629 | 1.03E-06 | Lep-L |
| cDC2 | CDC42EP3 | 2.60E-09 | 1.035782173 | 0.63 | 0.571 | 5.11E-05 | Lep-L |
| cDC2 | NOTCH2NLC | 1.16E-06 | 1.031671324 | 0.292 | 0.192 | 0.022861245 | Lep-L |
| cDC2 | RBFOX2 | 1.92E-10 | 1.026115843 | 0.654 | 0.616 | 3.78E-06 | Lep-L |
| cDC2 | CTTNBP2NL | 1.24E-12 | 1.020956684 | 0.587 | 0.482 | 2.43E-08 | Lep-L |
| cDC2 | HSPA1A | 5.03E-25 | 0.99619714 | 0.988 | 0.956 | 9.90E-21 | Lep-L |
| cDC2 | TSPYL2 | 7.47E-09 | 0.993878024 | 0.527 | 0.412 | 0.000147043 | Lep-L |
| cDC2 | NFKB2 | 8.45E-11 | 0.99377944 | 0.533 | 0.425 | 1.66E-06 | Lep-L |
| cDC2 | BAG3 | 9.98E-11 | 0.9917693 | 0.705 | 0.623 | 1.96E-06 | Lep-L |
| cDC2 | RIT1 | 4.06E-09 | 0.98761893 | 0.488 | 0.4 | 7.99E-05 | Lep-L |
| cDC2 | PTTG1 | 1.80E-08 | 0.986697464 | 0.551 | 0.477 | 0.000354763 | Lep-L |
| cDC2 | MARCKS | 1.07E-11 | 0.986411873 | 0.681 | 0.597 | 2.10E-07 | Lep-L |
| cDC2 | SAT1 | 9.31E-34 | 0.971767163 | 0.982 | 0.991 | 1.83E-29 | Lep-L |
| cDC2 | HSPD1 | 6.98E-35 | 0.962928012 | 0.979 | 0.934 | 1.37E-30 | Lep-L |
| cDC2 | CXXC5 | 5.50E-07 | 0.958607838 | 0.325 | 0.228 | 0.010826853 | Lep-L |
| cDC2 | TNIP1 | 2.51E-09 | 0.957271086 | 0.527 | 0.438 | 4.94E-05 | Lep-L |
| cDC2 | DDIT4 | 2.01E-13 | 0.945011652 | 0.756 | 0.637 | 3.95E-09 | Lep-L |
| cDC2 | CEBPB | 2.00E-08 | 0.942039662 | 0.684 | 0.681 | 0.00039276 | Lep-L |
| cDC2 | DNAJB1 | 1.56E-07 | 0.935477381 | 0.901 | 0.91 | 0.003068699 | Lep-L |
| cDC2 | PPP1R18 | 2.43E-14 | 0.934783408 | 0.717 | 0.673 | 4.78E-10 | Lep-L |
| cDC2 | MMP9 | 1.18E-06 | 0.93357467 | 0.533 | 0.439 | 0.023267906 | Lep-L |
| cDC2 | PRNP | 1.20E-07 | 0.933476315 | 0.59 | 0.531 | 0.002368365 | Lep-L |
| cDC2 | STIP1 | 3.79E-16 | 0.932767084 | 0.657 | 0.541 | 7.47E-12 | Lep-L |
| cDC2 | CXCL2 | 1.03E-06 | 0.926715419 | 0.545 | 0.447 | 0.020269835 | Lep-L |
| cDC2 | EZR | 8.80E-21 | 0.903078783 | 0.922 | 0.885 | 1.73E-16 | Lep-L |
| cDC2 | MRPS6 | 1.53E-09 | 0.879646963 | 0.602 | 0.552 | 3.01E-05 | Lep-L |
| cDC2 | CD9 | 1.44E-07 | 0.878653403 | 0.443 | 0.333 | 0.002842757 | Lep-L |
| cDC2 | OPN3 | 6.11E-09 | 0.868373265 | 0.503 | 0.397 | 0.000120322 | Lep-L |
| cDC2 | CACYBP | 6.87E-17 | 0.866883854 | 0.792 | 0.747 | 1.35E-12 | Lep-L |
| cDC2 | FCGR2A | 9.79E-15 | 0.865836793 | 0.777 | 0.745 | 1.93E-10 | Lep-L |
| cDC2 | PPTC7 | 2.61E-08 | 0.86431302 | 0.476 | 0.382 | 0.000514369 | Lep-L |
| cDC2 | TIMP1 | 8.34E-14 | 0.861589274 | 0.904 | 0.878 | 1.64E-09 | Lep-L |
| cDC2 | STK4 | 5.78E-15 | 0.844128319 | 0.801 | 0.775 | 1.14E-10 | Lep-L |
| cDC2 | HSPH1 | 1.86E-24 | 0.822582162 | 0.91 | 0.855 | 3.67E-20 | Lep-L |

| | | | | | | | |
|---|---|---|---|---|---|---|---|
| cDC2 | ANKRD10 | 1.19E-10 | 0.821866545 | 0.596 | 0.508 | 2.34E-06 | Lep-L |
| cDC2 | BTG3 | 2.51E-08 | 0.820837396 | 0.542 | 0.468 | 0.000494017 | Lep-L |
| cDC2 | RARA | 1.85E-08 | 0.817812199 | 0.566 | 0.489 | 0.000365042 | Lep-L |
| cDC2 | ETS2 | 1.02E-10 | 0.813165065 | 0.708 | 0.67 | 2.01E-06 | Lep-L |
| cDC2 | G3BP1 | 3.13E-08 | 0.803181659 | 0.623 | 0.582 | 0.000615829 | Lep-L |
| cDC2 | TCP1 | 2.75E-17 | 0.801930552 | 0.768 | 0.711 | 5.41E-13 | Lep-L |
| cDC2 | IL10RA | 4.04E-10 | 0.794906221 | 0.717 | 0.694 | 7.95E-06 | Lep-L |
| cDC2 | RAB8B | 3.31E-09 | 0.792835365 | 0.693 | 0.678 | 6.52E-05 | Lep-L |
| cDC2 | AHSA1 | 1.04E-08 | 0.785157306 | 0.617 | 0.575 | 0.000204231 | Lep-L |
| cDC2 | METRNL | 1.98E-07 | 0.783162738 | 0.738 | 0.768 | 0.003893523 | Lep-L |
| cDC2 | PNPLA8 | 1.85E-07 | 0.782714156 | 0.651 | 0.64 | 0.003644399 | Lep-L |
| cDC2 | DOK2 | 6.48E-08 | 0.775413773 | 0.666 | 0.649 | 0.001276465 | Lep-L |
| cDC2 | HSP90AB1 | 8.69E-26 | 0.774899779 | 0.994 | 0.983 | 1.71E-21 | Lep-L |
| cDC2 | SQSTM1 | 4.55E-22 | 0.774333003 | 0.937 | 0.895 | 8.96E-18 | Lep-L |
| cDC2 | CAPN2 | 7.59E-07 | 0.771414384 | 0.614 | 0.602 | 0.014941062 | Lep-L |
| cDC2 | YY1 | 1.62E-11 | 0.768832703 | 0.702 | 0.664 | 3.18E-07 | Lep-L |
| cDC2 | CHORDC1 | 1.10E-09 | 0.765494757 | 0.69 | 0.634 | 2.17E-05 | Lep-L |
| cDC2 | HBP1 | 2.35E-09 | 0.760430262 | 0.569 | 0.511 | 4.63E-05 | Lep-L |
| cDC2 | EEF2 | 1.44E-19 | 0.749564354 | 0.87 | 0.904 | 2.84E-15 | Lep-L |
| cDC2 | CD44 | 1.01E-14 | 0.736888237 | 0.955 | 0.933 | 2.00E-10 | Lep-L |
| cDC2 | HSPE1 | 1.03E-13 | 0.736681752 | 0.937 | 0.943 | 2.02E-09 | Lep-L |
| cDC2 | HSPA8 | 1.24E-08 | 0.725047594 | 0.961 | 0.974 | 0.000243185 | Lep-L |
| cDC2 | NAA50 | 8.43E-08 | 0.711339957 | 0.575 | 0.505 | 0.001659338 | Lep-L |
| cDC2 | PPP4R2 | 3.19E-07 | 0.710637915 | 0.536 | 0.466 | 0.00628123 | Lep-L |
| cDC2 | HLA-DRB5 | 2.29E-25 | 0.703711657 | 0.907 | 0.784 | 4.51E-21 | Lep-L |
| cDC2 | TNFAIP3 | 3.74E-12 | 0.702591866 | 0.88 | 0.862 | 7.36E-08 | Lep-L |
| cDC2 | HNRNPA0 | 5.08E-09 | 0.699344456 | 0.681 | 0.674 | 1.00E-04 | Lep-L |
| cDC2 | SDCBP | 1.13E-14 | 0.698374188 | 0.852 | 0.86 | 2.23E-10 | Lep-L |
| cDC2 | ALDOA | 2.46E-07 | 0.691885469 | 0.512 | 0.413 | 0.004835021 | Lep-L |
| cDC2 | SNHG7 | 1.11E-07 | 0.689505328 | 0.608 | 0.565 | 0.002186946 | Lep-L |
| cDC2 | MGAT1 | 1.91E-08 | 0.682963021 | 0.753 | 0.786 | 0.000376838 | Lep-L |
| cDC2 | MYO9B | 1.28E-09 | 0.676670948 | 0.705 | 0.666 | 2.53E-05 | Lep-L |
| cDC2 | EMP3 | 3.40E-18 | 0.676485793 | 0.931 | 0.919 | 6.69E-14 | Lep-L |
| cDC2 | CYTIP | 1.68E-11 | 0.675952224 | 0.828 | 0.832 | 3.31E-07 | Lep-L |
| cDC2 | PTGES3 | 4.29E-16 | 0.672191396 | 0.87 | 0.868 | 8.45E-12 | Lep-L |
| cDC2 | ZNF706 | 1.02E-08 | 0.669931204 | 0.708 | 0.72 | 0.000201463 | Lep-L |
| cDC2 | FXR1 | 1.97E-06 | 0.668379755 | 0.608 | 0.593 | 0.038864601 | Lep-L |
| cDC2 | HECA | 1.26E-06 | 0.66634257 | 0.636 | 0.573 | 0.024722696 | Lep-L |
| cDC2 | IER3 | 9.31E-15 | 0.651454489 | 0.886 | 0.841 | 1.83E-10 | Lep-L |
| cDC2 | MCL1 | 6.73E-12 | 0.63507168 | 0.901 | 0.905 | 1.33E-07 | Lep-L |
| cDC2 | NINJ1 | 4.20E-11 | 0.63144529 | 0.777 | 0.737 | 8.28E-07 | Lep-L |
| cDC2 | BIRC3 | 2.86E-08 | 0.627287401 | 0.762 | 0.719 | 0.000562544 | Lep-L |
| cDC2 | PPP2CA | 7.74E-07 | 0.624316464 | 0.596 | 0.577 | 0.015242154 | Lep-L |
| cDC2 | FTL | 1.91E-14 | 0.621568772 | 0.997 | 0.991 | 3.77E-10 | Lep-L |
| cDC2 | HSP90AA1 | 1.01E-19 | 0.614725884 | 0.994 | 0.987 | 1.99E-15 | Lep-L |
| cDC2 | GLA | 6.33E-08 | 0.611822815 | 0.75 | 0.731 | 0.001246677 | Lep-L |
| cDC2 | INSIG1 | 9.45E-10 | 0.60775437 | 0.858 | 0.834 | 1.86E-05 | Lep-L |
| cDC2 | CD68 | 1.94E-07 | 0.599734465 | 0.714 | 0.666 | 0.003818187 | Lep-L |
| cDC2 | MTCH1 | 1.03E-07 | 0.592399962 | 0.684 | 0.7 | 0.002031992 | Lep-L |
| cDC2 | DNAJA1 | 1.91E-06 | 0.590022658 | 0.934 | 0.959 | 0.037556947 | Lep-L |
| cDC2 | ARL8B | 1.84E-11 | 0.585336896 | 0.813 | 0.811 | 3.63E-07 | Lep-L |
| cDC2 | B4GALT1 | 1.87E-06 | 0.584949986 | 0.696 | 0.687 | 0.036783365 | Lep-L |
| cDC2 | NFKBIA | 6.82E-21 | 0.581596107 | 0.967 | 0.955 | 1.34E-16 | Lep-L |
| cDC2 | UBE2R2 | 4.80E-11 | 0.571499412 | 0.72 | 0.666 | 9.45E-07 | Lep-L |
| cDC2 | FTH1 | 1.37E-22 | 0.570606975 | 1 | 0.996 | 2.70E-18 | Lep-L |
| cDC2 | LMNA | 2.60E-09 | 0.567301978 | 0.892 | 0.837 | 5.12E-05 | Lep-L |

| | | | | | | | |
|---|---|---|---|---|---|---|---|
| cDC2 | CTSZ | 2.62E-10 | 0.555517176 | 0.931 | 0.956 | 5.16E-06 | Lep-L |
| cDC2 | ETV3 | 7.03E-10 | 0.554610337 | 0.753 | 0.686 | 1.38E-05 | Lep-L |
| cDC2 | ST13 | 1.14E-12 | 0.551228914 | 0.828 | 0.828 | 2.24E-08 | Lep-L |
| cDC2 | LRRFIP2 | 5.34E-07 | 0.545277641 | 0.717 | 0.708 | 0.010516826 | Lep-L |
| cDC2 | ALCAM | 1.32E-06 | 0.528254094 | 0.801 | 0.77 | 0.026013588 | Lep-L |
| cDC2 | VAPA | 8.11E-07 | 0.527142617 | 0.729 | 0.762 | 0.015970348 | Lep-L |
| cDC2 | H2AFZ | 1.91E-09 | 0.509099635 | 0.913 | 0.934 | 3.76E-05 | Lep-L |
| cDC2 | UBD | 2.11E-25 | 2.176162947 | 0.288 | 0.12 | 4.15E-21 | Lep-T |
| cDC2 | BEST11 | 2.87E-29 | 1.996328438 | 0.353 | 0.161 | 5.65E-25 | Lep-T |
| cDC2 | ALDOA1 | 9.64E-45 | 1.944913474 | 0.563 | 0.34 | 1.90E-40 | Lep-T |
| cDC2 | KCNMA1 | 5.73E-18 | 1.551348861 | 0.31 | 0.168 | 1.13E-13 | Lep-T |
| cDC2 | NFKBIA1 | 1.17E-55 | 1.542146538 | 0.984 | 0.939 | 2.30E-51 | Lep-T |
| cDC2 | ZFP36 | 2.48E-35 | 1.447716275 | 0.976 | 0.93 | 4.89E-31 | Lep-T |
| cDC2 | RBFOX21 | 4.11E-52 | 1.380598409 | 0.753 | 0.536 | 8.09E-48 | Lep-T |
| cDC2 | TNF | 2.25E-22 | 1.361444775 | 0.525 | 0.359 | 4.42E-18 | Lep-T |
| cDC2 | CD74 | 1.17E-58 | 1.266328199 | 0.997 | 0.997 | 2.30E-54 | Lep-T |
| cDC2 | CDC42EP2 | 7.71E-17 | 1.235127056 | 0.358 | 0.209 | 1.52E-12 | Lep-T |
| cDC2 | APOO | 1.75E-21 | 1.234073191 | 0.468 | 0.315 | 3.44E-17 | Lep-T |
| cDC2 | NLRP3 | 2.07E-19 | 1.230809296 | 0.554 | 0.424 | 4.08E-15 | Lep-T |
| cDC2 | PNRC1 | 3.73E-88 | 1.220361057 | 0.993 | 0.966 | 7.34E-84 | Lep-T |
| cDC2 | ICAM1 | 2.02E-29 | 1.195385121 | 0.79 | 0.7 | 3.97E-25 | Lep-T |
| cDC2 | CDKN1A1 | 1.56E-66 | 1.120864362 | 0.932 | 0.87 | 3.08E-62 | Lep-T |
| cDC2 | BCL31 | 5.57E-27 | 1.118710466 | 0.659 | 0.514 | 1.10E-22 | Lep-T |
| cDC2 | NRROS | 4.03E-09 | 1.064996167 | 0.255 | 0.162 | 7.94E-05 | Lep-T |
| cDC2 | HCAR3 | 1.02E-17 | 1.058958318 | 0.434 | 0.279 | 2.01E-13 | Lep-T |
| cDC2 | CXCL81 | 3.52E-20 | 1.049751987 | 0.753 | 0.614 | 6.93E-16 | Lep-T |
| cDC2 | IL1B | 1.52E-25 | 1.046849406 | 0.858 | 0.805 | 3.00E-21 | Lep-T |
| cDC2 | FCN1 | 3.58E-11 | 1.045668705 | 0.258 | 0.148 | 7.05E-07 | Lep-T |
| cDC2 | HLA-DQA2 | 2.32E-34 | 1.036124332 | 0.729 | 0.579 | 4.57E-30 | Lep-T |
| cDC2 | GPR84 | 6.71E-07 | 0.995282079 | 0.267 | 0.187 | 0.013199029 | Lep-T |
| cDC2 | RAB24 | 2.04E-08 | 0.993445044 | 0.346 | 0.261 | 0.000402114 | Lep-T |
| cDC2 | EMD1 | 5.72E-17 | 0.971332414 | 0.477 | 0.338 | 1.13E-12 | Lep-T |
| cDC2 | GPS1 | 1.70E-07 | 0.964662906 | 0.273 | 0.193 | 0.003356078 | Lep-T |
| cDC2 | U2AF1 | 7.83E-09 | 0.938504581 | 0.263 | 0.171 | 0.00015413 | Lep-T |
| cDC2 | CSRNP1 | 3.61E-39 | 0.936466876 | 0.779 | 0.607 | 7.10E-35 | Lep-T |
| cDC2 | CFP1 | 1.19E-17 | 0.934218464 | 0.515 | 0.361 | 2.34E-13 | Lep-T |
| cDC2 | IER31 | 1.13E-11 | 0.921450878 | 0.874 | 0.831 | 2.22E-07 | Lep-T |
| cDC2 | G0S2 | 1.54E-12 | 0.918830125 | 0.744 | 0.661 | 3.03E-08 | Lep-T |
| cDC2 | DUSP1 | 1.61E-24 | 0.916933725 | 0.976 | 0.959 | 3.17E-20 | Lep-T |
| cDC2 | ZNF581 | 1.23E-09 | 0.892380259 | 0.282 | 0.182 | 2.41E-05 | Lep-T |
| cDC2 | PLAUR | 4.58E-37 | 0.881397051 | 0.924 | 0.913 | 9.02E-33 | Lep-T |
| cDC2 | SERPING1 | 2.72E-10 | 0.871243311 | 0.442 | 0.349 | 5.35E-06 | Lep-T |
| cDC2 | MGLL1 | 3.58E-15 | 0.870080092 | 0.348 | 0.204 | 7.04E-11 | Lep-T |
| cDC2 | ITGB2-AS1 | 9.20E-10 | 0.86504648 | 0.291 | 0.19 | 1.81E-05 | Lep-T |
| cDC2 | TRAF11 | 5.26E-28 | 0.858395445 | 0.787 | 0.653 | 1.04E-23 | Lep-T |
| cDC2 | FSCN1 | 1.21E-27 | 0.857601476 | 0.664 | 0.452 | 2.39E-23 | Lep-T |
| cDC2 | NINJ11 | 2.67E-18 | 0.852214025 | 0.795 | 0.71 | 5.26E-14 | Lep-T |
| cDC2 | ERGIC3 | 2.17E-13 | 0.843132937 | 0.476 | 0.361 | 4.28E-09 | Lep-T |
| cDC2 | ALAS1 | 1.23E-08 | 0.840276347 | 0.424 | 0.328 | 0.000242629 | Lep-T |
| cDC2 | LILRB21 | 8.21E-11 | 0.839135247 | 0.431 | 0.319 | 1.62E-06 | Lep-T |
| cDC2 | ATP5PB | 1.62E-15 | 0.836780892 | 0.604 | 0.517 | 3.20E-11 | Lep-T |
| cDC2 | PLD4 | 2.82E-09 | 0.818418387 | 0.311 | 0.206 | 5.55E-05 | Lep-T |
| cDC2 | CST3 | 7.55E-44 | 0.811728344 | 0.994 | 0.984 | 1.49E-39 | Lep-T |
| cDC2 | INSIG11 | 6.29E-24 | 0.802814037 | 0.871 | 0.817 | 1.24E-19 | Lep-T |
| cDC2 | MIR155HG1 | 2.23E-25 | 0.799753904 | 0.712 | 0.518 | 4.39E-21 | Lep-T |
| cDC2 | TSPAN331 | 6.81E-17 | 0.794837666 | 0.705 | 0.621 | 1.34E-12 | Lep-T |

| | | | | | | | |
|---|---|---|---|---|---|---|---|
| cDC2 | IL1RN | 1.04E-09 | 0.793360084 | 0.639 | 0.574 | 2.04E-05 | Lep-T |
| cDC2 | TNFAIP31 | 1.09E-33 | 0.789346465 | 0.921 | 0.829 | 2.14E-29 | Lep-T |
| cDC2 | BIRC31 | 1.47E-25 | 0.782216841 | 0.802 | 0.675 | 2.89E-21 | Lep-T |
| cDC2 | BCL2A1 | 1.55E-21 | 0.780761394 | 0.918 | 0.882 | 3.06E-17 | Lep-T |
| cDC2 | GEM | 1.08E-08 | 0.775326958 | 0.516 | 0.427 | 0.000213305 | Lep-T |
| cDC2 | CD1E | 4.23E-19 | 0.774781308 | 0.56 | 0.377 | 8.32E-15 | Lep-T |
| cDC2 | CRTAP1 | 8.45E-13 | 0.755477744 | 0.548 | 0.453 | 1.66E-08 | Lep-T |
| cDC2 | ZFP36L11 | 4.45E-34 | 0.751845193 | 0.878 | 0.793 | 8.76E-30 | Lep-T |
| cDC2 | PPP1R15A1 | 3.72E-28 | 0.75086918 | 0.953 | 0.925 | 7.32E-24 | Lep-T |
| cDC2 | IRF8 | 7.62E-24 | 0.747751501 | 0.805 | 0.696 | 1.50E-19 | Lep-T |
| cDC2 | TPRA11 | 1.85E-07 | 0.744273523 | 0.478 | 0.399 | 0.0036444 | Lep-T |
| cDC2 | SYNGR2 | 1.01E-19 | 0.738981199 | 0.803 | 0.785 | 1.98E-15 | Lep-T |
| cDC2 | ZYX | 1.63E-13 | 0.738002823 | 0.715 | 0.653 | 3.20E-09 | Lep-T |
| cDC2 | IMPDH2 | 1.54E-09 | 0.737134714 | 0.382 | 0.278 | 3.04E-05 | Lep-T |
| cDC2 | CXCL21 | 4.12E-07 | 0.736519297 | 0.514 | 0.429 | 0.008115751 | Lep-T |
| cDC2 | SEC13 | 7.92E-11 | 0.733379052 | 0.446 | 0.337 | 1.56E-06 | Lep-T |
| cDC2 | RASSF4 | 1.88E-17 | 0.702437322 | 0.692 | 0.598 | 3.71E-13 | Lep-T |
| cDC2 | SLC2A3 | 1.48E-19 | 0.699835503 | 0.831 | 0.789 | 2.90E-15 | Lep-T |
| cDC2 | AVPI1 | 4.48E-08 | 0.695786724 | 0.523 | 0.453 | 0.000881114 | Lep-T |
| cDC2 | MMP91 | 2.35E-16 | 0.680891514 | 0.548 | 0.391 | 4.63E-12 | Lep-T |
| cDC2 | PPDPF | 3.76E-07 | 0.679676463 | 0.728 | 0.718 | 0.007405431 | Lep-T |
| cDC2 | TAGLN2 | 9.33E-26 | 0.650123033 | 0.939 | 0.898 | 1.84E-21 | Lep-T |
| cDC2 | DUSP5 | 4.64E-18 | 0.638042295 | 0.841 | 0.749 | 9.14E-14 | Lep-T |
| cDC2 | IRF2BP2 | 1.48E-12 | 0.637883295 | 0.68 | 0.63 | 2.92E-08 | Lep-T |
| cDC2 | LMNA1 | 9.29E-21 | 0.635872396 | 0.887 | 0.818 | 1.83E-16 | Lep-T |
| cDC2 | PNPLA2 | 5.01E-07 | 0.634959858 | 0.297 | 0.214 | 0.009855327 | Lep-T |
| cDC2 | CD1C | 1.44E-18 | 0.634083471 | 0.706 | 0.538 | 2.83E-14 | Lep-T |
| cDC2 | GRASP | 5.52E-12 | 0.631641429 | 0.722 | 0.691 | 1.09E-07 | Lep-T |
| cDC2 | AFDN | 2.06E-08 | 0.625728517 | 0.453 | 0.364 | 0.000405966 | Lep-T |
| cDC2 | NFKBIE1 | 7.77E-09 | 0.623353612 | 0.66 | 0.61 | 0.00015291 | Lep-T |
| cDC2 | EEF21 | 6.35E-43 | 0.620556292 | 0.933 | 0.877 | 1.25E-38 | Lep-T |
| cDC2 | CNBP | 4.74E-20 | 0.619702364 | 0.847 | 0.798 | 9.33E-16 | Lep-T |
| cDC2 | CD83 | 4.49E-41 | 0.603124593 | 0.991 | 0.974 | 8.85E-37 | Lep-T |
| cDC2 | SAT11 | 1.24E-16 | 0.599045368 | 0.995 | 0.986 | 2.43E-12 | Lep-T |
| cDC2 | JUNB | 1.75E-09 | 0.591742082 | 0.975 | 0.954 | 3.45E-05 | Lep-T |
| cDC2 | GNB2 | 7.19E-13 | 0.588090318 | 0.744 | 0.724 | 1.41E-08 | Lep-T |
| cDC2 | DUSP2 | 3.72E-12 | 0.587520007 | 0.917 | 0.859 | 7.33E-08 | Lep-T |
| cDC2 | ZC3H12A1 | 8.40E-21 | 0.58742096 | 0.682 | 0.527 | 1.65E-16 | Lep-T |
| cDC2 | MAP2K31 | 5.78E-20 | 0.575079563 | 0.81 | 0.72 | 1.14E-15 | Lep-T |
| cDC2 | PIM3 | 3.29E-09 | 0.573738641 | 0.789 | 0.806 | 6.47E-05 | Lep-T |
| cDC2 | C5AR11 | 2.35E-07 | 0.572654472 | 0.341 | 0.257 | 0.004630664 | Lep-T |
| cDC2 | DAZAP2 | 1.89E-08 | 0.567227308 | 0.644 | 0.591 | 0.000372211 | Lep-T |
| cDC2 | PTTG11 | 9.52E-07 | 0.564031325 | 0.53 | 0.461 | 0.018747679 | Lep-T |
| cDC2 | PSAP | 5.56E-23 | 0.562790433 | 0.896 | 0.845 | 1.09E-18 | Lep-T |
| cDC2 | TIMP11 | 1.22E-13 | 0.562535199 | 0.903 | 0.868 | 2.40E-09 | Lep-T |
| cDC2 | DNAJB11 | 4.85E-09 | 0.558344101 | 0.917 | 0.904 | 9.55E-05 | Lep-T |
| cDC2 | BHLHE40 | 2.32E-14 | 0.558192267 | 0.85 | 0.819 | 4.56E-10 | Lep-T |
| cDC2 | SMAD31 | 1.74E-07 | 0.54496209 | 0.354 | 0.259 | 0.003425264 | Lep-T |
| cDC2 | NR4A1 | 3.27E-29 | 0.543541311 | 0.909 | 0.859 | 6.43E-25 | Lep-T |
| cDC2 | PLEK | 1.19E-18 | 0.536432853 | 0.932 | 0.908 | 2.34E-14 | Lep-T |
| cDC2 | NFKB21 | 1.26E-06 | 0.525419488 | 0.479 | 0.417 | 0.024793536 | Lep-T |
| cDC2 | SF3B2 | 1.37E-09 | 0.522244695 | 0.563 | 0.488 | 2.70E-05 | Lep-T |
| cDC2 | TXN | 8.90E-09 | 0.520730559 | 0.939 | 0.92 | 0.000175219 | Lep-T |
| cDC2 | CD55 | 3.04E-14 | 0.517291539 | 0.857 | 0.828 | 5.98E-10 | Lep-T |
| cDC2 | GBP5 | 5.20E-11 | 0.5149087 | 0.599 | 0.492 | 1.02E-06 | Lep-T |
| cDC2 | CDC42EP31 | 8.10E-15 | 0.503171717 | 0.66 | 0.527 | 1.59E-10 | Lep-T |

| | | | | | | | |
|---|---|---|---|---|---|---|---|
| cDC2 | HLA-DMB | 8.78E-19 | 0.5021027 | 0.865 | 0.846 | 1.73E-14 | Lep-T |
| cDC2 | EIF4E | 9.21E-07 | 0.501560606 | 0.618 | 0.546 | 0.018128237 | Lep-T |
| cDC2 | CEACAM4 | 1.51E-53 | 2.287498773 | 0.458 | 0.145 | 2.97E-49 | Mari |
| cDC2 | ANK3 | 5.55E-29 | 2.062251872 | 0.31 | 0.108 | 1.09E-24 | Mari |
| cDC2 | KRT161 | 1.65E-35 | 2.020192118 | 0.389 | 0.148 | 3.25E-31 | Mari |
| cDC2 | S100A91 | 4.14E-46 | 1.99412575 | 0.704 | 0.368 | 8.16E-42 | Mari |
| cDC2 | S100A81 | 9.15E-43 | 1.809410965 | 0.627 | 0.306 | 1.80E-38 | Mari |
| cDC2 | CCL3L1 | 7.73E-19 | 1.779293984 | 0.715 | 0.569 | 1.52E-14 | Mari |
| cDC2 | CD14 | 2.84E-26 | 1.774379941 | 0.514 | 0.285 | 5.60E-22 | Mari |
| cDC2 | HLA-DRB51 | 2.05E-35 | 1.771548305 | 0.861 | 0.788 | 4.03E-31 | Mari |
| cDC2 | KRT6B1 | 2.98E-26 | 1.751726988 | 0.275 | 0.093 | 5.86E-22 | Mari |
| cDC2 | PLAC81 | 3.78E-27 | 1.717131311 | 0.375 | 0.151 | 7.44E-23 | Mari |
| cDC2 | CCL4L2 | 1.29E-15 | 1.692740938 | 0.674 | 0.562 | 2.53E-11 | Mari |
| cDC2 | MSRB2 | 2.58E-48 | 1.648052549 | 0.553 | 0.227 | 5.07E-44 | Mari |
| cDC2 | MS4A4A | 6.39E-28 | 1.596182977 | 0.403 | 0.175 | 1.26E-23 | Mari |
| cDC2 | PPP1R14A1 | 2.72E-18 | 1.586654042 | 0.368 | 0.184 | 5.36E-14 | Mari |
| cDC2 | FCGR3A | 3.01E-16 | 1.528907472 | 0.338 | 0.175 | 5.92E-12 | Mari |
| cDC2 | FCGR1B | 4.90E-23 | 1.488085699 | 0.435 | 0.221 | 9.64E-19 | Mari |
| cDC2 | GNG10 | 2.82E-72 | 1.468721944 | 0.866 | 0.558 | 5.54E-68 | Mari |
| cDC2 | MSR1 | 1.44E-13 | 1.467512832 | 0.387 | 0.234 | 2.83E-09 | Mari |
| cDC2 | CCDC167 | 3.27E-27 | 1.444506673 | 0.407 | 0.184 | 6.44E-23 | Mari |
| cDC2 | TMEM163 | 5.85E-27 | 1.421226368 | 0.303 | 0.104 | 1.15E-22 | Mari |
| cDC2 | FCGR1A | 6.75E-28 | 1.393818403 | 0.593 | 0.35 | 1.33E-23 | Mari |
| cDC2 | GIMAP4 | 2.44E-16 | 1.376878159 | 0.368 | 0.199 | 4.79E-12 | Mari |
| cDC2 | GCA | 1.79E-56 | 1.346365756 | 0.833 | 0.511 | 3.53E-52 | Mari |
| cDC2 | VPS25 | 6.48E-21 | 1.33342876 | 0.382 | 0.188 | 1.28E-16 | Mari |
| cDC2 | POLE4 | 7.65E-40 | 1.315288755 | 0.65 | 0.367 | 1.51E-35 | Mari |
| cDC2 | AIF1 | 6.07E-85 | 1.309582335 | 0.968 | 0.893 | 1.19E-80 | Mari |
| cDC2 | CCDC200 | 6.75E-15 | 1.298454294 | 0.363 | 0.204 | 1.33E-10 | Mari |
| cDC2 | CD691 | 1.84E-39 | 1.29549377 | 0.755 | 0.472 | 3.62E-35 | Mari |
| cDC2 | TCTN3 | 3.59E-13 | 1.267676173 | 0.398 | 0.246 | 7.06E-09 | Mari |
| cDC2 | NDUFC2 | 7.26E-60 | 1.259814689 | 0.88 | 0.555 | 1.43E-55 | Mari |
| cDC2 | RWDD3 | 1.70E-15 | 1.253893304 | 0.294 | 0.144 | 3.35E-11 | Mari |
| cDC2 | TYMP1 | 1.24E-89 | 1.251429101 | 0.991 | 0.848 | 2.43E-85 | Mari |
| cDC2 | DTYMK | 1.09E-14 | 1.235381506 | 0.292 | 0.144 | 2.14E-10 | Mari |
| cDC2 | LILRA5 | 1.66E-16 | 1.226242428 | 0.252 | 0.102 | 3.27E-12 | Mari |
| cDC2 | PRELID1 | 3.20E-77 | 1.219637904 | 0.961 | 0.842 | 6.29E-73 | Mari |
| cDC2 | TIMM8B | 9.13E-39 | 1.2149768 | 0.701 | 0.412 | 1.80E-34 | Mari |
| cDC2 | VAMP5 | 5.55E-42 | 1.211066032 | 0.85 | 0.629 | 1.09E-37 | Mari |
| cDC2 | PLAU | 9.98E-17 | 1.19658307 | 0.491 | 0.31 | 1.96E-12 | Mari |
| cDC2 | DDX60L | 3.56E-13 | 1.195338367 | 0.287 | 0.144 | 7.02E-09 | Mari |
| cDC2 | KRT6A1 | 1.40E-14 | 1.188772897 | 0.289 | 0.147 | 2.76E-10 | Mari |
| cDC2 | COMMD8 | 2.07E-31 | 1.187630602 | 0.581 | 0.321 | 4.07E-27 | Mari |
| cDC2 | IFI30 | 5.79E-99 | 1.184581466 | 0.991 | 0.962 | 1.14E-94 | Mari |
| cDC2 | IER2 | 5.17E-43 | 1.183687837 | 0.954 | 0.841 | 1.02E-38 | Mari |
| cDC2 | ACTR6 | 2.36E-12 | 1.179852661 | 0.31 | 0.173 | 4.64E-08 | Mari |
| cDC2 | MRPL58 | 1.28E-18 | 1.171741146 | 0.38 | 0.196 | 2.52E-14 | Mari |
| cDC2 | SATB1-AS11 | 1.83E-23 | 1.169734618 | 0.5 | 0.265 | 3.60E-19 | Mari |
| cDC2 | UCHL3 | 4.57E-28 | 1.163011525 | 0.567 | 0.318 | 8.99E-24 | Mari |
| cDC2 | PCED1B-AS1 | 2.04E-11 | 1.161576492 | 0.262 | 0.138 | 4.01E-07 | Mari |
| cDC2 | PPA11 | 1.97E-45 | 1.159882893 | 0.954 | 0.776 | 3.87E-41 | Mari |
| cDC2 | SCO2 | 6.08E-42 | 1.153886627 | 0.792 | 0.486 | 1.20E-37 | Mari |
| cDC2 | LRRC25 | 1.58E-13 | 1.144697792 | 0.352 | 0.202 | 3.12E-09 | Mari |
| cDC2 | GTF3C6 | 6.65E-38 | 1.13608485 | 0.736 | 0.461 | 1.31E-33 | Mari |
| cDC2 | MGAT4A | 1.25E-10 | 1.125328772 | 0.278 | 0.157 | 2.45E-06 | Mari |
| cDC2 | MEF2C | 2.09E-45 | 1.124064636 | 0.856 | 0.584 | 4.12E-41 | Mari |

| | | | | | | | |
|---|---|---|---|---|---|---|---|
| cDC2 | ALG5 | 9.67E-22 | 1.121365726 | 0.468 | 0.259 | 1.90E-17 | Mari |
| cDC2 | TXNDC171 | 1.70E-43 | 1.118812804 | 0.785 | 0.492 | 3.35E-39 | Mari |
| cDC2 | IFITM3 | 1.06E-64 | 1.117210749 | 0.963 | 0.904 | 2.09E-60 | Mari |
| cDC2 | FCER1A | 2.13E-06 | 1.115434889 | 0.514 | 0.474 | 0.041915791 | Mari |
| cDC2 | FAM89B1 | 7.24E-29 | 1.112660586 | 0.674 | 0.425 | 1.43E-24 | Mari |
| cDC2 | 2-Mar | 9.15E-21 | 1.109672944 | 0.556 | 0.358 | 1.80E-16 | Mari |
| cDC2 | SDF2L1 | 2.57E-24 | 1.10954225 | 0.676 | 0.46 | 5.07E-20 | Mari |
| cDC2 | CYBC1 | 2.59E-26 | 1.107759705 | 0.602 | 0.362 | 5.09E-22 | Mari |
| cDC2 | MALSU1 | 5.25E-13 | 1.103420205 | 0.336 | 0.19 | 1.03E-08 | Mari |
| cDC2 | DCXR | 1.75E-35 | 1.100153306 | 0.708 | 0.438 | 3.45E-31 | Mari |
| cDC2 | ISCA21 | 8.62E-11 | 1.099081175 | 0.301 | 0.18 | 1.70E-06 | Mari |
| cDC2 | GALK1 | 1.03E-20 | 1.09903779 | 0.493 | 0.292 | 2.02E-16 | Mari |
| cDC2 | RBKS | 9.58E-30 | 1.094962086 | 0.637 | 0.373 | 1.89E-25 | Mari |
| cDC2 | RNASEH2C | 2.53E-23 | 1.090662047 | 0.546 | 0.325 | 4.99E-19 | Mari |
| cDC2 | MX1 | 2.27E-18 | 1.088927273 | 0.491 | 0.29 | 4.47E-14 | Mari |
| cDC2 | FCER1G | 2.94E-75 | 1.088466695 | 0.972 | 0.932 | 5.78E-71 | Mari |
| cDC2 | MNDA | 7.70E-51 | 1.082664594 | 0.912 | 0.681 | 1.52E-46 | Mari |
| cDC2 | SEC61G | 1.22E-55 | 1.079145522 | 0.926 | 0.721 | 2.40E-51 | Mari |
| cDC2 | TMEM126A | 4.13E-23 | 1.078777288 | 0.528 | 0.31 | 8.13E-19 | Mari |
| cDC2 | S100B | 1.73E-14 | 1.075712241 | 0.435 | 0.25 | 3.41E-10 | Mari |
| cDC2 | SRP9 | 9.60E-44 | 1.075425339 | 0.806 | 0.538 | 1.89E-39 | Mari |
| cDC2 | CRELD2 | 4.17E-18 | 1.07476494 | 0.47 | 0.276 | 8.21E-14 | Mari |
| cDC2 | SNAPC51 | 4.17E-07 | 1.073174258 | 0.252 | 0.164 | 0.008200608 | Mari |
| cDC2 | HPS3 | 4.82E-15 | 1.07017594 | 0.47 | 0.296 | 9.50E-11 | Mari |
| cDC2 | ZSWIM7 | 9.03E-26 | 1.068238901 | 0.611 | 0.384 | 1.78E-21 | Mari |
| cDC2 | CIB1 | 4.10E-58 | 1.063044256 | 0.94 | 0.754 | 8.06E-54 | Mari |
| cDC2 | SLC35B1 | 2.72E-16 | 1.061837445 | 0.414 | 0.238 | 5.36E-12 | Mari |
| cDC2 | ZNF511 | 3.34E-18 | 1.061608479 | 0.463 | 0.279 | 6.58E-14 | Mari |
| cDC2 | GAS6 | 1.56E-34 | 1.061536928 | 0.757 | 0.512 | 3.08E-30 | Mari |
| cDC2 | COPZ1 | 8.36E-25 | 1.06116556 | 0.586 | 0.364 | 1.64E-20 | Mari |
| cDC2 | NIPSNAP3A | 1.93E-08 | 1.060601968 | 0.294 | 0.185 | 0.000380483 | Mari |
| cDC2 | IL10RB-DT | 2.55E-10 | 1.053769458 | 0.262 | 0.142 | 5.01E-06 | Mari |
| cDC2 | NDUFB2 | 4.33E-68 | 1.05113834 | 0.951 | 0.73 | 8.52E-64 | Mari |
| cDC2 | MRPS231 | 8.01E-22 | 1.042222241 | 0.477 | 0.259 | 1.58E-17 | Mari |
| cDC2 | ODC1 | 4.85E-10 | 1.04170579 | 0.424 | 0.301 | 9.55E-06 | Mari |
| cDC2 | EXOSC3 | 9.60E-10 | 1.03840341 | 0.285 | 0.168 | 1.89E-05 | Mari |
| cDC2 | BORCS8 | 9.87E-11 | 1.037371817 | 0.299 | 0.176 | 1.94E-06 | Mari |
| cDC2 | LAMTOR1 | 4.87E-49 | 1.037361432 | 0.882 | 0.662 | 9.59E-45 | Mari |
| cDC2 | CHCHD1 | 6.73E-20 | 1.033418019 | 0.509 | 0.311 | 1.32E-15 | Mari |
| cDC2 | UCP2 | 1.34E-42 | 1.033187961 | 0.875 | 0.657 | 2.64E-38 | Mari |
| cDC2 | ASGR1 | 1.92E-10 | 1.031274261 | 0.269 | 0.145 | 3.77E-06 | Mari |
| cDC2 | RGS18 | 2.03E-15 | 1.030539122 | 0.435 | 0.262 | 3.99E-11 | Mari |
| cDC2 | IMPA2 | 6.70E-13 | 1.029937734 | 0.303 | 0.161 | 1.32E-08 | Mari |
| cDC2 | ARL2 | 9.92E-18 | 1.029253695 | 0.472 | 0.291 | 1.95E-13 | Mari |
| cDC2 | OSM | 5.05E-11 | 1.027607834 | 0.37 | 0.231 | 9.94E-07 | Mari |
| cDC2 | ETFB | 2.57E-33 | 1.02715361 | 0.701 | 0.424 | 5.05E-29 | Mari |
| cDC2 | UQCR11 | 1.38E-71 | 1.025525342 | 0.954 | 0.771 | 2.72E-67 | Mari |
| cDC2 | TM2D2 | 2.34E-15 | 1.025508694 | 0.456 | 0.275 | 4.60E-11 | Mari |
| cDC2 | LDLRAD3 | 1.99E-08 | 1.025339267 | 0.273 | 0.163 | 0.000392458 | Mari |
| cDC2 | HIGD1A | 1.11E-20 | 1.020036647 | 0.512 | 0.302 | 2.19E-16 | Mari |
| cDC2 | LSM10 | 9.00E-23 | 1.019747286 | 0.609 | 0.396 | 1.77E-18 | Mari |
| cDC2 | MRPS10 | 4.02E-13 | 1.018218723 | 0.384 | 0.235 | 7.91E-09 | Mari |
| cDC2 | PTPMT1 | 1.96E-16 | 1.017282313 | 0.424 | 0.249 | 3.85E-12 | Mari |
| cDC2 | SDSL | 1.40E-12 | 1.016732702 | 0.324 | 0.18 | 2.75E-08 | Mari |
| cDC2 | ABI31 | 3.27E-24 | 1.015642077 | 0.62 | 0.387 | 6.45E-20 | Mari |
| cDC2 | MRPL28 | 4.84E-21 | 1.01540937 | 0.502 | 0.289 | 9.52E-17 | Mari |

| | | | | | | | |
|---|---|---|---|---|---|---|---|
| cDC2 | YIF1A | 7.08E-18 | 1.014670442 | 0.514 | 0.32 | 1.39E-13 | Mari |
| cDC2 | PDHB | 6.45E-15 | 1.012987252 | 0.403 | 0.234 | 1.27E-10 | Mari |
| cDC2 | C1QB | 9.32E-07 | 1.008132562 | 0.62 | 0.593 | 0.018343425 | Mari |
| cDC2 | NDUFAF8 | 3.62E-19 | 1.007289714 | 0.488 | 0.289 | 7.12E-15 | Mari |
| cDC2 | CHMP2A1 | 8.86E-41 | 1.005304862 | 0.808 | 0.539 | 1.74E-36 | Mari |
| cDC2 | SFT2D1 | 4.48E-36 | 0.997957967 | 0.778 | 0.513 | 8.83E-32 | Mari |
| cDC2 | THOC7 | 2.91E-28 | 0.997867087 | 0.657 | 0.417 | 5.73E-24 | Mari |
| cDC2 | SMCO4 | 4.68E-36 | 0.995294711 | 0.833 | 0.581 | 9.21E-32 | Mari |
| cDC2 | FOS | 5.59E-49 | 0.993690917 | 0.956 | 0.823 | 1.10E-44 | Mari |
| cDC2 | DYNLT1 | 5.85E-36 | 0.993442475 | 0.743 | 0.469 | 1.15E-31 | Mari |
| cDC2 | GSDMD | 5.44E-19 | 0.993329304 | 0.553 | 0.359 | 1.07E-14 | Mari |
| cDC2 | FDPS | 6.34E-20 | 0.990641485 | 0.544 | 0.337 | 1.25E-15 | Mari |
| cDC2 | LSM6 | 8.25E-29 | 0.990029678 | 0.701 | 0.453 | 1.62E-24 | Mari |
| cDC2 | KRT141 | 3.96E-14 | 0.989733042 | 0.461 | 0.319 | 7.79E-10 | Mari |
| cDC2 | MRPL20 | 4.51E-31 | 0.989619993 | 0.771 | 0.514 | 8.87E-27 | Mari |
| cDC2 | NOL12 | 5.17E-10 | 0.988989351 | 0.306 | 0.182 | 1.02E-05 | Mari |
| cDC2 | SNF8 | 1.97E-33 | 0.988964048 | 0.755 | 0.491 | 3.87E-29 | Mari |
| cDC2 | ASCL2 | 2.39E-11 | 0.988514812 | 0.34 | 0.206 | 4.71E-07 | Mari |
| cDC2 | NMI | 5.52E-26 | 0.988096472 | 0.653 | 0.412 | 1.09E-21 | Mari |
| cDC2 | IER3IP1 | 5.00E-20 | 0.987787649 | 0.505 | 0.299 | 9.84E-16 | Mari |
| cDC2 | TRAPPC1 | 1.55E-38 | 0.986793235 | 0.808 | 0.531 | 3.05E-34 | Mari |
| cDC2 | CALHM6 | 2.42E-36 | 0.986493932 | 0.907 | 0.776 | 4.76E-32 | Mari |
| cDC2 | YDJC | 1.35E-12 | 0.985189227 | 0.361 | 0.209 | 2.66E-08 | Mari |
| cDC2 | BST2 | 1.69E-47 | 0.984178433 | 0.891 | 0.707 | 3.32E-43 | Mari |
| cDC2 | ZBTB8OS | 1.61E-14 | 0.9838414 | 0.403 | 0.239 | 3.16E-10 | Mari |
| cDC2 | CYBA | 1.73E-105 | 0.982042527 | 0.988 | 0.963 | 3.41E-101 | Mari |
| cDC2 | CKLF1 | 2.95E-39 | 0.980239474 | 0.903 | 0.694 | 5.80E-35 | Mari |
| cDC2 | TRABD | 3.46E-22 | 0.97940534 | 0.653 | 0.426 | 6.82E-18 | Mari |
| cDC2 | NUBP2 | 4.04E-13 | 0.97778602 | 0.391 | 0.241 | 7.95E-09 | Mari |
| cDC2 | JOSD2 | 9.14E-18 | 0.977709946 | 0.532 | 0.346 | 1.80E-13 | Mari |
| cDC2 | GMFG | 4.96E-56 | 0.977600056 | 0.924 | 0.736 | 9.77E-52 | Mari |
| cDC2 | COA5 | 4.67E-07 | 0.973913038 | 0.285 | 0.192 | 0.009184703 | Mari |
| cDC2 | CCNDBP1 | 8.85E-16 | 0.973827175 | 0.463 | 0.291 | 1.74E-11 | Mari |
| cDC2 | COPS7A | 1.32E-07 | 0.968843098 | 0.25 | 0.155 | 0.002591781 | Mari |
| cDC2 | POP4 | 3.41E-15 | 0.967013954 | 0.396 | 0.227 | 6.72E-11 | Mari |
| cDC2 | NDUFB7 | 9.13E-38 | 0.966839142 | 0.806 | 0.571 | 1.80E-33 | Mari |
| cDC2 | ARHGDIB | 6.07E-59 | 0.966118662 | 0.954 | 0.86 | 1.19E-54 | Mari |
| cDC2 | CLEC12A | 2.57E-08 | 0.965091682 | 0.329 | 0.214 | 0.000505373 | Mari |
| cDC2 | BLOC1S1 | 4.47E-57 | 0.964564213 | 0.944 | 0.744 | 8.81E-53 | Mari |
| cDC2 | MYL12A | 4.80E-65 | 0.964028314 | 0.975 | 0.925 | 9.45E-61 | Mari |
| cDC2 | POLR2I | 2.23E-19 | 0.95853604 | 0.514 | 0.31 | 4.39E-15 | Mari |
| cDC2 | NUDT22 | 4.34E-14 | 0.957325154 | 0.481 | 0.325 | 8.54E-10 | Mari |
| cDC2 | BLOC1S2 | 4.24E-19 | 0.956847201 | 0.586 | 0.387 | 8.35E-15 | Mari |
| cDC2 | COMMD7 | 7.06E-22 | 0.953921477 | 0.562 | 0.351 | 1.39E-17 | Mari |
| cDC2 | MRPL19 | 1.52E-09 | 0.953154684 | 0.322 | 0.203 | 3.00E-05 | Mari |
| cDC2 | METTL23 | 3.58E-17 | 0.952801821 | 0.493 | 0.302 | 7.05E-13 | Mari |
| cDC2 | SDHAF3 | 7.12E-10 | 0.952579886 | 0.255 | 0.14 | 1.40E-05 | Mari |
| cDC2 | SSU72 | 5.46E-26 | 0.949868139 | 0.708 | 0.505 | 1.08E-21 | Mari |
| cDC2 | TOR2A | 2.06E-07 | 0.949127597 | 0.287 | 0.19 | 0.004045336 | Mari |
| cDC2 | METTL5 | 3.39E-19 | 0.942660029 | 0.505 | 0.308 | 6.67E-15 | Mari |
| cDC2 | ANAPC15 | 1.25E-15 | 0.942288108 | 0.44 | 0.26 | 2.47E-11 | Mari |
| cDC2 | PRDX5 | 2.48E-44 | 0.942129387 | 0.873 | 0.654 | 4.89E-40 | Mari |
| cDC2 | GOSR2 | 1.58E-07 | 0.941339202 | 0.282 | 0.183 | 0.00310523 | Mari |
| cDC2 | ASL | 2.69E-10 | 0.941244314 | 0.336 | 0.205 | 5.30E-06 | Mari |
| cDC2 | MED8 | 1.85E-14 | 0.938919711 | 0.472 | 0.296 | 3.65E-10 | Mari |
| cDC2 | IFI44 | 7.92E-08 | 0.938751281 | 0.252 | 0.149 | 0.001558363 | Mari |

| | | | | | | | |
|---|---|---|---|---|---|---|---|
| cDC2 | TRIAP1 | 8.43E-10 | 0.935900517 | 0.294 | 0.172 | 1.66E-05 | Mari |
| cDC2 | UBE2V2 | 6.11E-21 | 0.934238096 | 0.514 | 0.294 | 1.20E-16 | Mari |
| cDC2 | MTLN | 3.17E-10 | 0.93301589 | 0.35 | 0.227 | 6.24E-06 | Mari |
| cDC2 | MRPS18C | 9.55E-19 | 0.932164307 | 0.521 | 0.318 | 1.88E-14 | Mari |
| cDC2 | RNF135 | 1.90E-08 | 0.922304009 | 0.275 | 0.169 | 0.000373392 | Mari |
| cDC2 | MRPL131 | 4.95E-19 | 0.920700751 | 0.505 | 0.308 | 9.75E-15 | Mari |
| cDC2 | SPCS3 | 2.90E-32 | 0.920601056 | 0.803 | 0.561 | 5.71E-28 | Mari |
| cDC2 | LSM5 | 3.85E-31 | 0.920550352 | 0.738 | 0.494 | 7.58E-27 | Mari |
| cDC2 | LLPH | 1.06E-13 | 0.919930638 | 0.424 | 0.266 | 2.08E-09 | Mari |
| cDC2 | SMIM7 | 2.50E-17 | 0.916100623 | 0.553 | 0.367 | 4.92E-13 | Mari |
| cDC2 | TNFSF101 | 1.70E-25 | 0.916005183 | 0.759 | 0.541 | 3.34E-21 | Mari |
| cDC2 | MAGOHB | 3.94E-08 | 0.915581942 | 0.312 | 0.204 | 0.000775464 | Mari |
| cDC2 | LYZ | 4.82E-34 | 0.915469812 | 0.991 | 0.966 | 9.48E-30 | Mari |
| cDC2 | ATP5MF | 1.36E-59 | 0.915379204 | 0.949 | 0.776 | 2.68E-55 | Mari |
| cDC2 | GTF2H5 | 4.27E-15 | 0.913936981 | 0.509 | 0.345 | 8.41E-11 | Mari |
| cDC2 | PIN1 | 2.39E-17 | 0.912760381 | 0.581 | 0.391 | 4.70E-13 | Mari |
| cDC2 | NDUFAB11 | 1.80E-33 | 0.91218371 | 0.801 | 0.547 | 3.54E-29 | Mari |
| cDC2 | MRPL27 | 5.95E-14 | 0.911815167 | 0.414 | 0.255 | 1.17E-09 | Mari |
| cDC2 | CZIB | 1.01E-11 | 0.90939018 | 0.359 | 0.217 | 1.99E-07 | Mari |
| cDC2 | FAM207A | 2.59E-09 | 0.906765797 | 0.324 | 0.204 | 5.10E-05 | Mari |
| cDC2 | NCBP2AS2 | 1.84E-09 | 0.903414422 | 0.336 | 0.215 | 3.61E-05 | Mari |
| cDC2 | MS4A7 | 4.39E-25 | 0.902637031 | 0.706 | 0.467 | 8.64E-21 | Mari |
| cDC2 | MVB12A | 2.55E-10 | 0.901236265 | 0.324 | 0.195 | 5.01E-06 | Mari |
| cDC2 | CYB561D2 | 1.79E-14 | 0.900383148 | 0.38 | 0.218 | 3.52E-10 | Mari |
| cDC2 | PSMA4 | 3.26E-34 | 0.899131838 | 0.85 | 0.622 | 6.41E-30 | Mari |
| cDC2 | CALCRL | 1.35E-15 | 0.898001167 | 0.414 | 0.234 | 2.65E-11 | Mari |
| cDC2 | RBX1 | 1.23E-50 | 0.897811259 | 0.926 | 0.72 | 2.41E-46 | Mari |
| cDC2 | SVBP | 4.74E-16 | 0.897653526 | 0.44 | 0.259 | 9.34E-12 | Mari |
| cDC2 | TSEN15 | 3.77E-09 | 0.89644434 | 0.271 | 0.154 | 7.41E-05 | Mari |
| cDC2 | ANTKMT | 4.99E-16 | 0.896064284 | 0.519 | 0.337 | 9.83E-12 | Mari |
| cDC2 | PET100 | 5.73E-29 | 0.894774161 | 0.782 | 0.552 | 1.13E-24 | Mari |
| cDC2 | COPS9 | 4.18E-35 | 0.894527176 | 0.826 | 0.547 | 8.22E-31 | Mari |
| cDC2 | PLSCR1 | 2.75E-47 | 0.894440916 | 0.968 | 0.746 | 5.41E-43 | Mari |
| cDC2 | FPR1 | 1.67E-26 | 0.89341442 | 0.771 | 0.514 | 3.28E-22 | Mari |
| cDC2 | H3F3B | 4.86E-58 | 0.892966162 | 0.995 | 0.988 | 9.57E-54 | Mari |
| cDC2 | ARPC1B | 7.62E-71 | 0.888264059 | 0.963 | 0.886 | 1.50E-66 | Mari |
| cDC2 | S100A11 | 3.05E-85 | 0.887035078 | 0.986 | 0.971 | 6.00E-81 | Mari |
| cDC2 | IFITM2 | 4.84E-36 | 0.882994195 | 0.898 | 0.745 | 9.52E-32 | Mari |
| cDC2 | THYN1 | 1.35E-11 | 0.88152152 | 0.384 | 0.237 | 2.65E-07 | Mari |
| cDC2 | TIMM17B | 2.10E-13 | 0.881156868 | 0.512 | 0.346 | 4.13E-09 | Mari |
| cDC2 | SLC50A1 | 5.53E-16 | 0.879227794 | 0.493 | 0.314 | 1.09E-11 | Mari |
| cDC2 | CORO1A | 2.43E-41 | 0.878405963 | 0.942 | 0.737 | 4.79E-37 | Mari |
| cDC2 | P2RY131 | 3.41E-16 | 0.875993916 | 0.502 | 0.305 | 6.70E-12 | Mari |
| cDC2 | DUSP23 | 6.18E-19 | 0.874382481 | 0.579 | 0.387 | 1.22E-14 | Mari |
| cDC2 | TRADD | 8.01E-15 | 0.874027877 | 0.525 | 0.36 | 1.58E-10 | Mari |
| cDC2 | ADPRH | 5.01E-08 | 0.872960287 | 0.299 | 0.189 | 0.000986302 | Mari |
| cDC2 | NDUFA13 | 3.10E-62 | 0.872911975 | 0.968 | 0.807 | 6.11E-58 | Mari |
| cDC2 | EIF4EBP1 | 1.88E-21 | 0.871660083 | 0.662 | 0.453 | 3.71E-17 | Mari |
| cDC2 | CHMP4A | 6.67E-30 | 0.871556344 | 0.752 | 0.506 | 1.31E-25 | Mari |
| cDC2 | LYL1 | 5.05E-10 | 0.871274306 | 0.382 | 0.248 | 9.94E-06 | Mari |
| cDC2 | NUDT1 | 1.11E-26 | 0.871141931 | 0.685 | 0.442 | 2.18E-22 | Mari |
| cDC2 | ADAM28 | 1.17E-12 | 0.868223765 | 0.472 | 0.301 | 2.30E-08 | Mari |
| cDC2 | TXNDC9 | 1.14E-10 | 0.867784141 | 0.384 | 0.248 | 2.25E-06 | Mari |
| cDC2 | SCIMP1 | 2.04E-15 | 0.867118531 | 0.47 | 0.29 | 4.02E-11 | Mari |
| cDC2 | PARK7 | 7.02E-50 | 0.866416556 | 0.933 | 0.782 | 1.38E-45 | Mari |
| cDC2 | TP53TG1 | 2.88E-09 | 0.864150192 | 0.28 | 0.161 | 5.66E-05 | Mari |

| | | | | | | | |
|---|---|---|---|---|---|---|---|
| cDC2 | XRCC41 | 5.13E-08 | 0.863485857 | 0.273 | 0.167 | 0.00100915 | Mari |
| cDC2 | BOLA2 | 7.02E-15 | 0.860261524 | 0.454 | 0.275 | 1.38E-10 | Mari |
| cDC2 | CENPX | 1.53E-23 | 0.859985131 | 0.662 | 0.442 | 3.00E-19 | Mari |
| cDC2 | ID3 | 2.91E-10 | 0.858482654 | 0.576 | 0.435 | 5.73E-06 | Mari |
| cDC2 | PLAAT4 | 1.29E-18 | 0.856168205 | 0.697 | 0.552 | 2.54E-14 | Mari |
| cDC2 | DGUOK | 1.07E-26 | 0.854594424 | 0.736 | 0.496 | 2.10E-22 | Mari |
| cDC2 | TMEM141 | 2.86E-14 | 0.854199431 | 0.481 | 0.313 | 5.62E-10 | Mari |
| cDC2 | PSME2 | 2.42E-48 | 0.853737943 | 0.968 | 0.883 | 4.77E-44 | Mari |
| cDC2 | ERI3 | 1.64E-09 | 0.853249439 | 0.324 | 0.202 | 3.24E-05 | Mari |
| cDC2 | JPT1 | 7.79E-29 | 0.850861112 | 0.785 | 0.554 | 1.53E-24 | Mari |
| cDC2 | NDUFA4 | 1.76E-61 | 0.850840068 | 0.954 | 0.863 | 3.47E-57 | Mari |
| cDC2 | TMEM50B | 7.48E-10 | 0.848942554 | 0.338 | 0.208 | 1.47E-05 | Mari |
| cDC2 | UBLCP1 | 1.24E-08 | 0.848613452 | 0.336 | 0.219 | 0.000244764 | Mari |
| cDC2 | AP1S1 | 1.38E-07 | 0.848478343 | 0.315 | 0.213 | 0.002712182 | Mari |
| cDC2 | EVI2A | 8.06E-10 | 0.846798847 | 0.556 | 0.408 | 1.59E-05 | Mari |
| cDC2 | NME4 | 9.87E-11 | 0.846274698 | 0.384 | 0.241 | 1.94E-06 | Mari |
| cDC2 | DERA | 3.81E-15 | 0.846257156 | 0.47 | 0.287 | 7.50E-11 | Mari |
| cDC2 | MIA2 | 2.62E-08 | 0.845398884 | 0.359 | 0.245 | 0.000515328 | Mari |
| cDC2 | IFI27L2 | 1.88E-21 | 0.843937429 | 0.669 | 0.45 | 3.70E-17 | Mari |
| cDC2 | SLC25A39 | 2.00E-13 | 0.842620854 | 0.486 | 0.327 | 3.94E-09 | Mari |
| cDC2 | PSMB2 | 1.90E-24 | 0.841723947 | 0.757 | 0.548 | 3.74E-20 | Mari |
| cDC2 | UNC50 | 2.15E-10 | 0.841640842 | 0.352 | 0.221 | 4.23E-06 | Mari |
| cDC2 | SNX17 | 3.50E-21 | 0.841461153 | 0.671 | 0.452 | 6.90E-17 | Mari |
| cDC2 | RGS19 | 2.55E-27 | 0.840688694 | 0.785 | 0.556 | 5.02E-23 | Mari |
| cDC2 | HRH2 | 3.29E-09 | 0.84006589 | 0.338 | 0.218 | 6.49E-05 | Mari |
| cDC2 | NDUFS6 | 5.64E-33 | 0.839200782 | 0.859 | 0.631 | 1.11E-28 | Mari |
| cDC2 | PSMB9 | 1.10E-45 | 0.837682572 | 0.933 | 0.822 | 2.17E-41 | Mari |
| cDC2 | CHCHD5 | 7.36E-19 | 0.836473115 | 0.616 | 0.403 | 1.45E-14 | Mari |
| cDC2 | CASP4 | 8.75E-33 | 0.835440574 | 0.843 | 0.645 | 1.72E-28 | Mari |
| cDC2 | CDC123 | 8.51E-15 | 0.834510474 | 0.519 | 0.348 | 1.67E-10 | Mari |
| cDC2 | CCND3 | 1.38E-15 | 0.833528343 | 0.606 | 0.413 | 2.73E-11 | Mari |
| cDC2 | MCRS1 | 1.12E-06 | 0.832728941 | 0.292 | 0.198 | 0.021958308 | Mari |
| cDC2 | TOMM5 | 1.15E-23 | 0.832273614 | 0.681 | 0.447 | 2.27E-19 | Mari |
| cDC2 | NHP2 | 1.73E-17 | 0.831852996 | 0.63 | 0.45 | 3.40E-13 | Mari |
| cDC2 | TSSC4 | 1.21E-06 | 0.830869291 | 0.306 | 0.21 | 0.023748808 | Mari |
| cDC2 | LAMTOR2 | 1.37E-30 | 0.829227365 | 0.826 | 0.624 | 2.70E-26 | Mari |
| cDC2 | HINT2 | 1.93E-10 | 0.828796356 | 0.428 | 0.297 | 3.81E-06 | Mari |
| cDC2 | MRPL23 | 3.22E-16 | 0.827873857 | 0.537 | 0.365 | 6.34E-12 | Mari |
| cDC2 | SLIRP | 2.49E-22 | 0.827601132 | 0.688 | 0.468 | 4.90E-18 | Mari |
| cDC2 | SCAMP2 | 3.58E-15 | 0.826692179 | 0.558 | 0.397 | 7.05E-11 | Mari |
| cDC2 | TMEM230 | 3.23E-22 | 0.826552815 | 0.697 | 0.481 | 6.36E-18 | Mari |
| cDC2 | SUMO3 | 2.15E-22 | 0.82651017 | 0.736 | 0.525 | 4.24E-18 | Mari |
| cDC2 | TNFSF13 | 6.97E-13 | 0.824736355 | 0.519 | 0.356 | 1.37E-08 | Mari |
| cDC2 | C1D | 5.64E-15 | 0.824092075 | 0.525 | 0.355 | 1.11E-10 | Mari |
| cDC2 | GMFB | 8.55E-08 | 0.823828323 | 0.28 | 0.18 | 0.001683247 | Mari |
| cDC2 | PAFAH1B3 | 2.94E-10 | 0.823627792 | 0.384 | 0.249 | 5.78E-06 | Mari |
| cDC2 | CHD9 | 6.02E-20 | 0.823605516 | 0.632 | 0.431 | 1.18E-15 | Mari |
| cDC2 | UQCR10 | 5.39E-38 | 0.822285342 | 0.903 | 0.694 | 1.06E-33 | Mari |
| cDC2 | DTWD1 | 3.36E-07 | 0.821575646 | 0.259 | 0.162 | 0.006620003 | Mari |
| cDC2 | YWHAH1 | 3.31E-23 | 0.820601348 | 0.806 | 0.584 | 6.51E-19 | Mari |
| cDC2 | PSMD13 | 4.90E-17 | 0.819112748 | 0.63 | 0.45 | 9.64E-13 | Mari |
| cDC2 | PEX16 | 3.47E-08 | 0.818821875 | 0.336 | 0.225 | 0.000683853 | Mari |
| cDC2 | MRPL3 | 3.58E-15 | 0.818651127 | 0.56 | 0.381 | 7.05E-11 | Mari |
| cDC2 | AKR7A2 | 2.93E-11 | 0.818183769 | 0.477 | 0.337 | 5.76E-07 | Mari |
| cDC2 | RPP30 | 1.81E-07 | 0.815456189 | 0.326 | 0.219 | 0.003562367 | Mari |
| cDC2 | MS4A6A | 1.78E-15 | 0.81510727 | 0.773 | 0.712 | 3.51E-11 | Mari |

| | | | | | | | |
|---|---|---|---|---|---|---|---|
| cDC2 | HSBP1 | 1.35E-29 | 0.814645283 | 0.824 | 0.632 | 2.66E-25 | Mari |
| cDC2 | PSMA5 | 6.43E-30 | 0.814442609 | 0.838 | 0.647 | 1.27E-25 | Mari |
| cDC2 | UQCC2 | 2.04E-11 | 0.813871879 | 0.454 | 0.305 | 4.02E-07 | Mari |
| cDC2 | ATP5ME | 3.23E-36 | 0.8122198 | 0.91 | 0.699 | 6.36E-32 | Mari |
| cDC2 | POMP | 4.58E-48 | 0.811714989 | 0.94 | 0.858 | 9.02E-44 | Mari |
| cDC2 | SSBP1 | 2.16E-32 | 0.811484969 | 0.85 | 0.628 | 4.25E-28 | Mari |
| cDC2 | PARVG | 1.13E-19 | 0.810745942 | 0.641 | 0.423 | 2.23E-15 | Mari |
| cDC2 | PPCDC1 | 6.50E-11 | 0.81044538 | 0.322 | 0.185 | 1.28E-06 | Mari |
| cDC2 | VBP1 | 2.53E-12 | 0.810350681 | 0.472 | 0.309 | 4.98E-08 | Mari |
| cDC2 | HES4 | 3.21E-10 | 0.809935734 | 0.44 | 0.294 | 6.33E-06 | Mari |
| cDC2 | NDUFA3 | 4.44E-26 | 0.809414683 | 0.741 | 0.51 | 8.75E-22 | Mari |
| cDC2 | VAMP8 | 4.79E-52 | 0.808719513 | 0.949 | 0.86 | 9.43E-48 | Mari |
| cDC2 | RER1 | 2.41E-25 | 0.807190492 | 0.743 | 0.536 | 4.74E-21 | Mari |
| cDC2 | MEIKIN1 | 3.51E-07 | 0.806486314 | 0.271 | 0.166 | 0.006903924 | Mari |
| cDC2 | CFL1 | 1.33E-77 | 0.806215533 | 0.979 | 0.964 | 2.62E-73 | Mari |
| cDC2 | TBCB | 4.49E-26 | 0.805861173 | 0.775 | 0.548 | 8.83E-22 | Mari |
| cDC2 | SELENOH | 1.03E-33 | 0.805022211 | 0.903 | 0.71 | 2.03E-29 | Mari |
| cDC2 | NSMCE3 | 5.65E-08 | 0.804328978 | 0.333 | 0.219 | 0.001111847 | Mari |
| cDC2 | SRBD11 | 1.71E-06 | 0.804024201 | 0.287 | 0.193 | 0.033751727 | Mari |
| cDC2 | OSTC | 8.00E-34 | 0.803704074 | 0.861 | 0.689 | 1.57E-29 | Mari |
| cDC2 | DCPS | 2.51E-06 | 0.803040039 | 0.282 | 0.193 | 0.049330663 | Mari |
| cDC2 | LYSMD2 | 3.19E-24 | 0.803009513 | 0.796 | 0.577 | 6.27E-20 | Mari |
| cDC2 | TSTA3 | 3.58E-08 | 0.802589863 | 0.359 | 0.246 | 0.00070521 | Mari |
| cDC2 | PSMB8-AS1 | 1.82E-07 | 0.802415573 | 0.269 | 0.168 | 0.003577817 | Mari |
| cDC2 | DNAJC131 | 1.88E-07 | 0.802368152 | 0.359 | 0.249 | 0.003706103 | Mari |
| cDC2 | DBNL | 6.88E-22 | 0.802100634 | 0.729 | 0.501 | 1.35E-17 | Mari |
| cDC2 | CLTB | 7.66E-30 | 0.801845424 | 0.819 | 0.584 | 1.51E-25 | Mari |
| cDC2 | LY96 | 3.77E-20 | 0.801249757 | 0.722 | 0.542 | 7.41E-16 | Mari |
| cDC2 | PPCS | 1.34E-17 | 0.800592013 | 0.553 | 0.36 | 2.64E-13 | Mari |
| cDC2 | MRPS11 | 5.71E-10 | 0.800422137 | 0.368 | 0.232 | 1.12E-05 | Mari |
| cDC2 | R3HCC1 | 1.29E-06 | 0.799617692 | 0.252 | 0.161 | 0.025352922 | Mari |
| cDC2 | STMP1 | 2.19E-22 | 0.799510632 | 0.699 | 0.495 | 4.32E-18 | Mari |
| cDC2 | FCGR2B | 5.00E-17 | 0.799502343 | 0.75 | 0.624 | 9.84E-13 | Mari |
| cDC2 | AURKAIP1 | 1.22E-31 | 0.799325473 | 0.852 | 0.647 | 2.40E-27 | Mari |
| cDC2 | TMEM173 | 6.10E-07 | 0.799164392 | 0.299 | 0.199 | 0.01199843 | Mari |
| cDC2 | MRPL2 | 2.42E-06 | 0.798818194 | 0.275 | 0.186 | 0.047572788 | Mari |
| cDC2 | GET3 | 3.86E-15 | 0.79834679 | 0.574 | 0.397 | 7.60E-11 | Mari |
| cDC2 | AP2S1 | 5.52E-51 | 0.79833796 | 0.949 | 0.826 | 1.09E-46 | Mari |
| cDC2 | ORMDL2 | 7.53E-09 | 0.798246953 | 0.389 | 0.271 | 0.000148318 | Mari |
| cDC2 | COX14 | 1.76E-24 | 0.795849812 | 0.731 | 0.504 | 3.46E-20 | Mari |
| cDC2 | CD300E | 2.35E-09 | 0.795370016 | 0.282 | 0.162 | 4.62E-05 | Mari |
| cDC2 | EMC6 | 1.41E-12 | 0.794303828 | 0.463 | 0.314 | 2.78E-08 | Mari |
| cDC2 | SNRPG | 3.43E-39 | 0.793855929 | 0.903 | 0.72 | 6.75E-35 | Mari |
| cDC2 | MRPL40 | 1.21E-13 | 0.793210202 | 0.486 | 0.321 | 2.37E-09 | Mari |
| cDC2 | DPM2 | 1.67E-11 | 0.792729959 | 0.354 | 0.206 | 3.29E-07 | Mari |
| cDC2 | TRMT1 | 6.80E-14 | 0.790472627 | 0.484 | 0.315 | 1.34E-09 | Mari |
| cDC2 | SRM | 1.01E-14 | 0.78995564 | 0.519 | 0.341 | 2.00E-10 | Mari |
| cDC2 | TMEM167A | 1.84E-20 | 0.78782462 | 0.676 | 0.479 | 3.62E-16 | Mari |
| cDC2 | ITGA41 | 6.89E-24 | 0.787724371 | 0.727 | 0.493 | 1.36E-19 | Mari |
| cDC2 | PARL | 2.05E-16 | 0.787053824 | 0.59 | 0.403 | 4.04E-12 | Mari |
| cDC2 | SPCS2 | 7.03E-29 | 0.786016519 | 0.826 | 0.605 | 1.38E-24 | Mari |
| cDC2 | UBL5 | 1.65E-48 | 0.785597279 | 0.942 | 0.832 | 3.24E-44 | Mari |
| cDC2 | SEC11C | 6.03E-15 | 0.785520728 | 0.613 | 0.437 | 1.19E-10 | Mari |
| cDC2 | SLC39A101 | 4.18E-13 | 0.78484305 | 0.475 | 0.31 | 8.23E-09 | Mari |
| cDC2 | TPRKB | 4.03E-21 | 0.783524411 | 0.637 | 0.416 | 7.92E-17 | Mari |
| cDC2 | ATP5PF | 3.73E-46 | 0.783200774 | 0.926 | 0.76 | 7.34E-42 | Mari |

| | | | | | | | |
|---|---|---|---|---|---|---|---|
| cDC2 | BAX | 1.33E-25 | 0.780880984 | 0.796 | 0.572 | 2.62E-21 | Mari |
| cDC2 | SHKBP1 | 3.52E-15 | 0.780851448 | 0.525 | 0.338 | 6.94E-11 | Mari |
| cDC2 | HMGN1 | 6.71E-37 | 0.780396263 | 0.921 | 0.758 | 1.32E-32 | Mari |
| cDC2 | FBXO6 | 9.49E-15 | 0.776581169 | 0.576 | 0.395 | 1.87E-10 | Mari |
| cDC2 | FIBP | 1.57E-19 | 0.775993716 | 0.685 | 0.492 | 3.08E-15 | Mari |
| cDC2 | SLC12A9 | 2.56E-12 | 0.775608445 | 0.438 | 0.28 | 5.05E-08 | Mari |
| cDC2 | SRSF9 | 2.27E-36 | 0.773684308 | 0.905 | 0.747 | 4.46E-32 | Mari |
| cDC2 | TSN | 1.11E-06 | 0.772920944 | 0.292 | 0.197 | 0.021916973 | Mari |
| cDC2 | LSM4 | 2.55E-22 | 0.772886799 | 0.731 | 0.518 | 5.02E-18 | Mari |
| cDC2 | NEDD8 | 7.90E-32 | 0.772767187 | 0.884 | 0.69 | 1.56E-27 | Mari |
| cDC2 | NDUFB1 | 6.26E-29 | 0.772179686 | 0.843 | 0.617 | 1.23E-24 | Mari |
| cDC2 | TTC1 | 2.06E-11 | 0.771817292 | 0.463 | 0.31 | 4.06E-07 | Mari |
| cDC2 | SEM1 | 3.36E-36 | 0.771749895 | 0.907 | 0.737 | 6.60E-32 | Mari |
| cDC2 | JAGN1 | 9.65E-09 | 0.771747109 | 0.361 | 0.242 | 0.000189859 | Mari |
| cDC2 | MRPL51 | 3.89E-25 | 0.771658355 | 0.771 | 0.532 | 7.66E-21 | Mari |
| cDC2 | SERPINA1 | 1.57E-21 | 0.771139582 | 0.794 | 0.662 | 3.10E-17 | Mari |
| cDC2 | DERL2 | 1.23E-11 | 0.770832035 | 0.421 | 0.27 | 2.43E-07 | Mari |
| cDC2 | ZNHIT1 | 7.43E-23 | 0.769776295 | 0.748 | 0.56 | 1.46E-18 | Mari |
| cDC2 | NAPRT | 1.02E-08 | 0.769556304 | 0.389 | 0.265 | 0.000201403 | Mari |
| cDC2 | ISG15 | 5.92E-26 | 0.769147046 | 0.843 | 0.636 | 1.17E-21 | Mari |
| cDC2 | COPE | 5.95E-45 | 0.768717256 | 0.951 | 0.789 | 1.17E-40 | Mari |
| cDC2 | EIF2AK2 | 1.56E-09 | 0.767610187 | 0.394 | 0.258 | 3.07E-05 | Mari |
| cDC2 | ZDHHC121 | 4.39E-12 | 0.766965443 | 0.407 | 0.255 | 8.64E-08 | Mari |
| cDC2 | NDUFB3 | 4.03E-22 | 0.766473931 | 0.736 | 0.513 | 7.94E-18 | Mari |
| cDC2 | NUDT16 | 7.39E-11 | 0.766069719 | 0.449 | 0.31 | 1.45E-06 | Mari |
| cDC2 | CPQ1 | 2.44E-16 | 0.766063594 | 0.611 | 0.404 | 4.80E-12 | Mari |
| cDC2 | EBNA1BP2 | 6.37E-07 | 0.766019951 | 0.329 | 0.227 | 0.012528879 | Mari |
| cDC2 | POLR2J | 2.09E-17 | 0.765568119 | 0.616 | 0.431 | 4.11E-13 | Mari |
| cDC2 | DHRS4L2 | 6.04E-08 | 0.76531535 | 0.377 | 0.266 | 0.001189249 | Mari |
| cDC2 | SPINT21 | 4.61E-25 | 0.76495872 | 0.843 | 0.628 | 9.07E-21 | Mari |
| cDC2 | RAMAC | 4.20E-09 | 0.763829195 | 0.31 | 0.187 | 8.28E-05 | Mari |
| cDC2 | TMEM134 | 7.32E-10 | 0.763474533 | 0.419 | 0.287 | 1.44E-05 | Mari |
| cDC2 | TST | 9.62E-07 | 0.763020434 | 0.301 | 0.204 | 0.018945643 | Mari |
| cDC2 | RASGRP2 | 5.98E-08 | 0.762790763 | 0.299 | 0.184 | 0.001177685 | Mari |
| cDC2 | SPIDR | 1.04E-11 | 0.762408589 | 0.558 | 0.399 | 2.05E-07 | Mari |
| cDC2 | SNRPB2 | 1.32E-23 | 0.761869627 | 0.748 | 0.504 | 2.61E-19 | Mari |
| cDC2 | MIEN1 | 1.48E-14 | 0.761717838 | 0.463 | 0.289 | 2.92E-10 | Mari |
| cDC2 | MRPS26 | 2.26E-06 | 0.760394941 | 0.312 | 0.223 | 0.04449656 | Mari |
| cDC2 | COX7A2 | 4.83E-39 | 0.760147146 | 0.94 | 0.81 | 9.50E-35 | Mari |
| cDC2 | MRPL52 | 1.14E-22 | 0.758436412 | 0.773 | 0.588 | 2.24E-18 | Mari |
| cDC2 | STOML2 | 1.35E-14 | 0.757950934 | 0.542 | 0.363 | 2.66E-10 | Mari |
| cDC2 | FKBP1A | 3.27E-35 | 0.757630495 | 0.905 | 0.734 | 6.44E-31 | Mari |
| cDC2 | MRPS12 | 1.73E-13 | 0.757625809 | 0.519 | 0.352 | 3.41E-09 | Mari |
| cDC2 | CD86 | 8.59E-26 | 0.757530176 | 0.942 | 0.813 | 1.69E-21 | Mari |
| cDC2 | ARPC1A | 1.81E-08 | 0.756108995 | 0.387 | 0.263 | 0.00035647 | Mari |
| cDC2 | SERF2 | 6.40E-90 | 0.755920673 | 0.988 | 0.97 | 1.26E-85 | Mari |
| cDC2 | MRPL55 | 4.66E-15 | 0.754160458 | 0.593 | 0.417 | 9.18E-11 | Mari |
| cDC2 | DHRS4 | 6.45E-07 | 0.753297393 | 0.347 | 0.248 | 0.012691465 | Mari |
| cDC2 | DUSP12 | 1.03E-06 | 0.752731635 | 0.262 | 0.164 | 0.020352604 | Mari |
| cDC2 | PYURF | 2.19E-17 | 0.751875829 | 0.576 | 0.385 | 4.30E-13 | Mari |
| cDC2 | 11-Mar | 8.64E-21 | 0.751810241 | 0.676 | 0.428 | 1.70E-16 | Mari |
| cDC2 | MRPL57 | 2.17E-18 | 0.75118486 | 0.683 | 0.479 | 4.26E-14 | Mari |
| cDC2 | AK6 | 8.22E-08 | 0.750455423 | 0.289 | 0.181 | 0.001617996 | Mari |
| cDC2 | SDHAF2 | 1.37E-09 | 0.749555069 | 0.391 | 0.259 | 2.69E-05 | Mari |
| cDC2 | HSPB111 | 4.23E-16 | 0.749385332 | 0.604 | 0.418 | 8.32E-12 | Mari |
| cDC2 | ZMAT5 | 3.09E-08 | 0.748238968 | 0.301 | 0.188 | 0.000607972 | Mari |

| | | | | | | | |
|---|---|---|---|---|---|---|---|
| cDC2 | SLC35C21 | 1.20E-06 | 0.748108989 | 0.303 | 0.205 | 0.023578767 | Mari |
| cDC2 | OXLD1 | 8.74E-07 | 0.748085699 | 0.255 | 0.162 | 0.01720655 | Mari |
| cDC2 | SNX6 | 8.25E-21 | 0.747633768 | 0.808 | 0.638 | 1.62E-16 | Mari |
| cDC2 | TOMM6 | 5.38E-29 | 0.747200535 | 0.882 | 0.717 | 1.06E-24 | Mari |
| cDC2 | GADD45GIP1 | 4.02E-22 | 0.746758434 | 0.748 | 0.561 | 7.90E-18 | Mari |
| cDC2 | POP5 | 2.26E-07 | 0.746681454 | 0.294 | 0.189 | 0.004458156 | Mari |
| cDC2 | CNPY2 | 5.78E-13 | 0.746419199 | 0.537 | 0.373 | 1.14E-08 | Mari |
| cDC2 | PSENEN | 1.91E-16 | 0.745180598 | 0.66 | 0.475 | 3.76E-12 | Mari |
| cDC2 | ITGAE | 2.80E-10 | 0.744951957 | 0.359 | 0.223 | 5.52E-06 | Mari |
| cDC2 | MICOS10 | 2.85E-31 | 0.74397288 | 0.894 | 0.725 | 5.62E-27 | Mari |
| cDC2 | PSMB1 | 1.62E-38 | 0.743410265 | 0.933 | 0.795 | 3.18E-34 | Mari |
| cDC2 | EIF5A | 1.18E-26 | 0.743135981 | 0.882 | 0.74 | 2.33E-22 | Mari |
| cDC2 | PPIB | 2.31E-40 | 0.742569655 | 0.947 | 0.879 | 4.55E-36 | Mari |
| cDC2 | PDIA6 | 2.84E-20 | 0.741966191 | 0.741 | 0.546 | 5.59E-16 | Mari |
| cDC2 | TMEM126B | 1.21E-08 | 0.741385595 | 0.352 | 0.234 | 0.000237529 | Mari |
| cDC2 | NSMCE4A | 8.03E-09 | 0.738902313 | 0.373 | 0.251 | 0.000158011 | Mari |
| cDC2 | SNRNP40 | 1.05E-10 | 0.738465847 | 0.431 | 0.287 | 2.06E-06 | Mari |
| cDC2 | SS18L21 | 1.10E-13 | 0.738005038 | 0.465 | 0.299 | 2.16E-09 | Mari |
| cDC2 | SELENOT | 2.95E-21 | 0.737755569 | 0.729 | 0.51 | 5.80E-17 | Mari |
| cDC2 | MPV17 | 1.08E-13 | 0.737269609 | 0.507 | 0.341 | 2.13E-09 | Mari |
| cDC2 | SLC25A11 | 4.46E-11 | 0.734630941 | 0.493 | 0.342 | 8.78E-07 | Mari |
| cDC2 | PPP1R7 | 3.71E-13 | 0.734448945 | 0.539 | 0.375 | 7.30E-09 | Mari |
| cDC2 | COA1 | 5.72E-07 | 0.733911159 | 0.387 | 0.287 | 0.011266215 | Mari |
| cDC2 | SURF1 | 5.90E-18 | 0.733621323 | 0.632 | 0.424 | 1.16E-13 | Mari |
| cDC2 | PSMA2 | 1.91E-30 | 0.732507093 | 0.896 | 0.751 | 3.76E-26 | Mari |
| cDC2 | DAD1 | 6.47E-26 | 0.731942311 | 0.803 | 0.615 | 1.27E-21 | Mari |
| cDC2 | DCTN3 | 7.02E-22 | 0.731628492 | 0.729 | 0.517 | 1.38E-17 | Mari |
| cDC2 | TUBA1B | 1.76E-48 | 0.731522843 | 0.979 | 0.942 | 3.47E-44 | Mari |
| cDC2 | P2RY61 | 2.45E-06 | 0.731133248 | 0.301 | 0.204 | 0.048229168 | Mari |
| cDC2 | TPGS2 | 3.71E-08 | 0.731010252 | 0.296 | 0.182 | 0.000731211 | Mari |
| cDC2 | SCP2 | 2.43E-19 | 0.730368448 | 0.748 | 0.545 | 4.79E-15 | Mari |
| cDC2 | TMCO1 | 2.63E-13 | 0.730171024 | 0.553 | 0.386 | 5.17E-09 | Mari |
| cDC2 | CD2BP2 | 1.30E-07 | 0.729031603 | 0.384 | 0.279 | 0.00256836 | Mari |
| cDC2 | MCTS1 | 8.95E-11 | 0.728865504 | 0.479 | 0.338 | 1.76E-06 | Mari |
| cDC2 | EDF1 | 1.51E-54 | 0.728554527 | 0.972 | 0.862 | 2.98E-50 | Mari |
| cDC2 | MRPL22 | 3.46E-13 | 0.72821984 | 0.465 | 0.292 | 6.80E-09 | Mari |
| cDC2 | PHPT1 | 1.27E-26 | 0.728019212 | 0.852 | 0.672 | 2.50E-22 | Mari |
| cDC2 | TMEM109 | 1.81E-13 | 0.727934143 | 0.586 | 0.414 | 3.57E-09 | Mari |
| cDC2 | DTD1 | 1.11E-11 | 0.726894179 | 0.447 | 0.284 | 2.18E-07 | Mari |
| cDC2 | TMBIM4 | 5.29E-26 | 0.726712291 | 0.845 | 0.639 | 1.04E-21 | Mari |
| cDC2 | ANAPC13 | 1.69E-06 | 0.723900106 | 0.299 | 0.2 | 0.033247242 | Mari |
| cDC2 | POLR3GL | 2.22E-08 | 0.722422285 | 0.373 | 0.255 | 0.000436138 | Mari |
| cDC2 | S100A4 | 1.21E-24 | 0.722302668 | 0.963 | 0.906 | 2.38E-20 | Mari |
| cDC2 | LSM2 | 5.08E-15 | 0.721447118 | 0.579 | 0.393 | 1.00E-10 | Mari |
| cDC2 | SUMO1 | 1.02E-24 | 0.720659931 | 0.826 | 0.622 | 2.01E-20 | Mari |
| cDC2 | SCAND1 | 7.84E-23 | 0.720107975 | 0.782 | 0.574 | 1.54E-18 | Mari |
| cDC2 | EAPP1 | 4.92E-09 | 0.720106476 | 0.373 | 0.245 | 9.68E-05 | Mari |
| cDC2 | MRPL33 | 3.82E-20 | 0.720054881 | 0.715 | 0.484 | 7.51E-16 | Mari |
| cDC2 | LILRB1 | 3.19E-08 | 0.719939371 | 0.338 | 0.216 | 0.000628604 | Mari |
| cDC2 | UBE2L3 | 5.90E-28 | 0.719939177 | 0.826 | 0.641 | 1.16E-23 | Mari |
| cDC2 | JKAMP | 5.16E-10 | 0.719835933 | 0.315 | 0.19 | 1.02E-05 | Mari |
| cDC2 | ARL1 | 1.56E-09 | 0.719591633 | 0.377 | 0.245 | 3.07E-05 | Mari |
| cDC2 | SNRPA1 | 2.58E-17 | 0.719522667 | 0.604 | 0.386 | 5.08E-13 | Mari |
| cDC2 | HTATIP2 | 1.08E-06 | 0.718651782 | 0.398 | 0.284 | 0.02128615 | Mari |
| cDC2 | MYL12B | 2.11E-53 | 0.717990752 | 0.956 | 0.887 | 4.15E-49 | Mari |
| cDC2 | COX161 | 2.88E-11 | 0.717543651 | 0.53 | 0.389 | 5.66E-07 | Mari |

| | | | | | | | |
|---|---|---|---|---|---|---|---|
| cDC2 | SNRNP25 | 3.68E-11 | 0.715909278 | 0.509 | 0.36 | 7.24E-07 | Mari |
| cDC2 | SASH3 | 2.82E-08 | 0.715530348 | 0.356 | 0.24 | 0.000554421 | Mari |
| cDC2 | NAA20 | 2.34E-11 | 0.714102456 | 0.53 | 0.377 | 4.61E-07 | Mari |
| cDC2 | GPN3 | 1.63E-07 | 0.713203002 | 0.336 | 0.225 | 0.003202363 | Mari |
| cDC2 | TMEM160 | 9.33E-16 | 0.713123462 | 0.65 | 0.488 | 1.84E-11 | Mari |
| cDC2 | BOLA31 | 7.70E-08 | 0.713080908 | 0.382 | 0.269 | 0.001515756 | Mari |
| cDC2 | BUD31 | 6.91E-20 | 0.711548947 | 0.745 | 0.545 | 1.36E-15 | Mari |
| cDC2 | COMMD3 | 9.83E-09 | 0.709830926 | 0.477 | 0.361 | 0.000193439 | Mari |
| cDC2 | TMX1 | 8.09E-08 | 0.709662889 | 0.417 | 0.307 | 0.00159153 | Mari |
| cDC2 | ATP6V0B | 3.46E-44 | 0.709591143 | 0.935 | 0.836 | 6.82E-40 | Mari |
| cDC2 | COX17 | 5.67E-24 | 0.708630627 | 0.796 | 0.563 | 1.12E-19 | Mari |
| cDC2 | BBIP1 | 3.93E-08 | 0.708123648 | 0.294 | 0.183 | 0.000774476 | Mari |
| cDC2 | GTF3A | 6.92E-16 | 0.707142277 | 0.708 | 0.541 | 1.36E-11 | Mari |
| cDC2 | GTF2A2 | 3.38E-15 | 0.706695812 | 0.692 | 0.525 | 6.65E-11 | Mari |
| cDC2 | PSMA3 | 1.66E-20 | 0.706567527 | 0.787 | 0.597 | 3.26E-16 | Mari |
| cDC2 | BANF1 | 1.31E-22 | 0.705964208 | 0.789 | 0.607 | 2.57E-18 | Mari |
| cDC2 | MIIP | 5.88E-07 | 0.704526231 | 0.287 | 0.186 | 0.011581754 | Mari |
| cDC2 | IFI35 | 3.09E-09 | 0.703776105 | 0.447 | 0.32 | 6.09E-05 | Mari |
| cDC2 | FAM50A | 3.99E-16 | 0.702328504 | 0.627 | 0.445 | 7.85E-12 | Mari |
| cDC2 | MRPL12 | 1.13E-09 | 0.700303661 | 0.433 | 0.296 | 2.23E-05 | Mari |
| cDC2 | SMIM19 | 1.81E-10 | 0.699814619 | 0.493 | 0.351 | 3.57E-06 | Mari |
| cDC2 | TTC33 | 1.58E-06 | 0.699684173 | 0.396 | 0.288 | 0.031157872 | Mari |
| cDC2 | MYDGF | 3.27E-24 | 0.699301041 | 0.843 | 0.647 | 6.43E-20 | Mari |
| cDC2 | COMMD4 | 1.03E-09 | 0.699241095 | 0.461 | 0.327 | 2.03E-05 | Mari |
| cDC2 | FAM174C | 8.97E-15 | 0.699003098 | 0.618 | 0.448 | 1.77E-10 | Mari |
| cDC2 | TXNDC12 | 5.16E-13 | 0.69881077 | 0.553 | 0.388 | 1.01E-08 | Mari |
| cDC2 | SDAD1 | 2.42E-07 | 0.698348932 | 0.319 | 0.212 | 0.004765071 | Mari |
| cDC2 | TRMT112 | 9.40E-27 | 0.69832384 | 0.847 | 0.666 | 1.85E-22 | Mari |
| cDC2 | RNFT1 | 1.12E-06 | 0.698283424 | 0.282 | 0.184 | 0.022110486 | Mari |
| cDC2 | SIVA1 | 1.64E-20 | 0.697450895 | 0.734 | 0.524 | 3.23E-16 | Mari |
| cDC2 | SNAPIN | 5.62E-09 | 0.697443763 | 0.329 | 0.208 | 0.000110589 | Mari |
| cDC2 | ARL16 | 1.04E-06 | 0.697324056 | 0.308 | 0.208 | 0.020441925 | Mari |
| cDC2 | PEX2 | 5.51E-08 | 0.695221269 | 0.301 | 0.192 | 0.001085478 | Mari |
| cDC2 | ATP5MPL1 | 2.44E-22 | 0.694845093 | 0.829 | 0.618 | 4.80E-18 | Mari |
| cDC2 | CAPZA2 | 3.85E-20 | 0.692118287 | 0.782 | 0.618 | 7.58E-16 | Mari |
| cDC2 | TMEM208 | 5.61E-18 | 0.691938337 | 0.593 | 0.384 | 1.10E-13 | Mari |
| cDC2 | VPS29 | 4.27E-31 | 0.690275809 | 0.889 | 0.736 | 8.40E-27 | Mari |
| cDC2 | CCS | 5.41E-08 | 0.690146596 | 0.377 | 0.258 | 0.001065876 | Mari |
| cDC2 | CDC42 | 1.25E-49 | 0.689900694 | 0.965 | 0.898 | 2.47E-45 | Mari |
| cDC2 | TUBA1A | 1.67E-30 | 0.689714897 | 0.961 | 0.837 | 3.29E-26 | Mari |
| cDC2 | AIMP1 | 2.65E-14 | 0.689467354 | 0.6 | 0.419 | 5.21E-10 | Mari |
| cDC2 | MRPL41 | 1.28E-20 | 0.688775937 | 0.722 | 0.548 | 2.52E-16 | Mari |
| cDC2 | MIF4GD | 7.25E-12 | 0.688752504 | 0.583 | 0.428 | 1.43E-07 | Mari |
| cDC2 | WASHC3 | 2.43E-12 | 0.688624589 | 0.502 | 0.336 | 4.79E-08 | Mari |
| cDC2 | TMEM59 | 4.12E-21 | 0.688581441 | 0.799 | 0.615 | 8.10E-17 | Mari |
| cDC2 | CD300A | 2.08E-17 | 0.688183205 | 0.618 | 0.4 | 4.10E-13 | Mari |
| cDC2 | ERH | 5.70E-24 | 0.686911715 | 0.831 | 0.622 | 1.12E-19 | Mari |
| cDC2 | ICAM3 | 5.76E-08 | 0.686646543 | 0.387 | 0.259 | 0.001133369 | Mari |
| cDC2 | ATXN10 | 2.92E-13 | 0.685087421 | 0.516 | 0.343 | 5.75E-09 | Mari |
| cDC2 | COA4 | 5.01E-12 | 0.684805126 | 0.512 | 0.358 | 9.86E-08 | Mari |
| cDC2 | TRAPPC5 | 4.14E-21 | 0.684222261 | 0.789 | 0.603 | 8.15E-17 | Mari |
| cDC2 | TSR3 | 3.70E-07 | 0.683913334 | 0.382 | 0.276 | 0.007281737 | Mari |
| cDC2 | DYNLRB1 | 4.26E-22 | 0.683109442 | 0.762 | 0.543 | 8.39E-18 | Mari |
| cDC2 | HSD17B10 | 6.15E-13 | 0.682588623 | 0.572 | 0.403 | 1.21E-08 | Mari |
| cDC2 | S100A71 | 1.51E-09 | 0.682177236 | 0.296 | 0.171 | 2.98E-05 | Mari |
| cDC2 | GPAA1 | 7.54E-08 | 0.680606523 | 0.345 | 0.234 | 0.001484738 | Mari |

| | | | | | | | |
|---|---|---|---|---|---|---|---|
| cDC2 | VPS28 | 4.65E-27 | 0.68008498 | 0.875 | 0.682 | 9.15E-23 | Mari |
| cDC2 | ECHS11 | 3.86E-19 | 0.679773301 | 0.713 | 0.503 | 7.60E-15 | Mari |
| cDC2 | CIAO2B | 1.26E-24 | 0.679557906 | 0.836 | 0.642 | 2.48E-20 | Mari |
| cDC2 | TMEM147 | 3.40E-13 | 0.677857138 | 0.537 | 0.367 | 6.70E-09 | Mari |
| cDC2 | SERPINF1 | 4.36E-13 | 0.677573256 | 0.775 | 0.616 | 8.58E-09 | Mari |
| cDC2 | PSMB3 | 1.18E-31 | 0.67734536 | 0.917 | 0.769 | 2.32E-27 | Mari |
| cDC2 | POLR2K | 3.62E-13 | 0.677215797 | 0.535 | 0.371 | 7.12E-09 | Mari |
| cDC2 | BCKDK | 2.04E-08 | 0.676489641 | 0.449 | 0.331 | 0.000401104 | Mari |
| cDC2 | UROS | 1.59E-08 | 0.676460606 | 0.4 | 0.283 | 0.000313325 | Mari |
| cDC2 | UQCRQ | 2.39E-23 | 0.67615315 | 0.833 | 0.637 | 4.71E-19 | Mari |
| cDC2 | ATP5MC1 | 6.84E-20 | 0.676008568 | 0.782 | 0.595 | 1.35E-15 | Mari |
| cDC2 | ISOC21 | 3.08E-10 | 0.675529141 | 0.512 | 0.358 | 6.07E-06 | Mari |
| cDC2 | UQCRFS1 | 7.22E-20 | 0.67433987 | 0.759 | 0.578 | 1.42E-15 | Mari |
| cDC2 | MRPL54 | 8.81E-14 | 0.673914675 | 0.66 | 0.496 | 1.73E-09 | Mari |
| cDC2 | TXNL4A | 1.42E-19 | 0.673892653 | 0.757 | 0.563 | 2.80E-15 | Mari |
| cDC2 | NDUFC1 | 1.10E-13 | 0.673754145 | 0.567 | 0.396 | 2.16E-09 | Mari |
| cDC2 | ESD | 6.45E-14 | 0.673582603 | 0.65 | 0.491 | 1.27E-09 | Mari |
| cDC2 | GZF1 | 2.36E-09 | 0.67304663 | 0.463 | 0.318 | 4.66E-05 | Mari |
| cDC2 | SIGIRR | 2.43E-06 | 0.67301689 | 0.326 | 0.228 | 0.047769255 | Mari |
| cDC2 | PFN1 | 2.36E-60 | 0.6727312 | 0.981 | 0.961 | 4.64E-56 | Mari |
| cDC2 | VPS35 | 1.00E-18 | 0.671608365 | 0.778 | 0.589 | 1.97E-14 | Mari |
| cDC2 | IQCB11 | 4.11E-09 | 0.671400184 | 0.34 | 0.214 | 8.09E-05 | Mari |
| cDC2 | ITGB1BP11 | 1.56E-10 | 0.67115994 | 0.481 | 0.337 | 3.07E-06 | Mari |
| cDC2 | YPEL5 | 2.87E-16 | 0.671092839 | 0.812 | 0.638 | 5.65E-12 | Mari |
| cDC2 | SRI | 3.89E-17 | 0.670652175 | 0.711 | 0.531 | 7.65E-13 | Mari |
| cDC2 | SUB1 | 1.07E-35 | 0.669644586 | 0.972 | 0.894 | 2.10E-31 | Mari |
| cDC2 | IFITM1 | 7.16E-25 | 0.668979617 | 0.53 | 0.269 | 1.41E-20 | Mari |
| cDC2 | NUDT5 | 4.97E-08 | 0.66821065 | 0.407 | 0.289 | 0.000978646 | Mari |
| cDC2 | PDCD6 | 1.59E-19 | 0.667429701 | 0.743 | 0.52 | 3.14E-15 | Mari |
| cDC2 | OSTF1 | 3.89E-20 | 0.667022543 | 0.794 | 0.618 | 7.65E-16 | Mari |
| cDC2 | MSRB1 | 7.68E-07 | 0.666528188 | 0.31 | 0.208 | 0.015124123 | Mari |
| cDC2 | MTIF3 | 4.78E-09 | 0.666062031 | 0.458 | 0.334 | 9.41E-05 | Mari |
| cDC2 | TYROBP | 5.24E-50 | 0.664419196 | 0.986 | 0.968 | 1.03E-45 | Mari |
| cDC2 | SRA1 | 8.89E-21 | 0.663419873 | 0.755 | 0.561 | 1.75E-16 | Mari |
| cDC2 | PSMB6 | 1.97E-27 | 0.661711454 | 0.87 | 0.717 | 3.89E-23 | Mari |
| cDC2 | TLR1 | 4.50E-13 | 0.661352677 | 0.519 | 0.351 | 8.87E-09 | Mari |
| cDC2 | COPS6 | 2.51E-09 | 0.661162648 | 0.486 | 0.356 | 4.93E-05 | Mari |
| cDC2 | PLEKHJ1 | 3.24E-09 | 0.660526565 | 0.438 | 0.309 | 6.38E-05 | Mari |
| cDC2 | B3GAT3 | 1.53E-07 | 0.66008504 | 0.326 | 0.215 | 0.003010614 | Mari |
| cDC2 | HCCS | 1.94E-11 | 0.66007759 | 0.414 | 0.257 | 3.82E-07 | Mari |
| cDC2 | GLIPR2 | 3.58E-19 | 0.659872028 | 0.863 | 0.695 | 7.04E-15 | Mari |
| cDC2 | COMMD9 | 8.22E-14 | 0.659689853 | 0.569 | 0.394 | 1.62E-09 | Mari |
| cDC2 | SMIM151 | 2.42E-07 | 0.65901145 | 0.322 | 0.218 | 0.004756914 | Mari |
| cDC2 | PHF11 | 7.00E-08 | 0.659001299 | 0.435 | 0.319 | 0.001377596 | Mari |
| cDC2 | SLC39A3 | 6.21E-09 | 0.658091705 | 0.347 | 0.223 | 0.000122171 | Mari |
| cDC2 | ATP5F1E | 5.90E-80 | 0.657012302 | 0.984 | 0.979 | 1.16E-75 | Mari |
| cDC2 | FUOM | 2.81E-18 | 0.656427474 | 0.745 | 0.553 | 5.53E-14 | Mari |
| cDC2 | TM6SF1 | 1.03E-08 | 0.65594928 | 0.391 | 0.263 | 0.000202215 | Mari |
| cDC2 | PPP4C | 1.36E-19 | 0.655897352 | 0.748 | 0.562 | 2.68E-15 | Mari |
| cDC2 | STUB1 | 4.11E-12 | 0.65575468 | 0.588 | 0.436 | 8.10E-08 | Mari |
| cDC2 | TMED9 | 5.29E-14 | 0.655702349 | 0.671 | 0.498 | 1.04E-09 | Mari |
| cDC2 | PRKAG11 | 2.12E-06 | 0.655514558 | 0.352 | 0.252 | 0.041708143 | Mari |
| cDC2 | TIMM10 | 1.56E-14 | 0.655495205 | 0.627 | 0.443 | 3.07E-10 | Mari |
| cDC2 | MRPS71 | 3.64E-10 | 0.655378818 | 0.509 | 0.369 | 7.16E-06 | Mari |
| cDC2 | TXN2 | 6.55E-17 | 0.654911857 | 0.697 | 0.509 | 1.29E-12 | Mari |
| cDC2 | GSTO1 | 6.41E-33 | 0.653519738 | 0.938 | 0.816 | 1.26E-28 | Mari |

| | | | | | | | |
|---|---|---|---|---|---|---|---|
| cDC2 | NDUFA9 | 3.46E-10 | 0.653039296 | 0.477 | 0.34 | 6.80E-06 | Mari |
| cDC2 | NDUFA1 | 3.93E-32 | 0.652448498 | 0.921 | 0.748 | 7.74E-28 | Mari |
| cDC2 | PIN4 | 7.62E-09 | 0.65209578 | 0.405 | 0.271 | 0.000150073 | Mari |
| cDC2 | ARPC3 | 5.70E-64 | 0.651415671 | 0.963 | 0.953 | 1.12E-59 | Mari |
| cDC2 | MYL6 | 3.11E-55 | 0.65095474 | 0.984 | 0.964 | 6.12E-51 | Mari |
| cDC2 | TMEM2731 | 1.62E-06 | 0.649661362 | 0.405 | 0.294 | 0.031925689 | Mari |
| cDC2 | NDUFV3 | 3.61E-11 | 0.649099035 | 0.525 | 0.367 | 7.11E-07 | Mari |
| cDC2 | LAMTOR4 | 1.64E-30 | 0.648471927 | 0.914 | 0.786 | 3.23E-26 | Mari |
| cDC2 | TMSB10 | 2.14E-67 | 0.648338773 | 0.995 | 0.995 | 4.21E-63 | Mari |
| cDC2 | CAPG | 6.53E-20 | 0.648191302 | 0.889 | 0.777 | 1.29E-15 | Mari |
| cDC2 | FLOT1 | 3.85E-18 | 0.648079829 | 0.711 | 0.516 | 7.58E-14 | Mari |
| cDC2 | NOL7 | 3.21E-14 | 0.64765309 | 0.713 | 0.546 | 6.31E-10 | Mari |
| cDC2 | BCAS2 | 1.07E-12 | 0.647600664 | 0.657 | 0.516 | 2.10E-08 | Mari |
| cDC2 | VTI1B | 1.02E-06 | 0.64701333 | 0.444 | 0.348 | 0.020032358 | Mari |
| cDC2 | SEC61B | 9.84E-34 | 0.646984329 | 0.944 | 0.88 | 1.94E-29 | Mari |
| cDC2 | TMEM35B | 7.84E-09 | 0.646845597 | 0.405 | 0.275 | 0.000154242 | Mari |
| cDC2 | SAMHD1 | 7.70E-22 | 0.646709006 | 0.884 | 0.701 | 1.52E-17 | Mari |
| cDC2 | FCGRT | 4.44E-18 | 0.646451391 | 0.877 | 0.761 | 8.74E-14 | Mari |
| cDC2 | TAF9 | 1.45E-09 | 0.646269716 | 0.449 | 0.314 | 2.86E-05 | Mari |
| cDC2 | POLR2L | 9.00E-25 | 0.645742148 | 0.868 | 0.688 | 1.77E-20 | Mari |
| cDC2 | MICU2 | 2.11E-07 | 0.644650303 | 0.366 | 0.253 | 0.004159643 | Mari |
| cDC2 | LY86 | 2.40E-11 | 0.644546561 | 0.644 | 0.508 | 4.72E-07 | Mari |
| cDC2 | EXOSC7 | 1.79E-06 | 0.643224096 | 0.301 | 0.206 | 0.035302512 | Mari |
| cDC2 | SNRPD1 | 3.70E-18 | 0.643162981 | 0.757 | 0.56 | 7.29E-14 | Mari |
| cDC2 | PSMC4 | 4.64E-13 | 0.642896297 | 0.664 | 0.504 | 9.13E-09 | Mari |
| cDC2 | NT5C | 6.11E-16 | 0.64089568 | 0.688 | 0.503 | 1.20E-11 | Mari |
| cDC2 | ACP1 | 1.90E-14 | 0.640417953 | 0.637 | 0.454 | 3.73E-10 | Mari |
| cDC2 | FAM32A | 1.83E-11 | 0.639803549 | 0.535 | 0.382 | 3.60E-07 | Mari |
| cDC2 | FAM49B1 | 1.74E-29 | 0.638592004 | 0.951 | 0.786 | 3.43E-25 | Mari |
| cDC2 | PRDX21 | 4.75E-09 | 0.638368536 | 0.438 | 0.303 | 9.35E-05 | Mari |
| cDC2 | TMA7 | 9.25E-43 | 0.636819027 | 0.968 | 0.875 | 1.82E-38 | Mari |
| cDC2 | MRPL43 | 1.91E-12 | 0.636806615 | 0.569 | 0.405 | 3.75E-08 | Mari |
| cDC2 | UFD1 | 6.64E-09 | 0.636789232 | 0.539 | 0.404 | 0.000130661 | Mari |
| cDC2 | PDCD10 | 1.44E-13 | 0.635836205 | 0.613 | 0.442 | 2.84E-09 | Mari |
| cDC2 | PTPRC | 8.34E-31 | 0.635697016 | 0.977 | 0.889 | 1.64E-26 | Mari |
| cDC2 | IFI6 | 4.57E-07 | 0.635520433 | 0.433 | 0.329 | 0.008988913 | Mari |
| cDC2 | SRSF7 | 1.79E-24 | 0.635372111 | 0.933 | 0.763 | 3.53E-20 | Mari |
| cDC2 | RPA3 | 1.31E-10 | 0.635267937 | 0.486 | 0.328 | 2.57E-06 | Mari |
| cDC2 | RABAC1 | 7.57E-20 | 0.635193786 | 0.755 | 0.555 | 1.49E-15 | Mari |
| cDC2 | CARD16 | 6.98E-30 | 0.634929542 | 0.919 | 0.777 | 1.37E-25 | Mari |
| cDC2 | LMAN2 | 1.77E-12 | 0.634353999 | 0.69 | 0.534 | 3.47E-08 | Mari |
| cDC2 | LARP71 | 1.03E-08 | 0.634276949 | 0.419 | 0.292 | 0.000203328 | Mari |
| cDC2 | STXBP2 | 5.15E-10 | 0.633649627 | 0.565 | 0.413 | 1.01E-05 | Mari |
| cDC2 | SLC66A3 | 7.68E-10 | 0.633307173 | 0.488 | 0.348 | 1.51E-05 | Mari |
| cDC2 | LYRM2 | 2.42E-06 | 0.632996495 | 0.356 | 0.257 | 0.047643729 | Mari |
| cDC2 | PKIB | 1.90E-13 | 0.632194968 | 0.623 | 0.441 | 3.74E-09 | Mari |
| cDC2 | LSM8 | 2.32E-15 | 0.63109493 | 0.704 | 0.531 | 4.56E-11 | Mari |
| cDC2 | SRSF5 | 6.22E-26 | 0.630680293 | 0.87 | 0.651 | 1.22E-21 | Mari |
| cDC2 | PSMC5 | 8.59E-16 | 0.62991417 | 0.704 | 0.522 | 1.69E-11 | Mari |
| cDC2 | TWF2 | 4.43E-19 | 0.62973715 | 0.785 | 0.598 | 8.72E-15 | Mari |
| cDC2 | PSMD8 | 7.24E-21 | 0.628390187 | 0.859 | 0.713 | 1.43E-16 | Mari |
| cDC2 | HAT1 | 3.91E-10 | 0.627965924 | 0.414 | 0.275 | 7.70E-06 | Mari |
| cDC2 | TMEM179B | 4.87E-14 | 0.627961735 | 0.613 | 0.413 | 9.59E-10 | Mari |
| cDC2 | EEF1E1 | 1.17E-07 | 0.627480925 | 0.394 | 0.276 | 0.002307355 | Mari |
| cDC2 | MRPS16 | 4.08E-07 | 0.627194097 | 0.405 | 0.303 | 0.008028457 | Mari |
| cDC2 | ENTPD1-AS11 | 4.80E-07 | 0.627179035 | 0.477 | 0.356 | 0.009450553 | Mari |

| | | | | | | | |
|---|---|---|---|---|---|---|---|
| cDC2 | NDUFS8 | 3.60E-21 | 0.625472018 | 0.812 | 0.597 | 7.09E-17 | Mari |
| cDC2 | EXOSC8 | 3.33E-08 | 0.625143037 | 0.356 | 0.236 | 0.000654928 | Mari |
| cDC2 | ATF3 | 1.02E-19 | 0.624593441 | 0.935 | 0.782 | 2.01E-15 | Mari |
| cDC2 | HPRT1 | 1.56E-13 | 0.624025093 | 0.613 | 0.44 | 3.07E-09 | Mari |
| cDC2 | PYCARD | 2.56E-26 | 0.623712266 | 0.931 | 0.839 | 5.04E-22 | Mari |
| cDC2 | XPA | 6.80E-08 | 0.622669421 | 0.296 | 0.188 | 0.001339258 | Mari |
| cDC2 | STK17B | 1.34E-20 | 0.62166251 | 0.931 | 0.76 | 2.63E-16 | Mari |
| cDC2 | DGCR6L | 2.14E-06 | 0.620883725 | 0.375 | 0.267 | 0.042081859 | Mari |
| cDC2 | DYNC1I21 | 1.91E-08 | 0.620186396 | 0.447 | 0.319 | 0.000375738 | Mari |
| cDC2 | TSPO | 5.26E-25 | 0.619666782 | 0.903 | 0.827 | 1.03E-20 | Mari |
| cDC2 | HMGN3 | 4.47E-19 | 0.61887223 | 0.824 | 0.651 | 8.80E-15 | Mari |
| cDC2 | NOP10 | 5.88E-21 | 0.618458676 | 0.817 | 0.647 | 1.16E-16 | Mari |
| cDC2 | MOB1A | 1.74E-26 | 0.618199716 | 0.905 | 0.767 | 3.42E-22 | Mari |
| cDC2 | GNB41 | 1.57E-08 | 0.618188415 | 0.539 | 0.399 | 0.000309409 | Mari |
| cDC2 | JTB | 7.92E-14 | 0.617618445 | 0.671 | 0.525 | 1.56E-09 | Mari |
| cDC2 | BCL7C | 1.06E-10 | 0.617517407 | 0.505 | 0.357 | 2.08E-06 | Mari |
| cDC2 | IAH1 | 2.07E-13 | 0.617133938 | 0.639 | 0.458 | 4.06E-09 | Mari |
| cDC2 | DPY30 | 2.98E-09 | 0.616522725 | 0.4 | 0.271 | 5.87E-05 | Mari |
| cDC2 | PLGRKT1 | 5.69E-11 | 0.616463172 | 0.537 | 0.38 | 1.12E-06 | Mari |
| cDC2 | TFPT | 2.30E-06 | 0.61531962 | 0.377 | 0.273 | 0.045295156 | Mari |
| cDC2 | MCRIP1 | 8.64E-10 | 0.614905442 | 0.556 | 0.421 | 1.70E-05 | Mari |
| cDC2 | EIF4E2 | 1.30E-13 | 0.614892073 | 0.683 | 0.508 | 2.57E-09 | Mari |
| cDC2 | PSMB8 | 1.24E-22 | 0.614723379 | 0.896 | 0.768 | 2.44E-18 | Mari |
| cDC2 | CEBPZOS | 2.10E-06 | 0.613849448 | 0.324 | 0.22 | 0.041275417 | Mari |
| cDC2 | RNF166 | 7.18E-07 | 0.611582864 | 0.458 | 0.356 | 0.014131225 | Mari |
| cDC2 | PSMG2 | 3.33E-16 | 0.611132894 | 0.755 | 0.588 | 6.55E-12 | Mari |
| cDC2 | DCTPP11 | 6.07E-07 | 0.610808985 | 0.419 | 0.307 | 0.011950292 | Mari |
| cDC2 | RANBP1 | 1.03E-16 | 0.610620241 | 0.738 | 0.547 | 2.04E-12 | Mari |
| cDC2 | TRAPPC6A | 1.10E-12 | 0.610176591 | 0.574 | 0.404 | 2.16E-08 | Mari |
| cDC2 | IMP4 | 3.03E-08 | 0.610075743 | 0.449 | 0.316 | 0.000596695 | Mari |
| cDC2 | NDUFB5 | 5.48E-19 | 0.608362577 | 0.808 | 0.63 | 1.08E-14 | Mari |
| cDC2 | MRPL21 | 4.08E-08 | 0.608133103 | 0.475 | 0.35 | 0.000803526 | Mari |
| cDC2 | SLC8A11 | 2.07E-24 | 0.607937233 | 0.958 | 0.782 | 4.08E-20 | Mari |
| cDC2 | GPR651 | 3.12E-12 | 0.606869283 | 0.528 | 0.35 | 6.15E-08 | Mari |
| cDC2 | CYBB | 1.07E-09 | 0.606426371 | 0.734 | 0.624 | 2.10E-05 | Mari |
| cDC2 | NTMT1 | 4.59E-09 | 0.605875236 | 0.481 | 0.355 | 9.04E-05 | Mari |
| cDC2 | NDUFS5 | 2.07E-22 | 0.605777533 | 0.861 | 0.697 | 4.08E-18 | Mari |
| cDC2 | NIPSNAP2 | 3.76E-07 | 0.604368132 | 0.442 | 0.33 | 0.0074095 | Mari |
| cDC2 | NCF4 | 5.43E-10 | 0.604038575 | 0.428 | 0.28 | 1.07E-05 | Mari |
| cDC2 | SRP191 | 1.17E-15 | 0.603920811 | 0.685 | 0.484 | 2.30E-11 | Mari |
| cDC2 | PDCL3 | 2.37E-14 | 0.603680827 | 0.609 | 0.434 | 4.67E-10 | Mari |
| cDC2 | LAMTOR5 | 1.84E-12 | 0.603338948 | 0.706 | 0.538 | 3.63E-08 | Mari |
| cDC2 | EPSTI1 | 4.33E-18 | 0.602762355 | 0.817 | 0.623 | 8.53E-14 | Mari |
| cDC2 | CLNS1A | 1.65E-14 | 0.602570901 | 0.722 | 0.558 | 3.24E-10 | Mari |
| cDC2 | PAXX | 5.29E-14 | 0.602510939 | 0.688 | 0.521 | 1.04E-09 | Mari |
| cDC2 | NME1 | 1.31E-11 | 0.601877647 | 0.6 | 0.448 | 2.57E-07 | Mari |
| cDC2 | APRT | 1.43E-29 | 0.601855236 | 0.933 | 0.807 | 2.81E-25 | Mari |
| cDC2 | SP1401 | 1.04E-06 | 0.601551507 | 0.368 | 0.255 | 0.020559363 | Mari |
| cDC2 | IFI16 | 3.66E-18 | 0.601401712 | 0.815 | 0.623 | 7.21E-14 | Mari |
| cDC2 | NDUFB10 | 6.55E-22 | 0.601292789 | 0.847 | 0.66 | 1.29E-17 | Mari |
| cDC2 | COMMD2 | 8.52E-11 | 0.600849862 | 0.387 | 0.233 | 1.68E-06 | Mari |
| cDC2 | TMEM256 | 1.01E-13 | 0.600524306 | 0.681 | 0.533 | 2.00E-09 | Mari |
| cDC2 | WDR18 | 8.01E-07 | 0.599421385 | 0.273 | 0.174 | 0.015765499 | Mari |
| cDC2 | SSBP4 | 1.28E-07 | 0.59841639 | 0.475 | 0.358 | 0.002516004 | Mari |
| cDC2 | SNRPE | 3.33E-14 | 0.597541527 | 0.699 | 0.534 | 6.56E-10 | Mari |
| cDC2 | PSTPIP1 | 6.24E-08 | 0.597516726 | 0.458 | 0.336 | 0.001228271 | Mari |

| | | | | | | | |
|---|---|---|---|---|---|---|---|
| cDC2 | HCK | 4.32E-14 | 0.59577213 | 0.803 | 0.639 | 8.51E-10 | Mari |
| cDC2 | MCM5 | 1.10E-06 | 0.594755227 | 0.347 | 0.244 | 0.02170182 | Mari |
| cDC2 | PFDN11 | 2.92E-12 | 0.591854388 | 0.56 | 0.384 | 5.74E-08 | Mari |
| cDC2 | SNHG9 | 2.23E-09 | 0.591415578 | 0.301 | 0.179 | 4.39E-05 | Mari |
| cDC2 | ETV7 | 1.92E-06 | 0.59097811 | 0.312 | 0.212 | 0.03785356 | Mari |
| cDC2 | COX8A | 9.18E-32 | 0.590716868 | 0.942 | 0.872 | 1.81E-27 | Mari |
| cDC2 | LEPROTL1 | 2.37E-13 | 0.590063324 | 0.618 | 0.436 | 4.67E-09 | Mari |
| cDC2 | NDUFA12 | 9.51E-19 | 0.589478323 | 0.875 | 0.72 | 1.87E-14 | Mari |
| cDC2 | MRPS28 | 1.01E-07 | 0.589301638 | 0.419 | 0.305 | 0.001989052 | Mari |
| cDC2 | MTCH2 | 1.47E-09 | 0.58888372 | 0.519 | 0.374 | 2.88E-05 | Mari |
| cDC2 | NDUFB8 | 1.20E-22 | 0.588628407 | 0.896 | 0.711 | 2.36E-18 | Mari |
| cDC2 | PHB | 2.92E-12 | 0.587945901 | 0.676 | 0.521 | 5.74E-08 | Mari |
| cDC2 | SP110 | 8.76E-15 | 0.587475331 | 0.674 | 0.468 | 1.73E-10 | Mari |
| cDC2 | CISD2 | 1.03E-10 | 0.586915859 | 0.586 | 0.435 | 2.04E-06 | Mari |
| cDC2 | CCDC124 | 4.60E-07 | 0.586673861 | 0.44 | 0.329 | 0.009054235 | Mari |
| cDC2 | NDUFS21 | 2.65E-11 | 0.586082104 | 0.593 | 0.436 | 5.22E-07 | Mari |
| cDC2 | HLA-DMA | 8.12E-37 | 0.585971244 | 0.975 | 0.962 | 1.60E-32 | Mari |
| cDC2 | FUNDC2 | 5.82E-11 | 0.58519907 | 0.558 | 0.402 | 1.15E-06 | Mari |
| cDC2 | COMMD101 | 2.88E-14 | 0.584523095 | 0.583 | 0.395 | 5.67E-10 | Mari |
| cDC2 | ELOB | 1.94E-30 | 0.584428516 | 0.949 | 0.825 | 3.82E-26 | Mari |
| cDC2 | MED10 | 2.73E-11 | 0.584190258 | 0.537 | 0.382 | 5.37E-07 | Mari |
| cDC2 | ATP5MD | 6.83E-22 | 0.583777305 | 0.9 | 0.776 | 1.35E-17 | Mari |
| cDC2 | NQO2 | 6.12E-20 | 0.583409049 | 0.738 | 0.535 | 1.21E-15 | Mari |
| cDC2 | PPIE | 1.94E-06 | 0.583265017 | 0.308 | 0.208 | 0.03818752 | Mari |
| cDC2 | COPS5 | 2.13E-08 | 0.582996351 | 0.475 | 0.344 | 0.000418661 | Mari |
| cDC2 | EAF21 | 6.13E-11 | 0.582806499 | 0.5 | 0.341 | 1.21E-06 | Mari |
| cDC2 | PRPF31 | 8.29E-07 | 0.582753032 | 0.38 | 0.274 | 0.016315483 | Mari |
| cDC2 | HAX1 | 1.22E-17 | 0.581930066 | 0.662 | 0.452 | 2.41E-13 | Mari |
| cDC2 | ENTPD11 | 2.26E-19 | 0.579659686 | 0.743 | 0.516 | 4.44E-15 | Mari |
| cDC2 | SKA2 | 4.57E-07 | 0.579454732 | 0.299 | 0.196 | 0.008992415 | Mari |
| cDC2 | AKR1A1 | 8.24E-17 | 0.579113745 | 0.833 | 0.693 | 1.62E-12 | Mari |
| cDC2 | RBCK1 | 9.20E-09 | 0.578805747 | 0.56 | 0.429 | 0.000181137 | Mari |
| cDC2 | AOAH | 1.64E-12 | 0.578560961 | 0.634 | 0.453 | 3.24E-08 | Mari |
| cDC2 | IL10RB | 7.90E-09 | 0.578360733 | 0.442 | 0.31 | 0.000155544 | Mari |
| cDC2 | DHRS7 | 3.48E-12 | 0.578132704 | 0.588 | 0.423 | 6.85E-08 | Mari |
| cDC2 | SERPINB1 | 1.77E-26 | 0.57803064 | 0.961 | 0.859 | 3.49E-22 | Mari |
| cDC2 | CMC1 | 5.93E-07 | 0.57764497 | 0.426 | 0.312 | 0.011673676 | Mari |
| cDC2 | SNX2 | 1.87E-14 | 0.577242155 | 0.697 | 0.527 | 3.68E-10 | Mari |
| cDC2 | RBIS | 3.02E-20 | 0.575663517 | 0.826 | 0.655 | 5.95E-16 | Mari |
| cDC2 | ECH1 | 8.51E-11 | 0.57547286 | 0.602 | 0.452 | 1.68E-06 | Mari |
| cDC2 | IRF9 | 5.85E-13 | 0.575400614 | 0.597 | 0.424 | 1.15E-08 | Mari |
| cDC2 | TRAPPC2L | 4.74E-07 | 0.574665086 | 0.449 | 0.347 | 0.009326921 | Mari |
| cDC2 | HEIH | 4.82E-08 | 0.574319575 | 0.484 | 0.36 | 0.000949504 | Mari |
| cDC2 | DDX39B | 6.94E-13 | 0.573968089 | 0.706 | 0.541 | 1.37E-08 | Mari |
| cDC2 | MRPL15 | 7.26E-08 | 0.57389177 | 0.396 | 0.277 | 0.001429749 | Mari |
| cDC2 | SLC25A3 | 8.25E-31 | 0.573663633 | 0.951 | 0.876 | 1.62E-26 | Mari |
| cDC2 | TMEM14B | 1.27E-16 | 0.573573643 | 0.769 | 0.567 | 2.50E-12 | Mari |
| cDC2 | MT-ND4L | 1.24E-28 | 0.573250942 | 0.949 | 0.853 | 2.45E-24 | Mari |
| cDC2 | IGFBP7 | 1.13E-06 | 0.573214322 | 0.481 | 0.369 | 0.022268014 | Mari |
| cDC2 | FAM102B | 9.57E-11 | 0.5729501 | 0.618 | 0.447 | 1.88E-06 | Mari |
| cDC2 | CD48 | 6.21E-19 | 0.572825163 | 0.861 | 0.677 | 1.22E-14 | Mari |
| cDC2 | FPR31 | 2.00E-13 | 0.572715152 | 0.738 | 0.563 | 3.94E-09 | Mari |
| cDC2 | PSMB10 | 5.51E-19 | 0.572246059 | 0.891 | 0.689 | 1.08E-14 | Mari |
| cDC2 | ANAPC16 | 3.35E-12 | 0.572048573 | 0.644 | 0.481 | 6.59E-08 | Mari |
| cDC2 | NDUFS3 | 8.40E-10 | 0.571932508 | 0.565 | 0.42 | 1.65E-05 | Mari |
| cDC2 | CCDC50 | 2.01E-08 | 0.571742273 | 0.611 | 0.479 | 0.000395285 | Mari |

| | | | | | | | |
|---|---|---|---|---|---|---|---|
| cDC2 | GABARAP | 7.88E-41 | 0.570980939 | 0.968 | 0.946 | 1.55E-36 | Mari |
| cDC2 | ROMO1 | 7.19E-16 | 0.569497293 | 0.731 | 0.565 | 1.42E-11 | Mari |
| cDC2 | CIAO2A | 2.79E-14 | 0.569438042 | 0.671 | 0.492 | 5.49E-10 | Mari |
| cDC2 | RAB32 | 6.20E-11 | 0.569339425 | 0.708 | 0.59 | 1.22E-06 | Mari |
| cDC2 | TMSB4X | 2.14E-42 | 0.569206124 | 1 | 0.992 | 4.21E-38 | Mari |
| cDC2 | POLD4 | 4.48E-15 | 0.568621416 | 0.787 | 0.641 | 8.81E-11 | Mari |
| cDC2 | EIF2S1 | 2.86E-07 | 0.567362485 | 0.507 | 0.395 | 0.005620342 | Mari |
| cDC2 | HLA-DRB1 | 5.82E-30 | 0.566841914 | 1 | 0.993 | 1.15E-25 | Mari |
| cDC2 | OSGEP | 3.10E-08 | 0.566818933 | 0.454 | 0.334 | 0.0006104 | Mari |
| cDC2 | SSNA1 | 4.21E-12 | 0.566572384 | 0.646 | 0.507 | 8.28E-08 | Mari |
| cDC2 | GLRX3 | 7.69E-10 | 0.566284721 | 0.613 | 0.478 | 1.51E-05 | Mari |
| cDC2 | SF3B6 | 2.36E-19 | 0.565221846 | 0.806 | 0.64 | 4.65E-15 | Mari |
| cDC2 | EFHD2 | 6.72E-21 | 0.565069014 | 0.947 | 0.882 | 1.32E-16 | Mari |
| cDC2 | MGST2 | 7.13E-08 | 0.564645035 | 0.537 | 0.416 | 0.00140368 | Mari |
| cDC2 | NDUFAF3 | 5.57E-09 | 0.564206555 | 0.567 | 0.433 | 0.000109685 | Mari |
| cDC2 | PSMA1 | 4.17E-18 | 0.562260907 | 0.863 | 0.712 | 8.21E-14 | Mari |
| cDC2 | CAT | 1.06E-07 | 0.561725416 | 0.59 | 0.458 | 0.00208147 | Mari |
| cDC2 | ABRACL | 3.69E-12 | 0.561417664 | 0.641 | 0.487 | 7.25E-08 | Mari |
| cDC2 | ARF5 | 3.79E-21 | 0.561088856 | 0.861 | 0.743 | 7.46E-17 | Mari |
| cDC2 | POLR2F | 1.82E-07 | 0.559895489 | 0.454 | 0.337 | 0.003591644 | Mari |
| cDC2 | CHMP3 | 2.59E-07 | 0.559809907 | 0.486 | 0.368 | 0.005095412 | Mari |
| cDC2 | ARHGDIA | 1.13E-14 | 0.559733628 | 0.787 | 0.621 | 2.23E-10 | Mari |
| cDC2 | COA8 | 4.06E-07 | 0.559648497 | 0.375 | 0.258 | 0.007988323 | Mari |
| cDC2 | OCIAD1 | 6.30E-14 | 0.557768836 | 0.711 | 0.522 | 1.24E-09 | Mari |
| cDC2 | TEX141 | 9.87E-08 | 0.557118697 | 0.595 | 0.45 | 0.001942171 | Mari |
| cDC2 | SMIM12 | 6.97E-08 | 0.556071077 | 0.428 | 0.306 | 0.001372199 | Mari |
| cDC2 | SSR4 | 3.82E-29 | 0.556042777 | 0.935 | 0.855 | 7.53E-25 | Mari |
| cDC2 | MDH2 | 1.83E-17 | 0.555808711 | 0.794 | 0.629 | 3.60E-13 | Mari |
| cDC2 | VMA21 | 7.86E-12 | 0.555763508 | 0.6 | 0.431 | 1.55E-07 | Mari |
| cDC2 | PSMB7 | 9.51E-15 | 0.555417539 | 0.731 | 0.544 | 1.87E-10 | Mari |
| cDC2 | RUBCNL | 1.77E-06 | 0.555101572 | 0.266 | 0.166 | 0.034897742 | Mari |
| cDC2 | MRPL11 | 2.24E-10 | 0.554696664 | 0.593 | 0.448 | 4.42E-06 | Mari |
| cDC2 | PAIP2 | 3.81E-14 | 0.55458752 | 0.769 | 0.603 | 7.50E-10 | Mari |
| cDC2 | IFT20 | 1.48E-07 | 0.55414256 | 0.472 | 0.354 | 0.002915246 | Mari |
| cDC2 | SDHD | 4.98E-13 | 0.553812741 | 0.685 | 0.51 | 9.80E-09 | Mari |
| cDC2 | MCUB | 4.24E-09 | 0.552281693 | 0.6 | 0.472 | 8.34E-05 | Mari |
| cDC2 | BTF3L4 | 4.81E-11 | 0.55116317 | 0.583 | 0.432 | 9.48E-07 | Mari |
| cDC2 | GSTK1 | 2.86E-20 | 0.5500139 | 0.896 | 0.777 | 5.62E-16 | Mari |
| cDC2 | LSM7 | 5.86E-19 | 0.549383887 | 0.856 | 0.733 | 1.15E-14 | Mari |
| cDC2 | CEBPD | 6.22E-22 | 0.548977409 | 0.931 | 0.789 | 1.22E-17 | Mari |
| cDC2 | FKBP3 | 2.72E-07 | 0.548463344 | 0.414 | 0.296 | 0.005361248 | Mari |
| cDC2 | SNRPC | 2.81E-09 | 0.547854037 | 0.519 | 0.377 | 5.53E-05 | Mari |
| cDC2 | COTL1 | 7.34E-33 | 0.54751095 | 0.977 | 0.962 | 1.44E-28 | Mari |
| cDC2 | CNDP21 | 1.54E-10 | 0.546955301 | 0.669 | 0.526 | 3.03E-06 | Mari |
| cDC2 | SPCS1 | 5.41E-20 | 0.546897782 | 0.831 | 0.67 | 1.07E-15 | Mari |
| cDC2 | ATRAID | 9.65E-12 | 0.546704068 | 0.586 | 0.432 | 1.90E-07 | Mari |
| cDC2 | ATP5MC3 | 2.21E-24 | 0.545342991 | 0.928 | 0.827 | 4.35E-20 | Mari |
| cDC2 | CALM3 | 1.86E-09 | 0.543685823 | 0.646 | 0.509 | 3.65E-05 | Mari |
| cDC2 | CHURC1 | 3.16E-12 | 0.543513623 | 0.729 | 0.576 | 6.23E-08 | Mari |
| cDC2 | COMMD1 | 1.72E-14 | 0.542050913 | 0.694 | 0.484 | 3.39E-10 | Mari |
| cDC2 | ATP9B1 | 3.43E-07 | 0.54162816 | 0.539 | 0.41 | 0.006756736 | Mari |
| cDC2 | DDRGK1 | 3.35E-07 | 0.541280714 | 0.424 | 0.31 | 0.006591776 | Mari |
| cDC2 | LST1 | 3.45E-30 | 0.541214563 | 0.965 | 0.896 | 6.80E-26 | Mari |
| cDC2 | EIF2S21 | 5.72E-11 | 0.54106877 | 0.671 | 0.506 | 1.12E-06 | Mari |
| cDC2 | ATP6V0E1 | 4.58E-24 | 0.540833947 | 0.919 | 0.827 | 9.02E-20 | Mari |
| cDC2 | RRP36 | 1.04E-06 | 0.540713123 | 0.389 | 0.281 | 0.020525181 | Mari |

| | | | | | | | |
|---|---|---|---|---|---|---|---|
| cDC2 | NXT1 | 3.82E-13 | 0.539208898 | 0.604 | 0.415 | 7.52E-09 | Mari |
| cDC2 | TMEM258 | 1.21E-20 | 0.536563528 | 0.898 | 0.745 | 2.38E-16 | Mari |
| cDC2 | ATOX1 | 5.59E-12 | 0.536307921 | 0.87 | 0.761 | 1.10E-07 | Mari |
| cDC2 | CHMP5 | 1.60E-09 | 0.535056803 | 0.644 | 0.504 | 3.15E-05 | Mari |
| cDC2 | PSMA7 | 4.79E-25 | 0.534074139 | 0.942 | 0.838 | 9.43E-21 | Mari |
| cDC2 | PGLS | 2.99E-24 | 0.533808209 | 0.914 | 0.787 | 5.88E-20 | Mari |
| cDC2 | CNPY3 | 3.62E-10 | 0.533391268 | 0.641 | 0.488 | 7.12E-06 | Mari |
| cDC2 | CHCHD7 | 2.93E-09 | 0.533023697 | 0.481 | 0.345 | 5.77E-05 | Mari |
| cDC2 | LSM3 | 5.38E-11 | 0.5324357 | 0.655 | 0.503 | 1.06E-06 | Mari |
| cDC2 | ATP5IF1 | 3.87E-16 | 0.530920618 | 0.789 | 0.62 | 7.62E-12 | Mari |
| cDC2 | CCT8 | 2.89E-15 | 0.530155593 | 0.787 | 0.634 | 5.70E-11 | Mari |
| cDC2 | TIMM13 | 1.09E-09 | 0.530082639 | 0.63 | 0.501 | 2.14E-05 | Mari |
| cDC2 | BUD23 | 1.35E-10 | 0.528965697 | 0.653 | 0.503 | 2.65E-06 | Mari |
| cDC2 | MRPS34 | 9.01E-13 | 0.527859223 | 0.623 | 0.438 | 1.77E-08 | Mari |
| cDC2 | ATP6V1F | 1.26E-18 | 0.52780148 | 0.917 | 0.794 | 2.48E-14 | Mari |
| cDC2 | PUF60 | 1.69E-09 | 0.527359425 | 0.532 | 0.386 | 3.33E-05 | Mari |
| cDC2 | VSIR | 4.46E-11 | 0.527048942 | 0.713 | 0.584 | 8.79E-07 | Mari |
| cDC2 | SMDT1 | 4.07E-19 | 0.526785508 | 0.905 | 0.792 | 8.01E-15 | Mari |
| cDC2 | TALDO1 | 2.89E-17 | 0.526427855 | 0.861 | 0.76 | 5.68E-13 | Mari |
| cDC2 | OST4 | 1.89E-27 | 0.525439891 | 0.935 | 0.854 | 3.72E-23 | Mari |
| cDC2 | NDUFV2 | 1.31E-26 | 0.52543385 | 0.919 | 0.822 | 2.58E-22 | Mari |
| cDC2 | CMTM71 | 8.03E-12 | 0.52535422 | 0.657 | 0.477 | 1.58E-07 | Mari |
| cDC2 | ACTR10 | 2.47E-06 | 0.524140759 | 0.449 | 0.346 | 0.048558999 | Mari |
| cDC2 | FAM192A1 | 8.83E-07 | 0.524073248 | 0.428 | 0.315 | 0.017375481 | Mari |
| cDC2 | VKORC1 | 2.67E-08 | 0.523167658 | 0.495 | 0.362 | 0.000526493 | Mari |
| cDC2 | DNAJC15 | 5.21E-17 | 0.522761653 | 0.822 | 0.642 | 1.03E-12 | Mari |
| cDC2 | DBNDD2 | 1.79E-07 | 0.5227101 | 0.6 | 0.489 | 0.003526291 | Mari |
| cDC2 | ARPC4 | 1.39E-14 | 0.522154951 | 0.831 | 0.661 | 2.73E-10 | Mari |
| cDC2 | VAMP2 | 1.12E-08 | 0.519540346 | 0.47 | 0.337 | 0.000219613 | Mari |
| cDC2 | LPCAT2 | 4.77E-12 | 0.519441235 | 0.676 | 0.5 | 9.40E-08 | Mari |
| cDC2 | ARF6 | 3.73E-13 | 0.51941488 | 0.854 | 0.715 | 7.34E-09 | Mari |
| cDC2 | CWC15 | 1.14E-10 | 0.518659714 | 0.546 | 0.39 | 2.25E-06 | Mari |
| cDC2 | HLA-DQB1 | 1.69E-29 | 0.518591548 | 0.991 | 0.977 | 3.32E-25 | Mari |
| cDC2 | SNHG12 | 1.78E-06 | 0.5180118 | 0.481 | 0.376 | 0.035058045 | Mari |
| cDC2 | ACAA1 | 6.55E-14 | 0.517076981 | 0.78 | 0.617 | 1.29E-09 | Mari |
| cDC2 | MICOS13 | 2.23E-15 | 0.516946994 | 0.815 | 0.637 | 4.38E-11 | Mari |
| cDC2 | MPC2 | 1.05E-10 | 0.515960826 | 0.604 | 0.451 | 2.07E-06 | Mari |
| cDC2 | PCMT1 | 1.04E-08 | 0.515711136 | 0.6 | 0.464 | 0.000205136 | Mari |
| cDC2 | MRPL471 | 4.25E-08 | 0.51504402 | 0.525 | 0.389 | 0.000837253 | Mari |
| cDC2 | SRP14 | 1.09E-34 | 0.514716114 | 0.956 | 0.913 | 2.15E-30 | Mari |
| cDC2 | COX6C | 3.22E-24 | 0.513594071 | 0.94 | 0.832 | 6.35E-20 | Mari |
| cDC2 | MEF2A1 | 3.70E-18 | 0.510950746 | 0.87 | 0.678 | 7.29E-14 | Mari |
| cDC2 | SELENOW | 2.59E-12 | 0.510264729 | 0.718 | 0.576 | 5.11E-08 | Mari |
| cDC2 | KRTCAP2 | 1.19E-16 | 0.509389974 | 0.833 | 0.671 | 2.35E-12 | Mari |
| cDC2 | COMMD6 | 1.02E-25 | 0.508547635 | 0.933 | 0.86 | 2.00E-21 | Mari |
| cDC2 | MRFAP1 | 8.22E-09 | 0.50847077 | 0.588 | 0.452 | 0.000161866 | Mari |
| cDC2 | NDUFA2 | 5.66E-16 | 0.507771974 | 0.796 | 0.611 | 1.11E-11 | Mari |
| cDC2 | LRPAP1 | 1.92E-09 | 0.507379541 | 0.56 | 0.407 | 3.77E-05 | Mari |
| cDC2 | SNRPF | 1.24E-18 | 0.506952138 | 0.859 | 0.686 | 2.45E-14 | Mari |
| cDC2 | FIS1 | 2.40E-15 | 0.506023302 | 0.817 | 0.66 | 4.72E-11 | Mari |
| cDC2 | TMED10 | 2.89E-08 | 0.505592491 | 0.699 | 0.576 | 0.000568126 | Mari |
| cDC2 | SH3BGRL3 | 1.08E-30 | 0.505271966 | 0.979 | 0.968 | 2.12E-26 | Mari |
| cDC2 | MFSD10 | 2.14E-06 | 0.503956025 | 0.382 | 0.275 | 0.042162747 | Mari |
| cDC2 | HLA-DPA1 | 3.10E-28 | 0.503681048 | 0.998 | 0.984 | 6.11E-24 | Mari |
| cDC2 | DPYSL2 | 3.35E-13 | 0.501890281 | 0.752 | 0.582 | 6.59E-09 | Mari |
| cDC2 | EIF6 | 3.49E-11 | 0.500616401 | 0.731 | 0.605 | 6.87E-07 | Mari |

| | | | | | | | |
|---|---|---|---|---|---|---|---|
| cDC2 | MMP12 | 7.25E-39 | 3.634403276 | 0.262 | 0.065 | 1.43E-34 | Tb |
| cDC2 | CCL17 | 2.45E-30 | 3.291621515 | 0.25 | 0.073 | 4.82E-26 | Tb |
| cDC2 | KRT1 | 1.99E-29 | 3.153254269 | 0.277 | 0.097 | 3.91E-25 | Tb |
| cDC2 | NPHS1 | 1.32E-36 | 3.045274789 | 0.333 | 0.115 | 2.59E-32 | Tb |
| cDC2 | AQP3 | 1.11E-55 | 2.446616354 | 0.368 | 0.089 | 2.18E-51 | Tb |
| cDC2 | S100A7 | 4.14E-62 | 2.415127645 | 0.431 | 0.113 | 8.16E-58 | Tb |
| cDC2 | KRTDAP | 1.02E-28 | 2.253013413 | 0.321 | 0.124 | 2.00E-24 | Tb |
| cDC2 | FBP1 | 7.79E-23 | 2.204595564 | 0.613 | 0.442 | 1.53E-18 | Tb |
| cDC2 | KRT6A | 2.71E-62 | 2.154036042 | 0.408 | 0.093 | 5.34E-58 | Tb |
| cDC2 | GCNT2 | 1.11E-16 | 2.13913858 | 0.25 | 0.109 | 2.18E-12 | Tb |
| cDC2 | TAF4B | 1.04E-29 | 2.138919476 | 0.426 | 0.19 | 2.05E-25 | Tb |
| cDC2 | HOMER1 | 2.38E-57 | 2.051975791 | 0.583 | 0.245 | 4.68E-53 | Tb |
| cDC2 | ARL17A | 5.89E-34 | 2.048006477 | 0.271 | 0.074 | 1.16E-29 | Tb |
| cDC2 | CD1A | 7.77E-36 | 2.007181807 | 0.358 | 0.127 | 1.53E-31 | Tb |
| cDC2 | ADAM19 | 1.96E-58 | 1.989219972 | 0.616 | 0.263 | 3.87E-54 | Tb |
| cDC2 | NARS2 | 5.13E-22 | 1.903545634 | 0.34 | 0.154 | 1.01E-17 | Tb |
| cDC2 | MIR99AHG | 1.77E-25 | 1.884728961 | 0.276 | 0.099 | 3.48E-21 | Tb |
| cDC2 | DMKN | 2.70E-18 | 1.850534065 | 0.258 | 0.113 | 5.32E-14 | Tb |
| cDC2 | RAB30 | 8.27E-43 | 1.836595614 | 0.394 | 0.128 | 1.63E-38 | Tb |
| cDC2 | ZNF366 | 4.51E-35 | 1.833630648 | 0.34 | 0.113 | 8.88E-31 | Tb |
| cDC2 | PERP | 2.87E-29 | 1.718593729 | 0.271 | 0.086 | 5.65E-25 | Tb |
| cDC2 | ZNF516 | 2.29E-41 | 1.71823837 | 0.579 | 0.27 | 4.50E-37 | Tb |
| cDC2 | CCR6 | 2.35E-24 | 1.710651614 | 0.298 | 0.116 | 4.63E-20 | Tb |
| cDC2 | KRT6B | 2.85E-29 | 1.649994925 | 0.269 | 0.08 | 5.61E-25 | Tb |
| cDC2 | SLC35F1 | 1.22E-28 | 1.643538937 | 0.363 | 0.153 | 2.39E-24 | Tb |
| cDC2 | MCF2L2 | 8.68E-37 | 1.641167494 | 0.569 | 0.285 | 1.71E-32 | Tb |
| cDC2 | RASAL2 | 5.00E-42 | 1.623279291 | 0.545 | 0.251 | 9.84E-38 | Tb |
| cDC2 | CD226 | 1.30E-34 | 1.609785242 | 0.431 | 0.181 | 2.56E-30 | Tb |
| cDC2 | KRT5 | 1.83E-26 | 1.602448594 | 0.281 | 0.097 | 3.59E-22 | Tb |
| cDC2 | QPCTL | 1.92E-28 | 1.599774239 | 0.258 | 0.078 | 3.79E-24 | Tb |
| cDC2 | HES1 | 6.20E-31 | 1.597481146 | 0.393 | 0.161 | 1.22E-26 | Tb |
| cDC2 | PSMD5 | 8.90E-14 | 1.589532508 | 0.284 | 0.145 | 1.75E-09 | Tb |
| cDC2 | RHOH | 2.90E-43 | 1.584603599 | 0.585 | 0.277 | 5.71E-39 | Tb |
| cDC2 | PHLPP1 | 9.02E-33 | 1.571789423 | 0.6 | 0.341 | 1.77E-28 | Tb |
| cDC2 | PFKP | 4.11E-33 | 1.565766761 | 0.518 | 0.263 | 8.10E-29 | Tb |
| cDC2 | NEK1 | 3.74E-19 | 1.561578989 | 0.407 | 0.211 | 7.36E-15 | Tb |
| cDC2 | PPP1R16B | 7.64E-17 | 1.547498432 | 0.255 | 0.112 | 1.50E-12 | Tb |
| cDC2 | GGT5 | 9.28E-42 | 1.539478396 | 0.314 | 0.08 | 1.83E-37 | Tb |
| cDC2 | SPG11 | 7.95E-36 | 1.536974708 | 0.51 | 0.232 | 1.57E-31 | Tb |
| cDC2 | PRAG1 | 4.74E-31 | 1.527257575 | 0.291 | 0.09 | 9.34E-27 | Tb |
| cDC2 | SFN | 1.45E-29 | 1.510797686 | 0.349 | 0.131 | 2.85E-25 | Tb |
| cDC2 | GYS1 | 2.34E-24 | 1.507948619 | 0.26 | 0.088 | 4.60E-20 | Tb |
| cDC2 | MIR3945HG | 3.81E-24 | 1.503749867 | 0.375 | 0.174 | 7.50E-20 | Tb |
| cDC2 | ARAP2 | 8.35E-31 | 1.471202848 | 0.59 | 0.323 | 1.64E-26 | Tb |
| cDC2 | PRRG4 | 1.64E-22 | 1.470804238 | 0.265 | 0.098 | 3.23E-18 | Tb |
| cDC2 | ZNF10 | 6.44E-28 | 1.462614238 | 0.386 | 0.161 | 1.27E-23 | Tb |
| cDC2 | LIPA | 3.26E-11 | 1.45641395 | 0.316 | 0.186 | 6.42E-07 | Tb |
| cDC2 | JMY | 1.35E-18 | 1.449361575 | 0.518 | 0.319 | 2.66E-14 | Tb |
| cDC2 | BATF3 | 4.17E-17 | 1.443253497 | 0.267 | 0.116 | 8.21E-13 | Tb |
| cDC2 | AZIN1-AS1 | 5.68E-19 | 1.438977 | 0.332 | 0.156 | 1.12E-14 | Tb |
| cDC2 | WDFY4 | 6.41E-44 | 1.437765057 | 0.611 | 0.279 | 1.26E-39 | Tb |
| cDC2 | RGS13 | 1.51E-16 | 1.435305603 | 0.284 | 0.137 | 2.97E-12 | Tb |
| cDC2 | INHBA | 1.73E-13 | 1.428303515 | 0.461 | 0.317 | 3.40E-09 | Tb |
| cDC2 | KRT14 | 4.71E-65 | 1.42790675 | 0.654 | 0.238 | 9.26E-61 | Tb |
| cDC2 | TMTC2 | 2.58E-15 | 1.419749367 | 0.305 | 0.15 | 5.08E-11 | Tb |
| cDC2 | LRRK1 | 3.93E-33 | 1.414431979 | 0.569 | 0.292 | 7.74E-29 | Tb |

| | | | | | | | |
|---|---|---|---|---|---|---|---|
| cDC2 | DENND1B | 5.65E-67 | 1.412707487 | 0.909 | 0.582 | 1.11E-62 | Tb |
| cDC2 | UBAP2 | 4.51E-34 | 1.407491699 | 0.471 | 0.2 | 8.87E-30 | Tb |
| cDC2 | MT-CO3 | 3.54E-86 | 1.406448303 | 0.983 | 0.898 | 6.97E-82 | Tb |
| cDC2 | IL1RAP | 1.06E-34 | 1.403765647 | 0.457 | 0.192 | 2.10E-30 | Tb |
| cDC2 | ZBTB46 | 2.21E-31 | 1.398835829 | 0.543 | 0.278 | 4.35E-27 | Tb |
| cDC2 | KAT2B | 3.74E-31 | 1.397260437 | 0.55 | 0.278 | 7.35E-27 | Tb |
| cDC2 | XIST | 4.81E-49 | 1.391681234 | 0.578 | 0.235 | 9.46E-45 | Tb |
| cDC2 | ZNF609 | 3.69E-30 | 1.391612818 | 0.471 | 0.221 | 7.27E-26 | Tb |
| cDC2 | EPOP | 2.04E-27 | 1.38418877 | 0.269 | 0.085 | 4.01E-23 | Tb |
| cDC2 | CLCN3 | 5.84E-18 | 1.383277275 | 0.414 | 0.22 | 1.15E-13 | Tb |
| cDC2 | MT-ATP6 | 7.28E-93 | 1.378105737 | 0.99 | 0.881 | 1.43E-88 | Tb |
| cDC2 | CLEC16A | 4.59E-30 | 1.376407819 | 0.541 | 0.271 | 9.04E-26 | Tb |
| cDC2 | ST6GAL1 | 1.17E-26 | 1.376266904 | 0.422 | 0.189 | 2.31E-22 | Tb |
| cDC2 | IQCG | 3.37E-21 | 1.366474245 | 0.305 | 0.129 | 6.63E-17 | Tb |
| cDC2 | UST | 1.04E-35 | 1.358387315 | 0.717 | 0.45 | 2.04E-31 | Tb |
| cDC2 | MGA | 1.14E-17 | 1.357828026 | 0.291 | 0.132 | 2.24E-13 | Tb |
| cDC2 | P2RY14 | 6.09E-29 | 1.349951656 | 0.565 | 0.301 | 1.20E-24 | Tb |
| cDC2 | KRT17 | 1.26E-28 | 1.349799879 | 0.26 | 0.077 | 2.48E-24 | Tb |
| cDC2 | CLIC4 | 3.41E-30 | 1.348284973 | 0.67 | 0.414 | 6.72E-26 | Tb |
| cDC2 | L3MBTL4 | 4.31E-14 | 1.344306705 | 0.284 | 0.14 | 8.48E-10 | Tb |
| cDC2 | DLEU2 | 1.36E-40 | 1.334309825 | 0.825 | 0.562 | 2.67E-36 | Tb |
| cDC2 | MCU | 1.53E-18 | 1.333786274 | 0.353 | 0.169 | 3.01E-14 | Tb |
| cDC2 | TMEM184C | 6.22E-23 | 1.332391903 | 0.375 | 0.172 | 1.23E-18 | Tb |
| cDC2 | TSPOAP1 | 1.65E-25 | 1.325224701 | 0.279 | 0.097 | 3.24E-21 | Tb |
| cDC2 | CARD11 | 1.49E-20 | 1.324880505 | 0.279 | 0.111 | 2.93E-16 | Tb |
| cDC2 | TFCP2 | 1.20E-28 | 1.315160145 | 0.407 | 0.178 | 2.36E-24 | Tb |
| cDC2 | FOXO1 | 4.94E-15 | 1.313345258 | 0.44 | 0.263 | 9.72E-11 | Tb |
| cDC2 | VAC14 | 1.71E-31 | 1.312192564 | 0.478 | 0.214 | 3.37E-27 | Tb |
| cDC2 | ABL1 | 5.35E-18 | 1.311861567 | 0.34 | 0.167 | 1.05E-13 | Tb |
| cDC2 | GAB1 | 3.49E-12 | 1.305861836 | 0.253 | 0.127 | 6.86E-08 | Tb |
| cDC2 | SOCS1 | 6.76E-31 | 1.305287982 | 0.661 | 0.384 | 1.33E-26 | Tb |
| cDC2 | KRT16 | 1.35E-42 | 1.305087018 | 0.403 | 0.123 | 2.66E-38 | Tb |
| cDC2 | CISH | 1.11E-34 | 1.30163292 | 0.351 | 0.113 | 2.19E-30 | Tb |
| cDC2 | WDR37 | 3.66E-35 | 1.300264273 | 0.51 | 0.226 | 7.21E-31 | Tb |
| cDC2 | NSMCE2 | 6.43E-41 | 1.299536526 | 0.672 | 0.36 | 1.27E-36 | Tb |
| cDC2 | BRCA2 | 3.40E-23 | 1.298871505 | 0.436 | 0.218 | 6.69E-19 | Tb |
| cDC2 | HS3ST3B1 | 3.27E-13 | 1.295279698 | 0.272 | 0.14 | 6.44E-09 | Tb |
| cDC2 | TXLNA | 2.70E-27 | 1.291713043 | 0.391 | 0.162 | 5.31E-23 | Tb |
| cDC2 | PPARD | 3.56E-42 | 1.291206584 | 0.707 | 0.379 | 7.01E-38 | Tb |
| cDC2 | CEP192 | 1.16E-19 | 1.290252354 | 0.284 | 0.118 | 2.28E-15 | Tb |
| cDC2 | RNF19A | 1.37E-29 | 1.287463263 | 0.583 | 0.317 | 2.69E-25 | Tb |
| cDC2 | S100A2 | 1.83E-41 | 1.286085002 | 0.351 | 0.099 | 3.61E-37 | Tb |
| cDC2 | SLC39A9 | 2.25E-22 | 1.284397625 | 0.277 | 0.104 | 4.43E-18 | Tb |
| cDC2 | FTO | 4.10E-16 | 1.28119321 | 0.298 | 0.138 | 8.07E-12 | Tb |
| cDC2 | PALD1 | 1.03E-30 | 1.280772612 | 0.415 | 0.169 | 2.02E-26 | Tb |
| cDC2 | KCNK6 | 1.10E-40 | 1.277791824 | 0.503 | 0.206 | 2.17E-36 | Tb |
| cDC2 | GBF1 | 2.29E-38 | 1.276838573 | 0.483 | 0.199 | 4.50E-34 | Tb |
| cDC2 | SLC33A1 | 4.95E-17 | 1.275234916 | 0.286 | 0.128 | 9.74E-13 | Tb |
| cDC2 | SFMBT2 | 1.60E-40 | 1.274641832 | 0.693 | 0.362 | 3.15E-36 | Tb |
| cDC2 | VASH1 | 1.64E-40 | 1.272938975 | 0.578 | 0.262 | 3.23E-36 | Tb |
| cDC2 | LONP1 | 5.47E-19 | 1.271577115 | 0.25 | 0.097 | 1.08E-14 | Tb |
| cDC2 | ITGAM | 7.28E-11 | 1.266908184 | 0.325 | 0.201 | 1.43E-06 | Tb |
| cDC2 | S100A8 | 5.22E-80 | 1.266841552 | 0.705 | 0.251 | 1.03E-75 | Tb |
| cDC2 | UBXN2A | 1.96E-28 | 1.265208967 | 0.473 | 0.229 | 3.86E-24 | Tb |
| cDC2 | CYTH3 | 4.24E-15 | 1.261176769 | 0.29 | 0.137 | 8.34E-11 | Tb |
| cDC2 | HDAC4 | 2.32E-33 | 1.257118535 | 0.494 | 0.214 | 4.57E-29 | Tb |

| | | | | | | | |
|---|---|---|---|---|---|---|---|
| cDC2 | NLRC5 | 4.46E-28 | 1.253962737 | 0.436 | 0.197 | 8.79E-24 | Tb |
| cDC2 | LRRC23 | 2.12E-26 | 1.251823975 | 0.48 | 0.235 | 4.18E-22 | Tb |
| cDC2 | XKR6 | 1.40E-22 | 1.251590977 | 0.26 | 0.092 | 2.75E-18 | Tb |
| cDC2 | IPCEF1 | 1.54E-24 | 1.25129074 | 0.462 | 0.239 | 3.03E-20 | Tb |
| cDC2 | NCR3LG1 | 2.27E-26 | 1.248192383 | 0.407 | 0.181 | 4.46E-22 | Tb |
| cDC2 | ARIH1 | 1.46E-36 | 1.245461654 | 0.88 | 0.647 | 2.88E-32 | Tb |
| cDC2 | ELOVL7 | 8.72E-14 | 1.242311196 | 0.302 | 0.158 | 1.72E-09 | Tb |
| cDC2 | VPS13B | 1.38E-40 | 1.235556886 | 0.649 | 0.336 | 2.72E-36 | Tb |
| cDC2 | TRIM38 | 2.86E-43 | 1.235180104 | 0.693 | 0.37 | 5.62E-39 | Tb |
| cDC2 | PRMT9 | 1.60E-37 | 1.2347937 | 0.698 | 0.396 | 3.15E-33 | Tb |
| cDC2 | TRAK1 | 9.23E-30 | 1.234239418 | 0.709 | 0.459 | 1.82E-25 | Tb |
| cDC2 | ENPP2 | 3.24E-26 | 1.230113528 | 0.354 | 0.144 | 6.39E-22 | Tb |
| cDC2 | SLC27A3 | 5.74E-20 | 1.229824519 | 0.368 | 0.181 | 1.13E-15 | Tb |
| cDC2 | MT-CYB | 5.33E-58 | 1.228516224 | 0.974 | 0.864 | 1.05E-53 | Tb |
| cDC2 | VDR | 2.85E-33 | 1.225888076 | 0.56 | 0.266 | 5.61E-29 | Tb |
| cDC2 | PDCD1LG2 | 9.36E-23 | 1.22067861 | 0.356 | 0.157 | 1.84E-18 | Tb |
| cDC2 | KIF1B | 3.64E-30 | 1.217286551 | 0.625 | 0.369 | 7.17E-26 | Tb |
| cDC2 | KLHL2 | 1.08E-19 | 1.217215427 | 0.361 | 0.169 | 2.13E-15 | Tb |
| cDC2 | PLPP1 | 1.84E-17 | 1.21279528 | 0.279 | 0.119 | 3.63E-13 | Tb |
| cDC2 | TRABD2A | 4.02E-22 | 1.21078528 | 0.262 | 0.094 | 7.92E-18 | Tb |
| cDC2 | CASS4 | 1.44E-24 | 1.210088343 | 0.454 | 0.229 | 2.84E-20 | Tb |
| cDC2 | MOB3B | 1.22E-39 | 1.206679733 | 0.759 | 0.469 | 2.41E-35 | Tb |
| cDC2 | APOL2 | 2.04E-15 | 1.205659972 | 0.255 | 0.113 | 4.01E-11 | Tb |
| cDC2 | EEF2K | 5.59E-20 | 1.20298264 | 0.339 | 0.156 | 1.10E-15 | Tb |
| cDC2 | FAM222B | 9.86E-19 | 1.202272433 | 0.255 | 0.102 | 1.94E-14 | Tb |
| cDC2 | APOL4 | 2.09E-31 | 1.196210055 | 0.354 | 0.122 | 4.11E-27 | Tb |
| cDC2 | TCOF1 | 3.51E-13 | 1.194810817 | 0.307 | 0.16 | 6.91E-09 | Tb |
| cDC2 | SLA | 4.60E-26 | 1.194084261 | 0.393 | 0.169 | 9.05E-22 | Tb |
| cDC2 | ABR | 5.53E-32 | 1.187298682 | 0.796 | 0.545 | 1.09E-27 | Tb |
| cDC2 | PHTF2 | 1.17E-21 | 1.181847335 | 0.492 | 0.268 | 2.31E-17 | Tb |
| cDC2 | PHTF1 | 5.25E-18 | 1.181127408 | 0.37 | 0.188 | 1.03E-13 | Tb |
| cDC2 | STAT4 | 3.28E-23 | 1.180725411 | 0.534 | 0.312 | 6.45E-19 | Tb |
| cDC2 | RNF122 | 4.85E-19 | 1.18008494 | 0.274 | 0.115 | 9.55E-15 | Tb |
| cDC2 | LRRC8D | 1.37E-22 | 1.179724 | 0.398 | 0.191 | 2.70E-18 | Tb |
| cDC2 | KDM6A | 1.13E-32 | 1.177900945 | 0.66 | 0.369 | 2.22E-28 | Tb |
| cDC2 | TNS3 | 4.93E-34 | 1.177385034 | 0.555 | 0.275 | 9.71E-30 | Tb |
| cDC2 | DENND4A | 5.38E-23 | 1.176693311 | 0.85 | 0.695 | 1.06E-18 | Tb |
| cDC2 | OXSM | 1.80E-18 | 1.170715894 | 0.33 | 0.151 | 3.55E-14 | Tb |
| cDC2 | MYO9A | 3.79E-19 | 1.170571839 | 0.312 | 0.14 | 7.45E-15 | Tb |
| cDC2 | NFX1 | 5.81E-23 | 1.170002392 | 0.429 | 0.207 | 1.14E-18 | Tb |
| cDC2 | PATL2 | 2.35E-22 | 1.168547373 | 0.311 | 0.125 | 4.62E-18 | Tb |
| cDC2 | ILF3-DT | 1.64E-18 | 1.167602504 | 0.347 | 0.163 | 3.22E-14 | Tb |
| cDC2 | LRBA | 2.33E-24 | 1.166183405 | 0.508 | 0.264 | 4.59E-20 | Tb |
| cDC2 | ST3GAL5 | 2.26E-35 | 1.161372324 | 0.529 | 0.232 | 4.46E-31 | Tb |
| cDC2 | KRT10 | 3.01E-16 | 1.157347867 | 0.726 | 0.509 | 5.92E-12 | Tb |
| cDC2 | PPARGC1B | 8.10E-21 | 1.156136858 | 0.265 | 0.103 | 1.59E-16 | Tb |
| cDC2 | CENPP | 2.72E-14 | 1.152968392 | 0.265 | 0.127 | 5.35E-10 | Tb |
| cDC2 | RALGAPA1 | 3.35E-24 | 1.150845121 | 0.628 | 0.388 | 6.59E-20 | Tb |
| cDC2 | IRS2 | 7.17E-15 | 1.15042855 | 0.318 | 0.164 | 1.41E-10 | Tb |
| cDC2 | SPDYA | 1.14E-15 | 1.149040132 | 0.332 | 0.175 | 2.25E-11 | Tb |
| cDC2 | SAMD4A | 2.88E-13 | 1.147855342 | 0.267 | 0.131 | 5.67E-09 | Tb |
| cDC2 | LYRM1 | 2.81E-23 | 1.147403028 | 0.309 | 0.119 | 5.53E-19 | Tb |
| cDC2 | SLC39A11 | 1.34E-25 | 1.145093929 | 0.429 | 0.2 | 2.63E-21 | Tb |
| cDC2 | WDPCP | 1.64E-23 | 1.140854464 | 0.297 | 0.112 | 3.22E-19 | Tb |
| cDC2 | ZNF710 | 3.70E-28 | 1.136748252 | 0.642 | 0.371 | 7.28E-24 | Tb |
| cDC2 | PLEKHG2 | 9.17E-16 | 1.134289826 | 0.415 | 0.242 | 1.81E-11 | Tb |

| | | | | | | | |
|---|---|---|---|---|---|---|---|
| cDC2 | YTHDC2 | 4.90E-25 | 1.132025711 | 0.567 | 0.305 | 9.65E-21 | Tb |
| cDC2 | TSPAN14 | 1.78E-22 | 1.131708709 | 0.366 | 0.163 | 3.51E-18 | Tb |
| cDC2 | SOAT1 | 1.20E-29 | 1.130197571 | 0.574 | 0.29 | 2.36E-25 | Tb |
| cDC2 | SCAF8 | 1.05E-30 | 1.129151179 | 0.447 | 0.184 | 2.06E-26 | Tb |
| cDC2 | TET2 | 1.09E-33 | 1.126713115 | 0.923 | 0.751 | 2.14E-29 | Tb |
| cDC2 | SMG6 | 7.27E-24 | 1.12656378 | 0.417 | 0.192 | 1.43E-19 | Tb |
| cDC2 | KLHL15 | 6.06E-15 | 1.126406166 | 0.332 | 0.167 | 1.19E-10 | Tb |
| cDC2 | ST3GAL2 | 3.07E-26 | 1.125708619 | 0.382 | 0.161 | 6.04E-22 | Tb |
| cDC2 | CERS5 | 2.14E-26 | 1.122699461 | 0.365 | 0.143 | 4.21E-22 | Tb |
| cDC2 | INTS6L | 2.98E-22 | 1.122048007 | 0.326 | 0.139 | 5.86E-18 | Tb |
| cDC2 | BRAF | 1.03E-24 | 1.121461971 | 0.714 | 0.482 | 2.03E-20 | Tb |
| cDC2 | FBXO34 | 5.81E-32 | 1.119741608 | 0.658 | 0.374 | 1.14E-27 | Tb |
| cDC2 | ADD3 | 4.13E-10 | 1.119360934 | 0.283 | 0.161 | 8.12E-06 | Tb |
| cDC2 | RGS12 | 6.62E-23 | 1.118887551 | 0.372 | 0.163 | 1.30E-18 | Tb |
| cDC2 | SLC1A4 | 5.28E-16 | 1.118725729 | 0.333 | 0.167 | 1.04E-11 | Tb |
| cDC2 | CLSTN1 | 6.06E-18 | 1.118513511 | 0.269 | 0.11 | 1.19E-13 | Tb |
| cDC2 | NLK | 1.18E-19 | 1.117941953 | 0.304 | 0.132 | 2.33E-15 | Tb |
| cDC2 | INHBA-AS1 | 5.17E-17 | 1.116586481 | 0.325 | 0.154 | 1.02E-12 | Tb |
| cDC2 | FOXK2 | 1.30E-18 | 1.116566298 | 0.459 | 0.248 | 2.57E-14 | Tb |
| cDC2 | SGPL1 | 5.54E-28 | 1.116494853 | 0.435 | 0.189 | 1.09E-23 | Tb |
| cDC2 | GOT1 | 1.18E-14 | 1.114488731 | 0.265 | 0.12 | 2.33E-10 | Tb |
| cDC2 | MX2 | 3.10E-13 | 1.111227541 | 0.508 | 0.335 | 6.10E-09 | Tb |
| cDC2 | 3-Mar | 2.11E-10 | 1.109977157 | 0.295 | 0.164 | 4.15E-06 | Tb |
| cDC2 | ZCCHC2 | 1.90E-24 | 1.108296819 | 0.461 | 0.224 | 3.73E-20 | Tb |
| cDC2 | NLRP1 | 9.67E-17 | 1.108030156 | 0.447 | 0.249 | 1.90E-12 | Tb |
| cDC2 | FLT1 | 4.55E-26 | 1.107795974 | 0.794 | 0.596 | 8.96E-22 | Tb |
| cDC2 | ZEB1 | 3.93E-12 | 1.106184914 | 0.356 | 0.212 | 7.73E-08 | Tb |
| cDC2 | FIG4 | 5.88E-21 | 1.104395582 | 0.276 | 0.106 | 1.16E-16 | Tb |
| cDC2 | ALPK1 | 1.41E-27 | 1.102932141 | 0.569 | 0.311 | 2.78E-23 | Tb |
| cDC2 | SREBF2 | 6.52E-26 | 1.102869395 | 0.41 | 0.176 | 1.28E-21 | Tb |
| cDC2 | FAM214A | 3.28E-15 | 1.10226591 | 0.253 | 0.113 | 6.46E-11 | Tb |
| cDC2 | PRKD3 | 1.40E-20 | 1.10138932 | 0.529 | 0.3 | 2.75E-16 | Tb |
| cDC2 | EML4 | 1.09E-36 | 1.100883044 | 0.742 | 0.475 | 2.14E-32 | Tb |
| cDC2 | CD109 | 1.47E-12 | 1.100727099 | 0.473 | 0.312 | 2.90E-08 | Tb |
| cDC2 | EVL | 5.24E-23 | 1.098464616 | 0.593 | 0.366 | 1.03E-18 | Tb |
| cDC2 | MBTD1 | 1.61E-20 | 1.098420412 | 0.333 | 0.147 | 3.18E-16 | Tb |
| cDC2 | PLEKHA51 | 9.83E-10 | 1.096955657 | 0.401 | 0.26 | 1.93E-05 | Tb |
| cDC2 | BTBD7 | 1.47E-20 | 1.093814328 | 0.595 | 0.361 | 2.89E-16 | Tb |
| cDC2 | SYNJ1 | 5.31E-20 | 1.093515679 | 0.33 | 0.146 | 1.05E-15 | Tb |
| cDC2 | STX17-AS1 | 1.50E-16 | 1.092759097 | 0.454 | 0.26 | 2.95E-12 | Tb |
| cDC2 | EXT1 | 1.13E-21 | 1.09272268 | 0.529 | 0.297 | 2.23E-17 | Tb |
| cDC2 | PRKCA | 4.18E-27 | 1.089450227 | 0.567 | 0.308 | 8.22E-23 | Tb |
| cDC2 | MAN1A1 | 2.08E-32 | 1.088039419 | 0.761 | 0.478 | 4.10E-28 | Tb |
| cDC2 | TBL1X | 1.45E-16 | 1.087302312 | 0.424 | 0.233 | 2.85E-12 | Tb |
| cDC2 | ZNF141 | 4.15E-12 | 1.086629567 | 0.295 | 0.157 | 8.17E-08 | Tb |
| cDC2 | PICALM | 4.05E-47 | 1.085008611 | 0.934 | 0.757 | 7.97E-43 | Tb |
| cDC2 | SCLT1 | 3.39E-29 | 1.083878813 | 0.738 | 0.484 | 6.68E-25 | Tb |
| cDC2 | B3GNT5 | 3.87E-24 | 1.082955689 | 0.759 | 0.542 | 7.62E-20 | Tb |
| cDC2 | MICU1 | 5.02E-24 | 1.082387033 | 0.466 | 0.233 | 9.89E-20 | Tb |
| cDC2 | MRC1 | 1.67E-20 | 1.081647314 | 0.632 | 0.43 | 3.29E-16 | Tb |
| cDC2 | CSF2RB | 3.52E-28 | 1.080474759 | 0.506 | 0.24 | 6.93E-24 | Tb |
| cDC2 | MT-ND4 | 3.01E-85 | 1.080233644 | 0.991 | 0.924 | 5.92E-81 | Tb |
| cDC2 | ARHGAP10 | 1.12E-19 | 1.080021635 | 0.562 | 0.35 | 2.21E-15 | Tb |
| cDC2 | ATP11C | 2.71E-15 | 1.076584665 | 0.269 | 0.122 | 5.32E-11 | Tb |
| cDC2 | MSL2 | 4.37E-21 | 1.076376632 | 0.335 | 0.143 | 8.59E-17 | Tb |
| cDC2 | CRYBG1 | 1.62E-25 | 1.076271809 | 0.74 | 0.498 | 3.19E-21 | Tb |

| | | | | | | | |
|---|---|---|---|---|---|---|---|
| cDC2 | MREG | 8.18E-17 | 1.075980924 | 0.396 | 0.211 | 1.61E-12 | Tb |
| cDC2 | UBL3 | 5.01E-29 | 1.075025487 | 0.74 | 0.469 | 9.86E-25 | Tb |
| cDC2 | STRADA | 1.00E-16 | 1.074714622 | 0.267 | 0.115 | 1.97E-12 | Tb |
| cDC2 | CAMK1D | 1.04E-41 | 1.074572608 | 0.853 | 0.564 | 2.06E-37 | Tb |
| cDC2 | DUSP16 | 5.55E-24 | 1.074341587 | 0.426 | 0.199 | 1.09E-19 | Tb |
| cDC2 | DDX58 | 2.54E-15 | 1.073618167 | 0.333 | 0.171 | 5.01E-11 | Tb |
| cDC2 | UBASH3B | 3.48E-18 | 1.071911832 | 0.567 | 0.339 | 6.86E-14 | Tb |
| cDC2 | ARHGEF3 | 1.45E-22 | 1.068176705 | 0.375 | 0.169 | 2.86E-18 | Tb |
| cDC2 | ACSS2 | 3.04E-18 | 1.06798978 | 0.305 | 0.134 | 5.99E-14 | Tb |
| cDC2 | SCARF1 | 2.87E-26 | 1.066208347 | 0.45 | 0.209 | 5.65E-22 | Tb |
| cDC2 | SCFD2 | 2.19E-19 | 1.066146333 | 0.335 | 0.152 | 4.30E-15 | Tb |
| cDC2 | FBXL17 | 5.88E-24 | 1.0614818 | 0.478 | 0.238 | 1.16E-19 | Tb |
| cDC2 | PELP1 | 2.93E-17 | 1.058285994 | 0.25 | 0.1 | 5.77E-13 | Tb |
| cDC2 | WASHC5 | 1.25E-14 | 1.05818104 | 0.325 | 0.165 | 2.46E-10 | Tb |
| cDC2 | SEC23B | 5.67E-22 | 1.056724775 | 0.455 | 0.232 | 1.12E-17 | Tb |
| cDC2 | DNMBP | 5.70E-08 | 1.054552246 | 0.271 | 0.165 | 0.001122112 | Tb |
| cDC2 | IBTK | 1.52E-24 | 1.048314096 | 0.457 | 0.218 | 2.98E-20 | Tb |
| cDC2 | GMDS-DT | 1.54E-10 | 1.046251538 | 0.302 | 0.176 | 3.03E-06 | Tb |
| cDC2 | PHLDB1 | 6.06E-11 | 1.043525059 | 0.279 | 0.148 | 1.19E-06 | Tb |
| cDC2 | RIPK1 | 3.69E-27 | 1.042884432 | 0.583 | 0.319 | 7.27E-23 | Tb |
| cDC2 | RAB30-DT | 7.98E-22 | 1.041963512 | 0.325 | 0.134 | 1.57E-17 | Tb |
| cDC2 | CDK17 | 5.32E-18 | 1.038435957 | 0.48 | 0.269 | 1.05E-13 | Tb |
| cDC2 | ZMYND8 | 1.01E-22 | 1.038178534 | 0.389 | 0.174 | 1.99E-18 | Tb |
| cDC2 | WWOX | 4.69E-14 | 1.037944344 | 0.377 | 0.215 | 9.24E-10 | Tb |
| cDC2 | LIMS1 | 1.77E-71 | 1.036363217 | 0.955 | 0.764 | 3.49E-67 | Tb |
| cDC2 | MGAT5 | 1.72E-22 | 1.035627545 | 0.548 | 0.29 | 3.39E-18 | Tb |
| cDC2 | RPF2 | 8.17E-17 | 1.035515229 | 0.346 | 0.165 | 1.61E-12 | Tb |
| cDC2 | ERI1 | 9.04E-24 | 1.034899067 | 0.442 | 0.211 | 1.78E-19 | Tb |
| cDC2 | AXL | 3.80E-26 | 1.034675322 | 0.597 | 0.341 | 7.48E-22 | Tb |
| cDC2 | CCDC97 | 2.52E-21 | 1.032869814 | 0.349 | 0.153 | 4.95E-17 | Tb |
| cDC2 | TBC1D22A | 2.17E-31 | 1.032146575 | 0.756 | 0.494 | 4.28E-27 | Tb |
| cDC2 | HOMER2 | 2.08E-26 | 1.031667347 | 0.368 | 0.147 | 4.09E-22 | Tb |
| cDC2 | SPATS2L | 4.03E-27 | 1.030749148 | 0.757 | 0.496 | 7.94E-23 | Tb |
| cDC2 | SRGAP2C | 1.44E-24 | 1.030248587 | 0.403 | 0.188 | 2.83E-20 | Tb |
| cDC2 | SPRED1 | 3.87E-16 | 1.029239086 | 0.354 | 0.184 | 7.62E-12 | Tb |
| cDC2 | THOC1 | 3.29E-17 | 1.028679991 | 0.349 | 0.172 | 6.48E-13 | Tb |
| cDC2 | PITPNA | 1.06E-34 | 1.028599872 | 0.759 | 0.468 | 2.09E-30 | Tb |
| cDC2 | RCAN1 | 4.11E-07 | 1.026997823 | 0.264 | 0.159 | 0.008095914 | Tb |
| cDC2 | ATG7 | 1.78E-28 | 1.024763224 | 0.593 | 0.323 | 3.50E-24 | Tb |
| cDC2 | RUFY4 | 8.88E-23 | 1.024531668 | 0.344 | 0.143 | 1.75E-18 | Tb |
| cDC2 | CTDP1 | 1.33E-18 | 1.024269085 | 0.33 | 0.151 | 2.62E-14 | Tb |
| cDC2 | NT5C2 | 3.80E-28 | 1.023676908 | 0.557 | 0.282 | 7.48E-24 | Tb |
| cDC2 | ANAPC10 | 5.14E-20 | 1.020987148 | 0.429 | 0.211 | 1.01E-15 | Tb |
| cDC2 | SUSD6 | 7.11E-35 | 1.013652791 | 0.672 | 0.378 | 1.40E-30 | Tb |
| cDC2 | GALNT11 | 2.08E-18 | 1.013121055 | 0.389 | 0.191 | 4.10E-14 | Tb |
| cDC2 | NFATC3 | 4.59E-12 | 1.012430386 | 0.305 | 0.162 | 9.04E-08 | Tb |
| cDC2 | ESYT2 | 1.56E-23 | 1.010466347 | 0.812 | 0.634 | 3.07E-19 | Tb |
| cDC2 | F8A1 | 6.76E-16 | 1.010059473 | 0.257 | 0.111 | 1.33E-11 | Tb |
| cDC2 | FAM118A | 1.83E-18 | 1.009497963 | 0.403 | 0.202 | 3.59E-14 | Tb |
| cDC2 | CBFB | 1.17E-26 | 1.00904164 | 0.478 | 0.225 | 2.30E-22 | Tb |
| cDC2 | N4BP2 | 7.31E-23 | 1.007320137 | 0.442 | 0.21 | 1.44E-18 | Tb |
| cDC2 | TMEM131 | 4.20E-31 | 1.006825816 | 0.771 | 0.499 | 8.27E-27 | Tb |
| cDC2 | EZH2 | 4.85E-31 | 1.003413955 | 0.681 | 0.395 | 9.56E-27 | Tb |
| cDC2 | TAOK1 | 5.32E-27 | 1.001363642 | 0.579 | 0.295 | 1.05E-22 | Tb |
| cDC2 | MTHFR | 2.70E-24 | 1.001291375 | 0.506 | 0.255 | 5.31E-20 | Tb |
| cDC2 | STEAP1B | 1.44E-16 | 1.000929891 | 0.433 | 0.235 | 2.84E-12 | Tb |

| | | | | | | | |
|---|---|---|---|---|---|---|---|
| cDC2 | ASAP1 | 4.67E-35 | 0.998839743 | 0.913 | 0.742 | 9.19E-31 | Tb |
| cDC2 | POGZ | 1.90E-21 | 0.997221781 | 0.452 | 0.227 | 3.75E-17 | Tb |
| cDC2 | RCOR1 | 5.52E-23 | 0.99601554 | 0.815 | 0.611 | 1.09E-18 | Tb |
| cDC2 | ADNP2 | 1.03E-20 | 0.995240575 | 0.386 | 0.18 | 2.03E-16 | Tb |
| cDC2 | LRRK2 | 4.73E-18 | 0.995116298 | 0.379 | 0.191 | 9.31E-14 | Tb |
| cDC2 | CRY1 | 7.52E-13 | 0.995052402 | 0.316 | 0.169 | 1.48E-08 | Tb |
| cDC2 | CDCA4 | 7.32E-14 | 0.994936577 | 0.258 | 0.121 | 1.44E-09 | Tb |
| cDC2 | ERO1A | 2.31E-21 | 0.994776664 | 0.482 | 0.252 | 4.54E-17 | Tb |
| cDC2 | SUCNR1 | 3.61E-22 | 0.994590308 | 0.422 | 0.206 | 7.10E-18 | Tb |
| cDC2 | SYTL3 | 8.49E-16 | 0.993628482 | 0.438 | 0.263 | 1.67E-11 | Tb |
| cDC2 | FGD4 | 1.10E-20 | 0.993560328 | 0.616 | 0.398 | 2.16E-16 | Tb |
| cDC2 | SLC45A4 | 1.03E-17 | 0.9926811 | 0.408 | 0.216 | 2.04E-13 | Tb |
| cDC2 | GSK3B | 2.44E-22 | 0.992437763 | 0.618 | 0.382 | 4.81E-18 | Tb |
| cDC2 | SLC16A3 | 2.05E-30 | 0.991530543 | 0.618 | 0.334 | 4.03E-26 | Tb |
| cDC2 | BCL11A | 9.11E-25 | 0.990067086 | 0.625 | 0.374 | 1.79E-20 | Tb |
| cDC2 | FKBP5 | 1.99E-39 | 0.98695109 | 0.815 | 0.499 | 3.91E-35 | Tb |
| cDC2 | ZNF407 | 2.73E-25 | 0.986885548 | 0.489 | 0.242 | 5.38E-21 | Tb |
| cDC2 | CEP83 | 2.06E-18 | 0.986485655 | 0.508 | 0.291 | 4.06E-14 | Tb |
| cDC2 | INSR | 7.92E-19 | 0.985880462 | 0.305 | 0.137 | 1.56E-14 | Tb |
| cDC2 | N4BP1 | 1.39E-28 | 0.984893425 | 0.723 | 0.45 | 2.74E-24 | Tb |
| cDC2 | ELMSAN1 | 3.52E-26 | 0.984452837 | 0.529 | 0.27 | 6.94E-22 | Tb |
| cDC2 | ZHX2 | 4.58E-18 | 0.984418951 | 0.562 | 0.337 | 9.02E-14 | Tb |
| cDC2 | HAVCR2 | 1.23E-19 | 0.983669807 | 0.726 | 0.477 | 2.41E-15 | Tb |
| cDC2 | RALGAPB | 2.67E-10 | 0.983282765 | 0.26 | 0.135 | 5.26E-06 | Tb |
| cDC2 | GALC | 5.69E-10 | 0.98158706 | 0.269 | 0.15 | 1.12E-05 | Tb |
| cDC2 | EPB41L2 | 5.81E-21 | 0.980023508 | 0.64 | 0.409 | 1.14E-16 | Tb |
| cDC2 | NBEAL1 | 3.87E-17 | 0.979053904 | 0.738 | 0.52 | 7.61E-13 | Tb |
| cDC2 | KDM4C | 6.92E-34 | 0.978906296 | 0.712 | 0.409 | 1.36E-29 | Tb |
| cDC2 | KANSL1 | 8.82E-21 | 0.97813344 | 0.501 | 0.274 | 1.74E-16 | Tb |
| cDC2 | CERS6 | 4.38E-18 | 0.977132163 | 0.682 | 0.448 | 8.63E-14 | Tb |
| cDC2 | RASGRP3 | 2.45E-19 | 0.973843934 | 0.435 | 0.232 | 4.83E-15 | Tb |
| cDC2 | RHBDD1 | 2.05E-12 | 0.973430731 | 0.283 | 0.146 | 4.03E-08 | Tb |
| cDC2 | CDKAL1 | 5.33E-18 | 0.973079271 | 0.436 | 0.243 | 1.05E-13 | Tb |
| cDC2 | RASA1 | 1.91E-25 | 0.972810186 | 0.428 | 0.191 | 3.76E-21 | Tb |
| cDC2 | ELL2 | 4.61E-32 | 0.970391373 | 0.796 | 0.544 | 9.07E-28 | Tb |
| cDC2 | NUBPL | 2.79E-18 | 0.969987663 | 0.419 | 0.222 | 5.49E-14 | Tb |
| cDC2 | CD80 | 3.15E-18 | 0.969076299 | 0.529 | 0.319 | 6.19E-14 | Tb |
| cDC2 | USP25 | 2.60E-17 | 0.968107021 | 0.485 | 0.28 | 5.11E-13 | Tb |
| cDC2 | UBP1 | 2.70E-17 | 0.966486929 | 0.325 | 0.151 | 5.31E-13 | Tb |
| cDC2 | NFAM1 | 6.92E-18 | 0.965042459 | 0.337 | 0.159 | 1.36E-13 | Tb |
| cDC2 | TOPORS | 5.19E-22 | 0.963104536 | 0.419 | 0.199 | 1.02E-17 | Tb |
| cDC2 | CYTOR | 1.38E-32 | 0.961790985 | 0.867 | 0.661 | 2.71E-28 | Tb |
| cDC2 | SIPA1L3 | 1.09E-23 | 0.961655767 | 0.551 | 0.317 | 2.14E-19 | Tb |
| cDC2 | AP1S3 | 6.49E-11 | 0.960985814 | 0.323 | 0.188 | 1.28E-06 | Tb |
| cDC2 | IQCJ-SCHIP1 | 6.03E-07 | 0.958035427 | 0.26 | 0.166 | 0.011865958 | Tb |
| cDC2 | MT-ND1 | 3.84E-61 | 0.95673705 | 0.977 | 0.894 | 7.56E-57 | Tb |
| cDC2 | MT-CO1 | 7.18E-64 | 0.95595807 | 0.997 | 0.985 | 1.41E-59 | Tb |
| cDC2 | NFAT5 | 3.31E-26 | 0.955137074 | 0.869 | 0.725 | 6.52E-22 | Tb |
| cDC2 | SNX8 | 1.35E-33 | 0.955054106 | 0.841 | 0.622 | 2.65E-29 | Tb |
| cDC2 | TNRC18 | 8.37E-16 | 0.954545115 | 0.431 | 0.242 | 1.65E-11 | Tb |
| cDC2 | SLC12A2 | 5.13E-10 | 0.953466513 | 0.253 | 0.135 | 1.01E-05 | Tb |
| cDC2 | DLEU1 | 1.48E-25 | 0.952657183 | 0.476 | 0.232 | 2.91E-21 | Tb |
| cDC2 | THADA | 2.64E-17 | 0.952632728 | 0.387 | 0.197 | 5.19E-13 | Tb |
| cDC2 | CCND1 | 1.21E-20 | 0.952368723 | 0.429 | 0.216 | 2.39E-16 | Tb |
| cDC2 | MYO1C | 4.72E-18 | 0.952114028 | 0.302 | 0.133 | 9.29E-14 | Tb |
| cDC2 | EGLN3 | 1.31E-17 | 0.951626901 | 0.365 | 0.187 | 2.58E-13 | Tb |

| | | | | | | | |
|---|---|---|---|---|---|---|---|
| cDC2 | PRPSAP1 | 1.05E-15 | 0.951324122 | 0.342 | 0.169 | 2.07E-11 | Tb |
| cDC2 | MCCC1 | 9.02E-15 | 0.951209091 | 0.274 | 0.126 | 1.77E-10 | Tb |
| cDC2 | ZNF146 | 4.15E-17 | 0.951057327 | 0.272 | 0.115 | 8.17E-13 | Tb |
| cDC2 | RNF111 | 7.18E-18 | 0.950835193 | 0.445 | 0.233 | 1.41E-13 | Tb |
| cDC2 | RICTOR | 3.10E-18 | 0.950835082 | 0.405 | 0.206 | 6.09E-14 | Tb |
| cDC2 | DCLRE1C | 3.47E-20 | 0.950162361 | 0.293 | 0.117 | 6.82E-16 | Tb |
| cDC2 | DIP2B | 7.01E-22 | 0.948940025 | 0.494 | 0.257 | 1.38E-17 | Tb |
| cDC2 | METTL16 | 3.22E-15 | 0.948057369 | 0.26 | 0.116 | 6.34E-11 | Tb |
| cDC2 | WDR12 | 3.24E-13 | 0.947931706 | 0.255 | 0.117 | 6.37E-09 | Tb |
| cDC2 | MINK1 | 2.25E-19 | 0.947826682 | 0.358 | 0.167 | 4.42E-15 | Tb |
| cDC2 | WDR81 | 3.98E-16 | 0.947415622 | 0.269 | 0.118 | 7.84E-12 | Tb |
| cDC2 | ARFGEF2 | 2.48E-18 | 0.946986511 | 0.426 | 0.227 | 4.88E-14 | Tb |
| cDC2 | ELMO1 | 6.81E-25 | 0.946882292 | 0.824 | 0.665 | 1.34E-20 | Tb |
| cDC2 | TCF20 | 1.50E-15 | 0.944796677 | 0.325 | 0.16 | 2.95E-11 | Tb |
| cDC2 | MGAT2 | 1.26E-13 | 0.943994924 | 0.265 | 0.125 | 2.48E-09 | Tb |
| cDC2 | SPECC1L | 2.42E-18 | 0.94279253 | 0.33 | 0.15 | 4.77E-14 | Tb |
| cDC2 | ADAP2 | 6.52E-18 | 0.942526044 | 0.419 | 0.216 | 1.28E-13 | Tb |
| cDC2 | ATP2A2 | 1.16E-22 | 0.942493155 | 0.653 | 0.385 | 2.28E-18 | Tb |
| cDC2 | VPS9D1 | 1.92E-19 | 0.94244098 | 0.335 | 0.15 | 3.77E-15 | Tb |
| cDC2 | GNG7 | 5.93E-22 | 0.940418892 | 0.661 | 0.426 | 1.17E-17 | Tb |
| cDC2 | ACO2 | 6.35E-21 | 0.940253289 | 0.496 | 0.264 | 1.25E-16 | Tb |
| cDC2 | FCHSD2 | 1.82E-22 | 0.939605903 | 0.782 | 0.566 | 3.58E-18 | Tb |
| cDC2 | GGA3 | 5.31E-17 | 0.938959956 | 0.272 | 0.116 | 1.05E-12 | Tb |
| cDC2 | ITPRIPL2 | 1.76E-20 | 0.937492428 | 0.353 | 0.157 | 3.47E-16 | Tb |
| cDC2 | SLC38A1 | 2.15E-21 | 0.936636301 | 0.593 | 0.363 | 4.23E-17 | Tb |
| cDC2 | MAP3K13 | 5.46E-27 | 0.936125353 | 0.689 | 0.436 | 1.07E-22 | Tb |
| cDC2 | ADIPOR2 | 4.30E-18 | 0.93531372 | 0.41 | 0.214 | 8.46E-14 | Tb |
| cDC2 | MIB1 | 6.23E-17 | 0.93446368 | 0.288 | 0.126 | 1.23E-12 | Tb |
| cDC2 | TMEM39A | 3.26E-19 | 0.934444197 | 0.508 | 0.285 | 6.42E-15 | Tb |
| cDC2 | TP53 | 2.06E-12 | 0.933733469 | 0.264 | 0.131 | 4.05E-08 | Tb |
| cDC2 | INTS6 | 2.31E-20 | 0.933427648 | 0.724 | 0.508 | 4.55E-16 | Tb |
| cDC2 | NFATC1 | 1.04E-16 | 0.931971919 | 0.515 | 0.311 | 2.06E-12 | Tb |
| cDC2 | CTSC | 8.85E-18 | 0.931331095 | 0.798 | 0.614 | 1.74E-13 | Tb |
| cDC2 | GRK3 | 6.11E-23 | 0.931073699 | 0.689 | 0.446 | 1.20E-18 | Tb |
| cDC2 | AP3D1 | 7.62E-25 | 0.930457724 | 0.551 | 0.294 | 1.50E-20 | Tb |
| cDC2 | R3HDM2 | 6.03E-24 | 0.930347498 | 0.614 | 0.365 | 1.19E-19 | Tb |
| cDC2 | CHST11 | 8.01E-31 | 0.930259414 | 0.859 | 0.642 | 1.58E-26 | Tb |
| cDC2 | ARHGAP25 | 3.24E-18 | 0.929856322 | 0.541 | 0.323 | 6.38E-14 | Tb |
| cDC2 | CLCN6 | 8.65E-14 | 0.929266121 | 0.255 | 0.118 | 1.70E-09 | Tb |
| cDC2 | ULK4 | 1.10E-16 | 0.928710185 | 0.318 | 0.15 | 2.17E-12 | Tb |
| cDC2 | CCDC138 | 3.85E-21 | 0.928585051 | 0.363 | 0.167 | 7.58E-17 | Tb |
| cDC2 | RBMS1 | 3.46E-22 | 0.927726476 | 0.606 | 0.35 | 6.81E-18 | Tb |
| cDC2 | ABL2 | 9.97E-32 | 0.927530731 | 0.714 | 0.438 | 1.96E-27 | Tb |
| cDC2 | SEL1L3 | 5.08E-15 | 0.927239942 | 0.314 | 0.157 | 1.00E-10 | Tb |
| cDC2 | ZBTB44 | 4.74E-14 | 0.927049149 | 0.269 | 0.126 | 9.33E-10 | Tb |
| cDC2 | FAM172A | 2.42E-21 | 0.926946125 | 0.524 | 0.289 | 4.76E-17 | Tb |
| cDC2 | KIAA2026 | 1.06E-16 | 0.926491778 | 0.4 | 0.211 | 2.09E-12 | Tb |
| cDC2 | SIPA1L1 | 1.11E-33 | 0.926296095 | 0.953 | 0.789 | 2.18E-29 | Tb |
| cDC2 | CBFA2T2 | 9.79E-18 | 0.926027562 | 0.319 | 0.147 | 1.93E-13 | Tb |
| cDC2 | FAM133B | 1.51E-26 | 0.925238675 | 0.604 | 0.335 | 2.96E-22 | Tb |
| cDC2 | PDS5B | 1.05E-20 | 0.925144264 | 0.469 | 0.249 | 2.07E-16 | Tb |
| cDC2 | ATP1B3 | 5.21E-25 | 0.925092347 | 0.948 | 0.862 | 1.03E-20 | Tb |
| cDC2 | MSI2 | 2.77E-09 | 0.925024795 | 0.311 | 0.185 | 5.46E-05 | Tb |
| cDC2 | ZNF33A | 9.22E-12 | 0.923395881 | 0.328 | 0.181 | 1.81E-07 | Tb |
| cDC2 | CTNND1 | 3.81E-21 | 0.923008339 | 0.632 | 0.385 | 7.51E-17 | Tb |
| cDC2 | PMEPA1 | 3.52E-21 | 0.921409275 | 0.379 | 0.176 | 6.93E-17 | Tb |

| | | | | | | | |
|---|---|---|---|---|---|---|---|
| cDC2 | TEC | 6.56E-12 | 0.920592531 | 0.309 | 0.168 | 1.29E-07 | Tb |
| cDC2 | TENT4B | 2.83E-16 | 0.920475124 | 0.536 | 0.33 | 5.57E-12 | Tb |
| cDC2 | CEBPA | 1.06E-25 | 0.918522615 | 0.513 | 0.262 | 2.08E-21 | Tb |
| cDC2 | CASP8 | 3.67E-23 | 0.91806928 | 0.56 | 0.304 | 7.23E-19 | Tb |
| cDC2 | PHF19 | 4.19E-10 | 0.916877894 | 0.431 | 0.282 | 8.25E-06 | Tb |
| cDC2 | CHD8 | 5.03E-16 | 0.913745125 | 0.33 | 0.161 | 9.90E-12 | Tb |
| cDC2 | DMTF1 | 1.62E-09 | 0.913051163 | 0.253 | 0.137 | 3.19E-05 | Tb |
| cDC2 | VPS35L | 3.10E-14 | 0.912775925 | 0.265 | 0.121 | 6.09E-10 | Tb |
| cDC2 | BMPR2 | 2.07E-17 | 0.912286194 | 0.424 | 0.227 | 4.08E-13 | Tb |
| cDC2 | PTPN9 | 3.25E-11 | 0.910851168 | 0.335 | 0.188 | 6.39E-07 | Tb |
| cDC2 | RNF103 | 1.57E-16 | 0.910750193 | 0.337 | 0.165 | 3.09E-12 | Tb |
| cDC2 | CFLAR-AS1 | 3.06E-18 | 0.909499931 | 0.408 | 0.207 | 6.03E-14 | Tb |
| cDC2 | NOL10 | 1.59E-15 | 0.907816818 | 0.298 | 0.14 | 3.13E-11 | Tb |
| cDC2 | MALT1 | 1.05E-14 | 0.907730888 | 0.815 | 0.676 | 2.07E-10 | Tb |
| cDC2 | DIS3L2 | 5.16E-13 | 0.906780196 | 0.26 | 0.125 | 1.02E-08 | Tb |
| cDC2 | DDX10 | 1.42E-16 | 0.906649343 | 0.351 | 0.175 | 2.80E-12 | Tb |
| cDC2 | TRPM2 | 5.19E-14 | 0.90522572 | 0.309 | 0.159 | 1.02E-09 | Tb |
| cDC2 | MT-CO2 | 2.36E-70 | 0.903698226 | 0.991 | 0.979 | 4.65E-66 | Tb |
| cDC2 | INPP5D | 1.14E-16 | 0.903540953 | 0.503 | 0.286 | 2.25E-12 | Tb |
| cDC2 | SULF2 | 1.12E-22 | 0.902655488 | 0.745 | 0.527 | 2.20E-18 | Tb |
| cDC2 | ZSWIM6 | 1.48E-30 | 0.902333081 | 0.892 | 0.686 | 2.91E-26 | Tb |
| cDC2 | PAG1 | 2.06E-17 | 0.902227992 | 0.525 | 0.311 | 4.05E-13 | Tb |
| cDC2 | ARMH3 | 1.93E-25 | 0.901184143 | 0.48 | 0.229 | 3.80E-21 | Tb |
| cDC2 | USP32 | 2.84E-27 | 0.899181493 | 0.717 | 0.444 | 5.59E-23 | Tb |
| cDC2 | GATAD2B | 2.91E-23 | 0.898887854 | 0.527 | 0.281 | 5.73E-19 | Tb |
| cDC2 | IQCB1 | 3.77E-15 | 0.898724541 | 0.366 | 0.194 | 7.42E-11 | Tb |
| cDC2 | RUNX1 | 3.82E-28 | 0.898622627 | 0.841 | 0.629 | 7.52E-24 | Tb |
| cDC2 | SRGAP2 | 1.42E-29 | 0.898543743 | 0.642 | 0.366 | 2.79E-25 | Tb |
| cDC2 | QSOX1 | 7.52E-21 | 0.898141834 | 0.429 | 0.21 | 1.48E-16 | Tb |
| cDC2 | MTHFD1L | 4.87E-17 | 0.897500887 | 0.501 | 0.292 | 9.59E-13 | Tb |
| cDC2 | SPN | 2.94E-18 | 0.896428693 | 0.387 | 0.195 | 5.78E-14 | Tb |
| cDC2 | PDLIM7 | 8.05E-16 | 0.895898985 | 0.3 | 0.14 | 1.58E-11 | Tb |
| cDC2 | DENND4B | 1.73E-17 | 0.895445653 | 0.316 | 0.143 | 3.41E-13 | Tb |
| cDC2 | RARS | 3.86E-16 | 0.895340533 | 0.38 | 0.195 | 7.60E-12 | Tb |
| cDC2 | GPBP1L1 | 9.30E-25 | 0.894842339 | 0.574 | 0.299 | 1.83E-20 | Tb |
| cDC2 | PLXDC2 | 5.15E-37 | 0.894805246 | 0.864 | 0.6 | 1.01E-32 | Tb |
| cDC2 | ANKRD17 | 9.00E-28 | 0.894690263 | 0.679 | 0.398 | 1.77E-23 | Tb |
| cDC2 | TNFRSF10D | 4.00E-14 | 0.893657879 | 0.366 | 0.198 | 7.88E-10 | Tb |
| cDC2 | ANKIB1 | 1.58E-17 | 0.893534241 | 0.38 | 0.192 | 3.10E-13 | Tb |
| cDC2 | PDE7A | 7.82E-13 | 0.893436676 | 0.628 | 0.442 | 1.54E-08 | Tb |
| cDC2 | 8-Mar | 1.06E-17 | 0.893344683 | 0.29 | 0.126 | 2.10E-13 | Tb |
| cDC2 | CDR2 | 2.33E-18 | 0.892238406 | 0.531 | 0.308 | 4.59E-14 | Tb |
| cDC2 | SETDB2 | 1.15E-18 | 0.891045182 | 0.543 | 0.315 | 2.25E-14 | Tb |
| cDC2 | CIITA | 8.07E-24 | 0.890761414 | 0.878 | 0.714 | 1.59E-19 | Tb |
| cDC2 | CDK13 | 2.06E-24 | 0.890381985 | 0.585 | 0.33 | 4.05E-20 | Tb |
| cDC2 | SOS2 | 3.94E-15 | 0.890085723 | 0.38 | 0.203 | 7.76E-11 | Tb |
| cDC2 | FAF1 | 2.76E-22 | 0.889320554 | 0.569 | 0.332 | 5.43E-18 | Tb |
| cDC2 | FRS2 | 4.58E-20 | 0.887885844 | 0.351 | 0.156 | 9.02E-16 | Tb |
| cDC2 | EPC2 | 1.22E-10 | 0.887692089 | 0.33 | 0.188 | 2.41E-06 | Tb |
| cDC2 | USP37 | 5.35E-16 | 0.887617719 | 0.475 | 0.281 | 1.05E-11 | Tb |
| cDC2 | KSR1 | 5.23E-15 | 0.88691673 | 0.312 | 0.153 | 1.03E-10 | Tb |
| cDC2 | LAMP3 | 2.94E-23 | 0.886217944 | 0.6 | 0.361 | 5.78E-19 | Tb |
| cDC2 | LNPEP | 2.28E-15 | 0.885185867 | 0.452 | 0.251 | 4.48E-11 | Tb |
| cDC2 | ADAMTSL4-AS | 8.97E-22 | 0.885025705 | 0.487 | 0.257 | 1.76E-17 | Tb |
| cDC2 | NIBAN1 | 5.25E-24 | 0.883681446 | 0.763 | 0.553 | 1.03E-19 | Tb |
| cDC2 | CDK19 | 1.60E-17 | 0.883162879 | 0.373 | 0.184 | 3.14E-13 | Tb |

| | | | | | | | |
|---|---|---|---|---|---|---|---|
| cDC2 | ZCCHC7 | 1.75E-22 | 0.882708308 | 0.551 | 0.314 | 3.45E-18 | Tb |
| cDC2 | MTR | 3.18E-11 | 0.88262605 | 0.305 | 0.168 | 6.27E-07 | Tb |
| cDC2 | KCNK13 | 1.53E-13 | 0.881951734 | 0.302 | 0.154 | 3.01E-09 | Tb |
| cDC2 | PPP1R14A | 3.24E-15 | 0.881786442 | 0.34 | 0.178 | 6.38E-11 | Tb |
| cDC2 | HMGCR | 7.59E-11 | 0.880192612 | 0.258 | 0.132 | 1.49E-06 | Tb |
| cDC2 | SLC35E1 | 4.38E-19 | 0.879000601 | 0.34 | 0.151 | 8.62E-15 | Tb |
| cDC2 | MOSMO | 7.70E-14 | 0.878770199 | 0.267 | 0.124 | 1.52E-09 | Tb |
| cDC2 | APPBP2 | 3.49E-22 | 0.877781624 | 0.386 | 0.17 | 6.86E-18 | Tb |
| cDC2 | HMGCS1 | 4.45E-14 | 0.877665761 | 0.462 | 0.269 | 8.75E-10 | Tb |
| cDC2 | STIM2 | 1.55E-23 | 0.876959245 | 0.506 | 0.268 | 3.04E-19 | Tb |
| cDC2 | NEMP1 | 1.51E-16 | 0.876885603 | 0.323 | 0.15 | 2.98E-12 | Tb |
| cDC2 | CLASP2 | 2.49E-22 | 0.876290945 | 0.595 | 0.356 | 4.91E-18 | Tb |
| cDC2 | SECISBP2 | 8.23E-18 | 0.875983083 | 0.489 | 0.264 | 1.62E-13 | Tb |
| cDC2 | PCSK7 | 1.62E-18 | 0.875250555 | 0.305 | 0.132 | 3.19E-14 | Tb |
| cDC2 | CCDC18 | 1.21E-16 | 0.875178819 | 0.339 | 0.167 | 2.38E-12 | Tb |
| cDC2 | BCL2L14 | 2.86E-08 | 0.875159451 | 0.25 | 0.144 | 0.000563336 | Tb |
| cDC2 | ITFG1 | 4.93E-21 | 0.874569459 | 0.494 | 0.27 | 9.71E-17 | Tb |
| cDC2 | IGF2R | 6.07E-28 | 0.873544478 | 0.654 | 0.369 | 1.19E-23 | Tb |
| cDC2 | GCLC | 1.07E-13 | 0.873297264 | 0.298 | 0.147 | 2.11E-09 | Tb |
| cDC2 | SMYD3 | 6.40E-11 | 0.873176934 | 0.326 | 0.186 | 1.26E-06 | Tb |
| cDC2 | PPME1 | 2.68E-15 | 0.872538126 | 0.262 | 0.114 | 5.28E-11 | Tb |
| cDC2 | ARID2 | 9.31E-19 | 0.870213964 | 0.565 | 0.344 | 1.83E-14 | Tb |
| cDC2 | VAV1 | 1.94E-20 | 0.86973658 | 0.442 | 0.222 | 3.82E-16 | Tb |
| cDC2 | 1-Mar | 1.31E-18 | 0.869719084 | 0.64 | 0.419 | 2.58E-14 | Tb |
| cDC2 | EXOSC6 | 1.24E-09 | 0.869617083 | 0.258 | 0.141 | 2.45E-05 | Tb |
| cDC2 | ATR | 1.25E-16 | 0.869515403 | 0.38 | 0.194 | 2.47E-12 | Tb |
| cDC2 | SLC25A44 | 3.52E-14 | 0.869124801 | 0.267 | 0.124 | 6.92E-10 | Tb |
| cDC2 | MIR4435-2HG | 3.95E-25 | 0.869105342 | 0.672 | 0.411 | 7.78E-21 | Tb |
| cDC2 | SESN1 | 6.13E-14 | 0.869037722 | 0.37 | 0.2 | 1.21E-09 | Tb |
| cDC2 | SKI | 2.14E-16 | 0.868280156 | 0.422 | 0.227 | 4.21E-12 | Tb |
| cDC2 | NDUFS1 | 6.61E-17 | 0.868151616 | 0.305 | 0.139 | 1.30E-12 | Tb |
| cDC2 | CNOT6L | 1.23E-21 | 0.86806158 | 0.62 | 0.372 | 2.42E-17 | Tb |
| cDC2 | INTS3 | 1.28E-21 | 0.867743099 | 0.26 | 0.094 | 2.51E-17 | Tb |
| cDC2 | LATS2 | 1.89E-12 | 0.867576204 | 0.445 | 0.274 | 3.73E-08 | Tb |
| cDC2 | ZNF410 | 1.11E-17 | 0.86746072 | 0.471 | 0.257 | 2.19E-13 | Tb |
| cDC2 | ERCC8 | 6.56E-12 | 0.867278694 | 0.323 | 0.177 | 1.29E-07 | Tb |
| cDC2 | YME1L1 | 4.51E-22 | 0.867235337 | 0.839 | 0.635 | 8.87E-18 | Tb |
| cDC2 | DNAJC5 | 1.77E-17 | 0.866470812 | 0.478 | 0.262 | 3.49E-13 | Tb |
| cDC2 | SREBF1 | 2.96E-20 | 0.866443178 | 0.373 | 0.169 | 5.82E-16 | Tb |
| cDC2 | CYREN | 2.03E-12 | 0.866059232 | 0.314 | 0.167 | 4.00E-08 | Tb |
| cDC2 | TOP3A | 4.43E-14 | 0.864913547 | 0.269 | 0.126 | 8.71E-10 | Tb |
| cDC2 | XPO1 | 3.48E-25 | 0.864235396 | 0.719 | 0.447 | 6.86E-21 | Tb |
| cDC2 | SNX14 | 1.74E-18 | 0.863206609 | 0.4 | 0.2 | 3.43E-14 | Tb |
| cDC2 | ENTPD1 | 7.10E-18 | 0.863121313 | 0.714 | 0.506 | 1.40E-13 | Tb |
| cDC2 | PTPN7 | 1.47E-15 | 0.862708655 | 0.332 | 0.164 | 2.89E-11 | Tb |
| cDC2 | CCNYL1 | 3.09E-19 | 0.862247757 | 0.468 | 0.24 | 6.08E-15 | Tb |
| cDC2 | HPS5 | 1.31E-31 | 0.861378322 | 0.745 | 0.448 | 2.58E-27 | Tb |
| cDC2 | LPP | 3.14E-33 | 0.861352006 | 0.825 | 0.538 | 6.17E-29 | Tb |
| cDC2 | MR1 | 1.52E-13 | 0.860317964 | 0.323 | 0.167 | 2.99E-09 | Tb |
| cDC2 | KDM7A-DT | 4.68E-18 | 0.859891845 | 0.363 | 0.17 | 9.22E-14 | Tb |
| cDC2 | VAPB | 2.68E-15 | 0.859325116 | 0.283 | 0.126 | 5.27E-11 | Tb |
| cDC2 | SLC49A4 | 7.40E-23 | 0.859087371 | 0.48 | 0.245 | 1.46E-18 | Tb |
| cDC2 | ZKSCAN1 | 1.50E-15 | 0.858959804 | 0.326 | 0.159 | 2.95E-11 | Tb |
| cDC2 | STAT3 | 1.92E-28 | 0.857526773 | 0.846 | 0.618 | 3.77E-24 | Tb |
| cDC2 | AQR | 5.11E-15 | 0.857098377 | 0.349 | 0.18 | 1.01E-10 | Tb |
| cDC2 | FARP2 | 2.11E-16 | 0.856228246 | 0.489 | 0.279 | 4.15E-12 | Tb |

| | | | | | | | |
|---|---|---|---|---|---|---|---|
| cDC2 | PACS1 | 1.14E-19 | 0.856005327 | 0.553 | 0.322 | 2.24E-15 | Tb |
| cDC2 | PBRM1 | 8.67E-23 | 0.854888546 | 0.654 | 0.387 | 1.71E-18 | Tb |
| cDC2 | USP6NL | 4.99E-17 | 0.854886871 | 0.445 | 0.245 | 9.82E-13 | Tb |
| cDC2 | MROH1 | 2.17E-17 | 0.854463019 | 0.255 | 0.103 | 4.27E-13 | Tb |
| cDC2 | LRCH3 | 6.82E-25 | 0.854012173 | 0.405 | 0.176 | 1.34E-20 | Tb |
| cDC2 | ZXDC | 3.52E-15 | 0.853814419 | 0.302 | 0.143 | 6.93E-11 | Tb |
| cDC2 | MT-ND2 | 3.86E-52 | 0.853507066 | 0.981 | 0.868 | 7.60E-48 | Tb |
| cDC2 | THUMPD3 | 1.69E-11 | 0.853305799 | 0.262 | 0.132 | 3.33E-07 | Tb |
| cDC2 | TTC9C | 4.07E-18 | 0.851606606 | 0.323 | 0.144 | 8.02E-14 | Tb |
| cDC2 | KDM4B | 5.77E-19 | 0.851010116 | 0.429 | 0.219 | 1.14E-14 | Tb |
| cDC2 | SKIL | 1.59E-27 | 0.850155502 | 0.813 | 0.593 | 3.14E-23 | Tb |
| cDC2 | MFSD14B | 1.70E-16 | 0.850013808 | 0.433 | 0.234 | 3.35E-12 | Tb |
| cDC2 | TSFM | 5.75E-15 | 0.849207174 | 0.269 | 0.121 | 1.13E-10 | Tb |
| cDC2 | HDAC9 | 2.78E-16 | 0.847889899 | 0.688 | 0.541 | 5.48E-12 | Tb |
| cDC2 | TRPM7 | 7.09E-17 | 0.847624581 | 0.492 | 0.271 | 1.40E-12 | Tb |
| cDC2 | YAF2 | 1.78E-17 | 0.846861158 | 0.447 | 0.237 | 3.51E-13 | Tb |
| cDC2 | SLC38A10 | 7.45E-17 | 0.846815561 | 0.272 | 0.115 | 1.47E-12 | Tb |
| cDC2 | KLF7 | 1.06E-16 | 0.846407994 | 0.365 | 0.182 | 2.09E-12 | Tb |
| cDC2 | MLLT6 | 5.90E-13 | 0.846163535 | 0.325 | 0.17 | 1.16E-08 | Tb |
| cDC2 | GMDS | 8.09E-15 | 0.845417991 | 0.574 | 0.366 | 1.59E-10 | Tb |
| cDC2 | ACSL3 | 7.12E-19 | 0.845251327 | 0.773 | 0.531 | 1.40E-14 | Tb |
| cDC2 | ATP2C1 | 6.45E-20 | 0.844913919 | 0.796 | 0.558 | 1.27E-15 | Tb |
| cDC2 | SGMS1 | 7.91E-14 | 0.844660892 | 0.335 | 0.175 | 1.56E-09 | Tb |
| cDC2 | FPR3 | 5.26E-22 | 0.844246953 | 0.766 | 0.538 | 1.04E-17 | Tb |
| cDC2 | MFN1 | 6.29E-20 | 0.844106553 | 0.351 | 0.156 | 1.24E-15 | Tb |
| cDC2 | CHKA | 1.04E-16 | 0.843452432 | 0.489 | 0.289 | 2.04E-12 | Tb |
| cDC2 | GTF2A1 | 4.88E-13 | 0.843239435 | 0.443 | 0.258 | 9.61E-09 | Tb |
| cDC2 | POLR2J3 | 1.30E-17 | 0.842958585 | 0.335 | 0.156 | 2.56E-13 | Tb |
| cDC2 | CENPW | 4.21E-14 | 0.842849857 | 0.351 | 0.181 | 8.29E-10 | Tb |
| cDC2 | LDLRAD4 | 1.41E-25 | 0.842748626 | 0.834 | 0.664 | 2.78E-21 | Tb |
| cDC2 | GNAQ | 4.84E-32 | 0.842643513 | 0.874 | 0.642 | 9.53E-28 | Tb |
| cDC2 | PCYT1A | 1.04E-24 | 0.842409917 | 0.539 | 0.288 | 2.04E-20 | Tb |
| cDC2 | CSNK1G1 | 3.46E-14 | 0.841641272 | 0.316 | 0.161 | 6.82E-10 | Tb |
| cDC2 | LY75 | 7.41E-16 | 0.84143687 | 0.428 | 0.23 | 1.46E-11 | Tb |
| cDC2 | POU2F1 | 2.30E-09 | 0.840695556 | 0.302 | 0.174 | 4.52E-05 | Tb |
| cDC2 | SLC16A6 | 3.59E-12 | 0.840692008 | 0.269 | 0.136 | 7.06E-08 | Tb |
| cDC2 | PPP2R5C | 1.56E-23 | 0.840371617 | 0.654 | 0.379 | 3.07E-19 | Tb |
| cDC2 | FGD2 | 1.71E-18 | 0.840227515 | 0.527 | 0.297 | 3.36E-14 | Tb |
| cDC2 | RHOQ | 2.08E-23 | 0.84020481 | 0.754 | 0.507 | 4.09E-19 | Tb |
| cDC2 | PLAC8 | 2.37E-08 | 0.840127471 | 0.274 | 0.167 | 0.000467424 | Tb |
| cDC2 | SLC15A4 | 2.95E-12 | 0.839824345 | 0.394 | 0.223 | 5.81E-08 | Tb |
| cDC2 | FTX | 5.66E-25 | 0.839416249 | 0.571 | 0.311 | 1.11E-20 | Tb |
| cDC2 | TYMP | 4.85E-52 | 0.838939229 | 0.97 | 0.843 | 9.55E-48 | Tb |
| cDC2 | FNIP2 | 1.60E-34 | 0.838668751 | 0.913 | 0.716 | 3.15E-30 | Tb |
| cDC2 | TMEM181 | 5.42E-12 | 0.83795172 | 0.25 | 0.123 | 1.07E-07 | Tb |
| cDC2 | SNX29 | 3.16E-18 | 0.837907517 | 0.613 | 0.401 | 6.22E-14 | Tb |
| cDC2 | NRF1 | 6.24E-13 | 0.837040209 | 0.318 | 0.169 | 1.23E-08 | Tb |
| cDC2 | TRMO | 3.81E-14 | 0.836243284 | 0.321 | 0.16 | 7.49E-10 | Tb |
| cDC2 | DCAF10 | 9.86E-15 | 0.83564613 | 0.298 | 0.142 | 1.94E-10 | Tb |
| cDC2 | BABAM2 | 3.27E-18 | 0.835394447 | 0.62 | 0.4 | 6.44E-14 | Tb |
| cDC2 | SGMS2 | 1.21E-16 | 0.834543033 | 0.421 | 0.226 | 2.39E-12 | Tb |
| cDC2 | ALOX5AP | 4.98E-13 | 0.834124159 | 0.522 | 0.342 | 9.80E-09 | Tb |
| cDC2 | TRIM24 | 1.41E-08 | 0.834007426 | 0.298 | 0.176 | 0.000277088 | Tb |
| cDC2 | ATXN1 | 1.39E-17 | 0.833955532 | 0.667 | 0.447 | 2.74E-13 | Tb |
| cDC2 | RBM26 | 7.04E-25 | 0.8322089 | 0.473 | 0.218 | 1.38E-20 | Tb |
| cDC2 | NARF | 5.08E-07 | 0.831158042 | 0.45 | 0.297 | 0.00999166 | Tb |

| | | | | | | | |
|---|---|---|---|---|---|---|---|
| cDC2 | PDSS2 | 5.17E-19 | 0.829268834 | 0.307 | 0.131 | 1.02E-14 | Tb |
| cDC2 | ITPR1 | 1.21E-25 | 0.828605111 | 0.642 | 0.384 | 2.38E-21 | Tb |
| cDC2 | STAU2 | 2.37E-18 | 0.828408102 | 0.368 | 0.176 | 4.66E-14 | Tb |
| cDC2 | RNF38 | 2.14E-18 | 0.828186802 | 0.449 | 0.24 | 4.22E-14 | Tb |
| cDC2 | APTX | 1.37E-17 | 0.828088843 | 0.382 | 0.188 | 2.70E-13 | Tb |
| cDC2 | STAT6 | 9.50E-28 | 0.82784424 | 0.595 | 0.318 | 1.87E-23 | Tb |
| cDC2 | LMTK2 | 1.02E-13 | 0.827253164 | 0.253 | 0.117 | 2.00E-09 | Tb |
| cDC2 | ACOT13 | 1.86E-18 | 0.825169595 | 0.356 | 0.165 | 3.65E-14 | Tb |
| cDC2 | KAT6B | 2.28E-12 | 0.82452431 | 0.316 | 0.169 | 4.48E-08 | Tb |
| cDC2 | WBP1L | 1.81E-15 | 0.824253134 | 0.304 | 0.142 | 3.57E-11 | Tb |
| cDC2 | GLB1 | 3.50E-18 | 0.82280419 | 0.417 | 0.209 | 6.88E-14 | Tb |
| cDC2 | JAK3 | 1.38E-12 | 0.822614533 | 0.271 | 0.135 | 2.72E-08 | Tb |
| cDC2 | FRYL | 2.79E-19 | 0.822283384 | 0.625 | 0.375 | 5.48E-15 | Tb |
| cDC2 | TRAF3 | 7.43E-23 | 0.821999214 | 0.707 | 0.461 | 1.46E-18 | Tb |
| cDC2 | SCAPER | 1.45E-12 | 0.821499722 | 0.356 | 0.197 | 2.85E-08 | Tb |
| cDC2 | RAPGEF2 | 5.60E-18 | 0.821415838 | 0.67 | 0.474 | 1.10E-13 | Tb |
| cDC2 | PRKCE | 1.54E-09 | 0.820771141 | 0.52 | 0.374 | 3.04E-05 | Tb |
| cDC2 | NSD2 | 5.29E-14 | 0.820561076 | 0.269 | 0.126 | 1.04E-09 | Tb |
| cDC2 | INPP5A | 7.62E-12 | 0.820131411 | 0.412 | 0.249 | 1.50E-07 | Tb |
| cDC2 | FAF2 | 2.08E-16 | 0.820024572 | 0.304 | 0.139 | 4.10E-12 | Tb |
| cDC2 | PDIA4 | 2.51E-17 | 0.819899182 | 0.394 | 0.2 | 4.94E-13 | Tb |
| cDC2 | PPM1A | 1.46E-10 | 0.819241166 | 0.302 | 0.164 | 2.88E-06 | Tb |
| cDC2 | CREB5 | 1.30E-17 | 0.818925734 | 0.712 | 0.513 | 2.56E-13 | Tb |
| cDC2 | THUMPD3-AS1 | 7.50E-19 | 0.818585571 | 0.517 | 0.291 | 1.48E-14 | Tb |
| cDC2 | CDK6 | 3.68E-12 | 0.817414322 | 0.558 | 0.385 | 7.24E-08 | Tb |
| cDC2 | NRP1 | 1.87E-13 | 0.817298118 | 0.344 | 0.191 | 3.68E-09 | Tb |
| cDC2 | NETO2 | 6.32E-09 | 0.816427457 | 0.283 | 0.164 | 0.000124436 | Tb |
| cDC2 | BICRAL | 1.85E-15 | 0.815877508 | 0.288 | 0.133 | 3.64E-11 | Tb |
| cDC2 | UBAC2 | 9.15E-23 | 0.815866258 | 0.782 | 0.567 | 1.80E-18 | Tb |
| cDC2 | LRIF1 | 1.86E-28 | 0.815224348 | 0.705 | 0.415 | 3.66E-24 | Tb |
| cDC2 | UBE3C | 5.16E-17 | 0.813576353 | 0.408 | 0.209 | 1.02E-12 | Tb |
| cDC2 | MLLT10 | 5.61E-17 | 0.813400774 | 0.44 | 0.238 | 1.11E-12 | Tb |
| cDC2 | IARS | 2.05E-11 | 0.813120337 | 0.403 | 0.237 | 4.03E-07 | Tb |
| cDC2 | EPB41 | 1.74E-12 | 0.812356489 | 0.398 | 0.232 | 3.42E-08 | Tb |
| cDC2 | NAA15 | 9.57E-17 | 0.812295401 | 0.372 | 0.188 | 1.88E-12 | Tb |
| cDC2 | MAST2 | 3.84E-12 | 0.812211178 | 0.262 | 0.129 | 7.55E-08 | Tb |
| cDC2 | HERC3 | 2.12E-09 | 0.812198062 | 0.269 | 0.151 | 4.17E-05 | Tb |
| cDC2 | MTIF2 | 5.16E-17 | 0.811522306 | 0.307 | 0.138 | 1.01E-12 | Tb |
| cDC2 | ARHGAP18 | 1.84E-28 | 0.810951468 | 0.764 | 0.512 | 3.63E-24 | Tb |
| cDC2 | UTRN | 3.49E-26 | 0.810388011 | 0.784 | 0.521 | 6.86E-22 | Tb |
| cDC2 | GTF2F2 | 4.80E-19 | 0.810320156 | 0.525 | 0.295 | 9.44E-15 | Tb |
| cDC2 | STARD4 | 2.70E-08 | 0.810159407 | 0.302 | 0.183 | 0.000530589 | Tb |
| cDC2 | ZNF292 | 6.62E-22 | 0.810154832 | 0.578 | 0.325 | 1.30E-17 | Tb |
| cDC2 | ATXN7 | 8.36E-17 | 0.810016541 | 0.454 | 0.252 | 1.64E-12 | Tb |
| cDC2 | TUG1 | 5.80E-14 | 0.809432715 | 0.26 | 0.12 | 1.14E-09 | Tb |
| cDC2 | PPP3CC | 1.34E-12 | 0.809402509 | 0.356 | 0.195 | 2.64E-08 | Tb |
| cDC2 | SPRED2 | 2.15E-10 | 0.809139272 | 0.536 | 0.375 | 4.23E-06 | Tb |
| cDC2 | COX19 | 2.02E-15 | 0.809063434 | 0.421 | 0.224 | 3.97E-11 | Tb |
| cDC2 | CCNT1 | 1.90E-15 | 0.808989905 | 0.428 | 0.236 | 3.73E-11 | Tb |
| cDC2 | TBC1D17 | 1.49E-14 | 0.807708736 | 0.262 | 0.119 | 2.93E-10 | Tb |
| cDC2 | MT-ND3 | 4.57E-55 | 0.807638679 | 0.981 | 0.883 | 9.00E-51 | Tb |
| cDC2 | RPRD2 | 9.92E-13 | 0.807267505 | 0.264 | 0.127 | 1.95E-08 | Tb |
| cDC2 | UGP2 | 3.32E-22 | 0.807163549 | 0.634 | 0.377 | 6.53E-18 | Tb |
| cDC2 | PRORP | 6.93E-18 | 0.807086953 | 0.335 | 0.156 | 1.36E-13 | Tb |
| cDC2 | MTMR9 | 7.60E-09 | 0.807037576 | 0.291 | 0.167 | 0.00014962 | Tb |
| cDC2 | SLC7A1 | 8.63E-18 | 0.806205883 | 0.459 | 0.245 | 1.70E-13 | Tb |

| | | | | | | | |
|---|---|---|---|---|---|---|---|
| cDC2 | SPTAN1 | 6.36E-08 | 0.805678827 | 0.431 | 0.281 | 0.001251448 | Tb |
| cDC2 | RAPGEF1 | 1.82E-36 | 0.805553814 | 0.934 | 0.796 | 3.59E-32 | Tb |
| cDC2 | ZNF804A | 3.63E-10 | 0.805497288 | 0.384 | 0.248 | 7.14E-06 | Tb |
| cDC2 | OFD1 | 1.52E-12 | 0.805406265 | 0.431 | 0.265 | 2.98E-08 | Tb |
| cDC2 | OPA3 | 1.74E-17 | 0.805041549 | 0.44 | 0.23 | 3.43E-13 | Tb |
| cDC2 | SAP30 | 2.32E-06 | 0.804588657 | 0.309 | 0.196 | 0.045571333 | Tb |
| cDC2 | FBXO11 | 8.75E-24 | 0.804463603 | 0.817 | 0.596 | 1.72E-19 | Tb |
| cDC2 | FASTKD1 | 5.70E-15 | 0.804216638 | 0.257 | 0.113 | 1.12E-10 | Tb |
| cDC2 | GCC2 | 1.33E-15 | 0.804183834 | 0.551 | 0.339 | 2.61E-11 | Tb |
| cDC2 | RABGAP1L | 1.53E-18 | 0.804152938 | 0.635 | 0.39 | 3.02E-14 | Tb |
| cDC2 | AKAP10 | 7.77E-17 | 0.80388043 | 0.277 | 0.119 | 1.53E-12 | Tb |
| cDC2 | ACLY | 1.75E-14 | 0.803793149 | 0.34 | 0.17 | 3.45E-10 | Tb |
| cDC2 | DOCK8 | 2.73E-23 | 0.803737812 | 0.817 | 0.587 | 5.37E-19 | Tb |
| cDC2 | TENT4A | 3.88E-19 | 0.80334844 | 0.457 | 0.241 | 7.64E-15 | Tb |
| cDC2 | ELK4 | 3.55E-12 | 0.80280337 | 0.258 | 0.126 | 6.99E-08 | Tb |
| cDC2 | STK3 | 3.28E-19 | 0.802681311 | 0.579 | 0.35 | 6.47E-15 | Tb |
| cDC2 | FUBP3 | 1.29E-12 | 0.802641568 | 0.257 | 0.122 | 2.54E-08 | Tb |
| cDC2 | SLC31A1 | 5.32E-15 | 0.802520325 | 0.408 | 0.23 | 1.05E-10 | Tb |
| cDC2 | ZZEF1 | 9.78E-16 | 0.801845701 | 0.384 | 0.197 | 1.92E-11 | Tb |
| cDC2 | OSBPL3 | 6.86E-17 | 0.801607712 | 0.45 | 0.254 | 1.35E-12 | Tb |
| cDC2 | TMEM51 | 1.48E-13 | 0.800205574 | 0.38 | 0.211 | 2.91E-09 | Tb |
| cDC2 | TPT1-AS1 | 2.56E-16 | 0.799814431 | 0.271 | 0.116 | 5.03E-12 | Tb |
| cDC2 | USO1 | 2.26E-19 | 0.799763148 | 0.529 | 0.297 | 4.45E-15 | Tb |
| cDC2 | CAAP1 | 7.37E-17 | 0.7997393 | 0.349 | 0.169 | 1.45E-12 | Tb |
| cDC2 | SLC30A1 | 3.28E-09 | 0.799435507 | 0.274 | 0.156 | 6.46E-05 | Tb |
| cDC2 | AIMP2 | 2.89E-12 | 0.798319842 | 0.386 | 0.213 | 5.68E-08 | Tb |
| cDC2 | LARP1B | 4.36E-14 | 0.797972448 | 0.309 | 0.152 | 8.59E-10 | Tb |
| cDC2 | MBNL1 | 2.49E-30 | 0.797895344 | 0.939 | 0.801 | 4.91E-26 | Tb |
| cDC2 | ABHD3 | 2.44E-12 | 0.797671073 | 0.532 | 0.347 | 4.81E-08 | Tb |
| cDC2 | STAM2 | 1.06E-13 | 0.796227228 | 0.319 | 0.162 | 2.10E-09 | Tb |
| cDC2 | TRAPPC9 | 5.44E-21 | 0.795644112 | 0.499 | 0.265 | 1.07E-16 | Tb |
| cDC2 | PSMC1 | 3.35E-25 | 0.795381438 | 0.635 | 0.35 | 6.59E-21 | Tb |
| cDC2 | MAML3 | 1.13E-15 | 0.794430354 | 0.64 | 0.456 | 2.23E-11 | Tb |
| cDC2 | AFF1 | 8.16E-21 | 0.793868415 | 0.766 | 0.518 | 1.61E-16 | Tb |
| cDC2 | TRERF1 | 3.83E-11 | 0.791998289 | 0.363 | 0.213 | 7.53E-07 | Tb |
| cDC2 | IDI1 | 1.01E-19 | 0.791596657 | 0.71 | 0.478 | 1.98E-15 | Tb |
| cDC2 | ZMYM2 | 1.07E-11 | 0.790774401 | 0.632 | 0.446 | 2.10E-07 | Tb |
| cDC2 | PLA2G4A | 1.99E-11 | 0.79065807 | 0.305 | 0.169 | 3.92E-07 | Tb |
| cDC2 | TGFBR1 | 3.37E-15 | 0.790619352 | 0.515 | 0.308 | 6.64E-11 | Tb |
| cDC2 | KRIT1 | 4.00E-15 | 0.790583593 | 0.328 | 0.161 | 7.88E-11 | Tb |
| cDC2 | WDR70 | 4.14E-18 | 0.790305019 | 0.492 | 0.274 | 8.16E-14 | Tb |
| cDC2 | PPP2R5A | 3.11E-15 | 0.789920308 | 0.326 | 0.158 | 6.12E-11 | Tb |
| cDC2 | RIPOR3 | 2.83E-11 | 0.78934505 | 0.253 | 0.129 | 5.56E-07 | Tb |
| cDC2 | GNB4 | 2.47E-21 | 0.789184831 | 0.609 | 0.363 | 4.86E-17 | Tb |
| cDC2 | CCDC91 | 3.55E-23 | 0.788821884 | 0.623 | 0.352 | 7.00E-19 | Tb |
| cDC2 | MICAL2 | 2.82E-16 | 0.788541594 | 0.407 | 0.22 | 5.54E-12 | Tb |
| cDC2 | PPID | 2.55E-22 | 0.787332873 | 0.562 | 0.296 | 5.02E-18 | Tb |
| cDC2 | RRBP1 | 1.50E-17 | 0.787113738 | 0.705 | 0.468 | 2.96E-13 | Tb |
| cDC2 | TENT2 | 2.13E-28 | 0.78706918 | 0.691 | 0.384 | 4.20E-24 | Tb |
| cDC2 | EVI5 | 7.26E-18 | 0.786781791 | 0.571 | 0.341 | 1.43E-13 | Tb |
| cDC2 | SPPL3 | 2.56E-14 | 0.786458491 | 0.576 | 0.368 | 5.05E-10 | Tb |
| cDC2 | RSRC1 | 7.82E-21 | 0.785923741 | 0.555 | 0.306 | 1.54E-16 | Tb |
| cDC2 | SIK2 | 1.50E-11 | 0.785767503 | 0.433 | 0.265 | 2.96E-07 | Tb |
| cDC2 | MTX2 | 8.38E-13 | 0.785319912 | 0.323 | 0.169 | 1.65E-08 | Tb |
| cDC2 | TMEM131L | 1.27E-13 | 0.785239333 | 0.501 | 0.314 | 2.50E-09 | Tb |
| cDC2 | EFL1 | 2.93E-18 | 0.785123838 | 0.377 | 0.181 | 5.78E-14 | Tb |

| | | | | | | | |
|---|---|---|---|---|---|---|---|
| cDC2 | AKT1S1 | 4.16E-15 | 0.784875686 | 0.309 | 0.146 | 8.20E-11 | Tb |
| cDC2 | MFSD4B | 1.84E-14 | 0.784344064 | 0.475 | 0.273 | 3.63E-10 | Tb |
| cDC2 | FOXN3 | 7.12E-28 | 0.783268722 | 0.895 | 0.76 | 1.40E-23 | Tb |
| cDC2 | MT-ND5 | 5.46E-42 | 0.781606614 | 0.962 | 0.808 | 1.07E-37 | Tb |
| cDC2 | YARS | 1.49E-16 | 0.781520264 | 0.396 | 0.2 | 2.93E-12 | Tb |
| cDC2 | EXOC2 | 1.09E-12 | 0.780490942 | 0.267 | 0.129 | 2.15E-08 | Tb |
| cDC2 | AHCYL2 | 1.19E-14 | 0.779936078 | 0.33 | 0.169 | 2.35E-10 | Tb |
| cDC2 | MRPL35 | 1.07E-14 | 0.779884097 | 0.297 | 0.14 | 2.10E-10 | Tb |
| cDC2 | SKAP2 | 6.92E-26 | 0.77933192 | 0.738 | 0.469 | 1.36E-21 | Tb |
| cDC2 | NR4A3 | 7.36E-19 | 0.7792902 | 0.881 | 0.789 | 1.45E-14 | Tb |
| cDC2 | KLHL6 | 8.46E-17 | 0.779017992 | 0.726 | 0.515 | 1.66E-12 | Tb |
| cDC2 | FAM168A | 1.08E-18 | 0.778470744 | 0.372 | 0.176 | 2.12E-14 | Tb |
| cDC2 | TLE4 | 8.57E-12 | 0.778349747 | 0.569 | 0.384 | 1.69E-07 | Tb |
| cDC2 | TFEB | 1.73E-14 | 0.776882243 | 0.33 | 0.165 | 3.40E-10 | Tb |
| cDC2 | RYBP | 1.61E-19 | 0.776623516 | 0.684 | 0.449 | 3.17E-15 | Tb |
| cDC2 | ZNF124 | 1.23E-21 | 0.776264944 | 0.401 | 0.186 | 2.42E-17 | Tb |
| cDC2 | AKNA | 3.06E-10 | 0.775979447 | 0.389 | 0.24 | 6.03E-06 | Tb |
| cDC2 | PCNX1 | 1.03E-13 | 0.775487203 | 0.623 | 0.415 | 2.03E-09 | Tb |
| cDC2 | RSPRY1 | 5.17E-10 | 0.775436087 | 0.276 | 0.15 | 1.02E-05 | Tb |
| cDC2 | PPM1L | 3.98E-12 | 0.775410907 | 0.368 | 0.206 | 7.84E-08 | Tb |
| cDC2 | SAMD9L | 8.44E-27 | 0.775362817 | 0.62 | 0.335 | 1.66E-22 | Tb |
| cDC2 | ABTB2 | 9.37E-11 | 0.774672428 | 0.455 | 0.288 | 1.84E-06 | Tb |
| cDC2 | STIM1 | 2.35E-16 | 0.774211482 | 0.339 | 0.163 | 4.62E-12 | Tb |
| cDC2 | ANKRD11 | 2.15E-29 | 0.774156878 | 0.852 | 0.626 | 4.23E-25 | Tb |
| cDC2 | ARSG | 1.94E-11 | 0.773329912 | 0.251 | 0.124 | 3.82E-07 | Tb |
| cDC2 | DOCK5 | 9.75E-16 | 0.773054067 | 0.59 | 0.4 | 1.92E-11 | Tb |
| cDC2 | RERE | 2.90E-17 | 0.772982455 | 0.688 | 0.48 | 5.71E-13 | Tb |
| cDC2 | WDR44 | 3.48E-09 | 0.772780895 | 0.281 | 0.16 | 6.86E-05 | Tb |
| cDC2 | GTF2H1 | 1.30E-16 | 0.772489716 | 0.51 | 0.297 | 2.55E-12 | Tb |
| cDC2 | RFX7 | 8.39E-09 | 0.772061134 | 0.251 | 0.142 | 0.000165231 | Tb |
| cDC2 | SLC8A1 | 8.36E-33 | 0.771770203 | 0.941 | 0.773 | 1.64E-28 | Tb |
| cDC2 | ANKRD44 | 1.43E-26 | 0.771659438 | 0.789 | 0.549 | 2.81E-22 | Tb |
| cDC2 | CAMK1 | 3.54E-14 | 0.770597767 | 0.379 | 0.207 | 6.96E-10 | Tb |
| cDC2 | SMAD7 | 2.53E-25 | 0.770170008 | 0.581 | 0.3 | 4.98E-21 | Tb |
| cDC2 | MKLN1 | 4.15E-20 | 0.768688389 | 0.686 | 0.433 | 8.17E-16 | Tb |
| cDC2 | CDC42SE2 | 1.63E-22 | 0.767906024 | 0.86 | 0.643 | 3.22E-18 | Tb |
| cDC2 | ZC3HAV1 | 2.23E-19 | 0.767825056 | 0.817 | 0.599 | 4.38E-15 | Tb |
| cDC2 | GALNT1 | 2.31E-24 | 0.7675625 | 0.579 | 0.309 | 4.55E-20 | Tb |
| cDC2 | MED27 | 1.03E-08 | 0.7666501 | 0.265 | 0.149 | 0.000202099 | Tb |
| cDC2 | MFSD14C | 2.96E-18 | 0.766228599 | 0.429 | 0.219 | 5.83E-14 | Tb |
| cDC2 | ACBD5 | 1.65E-11 | 0.764503097 | 0.302 | 0.161 | 3.25E-07 | Tb |
| cDC2 | SELENOI | 3.35E-12 | 0.764237798 | 0.297 | 0.151 | 6.60E-08 | Tb |
| cDC2 | MITD1 | 8.50E-14 | 0.763836134 | 0.36 | 0.189 | 1.67E-09 | Tb |
| cDC2 | PLGRKT | 1.61E-19 | 0.763492183 | 0.581 | 0.351 | 3.17E-15 | Tb |
| cDC2 | PIAS2 | 4.97E-14 | 0.763355708 | 0.305 | 0.15 | 9.79E-10 | Tb |
| cDC2 | SLC9A7 | 2.11E-12 | 0.763230599 | 0.539 | 0.356 | 4.16E-08 | Tb |
| cDC2 | PARP11 | 1.14E-12 | 0.763147709 | 0.319 | 0.168 | 2.24E-08 | Tb |
| cDC2 | FBXW7 | 4.49E-20 | 0.763049082 | 0.733 | 0.517 | 8.83E-16 | Tb |
| cDC2 | TCF12 | 5.59E-24 | 0.762483653 | 0.682 | 0.423 | 1.10E-19 | Tb |
| cDC2 | MYOF | 9.18E-18 | 0.761873589 | 0.67 | 0.454 | 1.81E-13 | Tb |
| cDC2 | NCSTN | 3.30E-11 | 0.761561973 | 0.264 | 0.132 | 6.50E-07 | Tb |
| cDC2 | WDR55 | 8.41E-13 | 0.760705054 | 0.264 | 0.124 | 1.66E-08 | Tb |
| cDC2 | COP1 | 4.13E-26 | 0.760504799 | 0.716 | 0.438 | 8.13E-22 | Tb |
| cDC2 | AHCYL1 | 5.24E-18 | 0.760492751 | 0.503 | 0.283 | 1.03E-13 | Tb |
| cDC2 | TRNT1 | 2.88E-14 | 0.760052304 | 0.25 | 0.111 | 5.67E-10 | Tb |
| cDC2 | TEX14 | 2.38E-14 | 0.759873216 | 0.618 | 0.43 | 4.68E-10 | Tb |

| | | | | | | | |
|---|---|---|---|---|---|---|---|
| cDC2 | AP3S2 | 3.22E-14 | 0.759383699 | 0.339 | 0.172 | 6.35E-10 | Tb |
| cDC2 | PHACTR4 | 2.75E-15 | 0.759052044 | 0.51 | 0.306 | 5.42E-11 | Tb |
| cDC2 | CDYL | 8.20E-12 | 0.758795024 | 0.606 | 0.421 | 1.61E-07 | Tb |
| cDC2 | GOLPH3 | 6.77E-18 | 0.75812642 | 0.454 | 0.239 | 1.33E-13 | Tb |
| cDC2 | EXOC6 | 6.74E-15 | 0.758072508 | 0.366 | 0.198 | 1.33E-10 | Tb |
| cDC2 | SHLD2 | 3.40E-14 | 0.756843605 | 0.408 | 0.224 | 6.70E-10 | Tb |
| cDC2 | KCNMB1 | 4.70E-10 | 0.756568341 | 0.271 | 0.146 | 9.25E-06 | Tb |
| cDC2 | ITPR2 | 1.02E-27 | 0.756472808 | 0.717 | 0.436 | 2.00E-23 | Tb |
| cDC2 | PARP9 | 1.85E-21 | 0.755671291 | 0.527 | 0.281 | 3.65E-17 | Tb |
| cDC2 | USP9X | 3.84E-15 | 0.755564772 | 0.625 | 0.406 | 7.56E-11 | Tb |
| cDC2 | PKNOX1 | 1.12E-11 | 0.75552137 | 0.314 | 0.17 | 2.20E-07 | Tb |
| cDC2 | CLASP1 | 2.84E-16 | 0.755281336 | 0.382 | 0.197 | 5.59E-12 | Tb |
| cDC2 | GANAB | 1.12E-12 | 0.755166549 | 0.305 | 0.157 | 2.21E-08 | Tb |
| cDC2 | POLR2B | 3.38E-12 | 0.755162228 | 0.403 | 0.225 | 6.65E-08 | Tb |
| cDC2 | VPS8 | 1.97E-09 | 0.755027924 | 0.316 | 0.186 | 3.88E-05 | Tb |
| cDC2 | GPHN | 2.29E-10 | 0.754943844 | 0.295 | 0.162 | 4.52E-06 | Tb |
| cDC2 | LMBR1 | 6.92E-12 | 0.754453857 | 0.36 | 0.203 | 1.36E-07 | Tb |
| cDC2 | KANSL1L | 3.32E-12 | 0.752925324 | 0.391 | 0.218 | 6.54E-08 | Tb |
| cDC2 | SMURF2 | 2.44E-22 | 0.751522644 | 0.545 | 0.296 | 4.80E-18 | Tb |
| cDC2 | SNX13 | 7.41E-14 | 0.751264806 | 0.386 | 0.206 | 1.46E-09 | Tb |
| cDC2 | KDM3B | 1.41E-11 | 0.750526436 | 0.372 | 0.214 | 2.78E-07 | Tb |
| cDC2 | TASOR | 3.86E-15 | 0.75049823 | 0.373 | 0.197 | 7.61E-11 | Tb |
| cDC2 | SCIMP | 4.62E-14 | 0.749615515 | 0.466 | 0.276 | 9.09E-10 | Tb |
| cDC2 | NPLOC4 | 1.30E-23 | 0.749533855 | 0.48 | 0.23 | 2.57E-19 | Tb |
| cDC2 | PISD | 1.91E-09 | 0.748995917 | 0.305 | 0.174 | 3.77E-05 | Tb |
| cDC2 | RUBCN | 6.25E-08 | 0.74865638 | 0.318 | 0.194 | 0.001230879 | Tb |
| cDC2 | CD36 | 4.42E-08 | 0.748597324 | 0.274 | 0.164 | 0.000869983 | Tb |
| cDC2 | CPSF2 | 2.57E-14 | 0.748415318 | 0.277 | 0.129 | 5.06E-10 | Tb |
| cDC2 | MTMR6 | 2.82E-17 | 0.748301815 | 0.455 | 0.244 | 5.55E-13 | Tb |
| cDC2 | DENND10 | 6.02E-14 | 0.74757459 | 0.365 | 0.191 | 1.18E-09 | Tb |
| cDC2 | ELF2 | 2.26E-15 | 0.747394333 | 0.693 | 0.486 | 4.46E-11 | Tb |
| cDC2 | PCNX4 | 1.31E-10 | 0.746946082 | 0.339 | 0.197 | 2.58E-06 | Tb |
| cDC2 | PPCDC | 2.79E-09 | 0.746473425 | 0.307 | 0.178 | 5.50E-05 | Tb |
| cDC2 | AP5Z1 | 1.43E-17 | 0.746466882 | 0.387 | 0.194 | 2.81E-13 | Tb |
| cDC2 | CBLB | 4.85E-12 | 0.746074207 | 0.26 | 0.132 | 9.55E-08 | Tb |
| cDC2 | IL27RA | 8.11E-16 | 0.745673874 | 0.424 | 0.231 | 1.60E-11 | Tb |
| cDC2 | TNPO3 | 9.11E-14 | 0.745429014 | 0.377 | 0.2 | 1.79E-09 | Tb |
| cDC2 | ERBIN | 1.95E-23 | 0.745374126 | 0.7 | 0.445 | 3.84E-19 | Tb |
| cDC2 | PNPLA6 | 2.67E-14 | 0.745373192 | 0.347 | 0.176 | 5.26E-10 | Tb |
| cDC2 | OSCAR | 5.12E-16 | 0.74516765 | 0.309 | 0.142 | 1.01E-11 | Tb |
| cDC2 | BPTF | 7.07E-21 | 0.745093946 | 0.689 | 0.432 | 1.39E-16 | Tb |
| cDC2 | ATP11A | 1.09E-17 | 0.744730223 | 0.475 | 0.254 | 2.15E-13 | Tb |
| cDC2 | GPD2 | 3.54E-10 | 0.74440799 | 0.264 | 0.138 | 6.98E-06 | Tb |
| cDC2 | NIN | 5.28E-11 | 0.744188028 | 0.546 | 0.366 | 1.04E-06 | Tb |
| cDC2 | SENP5 | 4.66E-15 | 0.744186967 | 0.447 | 0.252 | 9.17E-11 | Tb |
| cDC2 | BCAS3 | 6.01E-12 | 0.74391264 | 0.36 | 0.208 | 1.18E-07 | Tb |
| cDC2 | COA6 | 3.15E-19 | 0.743236035 | 0.469 | 0.242 | 6.20E-15 | Tb |
| cDC2 | DLGAP4 | 7.54E-19 | 0.741557443 | 0.503 | 0.276 | 1.48E-14 | Tb |
| cDC2 | TBC1D5 | 4.54E-19 | 0.741458766 | 0.682 | 0.456 | 8.93E-15 | Tb |
| cDC2 | ALOX5 | 6.96E-22 | 0.741143772 | 0.705 | 0.452 | 1.37E-17 | Tb |
| cDC2 | PTRH2 | 5.42E-15 | 0.740808251 | 0.358 | 0.179 | 1.07E-10 | Tb |
| cDC2 | SLC30A7 | 8.52E-12 | 0.740750276 | 0.314 | 0.165 | 1.68E-07 | Tb |
| cDC2 | PDCD7 | 7.19E-13 | 0.740695635 | 0.326 | 0.17 | 1.41E-08 | Tb |
| cDC2 | PAN3 | 4.94E-21 | 0.740368951 | 0.59 | 0.343 | 9.73E-17 | Tb |
| cDC2 | INIP | 9.64E-20 | 0.73924157 | 0.361 | 0.161 | 1.90E-15 | Tb |
| cDC2 | CPB2-AS1 | 1.52E-10 | 0.738537517 | 0.363 | 0.211 | 3.00E-06 | Tb |

| | | | | | | | |
|---|---|---|---|---|---|---|---|
| cDC2 | PITPNB | 6.23E-17 | 0.738138326 | 0.562 | 0.331 | 1.23E-12 | Tb |
| cDC2 | DLG1 | 1.08E-15 | 0.738132016 | 0.36 | 0.181 | 2.13E-11 | Tb |
| cDC2 | MAP3K7 | 2.07E-08 | 0.738095254 | 0.26 | 0.146 | 0.000407923 | Tb |
| cDC2 | SEC31A | 3.96E-23 | 0.738027071 | 0.663 | 0.384 | 7.79E-19 | Tb |
| cDC2 | CASP10 | 1.23E-09 | 0.737857346 | 0.297 | 0.17 | 2.43E-05 | Tb |
| cDC2 | CEP128 | 4.53E-09 | 0.737265927 | 0.25 | 0.137 | 8.92E-05 | Tb |
| cDC2 | QRICH1 | 1.37E-14 | 0.736858483 | 0.312 | 0.153 | 2.70E-10 | Tb |
| cDC2 | SNAPC1 | 1.21E-06 | 0.736701407 | 0.309 | 0.2 | 0.023874158 | Tb |
| cDC2 | FMNL3 | 3.54E-16 | 0.736254777 | 0.36 | 0.178 | 6.97E-12 | Tb |
| cDC2 | TGIF2 | 5.99E-14 | 0.735983134 | 0.468 | 0.276 | 1.18E-09 | Tb |
| cDC2 | SLC20A2 | 4.75E-13 | 0.735761434 | 0.253 | 0.118 | 9.36E-09 | Tb |
| cDC2 | CNOT10 | 1.76E-10 | 0.735708965 | 0.356 | 0.21 | 3.47E-06 | Tb |
| cDC2 | RAB29 | 8.46E-11 | 0.735311312 | 0.384 | 0.234 | 1.67E-06 | Tb |
| cDC2 | XIAP | 2.09E-13 | 0.735290612 | 0.391 | 0.212 | 4.12E-09 | Tb |
| cDC2 | KYAT3 | 5.03E-12 | 0.734998266 | 0.255 | 0.122 | 9.91E-08 | Tb |
| cDC2 | SEMA4D | 1.30E-12 | 0.7342428 | 0.305 | 0.157 | 2.55E-08 | Tb |
| cDC2 | PHKB | 1.57E-10 | 0.733747943 | 0.363 | 0.214 | 3.10E-06 | Tb |
| cDC2 | GALM | 1.48E-14 | 0.732943578 | 0.508 | 0.318 | 2.92E-10 | Tb |
| cDC2 | LPIN2 | 8.67E-19 | 0.732831891 | 0.565 | 0.325 | 1.71E-14 | Tb |
| cDC2 | PIK3R5 | 8.46E-16 | 0.732807376 | 0.686 | 0.478 | 1.67E-11 | Tb |
| cDC2 | WRN | 4.79E-13 | 0.731701071 | 0.26 | 0.124 | 9.43E-09 | Tb |
| cDC2 | TOP1 | 6.99E-17 | 0.731694664 | 0.857 | 0.686 | 1.38E-12 | Tb |
| cDC2 | RABGEF1 | 4.28E-18 | 0.731522324 | 0.7 | 0.482 | 8.43E-14 | Tb |
| cDC2 | SRGAP2B | 1.37E-14 | 0.731515752 | 0.438 | 0.25 | 2.69E-10 | Tb |
| cDC2 | MTREX | 1.06E-11 | 0.730079128 | 0.358 | 0.202 | 2.09E-07 | Tb |
| cDC2 | MADD | 8.64E-14 | 0.729377091 | 0.251 | 0.113 | 1.70E-09 | Tb |
| cDC2 | NEK6 | 2.36E-16 | 0.728249162 | 0.433 | 0.232 | 4.65E-12 | Tb |
| cDC2 | ACSL1 | 1.31E-06 | 0.728135919 | 0.614 | 0.483 | 0.025812127 | Tb |
| cDC2 | BANP | 1.99E-14 | 0.727920048 | 0.675 | 0.472 | 3.92E-10 | Tb |
| cDC2 | SP140 | 1.99E-11 | 0.727557163 | 0.394 | 0.236 | 3.91E-07 | Tb |
| cDC2 | GAS2L3 | 3.67E-10 | 0.727197053 | 0.349 | 0.205 | 7.23E-06 | Tb |
| cDC2 | PFDN6 | 1.00E-13 | 0.726874345 | 0.321 | 0.159 | 1.98E-09 | Tb |
| cDC2 | SNRK | 1.13E-09 | 0.726001778 | 0.295 | 0.167 | 2.23E-05 | Tb |
| cDC2 | RAB3GAP2 | 4.43E-13 | 0.725182816 | 0.277 | 0.135 | 8.73E-09 | Tb |
| cDC2 | EXOC4 | 8.35E-23 | 0.725043726 | 0.665 | 0.407 | 1.64E-18 | Tb |
| cDC2 | INSIG2 | 7.36E-11 | 0.724491199 | 0.258 | 0.131 | 1.45E-06 | Tb |
| cDC2 | EMG1 | 4.05E-12 | 0.724437471 | 0.321 | 0.168 | 7.97E-08 | Tb |
| cDC2 | MBD5 | 3.23E-12 | 0.723494751 | 0.386 | 0.227 | 6.35E-08 | Tb |
| cDC2 | ZNF490 | 5.93E-15 | 0.72349261 | 0.271 | 0.121 | 1.17E-10 | Tb |
| cDC2 | PRKX | 1.14E-18 | 0.723112768 | 0.414 | 0.205 | 2.24E-14 | Tb |
| cDC2 | B4GALT4 | 2.42E-08 | 0.722703116 | 0.281 | 0.164 | 0.000476874 | Tb |
| cDC2 | ZBTB20 | 2.57E-14 | 0.722101755 | 0.557 | 0.368 | 5.05E-10 | Tb |
| cDC2 | RAD50 | 5.16E-16 | 0.722019038 | 0.391 | 0.202 | 1.02E-11 | Tb |
| cDC2 | CLOCK | 8.33E-07 | 0.721972531 | 0.29 | 0.182 | 0.016405002 | Tb |
| cDC2 | CCAR1 | 5.15E-21 | 0.721798817 | 0.55 | 0.301 | 1.01E-16 | Tb |
| cDC2 | PTK2 | 8.71E-10 | 0.721142632 | 0.3 | 0.171 | 1.72E-05 | Tb |
| cDC2 | NMT1 | 2.60E-20 | 0.720540974 | 0.496 | 0.253 | 5.13E-16 | Tb |
| cDC2 | COMMD10 | 2.13E-15 | 0.720252383 | 0.6 | 0.373 | 4.19E-11 | Tb |
| cDC2 | KAT6A | 1.95E-15 | 0.72007129 | 0.476 | 0.278 | 3.84E-11 | Tb |
| cDC2 | TMEM273 | 9.12E-14 | 0.719820399 | 0.454 | 0.268 | 1.80E-09 | Tb |
| cDC2 | IL4I1 | 3.01E-13 | 0.719433775 | 0.775 | 0.612 | 5.92E-09 | Tb |
| cDC2 | ATF6 | 9.22E-16 | 0.719362152 | 0.745 | 0.532 | 1.82E-11 | Tb |
| cDC2 | USP14 | 7.93E-13 | 0.71888926 | 0.504 | 0.308 | 1.56E-08 | Tb |
| cDC2 | CLCN5 | 1.02E-10 | 0.718140072 | 0.377 | 0.228 | 2.00E-06 | Tb |
| cDC2 | RNMT | 2.34E-22 | 0.71795056 | 0.665 | 0.381 | 4.60E-18 | Tb |
| cDC2 | ABI2 | 1.77E-09 | 0.717139485 | 0.297 | 0.17 | 3.48E-05 | Tb |

| | | | | | | | |
|---|---|---|---|---|---|---|---|
| cDC2 | RFTN1 | 9.26E-10 | 0.716817441 | 0.733 | 0.62 | 1.82E-05 | Tb |
| cDC2 | ITCH | 1.81E-18 | 0.716775948 | 0.471 | 0.248 | 3.56E-14 | Tb |
| cDC2 | FBXO28 | 5.29E-13 | 0.716355394 | 0.276 | 0.132 | 1.04E-08 | Tb |
| cDC2 | CHSY1 | 4.08E-11 | 0.716328664 | 0.349 | 0.201 | 8.03E-07 | Tb |
| cDC2 | ATF5 | 2.84E-11 | 0.716292674 | 0.756 | 0.614 | 5.58E-07 | Tb |
| cDC2 | RPRD1A | 1.15E-10 | 0.715929162 | 0.269 | 0.14 | 2.25E-06 | Tb |
| cDC2 | SF3B3 | 4.79E-15 | 0.71517163 | 0.314 | 0.15 | 9.44E-11 | Tb |
| cDC2 | EHMT1 | 7.04E-21 | 0.715000123 | 0.546 | 0.298 | 1.39E-16 | Tb |
| cDC2 | CEP135 | 9.45E-11 | 0.714930129 | 0.276 | 0.145 | 1.86E-06 | Tb |
| cDC2 | NRBF2 | 1.28E-23 | 0.713513633 | 0.609 | 0.344 | 2.52E-19 | Tb |
| cDC2 | SH2B3 | 1.49E-14 | 0.713412334 | 0.674 | 0.504 | 2.93E-10 | Tb |
| cDC2 | TTC3 | 7.04E-23 | 0.712378112 | 0.642 | 0.382 | 1.39E-18 | Tb |
| cDC2 | P2RY6 | 1.44E-11 | 0.712313735 | 0.33 | 0.186 | 2.84E-07 | Tb |
| cDC2 | CTBP2 | 3.94E-16 | 0.712284442 | 0.588 | 0.365 | 7.75E-12 | Tb |
| cDC2 | BTAF1 | 9.77E-16 | 0.712171956 | 0.574 | 0.365 | 1.92E-11 | Tb |
| cDC2 | DTNBP1 | 7.59E-22 | 0.712061742 | 0.642 | 0.377 | 1.49E-17 | Tb |
| cDC2 | NAPA | 1.04E-21 | 0.711779232 | 0.862 | 0.666 | 2.04E-17 | Tb |
| cDC2 | HIVEP2 | 2.44E-11 | 0.711404114 | 0.719 | 0.54 | 4.81E-07 | Tb |
| cDC2 | MSH3 | 1.67E-09 | 0.711253998 | 0.271 | 0.148 | 3.28E-05 | Tb |
| cDC2 | PXK | 4.39E-14 | 0.71100336 | 0.358 | 0.187 | 8.64E-10 | Tb |
| cDC2 | CD2AP | 8.17E-08 | 0.71033064 | 0.342 | 0.218 | 0.001608011 | Tb |
| cDC2 | RPTOR | 6.20E-16 | 0.710272763 | 0.375 | 0.188 | 1.22E-11 | Tb |
| cDC2 | PPP6R3 | 7.94E-17 | 0.710208743 | 0.647 | 0.412 | 1.56E-12 | Tb |
| cDC2 | ESCO1 | 8.34E-15 | 0.709990774 | 0.424 | 0.235 | 1.64E-10 | Tb |
| cDC2 | RAB3GAP1 | 3.91E-15 | 0.709990305 | 0.457 | 0.26 | 7.70E-11 | Tb |
| cDC2 | PLCG2 | 1.70E-14 | 0.709379941 | 0.487 | 0.278 | 3.35E-10 | Tb |
| cDC2 | NFATC2 | 1.37E-11 | 0.708821222 | 0.435 | 0.268 | 2.70E-07 | Tb |
| cDC2 | UBA6 | 9.89E-14 | 0.707491877 | 0.473 | 0.275 | 1.95E-09 | Tb |
| cDC2 | DESI1 | 1.28E-18 | 0.706460204 | 0.546 | 0.309 | 2.52E-14 | Tb |
| cDC2 | APOL6 | 1.03E-15 | 0.706252054 | 0.49 | 0.278 | 2.02E-11 | Tb |
| cDC2 | NOP56 | 1.41E-20 | 0.705760803 | 0.462 | 0.227 | 2.78E-16 | Tb |
| cDC2 | PFDN1 | 1.79E-15 | 0.704547758 | 0.571 | 0.365 | 3.53E-11 | Tb |
| cDC2 | PSEN1 | 3.77E-18 | 0.704134467 | 0.574 | 0.344 | 7.42E-14 | Tb |
| cDC2 | PTPN22 | 1.69E-06 | 0.70356565 | 0.384 | 0.26 | 0.033287104 | Tb |
| cDC2 | MFSD12 | 1.28E-11 | 0.70325117 | 0.576 | 0.389 | 2.53E-07 | Tb |
| cDC2 | UBAP2L | 1.23E-21 | 0.703226158 | 0.614 | 0.341 | 2.41E-17 | Tb |
| cDC2 | CSTF3 | 2.83E-14 | 0.703185896 | 0.283 | 0.132 | 5.56E-10 | Tb |
| cDC2 | SS18 | 1.70E-13 | 0.703057672 | 0.393 | 0.211 | 3.34E-09 | Tb |
| cDC2 | ESF1 | 2.70E-13 | 0.702395059 | 0.34 | 0.179 | 5.32E-09 | Tb |
| cDC2 | LIMK1 | 4.10E-19 | 0.701547504 | 0.377 | 0.173 | 8.08E-15 | Tb |
| cDC2 | USP34 | 1.30E-21 | 0.70124643 | 0.761 | 0.502 | 2.56E-17 | Tb |
| cDC2 | RNF213 | 3.27E-25 | 0.701167872 | 0.845 | 0.577 | 6.44E-21 | Tb |
| cDC2 | JPX | 1.29E-17 | 0.701083429 | 0.471 | 0.249 | 2.55E-13 | Tb |
| cDC2 | SEMA4A | 1.22E-16 | 0.700942164 | 0.569 | 0.339 | 2.41E-12 | Tb |
| cDC2 | ITGA4 | 7.66E-12 | 0.699316677 | 0.677 | 0.49 | 1.51E-07 | Tb |
| cDC2 | S100A9 | 2.48E-60 | 0.699298885 | 0.743 | 0.325 | 4.87E-56 | Tb |
| cDC2 | TAOK3 | 3.51E-23 | 0.699187791 | 0.895 | 0.747 | 6.90E-19 | Tb |
| cDC2 | IL15 | 1.25E-13 | 0.698927291 | 0.579 | 0.388 | 2.47E-09 | Tb |
| cDC2 | JAK2 | 1.02E-15 | 0.697526877 | 0.796 | 0.585 | 2.02E-11 | Tb |
| cDC2 | NSRP1 | 1.82E-17 | 0.697463922 | 0.501 | 0.274 | 3.58E-13 | Tb |
| cDC2 | POLR2A | 7.32E-16 | 0.697171181 | 0.539 | 0.331 | 1.44E-11 | Tb |
| cDC2 | CEP68 | 3.97E-14 | 0.697079325 | 0.431 | 0.242 | 7.80E-10 | Tb |
| cDC2 | PPA1 | 7.76E-29 | 0.697039875 | 0.894 | 0.781 | 1.53E-24 | Tb |
| cDC2 | POLG2 | 1.47E-07 | 0.696861722 | 0.305 | 0.191 | 0.002885645 | Tb |
| cDC2 | NUDCD1 | 2.22E-13 | 0.69660903 | 0.281 | 0.135 | 4.38E-09 | Tb |
| cDC2 | LRRC8C | 3.43E-11 | 0.696546664 | 0.503 | 0.328 | 6.74E-07 | Tb |

| | | | | | | | |
|---|---|---|---|---|---|---|---|
| cDC2 | PPM1D | 3.23E-16 | 0.694707162 | 0.429 | 0.227 | 6.36E-12 | Tb |
| cDC2 | ISCA2 | 2.18E-13 | 0.694477632 | 0.325 | 0.161 | 4.28E-09 | Tb |
| cDC2 | PPA2 | 4.54E-17 | 0.694453736 | 0.558 | 0.324 | 8.94E-13 | Tb |
| cDC2 | ELOVL5 | 2.36E-16 | 0.694057518 | 0.812 | 0.629 | 4.65E-12 | Tb |
| cDC2 | ATF7IP2 | 8.06E-16 | 0.693935372 | 0.469 | 0.271 | 1.59E-11 | Tb |
| cDC2 | SPINT2 | 7.50E-29 | 0.693520721 | 0.829 | 0.615 | 1.48E-24 | Tb |
| cDC2 | TIMM9 | 1.10E-12 | 0.692264424 | 0.288 | 0.142 | 2.17E-08 | Tb |
| cDC2 | FAM241A | 4.26E-12 | 0.692091745 | 0.56 | 0.373 | 8.38E-08 | Tb |
| cDC2 | RBM23 | 1.93E-17 | 0.691943034 | 0.44 | 0.23 | 3.80E-13 | Tb |
| cDC2 | SLC43A2 | 1.37E-14 | 0.691726587 | 0.658 | 0.449 | 2.70E-10 | Tb |
| cDC2 | MACF1 | 7.31E-24 | 0.691558047 | 0.749 | 0.475 | 1.44E-19 | Tb |
| cDC2 | GAB2 | 5.66E-19 | 0.691151772 | 0.716 | 0.484 | 1.11E-14 | Tb |
| cDC2 | SHB | 1.23E-10 | 0.691093084 | 0.403 | 0.243 | 2.42E-06 | Tb |
| cDC2 | PTPRA | 1.86E-13 | 0.691076848 | 0.424 | 0.238 | 3.66E-09 | Tb |
| cDC2 | AIDA | 5.58E-10 | 0.69105297 | 0.272 | 0.146 | 1.10E-05 | Tb |
| cDC2 | RHOT1 | 8.85E-14 | 0.690704719 | 0.429 | 0.248 | 1.74E-09 | Tb |
| cDC2 | ABCC1 | 1.01E-13 | 0.690668333 | 0.387 | 0.213 | 1.99E-09 | Tb |
| cDC2 | KPNB1 | 2.40E-22 | 0.690201342 | 0.682 | 0.423 | 4.72E-18 | Tb |
| cDC2 | TMEM176A | 4.59E-11 | 0.689660376 | 0.417 | 0.264 | 9.04E-07 | Tb |
| cDC2 | SLC27A4 | 6.91E-21 | 0.689368765 | 0.461 | 0.225 | 1.36E-16 | Tb |
| cDC2 | GPATCH2 | 3.93E-11 | 0.689062631 | 0.311 | 0.168 | 7.73E-07 | Tb |
| cDC2 | ZBTB21 | 4.08E-09 | 0.688928431 | 0.286 | 0.161 | 8.04E-05 | Tb |
| cDC2 | CPSF3 | 2.20E-12 | 0.6887324 | 0.288 | 0.145 | 4.33E-08 | Tb |
| cDC2 | MED13L | 7.80E-23 | 0.688229896 | 0.888 | 0.695 | 1.54E-18 | Tb |
| cDC2 | MRTO4 | 1.70E-11 | 0.687676707 | 0.328 | 0.181 | 3.34E-07 | Tb |
| cDC2 | ZC3H13 | 3.49E-20 | 0.68744289 | 0.527 | 0.276 | 6.87E-16 | Tb |
| cDC2 | MAPK8 | 3.85E-10 | 0.687356121 | 0.499 | 0.328 | 7.57E-06 | Tb |
| cDC2 | DNAJC13 | 6.37E-18 | 0.687131016 | 0.422 | 0.217 | 1.25E-13 | Tb |
| cDC2 | CNOT2 | 3.39E-13 | 0.686834028 | 0.686 | 0.485 | 6.68E-09 | Tb |
| cDC2 | RREB1 | 1.62E-17 | 0.686750698 | 0.574 | 0.345 | 3.18E-13 | Tb |
| cDC2 | PMM2 | 5.49E-12 | 0.686549441 | 0.34 | 0.182 | 1.08E-07 | Tb |
| cDC2 | CPEB2 | 4.41E-11 | 0.68639506 | 0.396 | 0.237 | 8.69E-07 | Tb |
| cDC2 | CUL1 | 3.88E-20 | 0.686305128 | 0.738 | 0.504 | 7.64E-16 | Tb |
| cDC2 | MAP3K5 | 1.61E-16 | 0.685964826 | 0.462 | 0.259 | 3.17E-12 | Tb |
| cDC2 | PPP2R5E | 1.16E-14 | 0.685422848 | 0.459 | 0.265 | 2.28E-10 | Tb |
| cDC2 | SND1 | 3.16E-18 | 0.685220933 | 0.74 | 0.483 | 6.21E-14 | Tb |
| cDC2 | ASH1L | 2.20E-18 | 0.685180608 | 0.75 | 0.517 | 4.34E-14 | Tb |
| cDC2 | AHI1 | 1.50E-10 | 0.684724204 | 0.375 | 0.219 | 2.95E-06 | Tb |
| cDC2 | CYB561A3 | 5.70E-09 | 0.684684895 | 0.3 | 0.17 | 0.000112209 | Tb |
| cDC2 | P2RY13 | 2.30E-14 | 0.684634695 | 0.485 | 0.294 | 4.54E-10 | Tb |
| cDC2 | KIF16B | 7.93E-12 | 0.684062625 | 0.279 | 0.142 | 1.56E-07 | Tb |
| cDC2 | TAF3 | 1.18E-12 | 0.683469095 | 0.304 | 0.154 | 2.32E-08 | Tb |
| cDC2 | SLC44A1 | 1.03E-08 | 0.682768334 | 0.311 | 0.181 | 0.000201884 | Tb |
| cDC2 | ARID1B | 6.36E-23 | 0.682202633 | 0.735 | 0.481 | 1.25E-18 | Tb |
| cDC2 | PIP5K1C | 1.22E-12 | 0.682115981 | 0.281 | 0.139 | 2.40E-08 | Tb |
| cDC2 | NSD3 | 2.79E-14 | 0.68203381 | 0.64 | 0.428 | 5.48E-10 | Tb |
| cDC2 | CLPTM1L | 5.14E-14 | 0.681617119 | 0.387 | 0.2 | 1.01E-09 | Tb |
| cDC2 | LARP1 | 1.19E-14 | 0.681116083 | 0.489 | 0.28 | 2.35E-10 | Tb |
| cDC2 | SLC7A5 | 1.05E-09 | 0.68090795 | 0.627 | 0.463 | 2.06E-05 | Tb |
| cDC2 | CPEB4 | 4.51E-20 | 0.68044318 | 0.635 | 0.395 | 8.88E-16 | Tb |
| cDC2 | SPEN | 3.94E-11 | 0.679981404 | 0.656 | 0.47 | 7.75E-07 | Tb |
| cDC2 | CAP1 | 5.94E-31 | 0.679775289 | 0.897 | 0.748 | 1.17E-26 | Tb |
| cDC2 | IRF4 | 3.25E-16 | 0.679404495 | 0.661 | 0.45 | 6.40E-12 | Tb |
| cDC2 | EYA3 | 1.23E-17 | 0.677930831 | 0.466 | 0.252 | 2.42E-13 | Tb |
| cDC2 | STRN3 | 1.70E-16 | 0.677224661 | 0.405 | 0.21 | 3.35E-12 | Tb |
| cDC2 | NKAP | 3.62E-15 | 0.676077657 | 0.372 | 0.189 | 7.13E-11 | Tb |

| | | | | | | | |
|---|---|---|---|---|---|---|---|
| cDC2 | RNF149 | 3.58E-10 | 0.676064681 | 0.81 | 0.682 | 7.05E-06 | Tb |
| cDC2 | NEK7 | 3.45E-12 | 0.676033953 | 0.459 | 0.278 | 6.80E-08 | Tb |
| cDC2 | MTMR12 | 2.28E-12 | 0.675846084 | 0.318 | 0.167 | 4.48E-08 | Tb |
| cDC2 | SMCHD1 | 1.43E-07 | 0.675724267 | 0.829 | 0.669 | 0.002814373 | Tb |
| cDC2 | CREBBP | 9.23E-16 | 0.675586608 | 0.625 | 0.392 | 1.82E-11 | Tb |
| cDC2 | ENTPD1-AS1 | 6.19E-20 | 0.675455038 | 0.56 | 0.316 | 1.22E-15 | Tb |
| cDC2 | EEA1 | 8.41E-16 | 0.675444588 | 0.569 | 0.35 | 1.66E-11 | Tb |
| cDC2 | LCOR | 9.06E-14 | 0.675254288 | 0.576 | 0.361 | 1.78E-09 | Tb |
| cDC2 | RASSF3 | 8.10E-15 | 0.675165149 | 0.771 | 0.569 | 1.59E-10 | Tb |
| cDC2 | RNASEH1 | 4.69E-13 | 0.674487396 | 0.325 | 0.166 | 9.23E-09 | Tb |
| cDC2 | EGR2 | 1.18E-23 | 0.674339352 | 0.461 | 0.221 | 2.33E-19 | Tb |
| cDC2 | CFDP1 | 4.42E-19 | 0.674080629 | 0.524 | 0.283 | 8.70E-15 | Tb |
| cDC2 | TRPS1 | 1.90E-16 | 0.673819136 | 0.768 | 0.553 | 3.75E-12 | Tb |
| cDC2 | ARSB | 4.96E-14 | 0.673072792 | 0.393 | 0.212 | 9.76E-10 | Tb |
| cDC2 | MAPK1 | 2.65E-16 | 0.672824594 | 0.628 | 0.396 | 5.21E-12 | Tb |
| cDC2 | GPAT4 | 2.39E-16 | 0.672791428 | 0.442 | 0.232 | 4.70E-12 | Tb |
| cDC2 | KIAA0040 | 7.50E-11 | 0.67277617 | 0.304 | 0.164 | 1.48E-06 | Tb |
| cDC2 | KDM3A | 7.80E-13 | 0.672690078 | 0.298 | 0.151 | 1.54E-08 | Tb |
| cDC2 | SAE1 | 2.01E-12 | 0.672263451 | 0.351 | 0.191 | 3.95E-08 | Tb |
| cDC2 | RAB11A | 6.47E-10 | 0.672172001 | 0.801 | 0.673 | 1.27E-05 | Tb |
| cDC2 | TRANK1 | 5.60E-10 | 0.672151227 | 0.25 | 0.132 | 1.10E-05 | Tb |
| cDC2 | NF1 | 1.12E-14 | 0.67184899 | 0.583 | 0.379 | 2.20E-10 | Tb |
| cDC2 | ALOX12-AS1 | 1.09E-13 | 0.670888182 | 0.281 | 0.134 | 2.14E-09 | Tb |
| cDC2 | ADPGK | 3.10E-17 | 0.670463571 | 0.52 | 0.294 | 6.09E-13 | Tb |
| cDC2 | PAK1 | 3.72E-14 | 0.670437599 | 0.909 | 0.787 | 7.32E-10 | Tb |
| cDC2 | CMAS | 4.64E-14 | 0.67007621 | 0.319 | 0.154 | 9.13E-10 | Tb |
| cDC2 | IDO2 | 3.15E-13 | 0.669353415 | 0.553 | 0.352 | 6.19E-09 | Tb |
| cDC2 | DIAPH2 | 3.56E-13 | 0.669194176 | 0.627 | 0.422 | 7.02E-09 | Tb |
| cDC2 | RXRA | 1.51E-09 | 0.669158477 | 0.379 | 0.227 | 2.98E-05 | Tb |
| cDC2 | CHFR | 2.70E-16 | 0.668948581 | 0.401 | 0.203 | 5.31E-12 | Tb |
| cDC2 | MAP3K2 | 2.53E-19 | 0.668542361 | 0.817 | 0.599 | 4.99E-15 | Tb |
| cDC2 | SLMAP | 2.75E-15 | 0.668434552 | 0.408 | 0.216 | 5.41E-11 | Tb |
| cDC2 | INO80D | 4.84E-14 | 0.668296188 | 0.339 | 0.173 | 9.52E-10 | Tb |
| cDC2 | GAK | 1.80E-14 | 0.667741227 | 0.513 | 0.3 | 3.55E-10 | Tb |
| cDC2 | FAM135A | 1.13E-14 | 0.667506303 | 0.403 | 0.224 | 2.23E-10 | Tb |
| cDC2 | CTNNAL1 | 9.82E-10 | 0.666875484 | 0.269 | 0.146 | 1.93E-05 | Tb |
| cDC2 | ZNF736 | 2.68E-09 | 0.666836932 | 0.251 | 0.135 | 5.28E-05 | Tb |
| cDC2 | DNAJC1 | 1.59E-11 | 0.666625102 | 0.597 | 0.404 | 3.14E-07 | Tb |
| cDC2 | INO80 | 1.78E-11 | 0.666584672 | 0.394 | 0.233 | 3.50E-07 | Tb |
| cDC2 | HNRNPUL1 | 1.99E-17 | 0.66624436 | 0.667 | 0.425 | 3.92E-13 | Tb |
| cDC2 | GNB1 | 4.07E-25 | 0.665236953 | 0.853 | 0.656 | 8.00E-21 | Tb |
| cDC2 | ABHD5 | 8.24E-09 | 0.66479314 | 0.379 | 0.232 | 0.000162219 | Tb |
| cDC2 | FAM89B | 1.02E-21 | 0.664590312 | 0.658 | 0.409 | 2.01E-17 | Tb |
| cDC2 | TRAF3IP3 | 1.48E-09 | 0.664374906 | 0.333 | 0.199 | 2.92E-05 | Tb |
| cDC2 | RAF1 | 2.98E-16 | 0.663715866 | 0.469 | 0.259 | 5.86E-12 | Tb |
| cDC2 | APOBR | 6.50E-19 | 0.663461055 | 0.396 | 0.188 | 1.28E-14 | Tb |
| cDC2 | NAF1 | 1.53E-10 | 0.663177951 | 0.546 | 0.378 | 3.02E-06 | Tb |
| cDC2 | ARL15 | 1.03E-13 | 0.663097059 | 0.459 | 0.271 | 2.03E-09 | Tb |
| cDC2 | ZCCHC17 | 1.20E-13 | 0.661738175 | 0.394 | 0.211 | 2.36E-09 | Tb |
| cDC2 | SUMF1 | 1.55E-13 | 0.661706788 | 0.538 | 0.339 | 3.05E-09 | Tb |
| cDC2 | GPR137B | 1.68E-11 | 0.661267314 | 0.768 | 0.619 | 3.31E-07 | Tb |
| cDC2 | NAA35 | 4.75E-11 | 0.66126431 | 0.281 | 0.147 | 9.35E-07 | Tb |
| cDC2 | UFL1 | 1.47E-14 | 0.661109113 | 0.391 | 0.208 | 2.90E-10 | Tb |
| cDC2 | AVL9 | 1.48E-14 | 0.661072732 | 0.351 | 0.178 | 2.91E-10 | Tb |
| cDC2 | DOCK7 | 2.38E-11 | 0.660919287 | 0.314 | 0.169 | 4.69E-07 | Tb |
| cDC2 | RIPOR2 | 3.84E-07 | 0.660911144 | 0.464 | 0.319 | 0.007553573 | Tb |

| | | | | | | | |
|---|---|---|---|---|---|---|---|
| cDC2 | SBNO2 | 8.00E-17 | 0.660169271 | 0.564 | 0.325 | 1.57E-12 | Tb |
| cDC2 | GAR1 | 2.89E-14 | 0.660063001 | 0.297 | 0.14 | 5.69E-10 | Tb |
| cDC2 | NEAT1 | 2.98E-38 | 0.658980718 | 0.981 | 0.915 | 5.87E-34 | Tb |
| cDC2 | RFX3 | 1.80E-10 | 0.658629439 | 0.29 | 0.159 | 3.55E-06 | Tb |
| cDC2 | UBE2E2 | 1.39E-27 | 0.658308601 | 0.918 | 0.763 | 2.74E-23 | Tb |
| cDC2 | MAP3K14 | 4.66E-13 | 0.658172403 | 0.372 | 0.202 | 9.17E-09 | Tb |
| cDC2 | ZNF721 | 1.14E-13 | 0.658153929 | 0.468 | 0.262 | 2.25E-09 | Tb |
| cDC2 | BET1 | 2.39E-16 | 0.658134685 | 0.281 | 0.122 | 4.70E-12 | Tb |
| cDC2 | NAA25 | 4.14E-09 | 0.657236873 | 0.276 | 0.152 | 8.15E-05 | Tb |
| cDC2 | LTV1 | 9.67E-15 | 0.656785746 | 0.391 | 0.208 | 1.90E-10 | Tb |
| cDC2 | TBC1D14 | 2.91E-11 | 0.656379335 | 0.353 | 0.197 | 5.73E-07 | Tb |
| cDC2 | RFFL | 2.77E-12 | 0.655689917 | 0.269 | 0.13 | 5.46E-08 | Tb |
| cDC2 | HIST1H2AC | 2.53E-09 | 0.654682329 | 0.311 | 0.18 | 4.98E-05 | Tb |
| cDC2 | PRR14L | 1.25E-15 | 0.654638984 | 0.347 | 0.172 | 2.46E-11 | Tb |
| cDC2 | ETV6 | 2.72E-20 | 0.654328574 | 0.941 | 0.868 | 5.34E-16 | Tb |
| cDC2 | TMEM176B | 2.17E-15 | 0.654130516 | 0.532 | 0.346 | 4.27E-11 | Tb |
| cDC2 | PARP12 | 2.05E-14 | 0.654109706 | 0.415 | 0.226 | 4.04E-10 | Tb |
| cDC2 | APC | 1.28E-11 | 0.653951424 | 0.36 | 0.199 | 2.51E-07 | Tb |
| cDC2 | BCL2L11 | 7.71E-14 | 0.65362985 | 0.658 | 0.442 | 1.52E-09 | Tb |
| cDC2 | PRDM1 | 1.43E-12 | 0.653464884 | 0.614 | 0.432 | 2.82E-08 | Tb |
| cDC2 | DMXL1 | 1.21E-14 | 0.653336212 | 0.517 | 0.31 | 2.38E-10 | Tb |
| cDC2 | AFF4 | 5.42E-19 | 0.652761991 | 0.836 | 0.679 | 1.07E-14 | Tb |
| cDC2 | GGA2 | 9.43E-12 | 0.65241532 | 0.379 | 0.221 | 1.86E-07 | Tb |
| cDC2 | FYN | 9.37E-10 | 0.652211344 | 0.689 | 0.513 | 1.84E-05 | Tb |
| cDC2 | ATP11B | 1.31E-17 | 0.651534556 | 0.545 | 0.321 | 2.58E-13 | Tb |
| cDC2 | USP36 | 5.57E-19 | 0.650861749 | 0.695 | 0.434 | 1.10E-14 | Tb |
| cDC2 | B3GNTL1 | 1.28E-14 | 0.650547289 | 0.265 | 0.118 | 2.52E-10 | Tb |
| cDC2 | GFPT1 | 3.12E-14 | 0.650219282 | 0.387 | 0.207 | 6.14E-10 | Tb |
| cDC2 | ZNF638 | 7.73E-17 | 0.650032542 | 0.595 | 0.362 | 1.52E-12 | Tb |
| cDC2 | NOP58 | 4.26E-14 | 0.647840669 | 0.639 | 0.433 | 8.38E-10 | Tb |
| cDC2 | CAPZA1 | 2.05E-25 | 0.647821603 | 0.808 | 0.548 | 4.04E-21 | Tb |
| cDC2 | DAPP1 | 1.49E-14 | 0.647488668 | 0.784 | 0.617 | 2.93E-10 | Tb |
| cDC2 | PCBD2 | 1.00E-14 | 0.647162723 | 0.264 | 0.116 | 1.97E-10 | Tb |
| cDC2 | UBR1 | 2.63E-09 | 0.646934648 | 0.398 | 0.249 | 5.18E-05 | Tb |
| cDC2 | ATG2A | 6.84E-11 | 0.646702774 | 0.276 | 0.143 | 1.35E-06 | Tb |
| cDC2 | PTAFR | 2.25E-14 | 0.64661202 | 0.445 | 0.259 | 4.42E-10 | Tb |
| cDC2 | UBE2E1 | 1.21E-13 | 0.646483121 | 0.728 | 0.545 | 2.38E-09 | Tb |
| cDC2 | UBE2O | 3.44E-12 | 0.646422732 | 0.342 | 0.187 | 6.78E-08 | Tb |
| cDC2 | SLC12A6 | 2.66E-10 | 0.646055526 | 0.4 | 0.245 | 5.24E-06 | Tb |
| cDC2 | YEATS2 | 5.85E-08 | 0.645074243 | 0.274 | 0.16 | 0.001151122 | Tb |
| cDC2 | SH3BP4 | 1.49E-10 | 0.644924392 | 0.344 | 0.195 | 2.93E-06 | Tb |
| cDC2 | AGO3 | 8.90E-18 | 0.644666149 | 0.578 | 0.344 | 1.75E-13 | Tb |
| cDC2 | PDPK1 | 7.87E-16 | 0.644554248 | 0.459 | 0.251 | 1.55E-11 | Tb |
| cDC2 | USP11 | 3.04E-12 | 0.642556219 | 0.297 | 0.154 | 5.98E-08 | Tb |
| cDC2 | SEC24B | 1.97E-12 | 0.64238474 | 0.394 | 0.221 | 3.88E-08 | Tb |
| cDC2 | WIPF2 | 4.49E-11 | 0.642254014 | 0.283 | 0.148 | 8.85E-07 | Tb |
| cDC2 | FLVCR2 | 1.22E-07 | 0.641023933 | 0.274 | 0.162 | 0.002409794 | Tb |
| cDC2 | RABEP2 | 1.77E-13 | 0.640559097 | 0.267 | 0.126 | 3.49E-09 | Tb |
| cDC2 | ESRRA | 2.03E-18 | 0.640555558 | 0.466 | 0.24 | 3.99E-14 | Tb |
| cDC2 | OGA | 4.63E-14 | 0.640430833 | 0.431 | 0.241 | 9.12E-10 | Tb |
| cDC2 | CMIP | 1.78E-15 | 0.640332907 | 0.78 | 0.574 | 3.50E-11 | Tb |
| cDC2 | CYB5R1 | 9.03E-10 | 0.640130568 | 0.253 | 0.131 | 1.78E-05 | Tb |
| cDC2 | MANF | 1.03E-17 | 0.640070914 | 0.567 | 0.336 | 2.03E-13 | Tb |
| cDC2 | RLF | 3.24E-13 | 0.639879725 | 0.719 | 0.529 | 6.38E-09 | Tb |
| cDC2 | SMG1 | 2.22E-17 | 0.639571918 | 0.756 | 0.531 | 4.37E-13 | Tb |
| cDC2 | MTMR3 | 1.56E-11 | 0.638890321 | 0.356 | 0.198 | 3.06E-07 | Tb |

| | | | | | | | |
|---|---|---|---|---|---|---|---|
| cDC2 | CASK | 2.37E-09 | 0.63854941 | 0.274 | 0.152 | 4.67E-05 | Tb |
| cDC2 | ZMPSTE24 | 1.67E-11 | 0.637987571 | 0.255 | 0.126 | 3.29E-07 | Tb |
| cDC2 | CNTLN | 6.76E-10 | 0.637719369 | 0.363 | 0.22 | 1.33E-05 | Tb |
| cDC2 | ZFX | 4.57E-13 | 0.637676757 | 0.44 | 0.254 | 9.00E-09 | Tb |
| cDC2 | RBMX2 | 1.20E-10 | 0.637587023 | 0.258 | 0.134 | 2.37E-06 | Tb |
| cDC2 | HEATR5B | 5.23E-10 | 0.63746125 | 0.346 | 0.196 | 1.03E-05 | Tb |
| cDC2 | NOC2L | 1.75E-08 | 0.636959844 | 0.267 | 0.151 | 0.000344456 | Tb |
| cDC2 | SMURF1 | 5.37E-14 | 0.636117925 | 0.407 | 0.228 | 1.06E-09 | Tb |
| cDC2 | CBR1 | 1.79E-13 | 0.635673288 | 0.461 | 0.269 | 3.52E-09 | Tb |
| cDC2 | SEC24D | 1.21E-06 | 0.635421124 | 0.412 | 0.287 | 0.023890129 | Tb |
| cDC2 | SORT1 | 3.55E-12 | 0.635383841 | 0.255 | 0.123 | 6.99E-08 | Tb |
| cDC2 | EIF2AK3 | 2.84E-09 | 0.634609354 | 0.679 | 0.505 | 5.58E-05 | Tb |
| cDC2 | TNFRSF10B | 1.42E-10 | 0.634326169 | 0.693 | 0.542 | 2.80E-06 | Tb |
| cDC2 | PARM1 | 3.90E-10 | 0.634306836 | 0.339 | 0.202 | 7.67E-06 | Tb |
| cDC2 | NCOA2 | 1.44E-11 | 0.634132097 | 0.417 | 0.249 | 2.84E-07 | Tb |
| cDC2 | MAEA | 1.50E-12 | 0.63322863 | 0.339 | 0.18 | 2.95E-08 | Tb |
| cDC2 | EIF4G3 | 1.24E-12 | 0.633196608 | 0.569 | 0.379 | 2.45E-08 | Tb |
| cDC2 | SCYL2 | 3.26E-14 | 0.633152014 | 0.478 | 0.27 | 6.41E-10 | Tb |
| cDC2 | ARID1A | 2.52E-20 | 0.633046816 | 0.504 | 0.262 | 4.97E-16 | Tb |
| cDC2 | SLC41A2 | 3.85E-09 | 0.632463425 | 0.513 | 0.347 | 7.59E-05 | Tb |
| cDC2 | SATB1-AS1 | 1.03E-12 | 0.632250926 | 0.442 | 0.265 | 2.02E-08 | Tb |
| cDC2 | IGSF6 | 9.06E-25 | 0.632150077 | 0.866 | 0.676 | 1.78E-20 | Tb |
| cDC2 | TCAIM | 1.34E-08 | 0.631625981 | 0.297 | 0.169 | 0.000264357 | Tb |
| cDC2 | UBE2W | 1.75E-12 | 0.631417436 | 0.677 | 0.513 | 3.45E-08 | Tb |
| cDC2 | SLAMF1 | 4.85E-08 | 0.631176648 | 0.295 | 0.18 | 0.000954399 | Tb |
| cDC2 | MAN1A2 | 3.11E-15 | 0.631125739 | 0.443 | 0.244 | 6.12E-11 | Tb |
| cDC2 | PSD3 | 3.69E-09 | 0.629829669 | 0.3 | 0.174 | 7.27E-05 | Tb |
| cDC2 | CHAF1A | 9.31E-15 | 0.62966513 | 0.426 | 0.238 | 1.83E-10 | Tb |
| cDC2 | OSTM1 | 2.12E-11 | 0.629541524 | 0.325 | 0.177 | 4.17E-07 | Tb |
| cDC2 | SRBD1 | 8.20E-10 | 0.629184367 | 0.311 | 0.177 | 1.61E-05 | Tb |
| cDC2 | LARP7 | 1.68E-15 | 0.629067254 | 0.473 | 0.262 | 3.30E-11 | Tb |
| cDC2 | R3HCC1L | 1.99E-11 | 0.62848658 | 0.354 | 0.199 | 3.91E-07 | Tb |
| cDC2 | ATP9B | 2.18E-10 | 0.628409574 | 0.572 | 0.387 | 4.30E-06 | Tb |
| cDC2 | LMNB1 | 1.34E-14 | 0.62768806 | 0.489 | 0.286 | 2.64E-10 | Tb |
| cDC2 | SMARCA4 | 3.07E-10 | 0.627679113 | 0.305 | 0.169 | 6.04E-06 | Tb |
| cDC2 | ANKRD13C | 1.66E-12 | 0.627399059 | 0.251 | 0.119 | 3.27E-08 | Tb |
| cDC2 | ATAD2B | 2.06E-15 | 0.627269104 | 0.644 | 0.414 | 4.06E-11 | Tb |
| cDC2 | AKAP13 | 2.52E-18 | 0.627191767 | 0.96 | 0.887 | 4.96E-14 | Tb |
| cDC2 | PLA2G4C | 1.66E-10 | 0.627151649 | 0.3 | 0.165 | 3.27E-06 | Tb |
| cDC2 | SPOP | 1.82E-13 | 0.627064779 | 0.475 | 0.28 | 3.58E-09 | Tb |
| cDC2 | GPATCH11 | 1.11E-09 | 0.626546787 | 0.297 | 0.162 | 2.18E-05 | Tb |
| cDC2 | TRAPPC8 | 6.95E-13 | 0.626352518 | 0.321 | 0.168 | 1.37E-08 | Tb |
| cDC2 | PPP6R2 | 8.06E-18 | 0.626148301 | 0.506 | 0.276 | 1.59E-13 | Tb |
| cDC2 | FOXP1 | 4.17E-11 | 0.626085592 | 0.77 | 0.621 | 8.20E-07 | Tb |
| cDC2 | ZBTB10 | 5.93E-09 | 0.625865342 | 0.586 | 0.434 | 0.000116779 | Tb |
| cDC2 | SLC30A9 | 7.75E-14 | 0.625744733 | 0.349 | 0.18 | 1.53E-09 | Tb |
| cDC2 | ZHX1 | 4.46E-10 | 0.625489886 | 0.295 | 0.16 | 8.77E-06 | Tb |
| cDC2 | SEPTIN6 | 7.82E-16 | 0.625474402 | 0.855 | 0.689 | 1.54E-11 | Tb |
| cDC2 | ANKRD22 | 8.13E-16 | 0.62534921 | 0.581 | 0.371 | 1.60E-11 | Tb |
| cDC2 | TXNIP | 3.73E-25 | 0.625092077 | 0.799 | 0.531 | 7.35E-21 | Tb |
| cDC2 | USP8 | 1.57E-17 | 0.624914541 | 0.555 | 0.316 | 3.08E-13 | Tb |
| cDC2 | LONP2 | 1.19E-14 | 0.624460561 | 0.464 | 0.26 | 2.34E-10 | Tb |
| cDC2 | TNFRSF21 | 1.16E-07 | 0.624430578 | 0.286 | 0.174 | 0.002291006 | Tb |
| cDC2 | SLC39A8 | 4.79E-10 | 0.624144321 | 0.506 | 0.344 | 9.43E-06 | Tb |
| cDC2 | AZI2 | 1.03E-06 | 0.623068749 | 0.267 | 0.163 | 0.020183337 | Tb |
| cDC2 | SLC7A111 | 2.34E-09 | 0.623044029 | 0.318 | 0.191 | 4.61E-05 | Tb |

| | | | | | | | |
|---|---|---|---|---|---|---|---|
| cDC2 | PLAA | 3.24E-14 | 0.622810336 | 0.36 | 0.187 | 6.38E-10 | Tb |
| cDC2 | IP6K1 | 6.10E-08 | 0.622746592 | 0.25 | 0.138 | 0.001200039 | Tb |
| cDC2 | ZBTB18 | 2.26E-08 | 0.622670151 | 0.25 | 0.137 | 0.000444466 | Tb |
| cDC2 | EED | 1.32E-10 | 0.622326285 | 0.394 | 0.233 | 2.59E-06 | Tb |
| cDC2 | CRLS1 | 2.68E-10 | 0.621119651 | 0.269 | 0.139 | 5.27E-06 | Tb |
| cDC2 | ZNF592 | 5.55E-08 | 0.620842335 | 0.255 | 0.147 | 0.001093437 | Tb |
| cDC2 | PRKAR2A | 1.46E-11 | 0.620826555 | 0.48 | 0.301 | 2.87E-07 | Tb |
| cDC2 | WDFY2 | 6.88E-14 | 0.620518095 | 0.602 | 0.409 | 1.35E-09 | Tb |
| cDC2 | PREP | 1.84E-08 | 0.620474178 | 0.265 | 0.149 | 0.000362269 | Tb |
| cDC2 | ST3GAL1 | 2.15E-16 | 0.62007581 | 0.501 | 0.288 | 4.22E-12 | Tb |
| cDC2 | TMA16 | 2.40E-10 | 0.619668162 | 0.3 | 0.162 | 4.72E-06 | Tb |
| cDC2 | CELF2 | 4.57E-21 | 0.619641355 | 0.855 | 0.64 | 9.00E-17 | Tb |
| cDC2 | KPNA6 | 6.64E-10 | 0.619249252 | 0.277 | 0.15 | 1.31E-05 | Tb |
| cDC2 | BICD2 | 2.82E-13 | 0.619185259 | 0.38 | 0.202 | 5.55E-09 | Tb |
| cDC2 | TBC1D9 | 3.78E-11 | 0.619087369 | 0.513 | 0.327 | 7.44E-07 | Tb |
| cDC2 | OTUD4 | 1.20E-12 | 0.618798552 | 0.504 | 0.314 | 2.37E-08 | Tb |
| cDC2 | TIMMDC1 | 3.05E-11 | 0.618486198 | 0.494 | 0.311 | 6.01E-07 | Tb |
| cDC2 | ADCY7 | 7.78E-16 | 0.618102793 | 0.614 | 0.379 | 1.53E-11 | Tb |
| cDC2 | SEC23IP | 3.19E-12 | 0.617330766 | 0.297 | 0.153 | 6.28E-08 | Tb |
| cDC2 | PIBF1 | 4.31E-10 | 0.61654884 | 0.265 | 0.141 | 8.49E-06 | Tb |
| cDC2 | LIMD1 | 3.81E-13 | 0.616273316 | 0.351 | 0.187 | 7.51E-09 | Tb |
| cDC2 | WDR7 | 1.10E-10 | 0.616227887 | 0.37 | 0.219 | 2.16E-06 | Tb |
| cDC2 | ACOX1 | 3.76E-11 | 0.615875662 | 0.29 | 0.153 | 7.40E-07 | Tb |
| cDC2 | PMF1 | 2.89E-12 | 0.615770451 | 0.457 | 0.269 | 5.69E-08 | Tb |
| cDC2 | LMO4 | 2.32E-16 | 0.615200842 | 0.627 | 0.396 | 4.57E-12 | Tb |
| cDC2 | RARS2 | 7.69E-11 | 0.614608665 | 0.305 | 0.166 | 1.51E-06 | Tb |
| cDC2 | AMBRA1 | 6.80E-17 | 0.614155848 | 0.538 | 0.306 | 1.34E-12 | Tb |
| cDC2 | BAZ2B | 2.96E-08 | 0.614035067 | 0.536 | 0.376 | 0.000583052 | Tb |
| cDC2 | KLHL18 | 8.92E-09 | 0.613923512 | 0.318 | 0.186 | 0.000175635 | Tb |
| cDC2 | ZC3H4 | 2.14E-17 | 0.613816189 | 0.316 | 0.139 | 4.22E-13 | Tb |
| cDC2 | CHML | 3.51E-11 | 0.613808822 | 0.379 | 0.22 | 6.91E-07 | Tb |
| cDC2 | B3GALNT2 | 3.72E-10 | 0.613533673 | 0.26 | 0.135 | 7.31E-06 | Tb |
| cDC2 | APBB1IP | 2.50E-13 | 0.613164355 | 0.723 | 0.511 | 4.91E-09 | Tb |
| cDC2 | SIN3A | 1.17E-14 | 0.613112968 | 0.339 | 0.167 | 2.30E-10 | Tb |
| cDC2 | FNDC3B | 1.43E-21 | 0.61299661 | 0.812 | 0.572 | 2.81E-17 | Tb |
| cDC2 | PARN | 3.06E-11 | 0.61283795 | 0.312 | 0.169 | 6.02E-07 | Tb |
| cDC2 | GAPVD1 | 1.78E-12 | 0.612557906 | 0.435 | 0.245 | 3.50E-08 | Tb |
| cDC2 | PSMD14 | 1.16E-14 | 0.612504484 | 0.634 | 0.392 | 2.29E-10 | Tb |
| cDC2 | PPP4R1 | 1.66E-06 | 0.612488442 | 0.25 | 0.153 | 0.032673079 | Tb |
| cDC2 | SIL1 | 4.22E-14 | 0.612452567 | 0.515 | 0.314 | 8.30E-10 | Tb |
| cDC2 | ACTN1 | 1.77E-09 | 0.612137447 | 0.558 | 0.399 | 3.49E-05 | Tb |
| cDC2 | ABRAXAS2 | 2.33E-10 | 0.611868323 | 0.262 | 0.137 | 4.59E-06 | Tb |
| cDC2 | TMCC1 | 1.58E-11 | 0.61184109 | 0.335 | 0.184 | 3.11E-07 | Tb |
| cDC2 | PCNA | 3.85E-08 | 0.611820559 | 0.272 | 0.157 | 0.000757405 | Tb |
| cDC2 | NOC3L | 7.59E-11 | 0.611797863 | 0.255 | 0.129 | 1.49E-06 | Tb |
| cDC2 | BRD1 | 4.33E-09 | 0.611248145 | 0.298 | 0.17 | 8.53E-05 | Tb |
| cDC2 | LYPLAL1 | 5.45E-10 | 0.61107776 | 0.314 | 0.179 | 1.07E-05 | Tb |
| cDC2 | ASB3 | 4.60E-08 | 0.610666082 | 0.347 | 0.214 | 0.000904567 | Tb |
| cDC2 | CUL3 | 4.01E-17 | 0.610169498 | 0.628 | 0.382 | 7.89E-13 | Tb |
| cDC2 | NCOA3 | 7.01E-13 | 0.610162523 | 0.689 | 0.49 | 1.38E-08 | Tb |
| cDC2 | PPP4R3A | 8.75E-15 | 0.609558945 | 0.754 | 0.516 | 1.72E-10 | Tb |
| cDC2 | SGK3 | 3.32E-17 | 0.609101352 | 0.616 | 0.377 | 6.54E-13 | Tb |
| cDC2 | TMEM33 | 4.24E-16 | 0.609041234 | 0.36 | 0.172 | 8.34E-12 | Tb |
| cDC2 | SMIM15 | 4.41E-10 | 0.608556906 | 0.349 | 0.199 | 8.68E-06 | Tb |
| cDC2 | TNFSF10 | 3.01E-18 | 0.608239357 | 0.763 | 0.521 | 5.92E-14 | Tb |
| cDC2 | GNL1 | 1.18E-08 | 0.608082821 | 0.332 | 0.197 | 0.000231902 | Tb |

| | | | | | | | |
|---|---|---|---|---|---|---|---|
| cDC2 | TMEM87A | 2.07E-13 | 0.607899381 | 0.466 | 0.265 | 4.07E-09 | Tb |
| cDC2 | CEMIP2 | 2.34E-08 | 0.607549718 | 0.4 | 0.261 | 0.000461103 | Tb |
| cDC2 | SUSD1 | 1.19E-07 | 0.607379618 | 0.332 | 0.21 | 0.00234825 | Tb |
| cDC2 | CCPG1 | 1.24E-10 | 0.607292491 | 0.356 | 0.2 | 2.44E-06 | Tb |
| cDC2 | PDXK | 5.47E-21 | 0.607088287 | 0.775 | 0.542 | 1.08E-16 | Tb |
| cDC2 | BIRC6 | 3.23E-17 | 0.606498445 | 0.74 | 0.501 | 6.36E-13 | Tb |
| cDC2 | USP22 | 3.61E-20 | 0.606229253 | 0.525 | 0.281 | 7.10E-16 | Tb |
| cDC2 | HERC1 | 6.70E-11 | 0.606224348 | 0.621 | 0.427 | 1.32E-06 | Tb |
| cDC2 | RPA2 | 4.07E-12 | 0.606054834 | 0.417 | 0.238 | 8.01E-08 | Tb |
| cDC2 | PPP3CA | 1.39E-17 | 0.605936956 | 0.628 | 0.406 | 2.73E-13 | Tb |
| cDC2 | FAM168B | 2.07E-10 | 0.605838966 | 0.333 | 0.187 | 4.08E-06 | Tb |
| cDC2 | VPS53 | 1.52E-18 | 0.60487089 | 0.445 | 0.228 | 3.00E-14 | Tb |
| cDC2 | STX18 | 6.37E-09 | 0.604864042 | 0.382 | 0.235 | 0.000125362 | Tb |
| cDC2 | FAM13B | 2.64E-10 | 0.604807959 | 0.487 | 0.314 | 5.20E-06 | Tb |
| cDC2 | PPP4R3B | 3.07E-12 | 0.604601136 | 0.459 | 0.273 | 6.04E-08 | Tb |
| cDC2 | TPK1 | 1.38E-10 | 0.604481455 | 0.438 | 0.267 | 2.73E-06 | Tb |
| cDC2 | STX12 | 2.40E-12 | 0.604417158 | 0.653 | 0.455 | 4.72E-08 | Tb |
| cDC2 | RBM6 | 3.36E-17 | 0.604212958 | 0.592 | 0.362 | 6.62E-13 | Tb |
| cDC2 | GPR65 | 3.04E-08 | 0.604025455 | 0.496 | 0.346 | 0.000598342 | Tb |
| cDC2 | FAM49B | 9.13E-30 | 0.603927221 | 0.939 | 0.776 | 1.80E-25 | Tb |
| cDC2 | LONRF1 | 8.55E-16 | 0.603845132 | 0.462 | 0.256 | 1.68E-11 | Tb |
| cDC2 | DNM1L | 3.67E-13 | 0.603723152 | 0.525 | 0.317 | 7.23E-09 | Tb |
| cDC2 | NARS | 2.21E-12 | 0.603680711 | 0.511 | 0.316 | 4.34E-08 | Tb |
| cDC2 | PHC3 | 4.43E-09 | 0.6036576 | 0.321 | 0.191 | 8.72E-05 | Tb |
| cDC2 | PHF14 | 7.06E-13 | 0.603277638 | 0.412 | 0.235 | 1.39E-08 | Tb |
| cDC2 | PHF21A | 1.48E-07 | 0.602864105 | 0.382 | 0.25 | 0.002915971 | Tb |
| cDC2 | RAB4B | 8.65E-14 | 0.601918214 | 0.476 | 0.279 | 1.70E-09 | Tb |
| cDC2 | DIAPH1 | 4.47E-13 | 0.601901195 | 0.686 | 0.471 | 8.79E-09 | Tb |
| cDC2 | NDUFA8 | 2.63E-12 | 0.601861079 | 0.452 | 0.264 | 5.17E-08 | Tb |
| cDC2 | AHCTF1 | 1.14E-09 | 0.601431769 | 0.379 | 0.226 | 2.25E-05 | Tb |
| cDC2 | NEMF | 6.47E-15 | 0.601312544 | 0.557 | 0.34 | 1.27E-10 | Tb |
| cDC2 | PSMD1 | 4.81E-17 | 0.60123634 | 0.611 | 0.361 | 9.46E-13 | Tb |
| cDC2 | RIOK1 | 5.58E-12 | 0.60110102 | 0.373 | 0.208 | 1.10E-07 | Tb |
| cDC2 | MRM2 | 2.36E-13 | 0.6010746 | 0.443 | 0.252 | 4.64E-09 | Tb |
| cDC2 | POLR3K | 6.54E-12 | 0.601025111 | 0.314 | 0.165 | 1.29E-07 | Tb |
| cDC2 | CRK | 3.14E-13 | 0.600996576 | 0.452 | 0.262 | 6.17E-09 | Tb |
| cDC2 | KARS | 4.30E-13 | 0.600914748 | 0.455 | 0.261 | 8.46E-09 | Tb |
| cDC2 | RAB5B | 1.86E-08 | 0.600786665 | 0.291 | 0.169 | 0.000366764 | Tb |
| cDC2 | FARS2 | 1.49E-15 | 0.60043741 | 0.373 | 0.191 | 2.93E-11 | Tb |
| cDC2 | VPS54 | 1.14E-12 | 0.600216502 | 0.274 | 0.135 | 2.25E-08 | Tb |
| cDC2 | NUB1 | 5.25E-16 | 0.599908916 | 0.736 | 0.528 | 1.03E-11 | Tb |
| cDC2 | NUS1 | 8.43E-11 | 0.599891808 | 0.3 | 0.162 | 1.66E-06 | Tb |
| cDC2 | DCTPP1 | 8.25E-18 | 0.599816616 | 0.501 | 0.269 | 1.62E-13 | Tb |
| cDC2 | DOCK10 | 5.48E-14 | 0.59914202 | 0.778 | 0.607 | 1.08E-09 | Tb |
| cDC2 | BCL2L13 | 1.14E-11 | 0.599033254 | 0.468 | 0.292 | 2.25E-07 | Tb |
| cDC2 | TAPT1 | 7.69E-12 | 0.598446096 | 0.344 | 0.192 | 1.51E-07 | Tb |
| cDC2 | PCF11 | 7.59E-12 | 0.598437502 | 0.494 | 0.3 | 1.49E-07 | Tb |
| cDC2 | TRIP11 | 2.82E-16 | 0.598413142 | 0.38 | 0.191 | 5.54E-12 | Tb |
| cDC2 | LARP4B | 4.92E-20 | 0.598349292 | 0.618 | 0.369 | 9.69E-16 | Tb |
| cDC2 | RNF34 | 3.19E-13 | 0.598120392 | 0.333 | 0.17 | 6.29E-09 | Tb |
| cDC2 | SPATS2 | 1.30E-15 | 0.597623313 | 0.513 | 0.3 | 2.56E-11 | Tb |
| cDC2 | CTDSP2 | 1.48E-12 | 0.597339873 | 0.368 | 0.198 | 2.91E-08 | Tb |
| cDC2 | TSG101 | 1.15E-11 | 0.597209589 | 0.501 | 0.304 | 2.26E-07 | Tb |
| cDC2 | MAP4 | 1.26E-12 | 0.596968596 | 0.592 | 0.387 | 2.47E-08 | Tb |
| cDC2 | HIVEP1 | 1.68E-08 | 0.596456514 | 0.681 | 0.549 | 0.000330864 | Tb |
| cDC2 | AP2B1 | 8.44E-15 | 0.596292404 | 0.506 | 0.29 | 1.66E-10 | Tb |

| | | | | | | | |
|---|---|---|---|---|---|---|---|
| cDC2 | FBRSL1 | 1.97E-15 | 0.596073654 | 0.442 | 0.234 | 3.88E-11 | Tb |
| cDC2 | EP400 | 3.72E-11 | 0.595624493 | 0.283 | 0.146 | 7.33E-07 | Tb |
| cDC2 | AGTPBP1 | 1.46E-10 | 0.595450619 | 0.553 | 0.382 | 2.88E-06 | Tb |
| cDC2 | HMGB2 | 4.08E-12 | 0.595222652 | 0.604 | 0.395 | 8.02E-08 | Tb |
| cDC2 | AP1G1 | 3.53E-19 | 0.595072413 | 0.539 | 0.291 | 6.95E-15 | Tb |
| cDC2 | NUTM2B-AS1 | 8.36E-12 | 0.595010653 | 0.48 | 0.295 | 1.65E-07 | Tb |
| cDC2 | NCOA1 | 3.42E-15 | 0.59399007 | 0.588 | 0.362 | 6.73E-11 | Tb |
| cDC2 | HERC2 | 1.52E-11 | 0.593296569 | 0.358 | 0.197 | 2.99E-07 | Tb |
| cDC2 | TTC37 | 1.88E-09 | 0.592970331 | 0.339 | 0.197 | 3.71E-05 | Tb |
| cDC2 | RANBP2 | 1.63E-10 | 0.592866995 | 0.766 | 0.593 | 3.20E-06 | Tb |
| cDC2 | ANKHD1 | 2.58E-18 | 0.592829517 | 0.808 | 0.589 | 5.08E-14 | Tb |
| cDC2 | WDR43 | 1.32E-12 | 0.592734601 | 0.396 | 0.218 | 2.59E-08 | Tb |
| cDC2 | TNRC6B | 2.14E-13 | 0.592005195 | 0.602 | 0.402 | 4.21E-09 | Tb |
| cDC2 | TRAFD1 | 3.56E-10 | 0.591551721 | 0.312 | 0.175 | 7.00E-06 | Tb |
| cDC2 | DCTN4 | 5.34E-13 | 0.591188439 | 0.494 | 0.296 | 1.05E-08 | Tb |
| cDC2 | ODR4 | 4.10E-11 | 0.59081161 | 0.305 | 0.164 | 8.06E-07 | Tb |
| cDC2 | TRIM69 | 2.48E-14 | 0.590127341 | 0.63 | 0.432 | 4.88E-10 | Tb |
| cDC2 | ZNF331 | 1.72E-10 | 0.58950688 | 0.796 | 0.697 | 3.38E-06 | Tb |
| cDC2 | WDFY1 | 3.00E-13 | 0.589332518 | 0.366 | 0.199 | 5.91E-09 | Tb |
| cDC2 | TAPBP | 2.71E-25 | 0.588834282 | 0.876 | 0.631 | 5.34E-21 | Tb |
| cDC2 | ZBTB1 | 4.24E-13 | 0.588667454 | 0.656 | 0.444 | 8.35E-09 | Tb |
| cDC2 | IL4R | 5.31E-14 | 0.588267365 | 0.49 | 0.291 | 1.05E-09 | Tb |
| cDC2 | NBAS | 3.68E-11 | 0.587710195 | 0.319 | 0.173 | 7.24E-07 | Tb |
| cDC2 | EIF2B3 | 2.27E-09 | 0.587367291 | 0.283 | 0.157 | 4.47E-05 | Tb |
| cDC2 | AFG3L2 | 1.26E-11 | 0.587255846 | 0.368 | 0.203 | 2.48E-07 | Tb |
| cDC2 | APOBEC3G | 4.72E-11 | 0.587206715 | 0.354 | 0.198 | 9.29E-07 | Tb |
| cDC2 | ADCY3 | 1.17E-11 | 0.58690425 | 0.318 | 0.172 | 2.30E-07 | Tb |
| cDC2 | OPA1 | 1.61E-13 | 0.586736751 | 0.45 | 0.256 | 3.16E-09 | Tb |
| cDC2 | TBL1XR1 | 2.86E-15 | 0.586676837 | 0.623 | 0.391 | 5.62E-11 | Tb |
| cDC2 | GOLGA5 | 2.00E-10 | 0.58589521 | 0.253 | 0.129 | 3.94E-06 | Tb |
| cDC2 | ERCC5 | 1.20E-09 | 0.585039558 | 0.325 | 0.186 | 2.36E-05 | Tb |
| cDC2 | OSGIN2 | 3.51E-14 | 0.584924 | 0.326 | 0.161 | 6.90E-10 | Tb |
| cDC2 | DCAF7 | 2.03E-13 | 0.584814839 | 0.503 | 0.305 | 4.00E-09 | Tb |
| cDC2 | EBAG9 | 1.38E-11 | 0.584683654 | 0.293 | 0.149 | 2.72E-07 | Tb |
| cDC2 | RAPGEF6 | 3.81E-14 | 0.584426953 | 0.632 | 0.406 | 7.49E-10 | Tb |
| cDC2 | LRPPRC | 1.08E-14 | 0.583644664 | 0.454 | 0.248 | 2.13E-10 | Tb |
| cDC2 | PPHLN1 | 1.47E-16 | 0.583540583 | 0.599 | 0.355 | 2.90E-12 | Tb |
| cDC2 | CD274 | 1.23E-11 | 0.583269922 | 0.415 | 0.256 | 2.43E-07 | Tb |
| cDC2 | TSC22D2 | 1.57E-11 | 0.583262854 | 0.675 | 0.484 | 3.09E-07 | Tb |
| cDC2 | ZFAND3 | 2.13E-24 | 0.583202273 | 0.876 | 0.686 | 4.19E-20 | Tb |
| cDC2 | CEP63 | 1.72E-07 | 0.582983948 | 0.318 | 0.197 | 0.003384961 | Tb |
| cDC2 | DNAJA4 | 3.05E-14 | 0.582747612 | 0.609 | 0.395 | 6.00E-10 | Tb |
| cDC2 | ISOC2 | 1.39E-13 | 0.582237296 | 0.524 | 0.34 | 2.74E-09 | Tb |
| cDC2 | ADK | 2.55E-12 | 0.581956267 | 0.59 | 0.414 | 5.02E-08 | Tb |
| cDC2 | ICE2 | 1.17E-14 | 0.581914587 | 0.318 | 0.154 | 2.31E-10 | Tb |
| cDC2 | KIZ-AS1 | 7.94E-08 | 0.581677384 | 0.267 | 0.157 | 0.001562666 | Tb |
| cDC2 | FNDC3A | 1.92E-14 | 0.581629584 | 0.691 | 0.498 | 3.77E-10 | Tb |
| cDC2 | WNK1 | 5.25E-16 | 0.581495926 | 0.716 | 0.489 | 1.03E-11 | Tb |
| cDC2 | SLC66A2 | 6.82E-14 | 0.581285474 | 0.688 | 0.49 | 1.34E-09 | Tb |
| cDC2 | EEFSEC | 3.68E-14 | 0.581196237 | 0.264 | 0.119 | 7.25E-10 | Tb |
| cDC2 | PIEZO1 | 3.85E-09 | 0.581053898 | 0.29 | 0.159 | 7.58E-05 | Tb |
| cDC2 | CXCL9 | 1.78E-14 | 0.581002409 | 0.541 | 0.355 | 3.50E-10 | Tb |
| cDC2 | DTNA | 1.14E-06 | 0.580950305 | 0.312 | 0.208 | 0.0223473 | Tb |
| cDC2 | PRDX2 | 2.23E-12 | 0.580551698 | 0.464 | 0.282 | 4.39E-08 | Tb |
| cDC2 | PACSIN2 | 1.49E-21 | 0.580248177 | 0.792 | 0.523 | 2.93E-17 | Tb |
| cDC2 | PDE8A | 3.47E-10 | 0.580153491 | 0.377 | 0.23 | 6.83E-06 | Tb |

| | | | | | | | |
|---|---|---|---|---|---|---|---|
| cDC2 | CCSER1 | 2.32E-09 | 0.579573511 | 0.831 | 0.722 | 4.56E-05 | Tb |
| cDC2 | ADNP | 2.63E-11 | 0.579342672 | 0.455 | 0.283 | 5.17E-07 | Tb |
| cDC2 | FOXK1 | 1.19E-10 | 0.579222979 | 0.328 | 0.182 | 2.34E-06 | Tb |
| cDC2 | IMMT | 2.12E-14 | 0.579159198 | 0.494 | 0.289 | 4.17E-10 | Tb |
| cDC2 | MGME1 | 3.31E-09 | 0.578048253 | 0.316 | 0.182 | 6.52E-05 | Tb |
| cDC2 | GART | 4.85E-10 | 0.578046947 | 0.26 | 0.137 | 9.54E-06 | Tb |
| cDC2 | GNPTAB | 9.14E-15 | 0.578007216 | 0.49 | 0.282 | 1.80E-10 | Tb |
| cDC2 | OTUD6B-AS1 | 4.57E-09 | 0.577967958 | 0.356 | 0.209 | 9.00E-05 | Tb |
| cDC2 | RCN2 | 2.60E-09 | 0.577951898 | 0.325 | 0.183 | 5.12E-05 | Tb |
| cDC2 | BACH1 | 1.65E-15 | 0.577815262 | 0.818 | 0.648 | 3.26E-11 | Tb |
| cDC2 | KIAA0232 | 1.39E-10 | 0.577675486 | 0.342 | 0.194 | 2.74E-06 | Tb |
| cDC2 | NADK | 6.07E-14 | 0.577609058 | 0.325 | 0.161 | 1.20E-09 | Tb |
| cDC2 | TRAPPC12 | 7.68E-15 | 0.576431817 | 0.363 | 0.182 | 1.51E-10 | Tb |
| cDC2 | RBPJ | 8.88E-16 | 0.576323871 | 0.855 | 0.713 | 1.75E-11 | Tb |
| cDC2 | BAZ1A | 1.91E-18 | 0.57591759 | 0.934 | 0.789 | 3.76E-14 | Tb |
| cDC2 | PRKAG1 | 1.49E-12 | 0.575894928 | 0.401 | 0.226 | 2.93E-08 | Tb |
| cDC2 | ZRSR2 | 3.92E-10 | 0.575623319 | 0.298 | 0.165 | 7.71E-06 | Tb |
| cDC2 | SUN2 | 1.07E-15 | 0.575602538 | 0.386 | 0.197 | 2.10E-11 | Tb |
| cDC2 | NSUN6 | 1.60E-17 | 0.575286489 | 0.583 | 0.354 | 3.15E-13 | Tb |
| cDC2 | SLC39A10 | 9.26E-13 | 0.574479211 | 0.482 | 0.294 | 1.82E-08 | Tb |
| cDC2 | NBR1 | 1.18E-12 | 0.573699539 | 0.295 | 0.146 | 2.33E-08 | Tb |
| cDC2 | HAGH | 4.02E-12 | 0.572499529 | 0.464 | 0.276 | 7.90E-08 | Tb |
| cDC2 | SENP6 | 1.06E-11 | 0.572125329 | 0.634 | 0.431 | 2.08E-07 | Tb |
| cDC2 | HDAC8 | 1.42E-08 | 0.572114451 | 0.302 | 0.18 | 0.000278746 | Tb |
| cDC2 | TAF1 | 4.35E-10 | 0.571831376 | 0.279 | 0.151 | 8.57E-06 | Tb |
| cDC2 | GPR160 | 9.02E-09 | 0.571611877 | 0.276 | 0.157 | 0.000177518 | Tb |
| cDC2 | VEZT | 3.19E-13 | 0.571002276 | 0.414 | 0.233 | 6.27E-09 | Tb |
| cDC2 | GOLIM4 | 5.98E-13 | 0.570834192 | 0.457 | 0.263 | 1.18E-08 | Tb |
| cDC2 | STAG1 | 1.71E-18 | 0.570820418 | 0.649 | 0.397 | 3.36E-14 | Tb |
| cDC2 | MASTL | 8.94E-12 | 0.570579948 | 0.466 | 0.287 | 1.76E-07 | Tb |
| cDC2 | TIMM10B | 1.12E-10 | 0.570454774 | 0.321 | 0.178 | 2.21E-06 | Tb |
| cDC2 | PBX3 | 1.91E-11 | 0.569871555 | 0.565 | 0.385 | 3.76E-07 | Tb |
| cDC2 | RIC1 | 1.36E-08 | 0.569749996 | 0.55 | 0.384 | 0.000267617 | Tb |
| cDC2 | HIP1 | 8.97E-10 | 0.569578124 | 0.606 | 0.452 | 1.77E-05 | Tb |
| cDC2 | EDEM3 | 1.30E-09 | 0.568929113 | 0.314 | 0.178 | 2.55E-05 | Tb |
| cDC2 | ILRUN | 1.92E-12 | 0.568736231 | 0.569 | 0.373 | 3.77E-08 | Tb |
| cDC2 | VCPIP1 | 1.84E-11 | 0.568525278 | 0.328 | 0.179 | 3.61E-07 | Tb |
| cDC2 | UBE2Q1 | 1.47E-11 | 0.568378374 | 0.351 | 0.194 | 2.89E-07 | Tb |
| cDC2 | CCDC47 | 1.24E-09 | 0.568308298 | 0.656 | 0.449 | 2.44E-05 | Tb |
| cDC2 | PSMD7 | 2.21E-17 | 0.568242157 | 0.724 | 0.493 | 4.35E-13 | Tb |
| cDC2 | XPO6 | 2.70E-11 | 0.568215057 | 0.366 | 0.206 | 5.32E-07 | Tb |
| cDC2 | YWHAH | 6.73E-18 | 0.567862678 | 0.791 | 0.57 | 1.32E-13 | Tb |
| cDC2 | TIAM1 | 4.43E-12 | 0.567544399 | 0.541 | 0.351 | 8.71E-08 | Tb |
| cDC2 | SNAPC5 | 1.02E-14 | 0.567170312 | 0.3 | 0.139 | 2.02E-10 | Tb |
| cDC2 | WDR82 | 1.31E-12 | 0.567059247 | 0.421 | 0.24 | 2.58E-08 | Tb |
| cDC2 | CGAS | 2.84E-14 | 0.566934871 | 0.571 | 0.354 | 5.58E-10 | Tb |
| cDC2 | SRFBP1 | 2.23E-10 | 0.566774639 | 0.401 | 0.237 | 4.39E-06 | Tb |
| cDC2 | PIK3C3 | 4.11E-11 | 0.566098161 | 0.3 | 0.16 | 8.09E-07 | Tb |
| cDC2 | RBM28 | 4.30E-08 | 0.566091898 | 0.265 | 0.153 | 0.000847359 | Tb |
| cDC2 | CELF11 | 1.35E-11 | 0.56584693 | 0.529 | 0.335 | 2.67E-07 | Tb |
| cDC2 | TDG | 1.44E-07 | 0.565749197 | 0.49 | 0.329 | 0.002828282 | Tb |
| cDC2 | IPMK | 2.06E-08 | 0.565607639 | 0.454 | 0.306 | 0.000405529 | Tb |
| cDC2 | LRRC41 | 2.20E-11 | 0.565141667 | 0.325 | 0.175 | 4.33E-07 | Tb |
| cDC2 | UIMC1 | 2.42E-10 | 0.565027823 | 0.337 | 0.188 | 4.76E-06 | Tb |
| cDC2 | ERC1 | 3.99E-20 | 0.564958296 | 0.682 | 0.428 | 7.86E-16 | Tb |
| cDC2 | ZMIZ1 | 1.59E-15 | 0.564647845 | 0.707 | 0.49 | 3.12E-11 | Tb |

| | | | | | | | |
|---|---|---|---|---|---|---|---|
| cDC2 | GOSR1 | 7.31E-11 | 0.564272732 | 0.44 | 0.267 | 1.44E-06 | Tb |
| cDC2 | SRP19 | 7.37E-18 | 0.563726723 | 0.714 | 0.456 | 1.45E-13 | Tb |
| cDC2 | MAPKAP1 | 2.49E-12 | 0.563306795 | 0.565 | 0.362 | 4.90E-08 | Tb |
| cDC2 | MAD1L1 | 1.81E-11 | 0.562828228 | 0.675 | 0.459 | 3.56E-07 | Tb |
| cDC2 | MRRF | 3.16E-10 | 0.56281135 | 0.319 | 0.178 | 6.22E-06 | Tb |
| cDC2 | DYRK1A | 1.90E-18 | 0.562259932 | 0.634 | 0.385 | 3.74E-14 | Tb |
| cDC2 | CCZ1 | 3.63E-12 | 0.562177654 | 0.328 | 0.175 | 7.14E-08 | Tb |
| cDC2 | NUP107 | 6.60E-09 | 0.561892539 | 0.276 | 0.154 | 0.000129937 | Tb |
| cDC2 | CNDP2 | 6.29E-20 | 0.561646885 | 0.745 | 0.487 | 1.24E-15 | Tb |
| cDC2 | SMC1A | 2.09E-10 | 0.561524187 | 0.457 | 0.279 | 4.10E-06 | Tb |
| cDC2 | ITGB1BP1 | 1.45E-13 | 0.561418858 | 0.524 | 0.309 | 2.86E-09 | Tb |
| cDC2 | PER1 | 2.73E-11 | 0.560995617 | 0.586 | 0.39 | 5.37E-07 | Tb |
| cDC2 | CCNG2 | 3.94E-10 | 0.560766949 | 0.387 | 0.239 | 7.75E-06 | Tb |
| cDC2 | PHF20L1 | 1.95E-10 | 0.560417845 | 0.551 | 0.37 | 3.84E-06 | Tb |
| cDC2 | SCFD1 | 2.27E-12 | 0.56036638 | 0.536 | 0.333 | 4.47E-08 | Tb |
| cDC2 | NEK9 | 1.79E-08 | 0.560247247 | 0.262 | 0.148 | 0.00035261 | Tb |
| cDC2 | OSBPL8 | 4.55E-15 | 0.560161164 | 0.852 | 0.71 | 8.96E-11 | Tb |
| cDC2 | ATP5MPL | 9.26E-24 | 0.560150908 | 0.836 | 0.597 | 1.82E-19 | Tb |
| cDC2 | PLEKHA2 | 1.36E-10 | 0.56013993 | 0.496 | 0.309 | 2.68E-06 | Tb |
| cDC2 | ARFIP1 | 2.11E-10 | 0.559984754 | 0.293 | 0.157 | 4.16E-06 | Tb |
| cDC2 | API5 | 3.02E-11 | 0.559757983 | 0.398 | 0.23 | 5.94E-07 | Tb |
| cDC2 | DLST | 2.94E-10 | 0.559345519 | 0.366 | 0.206 | 5.79E-06 | Tb |
| cDC2 | CARS2 | 1.47E-13 | 0.558897973 | 0.311 | 0.151 | 2.89E-09 | Tb |
| cDC2 | SLC1A5 | 1.25E-09 | 0.558824774 | 0.522 | 0.333 | 2.46E-05 | Tb |
| cDC2 | TAF2 | 7.51E-08 | 0.558362754 | 0.272 | 0.159 | 0.001478141 | Tb |
| cDC2 | NDUFS2 | 9.17E-15 | 0.55818389 | 0.63 | 0.409 | 1.81E-10 | Tb |
| cDC2 | CLASRP | 2.58E-12 | 0.557761197 | 0.462 | 0.268 | 5.08E-08 | Tb |
| cDC2 | EMC2 | 8.31E-10 | 0.557690557 | 0.323 | 0.183 | 1.64E-05 | Tb |
| cDC2 | LZIC | 9.19E-14 | 0.557459533 | 0.415 | 0.222 | 1.81E-09 | Tb |
| cDC2 | PARP8 | 8.72E-11 | 0.557421249 | 0.487 | 0.32 | 1.72E-06 | Tb |
| cDC2 | PDE4A | 1.58E-06 | 0.557149756 | 0.754 | 0.669 | 0.031107776 | Tb |
| cDC2 | EAF2 | 2.51E-08 | 0.557059472 | 0.494 | 0.33 | 0.000494038 | Tb |
| cDC2 | CPQ | 3.40E-07 | 0.556781024 | 0.557 | 0.406 | 0.006686809 | Tb |
| cDC2 | LIX1L | 1.87E-11 | 0.556585988 | 0.291 | 0.149 | 3.68E-07 | Tb |
| cDC2 | SAMD1 | 4.60E-11 | 0.55631894 | 0.272 | 0.136 | 9.06E-07 | Tb |
| cDC2 | FNIP1 | 1.40E-19 | 0.556308626 | 0.717 | 0.452 | 2.75E-15 | Tb |
| cDC2 | DDB1 | 1.34E-10 | 0.55600171 | 0.346 | 0.194 | 2.63E-06 | Tb |
| cDC2 | MRPL13 | 5.70E-12 | 0.555864165 | 0.496 | 0.294 | 1.12E-07 | Tb |
| cDC2 | ECHS1 | 2.26E-12 | 0.555750421 | 0.686 | 0.494 | 4.46E-08 | Tb |
| cDC2 | ADAR | 6.45E-10 | 0.555486761 | 0.562 | 0.368 | 1.27E-05 | Tb |
| cDC2 | ZC3H18 | 2.73E-09 | 0.555020592 | 0.405 | 0.246 | 5.37E-05 | Tb |
| cDC2 | AP2A2 | 9.71E-12 | 0.554896863 | 0.29 | 0.149 | 1.91E-07 | Tb |
| cDC2 | COG5 | 1.41E-14 | 0.55473817 | 0.532 | 0.313 | 2.77E-10 | Tb |
| cDC2 | UBA2 | 3.10E-16 | 0.554736187 | 0.524 | 0.295 | 6.10E-12 | Tb |
| cDC2 | CSF2RA | 3.10E-15 | 0.554723233 | 0.866 | 0.723 | 6.10E-11 | Tb |
| cDC2 | MEF2A | 3.03E-13 | 0.554508996 | 0.839 | 0.672 | 5.97E-09 | Tb |
| cDC2 | ZNF791 | 1.70E-09 | 0.554473886 | 0.323 | 0.188 | 3.36E-05 | Tb |
| cDC2 | DYNC1I2 | 4.17E-14 | 0.554422138 | 0.497 | 0.29 | 8.21E-10 | Tb |
| cDC2 | BMS1 | 1.33E-11 | 0.554398346 | 0.304 | 0.158 | 2.62E-07 | Tb |
| cDC2 | BZW2 | 3.07E-08 | 0.554341009 | 0.609 | 0.43 | 0.000604167 | Tb |
| cDC2 | ADAMTSL4 | 4.33E-12 | 0.554046032 | 0.281 | 0.142 | 8.52E-08 | Tb |
| cDC2 | ECPAS | 5.77E-15 | 0.554045323 | 0.478 | 0.269 | 1.14E-10 | Tb |
| cDC2 | UBTD2 | 3.07E-09 | 0.553859136 | 0.435 | 0.281 | 6.04E-05 | Tb |
| cDC2 | PALM2-AKAP2 | 1.87E-12 | 0.553810658 | 0.647 | 0.462 | 3.69E-08 | Tb |
| cDC2 | ARFGAP2 | 5.94E-10 | 0.553676114 | 0.328 | 0.184 | 1.17E-05 | Tb |
| cDC2 | STIMATE | 9.88E-10 | 0.553394244 | 0.25 | 0.129 | 1.95E-05 | Tb |

| | | | | | | | |
|---|---|---|---|---|---|---|---|
| cDC2 | THBD | 3.49E-11 | 0.553235491 | 0.431 | 0.265 | 6.87E-07 | Tb |
| cDC2 | IGF2BP2 | 1.01E-11 | 0.553122148 | 0.414 | 0.248 | 1.98E-07 | Tb |
| cDC2 | KDSR | 2.98E-16 | 0.553042328 | 0.464 | 0.249 | 5.86E-12 | Tb |
| cDC2 | GLG1 | 5.68E-11 | 0.552761404 | 0.487 | 0.309 | 1.12E-06 | Tb |
| cDC2 | FBXL20 | 3.29E-08 | 0.552614818 | 0.25 | 0.14 | 0.000647623 | Tb |
| cDC2 | SS18L2 | 2.44E-13 | 0.552021155 | 0.485 | 0.278 | 4.81E-09 | Tb |
| cDC2 | CKLF | 2.61E-17 | 0.551825114 | 0.864 | 0.689 | 5.13E-13 | Tb |
| cDC2 | SMARCC1 | 1.59E-12 | 0.551572009 | 0.572 | 0.37 | 3.13E-08 | Tb |
| cDC2 | LTB | 2.26E-08 | 0.550790191 | 0.476 | 0.335 | 0.000445657 | Tb |
| cDC2 | PTPRJ | 3.24E-19 | 0.550571465 | 0.759 | 0.57 | 6.38E-15 | Tb |
| cDC2 | DESI2 | 5.69E-09 | 0.550192404 | 0.379 | 0.229 | 0.000112098 | Tb |
| cDC2 | LILRB4 | 1.37E-07 | 0.550081838 | 0.529 | 0.398 | 0.002687337 | Tb |
| cDC2 | CIAPIN1 | 4.06E-11 | 0.549810337 | 0.328 | 0.177 | 7.98E-07 | Tb |
| cDC2 | MRPS33 | 1.10E-13 | 0.549504831 | 0.435 | 0.239 | 2.17E-09 | Tb |
| cDC2 | ZDHHC12 | 1.56E-11 | 0.54949653 | 0.419 | 0.238 | 3.08E-07 | Tb |
| cDC2 | PSME3 | 1.48E-10 | 0.5491485 | 0.281 | 0.147 | 2.91E-06 | Tb |
| cDC2 | EXOC5 | 3.34E-11 | 0.54828982 | 0.349 | 0.192 | 6.58E-07 | Tb |
| cDC2 | CD69 | 7.72E-19 | 0.548117324 | 0.705 | 0.465 | 1.52E-14 | Tb |
| cDC2 | MRTFA | 1.74E-12 | 0.548048991 | 0.682 | 0.478 | 3.43E-08 | Tb |
| cDC2 | DOCK2 | 1.45E-13 | 0.547935409 | 0.752 | 0.531 | 2.84E-09 | Tb |
| cDC2 | SRP54 | 6.81E-12 | 0.547922592 | 0.618 | 0.408 | 1.34E-07 | Tb |
| cDC2 | YTHDF3 | 4.10E-09 | 0.547608745 | 0.688 | 0.521 | 8.07E-05 | Tb |
| cDC2 | GATAD2A | 8.17E-12 | 0.54741914 | 0.604 | 0.407 | 1.61E-07 | Tb |
| cDC2 | NPIPB5 | 5.95E-11 | 0.547286569 | 0.314 | 0.169 | 1.17E-06 | Tb |
| cDC2 | UPF1 | 1.70E-10 | 0.547184634 | 0.258 | 0.131 | 3.34E-06 | Tb |
| cDC2 | SAMD4B | 1.08E-11 | 0.547156122 | 0.379 | 0.216 | 2.12E-07 | Tb |
| cDC2 | ZNFX1 | 1.14E-08 | 0.546750366 | 0.538 | 0.365 | 0.000224793 | Tb |
| cDC2 | PNKD | 2.05E-09 | 0.546690188 | 0.44 | 0.268 | 4.03E-05 | Tb |
| cDC2 | CYP51A1 | 2.21E-06 | 0.546608753 | 0.464 | 0.333 | 0.043515467 | Tb |
| cDC2 | MAP3K20 | 7.23E-11 | 0.546505898 | 0.398 | 0.229 | 1.42E-06 | Tb |
| cDC2 | HIPK1 | 6.85E-11 | 0.546385751 | 0.419 | 0.248 | 1.35E-06 | Tb |
| cDC2 | VPS13D | 5.73E-09 | 0.545942443 | 0.435 | 0.286 | 0.000112691 | Tb |
| cDC2 | PPP1R16A | 4.07E-12 | 0.545732974 | 0.286 | 0.142 | 8.01E-08 | Tb |
| cDC2 | ANXA7 | 3.75E-16 | 0.545700818 | 0.709 | 0.445 | 7.39E-12 | Tb |
| cDC2 | NELFB | 9.89E-08 | 0.54567385 | 0.264 | 0.153 | 0.001945953 | Tb |
| cDC2 | SLC35C2 | 1.20E-10 | 0.545617843 | 0.332 | 0.186 | 2.35E-06 | Tb |
| cDC2 | SSH1 | 8.87E-12 | 0.545602307 | 0.36 | 0.2 | 1.75E-07 | Tb |
| cDC2 | HDDC3 | 1.22E-13 | 0.54554322 | 0.267 | 0.121 | 2.39E-09 | Tb |
| cDC2 | MRPS7 | 1.36E-13 | 0.545152601 | 0.55 | 0.343 | 2.67E-09 | Tb |
| cDC2 | CLCN7 | 3.94E-10 | 0.545080427 | 0.332 | 0.186 | 7.75E-06 | Tb |
| cDC2 | NUDT14 | 7.00E-11 | 0.544936091 | 0.356 | 0.203 | 1.38E-06 | Tb |
| cDC2 | ABI3 | 3.42E-15 | 0.544913849 | 0.586 | 0.379 | 6.73E-11 | Tb |
| cDC2 | RAB11FIP1 | 2.94E-10 | 0.544606058 | 0.764 | 0.627 | 5.78E-06 | Tb |
| cDC2 | TRIM13 | 3.69E-11 | 0.544587027 | 0.377 | 0.214 | 7.27E-07 | Tb |
| cDC2 | TUBB6 | 6.70E-11 | 0.544090341 | 0.602 | 0.428 | 1.32E-06 | Tb |
| cDC2 | EXOC1 | 1.04E-13 | 0.544010941 | 0.382 | 0.203 | 2.04E-09 | Tb |
| cDC2 | TSPAN31 | 3.02E-13 | 0.543598738 | 0.277 | 0.129 | 5.94E-09 | Tb |
| cDC2 | ZFC3H1 | 7.50E-11 | 0.543211234 | 0.661 | 0.466 | 1.48E-06 | Tb |
| cDC2 | FZD2 | 3.59E-20 | 0.543048799 | 0.333 | 0.14 | 7.07E-16 | Tb |
| cDC2 | FAM120B | 6.19E-13 | 0.542227174 | 0.26 | 0.124 | 1.22E-08 | Tb |
| cDC2 | CTCF | 5.46E-12 | 0.541723055 | 0.36 | 0.196 | 1.07E-07 | Tb |
| cDC2 | TIFAB | 1.24E-10 | 0.541636452 | 0.325 | 0.183 | 2.44E-06 | Tb |
| cDC2 | NUP98 | 5.20E-13 | 0.541527599 | 0.7 | 0.488 | 1.02E-08 | Tb |
| cDC2 | RAMP1 | 2.71E-08 | 0.54145546 | 0.277 | 0.165 | 0.000532863 | Tb |
| cDC2 | CYP20A1 | 6.94E-07 | 0.540840547 | 0.361 | 0.238 | 0.013667875 | Tb |
| cDC2 | BOLA3 | 3.41E-10 | 0.54069641 | 0.41 | 0.249 | 6.72E-06 | Tb |

| | | | | | | | |
|---|---|---|---|---|---|---|---|
| cDC2 | PHACTR2 | 7.18E-10 | 0.540303774 | 0.551 | 0.374 | 1.41E-05 | Tb |
| cDC2 | ZDHHC5 | 1.45E-10 | 0.540106787 | 0.323 | 0.177 | 2.85E-06 | Tb |
| cDC2 | PDCD4 | 2.96E-10 | 0.539023119 | 0.44 | 0.267 | 5.83E-06 | Tb |
| cDC2 | EIF2S2 | 1.70E-18 | 0.53884311 | 0.733 | 0.471 | 3.35E-14 | Tb |
| cDC2 | CCDC25 | 2.87E-10 | 0.538588216 | 0.358 | 0.203 | 5.65E-06 | Tb |
| cDC2 | ABCE1 | 1.35E-12 | 0.537879108 | 0.506 | 0.306 | 2.65E-08 | Tb |
| cDC2 | TMEM165 | 5.95E-15 | 0.537693774 | 0.719 | 0.503 | 1.17E-10 | Tb |
| cDC2 | ARNTL2 | 4.97E-07 | 0.537486294 | 0.283 | 0.176 | 0.009773529 | Tb |
| cDC2 | SETX | 1.28E-15 | 0.537308157 | 0.682 | 0.441 | 2.51E-11 | Tb |
| cDC2 | RBM47 | 5.49E-13 | 0.536734362 | 0.827 | 0.656 | 1.08E-08 | Tb |
| cDC2 | CMTM7 | 3.21E-14 | 0.536718053 | 0.682 | 0.452 | 6.32E-10 | Tb |
| cDC2 | CIR1 | 4.19E-12 | 0.536287513 | 0.515 | 0.312 | 8.24E-08 | Tb |
| cDC2 | ETNK1 | 5.81E-17 | 0.535327558 | 0.658 | 0.416 | 1.14E-12 | Tb |
| cDC2 | HSD17B4 | 1.31E-09 | 0.535319112 | 0.373 | 0.221 | 2.58E-05 | Tb |
| cDC2 | SPIN1 | 2.47E-08 | 0.535127793 | 0.269 | 0.151 | 0.000485358 | Tb |
| cDC2 | MRPS23 | 2.19E-12 | 0.534985982 | 0.436 | 0.254 | 4.31E-08 | Tb |
| cDC2 | MYO1E | 5.82E-13 | 0.53487919 | 0.798 | 0.629 | 1.14E-08 | Tb |
| cDC2 | DCTN5 | 1.47E-12 | 0.534187348 | 0.267 | 0.127 | 2.90E-08 | Tb |
| cDC2 | ATP6V0A1 | 1.45E-12 | 0.534025885 | 0.494 | 0.304 | 2.86E-08 | Tb |
| cDC2 | SLC37A2 | 2.47E-07 | 0.533846477 | 0.311 | 0.192 | 0.004863122 | Tb |
| cDC2 | ARHGAP26 | 2.12E-11 | 0.533443015 | 0.782 | 0.606 | 4.17E-07 | Tb |
| cDC2 | PWP1 | 3.36E-10 | 0.53311258 | 0.358 | 0.207 | 6.61E-06 | Tb |
| cDC2 | SLC4A7 | 2.01E-12 | 0.532933803 | 0.529 | 0.327 | 3.95E-08 | Tb |
| cDC2 | EIF3J | 1.27E-15 | 0.53256544 | 0.654 | 0.402 | 2.50E-11 | Tb |
| cDC2 | PPP2CB | 4.11E-14 | 0.532330535 | 0.415 | 0.223 | 8.09E-10 | Tb |
| cDC2 | HECTD1 | 3.30E-15 | 0.531248844 | 0.515 | 0.297 | 6.49E-11 | Tb |
| cDC2 | ABCF1 | 2.55E-11 | 0.53117279 | 0.515 | 0.319 | 5.03E-07 | Tb |
| cDC2 | GRIPAP1 | 2.49E-11 | 0.531054243 | 0.428 | 0.251 | 4.90E-07 | Tb |
| cDC2 | KMT2C | 3.94E-18 | 0.530965229 | 0.792 | 0.568 | 7.75E-14 | Tb |
| cDC2 | TANK | 1.02E-10 | 0.530797129 | 0.855 | 0.748 | 2.00E-06 | Tb |
| cDC2 | STAT2 | 1.85E-11 | 0.530582148 | 0.475 | 0.29 | 3.64E-07 | Tb |
| cDC2 | DICER1 | 6.50E-12 | 0.530451276 | 0.44 | 0.257 | 1.28E-07 | Tb |
| cDC2 | LRR1 | 1.28E-09 | 0.530440021 | 0.293 | 0.164 | 2.53E-05 | Tb |
| cDC2 | PAICS | 8.03E-09 | 0.530410279 | 0.281 | 0.156 | 0.000158011 | Tb |
| cDC2 | SMC5 | 6.72E-13 | 0.530178576 | 0.565 | 0.362 | 1.32E-08 | Tb |
| cDC2 | KIF2A | 1.10E-09 | 0.529633601 | 0.548 | 0.377 | 2.17E-05 | Tb |
| cDC2 | NFIL3 | 4.33E-09 | 0.529432707 | 0.567 | 0.422 | 8.52E-05 | Tb |
| cDC2 | SLC8A1-AS1 | 6.15E-12 | 0.52929833 | 0.408 | 0.239 | 1.21E-07 | Tb |
| cDC2 | MYCBP2 | 6.36E-13 | 0.528988911 | 0.686 | 0.475 | 1.25E-08 | Tb |
| cDC2 | DGKH | 1.69E-09 | 0.528857824 | 0.304 | 0.172 | 3.34E-05 | Tb |
| cDC2 | FH | 3.97E-11 | 0.528647363 | 0.414 | 0.238 | 7.80E-07 | Tb |
| cDC2 | PLCL2 | 6.80E-10 | 0.528569999 | 0.433 | 0.273 | 1.34E-05 | Tb |
| cDC2 | GSAP | 1.28E-15 | 0.528502103 | 0.56 | 0.343 | 2.51E-11 | Tb |
| cDC2 | KIAA1143 | 1.24E-07 | 0.52847553 | 0.264 | 0.153 | 0.002432084 | Tb |
| cDC2 | SPTLC1 | 2.86E-09 | 0.528341771 | 0.38 | 0.228 | 5.63E-05 | Tb |
| cDC2 | GPCPD1 | 6.41E-16 | 0.527889193 | 0.846 | 0.654 | 1.26E-11 | Tb |
| cDC2 | USPL1 | 1.02E-08 | 0.527560371 | 0.593 | 0.415 | 0.000200406 | Tb |
| cDC2 | MYO1F | 2.07E-10 | 0.527381523 | 0.586 | 0.402 | 4.08E-06 | Tb |
| cDC2 | CISD1 | 1.63E-10 | 0.526960618 | 0.361 | 0.208 | 3.21E-06 | Tb |
| cDC2 | PRRC2B | 4.76E-09 | 0.526797853 | 0.387 | 0.239 | 9.36E-05 | Tb |
| cDC2 | PAGR1 | 1.97E-08 | 0.526576246 | 0.274 | 0.153 | 0.00038779 | Tb |
| cDC2 | COX16 | 3.70E-16 | 0.526312634 | 0.599 | 0.352 | 7.29E-12 | Tb |
| cDC2 | PRPF8 | 1.51E-10 | 0.526233071 | 0.482 | 0.297 | 2.98E-06 | Tb |
| cDC2 | NIFK | 1.52E-13 | 0.526157354 | 0.522 | 0.309 | 3.00E-09 | Tb |
| cDC2 | KATNBL1 | 1.46E-09 | 0.52586371 | 0.455 | 0.279 | 2.88E-05 | Tb |
| cDC2 | WDR59 | 1.27E-09 | 0.525669891 | 0.323 | 0.184 | 2.49E-05 | Tb |

| | | | | | | | |
|---|---|---|---|---|---|---|---|
| cDC2 | KDM5A | 1.61E-13 | 0.525606168 | 0.628 | 0.409 | 3.17E-09 | Tb |
| cDC2 | GSE1 | 1.77E-12 | 0.525527535 | 0.271 | 0.132 | 3.49E-08 | Tb |
| cDC2 | SEC14L1 | 6.50E-08 | 0.525485875 | 0.647 | 0.465 | 0.001280228 | Tb |
| cDC2 | UTP6 | 1.53E-13 | 0.525427346 | 0.475 | 0.271 | 3.02E-09 | Tb |
| cDC2 | HERC4 | 7.02E-10 | 0.525370626 | 0.387 | 0.236 | 1.38E-05 | Tb |
| cDC2 | BLOC1S4 | 4.74E-10 | 0.524849553 | 0.253 | 0.131 | 9.33E-06 | Tb |
| cDC2 | GBE1 | 1.47E-12 | 0.52446996 | 0.462 | 0.281 | 2.88E-08 | Tb |
| cDC2 | EAPP | 1.93E-13 | 0.524461718 | 0.405 | 0.223 | 3.79E-09 | Tb |
| cDC2 | CHMP2A | 1.13E-17 | 0.52442392 | 0.764 | 0.531 | 2.22E-13 | Tb |
| cDC2 | TOMM70 | 1.31E-08 | 0.524174212 | 0.265 | 0.147 | 0.000257592 | Tb |
| cDC2 | GDE1 | 1.71E-11 | 0.524082539 | 0.279 | 0.138 | 3.38E-07 | Tb |
| cDC2 | STX17 | 9.08E-08 | 0.523819549 | 0.328 | 0.201 | 0.00178682 | Tb |
| cDC2 | PUM1 | 7.38E-11 | 0.523356901 | 0.545 | 0.354 | 1.45E-06 | Tb |
| cDC2 | PPM1K | 8.30E-10 | 0.52333288 | 0.288 | 0.157 | 1.63E-05 | Tb |
| cDC2 | QPCT | 4.46E-07 | 0.52328909 | 0.541 | 0.429 | 0.00877509 | Tb |
| cDC2 | UBA5 | 2.64E-08 | 0.522501698 | 0.265 | 0.148 | 0.000519186 | Tb |
| cDC2 | UBE2K | 8.08E-13 | 0.52226856 | 0.627 | 0.427 | 1.59E-08 | Tb |
| cDC2 | N4BP2L2 | 9.25E-15 | 0.522230046 | 0.757 | 0.535 | 1.82E-10 | Tb |
| cDC2 | SSH2 | 3.50E-09 | 0.522183324 | 0.642 | 0.472 | 6.89E-05 | Tb |
| cDC2 | BMP2K | 4.60E-12 | 0.522003016 | 0.651 | 0.447 | 9.06E-08 | Tb |
| cDC2 | MKNK1 | 1.44E-10 | 0.52108959 | 0.417 | 0.253 | 2.83E-06 | Tb |
| cDC2 | REEP3 | 1.11E-09 | 0.521046237 | 0.438 | 0.278 | 2.19E-05 | Tb |
| cDC2 | NR3C1 | 1.11E-11 | 0.519873584 | 0.74 | 0.532 | 2.19E-07 | Tb |
| cDC2 | NFIC | 2.22E-07 | 0.519681899 | 0.393 | 0.249 | 0.00437386 | Tb |
| cDC2 | DENND1A | 9.31E-11 | 0.519327596 | 0.602 | 0.418 | 1.83E-06 | Tb |
| cDC2 | MTFP1 | 1.27E-07 | 0.519324437 | 0.253 | 0.146 | 0.00249847 | Tb |
| cDC2 | CREBRF | 4.18E-12 | 0.519235995 | 0.459 | 0.265 | 8.23E-08 | Tb |
| cDC2 | ZNF644 | 1.26E-11 | 0.519174505 | 0.604 | 0.417 | 2.48E-07 | Tb |
| cDC2 | ARHGAP22 | 1.08E-06 | 0.519160036 | 0.373 | 0.252 | 0.02125164 | Tb |
| cDC2 | CPM | 1.30E-11 | 0.51874771 | 0.454 | 0.283 | 2.56E-07 | Tb |
| cDC2 | ISCA1 | 2.75E-10 | 0.51857607 | 0.373 | 0.218 | 5.42E-06 | Tb |
| cDC2 | TMX2 | 2.81E-13 | 0.517730376 | 0.323 | 0.161 | 5.53E-09 | Tb |
| cDC2 | ZDHHC17 | 1.65E-07 | 0.517440128 | 0.311 | 0.19 | 0.003246981 | Tb |
| cDC2 | MON2 | 2.74E-10 | 0.51728804 | 0.396 | 0.24 | 5.39E-06 | Tb |
| cDC2 | BTBD9 | 1.53E-14 | 0.517243124 | 0.656 | 0.43 | 3.01E-10 | Tb |
| cDC2 | NIPA2 | 5.44E-11 | 0.516956506 | 0.442 | 0.266 | 1.07E-06 | Tb |
| cDC2 | POGLUT1 | 2.13E-11 | 0.516269173 | 0.373 | 0.211 | 4.19E-07 | Tb |
| cDC2 | HEXA | 4.90E-07 | 0.516055397 | 0.565 | 0.416 | 0.009639596 | Tb |
| cDC2 | MRPL47 | 4.05E-13 | 0.515965927 | 0.574 | 0.36 | 7.97E-09 | Tb |
| cDC2 | ERAP2 | 1.74E-11 | 0.515621925 | 0.366 | 0.203 | 3.43E-07 | Tb |
| cDC2 | IKZF1 | 2.07E-10 | 0.515616 | 0.668 | 0.512 | 4.06E-06 | Tb |
| cDC2 | TASOR2 | 5.67E-12 | 0.51548893 | 0.436 | 0.261 | 1.12E-07 | Tb |
| cDC2 | SH3GL1 | 1.54E-11 | 0.515322135 | 0.485 | 0.297 | 3.04E-07 | Tb |
| cDC2 | SLC1A3 | 2.08E-10 | 0.51525436 | 0.307 | 0.174 | 4.09E-06 | Tb |
| cDC2 | TRRAP | 3.44E-07 | 0.514857274 | 0.297 | 0.178 | 0.00676773 | Tb |
| cDC2 | NDUFAB1 | 9.02E-15 | 0.513912933 | 0.759 | 0.54 | 1.77E-10 | Tb |
| cDC2 | ERO1B | 2.37E-06 | 0.513884823 | 0.539 | 0.391 | 0.046740428 | Tb |
| cDC2 | FAR1 | 2.34E-10 | 0.513658956 | 0.551 | 0.364 | 4.61E-06 | Tb |
| cDC2 | XRN1 | 1.60E-10 | 0.513105448 | 0.668 | 0.463 | 3.14E-06 | Tb |
| cDC2 | GLOD4 | 4.81E-11 | 0.512882327 | 0.37 | 0.205 | 9.47E-07 | Tb |
| cDC2 | ASCC3 | 7.82E-13 | 0.512601104 | 0.548 | 0.343 | 1.54E-08 | Tb |
| cDC2 | TNRC6A | 4.20E-10 | 0.512577685 | 0.408 | 0.248 | 8.27E-06 | Tb |
| cDC2 | XRCC4 | 1.45E-06 | 0.512449707 | 0.262 | 0.162 | 0.028532012 | Tb |
| cDC2 | TBRG4 | 2.99E-10 | 0.512101362 | 0.304 | 0.163 | 5.88E-06 | Tb |
| cDC2 | KIAA0100 | 3.52E-13 | 0.511763002 | 0.424 | 0.24 | 6.94E-09 | Tb |
| cDC2 | TFRC | 6.66E-15 | 0.511420098 | 0.609 | 0.382 | 1.31E-10 | Tb |

| | | | | | | | |
|---|---|---|---|---|---|---|---|
| cDC2 | PTPN2 | 8.95E-14 | 0.511297598 | 0.808 | 0.627 | 1.76E-09 | Tb |
| cDC2 | AEBP2 | 3.27E-11 | 0.510816898 | 0.386 | 0.221 | 6.44E-07 | Tb |
| cDC2 | NRDC | 1.37E-12 | 0.510750237 | 0.747 | 0.528 | 2.69E-08 | Tb |
| cDC2 | FAM192A | 1.77E-11 | 0.510584369 | 0.475 | 0.289 | 3.47E-07 | Tb |
| cDC2 | HMBOX1 | 3.28E-11 | 0.510526324 | 0.461 | 0.281 | 6.46E-07 | Tb |
| cDC2 | NOCT | 2.25E-08 | 0.510338238 | 0.33 | 0.2 | 0.000442364 | Tb |
| cDC2 | DARS | 3.56E-10 | 0.509629061 | 0.55 | 0.365 | 7.01E-06 | Tb |
| cDC2 | KRCC1 | 6.82E-10 | 0.509482441 | 0.291 | 0.162 | 1.34E-05 | Tb |
| cDC2 | FAM193A | 4.47E-08 | 0.50883522 | 0.312 | 0.188 | 0.000879387 | Tb |
| cDC2 | KLF9 | 3.38E-10 | 0.508681302 | 0.36 | 0.203 | 6.65E-06 | Tb |
| cDC2 | HCAR2 | 5.06E-09 | 0.50854846 | 0.574 | 0.415 | 9.95E-05 | Tb |
| cDC2 | ACTN4 | 9.46E-13 | 0.508153545 | 0.674 | 0.453 | 1.86E-08 | Tb |
| cDC2 | PARP14 | 1.24E-16 | 0.508129932 | 0.822 | 0.602 | 2.44E-12 | Tb |
| cDC2 | RRP7A | 7.74E-10 | 0.508113696 | 0.489 | 0.306 | 1.52E-05 | Tb |
| cDC2 | SLCO3A1 | 4.81E-14 | 0.507757063 | 0.759 | 0.561 | 9.46E-10 | Tb |
| cDC2 | TSPOAP1-AS1 | 4.35E-11 | 0.506002064 | 0.291 | 0.154 | 8.56E-07 | Tb |
| cDC2 | KIAA1109 | 2.38E-08 | 0.505748752 | 0.414 | 0.268 | 0.000468093 | Tb |
| cDC2 | RBM33 | 8.98E-11 | 0.505326681 | 0.469 | 0.282 | 1.77E-06 | Tb |
| cDC2 | CPED1 | 6.07E-07 | 0.505206206 | 0.332 | 0.216 | 0.011947915 | Tb |
| cDC2 | HCP5 | 1.74E-10 | 0.505008439 | 0.339 | 0.189 | 3.42E-06 | Tb |
| cDC2 | ELL | 6.91E-15 | 0.504919334 | 0.403 | 0.209 | 1.36E-10 | Tb |
| cDC2 | SETD51 | 7.55E-15 | 0.504478527 | 0.557 | 0.343 | 1.49E-10 | Tb |
| cDC2 | CARM1 | 4.40E-10 | 0.503903746 | 0.264 | 0.135 | 8.65E-06 | Tb |
| cDC2 | TNPO1 | 1.41E-10 | 0.503822871 | 0.45 | 0.273 | 2.77E-06 | Tb |
| cDC2 | TERF1 | 3.47E-10 | 0.503436758 | 0.333 | 0.186 | 6.83E-06 | Tb |
| cDC2 | LYST | 5.11E-10 | 0.503326775 | 0.667 | 0.487 | 1.01E-05 | Tb |
| cDC2 | EPRS | 1.04E-17 | 0.50307388 | 0.52 | 0.279 | 2.04E-13 | Tb |
| cDC2 | MPHOSPH10 | 4.52E-10 | 0.502933278 | 0.354 | 0.199 | 8.89E-06 | Tb |
| cDC2 | TXNDC17 | 7.20E-16 | 0.501800417 | 0.724 | 0.488 | 1.42E-11 | Tb |
| cDC2 | PAM16 | 1.55E-09 | 0.500671819 | 0.356 | 0.207 | 3.05E-05 | Tb |
| cDC2 | SNX4 | 4.15E-09 | 0.500615507 | 0.353 | 0.207 | 8.16E-05 | Tb |
| cDC2 | KIF13A | 2.90E-08 | 0.500102399 | 0.258 | 0.148 | 0.00057135 | Tb |
| **Disease-associated genes in LAMP3$^{+}$ DC** | | | | | | | |
| LAMP3+ DC | EREG | 3.07E-07 | 4.265712869 | 0.311 | 0.088 | 0.006050513 | Lep-L |
| LAMP3+ DC | PID1 | 7.53E-12 | 3.85865853 | 0.289 | 0.041 | 1.48E-07 | Lep-L |
| LAMP3+ DC | SDS | 1.41E-08 | 3.801626093 | 0.267 | 0.052 | 0.000276608 | Lep-L |
| LAMP3+ DC | FMN1 | 8.36E-07 | 3.545126265 | 0.289 | 0.079 | 0.016461836 | Lep-L |
| LAMP3+ DC | OLR1 | 4.67E-07 | 3.496073284 | 0.267 | 0.065 | 0.009186411 | Lep-L |
| LAMP3+ DC | PDE6G | 3.51E-07 | 3.394636942 | 0.311 | 0.088 | 0.006913499 | Lep-L |
| LAMP3+ DC | RCAN1 | 1.05E-08 | 2.992396476 | 0.333 | 0.081 | 0.000206062 | Lep-L |
| LAMP3+ DC | FCGR2B | 3.78E-09 | 2.939712994 | 0.289 | 0.058 | 7.45E-05 | Lep-L |
| LAMP3+ DC | KMO | 1.15E-10 | 2.823796298 | 0.578 | 0.2 | 2.26E-06 | Lep-L |
| LAMP3+ DC | C5AR1 | 6.86E-10 | 2.71427531 | 0.578 | 0.241 | 1.35E-05 | Lep-L |
| LAMP3+ DC | IL1R2 | 2.22E-06 | 2.363404625 | 0.4 | 0.148 | 0.043639907 | Lep-L |
| LAMP3+ DC | GPR35 | 1.05E-07 | 2.184373972 | 0.356 | 0.099 | 0.002069993 | Lep-L |
| LAMP3+ DC | HLA-DRB5 | 1.46E-11 | 2.07818973 | 0.844 | 0.539 | 2.88E-07 | Lep-L |
| LAMP3+ DC | PRKACA | 1.43E-06 | 2.01117713 | 0.4 | 0.15 | 0.028128746 | Lep-L |
| LAMP3+ DC | SMIM3 | 5.77E-07 | 1.902751571 | 0.444 | 0.173 | 0.011354964 | Lep-L |
| LAMP3+ DC | FTL | 5.03E-11 | 1.881518 | 1 | 0.991 | 9.90E-07 | Lep-L |
| LAMP3+ DC | CXCL8 | 4.31E-07 | 1.786262732 | 0.644 | 0.362 | 0.008491464 | Lep-L |
| LAMP3+ DC | EMP3 | 1.63E-09 | 1.641614524 | 0.822 | 0.523 | 3.20E-05 | Lep-L |
| LAMP3+ DC | OPN3 | 4.64E-07 | 1.605275674 | 0.578 | 0.281 | 0.009128791 | Lep-L |
| LAMP3+ DC | CKS2 | 4.56E-07 | 1.542704312 | 0.756 | 0.575 | 0.008969718 | Lep-L |
| LAMP3+ DC | CD68 | 2.75E-07 | 1.370158389 | 0.644 | 0.333 | 0.005413213 | Lep-L |
| LAMP3+ DC | ANXA5 | 1.72E-09 | 1.335402331 | 0.956 | 0.741 | 3.38E-05 | Lep-L |
| LAMP3+ DC | SAT1 | 5.21E-11 | 1.323625412 | 1 | 0.991 | 1.03E-06 | Lep-L |

| | | | | | | | |
|---|---|---|---|---|---|---|---|
| LAMP3+ DC | PRDX1 | 1.19E-10 | 1.263045642 | 0.978 | 0.78 | 2.34E-06 | Lep-L |
| LAMP3+ DC | SDCBP | 3.38E-07 | 1.259881992 | 0.867 | 0.809 | 0.006647508 | Lep-L |
| LAMP3+ DC | ATP6V0C | 1.89E-08 | 1.132436408 | 0.889 | 0.719 | 0.000371828 | Lep-L |
| LAMP3+ DC | G0S2 | 1.04E-07 | 1.088070246 | 0.956 | 0.82 | 0.002055531 | Lep-L |
| LAMP3+ DC | CD44 | 9.00E-08 | 1.043598453 | 0.956 | 0.789 | 0.001771105 | Lep-L |
| LAMP3+ DC | ST13 | 2.33E-06 | 0.879754028 | 0.933 | 0.784 | 0.045830426 | Lep-L |
| LAMP3+ DC | FTH1 | 2.41E-07 | 0.792839551 | 1 | 0.995 | 0.004745599 | Lep-L |
| LAMP3+ DC | OAZ1 | 4.42E-09 | 0.77965721 | 1 | 0.915 | 8.69E-05 | Lep-L |
| LAMP3$^+$ DC | BCL2A1 | 1.20E-06 | 0.754927435 | 0.978 | 0.932 | 0.023602404 | Lep-L |
| LAMP3$^+$ DC | SQSTM1 | 4.79E-08 | 0.629351301 | 1 | 0.941 | 0.000941894 | Lep-L |
| LAMP3+ DC | CFD | 2.20E-10 | 2.260053136 | 0.271 | 0.079 | 4.34E-06 | Lep-T |
| LAMP3+ DC | ALDOA1 | 7.13E-07 | 2.233171075 | 0.339 | 0.19 | 0.014028166 | Lep-T |
| LAMP3+ DC | C1QA | 1.85E-06 | 2.08443957 | 0.385 | 0.214 | 0.036513486 | Lep-T |
| LAMP3+ DC | DCN | 6.58E-07 | 1.756507812 | 0.29 | 0.129 | 0.012957206 | Lep-T |
| LAMP3+ DC | RBFOX21 | 2.86E-10 | 1.706917402 | 0.697 | 0.52 | 5.63E-06 | Lep-T |
| LAMP3+ DC | MMP2 | 3.57E-08 | 1.693551834 | 0.276 | 0.103 | 0.000703342 | Lep-T |
| LAMP3+ DC | RTN1 | 6.02E-09 | 1.635263903 | 0.398 | 0.193 | 0.000118526 | Lep-T |
| LAMP3+ DC | MEAK7 | 8.39E-07 | 1.635026242 | 0.276 | 0.124 | 0.016505509 | Lep-T |
| LAMP3+ DC | FGL2 | 1.17E-07 | 1.552836075 | 0.335 | 0.166 | 0.002301009 | Lep-T |
| LAMP3+ DC | FCGR2A | 7.79E-07 | 1.526784748 | 0.376 | 0.203 | 0.015326267 | Lep-T |
| LAMP3+ DC | HIST2H2AC | 6.07E-08 | 1.466615468 | 0.285 | 0.113 | 0.001193993 | Lep-T |
| LAMP3+ DC | KCNMA1 | 5.89E-12 | 1.447906756 | 0.52 | 0.272 | 1.16E-07 | Lep-T |
| LAMP3+ DC | FYB1 | 1.10E-09 | 1.372822299 | 0.507 | 0.293 | 2.17E-05 | Lep-T |
| LAMP3+ DC | SNHG32 | 1.37E-06 | 1.294698192 | 0.498 | 0.356 | 0.027001975 | Lep-T |
| LAMP3+ DC | TSC22D3 | 1.06E-06 | 1.204247335 | 0.52 | 0.361 | 0.020827977 | Lep-T |
| LAMP3+ DC | HLA-DQB2 | 2.16E-07 | 1.192808495 | 0.498 | 0.327 | 0.00424547 | Lep-T |
| LAMP3+ DC | DMKN | 1.22E-08 | 1.177487754 | 0.308 | 0.113 | 0.000240306 | Lep-T |
| LAMP3+ DC | FRMD4A | 3.09E-07 | 1.175337519 | 0.498 | 0.301 | 0.006080978 | Lep-T |
| LAMP3+ DC | SNHG5 | 5.63E-08 | 1.055010059 | 0.719 | 0.72 | 0.001108436 | Lep-T |
| LAMP3+ DC | GRASP | 2.30E-07 | 1.043945114 | 0.751 | 0.712 | 0.004532813 | Lep-T |
| LAMP3+ DC | LMNA | 4.28E-09 | 1.042667379 | 0.665 | 0.47 | 8.43E-05 | Lep-T |
| LAMP3+ DC | KRT1 | 1.12E-10 | 1.027783138 | 0.312 | 0.092 | 2.20E-06 | Lep-T |
| LAMP3+ DC | PSAP1 | 5.95E-08 | 1.004415948 | 0.719 | 0.596 | 0.001171841 | Lep-T |
| LAMP3+ DC | NR4A1 | 1.40E-14 | 0.955497018 | 0.882 | 0.789 | 2.76E-10 | Lep-T |
| LAMP3+ DC | DUSP1 | 8.79E-09 | 0.887737672 | 0.973 | 0.921 | 0.000172952 | Lep-T |
| LAMP3+ DC | NR4A2 | 1.15E-06 | 0.886317159 | 0.733 | 0.675 | 0.022627554 | Lep-T |
| LAMP3+ DC | IL10RA | 2.77E-08 | 0.859556972 | 0.81 | 0.699 | 0.000545779 | Lep-T |
| LAMP3+ DC | CXCL14 | 1.43E-11 | 0.828779963 | 0.262 | 0.066 | 2.82E-07 | Lep-T |
| LAMP3+ DC | LGALS2 | 3.82E-08 | 0.803018756 | 0.869 | 0.784 | 0.000752005 | Lep-T |
| LAMP3+ DC | PNRC11 | 1.84E-20 | 0.784254246 | 1 | 0.979 | 3.63E-16 | Lep-T |
| LAMP3+ DC | TNFAIP3 | 5.13E-14 | 0.729130416 | 0.964 | 0.926 | 1.01E-09 | Lep-T |
| LAMP3+ DC | MCL1 | 8.36E-08 | 0.713771 | 0.819 | 0.839 | 0.001646538 | Lep-T |
| LAMP3+ DC | CTSZ1 | 4.80E-07 | 0.706933714 | 0.919 | 0.91 | 0.009456306 | Lep-T |
| LAMP3+ DC | CD441 | 1.11E-08 | 0.703298943 | 0.878 | 0.757 | 0.000217783 | Lep-T |
| LAMP3+ DC | EEF2 | 1.57E-09 | 0.695550442 | 0.851 | 0.876 | 3.08E-05 | Lep-T |
| LAMP3+ DC | DNAJB1 | 1.98E-06 | 0.646737192 | 0.91 | 0.828 | 0.03888427 | Lep-T |
| LAMP3+ DC | HSP90AA1 | 5.51E-09 | 0.593672757 | 0.982 | 0.992 | 0.000108537 | Lep-T |
| LAMP3+ DC | CCDC200 | 3.24E-07 | 2.546862222 | 0.305 | 0.139 | 0.006381235 | Mari |
| LAMP3+ DC | AOC1 | 8.56E-07 | 1.853819712 | 0.355 | 0.176 | 0.01684917 | Mari |
| LAMP3+ DC | ANK3 | 1.76E-08 | 1.702806291 | 0.418 | 0.207 | 0.000345773 | Mari |
| LAMP3+ DC | PDCL3 | 2.13E-06 | 1.585555861 | 0.44 | 0.29 | 0.041967988 | Mari |
| LAMP3+ DC | LRRC61 | 9.23E-08 | 1.510225672 | 0.255 | 0.092 | 0.00181718 | Mari |
| LAMP3+ DC | NPFFR1 | 1.78E-08 | 1.371238768 | 0.44 | 0.216 | 0.00035122 | Mari |
| LAMP3+ DC | PPA1 | 8.75E-18 | 1.333866846 | 0.901 | 0.856 | 1.72E-13 | Mari |
| LAMP3+ DC | PLGRKT | 1.03E-15 | 1.287237582 | 0.78 | 0.545 | 2.04E-11 | Mari |
| LAMP3+ DC | BLOC1S2 | 1.30E-09 | 1.242982613 | 0.56 | 0.325 | 2.55E-05 | Mari |

| | | | | | | | |
|---|---|---|---|---|---|---|---|
| LAMP3+ DC | CHMP4A | 1.05E-08 | 1.095892557 | 0.709 | 0.573 | 0.0002067 | Mari |
| LAMP3+ DC | TIMM10 | 5.88E-07 | 1.074671657 | 0.525 | 0.351 | 0.011567947 | Mari |
| LAMP3+ DC | IFI30 | 2.13E-09 | 1.052329423 | 0.936 | 0.937 | 4.19E-05 | Mari |
| LAMP3+ DC | NDUFC2 | 4.81E-10 | 1.022491214 | 0.695 | 0.497 | 9.47E-06 | Mari |
| LAMP3+ DC | MRPL11 | 1.72E-06 | 1.008671252 | 0.489 | 0.318 | 0.033948785 | Mari |
| LAMP3+ DC | MRPL52 | 2.47E-09 | 1.006615308 | 0.667 | 0.484 | 4.86E-05 | Mari |
| LAMP3+ DC | HMGN1 | 1.40E-11 | 1.00591888 | 0.837 | 0.804 | 2.76E-07 | Mari |
| LAMP3+ DC | IAH1 | 5.79E-07 | 1.004578597 | 0.624 | 0.46 | 0.011405226 | Mari |
| LAMP3+ DC | GTF2H5 | 1.46E-06 | 1.00086178 | 0.496 | 0.333 | 0.028795776 | Mari |
| LAMP3+ DC | VAMP5 | 1.76E-08 | 0.986528275 | 0.738 | 0.571 | 0.000345856 | Mari |
| LAMP3+ DC | IDO1 | 7.88E-10 | 0.985034924 | 0.865 | 0.821 | 1.55E-05 | Mari |
| LAMP3+ DC | ATOX1 | 3.72E-15 | 0.974926036 | 0.879 | 0.85 | 7.32E-11 | Mari |
| LAMP3+ DC | PSMA2 | 1.52E-11 | 0.969588923 | 0.823 | 0.769 | 2.99E-07 | Mari |
| LAMP3+ DC | CLNS1A | 9.31E-08 | 0.95986841 | 0.752 | 0.597 | 0.001832637 | Mari |
| LAMP3+ DC | TMEM176B | 1.17E-12 | 0.951171713 | 0.844 | 0.754 | 2.31E-08 | Mari |
| LAMP3+ DC | CYB5A | 3.65E-07 | 0.943592837 | 0.723 | 0.593 | 0.007191234 | Mari |
| LAMP3+ DC | NDUFA3 | 2.84E-07 | 0.94238664 | 0.624 | 0.471 | 0.005591927 | Mari |
| LAMP3+ DC | TYMP | 1.74E-22 | 0.934698072 | 0.986 | 0.932 | 3.43E-18 | Mari |
| LAMP3+ DC | NDUFB1 | 3.16E-11 | 0.931433143 | 0.738 | 0.588 | 6.23E-07 | Mari |
| LAMP3+ DC | NDUFAB1 | 1.46E-07 | 0.910168556 | 0.624 | 0.466 | 0.002866892 | Mari |
| LAMP3+ DC | MYL12A | 4.72E-10 | 0.905879294 | 0.858 | 0.83 | 9.29E-06 | Mari |
| LAMP3+ DC | CHMP2A | 3.90E-08 | 0.900225093 | 0.709 | 0.527 | 0.000766876 | Mari |
| LAMP3+ DC | ZNHIT1 | 2.38E-06 | 0.894616679 | 0.631 | 0.505 | 0.046859812 | Mari |
| LAMP3+ DC | DAD1 | 3.98E-07 | 0.894030783 | 0.624 | 0.449 | 0.007841757 | Mari |
| LAMP3+ DC | OSTF1 | 2.52E-11 | 0.893855205 | 0.887 | 0.797 | 4.96E-07 | Mari |
| LAMP3+ DC | PSMB9 | 5.59E-11 | 0.893824241 | 0.823 | 0.773 | 1.10E-06 | Mari |
| LAMP3+ DC | BLOC1S1 | 5.26E-08 | 0.880065559 | 0.766 | 0.688 | 0.001036077 | Mari |
| LAMP3+ DC | SUB1 | 1.82E-13 | 0.877366 | 0.908 | 0.917 | 3.58E-09 | Mari |
| LAMP3+ DC | COX7A2L | 1.65E-06 | 0.870969823 | 0.794 | 0.725 | 0.032476229 | Mari |
| LAMP3+ DC | MRPL51 | 1.56E-08 | 0.868882255 | 0.709 | 0.571 | 0.00030654 | Mari |
| LAMP3+ DC | AP2S1 | 5.28E-11 | 0.857746704 | 0.83 | 0.736 | 1.04E-06 | Mari |
| LAMP3+ DC | PET100 | 3.27E-07 | 0.855087679 | 0.695 | 0.586 | 0.006427736 | Mari |
| LAMP3+ DC | BUD31 | 3.24E-07 | 0.839942944 | 0.702 | 0.58 | 0.006375146 | Mari |
| LAMP3+ DC | RBX1 | 1.85E-06 | 0.836904809 | 0.716 | 0.675 | 0.036416565 | Mari |
| LAMP3+ DC | TMSB10 | 5.19E-20 | 0.832278148 | 0.993 | 0.998 | 1.02E-15 | Mari |
| LAMP3+ DC | UQCR11 | 6.21E-11 | 0.829815077 | 0.837 | 0.802 | 1.22E-06 | Mari |
| LAMP3+ DC | SEC61G | 7.15E-07 | 0.822615155 | 0.73 | 0.667 | 0.014064272 | Mari |
| LAMP3+ DC | TBC1D4 | 2.04E-06 | 0.82052466 | 0.915 | 0.778 | 0.04019483 | Mari |
| LAMP3+ DC | TMEM167A | 2.28E-06 | 0.817570392 | 0.652 | 0.484 | 0.044827783 | Mari |
| LAMP3+ DC | MRPL57 | 5.06E-07 | 0.816065005 | 0.603 | 0.416 | 0.009959977 | Mari |
| LAMP3+ DC | LSM8 | 3.31E-07 | 0.815726836 | 0.709 | 0.564 | 0.006523643 | Mari |
| LAMP3+ DC | TRADD | 3.62E-07 | 0.800713481 | 0.759 | 0.627 | 0.007128466 | Mari |
| LAMP3+ DC | SNX2 | 2.31E-07 | 0.79421427 | 0.688 | 0.508 | 0.00454671 | Mari |
| LAMP3+ DC | SUMO1 | 2.73E-07 | 0.779798154 | 0.723 | 0.577 | 0.005381183 | Mari |
| LAMP3+ DC | GSTP1 | 9.52E-09 | 0.779256179 | 0.901 | 0.93 | 0.000187473 | Mari |
| LAMP3+ DC | POLR2L | 1.34E-06 | 0.778499121 | 0.702 | 0.588 | 0.026305206 | Mari |
| LAMP3+ DC | BZW1 | 3.74E-10 | 0.774538887 | 0.858 | 0.8 | 7.35E-06 | Mari |
| LAMP3+ DC | JPT1 | 2.02E-08 | 0.774096095 | 0.78 | 0.671 | 0.000397809 | Mari |
| LAMP3+ DC | ATP5MF | 7.03E-08 | 0.770505063 | 0.78 | 0.686 | 0.001384159 | Mari |
| LAMP3+ DC | PSMG2 | 5.72E-08 | 0.765286904 | 0.723 | 0.571 | 0.001126903 | Mari |
| LAMP3+ DC | POMP | 4.89E-08 | 0.765150946 | 0.809 | 0.743 | 0.00096242 | Mari |
| LAMP3+ DC | HMGN3 | 1.90E-06 | 0.754706284 | 0.851 | 0.795 | 0.037434259 | Mari |
| LAMP3+ DC | ISCU | 4.16E-08 | 0.754372701 | 0.83 | 0.758 | 0.000818119 | Mari |
| LAMP3+ DC | ATP5ME | 1.43E-08 | 0.736665248 | 0.766 | 0.671 | 0.000282324 | Mari |
| LAMP3+ DC | MRPL54 | 4.99E-08 | 0.736019967 | 0.645 | 0.44 | 0.000982306 | Mari |
| LAMP3+ DC | NDUFA13 | 6.79E-11 | 0.725408466 | 0.837 | 0.749 | 1.34E-06 | Mari |

| | | | | | | | |
|---|---|---|---|---|---|---|---|
| LAMP3+ DC | SNRPG | 5.87E-08 | 0.719772106 | 0.766 | 0.656 | 0.001156023 | Mari |
| LAMP3+ DC | PARK7 | 7.74E-10 | 0.719628947 | 0.879 | 0.824 | 1.52E-05 | Mari |
| LAMP3+ DC | PPDPF | 3.02E-09 | 0.719386462 | 0.844 | 0.73 | 5.94E-05 | Mari |
| LAMP3+ DC | GPX4 | 3.45E-16 | 0.714880088 | 0.95 | 0.941 | 6.79E-12 | Mari |
| LAMP3+ DC | LAP3 | 5.63E-11 | 0.713288215 | 0.908 | 0.834 | 1.11E-06 | Mari |
| LAMP3+ DC | UQCR10 | 2.76E-07 | 0.703638257 | 0.78 | 0.706 | 0.005430778 | Mari |
| LAMP3+ DC | ATP6V0B | 4.55E-08 | 0.700278637 | 0.816 | 0.725 | 0.000896149 | Mari |
| LAMP3+ DC | COX6C | 1.31E-09 | 0.697966575 | 0.83 | 0.839 | 2.57E-05 | Mari |
| LAMP3+ DC | UBL5 | 3.99E-13 | 0.692079371 | 0.872 | 0.832 | 7.85E-09 | Mari |
| LAMP3+ DC | ATP6V0E1 | 3.18E-10 | 0.691921936 | 0.865 | 0.826 | 6.27E-06 | Mari |
| LAMP3+ DC | MYDGF | 3.44E-07 | 0.689719281 | 0.709 | 0.558 | 0.006773136 | Mari |
| LAMP3+ DC | H2AFZ1 | 1.55E-09 | 0.688494829 | 0.908 | 0.922 | 3.04E-05 | Mari |
| LAMP3+ DC | CFL1 | 5.51E-12 | 0.688383262 | 0.936 | 0.922 | 1.09E-07 | Mari |
| LAMP3+ DC | GNG5 | 2.54E-10 | 0.680241806 | 0.879 | 0.904 | 4.99E-06 | Mari |
| LAMP3+ DC | SUMO2 | 8.07E-13 | 0.679173275 | 0.851 | 0.874 | 1.59E-08 | Mari |
| LAMP3+ DC | MYL12B | 5.33E-09 | 0.669423664 | 0.851 | 0.861 | 0.000104909 | Mari |
| LAMP3+ DC | RAN | 2.13E-09 | 0.668763157 | 0.83 | 0.813 | 4.19E-05 | Mari |
| LAMP3+ DC | SRP14 | 1.63E-10 | 0.668742312 | 0.851 | 0.906 | 3.21E-06 | Mari |
| LAMP3+ DC | NEDD8 | 3.96E-07 | 0.668553038 | 0.773 | 0.673 | 0.007798528 | Mari |
| LAMP3+ DC | TMSB4X | 1.20E-08 | 0.65976023 | 0.972 | 0.993 | 0.00023669 | Mari |
| LAMP3+ DC | EDF1 | 5.53E-13 | 0.658072199 | 0.887 | 0.878 | 1.09E-08 | Mari |
| LAMP3+ DC | MAP3K131 | 2.09E-07 | 0.650880963 | 0.879 | 0.769 | 0.004109793 | Mari |
| LAMP3+ DC | COX7A2 | 6.57E-07 | 0.644477046 | 0.801 | 0.782 | 0.012927653 | Mari |
| LAMP3+ DC | DBI | 3.78E-09 | 0.643432556 | 0.844 | 0.885 | 7.43E-05 | Mari |
| LAMP3+ DC | NDUFV2 | 1.16E-07 | 0.641173493 | 0.844 | 0.791 | 0.002290758 | Mari |
| LAMP3+ DC | TUBB4B1 | 6.91E-07 | 0.640185805 | 0.865 | 0.806 | 0.013610747 | Mari |
| LAMP3+ DC | ATP5F1E | 8.28E-16 | 0.631982566 | 0.936 | 0.963 | 1.63E-11 | Mari |
| LAMP3+ DC | UXT | 1.38E-06 | 0.631827592 | 0.816 | 0.743 | 0.027254414 | Mari |
| LAMP3+ DC | ELOB | 3.26E-08 | 0.614603673 | 0.83 | 0.778 | 0.000641173 | Mari |
| LAMP3+ DC | GADD45B | 1.26E-06 | 0.613772418 | 0.887 | 0.913 | 0.024801293 | Mari |
| LAMP3+ DC | NDUFA1 | 9.79E-08 | 0.612152197 | 0.801 | 0.745 | 0.001927463 | Mari |
| LAMP3+ DC | TMA7 | 2.58E-07 | 0.607145472 | 0.816 | 0.808 | 0.005075418 | Mari |
| LAMP3+ DC | SLC25A3 | 5.52E-07 | 0.606457278 | 0.858 | 0.839 | 0.010870375 | Mari |
| LAMP3+ DC | CLTA | 1.16E-06 | 0.602489282 | 0.816 | 0.754 | 0.022798336 | Mari |
| LAMP3+ DC | PPIA | 4.43E-12 | 0.602038064 | 0.865 | 0.941 | 8.73E-08 | Mari |
| LAMP3+ DC | SAP18 | 6.92E-07 | 0.600547894 | 0.83 | 0.769 | 0.013612042 | Mari |
| LAMP3+ DC | PSMB1 | 9.84E-07 | 0.594014332 | 0.809 | 0.725 | 0.019375814 | Mari |
| LAMP3+ DC | MYL6 | 2.57E-09 | 0.591236579 | 0.922 | 0.956 | 5.06E-05 | Mari |
| LAMP3+ DC | SEC61B | 1.77E-06 | 0.587837506 | 0.901 | 0.889 | 0.034774325 | Mari |
| LAMP3+ DC | IER2 | 8.78E-08 | 0.577884821 | 0.922 | 0.904 | 0.001727502 | Mari |
| LAMP3+ DC | ITM2B | 1.32E-07 | 0.566806002 | 0.894 | 0.906 | 0.002590926 | Mari |
| LAMP3+ DC | HINT1 | 1.51E-06 | 0.553102186 | 0.879 | 0.919 | 0.02976186 | Mari |
| LAMP3+ DC | COX5B | 2.00E-06 | 0.530296505 | 0.858 | 0.806 | 0.039448967 | Mari |
| LAMP3+ DC | B2M | 3.11E-11 | 0.50476445 | 0.986 | 1 | 6.12E-07 | Mari |
| LAMP3+ DC | NPHS1 | 8.99E-19 | 3.663827661 | 0.461 | 0.17 | 1.77E-14 | Tb |
| LAMP3+ DC | S100A7 | 2.12E-35 | 3.366206233 | 0.528 | 0.076 | 4.17E-31 | Tb |
| LAMP3+ DC | KRT6A | 2.38E-31 | 3.013509263 | 0.456 | 0.057 | 4.69E-27 | Tb |
| LAMP3+ DC | S100A2 | 2.23E-33 | 2.935344939 | 0.523 | 0.081 | 4.39E-29 | Tb |
| LAMP3+ DC | PLEKHA5 | 7.58E-09 | 2.666240608 | 0.456 | 0.243 | 0.000149222 | Tb |
| LAMP3+ DC | KRT6B | 1.83E-23 | 2.600770742 | 0.358 | 0.044 | 3.59E-19 | Tb |
| LAMP3+ DC | PSMD5 | 2.52E-16 | 2.451862583 | 0.523 | 0.231 | 4.96E-12 | Tb |
| LAMP3+ DC | GCNT2 | 1.65E-18 | 2.451212606 | 0.585 | 0.246 | 3.25E-14 | Tb |
| LAMP3+ DC | IKZF4 | 2.49E-15 | 2.399966695 | 0.352 | 0.093 | 4.91E-11 | Tb |
| LAMP3+ DC | KRT17 | 4.36E-17 | 2.286082274 | 0.347 | 0.076 | 8.58E-13 | Tb |
| LAMP3+ DC | PDE4C | 3.96E-15 | 2.187635643 | 0.295 | 0.061 | 7.80E-11 | Tb |
| LAMP3+ DC | PPP1R14A | 5.23E-15 | 2.146879127 | 0.389 | 0.111 | 1.03E-10 | Tb |

| | | | | | | | |
|---|---|---|---|---|---|---|---|
| LAMP3+ DC | S100A8 | 8.06E-35 | 1.937180658 | 0.741 | 0.204 | 1.59E-30 | Tb |
| LAMP3+ DC | KRT16 | 3.19E-23 | 1.896987204 | 0.415 | 0.069 | 6.29E-19 | Tb |
| LAMP3+ DC | FAM118A | 1.10E-21 | 1.880332767 | 0.705 | 0.349 | 2.17E-17 | Tb |
| LAMP3+ DC | KRT14 | 5.11E-31 | 1.823395469 | 0.72 | 0.233 | 1.01E-26 | Tb |
| LAMP3+ DC | PHF19 | 7.37E-17 | 1.814288516 | 0.674 | 0.41 | 1.45E-12 | Tb |
| LAMP3+ DC | ARL17B | 4.55E-17 | 1.797762141 | 0.415 | 0.111 | 8.96E-13 | Tb |
| LAMP3+ DC | CD226 | 2.40E-08 | 1.789171868 | 0.29 | 0.111 | 0.000471814 | Tb |
| LAMP3+ DC | ZNF76 | 1.30E-11 | 1.782385496 | 0.358 | 0.123 | 2.56E-07 | Tb |
| LAMP3+ DC | LRRC37A2 | 1.74E-14 | 1.750627112 | 0.269 | 0.047 | 3.42E-10 | Tb |
| LAMP3+ DC | CYP7B1 | 4.20E-09 | 1.749823529 | 0.487 | 0.265 | 8.27E-05 | Tb |
| LAMP3+ DC | SFN | 1.28E-07 | 1.730626331 | 0.316 | 0.138 | 0.002523798 | Tb |
| LAMP3+ DC | RASGRP3 | 9.12E-17 | 1.722383707 | 0.642 | 0.314 | 1.80E-12 | Tb |
| LAMP3+ DC | SCLY | 2.33E-10 | 1.721232769 | 0.347 | 0.128 | 4.58E-06 | Tb |
| LAMP3+ DC | ARL17A | 8.52E-22 | 1.716149328 | 0.601 | 0.221 | 1.68E-17 | Tb |
| LAMP3+ DC | FAM222B | 3.76E-16 | 1.710180525 | 0.44 | 0.138 | 7.41E-12 | Tb |
| LAMP3+ DC | S100A9 | 2.50E-28 | 1.699121953 | 0.756 | 0.283 | 4.91E-24 | Tb |
| LAMP3+ DC | DCAF10 | 1.35E-07 | 1.689385668 | 0.29 | 0.12 | 0.002659463 | Tb |
| LAMP3+ DC | RAB30-DT | 2.06E-18 | 1.679767387 | 0.544 | 0.179 | 4.05E-14 | Tb |
| LAMP3+ DC | CDKL3 | 3.16E-08 | 1.662914925 | 0.275 | 0.098 | 0.000621714 | Tb |
| LAMP3+ DC | TTN-AS1 | 2.61E-16 | 1.65894632 | 0.544 | 0.204 | 5.14E-12 | Tb |
| LAMP3+ DC | GANC | 2.31E-08 | 1.642489404 | 0.301 | 0.115 | 0.00045478 | Tb |
| LAMP3+ DC | DSTYK | 2.93E-08 | 1.640920725 | 0.264 | 0.091 | 0.000577217 | Tb |
| LAMP3+ DC | NAA25 | 1.64E-13 | 1.625673528 | 0.658 | 0.373 | 3.23E-09 | Tb |
| LAMP3+ DC | TATDN3 | 2.63E-07 | 1.623814539 | 0.301 | 0.125 | 0.0051771 | Tb |
| LAMP3+ DC | PRAG1 | 2.10E-09 | 1.622366629 | 0.378 | 0.162 | 4.14E-05 | Tb |
| LAMP3+ DC | NUS1 | 4.29E-07 | 1.586000494 | 0.316 | 0.135 | 0.008435506 | Tb |
| LAMP3+ DC | BCL2L1 | 5.98E-15 | 1.576163831 | 0.57 | 0.248 | 1.18E-10 | Tb |
| LAMP3+ DC | KIAA1211 | 4.32E-08 | 1.573513481 | 0.363 | 0.165 | 0.0008502 | Tb |
| LAMP3+ DC | NINJ2 | 2.98E-09 | 1.550843252 | 0.285 | 0.096 | 5.87E-05 | Tb |
| LAMP3+ DC | CRLF2 | 1.19E-12 | 1.535269161 | 0.539 | 0.263 | 2.34E-08 | Tb |
| LAMP3+ DC | KCNN1 | 2.53E-12 | 1.517950548 | 0.456 | 0.184 | 4.97E-08 | Tb |
| LAMP3+ DC | PHLPP1 | 9.61E-18 | 1.509441591 | 0.71 | 0.351 | 1.89E-13 | Tb |
| LAMP3+ DC | FUT10 | 1.19E-13 | 1.508568351 | 0.342 | 0.091 | 2.34E-09 | Tb |
| LAMP3+ DC | CCR6 | 1.32E-10 | 1.490866527 | 0.492 | 0.248 | 2.59E-06 | Tb |
| LAMP3+ DC | RHOH | 1.11E-13 | 1.483641451 | 0.699 | 0.435 | 2.19E-09 | Tb |
| LAMP3+ DC | IQCN | 7.64E-09 | 1.465339694 | 0.311 | 0.115 | 0.000150351 | Tb |
| LAMP3+ DC | ARHGAP25 | 2.47E-15 | 1.460182414 | 0.736 | 0.489 | 4.86E-11 | Tb |
| LAMP3+ DC | PILRB | 6.73E-10 | 1.437692807 | 0.264 | 0.076 | 1.32E-05 | Tb |
| LAMP3+ DC | TOP3A | 1.89E-10 | 1.419206189 | 0.332 | 0.108 | 3.72E-06 | Tb |
| LAMP3+ DC | CLEC16A | 2.48E-18 | 1.418236797 | 0.824 | 0.521 | 4.88E-14 | Tb |
| LAMP3+ DC | CD69 | 4.58E-07 | 1.417530479 | 0.352 | 0.167 | 0.009018054 | Tb |
| LAMP3+ DC | ZER1 | 8.87E-08 | 1.414019399 | 0.311 | 0.128 | 0.001745317 | Tb |
| LAMP3+ DC | KRT5 | 1.16E-11 | 1.410397354 | 0.352 | 0.115 | 2.28E-07 | Tb |
| LAMP3+ DC | OSBPL3 | 1.27E-09 | 1.406113902 | 0.466 | 0.221 | 2.50E-05 | Tb |
| LAMP3+ DC | MOB3B | 6.46E-14 | 1.4004195 | 0.554 | 0.253 | 1.27E-09 | Tb |
| LAMP3+ DC | FBXO38 | 2.26E-08 | 1.394802573 | 0.275 | 0.093 | 0.000443946 | Tb |
| LAMP3+ DC | RGS13 | 3.10E-10 | 1.37813043 | 0.43 | 0.184 | 6.11E-06 | Tb |
| LAMP3+ DC | CCDC18 | 5.87E-07 | 1.358052774 | 0.29 | 0.123 | 0.011550978 | Tb |
| LAMP3+ DC | CBWD3 | 4.23E-12 | 1.350772153 | 0.446 | 0.172 | 8.33E-08 | Tb |
| LAMP3+ DC | P2RY14 | 8.39E-07 | 1.350479463 | 0.425 | 0.233 | 0.016510027 | Tb |
| LAMP3+ DC | PATL2 | 1.50E-13 | 1.340235747 | 0.43 | 0.147 | 2.96E-09 | Tb |
| LAMP3+ DC | LNCAROD | 3.88E-07 | 1.320379542 | 0.425 | 0.233 | 0.00763979 | Tb |
| LAMP3+ DC | ECE1 | 1.34E-12 | 1.31851628 | 0.477 | 0.192 | 2.63E-08 | Tb |
| LAMP3+ DC | PHF8 | 2.73E-09 | 1.313921121 | 0.306 | 0.106 | 5.37E-05 | Tb |
| LAMP3+ DC | ODC1 | 1.17E-13 | 1.313069468 | 0.451 | 0.162 | 2.29E-09 | Tb |
| LAMP3+ DC | SLC35F5 | 2.70E-07 | 1.293586404 | 0.399 | 0.199 | 0.005324033 | Tb |

| | | | | | | | |
|---|---|---|---|---|---|---|---|
| LAMP3+ DC | SPIN1 | 1.53E-07 | 1.279834138 | 0.306 | 0.123 | 0.003015447 | Tb |
| LAMP3+ DC | DENND1B | 1.61E-14 | 1.278013131 | 0.788 | 0.538 | 3.18E-10 | Tb |
| LAMP3+ DC | LRRC8C | 4.36E-08 | 1.274413017 | 0.404 | 0.199 | 0.000858745 | Tb |
| LAMP3+ DC | TSTD2 | 4.94E-08 | 1.273112746 | 0.275 | 0.101 | 0.000971922 | Tb |
| LAMP3+ DC | CLIC4 | 1.75E-08 | 1.265542716 | 0.653 | 0.472 | 0.000343711 | Tb |
| LAMP3+ DC | ADNP2 | 6.07E-09 | 1.262032662 | 0.394 | 0.172 | 0.000119428 | Tb |
| LAMP3+ DC | FGFR1OP | 1.48E-07 | 1.261920032 | 0.259 | 0.093 | 0.002914322 | Tb |
| LAMP3+ DC | L3MBTL4 | 5.71E-09 | 1.261675604 | 0.746 | 0.555 | 0.000112426 | Tb |
| LAMP3+ DC | HDAC2-AS2 | 5.76E-09 | 1.260233064 | 0.368 | 0.152 | 0.000113421 | Tb |
| LAMP3+ DC | EYA4 | 1.62E-06 | 1.259932199 | 0.275 | 0.118 | 0.031831615 | Tb |
| LAMP3+ DC | ZC3H8 | 1.59E-06 | 1.243497923 | 0.254 | 0.101 | 0.031242223 | Tb |
| LAMP3+ DC | STIM1 | 2.04E-09 | 1.243423908 | 0.42 | 0.189 | 4.02E-05 | Tb |
| LAMP3+ DC | STAT4 | 6.58E-10 | 1.24151943 | 0.663 | 0.437 | 1.30E-05 | Tb |
| LAMP3+ DC | ZNF124 | 1.95E-09 | 1.235529594 | 0.363 | 0.143 | 3.84E-05 | Tb |
| LAMP3+ DC | LRRC8D | 1.49E-10 | 1.235285093 | 0.596 | 0.344 | 2.94E-06 | Tb |
| LAMP3+ DC | ADAM19 | 4.36E-19 | 1.231862407 | 0.824 | 0.518 | 8.59E-15 | Tb |
| LAMP3+ DC | SLC5A6 | 2.74E-08 | 1.231069373 | 0.342 | 0.14 | 0.000539314 | Tb |
| LAMP3+ DC | IL3RA | 2.50E-11 | 1.229061715 | 0.617 | 0.339 | 4.92E-07 | Tb |
| LAMP3+ DC | ARID3A | 1.05E-08 | 1.228529216 | 0.523 | 0.29 | 0.000206593 | Tb |
| LAMP3+ DC | BICRAL | 4.12E-11 | 1.227433117 | 0.425 | 0.17 | 8.12E-07 | Tb |
| LAMP3+ DC | MCM9 | 4.97E-09 | 1.226699125 | 0.373 | 0.157 | 9.77E-05 | Tb |
| LAMP3+ DC | MCU | 1.41E-06 | 1.223902117 | 0.347 | 0.17 | 0.02778736 | Tb |
| LAMP3+ DC | NLK | 1.69E-11 | 1.222483522 | 0.394 | 0.145 | 3.33E-07 | Tb |
| LAMP3+ DC | AP1S3 | 3.26E-11 | 1.217738857 | 0.601 | 0.329 | 6.42E-07 | Tb |
| LAMP3+ DC | MFSD14C | 1.59E-11 | 1.217556266 | 0.523 | 0.243 | 3.14E-07 | Tb |
| LAMP3+ DC | PBX3 | 2.50E-06 | 1.216598597 | 0.565 | 0.373 | 0.049194953 | Tb |
| LAMP3+ DC | TRAK1 | 1.99E-16 | 1.210451517 | 0.762 | 0.42 | 3.93E-12 | Tb |
| LAMP3+ DC | STAU2 | 2.33E-07 | 1.205147292 | 0.337 | 0.15 | 0.004595825 | Tb |
| LAMP3+ DC | MVB12B | 4.69E-08 | 1.205002206 | 0.409 | 0.197 | 0.000924016 | Tb |
| LAMP3+ DC | SPECC1L | 1.06E-13 | 1.202124541 | 0.658 | 0.344 | 2.09E-09 | Tb |
| LAMP3+ DC | TBC1D17 | 9.84E-10 | 1.197608818 | 0.446 | 0.201 | 1.94E-05 | Tb |
| LAMP3+ DC | ZBTB38 | 6.55E-07 | 1.196209702 | 0.596 | 0.423 | 0.012896473 | Tb |
| LAMP3+ DC | STX12 | 1.81E-08 | 1.188797537 | 0.637 | 0.425 | 0.000356233 | Tb |
| LAMP3+ DC | ALOX12-AS1 | 1.14E-07 | 1.184572646 | 0.321 | 0.135 | 0.002239261 | Tb |
| LAMP3+ DC | AHCYL2 | 1.34E-07 | 1.180733611 | 0.56 | 0.349 | 0.002645059 | Tb |
| LAMP3+ DC | UBAP2 | 9.17E-12 | 1.177265698 | 0.461 | 0.179 | 1.80E-07 | Tb |
| LAMP3+ DC | CDR2 | 3.87E-11 | 1.164099869 | 0.642 | 0.396 | 7.62E-07 | Tb |
| LAMP3+ DC | HOMER1 | 9.36E-08 | 1.161542716 | 0.492 | 0.265 | 0.001842922 | Tb |
| LAMP3+ DC | PLA1A | 5.86E-09 | 1.16151275 | 0.301 | 0.103 | 0.000115418 | Tb |
| LAMP3+ DC | DIPK1A | 5.52E-11 | 1.160916727 | 0.632 | 0.359 | 1.09E-06 | Tb |
| LAMP3+ DC | ABHD3 | 4.35E-09 | 1.154612464 | 0.674 | 0.445 | 8.55E-05 | Tb |
| LAMP3+ DC | DENND5B | 2.81E-08 | 1.149764751 | 0.352 | 0.152 | 0.000552411 | Tb |
| LAMP3+ DC | ENTPD1 | 8.17E-07 | 1.140353818 | 0.363 | 0.174 | 0.016077054 | Tb |
| LAMP3+ DC | SERINC5 | 5.67E-14 | 1.139329132 | 0.684 | 0.366 | 1.12E-09 | Tb |
| LAMP3+ DC | UBXN2A | 2.61E-10 | 1.135423806 | 0.627 | 0.351 | 5.13E-06 | Tb |
| LAMP3+ DC | FBXL8 | 1.05E-06 | 1.12543118 | 0.326 | 0.147 | 0.020612784 | Tb |
| LAMP3+ DC | ST6GAL1 | 8.53E-08 | 1.125066781 | 0.44 | 0.224 | 0.001679281 | Tb |
| LAMP3+ DC | FAF2 | 1.68E-07 | 1.118160949 | 0.326 | 0.135 | 0.003306961 | Tb |
| LAMP3+ DC | KDM5B | 4.30E-10 | 1.116724368 | 0.591 | 0.344 | 8.46E-06 | Tb |
| LAMP3+ DC | WDR63 | 3.47E-10 | 1.11210673 | 0.477 | 0.221 | 6.83E-06 | Tb |
| LAMP3+ DC | ARSB | 9.68E-08 | 1.110431517 | 0.513 | 0.295 | 0.001905297 | Tb |
| LAMP3+ DC | CAMK1D | 7.11E-09 | 1.108865988 | 0.71 | 0.548 | 0.000139906 | Tb |
| LAMP3+ DC | STIM2 | 1.24E-09 | 1.104379311 | 0.534 | 0.278 | 2.43E-05 | Tb |
| LAMP3+ DC | ZNF140 | 3.22E-08 | 1.099347778 | 0.295 | 0.106 | 0.000634188 | Tb |
| LAMP3+ DC | LIMCH1 | 3.70E-09 | 1.098074397 | 0.689 | 0.528 | 7.28E-05 | Tb |
| LAMP3+ DC | ZNF131 | 4.70E-09 | 1.097671927 | 0.503 | 0.253 | 9.25E-05 | Tb |

| | | | | | | | |
|---|---|---|---|---|---|---|---|
| LAMP3+ DC | CDK6 | 9.05E-16 | 1.095110426 | 0.819 | 0.538 | 1.78E-11 | Tb |
| LAMP3+ DC | KLHL28 | 3.77E-11 | 1.091758243 | 0.591 | 0.302 | 7.41E-07 | Tb |
| LAMP3+ DC | CD80 | 5.74E-18 | 1.091372518 | 0.865 | 0.654 | 1.13E-13 | Tb |
| LAMP3+ DC | EEF2K | 1.96E-07 | 1.090667455 | 0.415 | 0.199 | 0.003859482 | Tb |
| LAMP3+ DC | N4BP2 | 4.98E-07 | 1.088655553 | 0.513 | 0.312 | 0.009801676 | Tb |
| LAMP3+ DC | CEP83 | 1.11E-13 | 1.083013726 | 0.658 | 0.332 | 2.18E-09 | Tb |
| LAMP3+ DC | CALCRL | 1.52E-06 | 1.080212828 | 0.446 | 0.263 | 0.029859319 | Tb |
| LAMP3+ DC | LONRF1 | 5.83E-09 | 1.075702956 | 0.523 | 0.275 | 0.000114694 | Tb |
| LAMP3+ DC | KRIT1 | 4.50E-10 | 1.067506514 | 0.451 | 0.192 | 8.86E-06 | Tb |
| LAMP3+ DC | GBF1 | 3.10E-11 | 1.064252541 | 0.58 | 0.305 | 6.10E-07 | Tb |
| LAMP3+ DC | TP53 | 1.24E-07 | 1.063552226 | 0.269 | 0.098 | 0.002437611 | Tb |
| LAMP3+ DC | ARID2 | 6.84E-11 | 1.061803385 | 0.793 | 0.597 | 1.35E-06 | Tb |
| LAMP3+ DC | ADAP2 | 8.72E-07 | 1.057334921 | 0.373 | 0.184 | 0.017163964 | Tb |
| LAMP3+ DC | ST8SIA1 | 2.27E-07 | 1.055222767 | 0.487 | 0.268 | 0.004470137 | Tb |
| LAMP3+ DC | MFSD4B | 5.60E-07 | 1.052370773 | 0.383 | 0.187 | 0.011018698 | Tb |
| LAMP3+ DC | PNO1 | 4.25E-08 | 1.049855053 | 0.342 | 0.143 | 0.000835921 | Tb |
| LAMP3+ DC | PRRC2B | 1.26E-09 | 1.0492783 | 0.528 | 0.268 | 2.48E-05 | Tb |
| LAMP3+ DC | PHTF1 | 3.79E-07 | 1.048358934 | 0.358 | 0.174 | 0.00746289 | Tb |
| LAMP3+ DC | SFMBT2 | 5.34E-10 | 1.048147987 | 0.663 | 0.4 | 1.05E-05 | Tb |
| LAMP3+ DC | SEC23B | 4.21E-10 | 1.047521297 | 0.492 | 0.233 | 8.28E-06 | Tb |
| LAMP3+ DC | MT-ATP6 | 5.21E-26 | 1.046358495 | 0.99 | 0.921 | 1.02E-21 | Tb |
| LAMP3+ DC | MPP5 | 2.64E-07 | 1.045959812 | 0.295 | 0.115 | 0.005205343 | Tb |
| LAMP3+ DC | BCL11A | 2.92E-09 | 1.045149353 | 0.658 | 0.437 | 5.75E-05 | Tb |
| LAMP3+ DC | POGZ | 2.41E-08 | 1.0417235 | 0.528 | 0.285 | 0.000474551 | Tb |
| LAMP3+ DC | ERCC6L2 | 1.83E-06 | 1.038363229 | 0.295 | 0.128 | 0.036086663 | Tb |
| LAMP3+ DC | ZNF609 | 2.01E-07 | 1.03393557 | 0.472 | 0.273 | 0.003957506 | Tb |
| LAMP3+ DC | SLC43A2 | 3.63E-08 | 1.030680687 | 0.575 | 0.346 | 0.00071512 | Tb |
| LAMP3+ DC | EAF2 | 5.66E-08 | 1.018050918 | 0.549 | 0.319 | 0.001114387 | Tb |
| LAMP3+ DC | WDR59 | 4.69E-11 | 1.012421693 | 0.611 | 0.312 | 9.23E-07 | Tb |
| LAMP3+ DC | ZNF516 | 1.54E-08 | 1.012025277 | 0.492 | 0.265 | 0.000302819 | Tb |
| LAMP3+ DC | SAMD9L | 1.52E-11 | 1.011523447 | 0.71 | 0.423 | 2.99E-07 | Tb |
| LAMP3+ DC | NRF1 | 2.50E-06 | 1.006305539 | 0.389 | 0.201 | 0.049121092 | Tb |
| LAMP3+ DC | ASL | 1.50E-06 | 1.005367044 | 0.399 | 0.194 | 0.029587193 | Tb |
| LAMP3+ DC | MCC | 4.07E-07 | 0.999067826 | 0.326 | 0.145 | 0.00801358 | Tb |
| LAMP3+ DC | WIPF2 | 8.33E-07 | 0.99850769 | 0.373 | 0.184 | 0.016390637 | Tb |
| LAMP3+ DC | UBAC2 | 5.11E-08 | 0.997999088 | 0.648 | 0.445 | 0.001006746 | Tb |
| LAMP3+ DC | ZNF430 | 1.83E-06 | 0.997329992 | 0.373 | 0.179 | 0.036103639 | Tb |
| LAMP3+ DC | LRCH3 | 3.90E-12 | 0.995133072 | 0.772 | 0.506 | 7.67E-08 | Tb |
| LAMP3+ DC | SPINT2 | 1.51E-14 | 0.994562555 | 0.839 | 0.55 | 2.97E-10 | Tb |
| LAMP3+ DC | FBXL17 | 3.07E-07 | 0.992914039 | 0.528 | 0.312 | 0.006033759 | Tb |
| LAMP3+ DC | SLAMF1 | 1.37E-07 | 0.98785983 | 0.549 | 0.346 | 0.002689256 | Tb |
| LAMP3+ DC | TOGARAM1 | 4.16E-10 | 0.986723853 | 0.58 | 0.312 | 8.19E-06 | Tb |
| LAMP3+ DC | NSMCE2 | 7.11E-08 | 0.980392112 | 0.741 | 0.521 | 0.001398946 | Tb |
| LAMP3+ DC | CBWD6 | 5.75E-09 | 0.979049746 | 0.347 | 0.133 | 0.000113095 | Tb |
| LAMP3+ DC | FBXO11 | 3.79E-10 | 0.978995574 | 0.834 | 0.649 | 7.45E-06 | Tb |
| LAMP3+ DC | SORBS1 | 2.02E-06 | 0.976530835 | 0.295 | 0.13 | 0.03980325 | Tb |
| LAMP3+ DC | SLC25A25 | 1.37E-11 | 0.971694891 | 0.679 | 0.364 | 2.70E-07 | Tb |
| LAMP3+ DC | ARL13B | 1.74E-06 | 0.971044859 | 0.326 | 0.152 | 0.034344542 | Tb |
| LAMP3+ DC | ABTB2 | 8.27E-11 | 0.967875827 | 0.819 | 0.609 | 1.63E-06 | Tb |
| LAMP3+ DC | SLC1A4 | 1.70E-07 | 0.967772196 | 0.301 | 0.118 | 0.003352806 | Tb |
| LAMP3+ DC | RABGAP1L | 1.95E-12 | 0.967738137 | 0.87 | 0.666 | 3.84E-08 | Tb |
| LAMP3+ DC | CBLB | 8.83E-17 | 0.965903141 | 0.777 | 0.435 | 1.74E-12 | Tb |
| LAMP3+ DC | DUSP16 | 5.11E-09 | 0.965349775 | 0.611 | 0.346 | 0.000100648 | Tb |
| LAMP3+ DC | INO80 | 1.77E-07 | 0.965187205 | 0.523 | 0.287 | 0.003490379 | Tb |
| LAMP3+ DC | GEN1 | 3.85E-08 | 0.963644271 | 0.28 | 0.098 | 0.000757082 | Tb |
| LAMP3+ DC | AEBP2 | 1.10E-07 | 0.962807134 | 0.617 | 0.396 | 0.002167159 | Tb |

| | | | | | | | |
|---|---|---|---|---|---|---|---|
| LAMP3+ DC | INPP5F | 6.41E-08 | 0.958910507 | 0.539 | 0.297 | 0.0012613 | Tb |
| LAMP3+ DC | CRK | 2.02E-07 | 0.958235719 | 0.492 | 0.278 | 0.003980258 | Tb |
| LAMP3+ DC | PSEN1 | 1.47E-10 | 0.957876948 | 0.663 | 0.369 | 2.89E-06 | Tb |
| LAMP3+ DC | TPD52 | 1.03E-07 | 0.956743209 | 0.492 | 0.265 | 0.002024314 | Tb |
| LAMP3+ DC | ALG2 | 6.81E-08 | 0.954921924 | 0.383 | 0.17 | 0.001340353 | Tb |
| LAMP3+ DC | MSL2 | 6.07E-09 | 0.953227576 | 0.373 | 0.155 | 0.000119563 | Tb |
| LAMP3+ DC | ZNF10 | 7.83E-07 | 0.952393548 | 0.358 | 0.165 | 0.015411865 | Tb |
| LAMP3+ DC | HOMER2 | 2.33E-08 | 0.951354432 | 0.425 | 0.199 | 0.00045946 | Tb |
| LAMP3+ DC | FLT3 | 2.52E-09 | 0.950662309 | 0.746 | 0.521 | 4.96E-05 | Tb |
| LAMP3+ DC | SPRED2 | 3.90E-17 | 0.950606264 | 0.912 | 0.683 | 7.67E-13 | Tb |
| LAMP3+ DC | PSMD6-AS2 | 2.24E-06 | 0.949754666 | 0.254 | 0.101 | 0.044165473 | Tb |
| LAMP3+ DC | NCR3LG1 | 8.71E-07 | 0.949045931 | 0.332 | 0.15 | 0.017145977 | Tb |
| LAMP3+ DC | DLEU2 | 2.40E-13 | 0.946574431 | 0.87 | 0.695 | 4.72E-09 | Tb |
| LAMP3+ DC | PIP5K1A | 2.87E-10 | 0.945326447 | 0.523 | 0.248 | 5.65E-06 | Tb |
| LAMP3+ DC | KLF7 | 2.55E-11 | 0.94524469 | 0.772 | 0.538 | 5.02E-07 | Tb |
| LAMP3+ DC | KANSL1L | 3.08E-08 | 0.943702129 | 0.482 | 0.243 | 0.000606349 | Tb |
| LAMP3+ DC | EPB41L2 | 9.34E-08 | 0.942799553 | 0.689 | 0.484 | 0.001837854 | Tb |
| LAMP3+ DC | PSIP1 | 6.17E-08 | 0.940478078 | 0.58 | 0.346 | 0.00121537 | Tb |
| LAMP3+ DC | ACER3 | 9.01E-07 | 0.939002276 | 0.549 | 0.359 | 0.017732958 | Tb |
| LAMP3+ DC | SH3KBP1 | 1.07E-11 | 0.938636887 | 0.829 | 0.585 | 2.11E-07 | Tb |
| LAMP3+ DC | DIP2B | 1.55E-08 | 0.934580279 | 0.461 | 0.236 | 0.000305585 | Tb |
| LAMP3+ DC | GAB2 | 2.97E-10 | 0.933197195 | 0.736 | 0.479 | 5.84E-06 | Tb |
| LAMP3+ DC | GALNT7 | 5.10E-07 | 0.933183223 | 0.415 | 0.214 | 0.010047876 | Tb |
| LAMP3+ DC | PITPNA | 4.12E-12 | 0.932338723 | 0.824 | 0.631 | 8.11E-08 | Tb |
| LAMP3+ DC | RAB30 | 4.32E-11 | 0.931974301 | 0.788 | 0.555 | 8.50E-07 | Tb |
| LAMP3+ DC | SMYD4 | 8.33E-07 | 0.930734752 | 0.264 | 0.106 | 0.016389982 | Tb |
| LAMP3+ DC | PRKCA | 2.22E-08 | 0.925361357 | 0.508 | 0.27 | 0.000436308 | Tb |
| LAMP3+ DC | ING5 | 1.63E-06 | 0.924165667 | 0.254 | 0.101 | 0.03201084 | Tb |
| LAMP3+ DC | XPO1 | 9.01E-11 | 0.92325118 | 0.725 | 0.437 | 1.77E-06 | Tb |
| LAMP3+ DC | UIMC1 | 1.03E-07 | 0.922545385 | 0.456 | 0.229 | 0.002020835 | Tb |
| LAMP3+ DC | NR2C2 | 3.00E-07 | 0.921008891 | 0.389 | 0.187 | 0.005914822 | Tb |
| LAMP3+ DC | SH2B3 | 2.96E-12 | 0.913833496 | 0.772 | 0.516 | 5.82E-08 | Tb |
| LAMP3+ DC | CPSF2 | 1.69E-06 | 0.910040784 | 0.301 | 0.13 | 0.033173545 | Tb |
| LAMP3+ DC | PLEKHG2 | 1.23E-07 | 0.909364421 | 0.508 | 0.31 | 0.00241427 | Tb |
| LAMP3+ DC | EZH2 | 1.63E-09 | 0.908360436 | 0.845 | 0.604 | 3.21E-05 | Tb |
| LAMP3+ DC | PDCD1LG2 | 2.88E-12 | 0.904715618 | 0.803 | 0.514 | 5.67E-08 | Tb |
| LAMP3+ DC | TSPOAP1-AS1 | 1.90E-10 | 0.904033187 | 0.549 | 0.273 | 3.73E-06 | Tb |
| LAMP3+ DC | SLC25A43 | 1.23E-06 | 0.902659751 | 0.321 | 0.145 | 0.024304964 | Tb |
| LAMP3+ DC | GRAMD1B | 4.65E-09 | 0.90071963 | 0.534 | 0.278 | 9.14E-05 | Tb |
| LAMP3+ DC | ATR | 7.01E-07 | 0.900292171 | 0.383 | 0.189 | 0.013799062 | Tb |
| LAMP3+ DC | THUMPD3-AS1 | 4.29E-11 | 0.897393805 | 0.627 | 0.332 | 8.45E-07 | Tb |
| LAMP3+ DC | GPR157 | 1.04E-11 | 0.892933236 | 0.912 | 0.767 | 2.05E-07 | Tb |
| LAMP3+ DC | FOXP1 | 1.09E-13 | 0.892438636 | 0.896 | 0.801 | 2.15E-09 | Tb |
| LAMP3+ DC | OGDH | 1.85E-07 | 0.891369824 | 0.642 | 0.459 | 0.003646183 | Tb |
| LAMP3+ DC | SLMAP | 3.73E-08 | 0.890597482 | 0.415 | 0.192 | 0.000734415 | Tb |
| LAMP3+ DC | DOCK8 | 7.61E-08 | 0.886568827 | 0.767 | 0.622 | 0.001497986 | Tb |
| LAMP3+ DC | TRAPPC9 | 3.98E-07 | 0.885540724 | 0.534 | 0.312 | 0.007839283 | Tb |
| LAMP3+ DC | STK38L | 4.46E-10 | 0.876833064 | 0.679 | 0.405 | 8.77E-06 | Tb |
| LAMP3+ DC | SP4 | 7.74E-07 | 0.876305739 | 0.295 | 0.123 | 0.015241141 | Tb |
| LAMP3+ DC | PICALM | 5.85E-16 | 0.874589451 | 0.912 | 0.681 | 1.15E-11 | Tb |
| LAMP3+ DC | RIPOR31 | 1.93E-07 | 0.873024322 | 0.539 | 0.297 | 0.003796163 | Tb |
| LAMP3+ DC | TMCC1 | 3.36E-07 | 0.869947572 | 0.497 | 0.283 | 0.006620904 | Tb |
| LAMP3+ DC | WDFY4 | 2.52E-06 | 0.869670494 | 0.591 | 0.383 | 0.049640933 | Tb |
| LAMP3+ DC | PDE7A | 3.57E-08 | 0.866645503 | 0.699 | 0.467 | 0.000702949 | Tb |
| LAMP3+ DC | SINHCAF | 1.66E-13 | 0.866312884 | 0.87 | 0.754 | 3.27E-09 | Tb |
| LAMP3+ DC | MAML1 | 4.00E-07 | 0.859229485 | 0.487 | 0.273 | 0.007874963 | Tb |

| | | | | | | | |
|---|---|---|---|---|---|---|---|
| LAMP3+ DC | CD86 | 3.41E-13 | 0.85665487 | 0.917 | 0.803 | 6.71E-09 | Tb |
| LAMP3+ DC | ATP11B | 1.03E-06 | 0.856207874 | 0.637 | 0.408 | 0.020204801 | Tb |
| LAMP3+ DC | PRRG4 | 1.93E-08 | 0.854522792 | 0.663 | 0.425 | 0.000380015 | Tb |
| LAMP3+ DC | UBE2H | 9.25E-10 | 0.85433282 | 0.762 | 0.56 | 1.82E-05 | Tb |
| LAMP3+ DC | HLX | 1.44E-09 | 0.853771661 | 0.451 | 0.197 | 2.84E-05 | Tb |
| LAMP3+ DC | SMURF1 | 1.12E-09 | 0.852789571 | 0.617 | 0.361 | 2.21E-05 | Tb |
| LAMP3+ DC | ZFPM1 | 5.48E-08 | 0.846910506 | 0.337 | 0.133 | 0.001078017 | Tb |
| LAMP3+ DC | MT-CO3 | 4.36E-18 | 0.845075824 | 0.979 | 0.946 | 8.58E-14 | Tb |
| LAMP3+ DC | TBL1XR1 | 2.09E-09 | 0.84379009 | 0.725 | 0.489 | 4.11E-05 | Tb |
| LAMP3+ DC | STX5 | 4.90E-11 | 0.84136656 | 0.591 | 0.297 | 9.64E-07 | Tb |
| LAMP3+ DC | RABGEF1 | 7.47E-09 | 0.840182923 | 0.663 | 0.43 | 0.000147121 | Tb |
| LAMP3+ DC | YEATS2 | 1.38E-11 | 0.837019488 | 0.44 | 0.17 | 2.72E-07 | Tb |
| LAMP3+ DC | RSPRY1 | 1.60E-06 | 0.834626407 | 0.389 | 0.194 | 0.03158335 | Tb |
| LAMP3+ DC | DOCK10 | 1.79E-14 | 0.832705848 | 0.933 | 0.789 | 3.52E-10 | Tb |
| LAMP3+ DC | TMCC2 | 1.88E-06 | 0.832704977 | 0.42 | 0.231 | 0.036952639 | Tb |
| LAMP3+ DC | RAD54L2 | 4.87E-07 | 0.829134936 | 0.29 | 0.113 | 0.009581561 | Tb |
| LAMP3+ DC | HSH2D | 6.78E-07 | 0.82732232 | 0.461 | 0.26 | 0.013346031 | Tb |
| LAMP3+ DC | RAP2A | 4.94E-07 | 0.827042168 | 0.689 | 0.499 | 0.009731838 | Tb |
| LAMP3+ DC | TGFBR1 | 9.84E-08 | 0.826530337 | 0.674 | 0.42 | 0.00193651 | Tb |
| LAMP3+ DC | GRAMD1A | 4.34E-07 | 0.826295388 | 0.731 | 0.595 | 0.008537161 | Tb |
| LAMP3+ DC | LUC7L2 | 3.05E-07 | 0.826272514 | 0.679 | 0.467 | 0.006009861 | Tb |
| LAMP3+ DC | ELOVL7 | 1.03E-07 | 0.824647537 | 0.539 | 0.297 | 0.002022592 | Tb |
| LAMP3+ DC | MAP3K14 | 5.49E-09 | 0.821000603 | 0.782 | 0.555 | 0.000108026 | Tb |
| LAMP3+ DC | BTLA | 1.19E-08 | 0.819699082 | 0.321 | 0.118 | 0.00023415 | Tb |
| LAMP3+ DC | SRGAP2 | 1.95E-11 | 0.819160695 | 0.736 | 0.425 | 3.85E-07 | Tb |
| LAMP3+ DC | ASXL2 | 2.17E-07 | 0.819062332 | 0.394 | 0.189 | 0.004267693 | Tb |
| LAMP3+ DC | ST3GAL2 | 7.62E-09 | 0.818203976 | 0.534 | 0.28 | 0.000150072 | Tb |
| LAMP3+ DC | FAM13B | 5.46E-08 | 0.817910779 | 0.601 | 0.361 | 0.001075269 | Tb |
| LAMP3+ DC | PAN3 | 3.74E-08 | 0.817909819 | 0.793 | 0.568 | 0.000735602 | Tb |
| LAMP3+ DC | MED27 | 4.84E-07 | 0.817001934 | 0.352 | 0.162 | 0.009534304 | Tb |
| LAMP3+ DC | ITFG1 | 7.93E-09 | 0.816934822 | 0.72 | 0.484 | 0.000156161 | Tb |
| LAMP3+ DC | NT5C2 | 6.63E-08 | 0.81686196 | 0.539 | 0.295 | 0.00130457 | Tb |
| LAMP3+ DC | AP2B1 | 6.94E-07 | 0.816243054 | 0.503 | 0.278 | 0.013660826 | Tb |
| LAMP3+ DC | NCOA3 | 1.46E-08 | 0.814272545 | 0.751 | 0.624 | 0.000286417 | Tb |
| LAMP3+ DC | PPP3CC | 6.47E-09 | 0.81181749 | 0.606 | 0.334 | 0.000127369 | Tb |
| LAMP3+ DC | ASMTL | 2.51E-08 | 0.811481177 | 0.663 | 0.423 | 0.000494222 | Tb |
| LAMP3+ DC | EBF4 | 1.73E-06 | 0.811221594 | 0.425 | 0.221 | 0.034029796 | Tb |
| LAMP3+ DC | SIPA1L1 | 2.13E-07 | 0.808991302 | 0.865 | 0.713 | 0.004187301 | Tb |
| LAMP3+ DC | EDARADD | 2.38E-08 | 0.808108792 | 0.767 | 0.553 | 0.000469 | Tb |
| LAMP3+ DC | SPECC1 | 4.41E-08 | 0.807462952 | 0.803 | 0.619 | 0.000867903 | Tb |
| LAMP3+ DC | ASAP1 | 7.00E-12 | 0.806692638 | 0.922 | 0.843 | 1.38E-07 | Tb |
| LAMP3+ DC | SRGAP2B | 4.98E-09 | 0.805674821 | 0.591 | 0.312 | 9.80E-05 | Tb |
| LAMP3+ DC | IL1R1 | 1.68E-06 | 0.803934252 | 0.425 | 0.229 | 0.032972782 | Tb |
| LAMP3+ DC | GPCPD1 | 1.68E-06 | 0.803637433 | 0.642 | 0.469 | 0.033156055 | Tb |
| LAMP3+ DC | ZCCHC2 | 4.37E-08 | 0.802396583 | 0.404 | 0.184 | 0.000859764 | Tb |
| LAMP3+ DC | SSH1 | 1.31E-06 | 0.801810425 | 0.554 | 0.327 | 0.025825527 | Tb |
| LAMP3+ DC | SENP6 | 1.20E-07 | 0.80159971 | 0.679 | 0.437 | 0.002357788 | Tb |
| LAMP3+ DC | CGAS | 4.07E-08 | 0.801044996 | 0.57 | 0.349 | 0.000802026 | Tb |
| LAMP3+ DC | MT-ND4 | 5.78E-21 | 0.800306718 | 0.979 | 0.953 | 1.14E-16 | Tb |
| LAMP3+ DC | DAPP1 | 2.24E-20 | 0.798718068 | 0.969 | 0.872 | 4.41E-16 | Tb |
| LAMP3+ DC | POLR2J3 | 1.48E-06 | 0.798552301 | 0.415 | 0.219 | 0.029101264 | Tb |
| LAMP3+ DC | NFAT5 | 1.02E-10 | 0.796357246 | 0.917 | 0.781 | 2.00E-06 | Tb |
| LAMP3+ DC | NOP58 | 5.11E-07 | 0.795601818 | 0.694 | 0.489 | 0.010063314 | Tb |
| LAMP3+ DC | UBE2E1 | 6.71E-09 | 0.793907781 | 0.777 | 0.617 | 0.000132037 | Tb |
| LAMP3+ DC | FARP2 | 7.46E-08 | 0.793003148 | 0.705 | 0.491 | 0.001467823 | Tb |
| LAMP3+ DC | WASL | 8.01E-07 | 0.79198997 | 0.508 | 0.283 | 0.015762786 | Tb |

| | | | | | | | |
|---|---|---|---|---|---|---|---|
| LAMP3+ DC | MSRB1 | 4.62E-07 | 0.791241037 | 0.451 | 0.231 | 0.009100564 | Tb |
| LAMP3+ DC | MT-CYB | 3.68E-14 | 0.790914794 | 0.959 | 0.931 | 7.24E-10 | Tb |
| LAMP3+ DC | PITPNC1 | 1.01E-06 | 0.790626702 | 0.658 | 0.435 | 0.019969345 | Tb |
| LAMP3+ DC | FNIP1 | 2.13E-08 | 0.790610837 | 0.653 | 0.398 | 0.000418499 | Tb |
| LAMP3+ DC | RGL1 | 1.90E-06 | 0.78795616 | 0.508 | 0.305 | 0.037302831 | Tb |
| LAMP3+ DC | RIC1 | 3.55E-09 | 0.78732671 | 0.762 | 0.563 | 6.99E-05 | Tb |
| LAMP3+ DC | TFCP2 | 4.40E-08 | 0.780320519 | 0.637 | 0.391 | 0.000865214 | Tb |
| LAMP3+ DC | RBM6 | 3.22E-08 | 0.77974262 | 0.648 | 0.398 | 0.00063458 | Tb |
| LAMP3+ DC | GNG2 | 6.62E-08 | 0.778730771 | 0.461 | 0.236 | 0.001303672 | Tb |
| LAMP3+ DC | LRRK1 | 1.18E-09 | 0.777416144 | 0.865 | 0.673 | 2.32E-05 | Tb |
| LAMP3+ DC | LIMS1 | 3.03E-09 | 0.77616253 | 0.803 | 0.538 | 5.96E-05 | Tb |
| LAMP3+ DC | HIVEP2 | 3.43E-08 | 0.775484309 | 0.834 | 0.693 | 0.000675363 | Tb |
| LAMP3+ DC | FOXO3 | 1.46E-07 | 0.773586098 | 0.622 | 0.383 | 0.002877469 | Tb |
| LAMP3+ DC | SIK2 | 3.12E-07 | 0.771683661 | 0.56 | 0.332 | 0.006142919 | Tb |
| LAMP3+ DC | TRNT1 | 7.41E-07 | 0.770148458 | 0.29 | 0.115 | 0.014580113 | Tb |
| LAMP3+ DC | SPEN | 1.63E-06 | 0.769831076 | 0.663 | 0.457 | 0.032097552 | Tb |
| LAMP3+ DC | AFF1 | 2.53E-08 | 0.768732379 | 0.741 | 0.496 | 0.000497427 | Tb |
| LAMP3+ DC | CPB2-AS1 | 4.85E-08 | 0.765177409 | 0.472 | 0.231 | 0.000953784 | Tb |
| LAMP3+ DC | ELMSAN1 | 8.06E-09 | 0.7646639 | 0.596 | 0.342 | 0.000158576 | Tb |
| LAMP3+ DC | GNPTAB | 2.59E-07 | 0.763765119 | 0.544 | 0.319 | 0.005092766 | Tb |
| LAMP3+ DC | MTMR6 | 1.80E-06 | 0.761404606 | 0.389 | 0.184 | 0.03551794 | Tb |
| LAMP3+ DC | CSTF3 | 1.40E-06 | 0.760405071 | 0.311 | 0.138 | 0.027501433 | Tb |
| LAMP3+ DC | ZHX2 | 1.15E-11 | 0.76009243 | 0.85 | 0.624 | 2.27E-07 | Tb |
| LAMP3+ DC | SOS1 | 5.68E-08 | 0.759459677 | 0.668 | 0.425 | 0.001118874 | Tb |
| LAMP3+ DC | MYO9A | 2.08E-06 | 0.759245191 | 0.58 | 0.378 | 0.040915715 | Tb |
| LAMP3+ DC | BCL6 | 3.23E-07 | 0.758480959 | 0.642 | 0.403 | 0.006367328 | Tb |
| LAMP3+ DC | KIF1B | 1.23E-07 | 0.756329919 | 0.565 | 0.332 | 0.002422916 | Tb |
| LAMP3+ DC | MALT1 | 5.39E-10 | 0.754664745 | 0.855 | 0.735 | 1.06E-05 | Tb |
| LAMP3+ DC | ADAMTSL4-AS | 2.37E-07 | 0.754614878 | 0.554 | 0.322 | 0.004674653 | Tb |
| LAMP3+ DC | CTNNA1 | 1.07E-07 | 0.754608962 | 0.705 | 0.491 | 0.002102919 | Tb |
| LAMP3+ DC | NFIL3 | 7.22E-07 | 0.751647066 | 0.663 | 0.455 | 0.014203832 | Tb |
| LAMP3+ DC | RNF19B | 1.01E-06 | 0.749131768 | 0.725 | 0.533 | 0.01992289 | Tb |
| LAMP3+ DC | TAB2 | 1.34E-10 | 0.748958528 | 0.824 | 0.582 | 2.65E-06 | Tb |
| LAMP3+ DC | XPO6 | 5.49E-08 | 0.747786358 | 0.534 | 0.29 | 0.00108089 | Tb |
| LAMP3+ DC | TRIM38 | 1.81E-10 | 0.747235195 | 0.715 | 0.43 | 3.56E-06 | Tb |
| LAMP3+ DC | P2RY10 | 1.43E-06 | 0.745119841 | 0.554 | 0.351 | 0.028098559 | Tb |
| LAMP3+ DC | CFLAR-AS1 | 1.60E-08 | 0.744694689 | 0.575 | 0.3 | 0.000315355 | Tb |
| LAMP3+ DC | MYO1C | 1.48E-08 | 0.744549566 | 0.518 | 0.256 | 0.000291469 | Tb |
| LAMP3+ DC | SETD5 | 5.04E-09 | 0.743404968 | 0.668 | 0.393 | 9.92E-05 | Tb |
| LAMP3+ DC | ENTPD1-AS1 | 8.19E-07 | 0.742806017 | 0.358 | 0.17 | 0.016111993 | Tb |
| LAMP3+ DC | ITCH | 4.08E-09 | 0.739692474 | 0.58 | 0.312 | 8.04E-05 | Tb |
| LAMP3+ DC | GALC | 7.63E-08 | 0.73862542 | 0.482 | 0.248 | 0.001501606 | Tb |
| LAMP3+ DC | ASCC3 | 7.64E-07 | 0.738423331 | 0.57 | 0.342 | 0.015036368 | Tb |
| LAMP3+ DC | PPM1K | 7.71E-09 | 0.738304481 | 0.767 | 0.526 | 0.000151825 | Tb |
| LAMP3+ DC | ARIH1 | 2.18E-10 | 0.738126773 | 0.834 | 0.654 | 4.28E-06 | Tb |
| LAMP3+ DC | CLASRP | 2.26E-06 | 0.738099312 | 0.591 | 0.354 | 0.044510855 | Tb |
| LAMP3+ DC | PPM1B | 9.97E-07 | 0.737772626 | 0.658 | 0.459 | 0.01962526 | Tb |
| LAMP3+ DC | CTC1 | 6.67E-07 | 0.735888443 | 0.295 | 0.12 | 0.013120998 | Tb |
| LAMP3+ DC | CLIP1 | 6.27E-07 | 0.735446891 | 0.492 | 0.283 | 0.012350897 | Tb |
| LAMP3+ DC | KDM6A | 3.25E-12 | 0.735419311 | 0.782 | 0.526 | 6.40E-08 | Tb |
| LAMP3+ DC | PAK1 | 4.39E-10 | 0.733615025 | 0.86 | 0.676 | 8.64E-06 | Tb |
| LAMP3+ DC | PARP8 | 7.43E-07 | 0.73240674 | 0.539 | 0.31 | 0.014635044 | Tb |
| LAMP3+ DC | CD58 | 3.62E-08 | 0.73216438 | 0.819 | 0.673 | 0.000711919 | Tb |
| LAMP3+ DC | TCF12 | 2.16E-09 | 0.731549394 | 0.756 | 0.518 | 4.26E-05 | Tb |
| LAMP3+ DC | ZMYM5 | 1.22E-06 | 0.728689719 | 0.528 | 0.297 | 0.023996746 | Tb |
| LAMP3+ DC | STAT6 | 3.51E-09 | 0.727780258 | 0.611 | 0.327 | 6.90E-05 | Tb |

| | | | | | | | |
|---|---|---|---|---|---|---|---|
| LAMP3+ DC | ITPR1 | 1.47E-07 | 0.724901725 | 0.596 | 0.356 | 0.002897725 | Tb |
| LAMP3+ DC | TCAIM | 1.15E-06 | 0.724466506 | 0.285 | 0.118 | 0.022594966 | Tb |
| LAMP3+ DC | SESN3 | 7.43E-07 | 0.722303232 | 0.689 | 0.479 | 0.014631188 | Tb |
| LAMP3+ DC | ESYT2 | 8.49E-11 | 0.719913301 | 0.85 | 0.676 | 1.67E-06 | Tb |
| LAMP3+ DC | TET3 | 4.49E-07 | 0.719234993 | 0.58 | 0.344 | 0.008846715 | Tb |
| LAMP3+ DC | WNK1 | 1.14E-10 | 0.719221444 | 0.85 | 0.661 | 2.25E-06 | Tb |
| LAMP3+ DC | TAOK1 | 2.29E-07 | 0.716197415 | 0.596 | 0.366 | 0.004514477 | Tb |
| LAMP3+ DC | NR6A1 | 2.89E-07 | 0.715644113 | 0.689 | 0.452 | 0.005695155 | Tb |
| LAMP3+ DC | BPTF | 1.28E-08 | 0.714324464 | 0.741 | 0.489 | 0.000251937 | Tb |
| LAMP3+ DC | ZEB1 | 8.81E-10 | 0.71430758 | 0.839 | 0.627 | 1.73E-05 | Tb |
| LAMP3+ DC | VPS8 | 1.34E-06 | 0.713712634 | 0.611 | 0.403 | 0.026453857 | Tb |
| LAMP3+ DC | UGCG | 2.20E-06 | 0.711962007 | 0.715 | 0.494 | 0.043212884 | Tb |
| LAMP3+ DC | KDM2A | 4.42E-07 | 0.710065834 | 0.85 | 0.661 | 0.008705327 | Tb |
| LAMP3+ DC | FAF1 | 7.21E-08 | 0.70969031 | 0.653 | 0.418 | 0.001420057 | Tb |
| LAMP3+ DC | IKZF1 | 2.10E-11 | 0.703305599 | 0.865 | 0.686 | 4.14E-07 | Tb |
| LAMP3+ DC | WDR7 | 1.05E-06 | 0.703057485 | 0.42 | 0.206 | 0.020735584 | Tb |
| LAMP3+ DC | SLC39A10 | 9.52E-07 | 0.701820671 | 0.347 | 0.16 | 0.018734365 | Tb |
| LAMP3+ DC | KDM2B | 8.77E-11 | 0.699297596 | 0.933 | 0.843 | 1.73E-06 | Tb |
| LAMP3+ DC | SLC33A1 | 1.61E-06 | 0.698227834 | 0.275 | 0.111 | 0.031627088 | Tb |
| LAMP3+ DC | APPBP2 | 1.02E-06 | 0.69549396 | 0.373 | 0.177 | 0.020038012 | Tb |
| LAMP3+ DC | AKNA | 1.63E-06 | 0.695089364 | 0.492 | 0.275 | 0.032154759 | Tb |
| LAMP3+ DC | FYN | 1.69E-08 | 0.688499442 | 0.684 | 0.437 | 0.000333406 | Tb |
| LAMP3+ DC | ZC3H7B | 5.02E-07 | 0.687233435 | 0.596 | 0.359 | 0.009877749 | Tb |
| LAMP3+ DC | TGIF2 | 1.15E-06 | 0.684992732 | 0.466 | 0.248 | 0.022682425 | Tb |
| LAMP3+ DC | R3HDM2 | 4.48E-08 | 0.684189257 | 0.71 | 0.469 | 0.000882565 | Tb |
| LAMP3+ DC | DENND4A | 2.48E-08 | 0.68266901 | 0.865 | 0.823 | 0.000488866 | Tb |
| LAMP3+ DC | AKT3 | 2.55E-07 | 0.681329466 | 0.699 | 0.457 | 0.00502208 | Tb |
| LAMP3+ DC | MTMR9 | 6.15E-07 | 0.681077032 | 0.451 | 0.236 | 0.012110229 | Tb |
| LAMP3+ DC | RABGAP1 | 3.19E-10 | 0.680553596 | 0.731 | 0.435 | 6.28E-06 | Tb |
| LAMP3+ DC | ABR | 2.49E-06 | 0.680530954 | 0.689 | 0.501 | 0.049017835 | Tb |
| LAMP3+ DC | ARID1A | 2.85E-07 | 0.680311505 | 0.591 | 0.356 | 0.005600683 | Tb |
| LAMP3+ DC | SZRD1 | 6.85E-08 | 0.676631269 | 0.591 | 0.322 | 0.001348088 | Tb |
| LAMP3+ DC | ADORA2A-AS1 | 1.07E-07 | 0.675038779 | 0.342 | 0.145 | 0.002098603 | Tb |
| LAMP3+ DC | ALPK1 | 1.85E-07 | 0.673077839 | 0.679 | 0.459 | 0.003639702 | Tb |
| LAMP3+ DC | GPR137B | 9.28E-09 | 0.671521307 | 0.922 | 0.828 | 0.000182697 | Tb |
| LAMP3+ DC | KLHL5 | 8.82E-07 | 0.670766868 | 0.508 | 0.287 | 0.017362193 | Tb |
| LAMP3+ DC | PDLIM5 | 1.25E-07 | 0.668895078 | 0.813 | 0.624 | 0.002453309 | Tb |
| LAMP3+ DC | CERT1 | 1.41E-06 | 0.667808158 | 0.591 | 0.388 | 0.027796303 | Tb |
| LAMP3+ DC | ELOVL5 | 9.86E-07 | 0.666924261 | 0.824 | 0.6 | 0.019408384 | Tb |
| LAMP3+ DC | KANSL1 | 3.13E-07 | 0.666011189 | 0.575 | 0.346 | 0.006157046 | Tb |
| LAMP3+ DC | FAM133B | 8.64E-09 | 0.663465031 | 0.632 | 0.354 | 0.000170103 | Tb |
| LAMP3+ DC | FNDC3B | 2.02E-08 | 0.65877791 | 0.865 | 0.747 | 0.00039856 | Tb |
| LAMP3+ DC | MOB3A | 1.71E-08 | 0.650984068 | 0.793 | 0.6 | 0.00033616 | Tb |
| LAMP3+ DC | P2RY8 | 1.38E-06 | 0.649192292 | 0.591 | 0.383 | 0.027093763 | Tb |
| LAMP3+ DC | TMEM39A | 2.72E-09 | 0.646990258 | 0.876 | 0.663 | 5.36E-05 | Tb |
| LAMP3+ DC | IGF2BP2 | 3.96E-09 | 0.645371308 | 0.658 | 0.393 | 7.79E-05 | Tb |
| LAMP3+ DC | STK10 | 7.59E-07 | 0.64370695 | 0.523 | 0.3 | 0.014947075 | Tb |
| LAMP3+ DC | SREBF2 | 1.76E-07 | 0.642845954 | 0.591 | 0.332 | 0.003473093 | Tb |
| LAMP3+ DC | IPMK | 1.36E-06 | 0.641717417 | 0.404 | 0.204 | 0.026695835 | Tb |
| LAMP3+ DC | UBE2W | 8.50E-07 | 0.639740845 | 0.746 | 0.545 | 0.016732409 | Tb |
| LAMP3+ DC | TENT4B | 1.99E-08 | 0.639569507 | 0.72 | 0.474 | 0.000392016 | Tb |
| LAMP3+ DC | ZNF260 | 1.22E-06 | 0.638726386 | 0.43 | 0.224 | 0.024023335 | Tb |
| LAMP3+ DC | MAPK8 | 2.91E-10 | 0.638331569 | 0.788 | 0.558 | 5.73E-06 | Tb |
| LAMP3+ DC | RUFY1 | 1.03E-06 | 0.636966372 | 0.56 | 0.332 | 0.020325064 | Tb |
| LAMP3+ DC | GSK3B | 2.16E-08 | 0.635658992 | 0.767 | 0.518 | 0.000424388 | Tb |
| LAMP3+ DC | RAD23B | 1.46E-07 | 0.634745002 | 0.591 | 0.342 | 0.002877041 | Tb |

| | | | | | | | |
|---|---|---|---|---|---|---|---|
| LAMP3+ DC | MT-CO1 | 6.76E-17 | 0.632441302 | 0.995 | 0.99 | 1.33E-12 | Tb |
| LAMP3+ DC | ZNFX1 | 7.96E-07 | 0.632418297 | 0.663 | 0.464 | 0.015672811 | Tb |
| LAMP3+ DC | CHP1 | 1.15E-06 | 0.628749969 | 0.622 | 0.41 | 0.022614657 | Tb |
| LAMP3+ DC | CIITA | 5.49E-07 | 0.628410417 | 0.803 | 0.661 | 0.01081136 | Tb |
| LAMP3+ DC | MREG | 1.93E-06 | 0.628303718 | 0.819 | 0.651 | 0.03794075 | Tb |
| LAMP3+ DC | SLC15A4 | 6.39E-07 | 0.627998675 | 0.632 | 0.396 | 0.012576688 | Tb |
| LAMP3+ DC | GATAD2B | 1.85E-08 | 0.627866152 | 0.642 | 0.388 | 0.000364562 | Tb |
| LAMP3+ DC | BMPR2 | 3.46E-07 | 0.624175009 | 0.513 | 0.295 | 0.006810742 | Tb |
| LAMP3+ DC | PTGIR | 6.94E-07 | 0.623360887 | 0.523 | 0.292 | 0.013666861 | Tb |
| LAMP3+ DC | ASH1L | 7.00E-07 | 0.622955193 | 0.86 | 0.705 | 0.013779536 | Tb |
| LAMP3+ DC | TANK | 5.02E-11 | 0.622591791 | 0.943 | 0.862 | 9.88E-07 | Tb |
| LAMP3+ DC | KCMF1 | 9.11E-07 | 0.622516945 | 0.679 | 0.452 | 0.017933592 | Tb |
| LAMP3+ DC | IKBKB | 1.62E-07 | 0.621619 | 0.415 | 0.189 | 0.00319324 | Tb |
| LAMP3+ DC | VMP1 | 1.05E-06 | 0.621345007 | 0.782 | 0.558 | 0.020714916 | Tb |
| LAMP3+ DC | CDC42SE2 | 1.39E-11 | 0.620221803 | 0.933 | 0.759 | 2.74E-07 | Tb |
| LAMP3+ DC | MBD2 | 7.48E-07 | 0.619283898 | 0.741 | 0.541 | 0.014716165 | Tb |
| LAMP3+ DC | EED | 1.08E-06 | 0.61827694 | 0.585 | 0.327 | 0.021350896 | Tb |
| LAMP3+ DC | WDR44 | 1.90E-06 | 0.616853914 | 0.539 | 0.307 | 0.037362877 | Tb |
| LAMP3+ DC | MAP3K13 | 2.42E-07 | 0.615284111 | 0.886 | 0.752 | 0.00476914 | Tb |
| LAMP3+ DC | ARFGEF2 | 1.42E-08 | 0.612243395 | 0.518 | 0.258 | 0.000280471 | Tb |
| LAMP3+ DC | LCOR | 2.04E-06 | 0.611402259 | 0.699 | 0.479 | 0.04024033 | Tb |
| LAMP3+ DC | SRSF4 | 4.92E-07 | 0.610789032 | 0.751 | 0.486 | 0.009687661 | Tb |
| LAMP3+ DC | CSF2RA | 2.79E-11 | 0.608022905 | 0.953 | 0.902 | 5.50E-07 | Tb |
| LAMP3+ DC | RFTN1 | 3.12E-13 | 0.605611358 | 0.979 | 0.889 | 6.15E-09 | Tb |
| LAMP3+ DC | SLC36A4 | 2.02E-06 | 0.601390768 | 0.43 | 0.224 | 0.039783137 | Tb |
| LAMP3+ DC | RUBCN | 2.20E-07 | 0.601290563 | 0.694 | 0.486 | 0.004322407 | Tb |
| LAMP3+ DC | RAB4B | 1.07E-06 | 0.600357738 | 0.508 | 0.28 | 0.021138611 | Tb |
| LAMP3+ DC | NR4A3 | 2.49E-09 | 0.597345078 | 0.907 | 0.767 | 4.91E-05 | Tb |
| LAMP3+ DC | ARMH3 | 2.25E-06 | 0.595129738 | 0.544 | 0.324 | 0.044326415 | Tb |
| LAMP3+ DC | STAT5A | 9.40E-07 | 0.590108513 | 0.731 | 0.536 | 0.018505062 | Tb |
| LAMP3+ DC | KPNA4 | 1.00E-07 | 0.589593048 | 0.725 | 0.477 | 0.001974313 | Tb |
| LAMP3+ DC | KMT2E | 5.84E-09 | 0.589187949 | 0.86 | 0.715 | 0.000114893 | Tb |
| LAMP3+ DC | PTPRJ | 1.61E-08 | 0.589124479 | 0.845 | 0.661 | 0.000317555 | Tb |
| LAMP3+ DC | RAPGEF1 | 2.78E-12 | 0.589037683 | 0.948 | 0.845 | 5.48E-08 | Tb |
| LAMP3+ DC | FCHSD2 | 1.33E-07 | 0.588162004 | 0.912 | 0.789 | 0.002625672 | Tb |
| LAMP3+ DC | TMEM131 | 3.08E-08 | 0.587086664 | 0.891 | 0.757 | 0.000606332 | Tb |
| LAMP3+ DC | MRTFA | 1.75E-09 | 0.582955126 | 0.922 | 0.791 | 3.45E-05 | Tb |
| LAMP3+ DC | LPP | 5.11E-08 | 0.581350744 | 0.855 | 0.617 | 0.00100531 | Tb |
| LAMP3+ DC | LRIF1 | 6.13E-09 | 0.580483052 | 0.772 | 0.506 | 0.000120755 | Tb |
| LAMP3+ DC | GNB1 | 1.21E-08 | 0.572679888 | 0.886 | 0.69 | 0.000238329 | Tb |
| LAMP3+ DC | LAMP3 | 1.65E-13 | 0.571500147 | 0.984 | 0.943 | 3.25E-09 | Tb |
| LAMP3+ DC | CELF2 | 1.17E-10 | 0.569727122 | 0.964 | 0.848 | 2.30E-06 | Tb |
| LAMP3+ DC | GNAQ | 2.25E-07 | 0.568501424 | 0.855 | 0.676 | 0.004420604 | Tb |
| LAMP3+ DC | MAP4 | 1.05E-07 | 0.568188088 | 0.839 | 0.646 | 0.002067099 | Tb |
| LAMP3+ DC | ATXN7 | 3.48E-08 | 0.566908447 | 0.731 | 0.494 | 0.000684293 | Tb |
| LAMP3+ DC | RNF121 | 1.55E-06 | 0.565650614 | 0.363 | 0.172 | 0.03056185 | Tb |
| LAMP3+ DC | TOMM34 | 8.10E-08 | 0.564452198 | 0.715 | 0.445 | 0.001595242 | Tb |
| LAMP3+ DC | ARHGEF2 | 3.63E-07 | 0.563849878 | 0.72 | 0.531 | 0.007137085 | Tb |
| LAMP3+ DC | TMEM63B | 5.27E-07 | 0.557894291 | 0.342 | 0.152 | 0.01037009 | Tb |
| LAMP3+ DC | SNX20 | 1.92E-07 | 0.55617801 | 0.508 | 0.268 | 0.003787686 | Tb |
| LAMP3+ DC | STAT3 | 4.25E-07 | 0.55548267 | 0.881 | 0.715 | 0.008367581 | Tb |
| LAMP3+ DC | PDE4B | 1.74E-07 | 0.55477334 | 0.886 | 0.722 | 0.003422112 | Tb |
| LAMP3+ DC | RUNX1 | 1.73E-06 | 0.553626346 | 0.715 | 0.523 | 0.034132152 | Tb |
| LAMP3+ DC | NIBAN1 | 4.01E-08 | 0.551259016 | 0.881 | 0.727 | 0.00078857 | Tb |
| LAMP3+ DC | MYH9 | 2.70E-09 | 0.551000359 | 0.86 | 0.663 | 5.31E-05 | Tb |
| LAMP3+ DC | FBXO34 | 2.00E-09 | 0.549336688 | 0.725 | 0.442 | 3.94E-05 | Tb |

| | | | | | | | |
|---|---|---|---|---|---|---|---|
| LAMP3+ DC | TBCEL | 8.77E-07 | 0.547305141 | 0.337 | 0.152 | 0.017255004 | Tb |
| LAMP3+ DC | ANKRD33B | 9.82E-09 | 0.546727643 | 0.922 | 0.776 | 0.000193216 | Tb |
| LAMP3+ DC | OGA | 3.64E-07 | 0.545116922 | 0.689 | 0.486 | 0.007170123 | Tb |
| LAMP3+ DC | MEF2A | 3.42E-07 | 0.542090311 | 0.839 | 0.658 | 0.006738598 | Tb |
| LAMP3+ DC | KDM4C | 1.20E-06 | 0.541991045 | 0.824 | 0.609 | 0.023701432 | Tb |
| LAMP3+ DC | MAP4K4 | 4.81E-11 | 0.540945189 | 0.943 | 0.771 | 9.47E-07 | Tb |
| LAMP3+ DC | MT-ND3 | 1.87E-15 | 0.54045462 | 0.99 | 0.953 | 3.68E-11 | Tb |
| LAMP3+ DC | GRK3 | 1.66E-07 | 0.540322559 | 0.912 | 0.776 | 0.003266016 | Tb |
| LAMP3+ DC | BACH1 | 1.70E-06 | 0.538197261 | 0.86 | 0.73 | 0.03337582 | Tb |
| LAMP3+ DC | CERS6 | 1.71E-08 | 0.537829771 | 0.943 | 0.892 | 0.000336084 | Tb |
| LAMP3+ DC | MT-CO2 | 1.74E-16 | 0.537359219 | 0.99 | 0.978 | 3.43E-12 | Tb |
| LAMP3+ DC | CAND1 | 6.90E-07 | 0.536683829 | 0.632 | 0.381 | 0.013585329 | Tb |
| LAMP3+ DC | USP36 | 5.93E-07 | 0.534043628 | 0.56 | 0.3 | 0.011675399 | Tb |
| LAMP3+ DC | MED13L | 9.70E-11 | 0.533852282 | 0.959 | 0.843 | 1.91E-06 | Tb |
| LAMP3+ DC | ANAPC10 | 1.38E-06 | 0.532515552 | 0.482 | 0.256 | 0.027163163 | Tb |
| LAMP3+ DC | N4BP1 | 1.44E-07 | 0.530362643 | 0.839 | 0.614 | 0.002833652 | Tb |
| LAMP3+ DC | TTYH2 | 1.48E-07 | 0.529300799 | 0.819 | 0.604 | 0.002907892 | Tb |
| LAMP3+ DC | HNRNPC | 3.79E-12 | 0.523461775 | 0.948 | 0.88 | 7.46E-08 | Tb |
| LAMP3+ DC | RYBP | 2.93E-07 | 0.523109188 | 0.767 | 0.511 | 0.005775205 | Tb |
| LAMP3+ DC | IDO2 | 1.12E-06 | 0.522428428 | 0.788 | 0.575 | 0.022096756 | Tb |
| LAMP3+ DC | ZNF217 | 4.04E-08 | 0.520527155 | 0.606 | 0.346 | 0.000794419 | Tb |
| LAMP3+ DC | DSE | 5.62E-08 | 0.520464666 | 0.715 | 0.445 | 0.001106019 | Tb |
| LAMP3+ DC | RAB10 | 2.15E-07 | 0.518560173 | 0.953 | 0.853 | 0.004239332 | Tb |
| LAMP3+ DC | GPR132 | 6.50E-07 | 0.515441148 | 0.865 | 0.71 | 0.012801368 | Tb |
| LAMP3+ DC | FTX | 4.82E-07 | 0.513383971 | 0.824 | 0.607 | 0.009480878 | Tb |
| LAMP3+ DC | SWAP70 | 1.91E-07 | 0.502704264 | 0.839 | 0.668 | 0.003750978 | Tb |
| **Disease-associated genes in pDC** | | | | | | | |
| pDC | HSPH1 | 5.21E-07 | 0.860962257 | 1 | 0.826 | 0.01025504 | Lep-L |
| pDC | HSPA1A | 1.78E-06 | 1.152788383 | 1 | 0.847 | 0.034994868 | Lep-L |
| pDC | FKBP4 | 1.03E-06 | 1.310489061 | 0.808 | 0.389 | 0.020315879 | Lep-L |
| pDC | CD83 | 9.34E-10 | 1.319655524 | 1 | 0.901 | 1.84E-05 | Lep-L |
| pDC | HSP90AA1 | 5.75E-09 | 1.34579683 | 1 | 0.989 | 0.000113113 | Lep-L |
| pDC | HSPD1 | 4.41E-09 | 1.385077777 | 1 | 0.847 | 8.68E-05 | Lep-L |
| pDC | HSP90AB1 | 2.83E-10 | 1.411490616 | 1 | 0.977 | 5.57E-06 | Lep-L |
| pDC | EZR | 8.34E-09 | 1.429332618 | 1 | 0.941 | 0.000164243 | Lep-L |
| pDC | HSPA6 | 6.82E-08 | 1.619861216 | 0.615 | 0.184 | 0.00134182 | Lep-L |
| pDC | USP9Y | 1.15E-06 | 1.670794812 | 0.654 | 0.227 | 0.022541492 | Lep-L |
| pDC | HSPB1 | 2.62E-09 | 1.703113607 | 0.962 | 0.564 | 5.16E-05 | Lep-L |
| pDC | TSPYL2 | 4.08E-07 | 1.739756381 | 0.923 | 0.61 | 0.008039678 | Lep-L |
| pDC | CHORDC1 | 3.69E-09 | 1.778012397 | 0.923 | 0.595 | 7.26E-05 | Lep-L |
| pDC | BMT2 | 1.40E-06 | 2.391099419 | 0.462 | 0.137 | 0.027518676 | Lep-L |
| pDC | RET | 2.46E-06 | 2.573457362 | 0.423 | 0.126 | 0.048331101 | Lep-L |
| pDC | TTTY14 | 6.44E-07 | 2.941298708 | 0.385 | 0.097 | 0.012681298 | Lep-L |
| pDC | APOE | 8.28E-15 | 3.356585425 | 0.731 | 0.185 | 1.63E-10 | Lep-L |
| pDC | BEND4 | 1.24E-08 | 3.686679217 | 0.346 | 0.063 | 0.000243508 | Lep-L |
| pDC | APOC1 | 7.35E-08 | 3.89882188 | 0.385 | 0.09 | 0.001446214 | Lep-L |
| pDC | CLEC7A | 2.65E-15 | 4.077646881 | 0.269 | 0.018 | 5.22E-11 | Lep-L |
| pDC | RNF149 | 7.48E-08 | 0.528921732 | 0.839 | 0.86 | 0.001472267 | Lep-T |
| pDC | CCR7 | 1.20E-08 | 0.56095282 | 0.366 | 0.174 | 0.000235574 | Lep-T |
| pDC | XBP1 | 1.89E-09 | 0.567166738 | 0.901 | 0.851 | 3.73E-05 | Lep-T |
| pDC | PPP1R15A | 2.65E-07 | 0.585539733 | 0.925 | 0.913 | 0.005207134 | Lep-T |
| pDC | SLC7A51 | 2.69E-07 | 0.59551695 | 0.882 | 0.907 | 0.005294776 | Lep-T |
| pDC | ANKRD11 | 4.49E-07 | 0.608762417 | 0.894 | 0.893 | 0.008846965 | Lep-T |
| pDC | REL1 | 4.45E-13 | 0.634141504 | 0.957 | 0.921 | 8.75E-09 | Lep-T |
| pDC | GPR1831 | 2.84E-16 | 0.649175219 | 0.969 | 0.949 | 5.60E-12 | Lep-T |
| pDC | HSP90B11 | 3.45E-11 | 0.667520485 | 0.981 | 0.974 | 6.79E-07 | Lep-T |

| | | | | | | | |
|---|---|---|---|---|---|---|---|
| pDC | NFKB1 | 1.70E-09 | 0.689957819 | 0.876 | 0.757 | 3.34E-05 | Lep-T |
| pDC | RAB11FIP1 | 8.73E-13 | 0.691127937 | 0.894 | 0.913 | 1.72E-08 | Lep-T |
| pDC | JUND1 | 7.22E-11 | 0.70922878 | 0.981 | 0.974 | 1.42E-06 | Lep-T |
| pDC | LDLRAD4 | 2.66E-08 | 0.73712876 | 0.876 | 0.894 | 0.000523483 | Lep-T |
| pDC | FYTTD1 | 5.15E-08 | 0.737231886 | 0.801 | 0.722 | 0.001013268 | Lep-T |
| pDC | PMEPA1 | 1.78E-10 | 0.825026823 | 0.795 | 0.681 | 3.50E-06 | Lep-T |
| pDC | SRGN1 | 9.28E-19 | 0.859052646 | 0.975 | 0.969 | 1.83E-14 | Lep-T |
| pDC | HSP90AB11 | 1.73E-13 | 0.869268074 | 0.988 | 0.975 | 3.40E-09 | Lep-T |
| pDC | GNA131 | 2.01E-09 | 0.878805936 | 0.789 | 0.618 | 3.95E-05 | Lep-T |
| pDC | IRF4 | 1.48E-11 | 0.882128511 | 0.907 | 0.875 | 2.91E-07 | Lep-T |
| pDC | CXCR4 | 1.33E-12 | 0.905494362 | 0.894 | 0.912 | 2.63E-08 | Lep-T |
| pDC | POLB1 | 1.31E-13 | 0.925212664 | 0.876 | 0.844 | 2.59E-09 | Lep-T |
| pDC | NR4A1 | 1.56E-09 | 0.936238221 | 0.888 | 0.838 | 3.06E-05 | Lep-T |
| pDC | MIR155HG | 1.62E-09 | 0.945023596 | 0.646 | 0.425 | 3.18E-05 | Lep-T |
| pDC | KHDRBS1 | 1.10E-07 | 0.957250037 | 0.708 | 0.56 | 0.002156077 | Lep-T |
| pDC | TGFB1 | 1.54E-07 | 0.988864384 | 0.807 | 0.729 | 0.003023071 | Lep-T |
| pDC | EZR1 | 2.24E-23 | 1.000963843 | 0.988 | 0.932 | 4.41E-19 | Lep-T |
| pDC | HLA-DQA21 | 8.59E-07 | 1.007032531 | 0.522 | 0.362 | 0.016906494 | Lep-T |
| pDC | HLA-DQA1 | 5.82E-13 | 1.008541776 | 0.95 | 0.943 | 1.15E-08 | Lep-T |
| pDC | GRASP | 1.39E-07 | 1.016360375 | 0.77 | 0.634 | 0.002733186 | Lep-T |
| pDC | HSPA5 | 1.49E-09 | 1.038251477 | 0.857 | 0.779 | 2.93E-05 | Lep-T |
| pDC | ICAM1 | 4.08E-07 | 1.058345432 | 0.615 | 0.466 | 0.008032281 | Lep-T |
| pDC | HNRNPA0 | 2.17E-06 | 1.070323213 | 0.683 | 0.596 | 0.042699209 | Lep-T |
| pDC | CD831 | 3.20E-19 | 1.073151645 | 0.944 | 0.894 | 6.30E-15 | Lep-T |
| pDC | PSAP | 2.51E-10 | 1.07334508 | 0.801 | 0.734 | 4.94E-06 | Lep-T |
| pDC | EEF2 | 1.68E-18 | 1.07631592 | 0.888 | 0.865 | 3.31E-14 | Lep-T |
| pDC | BRD4 | 2.08E-12 | 1.08304574 | 0.758 | 0.622 | 4.09E-08 | Lep-T |
| pDC | TSPYL21 | 7.43E-09 | 1.095546023 | 0.727 | 0.594 | 0.000146208 | Lep-T |
| pDC | DUSP5 | 7.40E-09 | 1.104362014 | 0.938 | 0.912 | 0.000145569 | Lep-T |
| pDC | CTSB | 6.29E-08 | 1.110193317 | 0.795 | 0.763 | 0.001238379 | Lep-T |
| pDC | SQSTM1 | 3.50E-18 | 1.179981147 | 0.845 | 0.684 | 6.88E-14 | Lep-T |
| pDC | SEC13 | 8.96E-08 | 1.19229392 | 0.503 | 0.319 | 0.001763688 | Lep-T |
| pDC | BHLHE40 | 3.87E-09 | 1.227657852 | 0.745 | 0.621 | 7.62E-05 | Lep-T |
| pDC | DDIT4 | 9.42E-10 | 1.239767022 | 0.826 | 0.74 | 1.85E-05 | Lep-T |
| pDC | IRF2BP2 | 7.21E-12 | 1.249000244 | 0.776 | 0.744 | 1.42E-07 | Lep-T |
| pDC | ATP5PB | 1.26E-06 | 1.2936752 | 0.46 | 0.304 | 0.024852173 | Lep-T |
| pDC | SEMA7A | 1.08E-09 | 1.339976578 | 0.559 | 0.388 | 2.13E-05 | Lep-T |
| pDC | KDM6B | 2.25E-23 | 1.358651325 | 0.857 | 0.607 | 4.42E-19 | Lep-T |
| pDC | MCL11 | 1.60E-24 | 1.388327903 | 0.932 | 0.824 | 3.15E-20 | Lep-T |
| pDC | PNRC1 | 3.96E-19 | 1.443041584 | 0.957 | 0.925 | 7.80E-15 | Lep-T |
| pDC | SYNGR2 | 1.45E-06 | 1.466962533 | 0.64 | 0.525 | 0.028444839 | Lep-T |
| pDC | RBM38 | 4.07E-09 | 1.488570703 | 0.59 | 0.406 | 8.01E-05 | Lep-T |
| pDC | SMIM3 | 4.86E-09 | 1.491959203 | 0.615 | 0.45 | 9.56E-05 | Lep-T |
| pDC | MARCKSL1 | 6.07E-07 | 1.499968166 | 0.59 | 0.429 | 0.011956637 | Lep-T |
| pDC | GNA15 | 1.34E-21 | 1.606538997 | 0.876 | 0.775 | 2.64E-17 | Lep-T |
| pDC | MKNK2 | 6.46E-11 | 1.681401785 | 0.652 | 0.45 | 1.27E-06 | Lep-T |
| pDC | NINJ1 | 1.55E-09 | 1.79899183 | 0.54 | 0.356 | 3.04E-05 | Lep-T |
| pDC | UBALD2 | 1.54E-09 | 1.80862369 | 0.596 | 0.409 | 3.02E-05 | Lep-T |
| pDC | BCL3 | 1.26E-10 | 1.893335884 | 0.516 | 0.307 | 2.48E-06 | Lep-T |
| pDC | CDKN1A | 9.72E-27 | 1.910360317 | 0.901 | 0.728 | 1.91E-22 | Lep-T |
| pDC | EMD | 3.60E-12 | 1.969006264 | 0.46 | 0.229 | 7.10E-08 | Lep-T |
| pDC | CSRNP1 | 4.61E-14 | 1.986088336 | 0.534 | 0.296 | 9.08E-10 | Lep-T |
| pDC | CD74 | 4.57E-27 | 1.986882443 | 1 | 0.999 | 9.00E-23 | Lep-T |
| pDC | PIM2 | 6.74E-13 | 2.007537185 | 0.497 | 0.263 | 1.33E-08 | Lep-T |
| pDC | NUAK2 | 2.21E-07 | 2.070987592 | 0.286 | 0.132 | 0.004343635 | Lep-T |
| pDC | NFKBIA | 2.20E-07 | 2.074368581 | 0.789 | 0.699 | 0.004339946 | Lep-T |

| | | | | | | | |
|---|---|---|---|---|---|---|---|
| pDC | U2AF1 | 8.75E-15 | 2.275370149 | 0.385 | 0.143 | 1.72E-10 | Lep-T |
| pDC | ZFP36 | 1.51E-20 | 2.609713166 | 0.975 | 0.941 | 2.97E-16 | Lep-T |
| pDC | SIK1B | 1.09E-12 | 2.805991528 | 0.311 | 0.103 | 2.15E-08 | Lep-T |
| pDC | GYPC | 1.10E-16 | 2.910171531 | 0.36 | 0.112 | 2.16E-12 | Lep-T |
| pDC | RBFOX2 | 1.29E-21 | 2.945401204 | 0.683 | 0.391 | 2.55E-17 | Lep-T |
| pDC | APOO | 3.72E-17 | 2.979387771 | 0.391 | 0.137 | 7.32E-13 | Lep-T |
| pDC | ALDOA | 1.93E-19 | 3.350948411 | 0.36 | 0.104 | 3.81E-15 | Lep-T |
| pDC | POMP | 8.56E-13 | 0.501175918 | 0.899 | 0.774 | 1.69E-08 | Mari |
| pDC | TSPAN13 | 9.44E-13 | 0.50341391 | 0.946 | 0.83 | 1.86E-08 | Mari |
| pDC | NME2 | 2.12E-18 | 0.503517469 | 0.976 | 0.907 | 4.17E-14 | Mari |
| pDC | CLECL1 | 6.16E-07 | 0.503716081 | 0.604 | 0.428 | 0.012126945 | Mari |
| pDC | GPSM3 | 3.33E-09 | 0.504238672 | 0.729 | 0.552 | 6.55E-05 | Mari |
| pDC | UFM1 | 3.09E-07 | 0.505873347 | 0.67 | 0.527 | 0.006090446 | Mari |
| pDC | UBE2D2 | 1.47E-11 | 0.507739999 | 0.854 | 0.721 | 2.89E-07 | Mari |
| pDC | TMEM258 | 4.05E-11 | 0.509295722 | 0.875 | 0.766 | 7.98E-07 | Mari |
| pDC | GPX4 | 2.81E-12 | 0.510078289 | 0.878 | 0.762 | 5.54E-08 | Mari |
| pDC | COX8A | 1.41E-13 | 0.512996403 | 0.914 | 0.804 | 2.77E-09 | Mari |
| pDC | SNRPG | 4.35E-12 | 0.518333955 | 0.893 | 0.756 | 8.56E-08 | Mari |
| pDC | UBE2L3 | 1.65E-06 | 0.51843562 | 0.69 | 0.578 | 0.032426892 | Mari |
| pDC | TXN | 3.74E-16 | 0.519379354 | 0.991 | 0.929 | 7.37E-12 | Mari |
| pDC | B2M | 2.55E-36 | 0.520482154 | 1 | 0.998 | 5.02E-32 | Mari |
| pDC | SIVA1 | 8.59E-11 | 0.523254626 | 0.854 | 0.715 | 1.69E-06 | Mari |
| pDC | STX7 | 2.82E-08 | 0.523445674 | 0.717 | 0.549 | 0.000555974 | Mari |
| pDC | FERMT3 | 3.56E-08 | 0.523485203 | 0.765 | 0.628 | 0.000701658 | Mari |
| pDC | HINT1 | 2.56E-20 | 0.5253307 | 0.976 | 0.937 | 5.04E-16 | Mari |
| pDC | SRSF5 | 6.31E-09 | 0.525545161 | 0.845 | 0.675 | 0.000124223 | Mari |
| pDC | SEM1 | 3.96E-10 | 0.533250999 | 0.81 | 0.703 | 7.79E-06 | Mari |
| pDC | ACYP2 | 2.53E-06 | 0.533353116 | 0.497 | 0.354 | 0.049705153 | Mari |
| pDC | PSMB3 | 7.94E-09 | 0.533531095 | 0.786 | 0.648 | 0.000156337 | Mari |
| pDC | KRTCAP2 | 6.04E-10 | 0.536472605 | 0.845 | 0.677 | 1.19E-05 | Mari |
| pDC | IGFLR1 | 6.65E-12 | 0.537870914 | 0.869 | 0.741 | 1.31E-07 | Mari |
| pDC | TAF10 | 2.72E-10 | 0.537873217 | 0.836 | 0.699 | 5.35E-06 | Mari |
| pDC | LSM8 | 5.76E-10 | 0.538147389 | 0.786 | 0.596 | 1.13E-05 | Mari |
| pDC | ZDHHC17 | 2.68E-08 | 0.538981308 | 0.753 | 0.594 | 0.000526591 | Mari |
| pDC | EIF1B | 2.72E-07 | 0.540427415 | 0.711 | 0.576 | 0.005359037 | Mari |
| pDC | MIF | 6.52E-15 | 0.540754606 | 0.917 | 0.808 | 1.28E-10 | Mari |
| pDC | CYTH4 | 1.10E-07 | 0.542836534 | 0.539 | 0.366 | 0.002158279 | Mari |
| pDC | APRT | 2.94E-14 | 0.543701337 | 0.914 | 0.798 | 5.78E-10 | Mari |
| pDC | CD164 | 1.18E-08 | 0.544187568 | 0.801 | 0.624 | 0.000232449 | Mari |
| pDC | RASD1 | 2.03E-07 | 0.545766686 | 0.646 | 0.497 | 0.004000333 | Mari |
| pDC | MRPS24 | 2.27E-09 | 0.54856682 | 0.625 | 0.444 | 4.46E-05 | Mari |
| pDC | ATOX1 | 1.34E-07 | 0.548813997 | 0.696 | 0.539 | 0.002644276 | Mari |
| pDC | FEZ2 | 2.35E-06 | 0.549228132 | 0.554 | 0.416 | 0.046287871 | Mari |
| pDC | TMSB10 | 7.30E-30 | 0.549890301 | 0.997 | 0.984 | 1.44E-25 | Mari |
| pDC | VDAC3 | 7.42E-08 | 0.553263476 | 0.664 | 0.533 | 0.001461355 | Mari |
| pDC | DPYSL2 | 1.26E-07 | 0.553705459 | 0.646 | 0.485 | 0.00247982 | Mari |
| pDC | NDUFB1 | 1.03E-10 | 0.554764835 | 0.798 | 0.614 | 2.03E-06 | Mari |
| pDC | UNC93B1 | 4.67E-07 | 0.556214548 | 0.711 | 0.562 | 0.009182923 | Mari |
| pDC | TMED10 | 1.59E-07 | 0.556412702 | 0.789 | 0.655 | 0.003121633 | Mari |
| pDC | PPIB | 1.18E-20 | 0.558491872 | 0.979 | 0.915 | 2.32E-16 | Mari |
| pDC | GABARAPL2 | 3.33E-15 | 0.559945867 | 0.887 | 0.727 | 6.56E-11 | Mari |
| pDC | TUFM | 1.43E-07 | 0.560448042 | 0.592 | 0.412 | 0.002813348 | Mari |
| pDC | CCL4L2 | 9.83E-08 | 0.56068842 | 0.39 | 0.23 | 0.001935494 | Mari |
| pDC | BUD23 | 4.70E-07 | 0.562606238 | 0.661 | 0.529 | 0.009247154 | Mari |
| pDC | ATP5PO | 2.40E-07 | 0.563893966 | 0.685 | 0.55 | 0.004719506 | Mari |
| pDC | SNRPF | 2.57E-09 | 0.564527049 | 0.729 | 0.606 | 5.06E-05 | Mari |

| | | | | | | | |
|---|---|---|---|---|---|---|---|
| pDC | OCIAD1 | 2.05E-08 | 0.564704597 | 0.702 | 0.523 | 0.000403424 | Mari |
| pDC | FKBP1A | 2.93E-07 | 0.568915514 | 0.676 | 0.55 | 0.005774542 | Mari |
| pDC | SELENOS | 7.26E-13 | 0.570047576 | 0.932 | 0.81 | 1.43E-08 | Mari |
| pDC | NDUFB4 | 3.99E-12 | 0.570563032 | 0.821 | 0.644 | 7.86E-08 | Mari |
| pDC | PRR13 | 7.03E-07 | 0.575031189 | 0.601 | 0.473 | 0.013832667 | Mari |
| pDC | ZNF428 | 7.79E-07 | 0.575622741 | 0.545 | 0.4 | 0.01532552 | Mari |
| pDC | COX6B1 | 3.93E-20 | 0.575995597 | 0.943 | 0.859 | 7.74E-16 | Mari |
| pDC | HLA-DMA | 9.03E-16 | 0.576677268 | 0.955 | 0.875 | 1.78E-11 | Mari |
| pDC | CCDC186 | 3.30E-12 | 0.576919231 | 0.857 | 0.681 | 6.50E-08 | Mari |
| pDC | ZBTB18 | 1.46E-06 | 0.579994919 | 0.673 | 0.531 | 0.028735068 | Mari |
| pDC | TRABD | 1.25E-06 | 0.5810455 | 0.646 | 0.533 | 0.02455196 | Mari |
| pDC | GUK1 | 3.25E-15 | 0.581751257 | 0.92 | 0.808 | 6.39E-11 | Mari |
| pDC | SSR2 | 4.34E-14 | 0.58405047 | 0.896 | 0.764 | 8.55E-10 | Mari |
| pDC | GLIPR1 | 4.84E-07 | 0.585265099 | 0.702 | 0.588 | 0.009525719 | Mari |
| pDC | TOMM6 | 2.41E-11 | 0.58546402 | 0.821 | 0.675 | 4.74E-07 | Mari |
| pDC | C1QBP | 1.88E-11 | 0.585675916 | 0.765 | 0.646 | 3.70E-07 | Mari |
| pDC | NENF | 6.53E-08 | 0.586115496 | 0.622 | 0.465 | 0.001284968 | Mari |
| pDC | ATP5MG | 8.64E-27 | 0.586231978 | 0.985 | 0.901 | 1.70E-22 | Mari |
| pDC | ATP5MPL | 1.81E-10 | 0.586253415 | 0.768 | 0.56 | 3.57E-06 | Mari |
| pDC | REEP5 | 1.18E-07 | 0.586689702 | 0.649 | 0.529 | 0.002322596 | Mari |
| pDC | TMED9 | 3.41E-09 | 0.588014208 | 0.705 | 0.554 | 6.71E-05 | Mari |
| pDC | TMEM50A | 1.10E-09 | 0.589884416 | 0.807 | 0.693 | 2.17E-05 | Mari |
| pDC | CFL1 | 1.55E-26 | 0.590283816 | 0.973 | 0.917 | 3.06E-22 | Mari |
| pDC | PSMB6 | 8.21E-10 | 0.590328245 | 0.771 | 0.61 | 1.62E-05 | Mari |
| pDC | CLTA | 4.53E-10 | 0.591669836 | 0.765 | 0.598 | 8.92E-06 | Mari |
| pDC | ARL5A | 2.29E-07 | 0.592653471 | 0.58 | 0.438 | 0.004506915 | Mari |
| pDC | HLA-DPA1 | 3.85E-26 | 0.594487866 | 0.994 | 0.947 | 7.58E-22 | Mari |
| pDC | SMARCB1 | 4.52E-07 | 0.595764499 | 0.634 | 0.475 | 0.008890623 | Mari |
| pDC | OTULINL | 1.24E-06 | 0.596064016 | 0.527 | 0.394 | 0.024344378 | Mari |
| pDC | SNRPD3 | 3.05E-07 | 0.596701261 | 0.592 | 0.444 | 0.006013421 | Mari |
| pDC | COMMD3 | 1.09E-06 | 0.598470108 | 0.53 | 0.384 | 0.021489546 | Mari |
| pDC | NDUFA1 | 4.96E-10 | 0.598816807 | 0.81 | 0.713 | 9.77E-06 | Mari |
| pDC | CDC40 | 5.09E-07 | 0.599196137 | 0.598 | 0.434 | 0.010009999 | Mari |
| pDC | MYL12B | 4.46E-24 | 0.60018436 | 0.976 | 0.897 | 8.78E-20 | Mari |
| pDC | BTG2 | 1.76E-17 | 0.600892638 | 0.923 | 0.812 | 3.47E-13 | Mari |
| pDC | NDUFB8 | 3.88E-14 | 0.601489488 | 0.878 | 0.733 | 7.64E-10 | Mari |
| pDC | COX6C | 3.45E-18 | 0.602255673 | 0.926 | 0.824 | 6.79E-14 | Mari |
| pDC | ALOX5AP | 6.67E-21 | 0.608229541 | 0.967 | 0.917 | 1.31E-16 | Mari |
| pDC | RABAC1 | 2.93E-12 | 0.608871084 | 0.818 | 0.626 | 5.77E-08 | Mari |
| pDC | COX14 | 1.86E-07 | 0.609780622 | 0.583 | 0.44 | 0.003652015 | Mari |
| pDC | SFT2D1 | 1.70E-09 | 0.610687544 | 0.723 | 0.584 | 3.34E-05 | Mari |
| pDC | SAMHD1 | 9.62E-10 | 0.611912154 | 0.726 | 0.56 | 1.89E-05 | Mari |
| pDC | GNG5 | 3.91E-16 | 0.61237949 | 0.92 | 0.804 | 7.69E-12 | Mari |
| pDC | DDOST | 2.70E-09 | 0.613854336 | 0.682 | 0.519 | 5.31E-05 | Mari |
| pDC | COMT | 2.22E-09 | 0.616213892 | 0.69 | 0.549 | 4.37E-05 | Mari |
| pDC | DECR1 | 1.74E-06 | 0.616910564 | 0.488 | 0.35 | 0.034295737 | Mari |
| pDC | PSMD9 | 1.98E-08 | 0.619138742 | 0.649 | 0.475 | 0.000389539 | Mari |
| pDC | VPS28 | 7.84E-07 | 0.619433361 | 0.592 | 0.461 | 0.015430823 | Mari |
| pDC | FABP5 | 4.11E-12 | 0.622207569 | 0.771 | 0.598 | 8.09E-08 | Mari |
| pDC | PEBP1 | 1.66E-12 | 0.62223287 | 0.854 | 0.695 | 3.27E-08 | Mari |
| pDC | ANAPC11 | 7.67E-09 | 0.622298654 | 0.64 | 0.471 | 0.000151042 | Mari |
| pDC | SPCS2 | 1.43E-13 | 0.623602274 | 0.875 | 0.729 | 2.81E-09 | Mari |
| pDC | COX5A | 5.19E-15 | 0.624002215 | 0.851 | 0.679 | 1.02E-10 | Mari |
| pDC | OSTC | 4.40E-14 | 0.624259459 | 0.896 | 0.774 | 8.66E-10 | Mari |
| pDC | PAIP1 | 1.02E-07 | 0.625717792 | 0.583 | 0.428 | 0.002004393 | Mari |
| pDC | RTRAF | 4.60E-13 | 0.627934091 | 0.827 | 0.657 | 9.06E-09 | Mari |

| | | | | | | | |
|---|---|---|---|---|---|---|---|
| pDC | SUB1 | 1.45E-24 | 0.628169309 | 0.976 | 0.911 | 2.85E-20 | Mari |
| pDC | MPG | 4.05E-07 | 0.628675285 | 0.595 | 0.44 | 0.007973062 | Mari |
| pDC | SUMO2 | 3.41E-24 | 0.628786486 | 0.961 | 0.863 | 6.71E-20 | Mari |
| pDC | NOP10 | 2.46E-10 | 0.629089992 | 0.732 | 0.558 | 4.85E-06 | Mari |
| pDC | LDHB | 5.16E-17 | 0.629611586 | 0.914 | 0.778 | 1.02E-12 | Mari |
| pDC | SERPINB1 | 3.11E-11 | 0.629966712 | 0.649 | 0.45 | 6.12E-07 | Mari |
| pDC | PTRHD1 | 1.28E-09 | 0.630065221 | 0.699 | 0.525 | 2.52E-05 | Mari |
| pDC | TMA7 | 2.88E-23 | 0.631829432 | 0.97 | 0.857 | 5.67E-19 | Mari |
| pDC | SH3BGRL3 | 3.09E-16 | 0.632055453 | 0.955 | 0.832 | 6.08E-12 | Mari |
| pDC | COX7A2 | 4.80E-21 | 0.63297554 | 0.943 | 0.836 | 9.45E-17 | Mari |
| pDC | UQCRFS1 | 2.63E-07 | 0.633391508 | 0.619 | 0.477 | 0.00517571 | Mari |
| pDC | ATP5MC3 | 6.22E-18 | 0.634910012 | 0.914 | 0.784 | 1.22E-13 | Mari |
| pDC | NDUFV2 | 1.57E-14 | 0.635134162 | 0.884 | 0.687 | 3.09E-10 | Mari |
| pDC | HMGN2 | 1.62E-13 | 0.635428796 | 0.884 | 0.77 | 3.19E-09 | Mari |
| pDC | EMB | 3.56E-07 | 0.636259111 | 0.583 | 0.438 | 0.00700335 | Mari |
| pDC | IMP3 | 9.71E-07 | 0.636313931 | 0.423 | 0.277 | 0.019119369 | Mari |
| pDC | CAT | 2.76E-08 | 0.637199029 | 0.64 | 0.505 | 0.000542449 | Mari |
| pDC | VDAC2 | 3.42E-10 | 0.638097272 | 0.75 | 0.602 | 6.73E-06 | Mari |
| pDC | UBL5 | 1.64E-22 | 0.639104753 | 0.961 | 0.832 | 3.24E-18 | Mari |
| pDC | MOB1A | 1.26E-13 | 0.640479986 | 0.866 | 0.697 | 2.47E-09 | Mari |
| pDC | CAPG | 8.70E-14 | 0.642209404 | 0.869 | 0.75 | 1.71E-09 | Mari |
| pDC | CRIP1 | 1.67E-14 | 0.643131211 | 0.932 | 0.778 | 3.29E-10 | Mari |
| pDC | DRAP1 | 1.09E-10 | 0.643236048 | 0.774 | 0.606 | 2.14E-06 | Mari |
| pDC | CYBB | 1.73E-13 | 0.643609917 | 0.893 | 0.754 | 3.40E-09 | Mari |
| pDC | CRYM | 4.81E-09 | 0.643864482 | 0.548 | 0.356 | 9.47E-05 | Mari |
| pDC | ANXA2 | 1.25E-07 | 0.644641105 | 0.551 | 0.388 | 0.002468769 | Mari |
| pDC | VAMP2 | 8.80E-07 | 0.645548371 | 0.551 | 0.402 | 0.017327317 | Mari |
| pDC | CD302 | 1.57E-07 | 0.645848691 | 0.542 | 0.386 | 0.003096502 | Mari |
| pDC | IGHM | 6.71E-09 | 0.645858407 | 0.676 | 0.511 | 0.00013204 | Mari |
| pDC | CD47 | 2.22E-11 | 0.646393613 | 0.821 | 0.655 | 4.38E-07 | Mari |
| pDC | RNH1 | 9.89E-08 | 0.646611626 | 0.589 | 0.436 | 0.00194731 | Mari |
| pDC | SPINT2 | 3.28E-12 | 0.649118701 | 0.774 | 0.59 | 6.45E-08 | Mari |
| pDC | PSMB1 | 9.01E-13 | 0.649512598 | 0.833 | 0.665 | 1.77E-08 | Mari |
| pDC | NAA38 | 3.06E-09 | 0.650741224 | 0.601 | 0.424 | 6.01E-05 | Mari |
| pDC | DDT | 1.48E-12 | 0.650819125 | 0.792 | 0.614 | 2.91E-08 | Mari |
| pDC | OST4 | 3.22E-25 | 0.651604811 | 0.961 | 0.869 | 6.33E-21 | Mari |
| pDC | ATP5F1D | 5.14E-16 | 0.651845857 | 0.908 | 0.745 | 1.01E-11 | Mari |
| pDC | HCST | 2.07E-13 | 0.652403294 | 0.848 | 0.705 | 4.08E-09 | Mari |
| pDC | OAZ1 | 3.32E-30 | 0.65252786 | 0.988 | 0.935 | 6.54E-26 | Mari |
| pDC | UBE2I | 6.36E-08 | 0.654070837 | 0.679 | 0.547 | 0.001252295 | Mari |
| pDC | MLF2 | 1.16E-06 | 0.655345982 | 0.491 | 0.35 | 0.022820491 | Mari |
| pDC | CYC1 | 1.98E-06 | 0.655556686 | 0.53 | 0.386 | 0.038976673 | Mari |
| pDC | IFI16 | 5.11E-09 | 0.656402825 | 0.744 | 0.598 | 0.000100634 | Mari |
| pDC | PSMB2 | 9.46E-10 | 0.656423933 | 0.661 | 0.467 | 1.86E-05 | Mari |
| pDC | MEF2C | 2.07E-17 | 0.656896871 | 0.938 | 0.768 | 4.08E-13 | Mari |
| pDC | MRPS21 | 5.97E-10 | 0.657271659 | 0.735 | 0.586 | 1.18E-05 | Mari |
| pDC | SDHD | 1.90E-06 | 0.658573765 | 0.5 | 0.362 | 0.037420511 | Mari |
| pDC | ALKBH7 | 8.11E-08 | 0.659027148 | 0.628 | 0.483 | 0.001596625 | Mari |
| pDC | NPC2 | 1.47E-20 | 0.660067702 | 0.973 | 0.855 | 2.89E-16 | Mari |
| pDC | STMN1 | 1.62E-14 | 0.660620142 | 0.863 | 0.691 | 3.19E-10 | Mari |
| pDC | TATDN3 | 1.33E-08 | 0.66222967 | 0.545 | 0.356 | 0.000262093 | Mari |
| pDC | CORO1A | 5.61E-14 | 0.663400484 | 0.899 | 0.754 | 1.10E-09 | Mari |
| pDC | MDH2 | 5.00E-10 | 0.665152762 | 0.708 | 0.535 | 9.84E-06 | Mari |
| pDC | COX5B | 1.82E-17 | 0.665518315 | 0.89 | 0.719 | 3.59E-13 | Mari |
| pDC | SMIM7 | 1.05E-06 | 0.66614798 | 0.557 | 0.41 | 0.020595395 | Mari |
| pDC | CALM2 | 8.14E-21 | 0.667212718 | 0.958 | 0.871 | 1.60E-16 | Mari |

| | | | | | | | |
|---|---|---|---|---|---|---|---|
| pDC | SLIRP | 1.12E-06 | 0.66759754 | 0.521 | 0.374 | 0.021957372 | Mari |
| pDC | PSMB9 | 1.11E-13 | 0.667702558 | 0.839 | 0.681 | 2.19E-09 | Mari |
| pDC | AIMP1 | 9.52E-07 | 0.668479784 | 0.494 | 0.36 | 0.018734008 | Mari |
| pDC | MRPS36 | 6.35E-07 | 0.66976873 | 0.458 | 0.303 | 0.012508973 | Mari |
| pDC | SNRPE | 1.93E-10 | 0.671054457 | 0.67 | 0.497 | 3.81E-06 | Mari |
| pDC | LSP1 | 1.18E-12 | 0.67169957 | 0.798 | 0.622 | 2.31E-08 | Mari |
| pDC | NDUFA7 | 1.31E-12 | 0.671783733 | 0.744 | 0.547 | 2.57E-08 | Mari |
| pDC | NAA10 | 1.55E-07 | 0.671961583 | 0.47 | 0.309 | 0.003055578 | Mari |
| pDC | CXXC5 | 1.55E-07 | 0.673032998 | 0.562 | 0.41 | 0.003045237 | Mari |
| pDC | SPCS1 | 6.22E-28 | 0.674333137 | 0.97 | 0.891 | 1.22E-23 | Mari |
| pDC | MRPL43 | 4.88E-07 | 0.675246602 | 0.438 | 0.281 | 0.009615045 | Mari |
| pDC | SCAND1 | 2.15E-09 | 0.676114626 | 0.708 | 0.552 | 4.23E-05 | Mari |
| pDC | CCT8 | 7.61E-12 | 0.680529986 | 0.744 | 0.596 | 1.50E-07 | Mari |
| pDC | TRAPPC5 | 1.30E-06 | 0.680806786 | 0.5 | 0.382 | 0.025623985 | Mari |
| pDC | MRPL41 | 8.86E-10 | 0.680992965 | 0.625 | 0.469 | 1.74E-05 | Mari |
| pDC | GPR65 | 2.46E-10 | 0.681284151 | 0.804 | 0.673 | 4.84E-06 | Mari |
| pDC | SRP14 | 1.10E-31 | 0.681759743 | 0.985 | 0.927 | 2.17E-27 | Mari |
| pDC | ATP5MD | 4.82E-16 | 0.682791158 | 0.869 | 0.715 | 9.49E-12 | Mari |
| pDC | PSMD8 | 2.51E-13 | 0.684851722 | 0.771 | 0.566 | 4.93E-09 | Mari |
| pDC | CYSLTR1 | 1.16E-06 | 0.685065209 | 0.551 | 0.422 | 0.022906037 | Mari |
| pDC | NDUFC1 | 6.25E-09 | 0.686683098 | 0.565 | 0.38 | 0.000123096 | Mari |
| pDC | BRI3 | 1.76E-17 | 0.686929468 | 0.899 | 0.754 | 3.47E-13 | Mari |
| pDC | TALDO1 | 1.79E-12 | 0.690042977 | 0.798 | 0.61 | 3.51E-08 | Mari |
| pDC | LILRA4 | 7.06E-18 | 0.690305562 | 0.923 | 0.786 | 1.39E-13 | Mari |
| pDC | MZT2A | 1.56E-11 | 0.690477127 | 0.649 | 0.442 | 3.08E-07 | Mari |
| pDC | BAX | 2.73E-10 | 0.69174664 | 0.673 | 0.477 | 5.38E-06 | Mari |
| pDC | DHRS7 | 2.22E-11 | 0.69261443 | 0.774 | 0.602 | 4.38E-07 | Mari |
| pDC | DPP7 | 5.54E-12 | 0.693787252 | 0.759 | 0.574 | 1.09E-07 | Mari |
| pDC | RAB4A | 8.35E-07 | 0.694362975 | 0.47 | 0.327 | 0.016432405 | Mari |
| pDC | EXOSC8 | 1.98E-06 | 0.695273556 | 0.393 | 0.251 | 0.03902942 | Mari |
| pDC | SERF2 | 6.07E-39 | 0.696208947 | 0.991 | 0.939 | 1.19E-34 | Mari |
| pDC | ACP1 | 6.19E-08 | 0.697370307 | 0.601 | 0.45 | 0.001218303 | Mari |
| pDC | SS18L2 | 1.78E-06 | 0.698862335 | 0.375 | 0.244 | 0.035009427 | Mari |
| pDC | NUDT1 | 1.27E-11 | 0.699287052 | 0.693 | 0.513 | 2.51E-07 | Mari |
| pDC | EMC7 | 1.57E-07 | 0.701273318 | 0.613 | 0.483 | 0.003088449 | Mari |
| pDC | ROMO1 | 4.85E-12 | 0.70280557 | 0.717 | 0.541 | 9.55E-08 | Mari |
| pDC | SNHG8 | 9.50E-14 | 0.703428946 | 0.896 | 0.796 | 1.87E-09 | Mari |
| pDC | ITM2C | 6.19E-28 | 0.703858374 | 0.979 | 0.885 | 1.22E-23 | Mari |
| pDC | TMEM167A | 1.65E-07 | 0.705973945 | 0.562 | 0.424 | 0.003247066 | Mari |
| pDC | SCAMP2 | 6.84E-07 | 0.707776474 | 0.551 | 0.416 | 0.013469616 | Mari |
| pDC | MED30 | 2.18E-08 | 0.707886818 | 0.571 | 0.422 | 0.000430041 | Mari |
| pDC | SSU72 | 1.10E-06 | 0.708382404 | 0.551 | 0.406 | 0.021581871 | Mari |
| pDC | ERP29 | 4.58E-26 | 0.708713959 | 0.964 | 0.871 | 9.02E-22 | Mari |
| pDC | MRPS12 | 9.24E-08 | 0.710110571 | 0.461 | 0.297 | 0.001818582 | Mari |
| pDC | GSTK1 | 1.42E-11 | 0.71043697 | 0.711 | 0.539 | 2.80E-07 | Mari |
| pDC | NDUFA3 | 1.21E-09 | 0.711107424 | 0.622 | 0.448 | 2.37E-05 | Mari |
| pDC | CHCHD10 | 1.96E-09 | 0.711881801 | 0.655 | 0.497 | 3.86E-05 | Mari |
| pDC | PAXX | 7.86E-12 | 0.712329781 | 0.878 | 0.75 | 1.55E-07 | Mari |
| pDC | DCTN3 | 4.37E-08 | 0.712926698 | 0.571 | 0.432 | 0.000859862 | Mari |
| pDC | UCP2 | 3.02E-16 | 0.713231514 | 0.875 | 0.693 | 5.95E-12 | Mari |
| pDC | NDUFS8 | 6.53E-11 | 0.714215479 | 0.634 | 0.455 | 1.29E-06 | Mari |
| pDC | LEPROTL1 | 4.68E-08 | 0.71520944 | 0.548 | 0.388 | 0.000922102 | Mari |
| pDC | PARK7 | 3.08E-31 | 0.716754285 | 0.973 | 0.913 | 6.05E-27 | Mari |
| pDC | ATRAID | 6.71E-12 | 0.718768238 | 0.604 | 0.406 | 1.32E-07 | Mari |
| pDC | PSENEN | 2.10E-09 | 0.718963874 | 0.625 | 0.455 | 4.13E-05 | Mari |
| pDC | ATP6V1G1 | 6.34E-18 | 0.719395854 | 0.884 | 0.729 | 1.25E-13 | Mari |

| | | | | | | | |
|---|---|---|---|---|---|---|---|
| pDC | PPP4C | 7.48E-07 | 0.721470734 | 0.515 | 0.37 | 0.014730226 | Mari |
| pDC | AURKAIP1 | 5.27E-14 | 0.722653439 | 0.771 | 0.58 | 1.04E-09 | Mari |
| pDC | CIB1 | 8.43E-09 | 0.723105033 | 0.595 | 0.436 | 0.000166013 | Mari |
| pDC | STUB1 | 5.29E-08 | 0.724198695 | 0.536 | 0.386 | 0.001040565 | Mari |
| pDC | 2-Mar | 1.63E-08 | 0.725361434 | 0.521 | 0.339 | 0.000320329 | Mari |
| pDC | SNHG9 | 3.55E-07 | 0.72607824 | 0.464 | 0.311 | 0.006982437 | Mari |
| pDC | TYROBP | 2.01E-25 | 0.726160058 | 0.976 | 0.881 | 3.96E-21 | Mari |
| pDC | NDUFA2 | 3.75E-13 | 0.726348713 | 0.729 | 0.554 | 7.39E-09 | Mari |
| pDC | OSTF1 | 9.83E-11 | 0.72667098 | 0.673 | 0.501 | 1.93E-06 | Mari |
| pDC | ZNHIT1 | 1.32E-09 | 0.72833704 | 0.64 | 0.477 | 2.60E-05 | Mari |
| pDC | MIF4GD | 6.25E-09 | 0.728815136 | 0.616 | 0.45 | 0.000123103 | Mari |
| pDC | LSM10 | 2.44E-06 | 0.730317533 | 0.473 | 0.349 | 0.048053186 | Mari |
| pDC | ANAPC16 | 3.12E-15 | 0.732411626 | 0.821 | 0.638 | 6.15E-11 | Mari |
| pDC | CD99 | 8.97E-15 | 0.733429821 | 0.899 | 0.774 | 1.77E-10 | Mari |
| pDC | DCPS | 7.82E-11 | 0.733702511 | 0.613 | 0.41 | 1.54E-06 | Mari |
| pDC | GSTO1 | 6.03E-10 | 0.734731209 | 0.64 | 0.457 | 1.19E-05 | Mari |
| pDC | IDH3A | 2.50E-13 | 0.736413769 | 0.729 | 0.547 | 4.91E-09 | Mari |
| pDC | PSMA4 | 6.79E-12 | 0.738729393 | 0.699 | 0.499 | 1.34E-07 | Mari |
| pDC | C1D | 1.95E-07 | 0.738754463 | 0.5 | 0.35 | 0.003838858 | Mari |
| pDC | IFNAR1 | 2.18E-07 | 0.739592838 | 0.568 | 0.432 | 0.004296485 | Mari |
| pDC | ATP5F1E | 1.11E-47 | 0.740275271 | 0.994 | 0.943 | 2.19E-43 | Mari |
| pDC | RNASET2 | 4.56E-31 | 0.740326199 | 0.994 | 0.913 | 8.98E-27 | Mari |
| pDC | ARPC3 | 1.82E-34 | 0.741218326 | 0.973 | 0.917 | 3.57E-30 | Mari |
| pDC | CCDC50 | 1.42E-28 | 0.741298391 | 0.973 | 0.875 | 2.79E-24 | Mari |
| pDC | ELOB | 1.54E-13 | 0.742359781 | 0.827 | 0.711 | 3.03E-09 | Mari |
| pDC | MRPL33 | 2.01E-06 | 0.742462865 | 0.503 | 0.37 | 0.039627411 | Mari |
| pDC | CDKN2D | 4.19E-14 | 0.742798608 | 0.738 | 0.517 | 8.24E-10 | Mari |
| pDC | SPCS3 | 1.36E-16 | 0.744179284 | 0.842 | 0.669 | 2.69E-12 | Mari |
| pDC | PSMA6 | 1.69E-15 | 0.745755585 | 0.866 | 0.723 | 3.33E-11 | Mari |
| pDC | MPC2 | 1.47E-07 | 0.745778236 | 0.548 | 0.412 | 0.002885214 | Mari |
| pDC | CCND3 | 1.19E-09 | 0.745835689 | 0.67 | 0.509 | 2.34E-05 | Mari |
| pDC | ETFB | 1.38E-09 | 0.748014798 | 0.577 | 0.4 | 2.72E-05 | Mari |
| pDC | POLR2K | 4.79E-09 | 0.748054473 | 0.613 | 0.424 | 9.43E-05 | Mari |
| pDC | NDUFS3 | 9.70E-07 | 0.748232529 | 0.497 | 0.368 | 0.019085447 | Mari |
| pDC | PLP2 | 9.50E-18 | 0.748939923 | 0.929 | 0.79 | 1.87E-13 | Mari |
| pDC | PSMB8 | 9.23E-17 | 0.749530163 | 0.848 | 0.669 | 1.82E-12 | Mari |
| pDC | STOML2 | 3.02E-09 | 0.750086569 | 0.509 | 0.325 | 5.94E-05 | Mari |
| pDC | MRPL52 | 3.84E-07 | 0.750790969 | 0.5 | 0.36 | 0.00755379 | Mari |
| pDC | ERH | 4.16E-19 | 0.754489278 | 0.848 | 0.63 | 8.19E-15 | Mari |
| pDC | ISCU | 4.34E-14 | 0.754598602 | 0.789 | 0.64 | 8.55E-10 | Mari |
| pDC | MRPL34 | 3.50E-07 | 0.755048889 | 0.491 | 0.356 | 0.006883829 | Mari |
| pDC | TMX1 | 8.16E-09 | 0.755638013 | 0.557 | 0.396 | 0.000160576 | Mari |
| pDC | PSMA2 | 3.91E-17 | 0.756287122 | 0.836 | 0.661 | 7.70E-13 | Mari |
| pDC | LYPLA1 | 5.69E-08 | 0.756424886 | 0.482 | 0.317 | 0.001120001 | Mari |
| pDC | RAC2 | 5.93E-17 | 0.75785129 | 0.851 | 0.683 | 1.17E-12 | Mari |
| pDC | APEX1 | 6.21E-12 | 0.758306471 | 0.762 | 0.622 | 1.22E-07 | Mari |
| pDC | ABRACL | 2.06E-14 | 0.759678241 | 0.774 | 0.594 | 4.06E-10 | Mari |
| pDC | SAT2 | 9.79E-07 | 0.762768359 | 0.414 | 0.281 | 0.01927507 | Mari |
| pDC | TXN2 | 5.43E-08 | 0.763898074 | 0.515 | 0.386 | 0.001069162 | Mari |
| pDC | SUMO3 | 1.01E-12 | 0.76560166 | 0.637 | 0.414 | 1.99E-08 | Mari |
| pDC | ATP5ME | 4.52E-15 | 0.766010365 | 0.75 | 0.572 | 8.91E-11 | Mari |
| pDC | NDUFA13 | 6.07E-20 | 0.766066444 | 0.881 | 0.685 | 1.19E-15 | Mari |
| pDC | DBNL | 2.74E-13 | 0.766756216 | 0.696 | 0.483 | 5.40E-09 | Mari |
| pDC | SEC61G | 9.92E-27 | 0.767045372 | 0.958 | 0.848 | 1.95E-22 | Mari |
| pDC | MZB1 | 7.28E-15 | 0.768833634 | 0.872 | 0.721 | 1.43E-10 | Mari |
| pDC | TMEM160 | 2.72E-11 | 0.769370608 | 0.592 | 0.4 | 5.35E-07 | Mari |

| | | | | | | | |
|---|---|---|---|---|---|---|---|
| pDC | TWF2 | 1.38E-07 | 0.771180527 | 0.518 | 0.364 | 0.002713364 | Mari |
| pDC | DERL3 | 5.42E-15 | 0.771320317 | 0.863 | 0.685 | 1.07E-10 | Mari |
| pDC | CHURC1 | 5.99E-11 | 0.772954035 | 0.702 | 0.519 | 1.18E-06 | Mari |
| pDC | LY96 | 5.01E-07 | 0.773775085 | 0.485 | 0.347 | 0.009865696 | Mari |
| pDC | CYBA | 9.99E-39 | 0.774056594 | 0.997 | 0.958 | 1.97E-34 | Mari |
| pDC | NDUFA12 | 6.01E-14 | 0.775666622 | 0.702 | 0.491 | 1.18E-09 | Mari |
| pDC | TLE5 | 2.96E-14 | 0.776244527 | 0.839 | 0.671 | 5.83E-10 | Mari |
| pDC | ITM2B | 3.84E-23 | 0.776868568 | 0.917 | 0.774 | 7.57E-19 | Mari |
| pDC | NEDD8 | 6.77E-19 | 0.777217647 | 0.854 | 0.675 | 1.33E-14 | Mari |
| pDC | TMEM179B | 3.76E-07 | 0.779797443 | 0.473 | 0.315 | 0.007396313 | Mari |
| pDC | MRPL36 | 4.92E-07 | 0.779811646 | 0.521 | 0.378 | 0.009676031 | Mari |
| pDC | PFDN1 | 1.19E-06 | 0.780978716 | 0.387 | 0.24 | 0.023398024 | Mari |
| pDC | SNF8 | 3.94E-08 | 0.781316461 | 0.595 | 0.451 | 0.000775603 | Mari |
| pDC | FIS1 | 1.46E-11 | 0.782080486 | 0.637 | 0.461 | 2.87E-07 | Mari |
| pDC | JTB | 9.46E-17 | 0.782154359 | 0.804 | 0.592 | 1.86E-12 | Mari |
| pDC | TMEM59 | 1.25E-19 | 0.782876151 | 0.884 | 0.697 | 2.47E-15 | Mari |
| pDC | NDUFB10 | 2.28E-16 | 0.783383359 | 0.789 | 0.58 | 4.49E-12 | Mari |
| pDC | TXNL4A | 1.47E-13 | 0.785049747 | 0.738 | 0.586 | 2.90E-09 | Mari |
| pDC | SNHG12 | 2.06E-06 | 0.790625932 | 0.375 | 0.234 | 0.040528123 | Mari |
| pDC | SELENOH | 1.23E-25 | 0.790962864 | 0.932 | 0.764 | 2.42E-21 | Mari |
| pDC | NDUFS6 | 3.05E-18 | 0.791837425 | 0.812 | 0.62 | 6.00E-14 | Mari |
| pDC | ORMDL1 | 7.72E-16 | 0.791884191 | 0.744 | 0.541 | 1.52E-11 | Mari |
| pDC | PHB | 4.14E-15 | 0.792255317 | 0.839 | 0.659 | 8.15E-11 | Mari |
| pDC | TMEM14B | 9.27E-17 | 0.79281701 | 0.801 | 0.574 | 1.83E-12 | Mari |
| pDC | MRPS34 | 8.46E-10 | 0.792887588 | 0.506 | 0.315 | 1.67E-05 | Mari |
| pDC | HMGN1 | 5.68E-23 | 0.793822928 | 0.932 | 0.737 | 1.12E-18 | Mari |
| pDC | TMEM126A | 1.48E-06 | 0.794071103 | 0.33 | 0.196 | 0.029168013 | Mari |
| pDC | ODC1 | 4.16E-11 | 0.794205506 | 0.854 | 0.717 | 8.19E-07 | Mari |
| pDC | GLRX5 | 3.96E-13 | 0.795044394 | 0.637 | 0.422 | 7.79E-09 | Mari |
| pDC | VAMP8 | 1.16E-33 | 0.795918502 | 0.967 | 0.879 | 2.29E-29 | Mari |
| pDC | PCED1B-AS1 | 7.72E-08 | 0.796084795 | 0.598 | 0.463 | 0.001518923 | Mari |
| pDC | LAMTOR5 | 8.56E-17 | 0.796990343 | 0.789 | 0.576 | 1.68E-12 | Mari |
| pDC | LMAN2 | 8.09E-14 | 0.797513997 | 0.693 | 0.477 | 1.59E-09 | Mari |
| pDC | UFC1 | 1.14E-14 | 0.797702723 | 0.774 | 0.608 | 2.24E-10 | Mari |
| pDC | SSBP1 | 4.93E-13 | 0.797924335 | 0.699 | 0.515 | 9.70E-09 | Mari |
| pDC | PPP2R5C | 1.16E-18 | 0.797989718 | 0.905 | 0.717 | 2.28E-14 | Mari |
| pDC | PDCD2 | 1.31E-07 | 0.798202251 | 0.455 | 0.305 | 0.002575947 | Mari |
| pDC | SELL | 4.11E-21 | 0.798633779 | 0.902 | 0.723 | 8.09E-17 | Mari |
| pDC | DAD1 | 2.72E-20 | 0.79931556 | 0.848 | 0.687 | 5.35E-16 | Mari |
| pDC | FOS | 2.08E-29 | 0.801636989 | 0.949 | 0.737 | 4.09E-25 | Mari |
| pDC | MGST3 | 1.28E-06 | 0.80192647 | 0.449 | 0.321 | 0.025104817 | Mari |
| pDC | THOC7 | 1.06E-09 | 0.804908172 | 0.509 | 0.319 | 2.09E-05 | Mari |
| pDC | LSM4 | 7.93E-13 | 0.805212935 | 0.673 | 0.453 | 1.56E-08 | Mari |
| pDC | ATP5MC1 | 1.23E-15 | 0.805352644 | 0.759 | 0.533 | 2.41E-11 | Mari |
| pDC | ITGAE | 4.29E-12 | 0.806470748 | 0.696 | 0.549 | 8.45E-08 | Mari |
| pDC | LSM3 | 7.31E-13 | 0.807596946 | 0.67 | 0.451 | 1.44E-08 | Mari |
| pDC | YIF1A | 1.32E-07 | 0.808869199 | 0.411 | 0.25 | 0.002598572 | Mari |
| pDC | POLR2L | 6.00E-17 | 0.809446084 | 0.824 | 0.681 | 1.18E-12 | Mari |
| pDC | AP2S1 | 7.17E-15 | 0.809562337 | 0.78 | 0.62 | 1.41E-10 | Mari |
| pDC | MRPL51 | 1.63E-12 | 0.812010182 | 0.679 | 0.489 | 3.20E-08 | Mari |
| pDC | BRK1 | 1.20E-20 | 0.813568565 | 0.833 | 0.612 | 2.35E-16 | Mari |
| pDC | CCDC85B | 8.42E-14 | 0.813876388 | 0.682 | 0.457 | 1.66E-09 | Mari |
| pDC | CCS | 1.83E-07 | 0.814382824 | 0.5 | 0.35 | 0.003610533 | Mari |
| pDC | ATP5F1C | 1.16E-17 | 0.816299356 | 0.836 | 0.624 | 2.27E-13 | Mari |
| pDC | PDAP1 | 2.21E-06 | 0.816635419 | 0.446 | 0.329 | 0.043440207 | Mari |
| pDC | CIAO2B | 1.34E-17 | 0.816801931 | 0.801 | 0.604 | 2.64E-13 | Mari |

| | | | | | | | |
|---|---|---|---|---|---|---|---|
| pDC | RGS19 | 1.04E-08 | 0.816848376 | 0.506 | 0.331 | 0.000203824 | Mari |
| pDC | TRMT112 | 2.10E-18 | 0.818532501 | 0.83 | 0.608 | 4.13E-14 | Mari |
| pDC | MRPL55 | 2.42E-06 | 0.819175569 | 0.435 | 0.299 | 0.047705493 | Mari |
| pDC | TIMM8B | 3.57E-11 | 0.819514008 | 0.622 | 0.432 | 7.03E-07 | Mari |
| pDC | TMEM14C | 1.21E-11 | 0.82404648 | 0.643 | 0.461 | 2.39E-07 | Mari |
| pDC | OCIAD2 | 1.19E-08 | 0.825406364 | 0.521 | 0.347 | 0.000233328 | Mari |
| pDC | SEC11A | 4.53E-16 | 0.825639354 | 0.783 | 0.58 | 8.92E-12 | Mari |
| pDC | PHF11 | 8.87E-07 | 0.826322668 | 0.414 | 0.273 | 0.017451274 | Mari |
| pDC | SDF2L1 | 9.24E-08 | 0.826351943 | 0.47 | 0.315 | 0.001818408 | Mari |
| pDC | PET100 | 8.25E-13 | 0.826834312 | 0.682 | 0.515 | 1.62E-08 | Mari |
| pDC | NDUFS7 | 5.29E-12 | 0.826842981 | 0.64 | 0.434 | 1.04E-07 | Mari |
| pDC | UQCRQ | 1.06E-18 | 0.826873538 | 0.786 | 0.568 | 2.09E-14 | Mari |
| pDC | ST6GALNAC4 | 4.92E-13 | 0.827197542 | 0.682 | 0.483 | 9.69E-09 | Mari |
| pDC | BST2 | 1.36E-20 | 0.827491547 | 0.881 | 0.737 | 2.67E-16 | Mari |
| pDC | UFD1 | 2.92E-08 | 0.828176232 | 0.506 | 0.345 | 0.00057498 | Mari |
| pDC | ATP5PF | 6.00E-22 | 0.828645834 | 0.863 | 0.661 | 1.18E-17 | Mari |
| pDC | PSMC5 | 9.51E-09 | 0.828730913 | 0.524 | 0.366 | 0.000187257 | Mari |
| pDC | TIMM13 | 1.66E-16 | 0.829483663 | 0.81 | 0.594 | 3.26E-12 | Mari |
| pDC | GADD45GIP1 | 1.42E-08 | 0.831155139 | 0.518 | 0.368 | 0.000280278 | Mari |
| pDC | GMFG | 1.05E-17 | 0.832475507 | 0.777 | 0.55 | 2.06E-13 | Mari |
| pDC | RPA3 | 5.17E-09 | 0.83569928 | 0.533 | 0.372 | 0.000101786 | Mari |
| pDC | FAM53C | 2.31E-09 | 0.836319734 | 0.577 | 0.396 | 4.54E-05 | Mari |
| pDC | GABARAP | 1.67E-35 | 0.836617845 | 0.973 | 0.889 | 3.30E-31 | Mari |
| pDC | RNASEH2C | 8.53E-10 | 0.836878515 | 0.548 | 0.374 | 1.68E-05 | Mari |
| pDC | HMGN3 | 9.67E-22 | 0.83836136 | 0.878 | 0.673 | 1.90E-17 | Mari |
| pDC | TSPO | 9.68E-09 | 0.83848293 | 0.515 | 0.364 | 0.000190486 | Mari |
| pDC | BAD | 1.96E-06 | 0.83888716 | 0.351 | 0.214 | 0.038510285 | Mari |
| pDC | LAMP5 | 3.35E-14 | 0.839158275 | 0.765 | 0.539 | 6.59E-10 | Mari |
| pDC | BOLA2 | 1.61E-08 | 0.840120524 | 0.482 | 0.315 | 0.000317625 | Mari |
| pDC | ATP5MF | 1.16E-24 | 0.840968662 | 0.89 | 0.729 | 2.29E-20 | Mari |
| pDC | ZBTB8OS | 1.30E-08 | 0.841249192 | 0.5 | 0.327 | 0.000255287 | Mari |
| pDC | UQCR10 | 4.62E-20 | 0.842466094 | 0.848 | 0.634 | 9.08E-16 | Mari |
| pDC | ARL4D | 3.86E-10 | 0.843196476 | 0.426 | 0.224 | 7.60E-06 | Mari |
| pDC | H3F3B | 4.68E-27 | 0.844480449 | 0.997 | 0.978 | 9.22E-23 | Mari |
| pDC | CNPY3 | 1.13E-19 | 0.845857255 | 0.848 | 0.671 | 2.23E-15 | Mari |
| pDC | SRSF9 | 2.55E-19 | 0.84818282 | 0.86 | 0.679 | 5.02E-15 | Mari |
| pDC | SNRNP25 | 8.76E-15 | 0.848936359 | 0.708 | 0.495 | 1.73E-10 | Mari |
| pDC | MRPL4 | 1.11E-08 | 0.849840525 | 0.429 | 0.257 | 0.000218995 | Mari |
| pDC | TOMM5 | 3.43E-13 | 0.84995868 | 0.685 | 0.479 | 6.75E-09 | Mari |
| pDC | UCHL3 | 2.39E-11 | 0.852111406 | 0.592 | 0.382 | 4.70E-07 | Mari |
| pDC | MRPL11 | 8.30E-08 | 0.852248391 | 0.417 | 0.259 | 0.001632947 | Mari |
| pDC | PRELID1 | 2.36E-23 | 0.853494479 | 0.86 | 0.6 | 4.64E-19 | Mari |
| pDC | BANF1 | 6.83E-10 | 0.857004275 | 0.557 | 0.38 | 1.35E-05 | Mari |
| pDC | DAB2 | 5.61E-10 | 0.857362223 | 0.5 | 0.305 | 1.10E-05 | Mari |
| pDC | PRKRA | 1.13E-06 | 0.858749067 | 0.414 | 0.271 | 0.022249494 | Mari |
| pDC | LSM5 | 2.34E-15 | 0.859002576 | 0.762 | 0.539 | 4.60E-11 | Mari |
| pDC | JCHAIN | 7.19E-33 | 0.862600914 | 0.988 | 0.931 | 1.42E-28 | Mari |
| pDC | SLC39A10 | 1.14E-11 | 0.863346674 | 0.562 | 0.349 | 2.24E-07 | Mari |
| pDC | COMMD6 | 3.15E-36 | 0.86411181 | 0.976 | 0.877 | 6.19E-32 | Mari |
| pDC | NDUFA4 | 1.57E-32 | 0.866420266 | 0.958 | 0.84 | 3.10E-28 | Mari |
| pDC | CHCHD5 | 3.54E-07 | 0.866845756 | 0.47 | 0.335 | 0.00696274 | Mari |
| pDC | VEGFB | 3.63E-24 | 0.867836447 | 0.929 | 0.764 | 7.15E-20 | Mari |
| pDC | USF2 | 8.76E-15 | 0.868688543 | 0.735 | 0.543 | 1.72E-10 | Mari |
| pDC | PDCD6 | 2.71E-15 | 0.869740708 | 0.688 | 0.455 | 5.33E-11 | Mari |
| pDC | CLIC3 | 6.74E-21 | 0.874165381 | 0.908 | 0.739 | 1.33E-16 | Mari |
| pDC | PHPT1 | 4.70E-16 | 0.875114042 | 0.771 | 0.564 | 9.24E-12 | Mari |

| | | | | | | | |
|---|---|---|---|---|---|---|---|
| pDC | PGLS | 1.34E-16 | 0.875681325 | 0.747 | 0.519 | 2.64E-12 | Mari |
| pDC | SOCS1 | 8.34E-16 | 0.875895713 | 0.705 | 0.465 | 1.64E-11 | Mari |
| pDC | ARF5 | 2.44E-11 | 0.876049432 | 0.631 | 0.465 | 4.81E-07 | Mari |
| pDC | PIN1 | 3.56E-10 | 0.876071162 | 0.539 | 0.368 | 7.01E-06 | Mari |
| pDC | GSTP1 | 2.37E-27 | 0.877344416 | 0.938 | 0.804 | 4.67E-23 | Mari |
| pDC | LAMTOR4 | 3.16E-19 | 0.877718665 | 0.78 | 0.564 | 6.23E-15 | Mari |
| pDC | SELENOT | 2.34E-15 | 0.884156026 | 0.741 | 0.533 | 4.61E-11 | Mari |
| pDC | KRT10 | 3.37E-23 | 0.885024537 | 0.845 | 0.606 | 6.64E-19 | Mari |
| pDC | DGUOK | 2.02E-13 | 0.885152412 | 0.652 | 0.467 | 3.99E-09 | Mari |
| pDC | ATP6V0B | 3.44E-30 | 0.886714016 | 0.952 | 0.756 | 6.77E-26 | Mari |
| pDC | HIST1H1C | 9.42E-09 | 0.888026586 | 0.524 | 0.341 | 0.000185427 | Mari |
| pDC | ADI1 | 6.03E-10 | 0.88861008 | 0.554 | 0.38 | 1.19E-05 | Mari |
| pDC | TMEM109 | 7.90E-12 | 0.893735871 | 0.607 | 0.396 | 1.55E-07 | Mari |
| pDC | SF3B6 | 3.24E-19 | 0.895789658 | 0.78 | 0.564 | 6.37E-15 | Mari |
| pDC | THYN1 | 7.01E-09 | 0.895905033 | 0.423 | 0.255 | 0.000138009 | Mari |
| pDC | TRAPPC6A | 1.03E-06 | 0.897778311 | 0.414 | 0.283 | 0.020328658 | Mari |
| pDC | AP2M1 | 1.07E-07 | 0.898624596 | 0.476 | 0.333 | 0.002097081 | Mari |
| pDC | TBCB | 1.73E-11 | 0.899353993 | 0.592 | 0.394 | 3.41E-07 | Mari |
| pDC | NDUFA9 | 1.49E-08 | 0.899839327 | 0.518 | 0.364 | 0.000293778 | Mari |
| pDC | AP1S2 | 6.77E-14 | 0.903439403 | 0.753 | 0.574 | 1.33E-09 | Mari |
| pDC | TMED3 | 2.57E-09 | 0.903543893 | 0.494 | 0.317 | 5.07E-05 | Mari |
| pDC | CHMP2A | 2.81E-12 | 0.903629985 | 0.592 | 0.394 | 5.54E-08 | Mari |
| pDC | GTF3C6 | 2.30E-10 | 0.903730469 | 0.533 | 0.366 | 4.53E-06 | Mari |
| pDC | LAMTOR2 | 2.98E-10 | 0.90556411 | 0.509 | 0.333 | 5.86E-06 | Mari |
| pDC | TRAPPC2L | 2.05E-09 | 0.905904979 | 0.488 | 0.321 | 4.03E-05 | Mari |
| pDC | EDF1 | 2.22E-34 | 0.90782601 | 0.961 | 0.812 | 4.37E-30 | Mari |
| pDC | GTF3A | 1.24E-22 | 0.908132237 | 0.842 | 0.624 | 2.45E-18 | Mari |
| pDC | VBP1 | 3.07E-08 | 0.908823744 | 0.432 | 0.275 | 0.000603962 | Mari |
| pDC | RBIS | 8.27E-19 | 0.912630783 | 0.815 | 0.63 | 1.63E-14 | Mari |
| pDC | GLCE | 2.04E-08 | 0.913334528 | 0.452 | 0.291 | 0.000401898 | Mari |
| pDC | MRPL57 | 5.87E-12 | 0.914127543 | 0.637 | 0.453 | 1.16E-07 | Mari |
| pDC | PTMS | 1.78E-16 | 0.915138214 | 0.75 | 0.531 | 3.51E-12 | Mari |
| pDC | RCN2 | 6.10E-08 | 0.91596393 | 0.405 | 0.246 | 0.001200658 | Mari |
| pDC | VPS29 | 1.53E-17 | 0.916115008 | 0.75 | 0.549 | 3.02E-13 | Mari |
| pDC | ARHGDIB | 3.77E-17 | 0.917253197 | 0.812 | 0.616 | 7.41E-13 | Mari |
| pDC | SMIM19 | 4.20E-11 | 0.918110279 | 0.518 | 0.325 | 8.27E-07 | Mari |
| pDC | PDCD5 | 4.79E-11 | 0.918232108 | 0.473 | 0.275 | 9.43E-07 | Mari |
| pDC | AKR7A2 | 2.57E-07 | 0.918492945 | 0.47 | 0.335 | 0.005067904 | Mari |
| pDC | RRP36 | 1.77E-06 | 0.918765559 | 0.292 | 0.16 | 0.034939023 | Mari |
| pDC | TXNDC17 | 1.36E-13 | 0.918912837 | 0.548 | 0.309 | 2.67E-09 | Mari |
| pDC | JPT1 | 6.88E-12 | 0.919018436 | 0.509 | 0.295 | 1.35E-07 | Mari |
| pDC | SMDT1 | 2.40E-21 | 0.91910027 | 0.833 | 0.596 | 4.73E-17 | Mari |
| pDC | CUTA | 4.76E-12 | 0.920394432 | 0.568 | 0.364 | 9.37E-08 | Mari |
| pDC | COX7A2L | 3.80E-22 | 0.924107548 | 0.845 | 0.675 | 7.48E-18 | Mari |
| pDC | ARPC1B | 2.72E-33 | 0.924184414 | 0.958 | 0.82 | 5.35E-29 | Mari |
| pDC | MIEN1 | 7.50E-07 | 0.926322786 | 0.399 | 0.251 | 0.014760268 | Mari |
| pDC | BLOC1S2 | 1.49E-08 | 0.927496378 | 0.452 | 0.285 | 0.000292821 | Mari |
| pDC | DCXR | 7.01E-09 | 0.928178264 | 0.47 | 0.305 | 0.000138064 | Mari |
| pDC | SRI | 4.55E-14 | 0.929410182 | 0.658 | 0.43 | 8.95E-10 | Mari |
| pDC | MDH1 | 1.90E-09 | 0.93101043 | 0.554 | 0.388 | 3.75E-05 | Mari |
| pDC | SMIM5 | 1.52E-11 | 0.931401121 | 0.542 | 0.325 | 2.98E-07 | Mari |
| pDC | PRKACB | 3.70E-07 | 0.932891955 | 0.446 | 0.307 | 0.007291644 | Mari |
| pDC | SEPTIN1 | 1.05E-12 | 0.934731213 | 0.607 | 0.384 | 2.06E-08 | Mari |
| pDC | SCT | 8.73E-22 | 0.935734018 | 0.917 | 0.764 | 1.72E-17 | Mari |
| pDC | CAPZA2 | 2.16E-14 | 0.935769455 | 0.607 | 0.364 | 4.26E-10 | Mari |
| pDC | SRM | 1.39E-07 | 0.936396831 | 0.435 | 0.295 | 0.002736264 | Mari |

| | | | | | | | |
|---|---|---|---|---|---|---|---|
| pDC | HSPB11 | 1.03E-12 | 0.939130671 | 0.568 | 0.356 | 2.02E-08 | Mari |
| pDC | METTL23 | 2.01E-09 | 0.940010141 | 0.461 | 0.287 | 3.95E-05 | Mari |
| pDC | GTF2H5 | 4.13E-10 | 0.94007373 | 0.47 | 0.287 | 8.14E-06 | Mari |
| pDC | ARL2 | 2.41E-06 | 0.940536512 | 0.333 | 0.21 | 0.047478491 | Mari |
| pDC | TMCO1 | 2.61E-14 | 0.940699177 | 0.676 | 0.477 | 5.14E-10 | Mari |
| pDC | HIST1H1E | 2.67E-07 | 0.942648133 | 0.348 | 0.21 | 0.005253364 | Mari |
| pDC | MCTS1 | 6.28E-10 | 0.943234374 | 0.497 | 0.315 | 1.24E-05 | Mari |
| pDC | SVIP | 1.17E-10 | 0.946468764 | 0.53 | 0.337 | 2.31E-06 | Mari |
| pDC | MYL12A | 1.25E-46 | 0.94723526 | 0.982 | 0.937 | 2.46E-42 | Mari |
| pDC | ZNF511 | 1.66E-07 | 0.948592749 | 0.354 | 0.214 | 0.003265405 | Mari |
| pDC | NHP2 | 9.88E-17 | 0.949653684 | 0.708 | 0.475 | 1.94E-12 | Mari |
| pDC | MRPL20 | 5.91E-16 | 0.950739496 | 0.72 | 0.515 | 1.16E-11 | Mari |
| pDC | FIBP | 2.46E-07 | 0.95152314 | 0.417 | 0.277 | 0.004843809 | Mari |
| pDC | DNAJC19 | 5.99E-12 | 0.951863905 | 0.512 | 0.309 | 1.18E-07 | Mari |
| pDC | PSMA3 | 1.51E-20 | 0.95248305 | 0.792 | 0.58 | 2.97E-16 | Mari |
| pDC | TMEM230 | 9.81E-17 | 0.953887072 | 0.693 | 0.45 | 1.93E-12 | Mari |
| pDC | GTF2A2 | 2.94E-18 | 0.955178954 | 0.708 | 0.475 | 5.79E-14 | Mari |
| pDC | NUDCD2 | 9.50E-09 | 0.958138004 | 0.464 | 0.299 | 0.000186962 | Mari |
| pDC | GLRX | 2.94E-13 | 0.958958595 | 0.568 | 0.352 | 5.78E-09 | Mari |
| pDC | PYCARD | 3.90E-24 | 0.959867497 | 0.878 | 0.657 | 7.67E-20 | Mari |
| pDC | UQCR11 | 1.44E-34 | 0.962868705 | 0.917 | 0.745 | 2.83E-30 | Mari |
| pDC | GCA | 4.76E-11 | 0.968439953 | 0.503 | 0.307 | 9.36E-07 | Mari |
| pDC | NCF4 | 1.38E-07 | 0.969072956 | 0.405 | 0.257 | 0.002710259 | Mari |
| pDC | CTSC | 5.48E-29 | 0.972230297 | 0.973 | 0.83 | 1.08E-24 | Mari |
| pDC | NDUFB7 | 9.23E-19 | 0.972413579 | 0.711 | 0.463 | 1.82E-14 | Mari |
| pDC | PUF60 | 6.43E-07 | 0.974120923 | 0.366 | 0.228 | 0.012658321 | Mari |
| pDC | RNASE6 | 1.10E-20 | 0.977073834 | 0.795 | 0.552 | 2.16E-16 | Mari |
| pDC | S100A4 | 1.04E-17 | 0.979663427 | 0.815 | 0.622 | 2.05E-13 | Mari |
| pDC | CYBC1 | 4.89E-09 | 0.98040864 | 0.461 | 0.299 | 9.63E-05 | Mari |
| pDC | ATP6V0E1 | 4.03E-33 | 0.981427921 | 0.94 | 0.768 | 7.93E-29 | Mari |
| pDC | S100A6 | 1.36E-33 | 0.985402934 | 0.976 | 0.889 | 2.68E-29 | Mari |
| pDC | MRPS18C | 1.53E-08 | 0.985622612 | 0.42 | 0.261 | 0.000301162 | Mari |
| pDC | NSMCE4A | 2.16E-07 | 0.988033261 | 0.366 | 0.22 | 0.004257836 | Mari |
| pDC | NIT2 | 2.01E-06 | 0.989642798 | 0.25 | 0.129 | 0.039620125 | Mari |
| pDC | MICOS13 | 2.03E-16 | 0.991920953 | 0.679 | 0.453 | 4.00E-12 | Mari |
| pDC | TMBIM4 | 8.57E-19 | 0.992518264 | 0.789 | 0.554 | 1.69E-14 | Mari |
| pDC | EIF4EBP3 | 1.49E-06 | 0.992589298 | 0.312 | 0.18 | 0.029421767 | Mari |
| pDC | SDHAF3 | 2.24E-16 | 0.996809367 | 0.649 | 0.378 | 4.40E-12 | Mari |
| pDC | ECH1 | 9.81E-10 | 0.997058986 | 0.473 | 0.285 | 1.93E-05 | Mari |
| pDC | CNPY2 | 1.23E-10 | 0.997214594 | 0.473 | 0.289 | 2.41E-06 | Mari |
| pDC | RER1 | 5.79E-19 | 0.997432718 | 0.717 | 0.481 | 1.14E-14 | Mari |
| pDC | NDUFAB1 | 1.07E-22 | 0.997774972 | 0.756 | 0.485 | 2.11E-18 | Mari |
| pDC | SSNA1 | 1.28E-12 | 0.998341198 | 0.58 | 0.4 | 2.51E-08 | Mari |
| pDC | PYURF | 6.38E-12 | 0.998849667 | 0.533 | 0.339 | 1.26E-07 | Mari |
| pDC | COPZ1 | 2.23E-12 | 1.000999248 | 0.533 | 0.329 | 4.39E-08 | Mari |
| pDC | PLAAT3 | 2.27E-20 | 1.002633493 | 0.801 | 0.549 | 4.46E-16 | Mari |
| pDC | POLR2C | 1.83E-06 | 1.003660295 | 0.289 | 0.16 | 0.036046706 | Mari |
| pDC | CCDC189 | 1.22E-08 | 1.004438612 | 0.31 | 0.152 | 0.000240143 | Mari |
| pDC | SNX17 | 2.58E-13 | 1.006884173 | 0.533 | 0.297 | 5.07E-09 | Mari |
| pDC | UQCC2 | 1.16E-08 | 1.007474173 | 0.393 | 0.23 | 0.000227863 | Mari |
| pDC | SCP2 | 4.79E-15 | 1.011471138 | 0.604 | 0.382 | 9.43E-11 | Mari |
| pDC | TIMMDC1 | 4.54E-07 | 1.012115854 | 0.36 | 0.22 | 0.008943437 | Mari |
| pDC | NDUFC2 | 1.26E-26 | 1.012955108 | 0.842 | 0.624 | 2.49E-22 | Mari |
| pDC | NDUFB5 | 3.02E-15 | 1.014379433 | 0.619 | 0.386 | 5.94E-11 | Mari |
| pDC | SRP9 | 3.67E-21 | 1.014982659 | 0.738 | 0.473 | 7.23E-17 | Mari |
| pDC | MRPL3 | 8.53E-10 | 1.017516476 | 0.432 | 0.251 | 1.68E-05 | Mari |

| | | | | | | | |
|---|---|---|---|---|---|---|---|
| pDC | PRDX5 | 3.72E-24 | 1.020856589 | 0.851 | 0.584 | 7.33E-20 | Mari |
| pDC | NDUFAF3 | 8.67E-12 | 1.021442443 | 0.527 | 0.325 | 1.71E-07 | Mari |
| pDC | TRAPPC1 | 4.95E-23 | 1.02328606 | 0.783 | 0.505 | 9.74E-19 | Mari |
| pDC | TXNDC9 | 1.39E-06 | 1.026541824 | 0.321 | 0.198 | 0.027321067 | Mari |
| pDC | TNNI2 | 1.45E-06 | 1.02725141 | 0.315 | 0.18 | 0.028568807 | Mari |
| pDC | FAM174C | 3.63E-13 | 1.027682794 | 0.545 | 0.327 | 7.15E-09 | Mari |
| pDC | NUTF2 | 6.14E-14 | 1.029649633 | 0.482 | 0.246 | 1.21E-09 | Mari |
| pDC | MEAF6 | 5.92E-08 | 1.029970153 | 0.378 | 0.22 | 0.001164519 | Mari |
| pDC | CIAO2A | 2.66E-17 | 1.02997383 | 0.705 | 0.467 | 5.24E-13 | Mari |
| pDC | NDUFB3 | 5.48E-18 | 1.030455444 | 0.655 | 0.412 | 1.08E-13 | Mari |
| pDC | STARD13 | 9.12E-08 | 1.030500936 | 0.28 | 0.133 | 0.001796124 | Mari |
| pDC | TTC13 | 5.18E-09 | 1.030568146 | 0.443 | 0.265 | 0.000101996 | Mari |
| pDC | RBX1 | 8.28E-36 | 1.034163252 | 0.923 | 0.721 | 1.63E-31 | Mari |
| pDC | DYNLRB1 | 1.38E-18 | 1.0354328 | 0.673 | 0.402 | 2.71E-14 | Mari |
| pDC | CHMP5 | 5.99E-15 | 1.036564915 | 0.661 | 0.455 | 1.18E-10 | Mari |
| pDC | SERPINF1 | 1.47E-38 | 1.036738197 | 0.982 | 0.869 | 2.89E-34 | Mari |
| pDC | EBNA1BP2 | 3.90E-07 | 1.038545079 | 0.28 | 0.145 | 0.007675708 | Mari |
| pDC | FAM89B | 1.06E-09 | 1.040464198 | 0.426 | 0.257 | 2.09E-05 | Mari |
| pDC | IRF7 | 6.34E-47 | 1.041902312 | 0.988 | 0.941 | 1.25E-42 | Mari |
| pDC | LYSMD2 | 4.42E-17 | 1.041953704 | 0.705 | 0.483 | 8.69E-13 | Mari |
| pDC | IAH1 | 1.86E-13 | 1.042456174 | 0.557 | 0.35 | 3.66E-09 | Mari |
| pDC | TMEM35B | 9.41E-08 | 1.048811523 | 0.387 | 0.24 | 0.001851667 | Mari |
| pDC | TMEM141 | 2.01E-09 | 1.05156528 | 0.512 | 0.35 | 3.96E-05 | Mari |
| pDC | CIB2 | 3.89E-15 | 1.053452696 | 0.625 | 0.386 | 7.67E-11 | Mari |
| pDC | LSM6 | 7.09E-12 | 1.05431785 | 0.491 | 0.289 | 1.39E-07 | Mari |
| pDC | NME4 | 6.02E-16 | 1.057229087 | 0.604 | 0.362 | 1.19E-11 | Mari |
| pDC | METTL5 | 7.03E-07 | 1.057868955 | 0.342 | 0.21 | 0.013832587 | Mari |
| pDC | SNRPN | 2.32E-21 | 1.058269652 | 0.807 | 0.552 | 4.56E-17 | Mari |
| pDC | PLCB4 | 5.53E-09 | 1.062800133 | 0.399 | 0.218 | 0.000108824 | Mari |
| pDC | ACAA1 | 1.22E-14 | 1.06281508 | 0.583 | 0.362 | 2.41E-10 | Mari |
| pDC | IMP4 | 1.00E-07 | 1.065007723 | 0.384 | 0.246 | 0.001973071 | Mari |
| pDC | PLGRKT | 4.36E-07 | 1.067066024 | 0.33 | 0.188 | 0.008588113 | Mari |
| pDC | CLEC4C | 6.45E-28 | 1.070023244 | 0.896 | 0.612 | 1.27E-23 | Mari |
| pDC | FCER1G | 1.17E-38 | 1.071668345 | 0.967 | 0.766 | 2.31E-34 | Mari |
| pDC | PLEKHJ1 | 1.00E-06 | 1.073479368 | 0.408 | 0.283 | 0.019774112 | Mari |
| pDC | CENPX | 3.12E-11 | 1.07412774 | 0.405 | 0.208 | 6.15E-07 | Mari |
| pDC | COMMD7 | 1.78E-06 | 1.077024706 | 0.295 | 0.17 | 0.035070136 | Mari |
| pDC | PRDX2 | 6.70E-07 | 1.080006231 | 0.318 | 0.186 | 0.013191288 | Mari |
| pDC | SMCO4 | 1.41E-14 | 1.081296776 | 0.536 | 0.301 | 2.77E-10 | Mari |
| pDC | NDUFS2 | 3.73E-10 | 1.085250604 | 0.485 | 0.323 | 7.35E-06 | Mari |
| pDC | LRRC26 | 1.27E-27 | 1.087176639 | 0.872 | 0.642 | 2.50E-23 | Mari |
| pDC | CD69 | 3.09E-10 | 1.090499672 | 0.598 | 0.426 | 6.07E-06 | Mari |
| pDC | POLR2I | 2.34E-13 | 1.090518751 | 0.53 | 0.331 | 4.61E-09 | Mari |
| pDC | IER2 | 8.91E-21 | 1.090704652 | 0.833 | 0.604 | 1.75E-16 | Mari |
| pDC | CUL4B | 1.37E-06 | 1.090827346 | 0.289 | 0.162 | 0.026994918 | Mari |
| pDC | GAS6 | 3.08E-40 | 1.093991006 | 0.946 | 0.737 | 6.06E-36 | Mari |
| pDC | DNASE2 | 8.53E-07 | 1.094169801 | 0.31 | 0.178 | 0.016794735 | Mari |
| pDC | SLC25A39 | 1.15E-06 | 1.094431993 | 0.283 | 0.156 | 0.022708652 | Mari |
| pDC | LAMTOR1 | 7.68E-21 | 1.095511914 | 0.726 | 0.501 | 1.51E-16 | Mari |
| pDC | MTCH2 | 2.41E-09 | 1.098026552 | 0.402 | 0.232 | 4.75E-05 | Mari |
| pDC | ARHGAP24 | 1.27E-10 | 1.101901529 | 0.324 | 0.139 | 2.50E-06 | Mari |
| pDC | NUBP2 | 1.33E-09 | 1.10627583 | 0.393 | 0.222 | 2.62E-05 | Mari |
| pDC | UROS | 4.00E-08 | 1.10694087 | 0.372 | 0.22 | 0.00078688 | Mari |
| pDC | MGAT4A | 8.16E-14 | 1.107725793 | 0.583 | 0.362 | 1.61E-09 | Mari |
| pDC | COPS9 | 2.75E-20 | 1.118802495 | 0.702 | 0.469 | 5.41E-16 | Mari |
| pDC | PPP1R14B | 2.57E-53 | 1.122415115 | 0.973 | 0.935 | 5.06E-49 | Mari |

| | | | | | | | |
|---|---|---|---|---|---|---|---|
| pDC | NDUFB2 | 2.19E-30 | 1.123550353 | 0.869 | 0.624 | 4.31E-26 | Mari |
| pDC | MICOS10 | 6.35E-27 | 1.124757988 | 0.812 | 0.541 | 1.25E-22 | Mari |
| pDC | GZMB | 7.04E-65 | 1.125117774 | 1 | 0.947 | 1.39E-60 | Mari |
| pDC | RASGRP2 | 4.53E-12 | 1.130702239 | 0.491 | 0.293 | 8.92E-08 | Mari |
| pDC | JKAMP | 1.19E-08 | 1.133989086 | 0.372 | 0.214 | 0.000234427 | Mari |
| pDC | UBLCP1 | 1.87E-06 | 1.135750863 | 0.33 | 0.208 | 0.036847646 | Mari |
| pDC | COA5 | 4.26E-08 | 1.138923103 | 0.339 | 0.192 | 0.000837776 | Mari |
| pDC | S100A11 | 3.09E-28 | 1.140540237 | 0.896 | 0.648 | 6.09E-24 | Mari |
| pDC | ZDHHC12 | 4.85E-09 | 1.142120465 | 0.366 | 0.214 | 9.55E-05 | Mari |
| pDC | NCBP2AS2 | 2.64E-08 | 1.143561521 | 0.351 | 0.2 | 0.000518987 | Mari |
| pDC | POLE4 | 5.66E-12 | 1.147784167 | 0.509 | 0.313 | 1.11E-07 | Mari |
| pDC | COMMD8 | 1.70E-08 | 1.150146983 | 0.354 | 0.192 | 0.000333992 | Mari |
| pDC | GAPT | 1.37E-17 | 1.155470328 | 0.682 | 0.434 | 2.70E-13 | Mari |
| pDC | TSTA3 | 1.14E-08 | 1.160257829 | 0.33 | 0.176 | 0.000224031 | Mari |
| pDC | CHCHD1 | 1.35E-10 | 1.164350583 | 0.405 | 0.232 | 2.66E-06 | Mari |
| pDC | EIF4EBP1 | 5.36E-08 | 1.16456044 | 0.345 | 0.196 | 0.001055432 | Mari |
| pDC | PLAC8 | 2.92E-51 | 1.167435057 | 0.982 | 0.889 | 5.74E-47 | Mari |
| pDC | COA4 | 4.41E-11 | 1.169956614 | 0.482 | 0.305 | 8.69E-07 | Mari |
| pDC | CYP4V2 | 1.41E-06 | 1.172550312 | 0.369 | 0.246 | 0.027790331 | Mari |
| pDC | PDLIM2 | 7.81E-09 | 1.175350917 | 0.348 | 0.19 | 0.000153677 | Mari |
| pDC | PDXP | 6.68E-11 | 1.180856409 | 0.357 | 0.168 | 1.32E-06 | Mari |
| pDC | MRPL42 | 3.36E-07 | 1.18786892 | 0.274 | 0.141 | 0.006611479 | Mari |
| pDC | CTSA | 1.66E-06 | 1.190374779 | 0.268 | 0.147 | 0.032634507 | Mari |
| pDC | S100A8 | 2.97E-09 | 1.194062713 | 0.473 | 0.315 | 5.84E-05 | Mari |
| pDC | KRT16 | 3.58E-08 | 1.195247592 | 0.304 | 0.158 | 0.000705372 | Mari |
| pDC | TMEM147 | 4.30E-12 | 1.197492212 | 0.429 | 0.23 | 8.46E-08 | Mari |
| pDC | STMP1 | 4.49E-13 | 1.210114338 | 0.485 | 0.273 | 8.84E-09 | Mari |
| pDC | FLOT1 | 1.19E-15 | 1.220452336 | 0.565 | 0.35 | 2.35E-11 | Mari |
| pDC | TM2D1 | 2.51E-11 | 1.255806963 | 0.435 | 0.246 | 4.94E-07 | Mari |
| pDC | NAE1 | 1.04E-09 | 1.258954389 | 0.283 | 0.125 | 2.05E-05 | Mari |
| pDC | DNAJC15 | 1.22E-20 | 1.263125581 | 0.652 | 0.392 | 2.41E-16 | Mari |
| pDC | MS4A6A | 1.09E-33 | 1.26525817 | 0.893 | 0.624 | 2.15E-29 | Mari |
| pDC | MRPL40 | 4.61E-11 | 1.267985309 | 0.402 | 0.218 | 9.06E-07 | Mari |
| pDC | RBCK1 | 1.09E-14 | 1.268585233 | 0.524 | 0.307 | 2.15E-10 | Mari |
| pDC | HINT2 | 2.88E-19 | 1.276087949 | 0.619 | 0.341 | 5.66E-15 | Mari |
| pDC | MTLN | 9.08E-08 | 1.28441183 | 0.301 | 0.16 | 0.001786923 | Mari |
| pDC | BLOC1S1 | 4.07E-27 | 1.287247593 | 0.714 | 0.4 | 8.00E-23 | Mari |
| pDC | CKLF | 1.22E-16 | 1.289120915 | 0.509 | 0.253 | 2.40E-12 | Mari |
| pDC | SUSD1 | 4.13E-17 | 1.290685584 | 0.607 | 0.349 | 8.12E-13 | Mari |
| pDC | ITGB1BP1 | 1.11E-09 | 1.294461038 | 0.348 | 0.18 | 2.18E-05 | Mari |
| pDC | PUS10 | 4.10E-08 | 1.306588278 | 0.53 | 0.37 | 0.000807156 | Mari |
| pDC | HIGD1A | 3.27E-34 | 1.314443505 | 0.872 | 0.598 | 6.44E-30 | Mari |
| pDC | PPP1R14A | 2.72E-08 | 1.318696042 | 0.369 | 0.208 | 0.000534564 | Mari |
| pDC | ISCA2 | 1.17E-12 | 1.341068073 | 0.393 | 0.188 | 2.29E-08 | Mari |
| pDC | GNG10 | 2.73E-15 | 1.34490348 | 0.524 | 0.293 | 5.38E-11 | Mari |
| pDC | IPCEF1 | 2.94E-08 | 1.346645143 | 0.277 | 0.129 | 0.00057925 | Mari |
| pDC | EEF1E1 | 1.22E-12 | 1.348926751 | 0.402 | 0.202 | 2.41E-08 | Mari |
| pDC | VPS25 | 2.08E-11 | 1.350741766 | 0.366 | 0.186 | 4.10E-07 | Mari |
| pDC | NUDT22 | 4.28E-24 | 1.356723478 | 0.679 | 0.406 | 8.42E-20 | Mari |
| pDC | TCL1A | 8.13E-08 | 1.37031517 | 0.467 | 0.337 | 0.001599414 | Mari |
| pDC | NDUFAF8 | 2.52E-20 | 1.382248412 | 0.58 | 0.319 | 4.96E-16 | Mari |
| pDC | MS4A4A | 2.26E-10 | 1.39914495 | 0.432 | 0.267 | 4.45E-06 | Mari |
| pDC | PTPMT1 | 2.52E-13 | 1.400615203 | 0.449 | 0.248 | 4.96E-09 | Mari |
| pDC | TLR10 | 3.15E-08 | 1.40279539 | 0.265 | 0.125 | 0.00061994 | Mari |
| pDC | NTAN1 | 1.03E-12 | 1.434736128 | 0.378 | 0.178 | 2.04E-08 | Mari |
| pDC | RGS18 | 1.68E-10 | 1.444737971 | 0.342 | 0.168 | 3.31E-06 | Mari |

| | | | | | | | |
|---|---|---|---|---|---|---|---|
| pDC | BIN1 | 2.94E-08 | 1.482905305 | 0.277 | 0.135 | 0.000578941 | Mari |
| pDC | PROC | 8.54E-17 | 1.511565649 | 0.521 | 0.281 | 1.68E-12 | Mari |
| pDC | PTP4A3 | 1.43E-17 | 1.53530387 | 0.429 | 0.172 | 2.81E-13 | Mari |
| pDC | SHD | 4.59E-23 | 1.594435255 | 0.619 | 0.327 | 9.03E-19 | Mari |
| pDC | ARRDC3 | 2.53E-19 | 1.604625175 | 0.58 | 0.329 | 4.98E-15 | Mari |
| pDC | OAS1 | 2.79E-20 | 1.643905376 | 0.494 | 0.212 | 5.49E-16 | Mari |
| pDC | DRD4 | 2.49E-19 | 1.724685388 | 0.539 | 0.267 | 4.89E-15 | Mari |
| pDC | SH2B31 | 5.38E-07 | 0.501105737 | 0.723 | 0.545 | 0.010592588 | Tb |
| pDC | SKIL | 2.06E-06 | 0.516461493 | 0.77 | 0.642 | 0.040479415 | Tb |
| pDC | JOSD1 | 2.84E-07 | 0.517633611 | 0.689 | 0.501 | 0.005595713 | Tb |
| pDC | AP3B1 | 1.63E-06 | 0.533345291 | 0.632 | 0.446 | 0.0321441 | Tb |
| pDC | CCDC88A | 1.36E-06 | 0.538844252 | 0.814 | 0.709 | 0.026772985 | Tb |
| pDC | SIK3 | 2.23E-06 | 0.539640418 | 0.761 | 0.62 | 0.043880658 | Tb |
| pDC | WDFY2 | 4.23E-07 | 0.540048055 | 0.648 | 0.476 | 0.008326973 | Tb |
| pDC | MAP4K4 | 9.85E-07 | 0.540622871 | 0.777 | 0.637 | 0.019393712 | Tb |
| pDC | GNAQ | 2.15E-06 | 0.546532162 | 0.692 | 0.545 | 0.042255749 | Tb |
| pDC | DNAJB12 | 1.70E-09 | 0.54901626 | 0.846 | 0.69 | 3.35E-05 | Tb |
| pDC | BBX | 1.43E-06 | 0.554158573 | 0.597 | 0.415 | 0.028051078 | Tb |
| pDC | ASH1L | 6.12E-07 | 0.568760052 | 0.623 | 0.468 | 0.012051057 | Tb |
| pDC | NEK7 | 4.18E-07 | 0.57387438 | 0.377 | 0.203 | 0.008223271 | Tb |
| pDC | TNPO3 | 1.72E-06 | 0.584983146 | 0.365 | 0.203 | 0.033937512 | Tb |
| pDC | UBE2H | 1.48E-06 | 0.609701232 | 0.638 | 0.449 | 0.029073354 | Tb |
| pDC | TBC1D22A | 3.00E-07 | 0.609747024 | 0.519 | 0.34 | 0.005910876 | Tb |
| pDC | CGAS | 1.78E-07 | 0.62253269 | 0.491 | 0.298 | 0.003509556 | Tb |
| pDC | SLC25A25 | 4.17E-08 | 0.628427868 | 0.362 | 0.174 | 0.000820371 | Tb |
| pDC | CARD11 | 4.98E-07 | 0.632790688 | 0.752 | 0.625 | 0.009811864 | Tb |
| pDC | PAG1 | 1.01E-07 | 0.63697856 | 0.77 | 0.627 | 0.001982133 | Tb |
| pDC | RBM47 | 2.31E-06 | 0.639721128 | 0.594 | 0.446 | 0.0454051 | Tb |
| pDC | RNF10 | 2.07E-06 | 0.640959585 | 0.437 | 0.26 | 0.04078267 | Tb |
| pDC | GYS1 | 1.49E-07 | 0.644614256 | 0.484 | 0.291 | 0.002939307 | Tb |
| pDC | RABGEF1 | 7.34E-07 | 0.647753783 | 0.447 | 0.272 | 0.014442501 | Tb |
| pDC | UPF3B | 5.75E-07 | 0.655240934 | 0.415 | 0.233 | 0.011324087 | Tb |
| pDC | RYBP | 1.01E-07 | 0.657686053 | 0.66 | 0.472 | 0.001986292 | Tb |
| pDC | HSPA1A1 | 1.27E-10 | 0.658245743 | 0.921 | 0.809 | 2.49E-06 | Tb |
| pDC | SFMBT2 | 3.78E-07 | 0.665221076 | 0.717 | 0.591 | 0.007435077 | Tb |
| pDC | PBRM1 | 9.15E-07 | 0.667284983 | 0.742 | 0.589 | 0.018009641 | Tb |
| pDC | ITCH | 1.38E-06 | 0.6680975 | 0.642 | 0.499 | 0.027154949 | Tb |
| pDC | MT-CYB | 7.93E-10 | 0.670762989 | 0.915 | 0.845 | 1.56E-05 | Tb |
| pDC | PRKCA | 5.96E-10 | 0.671822601 | 0.66 | 0.449 | 1.17E-05 | Tb |
| pDC | ARID1B | 1.83E-10 | 0.674293938 | 0.774 | 0.602 | 3.60E-06 | Tb |
| pDC | MT-ND3 | 1.33E-16 | 0.682054867 | 0.978 | 0.887 | 2.61E-12 | Tb |
| pDC | RHOH | 1.68E-06 | 0.686437059 | 0.654 | 0.524 | 0.033145494 | Tb |
| pDC | EIF2AK3 | 2.18E-09 | 0.690529373 | 0.503 | 0.285 | 4.29E-05 | Tb |
| pDC | TYMP | 1.18E-07 | 0.692510738 | 0.736 | 0.551 | 0.002326281 | Tb |
| pDC | RALGAPA1 | 2.37E-08 | 0.69389876 | 0.789 | 0.642 | 0.000465744 | Tb |
| pDC | ASB3 | 2.13E-06 | 0.696059165 | 0.343 | 0.185 | 0.041865405 | Tb |
| pDC | ADPGK | 9.37E-07 | 0.700740713 | 0.45 | 0.27 | 0.018443412 | Tb |
| pDC | RC3H1 | 8.20E-07 | 0.705228613 | 0.403 | 0.229 | 0.016139931 | Tb |
| pDC | HSPD11 | 5.27E-10 | 0.7100604 | 0.912 | 0.815 | 1.04E-05 | Tb |
| pDC | COP1 | 8.07E-07 | 0.719700174 | 0.535 | 0.377 | 0.015879116 | Tb |
| pDC | AHSA11 | 5.10E-08 | 0.721380779 | 0.623 | 0.447 | 0.001004211 | Tb |
| pDC | MT-CO3 | 1.48E-08 | 0.721914464 | 0.947 | 0.883 | 0.000290866 | Tb |
| pDC | RREB1 | 1.56E-06 | 0.722823342 | 0.645 | 0.509 | 0.030737186 | Tb |
| pDC | TRAF3 | 2.23E-09 | 0.736280751 | 0.726 | 0.528 | 4.38E-05 | Tb |
| pDC | RBM6 | 1.76E-07 | 0.740346232 | 0.645 | 0.489 | 0.003473307 | Tb |
| pDC | STIM1 | 2.39E-06 | 0.746001748 | 0.453 | 0.294 | 0.047032641 | Tb |

| | | | | | | | |
|---|---|---|---|---|---|---|---|
| pDC | DYRK1A | 6.01E-10 | 0.747772161 | 0.56 | 0.344 | 1.18E-05 | Tb |
| pDC | ELL21 | 1.82E-06 | 0.75054041 | 0.664 | 0.539 | 0.035823502 | Tb |
| pDC | CFLAR | 3.74E-07 | 0.760385592 | 0.645 | 0.499 | 0.007371591 | Tb |
| pDC | CEP192 | 1.59E-06 | 0.764528487 | 0.274 | 0.134 | 0.031389377 | Tb |
| pDC | MIR4435-2HG | 1.57E-08 | 0.769626052 | 0.594 | 0.396 | 0.000309463 | Tb |
| pDC | SRGAP2 | 4.28E-07 | 0.77643014 | 0.519 | 0.34 | 0.008434152 | Tb |
| pDC | KDM6A | 5.66E-09 | 0.777862115 | 0.569 | 0.371 | 0.000111399 | Tb |
| pDC | LPIN1 | 2.07E-09 | 0.779297639 | 0.651 | 0.438 | 4.07E-05 | Tb |
| pDC | PEAK1 | 9.22E-07 | 0.782040543 | 0.406 | 0.25 | 0.018148503 | Tb |
| pDC | PCSK7 | 1.90E-06 | 0.78567026 | 0.261 | 0.126 | 0.037456389 | Tb |
| pDC | KANSL1 | 2.47E-08 | 0.788075805 | 0.509 | 0.317 | 0.000485895 | Tb |
| pDC | IL6ST | 9.40E-09 | 0.790985695 | 0.638 | 0.43 | 0.000185061 | Tb |
| pDC | SWAP70 | 5.45E-07 | 0.792049965 | 0.27 | 0.126 | 0.0107347 | Tb |
| pDC | SLC9A7 | 1.88E-06 | 0.794005606 | 0.513 | 0.344 | 0.036996888 | Tb |
| pDC | MAP2K1 | 1.71E-09 | 0.798516269 | 0.547 | 0.333 | 3.37E-05 | Tb |
| pDC | CERS6 | 2.63E-13 | 0.800080298 | 0.739 | 0.556 | 5.17E-09 | Tb |
| pDC | CTDP1 | 1.44E-09 | 0.800261079 | 0.513 | 0.293 | 2.84E-05 | Tb |
| pDC | SDK2 | 2.48E-06 | 0.813935999 | 0.39 | 0.229 | 0.04887632 | Tb |
| pDC | CACYBP1 | 6.81E-07 | 0.819300191 | 0.66 | 0.514 | 0.013396826 | Tb |
| pDC | XIST | 2.00E-10 | 0.825421056 | 0.547 | 0.323 | 3.94E-06 | Tb |
| pDC | FTX | 2.71E-10 | 0.825424587 | 0.588 | 0.369 | 5.34E-06 | Tb |
| pDC | MYO9A | 8.54E-07 | 0.82868453 | 0.462 | 0.306 | 0.016804652 | Tb |
| pDC | ZNF518A | 1.49E-06 | 0.839365827 | 0.333 | 0.182 | 0.029354062 | Tb |
| pDC | SCAF4 | 8.69E-07 | 0.839624802 | 0.308 | 0.157 | 0.017096403 | Tb |
| pDC | FKBP41 | 6.57E-08 | 0.842646437 | 0.516 | 0.333 | 0.001292383 | Tb |
| pDC | FAM107B1 | 1.15E-07 | 0.854757139 | 0.733 | 0.579 | 0.002254212 | Tb |
| pDC | ARID2 | 2.84E-11 | 0.858121928 | 0.664 | 0.455 | 5.59E-07 | Tb |
| pDC | TRAK1 | 2.21E-06 | 0.866140979 | 0.418 | 0.268 | 0.043446595 | Tb |
| pDC | SPATS2 | 1.26E-07 | 0.867890821 | 0.484 | 0.283 | 0.002481239 | Tb |
| pDC | MT-ATP6 | 5.51E-16 | 0.868528289 | 0.984 | 0.904 | 1.08E-11 | Tb |
| pDC | GAB1 | 5.25E-09 | 0.86968989 | 0.701 | 0.554 | 0.000103347 | Tb |
| pDC | ZER1 | 3.68E-07 | 0.87585143 | 0.283 | 0.136 | 0.007250945 | Tb |
| pDC | XYLT1 | 1.46E-09 | 0.876900325 | 0.513 | 0.308 | 2.88E-05 | Tb |
| pDC | TNFAIP8 | 5.34E-08 | 0.878176453 | 0.582 | 0.4 | 0.001051863 | Tb |
| pDC | SLMAP | 4.75E-07 | 0.879654945 | 0.368 | 0.208 | 0.009342526 | Tb |
| pDC | PHACTR4 | 9.93E-07 | 0.887308936 | 0.327 | 0.176 | 0.019549257 | Tb |
| pDC | SMAD3 | 5.34E-08 | 0.887706991 | 0.34 | 0.168 | 0.001052057 | Tb |
| pDC | ATP6V0A1 | 7.41E-07 | 0.891275 | 0.343 | 0.191 | 0.014585448 | Tb |
| pDC | CLEC16A | 9.77E-07 | 0.891418636 | 0.349 | 0.193 | 0.019229245 | Tb |
| pDC | GLUL | 1.42E-07 | 0.893822275 | 0.399 | 0.228 | 0.002792099 | Tb |
| pDC | FAM118A | 2.14E-06 | 0.898887408 | 0.289 | 0.153 | 0.042196898 | Tb |
| pDC | PAK1 | 3.65E-10 | 0.902922564 | 0.642 | 0.413 | 7.19E-06 | Tb |
| pDC | GABPB1 | 1.13E-07 | 0.907817285 | 0.327 | 0.164 | 0.002224769 | Tb |
| pDC | MARK3 | 7.20E-07 | 0.911346074 | 0.437 | 0.281 | 0.014181694 | Tb |
| pDC | ETFDH | 9.49E-09 | 0.912109798 | 0.302 | 0.132 | 0.00018679 | Tb |
| pDC | THADA | 2.21E-08 | 0.91336745 | 0.462 | 0.272 | 0.000434503 | Tb |
| pDC | FOXN3 | 2.36E-06 | 0.922638427 | 0.525 | 0.373 | 0.046415912 | Tb |
| pDC | MED13L | 2.93E-15 | 0.926243032 | 0.84 | 0.665 | 5.76E-11 | Tb |
| pDC | PDE4A | 4.56E-08 | 0.92777183 | 0.553 | 0.373 | 0.000898214 | Tb |
| pDC | CALU | 4.84E-07 | 0.933898049 | 0.39 | 0.224 | 0.009529423 | Tb |
| pDC | DENND4A | 7.58E-14 | 0.945339156 | 0.836 | 0.642 | 1.49E-09 | Tb |
| pDC | CERS4 | 6.90E-07 | 0.998923427 | 0.267 | 0.132 | 0.01358099 | Tb |
| pDC | FGD2 | 7.32E-08 | 1.001347913 | 0.365 | 0.191 | 0.001441767 | Tb |
| pDC | S100A2 | 4.96E-08 | 1.004859585 | 0.28 | 0.122 | 0.000976809 | Tb |
| pDC | SCAF8 | 5.21E-08 | 1.016230437 | 0.362 | 0.184 | 0.001025885 | Tb |
| pDC | CCND2 | 6.10E-08 | 1.017298838 | 0.365 | 0.193 | 0.001201416 | Tb |

| | | | | | | | |
|---|---|---|---|---|---|---|---|
| pDC | PLCB1 | 5.78E-10 | 1.017721621 | 0.355 | 0.166 | 1.14E-05 | Tb |
| pDC | CUX2 | 1.69E-07 | 1.019648076 | 0.531 | 0.377 | 0.003329598 | Tb |
| pDC | CBFB | 1.86E-08 | 1.021722921 | 0.349 | 0.174 | 0.000366362 | Tb |
| pDC | ST3GAL1 | 3.36E-07 | 1.030487026 | 0.368 | 0.205 | 0.006606746 | Tb |
| pDC | RASA1 | 2.77E-08 | 1.034339378 | 0.406 | 0.231 | 0.000545448 | Tb |
| pDC | NLRC5 | 1.56E-06 | 1.035964393 | 0.314 | 0.168 | 0.030803045 | Tb |
| pDC | ABHD3 | 3.28E-12 | 1.036340684 | 0.484 | 0.245 | 6.46E-08 | Tb |
| pDC | CCDC6 | 3.53E-07 | 1.045722795 | 0.299 | 0.149 | 0.006938757 | Tb |
| pDC | CREB3L2 | 6.00E-09 | 1.056580327 | 0.503 | 0.306 | 0.000118076 | Tb |
| pDC | FBXO11 | 8.45E-12 | 1.056617259 | 0.619 | 0.419 | 1.66E-07 | Tb |
| pDC | NBEAL1 | 4.13E-11 | 1.05713434 | 0.557 | 0.333 | 8.13E-07 | Tb |
| pDC | KLHL20 | 6.06E-07 | 1.059163255 | 0.289 | 0.147 | 0.011930986 | Tb |
| pDC | BABAM2 | 1.16E-08 | 1.060012635 | 0.472 | 0.291 | 0.000228321 | Tb |
| pDC | ENTPD1-AS1 | 9.20E-07 | 1.067220316 | 0.261 | 0.126 | 0.018114192 | Tb |
| pDC | PHLPP1 | 4.39E-07 | 1.071528627 | 0.277 | 0.136 | 0.008636131 | Tb |
| pDC | ASCC1 | 1.18E-06 | 1.07791579 | 0.286 | 0.149 | 0.023129283 | Tb |
| pDC | WDR44 | 2.00E-06 | 1.083007813 | 0.252 | 0.122 | 0.039431629 | Tb |
| pDC | TFCP2 | 1.18E-07 | 1.084194218 | 0.311 | 0.153 | 0.002328085 | Tb |
| pDC | RASSF8 | 2.36E-07 | 1.087506396 | 0.277 | 0.128 | 0.004643389 | Tb |
| pDC | CNOT10 | 4.25E-07 | 1.095303427 | 0.277 | 0.134 | 0.008373772 | Tb |
| pDC | PWWP2A | 1.22E-07 | 1.113790799 | 0.274 | 0.126 | 0.002397585 | Tb |
| pDC | DGKD | 2.01E-06 | 1.125693111 | 0.358 | 0.21 | 0.039592024 | Tb |
| pDC | FAM222B | 3.42E-08 | 1.138965114 | 0.289 | 0.136 | 0.000673528 | Tb |
| pDC | BIRC3 | 9.61E-09 | 1.147601432 | 0.412 | 0.231 | 0.000189099 | Tb |
| pDC | PRKCE | 4.87E-08 | 1.167312111 | 0.403 | 0.235 | 0.000957793 | Tb |
| pDC | BMPR2 | 1.62E-09 | 1.175882645 | 0.387 | 0.197 | 3.19E-05 | Tb |
| pDC | MFN1 | 3.43E-08 | 1.191648826 | 0.311 | 0.151 | 0.000675318 | Tb |
| pDC | CLEC2D | 1.19E-06 | 1.195318346 | 0.349 | 0.199 | 0.023387686 | Tb |
| pDC | RAB12 | 1.16E-10 | 1.216409459 | 0.277 | 0.101 | 2.29E-06 | Tb |
| pDC | FAM193A | 1.38E-08 | 1.229989456 | 0.305 | 0.138 | 0.000271999 | Tb |
| pDC | CASP10 | 1.95E-08 | 1.232745258 | 0.336 | 0.164 | 0.000383996 | Tb |
| pDC | FAF2 | 4.31E-08 | 1.236192016 | 0.258 | 0.113 | 0.000847887 | Tb |
| pDC | PPP1R21 | 2.06E-08 | 1.241852361 | 0.261 | 0.111 | 0.000406032 | Tb |
| pDC | SPATA13 | 2.80E-12 | 1.243860769 | 0.472 | 0.241 | 5.52E-08 | Tb |
| pDC | B3GNT5 | 8.27E-07 | 1.270820841 | 0.412 | 0.268 | 0.01628496 | Tb |
| pDC | ABHD17B | 1.38E-07 | 1.278726759 | 0.333 | 0.176 | 0.00271369 | Tb |
| pDC | FNIP2 | 3.63E-08 | 1.287303863 | 0.365 | 0.201 | 0.000714482 | Tb |
| pDC | ZNF609 | 1.05E-10 | 1.350246902 | 0.371 | 0.176 | 2.06E-06 | Tb |
| pDC | RNF144B | 3.88E-11 | 1.352721486 | 0.5 | 0.272 | 7.64E-07 | Tb |
| pDC | HOMER1 | 1.63E-14 | 1.368392021 | 0.616 | 0.379 | 3.21E-10 | Tb |
| pDC | ABCC1 | 1.39E-08 | 1.379899783 | 0.274 | 0.119 | 0.000273561 | Tb |
| pDC | DIPK1A | 1.39E-10 | 1.428905373 | 0.34 | 0.155 | 2.74E-06 | Tb |
| pDC | CD180 | 7.97E-09 | 1.435823556 | 0.302 | 0.136 | 0.000156876 | Tb |
| pDC | KSR1 | 3.52E-10 | 1.438636521 | 0.277 | 0.107 | 6.92E-06 | Tb |
| pDC | TWISTNB | 4.45E-08 | 1.450828058 | 0.616 | 0.432 | 0.000874972 | Tb |
| pDC | PHLDB3 | 4.74E-12 | 1.465895271 | 0.333 | 0.134 | 9.33E-08 | Tb |
| pDC | ANKRD28 | 1.32E-13 | 1.475443515 | 0.557 | 0.342 | 2.60E-09 | Tb |
| pDC | SGK1 | 1.71E-07 | 1.519203296 | 0.292 | 0.147 | 0.003371274 | Tb |
| pDC | NAF1 | 3.04E-07 | 1.568494916 | 0.484 | 0.315 | 0.005993514 | Tb |
| pDC | RIPK2 | 1.12E-06 | 1.644691642 | 0.299 | 0.159 | 0.021959104 | Tb |
| pDC | AHR | 2.27E-07 | 1.703720094 | 0.399 | 0.235 | 0.004470174 | Tb |
| pDC | SESN1 | 1.16E-07 | 1.706224374 | 0.286 | 0.14 | 0.002282976 | Tb |
| pDC | SKI | 2.07E-12 | 1.744665928 | 0.296 | 0.103 | 4.08E-08 | Tb |
| pDC | TAF4B | 7.78E-10 | 1.815801223 | 0.336 | 0.155 | 1.53E-05 | Tb |
| pDC | SLC35F1 | 2.92E-57 | 4.175085982 | 0.566 | 0.088 | 5.75E-53 | Tb |
| **Disease-associated genes in Langerhans** | | | | | | | |

| | | | | | | | |
|---|---|---|---|---|---|---|---|
| Langerhans | GET4 | 3.41E-08 | 3.473024355 | 0.333 | 0.022 | 0.000670434 | Lep-L |
| Langerhans | HSPA1A | 1.21E-07 | 2.191802895 | 1 | 0.967 | 0.002385792 | Lep-L |
| Langerhans | EIF4A3 | 2.35E-07 | 2.822197909 | 1 | 0.749 | 0.004622022 | Lep-L |
| Langerhans | MCL1 | 1.59E-06 | 2.031715001 | 1 | 0.863 | 0.03129358 | Lep-L |
| Langerhans | HIST1H1E | 3.68E-07 | 2.316584265 | 0.582 | 0.282 | 0.007236143 | Lep-T |
| Langerhans | PPP1R15A1 | 4.20E-07 | 1.424874954 | 0.94 | 0.863 | 0.008267678 | Lep-T |
| Langerhans | PTGDS | 1.96E-06 | 2.297614113 | 0.343 | 0.076 | 0.038664724 | Lep-T |
| Langerhans | FOLR3 | 2.62E-14 | 6.956714366 | 0.364 | 0.006 | 5.16E-10 | Mari |
| Langerhans | TYROBP | 2.21E-11 | 1.44633153 | 1 | 0.943 | 4.35E-07 | Mari |
| Langerhans | HLA-DRB5 | 6.24E-11 | 3.871129868 | 1 | 0.659 | 1.23E-06 | Mari |
| Langerhans | PFN1 | 5.99E-10 | 1.27099958 | 1 | 0.926 | 1.18E-05 | Mari |
| Langerhans | S100A11 | 1.40E-09 | 1.232098457 | 1 | 0.972 | 2.75E-05 | Mari |
| Langerhans | IL7R | 1.67E-09 | 2.63464084 | 0.773 | 0.21 | 3.29E-05 | Mari |
| Langerhans | MS4A6A | 3.14E-09 | 2.150753497 | 0.909 | 0.369 | 6.18E-05 | Mari |
| Langerhans | ARHGDIB | 4.25E-09 | 1.536239114 | 1 | 0.875 | 8.36E-05 | Mari |
| Langerhans | AIF1 | 9.45E-09 | 1.407843935 | 1 | 0.949 | 0.000186094 | Mari |
| Langerhans | S100B | 1.06E-08 | 1.625902061 | 1 | 0.841 | 0.000209144 | Mari |
| Langerhans | CD14 | 1.12E-08 | 3.661537107 | 0.727 | 0.222 | 0.000221183 | Mari |
| Langerhans | FTL | 1.94E-08 | 1.426337401 | 1 | 0.989 | 0.000381045 | Mari |
| Langerhans | MYL12A | 2.02E-08 | 1.031244885 | 1 | 0.926 | 0.000397027 | Mari |
| Langerhans | GNG10 | 2.62E-08 | 1.757137308 | 0.909 | 0.67 | 0.000514931 | Mari |
| Langerhans | VAMP8 | 2.87E-08 | 1.409023981 | 1 | 0.886 | 0.000564056 | Mari |
| Langerhans | PRDX5 | 2.93E-08 | 1.40864694 | 1 | 0.653 | 0.000577331 | Mari |
| Langerhans | KRT16 | 3.80E-08 | 2.924674212 | 0.727 | 0.25 | 0.000747121 | Mari |
| Langerhans | C3AR1 | 5.70E-08 | 2.947807852 | 0.5 | 0.091 | 0.001121501 | Mari |
| Langerhans | SERF2 | 1.03E-07 | 0.906218391 | 1 | 0.966 | 0.002023166 | Mari |
| Langerhans | FAM89B | 1.20E-07 | 2.095115428 | 0.864 | 0.432 | 0.002355323 | Mari |
| Langerhans | SUMO2 | 1.47E-07 | 0.906712956 | 1 | 0.903 | 0.00289426 | Mari |
| Langerhans | HLA-DQB1 | 1.58E-07 | 1.04033797 | 1 | 0.989 | 0.003114231 | Mari |
| Langerhans | HLA-DRB1 | 2.86E-07 | 1.231440054 | 1 | 0.994 | 0.005635874 | Mari |
| Langerhans | CFL1 | 3.60E-07 | 1.153086524 | 1 | 0.955 | 0.007085646 | Mari |
| Langerhans | UQCR11 | 3.68E-07 | 1.141630821 | 1 | 0.812 | 0.007243373 | Mari |
| Langerhans | HLA-DPA1 | 3.76E-07 | 1.061544344 | 1 | 0.983 | 0.007392639 | Mari |
| Langerhans | IFI30 | 3.77E-07 | 1.600230701 | 1 | 0.875 | 0.007411172 | Mari |
| Langerhans | COTL1 | 4.29E-07 | 1.28849634 | 1 | 0.784 | 0.008453491 | Mari |
| Langerhans | C1QA | 5.21E-07 | 2.540089539 | 0.864 | 0.472 | 0.010255747 | Mari |
| Langerhans | ATP5PF | 5.89E-07 | 1.114937717 | 0.955 | 0.824 | 0.011593666 | Mari |
| Langerhans | SERPINB1 | 6.17E-07 | 1.226400461 | 0.955 | 0.585 | 0.012140979 | Mari |
| Langerhans | LDHA | 8.26E-07 | 1.027602487 | 1 | 0.892 | 0.016252538 | Mari |
| Langerhans | CDC42 | 9.35E-07 | 0.913116182 | 1 | 0.903 | 0.01839997 | Mari |
| Langerhans | TMSB4X | 1.04E-06 | 0.936739055 | 1 | 0.983 | 0.020501512 | Mari |
| Langerhans | C1QC | 1.19E-06 | 2.489030445 | 0.818 | 0.517 | 0.023480215 | Mari |
| Langerhans | SH3BGRL | 1.39E-06 | 0.968982791 | 0.955 | 0.898 | 0.027344873 | Mari |
| Langerhans | SMCO4 | 1.50E-06 | 1.512445681 | 0.909 | 0.636 | 0.02945082 | Mari |
| Langerhans | FCER1G | 1.57E-06 | 1.594733503 | 1 | 0.784 | 0.030929151 | Mari |
| Langerhans | IFITM2 | 1.67E-06 | 1.8782606 | 0.864 | 0.585 | 0.032804496 | Mari |
| Langerhans | TYMP | 1.87E-06 | 1.565806848 | 0.955 | 0.778 | 0.036779768 | Mari |
| Langerhans | POMP | 1.96E-06 | 0.941617454 | 0.955 | 0.824 | 0.038579652 | Mari |
| Langerhans | COX5A | 2.18E-06 | 1.044544849 | 1 | 0.795 | 0.042893913 | Mari |
| Langerhans | FCER1A | 2.51E-06 | 1.561481739 | 0.909 | 0.835 | 0.049441995 | Mari |
| Langerhans | MT-CO2 | 2.47E-13 | 1.074052686 | 1 | 0.952 | 4.86E-09 | Tb |
| Langerhans | MT-ND4 | 1.09E-12 | 1.030784907 | 0.989 | 0.952 | 2.15E-08 | Tb |
| Langerhans | MT-CO1 | 5.75E-12 | 1.021205981 | 1 | 0.981 | 1.13E-07 | Tb |
| Langerhans | MT-CO3 | 1.59E-11 | 1.151040787 | 1 | 0.904 | 3.12E-07 | Tb |
| Langerhans | MT-CYB | 5.12E-10 | 1.03483023 | 0.989 | 0.923 | 1.01E-05 | Tb |
| Langerhans | MT-ND3 | 3.12E-09 | 0.815581368 | 0.968 | 0.923 | 6.14E-05 | Tb |

| | | | | | | | |
|---|---|---|---|---|---|---|---|
| Langerhans | XIST | 5.57E-09 | 1.624342085 | 0.734 | 0.337 | 0.000109554 | Tb |
| Langerhans | S100A7 | 5.99E-09 | 1.562401325 | 0.564 | 0.173 | 0.000117962 | Tb |
| Langerhans | PDE4C | 6.15E-09 | 4.840654363 | 0.309 | 0.01 | 0.000121055 | Tb |
| Langerhans | MT-ATP6 | 8.49E-09 | 0.807459863 | 1 | 0.923 | 0.000167109 | Tb |
| Langerhans | CDH1 | 9.19E-09 | 1.269607604 | 0.819 | 0.452 | 0.000180903 | Tb |
| Langerhans | YME1L1 | 2.15E-08 | 1.556782006 | 0.819 | 0.615 | 0.000423341 | Tb |
| Langerhans | MSI2 | 3.80E-08 | 1.302578819 | 0.819 | 0.452 | 0.000747038 | Tb |
| Langerhans | ZBTB20 | 4.46E-08 | 1.334539828 | 0.883 | 0.567 | 0.000878219 | Tb |
| Langerhans | RGS13 | 5.03E-08 | 2.567545304 | 0.553 | 0.221 | 0.000989499 | Tb |
| Langerhans | S100A9 | 5.54E-08 | 1.246959824 | 0.734 | 0.413 | 0.001090963 | Tb |
| Langerhans | CD207 | 5.68E-08 | 1.244622315 | 0.83 | 0.635 | 0.001117903 | Tb |
| Langerhans | PIP5K1A | 6.20E-08 | 1.701461384 | 0.628 | 0.25 | 0.001220526 | Tb |
| Langerhans | TRIM69 | 8.12E-08 | 1.373971312 | 0.734 | 0.394 | 0.00159736 | Tb |
| Langerhans | RERE | 8.31E-08 | 1.311459919 | 0.809 | 0.558 | 0.001635899 | Tb |
| Langerhans | UBR5 | 9.52E-08 | 1.178412484 | 0.851 | 0.529 | 0.001873112 | Tb |
| Langerhans | SNX8 | 9.53E-08 | 0.844939025 | 0.883 | 0.702 | 0.001875848 | Tb |
| Langerhans | MT-ND2 | 1.02E-07 | 0.708600657 | 0.989 | 0.942 | 0.002001043 | Tb |
| Langerhans | S100A8 | 1.06E-07 | 1.752293659 | 0.681 | 0.375 | 0.002096329 | Tb |
| Langerhans | ELMO1 | 1.09E-07 | 0.725972774 | 0.968 | 0.846 | 0.002145372 | Tb |
| Langerhans | ITM2C | 2.28E-07 | 1.385293443 | 0.67 | 0.308 | 0.004483001 | Tb |
| Langerhans | MT-ND1 | 2.39E-07 | 0.60322783 | 0.989 | 0.894 | 0.00469547 | Tb |
| Langerhans | SH3D19 | 3.30E-07 | 2.112301116 | 0.468 | 0.144 | 0.006491033 | Tb |
| Langerhans | KRTDAP | 3.34E-07 | 1.810668991 | 0.574 | 0.26 | 0.006578559 | Tb |
| Langerhans | HPS5 | 3.87E-07 | 1.070793497 | 0.904 | 0.654 | 0.007614686 | Tb |
| Langerhans | ABL1 | 4.01E-07 | 1.701342196 | 0.521 | 0.183 | 0.007895074 | Tb |
| Langerhans | HIP1 | 4.13E-07 | 1.070539777 | 0.883 | 0.712 | 0.008123714 | Tb |
| Langerhans | LLGL2 | 4.53E-07 | 2.681942795 | 0.309 | 0.038 | 0.008924249 | Tb |
| Langerhans | KRT1 | 4.61E-07 | 2.418566432 | 0.553 | 0.279 | 0.009081218 | Tb |
| Langerhans | ARIH1 | 4.79E-07 | 0.981222343 | 0.936 | 0.702 | 0.009426784 | Tb |
| Langerhans | BRAF | 6.39E-07 | 0.980111729 | 0.819 | 0.519 | 0.012585499 | Tb |
| Langerhans | PAK1 | 7.42E-07 | 0.899434081 | 0.947 | 0.827 | 0.014609384 | Tb |
| Langerhans | UBAC2 | 7.76E-07 | 1.194559044 | 0.872 | 0.635 | 0.015272501 | Tb |
| Langerhans | PHTF1 | 8.58E-07 | 1.613554578 | 0.755 | 0.481 | 0.016893009 | Tb |
| Langerhans | ARID1B | 8.64E-07 | 0.966999761 | 0.787 | 0.471 | 0.017015693 | Tb |
| Langerhans | PDE7A | 1.15E-06 | 1.978568222 | 0.553 | 0.25 | 0.022727596 | Tb |
| Langerhans | HOMER1 | 1.34E-06 | 1.579570417 | 0.66 | 0.356 | 0.026397799 | Tb |
| Langerhans | ADK | 1.64E-06 | 0.972713118 | 0.84 | 0.577 | 0.032203023 | Tb |
| Langerhans | PBRM1 | 1.64E-06 | 1.445469762 | 0.745 | 0.471 | 0.032369355 | Tb |
| Langerhans | IDO2 | 1.68E-06 | 1.401359792 | 0.766 | 0.442 | 0.033023911 | Tb |
| Langerhans | NSMCE2 | 1.75E-06 | 1.329120686 | 0.745 | 0.49 | 0.03436106 | Tb |
| Langerhans | N4BP2L2 | 1.96E-06 | 0.707569544 | 0.862 | 0.644 | 0.038618935 | Tb |
| Langerhans | CCNT1 | 2.01E-06 | 1.402493637 | 0.489 | 0.163 | 0.039538121 | Tb |
| Langerhans | ZNRF1 | 2.20E-06 | 1.925841052 | 0.362 | 0.077 | 0.04330572 | Tb |
| Langerhans | CALML5 | 2.27E-06 | 1.138418166 | 0.447 | 0.154 | 0.044747337 | Tb |
| Langerhans | ZNF710 | 2.29E-06 | 1.107757281 | 0.787 | 0.519 | 0.04499808 | Tb |
| Langerhans | NUBPL | 2.50E-06 | 1.312772188 | 0.702 | 0.413 | 0.049246904 | Tb |

| Table S6 \| Disease-associated genes in macrophages | | | | | | | |
|---|---|---|---|---|---|---|---|
| **Disease-associated genes in all macrophages** | | | | | | | |
| **group** | **gene** | **p_val** | **avg_log2FC** | **pct.1** | **pct.2** | **p_val_adj** | **cluster** |
| allcell | HS3ST2 | 0 | 2.597177047 | 0.531 | 0.136 | 0 | Lep-L |
| allcell | CCR7 | 1.87E-213 | 2.473151522 | 0.307 | 0.13 | 3.67E-209 | Lep-L |
| allcell | CCL20 | 1.43E-212 | 2.417828104 | 0.373 | 0.188 | 2.81E-208 | Lep-L |
| allcell | SDS | 0 | 2.14919596 | 0.617 | 0.353 | 0 | Lep-L |
| allcell | PLIN2 | 0 | 2.147310746 | 0.875 | 0.691 | 0 | Lep-L |
| allcell | SLC19A2 | 8.35E-245 | 2.066015581 | 0.443 | 0.247 | 1.64E-240 | Lep-L |
| allcell | MIR3142HG | 2.14E-165 | 1.888506179 | 0.325 | 0.162 | 4.22E-161 | Lep-L |
| allcell | PPARG | 0 | 1.872803062 | 0.546 | 0.271 | 0 | Lep-L |
| allcell | CXCL8 | 0 | 1.843376235 | 0.865 | 0.644 | 0 | Lep-L |
| allcell | CYP27A1 | 0 | 1.800782261 | 0.612 | 0.269 | 0 | Lep-L |
| allcell | EREG | 2.37E-43 | 1.79519293 | 0.336 | 0.268 | 4.67E-39 | Lep-L |
| allcell | KLHL21 | 4.48E-271 | 1.776575089 | 0.416 | 0.201 | 8.81E-267 | Lep-L |
| allcell | RASGRP3 | 1.58E-290 | 1.74768956 | 0.546 | 0.322 | 3.11E-286 | Lep-L |
| allcell | SH3D21 | 1.78E-178 | 1.739994747 | 0.332 | 0.168 | 3.50E-174 | Lep-L |
| allcell | MMP19 | 1.70E-216 | 1.708751928 | 0.521 | 0.329 | 3.34E-212 | Lep-L |
| allcell | MFSD2A | 7.22E-279 | 1.681434322 | 0.464 | 0.25 | 1.42E-274 | Lep-L |
| allcell | FABP5 | 0 | 1.679779407 | 0.753 | 0.623 | 0 | Lep-L |
| allcell | KCNJ5 | 7.53E-149 | 1.668257357 | 0.251 | 0.109 | 1.48E-144 | Lep-L |
| allcell | APOC1 | 0 | 1.656818572 | 0.869 | 0.636 | 0 | Lep-L |
| allcell | PLPP3 | 2.19E-261 | 1.645850598 | 0.464 | 0.243 | 4.31E-257 | Lep-L |
| allcell | GPNMB | 0 | 1.632171371 | 0.845 | 0.577 | 0 | Lep-L |
| allcell | MGLL | 0 | 1.627618588 | 0.746 | 0.424 | 0 | Lep-L |
| allcell | SLC18B1 | 1.14E-183 | 1.619016767 | 0.283 | 0.116 | 2.25E-179 | Lep-L |
| allcell | LIPA | 0 | 1.605399742 | 0.717 | 0.506 | 0 | Lep-L |
| allcell | THAP2 | 3.05E-82 | 1.582478217 | 0.348 | 0.248 | 6.00E-78 | Lep-L |
| allcell | TPRG1 | 3.23E-99 | 1.566509626 | 0.258 | 0.14 | 6.35E-95 | Lep-L |
| allcell | CXCL3 | 0 | 1.554390342 | 0.817 | 0.587 | 0 | Lep-L |
| allcell | SDC4 | 1.63E-128 | 1.546352006 | 0.437 | 0.316 | 3.21E-124 | Lep-L |
| allcell | APBB3 | 7.15E-84 | 1.543402061 | 0.279 | 0.172 | 1.41E-79 | Lep-L |
| allcell | CDC42EP3 | 0 | 1.540602546 | 0.708 | 0.43 | 0 | Lep-L |
| allcell | CHCHD6 | 3.04E-294 | 1.510344026 | 0.464 | 0.238 | 5.99E-290 | Lep-L |
| allcell | ZNF703 | 9.35E-127 | 1.500717588 | 0.3 | 0.16 | 1.84E-122 | Lep-L |
| allcell | STMN1 | 6.50E-241 | 1.498911913 | 0.417 | 0.21 | 1.28E-236 | Lep-L |
| allcell | CMSS1 | 1.44E-44 | 1.488053298 | 0.425 | 0.373 | 2.84E-40 | Lep-L |
| allcell | TRIP10 | 1.13E-119 | 1.484651611 | 0.268 | 0.139 | 2.21E-115 | Lep-L |
| allcell | KMO | 0 | 1.474752034 | 0.69 | 0.49 | 0 | Lep-L |
| allcell | NBPF19 | 1.08E-137 | 1.449221084 | 0.384 | 0.243 | 2.13E-133 | Lep-L |
| allcell | APOE | 0 | 1.441339136 | 0.909 | 0.689 | 0 | Lep-L |
| allcell | MS4A14 | 1.14E-135 | 1.439076334 | 0.283 | 0.139 | 2.25E-131 | Lep-L |
| allcell | SMS | 0 | 1.398997311 | 0.746 | 0.569 | 0 | Lep-L |
| allcell | TREM2 | 1.82E-231 | 1.384962421 | 0.499 | 0.294 | 3.59E-227 | Lep-L |
| allcell | CSF1 | 1.88E-46 | 1.379850328 | 0.284 | 0.208 | 3.70E-42 | Lep-L |
| allcell | ZNF503 | 2.68E-107 | 1.378997135 | 0.253 | 0.129 | 5.27E-103 | Lep-L |
| allcell | DHRS3 | 3.29E-180 | 1.373236987 | 0.343 | 0.171 | 6.47E-176 | Lep-L |
| allcell | DDIT4 | 0 | 1.368600789 | 0.736 | 0.531 | 0 | Lep-L |
| allcell | OPN3 | 2.25E-200 | 1.367227365 | 0.514 | 0.346 | 4.44E-196 | Lep-L |
| allcell | ITPKB | 4.13E-189 | 1.365305191 | 0.404 | 0.22 | 8.13E-185 | Lep-L |
| allcell | PDE8A | 1.02E-235 | 1.338558792 | 0.591 | 0.393 | 2.01E-231 | Lep-L |
| allcell | SLC1A3 | 8.41E-276 | 1.322896046 | 0.734 | 0.573 | 1.66E-271 | Lep-L |
| allcell | GBE1 | 9.93E-153 | 1.318908479 | 0.565 | 0.419 | 1.96E-148 | Lep-L |
| allcell | SMPDL3A | 1.41E-148 | 1.312974425 | 0.453 | 0.308 | 2.77E-144 | Lep-L |
| allcell | AQP9 | 3.90E-93 | 1.303711686 | 0.509 | 0.407 | 7.68E-89 | Lep-L |

| | | | | | | | |
|---|---|---|---|---|---|---|---|
| allcell | CCRL2 | 2.23E-159 | 1.302319466 | 0.55 | 0.424 | 4.39E-155 | Lep-L |
| allcell | DDX3Y | 0 | 1.289298311 | 0.588 | 0.248 | 0 | Lep-L |
| allcell | KCNE1 | 3.72E-53 | 1.279738683 | 0.281 | 0.201 | 7.32E-49 | Lep-L |
| allcell | ABCC5 | 1.32E-106 | 1.279306647 | 0.285 | 0.159 | 2.60E-102 | Lep-L |
| allcell | DNAJC5B | 3.50E-64 | 1.27853559 | 0.265 | 0.177 | 6.88E-60 | Lep-L |
| allcell | MIR155HG | 8.39E-216 | 1.276007891 | 0.609 | 0.419 | 1.65E-211 | Lep-L |
| allcell | SCCPDH | 1.18E-107 | 1.260354974 | 0.338 | 0.218 | 2.32E-103 | Lep-L |
| allcell | PDXDC1 | 8.67E-196 | 1.244160795 | 0.558 | 0.399 | 1.71E-191 | Lep-L |
| allcell | TESK1 | 1.80E-82 | 1.243694806 | 0.302 | 0.199 | 3.55E-78 | Lep-L |
| allcell | SPSB1 | 3.44E-44 | 1.242183301 | 0.269 | 0.197 | 6.77E-40 | Lep-L |
| allcell | CHIT1 | 3.87E-120 | 1.218540282 | 0.302 | 0.166 | 7.62E-116 | Lep-L |
| allcell | CXCR4 | 0 | 1.218242299 | 0.795 | 0.652 | 0 | Lep-L |
| allcell | H2AFX | 1.30E-129 | 1.21567928 | 0.481 | 0.356 | 2.55E-125 | Lep-L |
| allcell | PHLDA1 | 9.73E-178 | 1.213616973 | 0.42 | 0.233 | 1.92E-173 | Lep-L |
| allcell | VAT1 | 4.57E-189 | 1.207927781 | 0.546 | 0.398 | 9.00E-185 | Lep-L |
| allcell | CXCL2 | 0 | 1.200590657 | 0.896 | 0.73 | 0 | Lep-L |
| allcell | CXCL1 | 1.71E-108 | 1.190698741 | 0.408 | 0.264 | 3.36E-104 | Lep-L |
| allcell | PDE4DIP | 0 | 1.189400683 | 0.825 | 0.621 | 0 | Lep-L |
| allcell | EBI3 | 3.23E-91 | 1.174056511 | 0.259 | 0.143 | 6.36E-87 | Lep-L |
| allcell | FABP3 | 1.28E-89 | 1.171145282 | 0.342 | 0.224 | 2.52E-85 | Lep-L |
| allcell | SMIM3 | 1.84E-144 | 1.169951792 | 0.5 | 0.372 | 3.63E-140 | Lep-L |
| allcell | C5AR1 | 0 | 1.157461044 | 0.787 | 0.657 | 0 | Lep-L |
| allcell | CSGALNACT2 | 7.62E-253 | 1.146889066 | 0.723 | 0.602 | 1.50E-248 | Lep-L |
| allcell | RBM38 | 1.62E-34 | 1.144687107 | 0.264 | 0.21 | 3.19E-30 | Lep-L |
| allcell | UGCG | 1.40E-219 | 1.140013333 | 0.655 | 0.517 | 2.76E-215 | Lep-L |
| allcell | SLC26A11 | 2.91E-53 | 1.139196071 | 0.284 | 0.205 | 5.74E-49 | Lep-L |
| allcell | SRGAP1 | 1.67E-33 | 1.133152629 | 0.355 | 0.292 | 3.28E-29 | Lep-L |
| allcell | CFD | 0 | 1.131812395 | 0.812 | 0.685 | 0 | Lep-L |
| allcell | RAB42 | 6.64E-58 | 1.128909693 | 0.27 | 0.185 | 1.31E-53 | Lep-L |
| allcell | SCD | 4.95E-54 | 1.128287594 | 0.364 | 0.288 | 9.75E-50 | Lep-L |
| allcell | HSPB1 | 1.06E-118 | 1.126870787 | 0.859 | 0.794 | 2.08E-114 | Lep-L |
| allcell | SLC29A1 | 2.23E-72 | 1.123149899 | 0.3 | 0.203 | 4.38E-68 | Lep-L |
| allcell | STAM | 2.44E-96 | 1.122961706 | 0.363 | 0.251 | 4.80E-92 | Lep-L |
| allcell | BCL2 | 1.88E-176 | 1.121613593 | 0.556 | 0.389 | 3.70E-172 | Lep-L |
| allcell | ADCY3 | 9.13E-43 | 1.11695318 | 0.325 | 0.255 | 1.80E-38 | Lep-L |
| allcell | DPEP2 | 6.43E-67 | 1.114904699 | 0.26 | 0.165 | 1.27E-62 | Lep-L |
| allcell | CCDC107 | 5.71E-220 | 1.110877225 | 0.659 | 0.556 | 1.12E-215 | Lep-L |
| allcell | PTGER2 | 1.91E-44 | 1.10876649 | 0.264 | 0.192 | 3.76E-40 | Lep-L |
| allcell | SNTB1 | 4.45E-58 | 1.100078511 | 0.391 | 0.308 | 8.76E-54 | Lep-L |
| allcell | ARRDC2 | 1.00E-54 | 1.086888504 | 0.348 | 0.272 | 1.97E-50 | Lep-L |
| allcell | PDK4 | 8.47E-190 | 1.086652304 | 0.347 | 0.154 | 1.67E-185 | Lep-L |
| allcell | PPP1R15B | 9.05E-248 | 1.085488528 | 0.723 | 0.578 | 1.78E-243 | Lep-L |
| allcell | BNIP3L | 5.20E-225 | 1.076133149 | 0.815 | 0.777 | 1.02E-220 | Lep-L |
| allcell | FNIP2 | 0 | 1.074630957 | 0.858 | 0.731 | 0 | Lep-L |
| allcell | TAGAP | 5.01E-131 | 1.073698186 | 0.714 | 0.639 | 9.87E-127 | Lep-L |
| allcell | SEPHS2 | 6.13E-128 | 1.071924801 | 0.632 | 0.537 | 1.21E-123 | Lep-L |
| allcell | TSC22D2 | 1.98E-173 | 1.064908958 | 0.713 | 0.582 | 3.90E-169 | Lep-L |
| allcell | TCEAL9 | 1.83E-79 | 1.064131871 | 0.373 | 0.268 | 3.59E-75 | Lep-L |
| allcell | TOM1L2 | 2.47E-37 | 1.046491061 | 0.287 | 0.224 | 4.86E-33 | Lep-L |
| allcell | CDK11B | 1.04E-07 | 1.04467657 | 0.325 | 0.325 | 0.00203996 | Lep-L |
| allcell | PI4K2A | 8.68E-94 | 1.028046741 | 0.485 | 0.396 | 1.71E-89 | Lep-L |
| allcell | A2M | 3.01E-191 | 1.024834592 | 0.628 | 0.468 | 5.92E-187 | Lep-L |
| allcell | BCL2A1 | 0 | 1.021986841 | 0.885 | 0.807 | 0 | Lep-L |
| allcell | HIPK2 | 5.42E-32 | 1.019232969 | 0.35 | 0.301 | 1.07E-27 | Lep-L |
| allcell | PHF1 | 1.19E-46 | 1.009990888 | 0.28 | 0.208 | 2.35E-42 | Lep-L |

| | | | | | | | |
|---|---|---|---|---|---|---|---|
| allcell | ADD3 | 3.68E-59 | 1.007395208 | 0.284 | 0.194 | 7.24E-55 | Lep-L |
| allcell | ADM | 9.61E-111 | 1.006632891 | 0.561 | 0.437 | 1.89E-106 | Lep-L |
| allcell | AVPI1 | 5.13E-123 | 1.00646609 | 0.536 | 0.412 | 1.01E-118 | Lep-L |
| allcell | NBPF14 | 2.49E-79 | 1.004564772 | 0.359 | 0.253 | 4.90E-75 | Lep-L |
| allcell | MAFF | 5.03E-192 | 1.003164401 | 0.662 | 0.526 | 9.91E-188 | Lep-L |
| allcell | HUS1 | 1.38E-36 | 1.002257429 | 0.31 | 0.253 | 2.72E-32 | Lep-L |
| allcell | UBD | 3.73E-174 | 3.125021668 | 0.433 | 0.227 | 7.35E-170 | Lep-T |
| allcell | CXCL14 | 8.63E-284 | 2.528337993 | 0.304 | 0.077 | 1.70E-279 | Lep-T |
| allcell | PLA2G2D | 3.58E-69 | 1.972170328 | 0.255 | 0.14 | 7.04E-65 | Lep-T |
| allcell | PTGDS | 4.63E-141 | 1.876513687 | 0.402 | 0.203 | 9.11E-137 | Lep-T |
| allcell | CST3 | 7.17E-215 | 1.85495304 | 0.974 | 0.966 | 1.41E-210 | Lep-T |
| allcell | DCN | 2.10E-76 | 1.752857686 | 0.262 | 0.133 | 4.13E-72 | Lep-T |
| allcell | UST | 8.74E-35 | 1.667452051 | 0.293 | 0.211 | 1.72E-30 | Lep-T |
| allcell | ALDOA | 2.64E-33 | 1.507175081 | 0.444 | 0.391 | 5.19E-29 | Lep-T |
| allcell | CCL2 | 1.27E-57 | 1.492530939 | 0.444 | 0.311 | 2.50E-53 | Lep-T |
| allcell | AFDN | 2.72E-108 | 1.416039042 | 0.364 | 0.196 | 5.36E-104 | Lep-T |
| allcell | HSD11B1 | 6.45E-47 | 1.354082036 | 0.26 | 0.164 | 1.27E-42 | Lep-T |
| allcell | PRKAR2B | 2.86E-95 | 1.334984277 | 0.314 | 0.165 | 5.63E-91 | Lep-T |
| allcell | MB21D2 | 1.73E-51 | 1.281689979 | 0.43 | 0.327 | 3.40E-47 | Lep-T |
| allcell | CD74 | 3.57E-261 | 1.271161411 | 0.99 | 0.994 | 7.03E-257 | Lep-T |
| allcell | LGALS2 | 1.05E-109 | 1.204308075 | 0.617 | 0.529 | 2.07E-105 | Lep-T |
| allcell | RNASE1 | 2.63E-23 | 1.177343406 | 0.455 | 0.418 | 5.18E-19 | Lep-T |
| allcell | F13A1 | 1.47E-109 | 1.150355796 | 0.452 | 0.27 | 2.89E-105 | Lep-T |
| allcell | TSPAN33 | 7.06E-67 | 1.138739664 | 0.5 | 0.381 | 1.39E-62 | Lep-T |
| allcell | FKBP2 | 2.43E-21 | 1.114763826 | 0.291 | 0.229 | 4.78E-17 | Lep-T |
| allcell | MVB12B | 7.38E-41 | 1.113080777 | 0.278 | 0.179 | 1.45E-36 | Lep-T |
| allcell | HLA-DQB2 | 1.15E-59 | 1.105986019 | 0.376 | 0.256 | 2.27E-55 | Lep-T |
| allcell | HDAC9 | 1.01E-06 | 1.099769944 | 0.27 | 0.244 | 0.019933037 | Lep-T |
| allcell | EMB | 1.51E-38 | 1.091416282 | 0.306 | 0.211 | 2.97E-34 | Lep-T |
| allcell | DHRS9 | 6.36E-09 | 1.085861982 | 0.272 | 0.24 | 0.000125226 | Lep-T |
| allcell | INPP5A | 5.71E-14 | 1.0681589 | 0.255 | 0.208 | 1.12E-09 | Lep-T |
| allcell | RFTN1 | 4.28E-50 | 1.047449199 | 0.465 | 0.375 | 8.42E-46 | Lep-T |
| allcell | RBFOX2 | 1.29E-58 | 1.042944102 | 0.558 | 0.453 | 2.54E-54 | Lep-T |
| allcell | CFP | 4.17E-27 | 1.037828603 | 0.281 | 0.204 | 8.21E-23 | Lep-T |
| allcell | STAB1 | 4.66E-23 | 1.021878367 | 0.393 | 0.337 | 9.18E-19 | Lep-T |
| allcell | PLCL1 | 2.47E-24 | 1.015020167 | 0.365 | 0.302 | 4.87E-20 | Lep-T |
| allcell | LPAR6 | 1.04E-25 | 1.007055878 | 0.408 | 0.326 | 2.06E-21 | Lep-T |
| allcell | FOLR3 | 0 | 5.676283412 | 0.386 | 0.024 | 0 | Mari |
| allcell | SPP1 | 3.35E-53 | 2.437998076 | 0.262 | 0.159 | 6.60E-49 | Mari |
| allcell | S100A8 | 0 | 2.29449628 | 0.893 | 0.407 | 0 | Mari |
| allcell | CASP5 | 1.76E-278 | 2.134665339 | 0.329 | 0.094 | 3.47E-274 | Mari |
| allcell | S100A9 | 0 | 1.761578372 | 0.955 | 0.698 | 0 | Mari |
| allcell | FCGR1B | 0 | 1.637933175 | 0.58 | 0.254 | 0 | Mari |
| allcell | PLTP | 2.39E-103 | 1.544221568 | 0.697 | 0.566 | 4.70E-99 | Mari |
| allcell | FCGR1A | 0 | 1.512682702 | 0.845 | 0.509 | 0 | Mari |
| allcell | CALHM6 | 0 | 1.500005303 | 0.901 | 0.615 | 0 | Mari |
| allcell | MGST1 | 1.38E-95 | 1.495554887 | 0.261 | 0.121 | 2.72E-91 | Mari |
| allcell | VAMP5 | 0 | 1.451839235 | 0.889 | 0.56 | 0 | Mari |
| allcell | FCGR3A | 0 | 1.399380463 | 0.882 | 0.562 | 0 | Mari |
| allcell | CORO1A | 1.09E-205 | 1.349470206 | 0.631 | 0.356 | 2.14E-201 | Mari |
| allcell | CCL8 | 3.08E-138 | 1.324380238 | 0.376 | 0.176 | 6.07E-134 | Mari |
| allcell | IL6 | 5.11E-117 | 1.323692962 | 0.308 | 0.14 | 1.01E-112 | Mari |
| allcell | TMEM173 | 2.92E-160 | 1.307642974 | 0.406 | 0.191 | 5.74E-156 | Mari |
| allcell | CD93 | 1.09E-110 | 1.295333995 | 0.283 | 0.128 | 2.15E-106 | Mari |
| allcell | OLFML2B | 1.14E-174 | 1.27414937 | 0.411 | 0.183 | 2.24E-170 | Mari |

| | | | | | | | |
|---|---|---|---|---|---|---|---|
| allcell | GIMAP6 | 1.04E-137 | 1.261418843 | 0.303 | 0.126 | 2.05E-133 | Mari |
| allcell | ETV7 | 6.79E-158 | 1.261126971 | 0.289 | 0.106 | 1.34E-153 | Mari |
| allcell | IL7R | 3.18E-170 | 1.251065797 | 0.453 | 0.217 | 6.26E-166 | Mari |
| allcell | TMEM163 | 8.78E-117 | 1.250801901 | 0.272 | 0.115 | 1.73E-112 | Mari |
| allcell | IFI44L | 8.79E-188 | 1.245166232 | 0.373 | 0.146 | 1.73E-183 | Mari |
| allcell | IFITM3 | 1.48E-225 | 1.231635455 | 0.909 | 0.722 | 2.91E-221 | Mari |
| allcell | TYMP | 0 | 1.210610274 | 0.983 | 0.831 | 0 | Mari |
| allcell | IDO1 | 1.41E-172 | 1.199494201 | 0.447 | 0.212 | 2.77E-168 | Mari |
| allcell | MRPL23 | 6.53E-240 | 1.18734964 | 0.55 | 0.257 | 1.29E-235 | Mari |
| allcell | MRPL58 | 6.04E-120 | 1.186649018 | 0.349 | 0.168 | 1.19E-115 | Mari |
| allcell | CCDC167 | 7.72E-142 | 1.18358581 | 0.347 | 0.154 | 1.52E-137 | Mari |
| allcell | HRH2 | 1.98E-157 | 1.181171813 | 0.427 | 0.206 | 3.89E-153 | Mari |
| allcell | CR1 | 8.11E-28 | 1.174344984 | 0.294 | 0.215 | 1.60E-23 | Mari |
| allcell | GBP1 | 0 | 1.149796713 | 0.839 | 0.505 | 0 | Mari |
| allcell | CD300E | 7.41E-144 | 1.148470708 | 0.308 | 0.123 | 1.46E-139 | Mari |
| allcell | MNDA | 1.59E-253 | 1.145828862 | 0.83 | 0.549 | 3.13E-249 | Mari |
| allcell | SDF2L1 | 2.14E-186 | 1.143230287 | 0.636 | 0.39 | 4.22E-182 | Mari |
| allcell | PSMB9 | 0 | 1.136825187 | 0.905 | 0.668 | 0 | Mari |
| allcell | SCO2 | 1.44E-263 | 1.134039843 | 0.728 | 0.411 | 2.83E-259 | Mari |
| allcell | DDX60L | 4.34E-146 | 1.128758519 | 0.47 | 0.245 | 8.54E-142 | Mari |
| allcell | CD14 | 0 | 1.120050303 | 0.944 | 0.803 | 0 | Mari |
| allcell | TNFSF10 | 0 | 1.114299409 | 0.763 | 0.423 | 0 | Mari |
| allcell | BST2 | 0 | 1.107462221 | 0.916 | 0.711 | 0 | Mari |
| allcell | CLEC11A | 8.03E-83 | 1.105380754 | 0.259 | 0.125 | 1.58E-78 | Mari |
| allcell | GALK1 | 4.90E-141 | 1.091692495 | 0.442 | 0.229 | 9.65E-137 | Mari |
| allcell | ANKRD22 | 6.33E-261 | 1.08768849 | 0.553 | 0.24 | 1.25E-256 | Mari |
| allcell | PRELID1 | 7.50E-295 | 1.084788006 | 0.903 | 0.691 | 1.48E-290 | Mari |
| allcell | IFIT3 | 5.58E-155 | 1.083887879 | 0.393 | 0.18 | 1.10E-150 | Mari |
| allcell | XRCC4 | 5.04E-117 | 1.073547276 | 0.351 | 0.17 | 9.93E-113 | Mari |
| allcell | CCR1 | 5.80E-166 | 1.073255928 | 0.523 | 0.278 | 1.14E-161 | Mari |
| allcell | CASP4 | 3.10E-292 | 1.071659933 | 0.821 | 0.529 | 6.09E-288 | Mari |
| allcell | PLAAT4 | 0 | 1.071363569 | 0.803 | 0.474 | 0 | Mari |
| allcell | PSME2 | 0 | 1.071034203 | 0.919 | 0.703 | 0 | Mari |
| allcell | CAMK1 | 8.62E-139 | 1.07038286 | 0.388 | 0.184 | 1.70E-134 | Mari |
| allcell | POLE4 | 4.95E-195 | 1.06812742 | 0.626 | 0.362 | 9.74E-191 | Mari |
| allcell | IFI35 | 8.63E-186 | 1.067948295 | 0.535 | 0.276 | 1.70E-181 | Mari |
| allcell | FAM89B | 7.03E-175 | 1.066787912 | 0.569 | 0.316 | 1.38E-170 | Mari |
| allcell | CXCL10 | 1.46E-150 | 1.066275771 | 0.539 | 0.3 | 2.87E-146 | Mari |
| allcell | RGS18 | 9.71E-62 | 1.060371006 | 0.252 | 0.137 | 1.91E-57 | Mari |
| allcell | ENPP2 | 3.18E-132 | 1.055425266 | 0.397 | 0.195 | 6.26E-128 | Mari |
| allcell | PCED1B-AS1 | 8.18E-95 | 1.046664726 | 0.311 | 0.156 | 1.61E-90 | Mari |
| allcell | JOSD2 | 2.00E-232 | 1.043787116 | 0.622 | 0.336 | 3.94E-228 | Mari |
| allcell | IFI30 | 0 | 1.042144038 | 0.991 | 0.941 | 0 | Mari |
| allcell | CRELD2 | 1.86E-125 | 1.038019138 | 0.423 | 0.223 | 3.66E-121 | Mari |
| allcell | SATB1-AS1 | 2.57E-72 | 1.031057957 | 0.321 | 0.181 | 5.05E-68 | Mari |
| allcell | VPS25 | 1.43E-151 | 1.017747798 | 0.465 | 0.24 | 2.82E-147 | Mari |
| allcell | FCER1G | 0 | 1.014222622 | 0.98 | 0.899 | 0 | Mari |
| allcell | S100A4 | 1.17E-238 | 1.013254141 | 0.942 | 0.82 | 2.31E-234 | Mari |
| allcell | APOL4 | 4.33E-118 | 1.008391737 | 0.33 | 0.154 | 8.52E-114 | Mari |
| allcell | TUBA1A | 1.12E-235 | 1.005667892 | 0.841 | 0.607 | 2.21E-231 | Mari |
| allcell | TREM1 | 8.25E-101 | 1.004683941 | 0.389 | 0.21 | 1.62E-96 | Mari |
| allcell | MSRB1 | 1.07E-116 | 1.000225187 | 0.384 | 0.197 | 2.11E-112 | Mari |
| allcell | CAMP | 3.51E-207 | 3.121586315 | 0.251 | 0.074 | 6.91E-203 | Tb |
| allcell | FN1 | 0 | 2.784608153 | 0.434 | 0.16 | 0 | Tb |
| allcell | SLC35F1 | 0 | 2.762905885 | 0.46 | 0.1 | 0 | Tb |

| | | | | | | | |
|---|---|---|---|---|---|---|---|
| allcell | GGT5 | 0 | 2.648150764 | 0.321 | 0.063 | 0 | Tb |
| allcell | MT1G | 6.60E-137 | 2.39558935 | 0.29 | 0.122 | 1.30E-132 | Tb |
| allcell | NRN1 | 1.88E-278 | 2.285825117 | 0.284 | 0.072 | 3.71E-274 | Tb |
| allcell | PDLIM4 | 5.30E-257 | 2.2263347 | 0.284 | 0.077 | 1.04E-252 | Tb |
| allcell | FBP1 | 3.11E-296 | 2.06616187 | 0.728 | 0.476 | 6.12E-292 | Tb |
| allcell | CXCL9 | 0 | 2.038385663 | 0.61 | 0.271 | 0 | Tb |
| allcell | S100A2 | 3.05E-283 | 1.949692298 | 0.304 | 0.077 | 6.00E-279 | Tb |
| allcell | IDO1 | 1.20E-212 | 1.787183535 | 0.448 | 0.206 | 2.36E-208 | Tb |
| allcell | MT1E | 3.18E-129 | 1.737150783 | 0.36 | 0.177 | 6.25E-125 | Tb |
| allcell | QPCT | 1.00E-262 | 1.728048671 | 0.475 | 0.202 | 1.97E-258 | Tb |
| allcell | HOMER1 | 2.77E-236 | 1.666993309 | 0.474 | 0.212 | 5.46E-232 | Tb |
| allcell | CISH | 2.14E-190 | 1.54831249 | 0.288 | 0.097 | 4.22E-186 | Tb |
| allcell | VDR | 1.07E-186 | 1.532386143 | 0.321 | 0.117 | 2.10E-182 | Tb |
| allcell | MT1X | 5.90E-137 | 1.478091264 | 0.56 | 0.345 | 1.16E-132 | Tb |
| allcell | CXCL10 | 4.85E-226 | 1.47515479 | 0.562 | 0.288 | 9.55E-222 | Tb |
| allcell | CTSK | 2.17E-42 | 1.467434987 | 0.294 | 0.194 | 4.28E-38 | Tb |
| allcell | SIGLEC10 | 5.01E-242 | 1.443601355 | 0.463 | 0.194 | 9.86E-238 | Tb |
| allcell | ANKRD22 | 3.24E-251 | 1.429830452 | 0.529 | 0.239 | 6.38E-247 | Tb |
| allcell | PPA1 | 0 | 1.42559112 | 0.81 | 0.479 | 0 | Tb |
| allcell | PTGDS | 5.55E-45 | 1.404402351 | 0.322 | 0.22 | 1.09E-40 | Tb |
| allcell | ENPP2 | 5.58E-237 | 1.400982071 | 0.442 | 0.179 | 1.10E-232 | Tb |
| allcell | APOL4 | 2.46E-229 | 1.398490641 | 0.378 | 0.137 | 4.85E-225 | Tb |
| allcell | MT1F | 5.63E-97 | 1.385232451 | 0.303 | 0.153 | 1.11E-92 | Tb |
| allcell | LYZ | 0 | 1.353762244 | 0.983 | 0.887 | 0 | Tb |
| allcell | FXYD2 | 1.26E-69 | 1.333589948 | 0.266 | 0.144 | 2.49E-65 | Tb |
| allcell | SFN | 7.74E-137 | 1.315898553 | 0.259 | 0.098 | 1.52E-132 | Tb |
| allcell | TSFM | 1.16E-160 | 1.296295526 | 0.383 | 0.173 | 2.29E-156 | Tb |
| allcell | FLT1 | 1.93E-114 | 1.293159291 | 0.451 | 0.264 | 3.79E-110 | Tb |
| allcell | USP30-AS1 | 3.78E-105 | 1.286993433 | 0.256 | 0.113 | 7.44E-101 | Tb |
| allcell | TMEM163 | 4.18E-122 | 1.285920856 | 0.269 | 0.112 | 8.23E-118 | Tb |
| allcell | HAPLN3 | 3.16E-221 | 1.282963188 | 0.457 | 0.197 | 6.22E-217 | Tb |
| allcell | TGM2 | 5.13E-130 | 1.277966659 | 0.367 | 0.18 | 1.01E-125 | Tb |
| allcell | TYMP | 0 | 1.251704121 | 0.979 | 0.828 | 0 | Tb |
| allcell | EGLN3 | 9.49E-158 | 1.212217227 | 0.374 | 0.169 | 1.87E-153 | Tb |
| allcell | MMP12 | 3.71E-225 | 1.204684087 | 0.286 | 0.083 | 7.30E-221 | Tb |
| allcell | CLEC4D | 1.05E-105 | 1.20043775 | 0.252 | 0.108 | 2.07E-101 | Tb |
| allcell | SUCNR1 | 3.82E-157 | 1.199999014 | 0.365 | 0.161 | 7.52E-153 | Tb |
| allcell | HSD11B1 | 2.79E-164 | 1.195718012 | 0.344 | 0.142 | 5.49E-160 | Tb |
| allcell | SEMA4D | 8.79E-200 | 1.187405033 | 0.503 | 0.242 | 1.73E-195 | Tb |
| allcell | TAF4B | 2.44E-53 | 1.176769264 | 0.326 | 0.203 | 4.81E-49 | Tb |
| allcell | CDA | 1.01E-109 | 1.174898742 | 0.319 | 0.155 | 1.99E-105 | Tb |
| allcell | CTSC | 1.22E-283 | 1.174474125 | 0.921 | 0.825 | 2.40E-279 | Tb |
| allcell | ACOT7 | 4.79E-199 | 1.17116591 | 0.43 | 0.189 | 9.43E-195 | Tb |
| allcell | MCF2L2 | 3.13E-119 | 1.16863236 | 0.426 | 0.233 | 6.17E-115 | Tb |
| allcell | PFKP | 4.74E-160 | 1.166385611 | 0.415 | 0.199 | 9.32E-156 | Tb |
| allcell | S100A8 | 0 | 1.165784927 | 0.837 | 0.411 | 0 | Tb |
| allcell | HS3ST3B1 | 8.89E-88 | 1.151359734 | 0.316 | 0.167 | 1.75E-83 | Tb |
| allcell | LIMK2 | 2.43E-170 | 1.144570828 | 0.48 | 0.24 | 4.79E-166 | Tb |
| allcell | IL3RA | 9.34E-295 | 1.129127308 | 0.654 | 0.322 | 1.84E-290 | Tb |
| allcell | MIR3945HG | 3.49E-89 | 1.127994919 | 0.429 | 0.259 | 6.87E-85 | Tb |
| allcell | SULT1C2 | 6.80E-89 | 1.125245102 | 0.302 | 0.159 | 1.34E-84 | Tb |
| allcell | IFIT3 | 1.29E-141 | 1.123906734 | 0.381 | 0.178 | 2.54E-137 | Tb |
| allcell | GBP5 | 1.19E-260 | 1.120405829 | 0.788 | 0.478 | 2.34E-256 | Tb |
| allcell | CD300E | 1.30E-104 | 1.104531456 | 0.279 | 0.126 | 2.55E-100 | Tb |
| allcell | RHOH | 5.24E-66 | 1.103198078 | 0.373 | 0.237 | 1.03E-61 | Tb |

| | | | | | | | |
|---|---|---|---|---|---|---|---|
| allcell | ERAP2 | 2.95E-161 | 1.084263237 | 0.353 | 0.147 | 5.80E-157 | Tb |
| allcell | VASH1 | 4.67E-147 | 1.082579494 | 0.378 | 0.176 | 9.19E-143 | Tb |
| allcell | CRISPLD2 | 2.58E-124 | 1.074939199 | 0.267 | 0.109 | 5.07E-120 | Tb |
| allcell | PDCD1LG2 | 4.08E-229 | 1.073955586 | 0.493 | 0.216 | 8.04E-225 | Tb |
| allcell | GBP1 | 0 | 1.066552777 | 0.843 | 0.497 | 0 | Tb |
| allcell | P2RY14 | 3.12E-193 | 1.066452766 | 0.325 | 0.115 | 6.14E-189 | Tb |
| allcell | CFB | 6.50E-131 | 1.0659381 | 0.27 | 0.107 | 1.28E-126 | Tb |
| allcell | SLC30A1 | 3.10E-129 | 1.049066121 | 0.365 | 0.174 | 6.11E-125 | Tb |
| allcell | CD300LF | 2.74E-140 | 1.045643247 | 0.456 | 0.24 | 5.40E-136 | Tb |
| allcell | PDSS1 | 4.19E-70 | 1.044178225 | 0.317 | 0.18 | 8.25E-66 | Tb |
| allcell | CCND2 | 4.84E-110 | 1.03939896 | 0.31 | 0.146 | 9.53E-106 | Tb |
| allcell | LRRK2 | 3.83E-125 | 1.035418831 | 0.452 | 0.245 | 7.54E-121 | Tb |
| allcell | P2RY13 | 3.59E-171 | 1.029335371 | 0.457 | 0.22 | 7.07E-167 | Tb |
| allcell | SNX20 | 1.35E-130 | 1.028982878 | 0.349 | 0.16 | 2.65E-126 | Tb |
| allcell | DENND6B | 9.16E-102 | 1.024056674 | 0.29 | 0.135 | 1.80E-97 | Tb |
| allcell | PLAAT4 | 3.97E-253 | 1.015899148 | 0.783 | 0.471 | 7.82E-249 | Tb |
| allcell | SMAD7 | 2.01E-135 | 1.014968989 | 0.482 | 0.263 | 3.96E-131 | Tb |
| allcell | PVT1 | 3.12E-119 | 1.014945699 | 0.275 | 0.116 | 6.15E-115 | Tb |
| allcell | TNFSF10 | 3.36E-283 | 1.0142942 | 0.755 | 0.417 | 6.61E-279 | Tb |
| **Disease-associated genes in Macro-FOLR2** | | | | | | | |
| Macro-FOLR2 | CHIT1 | 4.08E-89 | 3.033869681 | 0.312 | 0.09 | 8.03E-85 | Lep-L |
| Macro-FOLR2 | CYP27A1 | 2.87E-145 | 2.550826455 | 0.532 | 0.211 | 5.66E-141 | Lep-L |
| Macro-FOLR2 | FABP5 | 1.77E-113 | 2.52792952 | 0.672 | 0.495 | 3.48E-109 | Lep-L |
| Macro-FOLR2 | CHCHD6 | 8.02E-108 | 2.221689083 | 0.496 | 0.232 | 1.58E-103 | Lep-L |
| Macro-FOLR2 | CSF1 | 1.61E-48 | 2.154688364 | 0.273 | 0.113 | 3.16E-44 | Lep-L |
| Macro-FOLR2 | PLIN2 | 7.47E-219 | 2.069747517 | 0.843 | 0.694 | 1.47E-214 | Lep-L |
| Macro-FOLR2 | TPRG1 | 1.79E-60 | 2.041427116 | 0.259 | 0.084 | 3.53E-56 | Lep-L |
| Macro-FOLR2 | OTOA | 5.59E-152 | 2.004936975 | 0.655 | 0.352 | 1.10E-147 | Lep-L |
| Macro-FOLR2 | SDC4 | 3.75E-34 | 1.991525513 | 0.325 | 0.192 | 7.37E-30 | Lep-L |
| Macro-FOLR2 | CDC42EP3 | 7.08E-168 | 1.984739864 | 0.679 | 0.323 | 1.39E-163 | Lep-L |
| Macro-FOLR2 | SLC19A2 | 1.82E-64 | 1.969683714 | 0.458 | 0.257 | 3.58E-60 | Lep-L |
| Macro-FOLR2 | LIPA | 1.84E-152 | 1.889059374 | 0.739 | 0.57 | 3.62E-148 | Lep-L |
| Macro-FOLR2 | SDS | 1.09E-94 | 1.884174047 | 0.647 | 0.408 | 2.15E-90 | Lep-L |
| Macro-FOLR2 | DDIT4 | 2.21E-136 | 1.845040449 | 0.748 | 0.478 | 4.35E-132 | Lep-L |
| Macro-FOLR2 | LSP1 | 8.41E-74 | 1.768091646 | 0.392 | 0.168 | 1.65E-69 | Lep-L |
| Macro-FOLR2 | CXCL8 | 2.16E-121 | 1.67899672 | 0.795 | 0.579 | 4.25E-117 | Lep-L |
| Macro-FOLR2 | TREM2 | 1.90E-77 | 1.675295687 | 0.487 | 0.249 | 3.74E-73 | Lep-L |
| Macro-FOLR2 | RASGRP3 | 3.88E-66 | 1.658823642 | 0.473 | 0.257 | 7.63E-62 | Lep-L |
| Macro-FOLR2 | PDE6G | 4.66E-59 | 1.638771605 | 0.307 | 0.122 | 9.17E-55 | Lep-L |
| Macro-FOLR2 | DNAJC5B | 2.35E-24 | 1.623584017 | 0.266 | 0.159 | 4.62E-20 | Lep-L |
| Macro-FOLR2 | AVPI1 | 8.91E-61 | 1.61981597 | 0.484 | 0.287 | 1.75E-56 | Lep-L |
| Macro-FOLR2 | ZNF703 | 4.17E-41 | 1.599634785 | 0.338 | 0.175 | 8.21E-37 | Lep-L |
| Macro-FOLR2 | GPNMB | 2.49E-186 | 1.59786383 | 0.844 | 0.691 | 4.90E-182 | Lep-L |
| Macro-FOLR2 | APOC1 | 1.63E-147 | 1.566793925 | 0.84 | 0.697 | 3.20E-143 | Lep-L |
| Macro-FOLR2 | CCRL2 | 2.12E-51 | 1.552424076 | 0.521 | 0.36 | 4.18E-47 | Lep-L |
| Macro-FOLR2 | APBB3 | 3.85E-24 | 1.540202077 | 0.289 | 0.176 | 7.59E-20 | Lep-L |
| Macro-FOLR2 | IGKC | 2.37E-24 | 1.514524218 | 0.319 | 0.185 | 4.67E-20 | Lep-L |
| Macro-FOLR2 | KLHL21 | 9.80E-52 | 1.513222974 | 0.34 | 0.158 | 1.93E-47 | Lep-L |
| Macro-FOLR2 | PLPP3 | 4.44E-55 | 1.507620774 | 0.403 | 0.204 | 8.73E-51 | Lep-L |
| Macro-FOLR2 | CXCR4 | 6.78E-104 | 1.50460355 | 0.754 | 0.578 | 1.34E-99 | Lep-L |
| Macro-FOLR2 | CA2 | 3.46E-60 | 1.503305828 | 0.456 | 0.246 | 6.81E-56 | Lep-L |
| Macro-FOLR2 | SH3D21 | 4.97E-32 | 1.487406726 | 0.282 | 0.151 | 9.78E-28 | Lep-L |
| Macro-FOLR2 | BCL2A1 | 1.36E-125 | 1.482917879 | 0.808 | 0.64 | 2.67E-121 | Lep-L |
| Macro-FOLR2 | ADM | 3.71E-68 | 1.455053677 | 0.598 | 0.39 | 7.31E-64 | Lep-L |
| Macro-FOLR2 | MFSD2A | 1.83E-49 | 1.452380287 | 0.341 | 0.167 | 3.60E-45 | Lep-L |

| | | | | | | | |
|---|---|---|---|---|---|---|---|
| Macro-FOLR2 | HS3ST2 | 5.66E-136 | 1.446515351 | 0.59 | 0.233 | 1.11E-131 | Lep-L |
| Macro-FOLR2 | PPARG | 1.08E-96 | 1.436807444 | 0.554 | 0.279 | 2.13E-92 | Lep-L |
| Macro-FOLR2 | HSPA2 | 8.74E-43 | 1.430502597 | 0.369 | 0.203 | 1.72E-38 | Lep-L |
| Macro-FOLR2 | STMN1 | 3.03E-56 | 1.402487516 | 0.48 | 0.292 | 5.96E-52 | Lep-L |
| Macro-FOLR2 | APOE | 2.37E-152 | 1.397456111 | 0.881 | 0.78 | 4.67E-148 | Lep-L |
| Macro-FOLR2 | KMO | 1.51E-49 | 1.392871293 | 0.563 | 0.39 | 2.97E-45 | Lep-L |
| Macro-FOLR2 | DPEP2 | 4.26E-39 | 1.365493157 | 0.339 | 0.179 | 8.39E-35 | Lep-L |
| Macro-FOLR2 | PHLDA1 | 2.06E-50 | 1.360361642 | 0.376 | 0.183 | 4.06E-46 | Lep-L |
| Macro-FOLR2 | AQP9 | 1.59E-11 | 1.35541773 | 0.288 | 0.217 | 3.14E-07 | Lep-L |
| Macro-FOLR2 | TAGAP | 1.17E-53 | 1.350765896 | 0.72 | 0.6 | 2.30E-49 | Lep-L |
| Macro-FOLR2 | MGLL | 3.31E-109 | 1.329658442 | 0.714 | 0.479 | 6.52E-105 | Lep-L |
| Macro-FOLR2 | GBE1 | 7.12E-39 | 1.324224772 | 0.543 | 0.394 | 1.40E-34 | Lep-L |
| Macro-FOLR2 | BHLHE41 | 5.05E-54 | 1.294839202 | 0.46 | 0.257 | 9.93E-50 | Lep-L |
| Macro-FOLR2 | SEPHS2 | 1.87E-46 | 1.266914849 | 0.636 | 0.502 | 3.68E-42 | Lep-L |
| Macro-FOLR2 | MIR155HG | 1.17E-33 | 1.251504457 | 0.499 | 0.351 | 2.31E-29 | Lep-L |
| Macro-FOLR2 | CD52 | 4.82E-31 | 1.248433255 | 0.28 | 0.149 | 9.48E-27 | Lep-L |
| Macro-FOLR2 | CD82 | 1.42E-41 | 1.247167532 | 0.4 | 0.241 | 2.80E-37 | Lep-L |
| Macro-FOLR2 | SMIM3 | 2.40E-35 | 1.23788528 | 0.464 | 0.327 | 4.73E-31 | Lep-L |
| Macro-FOLR2 | HSPA1B | 7.09E-112 | 1.233016646 | 0.978 | 0.919 | 1.40E-107 | Lep-L |
| Macro-FOLR2 | ITGAX | 1.26E-56 | 1.232402804 | 0.603 | 0.413 | 2.47E-52 | Lep-L |
| Macro-FOLR2 | FGR | 8.01E-20 | 1.229114477 | 0.379 | 0.283 | 1.58E-15 | Lep-L |
| Macro-FOLR2 | UPP1 | 2.05E-51 | 1.209688705 | 0.65 | 0.513 | 4.04E-47 | Lep-L |
| Macro-FOLR2 | HSPA6 | 5.70E-113 | 1.206614048 | 0.816 | 0.574 | 1.12E-108 | Lep-L |
| Macro-FOLR2 | CDK11B | 2.39E-07 | 1.194712251 | 0.332 | 0.291 | 0.004708972 | Lep-L |
| Macro-FOLR2 | BCL2 | 6.82E-38 | 1.193984121 | 0.426 | 0.265 | 1.34E-33 | Lep-L |
| Macro-FOLR2 | ZNF503 | 2.71E-32 | 1.189766802 | 0.298 | 0.157 | 5.34E-28 | Lep-L |
| Macro-FOLR2 | CKS2 | 2.41E-84 | 1.159340002 | 0.793 | 0.655 | 4.74E-80 | Lep-L |
| Macro-FOLR2 | ADCY3 | 1.41E-17 | 1.155958036 | 0.259 | 0.165 | 2.78E-13 | Lep-L |
| Macro-FOLR2 | H2AFX | 9.83E-41 | 1.148587997 | 0.515 | 0.373 | 1.94E-36 | Lep-L |
| Macro-FOLR2 | ZC3H12A | 2.05E-46 | 1.142688408 | 0.584 | 0.407 | 4.04E-42 | Lep-L |
| Macro-FOLR2 | HSPB1 | 2.14E-71 | 1.139761256 | 0.921 | 0.831 | 4.22E-67 | Lep-L |
| Macro-FOLR2 | TWISTNB | 2.42E-24 | 1.128576308 | 0.556 | 0.46 | 4.76E-20 | Lep-L |
| Macro-FOLR2 | APOO | 1.98E-26 | 1.127013794 | 0.404 | 0.286 | 3.90E-22 | Lep-L |
| Macro-FOLR2 | EIF4A3 | 5.76E-76 | 1.126693264 | 0.826 | 0.741 | 1.13E-71 | Lep-L |
| Macro-FOLR2 | MS4A14 | 6.28E-25 | 1.105411059 | 0.304 | 0.179 | 1.24E-20 | Lep-L |
| Macro-FOLR2 | ARRDC2 | 2.15E-20 | 1.097532383 | 0.396 | 0.3 | 4.24E-16 | Lep-L |
| Macro-FOLR2 | PDE8A | 1.60E-31 | 1.097229803 | 0.467 | 0.321 | 3.15E-27 | Lep-L |
| Macro-FOLR2 | PDE4DIP | 2.97E-76 | 1.080455788 | 0.755 | 0.568 | 5.85E-72 | Lep-L |
| Macro-FOLR2 | PPP1R15A | 8.21E-116 | 1.079028723 | 0.948 | 0.892 | 1.62E-111 | Lep-L |
| Macro-FOLR2 | SLC26A11 | 4.85E-12 | 1.071100186 | 0.261 | 0.187 | 9.55E-08 | Lep-L |
| Macro-FOLR2 | SLC29A1 | 6.51E-25 | 1.0698729 | 0.34 | 0.223 | 1.28E-20 | Lep-L |
| Macro-FOLR2 | RBFOX2 | 3.36E-45 | 1.06061007 | 0.587 | 0.443 | 6.61E-41 | Lep-L |
| Macro-FOLR2 | KBTBD8 | 2.00E-26 | 1.052269219 | 0.346 | 0.216 | 3.95E-22 | Lep-L |
| Macro-FOLR2 | ALDOA | 7.90E-21 | 1.039117902 | 0.445 | 0.355 | 1.56E-16 | Lep-L |
| Macro-FOLR2 | KCNJ5 | 7.67E-29 | 1.032220402 | 0.324 | 0.184 | 1.51E-24 | Lep-L |
| Macro-FOLR2 | MMP19 | 7.21E-25 | 1.03159599 | 0.441 | 0.311 | 1.42E-20 | Lep-L |
| Macro-FOLR2 | SMS | 2.19E-69 | 1.031013976 | 0.726 | 0.63 | 4.31E-65 | Lep-L |
| Macro-FOLR2 | CSTB | 8.27E-107 | 1.026755012 | 0.926 | 0.887 | 1.63E-102 | Lep-L |
| Macro-FOLR2 | PRKCH | 5.69E-20 | 1.00055575 | 0.393 | 0.288 | 1.12E-15 | Lep-L |
| Macro-FOLR2 | UBD | 1.14E-34 | 3.036865728 | 0.438 | 0.291 | 2.24E-30 | Lep-T |
| Macro-FOLR2 | CXCL14 | 1.54E-74 | 2.520304673 | 0.344 | 0.115 | 3.03E-70 | Lep-T |
| Macro-FOLR2 | NAV2 | 2.49E-22 | 2.406102837 | 0.269 | 0.16 | 4.89E-18 | Lep-T |
| Macro-FOLR2 | CST3 | 6.27E-172 | 2.053102369 | 0.992 | 0.989 | 1.23E-167 | Lep-T |
| Macro-FOLR2 | SCN9A | 3.74E-29 | 1.8126549 | 0.287 | 0.15 | 7.36E-25 | Lep-T |
| Macro-FOLR2 | DCN | 1.64E-13 | 1.782288532 | 0.266 | 0.178 | 3.24E-09 | Lep-T |

| | | | | | | | |
|---|---|---|---|---|---|---|---|
| Macro-FOLR2 | ABCA6 | 3.91E-36 | 1.770664458 | 0.302 | 0.144 | 7.69E-32 | Lep-T |
| Macro-FOLR2 | PLA2G2D | 1.05E-18 | 1.744895842 | 0.297 | 0.192 | 2.07E-14 | Lep-T |
| Macro-FOLR2 | CCL2 | 1.27E-50 | 1.71607893 | 0.631 | 0.447 | 2.49E-46 | Lep-T |
| Macro-FOLR2 | GFRA2 | 1.14E-46 | 1.511056303 | 0.353 | 0.163 | 2.23E-42 | Lep-T |
| Macro-FOLR2 | PTGDS | 6.14E-43 | 1.48457686 | 0.386 | 0.198 | 1.21E-38 | Lep-T |
| Macro-FOLR2 | AFDN | 5.60E-17 | 1.440132162 | 0.294 | 0.194 | 1.10E-12 | Lep-T |
| Macro-FOLR2 | DHRS9 | 2.52E-13 | 1.397762145 | 0.365 | 0.29 | 4.96E-09 | Lep-T |
| Macro-FOLR2 | CCL13 | 5.63E-19 | 1.325225787 | 0.327 | 0.217 | 1.11E-14 | Lep-T |
| Macro-FOLR2 | PDE4D | 1.26E-09 | 1.298926235 | 0.332 | 0.264 | 2.48E-05 | Lep-T |
| Macro-FOLR2 | EMP1 | 2.28E-10 | 1.293881638 | 0.357 | 0.292 | 4.50E-06 | Lep-T |
| Macro-FOLR2 | PELI1 | 8.56E-10 | 1.275738719 | 0.616 | 0.653 | 1.69E-05 | Lep-T |
| Macro-FOLR2 | RTN1 | 1.54E-07 | 1.240480368 | 0.343 | 0.299 | 0.003037882 | Lep-T |
| Macro-FOLR2 | BEST11 | 3.83E-19 | 1.22402806 | 0.384 | 0.265 | 7.55E-15 | Lep-T |
| Macro-FOLR2 | VCAM1 | 3.50E-07 | 1.222956674 | 0.276 | 0.219 | 0.006886732 | Lep-T |
| Macro-FOLR2 | MT-ND6 | 5.86E-07 | 1.180539359 | 0.265 | 0.206 | 0.011528288 | Lep-T |
| Macro-FOLR2 | CFP | 5.76E-10 | 1.153130022 | 0.291 | 0.221 | 1.13E-05 | Lep-T |
| Macro-FOLR2 | ARHGAP6 | 3.47E-09 | 1.137279465 | 0.262 | 0.196 | 6.83E-05 | Lep-T |
| Macro-FOLR2 | CD74 | 9.84E-62 | 1.074665391 | 0.992 | 0.996 | 1.94E-57 | Lep-T |
| Macro-FOLR2 | COLEC12 | 9.88E-09 | 1.038155056 | 0.447 | 0.404 | 0.000194529 | Lep-T |
| Macro-FOLR2 | SPATS2L | 3.43E-07 | 1.002022112 | 0.544 | 0.568 | 0.006752619 | Lep-T |
| Macro-FOLR2 | FOLR3 | 1.30E-227 | 6.054982082 | 0.326 | 0.011 | 2.55E-223 | Mari |
| Macro-FOLR2 | KRT161 | 2.43E-119 | 3.22715043 | 0.346 | 0.072 | 4.77E-115 | Mari |
| Macro-FOLR2 | S100A71 | 5.54E-94 | 2.806235368 | 0.303 | 0.07 | 1.09E-89 | Mari |
| Macro-FOLR2 | ANK3 | 3.62E-91 | 2.774825089 | 0.284 | 0.059 | 7.13E-87 | Mari |
| Macro-FOLR2 | CD93 | 6.80E-122 | 2.566236242 | 0.395 | 0.092 | 1.34E-117 | Mari |
| Macro-FOLR2 | S100A81 | 4.59E-288 | 2.379090775 | 0.869 | 0.267 | 9.03E-284 | Mari |
| Macro-FOLR2 | CASP51 | 1.75E-112 | 2.361124287 | 0.376 | 0.088 | 3.44E-108 | Mari |
| Macro-FOLR2 | KRT6A1 | 8.94E-59 | 2.288424397 | 0.257 | 0.078 | 1.76E-54 | Mari |
| Macro-FOLR2 | NNMT | 1.99E-45 | 1.973030079 | 0.263 | 0.095 | 3.92E-41 | Mari |
| Macro-FOLR2 | VNN2 | 4.93E-44 | 1.919282028 | 0.254 | 0.091 | 9.71E-40 | Mari |
| Macro-FOLR2 | TMEM1631 | 6.70E-72 | 1.918695637 | 0.332 | 0.101 | 1.32E-67 | Mari |
| Macro-FOLR2 | IL7 | 3.63E-68 | 1.902443816 | 0.338 | 0.11 | 7.14E-64 | Mari |
| Macro-FOLR2 | CR1 | 1.27E-67 | 1.880473121 | 0.522 | 0.264 | 2.50E-63 | Mari |
| Macro-FOLR2 | S100A91 | 9.40E-233 | 1.77379401 | 0.972 | 0.626 | 1.85E-228 | Mari |
| Macro-FOLR2 | SPATA5L1 | 3.15E-29 | 1.766988565 | 0.253 | 0.115 | 6.20E-25 | Mari |
| Macro-FOLR2 | MYO1G1 | 1.75E-71 | 1.713148766 | 0.363 | 0.119 | 3.45E-67 | Mari |
| Macro-FOLR2 | MT1E | 7.23E-55 | 1.707542003 | 0.33 | 0.125 | 1.42E-50 | Mari |
| Macro-FOLR2 | FCGR1B1 | 5.07E-129 | 1.67756424 | 0.667 | 0.268 | 9.99E-125 | Mari |
| Macro-FOLR2 | GAL3ST41 | 9.94E-51 | 1.652641237 | 0.264 | 0.085 | 1.96E-46 | Mari |
| Macro-FOLR2 | FCGR1A1 | 2.19E-190 | 1.636293603 | 0.894 | 0.491 | 4.32E-186 | Mari |
| Macro-FOLR2 | ETV7 | 1.07E-54 | 1.613851365 | 0.254 | 0.073 | 2.11E-50 | Mari |
| Macro-FOLR2 | KRT141 | 3.20E-63 | 1.586915646 | 0.418 | 0.186 | 6.30E-59 | Mari |
| Macro-FOLR2 | PLTP | 4.02E-104 | 1.550640289 | 0.968 | 0.833 | 7.92E-100 | Mari |
| Macro-FOLR2 | OLFML2B1 | 3.63E-119 | 1.505095666 | 0.628 | 0.243 | 7.15E-115 | Mari |
| Macro-FOLR2 | ADAMTS2 | 5.70E-36 | 1.492180359 | 0.251 | 0.099 | 1.12E-31 | Mari |
| Macro-FOLR2 | IL7R1 | 4.82E-53 | 1.457156125 | 0.368 | 0.149 | 9.49E-49 | Mari |
| Macro-FOLR2 | FCGR3A1 | 5.94E-197 | 1.427008533 | 0.958 | 0.586 | 1.17E-192 | Mari |
| Macro-FOLR2 | IGSF21 | 2.69E-68 | 1.382941356 | 0.314 | 0.09 | 5.30E-64 | Mari |
| Macro-FOLR2 | TLR81 | 7.64E-49 | 1.375164012 | 0.397 | 0.179 | 1.50E-44 | Mari |
| Macro-FOLR2 | PCNX11 | 2.04E-106 | 1.341944454 | 0.67 | 0.291 | 4.01E-102 | Mari |
| Macro-FOLR2 | VAMP51 | 1.09E-151 | 1.339439924 | 0.9 | 0.567 | 2.15E-147 | Mari |
| Macro-FOLR2 | TMEM173 | 6.07E-83 | 1.314938206 | 0.529 | 0.216 | 1.19E-78 | Mari |
| Macro-FOLR2 | GIMAP61 | 1.69E-70 | 1.310689815 | 0.431 | 0.163 | 3.33E-66 | Mari |
| Macro-FOLR2 | TLR7 | 9.55E-46 | 1.308626512 | 0.359 | 0.159 | 1.88E-41 | Mari |
| Macro-FOLR2 | CLEC12A1 | 4.31E-98 | 1.303559765 | 0.678 | 0.311 | 8.48E-94 | Mari |

| | | | | | | | |
|---|---|---|---|---|---|---|---|
| Macro-FOLR2 | FAM89B | 1.82E-76 | 1.297321064 | 0.571 | 0.268 | 3.59E-72 | Mari |
| Macro-FOLR2 | 46082 | 1.28E-113 | 1.285385087 | 0.846 | 0.454 | 2.52E-109 | Mari |
| Macro-FOLR2 | DNAJC13 | 3.37E-67 | 1.281602042 | 0.512 | 0.234 | 6.63E-63 | Mari |
| Macro-FOLR2 | ODC1 | 8.25E-38 | 1.26686971 | 0.507 | 0.297 | 1.62E-33 | Mari |
| Macro-FOLR2 | CALHM61 | 2.45E-119 | 1.261435125 | 0.913 | 0.632 | 4.83E-115 | Mari |
| Macro-FOLR2 | CCDC167 | 1.25E-49 | 1.256233884 | 0.356 | 0.143 | 2.46E-45 | Mari |
| Macro-FOLR2 | IFI44L | 6.18E-77 | 1.240413659 | 0.469 | 0.17 | 1.22E-72 | Mari |
| Macro-FOLR2 | CORO1A | 5.20E-69 | 1.239247085 | 0.599 | 0.295 | 1.02E-64 | Mari |
| Macro-FOLR2 | ANKRD221 | 6.52E-67 | 1.227593191 | 0.425 | 0.162 | 1.28E-62 | Mari |
| Macro-FOLR2 | ENPP21 | 8.42E-81 | 1.204088275 | 0.625 | 0.278 | 1.66E-76 | Mari |
| Macro-FOLR2 | GNG101 | 1.43E-140 | 1.20342458 | 0.92 | 0.624 | 2.82E-136 | Mari |
| Macro-FOLR2 | NIBAN1 | 2.60E-33 | 1.203327944 | 0.354 | 0.18 | 5.12E-29 | Mari |
| Macro-FOLR2 | TYMP1 | 2.71E-178 | 1.200643788 | 0.978 | 0.786 | 5.33E-174 | Mari |
| Macro-FOLR2 | GCHFR1 | 7.84E-20 | 1.199856356 | 0.41 | 0.282 | 1.54E-15 | Mari |
| Macro-FOLR2 | MNDA1 | 1.37E-131 | 1.196496418 | 0.88 | 0.501 | 2.70E-127 | Mari |
| Macro-FOLR2 | PCED1B | 7.55E-43 | 1.184065154 | 0.31 | 0.125 | 1.49E-38 | Mari |
| Macro-FOLR2 | IL2RA1 | 2.09E-82 | 1.182840468 | 0.527 | 0.206 | 4.12E-78 | Mari |
| Macro-FOLR2 | VSIG4 | 1.27E-77 | 1.17907209 | 0.798 | 0.536 | 2.50E-73 | Mari |
| Macro-FOLR2 | IFIT31 | 5.20E-56 | 1.178840587 | 0.361 | 0.14 | 1.02E-51 | Mari |
| Macro-FOLR2 | CASP41 | 1.22E-118 | 1.176721412 | 0.854 | 0.496 | 2.41E-114 | Mari |
| Macro-FOLR2 | EIF4G3 | 6.09E-67 | 1.171012059 | 0.676 | 0.38 | 1.20E-62 | Mari |
| Macro-FOLR2 | IL6 | 2.28E-27 | 1.169975825 | 0.264 | 0.12 | 4.49E-23 | Mari |
| Macro-FOLR2 | TNFSF101 | 6.49E-112 | 1.169928012 | 0.816 | 0.439 | 1.28E-107 | Mari |
| Macro-FOLR2 | TUBA1A | 4.03E-83 | 1.140685671 | 0.84 | 0.567 | 7.92E-79 | Mari |
| Macro-FOLR2 | CD691 | 2.78E-25 | 1.139164273 | 0.28 | 0.138 | 5.48E-21 | Mari |
| Macro-FOLR2 | GPR85 | 6.84E-26 | 1.12802178 | 0.257 | 0.121 | 1.35E-21 | Mari |
| Macro-FOLR2 | GBP11 | 1.20E-102 | 1.125235941 | 0.794 | 0.422 | 2.36E-98 | Mari |
| Macro-FOLR2 | BATF | 3.48E-49 | 1.118948344 | 0.369 | 0.153 | 6.86E-45 | Mari |
| Macro-FOLR2 | ITGA4 | 9.55E-48 | 1.114771395 | 0.425 | 0.198 | 1.88E-43 | Mari |
| Macro-FOLR2 | MICAL2 | 2.31E-37 | 1.108369581 | 0.251 | 0.096 | 4.54E-33 | Mari |
| Macro-FOLR2 | CCL8 | 3.50E-42 | 1.106695412 | 0.538 | 0.303 | 6.89E-38 | Mari |
| Macro-FOLR2 | MRPL58 | 4.82E-42 | 1.106080315 | 0.365 | 0.163 | 9.49E-38 | Mari |
| Macro-FOLR2 | RFK | 3.67E-46 | 1.104664741 | 0.382 | 0.168 | 7.22E-42 | Mari |
| Macro-FOLR2 | IFITM3 | 2.60E-95 | 1.104459437 | 0.966 | 0.766 | 5.12E-91 | Mari |
| Macro-FOLR2 | FCER1G1 | 1.96E-197 | 1.103162751 | 0.993 | 0.888 | 3.86E-193 | Mari |
| Macro-FOLR2 | SIGLEC101 | 1.66E-34 | 1.093444889 | 0.376 | 0.192 | 3.26E-30 | Mari |
| Macro-FOLR2 | SCO21 | 6.62E-86 | 1.093045377 | 0.709 | 0.363 | 1.30E-81 | Mari |
| Macro-FOLR2 | HRH21 | 1.88E-52 | 1.092471706 | 0.482 | 0.236 | 3.71E-48 | Mari |
| Macro-FOLR2 | DTX3L | 5.03E-43 | 1.08536245 | 0.448 | 0.227 | 9.90E-39 | Mari |
| Macro-FOLR2 | CCR1 | 7.25E-67 | 1.079373232 | 0.593 | 0.295 | 1.43E-62 | Mari |
| Macro-FOLR2 | PARP9 | 1.41E-68 | 1.076268944 | 0.519 | 0.229 | 2.78E-64 | Mari |
| Macro-FOLR2 | BST2 | 2.26E-158 | 1.075348742 | 0.959 | 0.726 | 4.45E-154 | Mari |
| Macro-FOLR2 | SMCO41 | 1.82E-77 | 1.053486201 | 0.721 | 0.431 | 3.59E-73 | Mari |
| Macro-FOLR2 | STK38L | 4.92E-39 | 1.046283352 | 0.438 | 0.227 | 9.69E-35 | Mari |
| Macro-FOLR2 | MRPL42 | 2.01E-28 | 1.045667291 | 0.3 | 0.146 | 3.95E-24 | Mari |
| Macro-FOLR2 | GIMAP41 | 1.09E-98 | 1.045512438 | 0.804 | 0.445 | 2.15E-94 | Mari |
| Macro-FOLR2 | IFI301 | 7.67E-104 | 1.044665214 | 0.996 | 0.925 | 1.51E-99 | Mari |
| Macro-FOLR2 | MSRB11 | 2.03E-46 | 1.044547767 | 0.391 | 0.174 | 3.99E-42 | Mari |
| Macro-FOLR2 | ADGRE21 | 6.25E-43 | 1.044415065 | 0.52 | 0.285 | 1.23E-38 | Mari |
| Macro-FOLR2 | GIMAP71 | 6.76E-99 | 1.043952463 | 0.657 | 0.272 | 1.33E-94 | Mari |
| Macro-FOLR2 | MRPL23 | 3.85E-70 | 1.042415485 | 0.568 | 0.264 | 7.58E-66 | Mari |
| Macro-FOLR2 | VPS25 | 8.05E-52 | 1.039585484 | 0.485 | 0.237 | 1.58E-47 | Mari |
| Macro-FOLR2 | MT1X | 1.15E-44 | 1.030188719 | 0.59 | 0.345 | 2.27E-40 | Mari |
| Macro-FOLR2 | ZDHHC12 | 6.13E-59 | 1.029098722 | 0.544 | 0.28 | 1.21E-54 | Mari |
| Macro-FOLR2 | NOP10 | 1.04E-90 | 1.028365592 | 0.868 | 0.652 | 2.05E-86 | Mari |

| | | | | | | | |
|---|---|---|---|---|---|---|---|
| Macro-FOLR2 | RGS18 | 1.44E-36 | 1.023360152 | 0.353 | 0.167 | 2.83E-32 | Mari |
| Macro-FOLR2 | BLOC1S1 | 2.06E-135 | 1.016762129 | 0.943 | 0.674 | 4.06E-131 | Mari |
| Macro-FOLR2 | ACTR61 | 7.38E-28 | 1.016731899 | 0.331 | 0.173 | 1.45E-23 | Mari |
| Macro-FOLR2 | XRCC4 | 3.43E-36 | 1.014271764 | 0.337 | 0.154 | 6.75E-32 | Mari |
| Macro-FOLR2 | DYNLT11 | 7.68E-99 | 1.013194087 | 0.837 | 0.507 | 1.51E-94 | Mari |
| Macro-FOLR2 | DSE | 3.41E-90 | 1.012090628 | 0.882 | 0.543 | 6.70E-86 | Mari |
| Macro-FOLR2 | CD141 | 7.76E-142 | 1.006786163 | 0.994 | 0.867 | 1.53E-137 | Mari |
| Macro-FOLR2 | ENPP4 | 1.19E-27 | 1.005920412 | 0.275 | 0.127 | 2.35E-23 | Mari |
| Macro-FOLR2 | FCHSD2 | 7.97E-58 | 1.004788073 | 0.741 | 0.446 | 1.57E-53 | Mari |
| Macro-FOLR2 | HLA-DRB5 | 1.05E-30 | 1.003688102 | 0.888 | 0.748 | 2.06E-26 | Mari |
| Macro-FOLR2 | PLSCR1 | 3.43E-93 | 1.001445971 | 0.9 | 0.58 | 6.75E-89 | Mari |
| Macro-FOLR2 | FN1 | 1.10E-82 | 3.697969869 | 0.379 | 0.112 | 2.17E-78 | Tb |
| Macro-FOLR2 | GGT5 | 2.57E-143 | 3.557631768 | 0.306 | 0.032 | 5.06E-139 | Tb |
| Macro-FOLR2 | SLC35F1 | 2.96E-154 | 2.379726696 | 0.471 | 0.092 | 5.82E-150 | Tb |
| Macro-FOLR2 | FBP1 | 1.81E-66 | 2.326131592 | 0.632 | 0.323 | 3.55E-62 | Tb |
| Macro-FOLR2 | CXCL9 | 5.51E-124 | 2.211259083 | 0.657 | 0.235 | 1.08E-119 | Tb |
| Macro-FOLR2 | PTGDS1 | 4.95E-22 | 1.995364628 | 0.365 | 0.22 | 9.75E-18 | Tb |
| Macro-FOLR2 | SIGLEC10 | 8.07E-129 | 1.995088499 | 0.572 | 0.164 | 1.59E-124 | Tb |
| Macro-FOLR2 | CCL18 | 7.91E-17 | 1.978703831 | 0.669 | 0.582 | 1.56E-12 | Tb |
| Macro-FOLR2 | IL2RA | 1.65E-95 | 1.972990995 | 0.568 | 0.215 | 3.24E-91 | Tb |
| Macro-FOLR2 | SIGLEC12 | 5.46E-75 | 1.936158748 | 0.278 | 0.058 | 1.07E-70 | Tb |
| Macro-FOLR2 | LYZ | 7.98E-144 | 1.91795251 | 0.989 | 0.846 | 1.57E-139 | Tb |
| Macro-FOLR2 | QPCT | 5.35E-57 | 1.915493117 | 0.315 | 0.098 | 1.05E-52 | Tb |
| Macro-FOLR2 | SLC16A10 | 4.86E-48 | 1.873821637 | 0.534 | 0.283 | 9.56E-44 | Tb |
| Macro-FOLR2 | TGM2 | 2.99E-52 | 1.805629089 | 0.358 | 0.131 | 5.89E-48 | Tb |
| Macro-FOLR2 | IL7R | 2.89E-66 | 1.788656556 | 0.427 | 0.149 | 5.68E-62 | Tb |
| Macro-FOLR2 | CHI3L1 | 2.45E-27 | 1.788399129 | 0.3 | 0.145 | 4.81E-23 | Tb |
| Macro-FOLR2 | DNAI2 | 1.36E-52 | 1.736612613 | 0.332 | 0.113 | 2.67E-48 | Tb |
| Macro-FOLR2 | HS3ST3B1 | 6.91E-35 | 1.654005213 | 0.268 | 0.102 | 1.36E-30 | Tb |
| Macro-FOLR2 | SULT1C2 | 1.44E-39 | 1.618546123 | 0.295 | 0.114 | 2.84E-35 | Tb |
| Macro-FOLR2 | KRT17 | 4.87E-63 | 1.599886049 | 0.283 | 0.07 | 9.58E-59 | Tb |
| Macro-FOLR2 | PPA1 | 1.24E-114 | 1.59613035 | 0.756 | 0.338 | 2.44E-110 | Tb |
| Macro-FOLR2 | S100A8 | 1.72E-125 | 1.590690695 | 0.785 | 0.317 | 3.38E-121 | Tb |
| Macro-FOLR2 | KRT6A | 4.94E-72 | 1.58075117 | 0.317 | 0.076 | 9.72E-68 | Tb |
| Macro-FOLR2 | STAT4 | 3.27E-53 | 1.578788835 | 0.405 | 0.159 | 6.44E-49 | Tb |
| Macro-FOLR2 | HOMER1 | 7.57E-70 | 1.570254966 | 0.492 | 0.194 | 1.49E-65 | Tb |
| Macro-FOLR2 | GAL3ST4 | 3.03E-45 | 1.54584969 | 0.28 | 0.091 | 5.96E-41 | Tb |
| Macro-FOLR2 | ADAM8 | 3.35E-72 | 1.514610946 | 0.451 | 0.157 | 6.60E-68 | Tb |
| Macro-FOLR2 | VASH1 | 1.03E-69 | 1.511626541 | 0.402 | 0.13 | 2.03E-65 | Tb |
| Macro-FOLR2 | HAPLN3 | 5.08E-80 | 1.492154046 | 0.471 | 0.154 | 9.99E-76 | Tb |
| Macro-FOLR2 | TYMP | 4.30E-186 | 1.458428272 | 0.993 | 0.794 | 8.47E-182 | Tb |
| Macro-FOLR2 | ACOT7 | 1.65E-90 | 1.449316562 | 0.48 | 0.149 | 3.24E-86 | Tb |
| Macro-FOLR2 | TMEM163 | 8.66E-52 | 1.447981759 | 0.335 | 0.113 | 1.70E-47 | Tb |
| Macro-FOLR2 | CXCL10 | 1.42E-71 | 1.428639583 | 0.595 | 0.259 | 2.80E-67 | Tb |
| Macro-FOLR2 | SUCNR1 | 9.79E-67 | 1.425298554 | 0.431 | 0.153 | 1.93E-62 | Tb |
| Macro-FOLR2 | PKD2 | 2.23E-43 | 1.416447035 | 0.26 | 0.082 | 4.40E-39 | Tb |
| Macro-FOLR2 | USP30-AS1 | 5.10E-37 | 1.405330718 | 0.267 | 0.096 | 1.00E-32 | Tb |
| Macro-FOLR2 | ANKRD22 | 2.23E-55 | 1.400239367 | 0.438 | 0.174 | 4.40E-51 | Tb |
| Macro-FOLR2 | SGPL1 | 1.02E-91 | 1.396069572 | 0.714 | 0.337 | 2.00E-87 | Tb |
| Macro-FOLR2 | SATB1-AS1 | 1.63E-31 | 1.395585293 | 0.357 | 0.17 | 3.22E-27 | Tb |
| Macro-FOLR2 | JAK3 | 7.92E-38 | 1.38429747 | 0.271 | 0.096 | 1.56E-33 | Tb |
| Macro-FOLR2 | S100A2 | 3.72E-72 | 1.380774129 | 0.326 | 0.081 | 7.32E-68 | Tb |
| Macro-FOLR2 | ENPP2 | 1.25E-111 | 1.359416956 | 0.717 | 0.279 | 2.47E-107 | Tb |
| Macro-FOLR2 | ANOS1 | 2.98E-34 | 1.355107337 | 0.268 | 0.101 | 5.86E-30 | Tb |
| Macro-FOLR2 | APOL4 | 3.28E-67 | 1.34699739 | 0.404 | 0.131 | 6.45E-63 | Tb |

| | | | | | | | |
|---|---|---|---|---|---|---|---|
| Macro-FOLR2 | SLC2A6 | 4.79E-56 | 1.329534493 | 0.445 | 0.176 | 9.43E-52 | Tb |
| Macro-FOLR2 | CD274 | 2.41E-59 | 1.318480024 | 0.446 | 0.171 | 4.75E-55 | Tb |
| Macro-FOLR2 | TMEM39A | 8.18E-37 | 1.314310032 | 0.354 | 0.153 | 1.61E-32 | Tb |
| Macro-FOLR2 | ITPR1 | 4.73E-62 | 1.303237946 | 0.549 | 0.247 | 9.30E-58 | Tb |
| Macro-FOLR2 | GBP1 | 4.07E-103 | 1.301673584 | 0.863 | 0.429 | 8.02E-99 | Tb |
| Macro-FOLR2 | CTSC | 3.88E-163 | 1.298673647 | 0.99 | 0.869 | 7.63E-159 | Tb |
| Macro-FOLR2 | PFKP | 8.63E-60 | 1.296886575 | 0.382 | 0.13 | 1.70E-55 | Tb |
| Macro-FOLR2 | CD300LF | 1.38E-55 | 1.276730385 | 0.458 | 0.187 | 2.71E-51 | Tb |
| Macro-FOLR2 | CISH | 1.18E-51 | 1.269286246 | 0.306 | 0.098 | 2.32E-47 | Tb |
| Macro-FOLR2 | LIG1 | 9.17E-37 | 1.25603073 | 0.265 | 0.095 | 1.81E-32 | Tb |
| Macro-FOLR2 | GTF2IRD1 | 5.01E-37 | 1.249440719 | 0.419 | 0.204 | 9.86E-33 | Tb |
| Macro-FOLR2 | ACE | 9.44E-57 | 1.243215404 | 0.391 | 0.14 | 1.86E-52 | Tb |
| Macro-FOLR2 | SERINC2 | 7.37E-49 | 1.236505968 | 0.29 | 0.093 | 1.45E-44 | Tb |
| Macro-FOLR2 | CD40 | 3.16E-101 | 1.229816885 | 0.776 | 0.364 | 6.21E-97 | Tb |
| Macro-FOLR2 | SPART-AS1 | 1.30E-25 | 1.221571887 | 0.313 | 0.15 | 2.56E-21 | Tb |
| Macro-FOLR2 | APOL3 | 9.25E-62 | 1.216679081 | 0.469 | 0.181 | 1.82E-57 | Tb |
| Macro-FOLR2 | LIMK2 | 7.55E-63 | 1.195198728 | 0.487 | 0.189 | 1.49E-58 | Tb |
| Macro-FOLR2 | IL21R | 1.43E-50 | 1.193916678 | 0.371 | 0.135 | 2.82E-46 | Tb |
| Macro-FOLR2 | WNT2B | 1.51E-25 | 1.19011979 | 0.26 | 0.113 | 2.98E-21 | Tb |
| Macro-FOLR2 | SLAMF7 | 1.28E-80 | 1.16782176 | 0.695 | 0.329 | 2.53E-76 | Tb |
| Macro-FOLR2 | DNASE2 | 7.58E-83 | 1.165902792 | 0.845 | 0.508 | 1.49E-78 | Tb |
| Macro-FOLR2 | NFAM1 | 2.75E-67 | 1.165565869 | 0.606 | 0.278 | 5.41E-63 | Tb |
| Macro-FOLR2 | P2RY13 | 2.06E-65 | 1.162599555 | 0.58 | 0.258 | 4.05E-61 | Tb |
| Macro-FOLR2 | TSFM | 8.91E-45 | 1.158242497 | 0.361 | 0.139 | 1.75E-40 | Tb |
| Macro-FOLR2 | MMP14 | 4.78E-80 | 1.145541862 | 0.648 | 0.278 | 9.41E-76 | Tb |
| Macro-FOLR2 | FLT1 | 2.38E-19 | 1.142939786 | 0.272 | 0.136 | 4.68E-15 | Tb |
| Macro-FOLR2 | SMAD7 | 3.12E-48 | 1.142674864 | 0.438 | 0.185 | 6.14E-44 | Tb |
| Macro-FOLR2 | ATF7IP2 | 2.55E-47 | 1.140349213 | 0.402 | 0.163 | 5.03E-43 | Tb |
| Macro-FOLR2 | FCGR3A | 1.03E-82 | 1.131343372 | 0.906 | 0.617 | 2.02E-78 | Tb |
| Macro-FOLR2 | RFTN1 | 2.36E-51 | 1.131166742 | 0.539 | 0.254 | 4.65E-47 | Tb |
| Macro-FOLR2 | SIRPB2 | 1.66E-71 | 1.113686875 | 0.312 | 0.075 | 3.26E-67 | Tb |
| Macro-FOLR2 | IL32 | 1.58E-68 | 1.103460284 | 0.74 | 0.406 | 3.11E-64 | Tb |
| Macro-FOLR2 | S100A9 | 3.25E-68 | 1.095185718 | 0.93 | 0.654 | 6.39E-64 | Tb |
| Macro-FOLR2 | MRC2 | 4.21E-27 | 1.091585654 | 0.25 | 0.101 | 8.29E-23 | Tb |
| Macro-FOLR2 | PLAAT4 | 1.17E-75 | 1.089919037 | 0.866 | 0.511 | 2.31E-71 | Tb |
| Macro-FOLR2 | GIMAP7 | 5.04E-72 | 1.088476072 | 0.661 | 0.292 | 9.91E-68 | Tb |
| Macro-FOLR2 | DUSP16 | 3.29E-37 | 1.082545091 | 0.435 | 0.211 | 6.47E-33 | Tb |
| Macro-FOLR2 | PTAFR | 2.84E-77 | 1.078736549 | 0.731 | 0.36 | 5.58E-73 | Tb |
| Macro-FOLR2 | PDCD1LG2 | 1.61E-87 | 1.069833486 | 0.661 | 0.266 | 3.17E-83 | Tb |
| Macro-FOLR2 | FAM135A | 3.75E-48 | 1.060866734 | 0.402 | 0.158 | 7.38E-44 | Tb |
| Macro-FOLR2 | ABCC3 | 1.37E-66 | 1.056087976 | 0.487 | 0.183 | 2.70E-62 | Tb |
| Macro-FOLR2 | TRAF2 | 5.57E-40 | 1.052469769 | 0.305 | 0.112 | 1.10E-35 | Tb |
| Macro-FOLR2 | IKBKE | 6.62E-45 | 1.050753687 | 0.376 | 0.147 | 1.30E-40 | Tb |
| Macro-FOLR2 | ADAMDEC1 | 1.05E-88 | 1.049203448 | 0.84 | 0.45 | 2.07E-84 | Tb |
| Macro-FOLR2 | MAGOHB | 2.37E-30 | 1.045316919 | 0.338 | 0.151 | 4.66E-26 | Tb |
| Macro-FOLR2 | NFIL3 | 5.47E-36 | 1.04509683 | 0.59 | 0.366 | 1.08E-31 | Tb |
| Macro-FOLR2 | CSF2RB | 1.14E-54 | 1.04462146 | 0.535 | 0.238 | 2.25E-50 | Tb |
| Macro-FOLR2 | CDKN2B | 5.13E-32 | 1.044027044 | 0.257 | 0.096 | 1.01E-27 | Tb |
| Macro-FOLR2 | TMIGD31 | 7.51E-12 | 1.043693865 | 0.327 | 0.219 | 1.48E-07 | Tb |
| Macro-FOLR2 | CCDC138 | 1.47E-36 | 1.037523809 | 0.449 | 0.231 | 2.89E-32 | Tb |
| Macro-FOLR2 | C1RL | 8.88E-41 | 1.031132667 | 0.253 | 0.081 | 1.75E-36 | Tb |
| Macro-FOLR2 | ATP13A2 | 4.10E-42 | 1.02980832 | 0.447 | 0.203 | 8.07E-38 | Tb |
| Macro-FOLR2 | MT-ATP6 | 2.90E-92 | 1.028565824 | 0.997 | 0.93 | 5.71E-88 | Tb |
| Macro-FOLR2 | TJP2 | 4.91E-26 | 1.025296607 | 0.265 | 0.114 | 9.66E-22 | Tb |
| Macro-FOLR2 | SEMA4D | 1.74E-55 | 1.02488706 | 0.458 | 0.185 | 3.42E-51 | Tb |

| | | | | | | | |
|---|---|---|---|---|---|---|---|
| Macro-FOLR2 | RHOH | 3.19E-16 | 1.018639574 | 0.254 | 0.134 | 6.29E-12 | Tb |
| Macro-FOLR2 | ORAI2 | 4.04E-44 | 1.018410217 | 0.36 | 0.139 | 7.95E-40 | Tb |
| Macro-FOLR2 | PTPN22 | 1.95E-27 | 1.017257595 | 0.274 | 0.115 | 3.83E-23 | Tb |
| Macro-FOLR2 | FOXK1 | 6.68E-27 | 1.012835419 | 0.312 | 0.144 | 1.32E-22 | Tb |
| Macro-FOLR2 | LRRK2 | 2.42E-42 | 1.01114571 | 0.436 | 0.195 | 4.77E-38 | Tb |
| Macro-FOLR2 | RALA | 7.66E-55 | 1.009267132 | 0.75 | 0.454 | 1.51E-50 | Tb |
| Macro-FOLR2 | ACVRL1 | 9.48E-33 | 1.004072597 | 0.261 | 0.098 | 1.87E-28 | Tb |
| Macro-FOLR2 | SCO2 | 4.03E-71 | 1.001223156 | 0.758 | 0.373 | 7.94E-67 | Tb |
| **Disease-associated genes in Macro-TREM2** | | | | | | | |
| Macro-TREM2 | TTTY14 | 7.43E-94 | 3.839588693 | 0.25 | 0.021 | 1.46E-89 | Lep-L |
| Macro-TREM2 | ALKAL2 | 3.34E-93 | 3.751099572 | 0.302 | 0.057 | 6.58E-89 | Lep-L |
| Macro-TREM2 | SDS | 2.05E-154 | 3.015050751 | 0.633 | 0.295 | 4.04E-150 | Lep-L |
| Macro-TREM2 | CAPN3 | 8.17E-93 | 2.995527507 | 0.334 | 0.082 | 1.61E-88 | Lep-L |
| Macro-TREM2 | CCR7 | 1.33E-66 | 2.816777871 | 0.357 | 0.142 | 2.62E-62 | Lep-L |
| Macro-TREM2 | TSPAN10 | 9.02E-69 | 2.790015195 | 0.298 | 0.088 | 1.78E-64 | Lep-L |
| Macro-TREM2 | TMIGD3 | 3.82E-96 | 2.704631063 | 0.306 | 0.05 | 7.51E-92 | Lep-L |
| Macro-TREM2 | ZNF703 | 2.53E-87 | 2.637901413 | 0.327 | 0.081 | 4.99E-83 | Lep-L |
| Macro-TREM2 | PLIN2 | 3.39E-289 | 2.631582118 | 0.856 | 0.618 | 6.67E-285 | Lep-L |
| Macro-TREM2 | DDX3Y | 3.28E-230 | 2.561259605 | 0.585 | 0.084 | 6.45E-226 | Lep-L |
| Macro-TREM2 | ZNF503 | 9.45E-70 | 2.533135333 | 0.275 | 0.068 | 1.86E-65 | Lep-L |
| Macro-TREM2 | CDC42EP3 | 5.97E-210 | 2.528231881 | 0.714 | 0.299 | 1.18E-205 | Lep-L |
| Macro-TREM2 | CCL20 | 1.15E-39 | 2.476626079 | 0.321 | 0.161 | 2.26E-35 | Lep-L |
| Macro-TREM2 | PPARG | 2.34E-126 | 2.475796847 | 0.55 | 0.231 | 4.61E-122 | Lep-L |
| Macro-TREM2 | KLHL21 | 2.73E-99 | 2.449398154 | 0.451 | 0.182 | 5.38E-95 | Lep-L |
| Macro-TREM2 | PDK4 | 2.18E-106 | 2.431284861 | 0.324 | 0.048 | 4.28E-102 | Lep-L |
| Macro-TREM2 | HS3ST2 | 5.82E-185 | 2.399839435 | 0.628 | 0.187 | 1.15E-180 | Lep-L |
| Macro-TREM2 | SLC19A2 | 9.39E-112 | 2.374131578 | 0.516 | 0.222 | 1.85E-107 | Lep-L |
| Macro-TREM2 | MMP19 | 7.18E-74 | 2.339943252 | 0.419 | 0.186 | 1.41E-69 | Lep-L |
| Macro-TREM2 | MS4A14 | 4.23E-63 | 2.311055271 | 0.265 | 0.073 | 8.32E-59 | Lep-L |
| Macro-TREM2 | PKD2L1 | 4.43E-68 | 2.219307784 | 0.354 | 0.138 | 8.72E-64 | Lep-L |
| Macro-TREM2 | APBB3 | 1.10E-42 | 2.202114726 | 0.265 | 0.109 | 2.17E-38 | Lep-L |
| Macro-TREM2 | CXCL8 | 6.72E-187 | 2.150548674 | 0.821 | 0.524 | 1.32E-182 | Lep-L |
| Macro-TREM2 | BIRC7 | 1.15E-53 | 2.142449553 | 0.276 | 0.094 | 2.27E-49 | Lep-L |
| Macro-TREM2 | LY9 | 1.58E-39 | 2.092757626 | 0.254 | 0.105 | 3.12E-35 | Lep-L |
| Macro-TREM2 | OTOA | 1.01E-180 | 2.086277574 | 0.741 | 0.409 | 1.98E-176 | Lep-L |
| Macro-TREM2 | GPR183 | 1.07E-223 | 2.073438267 | 0.828 | 0.499 | 2.11E-219 | Lep-L |
| Macro-TREM2 | HES4 | 4.17E-44 | 2.031480011 | 0.288 | 0.122 | 8.21E-40 | Lep-L |
| Macro-TREM2 | MFSD2A | 1.83E-81 | 2.015593151 | 0.491 | 0.258 | 3.60E-77 | Lep-L |
| Macro-TREM2 | CD36 | 7.88E-80 | 1.994020501 | 0.259 | 0.039 | 1.55E-75 | Lep-L |
| Macro-TREM2 | CFD | 1.22E-235 | 1.99398813 | 0.85 | 0.568 | 2.40E-231 | Lep-L |
| Macro-TREM2 | CXCR4 | 1.93E-183 | 1.970657437 | 0.801 | 0.539 | 3.80E-179 | Lep-L |
| Macro-TREM2 | SMPDL3A | 2.79E-83 | 1.965643106 | 0.512 | 0.272 | 5.49E-79 | Lep-L |
| Macro-TREM2 | HSPA2 | 1.14E-36 | 1.938830854 | 0.297 | 0.149 | 2.24E-32 | Lep-L |
| Macro-TREM2 | PHACTR1 | 1.91E-167 | 1.908262717 | 0.746 | 0.426 | 3.77E-163 | Lep-L |
| Macro-TREM2 | TM4SF19 | 4.21E-33 | 1.893705854 | 0.329 | 0.185 | 8.30E-29 | Lep-L |
| Macro-TREM2 | CXCL1 | 9.20E-62 | 1.870703503 | 0.416 | 0.187 | 1.81E-57 | Lep-L |
| Macro-TREM2 | SLC18B1 | 4.97E-68 | 1.86359813 | 0.352 | 0.129 | 9.78E-64 | Lep-L |
| Macro-TREM2 | FEM1C | 1.90E-45 | 1.861872959 | 0.33 | 0.16 | 3.75E-41 | Lep-L |
| Macro-TREM2 | CXCL3 | 2.26E-151 | 1.851574378 | 0.771 | 0.426 | 4.45E-147 | Lep-L |
| Macro-TREM2 | ABCC5 | 5.40E-42 | 1.847529935 | 0.28 | 0.12 | 1.06E-37 | Lep-L |
| Macro-TREM2 | SNHG25 | 1.74E-31 | 1.843064339 | 0.366 | 0.227 | 3.42E-27 | Lep-L |
| Macro-TREM2 | OPN3 | 1.77E-80 | 1.842709432 | 0.529 | 0.307 | 3.48E-76 | Lep-L |
| Macro-TREM2 | CXCL2 | 2.55E-177 | 1.792534984 | 0.861 | 0.56 | 5.02E-173 | Lep-L |
| Macro-TREM2 | BCL2 | 4.30E-75 | 1.778302456 | 0.497 | 0.264 | 8.47E-71 | Lep-L |
| Macro-TREM2 | CKS2 | 9.54E-154 | 1.774537932 | 0.769 | 0.517 | 1.88E-149 | Lep-L |

| | | | | | | | |
|---|---|---|---|---|---|---|---|
| Macro-TREM2 | TAGAP | 1.21E-113 | 1.774026112 | 0.729 | 0.482 | 2.37E-109 | Lep-L |
| Macro-TREM2 | CMSS1 | 3.93E-23 | 1.768788749 | 0.41 | 0.314 | 7.73E-19 | Lep-L |
| Macro-TREM2 | C5AR1 | 8.11E-125 | 1.768377354 | 0.734 | 0.506 | 1.60E-120 | Lep-L |
| Macro-TREM2 | H2AFX | 7.70E-75 | 1.748651484 | 0.508 | 0.28 | 1.51E-70 | Lep-L |
| Macro-TREM2 | PDE6G | 6.58E-70 | 1.71632079 | 0.387 | 0.154 | 1.30E-65 | Lep-L |
| Macro-TREM2 | SERPINH1 | 3.60E-31 | 1.716176096 | 0.334 | 0.197 | 7.09E-27 | Lep-L |
| Macro-TREM2 | PHLDA1 | 6.75E-58 | 1.713126116 | 0.34 | 0.136 | 1.33E-53 | Lep-L |
| Macro-TREM2 | KMO | 1.12E-86 | 1.691288124 | 0.662 | 0.45 | 2.20E-82 | Lep-L |
| Macro-TREM2 | MAP4K4 | 2.51E-47 | 1.684541462 | 0.439 | 0.252 | 4.94E-43 | Lep-L |
| Macro-TREM2 | KCNJ5 | 3.15E-37 | 1.677678419 | 0.311 | 0.158 | 6.21E-33 | Lep-L |
| Macro-TREM2 | ITPKB | 5.59E-55 | 1.658153116 | 0.4 | 0.202 | 1.10E-50 | Lep-L |
| Macro-TREM2 | MIR155HG | 1.53E-68 | 1.64645268 | 0.51 | 0.269 | 3.01E-64 | Lep-L |
| Macro-TREM2 | LFNG | 3.41E-36 | 1.640065942 | 0.273 | 0.126 | 6.72E-32 | Lep-L |
| Macro-TREM2 | TSC22D2 | 1.28E-99 | 1.638749573 | 0.647 | 0.39 | 2.52E-95 | Lep-L |
| Macro-TREM2 | ERF | 3.94E-40 | 1.623776103 | 0.276 | 0.121 | 7.75E-36 | Lep-L |
| Macro-TREM2 | SH3D21 | 1.36E-39 | 1.620183848 | 0.354 | 0.201 | 2.69E-35 | Lep-L |
| Macro-TREM2 | DDIT4 | 1.70E-108 | 1.614321601 | 0.685 | 0.395 | 3.35E-104 | Lep-L |
| Macro-TREM2 | HSPA1B | 2.18E-106 | 1.607111067 | 0.901 | 0.699 | 4.29E-102 | Lep-L |
| Macro-TREM2 | MAFF | 3.32E-92 | 1.606072084 | 0.645 | 0.413 | 6.54E-88 | Lep-L |
| Macro-TREM2 | EMP1 | 1.10E-47 | 1.598706609 | 0.312 | 0.133 | 2.17E-43 | Lep-L |
| Macro-TREM2 | REV3L | 1.86E-73 | 1.598418812 | 0.537 | 0.311 | 3.67E-69 | Lep-L |
| Macro-TREM2 | STMN1 | 6.48E-114 | 1.594678557 | 0.435 | 0.117 | 1.28E-109 | Lep-L |
| Macro-TREM2 | TBC1D8 | 2.69E-38 | 1.591974411 | 0.336 | 0.18 | 5.30E-34 | Lep-L |
| Macro-TREM2 | KBTBD8 | 2.82E-48 | 1.585210945 | 0.341 | 0.162 | 5.56E-44 | Lep-L |
| Macro-TREM2 | SHB | 4.17E-26 | 1.582345241 | 0.329 | 0.211 | 8.21E-22 | Lep-L |
| Macro-TREM2 | LPCAT1 | 8.85E-42 | 1.577460273 | 0.398 | 0.236 | 1.74E-37 | Lep-L |
| Macro-TREM2 | GPAT3 | 5.12E-35 | 1.575939291 | 0.267 | 0.121 | 1.01E-30 | Lep-L |
| Macro-TREM2 | PFKFB3 | 2.84E-99 | 1.571061098 | 0.65 | 0.387 | 5.60E-95 | Lep-L |
| Macro-TREM2 | MAP2K3 | 6.10E-88 | 1.569275468 | 0.645 | 0.449 | 1.20E-83 | Lep-L |
| Macro-TREM2 | TPRA1 | 7.67E-52 | 1.567712436 | 0.531 | 0.371 | 1.51E-47 | Lep-L |
| Macro-TREM2 | HUS1 | 1.63E-22 | 1.561967898 | 0.333 | 0.232 | 3.22E-18 | Lep-L |
| Macro-TREM2 | ARRDC2 | 3.19E-41 | 1.55597267 | 0.381 | 0.224 | 6.28E-37 | Lep-L |
| Macro-TREM2 | ADM | 9.65E-60 | 1.555553051 | 0.566 | 0.354 | 1.90E-55 | Lep-L |
| Macro-TREM2 | GBE1 | 6.33E-47 | 1.549581945 | 0.533 | 0.368 | 1.25E-42 | Lep-L |
| Macro-TREM2 | KLF4 | 1.79E-99 | 1.542796978 | 0.729 | 0.505 | 3.52E-95 | Lep-L |
| Macro-TREM2 | NBPF14 | 1.59E-34 | 1.535380886 | 0.314 | 0.172 | 3.13E-30 | Lep-L |
| Macro-TREM2 | ETV3 | 1.66E-71 | 1.5329463 | 0.504 | 0.283 | 3.27E-67 | Lep-L |
| Macro-TREM2 | SNHG12 | 2.01E-82 | 1.531933052 | 0.648 | 0.453 | 3.95E-78 | Lep-L |
| Macro-TREM2 | HSPA6 | 5.47E-99 | 1.528584579 | 0.673 | 0.359 | 1.08E-94 | Lep-L |
| Macro-TREM2 | FABP5 | 9.34E-91 | 1.520019478 | 0.78 | 0.647 | 1.84E-86 | Lep-L |
| Macro-TREM2 | TSPYL2 | 2.81E-48 | 1.514268802 | 0.348 | 0.156 | 5.53E-44 | Lep-L |
| Macro-TREM2 | CCRL2 | 1.03E-67 | 1.510582574 | 0.598 | 0.413 | 2.03E-63 | Lep-L |
| Macro-TREM2 | KDM6B | 1.13E-104 | 1.50744908 | 0.645 | 0.382 | 2.23E-100 | Lep-L |
| Macro-TREM2 | MAP3K8 | 1.78E-153 | 1.502816731 | 0.807 | 0.584 | 3.51E-149 | Lep-L |
| Macro-TREM2 | UGCG | 6.59E-98 | 1.502081887 | 0.676 | 0.475 | 1.30E-93 | Lep-L |
| Macro-TREM2 | EZR | 2.68E-83 | 1.499347639 | 0.706 | 0.52 | 5.27E-79 | Lep-L |
| Macro-TREM2 | TIPARP | 6.73E-86 | 1.494170766 | 0.642 | 0.42 | 1.32E-81 | Lep-L |
| Macro-TREM2 | LILRA2 | 1.31E-35 | 1.4924209 | 0.256 | 0.113 | 2.58E-31 | Lep-L |
| Macro-TREM2 | BRD2 | 9.81E-112 | 1.4670907 | 0.749 | 0.538 | 1.93E-107 | Lep-L |
| Macro-TREM2 | FKBP4 | 4.77E-58 | 1.463420343 | 0.437 | 0.231 | 9.38E-54 | Lep-L |
| Macro-TREM2 | USP37 | 7.59E-35 | 1.462359639 | 0.384 | 0.239 | 1.49E-30 | Lep-L |
| Macro-TREM2 | CD83 | 6.39E-193 | 1.454305271 | 0.909 | 0.731 | 1.26E-188 | Lep-L |
| Macro-TREM2 | BAG3 | 9.15E-74 | 1.444041504 | 0.643 | 0.416 | 1.80E-69 | Lep-L |
| Macro-TREM2 | VPS37B | 1.90E-39 | 1.440669931 | 0.5 | 0.357 | 3.73E-35 | Lep-L |
| Macro-TREM2 | CREM | 2.54E-111 | 1.429328402 | 0.741 | 0.535 | 5.00E-107 | Lep-L |

| | | | | | | | |
|---|---|---|---|---|---|---|---|
| Macro-TREM2 | CD300A | 1.19E-75 | 1.427169303 | 0.519 | 0.265 | 2.34E-71 | Lep-L |
| Macro-TREM2 | SPSB1 | 6.79E-18 | 1.420790758 | 0.381 | 0.296 | 1.34E-13 | Lep-L |
| Macro-TREM2 | MKNK2 | 1.17E-55 | 1.41902666 | 0.511 | 0.335 | 2.31E-51 | Lep-L |
| Macro-TREM2 | STAM | 1.26E-32 | 1.417170591 | 0.37 | 0.24 | 2.49E-28 | Lep-L |
| Macro-TREM2 | BEST1 | 4.90E-34 | 1.416966949 | 0.401 | 0.253 | 9.65E-30 | Lep-L |
| Macro-TREM2 | HSPB1 | 1.80E-64 | 1.415903669 | 0.874 | 0.724 | 3.55E-60 | Lep-L |
| Macro-TREM2 | VEGFA | 2.55E-73 | 1.415540559 | 0.609 | 0.394 | 5.02E-69 | Lep-L |
| Macro-TREM2 | PAF1 | 2.05E-21 | 1.407936519 | 0.281 | 0.178 | 4.04E-17 | Lep-L |
| Macro-TREM2 | CSGALNACT2 | 6.22E-70 | 1.406566913 | 0.655 | 0.487 | 1.22E-65 | Lep-L |
| Macro-TREM2 | B3GNT2 | 2.13E-35 | 1.404296924 | 0.471 | 0.35 | 4.20E-31 | Lep-L |
| Macro-TREM2 | CCDC107 | 2.02E-96 | 1.403928483 | 0.705 | 0.527 | 3.97E-92 | Lep-L |
| Macro-TREM2 | EIF4A3 | 2.00E-101 | 1.402572951 | 0.806 | 0.631 | 3.93E-97 | Lep-L |
| Macro-TREM2 | PDE7A | 1.37E-43 | 1.400992503 | 0.442 | 0.275 | 2.70E-39 | Lep-L |
| Macro-TREM2 | BCL2A1 | 5.82E-152 | 1.39441306 | 0.864 | 0.737 | 1.15E-147 | Lep-L |
| Macro-TREM2 | THAP2 | 8.80E-21 | 1.392616864 | 0.28 | 0.174 | 1.73E-16 | Lep-L |
| Macro-TREM2 | MEF2D | 3.92E-33 | 1.389248962 | 0.37 | 0.233 | 7.71E-29 | Lep-L |
| Macro-TREM2 | FNIP2 | 8.50E-118 | 1.383532643 | 0.798 | 0.574 | 1.67E-113 | Lep-L |
| Macro-TREM2 | TGIF1 | 6.78E-83 | 1.383239902 | 0.637 | 0.424 | 1.33E-78 | Lep-L |
| Macro-TREM2 | UAP1 | 1.34E-31 | 1.381718923 | 0.404 | 0.264 | 2.63E-27 | Lep-L |
| Macro-TREM2 | CSF1 | 1.06E-21 | 1.381257272 | 0.391 | 0.288 | 2.10E-17 | Lep-L |
| Macro-TREM2 | BLVRB | 1.22E-192 | 1.377548541 | 0.861 | 0.757 | 2.40E-188 | Lep-L |
| Macro-TREM2 | CBX4 | 6.78E-12 | 1.36742284 | 0.287 | 0.221 | 1.33E-07 | Lep-L |
| Macro-TREM2 | SLFN11 | 1.26E-31 | 1.359822513 | 0.415 | 0.287 | 2.48E-27 | Lep-L |
| Macro-TREM2 | ATP1B1 | 5.07E-80 | 1.351616065 | 0.713 | 0.545 | 9.97E-76 | Lep-L |
| Macro-TREM2 | TWISTNB | 1.37E-30 | 1.347850198 | 0.575 | 0.46 | 2.70E-26 | Lep-L |
| Macro-TREM2 | RASGRP3 | 6.01E-41 | 1.34467257 | 0.508 | 0.351 | 1.18E-36 | Lep-L |
| Macro-TREM2 | SDC4 | 8.43E-25 | 1.342238178 | 0.479 | 0.388 | 1.66E-20 | Lep-L |
| Macro-TREM2 | TCEAL9 | 1.19E-26 | 1.340504713 | 0.345 | 0.217 | 2.33E-22 | Lep-L |
| Macro-TREM2 | RBM38 | 3.12E-22 | 1.336545524 | 0.336 | 0.238 | 6.15E-18 | Lep-L |
| Macro-TREM2 | GADD45B | 1.30E-105 | 1.336162553 | 0.907 | 0.758 | 2.56E-101 | Lep-L |
| Macro-TREM2 | HERPUD1 | 1.73E-107 | 1.334135622 | 0.822 | 0.707 | 3.40E-103 | Lep-L |
| Macro-TREM2 | BAIAP2 | 2.50E-19 | 1.332268288 | 0.304 | 0.205 | 4.91E-15 | Lep-L |
| Macro-TREM2 | TESK1 | 3.31E-29 | 1.324254197 | 0.307 | 0.182 | 6.51E-25 | Lep-L |
| Macro-TREM2 | TLE3 | 5.28E-31 | 1.31387185 | 0.356 | 0.224 | 1.04E-26 | Lep-L |
| Macro-TREM2 | DEXI | 2.30E-33 | 1.303063458 | 0.326 | 0.183 | 4.53E-29 | Lep-L |
| Macro-TREM2 | PPP1R15B | 1.50E-75 | 1.301084653 | 0.642 | 0.438 | 2.95E-71 | Lep-L |
| Macro-TREM2 | AVPI1 | 1.52E-62 | 1.295546519 | 0.627 | 0.448 | 2.99E-58 | Lep-L |
| Macro-TREM2 | SPTBN1 | 4.05E-43 | 1.294949593 | 0.312 | 0.138 | 7.98E-39 | Lep-L |
| Macro-TREM2 | SMIM3 | 8.15E-42 | 1.293699604 | 0.516 | 0.382 | 1.60E-37 | Lep-L |
| Macro-TREM2 | MAFB | 1.02E-57 | 1.293417372 | 0.784 | 0.653 | 2.02E-53 | Lep-L |
| Macro-TREM2 | PIK3R5 | 3.02E-43 | 1.292321396 | 0.488 | 0.326 | 5.94E-39 | Lep-L |
| Macro-TREM2 | CDKN1C | 8.76E-19 | 1.285425966 | 0.251 | 0.145 | 1.72E-14 | Lep-L |
| Macro-TREM2 | CDKN1A | 2.87E-82 | 1.283776353 | 0.762 | 0.567 | 5.64E-78 | Lep-L |
| Macro-TREM2 | SLC1A3 | 2.22E-57 | 1.282663806 | 0.734 | 0.58 | 4.37E-53 | Lep-L |
| Macro-TREM2 | ANKRD37 | 9.06E-48 | 1.282539854 | 0.527 | 0.359 | 1.78E-43 | Lep-L |
| Macro-TREM2 | PNP | 1.16E-24 | 1.28200679 | 0.418 | 0.313 | 2.29E-20 | Lep-L |
| Macro-TREM2 | GNG7 | 1.09E-25 | 1.28200328 | 0.348 | 0.221 | 2.15E-21 | Lep-L |
| Macro-TREM2 | RGS1 | 1.51E-54 | 1.27968873 | 0.745 | 0.57 | 2.97E-50 | Lep-L |
| Macro-TREM2 | HSPA1A | 1.60E-159 | 1.276775969 | 0.952 | 0.836 | 3.16E-155 | Lep-L |
| Macro-TREM2 | RARA | 9.30E-46 | 1.275792688 | 0.386 | 0.213 | 1.83E-41 | Lep-L |
| Macro-TREM2 | MGLL | 1.15E-119 | 1.275520189 | 0.765 | 0.498 | 2.27E-115 | Lep-L |
| Macro-TREM2 | TSC22D3 | 1.27E-42 | 1.273623776 | 0.654 | 0.544 | 2.50E-38 | Lep-L |
| Macro-TREM2 | EIF2AK3 | 3.47E-59 | 1.270026564 | 0.548 | 0.332 | 6.84E-55 | Lep-L |
| Macro-TREM2 | HIVEP2 | 1.57E-55 | 1.268155853 | 0.482 | 0.288 | 3.08E-51 | Lep-L |
| Macro-TREM2 | DEDD2 | 6.01E-71 | 1.261759421 | 0.59 | 0.394 | 1.18E-66 | Lep-L |

| | | | | | | | |
|---|---|---|---|---|---|---|---|
| Macro-TREM2 | FOSB | 3.67E-92 | 1.259901347 | 0.759 | 0.558 | 7.23E-88 | Lep-L |
| Macro-TREM2 | MAP7D1 | 4.92E-47 | 1.258550743 | 0.57 | 0.44 | 9.69E-43 | Lep-L |
| Macro-TREM2 | DNAJB9 | 4.99E-57 | 1.254948462 | 0.51 | 0.334 | 9.83E-53 | Lep-L |
| Macro-TREM2 | SCCPDH | 5.89E-45 | 1.254745865 | 0.446 | 0.287 | 1.16E-40 | Lep-L |
| Macro-TREM2 | CCSER1 | 5.59E-45 | 1.253355942 | 0.314 | 0.132 | 1.10E-40 | Lep-L |
| Macro-TREM2 | NBPF19 | 1.19E-16 | 1.252382741 | 0.29 | 0.202 | 2.35E-12 | Lep-L |
| Macro-TREM2 | TOB2 | 2.01E-35 | 1.235458865 | 0.342 | 0.193 | 3.96E-31 | Lep-L |
| Macro-TREM2 | CITED2 | 3.18E-26 | 1.235275387 | 0.6 | 0.514 | 6.26E-22 | Lep-L |
| Macro-TREM2 | CSRNP1 | 5.32E-56 | 1.229457907 | 0.578 | 0.396 | 1.05E-51 | Lep-L |
| Macro-TREM2 | PDXDC1 | 7.14E-43 | 1.229376962 | 0.52 | 0.37 | 1.40E-38 | Lep-L |
| Macro-TREM2 | PTGS2 | 1.54E-21 | 1.228781265 | 0.295 | 0.181 | 3.02E-17 | Lep-L |
| Macro-TREM2 | NFKBID | 1.59E-44 | 1.228742432 | 0.479 | 0.313 | 3.13E-40 | Lep-L |
| Macro-TREM2 | EHD1 | 3.73E-35 | 1.222373044 | 0.472 | 0.336 | 7.34E-31 | Lep-L |
| Macro-TREM2 | PPP1R15A | 3.80E-122 | 1.221797722 | 0.896 | 0.71 | 7.49E-118 | Lep-L |
| Macro-TREM2 | USP38 | 1.18E-12 | 1.219552627 | 0.281 | 0.213 | 2.32E-08 | Lep-L |
| Macro-TREM2 | SERTAD2 | 3.39E-33 | 1.216487279 | 0.442 | 0.305 | 6.66E-29 | Lep-L |
| Macro-TREM2 | MAPRE2 | 5.80E-39 | 1.215367298 | 0.51 | 0.356 | 1.14E-34 | Lep-L |
| Macro-TREM2 | PTGER2 | 5.36E-12 | 1.212219138 | 0.258 | 0.187 | 1.06E-07 | Lep-L |
| Macro-TREM2 | FEM1B | 2.09E-16 | 1.211385093 | 0.344 | 0.262 | 4.11E-12 | Lep-L |
| Macro-TREM2 | CA2 | 7.10E-52 | 1.206765454 | 0.53 | 0.328 | 1.40E-47 | Lep-L |
| Macro-TREM2 | RAB20 | 8.22E-78 | 1.20546694 | 0.743 | 0.581 | 1.62E-73 | Lep-L |
| Macro-TREM2 | BCL6 | 3.46E-31 | 1.204788228 | 0.518 | 0.412 | 6.81E-27 | Lep-L |
| Macro-TREM2 | DNAJA1 | 1.23E-153 | 1.197944762 | 0.872 | 0.774 | 2.43E-149 | Lep-L |
| Macro-TREM2 | IER3 | 1.82E-105 | 1.196698277 | 0.878 | 0.7 | 3.59E-101 | Lep-L |
| Macro-TREM2 | PRKAG2-AS1 | 3.93E-29 | 1.196246244 | 0.251 | 0.125 | 7.74E-25 | Lep-L |
| Macro-TREM2 | IRS2 | 1.16E-31 | 1.191661942 | 0.411 | 0.275 | 2.29E-27 | Lep-L |
| Macro-TREM2 | DHRS3 | 4.57E-41 | 1.189378859 | 0.402 | 0.239 | 8.99E-37 | Lep-L |
| Macro-TREM2 | AGPAT4 | 2.57E-19 | 1.188034852 | 0.353 | 0.251 | 5.06E-15 | Lep-L |
| Macro-TREM2 | CLN6 | 1.11E-19 | 1.187965235 | 0.323 | 0.226 | 2.19E-15 | Lep-L |
| Macro-TREM2 | ARL4A | 2.35E-83 | 1.185015829 | 0.67 | 0.448 | 4.63E-79 | Lep-L |
| Macro-TREM2 | MIDN | 8.95E-61 | 1.185000596 | 0.575 | 0.39 | 1.76E-56 | Lep-L |
| Macro-TREM2 | APOC1 | 4.58E-160 | 1.18419017 | 0.925 | 0.82 | 9.02E-156 | Lep-L |
| Macro-TREM2 | TREM2 | 6.43E-61 | 1.18391478 | 0.655 | 0.461 | 1.27E-56 | Lep-L |
| Macro-TREM2 | AHCTF1 | 2.30E-14 | 1.179692335 | 0.281 | 0.205 | 4.53E-10 | Lep-L |
| Macro-TREM2 | PDE8A | 5.48E-55 | 1.179240859 | 0.588 | 0.386 | 1.08E-50 | Lep-L |
| Macro-TREM2 | TENT5A | 1.87E-30 | 1.179035929 | 0.507 | 0.385 | 3.68E-26 | Lep-L |
| Macro-TREM2 | SMS | 8.60E-103 | 1.173137496 | 0.757 | 0.598 | 1.69E-98 | Lep-L |
| Macro-TREM2 | MIR3142HG | 2.89E-14 | 1.171740415 | 0.352 | 0.256 | 5.69E-10 | Lep-L |
| Macro-TREM2 | HMOX1 | 8.48E-63 | 1.171623387 | 0.774 | 0.664 | 1.67E-58 | Lep-L |
| Macro-TREM2 | ICAM1 | 3.36E-55 | 1.171606597 | 0.749 | 0.608 | 6.62E-51 | Lep-L |
| Macro-TREM2 | CTSF | 5.01E-19 | 1.169038487 | 0.25 | 0.154 | 9.87E-15 | Lep-L |
| Macro-TREM2 | CHORDC1 | 2.45E-59 | 1.166983461 | 0.576 | 0.386 | 4.82E-55 | Lep-L |
| Macro-TREM2 | NR4A2 | 1.72E-79 | 1.163479541 | 0.832 | 0.64 | 3.38E-75 | Lep-L |
| Macro-TREM2 | MYO9B | 6.69E-81 | 1.158283757 | 0.739 | 0.553 | 1.32E-76 | Lep-L |
| Macro-TREM2 | CHKA | 3.63E-37 | 1.156767436 | 0.494 | 0.356 | 7.14E-33 | Lep-L |
| Macro-TREM2 | EEPD1 | 1.73E-22 | 1.153163464 | 0.268 | 0.155 | 3.40E-18 | Lep-L |
| Macro-TREM2 | PER1 | 1.04E-20 | 1.152963196 | 0.372 | 0.267 | 2.04E-16 | Lep-L |
| Macro-TREM2 | METRNL | 9.74E-51 | 1.137209688 | 0.643 | 0.522 | 1.92E-46 | Lep-L |
| Macro-TREM2 | NUP214 | 3.14E-31 | 1.135830981 | 0.424 | 0.302 | 6.19E-27 | Lep-L |
| Macro-TREM2 | REL | 1.03E-103 | 1.133280889 | 0.842 | 0.647 | 2.02E-99 | Lep-L |
| Macro-TREM2 | PLPP3 | 9.97E-54 | 1.133207644 | 0.567 | 0.37 | 1.96E-49 | Lep-L |
| Macro-TREM2 | HOMER3 | 4.41E-44 | 1.133159281 | 0.35 | 0.173 | 8.68E-40 | Lep-L |
| Macro-TREM2 | JMJD6 | 2.06E-56 | 1.13293859 | 0.538 | 0.364 | 4.06E-52 | Lep-L |
| Macro-TREM2 | ZC3HAV1 | 2.14E-37 | 1.126702667 | 0.495 | 0.347 | 4.21E-33 | Lep-L |
| Macro-TREM2 | CD55 | 3.92E-34 | 1.126571808 | 0.61 | 0.486 | 7.71E-30 | Lep-L |

| | | | | | | | |
|---|---|---|---|---|---|---|---|
| Macro-TREM2 | PNPLA8 | 1.95E-55 | 1.126445251 | 0.621 | 0.469 | 3.84E-51 | Lep-L |
| Macro-TREM2 | IL1B | 9.71E-30 | 1.125101096 | 0.506 | 0.363 | 1.91E-25 | Lep-L |
| Macro-TREM2 | MCOLN1 | 1.17E-52 | 1.124050848 | 0.56 | 0.394 | 2.31E-48 | Lep-L |
| Macro-TREM2 | ZC3H12A | 5.50E-57 | 1.123866895 | 0.654 | 0.46 | 1.08E-52 | Lep-L |
| Macro-TREM2 | TNFSF9 | 2.83E-27 | 1.123018398 | 0.391 | 0.25 | 5.56E-23 | Lep-L |
| Macro-TREM2 | AIG1 | 1.08E-50 | 1.116172215 | 0.354 | 0.164 | 2.12E-46 | Lep-L |
| Macro-TREM2 | MAT2A | 7.18E-20 | 1.115009328 | 0.491 | 0.422 | 1.41E-15 | Lep-L |
| Macro-TREM2 | BNIP3L | 5.29E-81 | 1.114279693 | 0.782 | 0.668 | 1.04E-76 | Lep-L |
| Macro-TREM2 | TUBB2A | 1.04E-32 | 1.111213283 | 0.355 | 0.203 | 2.05E-28 | Lep-L |
| Macro-TREM2 | SEC24A | 2.84E-11 | 1.110510866 | 0.273 | 0.209 | 5.59E-07 | Lep-L |
| Macro-TREM2 | SGK1 | 1.80E-86 | 1.107230808 | 0.867 | 0.727 | 3.55E-82 | Lep-L |
| Macro-TREM2 | CDK11B | 7.36E-08 | 1.105953509 | 0.335 | 0.299 | 0.001448175 | Lep-L |
| Macro-TREM2 | RB1CC1 | 7.58E-29 | 1.104919196 | 0.434 | 0.318 | 1.49E-24 | Lep-L |
| Macro-TREM2 | NRP2 | 1.22E-78 | 1.103471058 | 0.78 | 0.584 | 2.39E-74 | Lep-L |
| Macro-TREM2 | VAT1 | 2.00E-51 | 1.102640728 | 0.661 | 0.535 | 3.95E-47 | Lep-L |
| Macro-TREM2 | ARHGAP24 | 2.58E-15 | 1.100827927 | 0.324 | 0.228 | 5.07E-11 | Lep-L |
| Macro-TREM2 | HSPH1 | 3.46E-126 | 1.100369398 | 0.85 | 0.604 | 6.81E-122 | Lep-L |
| Macro-TREM2 | ABCG1 | 2.45E-38 | 1.096898127 | 0.533 | 0.386 | 4.82E-34 | Lep-L |
| Macro-TREM2 | SERPINB9 | 1.13E-45 | 1.096070563 | 0.553 | 0.391 | 2.22E-41 | Lep-L |
| Macro-TREM2 | MS4A7 | 1.12E-105 | 1.093943656 | 0.803 | 0.595 | 2.20E-101 | Lep-L |
| Macro-TREM2 | ARID3A | 7.46E-15 | 1.093433744 | 0.311 | 0.233 | 1.47E-10 | Lep-L |
| Macro-TREM2 | HECA | 8.17E-31 | 1.091209872 | 0.531 | 0.393 | 1.61E-26 | Lep-L |
| Macro-TREM2 | PELI1 | 8.63E-76 | 1.089103919 | 0.575 | 0.334 | 1.70E-71 | Lep-L |
| Macro-TREM2 | SH3BP5 | 1.13E-73 | 1.08788378 | 0.731 | 0.539 | 2.22E-69 | Lep-L |
| Macro-TREM2 | POM121 | 3.52E-11 | 1.086723433 | 0.349 | 0.285 | 6.92E-07 | Lep-L |
| Macro-TREM2 | UBE2S | 6.94E-61 | 1.082464274 | 0.694 | 0.541 | 1.37E-56 | Lep-L |
| Macro-TREM2 | ATP1B3 | 3.56E-74 | 1.082406553 | 0.797 | 0.668 | 7.02E-70 | Lep-L |
| Macro-TREM2 | PDE4DIP | 1.02E-95 | 1.080102699 | 0.8 | 0.612 | 2.01E-91 | Lep-L |
| Macro-TREM2 | FOSL2 | 1.34E-53 | 1.079607987 | 0.577 | 0.408 | 2.64E-49 | Lep-L |
| Macro-TREM2 | NOTCH2 | 1.94E-47 | 1.078001326 | 0.569 | 0.41 | 3.82E-43 | Lep-L |
| Macro-TREM2 | CPEB2 | 1.92E-12 | 1.076744745 | 0.33 | 0.256 | 3.78E-08 | Lep-L |
| Macro-TREM2 | CHD2 | 1.07E-63 | 1.07227139 | 0.697 | 0.533 | 2.11E-59 | Lep-L |
| Macro-TREM2 | SNCA | 6.29E-37 | 1.071426144 | 0.306 | 0.144 | 1.24E-32 | Lep-L |
| Macro-TREM2 | JDP2 | 1.31E-32 | 1.068963532 | 0.438 | 0.303 | 2.57E-28 | Lep-L |
| Macro-TREM2 | ARL5B | 1.20E-46 | 1.064539573 | 0.502 | 0.308 | 2.36E-42 | Lep-L |
| Macro-TREM2 | ZFP36L1 | 2.66E-51 | 1.063198471 | 0.773 | 0.648 | 5.23E-47 | Lep-L |
| Macro-TREM2 | MAFG | 1.07E-17 | 1.063107233 | 0.354 | 0.274 | 2.11E-13 | Lep-L |
| Macro-TREM2 | NR1D2 | 3.54E-09 | 1.059812358 | 0.254 | 0.198 | 6.97E-05 | Lep-L |
| Macro-TREM2 | ALG13 | 8.30E-17 | 1.058209255 | 0.452 | 0.392 | 1.63E-12 | Lep-L |
| Macro-TREM2 | DNAJB1 | 2.07E-96 | 1.056752253 | 0.912 | 0.731 | 4.07E-92 | Lep-L |
| Macro-TREM2 | PIK3IP1 | 1.83E-19 | 1.056581263 | 0.399 | 0.307 | 3.60E-15 | Lep-L |
| Macro-TREM2 | MYLIP | 7.01E-18 | 1.054615096 | 0.426 | 0.336 | 1.38E-13 | Lep-L |
| Macro-TREM2 | ABCA1 | 9.40E-82 | 1.047931681 | 0.771 | 0.576 | 1.85E-77 | Lep-L |
| Macro-TREM2 | MCL1 | 1.11E-100 | 1.047422075 | 0.82 | 0.657 | 2.18E-96 | Lep-L |
| Macro-TREM2 | KLF3 | 6.10E-32 | 1.046848644 | 0.391 | 0.257 | 1.20E-27 | Lep-L |
| Macro-TREM2 | SAMSN1 | 2.34E-38 | 1.041675448 | 0.597 | 0.458 | 4.60E-34 | Lep-L |
| Macro-TREM2 | SNAPC1 | 8.34E-29 | 1.039336657 | 0.443 | 0.31 | 1.64E-24 | Lep-L |
| Macro-TREM2 | NR4A1 | 1.29E-40 | 1.039266026 | 0.706 | 0.552 | 2.54E-36 | Lep-L |
| Macro-TREM2 | AREG | 2.50E-15 | 1.039149 | 0.368 | 0.266 | 4.92E-11 | Lep-L |
| Macro-TREM2 | CLN8 | 4.66E-23 | 1.039056787 | 0.425 | 0.326 | 9.18E-19 | Lep-L |
| Macro-TREM2 | NOTCH2NLC | 3.33E-13 | 1.033039225 | 0.323 | 0.247 | 6.56E-09 | Lep-L |
| Macro-TREM2 | SAT1 | 1.02E-204 | 1.029045915 | 0.988 | 0.955 | 2.00E-200 | Lep-L |
| Macro-TREM2 | TCN2 | 4.05E-38 | 1.028016734 | 0.612 | 0.513 | 7.97E-34 | Lep-L |
| Macro-TREM2 | BANP | 9.99E-25 | 1.027477773 | 0.439 | 0.327 | 1.97E-20 | Lep-L |
| Macro-TREM2 | STK40 | 9.87E-37 | 1.027434926 | 0.432 | 0.287 | 1.94E-32 | Lep-L |

| | | | | | | | |
|---|---|---|---|---|---|---|---|
| Macro-TREM2 | ID2 | 1.23E-62 | 1.027041802 | 0.678 | 0.488 | 2.43E-58 | Lep-L |
| Macro-TREM2 | ADD3 | 4.31E-17 | 1.025935593 | 0.282 | 0.186 | 8.48E-13 | Lep-L |
| Macro-TREM2 | TRIB1 | 2.14E-23 | 1.024735828 | 0.421 | 0.314 | 4.21E-19 | Lep-L |
| Macro-TREM2 | IFRD1 | 1.02E-34 | 1.023254428 | 0.618 | 0.498 | 2.01E-30 | Lep-L |
| Macro-TREM2 | DHX15 | 2.03E-07 | 1.020410861 | 0.319 | 0.285 | 0.003992362 | Lep-L |
| Macro-TREM2 | TUBB4B | 1.00E-72 | 1.016278691 | 0.775 | 0.632 | 1.98E-68 | Lep-L |
| Macro-TREM2 | KLF6 | 9.73E-86 | 1.016182621 | 0.908 | 0.756 | 1.92E-81 | Lep-L |
| Macro-TREM2 | PABPC4 | 1.05E-83 | 1.015725528 | 0.739 | 0.576 | 2.06E-79 | Lep-L |
| Macro-TREM2 | STARD3NL | 9.69E-23 | 1.012374777 | 0.503 | 0.415 | 1.91E-18 | Lep-L |
| Macro-TREM2 | SESN1 | 1.17E-11 | 1.010501863 | 0.309 | 0.235 | 2.30E-07 | Lep-L |
| Macro-TREM2 | STX3 | 4.99E-13 | 1.007208414 | 0.348 | 0.288 | 9.82E-09 | Lep-L |
| Macro-TREM2 | RAB42 | 4.35E-23 | 1.004783479 | 0.386 | 0.271 | 8.57E-19 | Lep-L |
| Macro-TREM2 | DNAJB4 | 1.91E-36 | 1.004010342 | 0.367 | 0.209 | 3.77E-32 | Lep-L |
| Macro-TREM2 | CDR2 | 4.84E-15 | 1.002817699 | 0.313 | 0.231 | 9.52E-11 | Lep-L |
| Macro-TREM2 | LIPA | 1.17E-82 | 1.001783894 | 0.813 | 0.652 | 2.30E-78 | Lep-L |
| Macro-TREM2 | RHOB | 2.13E-19 | 1.000687341 | 0.57 | 0.478 | 4.20E-15 | Lep-L |
| Macro-TREM2 | UBD | 1.15E-68 | 3.059422916 | 0.539 | 0.255 | 2.27E-64 | Lep-T |
| Macro-TREM2 | PLA2G2D | 4.02E-31 | 2.116121154 | 0.446 | 0.272 | 7.92E-27 | Lep-T |
| Macro-TREM2 | PTGDS | 5.85E-59 | 2.082724692 | 0.618 | 0.364 | 1.15E-54 | Lep-T |
| Macro-TREM2 | IGKC | 5.82E-21 | 1.911639038 | 0.126 | 0.292 | 1.15E-16 | Lep-T |
| Macro-TREM2 | LGALS2 | 1.70E-100 | 1.846339036 | 0.757 | 0.51 | 3.35E-96 | Lep-T |
| Macro-TREM2 | APOL3 | 3.43E-21 | 1.778572824 | 0.363 | 0.216 | 6.75E-17 | Lep-T |
| Macro-TREM2 | TSPAN33 | 1.40E-41 | 1.767966375 | 0.514 | 0.308 | 2.76E-37 | Lep-T |
| Macro-TREM2 | TNFRSF4 | 1.14E-42 | 1.736278541 | 0.342 | 0.132 | 2.24E-38 | Lep-T |
| Macro-TREM2 | HSD11B1 | 4.41E-34 | 1.702159384 | 0.412 | 0.218 | 8.69E-30 | Lep-T |
| Macro-TREM2 | CST3 | 3.14E-47 | 1.682929278 | 0.972 | 0.967 | 6.18E-43 | Lep-T |
| Macro-TREM2 | GSTM4 | 4.28E-16 | 1.631303029 | 0.339 | 0.215 | 8.42E-12 | Lep-T |
| Macro-TREM2 | PRKAR2B | 3.99E-39 | 1.611387529 | 0.35 | 0.151 | 7.85E-35 | Lep-T |
| Macro-TREM2 | RTN1 | 2.28E-15 | 1.554744866 | 0.295 | 0.17 | 4.49E-11 | Lep-T |
| Macro-TREM2 | TIFAB | 5.17E-15 | 1.49017995 | 0.313 | 0.189 | 1.02E-10 | Lep-T |
| Macro-TREM2 | CHI3L1 | 3.85E-34 | 1.46046358 | 0.627 | 0.513 | 7.59E-30 | Lep-T |
| Macro-TREM2 | WARS | 1.23E-42 | 1.430512848 | 0.677 | 0.598 | 2.42E-38 | Lep-T |
| Macro-TREM2 | NUDT21 | 6.78E-14 | 1.419342616 | 0.317 | 0.194 | 1.34E-09 | Lep-T |
| Macro-TREM2 | RFTN1 | 1.24E-12 | 1.412479496 | 0.401 | 0.313 | 2.43E-08 | Lep-T |
| Macro-TREM2 | ALDOA | 8.88E-07 | 1.405609531 | 0.504 | 0.515 | 0.017473847 | Lep-T |
| Macro-TREM2 | AFDN | 1.92E-19 | 1.383040526 | 0.304 | 0.161 | 3.77E-15 | Lep-T |
| Macro-TREM2 | LIMD2 | 5.10E-18 | 1.367631731 | 0.496 | 0.369 | 1.00E-13 | Lep-T |
| Macro-TREM2 | CDA | 4.96E-20 | 1.325928731 | 0.39 | 0.244 | 9.77E-16 | Lep-T |
| Macro-TREM2 | CXCL10 | 6.96E-15 | 1.3086855 | 0.308 | 0.181 | 1.37E-10 | Lep-T |
| Macro-TREM2 | ACOT7 | 6.99E-25 | 1.301023012 | 0.441 | 0.285 | 1.38E-20 | Lep-T |
| Macro-TREM2 | GBP5 | 2.76E-29 | 1.298849316 | 0.604 | 0.459 | 5.43E-25 | Lep-T |
| Macro-TREM2 | CXCL9 | 3.74E-24 | 1.288452423 | 0.397 | 0.214 | 7.36E-20 | Lep-T |
| Macro-TREM2 | POLR2G | 3.65E-14 | 1.275917089 | 0.407 | 0.285 | 7.18E-10 | Lep-T |
| Macro-TREM2 | HLA-DQB2 | 4.41E-17 | 1.271978477 | 0.447 | 0.328 | 8.68E-13 | Lep-T |
| Macro-TREM2 | SLAMF7 | 4.38E-13 | 1.235000584 | 0.53 | 0.519 | 8.62E-09 | Lep-T |
| Macro-TREM2 | COL6A2 | 3.61E-11 | 1.232222003 | 0.281 | 0.181 | 7.11E-07 | Lep-T |
| Macro-TREM2 | CD40 | 1.33E-15 | 1.223000143 | 0.527 | 0.477 | 2.62E-11 | Lep-T |
| Macro-TREM2 | IGSF6 | 1.51E-32 | 1.209537959 | 0.759 | 0.81 | 2.98E-28 | Lep-T |
| Macro-TREM2 | CD74 | 4.70E-90 | 1.207356441 | 0.99 | 0.993 | 9.26E-86 | Lep-T |
| Macro-TREM2 | TLR8 | 2.47E-12 | 1.201660156 | 0.301 | 0.192 | 4.86E-08 | Lep-T |
| Macro-TREM2 | ATP5PB | 4.90E-20 | 1.198655949 | 0.551 | 0.474 | 9.65E-16 | Lep-T |
| Macro-TREM2 | CRIP1 | 1.88E-25 | 1.15656087 | 0.618 | 0.45 | 3.70E-21 | Lep-T |
| Macro-TREM2 | SPN | 1.65E-08 | 1.133925436 | 0.292 | 0.206 | 0.000325667 | Lep-T |
| Macro-TREM2 | SMIM29 | 4.73E-09 | 1.123575133 | 0.284 | 0.191 | 9.32E-05 | Lep-T |
| Macro-TREM2 | STAT1 | 6.86E-36 | 1.105926996 | 0.73 | 0.659 | 1.35E-31 | Lep-T |

| | | | | | | | |
|---|---|---|---|---|---|---|---|
| Macro-TREM2 | TNFSF10 | 8.53E-12 | 1.105172176 | 0.468 | 0.371 | 1.68E-07 | Lep-T |
| Macro-TREM2 | TNFAIP8L2 | 6.80E-12 | 1.104824518 | 0.27 | 0.163 | 1.34E-07 | Lep-T |
| Macro-TREM2 | SCNM1 | 9.71E-07 | 1.10396795 | 0.31 | 0.235 | 0.019122354 | Lep-T |
| Macro-TREM2 | GBP4 | 6.05E-10 | 1.09606608 | 0.422 | 0.338 | 1.19E-05 | Lep-T |
| Macro-TREM2 | GBP1 | 8.43E-31 | 1.084141576 | 0.612 | 0.427 | 1.66E-26 | Lep-T |
| Macro-TREM2 | CLEC12A | 2.75E-09 | 1.079219161 | 0.393 | 0.305 | 5.41E-05 | Lep-T |
| Macro-TREM2 | IL2RG | 2.92E-10 | 1.0567081 | 0.465 | 0.377 | 5.75E-06 | Lep-T |
| Macro-TREM2 | HLA-DMB | 1.21E-49 | 1.034842567 | 0.756 | 0.788 | 2.39E-45 | Lep-T |
| Macro-TREM2 | P2RY13 | 2.23E-07 | 1.029670889 | 0.292 | 0.209 | 0.004398552 | Lep-T |
| Macro-TREM2 | SNAP23 | 5.21E-10 | 1.019640322 | 0.539 | 0.506 | 1.02E-05 | Lep-T |
| Macro-TREM2 | PSME1 | 3.82E-21 | 1.018310888 | 0.726 | 0.704 | 7.53E-17 | Lep-T |
| Macro-TREM2 | NR1H3 | 1.15E-10 | 1.014275257 | 0.539 | 0.548 | 2.27E-06 | Lep-T |
| Macro-TREM2 | FOLR3 | 1.38E-165 | 4.976732596 | 0.325 | 0.019 | 2.72E-161 | Mari |
| Macro-TREM2 | MMP121 | 1.44E-68 | 3.139826908 | 0.408 | 0.114 | 2.84E-64 | Mari |
| Macro-TREM2 | S100A81 | 1.04E-190 | 2.96080747 | 0.9 | 0.351 | 2.05E-186 | Mari |
| Macro-TREM2 | SPP1 | 6.94E-18 | 2.389519346 | 0.389 | 0.228 | 1.37E-13 | Mari |
| Macro-TREM2 | IL7R1 | 7.46E-98 | 2.264658715 | 0.541 | 0.142 | 1.47E-93 | Mari |
| Macro-TREM2 | MGST1 | 8.26E-63 | 2.252060486 | 0.45 | 0.148 | 1.63E-58 | Mari |
| Macro-TREM2 | IDO11 | 5.57E-113 | 2.147246421 | 0.552 | 0.131 | 1.10E-108 | Mari |
| Macro-TREM2 | INHBA | 3.54E-51 | 2.124242486 | 0.505 | 0.213 | 6.97E-47 | Mari |
| Macro-TREM2 | CCL8 | 7.18E-39 | 2.096360557 | 0.31 | 0.1 | 1.41E-34 | Mari |
| Macro-TREM2 | IL6 | 1.41E-52 | 1.97987157 | 0.376 | 0.114 | 2.77E-48 | Mari |
| Macro-TREM2 | IL2RA | 8.07E-56 | 1.892299737 | 0.352 | 0.095 | 1.59E-51 | Mari |
| Macro-TREM2 | S100A91 | 6.40E-122 | 1.884000733 | 0.938 | 0.711 | 1.26E-117 | Mari |
| Macro-TREM2 | FCGR3A1 | 2.52E-100 | 1.827547034 | 0.841 | 0.439 | 4.96E-96 | Mari |
| Macro-TREM2 | CCND1 | 1.15E-35 | 1.827200924 | 0.295 | 0.097 | 2.26E-31 | Mari |
| Macro-TREM2 | PLTP | 2.24E-22 | 1.81572931 | 0.739 | 0.555 | 4.41E-18 | Mari |
| Macro-TREM2 | CALHM62 | 1.79E-77 | 1.730709417 | 0.843 | 0.525 | 3.52E-73 | Mari |
| Macro-TREM2 | ANKRD221 | 5.29E-99 | 1.714600763 | 0.493 | 0.112 | 1.04E-94 | Mari |
| Macro-TREM2 | OLFML2B | 7.75E-38 | 1.711283085 | 0.338 | 0.115 | 1.53E-33 | Mari |
| Macro-TREM2 | SCN1B | 8.45E-11 | 1.697459532 | 0.301 | 0.173 | 1.66E-06 | Mari |
| Macro-TREM2 | CLEC11A | 5.77E-27 | 1.633757562 | 0.257 | 0.091 | 1.14E-22 | Mari |
| Macro-TREM2 | RNASE1 | 1.12E-10 | 1.63065481 | 0.41 | 0.262 | 2.21E-06 | Mari |
| Macro-TREM2 | VAMP52 | 3.40E-89 | 1.575763159 | 0.851 | 0.503 | 6.69E-85 | Mari |
| Macro-TREM2 | SLC7A11 | 1.95E-32 | 1.574590285 | 0.348 | 0.139 | 3.83E-28 | Mari |
| Macro-TREM2 | TYMP1 | 4.10E-139 | 1.551846342 | 0.966 | 0.811 | 8.06E-135 | Mari |
| Macro-TREM2 | ETV7 | 3.73E-61 | 1.535619183 | 0.323 | 0.074 | 7.34E-57 | Mari |
| Macro-TREM2 | MYO1G | 1.41E-47 | 1.529957389 | 0.399 | 0.128 | 2.78E-43 | Mari |
| Macro-TREM2 | MSC | 1.39E-35 | 1.519144066 | 0.331 | 0.119 | 2.74E-31 | Mari |
| Macro-TREM2 | CAMK1 | 5.44E-37 | 1.518369557 | 0.321 | 0.107 | 1.07E-32 | Mari |
| Macro-TREM2 | GBP12 | 7.39E-73 | 1.47584904 | 0.781 | 0.415 | 1.45E-68 | Mari |
| Macro-TREM2 | MRPL58 | 3.45E-24 | 1.462165174 | 0.386 | 0.176 | 6.80E-20 | Mari |
| Macro-TREM2 | VMO1 | 1.12E-24 | 1.453419199 | 0.433 | 0.219 | 2.21E-20 | Mari |
| Macro-TREM2 | ALG3 | 3.53E-20 | 1.447578906 | 0.344 | 0.162 | 6.95E-16 | Mari |
| Macro-TREM2 | PRDX21 | 2.39E-30 | 1.425844361 | 0.397 | 0.171 | 4.70E-26 | Mari |
| Macro-TREM2 | HAPLN31 | 1.19E-47 | 1.425717833 | 0.395 | 0.128 | 2.34E-43 | Mari |
| Macro-TREM2 | PPA11 | 1.03E-63 | 1.413904042 | 0.783 | 0.46 | 2.03E-59 | Mari |
| Macro-TREM2 | FCGR1B | 5.19E-30 | 1.382454883 | 0.456 | 0.21 | 1.02E-25 | Mari |
| Macro-TREM2 | STAC3 | 3.39E-29 | 1.368695771 | 0.486 | 0.243 | 6.67E-25 | Mari |
| Macro-TREM2 | FCGR1A1 | 3.55E-60 | 1.35840694 | 0.781 | 0.512 | 6.99E-56 | Mari |
| Macro-TREM2 | NKG7 | 2.71E-32 | 1.352641296 | 0.406 | 0.175 | 5.34E-28 | Mari |
| Macro-TREM2 | SLC39A81 | 1.50E-43 | 1.345986352 | 0.556 | 0.274 | 2.96E-39 | Mari |
| Macro-TREM2 | BST2 | 2.24E-80 | 1.3182372 | 0.883 | 0.689 | 4.41E-76 | Mari |
| Macro-TREM2 | MSMO1 | 1.40E-29 | 1.317725782 | 0.301 | 0.113 | 2.76E-25 | Mari |
| Macro-TREM2 | TMEM173 | 1.43E-14 | 1.312825612 | 0.261 | 0.126 | 2.82E-10 | Mari |

| | | | | | | | |
|---|---|---|---|---|---|---|---|
| Macro-TREM2 | CCL18 | 1.34E-13 | 1.312355897 | 0.724 | 0.631 | 2.64E-09 | Mari |
| Macro-TREM2 | IFIT31 | 2.10E-56 | 1.31123512 | 0.527 | 0.197 | 4.13E-52 | Mari |
| Macro-TREM2 | SLC11A11 | 3.75E-56 | 1.301099274 | 0.569 | 0.23 | 7.38E-52 | Mari |
| Macro-TREM2 | ROGDI | 9.87E-23 | 1.296252868 | 0.365 | 0.165 | 1.94E-18 | Mari |
| Macro-TREM2 | GAS6 | 6.42E-12 | 1.287588857 | 0.367 | 0.218 | 1.26E-07 | Mari |
| Macro-TREM2 | AIFM21 | 6.72E-39 | 1.285833524 | 0.423 | 0.169 | 1.32E-34 | Mari |
| Macro-TREM2 | PDSS11 | 2.23E-29 | 1.2815967 | 0.35 | 0.141 | 4.40E-25 | Mari |
| Macro-TREM2 | RRAS1 | 2.30E-55 | 1.280395237 | 0.705 | 0.378 | 4.52E-51 | Mari |
| Macro-TREM2 | HK31 | 8.67E-52 | 1.269660543 | 0.461 | 0.16 | 1.71E-47 | Mari |
| Macro-TREM2 | CLEC4D | 1.85E-33 | 1.266485134 | 0.293 | 0.097 | 3.65E-29 | Mari |
| Macro-TREM2 | IL4I11 | 1.74E-40 | 1.254363144 | 0.707 | 0.456 | 3.42E-36 | Mari |
| Macro-TREM2 | TREM11 | 1.16E-29 | 1.248367009 | 0.384 | 0.163 | 2.28E-25 | Mari |
| Macro-TREM2 | XRCC4 | 3.49E-26 | 1.228865322 | 0.355 | 0.15 | 6.86E-22 | Mari |
| Macro-TREM2 | CORO1A | 2.42E-12 | 1.226715391 | 0.435 | 0.264 | 4.77E-08 | Mari |
| Macro-TREM2 | NME4 | 4.43E-07 | 1.225959658 | 0.399 | 0.273 | 0.00871286 | Mari |
| Macro-TREM2 | HRH2 | 3.44E-52 | 1.218557515 | 0.41 | 0.129 | 6.76E-48 | Mari |
| Macro-TREM2 | MRPL23 | 6.07E-41 | 1.21531715 | 0.565 | 0.254 | 1.19E-36 | Mari |
| Macro-TREM2 | CFB1 | 4.11E-29 | 1.213865929 | 0.318 | 0.122 | 8.09E-25 | Mari |
| Macro-TREM2 | CASP41 | 2.71E-46 | 1.206112609 | 0.754 | 0.471 | 5.34E-42 | Mari |
| Macro-TREM2 | IFI35 | 6.82E-40 | 1.204612914 | 0.554 | 0.254 | 1.34E-35 | Mari |
| Macro-TREM2 | OSM1 | 9.38E-24 | 1.204149361 | 0.459 | 0.246 | 1.85E-19 | Mari |
| Macro-TREM2 | SIGLEC101 | 4.25E-39 | 1.196000881 | 0.463 | 0.189 | 8.37E-35 | Mari |
| Macro-TREM2 | PSMB92 | 9.96E-72 | 1.192694889 | 0.873 | 0.663 | 1.96E-67 | Mari |
| Macro-TREM2 | INHBA-AS1 | 1.69E-21 | 1.190436617 | 0.28 | 0.116 | 3.32E-17 | Mari |
| Macro-TREM2 | MVB12A | 5.46E-29 | 1.181617762 | 0.51 | 0.254 | 1.07E-24 | Mari |
| Macro-TREM2 | TMEM1631 | 8.48E-32 | 1.181331833 | 0.348 | 0.137 | 1.67E-27 | Mari |
| Macro-TREM2 | RAB341 | 1.15E-47 | 1.176043506 | 0.65 | 0.321 | 2.26E-43 | Mari |
| Macro-TREM2 | ADAM81 | 1.16E-36 | 1.175556621 | 0.45 | 0.188 | 2.28E-32 | Mari |
| Macro-TREM2 | SPRED11 | 5.36E-38 | 1.165310897 | 0.501 | 0.222 | 1.05E-33 | Mari |
| Macro-TREM2 | CD141 | 6.34E-43 | 1.160593707 | 0.877 | 0.779 | 1.25E-38 | Mari |
| Macro-TREM2 | FBXO6 | 1.41E-47 | 1.157040549 | 0.652 | 0.34 | 2.78E-43 | Mari |
| Macro-TREM2 | CP1 | 3.00E-47 | 1.152038871 | 0.416 | 0.14 | 5.90E-43 | Mari |
| Macro-TREM2 | PLAAT42 | 4.27E-52 | 1.150459179 | 0.764 | 0.453 | 8.41E-48 | Mari |
| Macro-TREM2 | ATF7IP2 | 9.28E-10 | 1.14922911 | 0.274 | 0.162 | 1.83E-05 | Mari |
| Macro-TREM2 | SUCNR11 | 4.31E-32 | 1.142643963 | 0.414 | 0.176 | 8.48E-28 | Mari |
| Macro-TREM2 | APOL41 | 1.43E-35 | 1.139422162 | 0.38 | 0.146 | 2.81E-31 | Mari |
| Macro-TREM2 | PDCD1LG21 | 1.18E-37 | 1.135914698 | 0.476 | 0.206 | 2.33E-33 | Mari |
| Macro-TREM2 | EDEM2 | 9.15E-31 | 1.12981985 | 0.558 | 0.286 | 1.80E-26 | Mari |
| Macro-TREM2 | MTF1 | 1.41E-31 | 1.126275226 | 0.331 | 0.126 | 2.78E-27 | Mari |
| Macro-TREM2 | GPR84 | 3.10E-28 | 1.120397551 | 0.403 | 0.18 | 6.09E-24 | Mari |
| Macro-TREM2 | DTYMK | 8.33E-22 | 1.118001364 | 0.359 | 0.163 | 1.64E-17 | Mari |
| Macro-TREM2 | TUBA1A | 5.20E-39 | 1.110686903 | 0.837 | 0.609 | 1.02E-34 | Mari |
| Macro-TREM2 | ARMT1 | 1.46E-16 | 1.104410814 | 0.289 | 0.134 | 2.88E-12 | Mari |
| Macro-TREM2 | TMEM150A | 9.26E-15 | 1.102125936 | 0.259 | 0.122 | 1.82E-10 | Mari |
| Macro-TREM2 | GIMAP71 | 5.93E-46 | 1.101406156 | 0.399 | 0.134 | 1.17E-41 | Mari |
| Macro-TREM2 | FAM89B1 | 8.52E-30 | 1.100011296 | 0.648 | 0.366 | 1.68E-25 | Mari |
| Macro-TREM2 | CD471 | 2.37E-49 | 1.098546361 | 0.775 | 0.552 | 4.67E-45 | Mari |
| Macro-TREM2 | SDF2L11 | 1.33E-29 | 1.093407746 | 0.656 | 0.389 | 2.62E-25 | Mari |
| Macro-TREM2 | IFI30 | 2.52E-94 | 1.09205309 | 0.972 | 0.939 | 4.95E-90 | Mari |
| Macro-TREM2 | HSD3B7 | 1.11E-22 | 1.087632045 | 0.446 | 0.23 | 2.18E-18 | Mari |
| Macro-TREM2 | PSTPIP11 | 4.62E-45 | 1.087146823 | 0.469 | 0.175 | 9.10E-41 | Mari |
| Macro-TREM2 | GNG101 | 4.13E-48 | 1.083199733 | 0.875 | 0.659 | 8.13E-44 | Mari |
| Macro-TREM2 | C1QB1 | 9.82E-58 | 1.08234731 | 0.921 | 0.806 | 1.93E-53 | Mari |
| Macro-TREM2 | DDX49 | 2.46E-12 | 1.080765967 | 0.301 | 0.161 | 4.85E-08 | Mari |
| Macro-TREM2 | TMEM38B | 3.12E-35 | 1.079021301 | 0.58 | 0.308 | 6.15E-31 | Mari |

| | | | | | | | |
|---|---|---|---|---|---|---|---|
| Macro-TREM2 | TSFM1 | 1.21E-41 | 1.076951127 | 0.51 | 0.218 | 2.38E-37 | Mari |
| Macro-TREM2 | DYNC1I2 | 1.58E-34 | 1.076447481 | 0.62 | 0.335 | 3.11E-30 | Mari |
| Macro-TREM2 | PSME22 | 1.04E-71 | 1.074895906 | 0.915 | 0.739 | 2.05E-67 | Mari |
| Macro-TREM2 | TSTA3 | 1.08E-19 | 1.074067191 | 0.382 | 0.187 | 2.12E-15 | Mari |
| Macro-TREM2 | S100A61 | 5.31E-88 | 1.068580175 | 0.979 | 0.916 | 1.05E-83 | Mari |
| Macro-TREM2 | TMEM109 | 1.68E-25 | 1.068211917 | 0.558 | 0.312 | 3.30E-21 | Mari |
| Macro-TREM2 | SMCO41 | 6.57E-43 | 1.065447223 | 0.758 | 0.478 | 1.29E-38 | Mari |
| Macro-TREM2 | FDPS | 2.67E-26 | 1.065053382 | 0.573 | 0.325 | 5.25E-22 | Mari |
| Macro-TREM2 | SLC7A81 | 1.72E-19 | 1.063506745 | 0.276 | 0.123 | 3.38E-15 | Mari |
| Macro-TREM2 | MT1F1 | 6.36E-23 | 1.059950537 | 0.333 | 0.146 | 1.25E-18 | Mari |
| Macro-TREM2 | TMCO1 | 1.65E-39 | 1.057607123 | 0.711 | 0.436 | 3.25E-35 | Mari |
| Macro-TREM2 | JOSD21 | 9.53E-49 | 1.050212536 | 0.643 | 0.315 | 1.88E-44 | Mari |
| Macro-TREM2 | CLDN231 | 1.85E-21 | 1.050109612 | 0.268 | 0.111 | 3.65E-17 | Mari |
| Macro-TREM2 | LAP31 | 4.16E-66 | 1.045793251 | 0.887 | 0.722 | 8.18E-62 | Mari |
| Macro-TREM2 | S100A10 | 2.37E-75 | 1.045070386 | 0.955 | 0.897 | 4.67E-71 | Mari |
| Macro-TREM2 | MT1E1 | 2.94E-20 | 1.044159472 | 0.321 | 0.151 | 5.78E-16 | Mari |
| Macro-TREM2 | MRPL28 | 7.25E-29 | 1.040517596 | 0.552 | 0.29 | 1.43E-24 | Mari |
| Macro-TREM2 | IL1RN | 2.21E-31 | 1.036183378 | 0.563 | 0.307 | 4.35E-27 | Mari |
| Macro-TREM2 | ZDHHC121 | 8.43E-37 | 1.036115338 | 0.586 | 0.295 | 1.66E-32 | Mari |
| Macro-TREM2 | CYBC1 | 3.77E-35 | 1.032381409 | 0.694 | 0.435 | 7.41E-31 | Mari |
| Macro-TREM2 | GIPC1 | 1.47E-21 | 1.028960741 | 0.433 | 0.226 | 2.90E-17 | Mari |
| Macro-TREM2 | NOP10 | 1.50E-63 | 1.02861038 | 0.854 | 0.705 | 2.96E-59 | Mari |
| Macro-TREM2 | CCDC167 | 8.09E-24 | 1.024039194 | 0.342 | 0.149 | 1.59E-19 | Mari |
| Macro-TREM2 | CAVIN31 | 1.58E-23 | 1.024023212 | 0.282 | 0.114 | 3.11E-19 | Mari |
| Macro-TREM2 | PMP22 | 1.24E-19 | 1.023270046 | 0.524 | 0.309 | 2.44E-15 | Mari |
| Macro-TREM2 | FIBP | 1.05E-23 | 1.022369533 | 0.631 | 0.376 | 2.06E-19 | Mari |
| Macro-TREM2 | NQO1 | 3.95E-18 | 1.021745677 | 0.28 | 0.127 | 7.77E-14 | Mari |
| Macro-TREM2 | GIMAP11 | 1.25E-45 | 1.019816056 | 0.537 | 0.225 | 2.46E-41 | Mari |
| Macro-TREM2 | AGTRAP1 | 8.88E-42 | 1.018955828 | 0.76 | 0.532 | 1.75E-37 | Mari |
| Macro-TREM2 | IFITM31 | 3.93E-17 | 1.016793168 | 0.858 | 0.659 | 7.74E-13 | Mari |
| Macro-TREM2 | IFI44L | 1.05E-22 | 1.016334465 | 0.255 | 0.095 | 2.07E-18 | Mari |
| Macro-TREM2 | HLA-DQA21 | 5.52E-13 | 1.014086156 | 0.728 | 0.662 | 1.09E-08 | Mari |
| Macro-TREM2 | CSF2RB1 | 4.27E-27 | 1.011651858 | 0.456 | 0.218 | 8.41E-23 | Mari |
| Macro-TREM2 | ENPP4 | 1.45E-29 | 1.010464236 | 0.257 | 0.084 | 2.84E-25 | Mari |
| Macro-TREM2 | CNIH41 | 3.93E-41 | 1.009913678 | 0.69 | 0.409 | 7.73E-37 | Mari |
| Macro-TREM2 | NUBP21 | 2.88E-15 | 1.002099815 | 0.429 | 0.238 | 5.68E-11 | Mari |
| Macro-TREM2 | FN1 | 3.62E-104 | 3.66811995 | 0.409 | 0.101 | 7.13E-100 | Tb |
| Macro-TREM2 | SLC35F1 | 3.35E-85 | 3.296085951 | 0.341 | 0.085 | 6.60E-81 | Tb |
| Macro-TREM2 | CAMP | 3.96E-73 | 2.996546017 | 0.408 | 0.136 | 7.80E-69 | Tb |
| Macro-TREM2 | PPIC | 1.57E-121 | 2.746400965 | 0.375 | 0.064 | 3.09E-117 | Tb |
| Macro-TREM2 | PDLIM4 | 3.28E-140 | 2.640842753 | 0.485 | 0.104 | 6.46E-136 | Tb |
| Macro-TREM2 | NRN1 | 6.59E-133 | 2.585155708 | 0.429 | 0.078 | 1.30E-128 | Tb |
| Macro-TREM2 | PHLDA3 | 1.95E-92 | 2.511162523 | 0.426 | 0.118 | 3.84E-88 | Tb |
| Macro-TREM2 | LRRC61 | 3.41E-52 | 2.27890369 | 0.283 | 0.081 | 6.70E-48 | Tb |
| Macro-TREM2 | FBP1 | 3.67E-149 | 2.264037498 | 0.87 | 0.6 | 7.23E-145 | Tb |
| Macro-TREM2 | FGD5 | 7.92E-81 | 2.259822571 | 0.264 | 0.045 | 1.56E-76 | Tb |
| Macro-TREM2 | CP | 2.19E-115 | 2.063454892 | 0.462 | 0.108 | 4.32E-111 | Tb |
| Macro-TREM2 | FXYD2 | 8.04E-74 | 2.058867931 | 0.511 | 0.207 | 1.58E-69 | Tb |
| Macro-TREM2 | QPCT | 3.92E-77 | 1.971597934 | 0.521 | 0.199 | 7.72E-73 | Tb |
| Macro-TREM2 | HRK | 3.75E-78 | 1.966653992 | 0.321 | 0.072 | 7.38E-74 | Tb |
| Macro-TREM2 | CTSK1 | 1.67E-55 | 1.957430047 | 0.574 | 0.31 | 3.30E-51 | Tb |
| Macro-TREM2 | MT1X | 2.39E-24 | 1.904814121 | 0.463 | 0.28 | 4.71E-20 | Tb |
| Macro-TREM2 | CYP27B1 | 1.64E-47 | 1.852414567 | 0.316 | 0.107 | 3.24E-43 | Tb |
| Macro-TREM2 | KRT14 | 6.17E-63 | 1.830776752 | 0.376 | 0.117 | 1.21E-58 | Tb |
| Macro-TREM2 | CISH | 4.55E-37 | 1.829390838 | 0.265 | 0.092 | 8.96E-33 | Tb |

| | | | | | | | |
|---|---|---|---|---|---|---|---|
| Macro-TREM2 | LYZ | 3.23E-187 | 1.817154976 | 0.986 | 0.907 | 6.35E-183 | Tb |
| Macro-TREM2 | HOMER1 | 6.14E-43 | 1.809025349 | 0.36 | 0.149 | 1.21E-38 | Tb |
| Macro-TREM2 | EGLN3 | 8.48E-71 | 1.77920792 | 0.489 | 0.188 | 1.67E-66 | Tb |
| Macro-TREM2 | TMEM163 | 3.44E-52 | 1.772377294 | 0.349 | 0.12 | 6.77E-48 | Tb |
| Macro-TREM2 | CLDN23 | 6.33E-47 | 1.740201505 | 0.293 | 0.093 | 1.25E-42 | Tb |
| Macro-TREM2 | PVT1 | 2.11E-49 | 1.610739191 | 0.291 | 0.089 | 4.15E-45 | Tb |
| Macro-TREM2 | COL6A21 | 4.00E-33 | 1.608625711 | 0.348 | 0.164 | 7.86E-29 | Tb |
| Macro-TREM2 | ACO1 | 6.95E-66 | 1.598114978 | 0.595 | 0.295 | 1.37E-61 | Tb |
| Macro-TREM2 | USP30-AS11 | 1.59E-54 | 1.548229723 | 0.435 | 0.173 | 3.14E-50 | Tb |
| Macro-TREM2 | CDA1 | 5.54E-64 | 1.528153104 | 0.512 | 0.214 | 1.09E-59 | Tb |
| Macro-TREM2 | HPGD | 4.37E-27 | 1.511511424 | 0.266 | 0.113 | 8.61E-23 | Tb |
| Macro-TREM2 | PTGDS1 | 2.29E-45 | 1.503510542 | 0.595 | 0.365 | 4.50E-41 | Tb |
| Macro-TREM2 | TSFM | 5.06E-59 | 1.495807122 | 0.485 | 0.201 | 9.97E-55 | Tb |
| Macro-TREM2 | CA11 | 1.07E-44 | 1.493266654 | 0.327 | 0.117 | 2.11E-40 | Tb |
| Macro-TREM2 | MMP12 | 3.32E-90 | 1.471291922 | 0.393 | 0.094 | 6.54E-86 | Tb |
| Macro-TREM2 | APOC2 | 1.74E-15 | 1.457899573 | 0.321 | 0.201 | 3.43E-11 | Tb |
| Macro-TREM2 | MT1E | 5.87E-18 | 1.422851288 | 0.279 | 0.147 | 1.16E-13 | Tb |
| Macro-TREM2 | PKD2 | 3.57E-42 | 1.420284873 | 0.365 | 0.149 | 7.04E-38 | Tb |
| Macro-TREM2 | ANK2 | 4.47E-49 | 1.399967759 | 0.279 | 0.083 | 8.81E-45 | Tb |
| Macro-TREM2 | SIGLEC10 | 2.09E-41 | 1.398082356 | 0.405 | 0.18 | 4.12E-37 | Tb |
| Macro-TREM2 | HK3 | 7.17E-49 | 1.393048407 | 0.393 | 0.152 | 1.41E-44 | Tb |
| Macro-TREM2 | ZNF880 | 1.44E-29 | 1.367506289 | 0.258 | 0.099 | 2.83E-25 | Tb |
| Macro-TREM2 | ANKRD22 | 1.08E-21 | 1.364011048 | 0.276 | 0.131 | 2.13E-17 | Tb |
| Macro-TREM2 | IDO1 | 5.76E-16 | 1.360206372 | 0.283 | 0.159 | 1.13E-11 | Tb |
| Macro-TREM2 | S100A8 | 3.07E-102 | 1.350210478 | 0.762 | 0.339 | 6.05E-98 | Tb |
| Macro-TREM2 | PFKP1 | 2.00E-52 | 1.348379162 | 0.457 | 0.197 | 3.94E-48 | Tb |
| Macro-TREM2 | HSD11B11 | 1.16E-82 | 1.345101098 | 0.538 | 0.186 | 2.28E-78 | Tb |
| Macro-TREM2 | CXCL91 | 1.84E-14 | 1.328480739 | 0.348 | 0.222 | 3.63E-10 | Tb |
| Macro-TREM2 | PNPLA4 | 1.30E-24 | 1.322482272 | 0.28 | 0.125 | 2.55E-20 | Tb |
| Macro-TREM2 | DNAI2 | 1.53E-51 | 1.320283794 | 0.352 | 0.122 | 3.01E-47 | Tb |
| Macro-TREM2 | DENND6B | 5.93E-22 | 1.30426767 | 0.258 | 0.118 | 1.17E-17 | Tb |
| Macro-TREM2 | MRC1 | 6.56E-65 | 1.285331919 | 0.621 | 0.316 | 1.29E-60 | Tb |
| Macro-TREM2 | ALOX5 | 9.70E-62 | 1.27018037 | 0.63 | 0.342 | 1.91E-57 | Tb |
| Macro-TREM2 | PFDN1 | 7.42E-51 | 1.269770759 | 0.615 | 0.345 | 1.46E-46 | Tb |
| Macro-TREM2 | MOB3B | 9.86E-37 | 1.263629708 | 0.4 | 0.187 | 1.94E-32 | Tb |
| Macro-TREM2 | MSC-AS1 | 1.21E-39 | 1.259889222 | 0.364 | 0.152 | 2.39E-35 | Tb |
| Macro-TREM2 | COL6A11 | 1.57E-36 | 1.255627038 | 0.368 | 0.162 | 3.09E-32 | Tb |
| Macro-TREM2 | NIBAN2 | 7.38E-33 | 1.25518063 | 0.302 | 0.121 | 1.45E-28 | Tb |
| Macro-TREM2 | STEAP31 | 4.14E-52 | 1.254737088 | 0.452 | 0.187 | 8.15E-48 | Tb |
| Macro-TREM2 | SULT1C2 | 5.51E-53 | 1.25365672 | 0.541 | 0.274 | 1.09E-48 | Tb |
| Macro-TREM2 | C1RL | 8.97E-30 | 1.251853929 | 0.258 | 0.1 | 1.76E-25 | Tb |
| Macro-TREM2 | PPA1 | 4.64E-49 | 1.247134584 | 0.71 | 0.45 | 9.14E-45 | Tb |
| Macro-TREM2 | SMUG1 | 1.09E-21 | 1.234995144 | 0.251 | 0.112 | 2.15E-17 | Tb |
| Macro-TREM2 | ANOS1 | 8.91E-25 | 1.234176151 | 0.254 | 0.106 | 1.75E-20 | Tb |
| Macro-TREM2 | SLC27A3 | 4.07E-38 | 1.223861408 | 0.367 | 0.153 | 8.01E-34 | Tb |
| Macro-TREM2 | TAX1BP3 | 2.34E-44 | 1.220996089 | 0.445 | 0.198 | 4.61E-40 | Tb |
| Macro-TREM2 | CCL51 | 1.87E-23 | 1.218525569 | 0.5 | 0.317 | 3.68E-19 | Tb |
| Macro-TREM2 | PDCD1LG2 | 1.31E-58 | 1.21318137 | 0.467 | 0.186 | 2.58E-54 | Tb |
| Macro-TREM2 | APOL4 | 2.57E-46 | 1.203611006 | 0.357 | 0.132 | 5.06E-42 | Tb |
| Macro-TREM2 | GPR155 | 2.65E-33 | 1.194304184 | 0.332 | 0.143 | 5.21E-29 | Tb |
| Macro-TREM2 | CNIH3 | 1.71E-40 | 1.19209457 | 0.28 | 0.096 | 3.38E-36 | Tb |
| Macro-TREM2 | XIST | 1.01E-101 | 1.18994542 | 0.551 | 0.162 | 1.99E-97 | Tb |
| Macro-TREM2 | TYMP | 4.87E-121 | 1.188895878 | 0.937 | 0.805 | 9.58E-117 | Tb |
| Macro-TREM2 | IFIT3 | 3.59E-47 | 1.183394458 | 0.446 | 0.189 | 7.07E-43 | Tb |
| Macro-TREM2 | NUPR1 | 4.21E-52 | 1.182173305 | 0.827 | 0.749 | 8.28E-48 | Tb |

| | | | | | | | |
|---|---|---|---|---|---|---|---|
| Macro-TREM2 | MATK | 3.00E-39 | 1.178185026 | 0.288 | 0.102 | 5.91E-35 | Tb |
| Macro-TREM2 | CHI3L11 | 2.33E-73 | 1.176395434 | 0.76 | 0.48 | 4.58E-69 | Tb |
| Macro-TREM2 | CAVIN3 | 1.19E-32 | 1.170694387 | 0.272 | 0.103 | 2.34E-28 | Tb |
| Macro-TREM2 | TGM2 | 2.86E-31 | 1.168525871 | 0.431 | 0.224 | 5.62E-27 | Tb |
| Macro-TREM2 | APOL2 | 9.51E-41 | 1.168483981 | 0.348 | 0.134 | 1.87E-36 | Tb |
| Macro-TREM2 | METTL1 | 1.02E-25 | 1.148186177 | 0.264 | 0.112 | 2.01E-21 | Tb |
| Macro-TREM2 | AP1S1 | 7.07E-32 | 1.147507405 | 0.475 | 0.253 | 1.39E-27 | Tb |
| Macro-TREM2 | SNX20 | 2.50E-27 | 1.14505641 | 0.294 | 0.127 | 4.92E-23 | Tb |
| Macro-TREM2 | FHAD1 | 1.98E-35 | 1.136868336 | 0.295 | 0.112 | 3.91E-31 | Tb |
| Macro-TREM2 | ASCL21 | 1.32E-34 | 1.128928302 | 0.424 | 0.21 | 2.60E-30 | Tb |
| Macro-TREM2 | ACOT71 | 1.40E-57 | 1.125816055 | 0.552 | 0.257 | 2.75E-53 | Tb |
| Macro-TREM2 | SCO1 | 3.09E-20 | 1.11715021 | 0.298 | 0.149 | 6.08E-16 | Tb |
| Macro-TREM2 | FHL3 | 2.72E-35 | 1.116329456 | 0.312 | 0.122 | 5.36E-31 | Tb |
| Macro-TREM2 | ATR | 3.84E-30 | 1.113063223 | 0.28 | 0.111 | 7.55E-26 | Tb |
| Macro-TREM2 | PRXL2A1 | 1.54E-49 | 1.103648411 | 0.628 | 0.381 | 3.04E-45 | Tb |
| Macro-TREM2 | GCHFR | 2.44E-28 | 1.09763481 | 0.628 | 0.445 | 4.80E-24 | Tb |
| Macro-TREM2 | MT1F | 1.12E-17 | 1.095794575 | 0.276 | 0.144 | 2.21E-13 | Tb |
| Macro-TREM2 | CLU1 | 2.66E-11 | 1.095074005 | 0.382 | 0.262 | 5.24E-07 | Tb |
| Macro-TREM2 | IL3RA1 | 3.41E-59 | 1.095073317 | 0.562 | 0.261 | 6.72E-55 | Tb |
| Macro-TREM2 | RNF217 | 2.04E-19 | 1.09482061 | 0.288 | 0.146 | 4.01E-15 | Tb |
| Macro-TREM2 | FAIM | 1.19E-19 | 1.094708639 | 0.331 | 0.181 | 2.34E-15 | Tb |
| Macro-TREM2 | RAB13 | 4.09E-48 | 1.093337512 | 0.761 | 0.592 | 8.05E-44 | Tb |
| Macro-TREM2 | LTV1 | 6.13E-23 | 1.091547019 | 0.275 | 0.124 | 1.21E-18 | Tb |
| Macro-TREM2 | ZNF618 | 2.95E-42 | 1.088547932 | 0.312 | 0.11 | 5.80E-38 | Tb |
| Macro-TREM2 | CBR3 | 3.33E-29 | 1.087395696 | 0.339 | 0.155 | 6.55E-25 | Tb |
| Macro-TREM2 | ALDH21 | 5.85E-80 | 1.084605554 | 0.871 | 0.747 | 1.15E-75 | Tb |
| Macro-TREM2 | S100A13 | 2.56E-57 | 1.083931213 | 0.486 | 0.205 | 5.05E-53 | Tb |
| Macro-TREM2 | MED6 | 6.03E-12 | 1.081487361 | 0.271 | 0.156 | 1.19E-07 | Tb |
| Macro-TREM2 | CIR11 | 3.56E-39 | 1.07921501 | 0.659 | 0.413 | 7.01E-35 | Tb |
| Macro-TREM2 | MAGOHB | 1.12E-21 | 1.07677057 | 0.295 | 0.144 | 2.21E-17 | Tb |
| Macro-TREM2 | SLC12A5 | 2.88E-21 | 1.075138776 | 0.302 | 0.148 | 5.67E-17 | Tb |
| Macro-TREM2 | GSN1 | 7.73E-75 | 1.073271543 | 0.864 | 0.669 | 1.52E-70 | Tb |
| Macro-TREM2 | SLC30A1 | 6.29E-24 | 1.068702448 | 0.308 | 0.147 | 1.24E-19 | Tb |
| Macro-TREM2 | LIMK1 | 2.43E-28 | 1.068101251 | 0.431 | 0.229 | 4.78E-24 | Tb |
| Macro-TREM2 | RASAL2 | 1.25E-41 | 1.066861389 | 0.589 | 0.342 | 2.46E-37 | Tb |
| Macro-TREM2 | CCDC115 | 3.29E-37 | 1.065352823 | 0.591 | 0.357 | 6.48E-33 | Tb |
| Macro-TREM2 | ITGAM | 3.54E-39 | 1.060608703 | 0.526 | 0.285 | 6.96E-35 | Tb |
| Macro-TREM2 | PRDX2 | 4.10E-23 | 1.056741175 | 0.33 | 0.168 | 8.06E-19 | Tb |
| Macro-TREM2 | CLYBL | 1.06E-20 | 1.055781097 | 0.257 | 0.12 | 2.09E-16 | Tb |
| Macro-TREM2 | MNDA | 2.72E-54 | 1.053828174 | 0.762 | 0.505 | 5.36E-50 | Tb |
| Macro-TREM2 | ARL11 | 1.58E-42 | 1.0521609 | 0.42 | 0.177 | 3.10E-38 | Tb |
| Macro-TREM2 | HYAL2 | 1.94E-29 | 1.046559024 | 0.342 | 0.157 | 3.83E-25 | Tb |
| Macro-TREM2 | ZMAT3 | 9.87E-22 | 1.045434215 | 0.282 | 0.136 | 1.94E-17 | Tb |
| Macro-TREM2 | ADAMDEC11 | 6.94E-56 | 1.04485634 | 0.74 | 0.486 | 1.37E-51 | Tb |
| Macro-TREM2 | MRS2 | 1.04E-16 | 1.039945759 | 0.273 | 0.139 | 2.04E-12 | Tb |
| Macro-TREM2 | CEBPA | 1.75E-32 | 1.031517677 | 0.622 | 0.403 | 3.45E-28 | Tb |
| Macro-TREM2 | PDLIM7 | 3.22E-20 | 1.030579488 | 0.25 | 0.117 | 6.34E-16 | Tb |
| Macro-TREM2 | ZNF32 | 8.91E-19 | 1.030300349 | 0.363 | 0.198 | 1.75E-14 | Tb |
| Macro-TREM2 | COQ7 | 2.53E-21 | 1.025269536 | 0.265 | 0.125 | 4.97E-17 | Tb |
| Macro-TREM2 | ERAP2 | 1.08E-39 | 1.023329705 | 0.294 | 0.104 | 2.12E-35 | Tb |
| Macro-TREM2 | SCPEP11 | 1.15E-48 | 1.023024535 | 0.78 | 0.663 | 2.26E-44 | Tb |
| Macro-TREM2 | SUCNR1 | 6.00E-34 | 1.022193204 | 0.371 | 0.166 | 1.18E-29 | Tb |
| Macro-TREM2 | NEK61 | 3.07E-42 | 1.021828868 | 0.574 | 0.322 | 6.05E-38 | Tb |
| Macro-TREM2 | MT-ATP6 | 1.03E-63 | 1.018965638 | 0.944 | 0.893 | 2.02E-59 | Tb |
| Macro-TREM2 | COQ9 | 8.95E-25 | 1.01572946 | 0.331 | 0.161 | 1.76E-20 | Tb |

| | | | | | | | |
|---|---|---|---|---|---|---|---|
| Macro-TREM2 | IKBKE1 | 3.91E-24 | 1.010468627 | 0.407 | 0.218 | 7.70E-20 | Tb |
| Macro-TREM2 | NUB1 | 2.86E-33 | 1.009193498 | 0.622 | 0.399 | 5.63E-29 | Tb |
| Macro-TREM2 | SSBP3 | 1.21E-16 | 1.006555063 | 0.272 | 0.144 | 2.39E-12 | Tb |
| Macro-TREM2 | STIM2 | 1.19E-22 | 1.005431016 | 0.297 | 0.142 | 2.35E-18 | Tb |
| Macro-TREM2 | TST | 9.28E-25 | 1.001676719 | 0.324 | 0.154 | 1.83E-20 | Tb |
| **Disease-associated genes in Macro-THBS1** | | | | | | | |
| Macro-THBS1 | OTOA | 9.74E-131 | 4.470461637 | 0.42 | 0.044 | 1.92E-126 | Lep-L |
| Macro-THBS1 | APOE | 6.56E-224 | 4.023836698 | 0.841 | 0.347 | 1.29E-219 | Lep-L |
| Macro-THBS1 | TFPI | 2.93E-77 | 3.782467675 | 0.334 | 0.063 | 5.76E-73 | Lep-L |
| Macro-THBS1 | GPNMB | 7.25E-231 | 3.571047713 | 0.791 | 0.212 | 1.43E-226 | Lep-L |
| Macro-THBS1 | APOC1 | 3.19E-185 | 3.501146844 | 0.77 | 0.277 | 6.28E-181 | Lep-L |
| Macro-THBS1 | CYP27A1 | 9.11E-128 | 3.461782712 | 0.467 | 0.071 | 1.79E-123 | Lep-L |
| Macro-THBS1 | SELENOP | 6.07E-91 | 3.083380714 | 0.392 | 0.073 | 1.20E-86 | Lep-L |
| Macro-THBS1 | HSD17B14 | 6.82E-55 | 2.851985873 | 0.28 | 0.066 | 1.34E-50 | Lep-L |
| Macro-THBS1 | CCL20 | 3.45E-142 | 2.774966358 | 0.733 | 0.306 | 6.80E-138 | Lep-L |
| Macro-THBS1 | RARRES1 | 1.48E-76 | 2.724477533 | 0.398 | 0.104 | 2.92E-72 | Lep-L |
| Macro-THBS1 | SGPP2 | 7.53E-89 | 2.604390624 | 0.397 | 0.08 | 1.48E-84 | Lep-L |
| Macro-THBS1 | LIPA | 4.21E-103 | 2.442610549 | 0.609 | 0.254 | 8.29E-99 | Lep-L |
| Macro-THBS1 | CCR7 | 3.93E-41 | 2.315268006 | 0.347 | 0.139 | 7.73E-37 | Lep-L |
| Macro-THBS1 | VAT1 | 7.00E-74 | 2.259131338 | 0.44 | 0.143 | 1.38E-69 | Lep-L |
| Macro-THBS1 | TREM2 | 1.72E-60 | 2.24609343 | 0.372 | 0.115 | 3.39E-56 | Lep-L |
| Macro-THBS1 | TNFAIP6 | 4.70E-113 | 2.208920038 | 0.738 | 0.365 | 9.24E-109 | Lep-L |
| Macro-THBS1 | EBI3 | 2.59E-68 | 2.187021387 | 0.416 | 0.129 | 5.09E-64 | Lep-L |
| Macro-THBS1 | THBS1 | 3.04E-35 | 2.184027574 | 0.521 | 0.325 | 5.98E-31 | Lep-L |
| Macro-THBS1 | NKG7 | 2.32E-35 | 2.178943398 | 0.259 | 0.085 | 4.57E-31 | Lep-L |
| Macro-THBS1 | MMP19 | 2.28E-76 | 2.102501268 | 0.685 | 0.382 | 4.48E-72 | Lep-L |
| Macro-THBS1 | LSS | 2.89E-63 | 2.077257905 | 0.463 | 0.181 | 5.68E-59 | Lep-L |
| Macro-THBS1 | CXCL1 | 6.70E-48 | 2.052514855 | 0.422 | 0.174 | 1.32E-43 | Lep-L |
| Macro-THBS1 | SRGAP1 | 2.45E-73 | 2.028830841 | 0.632 | 0.328 | 4.83E-69 | Lep-L |
| Macro-THBS1 | MIR3142HG | 2.48E-53 | 2.011656921 | 0.36 | 0.113 | 4.89E-49 | Lep-L |
| Macro-THBS1 | EREG | 2.81E-110 | 2.007813993 | 0.848 | 0.584 | 5.52E-106 | Lep-L |
| Macro-THBS1 | TRIP10 | 2.03E-85 | 2.002869361 | 0.508 | 0.167 | 3.99E-81 | Lep-L |
| Macro-THBS1 | CXCL3 | 3.28E-155 | 1.994281653 | 0.915 | 0.566 | 6.45E-151 | Lep-L |
| Macro-THBS1 | MGLL | 1.34E-146 | 1.983920179 | 0.748 | 0.267 | 2.64E-142 | Lep-L |
| Macro-THBS1 | CD9 | 2.67E-84 | 1.979358416 | 0.59 | 0.26 | 5.25E-80 | Lep-L |
| Macro-THBS1 | SDC4 | 2.25E-55 | 1.933629288 | 0.516 | 0.262 | 4.43E-51 | Lep-L |
| Macro-THBS1 | IL23A | 3.88E-21 | 1.93027085 | 0.28 | 0.142 | 7.64E-17 | Lep-L |
| Macro-THBS1 | TNIP3 | 9.23E-41 | 1.92002558 | 0.407 | 0.185 | 1.82E-36 | Lep-L |
| Macro-THBS1 | NBPF19 | 6.82E-107 | 1.899547838 | 0.67 | 0.273 | 1.34E-102 | Lep-L |
| Macro-THBS1 | TDP2 | 7.82E-38 | 1.898639924 | 0.331 | 0.134 | 1.54E-33 | Lep-L |
| Macro-THBS1 | SMPDL3A | 1.03E-44 | 1.891851467 | 0.336 | 0.12 | 2.03E-40 | Lep-L |
| Macro-THBS1 | SMS | 8.24E-151 | 1.889881896 | 0.822 | 0.421 | 1.62E-146 | Lep-L |
| Macro-THBS1 | A2M | 2.04E-63 | 1.867685595 | 0.446 | 0.164 | 4.02E-59 | Lep-L |
| Macro-THBS1 | FOLR2 | 2.71E-63 | 1.83659401 | 0.457 | 0.167 | 5.34E-59 | Lep-L |
| Macro-THBS1 | RASGRP3 | 3.91E-75 | 1.831771374 | 0.577 | 0.241 | 7.69E-71 | Lep-L |
| Macro-THBS1 | CXCL8 | 2.65E-206 | 1.799459283 | 0.983 | 0.745 | 5.22E-202 | Lep-L |
| Macro-THBS1 | ADA | 2.57E-81 | 1.793375549 | 0.616 | 0.267 | 5.07E-77 | Lep-L |
| Macro-THBS1 | FILIP1L | 2.10E-31 | 1.764114815 | 0.473 | 0.284 | 4.14E-27 | Lep-L |
| Macro-THBS1 | BNIP3L | 1.15E-108 | 1.763345048 | 0.934 | 0.792 | 2.27E-104 | Lep-L |
| Macro-THBS1 | EPS15L1 | 3.93E-32 | 1.762857324 | 0.272 | 0.099 | 7.74E-28 | Lep-L |
| Macro-THBS1 | GK-AS1 | 1.86E-41 | 1.753572495 | 0.456 | 0.232 | 3.66E-37 | Lep-L |
| Macro-THBS1 | PLPP3 | 1.64E-50 | 1.752919708 | 0.336 | 0.102 | 3.22E-46 | Lep-L |
| Macro-THBS1 | SNTB1 | 5.65E-45 | 1.751781189 | 0.469 | 0.233 | 1.11E-40 | Lep-L |
| Macro-THBS1 | SDS | 1.15E-42 | 1.74638706 | 0.507 | 0.276 | 2.27E-38 | Lep-L |
| Macro-THBS1 | SPARC | 3.91E-29 | 1.739710489 | 0.261 | 0.102 | 7.70E-25 | Lep-L |

| | | | | | | | |
|---|---|---|---|---|---|---|---|
| Macro-THBS1 | SLC1A3 | 7.99E-85 | 1.736335707 | 0.739 | 0.417 | 1.57E-80 | Lep-L |
| Macro-THBS1 | AP3B1 | 2.10E-54 | 1.721566079 | 0.732 | 0.551 | 4.13E-50 | Lep-L |
| Macro-THBS1 | CTSD | 8.90E-147 | 1.720036945 | 0.93 | 0.731 | 1.75E-142 | Lep-L |
| Macro-THBS1 | PAM | 5.98E-43 | 1.705782187 | 0.379 | 0.15 | 1.18E-38 | Lep-L |
| Macro-THBS1 | CHST7 | 2.44E-69 | 1.701449974 | 0.434 | 0.13 | 4.80E-65 | Lep-L |
| Macro-THBS1 | PHF1 | 5.19E-52 | 1.674479088 | 0.433 | 0.181 | 1.02E-47 | Lep-L |
| Macro-THBS1 | SH3D21 | 1.16E-34 | 1.670675645 | 0.31 | 0.122 | 2.29E-30 | Lep-L |
| Macro-THBS1 | CD82 | 2.14E-58 | 1.670239218 | 0.495 | 0.213 | 4.21E-54 | Lep-L |
| Macro-THBS1 | FABP5 | 7.73E-73 | 1.6579003 | 0.819 | 0.627 | 1.52E-68 | Lep-L |
| Macro-THBS1 | ALDH1A1 | 9.49E-43 | 1.651453426 | 0.438 | 0.211 | 1.87E-38 | Lep-L |
| Macro-THBS1 | MPZL1 | 4.51E-35 | 1.647093811 | 0.446 | 0.247 | 8.88E-31 | Lep-L |
| Macro-THBS1 | ACSL4 | 1.57E-41 | 1.63492515 | 0.685 | 0.548 | 3.09E-37 | Lep-L |
| Macro-THBS1 | NR6A1 | 1.99E-49 | 1.622718568 | 0.404 | 0.152 | 3.92E-45 | Lep-L |
| Macro-THBS1 | SLC30A4 | 5.14E-28 | 1.616754788 | 0.291 | 0.127 | 1.01E-23 | Lep-L |
| Macro-THBS1 | HIPK2 | 1.37E-51 | 1.611612053 | 0.578 | 0.322 | 2.70E-47 | Lep-L |
| Macro-THBS1 | THAP2 | 3.60E-47 | 1.607848424 | 0.574 | 0.338 | 7.08E-43 | Lep-L |
| Macro-THBS1 | SLAMF1 | 2.56E-18 | 1.605832393 | 0.364 | 0.236 | 5.04E-14 | Lep-L |
| Macro-THBS1 | LGMN | 2.30E-61 | 1.591929857 | 0.663 | 0.399 | 4.52E-57 | Lep-L |
| Macro-THBS1 | CBLB | 1.43E-22 | 1.58679191 | 0.279 | 0.134 | 2.81E-18 | Lep-L |
| Macro-THBS1 | NR1H3 | 6.08E-33 | 1.569724452 | 0.38 | 0.185 | 1.20E-28 | Lep-L |
| Macro-THBS1 | MAP1LC3A | 1.02E-20 | 1.559392084 | 0.271 | 0.134 | 2.02E-16 | Lep-L |
| Macro-THBS1 | PTGER2 | 1.08E-38 | 1.551818797 | 0.402 | 0.185 | 2.13E-34 | Lep-L |
| Macro-THBS1 | CD59 | 2.71E-49 | 1.550804984 | 0.517 | 0.279 | 5.34E-45 | Lep-L |
| Macro-THBS1 | ITGB8 | 6.25E-30 | 1.539008391 | 0.346 | 0.162 | 1.23E-25 | Lep-L |
| Macro-THBS1 | IL18 | 2.09E-94 | 1.538187264 | 0.771 | 0.449 | 4.12E-90 | Lep-L |
| Macro-THBS1 | MTURN | 1.76E-21 | 1.535540684 | 0.254 | 0.119 | 3.46E-17 | Lep-L |
| Macro-THBS1 | CD72 | 8.57E-48 | 1.530547496 | 0.549 | 0.318 | 1.69E-43 | Lep-L |
| Macro-THBS1 | CREG1 | 3.11E-104 | 1.529819459 | 0.833 | 0.584 | 6.12E-100 | Lep-L |
| Macro-THBS1 | TXNRD1 | 3.72E-87 | 1.523510791 | 0.81 | 0.542 | 7.32E-83 | Lep-L |
| Macro-THBS1 | PDXDC1 | 1.26E-48 | 1.517722672 | 0.482 | 0.232 | 2.48E-44 | Lep-L |
| Macro-THBS1 | DENND4C | 6.81E-48 | 1.516402266 | 0.45 | 0.202 | 1.34E-43 | Lep-L |
| Macro-THBS1 | SCCPDH | 6.45E-26 | 1.516151019 | 0.27 | 0.118 | 1.27E-21 | Lep-L |
| Macro-THBS1 | PLIN2 | 2.73E-106 | 1.508848249 | 0.901 | 0.661 | 5.36E-102 | Lep-L |
| Macro-THBS1 | KDM5B | 5.18E-36 | 1.503228192 | 0.372 | 0.166 | 1.02E-31 | Lep-L |
| Macro-THBS1 | KMO | 1.31E-108 | 1.498095845 | 0.846 | 0.498 | 2.59E-104 | Lep-L |
| Macro-THBS1 | WIPI1 | 4.61E-27 | 1.491741426 | 0.34 | 0.173 | 9.07E-23 | Lep-L |
| Macro-THBS1 | PTGIR | 6.94E-31 | 1.486756306 | 0.285 | 0.109 | 1.37E-26 | Lep-L |
| Macro-THBS1 | RDX | 2.53E-71 | 1.486654887 | 0.729 | 0.445 | 4.98E-67 | Lep-L |
| Macro-THBS1 | STMN1 | 1.09E-25 | 1.484954216 | 0.321 | 0.16 | 2.14E-21 | Lep-L |
| Macro-THBS1 | CMSS1 | 7.31E-34 | 1.482305824 | 0.534 | 0.339 | 1.44E-29 | Lep-L |
| Macro-THBS1 | VSIG4 | 3.68E-57 | 1.476162199 | 0.493 | 0.204 | 7.23E-53 | Lep-L |
| Macro-THBS1 | TNFRSF4 | 2.47E-24 | 1.475437144 | 0.27 | 0.116 | 4.86E-20 | Lep-L |
| Macro-THBS1 | KCNE1 | 1.92E-31 | 1.466034125 | 0.362 | 0.172 | 3.78E-27 | Lep-L |
| Macro-THBS1 | MARCKS | 1.16E-141 | 1.454406223 | 0.964 | 0.794 | 2.29E-137 | Lep-L |
| Macro-THBS1 | MAPK13 | 1.77E-23 | 1.451004789 | 0.275 | 0.128 | 3.48E-19 | Lep-L |
| Macro-THBS1 | STAT5A | 6.47E-52 | 1.442317809 | 0.61 | 0.374 | 1.27E-47 | Lep-L |
| Macro-THBS1 | MS4A7 | 3.52E-120 | 1.439247708 | 0.871 | 0.602 | 6.92E-116 | Lep-L |
| Macro-THBS1 | ZNF281 | 2.04E-33 | 1.421357753 | 0.386 | 0.189 | 4.01E-29 | Lep-L |
| Macro-THBS1 | INF2 | 6.33E-36 | 1.418214986 | 0.384 | 0.181 | 1.25E-31 | Lep-L |
| Macro-THBS1 | FUCA1 | 2.36E-25 | 1.412419113 | 0.282 | 0.13 | 4.65E-21 | Lep-L |
| Macro-THBS1 | FTX | 4.64E-51 | 1.404550454 | 0.662 | 0.416 | 9.13E-47 | Lep-L |
| Macro-THBS1 | MFSD2A | 3.21E-50 | 1.391548018 | 0.538 | 0.281 | 6.31E-46 | Lep-L |
| Macro-THBS1 | DNAAF1 | 1.66E-17 | 1.390736869 | 0.333 | 0.203 | 3.27E-13 | Lep-L |
| Macro-THBS1 | RNF144B | 2.19E-58 | 1.388702892 | 0.819 | 0.642 | 4.31E-54 | Lep-L |
| Macro-THBS1 | CCNG1 | 1.78E-29 | 1.381433902 | 0.331 | 0.154 | 3.51E-25 | Lep-L |

| | | | | | | | |
|---|---|---|---|---|---|---|---|
| Macro-THBS1 | SEC22B | 4.65E-67 | 1.377088985 | 0.675 | 0.389 | 9.15E-63 | Lep-L |
| Macro-THBS1 | STAM | 2.48E-24 | 1.36984541 | 0.328 | 0.168 | 4.87E-20 | Lep-L |
| Macro-THBS1 | PI4K2A | 3.37E-44 | 1.369602976 | 0.502 | 0.271 | 6.63E-40 | Lep-L |
| Macro-THBS1 | ATP6V1C1 | 4.60E-91 | 1.367987197 | 0.797 | 0.495 | 9.06E-87 | Lep-L |
| Macro-THBS1 | CCSER2 | 6.30E-31 | 1.367771155 | 0.505 | 0.326 | 1.24E-26 | Lep-L |
| Macro-THBS1 | RB1CC1 | 1.90E-69 | 1.365177184 | 0.702 | 0.433 | 3.75E-65 | Lep-L |
| Macro-THBS1 | ITPKB | 2.06E-32 | 1.361960325 | 0.371 | 0.173 | 4.06E-28 | Lep-L |
| Macro-THBS1 | TESK1 | 5.87E-34 | 1.353145547 | 0.392 | 0.191 | 1.15E-29 | Lep-L |
| Macro-THBS1 | AQP9 | 5.39E-86 | 1.346967661 | 0.864 | 0.574 | 1.06E-81 | Lep-L |
| Macro-THBS1 | CYB5A | 1.78E-44 | 1.346154615 | 0.532 | 0.295 | 3.50E-40 | Lep-L |
| Macro-THBS1 | MAFG | 5.39E-70 | 1.344575181 | 0.657 | 0.339 | 1.06E-65 | Lep-L |
| Macro-THBS1 | TCEAL9 | 1.27E-38 | 1.342633441 | 0.4 | 0.181 | 2.50E-34 | Lep-L |
| Macro-THBS1 | SRPK1 | 1.27E-29 | 1.342202444 | 0.373 | 0.19 | 2.51E-25 | Lep-L |
| Macro-THBS1 | NCEH1 | 2.12E-22 | 1.339806737 | 0.394 | 0.237 | 4.17E-18 | Lep-L |
| Macro-THBS1 | PLD1 | 2.93E-46 | 1.329151587 | 0.469 | 0.222 | 5.77E-42 | Lep-L |
| Macro-THBS1 | ALCAM | 7.92E-102 | 1.327770278 | 0.897 | 0.589 | 1.56E-97 | Lep-L |
| Macro-THBS1 | HOMER3 | 1.21E-34 | 1.325203261 | 0.357 | 0.154 | 2.39E-30 | Lep-L |
| Macro-THBS1 | ANKUB1 | 1.13E-32 | 1.325055692 | 0.477 | 0.281 | 2.22E-28 | Lep-L |
| Macro-THBS1 | FAM177B | 1.13E-27 | 1.316055498 | 0.347 | 0.174 | 2.23E-23 | Lep-L |
| Macro-THBS1 | CXXC5 | 8.52E-34 | 1.313492695 | 0.416 | 0.213 | 1.68E-29 | Lep-L |
| Macro-THBS1 | FBXO42 | 8.47E-28 | 1.30990103 | 0.353 | 0.178 | 1.67E-23 | Lep-L |
| Macro-THBS1 | FEM1B | 3.64E-24 | 1.309714545 | 0.334 | 0.176 | 7.16E-20 | Lep-L |
| Macro-THBS1 | PXDC1 | 3.96E-21 | 1.301519777 | 0.331 | 0.184 | 7.79E-17 | Lep-L |
| Macro-THBS1 | SLC7A11 | 4.15E-31 | 1.299456676 | 0.375 | 0.174 | 8.17E-27 | Lep-L |
| Macro-THBS1 | BCL2L1 | 4.02E-18 | 1.298689677 | 0.288 | 0.158 | 7.90E-14 | Lep-L |
| Macro-THBS1 | PDE8A | 1.37E-70 | 1.292788033 | 0.689 | 0.353 | 2.69E-66 | Lep-L |
| Macro-THBS1 | RAB9A | 1.36E-54 | 1.292601365 | 0.603 | 0.325 | 2.68E-50 | Lep-L |
| Macro-THBS1 | RAB12 | 1.04E-40 | 1.291602543 | 0.492 | 0.263 | 2.05E-36 | Lep-L |
| Macro-THBS1 | UBR4 | 1.40E-44 | 1.288847197 | 0.526 | 0.283 | 2.75E-40 | Lep-L |
| Macro-THBS1 | FAM3C | 6.56E-33 | 1.28849083 | 0.457 | 0.258 | 1.29E-28 | Lep-L |
| Macro-THBS1 | NPL | 6.52E-63 | 1.287954566 | 0.644 | 0.365 | 1.28E-58 | Lep-L |
| Macro-THBS1 | KLHL21 | 5.58E-33 | 1.287937182 | 0.417 | 0.21 | 1.10E-28 | Lep-L |
| Macro-THBS1 | NTAN1 | 7.36E-33 | 1.281448425 | 0.436 | 0.237 | 1.45E-28 | Lep-L |
| Macro-THBS1 | CD84 | 1.20E-45 | 1.277647816 | 0.551 | 0.304 | 2.35E-41 | Lep-L |
| Macro-THBS1 | EIF1B | 4.39E-81 | 1.263258608 | 0.812 | 0.575 | 8.64E-77 | Lep-L |
| Macro-THBS1 | GRAMD1A | 4.23E-63 | 1.259731716 | 0.712 | 0.409 | 8.33E-59 | Lep-L |
| Macro-THBS1 | ACP5 | 7.13E-55 | 1.258562068 | 0.652 | 0.404 | 1.40E-50 | Lep-L |
| Macro-THBS1 | ANKLE2 | 2.81E-56 | 1.249049685 | 0.628 | 0.36 | 5.52E-52 | Lep-L |
| Macro-THBS1 | BTG1 | 1.47E-160 | 1.24227768 | 0.991 | 0.957 | 2.90E-156 | Lep-L |
| Macro-THBS1 | CCDC107 | 1.63E-58 | 1.23914865 | 0.691 | 0.45 | 3.21E-54 | Lep-L |
| Macro-THBS1 | DCAF6 | 1.14E-26 | 1.230234227 | 0.403 | 0.223 | 2.24E-22 | Lep-L |
| Macro-THBS1 | HSPA13 | 2.07E-19 | 1.229887679 | 0.289 | 0.151 | 4.08E-15 | Lep-L |
| Macro-THBS1 | RFFL | 1.22E-20 | 1.229428571 | 0.339 | 0.197 | 2.41E-16 | Lep-L |
| Macro-THBS1 | STK4 | 1.26E-80 | 1.224585322 | 0.842 | 0.604 | 2.47E-76 | Lep-L |
| Macro-THBS1 | ARID3A | 4.87E-47 | 1.22058655 | 0.524 | 0.267 | 9.59E-43 | Lep-L |
| Macro-THBS1 | CYP1B1 | 1.18E-16 | 1.219235243 | 0.25 | 0.125 | 2.31E-12 | Lep-L |
| Macro-THBS1 | ZNF638 | 1.40E-26 | 1.212922403 | 0.549 | 0.379 | 2.76E-22 | Lep-L |
| Macro-THBS1 | ASAH1 | 1.11E-146 | 1.212742099 | 0.941 | 0.801 | 2.18E-142 | Lep-L |
| Macro-THBS1 | DAPK1 | 1.46E-63 | 1.209615676 | 0.762 | 0.497 | 2.88E-59 | Lep-L |
| Macro-THBS1 | TNIP1 | 6.53E-66 | 1.208589539 | 0.779 | 0.536 | 1.29E-61 | Lep-L |
| Macro-THBS1 | MCRIP2 | 8.70E-29 | 1.204226849 | 0.406 | 0.219 | 1.71E-24 | Lep-L |
| Macro-THBS1 | PLTP | 5.15E-57 | 1.202728338 | 0.464 | 0.174 | 1.01E-52 | Lep-L |
| Macro-THBS1 | DIP2A | 6.95E-16 | 1.198007794 | 0.255 | 0.137 | 1.37E-11 | Lep-L |
| Macro-THBS1 | SEC62 | 1.04E-79 | 1.194667868 | 0.825 | 0.581 | 2.06E-75 | Lep-L |
| Macro-THBS1 | UBXN7 | 3.20E-27 | 1.192017961 | 0.368 | 0.19 | 6.30E-23 | Lep-L |

| | | | | | | | |
|---|---|---|---|---|---|---|---|
| Macro-THBS1 | CLN8 | 2.07E-32 | 1.191296964 | 0.485 | 0.284 | 4.07E-28 | Lep-L |
| Macro-THBS1 | MS4A14 | 1.62E-21 | 1.176504064 | 0.279 | 0.132 | 3.19E-17 | Lep-L |
| Macro-THBS1 | TOM1L2 | 7.50E-17 | 1.170254882 | 0.282 | 0.155 | 1.48E-12 | Lep-L |
| Macro-THBS1 | TNS1 | 1.85E-19 | 1.161887786 | 0.279 | 0.139 | 3.63E-15 | Lep-L |
| Macro-THBS1 | CTTNBP2NL | 1.43E-33 | 1.159935843 | 0.532 | 0.339 | 2.82E-29 | Lep-L |
| Macro-THBS1 | AMPD3 | 1.48E-78 | 1.159546433 | 0.783 | 0.443 | 2.91E-74 | Lep-L |
| Macro-THBS1 | LRCH3 | 9.18E-24 | 1.158011239 | 0.386 | 0.219 | 1.81E-19 | Lep-L |
| Macro-THBS1 | TSC22D1 | 1.82E-55 | 1.155899875 | 0.684 | 0.414 | 3.57E-51 | Lep-L |
| Macro-THBS1 | TNIK | 6.53E-27 | 1.155594524 | 0.316 | 0.14 | 1.29E-22 | Lep-L |
| Macro-THBS1 | AKR1B1 | 1.87E-23 | 1.154262335 | 0.385 | 0.225 | 3.68E-19 | Lep-L |
| Macro-THBS1 | BMT2 | 5.82E-19 | 1.15215489 | 0.304 | 0.163 | 1.15E-14 | Lep-L |
| Macro-THBS1 | ADD3 | 2.69E-18 | 1.149599355 | 0.303 | 0.164 | 5.30E-14 | Lep-L |
| Macro-THBS1 | PLD3 | 1.88E-68 | 1.147300735 | 0.743 | 0.452 | 3.71E-64 | Lep-L |
| Macro-THBS1 | LRP1 | 3.89E-57 | 1.142334152 | 0.679 | 0.412 | 7.65E-53 | Lep-L |
| Macro-THBS1 | CSGALNACT2 | 6.02E-95 | 1.142128844 | 0.906 | 0.682 | 1.18E-90 | Lep-L |
| Macro-THBS1 | ADORA2A | 1.19E-28 | 1.139622376 | 0.298 | 0.125 | 2.34E-24 | Lep-L |
| Macro-THBS1 | CLCF1 | 1.15E-32 | 1.138713101 | 0.292 | 0.109 | 2.27E-28 | Lep-L |
| Macro-THBS1 | GCLM | 2.04E-29 | 1.138609588 | 0.44 | 0.24 | 4.01E-25 | Lep-L |
| Macro-THBS1 | PDE4DIP | 7.75E-112 | 1.135674967 | 0.922 | 0.565 | 1.52E-107 | Lep-L |
| Macro-THBS1 | ATP6V1D | 7.56E-30 | 1.131983825 | 0.518 | 0.339 | 1.49E-25 | Lep-L |
| Macro-THBS1 | RNASE1 | 3.00E-34 | 1.124737818 | 0.343 | 0.143 | 5.90E-30 | Lep-L |
| Macro-THBS1 | ZNRF2 | 2.48E-17 | 1.120886379 | 0.572 | 0.446 | 4.88E-13 | Lep-L |
| Macro-THBS1 | ZFYVE26 | 2.32E-16 | 1.119651732 | 0.254 | 0.132 | 4.56E-12 | Lep-L |
| Macro-THBS1 | CHIC2 | 2.15E-52 | 1.119545204 | 0.679 | 0.42 | 4.23E-48 | Lep-L |
| Macro-THBS1 | SERPINB9 | 1.53E-90 | 1.119115967 | 0.968 | 0.884 | 3.01E-86 | Lep-L |
| Macro-THBS1 | TMCO3 | 2.69E-21 | 1.118789497 | 0.337 | 0.185 | 5.30E-17 | Lep-L |
| Macro-THBS1 | BASP1 | 9.32E-94 | 1.118729391 | 0.905 | 0.739 | 1.83E-89 | Lep-L |
| Macro-THBS1 | AGPS | 3.27E-19 | 1.117973369 | 0.408 | 0.266 | 6.43E-15 | Lep-L |
| Macro-THBS1 | OPN3 | 4.31E-42 | 1.115691871 | 0.578 | 0.33 | 8.48E-38 | Lep-L |
| Macro-THBS1 | SLC11A2 | 2.81E-44 | 1.115214359 | 0.57 | 0.319 | 5.54E-40 | Lep-L |
| Macro-THBS1 | MS4A4A | 1.10E-46 | 1.114868453 | 0.613 | 0.354 | 2.17E-42 | Lep-L |
| Macro-THBS1 | HLA-DRB5 | 1.54E-90 | 1.114865267 | 0.896 | 0.687 | 3.03E-86 | Lep-L |
| Macro-THBS1 | NRP2 | 2.21E-82 | 1.11303839 | 0.856 | 0.562 | 4.35E-78 | Lep-L |
| Macro-THBS1 | CTSL | 1.04E-92 | 1.111073656 | 0.923 | 0.736 | 2.05E-88 | Lep-L |
| Macro-THBS1 | ZFYVE28 | 2.61E-18 | 1.110496221 | 0.291 | 0.158 | 5.14E-14 | Lep-L |
| Macro-THBS1 | RIN3 | 1.23E-42 | 1.106011279 | 0.687 | 0.458 | 2.43E-38 | Lep-L |
| Macro-THBS1 | ITSN1 | 6.14E-20 | 1.105221259 | 0.266 | 0.128 | 1.21E-15 | Lep-L |
| Macro-THBS1 | IGKC | 9.33E-21 | 1.099939768 | 0.308 | 0.143 | 1.84E-16 | Lep-L |
| Macro-THBS1 | RRAGC | 1.82E-24 | 1.097916239 | 0.44 | 0.28 | 3.59E-20 | Lep-L |
| Macro-THBS1 | HSD17B4 | 3.20E-30 | 1.097372461 | 0.496 | 0.306 | 6.30E-26 | Lep-L |
| Macro-THBS1 | TMEM189 | 2.20E-23 | 1.096210278 | 0.435 | 0.272 | 4.32E-19 | Lep-L |
| Macro-THBS1 | RAD21 | 3.19E-31 | 1.092748752 | 0.544 | 0.361 | 6.28E-27 | Lep-L |
| Macro-THBS1 | OAZ2 | 2.82E-48 | 1.092389028 | 0.65 | 0.414 | 5.55E-44 | Lep-L |
| Macro-THBS1 | MIR155HG | 6.01E-68 | 1.091639247 | 0.805 | 0.502 | 1.18E-63 | Lep-L |
| Macro-THBS1 | MAML2 | 8.21E-26 | 1.089977961 | 0.769 | 0.643 | 1.62E-21 | Lep-L |
| Macro-THBS1 | UGCG | 2.05E-36 | 1.083755942 | 0.579 | 0.374 | 4.03E-32 | Lep-L |
| Macro-THBS1 | MCFD2 | 3.14E-24 | 1.083658512 | 0.405 | 0.237 | 6.18E-20 | Lep-L |
| Macro-THBS1 | KMT2A | 9.61E-29 | 1.079318587 | 0.458 | 0.269 | 1.89E-24 | Lep-L |
| Macro-THBS1 | CERT1 | 1.92E-51 | 1.078150795 | 0.735 | 0.505 | 3.77E-47 | Lep-L |
| Macro-THBS1 | CLOCK | 3.68E-10 | 1.075899639 | 0.251 | 0.164 | 7.25E-06 | Lep-L |
| Macro-THBS1 | HECTD1 | 3.05E-45 | 1.073183357 | 0.601 | 0.36 | 6.00E-41 | Lep-L |
| Macro-THBS1 | STK17A | 1.93E-35 | 1.0689275 | 0.532 | 0.309 | 3.79E-31 | Lep-L |
| Macro-THBS1 | EMSY | 1.63E-16 | 1.063918354 | 0.252 | 0.131 | 3.21E-12 | Lep-L |
| Macro-THBS1 | PSD3 | 1.52E-24 | 1.062546793 | 0.4 | 0.228 | 2.99E-20 | Lep-L |
| Macro-THBS1 | RIT1 | 2.11E-51 | 1.062330703 | 0.679 | 0.415 | 4.16E-47 | Lep-L |

| | | | | | | | |
|---|---|---|---|---|---|---|---|
| Macro-THBS1 | ANKRD10 | 2.85E-43 | 1.059841742 | 0.684 | 0.466 | 5.60E-39 | Lep-L |
| Macro-THBS1 | RCAN1 | 9.35E-28 | 1.058479701 | 0.371 | 0.19 | 1.84E-23 | Lep-L |
| Macro-THBS1 | NBN | 4.81E-39 | 1.057475385 | 0.708 | 0.532 | 9.46E-35 | Lep-L |
| Macro-THBS1 | ARHGEF2 | 6.70E-48 | 1.05605073 | 0.658 | 0.397 | 1.32E-43 | Lep-L |
| Macro-THBS1 | RAB8B | 1.03E-82 | 1.048490491 | 0.896 | 0.669 | 2.02E-78 | Lep-L |
| Macro-THBS1 | CSTB | 3.22E-71 | 1.04626739 | 0.942 | 0.871 | 6.34E-67 | Lep-L |
| Macro-THBS1 | SLC6A6 | 2.75E-36 | 1.043564722 | 0.603 | 0.402 | 5.41E-32 | Lep-L |
| Macro-THBS1 | PPP1R15B | 3.93E-72 | 1.039742155 | 0.839 | 0.614 | 7.74E-68 | Lep-L |
| Macro-THBS1 | GM2A | 8.77E-35 | 1.038934596 | 0.615 | 0.427 | 1.73E-30 | Lep-L |
| Macro-THBS1 | AKIRIN1 | 1.59E-30 | 1.03088011 | 0.546 | 0.358 | 3.13E-26 | Lep-L |
| Macro-THBS1 | TSPAN4 | 8.32E-38 | 1.030779376 | 0.552 | 0.319 | 1.64E-33 | Lep-L |
| Macro-THBS1 | INSIG2 | 2.00E-18 | 1.029034089 | 0.309 | 0.168 | 3.94E-14 | Lep-L |
| Macro-THBS1 | FTL | 7.21E-134 | 1.026349101 | 0.999 | 0.997 | 1.42E-129 | Lep-L |
| Macro-THBS1 | ASPH | 1.10E-19 | 1.025649356 | 0.554 | 0.414 | 2.17E-15 | Lep-L |
| Macro-THBS1 | HSDL2 | 5.41E-14 | 1.023683272 | 0.261 | 0.148 | 1.06E-09 | Lep-L |
| Macro-THBS1 | C5AR1 | 1.64E-73 | 1.023060178 | 0.934 | 0.782 | 3.23E-69 | Lep-L |
| Macro-THBS1 | IRAK1 | 8.13E-22 | 1.020431013 | 0.432 | 0.273 | 1.60E-17 | Lep-L |
| Macro-THBS1 | TNFAIP2 | 5.89E-61 | 1.017368877 | 0.923 | 0.778 | 1.16E-56 | Lep-L |
| Macro-THBS1 | CRYL1 | 1.50E-15 | 1.016393958 | 0.296 | 0.172 | 2.96E-11 | Lep-L |
| Macro-THBS1 | PABPN1 | 1.31E-34 | 1.013910671 | 0.572 | 0.358 | 2.57E-30 | Lep-L |
| Macro-THBS1 | TTYH3 | 1.48E-33 | 1.013837093 | 0.61 | 0.417 | 2.91E-29 | Lep-L |
| Macro-THBS1 | GPR35 | 3.04E-24 | 1.01306714 | 0.336 | 0.164 | 5.98E-20 | Lep-L |
| Macro-THBS1 | BANP | 1.42E-50 | 1.00940004 | 0.733 | 0.465 | 2.80E-46 | Lep-L |
| Macro-THBS1 | IP6K2 | 7.55E-16 | 1.00901375 | 0.281 | 0.157 | 1.49E-11 | Lep-L |
| Macro-THBS1 | GPR107 | 5.17E-25 | 1.007415876 | 0.436 | 0.258 | 1.02E-20 | Lep-L |
| Macro-THBS1 | PIK3CB | 1.51E-26 | 1.0052847 | 0.545 | 0.378 | 2.98E-22 | Lep-L |
| Macro-THBS1 | PLEKHM2 | 2.07E-53 | 1.004601032 | 0.704 | 0.454 | 4.08E-49 | Lep-L |
| Macro-THBS1 | MPP1 | 3.07E-54 | 1.00280628 | 0.75 | 0.505 | 6.05E-50 | Lep-L |
| Macro-THBS1 | PIK3R5 | 2.68E-57 | 1.002296731 | 0.828 | 0.622 | 5.27E-53 | Lep-L |
| Macro-THBS1 | ARL8A | 4.04E-35 | 1.001632032 | 0.584 | 0.376 | 7.96E-31 | Lep-L |
| Macro-THBS1 | EVA1B | 1.40E-26 | 1.001489463 | 0.438 | 0.255 | 2.75E-22 | Lep-L |
| Macro-THBS1 | UBD | 3.08E-30 | 3.45983141 | 0.271 | 0.086 | 6.06E-26 | Lep-T |
| Macro-THBS1 | CENPU | 3.65E-68 | 3.420419297 | 0.369 | 0.077 | 7.18E-64 | Lep-T |
| Macro-THBS1 | NCAPG2 | 1.15E-46 | 3.165668198 | 0.301 | 0.077 | 2.26E-42 | Lep-T |
| Macro-THBS1 | CCL19 | 1.81E-69 | 2.950636006 | 0.376 | 0.073 | 3.55E-65 | Lep-T |
| Macro-THBS1 | ALDOA | 9.89E-24 | 2.604826147 | 0.443 | 0.253 | 1.95E-19 | Lep-T |
| Macro-THBS1 | CCDC138 | 2.45E-30 | 2.602960417 | 0.417 | 0.2 | 4.82E-26 | Lep-T |
| Macro-THBS1 | UST | 3.67E-27 | 2.220368967 | 0.426 | 0.212 | 7.21E-23 | Lep-T |
| Macro-THBS1 | ASTL | 8.22E-51 | 2.174199856 | 0.354 | 0.091 | 1.62E-46 | Lep-T |
| Macro-THBS1 | RBFOX2 | 4.48E-31 | 2.030939294 | 0.622 | 0.358 | 8.82E-27 | Lep-T |
| Macro-THBS1 | HIST2H2AC | 1.55E-44 | 2.009228021 | 0.29 | 0.067 | 3.06E-40 | Lep-T |
| Macro-THBS1 | DNAI2 | 1.29E-36 | 1.889322427 | 0.371 | 0.128 | 2.53E-32 | Lep-T |
| Macro-THBS1 | HLA-DQB2 | 2.38E-28 | 1.800418446 | 0.317 | 0.115 | 4.68E-24 | Lep-T |
| Macro-THBS1 | TVP23A | 4.17E-23 | 1.706567136 | 0.308 | 0.126 | 8.21E-19 | Lep-T |
| Macro-THBS1 | KSR1 | 9.63E-18 | 1.681212854 | 0.338 | 0.168 | 1.90E-13 | Lep-T |
| Macro-THBS1 | FSCN1 | 1.50E-22 | 1.660336315 | 0.338 | 0.151 | 2.96E-18 | Lep-T |
| Macro-THBS1 | RIPOR3 | 8.34E-24 | 1.575685537 | 0.356 | 0.16 | 1.64E-19 | Lep-T |
| Macro-THBS1 | PRKAR2B | 4.28E-28 | 1.572162985 | 0.286 | 0.095 | 8.42E-24 | Lep-T |
| Macro-THBS1 | ZFP36 | 1.61E-20 | 1.542561077 | 0.965 | 0.885 | 3.17E-16 | Lep-T |
| Macro-THBS1 | BEST1 | 2.04E-19 | 1.53636416 | 0.354 | 0.181 | 4.02E-15 | Lep-T |
| Macro-THBS1 | MTHFD1L | 1.78E-17 | 1.530144115 | 0.295 | 0.134 | 3.51E-13 | Lep-T |
| Macro-THBS1 | CXCL14 | 8.00E-33 | 1.529662922 | 0.258 | 0.067 | 1.57E-28 | Lep-T |
| Macro-THBS1 | CD74 | 9.37E-56 | 1.528747056 | 0.991 | 0.993 | 1.84E-51 | Lep-T |
| Macro-THBS1 | AFDN | 8.85E-37 | 1.502091575 | 0.367 | 0.121 | 1.74E-32 | Lep-T |
| Macro-THBS1 | POGLUT1 | 1.37E-18 | 1.46593136 | 0.269 | 0.112 | 2.70E-14 | Lep-T |

| | | | | | | | |
|---|---|---|---|---|---|---|---|
| Macro-THBS1 | VPS13D | 7.18E-10 | 1.464501478 | 0.474 | 0.36 | 1.41E-05 | Lep-T |
| Macro-THBS1 | POC1B | 1.73E-07 | 1.459178773 | 0.293 | 0.192 | 0.003407534 | Lep-T |
| Macro-THBS1 | INPP5A | 4.83E-24 | 1.444201275 | 0.563 | 0.384 | 9.51E-20 | Lep-T |
| Macro-THBS1 | CLEC16A | 1.35E-12 | 1.424301171 | 0.38 | 0.23 | 2.66E-08 | Lep-T |
| Macro-THBS1 | CLIC2 | 2.15E-26 | 1.36382285 | 0.563 | 0.319 | 4.24E-22 | Lep-T |
| Macro-THBS1 | RTN1 | 2.74E-38 | 1.358062148 | 0.576 | 0.275 | 5.39E-34 | Lep-T |
| Macro-THBS1 | HLA-DQA2 | 1.50E-51 | 1.350362912 | 0.79 | 0.443 | 2.96E-47 | Lep-T |
| Macro-THBS1 | SERINC5 | 6.59E-10 | 1.337403769 | 0.585 | 0.509 | 1.30E-05 | Lep-T |
| Macro-THBS1 | ARHGAP26 | 2.92E-09 | 1.31708334 | 0.762 | 0.818 | 5.75E-05 | Lep-T |
| Macro-THBS1 | SLC25A51 | 4.78E-18 | 1.313688726 | 0.347 | 0.17 | 9.41E-14 | Lep-T |
| Macro-THBS1 | RAB7B | 2.21E-15 | 1.301902489 | 0.328 | 0.165 | 4.35E-11 | Lep-T |
| Macro-THBS1 | USP36 | 4.48E-21 | 1.298411739 | 0.653 | 0.531 | 8.81E-17 | Lep-T |
| Macro-THBS1 | RAB24 | 8.91E-08 | 1.289516133 | 0.286 | 0.181 | 0.001754757 | Lep-T |
| Macro-THBS1 | LAMP3 | 1.06E-19 | 1.246141779 | 0.314 | 0.137 | 2.09E-15 | Lep-T |
| Macro-THBS1 | SPN | 2.24E-19 | 1.243168559 | 0.255 | 0.098 | 4.41E-15 | Lep-T |
| Macro-THBS1 | FAM117B | 7.94E-13 | 1.232956138 | 0.26 | 0.122 | 1.56E-08 | Lep-T |
| Macro-THBS1 | DMKN | 9.26E-33 | 1.225626715 | 0.275 | 0.075 | 1.82E-28 | Lep-T |
| Macro-THBS1 | CASS4 | 3.57E-26 | 1.222009416 | 0.31 | 0.112 | 7.02E-22 | Lep-T |
| Macro-THBS1 | APOBEC3A | 3.14E-14 | 1.220625473 | 0.537 | 0.366 | 6.18E-10 | Lep-T |
| Macro-THBS1 | TTC7B | 2.08E-17 | 1.211559928 | 0.301 | 0.137 | 4.08E-13 | Lep-T |
| Macro-THBS1 | MB21D2 | 2.50E-19 | 1.207965792 | 0.561 | 0.361 | 4.92E-15 | Lep-T |
| Macro-THBS1 | FARS2 | 1.79E-13 | 1.162472575 | 0.279 | 0.137 | 3.52E-09 | Lep-T |
| Macro-THBS1 | CBFA2T3 | 3.02E-15 | 1.162395022 | 0.255 | 0.109 | 5.95E-11 | Lep-T |
| Macro-THBS1 | ADD1 | 3.77E-12 | 1.134830725 | 0.448 | 0.285 | 7.42E-08 | Lep-T |
| Macro-THBS1 | CDC42EP2 | 9.76E-25 | 1.134743952 | 0.642 | 0.387 | 1.92E-20 | Lep-T |
| Macro-THBS1 | SIDT2 | 2.03E-15 | 1.132042907 | 0.341 | 0.177 | 4.00E-11 | Lep-T |
| Macro-THBS1 | U2AF1 | 5.82E-17 | 1.130646571 | 0.277 | 0.119 | 1.15E-12 | Lep-T |
| Macro-THBS1 | SLC9A7 | 2.31E-07 | 1.127912566 | 0.275 | 0.169 | 0.004553942 | Lep-T |
| Macro-THBS1 | SMAD3 | 9.78E-10 | 1.124513535 | 0.496 | 0.354 | 1.93E-05 | Lep-T |
| Macro-THBS1 | FRY | 1.31E-09 | 1.124391577 | 0.262 | 0.144 | 2.57E-05 | Lep-T |
| Macro-THBS1 | SYAP1 | 1.78E-21 | 1.110298456 | 0.723 | 0.606 | 3.51E-17 | Lep-T |
| Macro-THBS1 | SHB | 1.97E-16 | 1.10136859 | 0.496 | 0.313 | 3.88E-12 | Lep-T |
| Macro-THBS1 | NFATC1 | 1.47E-19 | 1.097458007 | 0.493 | 0.292 | 2.89E-15 | Lep-T |
| Macro-THBS1 | SULF2 | 5.95E-23 | 1.096604234 | 0.662 | 0.463 | 1.17E-18 | Lep-T |
| Macro-THBS1 | TNF | 3.28E-07 | 1.094554176 | 0.443 | 0.333 | 0.00644959 | Lep-T |
| Macro-THBS1 | FBXO34 | 1.26E-07 | 1.092857623 | 0.533 | 0.464 | 0.002478851 | Lep-T |
| Macro-THBS1 | DNAH17 | 2.70E-11 | 1.09214678 | 0.323 | 0.183 | 5.31E-07 | Lep-T |
| Macro-THBS1 | MAP4K3 | 2.95E-09 | 1.091400377 | 0.517 | 0.402 | 5.82E-05 | Lep-T |
| Macro-THBS1 | CLEC10A | 3.85E-30 | 1.088610359 | 0.659 | 0.422 | 7.57E-26 | Lep-T |
| Macro-THBS1 | BCL3 | 1.99E-19 | 1.088590251 | 0.686 | 0.486 | 3.91E-15 | Lep-T |
| Macro-THBS1 | SEMA4A | 1.53E-15 | 1.088571718 | 0.539 | 0.367 | 3.01E-11 | Lep-T |
| Macro-THBS1 | CHAF1A | 7.81E-12 | 1.070039357 | 0.255 | 0.125 | 1.54E-07 | Lep-T |
| Macro-THBS1 | MAD1L1 | 9.74E-18 | 1.060788517 | 0.52 | 0.323 | 1.92E-13 | Lep-T |
| Macro-THBS1 | PDE2A | 1.97E-12 | 1.054467866 | 0.273 | 0.136 | 3.88E-08 | Lep-T |
| Macro-THBS1 | ESYT2 | 3.70E-19 | 1.047853106 | 0.721 | 0.615 | 7.29E-15 | Lep-T |
| Macro-THBS1 | ZNF331 | 2.91E-24 | 1.045390503 | 0.819 | 0.744 | 5.73E-20 | Lep-T |
| Macro-THBS1 | NFKBID | 6.34E-22 | 1.044992246 | 0.712 | 0.525 | 1.25E-17 | Lep-T |
| Macro-THBS1 | INSIG1 | 2.87E-18 | 1.042147753 | 0.854 | 0.727 | 5.65E-14 | Lep-T |
| Macro-THBS1 | TNFRSF21 | 1.02E-14 | 1.035727203 | 0.266 | 0.125 | 2.01E-10 | Lep-T |
| Macro-THBS1 | UBASH3B | 1.21E-06 | 1.035447637 | 0.498 | 0.417 | 0.023732125 | Lep-T |
| Macro-THBS1 | DCN | 5.40E-16 | 1.033244441 | 0.277 | 0.12 | 1.06E-11 | Lep-T |
| Macro-THBS1 | GLUD1 | 7.26E-20 | 1.029338607 | 0.64 | 0.422 | 1.43E-15 | Lep-T |
| Macro-THBS1 | SPDYA | 1.72E-09 | 1.026153212 | 0.384 | 0.259 | 3.38E-05 | Lep-T |
| Macro-THBS1 | RCOR1 | 8.85E-15 | 1.025859703 | 0.729 | 0.636 | 1.74E-10 | Lep-T |
| Macro-THBS1 | SEPTIN11 | 5.28E-13 | 1.018386284 | 0.367 | 0.212 | 1.04E-08 | Lep-T |

| | | | | | | | |
|---|---|---|---|---|---|---|---|
| Macro-THBS1 | NRARP | 7.02E-24 | 1.01623471 | 0.469 | 0.237 | 1.38E-19 | Lep-T |
| Macro-THBS1 | INPP5K | 9.96E-11 | 1.0129316 | 0.253 | 0.13 | 1.96E-06 | Lep-T |
| Macro-THBS1 | NMD3 | 9.21E-11 | 1.009192707 | 0.275 | 0.141 | 1.81E-06 | Lep-T |
| Macro-THBS1 | CYTH1 | 3.16E-18 | 1.002220674 | 0.858 | 0.81 | 6.22E-14 | Lep-T |
| Macro-THBS1 | FOLR3 | 8.89E-148 | 5.603617867 | 0.458 | 0.046 | 1.75E-143 | Mari |
| Macro-THBS1 | CCL4L2 | 6.14E-37 | 2.698633436 | 0.657 | 0.427 | 1.21E-32 | Mari |
| Macro-THBS1 | S100A81 | 2.55E-132 | 2.361056094 | 0.968 | 0.623 | 5.03E-128 | Mari |
| Macro-THBS1 | CCL3L1 | 5.90E-33 | 2.203211811 | 0.753 | 0.581 | 1.16E-28 | Mari |
| Macro-THBS1 | CASP5 | 3.26E-34 | 2.115816968 | 0.281 | 0.09 | 6.42E-30 | Mari |
| Macro-THBS1 | S100A12 | 2.73E-45 | 2.070545738 | 0.44 | 0.161 | 5.38E-41 | Mari |
| Macro-THBS1 | VCAN | 1.91E-53 | 2.06209426 | 0.604 | 0.283 | 3.77E-49 | Mari |
| Macro-THBS1 | CCL2 | 2.20E-10 | 1.940634857 | 0.311 | 0.2 | 4.34E-06 | Mari |
| Macro-THBS1 | S100A9 | 5.71E-98 | 1.926483133 | 0.987 | 0.82 | 1.12E-93 | Mari |
| Macro-THBS1 | PLAC8 | 2.05E-27 | 1.865456758 | 0.308 | 0.123 | 4.03E-23 | Mari |
| Macro-THBS1 | IER2 | 3.65E-63 | 1.835070062 | 0.908 | 0.73 | 7.19E-59 | Mari |
| Macro-THBS1 | KRT161 | 8.30E-21 | 1.741610209 | 0.272 | 0.121 | 1.63E-16 | Mari |
| Macro-THBS1 | CD691 | 6.78E-47 | 1.731953355 | 0.536 | 0.242 | 1.33E-42 | Mari |
| Macro-THBS1 | MNDA | 6.35E-55 | 1.638399975 | 0.813 | 0.567 | 1.25E-50 | Mari |
| Macro-THBS1 | ICAM3 | 6.82E-17 | 1.631989022 | 0.253 | 0.116 | 1.34E-12 | Mari |
| Macro-THBS1 | FCGR1B1 | 6.85E-45 | 1.624123597 | 0.517 | 0.234 | 1.35E-40 | Mari |
| Macro-THBS1 | CD14 | 3.24E-73 | 1.56852244 | 0.928 | 0.755 | 6.37E-69 | Mari |
| Macro-THBS1 | MSRB11 | 2.07E-23 | 1.540771684 | 0.351 | 0.17 | 4.08E-19 | Mari |
| Macro-THBS1 | NRG1 | 4.41E-20 | 1.525223495 | 0.255 | 0.106 | 8.68E-16 | Mari |
| Macro-THBS1 | MEF2C | 7.68E-35 | 1.517718615 | 0.609 | 0.4 | 1.51E-30 | Mari |
| Macro-THBS1 | CCR11 | 1.74E-25 | 1.516346029 | 0.379 | 0.186 | 3.42E-21 | Mari |
| Macro-THBS1 | IFI44L | 1.07E-28 | 1.507979043 | 0.332 | 0.134 | 2.10E-24 | Mari |
| Macro-THBS1 | CORO1A | 3.47E-59 | 1.49050727 | 0.783 | 0.533 | 6.83E-55 | Mari |
| Macro-THBS1 | VAMP51 | 6.76E-84 | 1.485983349 | 0.877 | 0.593 | 1.33E-79 | Mari |
| Macro-THBS1 | IER5L1 | 3.76E-26 | 1.477190054 | 0.457 | 0.247 | 7.40E-22 | Mari |
| Macro-THBS1 | FCGR1A | 6.11E-68 | 1.464218876 | 0.8 | 0.485 | 1.20E-63 | Mari |
| Macro-THBS1 | SNHG9 | 2.74E-17 | 1.46105196 | 0.258 | 0.12 | 5.38E-13 | Mari |
| Macro-THBS1 | LILRA2 | 2.55E-14 | 1.443302107 | 0.26 | 0.133 | 5.02E-10 | Mari |
| Macro-THBS1 | RASGRP2 | 6.11E-18 | 1.429206325 | 0.275 | 0.129 | 1.20E-13 | Mari |
| Macro-THBS1 | POLE4 | 3.59E-43 | 1.423401783 | 0.619 | 0.352 | 7.06E-39 | Mari |
| Macro-THBS1 | DDX60L | 5.29E-31 | 1.423209975 | 0.558 | 0.316 | 1.04E-26 | Mari |
| Macro-THBS1 | INHBA-AS1 | 6.55E-16 | 1.41810413 | 0.304 | 0.161 | 1.29E-11 | Mari |
| Macro-THBS1 | FOS | 6.79E-60 | 1.402872102 | 0.915 | 0.789 | 1.34E-55 | Mari |
| Macro-THBS1 | XRCC4 | 9.75E-19 | 1.402294496 | 0.306 | 0.15 | 1.92E-14 | Mari |
| Macro-THBS1 | S100A4 | 7.03E-83 | 1.38754627 | 0.962 | 0.914 | 1.38E-78 | Mari |
| Macro-THBS1 | CDKN2D | 4.77E-18 | 1.38406128 | 0.285 | 0.136 | 9.39E-14 | Mari |
| Macro-THBS1 | MSRB2 | 2.17E-25 | 1.372730651 | 0.458 | 0.267 | 4.27E-21 | Mari |
| Macro-THBS1 | CALHM61 | 4.46E-76 | 1.370158646 | 0.915 | 0.674 | 8.77E-72 | Mari |
| Macro-THBS1 | TMEM173 | 5.63E-18 | 1.368580695 | 0.332 | 0.178 | 1.11E-13 | Mari |
| Macro-THBS1 | TMEM170B | 7.76E-12 | 1.367600376 | 0.302 | 0.187 | 1.53E-07 | Mari |
| Macro-THBS1 | CRELD2 | 3.52E-14 | 1.364173058 | 0.3 | 0.17 | 6.93E-10 | Mari |
| Macro-THBS1 | HRH21 | 3.95E-24 | 1.346675537 | 0.391 | 0.196 | 7.77E-20 | Mari |
| Macro-THBS1 | LILRA5 | 1.54E-17 | 1.334618087 | 0.357 | 0.202 | 3.02E-13 | Mari |
| Macro-THBS1 | MALSU1 | 3.90E-12 | 1.334027415 | 0.251 | 0.138 | 7.68E-08 | Mari |
| Macro-THBS1 | SCO21 | 1.20E-51 | 1.329205679 | 0.743 | 0.44 | 2.37E-47 | Mari |
| Macro-THBS1 | IFITM3 | 2.80E-53 | 1.325355374 | 0.909 | 0.796 | 5.51E-49 | Mari |
| Macro-THBS1 | MRPS18C | 9.81E-35 | 1.321879727 | 0.504 | 0.262 | 1.93E-30 | Mari |
| Macro-THBS1 | SATB1-AS11 | 8.68E-18 | 1.321791184 | 0.381 | 0.215 | 1.71E-13 | Mari |
| Macro-THBS1 | PRELID11 | 7.67E-83 | 1.320833214 | 0.945 | 0.776 | 1.51E-78 | Mari |
| Macro-THBS1 | SMIM15 | 1.85E-17 | 1.315514528 | 0.296 | 0.149 | 3.64E-13 | Mari |
| Macro-THBS1 | STK38L | 5.86E-12 | 1.313112441 | 0.447 | 0.322 | 1.15E-07 | Mari |

| | | | | | | | |
|---|---|---|---|---|---|---|---|
| Macro-THBS1 | GALK1 | 1.39E-21 | 1.311525998 | 0.379 | 0.203 | 2.74E-17 | Mari |
| Macro-THBS1 | TNFSF101 | 8.25E-54 | 1.308799395 | 0.728 | 0.4 | 1.62E-49 | Mari |
| Macro-THBS1 | SHISA5 | 3.94E-15 | 1.308712022 | 0.304 | 0.164 | 7.76E-11 | Mari |
| Macro-THBS1 | SCIMP | 2.34E-23 | 1.29962161 | 0.442 | 0.248 | 4.60E-19 | Mari |
| Macro-THBS1 | FAM89B1 | 2.04E-22 | 1.293547142 | 0.453 | 0.265 | 4.01E-18 | Mari |
| Macro-THBS1 | NUDT16 | 1.73E-21 | 1.292794608 | 0.398 | 0.22 | 3.41E-17 | Mari |
| Macro-THBS1 | GIMAP41 | 4.41E-31 | 1.292645528 | 0.517 | 0.286 | 8.68E-27 | Mari |
| Macro-THBS1 | 46092 | 6.31E-37 | 1.289039772 | 0.606 | 0.343 | 1.24E-32 | Mari |
| Macro-THBS1 | P2RY131 | 2.74E-16 | 1.28838626 | 0.302 | 0.157 | 5.39E-12 | Mari |
| Macro-THBS1 | GCA1 | 4.85E-30 | 1.282507797 | 0.664 | 0.467 | 9.54E-26 | Mari |
| Macro-THBS1 | SMIM25 | 1.75E-23 | 1.27818875 | 0.521 | 0.326 | 3.45E-19 | Mari |
| Macro-THBS1 | FCGR3A | 2.38E-36 | 1.27721123 | 0.725 | 0.58 | 4.68E-32 | Mari |
| Macro-THBS1 | VPS9D11 | 1.72E-12 | 1.268861553 | 0.277 | 0.156 | 3.38E-08 | Mari |
| Macro-THBS1 | VPS25 | 2.59E-15 | 1.265999434 | 0.3 | 0.161 | 5.09E-11 | Mari |
| Macro-THBS1 | CYBC1 | 1.54E-26 | 1.264386296 | 0.5 | 0.299 | 3.02E-22 | Mari |
| Macro-THBS1 | COA4 | 7.17E-18 | 1.262881321 | 0.353 | 0.197 | 1.41E-13 | Mari |
| Macro-THBS1 | PSMB91 | 5.66E-80 | 1.252927808 | 0.894 | 0.689 | 1.11E-75 | Mari |
| Macro-THBS1 | TUBA1A | 5.53E-46 | 1.248733068 | 0.813 | 0.601 | 1.09E-41 | Mari |
| Macro-THBS1 | SULT1A1 | 5.23E-10 | 1.248400911 | 0.258 | 0.154 | 1.03E-05 | Mari |
| Macro-THBS1 | LMNB11 | 4.35E-17 | 1.246963503 | 0.328 | 0.173 | 8.57E-13 | Mari |
| Macro-THBS1 | TIMM8B | 2.84E-41 | 1.244917969 | 0.649 | 0.386 | 5.60E-37 | Mari |
| Macro-THBS1 | MRPL23 | 9.49E-27 | 1.231845447 | 0.426 | 0.216 | 1.87E-22 | Mari |
| Macro-THBS1 | NMI | 3.47E-23 | 1.227948639 | 0.485 | 0.29 | 6.82E-19 | Mari |
| Macro-THBS1 | PSME21 | 2.68E-73 | 1.215478788 | 0.911 | 0.726 | 5.28E-69 | Mari |
| Macro-THBS1 | IFI351 | 2.43E-23 | 1.211025731 | 0.442 | 0.246 | 4.79E-19 | Mari |
| Macro-THBS1 | CCDC167 | 8.55E-15 | 1.208769882 | 0.257 | 0.125 | 1.68E-10 | Mari |
| Macro-THBS1 | IFI30 | 1.19E-112 | 1.202182465 | 0.996 | 0.964 | 2.34E-108 | Mari |
| Macro-THBS1 | DUSP23 | 7.74E-26 | 1.200884597 | 0.489 | 0.291 | 1.52E-21 | Mari |
| Macro-THBS1 | ZDHHC12 | 5.51E-14 | 1.200435625 | 0.304 | 0.173 | 1.09E-09 | Mari |
| Macro-THBS1 | TLR81 | 3.29E-14 | 1.199910206 | 0.306 | 0.169 | 6.47E-10 | Mari |
| Macro-THBS1 | MRPL27 | 4.49E-16 | 1.195953261 | 0.338 | 0.188 | 8.84E-12 | Mari |
| Macro-THBS1 | ARHGDIB | 5.41E-61 | 1.195263924 | 0.881 | 0.717 | 1.06E-56 | Mari |
| Macro-THBS1 | LRRC251 | 1.62E-15 | 1.191288463 | 0.357 | 0.211 | 3.19E-11 | Mari |
| Macro-THBS1 | MRPL3 | 2.13E-13 | 1.190970859 | 0.321 | 0.188 | 4.18E-09 | Mari |
| Macro-THBS1 | RAC2 | 2.42E-31 | 1.176683034 | 0.583 | 0.339 | 4.76E-27 | Mari |
| Macro-THBS1 | CXCL101 | 3.17E-22 | 1.17278351 | 0.617 | 0.403 | 6.23E-18 | Mari |
| Macro-THBS1 | PLAAT41 | 7.57E-47 | 1.172571621 | 0.74 | 0.464 | 1.49E-42 | Mari |
| Macro-THBS1 | OSM1 | 1.09E-21 | 1.172080702 | 0.5 | 0.311 | 2.14E-17 | Mari |
| Macro-THBS1 | SVBP | 2.44E-17 | 1.169052141 | 0.362 | 0.202 | 4.81E-13 | Mari |
| Macro-THBS1 | MTLN | 1.78E-15 | 1.166782352 | 0.323 | 0.178 | 3.51E-11 | Mari |
| Macro-THBS1 | TXNDC17 | 2.63E-37 | 1.162395959 | 0.645 | 0.436 | 5.18E-33 | Mari |
| Macro-THBS1 | NDUFAF8 | 8.97E-21 | 1.157733658 | 0.398 | 0.221 | 1.77E-16 | Mari |
| Macro-THBS1 | AIF1 | 2.79E-85 | 1.149608136 | 0.985 | 0.943 | 5.50E-81 | Mari |
| Macro-THBS1 | COMMD8 | 3.57E-15 | 1.149013683 | 0.377 | 0.237 | 7.02E-11 | Mari |
| Macro-THBS1 | MRPS23 | 1.96E-14 | 1.14756208 | 0.308 | 0.173 | 3.85E-10 | Mari |
| Macro-THBS1 | BORCS8 | 5.64E-10 | 1.145227896 | 0.251 | 0.148 | 1.11E-05 | Mari |
| Macro-THBS1 | NDUFB7 | 6.87E-44 | 1.142860764 | 0.745 | 0.515 | 1.35E-39 | Mari |
| Macro-THBS1 | S100A6 | 2.66E-116 | 1.142467914 | 0.991 | 0.97 | 5.24E-112 | Mari |
| Macro-THBS1 | CDA | 5.29E-09 | 1.138783471 | 0.258 | 0.159 | 0.00010407 | Mari |
| Macro-THBS1 | NME1 | 7.11E-18 | 1.137381563 | 0.445 | 0.283 | 1.40E-13 | Mari |
| Macro-THBS1 | MRPS11 | 1.39E-12 | 1.136581997 | 0.296 | 0.172 | 2.73E-08 | Mari |
| Macro-THBS1 | MRPL28 | 3.07E-19 | 1.133540356 | 0.396 | 0.225 | 6.05E-15 | Mari |
| Macro-THBS1 | TYMP1 | 9.52E-94 | 1.132060217 | 0.985 | 0.917 | 1.87E-89 | Mari |
| Macro-THBS1 | RGS18 | 6.05E-11 | 1.129795701 | 0.253 | 0.145 | 1.19E-06 | Mari |
| Macro-THBS1 | BCL7C | 2.06E-13 | 1.129281753 | 0.362 | 0.229 | 4.05E-09 | Mari |

| | | | | | | | |
|---|---|---|---|---|---|---|---|
| Macro-THBS1 | TTC33 | 5.80E-10 | 1.128551201 | 0.325 | 0.214 | 1.14E-05 | Mari |
| Macro-THBS1 | STXBP2 | 9.40E-27 | 1.125180869 | 0.675 | 0.491 | 1.85E-22 | Mari |
| Macro-THBS1 | PLP2 | 2.15E-25 | 1.123733933 | 0.538 | 0.337 | 4.24E-21 | Mari |
| Macro-THBS1 | CD36 | 2.05E-10 | 1.123681719 | 0.372 | 0.263 | 4.03E-06 | Mari |
| Macro-THBS1 | SLC39A10 | 3.90E-09 | 1.123137842 | 0.275 | 0.176 | 7.68E-05 | Mari |
| Macro-THBS1 | CENPX | 1.09E-23 | 1.121341966 | 0.508 | 0.316 | 2.15E-19 | Mari |
| Macro-THBS1 | NDUFC2 | 1.24E-42 | 1.118915483 | 0.742 | 0.518 | 2.45E-38 | Mari |
| Macro-THBS1 | MX1 | 2.87E-16 | 1.117287918 | 0.387 | 0.23 | 5.65E-12 | Mari |
| Macro-THBS1 | LAMTOR1 | 1.44E-41 | 1.113172342 | 0.791 | 0.594 | 2.83E-37 | Mari |
| Macro-THBS1 | TRAPPC1 | 1.30E-28 | 1.113010108 | 0.615 | 0.399 | 2.55E-24 | Mari |
| Macro-THBS1 | ZNF593 | 8.40E-11 | 1.107962184 | 0.279 | 0.168 | 1.65E-06 | Mari |
| Macro-THBS1 | LDLRAD3 | 3.19E-15 | 1.107867726 | 0.321 | 0.175 | 6.28E-11 | Mari |
| Macro-THBS1 | ARL2 | 5.29E-11 | 1.105502451 | 0.323 | 0.21 | 1.04E-06 | Mari |
| Macro-THBS1 | ASCL2 | 1.02E-15 | 1.103042959 | 0.279 | 0.137 | 2.01E-11 | Mari |
| Macro-THBS1 | CRIP11 | 5.57E-39 | 1.101384149 | 0.817 | 0.587 | 1.10E-34 | Mari |
| Macro-THBS1 | CHMP2A | 2.61E-45 | 1.098795376 | 0.77 | 0.533 | 5.13E-41 | Mari |
| Macro-THBS1 | MRPS121 | 7.42E-15 | 1.094143272 | 0.37 | 0.225 | 1.46E-10 | Mari |
| Macro-THBS1 | TREM1 | 2.02E-24 | 1.091734402 | 0.551 | 0.334 | 3.98E-20 | Mari |
| Macro-THBS1 | COA3 | 7.91E-10 | 1.087617826 | 0.257 | 0.157 | 1.56E-05 | Mari |
| Macro-THBS1 | JOSD2 | 3.93E-21 | 1.085289195 | 0.479 | 0.3 | 7.73E-17 | Mari |
| Macro-THBS1 | UQCC2 | 2.58E-19 | 1.082528859 | 0.419 | 0.242 | 5.08E-15 | Mari |
| Macro-THBS1 | B3GAT3 | 5.29E-11 | 1.082099491 | 0.277 | 0.166 | 1.04E-06 | Mari |
| Macro-THBS1 | OAS1 | 1.48E-13 | 1.081816105 | 0.311 | 0.172 | 2.92E-09 | Mari |
| Macro-THBS1 | IFITM1 | 6.73E-25 | 1.079058954 | 0.298 | 0.117 | 1.32E-20 | Mari |
| Macro-THBS1 | S100A71 | 7.87E-13 | 1.076252347 | 0.251 | 0.132 | 1.55E-08 | Mari |
| Macro-THBS1 | LSM6 | 3.42E-23 | 1.07579149 | 0.54 | 0.345 | 6.74E-19 | Mari |
| Macro-THBS1 | MRPL40 | 1.38E-19 | 1.074020098 | 0.434 | 0.257 | 2.71E-15 | Mari |
| Macro-THBS1 | CZIB | 8.19E-09 | 1.070481018 | 0.251 | 0.156 | 0.00016127 | Mari |
| Macro-THBS1 | FAM207A | 3.56E-08 | 1.069525014 | 0.268 | 0.176 | 0.000701137 | Mari |
| Macro-THBS1 | SEC61G | 3.50E-53 | 1.068737667 | 0.874 | 0.692 | 6.88E-49 | Mari |
| Macro-THBS1 | THOC7 | 1.20E-18 | 1.067247878 | 0.468 | 0.303 | 2.37E-14 | Mari |
| Macro-THBS1 | CKLF | 4.36E-23 | 1.065591973 | 0.653 | 0.478 | 8.58E-19 | Mari |
| Macro-THBS1 | ZNF511 | 9.10E-16 | 1.06267537 | 0.394 | 0.244 | 1.79E-11 | Mari |
| Macro-THBS1 | RGS19 | 3.24E-20 | 1.061947144 | 0.572 | 0.399 | 6.37E-16 | Mari |
| Macro-THBS1 | DCXR | 4.75E-20 | 1.060659128 | 0.468 | 0.287 | 9.34E-16 | Mari |
| Macro-THBS1 | ANTKMT | 6.91E-18 | 1.058085689 | 0.417 | 0.253 | 1.36E-13 | Mari |
| Macro-THBS1 | MRPL20 | 4.01E-29 | 1.053998131 | 0.645 | 0.455 | 7.90E-25 | Mari |
| Macro-THBS1 | PYURF | 1.21E-17 | 1.051922733 | 0.415 | 0.25 | 2.37E-13 | Mari |
| Macro-THBS1 | GAS6 | 5.82E-13 | 1.049065391 | 0.375 | 0.242 | 1.15E-08 | Mari |
| Macro-THBS1 | MRPL52 | 1.16E-26 | 1.045424636 | 0.628 | 0.453 | 2.27E-22 | Mari |
| Macro-THBS1 | DENND101 | 3.35E-11 | 1.045411478 | 0.357 | 0.234 | 6.59E-07 | Mari |
| Macro-THBS1 | NUDT22 | 5.44E-14 | 1.043932746 | 0.377 | 0.238 | 1.07E-09 | Mari |
| Macro-THBS1 | BOLA3 | 1.71E-10 | 1.041499665 | 0.257 | 0.147 | 3.36E-06 | Mari |
| Macro-THBS1 | NCBP2AS2 | 4.94E-11 | 1.038702297 | 0.251 | 0.141 | 9.72E-07 | Mari |
| Macro-THBS1 | SNX17 | 2.13E-22 | 1.037440514 | 0.511 | 0.325 | 4.20E-18 | Mari |
| Macro-THBS1 | NHP2 | 1.19E-19 | 1.037058143 | 0.447 | 0.267 | 2.34E-15 | Mari |
| Macro-THBS1 | SNRPC | 5.74E-17 | 1.035557843 | 0.391 | 0.232 | 1.13E-12 | Mari |
| Macro-THBS1 | IFI27L2 | 5.08E-28 | 1.034345077 | 0.623 | 0.421 | 1.00E-23 | Mari |
| Macro-THBS1 | SRM | 2.48E-13 | 1.032463416 | 0.349 | 0.216 | 4.88E-09 | Mari |
| Macro-THBS1 | EIF4EBP1 | 4.48E-25 | 1.032072823 | 0.562 | 0.365 | 8.82E-21 | Mari |
| Macro-THBS1 | CLEC5A1 | 1.92E-10 | 1.03139635 | 0.315 | 0.197 | 3.78E-06 | Mari |
| Macro-THBS1 | ANAPC15 | 1.75E-13 | 1.027480695 | 0.338 | 0.203 | 3.44E-09 | Mari |
| Macro-THBS1 | PEX16 | 2.45E-10 | 1.027051224 | 0.264 | 0.154 | 4.82E-06 | Mari |
| Macro-THBS1 | RIPOR21 | 1.79E-14 | 1.026101141 | 0.308 | 0.163 | 3.53E-10 | Mari |
| Macro-THBS1 | BTG2 | 4.35E-32 | 1.026022648 | 0.721 | 0.484 | 8.57E-28 | Mari |

| | | | | | | | |
|---|---|---|---|---|---|---|---|
| Macro-THBS1 | MTX1 | 3.78E-11 | 1.025893865 | 0.294 | 0.177 | 7.44E-07 | Mari |
| Macro-THBS1 | UQCR11 | 2.33E-69 | 1.022346142 | 0.921 | 0.735 | 4.60E-65 | Mari |
| Macro-THBS1 | CIB1 | 9.88E-46 | 1.022080518 | 0.851 | 0.666 | 1.94E-41 | Mari |
| Macro-THBS1 | CYBA | 1.07E-120 | 1.015478364 | 0.992 | 0.971 | 2.10E-116 | Mari |
| Macro-THBS1 | NIFK1 | 2.51E-12 | 1.014723924 | 0.332 | 0.204 | 4.94E-08 | Mari |
| Macro-THBS1 | FCGR2B | 8.02E-16 | 1.013933962 | 0.515 | 0.374 | 1.58E-11 | Mari |
| Macro-THBS1 | ARF5 | 4.07E-32 | 1.013033309 | 0.689 | 0.492 | 8.01E-28 | Mari |
| Macro-THBS1 | TSPO | 3.39E-73 | 1.009042498 | 0.949 | 0.882 | 6.67E-69 | Mari |
| Macro-THBS1 | S100A10 | 5.89E-64 | 1.005579318 | 0.985 | 0.947 | 1.16E-59 | Mari |
| Macro-THBS1 | PYCARD | 3.50E-56 | 1.005482257 | 0.898 | 0.756 | 6.90E-52 | Mari |
| Macro-THBS1 | DHRS4 | 2.50E-12 | 1.003335592 | 0.33 | 0.201 | 4.92E-08 | Mari |
| Macro-THBS1 | ORMDL2 | 1.42E-12 | 1.003064242 | 0.343 | 0.211 | 2.80E-08 | Mari |
| Macro-THBS1 | PFN1 | 1.05E-97 | 1.000594601 | 0.987 | 0.934 | 2.07E-93 | Mari |
| Macro-THBS1 | CARD16 | 1.06E-51 | 1.000449791 | 0.885 | 0.672 | 2.08E-47 | Mari |
| Macro-THBS1 | SLC35F1 | 1.17E-104 | 2.924638255 | 0.489 | 0.106 | 2.30E-100 | Tb |
| Macro-THBS1 | KRT17 | 5.71E-44 | 2.563495776 | 0.288 | 0.075 | 1.12E-39 | Tb |
| Macro-THBS1 | GGT5 | 9.13E-57 | 2.382639746 | 0.34 | 0.086 | 1.80E-52 | Tb |
| Macro-THBS1 | KRT6A | 1.80E-55 | 2.325285891 | 0.312 | 0.07 | 3.54E-51 | Tb |
| Macro-THBS1 | KRT6B | 6.77E-54 | 2.258596194 | 0.28 | 0.055 | 1.33E-49 | Tb |
| Macro-THBS1 | S100A2 | 3.32E-42 | 2.155073172 | 0.316 | 0.095 | 6.54E-38 | Tb |
| Macro-THBS1 | TAF4B | 1.70E-13 | 2.09532362 | 0.293 | 0.173 | 3.35E-09 | Tb |
| Macro-THBS1 | HOMER1 | 2.36E-31 | 1.875639005 | 0.443 | 0.225 | 4.65E-27 | Tb |
| Macro-THBS1 | CXCL9 | 2.75E-66 | 1.83073782 | 0.642 | 0.29 | 5.41E-62 | Tb |
| Macro-THBS1 | MCF2L2 | 8.36E-36 | 1.812044209 | 0.496 | 0.26 | 1.65E-31 | Tb |
| Macro-THBS1 | ERI1 | 2.51E-26 | 1.811340243 | 0.419 | 0.227 | 4.94E-22 | Tb |
| Macro-THBS1 | FN1 | 1.30E-17 | 1.810924554 | 0.343 | 0.193 | 2.56E-13 | Tb |
| Macro-THBS1 | B3GNT5 | 1.05E-60 | 1.801852073 | 0.767 | 0.49 | 2.06E-56 | Tb |
| Macro-THBS1 | APOL4 | 1.23E-33 | 1.754013056 | 0.309 | 0.109 | 2.43E-29 | Tb |
| Macro-THBS1 | S100A7 | 1.45E-20 | 1.737397433 | 0.276 | 0.122 | 2.85E-16 | Tb |
| Macro-THBS1 | KRT14 | 1.83E-50 | 1.641087812 | 0.537 | 0.233 | 3.60E-46 | Tb |
| Macro-THBS1 | IRF4 | 1.82E-28 | 1.635872142 | 0.343 | 0.147 | 3.58E-24 | Tb |
| Macro-THBS1 | GNG7 | 2.18E-16 | 1.628525489 | 0.364 | 0.221 | 4.30E-12 | Tb |
| Macro-THBS1 | KRT16 | 5.15E-46 | 1.623052145 | 0.343 | 0.096 | 1.01E-41 | Tb |
| Macro-THBS1 | FPR2 | 1.40E-18 | 1.579276439 | 0.384 | 0.23 | 2.76E-14 | Tb |
| Macro-THBS1 | THBD | 6.65E-48 | 1.497227253 | 0.578 | 0.291 | 1.31E-43 | Tb |
| Macro-THBS1 | CXCL10 | 6.27E-65 | 1.458626701 | 0.727 | 0.365 | 1.24E-60 | Tb |
| Macro-THBS1 | SAP30 | 6.35E-10 | 1.446350135 | 0.338 | 0.23 | 1.25E-05 | Tb |
| Macro-THBS1 | IL1RAP | 3.21E-18 | 1.385605309 | 0.338 | 0.179 | 6.32E-14 | Tb |
| Macro-THBS1 | N4BP2 | 5.54E-14 | 1.382716114 | 0.3 | 0.17 | 1.09E-09 | Tb |
| Macro-THBS1 | IFIT3 | 4.33E-18 | 1.376156342 | 0.273 | 0.126 | 8.53E-14 | Tb |
| Macro-THBS1 | FBP1 | 1.35E-08 | 1.374872965 | 0.544 | 0.471 | 0.000266122 | Tb |
| Macro-THBS1 | HAVCR2 | 1.41E-26 | 1.371784995 | 0.583 | 0.39 | 2.77E-22 | Tb |
| Macro-THBS1 | NKRF | 9.22E-27 | 1.362687584 | 0.42 | 0.213 | 1.81E-22 | Tb |
| Macro-THBS1 | GBP5 | 3.71E-58 | 1.361614166 | 0.825 | 0.545 | 7.29E-54 | Tb |
| Macro-THBS1 | MT1E | 8.38E-11 | 1.359633394 | 0.285 | 0.175 | 1.65E-06 | Tb |
| Macro-THBS1 | KLF9 | 1.54E-13 | 1.355436525 | 0.305 | 0.177 | 3.03E-09 | Tb |
| Macro-THBS1 | ANKRD22 | 1.47E-48 | 1.344415975 | 0.648 | 0.332 | 2.90E-44 | Tb |
| Macro-THBS1 | ETV7 | 3.37E-17 | 1.339451086 | 0.261 | 0.125 | 6.64E-13 | Tb |
| Macro-THBS1 | IDO1 | 8.64E-42 | 1.337017668 | 0.664 | 0.406 | 1.70E-37 | Tb |
| Macro-THBS1 | IFIT2 | 1.71E-15 | 1.334011997 | 0.312 | 0.171 | 3.37E-11 | Tb |
| Macro-THBS1 | SFN | 4.39E-13 | 1.326202353 | 0.257 | 0.137 | 8.65E-09 | Tb |
| Macro-THBS1 | VDR1 | 1.59E-27 | 1.314856921 | 0.427 | 0.211 | 3.14E-23 | Tb |
| Macro-THBS1 | FLT11 | 1.71E-30 | 1.312642532 | 0.628 | 0.417 | 3.36E-26 | Tb |
| Macro-THBS1 | SPATS2L1 | 4.24E-15 | 1.311671729 | 0.456 | 0.308 | 8.35E-11 | Tb |
| Macro-THBS1 | TXNIP | 1.07E-46 | 1.309465628 | 0.746 | 0.493 | 2.11E-42 | Tb |

| | | | | | | | |
|---|---|---|---|---|---|---|---|
| Macro-THBS1 | NLRC5 | 2.58E-24 | 1.301473962 | 0.4 | 0.205 | 5.08E-20 | Tb |
| Macro-THBS1 | DNAJA41 | 5.23E-27 | 1.299685549 | 0.53 | 0.318 | 1.03E-22 | Tb |
| Macro-THBS1 | TMEM273 | 1.72E-16 | 1.299485349 | 0.281 | 0.141 | 3.39E-12 | Tb |
| Macro-THBS1 | SOCS1 | 7.05E-25 | 1.297190482 | 0.54 | 0.351 | 1.39E-20 | Tb |
| Macro-THBS1 | PPA1 | 6.64E-71 | 1.292164592 | 0.864 | 0.632 | 1.31E-66 | Tb |
| Macro-THBS1 | MRC11 | 3.57E-13 | 1.290496866 | 0.503 | 0.393 | 7.02E-09 | Tb |
| Macro-THBS1 | RHOH | 1.12E-22 | 1.281516777 | 0.533 | 0.358 | 2.20E-18 | Tb |
| Macro-THBS1 | ERAP2 | 3.73E-15 | 1.244678451 | 0.293 | 0.155 | 7.34E-11 | Tb |
| Macro-THBS1 | ZFAND2A | 1.41E-16 | 1.240864033 | 0.652 | 0.523 | 2.77E-12 | Tb |
| Macro-THBS1 | NARS2 | 1.25E-09 | 1.232480737 | 0.259 | 0.158 | 2.45E-05 | Tb |
| Macro-THBS1 | ENPP2 | 6.61E-21 | 1.226923405 | 0.286 | 0.126 | 1.30E-16 | Tb |
| Macro-THBS1 | IER5 | 2.51E-58 | 1.226751797 | 0.878 | 0.694 | 4.95E-54 | Tb |
| Macro-THBS1 | IL3RA | 3.40E-33 | 1.22660723 | 0.645 | 0.413 | 6.69E-29 | Tb |
| Macro-THBS1 | SMAD7 | 1.40E-22 | 1.220766415 | 0.479 | 0.293 | 2.76E-18 | Tb |
| Macro-THBS1 | EYA3 | 2.74E-12 | 1.213341647 | 0.367 | 0.242 | 5.39E-08 | Tb |
| Macro-THBS1 | DUSP4 | 3.57E-08 | 1.211136093 | 0.312 | 0.222 | 0.000703469 | Tb |
| Macro-THBS1 | SKI1 | 1.17E-25 | 1.204628792 | 0.479 | 0.275 | 2.30E-21 | Tb |
| Macro-THBS1 | CD69 | 2.83E-46 | 1.202992 | 0.542 | 0.233 | 5.58E-42 | Tb |
| Macro-THBS1 | PDSS1 | 9.68E-09 | 1.197229681 | 0.316 | 0.219 | 0.00019059 | Tb |
| Macro-THBS1 | HSPA1B | 1.04E-86 | 1.197203383 | 0.959 | 0.723 | 2.05E-82 | Tb |
| Macro-THBS1 | LDLR1 | 1.51E-12 | 1.192865853 | 0.346 | 0.21 | 2.97E-08 | Tb |
| Macro-THBS1 | FKBP5 | 9.62E-23 | 1.184005374 | 0.648 | 0.486 | 1.89E-18 | Tb |
| Macro-THBS1 | TNFSF10 | 3.66E-54 | 1.183640874 | 0.722 | 0.393 | 7.21E-50 | Tb |
| Macro-THBS1 | GZF1 | 3.16E-19 | 1.183010472 | 0.388 | 0.216 | 6.22E-15 | Tb |
| Macro-THBS1 | MAML3 | 6.59E-25 | 1.18188442 | 0.578 | 0.38 | 1.30E-20 | Tb |
| Macro-THBS1 | SEMA4D | 1.15E-21 | 1.181723971 | 0.463 | 0.281 | 2.27E-17 | Tb |
| Macro-THBS1 | HES1 | 7.75E-17 | 1.172990126 | 0.446 | 0.281 | 1.53E-12 | Tb |
| Macro-THBS1 | EGR2 | 7.09E-14 | 1.165379858 | 0.257 | 0.132 | 1.40E-09 | Tb |
| Macro-THBS1 | CXCL111 | 9.14E-21 | 1.165336144 | 0.293 | 0.133 | 1.80E-16 | Tb |
| Macro-THBS1 | KLHL6 | 9.46E-24 | 1.163026981 | 0.609 | 0.425 | 1.86E-19 | Tb |
| Macro-THBS1 | CASP8 | 1.56E-10 | 1.157542928 | 0.309 | 0.195 | 3.07E-06 | Tb |
| Macro-THBS1 | CEBPA | 5.14E-17 | 1.155748916 | 0.386 | 0.23 | 1.01E-12 | Tb |
| Macro-THBS1 | NETO2 | 3.76E-09 | 1.149955481 | 0.36 | 0.255 | 7.39E-05 | Tb |
| Macro-THBS1 | BAG3 | 4.96E-42 | 1.148139766 | 0.75 | 0.519 | 9.75E-38 | Tb |
| Macro-THBS1 | P2RY13 | 1.11E-14 | 1.147129161 | 0.293 | 0.156 | 2.19E-10 | Tb |
| Macro-THBS1 | SLC30A1 | 1.73E-10 | 1.146498557 | 0.259 | 0.152 | 3.40E-06 | Tb |
| Macro-THBS1 | HBEGF1 | 8.79E-47 | 1.145404812 | 0.772 | 0.5 | 1.73E-42 | Tb |
| Macro-THBS1 | ZC3HAV1 | 5.52E-39 | 1.140567039 | 0.774 | 0.563 | 1.09E-34 | Tb |
| Macro-THBS1 | CD300LF | 1.74E-12 | 1.136253819 | 0.326 | 0.204 | 3.43E-08 | Tb |
| Macro-THBS1 | MT1X | 9.89E-11 | 1.125330043 | 0.439 | 0.327 | 1.95E-06 | Tb |
| Macro-THBS1 | HDAC4 | 4.82E-24 | 1.117770199 | 0.458 | 0.25 | 9.48E-20 | Tb |
| Macro-THBS1 | ZMYND8 | 8.23E-11 | 1.116999683 | 0.268 | 0.156 | 1.62E-06 | Tb |
| Macro-THBS1 | MIR3945HG | 3.11E-09 | 1.114269008 | 0.503 | 0.419 | 6.12E-05 | Tb |
| Macro-THBS1 | ELMSAN1 | 3.98E-14 | 1.112256665 | 0.35 | 0.212 | 7.84E-10 | Tb |
| Macro-THBS1 | USPL1 | 6.64E-16 | 1.110448249 | 0.444 | 0.287 | 1.31E-11 | Tb |
| Macro-THBS1 | IL15 | 2.65E-17 | 1.108433285 | 0.506 | 0.343 | 5.22E-13 | Tb |
| Macro-THBS1 | SCLT11 | 1.22E-25 | 1.106115723 | 0.592 | 0.384 | 2.41E-21 | Tb |
| Macro-THBS1 | LRRK2 | 6.05E-18 | 1.102191049 | 0.458 | 0.292 | 1.19E-13 | Tb |
| Macro-THBS1 | VASH1 | 2.46E-12 | 1.097288129 | 0.302 | 0.182 | 4.85E-08 | Tb |
| Macro-THBS1 | DNAJB1 | 1.38E-69 | 1.095182053 | 0.952 | 0.827 | 2.71E-65 | Tb |
| Macro-THBS1 | JMY | 6.60E-13 | 1.09352224 | 0.563 | 0.452 | 1.30E-08 | Tb |
| Macro-THBS1 | TENT4B | 4.35E-12 | 1.090919934 | 0.489 | 0.37 | 8.57E-08 | Tb |
| Macro-THBS1 | TRIM25 | 1.75E-31 | 1.088793504 | 0.575 | 0.343 | 3.45E-27 | Tb |
| Macro-THBS1 | DENND1B1 | 3.52E-17 | 1.088791151 | 0.551 | 0.397 | 6.92E-13 | Tb |
| Macro-THBS1 | SLC45A41 | 2.23E-17 | 1.059496621 | 0.388 | 0.221 | 4.39E-13 | Tb |

| | | | | | | | |
|---|---|---|---|---|---|---|---|
| Macro-THBS1 | TYMP | 9.97E-100 | 1.048924632 | 0.988 | 0.914 | 1.96E-95 | Tb |
| Macro-THBS1 | ATP2A2 | 6.04E-15 | 1.046406747 | 0.619 | 0.515 | 1.19E-10 | Tb |
| Macro-THBS1 | KIF2A | 1.62E-10 | 1.045897098 | 0.389 | 0.266 | 3.18E-06 | Tb |
| Macro-THBS1 | BTBD9 | 9.52E-12 | 1.045811259 | 0.612 | 0.498 | 1.87E-07 | Tb |
| Macro-THBS1 | QPCT | 3.04E-12 | 1.041818203 | 0.376 | 0.248 | 5.98E-08 | Tb |
| Macro-THBS1 | CHSY1 | 9.48E-13 | 1.037058032 | 0.532 | 0.394 | 1.87E-08 | Tb |
| Macro-THBS1 | NAF1 | 8.69E-10 | 1.036872844 | 0.549 | 0.464 | 1.71E-05 | Tb |
| Macro-THBS1 | PRDM11 | 1.11E-20 | 1.033743852 | 0.58 | 0.407 | 2.19E-16 | Tb |
| Macro-THBS1 | RNF149 | 3.84E-34 | 1.021560891 | 0.846 | 0.728 | 7.56E-30 | Tb |
| Macro-THBS1 | MTHFR | 2.73E-11 | 1.019407551 | 0.357 | 0.235 | 5.37E-07 | Tb |
| Macro-THBS1 | SNX20 | 2.64E-13 | 1.019055649 | 0.319 | 0.188 | 5.19E-09 | Tb |
| Macro-THBS1 | MARCH3 | 2.23E-07 | 1.016101156 | 0.319 | 0.223 | 0.004387349 | Tb |
| Macro-THBS1 | TRIB1 | 2.43E-16 | 1.013140401 | 0.475 | 0.322 | 4.79E-12 | Tb |
| Macro-THBS1 | PPARD | 1.07E-24 | 1.011319713 | 0.642 | 0.473 | 2.11E-20 | Tb |
| Macro-THBS1 | SNX81 | 1.18E-19 | 1.011291144 | 0.597 | 0.444 | 2.32E-15 | Tb |
| Macro-THBS1 | HAPLN3 | 8.52E-22 | 1.010830841 | 0.444 | 0.253 | 1.68E-17 | Tb |
| Macro-THBS1 | PLAAT4 | 5.49E-39 | 1.009589007 | 0.722 | 0.461 | 1.08E-34 | Tb |
| Macro-THBS1 | CSF2RB | 1.23E-19 | 1.009554827 | 0.458 | 0.278 | 2.43E-15 | Tb |
| **Disease-associated genes in Macro-PPARG** | | | | | | | |
| Macro-PPARG | CCL20 | 5.99E-45 | 2.910503776 | 0.444 | 0.224 | 1.18E-40 | Lep-L |
| Macro-PPARG | HS3ST2 | 1.07E-111 | 2.642867055 | 0.621 | 0.178 | 2.10E-107 | Lep-L |
| Macro-PPARG | CCR7 | 3.83E-32 | 2.469929781 | 0.375 | 0.189 | 7.53E-28 | Lep-L |
| Macro-PPARG | EREG | 2.37E-22 | 2.357298632 | 0.421 | 0.272 | 4.66E-18 | Lep-L |
| Macro-PPARG | TFPI | 7.20E-28 | 2.300278287 | 0.261 | 0.097 | 1.42E-23 | Lep-L |
| Macro-PPARG | SLC18B1 | 7.16E-24 | 2.109584967 | 0.267 | 0.117 | 1.41E-19 | Lep-L |
| Macro-PPARG | CYP27A1 | 2.39E-77 | 2.083331109 | 0.592 | 0.27 | 4.71E-73 | Lep-L |
| Macro-PPARG | TRIP10 | 2.84E-32 | 2.077914893 | 0.31 | 0.128 | 5.59E-28 | Lep-L |
| Macro-PPARG | CXCL8 | 1.01E-148 | 1.998167125 | 0.935 | 0.757 | 2.00E-144 | Lep-L |
| Macro-PPARG | TPRG1 | 3.66E-28 | 1.886176654 | 0.342 | 0.164 | 7.21E-24 | Lep-L |
| Macro-PPARG | OTOA | 5.51E-68 | 1.877620272 | 0.565 | 0.28 | 1.08E-63 | Lep-L |
| Macro-PPARG | KLHL21 | 9.93E-57 | 1.863800365 | 0.511 | 0.239 | 1.95E-52 | Lep-L |
| Macro-PPARG | MIR3142HG | 1.10E-36 | 1.86354288 | 0.409 | 0.192 | 2.16E-32 | Lep-L |
| Macro-PPARG | CDC42EP3 | 1.34E-93 | 1.844967069 | 0.768 | 0.488 | 2.65E-89 | Lep-L |
| Macro-PPARG | BNC2 | 1.49E-29 | 1.842350083 | 0.269 | 0.094 | 2.93E-25 | Lep-L |
| Macro-PPARG | FABP5 | 2.28E-77 | 1.839122522 | 0.775 | 0.648 | 4.48E-73 | Lep-L |
| Macro-PPARG | PPARG | 5.46E-48 | 1.831553338 | 0.624 | 0.373 | 1.08E-43 | Lep-L |
| Macro-PPARG | AQP9 | 1.28E-48 | 1.807602943 | 0.615 | 0.399 | 2.51E-44 | Lep-L |
| Macro-PPARG | CSF1 | 3.60E-08 | 1.77115507 | 0.276 | 0.204 | 0.000707722 | Lep-L |
| Macro-PPARG | MGLL | 9.83E-106 | 1.765816987 | 0.783 | 0.454 | 1.93E-101 | Lep-L |
| Macro-PPARG | CHCHD6 | 6.94E-31 | 1.753350313 | 0.394 | 0.215 | 1.37E-26 | Lep-L |
| Macro-PPARG | MFSD2A | 9.80E-61 | 1.752423706 | 0.576 | 0.31 | 1.93E-56 | Lep-L |
| Macro-PPARG | PLIN2 | 3.25E-148 | 1.749998396 | 0.934 | 0.796 | 6.39E-144 | Lep-L |
| Macro-PPARG | THAP2 | 2.53E-29 | 1.738960444 | 0.515 | 0.37 | 4.97E-25 | Lep-L |
| Macro-PPARG | SPTBN1 | 1.31E-15 | 1.727686592 | 0.263 | 0.146 | 2.58E-11 | Lep-L |
| Macro-PPARG | STMN1 | 3.68E-28 | 1.727201404 | 0.4 | 0.232 | 7.25E-24 | Lep-L |
| Macro-PPARG | NBPF26 | 5.91E-12 | 1.722532352 | 0.266 | 0.175 | 1.16E-07 | Lep-L |
| Macro-PPARG | SLC19A2 | 1.08E-19 | 1.70628862 | 0.504 | 0.396 | 2.12E-15 | Lep-L |
| Macro-PPARG | GPNMB | 1.61E-111 | 1.662399097 | 0.878 | 0.631 | 3.16E-107 | Lep-L |
| Macro-PPARG | SDC4 | 1.98E-27 | 1.650853879 | 0.468 | 0.318 | 3.90E-23 | Lep-L |
| Macro-PPARG | RASGRP3 | 1.75E-58 | 1.648836936 | 0.706 | 0.457 | 3.45E-54 | Lep-L |
| Macro-PPARG | SH3D21 | 1.32E-28 | 1.639176604 | 0.406 | 0.229 | 2.59E-24 | Lep-L |
| Macro-PPARG | CHST7 | 7.36E-37 | 1.590854176 | 0.314 | 0.105 | 1.45E-32 | Lep-L |
| Macro-PPARG | SDS | 6.30E-42 | 1.552881251 | 0.703 | 0.526 | 1.24E-37 | Lep-L |
| Macro-PPARG | PLPP3 | 8.84E-37 | 1.521907959 | 0.546 | 0.322 | 1.74E-32 | Lep-L |
| Macro-PPARG | SMS | 2.78E-87 | 1.521763155 | 0.773 | 0.576 | 5.47E-83 | Lep-L |

| | | | | | | | |
|---|---|---|---|---|---|---|---|
| Macro-PPARG | CXCL3 | 1.69E-73 | 1.514662747 | 0.869 | 0.722 | 3.33E-69 | Lep-L |
| Macro-PPARG | BCL2A1 | 1.07E-119 | 1.511251765 | 0.92 | 0.848 | 2.11E-115 | Lep-L |
| Macro-PPARG | NBPF19 | 1.13E-40 | 1.501145931 | 0.509 | 0.3 | 2.23E-36 | Lep-L |
| Macro-PPARG | DHRS3 | 6.47E-26 | 1.483753587 | 0.319 | 0.152 | 1.27E-21 | Lep-L |
| Macro-PPARG | AP1S3 | 2.82E-13 | 1.458313024 | 0.269 | 0.161 | 5.56E-09 | Lep-L |
| Macro-PPARG | APOC1 | 1.57E-88 | 1.45647314 | 0.905 | 0.663 | 3.09E-84 | Lep-L |
| Macro-PPARG | OPN3 | 5.21E-33 | 1.456049422 | 0.577 | 0.431 | 1.02E-28 | Lep-L |
| Macro-PPARG | MIR155HG | 8.31E-55 | 1.453078224 | 0.717 | 0.494 | 1.64E-50 | Lep-L |
| Macro-PPARG | LIPA | 1.24E-52 | 1.43099427 | 0.681 | 0.506 | 2.45E-48 | Lep-L |
| Macro-PPARG | CMSS1 | 3.83E-14 | 1.428808296 | 0.58 | 0.526 | 7.53E-10 | Lep-L |
| Macro-PPARG | APOE | 2.01E-99 | 1.422209353 | 0.956 | 0.744 | 3.95E-95 | Lep-L |
| Macro-PPARG | DDIT4 | 9.45E-48 | 1.418788563 | 0.709 | 0.535 | 1.86E-43 | Lep-L |
| Macro-PPARG | CCRL2 | 1.45E-23 | 1.414573587 | 0.595 | 0.498 | 2.85E-19 | Lep-L |
| Macro-PPARG | EBI3 | 3.96E-15 | 1.396221172 | 0.251 | 0.136 | 7.79E-11 | Lep-L |
| Macro-PPARG | ZFY | 5.01E-31 | 1.392759161 | 0.258 | 0.077 | 9.87E-27 | Lep-L |
| Macro-PPARG | NFKBIA | 2.94E-124 | 1.388321946 | 0.987 | 0.972 | 5.80E-120 | Lep-L |
| Macro-PPARG | STAM | 9.00E-25 | 1.376509568 | 0.433 | 0.289 | 1.77E-20 | Lep-L |
| Macro-PPARG | KCNE1 | 1.13E-14 | 1.356335558 | 0.356 | 0.244 | 2.22E-10 | Lep-L |
| Macro-PPARG | FABP3 | 1.89E-08 | 1.350256568 | 0.278 | 0.2 | 0.000371377 | Lep-L |
| Macro-PPARG | ZNF703 | 5.42E-13 | 1.327344563 | 0.324 | 0.217 | 1.07E-08 | Lep-L |
| Macro-PPARG | SLC1A3 | 1.01E-51 | 1.317503442 | 0.814 | 0.666 | 1.98E-47 | Lep-L |
| Macro-PPARG | MS4A14 | 3.87E-16 | 1.290635779 | 0.337 | 0.208 | 7.61E-12 | Lep-L |
| Macro-PPARG | G0S2 | 3.98E-35 | 1.28323661 | 0.731 | 0.613 | 7.83E-31 | Lep-L |
| Macro-PPARG | IER3 | 1.02E-80 | 1.27585038 | 0.941 | 0.923 | 2.01E-76 | Lep-L |
| Macro-PPARG | MMP19 | 1.62E-30 | 1.274978577 | 0.686 | 0.533 | 3.18E-26 | Lep-L |
| Macro-PPARG | BCL2 | 4.90E-44 | 1.268227121 | 0.695 | 0.526 | 9.64E-40 | Lep-L |
| Macro-PPARG | TNFAIP6 | 6.97E-09 | 1.266848635 | 0.362 | 0.29 | 0.000137266 | Lep-L |
| Macro-PPARG | CXXC5 | 2.51E-14 | 1.258196231 | 0.347 | 0.244 | 4.94E-10 | Lep-L |
| Macro-PPARG | TESK1 | 8.31E-14 | 1.244752892 | 0.364 | 0.27 | 1.64E-09 | Lep-L |
| Macro-PPARG | SCCPDH | 2.54E-10 | 1.242643332 | 0.286 | 0.207 | 5.01E-06 | Lep-L |
| Macro-PPARG | FEM1C | 6.65E-11 | 1.236266951 | 0.392 | 0.317 | 1.31E-06 | Lep-L |
| Macro-PPARG | PPP1R15B | 9.94E-62 | 1.231918097 | 0.847 | 0.704 | 1.96E-57 | Lep-L |
| Macro-PPARG | PDE4DIP | 1.23E-85 | 1.227073126 | 0.932 | 0.751 | 2.41E-81 | Lep-L |
| Macro-PPARG | DDX3Y | 2.37E-58 | 1.222746892 | 0.641 | 0.307 | 4.66E-54 | Lep-L |
| Macro-PPARG | IL1B | 5.39E-21 | 1.212450074 | 0.711 | 0.626 | 1.06E-16 | Lep-L |
| Macro-PPARG | UPP1 | 1.17E-64 | 1.20724663 | 0.842 | 0.707 | 2.30E-60 | Lep-L |
| Macro-PPARG | TSC22D2 | 7.49E-36 | 1.206303704 | 0.84 | 0.74 | 1.47E-31 | Lep-L |
| Macro-PPARG | KMO | 8.91E-68 | 1.204792284 | 0.817 | 0.607 | 1.75E-63 | Lep-L |
| Macro-PPARG | GBE1 | 3.22E-37 | 1.191440295 | 0.689 | 0.491 | 6.33E-33 | Lep-L |
| Macro-PPARG | MAP7D1 | 2.23E-22 | 1.190812639 | 0.595 | 0.516 | 4.40E-18 | Lep-L |
| Macro-PPARG | PDE8A | 4.31E-35 | 1.186583156 | 0.71 | 0.527 | 8.48E-31 | Lep-L |
| Macro-PPARG | APBB3 | 6.75E-08 | 1.181720442 | 0.38 | 0.324 | 0.001328414 | Lep-L |
| Macro-PPARG | ABCC5 | 7.95E-14 | 1.174417256 | 0.312 | 0.199 | 1.56E-09 | Lep-L |
| Macro-PPARG | ADCY3 | 5.48E-10 | 1.164943912 | 0.42 | 0.342 | 1.08E-05 | Lep-L |
| Macro-PPARG | SMIM3 | 1.04E-28 | 1.163104451 | 0.572 | 0.429 | 2.05E-24 | Lep-L |
| Macro-PPARG | CSTB | 1.28E-61 | 1.162194402 | 0.936 | 0.851 | 2.52E-57 | Lep-L |
| Macro-PPARG | TDP2 | 2.18E-09 | 1.160909091 | 0.327 | 0.253 | 4.29E-05 | Lep-L |
| Macro-PPARG | UBXN7 | 1.01E-10 | 1.156349922 | 0.31 | 0.225 | 1.99E-06 | Lep-L |
| Macro-PPARG | CD52 | 2.95E-10 | 1.152296485 | 0.329 | 0.245 | 5.81E-06 | Lep-L |
| Macro-PPARG | PIK3R5 | 6.66E-30 | 1.146437204 | 0.686 | 0.57 | 1.31E-25 | Lep-L |
| Macro-PPARG | SCD | 5.32E-07 | 1.137512778 | 0.352 | 0.3 | 0.01047568 | Lep-L |
| Macro-PPARG | PTGER2 | 6.53E-08 | 1.136236214 | 0.272 | 0.203 | 0.001286113 | Lep-L |
| Macro-PPARG | NOTCH2NLC | 4.01E-27 | 1.13112524 | 0.567 | 0.406 | 7.90E-23 | Lep-L |
| Macro-PPARG | HIVEP2 | 3.50E-48 | 1.125995719 | 0.78 | 0.614 | 6.88E-44 | Lep-L |
| Macro-PPARG | CD82 | 5.54E-20 | 1.116711052 | 0.452 | 0.325 | 1.09E-15 | Lep-L |

| | | | | | | | |
|---|---|---|---|---|---|---|---|
| Macro-PPARG | RILPL2 | 3.12E-65 | 1.116077164 | 0.83 | 0.723 | 6.13E-61 | Lep-L |
| Macro-PPARG | MIR222HG | 4.62E-17 | 1.114228767 | 0.286 | 0.148 | 9.10E-13 | Lep-L |
| Macro-PPARG | SNHG7 | 4.52E-22 | 1.111514006 | 0.592 | 0.531 | 8.90E-18 | Lep-L |
| Macro-PPARG | SEPHS2 | 2.63E-25 | 1.108112948 | 0.655 | 0.552 | 5.18E-21 | Lep-L |
| Macro-PPARG | PTTG1 | 1.12E-10 | 1.103614923 | 0.322 | 0.232 | 2.20E-06 | Lep-L |
| Macro-PPARG | H2AFX | 3.24E-16 | 1.101474202 | 0.509 | 0.418 | 6.38E-12 | Lep-L |
| Macro-PPARG | B3GNT2 | 2.03E-12 | 1.09173166 | 0.506 | 0.455 | 4.01E-08 | Lep-L |
| Macro-PPARG | SLC26A11 | 3.98E-11 | 1.087452735 | 0.361 | 0.276 | 7.84E-07 | Lep-L |
| Macro-PPARG | ITPKB | 6.52E-25 | 1.080611491 | 0.52 | 0.36 | 1.28E-20 | Lep-L |
| Macro-PPARG | SERPINB9 | 2.05E-44 | 1.078093012 | 0.793 | 0.712 | 4.03E-40 | Lep-L |
| Macro-PPARG | SPSB1 | 1.29E-10 | 1.07552413 | 0.336 | 0.239 | 2.54E-06 | Lep-L |
| Macro-PPARG | CSGALNACT2 | 6.87E-63 | 1.074311642 | 0.832 | 0.731 | 1.35E-58 | Lep-L |
| Macro-PPARG | MAFG | 1.39E-22 | 1.074056473 | 0.493 | 0.367 | 2.73E-18 | Lep-L |
| Macro-PPARG | UGCG | 8.86E-39 | 1.073482639 | 0.766 | 0.635 | 1.74E-34 | Lep-L |
| Macro-PPARG | NBPF10 | 1.12E-09 | 1.069179304 | 0.304 | 0.221 | 2.21E-05 | Lep-L |
| Macro-PPARG | ADD3 | 1.25E-07 | 1.063979305 | 0.317 | 0.252 | 0.002455594 | Lep-L |
| Macro-PPARG | ZNF503 | 3.93E-10 | 1.062656553 | 0.266 | 0.173 | 7.74E-06 | Lep-L |
| Macro-PPARG | CXCR4 | 1.26E-53 | 1.059686157 | 0.85 | 0.732 | 2.48E-49 | Lep-L |
| Macro-PPARG | PI4K2A | 6.06E-20 | 1.053850947 | 0.576 | 0.496 | 1.19E-15 | Lep-L |
| Macro-PPARG | SRPK1 | 7.92E-07 | 1.047532668 | 0.278 | 0.221 | 0.015593389 | Lep-L |
| Macro-PPARG | RB1CC1 | 3.75E-25 | 1.044406669 | 0.606 | 0.511 | 7.39E-21 | Lep-L |
| Macro-PPARG | TTTY14 | 6.84E-26 | 1.041217041 | 0.34 | 0.153 | 1.35E-21 | Lep-L |
| Macro-PPARG | TRAF1 | 1.92E-16 | 1.041056357 | 0.52 | 0.441 | 3.77E-12 | Lep-L |
| Macro-PPARG | CFD | 1.77E-37 | 1.039666574 | 0.781 | 0.732 | 3.49E-33 | Lep-L |
| Macro-PPARG | ALDH1A1 | 9.35E-17 | 1.037159448 | 0.627 | 0.526 | 1.84E-12 | Lep-L |
| Macro-PPARG | SNTB1 | 1.17E-14 | 1.035583892 | 0.467 | 0.365 | 2.30E-10 | Lep-L |
| Macro-PPARG | EIF1B | 5.45E-33 | 1.035144666 | 0.679 | 0.595 | 1.07E-28 | Lep-L |
| Macro-PPARG | SMPDL3A | 1.60E-14 | 1.034104646 | 0.467 | 0.377 | 3.16E-10 | Lep-L |
| Macro-PPARG | THAP9-AS1 | 3.05E-07 | 1.023157053 | 0.355 | 0.306 | 0.005999343 | Lep-L |
| Macro-PPARG | STX3 | 1.03E-14 | 1.016316629 | 0.428 | 0.336 | 2.03E-10 | Lep-L |
| Macro-PPARG | CXCL2 | 4.52E-66 | 1.012484467 | 0.941 | 0.849 | 8.89E-62 | Lep-L |
| Macro-PPARG | GADD45B | 2.16E-33 | 1.011913279 | 0.922 | 0.909 | 4.25E-29 | Lep-L |
| Macro-PPARG | HIPK2 | 9.21E-10 | 1.011237232 | 0.419 | 0.357 | 1.81E-05 | Lep-L |
| Macro-PPARG | PHLDA1 | 2.60E-29 | 1.005615888 | 0.51 | 0.302 | 5.13E-25 | Lep-L |
| Macro-PPARG | PHF1 | 6.53E-08 | 1.005296129 | 0.319 | 0.255 | 0.001285098 | Lep-L |
| Macro-PPARG | TXNRD1 | 5.81E-36 | 1.00363676 | 0.761 | 0.663 | 1.14E-31 | Lep-L |
| Macro-PPARG | EHD1 | 1.31E-25 | 1.001444736 | 0.625 | 0.527 | 2.57E-21 | Lep-L |
| Macro-PPARG | CENPU | 2.69E-47 | 3.886273149 | 0.314 | 0.075 | 5.29E-43 | Lep-T |
| Macro-PPARG | CXCL14 | 4.43E-62 | 3.029282133 | 0.387 | 0.08 | 8.72E-58 | Lep-T |
| Macro-PPARG | FGFR1 | 5.88E-51 | 2.822926555 | 0.492 | 0.168 | 1.16E-46 | Lep-T |
| Macro-PPARG | UBD | 1.39E-11 | 2.500827986 | 0.276 | 0.141 | 2.73E-07 | Lep-T |
| Macro-PPARG | NCAPG2 | 7.26E-14 | 2.377754245 | 0.26 | 0.129 | 1.43E-09 | Lep-T |
| Macro-PPARG | DCN | 5.54E-27 | 2.325792697 | 0.384 | 0.154 | 1.09E-22 | Lep-T |
| Macro-PPARG | PTGDS | 3.43E-24 | 2.151198081 | 0.394 | 0.174 | 6.75E-20 | Lep-T |
| Macro-PPARG | WDR61 | 6.36E-15 | 1.982372544 | 0.41 | 0.225 | 1.25E-10 | Lep-T |
| Macro-PPARG | AFDN | 3.88E-35 | 1.961761659 | 0.456 | 0.182 | 7.64E-31 | Lep-T |
| Macro-PPARG | LPAR6 | 2.22E-22 | 1.936380838 | 0.559 | 0.333 | 4.36E-18 | Lep-T |
| Macro-PPARG | MGP | 3.15E-28 | 1.922283963 | 0.286 | 0.087 | 6.20E-24 | Lep-T |
| Macro-PPARG | UST | 2.74E-24 | 1.919390055 | 0.526 | 0.312 | 5.40E-20 | Lep-T |
| Macro-PPARG | CCL2 | 8.15E-14 | 1.911306204 | 0.464 | 0.278 | 1.60E-09 | Lep-T |
| Macro-PPARG | PID1 | 1.85E-14 | 1.875763528 | 0.469 | 0.324 | 3.65E-10 | Lep-T |
| Macro-PPARG | NAV2 | 3.59E-08 | 1.853109038 | 0.263 | 0.163 | 0.000707406 | Lep-T |
| Macro-PPARG | TVP23A | 5.02E-30 | 1.839174329 | 0.376 | 0.141 | 9.87E-26 | Lep-T |
| Macro-PPARG | GPAT3 | 8.66E-07 | 1.806978715 | 0.43 | 0.35 | 0.017039298 | Lep-T |
| Macro-PPARG | RTN1 | 1.33E-35 | 1.803156323 | 0.567 | 0.282 | 2.61E-31 | Lep-T |

| | | | | | | | |
|---|---|---|---|---|---|---|---|
| Macro-PPARG | IL21R | 1.04E-21 | 1.756601077 | 0.294 | 0.109 | 2.04E-17 | Lep-T |
| Macro-PPARG | PMEPA1 | 4.28E-14 | 1.721494166 | 0.284 | 0.135 | 8.42E-10 | Lep-T |
| Macro-PPARG | AUTS2 | 4.96E-08 | 1.721292878 | 0.253 | 0.153 | 0.000976827 | Lep-T |
| Macro-PPARG | CARMIL1 | 6.28E-07 | 1.709342315 | 0.309 | 0.194 | 0.012361669 | Lep-T |
| Macro-PPARG | KSR1 | 9.56E-11 | 1.651113592 | 0.294 | 0.158 | 1.88E-06 | Lep-T |
| Macro-PPARG | ACVR2A | 1.32E-11 | 1.636639651 | 0.286 | 0.149 | 2.60E-07 | Lep-T |
| Macro-PPARG | MB21D2 | 7.39E-35 | 1.590624789 | 0.686 | 0.414 | 1.46E-30 | Lep-T |
| Macro-PPARG | WDFY2 | 1.67E-10 | 1.549769967 | 0.582 | 0.438 | 3.28E-06 | Lep-T |
| Macro-PPARG | SLC9A9 | 1.04E-32 | 1.531331659 | 0.747 | 0.527 | 2.04E-28 | Lep-T |
| Macro-PPARG | POLA1 | 1.22E-14 | 1.513478554 | 0.294 | 0.138 | 2.39E-10 | Lep-T |
| Macro-PPARG | PDE4D | 5.83E-07 | 1.507013498 | 0.358 | 0.256 | 0.011469529 | Lep-T |
| Macro-PPARG | STAB1 | 9.98E-16 | 1.50487245 | 0.593 | 0.424 | 1.96E-11 | Lep-T |
| Macro-PPARG | U2AF1 | 1.53E-07 | 1.482953259 | 0.265 | 0.155 | 0.00300965 | Lep-T |
| Macro-PPARG | STX17-AS1 | 4.19E-10 | 1.481068226 | 0.302 | 0.17 | 8.24E-06 | Lep-T |
| Macro-PPARG | INPP5A | 9.81E-16 | 1.46259663 | 0.485 | 0.321 | 1.93E-11 | Lep-T |
| Macro-PPARG | RFTN1 | 1.51E-17 | 1.459857037 | 0.544 | 0.373 | 2.98E-13 | Lep-T |
| Macro-PPARG | F13A1 | 3.94E-25 | 1.44740407 | 0.544 | 0.313 | 7.77E-21 | Lep-T |
| Macro-PPARG | TNFRSF4 | 6.15E-12 | 1.40712874 | 0.253 | 0.122 | 1.21E-07 | Lep-T |
| Macro-PPARG | HIVEP3 | 3.11E-07 | 1.392839911 | 0.436 | 0.312 | 0.006114828 | Lep-T |
| Macro-PPARG | FGL2 | 2.80E-40 | 1.39072873 | 0.879 | 0.717 | 5.52E-36 | Lep-T |
| Macro-PPARG | PTPRM | 3.15E-07 | 1.390580193 | 0.371 | 0.262 | 0.006193965 | Lep-T |
| Macro-PPARG | AFF1 | 5.82E-22 | 1.384284524 | 0.863 | 0.762 | 1.15E-17 | Lep-T |
| Macro-PPARG | MMP2 | 3.69E-13 | 1.372068352 | 0.322 | 0.166 | 7.26E-09 | Lep-T |
| Macro-PPARG | MVB12B | 5.48E-12 | 1.360348693 | 0.356 | 0.197 | 1.08E-07 | Lep-T |
| Macro-PPARG | DPYD-AS1 | 2.64E-10 | 1.356310838 | 0.278 | 0.148 | 5.20E-06 | Lep-T |
| Macro-PPARG | PRKAR2B | 1.20E-11 | 1.338464644 | 0.284 | 0.145 | 2.35E-07 | Lep-T |
| Macro-PPARG | COL1A2 | 1.25E-07 | 1.328997828 | 0.268 | 0.156 | 0.002466871 | Lep-T |
| Macro-PPARG | SERINC5 | 2.90E-09 | 1.308082189 | 0.642 | 0.58 | 5.70E-05 | Lep-T |
| Macro-PPARG | DNAJA4 | 1.81E-06 | 1.291819502 | 0.549 | 0.458 | 0.035718196 | Lep-T |
| Macro-PPARG | HSPD1 | 2.05E-09 | 1.272331329 | 0.92 | 0.948 | 4.03E-05 | Lep-T |
| Macro-PPARG | SLC8A1-AS1 | 8.27E-12 | 1.265327764 | 0.296 | 0.152 | 1.63E-07 | Lep-T |
| Macro-PPARG | LGALS2 | 1.48E-07 | 1.250459355 | 0.485 | 0.426 | 0.002921998 | Lep-T |
| Macro-PPARG | PTK2 | 7.33E-08 | 1.243095099 | 0.289 | 0.173 | 0.001443757 | Lep-T |
| Macro-PPARG | CCDC170 | 4.13E-09 | 1.237519019 | 0.332 | 0.201 | 8.12E-05 | Lep-T |
| Macro-PPARG | TCF20 | 1.46E-08 | 1.23462725 | 0.338 | 0.219 | 0.000287586 | Lep-T |
| Macro-PPARG | RGL1 | 1.33E-14 | 1.22260447 | 0.704 | 0.537 | 2.63E-10 | Lep-T |
| Macro-PPARG | ME1 | 5.88E-13 | 1.203277207 | 0.601 | 0.436 | 1.16E-08 | Lep-T |
| Macro-PPARG | NRP1 | 1.53E-15 | 1.193652887 | 0.701 | 0.547 | 3.00E-11 | Lep-T |
| Macro-PPARG | TNFSF15 | 1.67E-07 | 1.183269198 | 0.356 | 0.238 | 0.00327972 | Lep-T |
| Macro-PPARG | TTC7B | 1.35E-10 | 1.156242442 | 0.415 | 0.27 | 2.66E-06 | Lep-T |
| Macro-PPARG | ANPEP | 2.17E-07 | 1.155960177 | 0.505 | 0.395 | 0.004275456 | Lep-T |
| Macro-PPARG | ATR | 1.64E-09 | 1.15410759 | 0.278 | 0.15 | 3.23E-05 | Lep-T |
| Macro-PPARG | LIMA1 | 6.32E-14 | 1.14458393 | 0.273 | 0.122 | 1.24E-09 | Lep-T |
| Macro-PPARG | DENND3 | 2.49E-08 | 1.143758097 | 0.423 | 0.297 | 0.000489571 | Lep-T |
| Macro-PPARG | SESN3 | 4.39E-08 | 1.137082059 | 0.356 | 0.225 | 0.000863598 | Lep-T |
| Macro-PPARG | WWC2 | 1.73E-06 | 1.134519521 | 0.268 | 0.167 | 0.033960168 | Lep-T |
| Macro-PPARG | CCL5 | 2.54E-07 | 1.131945569 | 0.392 | 0.264 | 0.004994816 | Lep-T |
| Macro-PPARG | ABCC1 | 8.14E-14 | 1.130581971 | 0.539 | 0.348 | 1.60E-09 | Lep-T |
| Macro-PPARG | MED13L | 2.03E-24 | 1.125629282 | 0.835 | 0.813 | 3.99E-20 | Lep-T |
| Macro-PPARG | SEPTIN11 | 1.05E-14 | 1.118581636 | 0.528 | 0.362 | 2.08E-10 | Lep-T |
| Macro-PPARG | DUSP5 | 1.98E-09 | 1.113106234 | 0.575 | 0.449 | 3.90E-05 | Lep-T |
| Macro-PPARG | CCDC146 | 7.10E-11 | 1.106544929 | 0.281 | 0.143 | 1.40E-06 | Lep-T |
| Macro-PPARG | CST3 | 1.42E-09 | 1.101406381 | 0.938 | 0.919 | 2.79E-05 | Lep-T |
| Macro-PPARG | GAS2L3 | 1.32E-08 | 1.101205343 | 0.474 | 0.338 | 0.000260317 | Lep-T |
| Macro-PPARG | HSP90AA1 | 1.89E-14 | 1.098709688 | 0.979 | 0.986 | 3.71E-10 | Lep-T |

| | | | | | | | |
|---|---|---|---|---|---|---|---|
| Macro-PPARG | IL3RA | 7.02E-23 | 1.087608359 | 0.606 | 0.354 | 1.38E-18 | Lep-T |
| Macro-PPARG | GLUD1 | 2.80E-10 | 1.079629697 | 0.678 | 0.542 | 5.51E-06 | Lep-T |
| Macro-PPARG | RNASE1 | 3.07E-08 | 1.07778615 | 0.562 | 0.476 | 0.000604128 | Lep-T |
| Macro-PPARG | TIAM1 | 4.44E-09 | 1.075097363 | 0.619 | 0.479 | 8.73E-05 | Lep-T |
| Macro-PPARG | ZDHHC14 | 5.13E-10 | 1.07435931 | 0.418 | 0.259 | 1.01E-05 | Lep-T |
| Macro-PPARG | MAP4K3 | 3.89E-14 | 1.067818214 | 0.655 | 0.5 | 7.67E-10 | Lep-T |
| Macro-PPARG | HLA-DQB2 | 2.80E-09 | 1.066501304 | 0.327 | 0.199 | 5.51E-05 | Lep-T |
| Macro-PPARG | ARHGAP18 | 2.44E-28 | 1.05316982 | 0.812 | 0.704 | 4.81E-24 | Lep-T |
| Macro-PPARG | HLA-DQA2 | 7.64E-15 | 1.05176339 | 0.673 | 0.562 | 1.50E-10 | Lep-T |
| Macro-PPARG | INSIG1 | 6.92E-22 | 1.050907452 | 0.915 | 0.828 | 1.36E-17 | Lep-T |
| Macro-PPARG | FMN1 | 1.18E-07 | 1.04559317 | 0.765 | 0.771 | 0.002314442 | Lep-T |
| Macro-PPARG | TFEC | 2.24E-14 | 1.041857662 | 0.737 | 0.718 | 4.41E-10 | Lep-T |
| Macro-PPARG | HPCAL1 | 9.80E-07 | 1.030417852 | 0.593 | 0.5 | 0.019296184 | Lep-T |
| Macro-PPARG | STARD13 | 1.02E-11 | 1.029901123 | 0.642 | 0.517 | 2.00E-07 | Lep-T |
| Macro-PPARG | ARL14EP | 7.22E-08 | 1.028408043 | 0.268 | 0.151 | 0.001421153 | Lep-T |
| Macro-PPARG | MT-ATP6 | 7.48E-28 | 1.02729656 | 0.941 | 0.971 | 1.47E-23 | Lep-T |
| Macro-PPARG | WWP1 | 1.84E-07 | 1.026175687 | 0.526 | 0.414 | 0.003627734 | Lep-T |
| Macro-PPARG | SNX29 | 2.70E-19 | 1.025722842 | 0.742 | 0.607 | 5.31E-15 | Lep-T |
| Macro-PPARG | OSBPL3 | 2.39E-07 | 1.023598521 | 0.461 | 0.332 | 0.004697284 | Lep-T |
| Macro-PPARG | MRC1 | 1.34E-30 | 1.016745349 | 0.784 | 0.578 | 2.64E-26 | Lep-T |
| Macro-PPARG | PPARGC1B | 1.72E-07 | 1.013887286 | 0.268 | 0.154 | 0.003393233 | Lep-T |
| Macro-PPARG | NR3C1 | 1.89E-13 | 1.002555841 | 0.773 | 0.694 | 3.73E-09 | Lep-T |
| Macro-PPARG | FOLR3 | 2.07E-129 | 6.688780794 | 0.339 | 0.012 | 4.08E-125 | Mari |
| Macro-PPARG | MMP121 | 2.19E-19 | 3.572764232 | 0.257 | 0.092 | 4.31E-15 | Mari |
| Macro-PPARG | ANK3 | 1.43E-39 | 3.289119419 | 0.253 | 0.047 | 2.81E-35 | Mari |
| Macro-PPARG | KRT161 | 3.66E-21 | 2.50776856 | 0.273 | 0.099 | 7.20E-17 | Mari |
| Macro-PPARG | FCGR1B1 | 9.60E-57 | 2.488129522 | 0.539 | 0.173 | 1.89E-52 | Mari |
| Macro-PPARG | CASP51 | 2.70E-27 | 2.460324316 | 0.306 | 0.098 | 5.31E-23 | Mari |
| Macro-PPARG | S100A81 | 1.49E-76 | 2.388341688 | 0.806 | 0.347 | 2.94E-72 | Mari |
| Macro-PPARG | CCL4L2 | 6.64E-15 | 2.097165214 | 0.73 | 0.58 | 1.31E-10 | Mari |
| Macro-PPARG | IGSF21 | 1.57E-29 | 2.071494547 | 0.273 | 0.071 | 3.10E-25 | Mari |
| Macro-PPARG | FCGR1A1 | 2.85E-62 | 1.975333982 | 0.78 | 0.41 | 5.61E-58 | Mari |
| Macro-PPARG | SPATA5L1 | 2.03E-11 | 1.862039039 | 0.296 | 0.155 | 3.99E-07 | Mari |
| Macro-PPARG | VAMP51 | 3.66E-60 | 1.839916559 | 0.819 | 0.465 | 7.20E-56 | Mari |
| Macro-PPARG | TMEM1631 | 6.21E-29 | 1.804758677 | 0.365 | 0.122 | 1.22E-24 | Mari |
| Macro-PPARG | IFITM3 | 1.25E-31 | 1.804167773 | 0.826 | 0.642 | 2.45E-27 | Mari |
| Macro-PPARG | CD300E1 | 4.69E-24 | 1.773476152 | 0.299 | 0.099 | 9.24E-20 | Mari |
| Macro-PPARG | APOL41 | 1.64E-25 | 1.747774647 | 0.306 | 0.098 | 3.22E-21 | Mari |
| Macro-PPARG | CALHM61 | 2.90E-59 | 1.720437464 | 0.875 | 0.549 | 5.71E-55 | Mari |
| Macro-PPARG | CD691 | 1.25E-22 | 1.717901001 | 0.391 | 0.168 | 2.46E-18 | Mari |
| Macro-PPARG | CD931 | 2.47E-22 | 1.698798423 | 0.375 | 0.155 | 4.86E-18 | Mari |
| Macro-PPARG | TNFSF101 | 1.03E-48 | 1.693866033 | 0.711 | 0.332 | 2.02E-44 | Mari |
| Macro-PPARG | CXCL101 | 4.09E-18 | 1.661066178 | 0.418 | 0.208 | 8.06E-14 | Mari |
| Macro-PPARG | TMEM1731 | 5.44E-23 | 1.658708897 | 0.401 | 0.174 | 1.07E-18 | Mari |
| Macro-PPARG | CCDC167 | 2.81E-18 | 1.652593805 | 0.286 | 0.11 | 5.53E-14 | Mari |
| Macro-PPARG | PLAAT41 | 7.58E-53 | 1.638689462 | 0.727 | 0.34 | 1.49E-48 | Mari |
| Macro-PPARG | CCL3L1 | 4.16E-09 | 1.63643957 | 0.799 | 0.746 | 8.19E-05 | Mari |
| Macro-PPARG | IDO11 | 1.77E-18 | 1.6359395 | 0.319 | 0.131 | 3.49E-14 | Mari |
| Macro-PPARG | PSME21 | 4.85E-57 | 1.616869011 | 0.842 | 0.543 | 9.55E-53 | Mari |
| Macro-PPARG | GBP11 | 8.73E-50 | 1.610556054 | 0.789 | 0.413 | 1.72E-45 | Mari |
| Macro-PPARG | PCED1B-AS1 | 1.93E-20 | 1.596217145 | 0.342 | 0.138 | 3.79E-16 | Mari |
| Macro-PPARG | IFI44L | 1.10E-28 | 1.59491271 | 0.388 | 0.137 | 2.16E-24 | Mari |
| Macro-PPARG | IFI351 | 9.03E-25 | 1.585280786 | 0.438 | 0.194 | 1.78E-20 | Mari |
| Macro-PPARG | GIMAP81 | 8.99E-18 | 1.57309656 | 0.312 | 0.131 | 1.77E-13 | Mari |
| Macro-PPARG | S100A91 | 2.37E-47 | 1.566142952 | 0.882 | 0.647 | 4.66E-43 | Mari |

| | | | | | | | |
|---|---|---|---|---|---|---|---|
| Macro-PPARG | SDF2L11 | 5.07E-29 | 1.549550425 | 0.582 | 0.299 | 9.98E-25 | Mari |
| Macro-PPARG | PLTP | 3.88E-22 | 1.513983382 | 0.75 | 0.566 | 7.64E-18 | Mari |
| Macro-PPARG | GIMAP71 | 8.82E-24 | 1.50510028 | 0.388 | 0.158 | 1.74E-19 | Mari |
| Macro-PPARG | VPS251 | 1.75E-20 | 1.500792004 | 0.375 | 0.164 | 3.45E-16 | Mari |
| Macro-PPARG | TMEM1091 | 8.15E-25 | 1.475123284 | 0.464 | 0.208 | 1.60E-20 | Mari |
| Macro-PPARG | PSMB91 | 4.57E-52 | 1.452395461 | 0.842 | 0.558 | 9.00E-48 | Mari |
| Macro-PPARG | OLFML2B1 | 8.56E-30 | 1.437092463 | 0.556 | 0.256 | 1.69E-25 | Mari |
| Macro-PPARG | CRELD21 | 6.88E-18 | 1.424431676 | 0.382 | 0.179 | 1.35E-13 | Mari |
| Macro-PPARG | TMEM170B | 3.79E-08 | 1.40563367 | 0.286 | 0.165 | 0.000746359 | Mari |
| Macro-PPARG | CORO1A | 1.93E-23 | 1.388856029 | 0.579 | 0.321 | 3.81E-19 | Mari |
| Macro-PPARG | FCGR3A1 | 1.68E-52 | 1.387002867 | 0.878 | 0.562 | 3.31E-48 | Mari |
| Macro-PPARG | ASCL21 | 2.13E-19 | 1.376961992 | 0.352 | 0.149 | 4.19E-15 | Mari |
| Macro-PPARG | TYMP1 | 3.91E-72 | 1.375032794 | 0.98 | 0.808 | 7.69E-68 | Mari |
| Macro-PPARG | F13A11 | 8.56E-22 | 1.35732892 | 0.559 | 0.32 | 1.69E-17 | Mari |
| Macro-PPARG | CD163L11 | 4.04E-12 | 1.356703363 | 0.283 | 0.134 | 7.95E-08 | Mari |
| Macro-PPARG | MICAL21 | 4.86E-09 | 1.351912442 | 0.25 | 0.13 | 9.57E-05 | Mari |
| Macro-PPARG | ZDHHC121 | 1.97E-18 | 1.347416392 | 0.428 | 0.216 | 3.87E-14 | Mari |
| Macro-PPARG | HRH21 | 7.45E-19 | 1.346481665 | 0.424 | 0.208 | 1.47E-14 | Mari |
| Macro-PPARG | INHBA-AS1 | 1.28E-09 | 1.34509269 | 0.27 | 0.137 | 2.52E-05 | Mari |
| Macro-PPARG | UQCC3 | 3.30E-14 | 1.344777996 | 0.299 | 0.136 | 6.50E-10 | Mari |
| Macro-PPARG | HS3ST1 | 7.45E-13 | 1.340835347 | 0.293 | 0.136 | 1.47E-08 | Mari |
| Macro-PPARG | IL7R1 | 1.94E-21 | 1.33971846 | 0.53 | 0.28 | 3.82E-17 | Mari |
| Macro-PPARG | GALK1 | 1.28E-12 | 1.336493226 | 0.336 | 0.174 | 2.51E-08 | Mari |
| Macro-PPARG | MRPS121 | 7.45E-20 | 1.33317829 | 0.451 | 0.222 | 1.47E-15 | Mari |
| Macro-PPARG | ANKRD221 | 3.29E-34 | 1.327634544 | 0.49 | 0.185 | 6.48E-30 | Mari |
| Macro-PPARG | FAM89B | 7.38E-21 | 1.326844605 | 0.477 | 0.237 | 1.45E-16 | Mari |
| Macro-PPARG | CCL81 | 3.34E-14 | 1.310105219 | 0.322 | 0.15 | 6.58E-10 | Mari |
| Macro-PPARG | XRCC4 | 7.38E-16 | 1.306844682 | 0.342 | 0.157 | 1.45E-11 | Mari |
| Macro-PPARG | POLE41 | 7.23E-15 | 1.306125954 | 0.503 | 0.311 | 1.42E-10 | Mari |
| Macro-PPARG | PDIA41 | 3.61E-14 | 1.305216404 | 0.441 | 0.255 | 7.11E-10 | Mari |
| Macro-PPARG | BST2 | 3.06E-48 | 1.303223399 | 0.875 | 0.666 | 6.03E-44 | Mari |
| Macro-PPARG | SERPINA11 | 4.61E-42 | 1.299635867 | 0.849 | 0.57 | 9.07E-38 | Mari |
| Macro-PPARG | MRPL23 | 1.17E-25 | 1.294580196 | 0.47 | 0.205 | 2.31E-21 | Mari |
| Macro-PPARG | TLR82 | 1.56E-16 | 1.292438304 | 0.398 | 0.198 | 3.08E-12 | Mari |
| Macro-PPARG | DNAJC13 | 8.95E-15 | 1.281648859 | 0.401 | 0.212 | 1.76E-10 | Mari |
| Macro-PPARG | TUBA1A | 4.29E-29 | 1.277442765 | 0.796 | 0.532 | 8.45E-25 | Mari |
| Macro-PPARG | YIF1A | 3.52E-20 | 1.273525731 | 0.438 | 0.212 | 6.93E-16 | Mari |
| Macro-PPARG | TSFM1 | 4.18E-12 | 1.270839242 | 0.273 | 0.128 | 8.23E-08 | Mari |
| Macro-PPARG | DCTPP1 | 3.22E-11 | 1.269331851 | 0.283 | 0.141 | 6.34E-07 | Mari |
| Macro-PPARG | S100A41 | 3.99E-28 | 1.254637113 | 0.901 | 0.746 | 7.85E-24 | Mari |
| Macro-PPARG | CAMK1 | 5.56E-14 | 1.25083079 | 0.322 | 0.154 | 1.10E-09 | Mari |
| Macro-PPARG | CENPX1 | 6.55E-19 | 1.250004616 | 0.52 | 0.291 | 1.29E-14 | Mari |
| Macro-PPARG | INHBA | 1.67E-06 | 1.244122393 | 0.444 | 0.328 | 0.032925497 | Mari |
| Macro-PPARG | MRPL42 | 2.96E-10 | 1.236797791 | 0.27 | 0.137 | 5.83E-06 | Mari |
| Macro-PPARG | FCGR2B | 3.87E-28 | 1.231529341 | 0.714 | 0.431 | 7.62E-24 | Mari |
| Macro-PPARG | PRELID11 | 2.91E-32 | 1.231162141 | 0.816 | 0.593 | 5.73E-28 | Mari |
| Macro-PPARG | MAGOHB1 | 3.59E-12 | 1.226858163 | 0.293 | 0.138 | 7.07E-08 | Mari |
| Macro-PPARG | CENPW | 2.94E-07 | 1.225381642 | 0.26 | 0.147 | 0.005792187 | Mari |
| Macro-PPARG | LAP31 | 4.56E-55 | 1.221081471 | 0.918 | 0.679 | 8.98E-51 | Mari |
| Macro-PPARG | NHP2 | 5.73E-17 | 1.218988458 | 0.421 | 0.217 | 1.13E-12 | Mari |
| Macro-PPARG | NMI1 | 1.37E-15 | 1.218242763 | 0.431 | 0.233 | 2.70E-11 | Mari |
| Macro-PPARG | HAPLN31 | 6.86E-18 | 1.215245722 | 0.368 | 0.167 | 1.35E-13 | Mari |
| Macro-PPARG | DCXR | 4.72E-19 | 1.214850395 | 0.474 | 0.243 | 9.30E-15 | Mari |
| Macro-PPARG | CASP41 | 1.32E-32 | 1.214279448 | 0.783 | 0.484 | 2.61E-28 | Mari |
| Macro-PPARG | SCO21 | 2.46E-33 | 1.212247271 | 0.766 | 0.442 | 4.84E-29 | Mari |

| | | | | | | | |
|---|---|---|---|---|---|---|---|
| Macro-PPARG | GIMAP61 | 1.36E-13 | 1.212157996 | 0.266 | 0.115 | 2.68E-09 | Mari |
| Macro-PPARG | NME11 | 8.83E-15 | 1.204581198 | 0.464 | 0.263 | 1.74E-10 | Mari |
| Macro-PPARG | RNF141 | 3.86E-09 | 1.19886892 | 0.336 | 0.194 | 7.59E-05 | Mari |
| Macro-PPARG | SLC35B1 | 1.22E-15 | 1.195540249 | 0.388 | 0.192 | 2.40E-11 | Mari |
| Macro-PPARG | SATB1-AS11 | 3.89E-12 | 1.193364593 | 0.395 | 0.223 | 7.66E-08 | Mari |
| Macro-PPARG | RGS18 | 4.62E-07 | 1.19120825 | 0.276 | 0.167 | 0.009086532 | Mari |
| Macro-PPARG | IFIT31 | 1.87E-14 | 1.191168852 | 0.266 | 0.11 | 3.69E-10 | Mari |
| Macro-PPARG | IL61 | 1.51E-08 | 1.190215726 | 0.286 | 0.161 | 0.000297168 | Mari |
| Macro-PPARG | CD141 | 9.49E-32 | 1.189612401 | 0.908 | 0.787 | 1.87E-27 | Mari |
| Macro-PPARG | MNDA1 | 4.83E-29 | 1.189443447 | 0.776 | 0.492 | 9.51E-25 | Mari |
| Macro-PPARG | RNASE11 | 4.29E-18 | 1.18680265 | 0.688 | 0.462 | 8.45E-14 | Mari |
| Macro-PPARG | ORMDL21 | 3.87E-17 | 1.185946697 | 0.378 | 0.179 | 7.62E-13 | Mari |
| Macro-PPARG | AIF11 | 6.90E-33 | 1.184429755 | 0.914 | 0.817 | 1.36E-28 | Mari |
| Macro-PPARG | KRT141 | 9.57E-07 | 1.179909088 | 0.332 | 0.237 | 0.018836264 | Mari |
| Macro-PPARG | GBP51 | 3.72E-33 | 1.175718363 | 0.714 | 0.395 | 7.32E-29 | Mari |
| Macro-PPARG | BANF11 | 5.49E-24 | 1.173600296 | 0.658 | 0.375 | 1.08E-19 | Mari |
| Macro-PPARG | DTX3L1 | 2.95E-23 | 1.173447488 | 0.457 | 0.205 | 5.81E-19 | Mari |
| Macro-PPARG | DHRS4 | 1.55E-15 | 1.17329569 | 0.408 | 0.212 | 3.04E-11 | Mari |
| Macro-PPARG | GADD45GIP1 | 1.17E-20 | 1.172970833 | 0.648 | 0.411 | 2.30E-16 | Mari |
| Macro-PPARG | JAK21 | 2.96E-22 | 1.168900908 | 0.655 | 0.379 | 5.82E-18 | Mari |
| Macro-PPARG | 46092 | 2.13E-24 | 1.163968528 | 0.704 | 0.408 | 4.19E-20 | Mari |
| Macro-PPARG | PSME11 | 1.23E-40 | 1.163859925 | 0.803 | 0.545 | 2.42E-36 | Mari |
| Macro-PPARG | PPIE | 1.75E-12 | 1.163802816 | 0.312 | 0.152 | 3.44E-08 | Mari |
| Macro-PPARG | ISCA21 | 8.43E-14 | 1.160281586 | 0.362 | 0.181 | 1.66E-09 | Mari |
| Macro-PPARG | CRIP11 | 7.57E-27 | 1.159843121 | 0.747 | 0.455 | 1.49E-22 | Mari |
| Macro-PPARG | TMEM147 | 2.55E-19 | 1.15705429 | 0.582 | 0.335 | 5.03E-15 | Mari |
| Macro-PPARG | PSTPIP1 | 1.37E-17 | 1.155282054 | 0.405 | 0.189 | 2.71E-13 | Mari |
| Macro-PPARG | MRPS18C | 7.21E-27 | 1.155119271 | 0.592 | 0.302 | 1.42E-22 | Mari |
| Macro-PPARG | IFITM2 | 4.32E-14 | 1.154147004 | 0.668 | 0.473 | 8.51E-10 | Mari |
| Macro-PPARG | GIMAP11 | 2.48E-20 | 1.15381493 | 0.516 | 0.269 | 4.88E-16 | Mari |
| Macro-PPARG | GIMAP41 | 1.82E-27 | 1.147114955 | 0.602 | 0.31 | 3.59E-23 | Mari |
| Macro-PPARG | COMMD8 | 8.21E-13 | 1.142465273 | 0.398 | 0.222 | 1.62E-08 | Mari |
| Macro-PPARG | UBE2L61 | 2.08E-21 | 1.140070594 | 0.592 | 0.339 | 4.09E-17 | Mari |
| Macro-PPARG | MRPL581 | 1.20E-11 | 1.137067188 | 0.28 | 0.134 | 2.36E-07 | Mari |
| Macro-PPARG | IFI301 | 3.04E-54 | 1.1349661 | 0.984 | 0.928 | 5.99E-50 | Mari |
| Macro-PPARG | PFN11 | 7.82E-48 | 1.131541516 | 0.911 | 0.809 | 1.54E-43 | Mari |
| Macro-PPARG | ARF5 | 1.72E-20 | 1.130950556 | 0.622 | 0.394 | 3.39E-16 | Mari |
| Macro-PPARG | TREM11 | 2.62E-10 | 1.127751224 | 0.316 | 0.169 | 5.16E-06 | Mari |
| Macro-PPARG | CYBA | 1.73E-65 | 1.125600852 | 0.967 | 0.872 | 3.41E-61 | Mari |
| Macro-PPARG | C1QC | 1.23E-36 | 1.120481731 | 0.911 | 0.789 | 2.41E-32 | Mari |
| Macro-PPARG | BST1 | 2.44E-12 | 1.117600583 | 0.326 | 0.159 | 4.81E-08 | Mari |
| Macro-PPARG | ALG5 | 2.04E-18 | 1.113749283 | 0.477 | 0.248 | 4.01E-14 | Mari |
| Macro-PPARG | SLC8A12 | 7.84E-40 | 1.113562442 | 0.964 | 0.718 | 1.54E-35 | Mari |
| Macro-PPARG | CCR11 | 1.95E-14 | 1.111180776 | 0.447 | 0.248 | 3.84E-10 | Mari |
| Macro-PPARG | DCUN1D5 | 1.74E-13 | 1.111135577 | 0.339 | 0.166 | 3.43E-09 | Mari |
| Macro-PPARG | SECTM11 | 2.69E-20 | 1.109809106 | 0.526 | 0.278 | 5.29E-16 | Mari |
| Macro-PPARG | GTF3C61 | 1.84E-21 | 1.109431574 | 0.595 | 0.354 | 3.62E-17 | Mari |
| Macro-PPARG | TXNDC111 | 1.30E-12 | 1.109341836 | 0.507 | 0.311 | 2.56E-08 | Mari |
| Macro-PPARG | MTLN1 | 2.96E-19 | 1.109043056 | 0.454 | 0.224 | 5.82E-15 | Mari |
| Macro-PPARG | MRPL52 | 9.88E-26 | 1.10605711 | 0.678 | 0.394 | 1.94E-21 | Mari |
| Macro-PPARG | ANTKMT | 7.96E-18 | 1.102512528 | 0.493 | 0.272 | 1.57E-13 | Mari |
| Macro-PPARG | SP1101 | 2.14E-22 | 1.100269166 | 0.546 | 0.275 | 4.22E-18 | Mari |
| Macro-PPARG | MEF2C1 | 6.17E-24 | 1.098771985 | 0.74 | 0.488 | 1.21E-19 | Mari |
| Macro-PPARG | SIGLEC101 | 3.13E-11 | 1.093171339 | 0.362 | 0.199 | 6.15E-07 | Mari |
| Macro-PPARG | JOSD21 | 7.19E-19 | 1.092565664 | 0.513 | 0.282 | 1.42E-14 | Mari |

| | | | | | | | |
|---|---|---|---|---|---|---|---|
| Macro-PPARG | TRAPPC12 | 3.44E-23 | 1.092162362 | 0.645 | 0.378 | 6.77E-19 | Mari |
| Macro-PPARG | SRP9 | 4.24E-23 | 1.091336492 | 0.664 | 0.401 | 8.36E-19 | Mari |
| Macro-PPARG | RRAS1 | 5.95E-20 | 1.090297186 | 0.536 | 0.29 | 1.17E-15 | Mari |
| Macro-PPARG | ARHGDIB1 | 3.81E-32 | 1.08834665 | 0.836 | 0.648 | 7.50E-28 | Mari |
| Macro-PPARG | COA41 | 4.28E-14 | 1.086442531 | 0.398 | 0.21 | 8.43E-10 | Mari |
| Macro-PPARG | COPS7A | 6.85E-08 | 1.085781284 | 0.263 | 0.147 | 0.001349158 | Mari |
| Macro-PPARG | PARP91 | 3.79E-18 | 1.085033673 | 0.434 | 0.211 | 7.47E-14 | Mari |
| Macro-PPARG | C1QB1 | 1.48E-42 | 1.078631721 | 0.944 | 0.803 | 2.92E-38 | Mari |
| Macro-PPARG | VSIG4 | 1.06E-15 | 1.075199834 | 0.612 | 0.406 | 2.08E-11 | Mari |
| Macro-PPARG | SNX6 | 8.49E-28 | 1.075083985 | 0.849 | 0.687 | 1.67E-23 | Mari |
| Macro-PPARG | TNFSF131 | 5.17E-23 | 1.074928723 | 0.707 | 0.476 | 1.02E-18 | Mari |
| Macro-PPARG | KDM1B | 8.37E-12 | 1.072641485 | 0.326 | 0.17 | 1.65E-07 | Mari |
| Macro-PPARG | NOL12 | 2.69E-10 | 1.072542426 | 0.263 | 0.129 | 5.29E-06 | Mari |
| Macro-PPARG | PPIB | 5.59E-36 | 1.072244894 | 0.872 | 0.69 | 1.10E-31 | Mari |
| Macro-PPARG | FPR1 | 4.71E-20 | 1.069075148 | 0.668 | 0.434 | 9.28E-16 | Mari |
| Macro-PPARG | STAT12 | 1.37E-42 | 1.065947155 | 0.905 | 0.585 | 2.70E-38 | Mari |
| Macro-PPARG | CHMP6 | 1.02E-10 | 1.061970378 | 0.257 | 0.122 | 2.00E-06 | Mari |
| Macro-PPARG | ITGA41 | 1.41E-13 | 1.061375827 | 0.418 | 0.233 | 2.78E-09 | Mari |
| Macro-PPARG | ZNF593 | 2.65E-14 | 1.061206241 | 0.365 | 0.182 | 5.23E-10 | Mari |
| Macro-PPARG | FBXO61 | 2.16E-14 | 1.059986504 | 0.411 | 0.222 | 4.25E-10 | Mari |
| Macro-PPARG | LIMA11 | 9.01E-09 | 1.058632136 | 0.25 | 0.131 | 0.000177433 | Mari |
| Macro-PPARG | MRPL121 | 4.27E-13 | 1.058288841 | 0.382 | 0.203 | 8.41E-09 | Mari |
| Macro-PPARG | ISOC21 | 2.28E-12 | 1.057731646 | 0.424 | 0.244 | 4.49E-08 | Mari |
| Macro-PPARG | IER2 | 8.61E-17 | 1.05662642 | 0.911 | 0.795 | 1.70E-12 | Mari |
| Macro-PPARG | PCNX12 | 7.40E-25 | 1.056601342 | 0.73 | 0.435 | 1.46E-20 | Mari |
| Macro-PPARG | ARL21 | 5.58E-14 | 1.056319276 | 0.474 | 0.281 | 1.10E-09 | Mari |
| Macro-PPARG | TMEM87B | 9.79E-11 | 1.055711389 | 0.283 | 0.139 | 1.93E-06 | Mari |
| Macro-PPARG | HIGD1A | 3.42E-11 | 1.05551579 | 0.352 | 0.19 | 6.73E-07 | Mari |
| Macro-PPARG | TMEM2081 | 3.19E-21 | 1.051050994 | 0.618 | 0.384 | 6.27E-17 | Mari |
| Macro-PPARG | S100A10 | 1.27E-31 | 1.050995212 | 0.947 | 0.816 | 2.50E-27 | Mari |
| Macro-PPARG | MSRB21 | 4.97E-18 | 1.050848982 | 0.549 | 0.308 | 9.78E-14 | Mari |
| Macro-PPARG | FIBP | 7.84E-20 | 1.048375194 | 0.562 | 0.312 | 1.54E-15 | Mari |
| Macro-PPARG | CHMP2A | 8.11E-24 | 1.047023613 | 0.707 | 0.464 | 1.60E-19 | Mari |
| Macro-PPARG | TSPO | 1.59E-38 | 1.046266386 | 0.908 | 0.739 | 3.13E-34 | Mari |
| Macro-PPARG | MYO1G1 | 1.61E-13 | 1.045974489 | 0.372 | 0.186 | 3.17E-09 | Mari |
| Macro-PPARG | NUDT161 | 7.88E-14 | 1.045019796 | 0.424 | 0.233 | 1.55E-09 | Mari |
| Macro-PPARG | EPSTI11 | 8.36E-41 | 1.044750527 | 0.855 | 0.538 | 1.65E-36 | Mari |
| Macro-PPARG | GCHFR | 1.68E-06 | 1.04155945 | 0.349 | 0.244 | 0.033158174 | Mari |
| Macro-PPARG | EEF1E1 | 2.78E-10 | 1.040527618 | 0.276 | 0.139 | 5.48E-06 | Mari |
| Macro-PPARG | NDUFC2 | 4.51E-29 | 1.038907063 | 0.763 | 0.525 | 8.87E-25 | Mari |
| Macro-PPARG | MRPL28 | 8.13E-12 | 1.036051825 | 0.391 | 0.218 | 1.60E-07 | Mari |
| Macro-PPARG | TMED31 | 1.53E-14 | 1.034975445 | 0.418 | 0.224 | 3.02E-10 | Mari |
| Macro-PPARG | EIF4EBP1 | 2.53E-20 | 1.034080064 | 0.632 | 0.37 | 4.98E-16 | Mari |
| Macro-PPARG | OSM1 | 4.66E-16 | 1.033960895 | 0.507 | 0.278 | 9.17E-12 | Mari |
| Macro-PPARG | SELENON | 1.24E-09 | 1.03290919 | 0.27 | 0.135 | 2.45E-05 | Mari |
| Macro-PPARG | SMCO41 | 1.07E-27 | 1.030289501 | 0.711 | 0.426 | 2.10E-23 | Mari |
| Macro-PPARG | RAB39A1 | 2.57E-13 | 1.029511524 | 0.336 | 0.16 | 5.06E-09 | Mari |
| Macro-PPARG | NDUFB31 | 1.45E-21 | 1.027404222 | 0.651 | 0.401 | 2.84E-17 | Mari |
| Macro-PPARG | RFK1 | 2.81E-11 | 1.026440919 | 0.312 | 0.16 | 5.52E-07 | Mari |
| Macro-PPARG | RNF166 | 1.76E-10 | 1.026376276 | 0.408 | 0.245 | 3.46E-06 | Mari |
| Macro-PPARG | SLC39A81 | 1.66E-08 | 1.025888522 | 0.582 | 0.448 | 0.000327158 | Mari |
| Macro-PPARG | PRDX2 | 1.38E-13 | 1.02423037 | 0.342 | 0.169 | 2.71E-09 | Mari |
| Macro-PPARG | BORCS8 | 1.67E-10 | 1.022305782 | 0.283 | 0.141 | 3.29E-06 | Mari |
| Macro-PPARG | CYB561D21 | 1.22E-13 | 1.021656472 | 0.434 | 0.237 | 2.40E-09 | Mari |
| Macro-PPARG | MX1 | 3.45E-15 | 1.021521371 | 0.47 | 0.26 | 6.80E-11 | Mari |

| | | | | | | | |
|---|---|---|---|---|---|---|---|
| Macro-PPARG | GCA | 2.80E-19 | 1.017206322 | 0.635 | 0.406 | 5.51E-15 | Mari |
| Macro-PPARG | FHL31 | 4.35E-07 | 1.016217064 | 0.25 | 0.142 | 0.008559731 | Mari |
| Macro-PPARG | ANKRD441 | 4.91E-23 | 1.015559795 | 0.717 | 0.418 | 9.67E-19 | Mari |
| Macro-PPARG | CIAO2A | 4.28E-25 | 1.014819849 | 0.697 | 0.443 | 8.43E-21 | Mari |
| Macro-PPARG | CD471 | 6.42E-31 | 1.012956075 | 0.809 | 0.545 | 1.26E-26 | Mari |
| Macro-PPARG | CIB1 | 2.34E-31 | 1.012780701 | 0.809 | 0.569 | 4.61E-27 | Mari |
| Macro-PPARG | SLC12A91 | 3.12E-09 | 1.012253635 | 0.349 | 0.201 | 6.15E-05 | Mari |
| Macro-PPARG | LDLRAD3 | 1.14E-07 | 1.011115206 | 0.253 | 0.138 | 0.002242962 | Mari |
| Macro-PPARG | SCNM11 | 1.12E-09 | 1.010483597 | 0.319 | 0.178 | 2.20E-05 | Mari |
| Macro-PPARG | ENTPD12 | 3.91E-15 | 1.006377432 | 0.533 | 0.314 | 7.70E-11 | Mari |
| Macro-PPARG | APOL61 | 2.19E-22 | 1.005896114 | 0.625 | 0.352 | 4.32E-18 | Mari |
| Macro-PPARG | IL10 | 3.91E-07 | 1.004762624 | 0.447 | 0.327 | 0.00768748 | Mari |
| Macro-PPARG | FAM207A | 5.12E-08 | 1.004148183 | 0.276 | 0.154 | 0.001008301 | Mari |
| Macro-PPARG | REEP41 | 4.87E-12 | 1.003808675 | 0.362 | 0.192 | 9.59E-08 | Mari |
| Macro-PPARG | PHF21A | 3.59E-10 | 1.003376706 | 0.451 | 0.282 | 7.07E-06 | Mari |
| Macro-PPARG | S100A61 | 1.50E-33 | 1.002425099 | 0.97 | 0.863 | 2.96E-29 | Mari |
| Macro-PPARG | GOSR2 | 3.63E-12 | 1.001929559 | 0.329 | 0.165 | 7.14E-08 | Mari |
| Macro-PPARG | SLC8A2 | 4.71E-72 | 3.812523227 | 0.379 | 0.072 | 9.27E-68 | Tb |
| Macro-PPARG | FN1 | 1.67E-51 | 3.271088265 | 0.441 | 0.139 | 3.29E-47 | Tb |
| Macro-PPARG | KRT6A | 3.39E-104 | 3.074679977 | 0.424 | 0.05 | 6.68E-100 | Tb |
| Macro-PPARG | GGT5 | 1.00E-70 | 2.990793201 | 0.335 | 0.051 | 1.98E-66 | Tb |
| Macro-PPARG | SIGLEC12 | 6.73E-69 | 2.91147995 | 0.283 | 0.033 | 1.33E-64 | Tb |
| Macro-PPARG | CXCL9 | 2.52E-100 | 2.824548173 | 0.665 | 0.178 | 4.95E-96 | Tb |
| Macro-PPARG | CCL18 | 9.66E-16 | 2.677463417 | 0.567 | 0.422 | 1.90E-11 | Tb |
| Macro-PPARG | IDO1 | 9.91E-35 | 2.462604258 | 0.35 | 0.115 | 1.95E-30 | Tb |
| Macro-PPARG | S100A7 | 5.24E-60 | 2.426022556 | 0.377 | 0.077 | 1.03E-55 | Tb |
| Macro-PPARG | ENPP2 | 2.90E-94 | 2.400737673 | 0.613 | 0.148 | 5.71E-90 | Tb |
| Macro-PPARG | KRT6B | 3.18E-62 | 2.363479513 | 0.273 | 0.034 | 6.25E-58 | Tb |
| Macro-PPARG | KRT17 | 7.11E-60 | 2.256951418 | 0.328 | 0.056 | 1.40E-55 | Tb |
| Macro-PPARG | KPTN | 4.42E-45 | 2.226863397 | 0.266 | 0.051 | 8.70E-41 | Tb |
| Macro-PPARG | SIRPB2 | 9.49E-41 | 2.134604144 | 0.273 | 0.059 | 1.87E-36 | Tb |
| Macro-PPARG | IL2RA | 5.94E-60 | 2.099608576 | 0.483 | 0.136 | 1.17E-55 | Tb |
| Macro-PPARG | AATK | 3.16E-50 | 2.087273058 | 0.325 | 0.068 | 6.21E-46 | Tb |
| Macro-PPARG | KRT5 | 2.80E-38 | 2.081055613 | 0.286 | 0.069 | 5.51E-34 | Tb |
| Macro-PPARG | SFN | 4.17E-34 | 2.041390892 | 0.318 | 0.093 | 8.20E-30 | Tb |
| Macro-PPARG | CXCL10 | 1.28E-54 | 1.996013081 | 0.522 | 0.177 | 2.53E-50 | Tb |
| Macro-PPARG | KRT16 | 8.07E-68 | 1.969133942 | 0.384 | 0.069 | 1.59E-63 | Tb |
| Macro-PPARG | S100A2 | 1.19E-63 | 1.943564202 | 0.394 | 0.079 | 2.34E-59 | Tb |
| Macro-PPARG | GBP5 | 8.92E-74 | 1.919625516 | 0.815 | 0.359 | 1.76E-69 | Tb |
| Macro-PPARG | SIGLEC10 | 3.96E-57 | 1.916691824 | 0.515 | 0.161 | 7.80E-53 | Tb |
| Macro-PPARG | P2RY14 | 6.39E-34 | 1.915332431 | 0.286 | 0.077 | 1.26E-29 | Tb |
| Macro-PPARG | SLC35F1 | 3.43E-48 | 1.914945604 | 0.387 | 0.105 | 6.75E-44 | Tb |
| Macro-PPARG | CD300E | 1.95E-34 | 1.897152798 | 0.308 | 0.087 | 3.83E-30 | Tb |
| Macro-PPARG | KRT14 | 3.50E-76 | 1.871679542 | 0.616 | 0.176 | 6.89E-72 | Tb |
| Macro-PPARG | MT1G | 8.60E-21 | 1.868700908 | 0.3 | 0.118 | 1.69E-16 | Tb |
| Macro-PPARG | S100A8 | 2.63E-89 | 1.844906431 | 0.83 | 0.32 | 5.19E-85 | Tb |
| Macro-PPARG | IFIT3 | 1.99E-22 | 1.833819427 | 0.281 | 0.1 | 3.91E-18 | Tb |
| Macro-PPARG | HOMER1 | 2.69E-57 | 1.783666286 | 0.665 | 0.27 | 5.29E-53 | Tb |
| Macro-PPARG | APOL4 | 4.68E-35 | 1.761585608 | 0.31 | 0.087 | 9.22E-31 | Tb |
| Macro-PPARG | ANKRD22 | 4.21E-51 | 1.756638992 | 0.507 | 0.166 | 8.29E-47 | Tb |
| Macro-PPARG | IL7R | 5.49E-38 | 1.753334839 | 0.567 | 0.26 | 1.08E-33 | Tb |
| Macro-PPARG | GIMAP7 | 1.32E-52 | 1.714160572 | 0.463 | 0.132 | 2.60E-48 | Tb |
| Macro-PPARG | QPCT | 1.25E-29 | 1.708363563 | 0.335 | 0.116 | 2.46E-25 | Tb |
| Macro-PPARG | FBP1 | 9.82E-28 | 1.668027514 | 0.643 | 0.382 | 1.93E-23 | Tb |
| Macro-PPARG | HS3ST3B1 | 3.54E-29 | 1.64093537 | 0.433 | 0.187 | 6.97E-25 | Tb |

| | | | | | | | |
|---|---|---|---|---|---|---|---|
| Macro-PPARG | JAK3 | 2.42E-24 | 1.635486199 | 0.283 | 0.096 | 4.76E-20 | Tb |
| Macro-PPARG | LIMK2 | 4.05E-38 | 1.610727069 | 0.525 | 0.214 | 7.98E-34 | Tb |
| Macro-PPARG | IFIT2 | 6.98E-23 | 1.598899475 | 0.394 | 0.167 | 1.37E-18 | Tb |
| Macro-PPARG | DNAI2 | 5.44E-26 | 1.587607662 | 0.406 | 0.17 | 1.07E-21 | Tb |
| Macro-PPARG | HAPLN3 | 4.36E-37 | 1.565355515 | 0.421 | 0.146 | 8.58E-33 | Tb |
| Macro-PPARG | TMEM163 | 2.05E-32 | 1.565001403 | 0.345 | 0.113 | 4.03E-28 | Tb |
| Macro-PPARG | VDR1 | 9.47E-41 | 1.544890142 | 0.426 | 0.137 | 1.86E-36 | Tb |
| Macro-PPARG | PLAAT4 | 3.67E-61 | 1.542198035 | 0.739 | 0.318 | 7.23E-57 | Tb |
| Macro-PPARG | CCL8 | 1.22E-29 | 1.525762265 | 0.365 | 0.133 | 2.40E-25 | Tb |
| Macro-PPARG | CTSC | 3.00E-89 | 1.513981681 | 0.968 | 0.795 | 5.90E-85 | Tb |
| Macro-PPARG | FAM198B-AS1 | 7.58E-32 | 1.511964029 | 0.384 | 0.134 | 1.49E-27 | Tb |
| Macro-PPARG | MT1E | 4.84E-23 | 1.50612124 | 0.369 | 0.156 | 9.53E-19 | Tb |
| Macro-PPARG | LYZ | 8.42E-64 | 1.50500616 | 0.975 | 0.821 | 1.66E-59 | Tb |
| Macro-PPARG | PPA1 | 2.67E-50 | 1.498775084 | 0.771 | 0.416 | 5.25E-46 | Tb |
| Macro-PPARG | GBP1 | 4.57E-68 | 1.495332169 | 0.823 | 0.388 | 8.99E-64 | Tb |
| Macro-PPARG | TGM2 | 8.22E-27 | 1.49051079 | 0.377 | 0.146 | 1.62E-22 | Tb |
| Macro-PPARG | P2RY6 | 5.19E-45 | 1.46591427 | 0.389 | 0.108 | 1.02E-40 | Tb |
| Macro-PPARG | ADAMTS17 | 2.97E-15 | 1.449003105 | 0.271 | 0.12 | 5.85E-11 | Tb |
| Macro-PPARG | CYSLTR2 | 1.02E-37 | 1.432604182 | 0.34 | 0.097 | 2.00E-33 | Tb |
| Macro-PPARG | TYMP | 3.52E-106 | 1.430102566 | 0.993 | 0.796 | 6.93E-102 | Tb |
| Macro-PPARG | FCGR3A | 3.23E-59 | 1.417265302 | 0.86 | 0.55 | 6.36E-55 | Tb |
| Macro-PPARG | CASS4 | 2.57E-17 | 1.415680268 | 0.256 | 0.1 | 5.06E-13 | Tb |
| Macro-PPARG | GIMAP6 | 6.42E-23 | 1.4153036 | 0.288 | 0.103 | 1.26E-18 | Tb |
| Macro-PPARG | PDCD1LG21 | 1.37E-46 | 1.405998235 | 0.507 | 0.172 | 2.69E-42 | Tb |
| Macro-PPARG | SAP30 | 3.05E-09 | 1.40104391 | 0.399 | 0.255 | 6.01E-05 | Tb |
| Macro-PPARG | DUSP4 | 5.09E-13 | 1.399900861 | 0.259 | 0.123 | 1.00E-08 | Tb |
| Macro-PPARG | SLC12A5 | 2.34E-31 | 1.395693632 | 0.283 | 0.078 | 4.61E-27 | Tb |
| Macro-PPARG | MAN1C1 | 9.19E-20 | 1.388506233 | 0.264 | 0.095 | 1.81E-15 | Tb |
| Macro-PPARG | P2RY131 | 1.90E-41 | 1.383230676 | 0.451 | 0.148 | 3.75E-37 | Tb |
| Macro-PPARG | GIMAP8 | 3.01E-32 | 1.381633051 | 0.352 | 0.114 | 5.93E-28 | Tb |
| Macro-PPARG | SLC16A6 | 5.83E-21 | 1.373744892 | 0.367 | 0.164 | 1.15E-16 | Tb |
| Macro-PPARG | APOL1 | 2.61E-31 | 1.370261515 | 0.32 | 0.099 | 5.14E-27 | Tb |
| Macro-PPARG | CRISPLD2 | 7.09E-30 | 1.365820828 | 0.342 | 0.114 | 1.40E-25 | Tb |
| Macro-PPARG | CD69 | 9.94E-43 | 1.363365104 | 0.448 | 0.146 | 1.96E-38 | Tb |
| Macro-PPARG | VPS9D1 | 1.49E-25 | 1.361970243 | 0.315 | 0.111 | 2.92E-21 | Tb |
| Macro-PPARG | MT1X | 5.33E-19 | 1.355029371 | 0.562 | 0.327 | 1.05E-14 | Tb |
| Macro-PPARG | CASP5 | 7.90E-35 | 1.342862425 | 0.313 | 0.086 | 1.56E-30 | Tb |
| Macro-PPARG | ACE1 | 3.69E-36 | 1.336046996 | 0.401 | 0.134 | 7.27E-32 | Tb |
| Macro-PPARG | CD209 | 9.69E-39 | 1.335057508 | 0.374 | 0.11 | 1.91E-34 | Tb |
| Macro-PPARG | TRAFD1 | 2.19E-22 | 1.328668711 | 0.328 | 0.13 | 4.31E-18 | Tb |
| Macro-PPARG | GBP4 | 1.86E-40 | 1.320919089 | 0.621 | 0.273 | 3.67E-36 | Tb |
| Macro-PPARG | S100A9 | 8.34E-48 | 1.319653863 | 0.938 | 0.625 | 1.64E-43 | Tb |
| Macro-PPARG | SDHC | 2.97E-19 | 1.31672193 | 0.478 | 0.249 | 5.85E-15 | Tb |
| Macro-PPARG | ERRFI1 | 1.29E-22 | 1.314707756 | 0.466 | 0.23 | 2.54E-18 | Tb |
| Macro-PPARG | WARS1 | 1.96E-57 | 1.314360875 | 0.884 | 0.515 | 3.86E-53 | Tb |
| Macro-PPARG | SAMD9L | 9.95E-39 | 1.307251149 | 0.505 | 0.193 | 1.96E-34 | Tb |
| Macro-PPARG | IL15 | 2.19E-25 | 1.306202246 | 0.589 | 0.302 | 4.31E-21 | Tb |
| Macro-PPARG | TMEM39A | 1.78E-17 | 1.306130285 | 0.379 | 0.189 | 3.50E-13 | Tb |
| Macro-PPARG | TSFM | 3.26E-17 | 1.305543813 | 0.283 | 0.118 | 6.42E-13 | Tb |
| Macro-PPARG | VASH1 | 1.02E-27 | 1.304798922 | 0.394 | 0.153 | 2.02E-23 | Tb |
| Macro-PPARG | IL6 | 2.30E-19 | 1.291481668 | 0.335 | 0.145 | 4.52E-15 | Tb |
| Macro-PPARG | TNFSF10 | 4.48E-54 | 1.284881962 | 0.719 | 0.311 | 8.83E-50 | Tb |
| Macro-PPARG | PTGDS1 | 4.17E-07 | 1.283214731 | 0.303 | 0.19 | 0.008206639 | Tb |
| Macro-PPARG | SLC16A10 | 3.99E-28 | 1.27847596 | 0.759 | 0.534 | 7.85E-24 | Tb |
| Macro-PPARG | HRH2 | 3.46E-36 | 1.266148174 | 0.483 | 0.186 | 6.82E-32 | Tb |

| | | | | | | | |
|---|---|---|---|---|---|---|---|
| Macro-PPARG | SLC39A8 | 7.14E-25 | 1.263718681 | 0.68 | 0.422 | 1.41E-20 | Tb |
| Macro-PPARG | KCNK6 | 2.77E-27 | 1.260672719 | 0.337 | 0.118 | 5.45E-23 | Tb |
| Macro-PPARG | LNCAROD | 8.96E-18 | 1.255130302 | 0.256 | 0.099 | 1.76E-13 | Tb |
| Macro-PPARG | ABCC3 | 9.66E-29 | 1.248318026 | 0.498 | 0.226 | 1.90E-24 | Tb |
| Macro-PPARG | ILF3-DT | 7.79E-25 | 1.23919478 | 0.515 | 0.259 | 1.53E-20 | Tb |
| Macro-PPARG | VAMP5 | 8.64E-55 | 1.231230507 | 0.818 | 0.448 | 1.70E-50 | Tb |
| Macro-PPARG | FBXO6 | 4.31E-29 | 1.229271552 | 0.473 | 0.201 | 8.48E-25 | Tb |
| Macro-PPARG | STAT11 | 1.09E-74 | 1.226751668 | 0.941 | 0.562 | 2.15E-70 | Tb |
| Macro-PPARG | SNX20 | 4.82E-27 | 1.226420976 | 0.33 | 0.115 | 9.49E-23 | Tb |
| Macro-PPARG | ERAP21 | 1.55E-34 | 1.224648914 | 0.404 | 0.138 | 3.04E-30 | Tb |
| Macro-PPARG | MMP14 | 1.50E-38 | 1.220927734 | 0.606 | 0.266 | 2.94E-34 | Tb |
| Macro-PPARG | NLRC5 | 1.41E-39 | 1.219684362 | 0.5 | 0.183 | 2.78E-35 | Tb |
| Macro-PPARG | CCND1 | 7.28E-20 | 1.219388124 | 0.278 | 0.106 | 1.43E-15 | Tb |
| Macro-PPARG | SEMA4D | 1.17E-30 | 1.217373583 | 0.515 | 0.223 | 2.30E-26 | Tb |
| Macro-PPARG | SERPING11 | 1.76E-53 | 1.213068353 | 0.828 | 0.467 | 3.46E-49 | Tb |
| Macro-PPARG | ZNF688 | 3.56E-17 | 1.211108877 | 0.256 | 0.099 | 7.01E-13 | Tb |
| Macro-PPARG | STRADA | 1.51E-14 | 1.199086419 | 0.261 | 0.113 | 2.98E-10 | Tb |
| Macro-PPARG | TGFBI1 | 3.60E-27 | 1.19591157 | 0.764 | 0.506 | 7.08E-23 | Tb |
| Macro-PPARG | PAPSS21 | 1.53E-26 | 1.194746183 | 0.781 | 0.585 | 3.02E-22 | Tb |
| Macro-PPARG | HBEGF1 | 3.16E-48 | 1.19335045 | 0.847 | 0.522 | 6.22E-44 | Tb |
| Macro-PPARG | ADAMTSL4 | 5.97E-24 | 1.191936715 | 0.283 | 0.096 | 1.17E-19 | Tb |
| Macro-PPARG | PRKAR2B1 | 7.07E-17 | 1.190462008 | 0.313 | 0.138 | 1.39E-12 | Tb |
| Macro-PPARG | SKI1 | 2.51E-40 | 1.179860843 | 0.631 | 0.296 | 4.94E-36 | Tb |
| Macro-PPARG | PTAFR | 5.34E-40 | 1.17865587 | 0.685 | 0.326 | 1.05E-35 | Tb |
| Macro-PPARG | LRRC25 | 3.72E-31 | 1.176708762 | 0.527 | 0.236 | 7.32E-27 | Tb |
| Macro-PPARG | FLT1 | 6.99E-12 | 1.174846758 | 0.433 | 0.272 | 1.38E-07 | Tb |
| Macro-PPARG | FCGR1A | 3.70E-53 | 1.174788304 | 0.778 | 0.391 | 7.27E-49 | Tb |
| Macro-PPARG | ALOX5 | 6.38E-40 | 1.174315522 | 0.702 | 0.344 | 1.26E-35 | Tb |
| Macro-PPARG | MRC11 | 1.19E-40 | 1.173807189 | 0.862 | 0.561 | 2.34E-36 | Tb |
| Macro-PPARG | PSME2 | 3.84E-61 | 1.173669332 | 0.877 | 0.521 | 7.55E-57 | Tb |
| Macro-PPARG | SUCNR1 | 7.07E-31 | 1.171482727 | 0.355 | 0.119 | 1.39E-26 | Tb |
| Macro-PPARG | MMP12 | 2.91E-51 | 1.166229848 | 0.33 | 0.069 | 5.72E-47 | Tb |
| Macro-PPARG | SYTL3 | 2.36E-21 | 1.160798344 | 0.451 | 0.227 | 4.65E-17 | Tb |
| Macro-PPARG | APOL31 | 4.95E-30 | 1.156910082 | 0.433 | 0.171 | 9.75E-26 | Tb |
| Macro-PPARG | PKD2 | 8.03E-14 | 1.153308576 | 0.313 | 0.149 | 1.58E-09 | Tb |
| Macro-PPARG | EEF2K1 | 2.26E-29 | 1.149595345 | 0.426 | 0.168 | 4.46E-25 | Tb |
| Macro-PPARG | RRAS | 5.13E-38 | 1.1465141 | 0.611 | 0.263 | 1.01E-33 | Tb |
| Macro-PPARG | N4BP2 | 1.13E-20 | 1.14549458 | 0.365 | 0.156 | 2.22E-16 | Tb |
| Macro-PPARG | ACOT71 | 5.18E-29 | 1.144725889 | 0.397 | 0.149 | 1.02E-24 | Tb |
| Macro-PPARG | SMAD7 | 9.49E-34 | 1.140466476 | 0.643 | 0.314 | 1.87E-29 | Tb |
| Macro-PPARG | UST1 | 7.93E-34 | 1.138135375 | 0.591 | 0.297 | 1.56E-29 | Tb |
| Macro-PPARG | SCO2 | 8.34E-37 | 1.137608765 | 0.761 | 0.426 | 1.64E-32 | Tb |
| Macro-PPARG | ASCL2 | 1.76E-24 | 1.136446783 | 0.357 | 0.138 | 3.46E-20 | Tb |
| Macro-PPARG | HIST1H1C | 4.37E-22 | 1.135596567 | 0.281 | 0.099 | 8.61E-18 | Tb |
| Macro-PPARG | PTPN22 | 2.24E-14 | 1.133981481 | 0.281 | 0.129 | 4.41E-10 | Tb |
| Macro-PPARG | CD300LF | 7.88E-20 | 1.126085507 | 0.421 | 0.198 | 1.55E-15 | Tb |
| Macro-PPARG | HELB | 1.80E-15 | 1.123728497 | 0.254 | 0.104 | 3.54E-11 | Tb |
| Macro-PPARG | ICE2 | 7.97E-18 | 1.122832379 | 0.352 | 0.161 | 1.57E-13 | Tb |
| Macro-PPARG | ZNF10 | 1.18E-26 | 1.118690986 | 0.534 | 0.275 | 2.33E-22 | Tb |
| Macro-PPARG | IL21R1 | 6.96E-24 | 1.118019364 | 0.3 | 0.106 | 1.37E-19 | Tb |
| Macro-PPARG | SULF21 | 4.38E-32 | 1.114048187 | 0.635 | 0.331 | 8.61E-28 | Tb |
| Macro-PPARG | BRCA2 | 8.37E-26 | 1.11081691 | 0.431 | 0.188 | 1.65E-21 | Tb |
| Macro-PPARG | IL32 | 1.10E-23 | 1.107945508 | 0.645 | 0.394 | 2.17E-19 | Tb |
| Macro-PPARG | MT1F | 7.96E-15 | 1.106608158 | 0.286 | 0.129 | 1.57E-10 | Tb |
| Macro-PPARG | PFDN1 | 1.93E-31 | 1.104673331 | 0.618 | 0.306 | 3.80E-27 | Tb |

| | | | | | | | |
|---|---|---|---|---|---|---|---|
| Macro-PPARG | SERPINF1 | 8.57E-32 | 1.104572715 | 0.712 | 0.415 | 1.69E-27 | Tb |
| Macro-PPARG | SULT1C2 | 1.62E-08 | 1.103617107 | 0.276 | 0.156 | 0.000319144 | Tb |
| Macro-PPARG | TRAPPC11 | 1.56E-12 | 1.093163466 | 0.286 | 0.14 | 3.06E-08 | Tb |
| Macro-PPARG | SLC1A4 | 1.89E-18 | 1.089795029 | 0.251 | 0.093 | 3.73E-14 | Tb |
| Macro-PPARG | MT-ATP61 | 4.07E-46 | 1.083641207 | 0.993 | 0.961 | 8.01E-42 | Tb |
| Macro-PPARG | NETO2 | 9.12E-09 | 1.08157731 | 0.377 | 0.239 | 0.00017955 | Tb |
| Macro-PPARG | STEAP1B | 1.77E-14 | 1.081482591 | 0.431 | 0.242 | 3.49E-10 | Tb |
| Macro-PPARG | LATS2 | 1.06E-16 | 1.080562449 | 0.648 | 0.415 | 2.09E-12 | Tb |
| Macro-PPARG | SPRED11 | 4.44E-34 | 1.079268809 | 0.734 | 0.4 | 8.74E-30 | Tb |
| Macro-PPARG | STK19 | 7.59E-10 | 1.078737126 | 0.288 | 0.155 | 1.49E-05 | Tb |
| Macro-PPARG | THBD | 1.42E-28 | 1.075810297 | 0.596 | 0.317 | 2.80E-24 | Tb |
| Macro-PPARG | KLF9 | 5.06E-10 | 1.07286939 | 0.36 | 0.207 | 9.95E-06 | Tb |
| Macro-PPARG | QPCTL | 1.03E-24 | 1.071976732 | 0.296 | 0.1 | 2.02E-20 | Tb |
| Macro-PPARG | ZMAT3 | 4.59E-15 | 1.070673616 | 0.276 | 0.12 | 9.04E-11 | Tb |
| Macro-PPARG | GIMAP4 | 1.37E-35 | 1.069747027 | 0.626 | 0.291 | 2.69E-31 | Tb |
| Macro-PPARG | MICAL2 | 1.32E-17 | 1.068610605 | 0.286 | 0.117 | 2.60E-13 | Tb |
| Macro-PPARG | RFTN11 | 1.84E-29 | 1.068586683 | 0.645 | 0.351 | 3.62E-25 | Tb |
| Macro-PPARG | SATB1-AS1 | 1.54E-20 | 1.067730289 | 0.429 | 0.207 | 3.04E-16 | Tb |
| Macro-PPARG | MED11 | 1.15E-13 | 1.066594533 | 0.266 | 0.12 | 2.27E-09 | Tb |
| Macro-PPARG | CD80 | 2.50E-21 | 1.063496794 | 0.48 | 0.247 | 4.93E-17 | Tb |
| Macro-PPARG | MGME1 | 6.87E-14 | 1.057703869 | 0.424 | 0.232 | 1.35E-09 | Tb |
| Macro-PPARG | SMURF2 | 1.26E-26 | 1.057669441 | 0.724 | 0.463 | 2.48E-22 | Tb |
| Macro-PPARG | FADS11 | 4.03E-22 | 1.057338663 | 0.416 | 0.185 | 7.93E-18 | Tb |
| Macro-PPARG | NFAM11 | 4.46E-30 | 1.056635868 | 0.594 | 0.292 | 8.77E-26 | Tb |
| Macro-PPARG | ORAI2 | 1.90E-20 | 1.055270288 | 0.347 | 0.148 | 3.74E-16 | Tb |
| Macro-PPARG | EML41 | 4.44E-41 | 1.054246293 | 0.865 | 0.555 | 8.74E-37 | Tb |
| Macro-PPARG | TXNIP | 4.47E-43 | 1.052832173 | 0.83 | 0.513 | 8.79E-39 | Tb |
| Macro-PPARG | MSRB1 | 1.66E-17 | 1.051832141 | 0.276 | 0.11 | 3.26E-13 | Tb |
| Macro-PPARG | TRMO | 4.32E-18 | 1.044023831 | 0.387 | 0.182 | 8.51E-14 | Tb |
| Macro-PPARG | IDO2 | 1.54E-14 | 1.042732496 | 0.411 | 0.227 | 3.02E-10 | Tb |
| Macro-PPARG | FHL3 | 3.01E-16 | 1.041348975 | 0.296 | 0.128 | 5.93E-12 | Tb |
| Macro-PPARG | TMEM176A | 2.28E-36 | 1.041174111 | 0.82 | 0.559 | 4.49E-32 | Tb |
| Macro-PPARG | PPARGC1B1 | 7.85E-17 | 1.038050763 | 0.323 | 0.142 | 1.54E-12 | Tb |
| Macro-PPARG | FCGR1B | 4.38E-31 | 1.037189557 | 0.446 | 0.173 | 8.62E-27 | Tb |
| Macro-PPARG | CLEC4A | 6.32E-28 | 1.035364061 | 0.569 | 0.277 | 1.24E-23 | Tb |
| Macro-PPARG | SECTM1 | 1.44E-29 | 1.033419133 | 0.559 | 0.259 | 2.83E-25 | Tb |
| Macro-PPARG | AIM2 | 8.60E-21 | 1.03328172 | 0.31 | 0.121 | 1.69E-16 | Tb |
| Macro-PPARG | TNFRSF211 | 4.54E-30 | 1.032617925 | 0.441 | 0.174 | 8.93E-26 | Tb |
| Macro-PPARG | LRRK2 | 2.52E-17 | 1.031851545 | 0.433 | 0.224 | 4.97E-13 | Tb |
| Macro-PPARG | NUB1 | 4.33E-26 | 1.030860566 | 0.626 | 0.332 | 8.53E-22 | Tb |
| Macro-PPARG | DENND10 | 3.91E-21 | 1.028627085 | 0.399 | 0.178 | 7.70E-17 | Tb |
| Macro-PPARG | APOL6 | 2.45E-38 | 1.020032174 | 0.7 | 0.323 | 4.82E-34 | Tb |
| Macro-PPARG | UBE2L6 | 3.26E-36 | 1.011935595 | 0.667 | 0.312 | 6.42E-32 | Tb |
| Macro-PPARG | ZNF32 | 3.21E-17 | 1.011808486 | 0.3 | 0.126 | 6.32E-13 | Tb |
| Macro-PPARG | IL3RA1 | 6.09E-46 | 1.009922039 | 0.729 | 0.327 | 1.20E-41 | Tb |
| Macro-PPARG | SEMA4A1 | 1.34E-32 | 1.007789317 | 0.771 | 0.491 | 2.64E-28 | Tb |
| Macro-PPARG | ANGEL1 | 4.77E-19 | 1.006970194 | 0.278 | 0.105 | 9.39E-15 | Tb |
| Macro-PPARG | ARSD | 1.03E-16 | 1.005222132 | 0.3 | 0.128 | 2.03E-12 | Tb |
| Macro-PPARG | RGS13 | 2.13E-21 | 1.004481493 | 0.303 | 0.114 | 4.18E-17 | Tb |
| Macro-PPARG | GABPB2 | 6.44E-23 | 1.004185717 | 0.264 | 0.086 | 1.27E-18 | Tb |
| Macro-PPARG | TNFSF13B | 1.35E-47 | 1.003634053 | 0.874 | 0.581 | 2.66E-43 | Tb |
| Macro-PPARG | STAC3 | 1.79E-18 | 1.002971887 | 0.387 | 0.178 | 3.52E-14 | Tb |
| Macro-PPARG | FLVCR2 | 1.60E-20 | 1.001338475 | 0.411 | 0.185 | 3.15E-16 | Tb |
| **Disease-associated genes in Macro-SPP1** | | | | | | | |
| Macro-SPP1 | HS3ST2 | 7.59E-47 | 4.627334097 | 0.35 | 0.058 | 1.49E-42 | Lep-L |

| | | | | | | | |
|---|---|---|---|---|---|---|---|
| Macro-SPP1 | SELENOP | 2.60E-33 | 2.979001311 | 0.36 | 0.099 | 5.11E-29 | Lep-L |
| Macro-SPP1 | NR6A1 | 8.16E-19 | 2.974793103 | 0.275 | 0.099 | 1.61E-14 | Lep-L |
| Macro-SPP1 | TTTY14 | 1.32E-30 | 2.960459047 | 0.257 | 0.045 | 2.60E-26 | Lep-L |
| Macro-SPP1 | STAB1 | 4.93E-22 | 2.933189398 | 0.278 | 0.088 | 9.71E-18 | Lep-L |
| Macro-SPP1 | SDS | 2.23E-29 | 2.773559226 | 0.505 | 0.254 | 4.39E-25 | Lep-L |
| Macro-SPP1 | OTOA | 1.89E-45 | 2.766185158 | 0.556 | 0.212 | 3.71E-41 | Lep-L |
| Macro-SPP1 | SGPP2 | 1.09E-20 | 2.749878768 | 0.287 | 0.096 | 2.15E-16 | Lep-L |
| Macro-SPP1 | TFPI | 1.05E-18 | 2.680161564 | 0.293 | 0.112 | 2.08E-14 | Lep-L |
| Macro-SPP1 | MMP19 | 3.48E-42 | 2.563316316 | 0.695 | 0.42 | 6.85E-38 | Lep-L |
| Macro-SPP1 | CCR7 | 6.04E-10 | 2.542085401 | 0.296 | 0.166 | 1.19E-05 | Lep-L |
| Macro-SPP1 | PLIN2 | 1.76E-92 | 2.341812262 | 0.955 | 0.869 | 3.46E-88 | Lep-L |
| Macro-SPP1 | DDX3Y | 2.49E-55 | 2.269685675 | 0.535 | 0.127 | 4.91E-51 | Lep-L |
| Macro-SPP1 | FOLR2 | 2.71E-31 | 2.256774021 | 0.387 | 0.119 | 5.34E-27 | Lep-L |
| Macro-SPP1 | DHRS3 | 1.86E-17 | 2.239515999 | 0.335 | 0.153 | 3.66E-13 | Lep-L |
| Macro-SPP1 | NBPF19 | 1.36E-23 | 2.239239178 | 0.514 | 0.316 | 2.68E-19 | Lep-L |
| Macro-SPP1 | CFD | 8.72E-48 | 2.224145621 | 0.767 | 0.565 | 1.72E-43 | Lep-L |
| Macro-SPP1 | PDK4 | 5.61E-18 | 2.212712512 | 0.29 | 0.107 | 1.10E-13 | Lep-L |
| Macro-SPP1 | PHACTR1 | 1.66E-42 | 2.207778501 | 0.731 | 0.514 | 3.27E-38 | Lep-L |
| Macro-SPP1 | SERPINB9 | 1.05E-41 | 2.139304822 | 0.822 | 0.672 | 2.07E-37 | Lep-L |
| Macro-SPP1 | CCL20 | 7.49E-27 | 2.086328161 | 0.637 | 0.394 | 1.47E-22 | Lep-L |
| Macro-SPP1 | MS4A14 | 9.65E-10 | 2.08224414 | 0.26 | 0.139 | 1.90E-05 | Lep-L |
| Macro-SPP1 | ITPKB | 6.94E-17 | 2.00222789 | 0.411 | 0.23 | 1.37E-12 | Lep-L |
| Macro-SPP1 | TESK1 | 1.30E-06 | 1.978229462 | 0.338 | 0.266 | 0.025639642 | Lep-L |
| Macro-SPP1 | NUP214 | 8.83E-21 | 1.971700572 | 0.568 | 0.448 | 1.74E-16 | Lep-L |
| Macro-SPP1 | CHST7 | 1.21E-13 | 1.951434887 | 0.308 | 0.152 | 2.39E-09 | Lep-L |
| Macro-SPP1 | IMPA2 | 2.90E-10 | 1.947074737 | 0.308 | 0.184 | 5.71E-06 | Lep-L |
| Macro-SPP1 | MAP4K4 | 3.55E-25 | 1.927642606 | 0.665 | 0.478 | 6.99E-21 | Lep-L |
| Macro-SPP1 | PPARG | 2.52E-16 | 1.922601379 | 0.538 | 0.36 | 4.97E-12 | Lep-L |
| Macro-SPP1 | ADD3 | 4.19E-08 | 1.91425298 | 0.284 | 0.177 | 0.000825094 | Lep-L |
| Macro-SPP1 | LPL | 2.81E-19 | 1.904383603 | 0.438 | 0.229 | 5.53E-15 | Lep-L |
| Macro-SPP1 | IGLC2 | 5.47E-13 | 1.904140058 | 0.278 | 0.123 | 1.08E-08 | Lep-L |
| Macro-SPP1 | CXCL8 | 3.97E-59 | 1.882210884 | 0.952 | 0.83 | 7.81E-55 | Lep-L |
| Macro-SPP1 | EREG | 2.55E-15 | 1.878418969 | 0.65 | 0.489 | 5.02E-11 | Lep-L |
| Macro-SPP1 | CYP27A1 | 2.57E-38 | 1.871484443 | 0.686 | 0.409 | 5.07E-34 | Lep-L |
| Macro-SPP1 | SH3D21 | 9.33E-11 | 1.868418124 | 0.363 | 0.235 | 1.84E-06 | Lep-L |
| Macro-SPP1 | LGMN | 5.24E-26 | 1.864255967 | 0.71 | 0.589 | 1.03E-21 | Lep-L |
| Macro-SPP1 | KLHL21 | 2.33E-18 | 1.856331125 | 0.514 | 0.357 | 4.58E-14 | Lep-L |
| Macro-SPP1 | STMN1 | 1.41E-18 | 1.856031195 | 0.402 | 0.199 | 2.78E-14 | Lep-L |
| Macro-SPP1 | PHLDA1 | 6.10E-24 | 1.83394525 | 0.604 | 0.367 | 1.20E-19 | Lep-L |
| Macro-SPP1 | HES4 | 1.38E-07 | 1.826414399 | 0.372 | 0.275 | 0.002719544 | Lep-L |
| Macro-SPP1 | MS4A6A | 8.31E-34 | 1.822581733 | 0.695 | 0.437 | 1.64E-29 | Lep-L |
| Macro-SPP1 | CXCL3 | 1.73E-50 | 1.821486032 | 0.9 | 0.622 | 3.40E-46 | Lep-L |
| Macro-SPP1 | MAFG | 4.13E-17 | 1.806825054 | 0.571 | 0.461 | 8.13E-13 | Lep-L |
| Macro-SPP1 | OGT | 7.24E-07 | 1.804827363 | 0.366 | 0.288 | 0.014243105 | Lep-L |
| Macro-SPP1 | RNASE1 | 3.31E-09 | 1.792641472 | 0.384 | 0.252 | 6.52E-05 | Lep-L |
| Macro-SPP1 | C5AR1 | 1.22E-52 | 1.758918011 | 0.903 | 0.772 | 2.40E-48 | Lep-L |
| Macro-SPP1 | MFSD2A | 2.29E-19 | 1.75538578 | 0.553 | 0.388 | 4.51E-15 | Lep-L |
| Macro-SPP1 | MARCKS | 3.82E-65 | 1.753567237 | 0.934 | 0.832 | 7.52E-61 | Lep-L |
| Macro-SPP1 | CHPT1 | 6.55E-11 | 1.720175294 | 0.396 | 0.276 | 1.29E-06 | Lep-L |
| Macro-SPP1 | LPCAT1 | 2.52E-12 | 1.691454234 | 0.474 | 0.371 | 4.96E-08 | Lep-L |
| Macro-SPP1 | BCAR3 | 2.64E-08 | 1.688473464 | 0.375 | 0.268 | 0.000519415 | Lep-L |
| Macro-SPP1 | RB1CC1 | 8.80E-16 | 1.681591345 | 0.616 | 0.547 | 1.73E-11 | Lep-L |
| Macro-SPP1 | MIR222HG | 6.38E-09 | 1.672630854 | 0.341 | 0.214 | 0.000125568 | Lep-L |
| Macro-SPP1 | CSGALNACT2 | 5.06E-48 | 1.641535565 | 0.867 | 0.786 | 9.96E-44 | Lep-L |
| Macro-SPP1 | FABP5 | 1.38E-36 | 1.637886562 | 0.906 | 0.844 | 2.71E-32 | Lep-L |

| | | | | | | | |
|---|---|---|---|---|---|---|---|
| Macro-SPP1 | TBC1D8 | 7.70E-17 | 1.620085805 | 0.532 | 0.365 | 1.52E-12 | Lep-L |
| Macro-SPP1 | GPR183 | 2.85E-52 | 1.619008806 | 0.927 | 0.737 | 5.61E-48 | Lep-L |
| Macro-SPP1 | EEPD1 | 2.21E-06 | 1.617116514 | 0.317 | 0.23 | 0.04346006 | Lep-L |
| Macro-SPP1 | PTGER2 | 9.84E-07 | 1.598291949 | 0.356 | 0.278 | 0.01936361 | Lep-L |
| Macro-SPP1 | NBPF14 | 1.93E-07 | 1.595259531 | 0.39 | 0.305 | 0.003789689 | Lep-L |
| Macro-SPP1 | MGLL | 5.66E-53 | 1.590553393 | 0.849 | 0.541 | 1.11E-48 | Lep-L |
| Macro-SPP1 | MIR155HG | 8.28E-26 | 1.588637328 | 0.758 | 0.596 | 1.63E-21 | Lep-L |
| Macro-SPP1 | AP1S3 | 8.23E-07 | 1.559718091 | 0.281 | 0.184 | 0.016203259 | Lep-L |
| Macro-SPP1 | STX3 | 8.14E-14 | 1.558207312 | 0.532 | 0.437 | 1.60E-09 | Lep-L |
| Macro-SPP1 | MIR3142HG | 2.34E-08 | 1.554523428 | 0.378 | 0.26 | 0.000461572 | Lep-L |
| Macro-SPP1 | MAP3K8 | 7.55E-46 | 1.549703236 | 0.915 | 0.847 | 1.49E-41 | Lep-L |
| Macro-SPP1 | SCCPDH | 3.34E-10 | 1.538666443 | 0.396 | 0.275 | 6.57E-06 | Lep-L |
| Macro-SPP1 | SAMSN1 | 1.53E-22 | 1.533526422 | 0.801 | 0.747 | 3.02E-18 | Lep-L |
| Macro-SPP1 | STK4 | 1.85E-15 | 1.526008869 | 0.665 | 0.633 | 3.64E-11 | Lep-L |
| Macro-SPP1 | MS4A7 | 1.32E-45 | 1.520968553 | 0.855 | 0.705 | 2.60E-41 | Lep-L |
| Macro-SPP1 | BASP1 | 9.61E-38 | 1.510440968 | 0.885 | 0.831 | 1.89E-33 | Lep-L |
| Macro-SPP1 | CDC42EP3 | 5.45E-25 | 1.510232741 | 0.752 | 0.615 | 1.07E-20 | Lep-L |
| Macro-SPP1 | BCL2 | 2.24E-19 | 1.495158049 | 0.64 | 0.513 | 4.41E-15 | Lep-L |
| Macro-SPP1 | PIK3R5 | 5.74E-27 | 1.489183042 | 0.746 | 0.601 | 1.13E-22 | Lep-L |
| Macro-SPP1 | LIPA | 7.02E-27 | 1.484716273 | 0.746 | 0.623 | 1.38E-22 | Lep-L |
| Macro-SPP1 | IL10 | 1.52E-07 | 1.470829437 | 0.272 | 0.16 | 0.00298942 | Lep-L |
| Macro-SPP1 | RILPL2 | 4.94E-32 | 1.469289031 | 0.87 | 0.879 | 9.72E-28 | Lep-L |
| Macro-SPP1 | ETV3 | 3.43E-15 | 1.460352543 | 0.586 | 0.491 | 6.75E-11 | Lep-L |
| Macro-SPP1 | RARA | 1.20E-06 | 1.458584886 | 0.435 | 0.394 | 0.023696353 | Lep-L |
| Macro-SPP1 | CXCL2 | 5.66E-43 | 1.454012999 | 0.943 | 0.753 | 1.11E-38 | Lep-L |
| Macro-SPP1 | RNF144B | 4.50E-11 | 1.444383859 | 0.725 | 0.748 | 8.86E-07 | Lep-L |
| Macro-SPP1 | GPNMB | 6.50E-51 | 1.444102717 | 0.927 | 0.859 | 1.28E-46 | Lep-L |
| Macro-SPP1 | STK40 | 1.68E-11 | 1.421928448 | 0.517 | 0.432 | 3.31E-07 | Lep-L |
| Macro-SPP1 | APOC1 | 6.09E-33 | 1.410839088 | 0.921 | 0.907 | 1.20E-28 | Lep-L |
| Macro-SPP1 | STAM | 1.56E-06 | 1.405156134 | 0.429 | 0.38 | 0.03064921 | Lep-L |
| Macro-SPP1 | ARID3A | 6.38E-08 | 1.399745507 | 0.444 | 0.377 | 0.001254978 | Lep-L |
| Macro-SPP1 | CXCR4 | 4.92E-23 | 1.382746833 | 0.813 | 0.795 | 9.68E-19 | Lep-L |
| Macro-SPP1 | FABP4 | 3.66E-34 | 1.376849438 | 0.341 | 0.075 | 7.19E-30 | Lep-L |
| Macro-SPP1 | MAFF | 5.47E-20 | 1.366087111 | 0.737 | 0.691 | 1.08E-15 | Lep-L |
| Macro-SPP1 | CD302 | 2.84E-21 | 1.359012941 | 0.659 | 0.542 | 5.59E-17 | Lep-L |
| Macro-SPP1 | UGCG | 7.02E-12 | 1.335067696 | 0.653 | 0.66 | 1.38E-07 | Lep-L |
| Macro-SPP1 | MSR1 | 4.02E-16 | 1.333498808 | 0.713 | 0.604 | 7.91E-12 | Lep-L |
| Macro-SPP1 | SNHG12 | 4.86E-13 | 1.318084941 | 0.631 | 0.584 | 9.57E-09 | Lep-L |
| Macro-SPP1 | APOE | 8.19E-41 | 1.31553015 | 0.973 | 0.898 | 1.61E-36 | Lep-L |
| Macro-SPP1 | GADD45B | 9.56E-11 | 1.29789197 | 0.934 | 0.959 | 1.88E-06 | Lep-L |
| Macro-SPP1 | PDXDC1 | 5.31E-09 | 1.289825178 | 0.52 | 0.476 | 0.000104439 | Lep-L |
| Macro-SPP1 | CHIC2 | 2.66E-14 | 1.288089938 | 0.637 | 0.596 | 5.23E-10 | Lep-L |
| Macro-SPP1 | NOTCH2NLC | 1.62E-09 | 1.285148644 | 0.538 | 0.466 | 3.18E-05 | Lep-L |
| Macro-SPP1 | RASGRP3 | 2.91E-12 | 1.281045616 | 0.634 | 0.545 | 5.74E-08 | Lep-L |
| Macro-SPP1 | TOB2 | 4.31E-07 | 1.275313162 | 0.438 | 0.383 | 0.008488411 | Lep-L |
| Macro-SPP1 | RAB9A | 2.42E-07 | 1.274029727 | 0.498 | 0.49 | 0.004756711 | Lep-L |
| Macro-SPP1 | KMO | 4.97E-23 | 1.270354923 | 0.785 | 0.728 | 9.79E-19 | Lep-L |
| Macro-SPP1 | DENND4C | 1.74E-07 | 1.267742017 | 0.529 | 0.494 | 0.00342869 | Lep-L |
| Macro-SPP1 | IER3 | 2.49E-40 | 1.26194085 | 0.964 | 0.934 | 4.91E-36 | Lep-L |
| Macro-SPP1 | DDAH2 | 1.22E-18 | 1.254562972 | 0.656 | 0.575 | 2.40E-14 | Lep-L |
| Macro-SPP1 | ATF4 | 5.20E-22 | 1.254407961 | 0.731 | 0.698 | 1.02E-17 | Lep-L |
| Macro-SPP1 | TRAF1 | 5.45E-10 | 1.248578663 | 0.653 | 0.626 | 1.07E-05 | Lep-L |
| Macro-SPP1 | AQP9 | 4.53E-11 | 1.247183461 | 0.792 | 0.824 | 8.92E-07 | Lep-L |
| Macro-SPP1 | CKS2 | 1.98E-23 | 1.244798923 | 0.81 | 0.789 | 3.89E-19 | Lep-L |
| Macro-SPP1 | SMS | 2.06E-23 | 1.235015746 | 0.764 | 0.763 | 4.05E-19 | Lep-L |

| | | | | | | | |
|---|---|---|---|---|---|---|---|
| Macro-SPP1 | PABPN1 | 8.11E-14 | 1.231393671 | 0.616 | 0.554 | 1.60E-09 | Lep-L |
| Macro-SPP1 | MAP2K3 | 6.66E-24 | 1.230853407 | 0.813 | 0.832 | 1.31E-19 | Lep-L |
| Macro-SPP1 | GRASP | 6.38E-10 | 1.222663743 | 0.489 | 0.383 | 1.26E-05 | Lep-L |
| Macro-SPP1 | TSC22D2 | 2.03E-21 | 1.220270469 | 0.804 | 0.713 | 3.99E-17 | Lep-L |
| Macro-SPP1 | SNHG7 | 2.16E-11 | 1.219161375 | 0.644 | 0.682 | 4.24E-07 | Lep-L |
| Macro-SPP1 | MXD1 | 2.01E-16 | 1.217207555 | 0.801 | 0.784 | 3.96E-12 | Lep-L |
| Macro-SPP1 | PDE7A | 6.77E-08 | 1.216441315 | 0.526 | 0.492 | 0.001333185 | Lep-L |
| Macro-SPP1 | HLA-DRB5 | 1.24E-30 | 1.213119829 | 0.843 | 0.746 | 2.43E-26 | Lep-L |
| Macro-SPP1 | HIVEP2 | 2.58E-22 | 1.208329856 | 0.758 | 0.636 | 5.08E-18 | Lep-L |
| Macro-SPP1 | REV3L | 9.21E-12 | 1.204320283 | 0.634 | 0.58 | 1.81E-07 | Lep-L |
| Macro-SPP1 | BCL2A1 | 1.44E-50 | 1.201884654 | 0.967 | 0.986 | 2.84E-46 | Lep-L |
| Macro-SPP1 | FOSB | 7.25E-11 | 1.201232439 | 0.77 | 0.797 | 1.43E-06 | Lep-L |
| Macro-SPP1 | PELI1 | 5.71E-15 | 1.196791596 | 0.674 | 0.57 | 1.12E-10 | Lep-L |
| Macro-SPP1 | DEDD2 | 9.41E-07 | 1.194172874 | 0.589 | 0.634 | 0.018529274 | Lep-L |
| Macro-SPP1 | VAT1 | 3.29E-11 | 1.192550217 | 0.616 | 0.597 | 6.47E-07 | Lep-L |
| Macro-SPP1 | SP3 | 3.89E-08 | 1.191951462 | 0.526 | 0.493 | 0.000766198 | Lep-L |
| Macro-SPP1 | ERGIC1 | 8.05E-10 | 1.191276325 | 0.547 | 0.514 | 1.59E-05 | Lep-L |
| Macro-SPP1 | SETD5 | 6.23E-08 | 1.191274959 | 0.55 | 0.534 | 0.001225675 | Lep-L |
| Macro-SPP1 | MGAT1 | 3.39E-53 | 1.161955083 | 0.927 | 0.932 | 6.67E-49 | Lep-L |
| Macro-SPP1 | ATP6V1C1 | 2.25E-12 | 1.159957455 | 0.668 | 0.696 | 4.44E-08 | Lep-L |
| Macro-SPP1 | EIF4E | 3.83E-18 | 1.155753714 | 0.743 | 0.69 | 7.53E-14 | Lep-L |
| Macro-SPP1 | OPN3 | 1.64E-09 | 1.15417521 | 0.589 | 0.565 | 3.22E-05 | Lep-L |
| Macro-SPP1 | CHD2 | 1.01E-27 | 1.154073519 | 0.876 | 0.846 | 1.99E-23 | Lep-L |
| Macro-SPP1 | PNPLA8 | 3.45E-20 | 1.153650954 | 0.813 | 0.818 | 6.79E-16 | Lep-L |
| Macro-SPP1 | HIF1A | 6.41E-34 | 1.15144652 | 0.891 | 0.878 | 1.26E-29 | Lep-L |
| Macro-SPP1 | ZFP36L1 | 9.26E-13 | 1.148018651 | 0.828 | 0.89 | 1.82E-08 | Lep-L |
| Macro-SPP1 | FAM107B | 4.42E-24 | 1.142427757 | 0.849 | 0.82 | 8.71E-20 | Lep-L |
| Macro-SPP1 | B4GALT5 | 1.12E-13 | 1.135838552 | 0.692 | 0.665 | 2.21E-09 | Lep-L |
| Macro-SPP1 | IFNAR2 | 2.95E-10 | 1.131209304 | 0.622 | 0.645 | 5.80E-06 | Lep-L |
| Macro-SPP1 | SNHG5 | 2.11E-20 | 1.127523172 | 0.779 | 0.724 | 4.15E-16 | Lep-L |
| Macro-SPP1 | CCDC107 | 7.14E-19 | 1.126181768 | 0.734 | 0.747 | 1.40E-14 | Lep-L |
| Macro-SPP1 | BNIP3L | 1.72E-18 | 1.125867051 | 0.879 | 0.929 | 3.39E-14 | Lep-L |
| Macro-SPP1 | CDKN1A | 5.38E-24 | 1.112603318 | 0.897 | 0.83 | 1.06E-19 | Lep-L |
| Macro-SPP1 | PABPC4 | 1.33E-25 | 1.108244196 | 0.807 | 0.805 | 2.61E-21 | Lep-L |
| Macro-SPP1 | FNIP2 | 3.40E-42 | 1.108030271 | 0.97 | 0.933 | 6.70E-38 | Lep-L |
| Macro-SPP1 | HECTD1 | 6.85E-13 | 1.107166038 | 0.644 | 0.606 | 1.35E-08 | Lep-L |
| Macro-SPP1 | ELF2 | 3.88E-11 | 1.093279517 | 0.659 | 0.656 | 7.63E-07 | Lep-L |
| Macro-SPP1 | MIR22HG | 8.13E-09 | 1.091244255 | 0.568 | 0.545 | 0.00016008 | Lep-L |
| Macro-SPP1 | MYO9B | 1.85E-29 | 1.091093263 | 0.873 | 0.844 | 3.64E-25 | Lep-L |
| Macro-SPP1 | SNTB1 | 1.47E-06 | 1.090850453 | 0.544 | 0.515 | 0.029004746 | Lep-L |
| Macro-SPP1 | TMEM123 | 1.63E-07 | 1.081190076 | 0.665 | 0.726 | 0.00320695 | Lep-L |
| Macro-SPP1 | ABCA1 | 4.88E-16 | 1.075928226 | 0.867 | 0.874 | 9.61E-12 | Lep-L |
| Macro-SPP1 | EHD1 | 5.16E-11 | 1.074618286 | 0.689 | 0.675 | 1.02E-06 | Lep-L |
| Macro-SPP1 | MAP7D1 | 1.71E-08 | 1.074051235 | 0.601 | 0.641 | 0.00033632 | Lep-L |
| Macro-SPP1 | ZFP91 | 1.44E-07 | 1.071865366 | 0.583 | 0.614 | 0.002842254 | Lep-L |
| Macro-SPP1 | CXCL1 | 1.32E-11 | 1.059495069 | 0.465 | 0.281 | 2.60E-07 | Lep-L |
| Macro-SPP1 | PDE8A | 5.59E-14 | 1.056593533 | 0.698 | 0.612 | 1.10E-09 | Lep-L |
| Macro-SPP1 | BRD2 | 7.61E-13 | 1.055378119 | 0.792 | 0.823 | 1.50E-08 | Lep-L |
| Macro-SPP1 | ZC3H12C | 1.97E-13 | 1.048875084 | 0.631 | 0.53 | 3.88E-09 | Lep-L |
| Macro-SPP1 | PEAK1 | 3.01E-08 | 1.048660911 | 0.613 | 0.596 | 0.00059223 | Lep-L |
| Macro-SPP1 | BANP | 1.13E-06 | 1.04722619 | 0.589 | 0.569 | 0.022191411 | Lep-L |
| Macro-SPP1 | BTG1 | 1.39E-32 | 1.047131772 | 0.967 | 0.988 | 2.74E-28 | Lep-L |
| Macro-SPP1 | SSH2 | 1.25E-10 | 1.046792642 | 0.628 | 0.6 | 2.46E-06 | Lep-L |
| Macro-SPP1 | SAT1 | 4.92E-47 | 1.042762441 | 0.994 | 0.998 | 9.69E-43 | Lep-L |
| Macro-SPP1 | CD300A | 7.14E-10 | 1.040130178 | 0.553 | 0.48 | 1.41E-05 | Lep-L |

| | | | | | | | |
|---|---|---|---|---|---|---|---|
| Macro-SPP1 | BTG3 | 1.35E-08 | 1.035120065 | 0.631 | 0.642 | 0.000266572 | Lep-L |
| Macro-SPP1 | PPP1R15B | 4.76E-16 | 1.032432895 | 0.767 | 0.751 | 9.37E-12 | Lep-L |
| Macro-SPP1 | KDM6B | 5.01E-17 | 1.029825414 | 0.782 | 0.723 | 9.87E-13 | Lep-L |
| Macro-SPP1 | WTAP | 1.74E-33 | 1.02856641 | 0.94 | 0.957 | 3.43E-29 | Lep-L |
| Macro-SPP1 | NOTCH2 | 3.03E-09 | 1.027871281 | 0.689 | 0.723 | 5.97E-05 | Lep-L |
| Macro-SPP1 | DNAJB4 | 7.75E-07 | 1.026978372 | 0.453 | 0.372 | 0.015255734 | Lep-L |
| Macro-SPP1 | CREM | 1.42E-12 | 1.024361002 | 0.776 | 0.803 | 2.80E-08 | Lep-L |
| Macro-SPP1 | PPIF | 2.07E-18 | 1.019388797 | 0.852 | 0.892 | 4.07E-14 | Lep-L |
| Macro-SPP1 | CD36 | 4.67E-48 | 1.014857224 | 0.468 | 0.107 | 9.19E-44 | Lep-L |
| Macro-SPP1 | UBE2R2 | 5.20E-21 | 1.005923204 | 0.785 | 0.756 | 1.02E-16 | Lep-L |
| Macro-SPP1 | OTUD1 | 2.47E-10 | 1.00343895 | 0.671 | 0.624 | 4.87E-06 | Lep-L |
| Macro-SPP1 | GP1BA | 3.44E-89 | 4.259429569 | 0.589 | 0.06 | 6.78E-85 | Lep-T |
| Macro-SPP1 | NCAPG2 | 2.19E-40 | 3.862170254 | 0.48 | 0.113 | 4.31E-36 | Lep-T |
| Macro-SPP1 | CENPU | 1.67E-45 | 3.391230028 | 0.503 | 0.107 | 3.28E-41 | Lep-T |
| Macro-SPP1 | SLC22A14 | 7.48E-36 | 3.332496114 | 0.28 | 0.033 | 1.47E-31 | Lep-T |
| Macro-SPP1 | SIGLEC15 | 1.99E-35 | 3.099721024 | 0.366 | 0.064 | 3.93E-31 | Lep-T |
| Macro-SPP1 | RASGRF1 | 3.99E-33 | 2.975975362 | 0.4 | 0.083 | 7.85E-29 | Lep-T |
| Macro-SPP1 | CCDC138 | 2.09E-26 | 2.973844302 | 0.617 | 0.306 | 4.12E-22 | Lep-T |
| Macro-SPP1 | NAPA-AS1 | 4.45E-31 | 2.773877789 | 0.269 | 0.035 | 8.75E-27 | Lep-T |
| Macro-SPP1 | HIST1H3D | 5.57E-28 | 2.658365362 | 0.297 | 0.051 | 1.10E-23 | Lep-T |
| Macro-SPP1 | INPP5A | 6.09E-26 | 2.53689642 | 0.463 | 0.149 | 1.20E-21 | Lep-T |
| Macro-SPP1 | NARS2 | 3.32E-31 | 2.321957311 | 0.64 | 0.257 | 6.54E-27 | Lep-T |
| Macro-SPP1 | RASGRP1 | 3.94E-44 | 2.287705922 | 0.657 | 0.181 | 7.75E-40 | Lep-T |
| Macro-SPP1 | CALD1 | 2.52E-27 | 2.270260725 | 0.257 | 0.037 | 4.96E-23 | Lep-T |
| Macro-SPP1 | SLC22A1 | 2.42E-25 | 2.253681552 | 0.337 | 0.074 | 4.76E-21 | Lep-T |
| Macro-SPP1 | RIPOR3 | 2.30E-34 | 2.235143896 | 0.446 | 0.099 | 4.52E-30 | Lep-T |
| Macro-SPP1 | NT5DC4 | 1.26E-28 | 2.225887963 | 0.366 | 0.078 | 2.48E-24 | Lep-T |
| Macro-SPP1 | DHCR24 | 7.10E-37 | 2.186266796 | 0.589 | 0.179 | 1.40E-32 | Lep-T |
| Macro-SPP1 | PIP5K1B | 1.11E-18 | 2.168073546 | 0.309 | 0.081 | 2.19E-14 | Lep-T |
| Macro-SPP1 | CXCL14 | 5.30E-44 | 2.164123998 | 0.423 | 0.061 | 1.04E-39 | Lep-T |
| Macro-SPP1 | NDRG2 | 8.34E-45 | 2.139770367 | 0.651 | 0.186 | 1.64E-40 | Lep-T |
| Macro-SPP1 | HIST1H2BC | 1.11E-21 | 2.130109031 | 0.331 | 0.083 | 2.18E-17 | Lep-T |
| Macro-SPP1 | PRKAR2B | 2.90E-46 | 2.096385204 | 0.703 | 0.21 | 5.70E-42 | Lep-T |
| Macro-SPP1 | NEK1 | 8.13E-12 | 2.077607412 | 0.52 | 0.285 | 1.60E-07 | Lep-T |
| Macro-SPP1 | COL6A1 | 1.34E-26 | 2.066240655 | 0.514 | 0.169 | 2.64E-22 | Lep-T |
| Macro-SPP1 | TLN2 | 1.09E-12 | 2.059466149 | 0.309 | 0.107 | 2.15E-08 | Lep-T |
| Macro-SPP1 | IL21R | 3.49E-31 | 2.047511271 | 0.474 | 0.122 | 6.88E-27 | Lep-T |
| Macro-SPP1 | HMGN5 | 2.79E-16 | 2.045840288 | 0.423 | 0.164 | 5.50E-12 | Lep-T |
| Macro-SPP1 | PLCL1 | 7.76E-28 | 2.034203124 | 0.686 | 0.301 | 1.53E-23 | Lep-T |
| Macro-SPP1 | CCL19 | 1.69E-41 | 2.030761012 | 0.451 | 0.078 | 3.33E-37 | Lep-T |
| Macro-SPP1 | RAMP1 | 2.13E-23 | 2.030628389 | 0.286 | 0.056 | 4.20E-19 | Lep-T |
| Macro-SPP1 | HIST2H2AC | 5.01E-42 | 2.011783797 | 0.554 | 0.125 | 9.87E-38 | Lep-T |
| Macro-SPP1 | ITGAD | 1.51E-18 | 2.000444581 | 0.263 | 0.061 | 2.97E-14 | Lep-T |
| Macro-SPP1 | ASB6 | 3.49E-23 | 1.995665335 | 0.417 | 0.119 | 6.87E-19 | Lep-T |
| Macro-SPP1 | ZG16B | 9.80E-30 | 1.96810737 | 0.417 | 0.096 | 1.93E-25 | Lep-T |
| Macro-SPP1 | CPSF1 | 1.95E-06 | 1.916453433 | 0.251 | 0.112 | 0.038406651 | Lep-T |
| Macro-SPP1 | FKBP5 | 7.14E-32 | 1.911208443 | 0.886 | 0.652 | 1.41E-27 | Lep-T |
| Macro-SPP1 | DIRC3 | 2.91E-15 | 1.900100134 | 0.377 | 0.143 | 5.73E-11 | Lep-T |
| Macro-SPP1 | NUP107 | 5.58E-12 | 1.896688267 | 0.463 | 0.23 | 1.10E-07 | Lep-T |
| Macro-SPP1 | PRIMPOL | 1.73E-18 | 1.858625211 | 0.291 | 0.076 | 3.40E-14 | Lep-T |
| Macro-SPP1 | CD1E | 4.39E-18 | 1.842936305 | 0.251 | 0.057 | 8.64E-14 | Lep-T |
| Macro-SPP1 | SPARC | 4.30E-34 | 1.838560744 | 0.749 | 0.355 | 8.47E-30 | Lep-T |
| Macro-SPP1 | ICA1 | 1.90E-25 | 1.826582468 | 0.377 | 0.09 | 3.73E-21 | Lep-T |
| Macro-SPP1 | FADS1 | 4.13E-36 | 1.816520277 | 0.777 | 0.338 | 8.13E-32 | Lep-T |
| Macro-SPP1 | MGP | 1.58E-37 | 1.813896337 | 0.343 | 0.046 | 3.12E-33 | Lep-T |

| | | | | | | | |
|---|---|---|---|---|---|---|---|
| Macro-SPP1 | RTN1 | 2.12E-36 | 1.773609323 | 0.823 | 0.382 | 4.18E-32 | Lep-T |
| Macro-SPP1 | UST | 2.15E-12 | 1.773019566 | 0.44 | 0.205 | 4.23E-08 | Lep-T |
| Macro-SPP1 | FGFR1 | 4.20E-20 | 1.772981904 | 0.537 | 0.215 | 8.28E-16 | Lep-T |
| Macro-SPP1 | DCN | 1.16E-28 | 1.747147166 | 0.451 | 0.107 | 2.28E-24 | Lep-T |
| Macro-SPP1 | VCAM1 | 6.02E-15 | 1.744050427 | 0.32 | 0.102 | 1.18E-10 | Lep-T |
| Macro-SPP1 | CLECL1 | 4.03E-16 | 1.738632201 | 0.389 | 0.138 | 7.94E-12 | Lep-T |
| Macro-SPP1 | SLC16A6 | 1.65E-21 | 1.735061989 | 0.554 | 0.225 | 3.25E-17 | Lep-T |
| Macro-SPP1 | ADGRD1 | 7.22E-17 | 1.706620871 | 0.263 | 0.066 | 1.42E-12 | Lep-T |
| Macro-SPP1 | SCAT1 | 4.64E-14 | 1.696743831 | 0.274 | 0.081 | 9.14E-10 | Lep-T |
| Macro-SPP1 | ESYT2 | 1.04E-22 | 1.691770885 | 0.834 | 0.701 | 2.04E-18 | Lep-T |
| Macro-SPP1 | TFDP2 | 1.45E-22 | 1.691698522 | 0.669 | 0.364 | 2.85E-18 | Lep-T |
| Macro-SPP1 | CUL9 | 2.30E-10 | 1.685680081 | 0.274 | 0.098 | 4.53E-06 | Lep-T |
| Macro-SPP1 | RAB11FIP3 | 4.81E-13 | 1.677637996 | 0.274 | 0.088 | 9.46E-09 | Lep-T |
| Macro-SPP1 | DEPDC5 | 5.63E-14 | 1.674618909 | 0.309 | 0.098 | 1.11E-09 | Lep-T |
| Macro-SPP1 | SPN | 4.40E-39 | 1.670382676 | 0.811 | 0.379 | 8.66E-35 | Lep-T |
| Macro-SPP1 | MT-CO3 | 2.18E-18 | 1.663927821 | 0.966 | 0.985 | 4.29E-14 | Lep-T |
| Macro-SPP1 | TTC7B | 1.37E-27 | 1.663775813 | 0.657 | 0.279 | 2.70E-23 | Lep-T |
| Macro-SPP1 | FZR1 | 1.50E-20 | 1.663163783 | 0.497 | 0.181 | 2.96E-16 | Lep-T |
| Macro-SPP1 | HSD11B1 | 3.97E-33 | 1.661671533 | 0.874 | 0.523 | 7.81E-29 | Lep-T |
| Macro-SPP1 | CYTH3 | 3.46E-20 | 1.659776499 | 0.526 | 0.211 | 6.81E-16 | Lep-T |
| Macro-SPP1 | TGFB2 | 1.54E-17 | 1.643113235 | 0.297 | 0.081 | 3.04E-13 | Lep-T |
| Macro-SPP1 | FADS2 | 8.36E-23 | 1.626926343 | 0.497 | 0.17 | 1.65E-18 | Lep-T |
| Macro-SPP1 | CHRNE | 1.20E-22 | 1.621051856 | 0.314 | 0.07 | 2.35E-18 | Lep-T |
| Macro-SPP1 | RNF213-AS1 | 2.09E-18 | 1.613618045 | 0.28 | 0.067 | 4.11E-14 | Lep-T |
| Macro-SPP1 | ACTA2 | 3.24E-26 | 1.609189997 | 0.331 | 0.066 | 6.38E-22 | Lep-T |
| Macro-SPP1 | BACE1 | 7.55E-10 | 1.607898712 | 0.286 | 0.111 | 1.49E-05 | Lep-T |
| Macro-SPP1 | POC1B | 2.20E-09 | 1.601542046 | 0.474 | 0.267 | 4.34E-05 | Lep-T |
| Macro-SPP1 | LAMP3 | 5.11E-28 | 1.590478631 | 0.514 | 0.149 | 1.01E-23 | Lep-T |
| Macro-SPP1 | GSTM4 | 4.65E-21 | 1.582306346 | 0.606 | 0.262 | 9.16E-17 | Lep-T |
| Macro-SPP1 | PPP5C | 2.29E-16 | 1.58038269 | 0.394 | 0.131 | 4.51E-12 | Lep-T |
| Macro-SPP1 | TVP23A | 1.46E-18 | 1.576433212 | 0.434 | 0.152 | 2.87E-14 | Lep-T |
| Macro-SPP1 | CRTAM | 7.03E-30 | 1.567665013 | 0.571 | 0.179 | 1.38E-25 | Lep-T |
| Macro-SPP1 | SPOUT1 | 9.52E-10 | 1.564197266 | 0.257 | 0.091 | 1.87E-05 | Lep-T |
| Macro-SPP1 | ZNF618 | 1.29E-20 | 1.555624736 | 0.491 | 0.174 | 2.55E-16 | Lep-T |
| Macro-SPP1 | SCLY | 2.35E-15 | 1.552664819 | 0.274 | 0.076 | 4.62E-11 | Lep-T |
| Macro-SPP1 | PTGDS | 7.23E-12 | 1.546675636 | 0.474 | 0.21 | 1.42E-07 | Lep-T |
| Macro-SPP1 | DDX28 | 1.19E-16 | 1.545617913 | 0.263 | 0.065 | 2.34E-12 | Lep-T |
| Macro-SPP1 | BEST1 | 8.32E-10 | 1.530374659 | 0.463 | 0.239 | 1.64E-05 | Lep-T |
| Macro-SPP1 | CYSLTR1 | 4.05E-14 | 1.529284644 | 0.326 | 0.105 | 7.98E-10 | Lep-T |
| Macro-SPP1 | ELOVL5 | 6.71E-23 | 1.528435527 | 0.874 | 0.723 | 1.32E-18 | Lep-T |
| Macro-SPP1 | ABR | 3.61E-23 | 1.524233328 | 0.897 | 0.816 | 7.11E-19 | Lep-T |
| Macro-SPP1 | ANKRD36B | 1.68E-12 | 1.523448559 | 0.257 | 0.08 | 3.31E-08 | Lep-T |
| Macro-SPP1 | KRTDAP | 3.80E-73 | 1.519392179 | 0.629 | 0.086 | 7.48E-69 | Lep-T |
| Macro-SPP1 | PTPRF | 9.70E-15 | 1.513549441 | 0.343 | 0.118 | 1.91E-10 | Lep-T |
| Macro-SPP1 | GAB2 | 4.56E-19 | 1.510738887 | 0.829 | 0.718 | 8.98E-15 | Lep-T |
| Macro-SPP1 | NFATC1 | 3.71E-27 | 1.498499671 | 0.766 | 0.389 | 7.31E-23 | Lep-T |
| Macro-SPP1 | MT-ATP6 | 1.63E-17 | 1.494284289 | 0.937 | 0.991 | 3.20E-13 | Lep-T |
| Macro-SPP1 | CYP27B1 | 3.57E-27 | 1.494267109 | 0.771 | 0.377 | 7.02E-23 | Lep-T |
| Macro-SPP1 | ALOX15B | 2.49E-19 | 1.491298318 | 0.509 | 0.196 | 4.89E-15 | Lep-T |
| Macro-SPP1 | PMEPA1 | 4.69E-17 | 1.490148133 | 0.577 | 0.282 | 9.23E-13 | Lep-T |
| Macro-SPP1 | DMKN | 8.46E-56 | 1.480580868 | 0.52 | 0.074 | 1.67E-51 | Lep-T |
| Macro-SPP1 | SCRN2 | 6.17E-17 | 1.479521801 | 0.303 | 0.081 | 1.21E-12 | Lep-T |
| Macro-SPP1 | SERINC5 | 1.62E-12 | 1.477287446 | 0.731 | 0.509 | 3.19E-08 | Lep-T |
| Macro-SPP1 | CENPN | 7.34E-15 | 1.477124944 | 0.36 | 0.12 | 1.44E-10 | Lep-T |
| Macro-SPP1 | ABCC4 | 4.55E-14 | 1.470925996 | 0.297 | 0.093 | 8.96E-10 | Lep-T |

| | | | | | | | |
|---|---|---|---|---|---|---|---|
| Macro-SPP1 | MMS19 | 2.33E-11 | 1.470316261 | 0.32 | 0.119 | 4.58E-07 | Lep-T |
| Macro-SPP1 | POLA1 | 2.44E-10 | 1.470136066 | 0.314 | 0.125 | 4.80E-06 | Lep-T |
| Macro-SPP1 | POMZP3 | 4.18E-10 | 1.468258652 | 0.4 | 0.184 | 8.24E-06 | Lep-T |
| Macro-SPP1 | MTHFD1L | 1.01E-10 | 1.463732728 | 0.383 | 0.17 | 1.99E-06 | Lep-T |
| Macro-SPP1 | PLA2G6 | 1.47E-13 | 1.463101947 | 0.389 | 0.138 | 2.89E-09 | Lep-T |
| Macro-SPP1 | PTPN9 | 1.18E-25 | 1.462666167 | 0.709 | 0.315 | 2.33E-21 | Lep-T |
| Macro-SPP1 | HIST1H1E | 7.18E-14 | 1.461916842 | 0.429 | 0.174 | 1.41E-09 | Lep-T |
| Macro-SPP1 | CHAF1A | 8.97E-26 | 1.455209472 | 0.611 | 0.224 | 1.77E-21 | Lep-T |
| Macro-SPP1 | NOTCH2NLA | 1.99E-10 | 1.454340629 | 0.269 | 0.099 | 3.91E-06 | Lep-T |
| Macro-SPP1 | CCL4L2 | 5.71E-29 | 1.454310463 | 0.937 | 0.699 | 1.12E-24 | Lep-T |
| Macro-SPP1 | TLR8 | 3.46E-21 | 1.44759213 | 0.697 | 0.341 | 6.81E-17 | Lep-T |
| Macro-SPP1 | COL3A1 | 2.46E-26 | 1.434019402 | 0.463 | 0.122 | 4.85E-22 | Lep-T |
| Macro-SPP1 | ACOT7 | 9.39E-35 | 1.429609312 | 0.869 | 0.45 | 1.85E-30 | Lep-T |
| Macro-SPP1 | P2RY13 | 1.03E-26 | 1.427918759 | 0.629 | 0.23 | 2.02E-22 | Lep-T |
| Macro-SPP1 | BDH1 | 5.71E-12 | 1.422722024 | 0.331 | 0.124 | 1.12E-07 | Lep-T |
| Macro-SPP1 | AVPI1 | 8.08E-22 | 1.417206468 | 0.88 | 0.679 | 1.59E-17 | Lep-T |
| Macro-SPP1 | ZBTB46 | 1.45E-14 | 1.40861427 | 0.4 | 0.151 | 2.86E-10 | Lep-T |
| Macro-SPP1 | FMN1 | 2.01E-10 | 1.405210355 | 0.749 | 0.632 | 3.96E-06 | Lep-T |
| Macro-SPP1 | ZNF366 | 4.05E-18 | 1.396457123 | 0.331 | 0.091 | 7.98E-14 | Lep-T |
| Macro-SPP1 | SEPTIN11 | 3.81E-30 | 1.391719601 | 0.863 | 0.55 | 7.50E-26 | Lep-T |
| Macro-SPP1 | DSTYK | 5.86E-15 | 1.384733114 | 0.394 | 0.139 | 1.15E-10 | Lep-T |
| Macro-SPP1 | RAP1GAP2 | 2.30E-15 | 1.383924153 | 0.371 | 0.131 | 4.53E-11 | Lep-T |
| Macro-SPP1 | ACAD9 | 2.69E-10 | 1.38296196 | 0.394 | 0.176 | 5.29E-06 | Lep-T |
| Macro-SPP1 | ATP5PB | 3.79E-44 | 1.374092336 | 0.931 | 0.619 | 7.45E-40 | Lep-T |
| Macro-SPP1 | SH3PXD2A | 6.29E-20 | 1.372615886 | 0.554 | 0.222 | 1.24E-15 | Lep-T |
| Macro-SPP1 | BCL11A | 1.19E-21 | 1.366892208 | 0.754 | 0.454 | 2.34E-17 | Lep-T |
| Macro-SPP1 | COL8A2 | 1.64E-18 | 1.364038911 | 0.417 | 0.135 | 3.22E-14 | Lep-T |
| Macro-SPP1 | SPDYA | 5.56E-12 | 1.362802039 | 0.411 | 0.189 | 1.09E-07 | Lep-T |
| Macro-SPP1 | COL6A2 | 1.94E-11 | 1.359734565 | 0.349 | 0.14 | 3.81E-07 | Lep-T |
| Macro-SPP1 | HSPBP1 | 1.94E-08 | 1.358827074 | 0.269 | 0.108 | 0.000381958 | Lep-T |
| Macro-SPP1 | XPC | 2.84E-23 | 1.357353962 | 0.663 | 0.301 | 5.58E-19 | Lep-T |
| Macro-SPP1 | MB21D2 | 6.35E-16 | 1.35721666 | 0.669 | 0.355 | 1.25E-11 | Lep-T |
| Macro-SPP1 | NAV1 | 2.85E-17 | 1.353980058 | 0.503 | 0.204 | 5.60E-13 | Lep-T |
| Macro-SPP1 | SLC8A1-AS1 | 1.09E-20 | 1.347259771 | 0.503 | 0.183 | 2.14E-16 | Lep-T |
| Macro-SPP1 | POLR2G | 3.68E-31 | 1.34538985 | 0.811 | 0.399 | 7.24E-27 | Lep-T |
| Macro-SPP1 | RFTN1 | 1.79E-29 | 1.344863707 | 0.903 | 0.586 | 3.52E-25 | Lep-T |
| Macro-SPP1 | SLC35E4 | 2.00E-08 | 1.343809876 | 0.269 | 0.109 | 0.000393877 | Lep-T |
| Macro-SPP1 | PPA2 | 1.70E-22 | 1.343198985 | 0.691 | 0.323 | 3.35E-18 | Lep-T |
| Macro-SPP1 | PRCP | 3.64E-25 | 1.338922898 | 0.909 | 0.636 | 7.16E-21 | Lep-T |
| Macro-SPP1 | ALG1 | 5.74E-16 | 1.334926216 | 0.291 | 0.079 | 1.13E-11 | Lep-T |
| Macro-SPP1 | MYO1B | 5.54E-17 | 1.334647059 | 0.326 | 0.095 | 1.09E-12 | Lep-T |
| Macro-SPP1 | SCUBE2 | 1.38E-06 | 1.334539854 | 0.343 | 0.182 | 0.027080237 | Lep-T |
| Macro-SPP1 | ABCA6 | 1.19E-13 | 1.330631789 | 0.343 | 0.12 | 2.35E-09 | Lep-T |
| Macro-SPP1 | GK-AS1 | 4.18E-23 | 1.329979711 | 0.566 | 0.202 | 8.23E-19 | Lep-T |
| Macro-SPP1 | ITGAX | 1.05E-28 | 1.329803637 | 0.92 | 0.812 | 2.07E-24 | Lep-T |
| Macro-SPP1 | AGTPBP1 | 7.89E-08 | 1.328055322 | 0.709 | 0.483 | 0.001552648 | Lep-T |
| Macro-SPP1 | TSPAN32 | 1.77E-13 | 1.324695783 | 0.377 | 0.141 | 3.48E-09 | Lep-T |
| Macro-SPP1 | HLA-DMB | 4.61E-42 | 1.320620359 | 0.954 | 0.796 | 9.08E-38 | Lep-T |
| Macro-SPP1 | FRY | 8.49E-08 | 1.315350963 | 0.371 | 0.188 | 0.001671645 | Lep-T |
| Macro-SPP1 | PCSK5 | 2.07E-07 | 1.314110904 | 0.469 | 0.286 | 0.004069307 | Lep-T |
| Macro-SPP1 | LDLR | 4.16E-16 | 1.313110043 | 0.617 | 0.304 | 8.20E-12 | Lep-T |
| Macro-SPP1 | TFEB | 4.25E-24 | 1.312153859 | 0.686 | 0.286 | 8.36E-20 | Lep-T |
| Macro-SPP1 | POLR3E | 3.69E-19 | 1.309860785 | 0.377 | 0.11 | 7.25E-15 | Lep-T |
| Macro-SPP1 | AARSD1 | 9.14E-12 | 1.308659302 | 0.343 | 0.132 | 1.80E-07 | Lep-T |
| Macro-SPP1 | VDR | 6.85E-17 | 1.301241572 | 0.56 | 0.257 | 1.35E-12 | Lep-T |

| | | | | | | | |
|---|---|---|---|---|---|---|---|
| Macro-SPP1 | DPAGT1 | 1.10E-06 | 1.300648481 | 0.28 | 0.125 | 0.021651224 | Lep-T |
| Macro-SPP1 | DPYD-AS1 | 2.22E-21 | 1.300375443 | 0.526 | 0.178 | 4.38E-17 | Lep-T |
| Macro-SPP1 | CLCN3 | 1.75E-11 | 1.299009778 | 0.623 | 0.369 | 3.44E-07 | Lep-T |
| Macro-SPP1 | BTG2 | 5.97E-19 | 1.298926528 | 0.857 | 0.648 | 1.17E-14 | Lep-T |
| Macro-SPP1 | NOS1AP | 2.99E-15 | 1.295899161 | 0.537 | 0.248 | 5.89E-11 | Lep-T |
| Macro-SPP1 | PSD3 | 9.80E-14 | 1.294585338 | 0.714 | 0.463 | 1.93E-09 | Lep-T |
| Macro-SPP1 | STX17-AS1 | 1.43E-17 | 1.291798095 | 0.474 | 0.176 | 2.82E-13 | Lep-T |
| Macro-SPP1 | ACSF3 | 2.59E-11 | 1.291609875 | 0.463 | 0.203 | 5.10E-07 | Lep-T |
| Macro-SPP1 | USP36 | 4.47E-12 | 1.288405622 | 0.726 | 0.564 | 8.80E-08 | Lep-T |
| Macro-SPP1 | HEATR6 | 4.03E-14 | 1.287117615 | 0.337 | 0.114 | 7.93E-10 | Lep-T |
| Macro-SPP1 | ZMPSTE24 | 6.23E-12 | 1.286056134 | 0.537 | 0.275 | 1.23E-07 | Lep-T |
| Macro-SPP1 | TEP1 | 3.65E-08 | 1.282887133 | 0.291 | 0.122 | 0.000718451 | Lep-T |
| Macro-SPP1 | EMD1 | 7.49E-27 | 1.280584337 | 0.777 | 0.34 | 1.47E-22 | Lep-T |
| Macro-SPP1 | ACE | 3.23E-13 | 1.280250706 | 0.56 | 0.267 | 6.36E-09 | Lep-T |
| Macro-SPP1 | IGSF6 | 1.10E-34 | 1.278167437 | 0.931 | 0.874 | 2.16E-30 | Lep-T |
| Macro-SPP1 | TANGO6 | 1.90E-19 | 1.272353093 | 0.4 | 0.121 | 3.73E-15 | Lep-T |
| Macro-SPP1 | TARS2 | 3.96E-16 | 1.267427822 | 0.291 | 0.078 | 7.80E-12 | Lep-T |
| Macro-SPP1 | MT-CYB | 4.67E-17 | 1.265978923 | 0.92 | 0.978 | 9.20E-13 | Lep-T |
| Macro-SPP1 | RBFOX2 | 9.74E-14 | 1.265129553 | 0.811 | 0.514 | 1.92E-09 | Lep-T |
| Macro-SPP1 | FASN | 8.05E-12 | 1.26356218 | 0.297 | 0.104 | 1.58E-07 | Lep-T |
| Macro-SPP1 | EEF2K | 8.24E-19 | 1.263542487 | 0.549 | 0.231 | 1.62E-14 | Lep-T |
| Macro-SPP1 | NUDT21 | 1.67E-19 | 1.2611113 | 0.594 | 0.243 | 3.29E-15 | Lep-T |
| Macro-SPP1 | TIMM21 | 3.50E-09 | 1.259069182 | 0.274 | 0.109 | 6.88E-05 | Lep-T |
| Macro-SPP1 | FOXK2 | 4.14E-13 | 1.259046386 | 0.709 | 0.428 | 8.15E-09 | Lep-T |
| Macro-SPP1 | CTDP1 | 5.90E-10 | 1.256320317 | 0.371 | 0.161 | 1.16E-05 | Lep-T |
| Macro-SPP1 | LGALS2 | 5.26E-40 | 1.253311319 | 0.971 | 0.785 | 1.04E-35 | Lep-T |
| Macro-SPP1 | SEL1L3 | 1.22E-17 | 1.253080867 | 0.52 | 0.213 | 2.40E-13 | Lep-T |
| Macro-SPP1 | TFEC | 7.07E-17 | 1.250921645 | 0.943 | 0.819 | 1.39E-12 | Lep-T |
| Macro-SPP1 | TNFRSF4 | 2.41E-12 | 1.249342638 | 0.389 | 0.162 | 4.74E-08 | Lep-T |
| Macro-SPP1 | FBXO34 | 4.72E-13 | 1.246693267 | 0.651 | 0.385 | 9.30E-09 | Lep-T |
| Macro-SPP1 | NXPE3 | 2.84E-10 | 1.241172619 | 0.274 | 0.099 | 5.59E-06 | Lep-T |
| Macro-SPP1 | NSUN4 | 1.53E-14 | 1.23944022 | 0.337 | 0.11 | 3.01E-10 | Lep-T |
| Macro-SPP1 | KCNK6 | 1.52E-14 | 1.239129528 | 0.497 | 0.211 | 2.99E-10 | Lep-T |
| Macro-SPP1 | MTHFD1 | 4.41E-20 | 1.239125468 | 0.606 | 0.238 | 8.68E-16 | Lep-T |
| Macro-SPP1 | SLC24A1 | 5.12E-16 | 1.237729083 | 0.354 | 0.113 | 1.01E-11 | Lep-T |
| Macro-SPP1 | SYNGR2 | 6.63E-23 | 1.235069573 | 0.909 | 0.72 | 1.31E-18 | Lep-T |
| Macro-SPP1 | TENT4B | 7.34E-14 | 1.234659979 | 0.629 | 0.35 | 1.45E-09 | Lep-T |
| Macro-SPP1 | SQLE | 4.00E-19 | 1.234337695 | 0.6 | 0.269 | 7.88E-15 | Lep-T |
| Macro-SPP1 | CD74 | 1.30E-14 | 1.232950187 | 1 | 1 | 2.55E-10 | Lep-T |
| Macro-SPP1 | POC5 | 7.83E-13 | 1.232188346 | 0.269 | 0.082 | 1.54E-08 | Lep-T |
| Macro-SPP1 | SLF2 | 9.59E-17 | 1.231597533 | 0.543 | 0.216 | 1.89E-12 | Lep-T |
| Macro-SPP1 | TAF5L | 1.01E-07 | 1.224816755 | 0.314 | 0.144 | 0.001990363 | Lep-T |
| Macro-SPP1 | SMC6 | 2.03E-14 | 1.224358866 | 0.571 | 0.265 | 4.00E-10 | Lep-T |
| Macro-SPP1 | GNA11 | 1.40E-16 | 1.221460137 | 0.326 | 0.093 | 2.76E-12 | Lep-T |
| Macro-SPP1 | CS | 1.37E-20 | 1.218218181 | 0.703 | 0.327 | 2.70E-16 | Lep-T |
| Macro-SPP1 | MVB12B | 2.17E-12 | 1.217723502 | 0.32 | 0.114 | 4.27E-08 | Lep-T |
| Macro-SPP1 | CNOT3 | 4.17E-10 | 1.214619107 | 0.286 | 0.105 | 8.21E-06 | Lep-T |
| Macro-SPP1 | POC1B-AS1 | 1.01E-10 | 1.214407956 | 0.291 | 0.109 | 1.99E-06 | Lep-T |
| Macro-SPP1 | DNAJB5 | 2.66E-08 | 1.211581068 | 0.269 | 0.113 | 0.000523696 | Lep-T |
| Macro-SPP1 | BHLHE40 | 8.68E-40 | 1.208984463 | 0.966 | 0.814 | 1.71E-35 | Lep-T |
| Macro-SPP1 | TMEM104 | 3.63E-20 | 1.208021198 | 0.429 | 0.132 | 7.14E-16 | Lep-T |
| Macro-SPP1 | CENPP | 2.44E-16 | 1.206571894 | 0.56 | 0.252 | 4.81E-12 | Lep-T |
| Macro-SPP1 | RASAL2 | 1.29E-13 | 1.206293908 | 0.863 | 0.743 | 2.54E-09 | Lep-T |
| Macro-SPP1 | PHF7 | 4.12E-07 | 1.204819285 | 0.251 | 0.11 | 0.0081085 | Lep-T |
| Macro-SPP1 | FDFT1 | 1.49E-22 | 1.203158513 | 0.743 | 0.39 | 2.92E-18 | Lep-T |

| | | | | | | | |
|---|---|---|---|---|---|---|---|
| Macro-SPP1 | NEK6 | 7.26E-29 | 1.202585016 | 0.897 | 0.599 | 1.43E-24 | Lep-T |
| Macro-SPP1 | SREBF2 | 2.13E-23 | 1.201159977 | 0.726 | 0.334 | 4.19E-19 | Lep-T |
| Macro-SPP1 | RAB31 | 8.75E-36 | 1.19658385 | 0.954 | 0.797 | 1.72E-31 | Lep-T |
| Macro-SPP1 | UBA7 | 1.69E-07 | 1.194642328 | 0.32 | 0.147 | 0.003333045 | Lep-T |
| Macro-SPP1 | TMEM150B | 3.91E-10 | 1.194531335 | 0.326 | 0.135 | 7.70E-06 | Lep-T |
| Macro-SPP1 | FBRSL1 | 6.51E-17 | 1.193449136 | 0.566 | 0.243 | 1.28E-12 | Lep-T |
| Macro-SPP1 | JUP | 1.17E-18 | 1.193041425 | 0.429 | 0.142 | 2.30E-14 | Lep-T |
| Macro-SPP1 | COL1A2 | 4.12E-10 | 1.19142848 | 0.326 | 0.126 | 8.11E-06 | Lep-T |
| Macro-SPP1 | EEF1AKNMT | 1.59E-09 | 1.18874477 | 0.286 | 0.11 | 3.12E-05 | Lep-T |
| Macro-SPP1 | TMEM69 | 6.34E-19 | 1.186708097 | 0.423 | 0.133 | 1.25E-14 | Lep-T |
| Macro-SPP1 | DCSTAMP | 1.29E-20 | 1.180044777 | 0.691 | 0.332 | 2.54E-16 | Lep-T |
| Macro-SPP1 | FYB1 | 2.34E-31 | 1.179372548 | 0.931 | 0.751 | 4.61E-27 | Lep-T |
| Macro-SPP1 | CPT2 | 2.74E-14 | 1.178304646 | 0.423 | 0.165 | 5.39E-10 | Lep-T |
| Macro-SPP1 | ZP3 | 3.32E-08 | 1.177791503 | 0.286 | 0.121 | 0.000654247 | Lep-T |
| Macro-SPP1 | TSPAN33 | 2.66E-16 | 1.177441713 | 0.823 | 0.508 | 5.24E-12 | Lep-T |
| Macro-SPP1 | ERCC3 | 2.19E-11 | 1.176730735 | 0.4 | 0.173 | 4.31E-07 | Lep-T |
| Macro-SPP1 | CLEC12A | 6.65E-23 | 1.176591522 | 0.84 | 0.53 | 1.31E-18 | Lep-T |
| Macro-SPP1 | ALS2 | 9.33E-13 | 1.175939951 | 0.251 | 0.072 | 1.84E-08 | Lep-T |
| Macro-SPP1 | NR1H3 | 3.18E-24 | 1.172047592 | 0.903 | 0.648 | 6.27E-20 | Lep-T |
| Macro-SPP1 | CLASP2 | 1.11E-10 | 1.171489244 | 0.737 | 0.485 | 2.18E-06 | Lep-T |
| Macro-SPP1 | TMEM41A | 1.44E-09 | 1.171305789 | 0.314 | 0.127 | 2.83E-05 | Lep-T |
| Macro-SPP1 | ZNF316 | 2.46E-16 | 1.170587467 | 0.469 | 0.179 | 4.85E-12 | Lep-T |
| Macro-SPP1 | RABGGTA | 7.32E-11 | 1.170156231 | 0.52 | 0.253 | 1.44E-06 | Lep-T |
| Macro-SPP1 | SEC13 | 7.32E-25 | 1.16916976 | 0.789 | 0.357 | 1.44E-20 | Lep-T |
| Macro-SPP1 | ERAP2 | 2.94E-21 | 1.166123557 | 0.566 | 0.213 | 5.79E-17 | Lep-T |
| Macro-SPP1 | PPARD | 1.08E-17 | 1.165585758 | 0.766 | 0.518 | 2.14E-13 | Lep-T |
| Macro-SPP1 | DOP1B | 9.63E-17 | 1.162528161 | 0.52 | 0.221 | 1.90E-12 | Lep-T |
| Macro-SPP1 | BLVRA | 3.11E-23 | 1.158163712 | 0.84 | 0.522 | 6.12E-19 | Lep-T |
| Macro-SPP1 | SLC28A3 | 6.13E-07 | 1.157991569 | 0.366 | 0.186 | 0.012068206 | Lep-T |
| Macro-SPP1 | SUSD3 | 2.30E-12 | 1.157712375 | 0.269 | 0.086 | 4.52E-08 | Lep-T |
| Macro-SPP1 | ATP9B | 2.39E-12 | 1.156734474 | 0.817 | 0.691 | 4.70E-08 | Lep-T |
| Macro-SPP1 | PRAM1 | 5.51E-20 | 1.156387978 | 0.543 | 0.203 | 1.08E-15 | Lep-T |
| Macro-SPP1 | FCHSD2 | 1.14E-20 | 1.155396919 | 0.857 | 0.575 | 2.24E-16 | Lep-T |
| Macro-SPP1 | ADO | 1.07E-19 | 1.153890337 | 0.52 | 0.19 | 2.10E-15 | Lep-T |
| Macro-SPP1 | SLC27A3 | 2.46E-19 | 1.153240591 | 0.76 | 0.447 | 4.84E-15 | Lep-T |
| Macro-SPP1 | TRAP1 | 1.91E-15 | 1.153171022 | 0.617 | 0.293 | 3.77E-11 | Lep-T |
| Macro-SPP1 | ABCB10 | 3.73E-13 | 1.151377245 | 0.32 | 0.11 | 7.34E-09 | Lep-T |
| Macro-SPP1 | PCK2 | 1.51E-14 | 1.147578544 | 0.6 | 0.301 | 2.97E-10 | Lep-T |
| Macro-SPP1 | SLC25A43 | 6.90E-10 | 1.146310128 | 0.377 | 0.169 | 1.36E-05 | Lep-T |
| Macro-SPP1 | ADAMTSL4 | 1.33E-20 | 1.145857288 | 0.611 | 0.246 | 2.62E-16 | Lep-T |
| Macro-SPP1 | AFG1L | 7.54E-11 | 1.145025831 | 0.314 | 0.121 | 1.49E-06 | Lep-T |
| Macro-SPP1 | HLA-DQB2 | 1.10E-17 | 1.143342091 | 0.623 | 0.274 | 2.17E-13 | Lep-T |
| Macro-SPP1 | CTNS | 3.23E-15 | 1.141949908 | 0.417 | 0.153 | 6.35E-11 | Lep-T |
| Macro-SPP1 | SLC8A1 | 4.53E-19 | 1.138623482 | 0.943 | 0.836 | 8.91E-15 | Lep-T |
| Macro-SPP1 | VPS53 | 3.89E-13 | 1.137916413 | 0.686 | 0.426 | 7.65E-09 | Lep-T |
| Macro-SPP1 | SSPN | 1.86E-14 | 1.136080143 | 0.606 | 0.3 | 3.66E-10 | Lep-T |
| Macro-SPP1 | NR4A3 | 8.48E-15 | 1.131809828 | 0.8 | 0.668 | 1.67E-10 | Lep-T |
| Macro-SPP1 | S1PR2 | 4.49E-14 | 1.131691902 | 0.349 | 0.119 | 8.83E-10 | Lep-T |
| Macro-SPP1 | MVD | 9.34E-19 | 1.131017718 | 0.611 | 0.272 | 1.84E-14 | Lep-T |
| Macro-SPP1 | KCNMA1 | 3.86E-24 | 1.130875127 | 0.914 | 0.668 | 7.60E-20 | Lep-T |
| Macro-SPP1 | ABCF3 | 1.52E-06 | 1.130217459 | 0.309 | 0.146 | 0.029857843 | Lep-T |
| Macro-SPP1 | IL27RA | 1.15E-22 | 1.127882927 | 0.709 | 0.319 | 2.27E-18 | Lep-T |
| Macro-SPP1 | SC5D | 2.36E-13 | 1.127752219 | 0.514 | 0.224 | 4.64E-09 | Lep-T |
| Macro-SPP1 | TRUB2 | 2.29E-15 | 1.122671082 | 0.394 | 0.135 | 4.50E-11 | Lep-T |
| Macro-SPP1 | GAS5 | 2.93E-15 | 1.118204799 | 0.697 | 0.429 | 5.77E-11 | Lep-T |

| | | | | | | | |
|---|---|---|---|---|---|---|---|
| Macro-SPP1 | ALPK1 | 1.06E-15 | 1.117250753 | 0.783 | 0.469 | 2.08E-11 | Lep-T |
| Macro-SPP1 | NUCKS1 | 2.02E-25 | 1.115634348 | 0.88 | 0.679 | 3.98E-21 | Lep-T |
| Macro-SPP1 | ANKRD26 | 4.06E-07 | 1.114770719 | 0.417 | 0.228 | 0.007988266 | Lep-T |
| Macro-SPP1 | MAPKAPK5 | 4.08E-14 | 1.114200907 | 0.6 | 0.28 | 8.02E-10 | Lep-T |
| Macro-SPP1 | TAF15 | 2.05E-16 | 1.113906488 | 0.703 | 0.366 | 4.04E-12 | Lep-T |
| Macro-SPP1 | MT-ND1 | 1.53E-19 | 1.112574738 | 0.96 | 0.986 | 3.02E-15 | Lep-T |
| Macro-SPP1 | SLC29A3 | 1.23E-19 | 1.11244805 | 0.703 | 0.349 | 2.43E-15 | Lep-T |
| Macro-SPP1 | ST3GAL5 | 1.33E-14 | 1.112272209 | 0.68 | 0.383 | 2.62E-10 | Lep-T |
| Macro-SPP1 | EYA3 | 1.70E-15 | 1.111980035 | 0.617 | 0.304 | 3.35E-11 | Lep-T |
| Macro-SPP1 | TULP4 | 1.23E-11 | 1.111683873 | 0.623 | 0.354 | 2.42E-07 | Lep-T |
| Macro-SPP1 | APEH | 8.85E-13 | 1.109900309 | 0.52 | 0.228 | 1.74E-08 | Lep-T |
| Macro-SPP1 | KSR1 | 1.01E-14 | 1.109180356 | 0.457 | 0.183 | 1.98E-10 | Lep-T |
| Macro-SPP1 | ARHGAP151 | 4.29E-15 | 1.108879853 | 0.857 | 0.664 | 8.45E-11 | Lep-T |
| Macro-SPP1 | USP20 | 1.01E-11 | 1.107908489 | 0.274 | 0.092 | 1.99E-07 | Lep-T |
| Macro-SPP1 | SYTL3 | 1.46E-13 | 1.107121993 | 0.474 | 0.203 | 2.87E-09 | Lep-T |
| Macro-SPP1 | SYAP1 | 2.03E-09 | 1.106264639 | 0.789 | 0.641 | 3.99E-05 | Lep-T |
| Macro-SPP1 | MANBA | 6.28E-19 | 1.102351763 | 0.88 | 0.816 | 1.24E-14 | Lep-T |
| Macro-SPP1 | NUP98 | 4.11E-15 | 1.101671238 | 0.817 | 0.625 | 8.09E-11 | Lep-T |
| Macro-SPP1 | ATP5F1A | 3.55E-35 | 1.100078382 | 0.926 | 0.727 | 6.98E-31 | Lep-T |
| Macro-SPP1 | NUP93 | 1.97E-19 | 1.100003514 | 0.583 | 0.237 | 3.89E-15 | Lep-T |
| Macro-SPP1 | ZNF185 | 1.88E-12 | 1.099604245 | 0.251 | 0.075 | 3.71E-08 | Lep-T |
| Macro-SPP1 | INSIG1 | 4.39E-16 | 1.09292182 | 0.954 | 0.773 | 8.65E-12 | Lep-T |
| Macro-SPP1 | ROCK1 | 3.52E-25 | 1.092659937 | 0.909 | 0.729 | 6.93E-21 | Lep-T |
| Macro-SPP1 | ZNF331 | 1.08E-18 | 1.09155993 | 0.909 | 0.804 | 2.12E-14 | Lep-T |
| Macro-SPP1 | HBEGF | 9.43E-15 | 1.091236614 | 0.869 | 0.743 | 1.86E-10 | Lep-T |
| Macro-SPP1 | HIST1H1C | 5.64E-07 | 1.090051419 | 0.337 | 0.179 | 0.01109194 | Lep-T |
| Macro-SPP1 | IFFO2 | 1.25E-13 | 1.087141668 | 0.48 | 0.203 | 2.46E-09 | Lep-T |
| Macro-SPP1 | PCNX1 | 1.76E-06 | 1.08684723 | 0.726 | 0.604 | 0.034735992 | Lep-T |
| Macro-SPP1 | GLMP | 9.61E-11 | 1.085080512 | 0.56 | 0.284 | 1.89E-06 | Lep-T |
| Macro-SPP1 | CBFA2T3 | 1.21E-10 | 1.082729953 | 0.303 | 0.115 | 2.39E-06 | Lep-T |
| Macro-SPP1 | MALT1 | 2.75E-08 | 1.079458941 | 0.663 | 0.462 | 0.000540733 | Lep-T |
| Macro-SPP1 | CHST11 | 5.22E-16 | 1.079290962 | 0.954 | 0.902 | 1.03E-11 | Lep-T |
| Macro-SPP1 | SRD5A1 | 5.73E-16 | 1.07408701 | 0.589 | 0.27 | 1.13E-11 | Lep-T |
| Macro-SPP1 | VPS13D | 2.14E-10 | 1.073965906 | 0.623 | 0.38 | 4.22E-06 | Lep-T |
| Macro-SPP1 | HYAL2 | 7.71E-15 | 1.069608275 | 0.583 | 0.273 | 1.52E-10 | Lep-T |
| Macro-SPP1 | EXOC4 | 6.70E-18 | 1.069066538 | 0.84 | 0.721 | 1.32E-13 | Lep-T |
| Macro-SPP1 | PRKCB | 2.79E-14 | 1.067669233 | 0.817 | 0.556 | 5.49E-10 | Lep-T |
| Macro-SPP1 | PERP | 1.36E-23 | 1.066704957 | 0.4 | 0.1 | 2.68E-19 | Lep-T |
| Macro-SPP1 | CORO2A | 3.32E-10 | 1.063621283 | 0.297 | 0.115 | 6.53E-06 | Lep-T |
| Macro-SPP1 | SIDT2 | 1.41E-21 | 1.063185284 | 0.674 | 0.287 | 2.78E-17 | Lep-T |
| Macro-SPP1 | PSEN2 | 5.09E-08 | 1.061987522 | 0.394 | 0.193 | 0.001001786 | Lep-T |
| Macro-SPP1 | PRXL2A | 3.09E-20 | 1.061566479 | 0.8 | 0.464 | 6.09E-16 | Lep-T |
| Macro-SPP1 | NAP1L1 | 3.43E-25 | 1.060432748 | 0.931 | 0.876 | 6.74E-21 | Lep-T |
| Macro-SPP1 | LTA4H | 2.16E-17 | 1.05665064 | 0.886 | 0.706 | 4.25E-13 | Lep-T |
| Macro-SPP1 | ST7 | 1.90E-08 | 1.056612257 | 0.274 | 0.113 | 0.000373205 | Lep-T |
| Macro-SPP1 | PLXDC2 | 3.29E-20 | 1.055664723 | 0.903 | 0.866 | 6.47E-16 | Lep-T |
| Macro-SPP1 | NRROS | 7.12E-12 | 1.052246021 | 0.417 | 0.177 | 1.40E-07 | Lep-T |
| Macro-SPP1 | TACC3 | 5.03E-07 | 1.049216427 | 0.326 | 0.151 | 0.009896966 | Lep-T |
| Macro-SPP1 | INPP5K | 5.23E-17 | 1.048211448 | 0.543 | 0.225 | 1.03E-12 | Lep-T |
| Macro-SPP1 | SF3B2 | 2.44E-25 | 1.046342344 | 0.88 | 0.626 | 4.80E-21 | Lep-T |
| Macro-SPP1 | TFRC | 2.43E-22 | 1.044934961 | 0.949 | 0.881 | 4.79E-18 | Lep-T |
| Macro-SPP1 | ARL14EP | 2.73E-12 | 1.043629087 | 0.497 | 0.236 | 5.38E-08 | Lep-T |
| Macro-SPP1 | BHLHE41 | 1.20E-22 | 1.041801457 | 0.891 | 0.692 | 2.35E-18 | Lep-T |
| Macro-SPP1 | SPATA5 | 3.16E-07 | 1.03980295 | 0.36 | 0.183 | 0.006212505 | Lep-T |
| Macro-SPP1 | CKAP5 | 8.47E-15 | 1.038754631 | 0.543 | 0.232 | 1.67E-10 | Lep-T |

| | | | | | | | |
|---|---|---|---|---|---|---|---|
| Macro-SPP1 | UTP3 | 3.16E-13 | 1.035823915 | 0.554 | 0.255 | 6.23E-09 | Lep-T |
| Macro-SPP1 | NFAM1 | 6.90E-16 | 1.035093304 | 0.703 | 0.383 | 1.36E-11 | Lep-T |
| Macro-SPP1 | SPRYD3 | 3.16E-10 | 1.0345307 | 0.297 | 0.11 | 6.22E-06 | Lep-T |
| Macro-SPP1 | GBA | 4.05E-17 | 1.034151756 | 0.646 | 0.323 | 7.97E-13 | Lep-T |
| Macro-SPP1 | SLC22A15 | 1.12E-10 | 1.033958597 | 0.463 | 0.219 | 2.21E-06 | Lep-T |
| Macro-SPP1 | RCOR1 | 1.07E-15 | 1.032849275 | 0.84 | 0.64 | 2.11E-11 | Lep-T |
| Macro-SPP1 | ADGRE5 | 1.91E-21 | 1.032345804 | 0.909 | 0.704 | 3.77E-17 | Lep-T |
| Macro-SPP1 | MSMO1 | 1.87E-16 | 1.03198892 | 0.543 | 0.236 | 3.68E-12 | Lep-T |
| Macro-SPP1 | SEMA4A | 5.94E-13 | 1.031819323 | 0.771 | 0.532 | 1.17E-08 | Lep-T |
| Macro-SPP1 | EOGT | 1.55E-17 | 1.031322909 | 0.446 | 0.155 | 3.05E-13 | Lep-T |
| Macro-SPP1 | UBASH3B | 4.24E-08 | 1.031175395 | 0.674 | 0.524 | 0.000834941 | Lep-T |
| Macro-SPP1 | COG2 | 1.43E-06 | 1.030771988 | 0.446 | 0.239 | 0.028212321 | Lep-T |
| Macro-SPP1 | NUDT3 | 1.15E-10 | 1.030525247 | 0.526 | 0.272 | 2.26E-06 | Lep-T |
| Macro-SPP1 | MAT2B | 3.99E-18 | 1.026775849 | 0.629 | 0.287 | 7.86E-14 | Lep-T |
| Macro-SPP1 | ATP6V0A2 | 2.45E-15 | 1.026052446 | 0.514 | 0.213 | 4.83E-11 | Lep-T |
| Macro-SPP1 | FARS2 | 1.71E-12 | 1.025969226 | 0.606 | 0.317 | 3.37E-08 | Lep-T |
| Macro-SPP1 | TMCO4 | 8.31E-18 | 1.025136174 | 0.543 | 0.214 | 1.64E-13 | Lep-T |
| Macro-SPP1 | SLC9A7 | 5.48E-09 | 1.025017887 | 0.537 | 0.307 | 0.00010793 | Lep-T |
| Macro-SPP1 | CRIM1 | 1.86E-08 | 1.024368918 | 0.709 | 0.502 | 0.000365728 | Lep-T |
| Macro-SPP1 | SNX29 | 2.17E-11 | 1.024170771 | 0.726 | 0.502 | 4.28E-07 | Lep-T |
| Macro-SPP1 | TANGO2 | 5.84E-11 | 1.023794798 | 0.429 | 0.191 | 1.15E-06 | Lep-T |
| Macro-SPP1 | STRADA | 7.20E-12 | 1.023763466 | 0.417 | 0.173 | 1.42E-07 | Lep-T |
| Macro-SPP1 | CALML5 | 1.43E-33 | 1.023487442 | 0.337 | 0.05 | 2.81E-29 | Lep-T |
| Macro-SPP1 | ADAT1 | 1.02E-10 | 1.023248897 | 0.297 | 0.109 | 2.00E-06 | Lep-T |
| Macro-SPP1 | TYW1 | 6.39E-07 | 1.022548786 | 0.469 | 0.253 | 0.01257156 | Lep-T |
| Macro-SPP1 | PEX26 | 1.18E-07 | 1.022527186 | 0.303 | 0.131 | 0.002331012 | Lep-T |
| Macro-SPP1 | MOB3B | 1.39E-12 | 1.022050394 | 0.771 | 0.525 | 2.73E-08 | Lep-T |
| Macro-SPP1 | POLR3D | 4.33E-09 | 1.021748543 | 0.394 | 0.177 | 8.52E-05 | Lep-T |
| Macro-SPP1 | MAP4K3 | 1.17E-07 | 1.021145187 | 0.743 | 0.532 | 0.002308635 | Lep-T |
| Macro-SPP1 | LTV1 | 8.79E-14 | 1.020470919 | 0.566 | 0.272 | 1.73E-09 | Lep-T |
| Macro-SPP1 | NUAK2 | 3.75E-07 | 1.019751087 | 0.383 | 0.189 | 0.007379802 | Lep-T |
| Macro-SPP1 | HVCN1 | 1.58E-15 | 1.019102765 | 0.771 | 0.478 | 3.11E-11 | Lep-T |
| Macro-SPP1 | NQO1 | 4.36E-07 | 1.0168081 | 0.349 | 0.179 | 0.008575159 | Lep-T |
| Macro-SPP1 | GPATCH4 | 1.01E-11 | 1.014729382 | 0.434 | 0.187 | 1.99E-07 | Lep-T |
| Macro-SPP1 | FGD5 | 1.58E-13 | 1.013425826 | 0.429 | 0.167 | 3.11E-09 | Lep-T |
| Macro-SPP1 | MASTL | 4.31E-11 | 1.01341078 | 0.709 | 0.465 | 8.49E-07 | Lep-T |
| Macro-SPP1 | CARD9 | 3.96E-15 | 1.012108725 | 0.526 | 0.214 | 7.79E-11 | Lep-T |
| Macro-SPP1 | RAD54L2 | 2.64E-08 | 1.011892125 | 0.303 | 0.129 | 0.000518683 | Lep-T |
| Macro-SPP1 | FARP2 | 2.92E-14 | 1.011670084 | 0.583 | 0.273 | 5.76E-10 | Lep-T |
| Macro-SPP1 | ATG7 | 3.58E-14 | 1.011257829 | 0.811 | 0.672 | 7.04E-10 | Lep-T |
| Macro-SPP1 | MMP2 | 1.33E-23 | 1.010263031 | 0.611 | 0.223 | 2.62E-19 | Lep-T |
| Macro-SPP1 | RNASEL | 1.89E-09 | 1.010233384 | 0.257 | 0.093 | 3.73E-05 | Lep-T |
| Macro-SPP1 | OXSR1 | 2.40E-07 | 1.009999981 | 0.817 | 0.727 | 0.004732051 | Lep-T |
| Macro-SPP1 | HMBS | 9.64E-10 | 1.008362623 | 0.4 | 0.174 | 1.90E-05 | Lep-T |
| Macro-SPP1 | TXLNA | 1.01E-12 | 1.008138469 | 0.583 | 0.302 | 2.00E-08 | Lep-T |
| Macro-SPP1 | NIPAL2 | 1.05E-09 | 1.008060807 | 0.383 | 0.169 | 2.07E-05 | Lep-T |
| Macro-SPP1 | MPND | 8.67E-18 | 1.004883479 | 0.663 | 0.332 | 1.71E-13 | Lep-T |
| Macro-SPP1 | HSF1 | 1.86E-06 | 1.004037639 | 0.566 | 0.325 | 0.036623338 | Lep-T |
| Macro-SPP1 | CCDC102B | 1.36E-16 | 1.00123435 | 0.549 | 0.237 | 2.67E-12 | Lep-T |
| Macro-SPP1 | FOLR3 | 4.80E-83 | 5.005315564 | 0.479 | 0.054 | 9.45E-79 | Mari |
| Macro-SPP1 | ITGB7 | 1.53E-40 | 2.687164558 | 0.357 | 0.076 | 3.02E-36 | Mari |
| Macro-SPP1 | SPP1 | 9.07E-41 | 2.231407307 | 0.812 | 0.585 | 1.79E-36 | Mari |
| Macro-SPP1 | CCL7 | 6.12E-22 | 2.146559469 | 0.293 | 0.092 | 1.20E-17 | Mari |
| Macro-SPP1 | MMP12 | 8.99E-19 | 1.85030103 | 0.536 | 0.339 | 1.77E-14 | Mari |
| Macro-SPP1 | SLAMF9 | 1.80E-43 | 1.78289601 | 0.652 | 0.287 | 3.55E-39 | Mari |

| | | | | | | | |
|---|---|---|---|---|---|---|---|
| Macro-SPP1 | CCL8 | 9.32E-22 | 1.704587873 | 0.4 | 0.163 | 1.83E-17 | Mari |
| Macro-SPP1 | FAH | 1.25E-20 | 1.66971279 | 0.46 | 0.229 | 2.46E-16 | Mari |
| Macro-SPP1 | CA12 | 2.79E-27 | 1.579462892 | 0.488 | 0.217 | 5.50E-23 | Mari |
| Macro-SPP1 | HLA-DQA21 | 2.97E-39 | 1.515137262 | 0.876 | 0.631 | 5.85E-35 | Mari |
| Macro-SPP1 | LAT | 2.38E-16 | 1.506824151 | 0.319 | 0.136 | 4.69E-12 | Mari |
| Macro-SPP1 | CES1 | 5.07E-23 | 1.4992958 | 0.362 | 0.132 | 9.98E-19 | Mari |
| Macro-SPP1 | NOL3 | 1.59E-16 | 1.494017173 | 0.362 | 0.169 | 3.13E-12 | Mari |
| Macro-SPP1 | IL27 | 3.54E-13 | 1.462309879 | 0.26 | 0.107 | 6.97E-09 | Mari |
| Macro-SPP1 | CLEC5A | 2.29E-30 | 1.407401055 | 0.683 | 0.391 | 4.51E-26 | Mari |
| Macro-SPP1 | RSAD2 | 1.58E-13 | 1.308902424 | 0.271 | 0.115 | 3.10E-09 | Mari |
| Macro-SPP1 | MGST1 | 7.99E-27 | 1.28225472 | 0.645 | 0.367 | 1.57E-22 | Mari |
| Macro-SPP1 | S100A8 | 3.55E-31 | 1.267682283 | 0.893 | 0.718 | 6.98E-27 | Mari |
| Macro-SPP1 | PLTP | 4.61E-32 | 1.259354802 | 0.795 | 0.529 | 9.07E-28 | Mari |
| Macro-SPP1 | SLC24A3 | 5.43E-12 | 1.238983105 | 0.279 | 0.128 | 1.07E-07 | Mari |
| Macro-SPP1 | TNFSF14 | 2.13E-11 | 1.221256541 | 0.429 | 0.26 | 4.19E-07 | Mari |
| Macro-SPP1 | NME4 | 4.62E-21 | 1.21193055 | 0.448 | 0.21 | 9.10E-17 | Mari |
| Macro-SPP1 | CP1 | 1.64E-23 | 1.208540547 | 0.571 | 0.313 | 3.23E-19 | Mari |
| Macro-SPP1 | LAMC1 | 6.98E-11 | 1.190826357 | 0.29 | 0.144 | 1.37E-06 | Mari |
| Macro-SPP1 | CCDC58 | 1.06E-09 | 1.185234248 | 0.255 | 0.128 | 2.09E-05 | Mari |
| Macro-SPP1 | SPR | 1.30E-11 | 1.160145533 | 0.379 | 0.217 | 2.56E-07 | Mari |
| Macro-SPP1 | CALHM6 | 3.06E-24 | 1.152530326 | 0.886 | 0.745 | 6.03E-20 | Mari |
| Macro-SPP1 | TWF1 | 1.08E-23 | 1.115435826 | 0.605 | 0.349 | 2.12E-19 | Mari |
| Macro-SPP1 | IGFBP2 | 1.68E-20 | 1.112919043 | 0.402 | 0.167 | 3.30E-16 | Mari |
| Macro-SPP1 | ZGLP1 | 2.39E-11 | 1.11038172 | 0.352 | 0.196 | 4.70E-07 | Mari |
| Macro-SPP1 | CCND1 | 3.81E-21 | 1.103365031 | 0.605 | 0.348 | 7.50E-17 | Mari |
| Macro-SPP1 | STXBP5-AS1 | 1.76E-12 | 1.095825664 | 0.281 | 0.127 | 3.47E-08 | Mari |
| Macro-SPP1 | CHI3L1 | 5.10E-28 | 1.077283259 | 0.864 | 0.617 | 1.00E-23 | Mari |
| Macro-SPP1 | IFIT3 | 5.32E-17 | 1.066807912 | 0.588 | 0.362 | 1.05E-12 | Mari |
| Macro-SPP1 | GBP1 | 2.80E-32 | 1.053077711 | 0.919 | 0.734 | 5.51E-28 | Mari |
| Macro-SPP1 | MRPL23 | 2.75E-24 | 1.039837242 | 0.681 | 0.41 | 5.42E-20 | Mari |
| Macro-SPP1 | DDX60L | 2.10E-19 | 1.006237072 | 0.652 | 0.426 | 4.13E-15 | Mari |
| Macro-SPP1 | CLDN231 | 6.79E-11 | 1.004486758 | 0.367 | 0.203 | 1.34E-06 | Mari |
| Macro-SPP1 | CAMP | 1.95E-38 | 2.99305156 | 0.539 | 0.227 | 3.85E-34 | Tb |
| Macro-SPP1 | KRT6A | 1.92E-29 | 2.815677031 | 0.265 | 0.056 | 3.77E-25 | Tb |
| Macro-SPP1 | SFN | 1.03E-24 | 2.535862509 | 0.325 | 0.11 | 2.03E-20 | Tb |
| Macro-SPP1 | SLC35F1 | 1.21E-58 | 2.448293571 | 0.536 | 0.154 | 2.38E-54 | Tb |
| Macro-SPP1 | S100A7 | 3.14E-23 | 2.401416504 | 0.272 | 0.076 | 6.19E-19 | Tb |
| Macro-SPP1 | KRT17 | 2.44E-25 | 2.218171334 | 0.256 | 0.06 | 4.80E-21 | Tb |
| Macro-SPP1 | S100A2 | 3.10E-27 | 2.204284677 | 0.289 | 0.075 | 6.10E-23 | Tb |
| Macro-SPP1 | IL1R2 | 1.56E-08 | 2.202877591 | 0.283 | 0.164 | 0.000307273 | Tb |
| Macro-SPP1 | GGT51 | 5.34E-26 | 2.147397612 | 0.38 | 0.144 | 1.05E-21 | Tb |
| Macro-SPP1 | MT1M | 7.68E-16 | 2.094605726 | 0.481 | 0.296 | 1.51E-11 | Tb |
| Macro-SPP1 | KRT16 | 3.73E-34 | 1.937879458 | 0.289 | 0.053 | 7.35E-30 | Tb |
| Macro-SPP1 | ILF3-DT | 1.34E-30 | 1.873745893 | 0.581 | 0.289 | 2.63E-26 | Tb |
| Macro-SPP1 | HOMER1 | 2.25E-37 | 1.847247296 | 0.623 | 0.305 | 4.43E-33 | Tb |
| Macro-SPP1 | PPIC | 4.51E-20 | 1.805766954 | 0.355 | 0.149 | 8.88E-16 | Tb |
| Macro-SPP1 | P2RY141 | 7.09E-18 | 1.802980175 | 0.272 | 0.098 | 1.40E-13 | Tb |
| Macro-SPP1 | TNC1 | 9.53E-14 | 1.788757134 | 0.26 | 0.108 | 1.88E-09 | Tb |
| Macro-SPP1 | DNAI2 | 3.42E-21 | 1.772982666 | 0.477 | 0.243 | 6.73E-17 | Tb |
| Macro-SPP1 | SIRPB2 | 2.45E-18 | 1.751377066 | 0.287 | 0.106 | 4.82E-14 | Tb |
| Macro-SPP1 | CXCL9 | 2.38E-19 | 1.740757469 | 0.642 | 0.424 | 4.69E-15 | Tb |
| Macro-SPP1 | MT1H | 4.82E-22 | 1.739540802 | 0.523 | 0.284 | 9.48E-18 | Tb |
| Macro-SPP1 | IGKC | 1.54E-08 | 1.72007334 | 0.338 | 0.221 | 0.00030215 | Tb |
| Macro-SPP1 | FN1 | 6.41E-21 | 1.696127863 | 0.592 | 0.37 | 1.26E-16 | Tb |
| Macro-SPP1 | MT1G | 9.24E-29 | 1.695156366 | 0.706 | 0.422 | 1.82E-24 | Tb |

| | | | | | | | |
|---|---|---|---|---|---|---|---|
| Macro-SPP1 | NRN1 | 1.26E-37 | 1.673895476 | 0.618 | 0.298 | 2.49E-33 | Tb |
| Macro-SPP1 | PGBD5 | 2.30E-13 | 1.633158075 | 0.263 | 0.111 | 4.53E-09 | Tb |
| Macro-SPP1 | KRT14 | 3.44E-34 | 1.62439672 | 0.468 | 0.174 | 6.77E-30 | Tb |
| Macro-SPP1 | C1QTNF1 | 6.02E-13 | 1.585495215 | 0.28 | 0.127 | 1.18E-08 | Tb |
| Macro-SPP1 | GRK5 | 7.61E-17 | 1.552578478 | 0.318 | 0.133 | 1.50E-12 | Tb |
| Macro-SPP1 | ZNF101 | 8.45E-19 | 1.543200046 | 0.439 | 0.214 | 1.66E-14 | Tb |
| Macro-SPP1 | KCNJ2 | 2.13E-12 | 1.532096125 | 0.283 | 0.132 | 4.20E-08 | Tb |
| Macro-SPP1 | GZF1 | 2.50E-18 | 1.507691295 | 0.373 | 0.168 | 4.92E-14 | Tb |
| Macro-SPP1 | CLEC1A | 9.19E-13 | 1.492136181 | 0.322 | 0.162 | 1.81E-08 | Tb |
| Macro-SPP1 | FGD51 | 1.82E-18 | 1.466970393 | 0.331 | 0.136 | 3.58E-14 | Tb |
| Macro-SPP1 | IL1RAP | 1.64E-11 | 1.46106072 | 0.316 | 0.164 | 3.22E-07 | Tb |
| Macro-SPP1 | ENPP21 | 1.21E-10 | 1.423399385 | 0.316 | 0.171 | 2.38E-06 | Tb |
| Macro-SPP1 | ST6GAL11 | 5.32E-21 | 1.416048653 | 0.437 | 0.204 | 1.05E-16 | Tb |
| Macro-SPP1 | AQP3 | 2.26E-16 | 1.404849721 | 0.364 | 0.175 | 4.44E-12 | Tb |
| Macro-SPP1 | MT1E | 4.64E-31 | 1.396327658 | 0.786 | 0.497 | 9.13E-27 | Tb |
| Macro-SPP1 | IL2RA | 5.96E-15 | 1.378590451 | 0.38 | 0.199 | 1.17E-10 | Tb |
| Macro-SPP1 | PPP1R13B | 4.67E-10 | 1.349425342 | 0.252 | 0.12 | 9.20E-06 | Tb |
| Macro-SPP1 | TAF4B | 2.62E-07 | 1.344343194 | 0.4 | 0.27 | 0.00515366 | Tb |
| Macro-SPP1 | AURKA | 2.10E-12 | 1.342033469 | 0.291 | 0.139 | 4.13E-08 | Tb |
| Macro-SPP1 | CD69 | 8.14E-23 | 1.323669882 | 0.499 | 0.244 | 1.60E-18 | Tb |
| Macro-SPP1 | ARL4D | 9.63E-18 | 1.320142579 | 0.488 | 0.271 | 1.90E-13 | Tb |
| Macro-SPP1 | ME1 | 1.83E-15 | 1.314844862 | 0.578 | 0.395 | 3.61E-11 | Tb |
| Macro-SPP1 | PTGDS1 | 1.04E-06 | 1.312765944 | 0.318 | 0.207 | 0.020488137 | Tb |
| Macro-SPP1 | FBP1 | 2.80E-57 | 1.309902837 | 0.989 | 0.879 | 5.50E-53 | Tb |
| Macro-SPP1 | THBD | 5.30E-20 | 1.304105813 | 0.49 | 0.259 | 1.04E-15 | Tb |
| Macro-SPP1 | CBR3 | 2.22E-21 | 1.303668718 | 0.444 | 0.213 | 4.37E-17 | Tb |
| Macro-SPP1 | HLA-DRB1 | 7.23E-79 | 1.297797308 | 0.998 | 0.996 | 1.42E-74 | Tb |
| Macro-SPP1 | QPCT | 1.60E-44 | 1.276673223 | 0.857 | 0.567 | 3.15E-40 | Tb |
| Macro-SPP1 | MCF2L2 | 2.99E-18 | 1.257913675 | 0.543 | 0.309 | 5.89E-14 | Tb |
| Macro-SPP1 | GMPR | 5.25E-18 | 1.249223331 | 0.375 | 0.17 | 1.03E-13 | Tb |
| Macro-SPP1 | GTF2IRD1 | 4.49E-12 | 1.223889352 | 0.539 | 0.37 | 8.83E-08 | Tb |
| Macro-SPP1 | CLEC10A | 8.93E-18 | 1.217260962 | 0.576 | 0.36 | 1.76E-13 | Tb |
| Macro-SPP1 | PRKAR1B | 1.99E-11 | 1.215963509 | 0.298 | 0.148 | 3.92E-07 | Tb |
| Macro-SPP1 | NSUN6 | 8.32E-12 | 1.213007532 | 0.404 | 0.24 | 1.64E-07 | Tb |
| Macro-SPP1 | SATB1-AS1 | 2.75E-08 | 1.210755422 | 0.263 | 0.143 | 0.000540437 | Tb |
| Macro-SPP1 | SLC28A31 | 1.05E-10 | 1.205028854 | 0.302 | 0.163 | 2.06E-06 | Tb |
| Macro-SPP1 | HSD11B1-AS1 | 1.49E-09 | 1.201091273 | 0.349 | 0.203 | 2.94E-05 | Tb |
| Macro-SPP1 | HYDIN | 1.23E-07 | 1.188236139 | 0.327 | 0.206 | 0.002425041 | Tb |
| Macro-SPP1 | DIXDC1 | 2.55E-09 | 1.178251486 | 0.406 | 0.265 | 5.02E-05 | Tb |
| Macro-SPP1 | LIG1 | 3.39E-11 | 1.174579598 | 0.386 | 0.217 | 6.66E-07 | Tb |
| Macro-SPP1 | LYRM1 | 2.33E-10 | 1.172054362 | 0.43 | 0.269 | 4.59E-06 | Tb |
| Macro-SPP1 | IDO2 | 1.16E-06 | 1.167992624 | 0.541 | 0.423 | 0.022746034 | Tb |
| Macro-SPP1 | HPGD | 4.86E-12 | 1.158233543 | 0.294 | 0.141 | 9.57E-08 | Tb |
| Macro-SPP1 | MT2A | 1.86E-31 | 1.14822274 | 0.991 | 0.924 | 3.67E-27 | Tb |
| Macro-SPP1 | CPED1 | 2.81E-14 | 1.147496092 | 0.572 | 0.384 | 5.53E-10 | Tb |
| Macro-SPP1 | CD300E | 1.18E-13 | 1.147370322 | 0.38 | 0.195 | 2.31E-09 | Tb |
| Macro-SPP1 | RGS16 | 1.68E-11 | 1.144567769 | 0.437 | 0.267 | 3.30E-07 | Tb |
| Macro-SPP1 | MT1X | 2.99E-33 | 1.144532474 | 0.881 | 0.626 | 5.89E-29 | Tb |
| Macro-SPP1 | VDR1 | 1.15E-14 | 1.143046252 | 0.428 | 0.23 | 2.26E-10 | Tb |
| Macro-SPP1 | AXL | 3.17E-13 | 1.138297946 | 0.382 | 0.203 | 6.24E-09 | Tb |
| Macro-SPP1 | PFKP | 6.18E-23 | 1.138286774 | 0.722 | 0.502 | 1.22E-18 | Tb |
| Macro-SPP1 | CASS41 | 2.09E-10 | 1.124219532 | 0.287 | 0.149 | 4.11E-06 | Tb |
| Macro-SPP1 | RNF217 | 2.03E-07 | 1.121555574 | 0.402 | 0.268 | 0.003996275 | Tb |
| Macro-SPP1 | MAP3K7CL | 2.94E-10 | 1.118069817 | 0.389 | 0.229 | 5.79E-06 | Tb |
| Macro-SPP1 | PDLIM4 | 3.80E-24 | 1.117840164 | 0.656 | 0.403 | 7.48E-20 | Tb |

| | | | | | | | |
|---|---|---|---|---|---|---|---|
| Macro-SPP1 | INHBA-AS1 | 1.22E-10 | 1.11723787 | 0.483 | 0.334 | 2.41E-06 | Tb |
| Macro-SPP1 | BRCA2 | 1.86E-07 | 1.112190522 | 0.342 | 0.224 | 0.00365987 | Tb |
| Macro-SPP1 | CTSC1 | 1.39E-25 | 1.097418829 | 0.974 | 0.903 | 2.73E-21 | Tb |
| Macro-SPP1 | LRRC23 | 1.66E-15 | 1.093649045 | 0.649 | 0.459 | 3.27E-11 | Tb |
| Macro-SPP1 | CRISPLD21 | 2.78E-13 | 1.093170108 | 0.428 | 0.244 | 5.48E-09 | Tb |
| Macro-SPP1 | PPARGC1B1 | 1.11E-23 | 1.084598003 | 0.629 | 0.368 | 2.19E-19 | Tb |
| Macro-SPP1 | STK19 | 5.90E-12 | 1.082553683 | 0.377 | 0.215 | 1.16E-07 | Tb |
| Macro-SPP1 | ANK21 | 4.68E-09 | 1.081839771 | 0.276 | 0.148 | 9.22E-05 | Tb |
| Macro-SPP1 | CFB1 | 2.55E-13 | 1.081359307 | 0.448 | 0.266 | 5.02E-09 | Tb |
| Macro-SPP1 | F3 | 8.27E-12 | 1.079611869 | 0.419 | 0.248 | 1.63E-07 | Tb |
| Macro-SPP1 | GIMAP7 | 1.40E-09 | 1.079609279 | 0.338 | 0.199 | 2.75E-05 | Tb |
| Macro-SPP1 | SULF2 | 2.78E-26 | 1.075695193 | 0.777 | 0.512 | 5.48E-22 | Tb |
| Macro-SPP1 | CXCL10 | 3.77E-07 | 1.072643701 | 0.479 | 0.355 | 0.007413043 | Tb |
| Macro-SPP1 | ALOX5 | 1.19E-30 | 1.070251671 | 0.837 | 0.611 | 2.35E-26 | Tb |
| Macro-SPP1 | MAP3K13 | 6.34E-14 | 1.068531941 | 0.704 | 0.528 | 1.25E-09 | Tb |
| Macro-SPP1 | SEMA4B | 3.14E-12 | 1.067650296 | 0.333 | 0.166 | 6.18E-08 | Tb |
| Macro-SPP1 | CCDC102B1 | 2.07E-13 | 1.066967457 | 0.397 | 0.217 | 4.07E-09 | Tb |
| Macro-SPP1 | CISH | 2.01E-16 | 1.049483327 | 0.424 | 0.214 | 3.96E-12 | Tb |
| Macro-SPP1 | INTS4 | 3.51E-07 | 1.046923129 | 0.269 | 0.157 | 0.006900304 | Tb |
| Macro-SPP1 | CCND2 | 3.96E-20 | 1.045722714 | 0.605 | 0.373 | 7.80E-16 | Tb |
| Macro-SPP1 | FLT11 | 3.06E-23 | 1.045127824 | 0.781 | 0.549 | 6.02E-19 | Tb |
| Macro-SPP1 | MT1F | 1.35E-19 | 1.01768537 | 0.72 | 0.504 | 2.65E-15 | Tb |
| Macro-SPP1 | COQ7 | 7.27E-07 | 1.017550436 | 0.263 | 0.154 | 0.014309304 | Tb |
| Macro-SPP1 | SLC49A4 | 4.67E-14 | 1.01510396 | 0.565 | 0.364 | 9.20E-10 | Tb |
| Macro-SPP1 | HLA-DQA1 | 2.43E-76 | 1.012295381 | 0.991 | 0.971 | 4.79E-72 | Tb |
| Macro-SPP1 | P2RY131 | 1.19E-11 | 1.010814247 | 0.397 | 0.224 | 2.35E-07 | Tb |
| Macro-SPP1 | CD300LF | 7.63E-24 | 1.004266459 | 0.607 | 0.33 | 1.50E-19 | Tb |
| **Disease-associated genes in Macro-ISG** | | | | | | | |
| Macro-ISG | OTOA | 1.47E-32 | 3.042948922 | 0.398 | 0.109 | 2.89E-28 | Lep-L |
| Macro-ISG | CYP27A1 | 5.59E-26 | 2.796890995 | 0.35 | 0.097 | 1.10E-21 | Lep-L |
| Macro-ISG | GPNMB | 1.06E-34 | 2.692879738 | 0.619 | 0.296 | 2.08E-30 | Lep-L |
| Macro-ISG | CHCHD6 | 7.44E-17 | 2.643601621 | 0.354 | 0.152 | 1.46E-12 | Lep-L |
| Macro-ISG | PPARG | 2.19E-18 | 2.387311043 | 0.367 | 0.146 | 4.31E-14 | Lep-L |
| Macro-ISG | SELENOP | 2.21E-47 | 2.325986649 | 0.549 | 0.137 | 4.35E-43 | Lep-L |
| Macro-ISG | LILRB5 | 5.83E-15 | 2.304343367 | 0.327 | 0.134 | 1.15E-10 | Lep-L |
| Macro-ISG | LIPA | 6.02E-22 | 2.276044068 | 0.597 | 0.395 | 1.19E-17 | Lep-L |
| Macro-ISG | APOE | 4.76E-35 | 2.252478784 | 0.77 | 0.467 | 9.36E-31 | Lep-L |
| Macro-ISG | LGMN | 2.87E-28 | 2.250151673 | 0.726 | 0.518 | 5.65E-24 | Lep-L |
| Macro-ISG | APOC1 | 5.10E-33 | 2.231935573 | 0.761 | 0.454 | 1.00E-28 | Lep-L |
| Macro-ISG | PLIN2 | 9.77E-26 | 2.210607112 | 0.752 | 0.612 | 1.92E-21 | Lep-L |
| Macro-ISG | MGLL | 1.98E-27 | 2.164105267 | 0.562 | 0.267 | 3.91E-23 | Lep-L |
| Macro-ISG | HSPA6 | 6.57E-26 | 1.988867143 | 0.774 | 0.556 | 1.29E-21 | Lep-L |
| Macro-ISG | VAT1 | 6.29E-09 | 1.844449525 | 0.332 | 0.198 | 0.000123871 | Lep-L |
| Macro-ISG | KLHL21 | 3.49E-08 | 1.834099492 | 0.274 | 0.144 | 0.000687791 | Lep-L |
| Macro-ISG | GBE1 | 2.36E-07 | 1.791903451 | 0.451 | 0.33 | 0.004648014 | Lep-L |
| Macro-ISG | STMN1 | 8.87E-10 | 1.729297862 | 0.354 | 0.2 | 1.75E-05 | Lep-L |
| Macro-ISG | CXCL8 | 2.49E-20 | 1.694621079 | 0.788 | 0.579 | 4.91E-16 | Lep-L |
| Macro-ISG | RASGRP3 | 7.43E-07 | 1.690633945 | 0.341 | 0.219 | 0.014630926 | Lep-L |
| Macro-ISG | HSPB1 | 9.41E-16 | 1.659925891 | 0.854 | 0.775 | 1.85E-11 | Lep-L |
| Macro-ISG | A2M | 3.90E-14 | 1.650237136 | 0.482 | 0.279 | 7.68E-10 | Lep-L |
| Macro-ISG | HSPA1B | 1.58E-22 | 1.628063054 | 0.965 | 0.878 | 3.11E-18 | Lep-L |
| Macro-ISG | SMS | 8.78E-18 | 1.626088848 | 0.615 | 0.445 | 1.73E-13 | Lep-L |
| Macro-ISG | CXCL3 | 1.52E-17 | 1.594204862 | 0.717 | 0.498 | 2.99E-13 | Lep-L |
| Macro-ISG | SDS | 4.87E-07 | 1.506679447 | 0.442 | 0.314 | 0.009590887 | Lep-L |
| Macro-ISG | DDIT4 | 9.52E-15 | 1.499715857 | 0.748 | 0.625 | 1.87E-10 | Lep-L |

| | | | | | | | |
|---|---|---|---|---|---|---|---|
| Macro-ISG | MYLIP | 1.86E-08 | 1.491759545 | 0.562 | 0.492 | 0.000366177 | Lep-L |
| Macro-ISG | ADM | 1.02E-09 | 1.439443265 | 0.593 | 0.452 | 2.00E-05 | Lep-L |
| Macro-ISG | DNAJB1 | 6.88E-22 | 1.389305979 | 0.938 | 0.889 | 1.35E-17 | Lep-L |
| Macro-ISG | CDC42EP3 | 4.06E-14 | 1.378311107 | 0.642 | 0.463 | 8.00E-10 | Lep-L |
| Macro-ISG | RBFOX2 | 1.73E-07 | 1.369559324 | 0.571 | 0.511 | 0.00341189 | Lep-L |
| Macro-ISG | BAG3 | 1.07E-12 | 1.363533203 | 0.704 | 0.559 | 2.10E-08 | Lep-L |
| Macro-ISG | EBI3 | 9.46E-07 | 1.341047084 | 0.358 | 0.226 | 0.018623596 | Lep-L |
| Macro-ISG | UGCG | 4.09E-11 | 1.337267701 | 0.593 | 0.468 | 8.04E-07 | Lep-L |
| Macro-ISG | ITPKB | 7.70E-08 | 1.304473291 | 0.274 | 0.141 | 0.001515096 | Lep-L |
| Macro-ISG | SLC1A3 | 1.13E-06 | 1.300676407 | 0.588 | 0.507 | 0.022243121 | Lep-L |
| Macro-ISG | CXCL2 | 9.03E-19 | 1.278109736 | 0.819 | 0.647 | 1.78E-14 | Lep-L |
| Macro-ISG | CFD | 1.71E-14 | 1.268233648 | 0.704 | 0.631 | 3.37E-10 | Lep-L |
| Macro-ISG | DDX3Y | 2.59E-13 | 1.244781654 | 0.549 | 0.315 | 5.09E-09 | Lep-L |
| Macro-ISG | CPM | 3.17E-07 | 1.240230159 | 0.527 | 0.444 | 0.006230098 | Lep-L |
| Macro-ISG | MIR155HG | 2.50E-10 | 1.237991912 | 0.615 | 0.455 | 4.92E-06 | Lep-L |
| Macro-ISG | PPP1R15A | 8.36E-30 | 1.231776358 | 0.965 | 0.909 | 1.65E-25 | Lep-L |
| Macro-ISG | CHKA | 1.15E-06 | 1.231286031 | 0.451 | 0.35 | 0.022621251 | Lep-L |
| Macro-ISG | HSPA1A | 2.21E-32 | 1.230293655 | 0.987 | 0.96 | 4.36E-28 | Lep-L |
| Macro-ISG | MAPRE2 | 2.11E-06 | 1.227481959 | 0.518 | 0.42 | 0.041502747 | Lep-L |
| Macro-ISG | NR4A1 | 1.32E-09 | 1.213930006 | 0.832 | 0.79 | 2.60E-05 | Lep-L |
| Macro-ISG | MAFF | 1.35E-07 | 1.204830067 | 0.602 | 0.526 | 0.002649854 | Lep-L |
| Macro-ISG | ABCA1 | 1.50E-13 | 1.20437245 | 0.792 | 0.734 | 2.96E-09 | Lep-L |
| Macro-ISG | ATP1B1 | 3.81E-13 | 1.193500066 | 0.695 | 0.626 | 7.49E-09 | Lep-L |
| Macro-ISG | PRNP | 3.50E-07 | 1.1879172 | 0.65 | 0.59 | 0.006881335 | Lep-L |
| Macro-ISG | RNASE1 | 2.88E-11 | 1.181511703 | 0.465 | 0.267 | 5.67E-07 | Lep-L |
| Macro-ISG | C5AR1 | 7.96E-10 | 1.169081809 | 0.695 | 0.623 | 1.57E-05 | Lep-L |
| Macro-ISG | ZFP36L1 | 1.97E-13 | 1.168560566 | 0.889 | 0.86 | 3.88E-09 | Lep-L |
| Macro-ISG | MYO9B | 3.22E-15 | 1.135953051 | 0.752 | 0.67 | 6.34E-11 | Lep-L |
| Macro-ISG | PDE4DIP | 1.86E-10 | 1.128827793 | 0.615 | 0.471 | 3.66E-06 | Lep-L |
| Macro-ISG | CSGALNACT2 | 1.21E-12 | 1.117768822 | 0.704 | 0.592 | 2.39E-08 | Lep-L |
| Macro-ISG | P4HA1 | 1.96E-06 | 1.107907206 | 0.544 | 0.468 | 0.038515463 | Lep-L |
| Macro-ISG | SGK1 | 3.54E-16 | 1.101807919 | 0.881 | 0.859 | 6.98E-12 | Lep-L |
| Macro-ISG | FCGR2A | 9.64E-11 | 1.094505012 | 0.872 | 0.892 | 1.90E-06 | Lep-L |
| Macro-ISG | CXCR4 | 4.02E-11 | 1.088613573 | 0.761 | 0.686 | 7.92E-07 | Lep-L |
| Macro-ISG | BRD2 | 5.27E-09 | 1.084695219 | 0.721 | 0.695 | 0.000103677 | Lep-L |
| Macro-ISG | NR4A2 | 1.41E-14 | 1.066895701 | 0.894 | 0.873 | 2.77E-10 | Lep-L |
| Macro-ISG | ZC3H12A | 1.17E-08 | 1.06050519 | 0.619 | 0.502 | 0.000229728 | Lep-L |
| Macro-ISG | GAS7 | 5.47E-07 | 1.044370047 | 0.566 | 0.466 | 0.01076183 | Lep-L |
| Macro-ISG | CD83 | 2.11E-30 | 1.042323008 | 0.956 | 0.941 | 4.16E-26 | Lep-L |
| Macro-ISG | FOSB | 3.21E-11 | 1.039297481 | 0.765 | 0.731 | 6.33E-07 | Lep-L |
| Macro-ISG | PDGFB | 2.53E-07 | 1.028386449 | 0.504 | 0.369 | 0.004988385 | Lep-L |
| Macro-ISG | MAP2K3 | 2.13E-06 | 1.019111534 | 0.611 | 0.588 | 0.041840184 | Lep-L |
| Macro-ISG | IER5 | 1.67E-09 | 1.016403768 | 0.792 | 0.73 | 3.30E-05 | Lep-L |
| Macro-ISG | FNIP2 | 3.07E-10 | 1.002586505 | 0.726 | 0.678 | 6.04E-06 | Lep-L |
| Macro-ISG | UBD | 3.33E-35 | 2.938996429 | 0.689 | 0.389 | 6.56E-31 | Lep-T |
| Macro-ISG | ALDOA | 1.21E-32 | 2.691228805 | 0.632 | 0.377 | 2.39E-28 | Lep-T |
| Macro-ISG | CCL19 | 4.38E-30 | 2.306894541 | 0.478 | 0.178 | 8.63E-26 | Lep-T |
| Macro-ISG | VCAM1 | 2.93E-26 | 2.111222158 | 0.406 | 0.14 | 5.76E-22 | Lep-T |
| Macro-ISG | PTGDS | 3.37E-17 | 2.067784096 | 0.267 | 0.086 | 6.63E-13 | Lep-T |
| Macro-ISG | ZFP361 | 1.06E-25 | 1.755575785 | 0.972 | 0.948 | 2.08E-21 | Lep-T |
| Macro-ISG | CD74 | 3.06E-56 | 1.748172214 | 1 | 0.999 | 6.02E-52 | Lep-T |
| Macro-ISG | ITGB2-AS1 | 3.22E-18 | 1.728402964 | 0.371 | 0.158 | 6.33E-14 | Lep-T |
| Macro-ISG | CFP | 4.17E-17 | 1.724697339 | 0.437 | 0.209 | 8.21E-13 | Lep-T |
| Macro-ISG | CXCL14 | 3.78E-19 | 1.624545698 | 0.261 | 0.071 | 7.44E-15 | Lep-T |
| Macro-ISG | HLA-DQB2 | 2.11E-17 | 1.606378272 | 0.453 | 0.228 | 4.15E-13 | Lep-T |

| | | | | | | | |
|---|---|---|---|---|---|---|---|
| Macro-ISG | RBFOX21 | 4.94E-39 | 1.600761153 | 0.767 | 0.445 | 9.72E-35 | Lep-T |
| Macro-ISG | TNF1 | 1.61E-07 | 1.534882371 | 0.569 | 0.453 | 0.003168314 | Lep-T |
| Macro-ISG | FSCN1 | 1.53E-12 | 1.5032383 | 0.443 | 0.261 | 3.01E-08 | Lep-T |
| Macro-ISG | HSD11B1 | 2.68E-13 | 1.480356038 | 0.415 | 0.227 | 5.27E-09 | Lep-T |
| Macro-ISG | CDC42EP2 | 2.62E-13 | 1.473056584 | 0.431 | 0.253 | 5.16E-09 | Lep-T |
| Macro-ISG | APOO | 2.21E-18 | 1.455045775 | 0.497 | 0.273 | 4.35E-14 | Lep-T |
| Macro-ISG | DUSP4 | 1.09E-12 | 1.438177355 | 0.255 | 0.098 | 2.15E-08 | Lep-T |
| Macro-ISG | EMD | 1.77E-24 | 1.437927996 | 0.553 | 0.268 | 3.48E-20 | Lep-T |
| Macro-ISG | U2AF1 | 9.35E-10 | 1.409909376 | 0.286 | 0.139 | 1.84E-05 | Lep-T |
| Macro-ISG | SIK1B | 6.73E-13 | 1.38774409 | 0.289 | 0.124 | 1.33E-08 | Lep-T |
| Macro-ISG | PNRC1 | 1.99E-44 | 1.362274573 | 0.981 | 0.962 | 3.91E-40 | Lep-T |
| Macro-ISG | MMP2 | 1.27E-08 | 1.357924253 | 0.258 | 0.129 | 0.000250683 | Lep-T |
| Macro-ISG | BCL3 | 1.51E-19 | 1.342940783 | 0.657 | 0.426 | 2.97E-15 | Lep-T |
| Macro-ISG | FKBP2 | 6.00E-09 | 1.340409831 | 0.358 | 0.22 | 0.000118197 | Lep-T |
| Macro-ISG | ERGIC3 | 1.09E-19 | 1.322826505 | 0.575 | 0.326 | 2.15E-15 | Lep-T |
| Macro-ISG | CDKN1A1 | 3.46E-26 | 1.316982427 | 0.868 | 0.799 | 6.82E-22 | Lep-T |
| Macro-ISG | FCN1 | 2.09E-18 | 1.316742513 | 0.563 | 0.326 | 4.12E-14 | Lep-T |
| Macro-ISG | HLA-DQA21 | 1.32E-27 | 1.297235401 | 0.745 | 0.486 | 2.60E-23 | Lep-T |
| Macro-ISG | SEC13 | 9.14E-13 | 1.275787657 | 0.459 | 0.271 | 1.80E-08 | Lep-T |
| Macro-ISG | NRROS | 3.47E-10 | 1.260742863 | 0.308 | 0.153 | 6.83E-06 | Lep-T |
| Macro-ISG | TNFRSF4 | 1.97E-09 | 1.244905486 | 0.314 | 0.166 | 3.89E-05 | Lep-T |
| Macro-ISG | CST3 | 2.85E-22 | 1.232974962 | 0.994 | 0.991 | 5.62E-18 | Lep-T |
| Macro-ISG | ICAM11 | 1.97E-23 | 1.21620977 | 0.868 | 0.796 | 3.89E-19 | Lep-T |
| Macro-ISG | ZNF581 | 4.75E-08 | 1.191368829 | 0.264 | 0.135 | 0.000934472 | Lep-T |
| Macro-ISG | INSIG1 | 1.23E-10 | 1.185782089 | 0.758 | 0.718 | 2.42E-06 | Lep-T |
| Macro-ISG | SYNGR2 | 1.06E-29 | 1.174178072 | 0.849 | 0.762 | 2.09E-25 | Lep-T |
| Macro-ISG | ATP5PB | 1.22E-28 | 1.170465683 | 0.726 | 0.479 | 2.41E-24 | Lep-T |
| Macro-ISG | MT-ND6 | 9.08E-09 | 1.165508898 | 0.314 | 0.169 | 0.000178706 | Lep-T |
| Macro-ISG | NR1H3 | 2.48E-11 | 1.15742736 | 0.519 | 0.362 | 4.89E-07 | Lep-T |
| Macro-ISG | TRAF1 | 4.84E-13 | 1.138407286 | 0.597 | 0.424 | 9.52E-09 | Lep-T |
| Macro-ISG | NFKBIA1 | 2.23E-16 | 1.11749671 | 0.981 | 0.967 | 4.40E-12 | Lep-T |
| Macro-ISG | AFDN | 8.06E-09 | 1.110320934 | 0.415 | 0.263 | 0.000158704 | Lep-T |
| Macro-ISG | MMP191 | 1.27E-12 | 1.103745074 | 0.421 | 0.234 | 2.50E-08 | Lep-T |
| Macro-ISG | TSPAN33 | 2.84E-14 | 1.098977717 | 0.72 | 0.568 | 5.59E-10 | Lep-T |
| Macro-ISG | INPP5K | 3.18E-08 | 1.089738619 | 0.274 | 0.139 | 0.000625782 | Lep-T |
| Macro-ISG | RAB24 | 3.85E-08 | 1.062383392 | 0.415 | 0.269 | 0.000757237 | Lep-T |
| Macro-ISG | LGALS2 | 1.34E-37 | 1.055756264 | 0.937 | 0.833 | 2.64E-33 | Lep-T |
| Macro-ISG | GPS1 | 1.64E-06 | 1.054176054 | 0.324 | 0.203 | 0.032332422 | Lep-T |
| Macro-ISG | ACOT7 | 5.49E-10 | 1.054068324 | 0.346 | 0.185 | 1.08E-05 | Lep-T |
| Macro-ISG | LMNA | 1.88E-18 | 1.053240513 | 0.83 | 0.653 | 3.70E-14 | Lep-T |
| Macro-ISG | CRTAP | 4.94E-13 | 1.046586094 | 0.654 | 0.478 | 9.72E-09 | Lep-T |
| Macro-ISG | GEM | 3.88E-09 | 1.035022933 | 0.443 | 0.278 | 7.63E-05 | Lep-T |
| Macro-ISG | POLR2G | 1.21E-14 | 1.029181075 | 0.547 | 0.317 | 2.39E-10 | Lep-T |
| Macro-ISG | ITGAX | 3.22E-10 | 1.009448884 | 0.629 | 0.513 | 6.34E-06 | Lep-T |
| Macro-ISG | PRKAR2B | 1.83E-09 | 1.004023914 | 0.412 | 0.246 | 3.61E-05 | Lep-T |
| Macro-ISG | FOLR3 | 1.23E-72 | 5.063602982 | 0.439 | 0.048 | 2.42E-68 | Mari |
| Macro-ISG | KRT16 | 3.38E-15 | 1.809840368 | 0.326 | 0.155 | 6.65E-11 | Mari |
| Macro-ISG | CASP5 | 2.12E-37 | 1.647846914 | 0.567 | 0.225 | 4.17E-33 | Mari |
| Macro-ISG | KRT6A1 | 2.14E-08 | 1.518722566 | 0.266 | 0.153 | 0.000421808 | Mari |
| Macro-ISG | RASGRP2 | 3.00E-11 | 1.407805129 | 0.272 | 0.128 | 5.90E-07 | Mari |
| Macro-ISG | CEACAM4 | 5.55E-24 | 1.37067394 | 0.496 | 0.229 | 1.09E-19 | Mari |
| Macro-ISG | SPATA5L1 | 6.41E-08 | 1.348484182 | 0.279 | 0.161 | 0.001260943 | Mari |
| Macro-ISG | PCED1B-AS1 | 1.94E-16 | 1.262595854 | 0.389 | 0.19 | 3.81E-12 | Mari |
| Macro-ISG | NME4 | 3.80E-10 | 1.260188786 | 0.292 | 0.153 | 7.49E-06 | Mari |
| Macro-ISG | CD14 | 6.95E-53 | 1.23947113 | 0.987 | 0.828 | 1.37E-48 | Mari |

| | | | | | | | |
|---|---|---|---|---|---|---|---|
| Macro-ISG | HLA-DRB5 | 1.20E-12 | 1.228298149 | 0.828 | 0.801 | 2.36E-08 | Mari |
| Macro-ISG | LILRA5 | 2.13E-12 | 1.222303234 | 0.324 | 0.154 | 4.20E-08 | Mari |
| Macro-ISG | ZEB2-AS1 | 1.55E-11 | 1.216294495 | 0.285 | 0.134 | 3.06E-07 | Mari |
| Macro-ISG | CCL3L1 | 1.20E-11 | 1.206431401 | 0.841 | 0.738 | 2.37E-07 | Mari |
| Macro-ISG | S100A91 | 3.39E-48 | 1.197066732 | 0.979 | 0.719 | 6.68E-44 | Mari |
| Macro-ISG | IL10 | 3.71E-16 | 1.189215084 | 0.41 | 0.204 | 7.30E-12 | Mari |
| Macro-ISG | F13A1 | 4.03E-24 | 1.185792402 | 0.574 | 0.306 | 7.94E-20 | Mari |
| Macro-ISG | VCAN | 3.33E-08 | 1.175041964 | 0.264 | 0.138 | 0.000654891 | Mari |
| Macro-ISG | RNF141 | 1.16E-10 | 1.165178542 | 0.319 | 0.167 | 2.28E-06 | Mari |
| Macro-ISG | UCP2 | 2.47E-40 | 1.164079192 | 0.896 | 0.668 | 4.87E-36 | Mari |
| Macro-ISG | IL7R | 1.40E-12 | 1.145093966 | 0.454 | 0.266 | 2.76E-08 | Mari |
| Macro-ISG | CORO1A | 3.84E-32 | 1.128801418 | 0.838 | 0.551 | 7.57E-28 | Mari |
| Macro-ISG | SATB1-AS1 | 1.38E-14 | 1.126885675 | 0.486 | 0.283 | 2.71E-10 | Mari |
| Macro-ISG | SDF2L1 | 1.35E-25 | 1.117482298 | 0.781 | 0.556 | 2.66E-21 | Mari |
| Macro-ISG | CARHSP1 | 2.48E-09 | 1.116975556 | 0.298 | 0.163 | 4.88E-05 | Mari |
| Macro-ISG | S100A7 | 1.61E-12 | 1.105127879 | 0.287 | 0.134 | 3.17E-08 | Mari |
| Macro-ISG | NOL12 | 3.19E-11 | 1.098932279 | 0.368 | 0.202 | 6.28E-07 | Mari |
| Macro-ISG | GALK1 | 2.80E-15 | 1.095674891 | 0.517 | 0.313 | 5.52E-11 | Mari |
| Macro-ISG | FCGR2B | 2.49E-19 | 1.078366003 | 0.765 | 0.558 | 4.91E-15 | Mari |
| Macro-ISG | NISCH | 2.92E-11 | 1.070385918 | 0.274 | 0.127 | 5.75E-07 | Mari |
| Macro-ISG | ABI3 | 4.80E-17 | 1.056444134 | 0.593 | 0.392 | 9.44E-13 | Mari |
| Macro-ISG | GAS6 | 1.49E-21 | 1.056362475 | 0.648 | 0.399 | 2.94E-17 | Mari |
| Macro-ISG | MYO1G | 2.22E-20 | 1.047707218 | 0.501 | 0.244 | 4.37E-16 | Mari |
| Macro-ISG | BOLA2 | 4.22E-15 | 1.03321831 | 0.449 | 0.247 | 8.30E-11 | Mari |
| Macro-ISG | CRELD2 | 4.91E-16 | 1.024341519 | 0.546 | 0.329 | 9.67E-12 | Mari |
| Macro-ISG | IMPA2 | 1.52E-07 | 1.014797736 | 0.251 | 0.136 | 0.002990868 | Mari |
| Macro-ISG | S100A81 | 3.87E-41 | 1.005538517 | 0.924 | 0.538 | 7.61E-37 | Mari |
| Macro-ISG | SLC35F1 | 3.29E-87 | 3.100430292 | 0.619 | 0.121 | 6.48E-83 | Tb |
| Macro-ISG | MT1G | 4.52E-17 | 2.368603584 | 0.25 | 0.083 | 8.90E-13 | Tb |
| Macro-ISG | FN1 | 9.15E-24 | 2.07577934 | 0.512 | 0.25 | 1.80E-19 | Tb |
| Macro-ISG | MAP3K7CL | 7.28E-23 | 1.900255275 | 0.383 | 0.152 | 1.43E-18 | Tb |
| Macro-ISG | KRT17 | 8.19E-14 | 1.771817166 | 0.262 | 0.101 | 1.61E-09 | Tb |
| Macro-ISG | GGT5 | 4.33E-35 | 1.660428175 | 0.512 | 0.19 | 8.52E-31 | Tb |
| Macro-ISG | MT1F | 1.19E-13 | 1.659607463 | 0.252 | 0.1 | 2.35E-09 | Tb |
| Macro-ISG | MCF2L2 | 4.58E-21 | 1.487037556 | 0.532 | 0.286 | 9.02E-17 | Tb |
| Macro-ISG | SOCS1 | 2.67E-31 | 1.474793403 | 0.791 | 0.52 | 5.25E-27 | Tb |
| Macro-ISG | TAF4B | 5.37E-11 | 1.472843831 | 0.369 | 0.207 | 1.06E-06 | Tb |
| Macro-ISG | PHLPP1 | 1.60E-08 | 1.454690048 | 0.405 | 0.262 | 0.00031495 | Tb |
| Macro-ISG | LRRK2 | 3.51E-26 | 1.444972082 | 0.621 | 0.324 | 6.90E-22 | Tb |
| Macro-ISG | ERI1 | 3.28E-16 | 1.409850161 | 0.444 | 0.23 | 6.46E-12 | Tb |
| Macro-ISG | MT1X | 1.11E-22 | 1.409228688 | 0.6 | 0.346 | 2.19E-18 | Tb |
| Macro-ISG | IDO1 | 7.88E-31 | 1.398277299 | 0.752 | 0.464 | 1.55E-26 | Tb |
| Macro-ISG | FBP1 | 7.72E-24 | 1.389302711 | 0.765 | 0.545 | 1.52E-19 | Tb |
| Macro-ISG | MT1E | 4.21E-14 | 1.386329223 | 0.359 | 0.176 | 8.28E-10 | Tb |
| Macro-ISG | TBCC | 2.22E-10 | 1.380431992 | 0.286 | 0.142 | 4.37E-06 | Tb |
| Macro-ISG | CRISPLD2 | 2.69E-13 | 1.38016916 | 0.269 | 0.111 | 5.30E-09 | Tb |
| Macro-ISG | CTSC | 1.40E-39 | 1.359881097 | 0.966 | 0.906 | 2.76E-35 | Tb |
| Macro-ISG | P2RY14 | 1.51E-21 | 1.312135591 | 0.529 | 0.263 | 2.97E-17 | Tb |
| Macro-ISG | CXCL9 | 6.47E-38 | 1.292255855 | 0.883 | 0.65 | 1.27E-33 | Tb |
| Macro-ISG | TMEM273 | 3.15E-12 | 1.275834771 | 0.308 | 0.143 | 6.19E-08 | Tb |
| Macro-ISG | SQLE | 1.19E-09 | 1.262965987 | 0.289 | 0.15 | 2.35E-05 | Tb |
| Macro-ISG | RIPOR2 | 7.38E-10 | 1.257835199 | 0.279 | 0.141 | 1.45E-05 | Tb |
| Macro-ISG | SEMA4D | 1.83E-21 | 1.209621814 | 0.59 | 0.329 | 3.61E-17 | Tb |
| Macro-ISG | S100A8 | 1.24E-38 | 1.208424885 | 0.883 | 0.544 | 2.44E-34 | Tb |
| Macro-ISG | EZH2 | 3.27E-15 | 1.208260453 | 0.566 | 0.353 | 6.44E-11 | Tb |

| | | | | | | | |
|---|---|---|---|---|---|---|---|
| Macro-ISG | RHOH | 1.36E-09 | 1.2051142 | 0.451 | 0.305 | 2.69E-05 | Tb |
| Macro-ISG | LMTK2 | 6.03E-11 | 1.192159178 | 0.265 | 0.12 | 1.19E-06 | Tb |
| Macro-ISG | HIST1H2AC | 1.13E-06 | 1.187447092 | 0.32 | 0.2 | 0.022206777 | Tb |
| Macro-ISG | ENPP2 | 3.39E-15 | 1.182726364 | 0.578 | 0.366 | 6.67E-11 | Tb |
| Macro-ISG | TERF1 | 5.01E-08 | 1.181080898 | 0.313 | 0.179 | 0.000986278 | Tb |
| Macro-ISG | HSPA1L | 3.04E-12 | 1.155065137 | 0.26 | 0.109 | 5.99E-08 | Tb |
| Macro-ISG | IL1RAP | 4.01E-09 | 1.150956095 | 0.267 | 0.134 | 7.89E-05 | Tb |
| Macro-ISG | TNFRSF10D | 9.83E-16 | 1.148180011 | 0.444 | 0.234 | 1.94E-11 | Tb |
| Macro-ISG | LRRC23 | 1.73E-23 | 1.147370866 | 0.553 | 0.287 | 3.41E-19 | Tb |
| Macro-ISG | SCARF1 | 2.36E-08 | 1.14598327 | 0.306 | 0.174 | 0.000464528 | Tb |
| Macro-ISG | LMNB1 | 9.26E-09 | 1.14454001 | 0.374 | 0.236 | 0.000182222 | Tb |
| Macro-ISG | S100A2 | 7.10E-13 | 1.14270759 | 0.286 | 0.119 | 1.40E-08 | Tb |
| Macro-ISG | BORCS6 | 1.02E-15 | 1.137232571 | 0.359 | 0.161 | 2.00E-11 | Tb |
| Macro-ISG | SLC30A1 | 1.64E-16 | 1.136699952 | 0.444 | 0.223 | 3.24E-12 | Tb |
| Macro-ISG | SUCNR1 | 1.68E-09 | 1.11708117 | 0.42 | 0.264 | 3.31E-05 | Tb |
| Macro-ISG | IL3RA | 2.75E-20 | 1.115654005 | 0.672 | 0.423 | 5.42E-16 | Tb |
| Macro-ISG | UGP2 | 4.96E-23 | 1.115378664 | 0.74 | 0.516 | 9.76E-19 | Tb |
| Macro-ISG | MFSD14B | 4.44E-15 | 1.114722969 | 0.553 | 0.324 | 8.74E-11 | Tb |
| Macro-ISG | BPGM | 1.94E-09 | 1.106690263 | 0.284 | 0.145 | 3.82E-05 | Tb |
| Macro-ISG | TRIM5 | 2.59E-10 | 1.104880303 | 0.262 | 0.121 | 5.09E-06 | Tb |
| Macro-ISG | FAM241A | 9.19E-19 | 1.088660232 | 0.692 | 0.44 | 1.81E-14 | Tb |
| Macro-ISG | LIMK2 | 7.43E-19 | 1.073811155 | 0.578 | 0.328 | 1.46E-14 | Tb |
| Macro-ISG | SLC39A8 | 2.77E-18 | 1.070949656 | 0.641 | 0.394 | 5.45E-14 | Tb |
| Macro-ISG | KIAA0319L | 3.63E-10 | 1.064213451 | 0.25 | 0.114 | 7.14E-06 | Tb |
| Macro-ISG | CCL8 | 2.03E-06 | 1.064049023 | 0.371 | 0.255 | 0.039876676 | Tb |
| Macro-ISG | HMGB2 | 9.85E-14 | 1.063167187 | 0.604 | 0.396 | 1.94E-09 | Tb |
| Macro-ISG | APOL1 | 1.72E-11 | 1.048794887 | 0.296 | 0.141 | 3.38E-07 | Tb |
| Macro-ISG | DENND10 | 2.84E-18 | 1.046404427 | 0.551 | 0.296 | 5.60E-14 | Tb |
| Macro-ISG | CISH | 3.57E-13 | 1.044745956 | 0.371 | 0.19 | 7.03E-09 | Tb |
| Macro-ISG | HINT3 | 5.48E-13 | 1.043456425 | 0.388 | 0.202 | 1.08E-08 | Tb |
| Macro-ISG | NUP50-DT | 5.94E-09 | 1.038766172 | 0.291 | 0.154 | 0.000116895 | Tb |
| Macro-ISG | PDSS1 | 6.84E-12 | 1.036075427 | 0.403 | 0.224 | 1.35E-07 | Tb |
| Macro-ISG | RHEBL1 | 6.55E-11 | 1.034064827 | 0.296 | 0.14 | 1.29E-06 | Tb |
| Macro-ISG | FLOT2 | 7.41E-08 | 1.031238324 | 0.286 | 0.161 | 0.001458965 | Tb |
| Macro-ISG | KCNJ2 | 1.63E-14 | 1.030389134 | 0.352 | 0.161 | 3.21E-10 | Tb |
| Macro-ISG | APOL2 | 6.57E-11 | 1.027893365 | 0.354 | 0.192 | 1.29E-06 | Tb |
| Macro-ISG | FPR2 | 1.35E-09 | 1.025336832 | 0.299 | 0.156 | 2.65E-05 | Tb |
| **Disease-associated genes in Macro-IER2** | | | | | | | |
| Macro-IER2 | TM4SF19 | 2.77E-15 | 4.105314 | 0.258 | 0.086 | 5.46E-11 | Lep-L |
| Macro-IER2 | HS3ST2 | 1.13E-51 | 3.92308234 | 0.541 | 0.114 | 2.23E-47 | Lep-L |
| Macro-IER2 | CYP27A1 | 1.41E-43 | 2.537872465 | 0.6 | 0.216 | 2.78E-39 | Lep-L |
| Macro-IER2 | PPARG | 4.45E-36 | 2.472302103 | 0.622 | 0.283 | 8.76E-32 | Lep-L |
| Macro-IER2 | OTOA | 2.39E-31 | 2.366515235 | 0.497 | 0.196 | 4.71E-27 | Lep-L |
| Macro-IER2 | PDK4 | 2.15E-29 | 2.288882077 | 0.526 | 0.2 | 4.24E-25 | Lep-L |
| Macro-IER2 | MIR3142HG | 7.90E-12 | 2.245778074 | 0.312 | 0.154 | 1.56E-07 | Lep-L |
| Macro-IER2 | PLIN2 | 1.31E-65 | 2.159755277 | 0.858 | 0.613 | 2.57E-61 | Lep-L |
| Macro-IER2 | COL4A2-AS2 | 2.61E-22 | 2.113515217 | 0.432 | 0.172 | 5.14E-18 | Lep-L |
| Macro-IER2 | BEST1 | 9.22E-08 | 2.061290835 | 0.315 | 0.2 | 0.001815747 | Lep-L |
| Macro-IER2 | DHRS3 | 4.33E-12 | 1.930704673 | 0.285 | 0.126 | 8.52E-08 | Lep-L |
| Macro-IER2 | PLPP3 | 7.43E-20 | 1.901293324 | 0.454 | 0.214 | 1.46E-15 | Lep-L |
| Macro-IER2 | APOC1 | 3.94E-50 | 1.87333856 | 0.861 | 0.515 | 7.75E-46 | Lep-L |
| Macro-IER2 | SLC19A2 | 1.38E-18 | 1.864349882 | 0.485 | 0.265 | 2.71E-14 | Lep-L |
| Macro-IER2 | CCL20 | 7.96E-08 | 1.858027685 | 0.285 | 0.158 | 0.001566402 | Lep-L |
| Macro-IER2 | SCD | 2.67E-12 | 1.82472981 | 0.386 | 0.222 | 5.26E-08 | Lep-L |
| Macro-IER2 | GBE1 | 5.97E-20 | 1.786408935 | 0.607 | 0.395 | 1.17E-15 | Lep-L |

| | | | | | | | |
|---|---|---|---|---|---|---|---|
| Macro-IER2 | LIPA | 1.97E-26 | 1.730100827 | 0.616 | 0.391 | 3.87E-22 | Lep-L |
| Macro-IER2 | GPNMB | 6.48E-46 | 1.724416158 | 0.764 | 0.411 | 1.28E-41 | Lep-L |
| Macro-IER2 | SDS | 9.98E-17 | 1.703122774 | 0.604 | 0.413 | 1.96E-12 | Lep-L |
| Macro-IER2 | APOE | 5.22E-43 | 1.648932046 | 0.895 | 0.599 | 1.03E-38 | Lep-L |
| Macro-IER2 | CXCL8 | 3.20E-36 | 1.63674722 | 0.834 | 0.637 | 6.30E-32 | Lep-L |
| Macro-IER2 | PDXDC1 | 4.72E-30 | 1.617539612 | 0.67 | 0.413 | 9.28E-26 | Lep-L |
| Macro-IER2 | COL4A2 | 8.25E-07 | 1.591826571 | 0.283 | 0.174 | 0.01624052 | Lep-L |
| Macro-IER2 | RASGRP3 | 3.19E-15 | 1.577408323 | 0.49 | 0.293 | 6.27E-11 | Lep-L |
| Macro-IER2 | AQP9 | 1.98E-11 | 1.570898623 | 0.505 | 0.345 | 3.89E-07 | Lep-L |
| Macro-IER2 | OLMALINC | 4.85E-08 | 1.532110735 | 0.29 | 0.168 | 0.000955658 | Lep-L |
| Macro-IER2 | CXCL1 | 4.52E-10 | 1.525001535 | 0.373 | 0.214 | 8.89E-06 | Lep-L |
| Macro-IER2 | DDIT4 | 1.31E-37 | 1.5144036 | 0.863 | 0.625 | 2.58E-33 | Lep-L |
| Macro-IER2 | TPRG1 | 9.58E-13 | 1.50487979 | 0.382 | 0.2 | 1.89E-08 | Lep-L |
| Macro-IER2 | CCRL2 | 1.19E-15 | 1.502080815 | 0.564 | 0.391 | 2.34E-11 | Lep-L |
| Macro-IER2 | PDE8A | 2.17E-18 | 1.499373123 | 0.568 | 0.361 | 4.26E-14 | Lep-L |
| Macro-IER2 | MERTK | 2.83E-07 | 1.497910769 | 0.276 | 0.16 | 0.005576861 | Lep-L |
| Macro-IER2 | ALDH1A1 | 4.42E-07 | 1.493668915 | 0.485 | 0.383 | 0.008692571 | Lep-L |
| Macro-IER2 | CXCL2 | 4.76E-37 | 1.480811158 | 0.877 | 0.701 | 9.38E-33 | Lep-L |
| Macro-IER2 | SEPHS2 | 3.98E-23 | 1.453144733 | 0.735 | 0.531 | 7.84E-19 | Lep-L |
| Macro-IER2 | CXCL3 | 1.54E-25 | 1.39658839 | 0.789 | 0.559 | 3.03E-21 | Lep-L |
| Macro-IER2 | ABCC5 | 6.19E-11 | 1.394063111 | 0.314 | 0.158 | 1.22E-06 | Lep-L |
| Macro-IER2 | SMS | 2.79E-19 | 1.386467679 | 0.6 | 0.445 | 5.50E-15 | Lep-L |
| Macro-IER2 | GADD45B | 1.52E-49 | 1.384000049 | 0.982 | 0.96 | 2.99E-45 | Lep-L |
| Macro-IER2 | ZNF703 | 5.98E-07 | 1.361269241 | 0.285 | 0.17 | 0.011777771 | Lep-L |
| Macro-IER2 | MMP19 | 1.44E-07 | 1.360169937 | 0.414 | 0.299 | 0.002826742 | Lep-L |
| Macro-IER2 | FABP3 | 3.94E-07 | 1.358498737 | 0.274 | 0.158 | 0.007763435 | Lep-L |
| Macro-IER2 | KMO | 1.31E-10 | 1.324773527 | 0.595 | 0.453 | 2.57E-06 | Lep-L |
| Macro-IER2 | ITPKB | 3.59E-10 | 1.324377603 | 0.378 | 0.224 | 7.07E-06 | Lep-L |
| Macro-IER2 | STAM | 5.06E-07 | 1.317098051 | 0.303 | 0.194 | 0.009964625 | Lep-L |
| Macro-IER2 | MGLL | 5.79E-29 | 1.309070424 | 0.688 | 0.383 | 1.14E-24 | Lep-L |
| Macro-IER2 | TCEAL9 | 1.46E-08 | 1.307766672 | 0.366 | 0.228 | 0.000286413 | Lep-L |
| Macro-IER2 | OPN3 | 2.17E-09 | 1.307128976 | 0.445 | 0.327 | 4.28E-05 | Lep-L |
| Macro-IER2 | PHACTR1 | 5.97E-30 | 1.261119356 | 0.841 | 0.649 | 1.17E-25 | Lep-L |
| Macro-IER2 | FNIP2 | 2.31E-30 | 1.259638532 | 0.861 | 0.693 | 4.54E-26 | Lep-L |
| Macro-IER2 | IL6R-AS1 | 9.62E-08 | 1.258446514 | 0.305 | 0.178 | 0.001894469 | Lep-L |
| Macro-IER2 | SLC1A3 | 1.28E-19 | 1.253041211 | 0.708 | 0.511 | 2.51E-15 | Lep-L |
| Macro-IER2 | DDX3Y | 7.96E-26 | 1.246637169 | 0.641 | 0.311 | 1.57E-21 | Lep-L |
| Macro-IER2 | ADTRP | 8.66E-07 | 1.241454104 | 0.328 | 0.21 | 0.01703683 | Lep-L |
| Macro-IER2 | ADCY3 | 9.38E-07 | 1.239753109 | 0.375 | 0.259 | 0.018460708 | Lep-L |
| Macro-IER2 | RGS1 | 1.14E-11 | 1.232206524 | 0.768 | 0.647 | 2.24E-07 | Lep-L |
| Macro-IER2 | C5AR1 | 2.73E-22 | 1.229358794 | 0.76 | 0.577 | 5.38E-18 | Lep-L |
| Macro-IER2 | CXCR4 | 1.12E-18 | 1.228741123 | 0.795 | 0.643 | 2.20E-14 | Lep-L |
| Macro-IER2 | FABP5 | 5.27E-11 | 1.224735386 | 0.695 | 0.635 | 1.04E-06 | Lep-L |
| Macro-IER2 | CFD | 1.20E-13 | 1.21431499 | 0.641 | 0.543 | 2.36E-09 | Lep-L |
| Macro-IER2 | JUN | 9.63E-31 | 1.201520409 | 0.993 | 0.96 | 1.90E-26 | Lep-L |
| Macro-IER2 | CSTB | 6.35E-27 | 1.182039631 | 0.883 | 0.774 | 1.25E-22 | Lep-L |
| Macro-IER2 | ADM | 2.95E-21 | 1.175688008 | 0.71 | 0.485 | 5.81E-17 | Lep-L |
| Macro-IER2 | RAB42 | 5.52E-08 | 1.162592231 | 0.272 | 0.152 | 0.0010874 | Lep-L |
| Macro-IER2 | MIR155HG | 2.17E-08 | 1.161392078 | 0.595 | 0.469 | 0.000426406 | Lep-L |
| Macro-IER2 | TSC22D2 | 5.20E-14 | 1.157818857 | 0.771 | 0.625 | 1.02E-09 | Lep-L |
| Macro-IER2 | TAGAP | 1.42E-12 | 1.152503923 | 0.762 | 0.655 | 2.80E-08 | Lep-L |
| Macro-IER2 | HMOX1 | 2.78E-12 | 1.142971564 | 0.723 | 0.625 | 5.46E-08 | Lep-L |
| Macro-IER2 | ZNF503 | 7.16E-09 | 1.141121864 | 0.283 | 0.142 | 0.000140968 | Lep-L |
| Macro-IER2 | HSPA6 | 2.71E-19 | 1.138810819 | 0.791 | 0.603 | 5.34E-15 | Lep-L |
| Macro-IER2 | HSPB1 | 7.31E-15 | 1.136320831 | 0.912 | 0.824 | 1.44E-10 | Lep-L |

| | | | | | | | |
|---|---|---|---|---|---|---|---|
| Macro-IER2 | NFKBIA | 1.59E-43 | 1.131375287 | 0.986 | 0.966 | 3.13E-39 | Lep-L |
| Macro-IER2 | OXSM | 6.41E-09 | 1.119296909 | 0.445 | 0.307 | 0.000126247 | Lep-L |
| Macro-IER2 | HSPA1B | 5.71E-31 | 1.107838834 | 0.98 | 0.938 | 1.12E-26 | Lep-L |
| Macro-IER2 | ASAH1 | 6.56E-30 | 1.10123296 | 0.843 | 0.701 | 1.29E-25 | Lep-L |
| Macro-IER2 | BLVRB | 1.05E-22 | 1.087589238 | 0.777 | 0.675 | 2.07E-18 | Lep-L |
| Macro-IER2 | ZFP36 | 3.95E-10 | 1.075610535 | 0.933 | 0.904 | 7.77E-06 | Lep-L |
| Macro-IER2 | IER3 | 2.34E-36 | 1.074163506 | 0.971 | 0.946 | 4.61E-32 | Lep-L |
| Macro-IER2 | HSPA1A | 4.22E-46 | 1.059378488 | 0.998 | 0.968 | 8.31E-42 | Lep-L |
| Macro-IER2 | CHCHD6 | 4.34E-10 | 1.049922419 | 0.373 | 0.22 | 8.54E-06 | Lep-L |
| Macro-IER2 | FOSB | 2.12E-24 | 1.045498676 | 0.872 | 0.758 | 4.18E-20 | Lep-L |
| Macro-IER2 | PPP1R15A | 7.74E-35 | 1.039599162 | 0.987 | 0.946 | 1.52E-30 | Lep-L |
| Macro-IER2 | PDE4DIP | 1.48E-17 | 1.033750347 | 0.703 | 0.513 | 2.92E-13 | Lep-L |
| Macro-IER2 | ABCA1 | 1.43E-16 | 1.030073571 | 0.838 | 0.687 | 2.82E-12 | Lep-L |
| Macro-IER2 | SH3BP5 | 2.69E-18 | 1.029366011 | 0.715 | 0.553 | 5.30E-14 | Lep-L |
| Macro-IER2 | TSC22D1 | 1.61E-08 | 1.023503513 | 0.674 | 0.561 | 0.000316251 | Lep-L |
| Macro-IER2 | CDC42EP3 | 1.91E-23 | 1.015408416 | 0.75 | 0.507 | 3.76E-19 | Lep-L |
| Macro-IER2 | CHMP1B | 5.74E-12 | 1.012092469 | 0.744 | 0.639 | 1.13E-07 | Lep-L |
| Macro-IER2 | UBD | 1.68E-10 | 3.337381922 | 0.335 | 0.148 | 3.30E-06 | Lep-T |
| Macro-IER2 | TVP23A | 7.88E-10 | 2.131092657 | 0.308 | 0.131 | 1.55E-05 | Lep-T |
| Macro-IER2 | IL21R | 1.30E-06 | 1.842594276 | 0.253 | 0.124 | 0.025559789 | Lep-T |
| Macro-IER2 | AFF3 | 2.07E-08 | 1.820554462 | 0.253 | 0.104 | 0.000408392 | Lep-T |
| Macro-IER2 | PTGDS | 3.25E-08 | 1.751370823 | 0.297 | 0.133 | 0.000640071 | Lep-T |
| Macro-IER2 | P2RY14 | 1.34E-09 | 1.680889823 | 0.264 | 0.1 | 2.65E-05 | Lep-T |
| Macro-IER2 | CXCL14 | 2.39E-19 | 1.678521347 | 0.308 | 0.072 | 4.71E-15 | Lep-T |
| Macro-IER2 | LGALS2 | 2.19E-10 | 1.669436115 | 0.56 | 0.435 | 4.30E-06 | Lep-T |
| Macro-IER2 | RTN1 | 4.64E-15 | 1.609846336 | 0.527 | 0.266 | 9.12E-11 | Lep-T |
| Macro-IER2 | AFDN | 4.82E-13 | 1.5424239 | 0.385 | 0.157 | 9.49E-09 | Lep-T |
| Macro-IER2 | PID1 | 8.43E-07 | 1.524000727 | 0.313 | 0.17 | 0.016602643 | Lep-T |
| Macro-IER2 | RFTN1 | 2.37E-06 | 1.506245842 | 0.418 | 0.279 | 0.046685532 | Lep-T |
| Macro-IER2 | MB21D2 | 2.17E-06 | 1.208319269 | 0.467 | 0.333 | 0.042654329 | Lep-T |
| Macro-IER2 | CCSER1 | 5.33E-09 | 1.181014343 | 0.588 | 0.384 | 0.000104881 | Lep-T |
| Macro-IER2 | MVB12B | 2.14E-06 | 1.159503405 | 0.28 | 0.134 | 0.042133203 | Lep-T |
| Macro-IER2 | SNAP23 | 1.63E-07 | 1.158310595 | 0.709 | 0.578 | 0.00320238 | Lep-T |
| Macro-IER2 | GBP4 | 1.00E-07 | 1.135038564 | 0.462 | 0.281 | 0.001968809 | Lep-T |
| Macro-IER2 | MRC1 | 2.15E-08 | 1.106515285 | 0.533 | 0.351 | 0.000422451 | Lep-T |
| Macro-IER2 | F13A1 | 1.36E-06 | 1.048972064 | 0.33 | 0.18 | 0.026866103 | Lep-T |
| Macro-IER2 | STAT1 | 8.49E-14 | 1.008985139 | 0.78 | 0.604 | 1.67E-09 | Lep-T |
| Macro-IER2 | FOLR3 | 9.24E-65 | 5.528057386 | 0.391 | 0.014 | 1.82E-60 | Mari |
| Macro-IER2 | S100A81 | 1.97E-37 | 2.28946611 | 0.848 | 0.347 | 3.89E-33 | Mari |
| Macro-IER2 | CASP5 | 5.26E-14 | 2.209570115 | 0.304 | 0.088 | 1.04E-09 | Mari |
| Macro-IER2 | CD93 | 2.27E-11 | 2.032811472 | 0.275 | 0.086 | 4.46E-07 | Mari |
| Macro-IER2 | FCGR1B1 | 5.87E-27 | 1.995088158 | 0.616 | 0.204 | 1.16E-22 | Mari |
| Macro-IER2 | FCGR1A1 | 4.60E-25 | 1.855164519 | 0.754 | 0.396 | 9.06E-21 | Mari |
| Macro-IER2 | CD300E | 5.14E-14 | 1.833874451 | 0.312 | 0.09 | 1.01E-09 | Mari |
| Macro-IER2 | VAMP51 | 1.17E-24 | 1.798762036 | 0.761 | 0.408 | 2.30E-20 | Mari |
| Macro-IER2 | SLC1A2 | 2.32E-08 | 1.784959323 | 0.304 | 0.128 | 0.000457233 | Mari |
| Macro-IER2 | KRT6B | 7.47E-20 | 1.78407535 | 0.275 | 0.051 | 1.47E-15 | Mari |
| Macro-IER2 | CALHM61 | 1.30E-25 | 1.750976937 | 0.804 | 0.483 | 2.55E-21 | Mari |
| Macro-IER2 | CAMK1 | 6.93E-16 | 1.736122991 | 0.399 | 0.13 | 1.36E-11 | Mari |
| Macro-IER2 | TMEM170B | 5.74E-08 | 1.680810643 | 0.297 | 0.129 | 0.001130619 | Mari |
| Macro-IER2 | SECTM11 | 6.72E-15 | 1.675044348 | 0.493 | 0.198 | 1.32E-10 | Mari |
| Macro-IER2 | MRPL231 | 2.71E-18 | 1.671564274 | 0.5 | 0.182 | 5.33E-14 | Mari |
| Macro-IER2 | F13A11 | 2.16E-12 | 1.659433612 | 0.42 | 0.174 | 4.25E-08 | Mari |
| Macro-IER2 | S100A41 | 2.08E-14 | 1.634763257 | 0.812 | 0.646 | 4.09E-10 | Mari |
| Macro-IER2 | IL7R1 | 1.62E-11 | 1.625937493 | 0.384 | 0.155 | 3.18E-07 | Mari |

| | | | | | | | |
|---|---|---|---|---|---|---|---|
| Macro-IER2 | SDF2L11 | 4.06E-14 | 1.619262115 | 0.543 | 0.25 | 7.99E-10 | Mari |
| Macro-IER2 | MARCO | 1.16E-07 | 1.588948106 | 0.471 | 0.271 | 0.002281006 | Mari |
| Macro-IER2 | C1QB | 1.69E-22 | 1.583375864 | 0.862 | 0.62 | 3.33E-18 | Mari |
| Macro-IER2 | KRT161 | 7.97E-15 | 1.569828275 | 0.304 | 0.085 | 1.57E-10 | Mari |
| Macro-IER2 | VSIG4 | 6.05E-14 | 1.551819445 | 0.572 | 0.291 | 1.19E-09 | Mari |
| Macro-IER2 | CXCL102 | 2.95E-13 | 1.546605932 | 0.609 | 0.318 | 5.81E-09 | Mari |
| Macro-IER2 | CCL81 | 5.40E-11 | 1.538348898 | 0.399 | 0.159 | 1.06E-06 | Mari |
| Macro-IER2 | EBPL | 3.05E-08 | 1.538345626 | 0.275 | 0.108 | 0.000599628 | Mari |
| Macro-IER2 | RNASE1 | 2.19E-14 | 1.53192332 | 0.514 | 0.221 | 4.30E-10 | Mari |
| Macro-IER2 | CCDC167 | 4.56E-09 | 1.524327675 | 0.275 | 0.104 | 8.97E-05 | Mari |
| Macro-IER2 | SPATA5L1 | 9.04E-08 | 1.517741117 | 0.268 | 0.11 | 0.001778881 | Mari |
| Macro-IER2 | MYO1G1 | 2.62E-16 | 1.514473154 | 0.399 | 0.124 | 5.15E-12 | Mari |
| Macro-IER2 | MRPL581 | 1.18E-08 | 1.51271013 | 0.312 | 0.129 | 0.000232441 | Mari |
| Macro-IER2 | GPR84 | 2.80E-10 | 1.512321865 | 0.413 | 0.184 | 5.50E-06 | Mari |
| Macro-IER2 | HRH21 | 1.16E-10 | 1.510856156 | 0.42 | 0.186 | 2.28E-06 | Mari |
| Macro-IER2 | IFI351 | 1.99E-12 | 1.496791796 | 0.449 | 0.186 | 3.92E-08 | Mari |
| Macro-IER2 | PLAAT41 | 2.10E-26 | 1.488916135 | 0.746 | 0.325 | 4.14E-22 | Mari |
| Macro-IER2 | TMEM1091 | 9.28E-13 | 1.472457986 | 0.493 | 0.219 | 1.83E-08 | Mari |
| Macro-IER2 | FCGR2B | 2.85E-18 | 1.467696122 | 0.703 | 0.362 | 5.61E-14 | Mari |
| Macro-IER2 | IFITM3 | 5.32E-14 | 1.466033537 | 0.783 | 0.579 | 1.05E-09 | Mari |
| Macro-IER2 | MRPL28 | 3.91E-09 | 1.447292695 | 0.391 | 0.186 | 7.70E-05 | Mari |
| Macro-IER2 | POLE41 | 2.87E-14 | 1.439229213 | 0.507 | 0.228 | 5.65E-10 | Mari |
| Macro-IER2 | SCO21 | 2.63E-19 | 1.431609416 | 0.71 | 0.353 | 5.18E-15 | Mari |
| Macro-IER2 | C1QC | 4.91E-18 | 1.425795898 | 0.826 | 0.616 | 9.66E-14 | Mari |
| Macro-IER2 | CORO1A | 6.51E-13 | 1.424841603 | 0.616 | 0.328 | 1.28E-08 | Mari |
| Macro-IER2 | MILR1 | 1.90E-07 | 1.423905308 | 0.326 | 0.147 | 0.003743382 | Mari |
| Macro-IER2 | TMEM173 | 3.99E-10 | 1.41873767 | 0.377 | 0.158 | 7.86E-06 | Mari |
| Macro-IER2 | FCER1G | 1.83E-28 | 1.417173517 | 0.92 | 0.746 | 3.60E-24 | Mari |
| Macro-IER2 | TSFM1 | 3.22E-07 | 1.415807882 | 0.297 | 0.133 | 0.006341682 | Mari |
| Macro-IER2 | HLA-DRB5 | 1.13E-10 | 1.415447293 | 0.826 | 0.659 | 2.23E-06 | Mari |
| Macro-IER2 | CRIP1 | 1.60E-16 | 1.411674666 | 0.754 | 0.414 | 3.15E-12 | Mari |
| Macro-IER2 | VSIR | 5.07E-19 | 1.41041803 | 0.775 | 0.41 | 9.99E-15 | Mari |
| Macro-IER2 | USB1 | 3.44E-10 | 1.405692789 | 0.326 | 0.124 | 6.78E-06 | Mari |
| Macro-IER2 | EMB | 1.11E-11 | 1.405348297 | 0.341 | 0.121 | 2.19E-07 | Mari |
| Macro-IER2 | GALK11 | 4.36E-10 | 1.401912792 | 0.377 | 0.16 | 8.59E-06 | Mari |
| Macro-IER2 | S100A91 | 6.35E-21 | 1.381380486 | 0.862 | 0.558 | 1.25E-16 | Mari |
| Macro-IER2 | INHBA | 1.44E-11 | 1.374092148 | 0.514 | 0.241 | 2.83E-07 | Mari |
| Macro-IER2 | NAPRT | 6.88E-09 | 1.357752752 | 0.413 | 0.205 | 0.000135459 | Mari |
| Macro-IER2 | BST2 | 1.72E-19 | 1.356021575 | 0.79 | 0.523 | 3.39E-15 | Mari |
| Macro-IER2 | FCGR3A1 | 9.85E-19 | 1.355875249 | 0.768 | 0.479 | 1.94E-14 | Mari |
| Macro-IER2 | PSMB31 | 2.13E-19 | 1.353442427 | 0.768 | 0.456 | 4.20E-15 | Mari |
| Macro-IER2 | TYMP1 | 8.37E-33 | 1.350508016 | 0.978 | 0.733 | 1.65E-28 | Mari |
| Macro-IER2 | SLC35B1 | 8.05E-09 | 1.348942335 | 0.355 | 0.16 | 0.000158514 | Mari |
| Macro-IER2 | PCED1B-AS1 | 1.16E-09 | 1.3480555 | 0.326 | 0.128 | 2.29E-05 | Mari |
| Macro-IER2 | COMMD4 | 1.10E-12 | 1.33863918 | 0.493 | 0.223 | 2.17E-08 | Mari |
| Macro-IER2 | RRAS1 | 2.09E-12 | 1.327223567 | 0.522 | 0.253 | 4.11E-08 | Mari |
| Macro-IER2 | PSME21 | 4.87E-21 | 1.325161591 | 0.833 | 0.538 | 9.59E-17 | Mari |
| Macro-IER2 | OLFML2B | 2.30E-08 | 1.325081539 | 0.362 | 0.163 | 0.000453174 | Mari |
| Macro-IER2 | TOMM40 | 5.40E-08 | 1.322104067 | 0.355 | 0.167 | 0.001063429 | Mari |
| Macro-IER2 | SATB1-AS1 | 1.14E-13 | 1.320698564 | 0.58 | 0.277 | 2.24E-09 | Mari |
| Macro-IER2 | SLC25A39 | 6.59E-12 | 1.307441991 | 0.478 | 0.221 | 1.30E-07 | Mari |
| Macro-IER2 | DDX60L | 9.50E-10 | 1.304154993 | 0.486 | 0.25 | 1.87E-05 | Mari |
| Macro-IER2 | TLR81 | 4.42E-09 | 1.28575025 | 0.377 | 0.166 | 8.70E-05 | Mari |
| Macro-IER2 | CYBA1 | 4.83E-35 | 1.285033231 | 0.957 | 0.791 | 9.51E-31 | Mari |
| Macro-IER2 | FBXO62 | 6.41E-11 | 1.28324179 | 0.471 | 0.219 | 1.26E-06 | Mari |

| | | | | | | | |
|---|---|---|---|---|---|---|---|
| Macro-IER2 | CD141 | 3.67E-20 | 1.270247597 | 0.899 | 0.677 | 7.22E-16 | Mari |
| Macro-IER2 | BST1 | 7.48E-07 | 1.256652287 | 0.312 | 0.147 | 0.014716058 | Mari |
| Macro-IER2 | MT1X1 | 2.17E-06 | 1.25555908 | 0.486 | 0.285 | 0.042716924 | Mari |
| Macro-IER2 | PLTP | 2.50E-11 | 1.251131728 | 0.667 | 0.415 | 4.93E-07 | Mari |
| Macro-IER2 | CCR1 | 6.73E-12 | 1.249504335 | 0.5 | 0.228 | 1.32E-07 | Mari |
| Macro-IER2 | S100A61 | 8.50E-21 | 1.249384844 | 0.949 | 0.791 | 1.67E-16 | Mari |
| Macro-IER2 | PCNX11 | 8.92E-15 | 1.246099429 | 0.601 | 0.286 | 1.76E-10 | Mari |
| Macro-IER2 | TEX41 | 2.03E-07 | 1.243837828 | 0.37 | 0.175 | 0.003994064 | Mari |
| Macro-IER2 | SBNO2 | 4.37E-08 | 1.243652171 | 0.435 | 0.218 | 0.000860226 | Mari |
| Macro-IER2 | ARL21 | 7.15E-10 | 1.239788519 | 0.493 | 0.252 | 1.41E-05 | Mari |
| Macro-IER2 | UBE2L61 | 2.63E-15 | 1.23729388 | 0.659 | 0.345 | 5.18E-11 | Mari |
| Macro-IER2 | MNDA1 | 1.20E-10 | 1.233528618 | 0.761 | 0.504 | 2.37E-06 | Mari |
| Macro-IER2 | PLSCR1 | 1.92E-14 | 1.229560296 | 0.797 | 0.547 | 3.79E-10 | Mari |
| Macro-IER2 | AIF1 | 2.00E-19 | 1.22506952 | 0.899 | 0.743 | 3.94E-15 | Mari |
| Macro-IER2 | HSD17B101 | 6.15E-09 | 1.223427181 | 0.406 | 0.188 | 0.000121125 | Mari |
| Macro-IER2 | APOL41 | 9.74E-09 | 1.21920823 | 0.283 | 0.11 | 0.000191759 | Mari |
| Macro-IER2 | MVB12A | 9.85E-08 | 1.217152607 | 0.384 | 0.183 | 0.001939181 | Mari |
| Macro-IER2 | ARHGDIB | 1.37E-12 | 1.2160094 | 0.783 | 0.574 | 2.70E-08 | Mari |
| Macro-IER2 | PRELID11 | 9.03E-15 | 1.212563263 | 0.775 | 0.522 | 1.78E-10 | Mari |
| Macro-IER2 | S100A101 | 2.86E-16 | 1.210216399 | 0.87 | 0.75 | 5.63E-12 | Mari |
| Macro-IER2 | SLC11A1 | 1.63E-10 | 1.203980255 | 0.5 | 0.257 | 3.21E-06 | Mari |
| Macro-IER2 | B4GALT7 | 1.22E-06 | 1.201811271 | 0.254 | 0.109 | 0.02401076 | Mari |
| Macro-IER2 | PDCD6 | 3.25E-11 | 1.201128433 | 0.674 | 0.389 | 6.41E-07 | Mari |
| Macro-IER2 | MS4A6A | 3.53E-09 | 1.198305048 | 0.768 | 0.638 | 6.96E-05 | Mari |
| Macro-IER2 | GBP11 | 8.46E-17 | 1.196930451 | 0.754 | 0.421 | 1.67E-12 | Mari |
| Macro-IER2 | PFN11 | 4.30E-22 | 1.18301537 | 0.884 | 0.749 | 8.47E-18 | Mari |
| Macro-IER2 | DUSP23 | 1.36E-12 | 1.179173398 | 0.565 | 0.276 | 2.68E-08 | Mari |
| Macro-IER2 | PTPMT11 | 3.92E-10 | 1.176798253 | 0.486 | 0.241 | 7.71E-06 | Mari |
| Macro-IER2 | ANKRD221 | 5.00E-19 | 1.176508402 | 0.536 | 0.186 | 9.85E-15 | Mari |
| Macro-IER2 | 46092 | 1.36E-12 | 1.17496856 | 0.71 | 0.42 | 2.67E-08 | Mari |
| Macro-IER2 | TXNDC171 | 1.71E-14 | 1.174463614 | 0.681 | 0.371 | 3.36E-10 | Mari |
| Macro-IER2 | SERPINA11 | 6.35E-17 | 1.169056704 | 0.812 | 0.527 | 1.25E-12 | Mari |
| Macro-IER2 | UQCC2 | 3.44E-08 | 1.16507831 | 0.486 | 0.27 | 0.000676167 | Mari |
| Macro-IER2 | EPSTI11 | 9.07E-21 | 1.160975594 | 0.848 | 0.473 | 1.79E-16 | Mari |
| Macro-IER2 | TMEM2081 | 1.81E-13 | 1.160841603 | 0.623 | 0.338 | 3.56E-09 | Mari |
| Macro-IER2 | TIMM17B | 3.74E-08 | 1.160737883 | 0.384 | 0.186 | 0.000735598 | Mari |
| Macro-IER2 | DCXR | 3.40E-10 | 1.160512939 | 0.457 | 0.214 | 6.69E-06 | Mari |
| Macro-IER2 | EXOSC1 | 1.07E-08 | 1.156056935 | 0.312 | 0.129 | 0.00021003 | Mari |
| Macro-IER2 | CD33 | 1.73E-07 | 1.155121362 | 0.391 | 0.194 | 0.003403166 | Mari |
| Macro-IER2 | BANF1 | 6.98E-12 | 1.153397487 | 0.645 | 0.366 | 1.37E-07 | Mari |
| Macro-IER2 | PSMB91 | 3.71E-18 | 1.150163062 | 0.783 | 0.525 | 7.30E-14 | Mari |
| Macro-IER2 | ZNF593 | 3.07E-08 | 1.150115147 | 0.355 | 0.164 | 0.000603875 | Mari |
| Macro-IER2 | COMMD8 | 1.38E-07 | 1.147515774 | 0.442 | 0.238 | 0.002715559 | Mari |
| Macro-IER2 | DYNLT11 | 2.99E-14 | 1.135562973 | 0.703 | 0.414 | 5.88E-10 | Mari |
| Macro-IER2 | 46083 | 8.22E-13 | 1.130344031 | 0.659 | 0.378 | 1.62E-08 | Mari |
| Macro-IER2 | FAM89B1 | 2.76E-09 | 1.128653453 | 0.435 | 0.216 | 5.44E-05 | Mari |
| Macro-IER2 | CENPX1 | 1.12E-09 | 1.127169381 | 0.507 | 0.269 | 2.21E-05 | Mari |
| Macro-IER2 | MRPL201 | 1.59E-12 | 1.125686482 | 0.652 | 0.367 | 3.13E-08 | Mari |
| Macro-IER2 | C1QA | 1.70E-16 | 1.123409302 | 0.848 | 0.705 | 3.35E-12 | Mari |
| Macro-IER2 | HLA-DRB11 | 1.29E-07 | 1.121379309 | 0.949 | 0.917 | 0.002534503 | Mari |
| Macro-IER2 | IFI30 | 8.23E-21 | 1.117964178 | 0.957 | 0.925 | 1.62E-16 | Mari |
| Macro-IER2 | NDUFA9 | 1.51E-09 | 1.117774448 | 0.478 | 0.24 | 2.97E-05 | Mari |
| Macro-IER2 | CASP41 | 3.96E-13 | 1.110122885 | 0.732 | 0.46 | 7.79E-09 | Mari |
| Macro-IER2 | CFL1 | 2.50E-23 | 1.108556363 | 0.935 | 0.772 | 4.92E-19 | Mari |
| Macro-IER2 | PARP91 | 2.95E-11 | 1.102780285 | 0.464 | 0.204 | 5.80E-07 | Mari |

| | | | | | | | |
|---|---|---|---|---|---|---|---|
| Macro-IER2 | CD163 | 2.85E-11 | 1.10163136 | 0.739 | 0.487 | 5.61E-07 | Mari |
| Macro-IER2 | JOSD21 | 6.69E-11 | 1.101269584 | 0.5 | 0.248 | 1.32E-06 | Mari |
| Macro-IER2 | SDSL | 1.53E-06 | 1.096780242 | 0.42 | 0.239 | 0.03021019 | Mari |
| Macro-IER2 | DBNL1 | 1.55E-10 | 1.093172668 | 0.616 | 0.348 | 3.04E-06 | Mari |
| Macro-IER2 | SLIRP | 3.60E-14 | 1.090622005 | 0.667 | 0.364 | 7.08E-10 | Mari |
| Macro-IER2 | SLC16A101 | 1.97E-08 | 1.086270071 | 0.587 | 0.336 | 0.000387507 | Mari |
| Macro-IER2 | CHMP2A1 | 1.05E-11 | 1.084973728 | 0.659 | 0.397 | 2.07E-07 | Mari |
| Macro-IER2 | COPZ11 | 1.00E-07 | 1.078634698 | 0.449 | 0.246 | 0.001977814 | Mari |
| Macro-IER2 | GSTK1 | 6.20E-13 | 1.073087186 | 0.775 | 0.536 | 1.22E-08 | Mari |
| Macro-IER2 | SNF8 | 1.43E-10 | 1.071471677 | 0.601 | 0.356 | 2.81E-06 | Mari |
| Macro-IER2 | GSDMD | 3.98E-09 | 1.071043393 | 0.572 | 0.302 | 7.83E-05 | Mari |
| Macro-IER2 | IGFLR1 | 1.01E-13 | 1.066414574 | 0.71 | 0.436 | 1.99E-09 | Mari |
| Macro-IER2 | DUSP3 | 1.56E-07 | 1.056304409 | 0.464 | 0.251 | 0.00306848 | Mari |
| Macro-IER2 | PIN1 | 4.46E-12 | 1.055356592 | 0.551 | 0.266 | 8.78E-08 | Mari |
| Macro-IER2 | NUDT16 | 4.48E-08 | 1.055067418 | 0.442 | 0.228 | 0.000881489 | Mari |
| Macro-IER2 | BLOC1S11 | 8.63E-15 | 1.0549779 | 0.783 | 0.551 | 1.70E-10 | Mari |
| Macro-IER2 | SNX6 | 6.26E-12 | 1.053646068 | 0.775 | 0.576 | 1.23E-07 | Mari |
| Macro-IER2 | PRDX51 | 6.27E-14 | 1.053390328 | 0.732 | 0.46 | 1.23E-09 | Mari |
| Macro-IER2 | TSPO1 | 1.51E-17 | 1.052267493 | 0.862 | 0.631 | 2.97E-13 | Mari |
| Macro-IER2 | PSMB101 | 7.83E-12 | 1.051986368 | 0.703 | 0.402 | 1.54E-07 | Mari |
| Macro-IER2 | IL10RB | 8.81E-11 | 1.051017889 | 0.609 | 0.34 | 1.73E-06 | Mari |
| Macro-IER2 | PPIB1 | 1.49E-15 | 1.048376206 | 0.848 | 0.607 | 2.93E-11 | Mari |
| Macro-IER2 | PYCARD1 | 4.52E-15 | 1.047276245 | 0.804 | 0.556 | 8.89E-11 | Mari |
| Macro-IER2 | ZNHIT1 | 3.84E-09 | 1.03483897 | 0.623 | 0.385 | 7.57E-05 | Mari |
| Macro-IER2 | FBXL15 | 8.48E-09 | 1.034337207 | 0.42 | 0.202 | 0.000166954 | Mari |
| Macro-IER2 | MICOS13 | 3.45E-10 | 1.031779041 | 0.645 | 0.414 | 6.78E-06 | Mari |
| Macro-IER2 | CYBC11 | 4.03E-08 | 1.03142462 | 0.558 | 0.345 | 0.000792296 | Mari |
| Macro-IER2 | LMAN2 | 3.41E-11 | 1.029181435 | 0.667 | 0.397 | 6.72E-07 | Mari |
| Macro-IER2 | CUEDC2 | 2.28E-08 | 1.028553755 | 0.478 | 0.257 | 0.000449447 | Mari |
| Macro-IER2 | HIGD1A1 | 6.41E-07 | 1.027827527 | 0.348 | 0.17 | 0.012617609 | Mari |
| Macro-IER2 | DSE1 | 1.52E-13 | 1.026623822 | 0.833 | 0.556 | 2.98E-09 | Mari |
| Macro-IER2 | PSMA51 | 6.68E-13 | 1.022160897 | 0.71 | 0.428 | 1.32E-08 | Mari |
| Macro-IER2 | EIF4E21 | 6.82E-09 | 1.02182445 | 0.507 | 0.283 | 0.000134196 | Mari |
| Macro-IER2 | GAS6 | 1.87E-08 | 1.017541087 | 0.493 | 0.273 | 0.000368215 | Mari |
| Macro-IER2 | ENPP21 | 1.48E-07 | 1.016969104 | 0.377 | 0.187 | 0.002907059 | Mari |
| Macro-IER2 | LTC4S | 2.52E-08 | 1.016961715 | 0.304 | 0.123 | 0.00049594 | Mari |
| Macro-IER2 | PSMB6 | 5.09E-13 | 1.014819481 | 0.739 | 0.475 | 1.00E-08 | Mari |
| Macro-IER2 | MIEN1 | 4.55E-10 | 1.014512231 | 0.493 | 0.249 | 8.95E-06 | Mari |
| Macro-IER2 | PSME11 | 3.57E-15 | 1.014250156 | 0.775 | 0.491 | 7.04E-11 | Mari |
| Macro-IER2 | AGPAT2 | 2.63E-10 | 1.013490116 | 0.543 | 0.294 | 5.18E-06 | Mari |
| Macro-IER2 | VAMP81 | 1.55E-20 | 1.013478411 | 0.884 | 0.665 | 3.04E-16 | Mari |
| Macro-IER2 | FIBP1 | 7.86E-07 | 1.012197078 | 0.471 | 0.276 | 0.015477621 | Mari |
| Macro-IER2 | KRT141 | 2.87E-08 | 1.010820684 | 0.406 | 0.21 | 0.000564894 | Mari |
| Macro-IER2 | PTPN21 | 3.77E-12 | 1.007104658 | 0.732 | 0.43 | 7.42E-08 | Mari |
| Macro-IER2 | GIMAP41 | 8.91E-12 | 1.00534035 | 0.594 | 0.319 | 1.75E-07 | Mari |
| Macro-IER2 | CIB11 | 9.33E-14 | 1.004670362 | 0.761 | 0.488 | 1.84E-09 | Mari |
| Macro-IER2 | TMEM147 | 8.54E-10 | 1.004239821 | 0.507 | 0.261 | 1.68E-05 | Mari |
| Macro-IER2 | SDHAF2 | 1.47E-07 | 1.002045772 | 0.377 | 0.186 | 0.002898761 | Mari |
| Macro-IER2 | KRT6A | 2.87E-26 | 3.314837029 | 0.313 | 0.055 | 5.66E-22 | Tb |
| Macro-IER2 | GGT5 | 6.77E-25 | 3.215359829 | 0.257 | 0.037 | 1.33E-20 | Tb |
| Macro-IER2 | IDO1 | 2.30E-21 | 2.761721978 | 0.436 | 0.146 | 4.54E-17 | Tb |
| Macro-IER2 | KRT16 | 8.06E-22 | 2.754461842 | 0.324 | 0.071 | 1.59E-17 | Tb |
| Macro-IER2 | CXCL9 | 6.90E-23 | 2.648363592 | 0.547 | 0.218 | 1.36E-18 | Tb |
| Macro-IER2 | S100A7 | 1.97E-13 | 2.536607781 | 0.274 | 0.083 | 3.88E-09 | Tb |
| Macro-IER2 | SLC35F1 | 1.39E-34 | 2.500960827 | 0.425 | 0.079 | 2.74E-30 | Tb |

| | | | | | | | |
|---|---|---|---|---|---|---|---|
| Macro-IER2 | S100A2 | 9.99E-18 | 2.403127218 | 0.296 | 0.075 | 1.97E-13 | Tb |
| Macro-IER2 | FN1 | 6.17E-12 | 2.276580962 | 0.453 | 0.231 | 1.21E-07 | Tb |
| Macro-IER2 | FBP1 | 2.62E-12 | 2.224405894 | 0.587 | 0.398 | 5.15E-08 | Tb |
| Macro-IER2 | RIPOR2 | 2.63E-08 | 2.202858829 | 0.263 | 0.11 | 0.000517065 | Tb |
| Macro-IER2 | KCNJ2 | 6.92E-12 | 2.065591441 | 0.257 | 0.08 | 1.36E-07 | Tb |
| Macro-IER2 | YAE1 | 2.45E-10 | 2.023398053 | 0.257 | 0.087 | 4.82E-06 | Tb |
| Macro-IER2 | P2RY141 | 1.47E-21 | 1.986578243 | 0.346 | 0.083 | 2.89E-17 | Tb |
| Macro-IER2 | VDR | 1.08E-14 | 1.982740372 | 0.318 | 0.101 | 2.13E-10 | Tb |
| Macro-IER2 | PPA1 | 6.05E-22 | 1.935217696 | 0.676 | 0.381 | 1.19E-17 | Tb |
| Macro-IER2 | ANKRD22 | 1.38E-14 | 1.892308524 | 0.447 | 0.187 | 2.73E-10 | Tb |
| Macro-IER2 | CISH | 6.91E-16 | 1.882917706 | 0.279 | 0.073 | 1.36E-11 | Tb |
| Macro-IER2 | APOL4 | 2.14E-16 | 1.830631529 | 0.324 | 0.094 | 4.22E-12 | Tb |
| Macro-IER2 | HAPLN3 | 4.99E-18 | 1.753244865 | 0.408 | 0.131 | 9.83E-14 | Tb |
| Macro-IER2 | CASS4 | 1.71E-12 | 1.740399898 | 0.296 | 0.098 | 3.37E-08 | Tb |
| Macro-IER2 | MMP12 | 1.22E-20 | 1.736760185 | 0.263 | 0.049 | 2.39E-16 | Tb |
| Macro-IER2 | ACSS2 | 1.02E-12 | 1.717658276 | 0.391 | 0.152 | 2.00E-08 | Tb |
| Macro-IER2 | KDM1B | 1.02E-06 | 1.707256435 | 0.33 | 0.167 | 0.020171689 | Tb |
| Macro-IER2 | QPCT | 2.16E-14 | 1.670933829 | 0.369 | 0.137 | 4.26E-10 | Tb |
| Macro-IER2 | TYMP | 1.09E-43 | 1.658406988 | 0.944 | 0.728 | 2.14E-39 | Tb |
| Macro-IER2 | PRKAR2B | 7.06E-07 | 1.652576098 | 0.313 | 0.153 | 0.013900906 | Tb |
| Macro-IER2 | LYRM1 | 3.70E-10 | 1.648968046 | 0.385 | 0.173 | 7.28E-06 | Tb |
| Macro-IER2 | S100A8 | 1.10E-35 | 1.643996623 | 0.804 | 0.333 | 2.16E-31 | Tb |
| Macro-IER2 | ERAP2 | 1.10E-14 | 1.629076498 | 0.38 | 0.131 | 2.17E-10 | Tb |
| Macro-IER2 | CXCL101 | 9.60E-16 | 1.567352814 | 0.592 | 0.307 | 1.89E-11 | Tb |
| Macro-IER2 | KRT14 | 3.88E-15 | 1.555027702 | 0.464 | 0.189 | 7.63E-11 | Tb |
| Macro-IER2 | MBOAT7 | 1.84E-06 | 1.535380481 | 0.369 | 0.194 | 0.036300034 | Tb |
| Macro-IER2 | TNFRSF10D | 3.17E-09 | 1.525163395 | 0.559 | 0.328 | 6.24E-05 | Tb |
| Macro-IER2 | TREM1 | 4.60E-10 | 1.512689301 | 0.33 | 0.133 | 9.06E-06 | Tb |
| Macro-IER2 | IRF4 | 2.12E-07 | 1.510780792 | 0.324 | 0.161 | 0.004173379 | Tb |
| Macro-IER2 | HS3ST3B1 | 3.82E-07 | 1.505794412 | 0.263 | 0.117 | 0.007525019 | Tb |
| Macro-IER2 | NMT1 | 2.24E-08 | 1.495141975 | 0.425 | 0.218 | 0.000440201 | Tb |
| Macro-IER2 | NIBAN2 | 1.10E-10 | 1.487360008 | 0.318 | 0.126 | 2.16E-06 | Tb |
| Macro-IER2 | ACOT71 | 3.50E-13 | 1.48275722 | 0.425 | 0.177 | 6.88E-09 | Tb |
| Macro-IER2 | GBP1 | 2.69E-23 | 1.465046103 | 0.76 | 0.405 | 5.30E-19 | Tb |
| Macro-IER2 | CYSLTR2 | 2.93E-10 | 1.44797284 | 0.257 | 0.087 | 5.78E-06 | Tb |
| Macro-IER2 | HOMER1 | 4.11E-09 | 1.446685661 | 0.464 | 0.275 | 8.09E-05 | Tb |
| Macro-IER2 | PLAAT4 | 1.29E-15 | 1.443283328 | 0.631 | 0.329 | 2.55E-11 | Tb |
| Macro-IER2 | ARHGEF3 | 1.61E-07 | 1.418250781 | 0.363 | 0.185 | 0.003163566 | Tb |
| Macro-IER2 | ENPP2 | 6.73E-16 | 1.406747414 | 0.441 | 0.165 | 1.32E-11 | Tb |
| Macro-IER2 | DENND6B | 4.93E-08 | 1.398086506 | 0.251 | 0.099 | 0.000969884 | Tb |
| Macro-IER2 | NUB1 | 1.18E-14 | 1.392143866 | 0.626 | 0.329 | 2.32E-10 | Tb |
| Macro-IER2 | TNFSF10 | 5.53E-22 | 1.380218496 | 0.698 | 0.337 | 1.09E-17 | Tb |
| Macro-IER2 | ARID5A | 2.03E-14 | 1.374878812 | 0.615 | 0.313 | 4.00E-10 | Tb |
| Macro-IER2 | IL2RA | 4.25E-08 | 1.372670445 | 0.296 | 0.131 | 0.000835843 | Tb |
| Macro-IER2 | SELENOM | 3.44E-07 | 1.366382532 | 0.291 | 0.131 | 0.006774314 | Tb |
| Macro-IER2 | TES | 6.53E-11 | 1.360816587 | 0.486 | 0.233 | 1.28E-06 | Tb |
| Macro-IER2 | S100A9 | 2.23E-21 | 1.348372001 | 0.86 | 0.544 | 4.38E-17 | Tb |
| Macro-IER2 | PFKP1 | 1.36E-16 | 1.339321518 | 0.402 | 0.138 | 2.67E-12 | Tb |
| Macro-IER2 | VCL | 8.23E-07 | 1.334916378 | 0.352 | 0.177 | 0.016199578 | Tb |
| Macro-IER2 | ASCL21 | 2.46E-12 | 1.334459548 | 0.408 | 0.169 | 4.85E-08 | Tb |
| Macro-IER2 | THBD | 4.28E-10 | 1.330206828 | 0.341 | 0.146 | 8.43E-06 | Tb |
| Macro-IER2 | NUP93 | 8.18E-07 | 1.318829121 | 0.263 | 0.114 | 0.016096501 | Tb |
| Macro-IER2 | RFX5 | 3.00E-08 | 1.316825825 | 0.251 | 0.096 | 0.000591225 | Tb |
| Macro-IER2 | P2RY6 | 3.62E-13 | 1.315806661 | 0.335 | 0.118 | 7.13E-09 | Tb |
| Macro-IER2 | APOBEC3G | 1.49E-08 | 1.31118977 | 0.318 | 0.139 | 0.000292516 | Tb |

| | | | | | | | |
|---|---|---|---|---|---|---|---|
| Macro-IER2 | SEMA4D | 5.51E-11 | 1.300725441 | 0.503 | 0.248 | 1.08E-06 | Tb |
| Macro-IER2 | CDC123 | 3.40E-08 | 1.298617814 | 0.363 | 0.17 | 0.000668867 | Tb |
| Macro-IER2 | KSR11 | 9.53E-09 | 1.294361863 | 0.318 | 0.136 | 0.000187515 | Tb |
| Macro-IER2 | ACLY | 7.75E-10 | 1.293401203 | 0.413 | 0.187 | 1.53E-05 | Tb |
| Macro-IER2 | PEF1 | 1.22E-06 | 1.286124145 | 0.263 | 0.118 | 0.024006652 | Tb |
| Macro-IER2 | FADS1 | 3.00E-10 | 1.282522838 | 0.335 | 0.138 | 5.90E-06 | Tb |
| Macro-IER2 | PSME2 | 2.83E-19 | 1.270439235 | 0.754 | 0.541 | 5.57E-15 | Tb |
| Macro-IER2 | ATR | 7.12E-11 | 1.264020182 | 0.363 | 0.151 | 1.40E-06 | Tb |
| Macro-IER2 | RNF19A | 1.66E-07 | 1.260613637 | 0.564 | 0.349 | 0.00327655 | Tb |
| Macro-IER2 | LYZ | 3.56E-15 | 1.259439572 | 0.877 | 0.759 | 7.01E-11 | Tb |
| Macro-IER2 | VAMP5 | 1.35E-17 | 1.255785681 | 0.704 | 0.403 | 2.67E-13 | Tb |
| Macro-IER2 | MFSD14B | 1.07E-08 | 1.239407748 | 0.531 | 0.311 | 0.000209738 | Tb |
| Macro-IER2 | MAP3K13 | 3.36E-07 | 1.227312585 | 0.453 | 0.257 | 0.006620322 | Tb |
| Macro-IER2 | HMGN4 | 3.82E-09 | 1.217029395 | 0.402 | 0.197 | 7.53E-05 | Tb |
| Macro-IER2 | INO80C | 9.33E-08 | 1.215055837 | 0.251 | 0.102 | 0.001837052 | Tb |
| Macro-IER2 | VAMP7 | 1.79E-06 | 1.186861769 | 0.263 | 0.12 | 0.035158524 | Tb |
| Macro-IER2 | VPS9D11 | 1.18E-08 | 1.182300167 | 0.341 | 0.157 | 0.000232009 | Tb |
| Macro-IER2 | SAMD9L | 4.71E-13 | 1.181812227 | 0.48 | 0.208 | 9.27E-09 | Tb |
| Macro-IER2 | IL4R | 1.84E-08 | 1.177706051 | 0.453 | 0.237 | 0.000362564 | Tb |
| Macro-IER2 | CD274 | 9.22E-17 | 1.177193809 | 0.492 | 0.185 | 1.82E-12 | Tb |
| Macro-IER2 | LAP3 | 1.15E-23 | 1.175887843 | 0.911 | 0.68 | 2.26E-19 | Tb |
| Macro-IER2 | CTSC | 1.14E-10 | 1.17345283 | 0.799 | 0.721 | 2.23E-06 | Tb |
| Macro-IER2 | ARHGAP30 | 1.23E-07 | 1.168682829 | 0.447 | 0.242 | 0.002424626 | Tb |
| Macro-IER2 | CSF2RB1 | 2.83E-09 | 1.165241575 | 0.413 | 0.198 | 5.57E-05 | Tb |
| Macro-IER2 | N4BP1 | 3.00E-08 | 1.160188367 | 0.615 | 0.409 | 0.000589912 | Tb |
| Macro-IER2 | SNX201 | 2.57E-09 | 1.147855494 | 0.335 | 0.142 | 5.06E-05 | Tb |
| Macro-IER2 | LY75 | 1.44E-07 | 1.138048115 | 0.374 | 0.191 | 0.002826601 | Tb |
| Macro-IER2 | GBP41 | 1.09E-14 | 1.137081192 | 0.559 | 0.262 | 2.15E-10 | Tb |
| Macro-IER2 | SREBF21 | 3.91E-07 | 1.133454616 | 0.324 | 0.157 | 0.007695681 | Tb |
| Macro-IER2 | APOL6 | 1.33E-08 | 1.129731052 | 0.542 | 0.305 | 0.000262541 | Tb |
| Macro-IER2 | BAK1 | 8.20E-07 | 1.12707008 | 0.279 | 0.129 | 0.016142203 | Tb |
| Macro-IER2 | CD69 | 6.14E-15 | 1.125302761 | 0.525 | 0.232 | 1.21E-10 | Tb |
| Macro-IER2 | MTHFD2 | 1.25E-07 | 1.122771345 | 0.732 | 0.504 | 0.002452166 | Tb |
| Macro-IER2 | SIGLEC10 | 3.80E-10 | 1.120794619 | 0.363 | 0.153 | 7.48E-06 | Tb |
| Macro-IER2 | GBP51 | 2.17E-17 | 1.106959262 | 0.743 | 0.402 | 4.28E-13 | Tb |
| Macro-IER2 | PDCD1LG21 | 2.83E-15 | 1.102558 | 0.408 | 0.146 | 5.56E-11 | Tb |
| Macro-IER2 | SLC39A8 | 9.15E-08 | 1.10130439 | 0.514 | 0.315 | 0.00180145 | Tb |
| Macro-IER2 | ADAMDEC1 | 1.44E-08 | 1.10120464 | 0.497 | 0.31 | 0.000283322 | Tb |
| Macro-IER2 | TGM2 | 2.20E-06 | 1.100800901 | 0.279 | 0.134 | 0.043207843 | Tb |
| Macro-IER2 | CD38 | 1.59E-09 | 1.100764661 | 0.413 | 0.205 | 3.14E-05 | Tb |
| Macro-IER2 | CCDC25 | 1.03E-10 | 1.100547553 | 0.374 | 0.153 | 2.03E-06 | Tb |
| Macro-IER2 | TAP11 | 5.30E-14 | 1.099234689 | 0.631 | 0.336 | 1.04E-09 | Tb |
| Macro-IER2 | RNF213 | 1.65E-11 | 1.098748569 | 0.883 | 0.655 | 3.24E-07 | Tb |
| Macro-IER2 | SCLT1 | 7.91E-11 | 1.095843351 | 0.654 | 0.398 | 1.56E-06 | Tb |
| Macro-IER2 | PVT1 | 1.14E-07 | 1.095592253 | 0.33 | 0.159 | 0.002242848 | Tb |
| Macro-IER2 | PSMC3 | 2.15E-08 | 1.093634009 | 0.447 | 0.241 | 0.000423981 | Tb |
| Macro-IER2 | PCNX1 | 1.20E-07 | 1.090061356 | 0.503 | 0.291 | 0.002366457 | Tb |
| Macro-IER2 | LRRC25 | 2.37E-08 | 1.082306048 | 0.458 | 0.253 | 0.000466068 | Tb |
| Macro-IER2 | GIMAP71 | 8.99E-11 | 1.080274406 | 0.385 | 0.162 | 1.77E-06 | Tb |
| Macro-IER2 | HNRNPLL | 1.85E-09 | 1.078879603 | 0.335 | 0.143 | 3.64E-05 | Tb |
| Macro-IER2 | SEPTIN6 | 6.09E-10 | 1.077750656 | 0.587 | 0.353 | 1.20E-05 | Tb |
| Macro-IER2 | CD401 | 6.56E-15 | 1.075065624 | 0.665 | 0.36 | 1.29E-10 | Tb |
| Macro-IER2 | ITPR1 | 1.42E-13 | 1.071207328 | 0.709 | 0.449 | 2.79E-09 | Tb |
| Macro-IER2 | PSMB10 | 1.05E-13 | 1.068566968 | 0.67 | 0.394 | 2.07E-09 | Tb |
| Macro-IER2 | SLC38A10 | 1.08E-06 | 1.062812618 | 0.313 | 0.147 | 0.02128436 | Tb |

| | | | | | | | |
|---|---|---|---|---|---|---|---|
| Macro-IER2 | WSB2 | 8.13E-08 | 1.061772384 | 0.324 | 0.145 | 0.001600455 | Tb |
| Macro-IER2 | ABTB2 | 6.71E-08 | 1.060379275 | 0.48 | 0.269 | 0.001321773 | Tb |
| Macro-IER2 | CALHM6 | 6.72E-14 | 1.059840765 | 0.726 | 0.483 | 1.32E-09 | Tb |
| Macro-IER2 | GZF1 | 1.77E-07 | 1.048812926 | 0.43 | 0.235 | 0.003493148 | Tb |
| Macro-IER2 | MAP2K1 | 1.09E-06 | 1.048417615 | 0.637 | 0.453 | 0.021513009 | Tb |
| Macro-IER2 | LIMK2 | 1.41E-10 | 1.045981997 | 0.475 | 0.231 | 2.78E-06 | Tb |
| Macro-IER2 | AK2 | 1.51E-09 | 1.041980216 | 0.48 | 0.235 | 2.96E-05 | Tb |
| Macro-IER2 | IMPAD1 | 8.48E-07 | 1.041290068 | 0.346 | 0.168 | 0.016696294 | Tb |
| Macro-IER2 | VRK2 | 2.10E-09 | 1.036966049 | 0.62 | 0.368 | 4.12E-05 | Tb |
| Macro-IER2 | TMX4 | 8.40E-07 | 1.028253857 | 0.447 | 0.251 | 0.016540614 | Tb |
| Macro-IER2 | AUH | 1.09E-06 | 1.023283117 | 0.263 | 0.117 | 0.021496961 | Tb |
| Macro-IER2 | TTC7A | 8.07E-07 | 1.020467215 | 0.425 | 0.24 | 0.01588549 | Tb |
| Macro-IER2 | IL3RA | 6.53E-14 | 1.01735697 | 0.62 | 0.326 | 1.28E-09 | Tb |
| Macro-IER2 | RAB31 | 2.61E-09 | 1.016384517 | 0.754 | 0.552 | 5.13E-05 | Tb |
| Macro-IER2 | AIMP2 | 1.90E-08 | 1.01474117 | 0.296 | 0.125 | 0.000374028 | Tb |
| Macro-IER2 | RAB13 | 2.43E-06 | 1.010611313 | 0.564 | 0.408 | 0.047810648 | Tb |
| Macro-IER2 | RNF411 | 1.51E-06 | 1.00639161 | 0.302 | 0.146 | 0.029732599 | Tb |

| Table S7 \| Disease-associated genes in NKs | | | | | | | |
|---|---|---|---|---|---|---|---|
| Disease-associated genes in all NK | | | | | | | |
| group | gene | p_val | avg_log2FC | pct.1 | pct.2 | p_val_adj | cluster |
| allcell | PRKY | 7.21E-76 | 2.30885135 | 0.273 | 0.074 | 1.42E-71 | Lep-L |
| allcell | DDX3Y | 3.92E-92 | 2.044175283 | 0.373 | 0.116 | 7.72E-88 | Lep-L |
| allcell | APOE | 1.90E-51 | 1.578342822 | 0.403 | 0.209 | 3.73E-47 | Lep-L |
| allcell | RBFOX2 | 1.22E-36 | 1.496868668 | 0.421 | 0.272 | 2.39E-32 | Lep-L |
| allcell | TNFRSF1B | 4.70E-45 | 1.404133456 | 0.474 | 0.306 | 9.26E-41 | Lep-L |
| allcell | HLA-DQB1 | 2.64E-34 | 1.398016765 | 0.383 | 0.233 | 5.20E-30 | Lep-L |
| allcell | ATP1B3 | 5.22E-85 | 1.387430962 | 0.721 | 0.512 | 1.03E-80 | Lep-L |
| allcell | PRDM1 | 7.34E-23 | 1.369769245 | 0.278 | 0.16 | 1.45E-18 | Lep-L |
| allcell | TLE1 | 5.81E-27 | 1.332814889 | 0.317 | 0.186 | 1.14E-22 | Lep-L |
| allcell | CNN2 | 1.60E-35 | 1.289138779 | 0.442 | 0.286 | 3.16E-31 | Lep-L |
| allcell | SLC2A3 | 1.96E-34 | 1.201828054 | 0.644 | 0.524 | 3.87E-30 | Lep-L |
| allcell | SPOCK2 | 2.64E-23 | 1.172479868 | 0.301 | 0.176 | 5.20E-19 | Lep-L |
| allcell | MEF2D | 3.92E-16 | 1.147138173 | 0.284 | 0.191 | 7.71E-12 | Lep-L |
| allcell | CDKN1A | 4.80E-19 | 1.143309003 | 0.357 | 0.25 | 9.45E-15 | Lep-L |
| allcell | ATXN1 | 5.35E-26 | 1.14265337 | 0.48 | 0.35 | 1.05E-21 | Lep-L |
| allcell | ZFP36L1 | 6.76E-13 | 1.134611168 | 0.614 | 0.583 | 1.33E-08 | Lep-L |
| allcell | EMD | 2.53E-13 | 1.101135415 | 0.262 | 0.178 | 4.97E-09 | Lep-L |
| allcell | INSIG1 | 1.42E-16 | 1.093001795 | 0.366 | 0.265 | 2.80E-12 | Lep-L |
| allcell | PIM3 | 9.56E-34 | 1.08327891 | 0.475 | 0.316 | 1.88E-29 | Lep-L |
| allcell | DUSP5 | 1.19E-17 | 1.073954087 | 0.441 | 0.349 | 2.34E-13 | Lep-L |
| allcell | TGFBR2 | 1.94E-26 | 1.073125785 | 0.452 | 0.319 | 3.82E-22 | Lep-L |
| allcell | CSRNP1 | 4.98E-20 | 1.057890866 | 0.459 | 0.362 | 9.80E-16 | Lep-L |
| allcell | RBM38 | 9.23E-25 | 1.049660762 | 0.425 | 0.301 | 1.82E-20 | Lep-L |
| allcell | ZC3H12A | 1.23E-10 | 1.047434466 | 0.408 | 0.351 | 2.42E-06 | Lep-L |
| allcell | UBALD2 | 2.09E-20 | 1.046292475 | 0.376 | 0.265 | 4.12E-16 | Lep-L |
| allcell | METRNL | 1.06E-39 | 1.038831052 | 0.638 | 0.512 | 2.08E-35 | Lep-L |
| allcell | CCL4L2 | 2.71E-32 | 1.037520888 | 0.473 | 0.303 | 5.32E-28 | Lep-L |
| allcell | TAP1 | 2.00E-24 | 1.035620345 | 0.37 | 0.243 | 3.95E-20 | Lep-L |
| allcell | BIRC3 | 1.49E-20 | 1.033596667 | 0.535 | 0.438 | 2.94E-16 | Lep-L |
| allcell | SPON2 | 1.55E-13 | 1.033397405 | 0.278 | 0.188 | 3.05E-09 | Lep-L |
| allcell | DOK2 | 4.74E-43 | 1.026616442 | 0.624 | 0.47 | 9.33E-39 | Lep-L |
| allcell | MAPKAPK2 | 8.49E-20 | 1.001249762 | 0.361 | 0.251 | 1.67E-15 | Lep-L |
| allcell | EIF2AK3 | 9.76E-13 | 0.99739696 | 0.279 | 0.194 | 1.92E-08 | Lep-L |
| allcell | VPS37B | 2.93E-58 | 0.986281545 | 0.748 | 0.584 | 5.77E-54 | Lep-L |
| allcell | LRMP | 2.31E-16 | 0.976815875 | 0.37 | 0.275 | 4.55E-12 | Lep-L |
| allcell | FKBP4 | 1.57E-33 | 0.974649761 | 0.475 | 0.308 | 3.09E-29 | Lep-L |
| allcell | FLNA | 3.61E-38 | 0.960931 | 0.579 | 0.43 | 7.11E-34 | Lep-L |
| allcell | BICDL1 | 2.08E-09 | 0.957790633 | 0.273 | 0.205 | 4.09E-05 | Lep-L |
| allcell | PRR7 | 9.82E-16 | 0.95638134 | 0.337 | 0.241 | 1.93E-11 | Lep-L |
| allcell | NR4A2 | 8.79E-32 | 0.951076425 | 0.798 | 0.726 | 1.73E-27 | Lep-L |
| allcell | RORA | 8.43E-33 | 0.947099739 | 0.606 | 0.473 | 1.66E-28 | Lep-L |
| allcell | EZR | 6.11E-52 | 0.944484201 | 0.812 | 0.694 | 1.20E-47 | Lep-L |
| allcell | TUBB | 1.86E-22 | 0.941526413 | 0.471 | 0.358 | 3.67E-18 | Lep-L |
| allcell | GNA13 | 8.91E-22 | 0.934948386 | 0.395 | 0.278 | 1.75E-17 | Lep-L |
| allcell | SMCHD1 | 1.12E-28 | 0.926414839 | 0.682 | 0.62 | 2.21E-24 | Lep-L |
| allcell | SLBP | 3.75E-17 | 0.914440293 | 0.33 | 0.226 | 7.37E-13 | Lep-L |
| allcell | TGIF1 | 1.95E-09 | 0.911867484 | 0.252 | 0.185 | 3.84E-05 | Lep-L |
| allcell | IL21R | 1.90E-10 | 0.908428074 | 0.311 | 0.243 | 3.74E-06 | Lep-L |
| allcell | ZBTB16 | 1.99E-14 | 0.896279738 | 0.385 | 0.291 | 3.92E-10 | Lep-L |
| allcell | TAGAP | 1.97E-27 | 0.894474003 | 0.657 | 0.543 | 3.88E-23 | Lep-L |
| allcell | SAT1 | 2.52E-33 | 0.885719974 | 0.801 | 0.715 | 4.97E-29 | Lep-L |
| allcell | SMAD3 | 2.34E-07 | 0.884604103 | 0.285 | 0.233 | 0.004606118 | Lep-L |
| allcell | FGFR1OP2 | 6.36E-23 | 0.88030961 | 0.539 | 0.44 | 1.25E-18 | Lep-L |

| | | | | | | | |
|---|---|---|---|---|---|---|---|
| allcell | DIP2A | 4.94E-09 | 0.878519555 | 0.296 | 0.233 | 9.72E-05 | Lep-L |
| allcell | MAN1A1 | 1.07E-15 | 0.864203094 | 0.36 | 0.259 | 2.10E-11 | Lep-L |
| allcell | NFIL3 | 5.68E-10 | 0.862052155 | 0.289 | 0.215 | 1.12E-05 | Lep-L |
| allcell | CLEC2B | 6.38E-51 | 0.857553432 | 0.836 | 0.774 | 1.26E-46 | Lep-L |
| allcell | GLUL | 2.60E-08 | 0.850063868 | 0.26 | 0.198 | 0.000511434 | Lep-L |
| allcell | FMNL1 | 1.49E-19 | 0.849650735 | 0.44 | 0.337 | 2.94E-15 | Lep-L |
| allcell | CD55 | 4.28E-16 | 0.849189026 | 0.596 | 0.531 | 8.43E-12 | Lep-L |
| allcell | HAVCR2 | 4.15E-09 | 0.844812323 | 0.289 | 0.225 | 8.17E-05 | Lep-L |
| allcell | MVD | 1.61E-10 | 0.83392176 | 0.253 | 0.178 | 3.18E-06 | Lep-L |
| allcell | PIM1 | 1.89E-11 | 0.826916617 | 0.304 | 0.224 | 3.71E-07 | Lep-L |
| allcell | MTCH1 | 8.16E-13 | 0.822569545 | 0.357 | 0.277 | 1.61E-08 | Lep-L |
| allcell | PTPRJ | 3.62E-07 | 0.81103097 | 0.37 | 0.328 | 0.00713466 | Lep-L |
| allcell | ATP2A2 | 8.44E-10 | 0.810722319 | 0.256 | 0.185 | 1.66E-05 | Lep-L |
| allcell | STARD7 | 1.34E-12 | 0.809605911 | 0.314 | 0.227 | 2.64E-08 | Lep-L |
| allcell | RNF149 | 4.79E-18 | 0.805418739 | 0.417 | 0.311 | 9.44E-14 | Lep-L |
| allcell | GABARAPL1 | 6.34E-19 | 0.801109833 | 0.513 | 0.418 | 1.25E-14 | Lep-L |
| allcell | NPC1 | 5.20E-10 | 0.799896375 | 0.255 | 0.181 | 1.02E-05 | Lep-L |
| allcell | GNG2 | 5.60E-39 | 0.799766227 | 0.685 | 0.512 | 1.10E-34 | Lep-L |
| allcell | RAB9A | 5.41E-09 | 0.797421347 | 0.274 | 0.208 | 0.000106511 | Lep-L |
| allcell | HSPA8 | 1.27E-46 | 0.797249668 | 0.952 | 0.92 | 2.50E-42 | Lep-L |
| allcell | CNOT6L | 1.11E-23 | 0.792863296 | 0.709 | 0.689 | 2.19E-19 | Lep-L |
| allcell | CEMIP2 | 1.36E-36 | 0.781390015 | 0.748 | 0.61 | 2.67E-32 | Lep-L |
| allcell | CYTIP | 2.44E-21 | 0.771243584 | 0.629 | 0.568 | 4.81E-17 | Lep-L |
| allcell | BRD2 | 1.51E-16 | 0.767981143 | 0.694 | 0.643 | 2.97E-12 | Lep-L |
| allcell | OGT | 1.83E-07 | 0.767926218 | 0.343 | 0.295 | 0.003596765 | Lep-L |
| allcell | HNRNPL | 1.25E-09 | 0.763760766 | 0.318 | 0.255 | 2.47E-05 | Lep-L |
| allcell | NUDT4 | 1.48E-06 | 0.763238612 | 0.271 | 0.221 | 0.029089755 | Lep-L |
| allcell | HIBCH | 8.92E-10 | 0.762188062 | 0.357 | 0.287 | 1.76E-05 | Lep-L |
| allcell | FTL | 2.01E-80 | 0.760018437 | 0.974 | 0.975 | 3.96E-76 | Lep-L |
| allcell | OAZ1 | 2.23E-42 | 0.744679311 | 0.851 | 0.823 | 4.38E-38 | Lep-L |
| allcell | GNB2 | 2.80E-10 | 0.73786403 | 0.266 | 0.192 | 5.50E-06 | Lep-L |
| allcell | RHOF | 6.61E-10 | 0.737400929 | 0.357 | 0.289 | 1.30E-05 | Lep-L |
| allcell | UBE2F | 1.12E-09 | 0.735740439 | 0.351 | 0.285 | 2.20E-05 | Lep-L |
| allcell | RASA3 | 1.34E-10 | 0.7343968 | 0.363 | 0.284 | 2.63E-06 | Lep-L |
| allcell | STIP1 | 2.22E-13 | 0.731783355 | 0.396 | 0.315 | 4.37E-09 | Lep-L |
| allcell | KPNA2 | 7.33E-18 | 0.728425713 | 0.411 | 0.298 | 1.44E-13 | Lep-L |
| allcell | YES1 | 2.35E-13 | 0.726888047 | 0.464 | 0.383 | 4.63E-09 | Lep-L |
| allcell | CDKN1B | 1.78E-10 | 0.726213381 | 0.369 | 0.296 | 3.50E-06 | Lep-L |
| allcell | PRKCE | 9.19E-07 | 0.725293294 | 0.287 | 0.231 | 0.018091932 | Lep-L |
| allcell | PDE4D | 2.27E-14 | 0.724093692 | 0.438 | 0.345 | 4.46E-10 | Lep-L |
| allcell | YY1 | 2.71E-14 | 0.724067929 | 0.431 | 0.357 | 5.33E-10 | Lep-L |
| allcell | NFATC2 | 1.49E-09 | 0.723430311 | 0.403 | 0.344 | 2.93E-05 | Lep-L |
| allcell | IFITM1 | 5.09E-17 | 0.722912684 | 0.623 | 0.565 | 1.00E-12 | Lep-L |
| allcell | BAZ1A | 4.35E-26 | 0.711830652 | 0.612 | 0.477 | 8.56E-22 | Lep-L |
| allcell | SPEN | 9.11E-08 | 0.708296163 | 0.292 | 0.231 | 0.001793889 | Lep-L |
| allcell | MAP2K3 | 1.25E-12 | 0.707872245 | 0.43 | 0.346 | 2.46E-08 | Lep-L |
| allcell | SERPINB9 | 1.95E-08 | 0.705816274 | 0.422 | 0.373 | 0.000382879 | Lep-L |
| allcell | PNRC1 | 2.14E-27 | 0.704432296 | 0.89 | 0.855 | 4.22E-23 | Lep-L |
| allcell | GBP5 | 5.47E-09 | 0.703951207 | 0.357 | 0.289 | 0.0001077 | Lep-L |
| allcell | TBX21 | 4.91E-09 | 0.701839697 | 0.302 | 0.232 | 9.66E-05 | Lep-L |
| allcell | NFAT5 | 9.88E-12 | 0.7008047 | 0.519 | 0.462 | 1.94E-07 | Lep-L |
| allcell | KDM6B | 1.98E-13 | 0.698708156 | 0.469 | 0.395 | 3.90E-09 | Lep-L |
| allcell | HSP90AB1 | 2.92E-61 | 0.698623749 | 0.947 | 0.923 | 5.74E-57 | Lep-L |
| allcell | STX11 | 1.17E-15 | 0.694734137 | 0.427 | 0.326 | 2.31E-11 | Lep-L |
| allcell | EIF4H | 9.48E-10 | 0.6933569 | 0.339 | 0.27 | 1.87E-05 | Lep-L |
| allcell | CLIC3 | 1.83E-10 | 0.692937492 | 0.414 | 0.338 | 3.61E-06 | Lep-L |

| | | | | | | | |
|---|---|---|---|---|---|---|---|
| allcell | ZEB2 | 2.28E-13 | 0.689488901 | 0.585 | 0.523 | 4.50E-09 | Lep-L |
| allcell | TGFB1 | 3.33E-25 | 0.687159307 | 0.784 | 0.729 | 6.55E-21 | Lep-L |
| allcell | RAPGEF1 | 3.65E-07 | 0.685565933 | 0.293 | 0.236 | 0.007178864 | Lep-L |
| allcell | EEF2 | 6.68E-23 | 0.685544054 | 0.653 | 0.59 | 1.31E-18 | Lep-L |
| allcell | SIAH2 | 5.34E-08 | 0.681221939 | 0.286 | 0.224 | 0.001051848 | Lep-L |
| allcell | ZHX2 | 1.32E-07 | 0.680420063 | 0.384 | 0.334 | 0.002593467 | Lep-L |
| allcell | CMC2 | 1.16E-07 | 0.679342677 | 0.309 | 0.251 | 0.002292552 | Lep-L |
| allcell | CSNK1D | 4.49E-11 | 0.675663991 | 0.349 | 0.275 | 8.83E-07 | Lep-L |
| allcell | NEAT1 | 4.57E-10 | 0.674901882 | 0.657 | 0.648 | 8.99E-06 | Lep-L |
| allcell | PTGER4 | 2.56E-09 | 0.674709189 | 0.444 | 0.383 | 5.05E-05 | Lep-L |
| allcell | PPP1R18 | 1.68E-09 | 0.674010266 | 0.37 | 0.3 | 3.31E-05 | Lep-L |
| allcell | HELZ | 2.01E-08 | 0.666265488 | 0.376 | 0.326 | 0.000396458 | Lep-L |
| allcell | PPP2R5C | 3.32E-09 | 0.664477962 | 0.545 | 0.521 | 6.54E-05 | Lep-L |
| allcell | HBP1 | 5.92E-15 | 0.663016382 | 0.478 | 0.392 | 1.17E-10 | Lep-L |
| allcell | FTH1 | 6.57E-16 | 0.654990349 | 0.965 | 0.968 | 1.29E-11 | Lep-L |
| allcell | IDS | 9.88E-09 | 0.65112959 | 0.404 | 0.346 | 0.000194552 | Lep-L |
| allcell | CYTOR | 1.00E-08 | 0.648335097 | 0.422 | 0.363 | 0.000197306 | Lep-L |
| allcell | CHD3 | 5.16E-08 | 0.648215529 | 0.305 | 0.246 | 0.001015552 | Lep-L |
| allcell | RHOH | 6.50E-12 | 0.644069823 | 0.533 | 0.467 | 1.28E-07 | Lep-L |
| allcell | FOSL2 | 2.15E-16 | 0.641094788 | 0.543 | 0.461 | 4.24E-12 | Lep-L |
| allcell | MYH9 | 8.63E-12 | 0.640069276 | 0.45 | 0.388 | 1.70E-07 | Lep-L |
| allcell | ATP5F1B | 2.50E-09 | 0.638504896 | 0.469 | 0.427 | 4.92E-05 | Lep-L |
| allcell | TPM4 | 3.58E-15 | 0.634149959 | 0.544 | 0.472 | 7.04E-11 | Lep-L |
| allcell | MKNK2 | 7.28E-10 | 0.626666598 | 0.411 | 0.336 | 1.43E-05 | Lep-L |
| allcell | IGF2R | 4.59E-08 | 0.620944055 | 0.379 | 0.323 | 0.000904287 | Lep-L |
| allcell | RUNX3 | 1.87E-28 | 0.617649985 | 0.773 | 0.726 | 3.67E-24 | Lep-L |
| allcell | PER1 | 1.61E-07 | 0.613141501 | 0.336 | 0.281 | 0.003173254 | Lep-L |
| allcell | LYST | 1.40E-09 | 0.610082847 | 0.559 | 0.508 | 2.76E-05 | Lep-L |
| allcell | NAA50 | 1.05E-06 | 0.603686942 | 0.324 | 0.272 | 0.020607622 | Lep-L |
| allcell | PLIN2 | 1.18E-14 | 0.593855708 | 0.368 | 0.26 | 2.31E-10 | Lep-L |
| allcell | LEPROTL1 | 2.22E-07 | 0.591303254 | 0.539 | 0.535 | 0.00437144 | Lep-L |
| allcell | CNOT2 | 7.04E-07 | 0.590800922 | 0.599 | 0.595 | 0.013852737 | Lep-L |
| allcell | CCNH | 6.89E-15 | 0.587377159 | 0.627 | 0.571 | 1.36E-10 | Lep-L |
| allcell | H2AFJ | 1.06E-07 | 0.583334182 | 0.453 | 0.411 | 0.002091212 | Lep-L |
| allcell | LITAF | 2.44E-25 | 0.581435438 | 0.737 | 0.653 | 4.81E-21 | Lep-L |
| allcell | PRKCH | 1.15E-09 | 0.581293224 | 0.679 | 0.658 | 2.26E-05 | Lep-L |
| allcell | UBE2A | 9.65E-10 | 0.574016978 | 0.43 | 0.364 | 1.90E-05 | Lep-L |
| allcell | MOB4 | 9.70E-11 | 0.573014327 | 0.5 | 0.45 | 1.91E-06 | Lep-L |
| allcell | PRF1 | 9.11E-12 | 0.56987502 | 0.557 | 0.491 | 1.79E-07 | Lep-L |
| allcell | PRNP | 2.87E-07 | 0.569678717 | 0.327 | 0.265 | 0.005658457 | Lep-L |
| allcell | HLA-DRB1 | 4.91E-17 | 0.568487251 | 0.588 | 0.486 | 9.66E-13 | Lep-L |
| allcell | IFI16 | 2.86E-07 | 0.565831169 | 0.49 | 0.465 | 0.005638183 | Lep-L |
| allcell | ATP6V0C | 7.45E-12 | 0.563291815 | 0.59 | 0.543 | 1.47E-07 | Lep-L |
| allcell | CHD2 | 4.69E-13 | 0.561835702 | 0.683 | 0.667 | 9.23E-09 | Lep-L |
| allcell | CCL3L1 | 6.83E-09 | 0.560587577 | 0.31 | 0.229 | 0.000134414 | Lep-L |
| allcell | HSP90B1 | 1.45E-14 | 0.557568951 | 0.634 | 0.579 | 2.86E-10 | Lep-L |
| allcell | RASA2 | 2.90E-11 | 0.551458816 | 0.539 | 0.477 | 5.70E-07 | Lep-L |
| allcell | CBLB | 1.55E-06 | 0.546341934 | 0.594 | 0.577 | 0.030436787 | Lep-L |
| allcell | MAPRE2 | 2.21E-07 | 0.546302356 | 0.518 | 0.477 | 0.004349794 | Lep-L |
| allcell | MCL1 | 1.08E-19 | 0.545534662 | 0.688 | 0.61 | 2.13E-15 | Lep-L |
| allcell | HECA | 5.20E-08 | 0.544630263 | 0.448 | 0.391 | 0.001024062 | Lep-L |
| allcell | LAPTM5 | 8.54E-07 | 0.538819203 | 0.485 | 0.458 | 0.016814865 | Lep-L |
| allcell | PTGES3 | 3.73E-20 | 0.536936908 | 0.708 | 0.665 | 7.34E-16 | Lep-L |
| allcell | ZFP36L2 | 1.75E-07 | 0.536648355 | 0.808 | 0.827 | 0.003443141 | Lep-L |
| allcell | ANP32A | 1.90E-06 | 0.535535768 | 0.346 | 0.298 | 0.03740394 | Lep-L |
| allcell | JUND | 1.36E-49 | 0.535181085 | 0.942 | 0.94 | 2.67E-45 | Lep-L |

| | | | | | | | |
|---|---|---|---|---|---|---|---|
| allcell | ARL6IP1 | 2.66E-12 | 0.531470444 | 0.551 | 0.488 | 5.24E-08 | Lep-L |
| allcell | ISG20 | 1.37E-12 | 0.531094831 | 0.556 | 0.502 | 2.70E-08 | Lep-L |
| allcell | HSP90AA1 | 5.04E-40 | 0.525056237 | 0.976 | 0.971 | 9.93E-36 | Lep-L |
| allcell | ZC3HAV1 | 5.42E-12 | 0.522890224 | 0.63 | 0.579 | 1.07E-07 | Lep-L |
| allcell | ELOVL5 | 4.79E-11 | 0.522205727 | 0.581 | 0.512 | 9.43E-07 | Lep-L |
| allcell | NINJ1 | 1.68E-06 | 0.513852864 | 0.287 | 0.23 | 0.033025013 | Lep-L |
| allcell | SYNE2 | 5.85E-13 | 0.5137782 | 0.512 | 0.415 | 1.15E-08 | Lep-L |
| allcell | HSPH1 | 1.41E-25 | 0.512960366 | 0.796 | 0.713 | 2.77E-21 | Lep-L |
| allcell | SLC3A2 | 8.15E-10 | 0.512714309 | 0.497 | 0.435 | 1.60E-05 | Lep-L |
| allcell | STK4 | 4.21E-17 | 0.510997377 | 0.71 | 0.676 | 8.29E-13 | Lep-L |
| allcell | RAB8B | 2.68E-21 | 0.50885488 | 0.767 | 0.725 | 5.28E-17 | Lep-L |
| allcell | YWHAZ | 3.99E-19 | 0.505911873 | 0.777 | 0.754 | 7.85E-15 | Lep-L |
| allcell | SLC7A5 | 3.22E-07 | 0.502322868 | 0.474 | 0.434 | 0.006329126 | Lep-L |
| allcell | RCOR1 | 2.89E-07 | 0.501795995 | 0.334 | 0.274 | 0.005684366 | Lep-L |
| allcell | XCL1 | 3.16E-36 | 1.539156297 | 0.71 | 0.569 | 6.22E-32 | Lep-T |
| allcell | HIST1H1E | 4.84E-51 | 1.32676202 | 0.377 | 0.167 | 9.53E-47 | Lep-T |
| allcell | IFITM11 | 1.49E-14 | 1.291661776 | 0.642 | 0.558 | 2.94E-10 | Lep-T |
| allcell | CST3 | 7.59E-32 | 1.202022172 | 0.27 | 0.123 | 1.49E-27 | Lep-T |
| allcell | HIST1H1C | 1.15E-34 | 1.163738103 | 0.312 | 0.148 | 2.26E-30 | Lep-T |
| allcell | CD74 | 1.53E-21 | 1.152766201 | 0.776 | 0.666 | 3.01E-17 | Lep-T |
| allcell | RBFOX21 | 2.71E-35 | 1.13954546 | 0.446 | 0.263 | 5.33E-31 | Lep-T |
| allcell | NIBAN1 | 1.73E-14 | 1.059449917 | 0.365 | 0.257 | 3.41E-10 | Lep-T |
| allcell | MT-CO31 | 2.72E-43 | 0.956945939 | 0.833 | 0.87 | 5.35E-39 | Lep-T |
| allcell | XCL2 | 3.43E-18 | 0.943854996 | 0.756 | 0.692 | 6.75E-14 | Lep-T |
| allcell | BCL3 | 7.79E-19 | 0.925949538 | 0.278 | 0.161 | 1.53E-14 | Lep-T |
| allcell | CCL4L21 | 1.51E-17 | 0.906562344 | 0.444 | 0.309 | 2.98E-13 | Lep-T |
| allcell | IFITM3 | 1.62E-21 | 0.887091497 | 0.484 | 0.329 | 3.19E-17 | Lep-T |
| allcell | EMD1 | 2.88E-24 | 0.867213022 | 0.306 | 0.165 | 5.66E-20 | Lep-T |
| allcell | DAZAP2 | 2.62E-27 | 0.84594138 | 0.405 | 0.234 | 5.16E-23 | Lep-T |
| allcell | CCL4 | 1.09E-11 | 0.826212821 | 0.692 | 0.638 | 2.14E-07 | Lep-T |
| allcell | IFITM2 | 1.14E-27 | 0.806326163 | 0.873 | 0.785 | 2.24E-23 | Lep-T |
| allcell | IL12RB2 | 4.40E-15 | 0.803147712 | 0.45 | 0.339 | 8.67E-11 | Lep-T |
| allcell | TSC22D3 | 1.20E-20 | 0.7978912 | 0.708 | 0.592 | 2.37E-16 | Lep-T |
| allcell | CRTAM | 1.54E-21 | 0.76461541 | 0.424 | 0.281 | 3.03E-17 | Lep-T |
| allcell | PRKX | 3.30E-15 | 0.757599532 | 0.457 | 0.346 | 6.50E-11 | Lep-T |
| allcell | XAF1 | 2.26E-19 | 0.744793178 | 0.274 | 0.149 | 4.45E-15 | Lep-T |
| allcell | BHLHE40 | 1.21E-33 | 0.735304326 | 0.651 | 0.468 | 2.38E-29 | Lep-T |
| allcell | MT-ND11 | 9.26E-40 | 0.725445856 | 0.8 | 0.733 | 1.82E-35 | Lep-T |
| allcell | RASGRP1 | 7.63E-17 | 0.709103355 | 0.298 | 0.177 | 1.50E-12 | Lep-T |
| allcell | AREG | 2.24E-11 | 0.701084778 | 0.65 | 0.595 | 4.41E-07 | Lep-T |
| allcell | PPP1CA | 7.11E-12 | 0.668700969 | 0.304 | 0.201 | 1.40E-07 | Lep-T |
| allcell | RGS1 | 4.04E-08 | 0.666970305 | 0.469 | 0.414 | 0.000794779 | Lep-T |
| allcell | USP36 | 2.57E-17 | 0.66320401 | 0.359 | 0.225 | 5.07E-13 | Lep-T |
| allcell | NFKBIZ | 3.29E-19 | 0.651697393 | 0.521 | 0.385 | 6.48E-15 | Lep-T |
| allcell | H1FX | 2.23E-12 | 0.641969329 | 0.308 | 0.209 | 4.40E-08 | Lep-T |
| allcell | ASAP1 | 2.08E-11 | 0.639502494 | 0.406 | 0.304 | 4.09E-07 | Lep-T |
| allcell | KLF6 | 4.25E-26 | 0.632464506 | 0.744 | 0.622 | 8.37E-22 | Lep-T |
| allcell | LPIN1 | 3.41E-13 | 0.627630776 | 0.298 | 0.193 | 6.71E-09 | Lep-T |
| allcell | STK39 | 1.62E-15 | 0.613918354 | 0.34 | 0.217 | 3.20E-11 | Lep-T |
| allcell | DUSP10 | 7.20E-18 | 0.609878601 | 0.338 | 0.209 | 1.42E-13 | Lep-T |
| allcell | EML4 | 2.58E-19 | 0.60876443 | 0.651 | 0.526 | 5.08E-15 | Lep-T |
| allcell | FBXO34 | 6.56E-12 | 0.608749028 | 0.501 | 0.396 | 1.29E-07 | Lep-T |
| allcell | MT-ATP61 | 1.41E-21 | 0.608028002 | 0.8 | 0.9 | 2.77E-17 | Lep-T |
| allcell | ID2 | 3.19E-12 | 0.607256767 | 0.625 | 0.556 | 6.27E-08 | Lep-T |
| allcell | RGS2 | 7.94E-12 | 0.605627539 | 0.447 | 0.334 | 1.56E-07 | Lep-T |
| allcell | CSRNP11 | 3.21E-13 | 0.603408585 | 0.477 | 0.356 | 6.32E-09 | Lep-T |

| | | | | | | | |
|---|---|---|---|---|---|---|---|
| allcell | VAV3 | 2.77E-15 | 0.595146498 | 0.498 | 0.375 | 5.45E-11 | Lep-T |
| allcell | KIR2DL4 | 1.15E-11 | 0.594784268 | 0.27 | 0.175 | 2.26E-07 | Lep-T |
| allcell | ICAM1 | 4.76E-11 | 0.592366256 | 0.308 | 0.21 | 9.36E-07 | Lep-T |
| allcell | GTF2F2 | 8.89E-12 | 0.588772827 | 0.251 | 0.156 | 1.75E-07 | Lep-T |
| allcell | ZFP36L11 | 2.15E-34 | 0.583787374 | 0.744 | 0.545 | 4.23E-30 | Lep-T |
| allcell | EPG5 | 2.96E-13 | 0.577241964 | 0.278 | 0.173 | 5.83E-09 | Lep-T |
| allcell | SLC2A31 | 1.20E-22 | 0.576875396 | 0.665 | 0.516 | 2.36E-18 | Lep-T |
| allcell | CXCR4 | 3.52E-17 | 0.571141083 | 0.735 | 0.667 | 6.93E-13 | Lep-T |
| allcell | PFKFB3 | 1.92E-12 | 0.557036576 | 0.314 | 0.205 | 3.77E-08 | Lep-T |
| allcell | MT-CYB1 | 1.50E-31 | 0.556609497 | 0.819 | 0.805 | 2.95E-27 | Lep-T |
| allcell | SLA | 6.48E-13 | 0.551316074 | 0.477 | 0.363 | 1.28E-08 | Lep-T |
| allcell | LCP2 | 8.87E-15 | 0.547158518 | 0.368 | 0.242 | 1.75E-10 | Lep-T |
| allcell | GAS5 | 2.22E-15 | 0.541711621 | 0.46 | 0.325 | 4.38E-11 | Lep-T |
| allcell | ADGRE5 | 1.29E-16 | 0.541558054 | 0.71 | 0.605 | 2.55E-12 | Lep-T |
| allcell | CD3E | 3.34E-09 | 0.530725699 | 0.346 | 0.253 | 6.57E-05 | Lep-T |
| allcell | MAFF | 9.52E-17 | 0.526671331 | 0.45 | 0.31 | 1.87E-12 | Lep-T |
| allcell | PNRC11 | 2.15E-28 | 0.524066484 | 0.922 | 0.845 | 4.24E-24 | Lep-T |
| allcell | MT-ND5 | 3.34E-24 | 0.521171561 | 0.739 | 0.637 | 6.57E-20 | Lep-T |
| allcell | SIPA1L1 | 1.81E-09 | 0.519000427 | 0.455 | 0.358 | 3.56E-05 | Lep-T |
| allcell | PHF20 | 4.71E-13 | 0.518986432 | 0.518 | 0.409 | 9.27E-09 | Lep-T |
| allcell | FOXO3 | 1.55E-08 | 0.518527434 | 0.315 | 0.226 | 0.000305381 | Lep-T |
| allcell | IGFBP7 | 1.20E-10 | 0.516240868 | 0.267 | 0.172 | 2.36E-06 | Lep-T |
| allcell | ABHD2 | 2.72E-11 | 0.50841824 | 0.399 | 0.291 | 5.36E-07 | Lep-T |
| allcell | RERE | 5.91E-09 | 0.502957866 | 0.409 | 0.319 | 0.000116293 | Lep-T |
| allcell | FYB1 | 2.04E-16 | 0.501365904 | 0.507 | 0.373 | 4.02E-12 | Lep-T |
| allcell | GZMA | 3.61E-30 | 0.982599116 | 0.713 | 0.633 | 7.10E-26 | Mari |
| allcell | TMIGD2 | 8.29E-15 | 0.981208511 | 0.255 | 0.166 | 1.63E-10 | Mari |
| allcell | TIMP1 | 1.60E-15 | 0.934240146 | 0.279 | 0.184 | 3.14E-11 | Mari |
| allcell | LYAR | 1.68E-14 | 0.924586739 | 0.27 | 0.177 | 3.31E-10 | Mari |
| allcell | NDUFC2 | 7.41E-29 | 0.919933522 | 0.392 | 0.247 | 1.46E-24 | Mari |
| allcell | GNLY | 3.18E-73 | 0.871974881 | 0.918 | 0.809 | 6.26E-69 | Mari |
| allcell | TRDC1 | 2.17E-31 | 0.867458421 | 0.655 | 0.543 | 4.26E-27 | Mari |
| allcell | DCXR | 1.71E-15 | 0.836916505 | 0.292 | 0.194 | 3.37E-11 | Mari |
| allcell | ATP5MC1 | 2.28E-13 | 0.805167519 | 0.255 | 0.171 | 4.49E-09 | Mari |
| allcell | MICOS13 | 6.70E-15 | 0.804169933 | 0.278 | 0.182 | 1.32E-10 | Mari |
| allcell | ATP5MF | 8.99E-31 | 0.802125964 | 0.449 | 0.293 | 1.77E-26 | Mari |
| allcell | LAMTOR1 | 2.35E-14 | 0.799162018 | 0.275 | 0.186 | 4.63E-10 | Mari |
| allcell | ATP5ME | 5.21E-30 | 0.784491541 | 0.481 | 0.324 | 1.03E-25 | Mari |
| allcell | KLRF1 | 6.07E-33 | 0.777104658 | 0.654 | 0.508 | 1.19E-28 | Mari |
| allcell | BST2 | 8.22E-30 | 0.770017152 | 0.527 | 0.393 | 1.62E-25 | Mari |
| allcell | PET100 | 6.67E-19 | 0.76895604 | 0.348 | 0.233 | 1.31E-14 | Mari |
| allcell | GIMAP4 | 5.89E-13 | 0.768104882 | 0.302 | 0.213 | 1.16E-08 | Mari |
| allcell | OSTC | 6.94E-19 | 0.76675462 | 0.367 | 0.25 | 1.37E-14 | Mari |
| allcell | TMEM14B | 1.07E-16 | 0.763585928 | 0.324 | 0.22 | 2.10E-12 | Mari |
| allcell | CD52 | 7.69E-25 | 0.759762513 | 0.516 | 0.374 | 1.51E-20 | Mari |
| allcell | ATP5MPL | 8.08E-18 | 0.750737977 | 0.332 | 0.225 | 1.59E-13 | Mari |
| allcell | HSPB11 | 2.88E-11 | 0.749942746 | 0.268 | 0.19 | 5.67E-07 | Mari |
| allcell | TSTD1 | 1.82E-17 | 0.747489823 | 0.334 | 0.227 | 3.57E-13 | Mari |
| allcell | PRDX2 | 1.26E-15 | 0.747405261 | 0.275 | 0.179 | 2.47E-11 | Mari |
| allcell | BLOC1S1 | 7.73E-18 | 0.737293907 | 0.373 | 0.264 | 1.52E-13 | Mari |
| allcell | HOPX | 3.21E-32 | 0.736940153 | 0.696 | 0.573 | 6.31E-28 | Mari |
| allcell | PSMB1 | 1.23E-34 | 0.731164594 | 0.569 | 0.406 | 2.42E-30 | Mari |
| allcell | DGUOK | 5.42E-18 | 0.729445206 | 0.381 | 0.269 | 1.07E-13 | Mari |
| allcell | MRPL33 | 1.34E-19 | 0.729112753 | 0.349 | 0.231 | 2.65E-15 | Mari |
| allcell | SEPTIN1 | 6.79E-17 | 0.728209327 | 0.341 | 0.233 | 1.34E-12 | Mari |
| allcell | REX1BD | 3.75E-15 | 0.725117973 | 0.364 | 0.267 | 7.38E-11 | Mari |

| | | | | | | | |
|---|---|---|---|---|---|---|---|
| allcell | TMCO1 | 6.86E-14 | 0.714978015 | 0.309 | 0.214 | 1.35E-09 | Mari |
| allcell | AP2S1 | 1.44E-12 | 0.712027559 | 0.283 | 0.197 | 2.84E-08 | Mari |
| allcell | TRAPPC1 | 2.30E-12 | 0.711361611 | 0.304 | 0.218 | 4.53E-08 | Mari |
| allcell | ATP5PF | 9.94E-24 | 0.709648055 | 0.47 | 0.343 | 1.96E-19 | Mari |
| allcell | SEC11C | 3.74E-15 | 0.709496697 | 0.297 | 0.195 | 7.37E-11 | Mari |
| allcell | TOMM5 | 3.45E-15 | 0.707024446 | 0.323 | 0.223 | 6.80E-11 | Mari |
| allcell | TMBIM4 | 9.35E-18 | 0.70628376 | 0.381 | 0.269 | 1.84E-13 | Mari |
| allcell | COPS9 | 3.39E-15 | 0.699852315 | 0.355 | 0.255 | 6.68E-11 | Mari |
| allcell | IRF9 | 1.62E-11 | 0.696723484 | 0.284 | 0.202 | 3.18E-07 | Mari |
| allcell | COX6B1 | 1.28E-32 | 0.696421778 | 0.596 | 0.445 | 2.51E-28 | Mari |
| allcell | RHOB | 2.17E-08 | 0.69395838 | 0.313 | 0.245 | 0.00042709 | Mari |
| allcell | COX7A2L | 6.09E-15 | 0.692560387 | 0.358 | 0.261 | 1.20E-10 | Mari |
| allcell | SEC61G | 9.40E-27 | 0.686624472 | 0.517 | 0.368 | 1.85E-22 | Mari |
| allcell | COX7A2 | 4.72E-35 | 0.686366759 | 0.636 | 0.476 | 9.28E-31 | Mari |
| allcell | ERH | 6.79E-21 | 0.683631465 | 0.448 | 0.324 | 1.34E-16 | Mari |
| allcell | CD2 | 6.80E-26 | 0.680338878 | 0.607 | 0.472 | 1.34E-21 | Mari |
| allcell | C1D | 9.33E-10 | 0.679216784 | 0.25 | 0.178 | 1.84E-05 | Mari |
| allcell | CNPY2 | 6.02E-10 | 0.676755843 | 0.25 | 0.177 | 1.19E-05 | Mari |
| allcell | MRPS34 | 8.39E-11 | 0.676673884 | 0.261 | 0.184 | 1.65E-06 | Mari |
| allcell | UQCR10 | 2.97E-20 | 0.676447289 | 0.426 | 0.301 | 5.86E-16 | Mari |
| allcell | VPS28 | 2.00E-15 | 0.675754853 | 0.389 | 0.291 | 3.93E-11 | Mari |
| allcell | RBIS | 1.72E-13 | 0.667589024 | 0.377 | 0.289 | 3.39E-09 | Mari |
| allcell | SNRPE | 1.95E-14 | 0.665821868 | 0.316 | 0.218 | 3.85E-10 | Mari |
| allcell | NDUFAB1 | 7.25E-19 | 0.664955965 | 0.334 | 0.214 | 1.43E-14 | Mari |
| allcell | RANBP1 | 2.66E-13 | 0.663760826 | 0.291 | 0.199 | 5.24E-09 | Mari |
| allcell | PSMB9 | 2.69E-26 | 0.656427756 | 0.624 | 0.498 | 5.29E-22 | Mari |
| allcell | UQCR11 | 9.75E-23 | 0.653780536 | 0.516 | 0.388 | 1.92E-18 | Mari |
| allcell | TOMM6 | 8.28E-16 | 0.648674951 | 0.381 | 0.275 | 1.63E-11 | Mari |
| allcell | RER1 | 1.53E-09 | 0.647962913 | 0.277 | 0.204 | 3.02E-05 | Mari |
| allcell | SNHG9 | 2.03E-11 | 0.647229345 | 0.268 | 0.183 | 4.00E-07 | Mari |
| allcell | NDUFB2 | 1.22E-24 | 0.642923324 | 0.553 | 0.417 | 2.40E-20 | Mari |
| allcell | TRBC2 | 6.64E-19 | 0.639975148 | 0.5 | 0.383 | 1.31E-14 | Mari |
| allcell | GMFG | 8.13E-24 | 0.637271351 | 0.561 | 0.441 | 1.60E-19 | Mari |
| allcell | HIKESHI | 1.56E-09 | 0.630244774 | 0.275 | 0.201 | 3.08E-05 | Mari |
| allcell | NDUFS6 | 1.27E-12 | 0.629543431 | 0.37 | 0.28 | 2.50E-08 | Mari |
| allcell | NDUFB7 | 9.86E-15 | 0.628983372 | 0.367 | 0.264 | 1.94E-10 | Mari |
| allcell | NDUFA4 | 5.54E-28 | 0.62851906 | 0.641 | 0.516 | 1.09E-23 | Mari |
| allcell | MGST3 | 4.50E-10 | 0.627094759 | 0.266 | 0.189 | 8.86E-06 | Mari |
| allcell | C1QBP | 7.04E-15 | 0.625027602 | 0.336 | 0.233 | 1.39E-10 | Mari |
| allcell | COX5B | 7.24E-24 | 0.62445174 | 0.536 | 0.405 | 1.42E-19 | Mari |
| allcell | ARF5 | 2.87E-12 | 0.61944685 | 0.305 | 0.212 | 5.65E-08 | Mari |
| allcell | SSBP1 | 3.95E-17 | 0.61503826 | 0.414 | 0.299 | 7.78E-13 | Mari |
| allcell | PHPT1 | 1.94E-10 | 0.614125116 | 0.276 | 0.195 | 3.83E-06 | Mari |
| allcell | OCIAD2 | 6.86E-12 | 0.613045037 | 0.342 | 0.256 | 1.35E-07 | Mari |
| allcell | MED10 | 8.38E-14 | 0.611609857 | 0.354 | 0.257 | 1.65E-09 | Mari |
| allcell | PDCD6 | 1.05E-10 | 0.610174039 | 0.312 | 0.233 | 2.07E-06 | Mari |
| allcell | GABARAPL2 | 1.66E-16 | 0.608042585 | 0.414 | 0.297 | 3.27E-12 | Mari |
| allcell | MRPL20 | 2.42E-10 | 0.604477857 | 0.305 | 0.226 | 4.76E-06 | Mari |
| allcell | MRPL54 | 1.20E-11 | 0.603341321 | 0.301 | 0.215 | 2.36E-07 | Mari |
| allcell | MPG | 4.55E-09 | 0.602957598 | 0.276 | 0.205 | 8.95E-05 | Mari |
| allcell | LSM2 | 3.30E-11 | 0.600828381 | 0.334 | 0.251 | 6.49E-07 | Mari |
| allcell | NEDD8 | 2.40E-12 | 0.596256079 | 0.416 | 0.333 | 4.72E-08 | Mari |
| allcell | SRP9 | 2.47E-11 | 0.59325999 | 0.342 | 0.26 | 4.86E-07 | Mari |
| allcell | MGAT4A | 1.76E-09 | 0.590497164 | 0.301 | 0.223 | 3.47E-05 | Mari |
| allcell | ANAPC16 | 2.12E-20 | 0.590103508 | 0.515 | 0.392 | 4.18E-16 | Mari |
| allcell | APRT | 2.05E-18 | 0.588954157 | 0.556 | 0.448 | 4.03E-14 | Mari |

| | | | | | | | |
|---|---|---|---|---|---|---|---|
| allcell | NDUFA3 | 2.27E-10 | 0.587840201 | 0.296 | 0.216 | 4.46E-06 | Mari |
| allcell | NDUFA13 | 9.44E-22 | 0.585904448 | 0.565 | 0.447 | 1.86E-17 | Mari |
| allcell | MYL12A | 1.02E-34 | 0.585609941 | 0.83 | 0.736 | 2.01E-30 | Mari |
| allcell | MICOS10 | 2.12E-15 | 0.58535527 | 0.404 | 0.303 | 4.18E-11 | Mari |
| allcell | SNRPB2 | 1.48E-11 | 0.583453161 | 0.327 | 0.24 | 2.92E-07 | Mari |
| allcell | SELENOH | 8.37E-15 | 0.582162869 | 0.454 | 0.353 | 1.65E-10 | Mari |
| allcell | GLRX5 | 5.67E-11 | 0.581456832 | 0.262 | 0.181 | 1.12E-06 | Mari |
| allcell | DCTN3 | 7.18E-13 | 0.577510131 | 0.336 | 0.241 | 1.41E-08 | Mari |
| allcell | TMEM160 | 1.05E-09 | 0.577144135 | 0.271 | 0.195 | 2.07E-05 | Mari |
| allcell | BANF1 | 2.72E-11 | 0.575853758 | 0.347 | 0.265 | 5.36E-07 | Mari |
| allcell | PHB | 2.95E-10 | 0.575811022 | 0.255 | 0.178 | 5.81E-06 | Mari |
| allcell | MRPS21 | 1.13E-10 | 0.574847504 | 0.312 | 0.229 | 2.22E-06 | Mari |
| allcell | FCER1G | 3.93E-18 | 0.573913719 | 0.634 | 0.548 | 7.75E-14 | Mari |
| allcell | DAD1 | 2.95E-13 | 0.57309727 | 0.408 | 0.314 | 5.81E-09 | Mari |
| allcell | TMED9 | 5.91E-08 | 0.570112585 | 0.277 | 0.212 | 0.001162647 | Mari |
| allcell | MED30 | 1.49E-08 | 0.569071646 | 0.259 | 0.191 | 0.000293279 | Mari |
| allcell | COX8A | 6.28E-26 | 0.568552534 | 0.565 | 0.413 | 1.24E-21 | Mari |
| allcell | MRPL51 | 1.79E-11 | 0.568501925 | 0.319 | 0.232 | 3.53E-07 | Mari |
| allcell | ITM2B | 6.45E-42 | 0.568297377 | 0.829 | 0.703 | 1.27E-37 | Mari |
| allcell | NT5C | 2.15E-08 | 0.567381099 | 0.295 | 0.228 | 0.000422537 | Mari |
| allcell | COX6C | 1.13E-23 | 0.564489187 | 0.603 | 0.477 | 2.22E-19 | Mari |
| allcell | NOP10 | 5.35E-15 | 0.563687519 | 0.39 | 0.284 | 1.05E-10 | Mari |
| allcell | SPCS1 | 1.51E-13 | 0.563207941 | 0.394 | 0.298 | 2.97E-09 | Mari |
| allcell | DYNLRB1 | 4.89E-08 | 0.563073209 | 0.294 | 0.23 | 0.0009626 | Mari |
| allcell | FAM89B | 2.13E-10 | 0.560914076 | 0.262 | 0.181 | 4.19E-06 | Mari |
| allcell | SSNA1 | 1.20E-08 | 0.559218914 | 0.276 | 0.206 | 0.000236447 | Mari |
| allcell | SELL | 2.67E-07 | 0.555160007 | 0.262 | 0.199 | 0.005264271 | Mari |
| allcell | ATP5MD | 2.79E-14 | 0.550238246 | 0.431 | 0.333 | 5.50E-10 | Mari |
| allcell | EIF3G | 1.56E-16 | 0.549385552 | 0.68 | 0.594 | 3.08E-12 | Mari |
| allcell | LMAN2 | 9.92E-12 | 0.548480768 | 0.346 | 0.256 | 1.95E-07 | Mari |
| allcell | RBX1 | 8.20E-12 | 0.547362425 | 0.413 | 0.329 | 1.61E-07 | Mari |
| allcell | CKLF | 2.11E-13 | 0.547234971 | 0.492 | 0.39 | 4.15E-09 | Mari |
| allcell | UQCRQ | 1.07E-08 | 0.545334562 | 0.345 | 0.277 | 0.000210581 | Mari |
| allcell | GTF2A2 | 2.63E-09 | 0.545223896 | 0.262 | 0.189 | 5.18E-05 | Mari |
| allcell | APOBEC3G | 5.93E-17 | 0.544456927 | 0.592 | 0.48 | 1.17E-12 | Mari |
| allcell | SYF2 | 4.11E-17 | 0.543971385 | 0.496 | 0.376 | 8.08E-13 | Mari |
| allcell | PSMB3 | 3.76E-10 | 0.543925798 | 0.341 | 0.266 | 7.40E-06 | Mari |
| allcell | MRPL52 | 4.50E-08 | 0.543908493 | 0.269 | 0.203 | 0.000886274 | Mari |
| allcell | YPEL3 | 4.37E-11 | 0.542638381 | 0.408 | 0.325 | 8.61E-07 | Mari |
| allcell | SNF8 | 3.98E-08 | 0.538880746 | 0.251 | 0.185 | 0.000783248 | Mari |
| allcell | PSMB10 | 1.34E-10 | 0.537704169 | 0.341 | 0.258 | 2.64E-06 | Mari |
| allcell | TBC1D10C | 9.39E-14 | 0.537025213 | 0.363 | 0.263 | 1.85E-09 | Mari |
| allcell | HMGN2 | 2.27E-11 | 0.536105561 | 0.46 | 0.372 | 4.46E-07 | Mari |
| allcell | MPC2 | 2.20E-09 | 0.535932291 | 0.317 | 0.239 | 4.33E-05 | Mari |
| allcell | COX17 | 1.78E-08 | 0.535899774 | 0.283 | 0.213 | 0.000350191 | Mari |
| allcell | LAMTOR4 | 4.00E-13 | 0.534919087 | 0.415 | 0.321 | 7.88E-09 | Mari |
| allcell | SNRPA1 | 3.26E-08 | 0.534901534 | 0.273 | 0.207 | 0.000641137 | Mari |
| allcell | LSM4 | 3.03E-07 | 0.534659998 | 0.271 | 0.21 | 0.005961355 | Mari |
| allcell | ATP5F1C | 2.23E-13 | 0.533376063 | 0.398 | 0.294 | 4.40E-09 | Mari |
| allcell | SPCS2 | 3.17E-15 | 0.531450378 | 0.46 | 0.353 | 6.23E-11 | Mari |
| allcell | SLC39A10 | 3.39E-13 | 0.531112187 | 0.402 | 0.302 | 6.68E-09 | Mari |
| allcell | GTF3A | 1.21E-13 | 0.529641203 | 0.427 | 0.33 | 2.38E-09 | Mari |
| allcell | ARPC3 | 2.84E-30 | 0.529378571 | 0.776 | 0.658 | 5.59E-26 | Mari |
| allcell | TRAF3IP3 | 3.84E-09 | 0.528777271 | 0.316 | 0.244 | 7.55E-05 | Mari |
| allcell | SERF2 | 6.99E-41 | 0.527260729 | 0.887 | 0.803 | 1.38E-36 | Mari |
| allcell | ARPC1B | 5.79E-16 | 0.52313575 | 0.503 | 0.393 | 1.14E-11 | Mari |

| | | | | | | | |
|---|---|---|---|---|---|---|---|
| allcell | GLIPR2 | 2.93E-10 | 0.52304015 | 0.377 | 0.296 | 5.77E-06 | Mari |
| allcell | HMGN3 | 5.78E-12 | 0.521833335 | 0.398 | 0.306 | 1.14E-07 | Mari |
| allcell | ATP5PD | 5.16E-09 | 0.520317319 | 0.288 | 0.215 | 0.000101662 | Mari |
| allcell | TMEM258 | 4.14E-15 | 0.519649756 | 0.425 | 0.319 | 8.16E-11 | Mari |
| allcell | TMA7 | 2.56E-34 | 0.51895394 | 0.8 | 0.665 | 5.04E-30 | Mari |
| allcell | POLR2L | 2.31E-18 | 0.518280309 | 0.531 | 0.405 | 4.54E-14 | Mari |
| allcell | NDUFA5 | 3.64E-08 | 0.517929558 | 0.279 | 0.211 | 0.000715599 | Mari |
| allcell | ATP5MC3 | 7.19E-14 | 0.515500357 | 0.487 | 0.395 | 1.41E-09 | Mari |
| allcell | SCP2 | 8.43E-09 | 0.514230835 | 0.306 | 0.233 | 0.000165856 | Mari |
| allcell | KRTCAP2 | 2.67E-13 | 0.512659371 | 0.417 | 0.319 | 5.26E-09 | Mari |
| allcell | ELOB | 4.84E-18 | 0.5117567 | 0.601 | 0.508 | 9.53E-14 | Mari |
| allcell | SF3B6 | 7.35E-18 | 0.511084093 | 0.478 | 0.355 | 1.45E-13 | Mari |
| allcell | NDUFA12 | 8.18E-14 | 0.510078117 | 0.39 | 0.291 | 1.61E-09 | Mari |
| allcell | MDH2 | 2.49E-07 | 0.508101987 | 0.282 | 0.222 | 0.004905549 | Mari |
| allcell | UQCRC2 | 1.84E-09 | 0.50751843 | 0.336 | 0.255 | 3.63E-05 | Mari |
| allcell | TBCB | 1.86E-09 | 0.506274315 | 0.325 | 0.25 | 3.66E-05 | Mari |
| allcell | ECH1 | 8.21E-09 | 0.506067942 | 0.283 | 0.21 | 0.000161583 | Mari |
| allcell | NDUFB1 | 9.11E-09 | 0.505508719 | 0.31 | 0.237 | 0.000179341 | Mari |
| allcell | PSMC5 | 1.64E-08 | 0.505149496 | 0.282 | 0.21 | 0.000323668 | Mari |
| allcell | KLRD1 | 2.96E-30 | 0.504878312 | 0.853 | 0.766 | 5.83E-26 | Mari |
| allcell | SSU72 | 1.10E-06 | 0.502896521 | 0.27 | 0.21 | 0.021712843 | Mari |
| allcell | SEM1 | 1.29E-10 | 0.502379318 | 0.368 | 0.283 | 2.53E-06 | Mari |
| allcell | SNRPD3 | 4.08E-09 | 0.501381646 | 0.286 | 0.211 | 8.03E-05 | Mari |
| allcell | ATP6V1G1 | 2.83E-17 | 0.500861367 | 0.529 | 0.413 | 5.57E-13 | Mari |
| allcell | IGKC | 8.05E-71 | 2.683897058 | 0.316 | 0.127 | 1.58E-66 | Tb |
| allcell | SLC35F1 | 4.03E-138 | 2.454620054 | 0.357 | 0.078 | 7.93E-134 | Tb |
| allcell | S100A9 | 3.10E-54 | 1.63782491 | 0.334 | 0.149 | 6.10E-50 | Tb |
| allcell | S100A8 | 2.20E-48 | 1.574565208 | 0.316 | 0.144 | 4.33E-44 | Tb |
| allcell | KRT14 | 3.24E-42 | 1.330612914 | 0.28 | 0.127 | 6.37E-38 | Tb |
| allcell | GLCCI1 | 1.17E-31 | 0.962179541 | 0.453 | 0.301 | 2.31E-27 | Tb |
| allcell | PRMT9 | 2.55E-25 | 0.937018334 | 0.382 | 0.249 | 5.01E-21 | Tb |
| allcell | OXNAD1 | 1.57E-18 | 0.832990466 | 0.25 | 0.149 | 3.09E-14 | Tb |
| allcell | PAK1 | 7.93E-13 | 0.831649496 | 0.3 | 0.216 | 1.56E-08 | Tb |
| allcell | TWISTNB | 4.53E-11 | 0.822762346 | 0.332 | 0.254 | 8.91E-07 | Tb |
| allcell | MT-ND3 | 4.27E-95 | 0.795342891 | 0.958 | 0.833 | 8.41E-91 | Tb |
| allcell | EGR1 | 5.20E-17 | 0.785445047 | 0.253 | 0.157 | 1.02E-12 | Tb |
| allcell | STRBP | 9.16E-17 | 0.762237108 | 0.285 | 0.185 | 1.80E-12 | Tb |
| allcell | MT-ATP6 | 1.99E-73 | 0.760902174 | 0.965 | 0.838 | 3.91E-69 | Tb |
| allcell | TSC22D1 | 1.59E-09 | 0.757569825 | 0.288 | 0.214 | 3.14E-05 | Tb |
| allcell | KRT10 | 2.51E-12 | 0.754291206 | 0.463 | 0.379 | 4.94E-08 | Tb |
| allcell | NFE2L3 | 1.24E-14 | 0.7476675 | 0.309 | 0.214 | 2.44E-10 | Tb |
| allcell | LDLRAD4 | 7.13E-13 | 0.742638825 | 0.268 | 0.184 | 1.40E-08 | Tb |
| allcell | XIST | 2.00E-28 | 0.736846397 | 0.373 | 0.223 | 3.93E-24 | Tb |
| allcell | RASA1 | 4.90E-16 | 0.721918632 | 0.333 | 0.23 | 9.65E-12 | Tb |
| allcell | RGS16 | 6.00E-08 | 0.716728223 | 0.263 | 0.2 | 0.001180263 | Tb |
| allcell | ZNF10 | 6.60E-19 | 0.70075156 | 0.338 | 0.222 | 1.30E-14 | Tb |
| allcell | HIP1 | 9.38E-15 | 0.698543148 | 0.299 | 0.204 | 1.85E-10 | Tb |
| allcell | RGCC | 8.93E-25 | 0.69501067 | 0.528 | 0.384 | 1.76E-20 | Tb |
| allcell | MT-CYB | 1.63E-34 | 0.694357869 | 0.867 | 0.782 | 3.22E-30 | Tb |
| allcell | LTB | 8.85E-08 | 0.694261552 | 0.27 | 0.206 | 0.00174147 | Tb |
| allcell | SMURF2 | 4.73E-19 | 0.678403975 | 0.379 | 0.26 | 9.30E-15 | Tb |
| allcell | NSUN6 | 5.24E-13 | 0.671957813 | 0.373 | 0.279 | 1.03E-08 | Tb |
| allcell | JPX | 2.29E-10 | 0.661762813 | 0.266 | 0.192 | 4.51E-06 | Tb |
| allcell | STX5 | 2.72E-14 | 0.656335211 | 0.289 | 0.192 | 5.36E-10 | Tb |
| allcell | NAF1 | 2.92E-08 | 0.651930343 | 0.264 | 0.198 | 0.000575312 | Tb |
| allcell | NEK1 | 1.60E-09 | 0.645450193 | 0.347 | 0.275 | 3.14E-05 | Tb |

| | | | | | | | |
|---|---|---|---|---|---|---|---|
| allcell | FOSB | 6.67E-21 | 0.635697382 | 0.484 | 0.354 | 1.31E-16 | Tb |
| allcell | DLEU2 | 1.78E-10 | 0.630115712 | 0.399 | 0.319 | 3.51E-06 | Tb |
| allcell | YME1L1 | 3.30E-19 | 0.624889874 | 0.592 | 0.486 | 6.49E-15 | Tb |
| allcell | MT-CO3 | 4.13E-41 | 0.622340144 | 0.924 | 0.833 | 8.12E-37 | Tb |
| allcell | GBE1 | 9.06E-07 | 0.60813325 | 0.275 | 0.214 | 0.017825191 | Tb |
| allcell | RAB11A | 6.49E-21 | 0.605275882 | 0.575 | 0.444 | 1.28E-16 | Tb |
| allcell | PPP1R14B | 9.94E-13 | 0.600047112 | 0.353 | 0.26 | 1.96E-08 | Tb |
| allcell | INTS6 | 1.10E-21 | 0.596926408 | 0.574 | 0.447 | 2.16E-17 | Tb |
| allcell | CCDC97 | 1.45E-09 | 0.595677557 | 0.25 | 0.176 | 2.85E-05 | Tb |
| allcell | ZNF638 | 1.39E-12 | 0.595461688 | 0.332 | 0.24 | 2.73E-08 | Tb |
| allcell | MT-ND1 | 2.60E-37 | 0.593301357 | 0.823 | 0.713 | 5.11E-33 | Tb |
| allcell | CASK | 7.18E-14 | 0.579805604 | 0.411 | 0.306 | 1.41E-09 | Tb |
| allcell | ZSWIM6 | 3.39E-13 | 0.577427227 | 0.546 | 0.46 | 6.68E-09 | Tb |
| allcell | PPP2R3C | 4.31E-08 | 0.577213212 | 0.26 | 0.196 | 0.00084802 | Tb |
| allcell | PBRM1 | 5.66E-08 | 0.567525126 | 0.279 | 0.214 | 0.00111458 | Tb |
| allcell | CAPG | 1.75E-10 | 0.565896052 | 0.335 | 0.251 | 3.44E-06 | Tb |
| allcell | ZBTB20 | 7.33E-17 | 0.565156198 | 0.53 | 0.416 | 1.44E-12 | Tb |
| allcell | TNFRSF18 | 9.71E-21 | 0.562156513 | 0.482 | 0.345 | 1.91E-16 | Tb |
| allcell | SSBP2 | 1.18E-07 | 0.556176091 | 0.3 | 0.232 | 0.002319799 | Tb |
| allcell | KDM5B | 9.94E-08 | 0.551353468 | 0.287 | 0.224 | 0.001955766 | Tb |
| allcell | GADD45G | 7.98E-07 | 0.550091296 | 0.299 | 0.239 | 0.015715593 | Tb |
| allcell | TYMP | 2.33E-08 | 0.54422253 | 0.257 | 0.19 | 0.000458612 | Tb |
| allcell | KLRC3 | 4.98E-18 | 0.539424971 | 0.436 | 0.31 | 9.81E-14 | Tb |
| allcell | HMGCS1 | 1.16E-08 | 0.534765219 | 0.264 | 0.192 | 0.000228772 | Tb |
| allcell | TNRC6C | 1.11E-07 | 0.534410952 | 0.29 | 0.223 | 0.002188028 | Tb |
| allcell | MT-ND4 | 3.39E-53 | 0.532785091 | 0.944 | 0.853 | 6.67E-49 | Tb |
| allcell | SGMS1 | 8.71E-08 | 0.52349216 | 0.25 | 0.184 | 0.001714135 | Tb |
| allcell | RABGEF1 | 1.01E-09 | 0.514382441 | 0.363 | 0.283 | 1.98E-05 | Tb |
| allcell | BRAF | 1.17E-09 | 0.513947472 | 0.549 | 0.468 | 2.31E-05 | Tb |
| allcell | TRDC | 4.65E-37 | 0.513863608 | 0.704 | 0.51 | 9.15E-33 | Tb |
| allcell | RBKS | 2.07E-07 | 0.512958356 | 0.382 | 0.313 | 0.004084043 | Tb |
| allcell | FTX | 4.77E-09 | 0.512907742 | 0.308 | 0.231 | 9.40E-05 | Tb |
| allcell | ARIH1 | 5.24E-17 | 0.508551533 | 0.607 | 0.5 | 1.03E-12 | Tb |
| allcell | MT-ND2 | 1.93E-33 | 0.508280516 | 0.864 | 0.767 | 3.81E-29 | Tb |
| allcell | HNRNPH1 | 1.08E-15 | 0.502479447 | 0.614 | 0.493 | 2.14E-11 | Tb |
| allcell | MGAT5 | 8.37E-10 | 0.500401373 | 0.314 | 0.233 | 1.65E-05 | Tb |
| **Disease-associated genes in TCF7 NK** | | | | | | | |
| TCF7 NK | TRAF4 | 7.65E-10 | 1.901388116 | 0.268 | 0.101 | 1.50E-05 | Lep-L |
| TCF7 NK | ALDOA | 1.91E-11 | 1.879416603 | 0.282 | 0.097 | 3.76E-07 | Lep-L |
| TCF7 NK | IGFBP4 | 2.42E-08 | 2.07327158 | 0.317 | 0.148 | 0.000476237 | Lep-L |
| TCF7 NK | PRKY | 9.72E-19 | 2.541636471 | 0.317 | 0.084 | 1.91E-14 | Lep-L |
| TCF7 NK | HLA-DQB1 | 7.13E-07 | 1.679181172 | 0.331 | 0.181 | 0.014026181 | Lep-L |
| TCF7 NK | CDKN1A | 1.39E-07 | 1.73254663 | 0.366 | 0.207 | 0.002727317 | Lep-L |
| TCF7 NK | STMN1 | 6.47E-07 | 1.221658404 | 0.373 | 0.213 | 0.012726055 | Lep-L |
| TCF7 NK | UBALD2 | 8.52E-07 | 1.563641279 | 0.38 | 0.233 | 0.016761247 | Lep-L |
| TCF7 NK | MAPKAPK2 | 2.25E-08 | 1.536883477 | 0.408 | 0.238 | 0.000442609 | Lep-L |
| TCF7 NK | STX11 | 1.92E-06 | 1.108522442 | 0.423 | 0.271 | 0.037779941 | Lep-L |
| TCF7 NK | CDKN1B | 2.01E-07 | 1.476741955 | 0.423 | 0.271 | 0.003960272 | Lep-L |
| TCF7 NK | PRR7 | 4.17E-11 | 1.631947777 | 0.423 | 0.209 | 8.20E-07 | Lep-L |
| TCF7 NK | RCOR1 | 1.91E-06 | 1.093835931 | 0.43 | 0.278 | 0.037584772 | Lep-L |
| TCF7 NK | GNA13 | 2.16E-06 | 1.106008797 | 0.43 | 0.279 | 0.042577273 | Lep-L |
| TCF7 NK | RBM38 | 7.02E-07 | 1.315208325 | 0.43 | 0.294 | 0.013825805 | Lep-L |
| TCF7 NK | APOE | 1.29E-12 | 1.951442935 | 0.437 | 0.216 | 2.53E-08 | Lep-L |
| TCF7 NK | DDX3Y | 6.55E-28 | 2.38467145 | 0.437 | 0.106 | 1.29E-23 | Lep-L |
| TCF7 NK | FKBP4 | 1.71E-07 | 1.003653107 | 0.472 | 0.289 | 0.003356919 | Lep-L |
| TCF7 NK | RHOF | 9.72E-08 | 1.218294504 | 0.472 | 0.306 | 0.001912389 | Lep-L |

| | | | | | | | |
|---|---|---|---|---|---|---|---|
| TCF7 NK | NINJ1 | 4.03E-09 | 1.075569611 | 0.479 | 0.273 | 7.94E-05 | Lep-L |
| TCF7 NK | CNN2 | 4.63E-11 | 1.726865775 | 0.493 | 0.289 | 9.12E-07 | Lep-L |
| TCF7 NK | BCL6 | 1.56E-08 | 1.486895975 | 0.5 | 0.343 | 0.000306176 | Lep-L |
| TCF7 NK | RBFOX2 | 9.63E-11 | 1.917579841 | 0.5 | 0.317 | 1.90E-06 | Lep-L |
| TCF7 NK | TUBB | 2.88E-08 | 1.55815112 | 0.528 | 0.381 | 0.000566103 | Lep-L |
| TCF7 NK | FLNA | 1.97E-09 | 1.222270912 | 0.542 | 0.369 | 3.88E-05 | Lep-L |
| TCF7 NK | DOK2 | 4.19E-08 | 1.274140661 | 0.556 | 0.417 | 0.00082484 | Lep-L |
| TCF7 NK | GABARAPL1 | 1.28E-07 | 1.370464082 | 0.556 | 0.432 | 0.002526326 | Lep-L |
| TCF7 NK | MKNK2 | 1.85E-10 | 1.291316926 | 0.563 | 0.35 | 3.64E-06 | Lep-L |
| TCF7 NK | ATP1B3 | 9.85E-10 | 1.531477997 | 0.592 | 0.436 | 1.94E-05 | Lep-L |
| TCF7 NK | BIRC3 | 2.26E-08 | 1.399396884 | 0.613 | 0.479 | 0.000444504 | Lep-L |
| TCF7 NK | GPX4 | 3.89E-07 | 0.912324065 | 0.655 | 0.546 | 0.007662174 | Lep-L |
| TCF7 NK | BAZ1A | 1.63E-10 | 1.072986311 | 0.662 | 0.455 | 3.21E-06 | Lep-L |
| TCF7 NK | PIM3 | 8.12E-18 | 1.563988649 | 0.676 | 0.371 | 1.60E-13 | Lep-L |
| TCF7 NK | TAGAP | 1.85E-06 | 0.836914667 | 0.683 | 0.569 | 0.036425267 | Lep-L |
| TCF7 NK | CYTIP | 1.30E-08 | 1.020986548 | 0.697 | 0.6 | 0.000255589 | Lep-L |
| TCF7 NK | TUBA4A | 2.22E-06 | 0.863551503 | 0.718 | 0.626 | 0.043771478 | Lep-L |
| TCF7 NK | NFE2L2 | 2.97E-08 | 0.752367136 | 0.725 | 0.62 | 0.000585002 | Lep-L |
| TCF7 NK | EIF4G2 | 3.08E-08 | 0.812443923 | 0.725 | 0.609 | 0.000605797 | Lep-L |
| TCF7 NK | EEF2 | 9.45E-08 | 0.899517147 | 0.725 | 0.657 | 0.001861107 | Lep-L |
| TCF7 NK | PTGES3 | 1.21E-06 | 0.682251232 | 0.754 | 0.674 | 0.02373555 | Lep-L |
| TCF7 NK | TUBA1B | 6.95E-08 | 0.840500933 | 0.761 | 0.669 | 0.001367089 | Lep-L |
| TCF7 NK | SMAP2 | 4.48E-10 | 0.87313629 | 0.761 | 0.615 | 8.83E-06 | Lep-L |
| TCF7 NK | METRNL | 2.59E-14 | 1.164345545 | 0.768 | 0.552 | 5.10E-10 | Lep-L |
| TCF7 NK | VPS37B | 7.15E-13 | 1.159371251 | 0.775 | 0.584 | 1.41E-08 | Lep-L |
| TCF7 NK | HNRNPDL | 1.45E-08 | 0.864419199 | 0.782 | 0.717 | 0.00028571 | Lep-L |
| TCF7 NK | TGFB1 | 1.50E-10 | 1.104714081 | 0.796 | 0.712 | 2.96E-06 | Lep-L |
| TCF7 NK | SAT1 | 4.39E-11 | 1.192663589 | 0.796 | 0.686 | 8.65E-07 | Lep-L |
| TCF7 NK | IL7R | 1.62E-17 | 1.297079661 | 0.796 | 0.594 | 3.18E-13 | Lep-L |
| TCF7 NK | CXCR4 | 5.60E-09 | 0.671643528 | 0.81 | 0.674 | 0.000110233 | Lep-L |
| TCF7 NK | RUNX3 | 3.83E-07 | 0.666403853 | 0.817 | 0.776 | 0.007536783 | Lep-L |
| TCF7 NK | YWHAZ | 6.24E-07 | 0.694035668 | 0.817 | 0.748 | 0.012285552 | Lep-L |
| TCF7 NK | YBX1 | 2.18E-06 | 0.593093103 | 0.824 | 0.759 | 0.042879206 | Lep-L |
| TCF7 NK | HSPH1 | 1.70E-07 | 0.56165017 | 0.838 | 0.725 | 0.003337529 | Lep-L |
| TCF7 NK | CD55 | 5.81E-19 | 1.516253254 | 0.852 | 0.705 | 1.14E-14 | Lep-L |
| TCF7 NK | CALM2 | 2.39E-07 | 0.651495994 | 0.866 | 0.792 | 0.004710096 | Lep-L |
| TCF7 NK | H2AFZ | 5.16E-09 | 0.780155542 | 0.866 | 0.762 | 0.000101539 | Lep-L |
| TCF7 NK | CLEC2B | 6.42E-11 | 0.855547219 | 0.866 | 0.802 | 1.26E-06 | Lep-L |
| TCF7 NK | NFKB1 | 5.83E-07 | 0.531412129 | 0.873 | 0.801 | 0.011476248 | Lep-L |
| TCF7 NK | PNRC1 | 2.15E-08 | 1.006537619 | 0.873 | 0.89 | 0.000423604 | Lep-L |
| TCF7 NK | ZFP36L2 | 1.64E-08 | 1.024392566 | 0.873 | 0.872 | 0.000323761 | Lep-L |
| TCF7 NK | EZR | 1.14E-20 | 1.450188888 | 0.873 | 0.694 | 2.24E-16 | Lep-L |
| TCF7 NK | EIF4A1 | 1.73E-06 | 0.595081415 | 0.88 | 0.802 | 0.034071043 | Lep-L |
| TCF7 NK | OAZ1 | 4.28E-14 | 0.891545997 | 0.88 | 0.823 | 8.43E-10 | Lep-L |
| TCF7 NK | PABPC1 | 6.84E-08 | 0.635929927 | 0.894 | 0.901 | 0.001346904 | Lep-L |
| TCF7 NK | HSPD1 | 5.83E-12 | 0.638117406 | 0.894 | 0.796 | 1.15E-07 | Lep-L |
| TCF7 NK | SQSTM1 | 2.57E-12 | 0.663949686 | 0.908 | 0.767 | 5.06E-08 | Lep-L |
| TCF7 NK | SARAF | 3.96E-17 | 0.873415705 | 0.915 | 0.846 | 7.80E-13 | Lep-L |
| TCF7 NK | VIM | 2.54E-10 | 0.892701607 | 0.915 | 0.863 | 5.00E-06 | Lep-L |
| TCF7 NK | REL | 1.10E-11 | 0.76240149 | 0.923 | 0.874 | 2.17E-07 | Lep-L |
| TCF7 NK | HSPA8 | 1.04E-13 | 0.967221271 | 0.944 | 0.917 | 2.04E-09 | Lep-L |
| TCF7 NK | HSP90AB1 | 4.57E-13 | 0.744389752 | 0.958 | 0.95 | 9.00E-09 | Lep-L |
| TCF7 NK | SRGN | 9.39E-12 | 0.526169377 | 0.972 | 0.965 | 1.85E-07 | Lep-L |
| TCF7 NK | EIF1 | 6.57E-13 | 0.54909119 | 0.972 | 0.979 | 1.29E-08 | Lep-L |
| TCF7 NK | FTL | 5.19E-18 | 0.972678148 | 0.972 | 0.98 | 1.02E-13 | Lep-L |
| TCF7 NK | FTH1 | 2.62E-11 | 1.155755576 | 0.972 | 0.977 | 5.15E-07 | Lep-L |

| | | | | | | | |
|---|---|---|---|---|---|---|---|
| TCF7 NK | BTG1 | 5.37E-10 | 0.726315061 | 0.979 | 0.988 | 1.06E-05 | Lep-L |
| TCF7 NK | JUND | 1.66E-20 | 0.872579026 | 0.979 | 0.952 | 3.27E-16 | Lep-L |
| TCF7 NK | ATP5PB | 1.28E-07 | 1.136244839 | 0.25 | 0.127 | 0.002519544 | Lep-T |
| TCF7 NK | IGFBP41 | 3.63E-07 | 1.010707522 | 0.253 | 0.137 | 0.007140992 | Lep-T |
| TCF7 NK | HIST1H1D | 2.29E-08 | 1.208578028 | 0.262 | 0.132 | 0.000451469 | Lep-T |
| TCF7 NK | ASTL | 1.37E-15 | 1.521498219 | 0.262 | 0.088 | 2.71E-11 | Lep-T |
| TCF7 NK | HIST2H2AC | 5.46E-12 | 1.406646016 | 0.265 | 0.109 | 1.08E-07 | Lep-T |
| TCF7 NK | SNX9 | 9.53E-09 | 0.981907523 | 0.277 | 0.138 | 0.000187674 | Lep-T |
| TCF7 NK | RASGRP1 | 3.42E-08 | 0.859108888 | 0.283 | 0.145 | 0.000672242 | Lep-T |
| TCF7 NK | XAF1 | 1.00E-09 | 1.005863455 | 0.307 | 0.149 | 1.98E-05 | Lep-T |
| TCF7 NK | DUSP10 | 8.53E-07 | 0.67616059 | 0.331 | 0.197 | 0.01679464 | Lep-T |
| TCF7 NK | HIST1H1C | 3.60E-15 | 1.337767054 | 0.337 | 0.141 | 7.09E-11 | Lep-T |
| TCF7 NK | H1FX | 4.09E-07 | 0.892879595 | 0.34 | 0.205 | 0.008052882 | Lep-T |
| TCF7 NK | BCL31 | 3.39E-16 | 1.471074006 | 0.367 | 0.158 | 6.67E-12 | Lep-T |
| TCF7 NK | EMD | 1.41E-16 | 1.362988895 | 0.373 | 0.158 | 2.78E-12 | Lep-T |
| TCF7 NK | MOB3A | 2.04E-09 | 0.926583339 | 0.38 | 0.211 | 4.02E-05 | Lep-T |
| TCF7 NK | ITGAX | 2.39E-11 | 0.929491052 | 0.38 | 0.195 | 4.70E-07 | Lep-T |
| TCF7 NK | BRD4 | 1.40E-06 | 0.653183407 | 0.395 | 0.256 | 0.027490295 | Lep-T |
| TCF7 NK | RGS2 | 2.05E-07 | 1.004080443 | 0.407 | 0.26 | 0.004027686 | Lep-T |
| TCF7 NK | CNN21 | 1.65E-07 | 0.660887706 | 0.434 | 0.269 | 0.003255074 | Lep-T |
| TCF7 NK | GSPT1 | 4.40E-07 | 0.811423646 | 0.437 | 0.29 | 0.008651289 | Lep-T |
| TCF7 NK | DAZAP2 | 6.99E-17 | 1.279221976 | 0.446 | 0.21 | 1.38E-12 | Lep-T |
| TCF7 NK | KIR2DL4 | 2.75E-07 | 0.625443077 | 0.47 | 0.308 | 0.005413143 | Lep-T |
| TCF7 NK | HIST1H1E | 1.93E-21 | 1.462739878 | 0.473 | 0.208 | 3.80E-17 | Lep-T |
| TCF7 NK | RELB | 1.08E-08 | 0.754971488 | 0.491 | 0.31 | 0.000213146 | Lep-T |
| TCF7 NK | RBFOX21 | 2.63E-17 | 1.47154316 | 0.515 | 0.276 | 5.18E-13 | Lep-T |
| TCF7 NK | CSRNP11 | 7.30E-09 | 0.884921637 | 0.524 | 0.353 | 0.000143762 | Lep-T |
| TCF7 NK | RHOH | 5.67E-07 | 0.676119164 | 0.548 | 0.403 | 0.011169031 | Lep-T |
| TCF7 NK | GAS5 | 1.40E-07 | 0.626633708 | 0.56 | 0.398 | 0.002747049 | Lep-T |
| TCF7 NK | ITGB2 | 7.30E-07 | 0.591875786 | 0.563 | 0.391 | 0.014378349 | Lep-T |
| TCF7 NK | NFKBIZ | 4.28E-12 | 0.955198347 | 0.569 | 0.368 | 8.42E-08 | Lep-T |
| TCF7 NK | PRKX | 2.83E-10 | 0.923216831 | 0.59 | 0.431 | 5.57E-06 | Lep-T |
| TCF7 NK | PHF20 | 2.64E-08 | 0.685656713 | 0.599 | 0.449 | 0.000520579 | Lep-T |
| TCF7 NK | CRTAM | 3.66E-08 | 0.558708597 | 0.605 | 0.438 | 0.000719741 | Lep-T |
| TCF7 NK | KLF2 | 5.61E-07 | 0.915152406 | 0.611 | 0.485 | 0.011050283 | Lep-T |
| TCF7 NK | SLC2A31 | 1.45E-11 | 0.946746849 | 0.657 | 0.46 | 2.85E-07 | Lep-T |
| TCF7 NK | IFITM3 | 9.67E-10 | 0.948437235 | 0.663 | 0.485 | 1.90E-05 | Lep-T |
| TCF7 NK | ARL4C | 9.63E-11 | 0.777887699 | 0.672 | 0.513 | 1.89E-06 | Lep-T |
| TCF7 NK | SYAP1 | 1.52E-09 | 0.586084238 | 0.678 | 0.483 | 2.99E-05 | Lep-T |
| TCF7 NK | LMNA1 | 9.83E-10 | 0.800354553 | 0.693 | 0.524 | 1.93E-05 | Lep-T |
| TCF7 NK | KLF6 | 8.71E-09 | 0.689268077 | 0.699 | 0.607 | 0.000171408 | Lep-T |
| TCF7 NK | METRNL1 | 1.84E-08 | 0.565315104 | 0.714 | 0.528 | 0.000361984 | Lep-T |
| TCF7 NK | ZFP36L1 | 3.54E-15 | 0.982472795 | 0.717 | 0.517 | 6.97E-11 | Lep-T |
| TCF7 NK | MT-ND5 | 6.29E-07 | 0.540324746 | 0.723 | 0.646 | 0.012379975 | Lep-T |
| TCF7 NK | EML4 | 1.30E-08 | 0.622490926 | 0.726 | 0.625 | 0.000256627 | Lep-T |
| TCF7 NK | HSPA5 | 5.51E-07 | 0.702516215 | 0.744 | 0.592 | 0.01084148 | Lep-T |
| TCF7 NK | BHLHE40 | 6.83E-15 | 0.800307655 | 0.747 | 0.556 | 1.34E-10 | Lep-T |
| TCF7 NK | EEF21 | 2.55E-08 | 0.718132664 | 0.768 | 0.629 | 0.000500977 | Lep-T |
| TCF7 NK | MCL11 | 9.79E-10 | 0.569761865 | 0.774 | 0.598 | 1.93E-05 | Lep-T |
| TCF7 NK | ADGRE5 | 2.95E-13 | 0.821975272 | 0.774 | 0.638 | 5.81E-09 | Lep-T |
| TCF7 NK | CD74 | 6.27E-11 | 1.449015168 | 0.777 | 0.681 | 1.23E-06 | Lep-T |
| TCF7 NK | CXCR41 | 1.94E-10 | 0.653052051 | 0.783 | 0.656 | 3.81E-06 | Lep-T |
| TCF7 NK | FAM177A1 | 8.08E-08 | 0.521866854 | 0.81 | 0.659 | 0.001590241 | Lep-T |
| TCF7 NK | LITAF1 | 1.41E-19 | 0.920798244 | 0.813 | 0.586 | 2.78E-15 | Lep-T |
| TCF7 NK | TSC22D3 | 4.45E-17 | 1.295927552 | 0.816 | 0.658 | 8.76E-13 | Lep-T |
| TCF7 NK | NR4A21 | 4.91E-18 | 0.780502109 | 0.828 | 0.672 | 9.66E-14 | Lep-T |

| | | | | | | | |
|---|---|---|---|---|---|---|---|
| TCF7 NK | MT-CO3 | 6.52E-12 | 0.67069447 | 0.84 | 0.911 | 1.28E-07 | Lep-T |
| TCF7 NK | AREG | 3.18E-07 | 0.624804939 | 0.867 | 0.833 | 0.006263438 | Lep-T |
| TCF7 NK | SQSTM11 | 1.40E-08 | 0.501447317 | 0.87 | 0.753 | 0.000275592 | Lep-T |
| TCF7 NK | ZFP36L21 | 8.73E-08 | 0.691999415 | 0.91 | 0.859 | 0.001718764 | Lep-T |
| TCF7 NK | XCL11 | 6.79E-15 | 1.467058198 | 0.91 | 0.842 | 1.34E-10 | Lep-T |
| TCF7 NK | XCL21 | 2.46E-06 | 0.955143021 | 0.916 | 0.86 | 0.048460845 | Lep-T |
| TCF7 NK | IFITM2 | 1.64E-19 | 1.146361704 | 0.925 | 0.825 | 3.22E-15 | Lep-T |
| TCF7 NK | PNRC11 | 1.14E-17 | 0.712137126 | 0.946 | 0.869 | 2.25E-13 | Lep-T |
| TCF7 NK | SRGN1 | 1.89E-17 | 0.662304772 | 0.979 | 0.961 | 3.72E-13 | Lep-T |
| TCF7 NK | LSM6 | 7.88E-07 | 1.025821296 | 0.254 | 0.146 | 0.015520364 | Mari |
| TCF7 NK | CYBC1 | 1.47E-07 | 0.844302845 | 0.259 | 0.141 | 0.0028914 | Mari |
| TCF7 NK | ATRAID | 3.75E-07 | 0.975175153 | 0.264 | 0.15 | 0.007387817 | Mari |
| TCF7 NK | NDUFS7 | 8.84E-07 | 0.952359005 | 0.267 | 0.159 | 0.01739643 | Mari |
| TCF7 NK | NDUFB5 | 1.11E-06 | 0.88840503 | 0.269 | 0.161 | 0.021877998 | Mari |
| TCF7 NK | PYURF | 1.23E-06 | 0.865586797 | 0.288 | 0.176 | 0.024113627 | Mari |
| TCF7 NK | ETFB | 6.34E-07 | 0.92505149 | 0.293 | 0.176 | 0.012475421 | Mari |
| TCF7 NK | PYHIN1 | 1.06E-12 | 1.4726247 | 0.303 | 0.145 | 2.08E-08 | Mari |
| TCF7 NK | HSPB11 | 1.87E-06 | 0.891471778 | 0.306 | 0.196 | 0.03672164 | Mari |
| TCF7 NK | DCXR | 3.80E-07 | 0.988276285 | 0.313 | 0.198 | 0.00748468 | Mari |
| TCF7 NK | TMIGD2 | 3.31E-08 | 1.064460445 | 0.316 | 0.187 | 0.000651785 | Mari |
| TCF7 NK | TRAPPC1 | 5.01E-07 | 0.765768706 | 0.319 | 0.199 | 0.009853175 | Mari |
| TCF7 NK | RER1 | 4.09E-09 | 1.10569231 | 0.319 | 0.186 | 8.05E-05 | Mari |
| TCF7 NK | IRF9 | 1.59E-08 | 1.021895451 | 0.321 | 0.187 | 0.000312417 | Mari |
| TCF7 NK | TXNL4A | 2.32E-06 | 0.693457214 | 0.329 | 0.215 | 0.045640272 | Mari |
| TCF7 NK | TMCO1 | 1.62E-07 | 0.827951021 | 0.342 | 0.213 | 0.003182735 | Mari |
| TCF7 NK | SEPTIN1 | 2.62E-08 | 0.981866849 | 0.345 | 0.215 | 0.000515929 | Mari |
| TCF7 NK | TOMM5 | 8.27E-07 | 0.799797617 | 0.352 | 0.238 | 0.016287883 | Mari |
| TCF7 NK | ATP5MPL | 2.52E-07 | 0.837921638 | 0.358 | 0.236 | 0.004967124 | Mari |
| TCF7 NK | TIMP1 | 1.56E-06 | 0.759353891 | 0.37 | 0.262 | 0.030683144 | Mari |
| TCF7 NK | BLOC1S1 | 8.79E-07 | 0.887586094 | 0.373 | 0.266 | 0.01730609 | Mari |
| TCF7 NK | MRPL33 | 2.19E-08 | 0.808179399 | 0.383 | 0.246 | 0.00043151 | Mari |
| TCF7 NK | TRAF3IP3 | 2.56E-09 | 0.884801289 | 0.386 | 0.246 | 5.03E-05 | Mari |
| TCF7 NK | TMEM14B | 2.47E-07 | 0.699685842 | 0.399 | 0.273 | 0.004855359 | Mari |
| TCF7 NK | COX7A2L | 8.45E-07 | 0.739163212 | 0.399 | 0.28 | 0.016632696 | Mari |
| TCF7 NK | MED10 | 1.17E-08 | 0.785088675 | 0.399 | 0.253 | 0.000230444 | Mari |
| TCF7 NK | OCIAD2 | 1.61E-06 | 0.696548731 | 0.404 | 0.289 | 0.031778431 | Mari |
| TCF7 NK | SNRPE | 4.77E-08 | 0.655619371 | 0.407 | 0.268 | 0.000939069 | Mari |
| TCF7 NK | NDUFS8 | 8.69E-07 | 0.721488236 | 0.412 | 0.285 | 0.01710153 | Mari |
| TCF7 NK | NDUFB7 | 2.87E-09 | 0.886783895 | 0.412 | 0.263 | 5.66E-05 | Mari |
| TCF7 NK | NDUFS6 | 2.00E-06 | 0.710732453 | 0.415 | 0.301 | 0.039350386 | Mari |
| TCF7 NK | UQCRC2 | 8.15E-07 | 0.696596556 | 0.425 | 0.293 | 0.0160343 | Mari |
| TCF7 NK | REX1BD | 4.88E-09 | 0.895946178 | 0.425 | 0.289 | 9.61E-05 | Mari |
| TCF7 NK | BANF1 | 9.34E-08 | 0.748674151 | 0.435 | 0.31 | 0.001838199 | Mari |
| TCF7 NK | C1QBP | 5.08E-08 | 0.628983459 | 0.438 | 0.29 | 0.000999486 | Mari |
| TCF7 NK | SSBP1 | 1.91E-07 | 0.6490455 | 0.443 | 0.31 | 0.003765306 | Mari |
| TCF7 NK | SPCS1 | 7.45E-07 | 0.652901611 | 0.443 | 0.326 | 0.014658284 | Mari |
| TCF7 NK | UQCR10 | 4.46E-07 | 0.631703881 | 0.446 | 0.32 | 0.00877925 | Mari |
| TCF7 NK | OSTC | 5.25E-10 | 0.909231766 | 0.446 | 0.291 | 1.03E-05 | Mari |
| TCF7 NK | TRBC2 | 6.46E-08 | 0.740668198 | 0.459 | 0.327 | 0.001272052 | Mari |
| TCF7 NK | GABARAPL2 | 1.03E-08 | 0.726159772 | 0.466 | 0.311 | 0.000202811 | Mari |
| TCF7 NK | NDUFC2 | 3.75E-17 | 1.176704447 | 0.469 | 0.25 | 7.38E-13 | Mari |
| TCF7 NK | ATP5F1C | 7.68E-07 | 0.640583414 | 0.474 | 0.339 | 0.01511929 | Mari |
| TCF7 NK | HMGN3 | 5.35E-09 | 0.718853386 | 0.474 | 0.323 | 0.000105227 | Mari |
| TCF7 NK | CIB1 | 3.52E-07 | 0.523602587 | 0.477 | 0.342 | 0.006926475 | Mari |
| TCF7 NK | GTF3A | 9.71E-07 | 0.577561788 | 0.482 | 0.365 | 0.019114079 | Mari |
| TCF7 NK | ATP5PF | 8.82E-07 | 0.660240717 | 0.492 | 0.383 | 0.017355896 | Mari |

| | | | | | | | |
|---|---|---|---|---|---|---|---|
| TCF7 NK | ARPC1B | 5.61E-08 | 0.703202255 | 0.508 | 0.37 | 0.001103692 | Mari |
| TCF7 NK | PAXX | 3.87E-08 | 0.619506682 | 0.513 | 0.373 | 0.00076118 | Mari |
| TCF7 NK | SLC39A10 | 3.44E-08 | 0.673296285 | 0.513 | 0.371 | 0.000677964 | Mari |
| TCF7 NK | ATP5MF | 7.24E-13 | 0.866356041 | 0.518 | 0.342 | 1.42E-08 | Mari |
| TCF7 NK | SF3B6 | 7.88E-07 | 0.503176198 | 0.521 | 0.387 | 0.015503641 | Mari |
| TCF7 NK | GZMA | 7.07E-09 | 1.145470176 | 0.523 | 0.402 | 0.000139118 | Mari |
| TCF7 NK | ATP5ME | 2.56E-14 | 0.919892269 | 0.531 | 0.335 | 5.05E-10 | Mari |
| TCF7 NK | PRELID1 | 1.06E-06 | 0.541115222 | 0.536 | 0.421 | 0.02092379 | Mari |
| TCF7 NK | APOBEC3G | 4.63E-07 | 0.546336736 | 0.544 | 0.413 | 0.009109635 | Mari |
| TCF7 NK | SMDT1 | 1.61E-07 | 0.618401761 | 0.544 | 0.402 | 0.003163043 | Mari |
| TCF7 NK | PRDX5 | 1.16E-08 | 0.550578721 | 0.547 | 0.388 | 0.000229209 | Mari |
| TCF7 NK | ANAPC16 | 7.06E-08 | 0.670111277 | 0.547 | 0.427 | 0.001389906 | Mari |
| TCF7 NK | ATP5MC3 | 3.66E-07 | 0.512620869 | 0.552 | 0.428 | 0.007206237 | Mari |
| TCF7 NK | PSMB8 | 1.18E-06 | 0.514330178 | 0.554 | 0.433 | 0.023248792 | Mari |
| TCF7 NK | NDUFB2 | 4.85E-09 | 0.656469466 | 0.562 | 0.407 | 9.54E-05 | Mari |
| TCF7 NK | SEC61G | 6.74E-09 | 0.592825771 | 0.565 | 0.401 | 0.000132629 | Mari |
| TCF7 NK | UQCR11 | 3.89E-09 | 0.69976111 | 0.565 | 0.427 | 7.66E-05 | Mari |
| TCF7 NK | COX5B | 6.66E-07 | 0.587614732 | 0.567 | 0.466 | 0.013106541 | Mari |
| TCF7 NK | PSMB1 | 4.23E-10 | 0.630856918 | 0.596 | 0.438 | 8.34E-06 | Mari |
| TCF7 NK | BST2 | 9.75E-17 | 0.947996724 | 0.598 | 0.417 | 1.92E-12 | Mari |
| TCF7 NK | COX8A | 1.14E-11 | 0.661441074 | 0.601 | 0.415 | 2.24E-07 | Mari |
| TCF7 NK | NDUFA13 | 5.54E-08 | 0.553028 | 0.604 | 0.475 | 0.001091258 | Mari |
| TCF7 NK | PSMB9 | 3.04E-07 | 0.607240786 | 0.604 | 0.492 | 0.00597485 | Mari |
| TCF7 NK | APRT | 1.46E-06 | 0.564572816 | 0.606 | 0.503 | 0.028686175 | Mari |
| TCF7 NK | GMFG | 8.47E-10 | 0.673204606 | 0.606 | 0.472 | 1.67E-05 | Mari |
| TCF7 NK | COX6B1 | 5.34E-11 | 0.735352601 | 0.627 | 0.468 | 1.05E-06 | Mari |
| TCF7 NK | DRAP1 | 5.48E-09 | 0.635245001 | 0.632 | 0.516 | 0.000107917 | Mari |
| TCF7 NK | HMGN1 | 7.37E-08 | 0.547371099 | 0.637 | 0.507 | 0.001450297 | Mari |
| TCF7 NK | COX6C | 6.97E-09 | 0.567339296 | 0.637 | 0.5 | 0.000137205 | Mari |
| TCF7 NK | ELOB | 5.62E-08 | 0.509764151 | 0.648 | 0.538 | 0.001105963 | Mari |
| TCF7 NK | ATP6V1G1 | 1.77E-09 | 0.547384909 | 0.655 | 0.482 | 3.48E-05 | Mari |
| TCF7 NK | PPIB | 7.24E-08 | 0.576806988 | 0.671 | 0.538 | 0.001424298 | Mari |
| TCF7 NK | KLRF1 | 1.23E-16 | 0.986346962 | 0.674 | 0.491 | 2.42E-12 | Mari |
| TCF7 NK | GSTP1 | 6.42E-08 | 0.557833284 | 0.679 | 0.553 | 0.001263089 | Mari |
| TCF7 NK | CD52 | 4.08E-08 | 0.590696918 | 0.687 | 0.54 | 0.000803213 | Mari |
| TCF7 NK | CORO1A | 2.96E-08 | 0.604222139 | 0.697 | 0.602 | 0.00058243 | Mari |
| TCF7 NK | COX7A2 | 2.34E-12 | 0.673848993 | 0.702 | 0.526 | 4.61E-08 | Mari |
| TCF7 NK | EIF3G | 3.97E-07 | 0.532123566 | 0.705 | 0.6 | 0.00781769 | Mari |
| TCF7 NK | NDUFA4 | 1.08E-12 | 0.696336622 | 0.705 | 0.549 | 2.12E-08 | Mari |
| TCF7 NK | MYL12B | 7.20E-10 | 0.5070071 | 0.731 | 0.626 | 1.42E-05 | Mari |
| TCF7 NK | SAMD3 | 4.05E-10 | 0.535315135 | 0.736 | 0.55 | 7.96E-06 | Mari |
| TCF7 NK | TRDC | 7.50E-14 | 0.902774705 | 0.736 | 0.611 | 1.48E-09 | Mari |
| TCF7 NK | CD2 | 5.97E-09 | 0.565834577 | 0.746 | 0.624 | 0.000117554 | Mari |
| TCF7 NK | FCER1G | 4.23E-08 | 0.575937337 | 0.754 | 0.674 | 0.000832951 | Mari |
| TCF7 NK | SEPTIN7 | 6.38E-10 | 0.512555806 | 0.772 | 0.69 | 1.26E-05 | Mari |
| TCF7 NK | HOPX | 9.98E-11 | 0.636810833 | 0.798 | 0.693 | 1.96E-06 | Mari |
| TCF7 NK | MYL12A | 1.62E-14 | 0.758611019 | 0.803 | 0.668 | 3.19E-10 | Mari |
| TCF7 NK | ARPC3 | 5.36E-15 | 0.60825387 | 0.816 | 0.658 | 1.05E-10 | Mari |
| TCF7 NK | TMA7 | 8.09E-14 | 0.523413778 | 0.826 | 0.654 | 1.59E-09 | Mari |
| TCF7 NK | SH3BGRL3 | 1.92E-12 | 0.559903629 | 0.876 | 0.765 | 3.78E-08 | Mari |
| TCF7 NK | ATP5F1E | 1.08E-19 | 0.530820434 | 0.953 | 0.873 | 2.13E-15 | Mari |
| TCF7 NK | GNLY | 4.49E-24 | 0.621892098 | 0.992 | 0.974 | 8.83E-20 | Mari |
| TCF7 NK | KRT14 | 3.13E-07 | 1.056509087 | 0.267 | 0.151 | 0.006169214 | Tb |
| TCF7 NK | MRTFA | 8.35E-07 | 0.858185474 | 0.276 | 0.162 | 0.016427153 | Tb |
| TCF7 NK | TMEM184C | 2.19E-07 | 1.157214987 | 0.276 | 0.157 | 0.004304005 | Tb |
| TCF7 NK | IGKC | 5.81E-19 | 2.63142145 | 0.314 | 0.13 | 1.14E-14 | Tb |

| | | | | | | | |
|---|---|---|---|---|---|---|---|
| TCF7 NK | SLC35F1 | 7.96E-32 | 2.503499884 | 0.325 | 0.078 | 1.57E-27 | Tb |
| TCF7 NK | PRMT9 | 1.73E-10 | 1.108162748 | 0.453 | 0.302 | 3.40E-06 | Tb |
| TCF7 NK | GLCCI1 | 6.08E-13 | 1.043935331 | 0.489 | 0.308 | 1.20E-08 | Tb |
| TCF7 NK | INTS6 | 1.67E-07 | 0.685791155 | 0.576 | 0.435 | 0.003284736 | Tb |
| TCF7 NK | ARIH1 | 1.71E-07 | 0.677879931 | 0.617 | 0.498 | 0.003373838 | Tb |
| TCF7 NK | HNRNPH1 | 1.67E-07 | 0.703384338 | 0.659 | 0.519 | 0.003291204 | Tb |
| TCF7 NK | MT-ND1 | 1.89E-09 | 0.563802789 | 0.85 | 0.798 | 3.73E-05 | Tb |
| TCF7 NK | MT-CYB | 2.32E-08 | 0.703883834 | 0.863 | 0.801 | 0.000456945 | Tb |
| TCF7 NK | MT-CO31 | 4.43E-07 | 0.599337601 | 0.924 | 0.877 | 0.008716127 | Tb |
| TCF7 NK | MT-ATP6 | 1.79E-15 | 0.670611425 | 0.955 | 0.851 | 3.52E-11 | Tb |
| TCF7 NK | MT-ND3 | 1.85E-31 | 0.785358019 | 0.978 | 0.865 | 3.63E-27 | Tb |
| **Disease-associated genes in GZMK NK** | | | | | | | |
| GMK NK | DDX3Y | 2.68E-28 | 2.108482757 | 0.384 | 0.108 | 5.27E-24 | Lep-L |
| GMK NK | SERPINH1 | 4.74E-15 | 1.969448251 | 0.298 | 0.115 | 9.32E-11 | Lep-L |
| GMK NK | ALDOA | 3.26E-19 | 1.914238562 | 0.351 | 0.128 | 6.42E-15 | Lep-L |
| GMK NK | TLE1 | 1.62E-16 | 1.828419684 | 0.343 | 0.141 | 3.18E-12 | Lep-L |
| GMK NK | SLC2A3 | 2.32E-21 | 1.789282152 | 0.71 | 0.526 | 4.57E-17 | Lep-L |
| GMK NK | EMD | 7.35E-12 | 1.755965935 | 0.318 | 0.152 | 1.45E-07 | Lep-L |
| GMK NK | APOO | 9.17E-14 | 1.750884454 | 0.282 | 0.108 | 1.81E-09 | Lep-L |
| GMK NK | CNN2 | 9.45E-21 | 1.729256122 | 0.535 | 0.291 | 1.86E-16 | Lep-L |
| GMK NK | FKBP4 | 2.06E-23 | 1.728439096 | 0.522 | 0.255 | 4.05E-19 | Lep-L |
| GMK NK | RBFOX2 | 1.88E-24 | 1.727684337 | 0.571 | 0.304 | 3.71E-20 | Lep-L |
| GMK NK | PTPRCAP | 4.50E-22 | 1.683371633 | 0.278 | 0.068 | 8.86E-18 | Lep-L |
| GMK NK | CSRNP1 | 2.05E-25 | 1.668725101 | 0.633 | 0.361 | 4.04E-21 | Lep-L |
| GMK NK | ZFP36L1 | 1.35E-08 | 1.667595419 | 0.657 | 0.635 | 0.000266522 | Lep-L |
| GMK NK | CXCR6 | 2.16E-11 | 1.637329345 | 0.314 | 0.148 | 4.26E-07 | Lep-L |
| GMK NK | ATP1B3 | 3.34E-31 | 1.63294591 | 0.739 | 0.461 | 6.57E-27 | Lep-L |
| GMK NK | CDKN1A | 1.77E-15 | 1.576130132 | 0.478 | 0.278 | 3.49E-11 | Lep-L |
| GMK NK | ZC3H12A | 1.71E-12 | 1.560150722 | 0.547 | 0.392 | 3.37E-08 | Lep-L |
| GMK NK | NR4A2 | 2.18E-24 | 1.538873415 | 0.837 | 0.738 | 4.28E-20 | Lep-L |
| GMK NK | PAF1 | 1.83E-08 | 1.523560487 | 0.282 | 0.151 | 0.000360794 | Lep-L |
| GMK NK | TAGAP | 1.16E-19 | 1.488242502 | 0.735 | 0.539 | 2.29E-15 | Lep-L |
| GMK NK | HLA-DQB1 | 8.48E-16 | 1.475132072 | 0.514 | 0.309 | 1.67E-11 | Lep-L |
| GMK NK | NR4A1 | 9.20E-14 | 1.447524621 | 0.771 | 0.748 | 1.81E-09 | Lep-L |
| GMK NK | MEF2D | 1.81E-07 | 1.422218078 | 0.31 | 0.188 | 0.003553626 | Lep-L |
| GMK NK | INSIG1 | 1.11E-09 | 1.419398709 | 0.424 | 0.266 | 2.19E-05 | Lep-L |
| GMK NK | HSPA8 | 1.00E-44 | 1.417110601 | 0.976 | 0.918 | 1.97E-40 | Lep-L |
| GMK NK | RBM38 | 4.78E-14 | 1.415725402 | 0.506 | 0.307 | 9.41E-10 | Lep-L |
| GMK NK | METRNL | 1.17E-10 | 1.414525651 | 0.555 | 0.447 | 2.31E-06 | Lep-L |
| GMK NK | HSPA1B | 1.85E-12 | 1.414122859 | 0.857 | 0.782 | 3.65E-08 | Lep-L |
| GMK NK | CCL4L2 | 5.83E-10 | 1.412028002 | 0.49 | 0.328 | 1.15E-05 | Lep-L |
| GMK NK | APOE | 1.85E-08 | 1.404945757 | 0.343 | 0.206 | 0.000365136 | Lep-L |
| GMK NK | ZFP36 | 1.94E-12 | 1.403593686 | 0.882 | 0.867 | 3.83E-08 | Lep-L |
| GMK NK | DUSP5 | 1.15E-12 | 1.392905778 | 0.588 | 0.43 | 2.26E-08 | Lep-L |
| GMK NK | EZR | 3.26E-27 | 1.386557911 | 0.833 | 0.663 | 6.42E-23 | Lep-L |
| GMK NK | CCL3L1 | 2.17E-06 | 1.383601701 | 0.42 | 0.287 | 0.042631404 | Lep-L |
| GMK NK | ICAM1 | 3.62E-07 | 1.376352723 | 0.376 | 0.258 | 0.007128588 | Lep-L |
| GMK NK | CD55 | 7.38E-12 | 1.346735598 | 0.604 | 0.456 | 1.45E-07 | Lep-L |
| GMK NK | UBALD2 | 1.29E-13 | 1.341156023 | 0.473 | 0.279 | 2.54E-09 | Lep-L |
| GMK NK | TGIF1 | 2.71E-09 | 1.340006718 | 0.347 | 0.202 | 5.33E-05 | Lep-L |
| GMK NK | VPS37B | 7.05E-24 | 1.338127287 | 0.739 | 0.526 | 1.39E-19 | Lep-L |
| GMK NK | DOK2 | 4.83E-18 | 1.331151946 | 0.641 | 0.446 | 9.51E-14 | Lep-L |
| GMK NK | BCL3 | 5.84E-07 | 1.321234973 | 0.298 | 0.178 | 0.011489468 | Lep-L |
| GMK NK | SAT1 | 4.28E-20 | 1.27974531 | 0.865 | 0.716 | 8.43E-16 | Lep-L |
| GMK NK | JMY | 1.12E-06 | 1.271501415 | 0.351 | 0.232 | 0.022109437 | Lep-L |
| GMK NK | HSP90AB1 | 9.16E-42 | 1.208662236 | 0.951 | 0.903 | 1.80E-37 | Lep-L |

| | | | | | | | |
|---|---|---|---|---|---|---|---|
| GMK NK | BRD2 | 5.25E-14 | 1.182972353 | 0.763 | 0.65 | 1.03E-09 | Lep-L |
| GMK NK | SLBP | 6.43E-08 | 1.167143878 | 0.367 | 0.231 | 0.001265374 | Lep-L |
| GMK NK | PIM3 | 1.48E-11 | 1.154278099 | 0.494 | 0.319 | 2.91E-07 | Lep-L |
| GMK NK | TGFBR2 | 1.37E-07 | 1.137993464 | 0.453 | 0.321 | 0.002695945 | Lep-L |
| GMK NK | PRR7 | 1.91E-06 | 1.130030124 | 0.371 | 0.26 | 0.037618174 | Lep-L |
| GMK NK | TNFRSF1B | 2.22E-06 | 1.116147779 | 0.355 | 0.254 | 0.043700024 | Lep-L |
| GMK NK | GNA13 | 1.51E-08 | 1.115642468 | 0.424 | 0.277 | 0.000296955 | Lep-L |
| GMK NK | STIP1 | 7.04E-09 | 1.111650753 | 0.42 | 0.278 | 0.000138477 | Lep-L |
| GMK NK | HSPH1 | 2.47E-21 | 1.079660276 | 0.824 | 0.657 | 4.85E-17 | Lep-L |
| GMK NK | TGFB1 | 5.37E-11 | 1.074857906 | 0.776 | 0.715 | 1.06E-06 | Lep-L |
| GMK NK | HBP1 | 2.25E-10 | 1.071683512 | 0.522 | 0.371 | 4.42E-06 | Lep-L |
| GMK NK | MAPKAPK2 | 3.22E-07 | 1.064134498 | 0.396 | 0.268 | 0.006342801 | Lep-L |
| GMK NK | CLEC2B | 1.58E-14 | 1.049245544 | 0.824 | 0.775 | 3.11E-10 | Lep-L |
| GMK NK | PER1 | 7.52E-08 | 1.045981121 | 0.392 | 0.258 | 0.001480135 | Lep-L |
| GMK NK | ATP5F1B | 1.27E-10 | 1.043384533 | 0.567 | 0.428 | 2.51E-06 | Lep-L |
| GMK NK | KPNA2 | 1.76E-10 | 1.04018885 | 0.494 | 0.318 | 3.46E-06 | Lep-L |
| GMK NK | HSPD1 | 1.06E-23 | 1.027951179 | 0.886 | 0.741 | 2.08E-19 | Lep-L |
| GMK NK | TUBB | 1.81E-08 | 1.023273612 | 0.506 | 0.381 | 0.000356817 | Lep-L |
| GMK NK | PNRC1 | 8.89E-17 | 1.018080257 | 0.902 | 0.857 | 1.75E-12 | Lep-L |
| GMK NK | PTGER4 | 2.97E-08 | 1.013415829 | 0.522 | 0.382 | 0.00058518 | Lep-L |
| GMK NK | HSP90AA1 | 5.49E-29 | 0.951267043 | 0.98 | 0.959 | 1.08E-24 | Lep-L |
| GMK NK | MCL1 | 8.73E-16 | 0.947927379 | 0.747 | 0.595 | 1.72E-11 | Lep-L |
| GMK NK | GABARAPL1 | 1.21E-06 | 0.947783556 | 0.527 | 0.422 | 0.02377444 | Lep-L |
| GMK NK | TUBA4A | 1.87E-10 | 0.941502057 | 0.714 | 0.604 | 3.69E-06 | Lep-L |
| GMK NK | HSPA1A | 4.00E-21 | 0.934539392 | 0.947 | 0.846 | 7.88E-17 | Lep-L |
| GMK NK | OAZ1 | 8.38E-13 | 0.932642342 | 0.82 | 0.854 | 1.65E-08 | Lep-L |
| GMK NK | FTH1 | 1.65E-06 | 0.92999931 | 0.976 | 0.982 | 0.032412747 | Lep-L |
| GMK NK | ARL6IP1 | 1.21E-07 | 0.916709182 | 0.588 | 0.487 | 0.002372432 | Lep-L |
| GMK NK | FTL | 1.33E-25 | 0.908652175 | 0.976 | 0.984 | 2.61E-21 | Lep-L |
| GMK NK | KDM6B | 5.86E-07 | 0.906856081 | 0.498 | 0.377 | 0.011530751 | Lep-L |
| GMK NK | ATP6V0C | 5.03E-08 | 0.881886904 | 0.629 | 0.554 | 0.000989571 | Lep-L |
| GMK NK | H2AFJ | 2.22E-06 | 0.84767379 | 0.539 | 0.435 | 0.043712353 | Lep-L |
| GMK NK | DNAJB1 | 1.28E-07 | 0.847279352 | 0.886 | 0.853 | 0.002519894 | Lep-L |
| GMK NK | MOB4 | 1.17E-08 | 0.839486641 | 0.563 | 0.442 | 0.000229591 | Lep-L |
| GMK NK | ZFAND2A | 2.29E-06 | 0.824418428 | 0.498 | 0.394 | 0.045114252 | Lep-L |
| GMK NK | PTGES3 | 1.98E-10 | 0.818910582 | 0.71 | 0.622 | 3.90E-06 | Lep-L |
| GMK NK | STX11 | 1.84E-06 | 0.809609555 | 0.449 | 0.329 | 0.03616064 | Lep-L |
| GMK NK | IFITM1 | 9.46E-07 | 0.801165804 | 0.698 | 0.631 | 0.018616298 | Lep-L |
| GMK NK | FAM177A1 | 1.29E-09 | 0.778293541 | 0.714 | 0.619 | 2.53E-05 | Lep-L |
| GMK NK | IFRD1 | 2.67E-07 | 0.774394199 | 0.592 | 0.492 | 0.005264359 | Lep-L |
| GMK NK | STMN1 | 2.52E-07 | 0.772995677 | 0.445 | 0.29 | 0.004959207 | Lep-L |
| GMK NK | JMJD6 | 4.18E-08 | 0.770202191 | 0.539 | 0.388 | 0.00082253 | Lep-L |
| GMK NK | SOD1 | 7.25E-09 | 0.761622126 | 0.702 | 0.608 | 0.000142682 | Lep-L |
| GMK NK | CEMIP2 | 2.06E-10 | 0.755601877 | 0.747 | 0.602 | 4.05E-06 | Lep-L |
| GMK NK | RUNX3 | 8.54E-10 | 0.752681327 | 0.763 | 0.659 | 1.68E-05 | Lep-L |
| GMK NK | TCP1 | 3.57E-07 | 0.751621547 | 0.555 | 0.45 | 0.007022112 | Lep-L |
| GMK NK | LITAF | 2.00E-07 | 0.750832062 | 0.665 | 0.563 | 0.003944586 | Lep-L |
| GMK NK | CACYBP | 7.58E-14 | 0.73860321 | 0.751 | 0.56 | 1.49E-09 | Lep-L |
| GMK NK | ARF6 | 1.00E-06 | 0.737120555 | 0.555 | 0.462 | 0.019763641 | Lep-L |
| GMK NK | JUND | 8.91E-29 | 0.717242714 | 0.976 | 0.942 | 1.75E-24 | Lep-L |
| GMK NK | LEPROTL1 | 1.64E-06 | 0.710708791 | 0.629 | 0.568 | 0.032262067 | Lep-L |
| GMK NK | SLC25A5 | 1.10E-07 | 0.705475558 | 0.735 | 0.647 | 0.002155737 | Lep-L |
| GMK NK | SLC3A2 | 2.18E-06 | 0.702396773 | 0.571 | 0.465 | 0.042903787 | Lep-L |
| GMK NK | MAP1LC3B | 1.14E-10 | 0.699426101 | 0.755 | 0.643 | 2.25E-06 | Lep-L |
| GMK NK | YWHAZ | 1.91E-07 | 0.690071548 | 0.767 | 0.738 | 0.00375359 | Lep-L |
| GMK NK | DNAJA1 | 8.61E-13 | 0.673249871 | 0.918 | 0.894 | 1.70E-08 | Lep-L |

| | | | | | | | |
|---|---|---|---|---|---|---|---|
| GMK NK | UBC | 7.35E-09 | 0.665938794 | 0.931 | 0.936 | 0.000144732 | Lep-L |
| GMK NK | DNAJB6 | 1.87E-08 | 0.663098688 | 0.804 | 0.725 | 0.000367966 | Lep-L |
| GMK NK | HSPA5 | 1.07E-06 | 0.652170629 | 0.661 | 0.601 | 0.021143771 | Lep-L |
| GMK NK | SELENOK | 2.80E-09 | 0.648676268 | 0.759 | 0.681 | 5.50E-05 | Lep-L |
| GMK NK | TUBA1B | 7.17E-07 | 0.617328532 | 0.702 | 0.648 | 0.014107363 | Lep-L |
| GMK NK | SFPQ | 4.45E-07 | 0.616765938 | 0.706 | 0.624 | 0.008752947 | Lep-L |
| GMK NK | RAB8B | 2.27E-06 | 0.603247087 | 0.718 | 0.679 | 0.044707052 | Lep-L |
| GMK NK | CCNH | 2.37E-06 | 0.599138259 | 0.673 | 0.592 | 0.046736557 | Lep-L |
| GMK NK | SQSTM1 | 2.42E-08 | 0.594352451 | 0.833 | 0.763 | 0.000475977 | Lep-L |
| GMK NK | RBM39 | 2.35E-08 | 0.579726191 | 0.816 | 0.767 | 0.000463426 | Lep-L |
| GMK NK | UBE2D3 | 6.63E-09 | 0.569407966 | 0.747 | 0.671 | 0.000130538 | Lep-L |
| GMK NK | PABPC1 | 2.97E-10 | 0.564340498 | 0.906 | 0.845 | 5.85E-06 | Lep-L |
| GMK NK | H2AFZ | 4.41E-09 | 0.562952118 | 0.853 | 0.768 | 8.69E-05 | Lep-L |
| GMK NK | REL | 1.96E-08 | 0.553000124 | 0.853 | 0.807 | 0.000385983 | Lep-L |
| GMK NK | DUSP2 | 1.92E-10 | 0.532651101 | 0.959 | 0.895 | 3.79E-06 | Lep-L |
| GMK NK | EIF4A1 | 2.04E-07 | 0.50347212 | 0.841 | 0.803 | 0.004010371 | Lep-L |
| GMK NK | XCL1 | 4.45E-20 | 1.651408147 | 0.834 | 0.675 | 8.76E-16 | Lep-T |
| GMK NK | CD74 | 1.57E-11 | 1.410376971 | 0.841 | 0.711 | 3.08E-07 | Lep-T |
| GMK NK | IFITM11 | 5.13E-09 | 1.391104277 | 0.716 | 0.621 | 0.000100958 | Lep-T |
| GMK NK | ALDOA1 | 3.84E-10 | 1.3416772 | 0.275 | 0.135 | 7.57E-06 | Lep-T |
| GMK NK | HIST1H1E | 1.07E-18 | 1.311404714 | 0.409 | 0.17 | 2.11E-14 | Lep-T |
| GMK NK | EPG5 | 4.84E-14 | 1.303556865 | 0.291 | 0.116 | 9.52E-10 | Lep-T |
| GMK NK | ASAP1 | 1.71E-09 | 1.251396092 | 0.428 | 0.267 | 3.36E-05 | Lep-T |
| GMK NK | RBFOX21 | 1.58E-13 | 1.240346472 | 0.509 | 0.304 | 3.12E-09 | Lep-T |
| GMK NK | HIST1H1C | 1.72E-10 | 1.224404618 | 0.331 | 0.17 | 3.38E-06 | Lep-T |
| GMK NK | CST3 | 1.17E-12 | 1.200817809 | 0.269 | 0.106 | 2.30E-08 | Lep-T |
| GMK NK | IL12RB2 | 5.99E-13 | 1.18841548 | 0.475 | 0.289 | 1.18E-08 | Lep-T |
| GMK NK | TNFSF9 | 8.50E-07 | 1.183639686 | 0.3 | 0.177 | 0.01673321 | Lep-T |
| GMK NK | 3-Mar | 5.37E-08 | 1.17775341 | 0.372 | 0.225 | 0.001057287 | Lep-T |
| GMK NK | NIBAN1 | 9.09E-13 | 1.155780686 | 0.406 | 0.213 | 1.79E-08 | Lep-T |
| GMK NK | CRTAM | 2.55E-11 | 1.129586038 | 0.453 | 0.278 | 5.03E-07 | Lep-T |
| GMK NK | MT-CO3 | 4.67E-11 | 1.120977799 | 0.822 | 0.875 | 9.18E-07 | Lep-T |
| GMK NK | GNG2 | 1.82E-12 | 1.104328012 | 0.634 | 0.486 | 3.58E-08 | Lep-T |
| GMK NK | NOCT | 2.48E-10 | 1.090542427 | 0.262 | 0.116 | 4.88E-06 | Lep-T |
| GMK NK | EMD1 | 9.34E-12 | 1.081786463 | 0.312 | 0.143 | 1.84E-07 | Lep-T |
| GMK NK | CCL4L21 | 6.00E-11 | 1.06280748 | 0.506 | 0.313 | 1.18E-06 | Lep-T |
| GMK NK | TFDP2 | 4.88E-09 | 1.060999542 | 0.35 | 0.194 | 9.61E-05 | Lep-T |
| GMK NK | HLA-DQA2 | 3.78E-16 | 1.031298838 | 0.25 | 0.076 | 7.43E-12 | Lep-T |
| GMK NK | BHLHE40 | 3.08E-17 | 1.005454667 | 0.659 | 0.419 | 6.07E-13 | Lep-T |
| GMK NK | PRKX | 6.48E-10 | 0.990933074 | 0.444 | 0.273 | 1.28E-05 | Lep-T |
| GMK NK | SIPA1L1 | 1.63E-08 | 0.987504741 | 0.469 | 0.318 | 0.000320206 | Lep-T |
| GMK NK | BCL31 | 1.83E-09 | 0.982336915 | 0.319 | 0.164 | 3.60E-05 | Lep-T |
| GMK NK | ITK | 1.40E-10 | 0.976152795 | 0.391 | 0.214 | 2.76E-06 | Lep-T |
| GMK NK | USP36 | 1.73E-11 | 0.956166793 | 0.366 | 0.18 | 3.40E-07 | Lep-T |
| GMK NK | NFKBIZ | 1.02E-11 | 0.941454773 | 0.575 | 0.39 | 2.00E-07 | Lep-T |
| GMK NK | XCL2 | 4.58E-11 | 0.939860465 | 0.834 | 0.77 | 9.02E-07 | Lep-T |
| GMK NK | IFITM3 | 4.40E-11 | 0.920402362 | 0.559 | 0.349 | 8.66E-07 | Lep-T |
| GMK NK | PFKFB3 | 6.31E-10 | 0.901656205 | 0.403 | 0.22 | 1.24E-05 | Lep-T |
| GMK NK | CCL4 | 4.90E-07 | 0.887616831 | 0.759 | 0.695 | 0.009643479 | Lep-T |
| GMK NK | AREG | 7.08E-09 | 0.881746057 | 0.716 | 0.589 | 0.000139302 | Lep-T |
| GMK NK | EML4 | 8.54E-14 | 0.880867417 | 0.697 | 0.499 | 1.68E-09 | Lep-T |
| GMK NK | ELL2 | 3.47E-10 | 0.878517416 | 0.584 | 0.419 | 6.82E-06 | Lep-T |
| GMK NK | FOXP1 | 2.01E-07 | 0.872671108 | 0.494 | 0.351 | 0.003963944 | Lep-T |
| GMK NK | TANK | 3.95E-08 | 0.868210993 | 0.466 | 0.316 | 0.000778185 | Lep-T |
| GMK NK | VAV3 | 3.37E-09 | 0.857819551 | 0.462 | 0.295 | 6.64E-05 | Lep-T |
| GMK NK | FBXO34 | 1.60E-09 | 0.856089932 | 0.503 | 0.328 | 3.16E-05 | Lep-T |

| | | | | | | | |
|---|---|---|---|---|---|---|---|
| GMK NK | GALC | 7.20E-08 | 0.850740156 | 0.259 | 0.128 | 0.001417576 | Lep-T |
| GMK NK | OSBPL3 | 5.62E-09 | 0.843420737 | 0.338 | 0.181 | 0.000110706 | Lep-T |
| GMK NK | IFITM2 | 6.82E-13 | 0.830840435 | 0.906 | 0.798 | 1.34E-08 | Lep-T |
| GMK NK | DPP8 | 3.59E-07 | 0.827517856 | 0.269 | 0.144 | 0.007056995 | Lep-T |
| GMK NK | FOXO3 | 2.17E-06 | 0.824719487 | 0.334 | 0.201 | 0.042707911 | Lep-T |
| GMK NK | PHLDA1 | 6.66E-10 | 0.807755091 | 0.428 | 0.244 | 1.31E-05 | Lep-T |
| GMK NK | PDE4D | 6.06E-10 | 0.800672512 | 0.438 | 0.248 | 1.19E-05 | Lep-T |
| GMK NK | TENT4B | 2.41E-08 | 0.797936203 | 0.359 | 0.202 | 0.000473985 | Lep-T |
| GMK NK | DAZAP2 | 2.32E-07 | 0.792893941 | 0.378 | 0.224 | 0.004574081 | Lep-T |
| GMK NK | RELB | 3.86E-10 | 0.790601764 | 0.475 | 0.283 | 7.59E-06 | Lep-T |
| GMK NK | STK39 | 2.17E-07 | 0.783605188 | 0.344 | 0.2 | 0.004279679 | Lep-T |
| GMK NK | CXCR4 | 5.76E-15 | 0.778924748 | 0.85 | 0.716 | 1.13E-10 | Lep-T |
| GMK NK | DUSP10 | 1.11E-09 | 0.765374601 | 0.397 | 0.22 | 2.18E-05 | Lep-T |
| GMK NK | MT-ND1 | 8.17E-12 | 0.760671797 | 0.791 | 0.758 | 1.61E-07 | Lep-T |
| GMK NK | OTULIN | 2.45E-06 | 0.750072551 | 0.447 | 0.308 | 0.048142404 | Lep-T |
| GMK NK | ZEB2 | 1.55E-08 | 0.741451988 | 0.547 | 0.388 | 0.000304777 | Lep-T |
| GMK NK | IQGAP2 | 6.96E-07 | 0.734017416 | 0.547 | 0.41 | 0.013706275 | Lep-T |
| GMK NK | TLK2 | 6.02E-07 | 0.732525733 | 0.25 | 0.129 | 0.011848603 | Lep-T |
| GMK NK | NFKB1 | 1.09E-12 | 0.731400641 | 0.794 | 0.633 | 2.14E-08 | Lep-T |
| GMK NK | KLF6 | 1.61E-09 | 0.731109066 | 0.781 | 0.634 | 3.17E-05 | Lep-T |
| GMK NK | HIPK1 | 6.43E-08 | 0.725631894 | 0.362 | 0.204 | 0.001264801 | Lep-T |
| GMK NK | CSRNP11 | 2.75E-07 | 0.715381444 | 0.525 | 0.373 | 0.005419635 | Lep-T |
| GMK NK | CUL3 | 5.71E-07 | 0.71272981 | 0.453 | 0.3 | 0.01124347 | Lep-T |
| GMK NK | RASGRP1 | 1.24E-07 | 0.703665463 | 0.331 | 0.183 | 0.002446463 | Lep-T |
| GMK NK | ABL1 | 8.42E-07 | 0.703128168 | 0.275 | 0.15 | 0.016576537 | Lep-T |
| GMK NK | DUSP16 | 1.85E-07 | 0.695598315 | 0.278 | 0.147 | 0.003636815 | Lep-T |
| GMK NK | FYB1 | 1.87E-11 | 0.690269087 | 0.616 | 0.409 | 3.68E-07 | Lep-T |
| GMK NK | MAFF | 4.14E-07 | 0.687408168 | 0.478 | 0.328 | 0.008149866 | Lep-T |
| GMK NK | RHOH1 | 9.84E-11 | 0.684837264 | 0.591 | 0.384 | 1.94E-06 | Lep-T |
| GMK NK | PDE7A | 1.43E-06 | 0.671988717 | 0.644 | 0.529 | 0.028224777 | Lep-T |
| GMK NK | TNFAIP31 | 1.95E-16 | 0.660976425 | 0.866 | 0.729 | 3.84E-12 | Lep-T |
| GMK NK | NUP98 | 1.01E-06 | 0.659050572 | 0.35 | 0.206 | 0.019946061 | Lep-T |
| GMK NK | YES1 | 1.07E-06 | 0.657058204 | 0.459 | 0.309 | 0.021090348 | Lep-T |
| GMK NK | ICAM11 | 6.20E-07 | 0.648664164 | 0.397 | 0.244 | 0.012200671 | Lep-T |
| GMK NK | FNBP1 | 1.53E-06 | 0.634094288 | 0.606 | 0.485 | 0.030192556 | Lep-T |
| GMK NK | MAP3K8 | 1.92E-06 | 0.627116636 | 0.659 | 0.529 | 0.037876937 | Lep-T |
| GMK NK | GPCPD1 | 6.44E-07 | 0.624031356 | 0.478 | 0.319 | 0.012684762 | Lep-T |
| GMK NK | PNRC11 | 8.84E-14 | 0.623902011 | 0.95 | 0.84 | 1.74E-09 | Lep-T |
| GMK NK | ARRDC3 | 2.30E-06 | 0.600712366 | 0.409 | 0.264 | 0.045302808 | Lep-T |
| GMK NK | IL21R1 | 2.19E-07 | 0.599819888 | 0.325 | 0.183 | 0.004302488 | Lep-T |
| GMK NK | SYTL3 | 1.67E-07 | 0.590061212 | 0.678 | 0.562 | 0.003290392 | Lep-T |
| GMK NK | SCML4 | 5.48E-07 | 0.586678372 | 0.394 | 0.24 | 0.010786526 | Lep-T |
| GMK NK | ATAD2B | 1.71E-07 | 0.584934977 | 0.303 | 0.161 | 0.003367767 | Lep-T |
| GMK NK | SPATA13 | 3.50E-07 | 0.579548135 | 0.331 | 0.185 | 0.006894175 | Lep-T |
| GMK NK | LITAF1 | 2.43E-10 | 0.571368959 | 0.731 | 0.538 | 4.79E-06 | Lep-T |
| GMK NK | TSC22D3 | 9.33E-07 | 0.564732166 | 0.753 | 0.625 | 0.018356133 | Lep-T |
| GMK NK | ERN1 | 9.02E-07 | 0.559815337 | 0.5 | 0.334 | 0.017763742 | Lep-T |
| GMK NK | MTDH | 1.65E-06 | 0.547622075 | 0.531 | 0.367 | 0.032438258 | Lep-T |
| GMK NK | SLC2A31 | 2.87E-12 | 0.546065025 | 0.713 | 0.513 | 5.65E-08 | Lep-T |
| GMK NK | SMAP2 | 3.34E-07 | 0.542761692 | 0.591 | 0.438 | 0.006572767 | Lep-T |
| GMK NK | UST | 5.66E-07 | 0.541886149 | 0.297 | 0.16 | 0.011136259 | Lep-T |
| GMK NK | FAM177A11 | 4.98E-10 | 0.537155805 | 0.775 | 0.596 | 9.81E-06 | Lep-T |
| GMK NK | TIPARP1 | 6.70E-08 | 0.533673245 | 0.516 | 0.341 | 0.001318691 | Lep-T |
| GMK NK | YPEL5 | 3.29E-07 | 0.531297546 | 0.722 | 0.565 | 0.006484828 | Lep-T |
| GMK NK | EEF21 | 1.86E-07 | 0.530926376 | 0.719 | 0.551 | 0.003652617 | Lep-T |
| GMK NK | SH3KBP1 | 2.34E-06 | 0.530618525 | 0.484 | 0.332 | 0.046126261 | Lep-T |

| | | | | | | | |
|---|---|---|---|---|---|---|---|
| GMK NK | PRRC2C | 1.58E-09 | 0.528171897 | 0.722 | 0.541 | 3.12E-05 | Lep-T |
| GMK NK | CMC1 | 1.04E-06 | 0.523650571 | 0.653 | 0.514 | 0.020568978 | Lep-T |
| GMK NK | SRGN | 4.03E-11 | 0.523349935 | 0.938 | 0.948 | 7.94E-07 | Lep-T |
| GMK NK | MT-CYB | 5.45E-08 | 0.522200113 | 0.803 | 0.826 | 0.001073188 | Lep-T |
| GMK NK | REL1 | 3.00E-14 | 0.513487523 | 0.925 | 0.783 | 5.90E-10 | Lep-T |
| GMK NK | IRF2BP2 | 5.90E-08 | 0.508269929 | 0.4 | 0.228 | 0.001160975 | Lep-T |
| GMK NK | SEC11C | 1.70E-11 | 1.161347966 | 0.336 | 0.176 | 3.35E-07 | Mari |
| GMK NK | GNLY | 4.61E-23 | 1.092062844 | 0.824 | 0.653 | 9.08E-19 | Mari |
| GMK NK | GZMA | 1.23E-14 | 1.072876305 | 0.801 | 0.739 | 2.42E-10 | Mari |
| GMK NK | MICOS13 | 5.57E-09 | 1.062627152 | 0.35 | 0.214 | 0.000109664 | Mari |
| GMK NK | NDUFS7 | 3.30E-08 | 1.058565054 | 0.294 | 0.172 | 0.000649124 | Mari |
| GMK NK | TMIGD2 | 1.95E-06 | 1.046194043 | 0.325 | 0.224 | 0.038461478 | Mari |
| GMK NK | TIMM8B | 4.51E-08 | 1.032758987 | 0.252 | 0.134 | 0.000887246 | Mari |
| GMK NK | TMBIM4 | 3.14E-11 | 1.031267362 | 0.429 | 0.269 | 6.17E-07 | Mari |
| GMK NK | LAMTOR1 | 4.11E-08 | 1.007380614 | 0.333 | 0.21 | 0.000809541 | Mari |
| GMK NK | NDUFAB1 | 1.69E-11 | 0.990573058 | 0.401 | 0.236 | 3.32E-07 | Mari |
| GMK NK | CAPN12 | 5.29E-08 | 0.969560946 | 0.345 | 0.222 | 0.001040948 | Mari |
| GMK NK | SMIM19 | 5.24E-07 | 0.969230426 | 0.263 | 0.153 | 0.010307852 | Mari |
| GMK NK | PIN1 | 7.96E-07 | 0.965416285 | 0.269 | 0.159 | 0.015664542 | Mari |
| GMK NK | RBIS | 1.85E-10 | 0.941396514 | 0.468 | 0.321 | 3.63E-06 | Mari |
| GMK NK | COPS9 | 1.91E-10 | 0.941183546 | 0.429 | 0.278 | 3.75E-06 | Mari |
| GMK NK | PET100 | 1.20E-13 | 0.938335408 | 0.451 | 0.256 | 2.35E-09 | Mari |
| GMK NK | KLRF1 | 3.63E-13 | 0.926248689 | 0.653 | 0.503 | 7.14E-09 | Mari |
| GMK NK | SEC61B | 4.61E-12 | 0.916934614 | 0.591 | 0.46 | 9.07E-08 | Mari |
| GMK NK | LSM4 | 1.04E-07 | 0.908480408 | 0.342 | 0.214 | 0.002046411 | Mari |
| GMK NK | LSM6 | 1.92E-06 | 0.907859578 | 0.28 | 0.174 | 0.037706946 | Mari |
| GMK NK | MRPL52 | 2.69E-08 | 0.905733881 | 0.37 | 0.24 | 0.000529845 | Mari |
| GMK NK | STMP1 | 4.92E-07 | 0.90485411 | 0.294 | 0.179 | 0.009687628 | Mari |
| GMK NK | CD52 | 9.53E-10 | 0.899960045 | 0.49 | 0.344 | 1.88E-05 | Mari |
| GMK NK | TMED9 | 1.26E-06 | 0.897182343 | 0.317 | 0.21 | 0.024736615 | Mari |
| GMK NK | TOMM5 | 1.24E-06 | 0.873827638 | 0.347 | 0.234 | 0.024389338 | Mari |
| GMK NK | OSTC | 3.01E-08 | 0.871454498 | 0.401 | 0.265 | 0.000591576 | Mari |
| GMK NK | PLAC8 | 4.02E-08 | 0.856592962 | 0.398 | 0.261 | 0.000790818 | Mari |
| GMK NK | NOP10 | 3.58E-09 | 0.848326185 | 0.431 | 0.29 | 7.05E-05 | Mari |
| GMK NK | NDUFB5 | 9.69E-07 | 0.847970474 | 0.266 | 0.154 | 0.019080465 | Mari |
| GMK NK | SEC61G | 3.64E-11 | 0.847554876 | 0.552 | 0.398 | 7.17E-07 | Mari |
| GMK NK | TSTD1 | 1.12E-08 | 0.845992677 | 0.409 | 0.273 | 0.000219827 | Mari |
| GMK NK | UQCR10 | 4.13E-11 | 0.832619104 | 0.493 | 0.328 | 8.13E-07 | Mari |
| GMK NK | TRDC1 | 3.62E-13 | 0.824856708 | 0.742 | 0.62 | 7.13E-09 | Mari |
| GMK NK | ERH | 1.11E-09 | 0.810470985 | 0.482 | 0.332 | 2.19E-05 | Mari |
| GMK NK | BLOC1S1 | 6.87E-08 | 0.80873818 | 0.426 | 0.299 | 0.001351613 | Mari |
| GMK NK | ATP5MPL | 1.41E-06 | 0.803744473 | 0.356 | 0.245 | 0.027777985 | Mari |
| GMK NK | PDCD6 | 1.81E-06 | 0.802887946 | 0.37 | 0.266 | 0.035697717 | Mari |
| GMK NK | POLR2L | 2.77E-11 | 0.800106798 | 0.574 | 0.412 | 5.45E-07 | Mari |
| GMK NK | NDUFS6 | 2.45E-07 | 0.796483171 | 0.423 | 0.301 | 0.004830837 | Mari |
| GMK NK | ATP5ME | 6.33E-09 | 0.795375318 | 0.504 | 0.363 | 0.00012457 | Mari |
| GMK NK | PSMB1 | 5.74E-14 | 0.789244872 | 0.627 | 0.452 | 1.13E-09 | Mari |
| GMK NK | COX7A2 | 1.09E-16 | 0.789207096 | 0.686 | 0.497 | 2.15E-12 | Mari |
| GMK NK | RBX1 | 5.23E-09 | 0.78633806 | 0.465 | 0.326 | 0.000102993 | Mari |
| GMK NK | PSMB9 | 2.31E-15 | 0.781845388 | 0.72 | 0.546 | 4.55E-11 | Mari |
| GMK NK | DGUOK | 6.67E-08 | 0.781627253 | 0.415 | 0.278 | 0.001313825 | Mari |
| GMK NK | REX1BD | 6.13E-08 | 0.781558535 | 0.429 | 0.301 | 0.001207241 | Mari |
| GMK NK | NDUFB7 | 3.68E-07 | 0.780175045 | 0.392 | 0.268 | 0.007239943 | Mari |
| GMK NK | TRAPPC1 | 2.39E-06 | 0.777172082 | 0.37 | 0.262 | 0.046978533 | Mari |
| GMK NK | MICOS10 | 1.79E-10 | 0.776819583 | 0.468 | 0.304 | 3.51E-06 | Mari |
| GMK NK | NDUFC2 | 1.11E-06 | 0.774708655 | 0.398 | 0.29 | 0.021927387 | Mari |

| | | | | | | | |
|---|---|---|---|---|---|---|---|
| GMK NK | TOMM6 | 4.40E-09 | 0.774142317 | 0.457 | 0.306 | 8.67E-05 | Mari |
| GMK NK | COX5B | 3.35E-12 | 0.769481476 | 0.583 | 0.415 | 6.59E-08 | Mari |
| GMK NK | ATP5MF | 9.07E-09 | 0.760575477 | 0.457 | 0.319 | 0.000178477 | Mari |
| GMK NK | COX6B1 | 2.57E-11 | 0.760139675 | 0.613 | 0.476 | 5.05E-07 | Mari |
| GMK NK | ATP5PF | 2.97E-09 | 0.757165338 | 0.527 | 0.398 | 5.84E-05 | Mari |
| GMK NK | SELENOH | 2.90E-09 | 0.75166916 | 0.535 | 0.391 | 5.71E-05 | Mari |
| GMK NK | COX7A2L | 4.72E-07 | 0.749131117 | 0.423 | 0.31 | 0.00929474 | Mari |
| GMK NK | DAD1 | 2.36E-07 | 0.746647655 | 0.482 | 0.361 | 0.004653793 | Mari |
| GMK NK | ATP5MD | 4.63E-09 | 0.746157446 | 0.524 | 0.393 | 9.12E-05 | Mari |
| GMK NK | HOPX | 9.29E-12 | 0.744758442 | 0.717 | 0.546 | 1.83E-07 | Mari |
| GMK NK | LMAN2 | 1.72E-06 | 0.739352321 | 0.381 | 0.267 | 0.033918705 | Mari |
| GMK NK | MED10 | 5.05E-07 | 0.732510069 | 0.392 | 0.275 | 0.009945582 | Mari |
| GMK NK | ARPC1B | 2.64E-09 | 0.727120821 | 0.571 | 0.432 | 5.19E-05 | Mari |
| GMK NK | GZMM | 1.06E-07 | 0.720107284 | 0.625 | 0.515 | 0.002093154 | Mari |
| GMK NK | NDUFB2 | 4.18E-09 | 0.719610029 | 0.552 | 0.426 | 8.24E-05 | Mari |
| GMK NK | SPCS2 | 3.28E-09 | 0.716980763 | 0.51 | 0.353 | 6.45E-05 | Mari |
| GMK NK | ELOB | 6.90E-12 | 0.715803796 | 0.667 | 0.531 | 1.36E-07 | Mari |
| GMK NK | TRBC2 | 5.37E-08 | 0.714182701 | 0.566 | 0.438 | 0.001057902 | Mari |
| GMK NK | TBC1D10C | 4.33E-07 | 0.702139696 | 0.417 | 0.29 | 0.008517861 | Mari |
| GMK NK | SEC11A | 2.60E-08 | 0.700532883 | 0.58 | 0.456 | 0.000510935 | Mari |
| GMK NK | SPCS1 | 1.86E-07 | 0.695841012 | 0.457 | 0.328 | 0.003660223 | Mari |
| GMK NK | PPIB | 7.71E-11 | 0.695588941 | 0.658 | 0.519 | 1.52E-06 | Mari |
| GMK NK | EIF5A | 2.29E-06 | 0.694274331 | 0.482 | 0.381 | 0.045090839 | Mari |
| GMK NK | NDUFA12 | 2.36E-06 | 0.690935238 | 0.415 | 0.31 | 0.04648562 | Mari |
| GMK NK | BST2 | 2.31E-09 | 0.689382147 | 0.577 | 0.445 | 4.55E-05 | Mari |
| GMK NK | PRELID1 | 4.84E-08 | 0.68822928 | 0.563 | 0.445 | 0.000952971 | Mari |
| GMK NK | COX8A | 9.23E-14 | 0.687817042 | 0.63 | 0.428 | 1.82E-09 | Mari |
| GMK NK | APRT | 8.59E-09 | 0.687632406 | 0.613 | 0.494 | 0.0001691 | Mari |
| GMK NK | VPS28 | 8.82E-07 | 0.687329086 | 0.443 | 0.329 | 0.017359138 | Mari |
| GMK NK | PAIP2 | 1.10E-07 | 0.681046905 | 0.493 | 0.358 | 0.002167445 | Mari |
| GMK NK | SF3B6 | 4.49E-10 | 0.680767026 | 0.532 | 0.373 | 8.84E-06 | Mari |
| GMK NK | EIF3G | 7.51E-11 | 0.676809298 | 0.765 | 0.634 | 1.48E-06 | Mari |
| GMK NK | OCIAD2 | 2.39E-07 | 0.674572245 | 0.443 | 0.32 | 0.004704682 | Mari |
| GMK NK | TMA7 | 2.99E-17 | 0.673755978 | 0.849 | 0.697 | 5.88E-13 | Mari |
| GMK NK | MYL12A | 1.60E-17 | 0.671725541 | 0.896 | 0.792 | 3.15E-13 | Mari |
| GMK NK | NDUFA13 | 2.05E-09 | 0.668888648 | 0.625 | 0.488 | 4.04E-05 | Mari |
| GMK NK | CKLF | 1.22E-07 | 0.665899225 | 0.591 | 0.474 | 0.002395858 | Mari |
| GMK NK | SERPINB1 | 1.53E-06 | 0.665688469 | 0.429 | 0.309 | 0.030099252 | Mari |
| GMK NK | TMEM258 | 3.45E-07 | 0.665594884 | 0.454 | 0.33 | 0.00679525 | Mari |
| GMK NK | REEP5 | 1.18E-06 | 0.647040196 | 0.471 | 0.346 | 0.023275012 | Mari |
| GMK NK | GMFG | 2.93E-07 | 0.632886928 | 0.588 | 0.481 | 0.005760415 | Mari |
| GMK NK | LAMTOR4 | 2.02E-06 | 0.624331708 | 0.476 | 0.367 | 0.039683436 | Mari |
| GMK NK | SH3BGRL3 | 1.29E-17 | 0.621739053 | 0.902 | 0.789 | 2.53E-13 | Mari |
| GMK NK | LIMD2 | 1.91E-06 | 0.618681765 | 0.56 | 0.454 | 0.037506066 | Mari |
| GMK NK | UQCR11 | 3.64E-09 | 0.610695881 | 0.591 | 0.435 | 7.16E-05 | Mari |
| GMK NK | COPE | 7.36E-07 | 0.608879032 | 0.515 | 0.41 | 0.014489364 | Mari |
| GMK NK | ARPC3 | 1.06E-13 | 0.608837067 | 0.838 | 0.7 | 2.09E-09 | Mari |
| GMK NK | SRSF9 | 4.67E-08 | 0.606088183 | 0.563 | 0.432 | 0.000918415 | Mari |
| GMK NK | NDUFA4 | 1.65E-10 | 0.601754563 | 0.695 | 0.567 | 3.26E-06 | Mari |
| GMK NK | PSMA1 | 3.41E-07 | 0.597264061 | 0.482 | 0.347 | 0.006706683 | Mari |
| GMK NK | ITM2B | 1.04E-13 | 0.595188297 | 0.868 | 0.753 | 2.05E-09 | Mari |
| GMK NK | CFL1 | 7.02E-19 | 0.592121864 | 0.922 | 0.844 | 1.38E-14 | Mari |
| GMK NK | HMGN1 | 1.71E-06 | 0.580245714 | 0.591 | 0.507 | 0.03365073 | Mari |
| GMK NK | CD7 | 1.02E-11 | 0.579734886 | 0.854 | 0.767 | 2.02E-07 | Mari |
| GMK NK | APOBEC3G | 6.88E-07 | 0.578611307 | 0.669 | 0.556 | 0.013539769 | Mari |
| GMK NK | SYF2 | 7.34E-07 | 0.569011065 | 0.524 | 0.384 | 0.014441379 | Mari |

| | | | | | | | |
|---|---|---|---|---|---|---|---|
| GMK NK | SSR4 | 9.69E-07 | 0.568789889 | 0.641 | 0.565 | 0.019064819 | Mari |
| GMK NK | SERF2 | 2.18E-13 | 0.56766421 | 0.894 | 0.802 | 4.29E-09 | Mari |
| GMK NK | SNHG9 | 1.56E-06 | 0.566623831 | 0.347 | 0.221 | 0.030688689 | Mari |
| GMK NK | TRMT112 | 3.35E-08 | 0.566442988 | 0.636 | 0.508 | 0.000659561 | Mari |
| GMK NK | S100A11 | 2.29E-10 | 0.564049605 | 0.697 | 0.552 | 4.51E-06 | Mari |
| GMK NK | ATP6V1G1 | 3.25E-07 | 0.559942716 | 0.555 | 0.437 | 0.006392802 | Mari |
| GMK NK | MYL12B | 8.34E-13 | 0.554947092 | 0.804 | 0.676 | 1.64E-08 | Mari |
| GMK NK | NME2 | 6.86E-08 | 0.54961964 | 0.748 | 0.642 | 0.00134978 | Mari |
| GMK NK | EIF3K | 2.97E-09 | 0.546120151 | 0.765 | 0.663 | 5.84E-05 | Mari |
| GMK NK | FCER1G | 1.86E-07 | 0.543338991 | 0.728 | 0.625 | 0.003656116 | Mari |
| GMK NK | CYBA | 2.29E-10 | 0.542146474 | 0.86 | 0.776 | 4.51E-06 | Mari |
| GMK NK | UBL5 | 3.37E-11 | 0.534708659 | 0.751 | 0.608 | 6.64E-07 | Mari |
| GMK NK | ANAPC16 | 1.65E-06 | 0.527996355 | 0.56 | 0.441 | 0.032385726 | Mari |
| GMK NK | CCDC85B | 6.60E-07 | 0.523607875 | 0.571 | 0.438 | 0.01299082 | Mari |
| GMK NK | EDF1 | 1.54E-07 | 0.52063867 | 0.697 | 0.612 | 0.003036301 | Mari |
| GMK NK | ATP5F1D | 1.39E-06 | 0.511491629 | 0.58 | 0.472 | 0.027414316 | Mari |
| GMK NK | COX6C | 1.47E-07 | 0.509355665 | 0.641 | 0.528 | 0.002895304 | Mari |
| GMK NK | CORO1A | 9.04E-08 | 0.501831963 | 0.779 | 0.686 | 0.001779823 | Mari |
| GMK NK | IGKC | 2.68E-15 | 2.461130405 | 0.26 | 0.114 | 5.27E-11 | Tb |
| GMK NK | SLC35F1 | 1.39E-46 | 2.436920009 | 0.384 | 0.078 | 2.74E-42 | Tb |
| GMK NK | ID1 | 2.71E-15 | 1.488129551 | 0.27 | 0.112 | 5.34E-11 | Tb |
| GMK NK | S100A9 | 3.34E-15 | 1.318037636 | 0.321 | 0.152 | 6.58E-11 | Tb |
| GMK NK | S100A8 | 8.02E-12 | 1.173792006 | 0.289 | 0.143 | 1.58E-07 | Tb |
| GMK NK | HIP1 | 9.36E-08 | 1.115964126 | 0.293 | 0.181 | 0.001842595 | Tb |
| GMK NK | KRT14 | 5.66E-09 | 1.091318072 | 0.252 | 0.133 | 0.000111327 | Tb |
| GMK NK | NFE2L3 | 1.22E-07 | 1.040972716 | 0.342 | 0.23 | 0.002401691 | Tb |
| GMK NK | SMURF2 | 5.91E-09 | 0.931083495 | 0.392 | 0.261 | 0.000116364 | Tb |
| GMK NK | MT-ND3 | 1.09E-38 | 0.926761668 | 0.966 | 0.839 | 2.14E-34 | Tb |
| GMK NK | XIST | 1.28E-08 | 0.902930599 | 0.327 | 0.2 | 0.000251367 | Tb |
| GMK NK | LYZ | 5.71E-07 | 0.875572839 | 0.27 | 0.167 | 0.011235883 | Tb |
| GMK NK | GLCCI1 | 4.49E-07 | 0.789754698 | 0.426 | 0.311 | 0.008843365 | Tb |
| GMK NK | HNRNPH1 | 1.10E-10 | 0.769966249 | 0.604 | 0.45 | 2.16E-06 | Tb |
| GMK NK | ZNF638 | 7.73E-07 | 0.763295143 | 0.337 | 0.223 | 0.015224307 | Tb |
| GMK NK | MT-ATP6 | 2.26E-28 | 0.728505046 | 0.977 | 0.821 | 4.46E-24 | Tb |
| GMK NK | TNFAIP8 | 6.59E-07 | 0.713144431 | 0.434 | 0.313 | 0.012966013 | Tb |
| GMK NK | ZBTB20 | 1.86E-06 | 0.701089099 | 0.512 | 0.41 | 0.03660535 | Tb |
| GMK NK | ZNF10 | 2.98E-07 | 0.70097631 | 0.317 | 0.197 | 0.005863911 | Tb |
| GMK NK | MT-ND4 | 1.72E-24 | 0.659110308 | 0.95 | 0.826 | 3.39E-20 | Tb |
| GMK NK | ARIH1 | 7.86E-08 | 0.646775359 | 0.564 | 0.437 | 0.001547888 | Tb |
| GMK NK | MT-CYB1 | 3.20E-12 | 0.639188053 | 0.88 | 0.787 | 6.31E-08 | Tb |
| GMK NK | PPP1R14B | 1.83E-06 | 0.622155132 | 0.457 | 0.341 | 0.036076339 | Tb |
| GMK NK | MT-ND2 | 5.61E-15 | 0.618459193 | 0.901 | 0.765 | 1.10E-10 | Tb |
| GMK NK | TXK | 2.52E-08 | 0.587175544 | 0.564 | 0.413 | 0.000496625 | Tb |
| GMK NK | FOSB | 6.60E-07 | 0.580318311 | 0.568 | 0.458 | 0.012999865 | Tb |
| GMK NK | MT-ATP8 | 1.29E-09 | 0.574344365 | 0.663 | 0.537 | 2.53E-05 | Tb |
| GMK NK | RAB11A | 2.03E-06 | 0.557936346 | 0.551 | 0.428 | 0.039981497 | Tb |
| GMK NK | MT-ND11 | 1.11E-15 | 0.556785458 | 0.876 | 0.703 | 2.18E-11 | Tb |
| GMK NK | TXNIP | 1.04E-08 | 0.546306865 | 0.7 | 0.572 | 0.00020559 | Tb |
| GMK NK | CD2 | 3.45E-07 | 0.523881488 | 0.618 | 0.502 | 0.006784738 | Tb |
| GMK NK | TRDC | 3.92E-16 | 0.50675847 | 0.792 | 0.57 | 7.71E-12 | Tb |
| **Disease-associated genes in FCGR3A NK** | | | | | | | |
| FCGR3A NK | FKBP5 | 1.01E-12 | 1.744733008 | 0.295 | 0.14 | 1.99E-08 | Lep-L |
| FCGR3A NK | ALDOA | 2.95E-17 | 1.742326302 | 0.305 | 0.121 | 5.81E-13 | Lep-L |
| FCGR3A NK | RBFOX2 | 5.57E-23 | 1.685598554 | 0.475 | 0.243 | 1.10E-18 | Lep-L |
| FCGR3A NK | PIM3 | 5.82E-10 | 1.383151507 | 0.36 | 0.217 | 1.15E-05 | Lep-L |
| FCGR3A NK | PTPRCAP | 5.94E-26 | 1.367936948 | 0.358 | 0.102 | 1.17E-21 | Lep-L |

| | | | | | | | |
|---|---|---|---|---|---|---|---|
| FCGR3A NK | MCOLN2 | 1.67E-07 | 1.273059937 | 0.29 | 0.171 | 0.003292598 | Lep-L |
| FCGR3A NK | DDX3Y | 7.51E-10 | 1.226068341 | 0.29 | 0.146 | 1.48E-05 | Lep-L |
| FCGR3A NK | DIP2A | 1.23E-11 | 1.158258554 | 0.415 | 0.253 | 2.42E-07 | Lep-L |
| FCGR3A NK | TNFRSF1B | 2.77E-13 | 1.102596075 | 0.554 | 0.394 | 5.45E-09 | Lep-L |
| FCGR3A NK | ATP1B3 | 6.00E-14 | 1.087469751 | 0.627 | 0.486 | 1.18E-09 | Lep-L |
| FCGR3A NK | SLC2A3 | 1.45E-13 | 1.053306334 | 0.648 | 0.487 | 2.85E-09 | Lep-L |
| FCGR3A NK | LRMP | 1.82E-07 | 1.023731538 | 0.339 | 0.22 | 0.003590202 | Lep-L |
| FCGR3A NK | ENC1 | 2.38E-07 | 1.020830144 | 0.295 | 0.179 | 0.004681468 | Lep-L |
| FCGR3A NK | KLRC1 | 1.41E-07 | 0.998748053 | 0.277 | 0.157 | 0.002768104 | Lep-L |
| FCGR3A NK | LYST | 4.57E-10 | 0.990238253 | 0.52 | 0.365 | 8.99E-06 | Lep-L |
| FCGR3A NK | PRDM1 | 2.39E-07 | 0.978820185 | 0.439 | 0.332 | 0.004705281 | Lep-L |
| FCGR3A NK | EEF2 | 1.73E-23 | 0.974760331 | 0.749 | 0.53 | 3.40E-19 | Lep-L |
| FCGR3A NK | TAP1 | 4.29E-08 | 0.941147292 | 0.376 | 0.247 | 0.000845393 | Lep-L |
| FCGR3A NK | ZFP36L1 | 2.14E-06 | 0.93478371 | 0.653 | 0.59 | 0.042128502 | Lep-L |
| FCGR3A NK | FMNL1 | 6.18E-11 | 0.93220143 | 0.483 | 0.334 | 1.22E-06 | Lep-L |
| FCGR3A NK | NR4A2 | 4.05E-11 | 0.928289483 | 0.77 | 0.687 | 7.98E-07 | Lep-L |
| FCGR3A NK | IFITM1 | 1.65E-18 | 0.885458547 | 0.747 | 0.577 | 3.25E-14 | Lep-L |
| FCGR3A NK | UBE2F | 3.24E-07 | 0.881905569 | 0.41 | 0.292 | 0.006387267 | Lep-L |
| FCGR3A NK | TUBB | 2.26E-12 | 0.863993396 | 0.548 | 0.363 | 4.44E-08 | Lep-L |
| FCGR3A NK | HNRNPL | 6.13E-08 | 0.857867906 | 0.413 | 0.289 | 0.001206985 | Lep-L |
| FCGR3A NK | GABARAPL1 | 2.03E-08 | 0.826889094 | 0.465 | 0.335 | 0.000399232 | Lep-L |
| FCGR3A NK | GNA13 | 1.68E-06 | 0.817697724 | 0.316 | 0.209 | 0.033093618 | Lep-L |
| FCGR3A NK | RASA2 | 1.00E-07 | 0.811976566 | 0.546 | 0.44 | 0.00197451 | Lep-L |
| FCGR3A NK | NPC1 | 7.43E-07 | 0.809738551 | 0.345 | 0.224 | 0.014621235 | Lep-L |
| FCGR3A NK | RHOH | 4.50E-08 | 0.775735882 | 0.601 | 0.507 | 0.000886024 | Lep-L |
| FCGR3A NK | CYTIP | 6.31E-10 | 0.742417626 | 0.674 | 0.571 | 1.24E-05 | Lep-L |
| FCGR3A NK | YY1 | 1.08E-07 | 0.715666418 | 0.46 | 0.344 | 0.002134774 | Lep-L |
| FCGR3A NK | CNBP | 1.85E-07 | 0.711345103 | 0.546 | 0.429 | 0.003643028 | Lep-L |
| FCGR3A NK | NFATC2 | 3.18E-08 | 0.705036777 | 0.574 | 0.461 | 0.000626772 | Lep-L |
| FCGR3A NK | RORA | 5.19E-08 | 0.703958482 | 0.679 | 0.603 | 0.001021486 | Lep-L |
| FCGR3A NK | IFITM2 | 1.57E-11 | 0.687957862 | 0.825 | 0.8 | 3.10E-07 | Lep-L |
| FCGR3A NK | SPON2 | 5.10E-09 | 0.669730416 | 0.648 | 0.507 | 0.000100442 | Lep-L |
| FCGR3A NK | PNRC1 | 1.09E-09 | 0.667659985 | 0.893 | 0.813 | 2.14E-05 | Lep-L |
| FCGR3A NK | VPS37B | 5.28E-10 | 0.667407486 | 0.663 | 0.543 | 1.04E-05 | Lep-L |
| FCGR3A NK | CLEC2B | 3.35E-13 | 0.651634386 | 0.825 | 0.756 | 6.59E-09 | Lep-L |
| FCGR3A NK | RUNX3 | 4.75E-09 | 0.621987736 | 0.76 | 0.702 | 9.35E-05 | Lep-L |
| FCGR3A NK | FLNA | 2.06E-08 | 0.611894895 | 0.674 | 0.572 | 0.000406016 | Lep-L |
| FCGR3A NK | CD74 | 1.42E-06 | 0.589208827 | 0.7 | 0.637 | 0.027931788 | Lep-L |
| FCGR3A NK | TSC22D3 | 1.16E-08 | 0.586826957 | 0.671 | 0.536 | 0.000228911 | Lep-L |
| FCGR3A NK | EZR | 1.60E-06 | 0.578464687 | 0.718 | 0.622 | 0.031471914 | Lep-L |
| FCGR3A NK | CCL4L2 | 2.11E-09 | 0.544928107 | 0.718 | 0.552 | 4.15E-05 | Lep-L |
| FCGR3A NK | XCL1 | 9.32E-14 | 2.082403585 | 0.352 | 0.163 | 1.83E-09 | Lep-T |
| FCGR3A NK | CST3 | 3.27E-16 | 1.924467742 | 0.293 | 0.106 | 6.44E-12 | Lep-T |
| FCGR3A NK | PTPRCAP1 | 2.16E-13 | 1.637514309 | 0.306 | 0.137 | 4.26E-09 | Lep-T |
| FCGR3A NK | PTGDS | 2.30E-11 | 1.457753142 | 0.371 | 0.186 | 4.52E-07 | Lep-T |
| FCGR3A NK | CD741 | 2.99E-07 | 0.93876334 | 0.756 | 0.624 | 0.005889675 | Lep-T |
| FCGR3A NK | PPP1CA | 7.84E-07 | 0.935274202 | 0.404 | 0.263 | 0.015434405 | Lep-T |
| FCGR3A NK | CCL4 | 7.00E-12 | 0.878695034 | 0.902 | 0.858 | 1.38E-07 | Lep-T |
| FCGR3A NK | LCP2 | 4.18E-08 | 0.870365057 | 0.391 | 0.234 | 0.000823567 | Lep-T |
| FCGR3A NK | CCL4L21 | 4.88E-08 | 0.835895983 | 0.707 | 0.567 | 0.000960272 | Lep-T |
| FCGR3A NK | CD3E | 4.13E-09 | 0.780925314 | 0.482 | 0.287 | 8.14E-05 | Lep-T |
| FCGR3A NK | ACTB | 3.72E-07 | 0.747322213 | 0.961 | 0.936 | 0.007324518 | Lep-T |
| FCGR3A NK | RBFOX21 | 6.99E-08 | 0.704099393 | 0.433 | 0.273 | 0.001375461 | Lep-T |
| FCGR3A NK | MT-ND1 | 5.91E-07 | 0.665778606 | 0.694 | 0.623 | 0.01163451 | Lep-T |
| FCGR3A NK | VAMP2 | 1.15E-08 | 0.665284934 | 0.557 | 0.37 | 0.000226782 | Lep-T |
| FCGR3A NK | ZFP36L11 | 4.16E-11 | 0.556090002 | 0.769 | 0.559 | 8.19E-07 | Lep-T |

| | | | | | | | |
|---|---|---|---|---|---|---|---|
| FCGR3A NK | KLRC2 | 1.90E-06 | 0.54744158 | 0.495 | 0.339 | 0.037476981 | Lep-T |
| FCGR3A NK | TYMP | 2.59E-07 | 1.314114654 | 0.25 | 0.14 | 0.005105002 | Mari |
| FCGR3A NK | TMEM14B | 2.04E-07 | 1.188427284 | 0.256 | 0.144 | 0.004007086 | Mari |
| FCGR3A NK | TRAPPC5 | 7.61E-08 | 1.147056023 | 0.253 | 0.136 | 0.00149697 | Mari |
| FCGR3A NK | GZMA | 1.37E-23 | 1.054313365 | 0.871 | 0.701 | 2.70E-19 | Mari |
| FCGR3A NK | VAMP5 | 5.87E-09 | 0.990911485 | 0.323 | 0.182 | 0.000115614 | Mari |
| FCGR3A NK | MRPL20 | 3.27E-07 | 0.905595046 | 0.334 | 0.205 | 0.006431502 | Mari |
| FCGR3A NK | CD21 | 3.74E-08 | 0.889352741 | 0.472 | 0.343 | 0.000735942 | Mari |
| FCGR3A NK | APOBEC3G | 3.18E-12 | 0.857108575 | 0.601 | 0.418 | 6.26E-08 | Mari |
| FCGR3A NK | TRDC | 1.25E-10 | 0.854992585 | 0.576 | 0.419 | 2.46E-06 | Mari |
| FCGR3A NK | ERH | 1.25E-09 | 0.822276118 | 0.449 | 0.294 | 2.47E-05 | Mari |
| FCGR3A NK | ATP5MF | 4.94E-09 | 0.769401507 | 0.421 | 0.263 | 9.72E-05 | Mari |
| FCGR3A NK | BST2 | 6.83E-07 | 0.747112415 | 0.444 | 0.335 | 0.013442237 | Mari |
| FCGR3A NK | SEC61G | 2.16E-09 | 0.731872138 | 0.48 | 0.32 | 4.26E-05 | Mari |
| FCGR3A NK | GNLY | 3.95E-19 | 0.728307211 | 0.963 | 0.937 | 7.77E-15 | Mari |
| FCGR3A NK | PSMB1 | 9.95E-10 | 0.709591046 | 0.517 | 0.36 | 1.96E-05 | Mari |
| FCGR3A NK | ATP5MPL | 2.38E-06 | 0.691866001 | 0.337 | 0.22 | 0.046905536 | Mari |
| FCGR3A NK | HOPX | 5.19E-09 | 0.670454499 | 0.697 | 0.593 | 0.000102244 | Mari |
| FCGR3A NK | ATP5ME | 5.82E-08 | 0.670005248 | 0.458 | 0.309 | 0.001145873 | Mari |
| FCGR3A NK | PSMB3 | 1.94E-06 | 0.655804906 | 0.357 | 0.229 | 0.038212094 | Mari |
| FCGR3A NK | GZMB | 2.88E-14 | 0.654634069 | 0.938 | 0.914 | 5.66E-10 | Mari |
| FCGR3A NK | ATP5PF | 4.01E-08 | 0.65015667 | 0.461 | 0.313 | 0.000789215 | Mari |
| FCGR3A NK | COX7A2 | 1.01E-08 | 0.645145922 | 0.567 | 0.428 | 0.00019813 | Mari |
| FCGR3A NK | NDUFB2 | 1.03E-08 | 0.642008413 | 0.59 | 0.455 | 0.000201825 | Mari |
| FCGR3A NK | COX6C | 3.46E-08 | 0.613618848 | 0.598 | 0.466 | 0.000680269 | Mari |
| FCGR3A NK | PSMB9 | 3.07E-07 | 0.598763679 | 0.635 | 0.529 | 0.006047099 | Mari |
| FCGR3A NK | GMFG | 8.62E-08 | 0.580053292 | 0.559 | 0.434 | 0.0016962 | Mari |
| FCGR3A NK | NDUFA4 | 4.40E-07 | 0.566686924 | 0.598 | 0.492 | 0.008668916 | Mari |
| FCGR3A NK | ITM2B | 1.42E-12 | 0.565807136 | 0.809 | 0.67 | 2.79E-08 | Mari |
| FCGR3A NK | TRMT112 | 5.35E-07 | 0.554097971 | 0.596 | 0.472 | 0.010527741 | Mari |
| FCGR3A NK | ATP6V0E1 | 1.70E-07 | 0.536853746 | 0.593 | 0.45 | 0.003345852 | Mari |
| FCGR3A NK | COX6B1 | 5.22E-10 | 0.535515568 | 0.632 | 0.461 | 1.03E-05 | Mari |
| FCGR3A NK | IGKC | 2.31E-45 | 3.982392005 | 0.396 | 0.081 | 4.54E-41 | Tb |
| FCGR3A NK | SLC35F1 | 5.42E-48 | 2.569654295 | 0.428 | 0.078 | 1.07E-43 | Tb |
| FCGR3A NK | S100A9 | 1.87E-26 | 2.233011863 | 0.367 | 0.106 | 3.67E-22 | Tb |
| FCGR3A NK | S100A8 | 9.26E-21 | 1.954883177 | 0.349 | 0.117 | 1.82E-16 | Tb |
| FCGR3A NK | HOMER1 | 9.78E-11 | 1.741422783 | 0.263 | 0.115 | 1.93E-06 | Tb |
| FCGR3A NK | KRT14 | 5.13E-14 | 1.372115262 | 0.299 | 0.116 | 1.01E-09 | Tb |
| FCGR3A NK | MYOM2 | 5.09E-11 | 1.264349462 | 0.388 | 0.211 | 1.00E-06 | Tb |
| FCGR3A NK | HCP5 | 1.72E-08 | 1.166933843 | 0.291 | 0.151 | 0.000338278 | Tb |
| FCGR3A NK | INTS6 | 1.79E-13 | 1.156602212 | 0.601 | 0.402 | 3.53E-09 | Tb |
| FCGR3A NK | XIST | 4.75E-13 | 1.089234254 | 0.385 | 0.179 | 9.35E-09 | Tb |
| FCGR3A NK | GLCCI1 | 1.31E-07 | 1.002085405 | 0.396 | 0.244 | 0.002582651 | Tb |
| FCGR3A NK | ZSWIM6 | 1.62E-08 | 0.991656842 | 0.554 | 0.405 | 0.0003195 | Tb |
| FCGR3A NK | RASA1 | 5.15E-07 | 0.92460951 | 0.284 | 0.155 | 0.010128078 | Tb |
| FCGR3A NK | RYBP | 3.50E-07 | 0.893725644 | 0.345 | 0.206 | 0.006885174 | Tb |
| FCGR3A NK | MT-ATP6 | 3.72E-19 | 0.881271116 | 0.986 | 0.813 | 7.32E-15 | Tb |
| FCGR3A NK | DOCK8 | 8.90E-12 | 0.829688807 | 0.543 | 0.326 | 1.75E-07 | Tb |
| FCGR3A NK | MT-CO31 | 3.40E-12 | 0.820729134 | 0.892 | 0.789 | 6.70E-08 | Tb |
| FCGR3A NK | ELL2 | 1.23E-06 | 0.782947704 | 0.475 | 0.338 | 0.024230189 | Tb |
| FCGR3A NK | MT-CYB | 7.08E-10 | 0.662637631 | 0.885 | 0.759 | 1.39E-05 | Tb |
| FCGR3A NK | JPT1 | 4.37E-07 | 0.616837507 | 0.457 | 0.289 | 0.008611719 | Tb |
| FCGR3A NK | MT-ND3 | 5.50E-14 | 0.559621527 | 0.932 | 0.775 | 1.08E-09 | Tb |
| FCGR3A NK | CD2 | 1.23E-06 | 0.508807131 | 0.504 | 0.344 | 0.024191933 | Tb |
| **Disease-associated genes in TNFRSF9 NK** | | | | | | | |
| TNFRSF9 NK | PRKY | 1.41E-37 | 2.714305125 | 0.417 | 0.076 | 2.78E-33 | Lep-L |

| | | | | | | | |
|---|---|---|---|---|---|---|---|
| TNFRSF9 NK | MCOLN2 | 5.47E-22 | 2.646521699 | 0.309 | 0.08 | 1.08E-17 | Lep-L |
| TNFRSF9 NK | DDX3Y | 5.20E-34 | 2.505240718 | 0.44 | 0.106 | 1.02E-29 | Lep-L |
| TNFRSF9 NK | APOC1 | 1.63E-19 | 2.18036739 | 0.296 | 0.082 | 3.22E-15 | Lep-L |
| TNFRSF9 NK | UTY | 1.17E-23 | 2.09050441 | 0.339 | 0.083 | 2.31E-19 | Lep-L |
| TNFRSF9 NK | APOE | 1.09E-32 | 1.887818564 | 0.59 | 0.24 | 2.15E-28 | Lep-L |
| TNFRSF9 NK | HLA-DQB1 | 3.53E-19 | 1.656618528 | 0.502 | 0.266 | 6.94E-15 | Lep-L |
| TNFRSF9 NK | TLE1 | 1.61E-12 | 1.483499169 | 0.44 | 0.255 | 3.17E-08 | Lep-L |
| TNFRSF9 NK | HLA-DRB1 | 4.78E-26 | 1.476475471 | 0.678 | 0.397 | 9.42E-22 | Lep-L |
| TNFRSF9 NK | WHRN | 2.46E-07 | 1.360228149 | 0.336 | 0.208 | 0.004836273 | Lep-L |
| TNFRSF9 NK | SPOCK2 | 5.99E-11 | 1.353080407 | 0.401 | 0.227 | 1.18E-06 | Lep-L |
| TNFRSF9 NK | HLA-DQA1 | 8.63E-09 | 1.345938273 | 0.293 | 0.151 | 0.000169819 | Lep-L |
| TNFRSF9 NK | CXCR6 | 1.34E-07 | 1.342616166 | 0.313 | 0.181 | 0.002635142 | Lep-L |
| TNFRSF9 NK | TNFRSF1B | 8.09E-13 | 1.322952915 | 0.55 | 0.391 | 1.59E-08 | Lep-L |
| TNFRSF9 NK | ATXN1 | 4.20E-17 | 1.281571875 | 0.694 | 0.495 | 8.27E-13 | Lep-L |
| TNFRSF9 NK | HAVCR2 | 4.11E-08 | 1.232347178 | 0.345 | 0.204 | 0.000809786 | Lep-L |
| TNFRSF9 NK | TAP1 | 1.01E-10 | 1.195587672 | 0.394 | 0.224 | 1.98E-06 | Lep-L |
| TNFRSF9 NK | RASA3 | 1.50E-07 | 1.187481859 | 0.391 | 0.254 | 0.002951087 | Lep-L |
| TNFRSF9 NK | TNFRSF9 | 3.97E-07 | 1.18697384 | 0.42 | 0.295 | 0.007820916 | Lep-L |
| TNFRSF9 NK | AKIRIN1 | 6.16E-09 | 1.158134037 | 0.42 | 0.278 | 0.000121305 | Lep-L |
| TNFRSF9 NK | METRNL | 2.80E-24 | 1.146674974 | 0.821 | 0.607 | 5.51E-20 | Lep-L |
| TNFRSF9 NK | FGFR1OP2 | 4.17E-16 | 1.134071018 | 0.678 | 0.503 | 8.20E-12 | Lep-L |
| TNFRSF9 NK | BICDL1 | 2.65E-11 | 1.11678458 | 0.505 | 0.32 | 5.22E-07 | Lep-L |
| TNFRSF9 NK | FTL | 2.76E-43 | 1.102180915 | 0.974 | 0.95 | 5.44E-39 | Lep-L |
| TNFRSF9 NK | CNN2 | 7.15E-08 | 1.024244657 | 0.466 | 0.326 | 0.001407536 | Lep-L |
| TNFRSF9 NK | BIRC3 | 2.37E-11 | 1.017231149 | 0.7 | 0.554 | 4.66E-07 | Lep-L |
| TNFRSF9 NK | FLNA | 2.08E-11 | 1.015941664 | 0.586 | 0.434 | 4.09E-07 | Lep-L |
| TNFRSF9 NK | TC2N | 8.17E-09 | 0.968094344 | 0.524 | 0.367 | 0.000160781 | Lep-L |
| TNFRSF9 NK | TGFBR2 | 1.65E-07 | 0.958014715 | 0.547 | 0.424 | 0.003238067 | Lep-L |
| TNFRSF9 NK | SLBP | 6.38E-09 | 0.955622086 | 0.417 | 0.258 | 0.000125539 | Lep-L |
| TNFRSF9 NK | PIM3 | 1.19E-09 | 0.945482959 | 0.511 | 0.353 | 2.33E-05 | Lep-L |
| TNFRSF9 NK | OAZ1 | 2.32E-25 | 0.943827861 | 0.886 | 0.793 | 4.57E-21 | Lep-L |
| TNFRSF9 NK | NEAT1 | 3.36E-12 | 0.940959935 | 0.798 | 0.724 | 6.61E-08 | Lep-L |
| TNFRSF9 NK | LYST | 4.18E-11 | 0.934706996 | 0.704 | 0.559 | 8.23E-07 | Lep-L |
| TNFRSF9 NK | CLEC2B | 2.18E-16 | 0.934191304 | 0.844 | 0.752 | 4.29E-12 | Lep-L |
| TNFRSF9 NK | SMCHD1 | 8.89E-17 | 0.930448238 | 0.814 | 0.687 | 1.75E-12 | Lep-L |
| TNFRSF9 NK | CDK6 | 1.30E-08 | 0.930064684 | 0.485 | 0.32 | 0.000255223 | Lep-L |
| TNFRSF9 NK | HIBCH | 3.63E-09 | 0.918060439 | 0.547 | 0.388 | 7.14E-05 | Lep-L |
| TNFRSF9 NK | CNOT2 | 5.87E-12 | 0.906746718 | 0.798 | 0.715 | 1.16E-07 | Lep-L |
| TNFRSF9 NK | GZMB | 8.16E-07 | 0.89935681 | 0.651 | 0.588 | 0.016063208 | Lep-L |
| TNFRSF9 NK | SAT1 | 3.60E-08 | 0.889362551 | 0.805 | 0.74 | 0.000708439 | Lep-L |
| TNFRSF9 NK | ATP1B3 | 1.95E-20 | 0.876254213 | 0.883 | 0.776 | 3.84E-16 | Lep-L |
| TNFRSF9 NK | DOK2 | 3.20E-11 | 0.874708377 | 0.681 | 0.533 | 6.30E-07 | Lep-L |
| TNFRSF9 NK | PRF1 | 2.00E-10 | 0.841228527 | 0.541 | 0.358 | 3.94E-06 | Lep-L |
| TNFRSF9 NK | CNOT6L | 1.26E-20 | 0.838309091 | 0.886 | 0.805 | 2.47E-16 | Lep-L |
| TNFRSF9 NK | LEPROTL1 | 5.64E-08 | 0.831188737 | 0.678 | 0.61 | 0.001110519 | Lep-L |
| TNFRSF9 NK | DIAPH1 | 1.69E-06 | 0.823817468 | 0.466 | 0.352 | 0.033186968 | Lep-L |
| TNFRSF9 NK | MAPKAPK2 | 1.90E-06 | 0.806404621 | 0.436 | 0.314 | 0.037305006 | Lep-L |
| TNFRSF9 NK | HLA-DRA | 7.43E-09 | 0.796498748 | 0.57 | 0.438 | 0.000146249 | Lep-L |
| TNFRSF9 NK | HSP90B1 | 8.51E-09 | 0.779591055 | 0.661 | 0.536 | 0.000167417 | Lep-L |
| TNFRSF9 NK | CD74 | 6.97E-07 | 0.778812753 | 0.694 | 0.625 | 0.013719295 | Lep-L |
| TNFRSF9 NK | ISG20 | 4.01E-11 | 0.776664223 | 0.632 | 0.48 | 7.89E-07 | Lep-L |
| TNFRSF9 NK | IFI16 | 1.70E-06 | 0.773503953 | 0.567 | 0.464 | 0.033412473 | Lep-L |
| TNFRSF9 NK | CYTIP | 2.57E-07 | 0.750926924 | 0.625 | 0.533 | 0.005055782 | Lep-L |
| TNFRSF9 NK | TPM4 | 2.40E-08 | 0.746249883 | 0.599 | 0.488 | 0.000472634 | Lep-L |
| TNFRSF9 NK | RORA | 1.20E-07 | 0.686197147 | 0.713 | 0.621 | 0.002352344 | Lep-L |
| TNFRSF9 NK | RASA2 | 3.42E-07 | 0.683663743 | 0.612 | 0.488 | 0.006730377 | Lep-L |

| | | | | | | | |
|---|---|---|---|---|---|---|---|
| TNFRSF9 NK | FKBP4 | 4.16E-08 | 0.683634285 | 0.619 | 0.468 | 0.000819106 | Lep-L |
| TNFRSF9 NK | STK4 | 9.42E-14 | 0.683081351 | 0.827 | 0.73 | 1.86E-09 | Lep-L |
| TNFRSF9 NK | SUB1 | 5.16E-07 | 0.680503213 | 0.671 | 0.631 | 0.010149646 | Lep-L |
| TNFRSF9 NK | VPS37B | 3.39E-09 | 0.649695874 | 0.85 | 0.749 | 6.67E-05 | Lep-L |
| TNFRSF9 NK | NFAT5 | 6.05E-07 | 0.646435519 | 0.713 | 0.633 | 0.011909725 | Lep-L |
| TNFRSF9 NK | ITGA4 | 4.98E-07 | 0.613460612 | 0.684 | 0.577 | 0.009808831 | Lep-L |
| TNFRSF9 NK | SAMSN1 | 7.58E-07 | 0.583585047 | 0.801 | 0.731 | 0.014929491 | Lep-L |
| TNFRSF9 NK | CHD2 | 1.07E-06 | 0.583175611 | 0.821 | 0.779 | 0.020985615 | Lep-L |
| TNFRSF9 NK | EZR | 3.05E-09 | 0.579768432 | 0.886 | 0.852 | 6.00E-05 | Lep-L |
| TNFRSF9 NK | CEMIP2 | 9.52E-10 | 0.576725034 | 0.876 | 0.808 | 1.87E-05 | Lep-L |
| TNFRSF9 NK | HNRNPDL | 7.03E-09 | 0.553330556 | 0.743 | 0.631 | 0.000138464 | Lep-L |
| TNFRSF9 NK | JUND | 6.29E-17 | 0.528404564 | 0.958 | 0.97 | 1.24E-12 | Lep-L |
| TNFRSF9 NK | SARAF | 2.25E-10 | 0.515795081 | 0.883 | 0.853 | 4.42E-06 | Lep-L |
| TNFRSF9 NK | ASTL | 1.16E-28 | 2.588915064 | 0.365 | 0.064 | 2.29E-24 | Lep-T |
| TNFRSF9 NK | HIST2H2AC | 7.73E-20 | 2.225379168 | 0.294 | 0.059 | 1.52E-15 | Lep-T |
| TNFRSF9 NK | SPDYA | 3.37E-15 | 1.967927639 | 0.265 | 0.064 | 6.64E-11 | Lep-T |
| TNFRSF9 NK | ENOSF1 | 1.63E-11 | 1.627840193 | 0.335 | 0.126 | 3.20E-07 | Lep-T |
| TNFRSF9 NK | ADAMTS17 | 2.70E-08 | 1.613303382 | 0.435 | 0.232 | 0.000531907 | Lep-T |
| TNFRSF9 NK | CLNK | 5.88E-10 | 1.534525263 | 0.253 | 0.085 | 1.16E-05 | Lep-T |
| TNFRSF9 NK | SBF2 | 1.66E-09 | 1.520500773 | 0.259 | 0.09 | 3.26E-05 | Lep-T |
| TNFRSF9 NK | VPS13D | 7.55E-10 | 1.450558045 | 0.353 | 0.145 | 1.49E-05 | Lep-T |
| TNFRSF9 NK | SLC9A9 | 1.34E-08 | 1.439108165 | 0.282 | 0.115 | 0.0002628 | Lep-T |
| TNFRSF9 NK | HIST1H1E | 8.58E-10 | 1.426743076 | 0.324 | 0.129 | 1.69E-05 | Lep-T |
| TNFRSF9 NK | MICAL2 | 1.68E-12 | 1.399444875 | 0.429 | 0.17 | 3.32E-08 | Lep-T |
| TNFRSF9 NK | ADAM28 | 2.53E-09 | 1.384927129 | 0.347 | 0.151 | 4.97E-05 | Lep-T |
| TNFRSF9 NK | KLHL5 | 5.42E-08 | 1.371955023 | 0.253 | 0.1 | 0.001065918 | Lep-T |
| TNFRSF9 NK | PLCB1 | 2.76E-10 | 1.342252894 | 0.618 | 0.387 | 5.42E-06 | Lep-T |
| TNFRSF9 NK | MT-CO3 | 2.78E-35 | 1.333221539 | 0.971 | 0.87 | 5.48E-31 | Lep-T |
| TNFRSF9 NK | SNX9 | 1.48E-15 | 1.332320649 | 0.465 | 0.173 | 2.91E-11 | Lep-T |
| TNFRSF9 NK | TBC1D5 | 6.38E-10 | 1.317700135 | 0.441 | 0.227 | 1.26E-05 | Lep-T |
| TNFRSF9 NK | XAF1 | 6.67E-10 | 1.307315356 | 0.347 | 0.143 | 1.31E-05 | Lep-T |
| TNFRSF9 NK | LDB2 | 1.31E-08 | 1.27491583 | 0.418 | 0.217 | 0.000257868 | Lep-T |
| TNFRSF9 NK | MT-ATP6 | 2.44E-35 | 1.270861803 | 0.976 | 0.889 | 4.80E-31 | Lep-T |
| TNFRSF9 NK | STK39 | 2.03E-07 | 1.263825503 | 0.388 | 0.199 | 0.003986157 | Lep-T |
| TNFRSF9 NK | HDAC4 | 3.92E-08 | 1.260584946 | 0.282 | 0.114 | 0.00077189 | Lep-T |
| TNFRSF9 NK | KCNQ5 | 3.38E-07 | 1.258178518 | 0.335 | 0.166 | 0.006652329 | Lep-T |
| TNFRSF9 NK | FLNB | 4.44E-08 | 1.247043694 | 0.329 | 0.148 | 0.000874322 | Lep-T |
| TNFRSF9 NK | ZMYND8 | 1.72E-10 | 1.239807958 | 0.265 | 0.085 | 3.39E-06 | Lep-T |
| TNFRSF9 NK | TTC7A | 1.52E-10 | 1.233160539 | 0.388 | 0.169 | 2.99E-06 | Lep-T |
| TNFRSF9 NK | PPP2R2B | 1.15E-07 | 1.228949739 | 0.376 | 0.19 | 0.002254288 | Lep-T |
| TNFRSF9 NK | RASGRP1 | 8.25E-07 | 1.20306374 | 0.371 | 0.194 | 0.016243307 | Lep-T |
| TNFRSF9 NK | MT-ND1 | 3.05E-27 | 1.201479421 | 0.947 | 0.741 | 6.00E-23 | Lep-T |
| TNFRSF9 NK | COMMD10 | 6.86E-07 | 1.191801992 | 0.282 | 0.128 | 0.013496385 | Lep-T |
| TNFRSF9 NK | SCML4 | 5.45E-11 | 1.189729545 | 0.435 | 0.194 | 1.07E-06 | Lep-T |
| TNFRSF9 NK | FOXP1 | 3.90E-12 | 1.160042404 | 0.676 | 0.397 | 7.68E-08 | Lep-T |
| TNFRSF9 NK | VAV3 | 4.95E-10 | 1.158382314 | 0.565 | 0.325 | 9.75E-06 | Lep-T |
| TNFRSF9 NK | CCL4L2 | 4.00E-07 | 1.147873466 | 0.429 | 0.242 | 0.007883186 | Lep-T |
| TNFRSF9 NK | SCLT1 | 3.23E-07 | 1.13673219 | 0.588 | 0.404 | 0.006364652 | Lep-T |
| TNFRSF9 NK | GTF2F2 | 6.52E-07 | 1.129740808 | 0.329 | 0.163 | 0.012833521 | Lep-T |
| TNFRSF9 NK | FUT8 | 3.04E-07 | 1.125566364 | 0.306 | 0.139 | 0.005981093 | Lep-T |
| TNFRSF9 NK | MT-CYB | 6.28E-30 | 1.125270127 | 0.959 | 0.77 | 1.24E-25 | Lep-T |
| TNFRSF9 NK | MT-ND5 | 2.78E-21 | 1.115371573 | 0.888 | 0.606 | 5.48E-17 | Lep-T |
| TNFRSF9 NK | MPP7 | 1.18E-08 | 1.096323497 | 0.335 | 0.145 | 0.000231699 | Lep-T |
| TNFRSF9 NK | TGFBR3 | 9.90E-10 | 1.094153909 | 0.447 | 0.217 | 1.95E-05 | Lep-T |
| TNFRSF9 NK | ZNF609 | 4.64E-08 | 1.090832274 | 0.376 | 0.181 | 0.000913806 | Lep-T |
| TNFRSF9 NK | HIST1H1C | 1.31E-10 | 1.085734791 | 0.365 | 0.141 | 2.57E-06 | Lep-T |

| | | | | | | | |
|---|---|---|---|---|---|---|---|
| TNFRSF9 NK | RUNX1 | 1.53E-08 | 1.083725721 | 0.571 | 0.328 | 0.000301776 | Lep-T |
| TNFRSF9 NK | PTPN12 | 1.33E-08 | 1.069583614 | 0.441 | 0.224 | 0.000262109 | Lep-T |
| TNFRSF9 NK | WDR70 | 1.51E-06 | 1.064156584 | 0.3 | 0.146 | 0.029665784 | Lep-T |
| TNFRSF9 NK | STAT1 | 1.67E-07 | 1.061099362 | 0.453 | 0.245 | 0.00328925 | Lep-T |
| TNFRSF9 NK | CDKAL1 | 4.70E-07 | 1.055207806 | 0.394 | 0.207 | 0.00925255 | Lep-T |
| TNFRSF9 NK | EXOC4 | 7.02E-08 | 1.053350892 | 0.518 | 0.3 | 0.001382043 | Lep-T |
| TNFRSF9 NK | RGS1 | 1.14E-06 | 1.037233917 | 0.612 | 0.464 | 0.022382987 | Lep-T |
| TNFRSF9 NK | LPIN1 | 1.16E-06 | 1.036259183 | 0.424 | 0.244 | 0.02276777 | Lep-T |
| TNFRSF9 NK | XIST | 5.77E-11 | 1.030218108 | 0.5 | 0.239 | 1.13E-06 | Lep-T |
| TNFRSF9 NK | ZCCHC7 | 6.98E-09 | 1.028481617 | 0.412 | 0.199 | 0.000137393 | Lep-T |
| TNFRSF9 NK | NOCT | 1.02E-07 | 1.023641208 | 0.365 | 0.176 | 0.002005566 | Lep-T |
| TNFRSF9 NK | NEDD9 | 1.46E-08 | 1.023222632 | 0.476 | 0.25 | 0.000287694 | Lep-T |
| TNFRSF9 NK | ATP8A1 | 2.54E-09 | 1.022671922 | 0.629 | 0.404 | 5.00E-05 | Lep-T |
| TNFRSF9 NK | JUN | 6.21E-09 | 1.021802702 | 0.729 | 0.562 | 0.000122197 | Lep-T |
| TNFRSF9 NK | ABHD2 | 1.45E-07 | 1.005734257 | 0.529 | 0.323 | 0.002858201 | Lep-T |
| TNFRSF9 NK | SPIDR | 1.42E-10 | 0.983114699 | 0.547 | 0.289 | 2.80E-06 | Lep-T |
| TNFRSF9 NK | IQGAP2 | 9.79E-10 | 0.980545338 | 0.665 | 0.439 | 1.93E-05 | Lep-T |
| TNFRSF9 NK | ODC1 | 1.57E-09 | 0.962678182 | 0.441 | 0.209 | 3.09E-05 | Lep-T |
| TNFRSF9 NK | ATP9B | 1.26E-07 | 0.961680313 | 0.412 | 0.215 | 0.002478272 | Lep-T |
| TNFRSF9 NK | RSRC1 | 3.99E-07 | 0.9514406 | 0.259 | 0.105 | 0.007848251 | Lep-T |
| TNFRSF9 NK | IL12RB2 | 1.11E-08 | 0.950390871 | 0.729 | 0.503 | 0.000218161 | Lep-T |
| TNFRSF9 NK | RGS2 | 7.49E-07 | 0.948891913 | 0.565 | 0.359 | 0.014751584 | Lep-T |
| TNFRSF9 NK | CASK | 5.26E-09 | 0.948521243 | 0.606 | 0.362 | 0.000103448 | Lep-T |
| TNFRSF9 NK | RABGAP1L | 8.92E-12 | 0.94519895 | 0.771 | 0.541 | 1.75E-07 | Lep-T |
| TNFRSF9 NK | FBXO34 | 1.03E-07 | 0.945088432 | 0.735 | 0.548 | 0.002019384 | Lep-T |
| TNFRSF9 NK | MT-ND2 | 6.83E-21 | 0.930539755 | 0.965 | 0.785 | 1.34E-16 | Lep-T |
| TNFRSF9 NK | PRKX | 8.37E-08 | 0.920084497 | 0.635 | 0.437 | 0.001646718 | Lep-T |
| TNFRSF9 NK | PIP4K2A | 7.00E-11 | 0.911674997 | 0.771 | 0.534 | 1.38E-06 | Lep-T |
| TNFRSF9 NK | MAP3K5 | 1.77E-06 | 0.909585923 | 0.347 | 0.173 | 0.034788696 | Lep-T |
| TNFRSF9 NK | RERE | 1.94E-08 | 0.904747329 | 0.594 | 0.398 | 0.000382575 | Lep-T |
| TNFRSF9 NK | CCDC69 | 3.64E-08 | 0.89386239 | 0.371 | 0.169 | 0.000716632 | Lep-T |
| TNFRSF9 NK | PHF20 | 2.13E-08 | 0.880221359 | 0.671 | 0.461 | 0.000419672 | Lep-T |
| TNFRSF9 NK | ZBTB20 | 1.34E-08 | 0.855680788 | 0.659 | 0.442 | 0.000262941 | Lep-T |
| TNFRSF9 NK | FOXN3 | 1.33E-07 | 0.852649921 | 0.712 | 0.506 | 0.002608484 | Lep-T |
| TNFRSF9 NK | KLF6 | 1.85E-08 | 0.848872798 | 0.753 | 0.557 | 0.000364757 | Lep-T |
| TNFRSF9 NK | ID2 | 2.24E-07 | 0.83361533 | 0.676 | 0.469 | 0.004410635 | Lep-T |
| TNFRSF9 NK | TSPAN5 | 1.91E-06 | 0.830669797 | 0.376 | 0.196 | 0.037655069 | Lep-T |
| TNFRSF9 NK | TAF15 | 7.42E-07 | 0.824135759 | 0.388 | 0.198 | 0.014597524 | Lep-T |
| TNFRSF9 NK | KSR1 | 7.90E-07 | 0.823074856 | 0.347 | 0.164 | 0.0155449 | Lep-T |
| TNFRSF9 NK | SKAP1 | 1.40E-09 | 0.817372124 | 0.712 | 0.502 | 2.75E-05 | Lep-T |
| TNFRSF9 NK | LRBA | 6.35E-08 | 0.813284317 | 0.635 | 0.403 | 0.001250728 | Lep-T |
| TNFRSF9 NK | VPS13B | 1.49E-06 | 0.809778627 | 0.524 | 0.314 | 0.029408091 | Lep-T |
| TNFRSF9 NK | PLXNC1 | 1.04E-06 | 0.800267566 | 0.388 | 0.198 | 0.020383057 | Lep-T |
| TNFRSF9 NK | GAS5 | 4.85E-08 | 0.786055516 | 0.565 | 0.307 | 0.000954834 | Lep-T |
| TNFRSF9 NK | XPO1 | 5.54E-08 | 0.776269381 | 0.512 | 0.272 | 0.001089864 | Lep-T |
| TNFRSF9 NK | NCOA2 | 9.33E-07 | 0.769752022 | 0.518 | 0.289 | 0.018368752 | Lep-T |
| TNFRSF9 NK | TCF12 | 4.34E-08 | 0.765647793 | 0.506 | 0.275 | 0.000855176 | Lep-T |
| TNFRSF9 NK | MICU2 | 4.66E-07 | 0.759004635 | 0.318 | 0.143 | 0.009165813 | Lep-T |
| TNFRSF9 NK | MYCBP2 | 1.55E-06 | 0.750878551 | 0.594 | 0.362 | 0.030461871 | Lep-T |
| TNFRSF9 NK | MT-CO1 | 2.17E-22 | 0.722833497 | 0.982 | 0.97 | 4.27E-18 | Lep-T |
| TNFRSF9 NK | EML4 | 1.92E-07 | 0.703480548 | 0.865 | 0.672 | 0.003776067 | Lep-T |
| TNFRSF9 NK | MT-ND4 | 2.26E-19 | 0.690829686 | 0.959 | 0.885 | 4.44E-15 | Lep-T |
| TNFRSF9 NK | ARIH1 | 3.50E-07 | 0.668317365 | 0.824 | 0.648 | 0.006893629 | Lep-T |
| TNFRSF9 NK | YTHDF3 | 2.20E-06 | 0.668253992 | 0.459 | 0.267 | 0.043375142 | Lep-T |
| TNFRSF9 NK | ARHGAP15 | 3.95E-07 | 0.666269768 | 0.765 | 0.581 | 0.007774108 | Lep-T |
| TNFRSF9 NK | DOCK2 | 1.14E-06 | 0.648118037 | 0.494 | 0.277 | 0.022425934 | Lep-T |

| | | | | | | | |
|---|---|---|---|---|---|---|---|
| TNFRSF9 NK | GOLGA4 | 2.48E-06 | 0.643834075 | 0.524 | 0.32 | 0.04879734 | Lep-T |
| TNFRSF9 NK | STAT4 | 8.69E-07 | 0.622279531 | 0.865 | 0.819 | 0.017104345 | Lep-T |
| TNFRSF9 NK | MT-CO2 | 1.79E-15 | 0.597191968 | 0.982 | 0.935 | 3.52E-11 | Lep-T |
| TNFRSF9 NK | CELF2 | 3.86E-07 | 0.539824197 | 0.712 | 0.458 | 0.007601108 | Lep-T |
| TNFRSF9 NK | NFKBIZ | 4.51E-07 | 0.52839957 | 0.706 | 0.491 | 0.008886368 | Lep-T |
| TNFRSF9 NK | ZNF600 | 2.14E-08 | 1.762154841 | 0.278 | 0.116 | 0.000420919 | Mari |
| TNFRSF9 NK | LYAR | 1.14E-07 | 1.666467716 | 0.259 | 0.109 | 0.002247103 | Mari |
| TNFRSF9 NK | TMCO1 | 1.28E-07 | 1.488350764 | 0.321 | 0.159 | 0.002526769 | Mari |
| TNFRSF9 NK | UTP11 | 7.91E-07 | 1.122847219 | 0.278 | 0.123 | 0.015572777 | Mari |
| TNFRSF9 NK | IER5L | 3.24E-07 | 1.042307921 | 0.401 | 0.212 | 0.006383029 | Mari |
| TNFRSF9 NK | MT-ATP8 | 1.05E-07 | 0.803998144 | 0.667 | 0.48 | 0.002065148 | Mari |
| TNFRSF9 NK | GNLY1 | 2.77E-07 | 0.626145293 | 0.846 | 0.677 | 0.005442856 | Mari |
| TNFRSF9 NK | HSPA1A | 7.18E-07 | 0.603467536 | 0.944 | 0.975 | 0.014124648 | Mari |
| TNFRSF9 NK | S100A8 | 1.29E-20 | 2.710038806 | 0.342 | 0.108 | 2.54E-16 | Tb |
| TNFRSF9 NK | S100A9 | 6.73E-26 | 2.649711544 | 0.403 | 0.122 | 1.32E-21 | Tb |
| TNFRSF9 NK | SLC35F1 | 1.22E-19 | 2.16940181 | 0.297 | 0.08 | 2.40E-15 | Tb |
| TNFRSF9 NK | KRT14 | 9.82E-20 | 2.064466133 | 0.327 | 0.102 | 1.93E-15 | Tb |
| TNFRSF9 NK | IGKC | 1.01E-07 | 1.881425114 | 0.339 | 0.218 | 0.001993328 | Tb |
| TNFRSF9 NK | HOMER1 | 4.98E-10 | 1.529632681 | 0.297 | 0.138 | 9.81E-06 | Tb |
| TNFRSF9 NK | KRT10 | 1.12E-09 | 1.448930116 | 0.47 | 0.307 | 2.21E-05 | Tb |
| TNFRSF9 NK | STX17-AS11 | 1.20E-07 | 1.157572209 | 0.282 | 0.144 | 0.002369658 | Tb |
| TNFRSF9 NK | RGCC | 5.88E-15 | 1.127799565 | 0.676 | 0.468 | 1.16E-10 | Tb |
| TNFRSF9 NK | CLCN3 | 1.00E-09 | 1.109447649 | 0.436 | 0.258 | 1.97E-05 | Tb |
| TNFRSF9 NK | OXNAD1 | 7.44E-07 | 0.989198991 | 0.27 | 0.144 | 0.01463794 | Tb |
| TNFRSF9 NK | DLEU1 | 9.30E-07 | 0.911156947 | 0.285 | 0.15 | 0.018308191 | Tb |
| TNFRSF9 NK | PAK1 | 6.04E-07 | 0.844701451 | 0.461 | 0.315 | 0.011881546 | Tb |
| TNFRSF9 NK | PRMT9 | 2.99E-08 | 0.832960058 | 0.494 | 0.311 | 0.000588693 | Tb |
| TNFRSF9 NK | KLRC3 | 3.98E-08 | 0.821998028 | 0.43 | 0.263 | 0.000783562 | Tb |
| TNFRSF9 NK | HOPX | 3.88E-07 | 0.820818985 | 0.542 | 0.388 | 0.007635339 | Tb |
| TNFRSF9 NK | XIST1 | 1.34E-07 | 0.748065934 | 0.394 | 0.228 | 0.002629428 | Tb |
| TNFRSF9 NK | TRDC | 2.51E-09 | 0.738988136 | 0.618 | 0.435 | 4.93E-05 | Tb |
| TNFRSF9 NK | FCER1G | 4.90E-09 | 0.734392657 | 0.648 | 0.469 | 9.65E-05 | Tb |
| TNFRSF9 NK | CTSW | 4.18E-07 | 0.704711143 | 0.673 | 0.537 | 0.008230619 | Tb |
| TNFRSF9 NK | MT-CYB1 | 1.34E-06 | 0.700130917 | 0.839 | 0.784 | 0.026391321 | Tb |
| TNFRSF9 NK | MT-ATP61 | 2.63E-10 | 0.695702859 | 0.942 | 0.884 | 5.17E-06 | Tb |
| TNFRSF9 NK | TNFRSF18 | 4.11E-07 | 0.689817332 | 0.461 | 0.294 | 0.008082335 | Tb |
| TNFRSF9 NK | MYL12A | 2.67E-10 | 0.685989333 | 0.721 | 0.543 | 5.26E-06 | Tb |
| TNFRSF9 NK | ACTB | 1.22E-15 | 0.658022289 | 0.906 | 0.786 | 2.39E-11 | Tb |
| TNFRSF9 NK | UBE2S | 8.78E-11 | 0.654941237 | 0.797 | 0.659 | 1.73E-06 | Tb |
| TNFRSF9 NK | GNLY | 2.58E-07 | 0.646925692 | 0.779 | 0.667 | 0.005071775 | Tb |
| TNFRSF9 NK | RAB11A | 5.35E-09 | 0.642720856 | 0.691 | 0.509 | 0.00010535 | Tb |
| TNFRSF9 NK | JUNB | 4.84E-07 | 0.590613117 | 0.9 | 0.839 | 0.009528677 | Tb |
| TNFRSF9 NK | TUBB4B | 8.25E-09 | 0.549457687 | 0.794 | 0.66 | 0.000162412 | Tb |
| TNFRSF9 NK | SH3BGRL3 | 2.05E-07 | 0.544069345 | 0.721 | 0.592 | 0.004030748 | Tb |
| TNFRSF9 NK | MT-ND3 | 3.78E-10 | 0.531137515 | 0.939 | 0.873 | 7.44E-06 | Tb |
| TNFRSF9 NK | MT-CO11 | 8.73E-11 | 0.513159557 | 0.976 | 0.97 | 1.72E-06 | Tb |
| TNFRSF9 NK | ACTG1 | 6.17E-09 | 0.506359585 | 0.876 | 0.784 | 0.000121452 | Tb |
